# Epigenomic insights into extreme longevity in the world’s oldest terrestrial animal, Jonathan

**DOI:** 10.1101/2025.02.05.636284

**Authors:** Benjamin Vaisvil, Daniel P. Schmitt, Angela Jones, Vinayak Kapatral, James M. Ford, Madison L. Taylor, Mathia Colwell, Jonathan Hollins, Sam Pascucci, Konstantin Weissenow, Burkhard Rost, Pascal Notin, Justin Gerlach, Thomas C. Terwilliger, Li-Wei Hung, Lars Juhl Jensen, Steve Horvath, Christopher Faulk, Yanjun Ma, Stephen W. Clark

## Abstract

Giant tortoises exhibit exceptional longevity, often exceeding the human lifespan. To understand the genomic and epigenomic basis of their longevity, awe analyzed the DNA sequence and methylome of Jonathan, an Aldabra giant tortoise (Aldabrachelys gigantea), estimated to be 192 years old. Relative to other giant tortoises (Aldabrachelys gigantea and Chelonoidis abingdonii), we found Jonathan has gene variants in pathways associated with aging, including DNA repair and telomere regulation. Consistent with his advanced age, Jonathan has significant age-related changes in DNA methylation and methylation entropy, compared with a 5-year-old Aldabra individual. Notably, we found that low entropy regions in Jonathan’s methylome were enriched for genes involved in the electron transport chain. This suggests that high-fidelity transcription of these genes may be crucial for extreme longevity. With this data, we propose a model for aging, that links efficient mitochondrial energy production with nuclear maintenance of low methylation entropy.

---

Recent studies on longevity used comparative genomic analysis to focus on long-lived animals including the naked mole rat, bowhead whale, Brandt’s bat, immortal jellyfish, and long-lived variants of the African turquoise killifish^1–5^. These findings led to a deeper understanding of the gene markers that slow aging and inhibit cancer development in long-lived species. For example, in the bowhead whale, non-synonymous single nucleotide variants (nsSNVs) were identified in the DNA repair machinery (*ERCC1*), epigenetic modifiers (*HDAC1/2*), as well as, duplications in cell cycle regulators (*PCNA*), and the MTOR pathway (*LAMTOR1*)^2^. In the naked mole rat, the genes *TOP2A*, *TEP1*, and *TERF1,* involved in alternative telomere lengthening and critical for longevity, were positively selected^1^. In Brandt’s bat, nsSNVs within the *GH/IGF1* axis involved in growth hormone and insulin-like growth factor were identified^3^. To further explore longevity pathways in long-lived species, we evaluated the genome of the world’s oldest known land animal, Jonathan, an Aldabra giant tortoise (*Aldabrachelys gigantea*), with an estimated age of 192 years (**Supplement Section 1.1, Extended Data Fig. 1-5)**. Giant tortoises have long been known to have long lives, and some species have been found to harbor longevity-associated genetic features^6^. However, no existing study has assessed whether giant tortoises delay or avoid the epigenetic changes that accrue with aging in diverse vertebrate lineages, or identified features potentially unique to giant tortoises like Jonathan, whose lifespan is exceptional, even relative to his conspecifics.

Giant tortoises exist in two genera, the Galapagos giant tortoise (*Chelonoidis niger* group) and the Aldabra giant tortoise (*Aldabrachelys gigantea*). The Galapagos tortoise is endemic to the Galapagos Islands in the Eastern Pacific Ocean, and the Aldabra tortoise is endemic to the Aldabra Atoll in the Western Indian Ocean. Both genera are listed as Vulnerable by the International Union for the Conservation of Nature (IUCN 2024). Three giant tortoises have been sequenced: Lonesome George, a member of the *Chelonoidis abingdonii* species, and two Aldabra giant tortoises (*Aldabrachelys gigantea)*^6,7^. Comparative analyses of these tortoise genomes identified longevity-associated nsSNVs in several genes, including *IGF1R* (insulin pathway), *DCLRE1B* (telomerase pathway), and *XRCC6* (DNA repair pathway)^6^.

Here, we report the genome sequences and comparative analysis of two additional Aldabra giant tortoises, Jonathan and Tank. Tank is a 36-year-old giant tortoise living in captivity in Florida, USA. In contrast, Jonathan lives in a natural ecosystem on the Island of Saint Helena in the South Atlantic Ocean (**Extended Data Fig. 6A**). A recent study using the Species360 Zoological Information Management System (ZIMS) and Bayesian survival trajectory analysis estimated the “adult” life expectancy and aging rates of 457 male Aldabra and 160 Galapagos tortoises^8^. They defined “adult life expectancy” to start after the “age of first reproduction” which they described as 25 years for Aldabra and 20 years for Galapagos. From this, they estimated the mean and upper limit of total life expectancy is 80.47/93.68 (+/-5.64 SD) years for Aldabra and 69.22/87.39 (+/-8.0 SD) years for Galapagos respectively. These data are consistent with previous reports of the life expectancy of Aldabra giant tortoises ranging between 65-90 years^6,9,10^. This suggests that Jonathan is approximately 100 years older than the upper limit of both the Aldabra and Galapagos tortoise and that his unique genome can serve as a model for studying the conserved biological processes of aging.

Epigenetic changes have been associated with maximum lifespan, especially in DNA methylation, where DNA methylation rates scale with maximum lifespan in mammals^11–15^. DNA methylation clocks have been developed for the long-lived naked mole rat and many other mammalian species^16–19^. Here, we present the DNA methylome of Jonathan and compare it to the methylome of a 5-year-old conspecific hatchling. Our analysis reveals age-related modulation of DNA methylation and methylation entropy, with significant enrichment of low-entropy methylation regions observed within the genomic loci of genes involved in the electron transport chain.

## Genome Sequence, assembly and annotation

Given Jonathan’s iconic status on St. Helena and concerns for his health, a blood draw for DNA sequencing was not permitted. Thus, we collected several buccal scrapes and saliva samples from Jonathan for gDNA isolation. Several attempts to isolate high molecular weight (HWM) gDNA from Jonathan’s samples failed to yield samples of quality sufficient for long-read sequencing, so we instead used short-read 2×150bp Illumina sequencing. We obtained approximately 1.1 billion reads for a total of 165 Gb (gigabases). A permitted blood draw from Tank yielded HMW gDNA, and we generated an Aldabra reference genome using long-read PacBio HiFi and OmniC Hi-C reads for large scaffolds.

Tank’s assembled genome consists of 211 scaffolds with a size of approximately 2.41 Gb (**Extended Data Fig. 7A,B and Supplementary Table 1**). These metrics are similar to those previously reported for assemblies of the Aldabra genome (2.37Gb) and the Chelonoidis genome from Lonesome George (2.3 Gb)^6,7^. Jonathan’s Illumina reads achieved a 93% alignment rate against Tank’s genome, with an average coverage depth of 60x. A total of 6.9 million high-quality variants were called from this alignment and were used to create a consensus reference genome for Jonathan with a size of approximately 2.41 Gb. Using this consensus assembly, 25,738 protein-coding genes were predicted of which 16,466 were functionally annotated (**Extended Data Fig. 7B**). The mitochondrial genome was assembled to a single contig and was 100% identical to the Aldabra reference mitochondrial genome (NC_028438.1) (**Supplementary Table 1**).

## Jonathan’s Biogeography

Current theory on the dispersal route of Aldabras “out of Africa” is that they first dispersed from the eastern coast of Africa to Madagascar, and from there to Granitic Seychelles and eventually to the Aldabra atoll (**Fig. 1A**)^20^. The Aldabra atoll is the westernmost island in Seychelles and is a protected UNESCO World Heritage Site that harbors around 100,000 Aldabra tortoises^21^. The Aldabra atoll comprises four islands (Malabar, Grande Terre, Polymnie, and Picard) (**Extended Data Fig. 6B**). Picard tortoises were harvested to extinction in the 1800s but were later repopulated by tortoises from Malabar and Grande Terre^22^. Interestingly, Polymnie Island harbors no tortoises despite a similar ecosystem and climate. Recent studies evaluated island genetic differentiation within the Aldabra population using low-coverage sequencing of 30 Aldabras tortoises (from islands Malabar and Grande Terre) and two (non-island) Zurich Zoo Aldabra tortoises (*Aldabrachelys gigantea)*, named Hermania and Maleika^7,23^. Two distinct groups were identified using principal component analysis (PCA) and 6 million single nucleotide variants (SNVs). Hermania and Maleika were grouped with the Grande Terre tortoises indicating their common origin.

**Figure 1.**
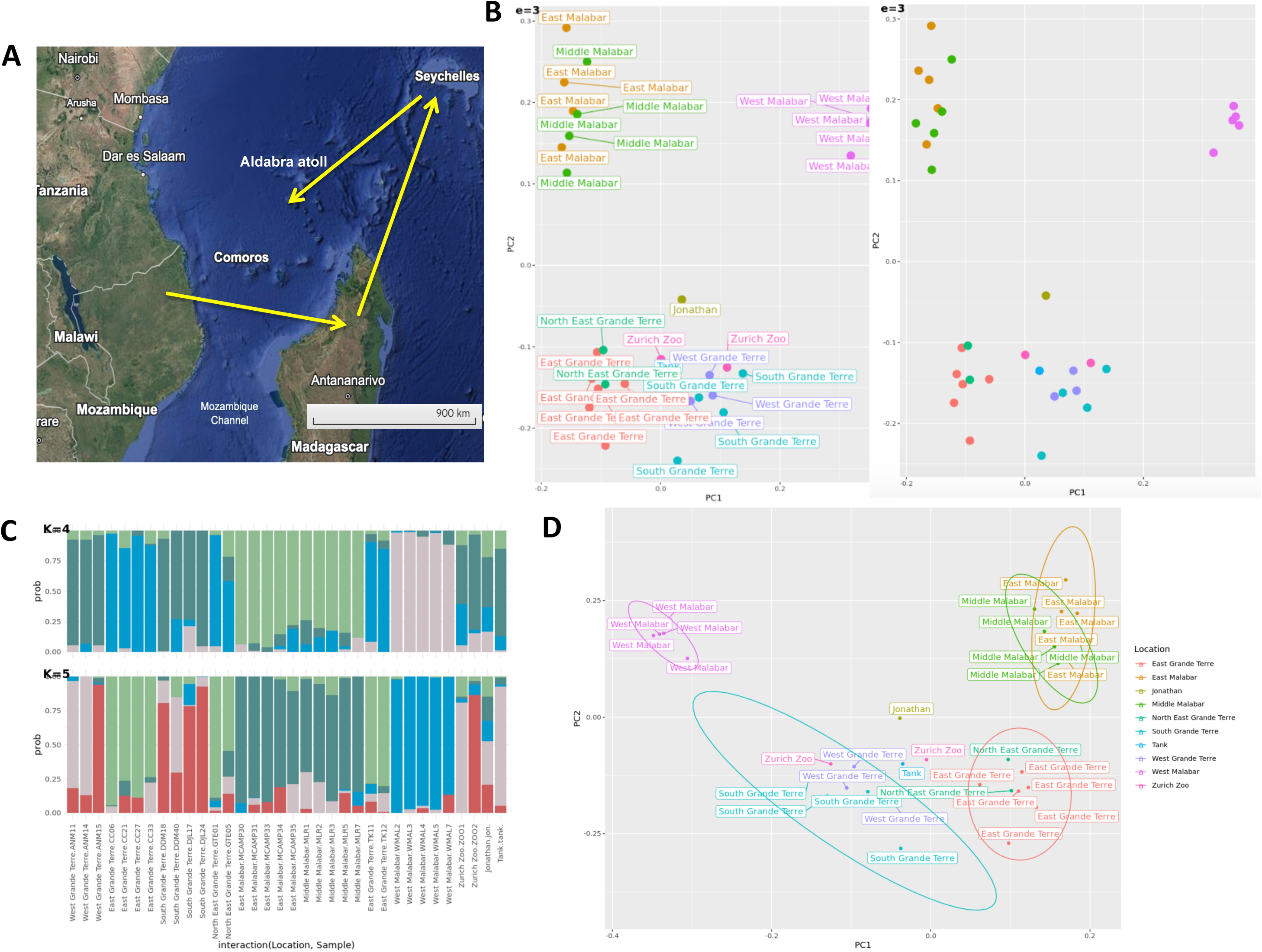
Jonathan’s Island origin location. **A.** The current theory on the dispersal route of Aldabras out of Africa^11^. **B, D**. Principal component analysis plot of 30 *Aldabrachelys gigantea* genomes from a previous study from the Islands of Malabar and Grande Terre of the Aldabra atoll. Included are the two Zurich Zoo Aldabras^10^. This study adds Tank and Jonathan. Principal components 1 and 2 represent 14.2% and 3.64% of the overall genetic variability, respectively. **C.** An ancestral population k-mer plot for 4 or 5 k-mers for each individual.

Given Jonathn’s exceptional age and the above Aldabra dispersal route, we used this group’s data to determine the potential birthplace of Jonathan and Tank. We generated variants for Jonathan, Tank, Lonesome George, and the other tortoises using Hermania as the reference genome^7^. From those variants, we computed a PCA. We found that Tank, like Hermania and Maleika, is close genetically to those Aldabras associated with the western part of Grande Terre (**Fig. 1B,D**) whereas, Jonathan was proximal to Grande Terre but further from it than either Tank, Hermania, or Maleika. We speculate this is explained by either Jonathan being removed from the Aldabra Atoll much earlier than the other tortoises or he was removed from Granitic Seychelles; the former is more likely, as it is thought that sailors completely extirpated the Aldabras present on Granitic Seychelles by the early 19th century (1800-1830)^24^. Thus, if Jonathan is 192 years old, his birth year would be 1832, making it less likely that he was removed from Granitic Seychelles, however, if he is older than 192, this is plausible. In comparison, we analyzed Lonesome George, and as expected, he falls outside the island chain in the PCA plot (**Extended Data Fig. 8**).

In addition to PCA, the previous studies used unsupervised Bayesian clustering and pairwise linkage analysis to assess the genetic structure of each island^7,23^. In an admixture plot where K = 4, Jonathan is closest to Hermania, Maleika, and an individual from South Grande Terre, but when K=5, Jonathan’s admixture is unique among the individuals sampled (**Fig. 1c**). Overall, these data show Jonathan has a contribution from an unknown source. However, it is impossible to conclude from these data if that unknown source is because the background population changed or because Jonathan was extirpated from a different island.

## Positively selected genes

To understand Jonathan’s specific evolutionary adaptations and survival, we analyzed our predicted gene set for signs of positive selection, i.e., with a ratio of nonsynonymous to synonymous mutations (dN/dS) greater than 1. Predicted genes from Jonathan were tested for positive selection as the foreground branch while other long-lived and short-lived species were set as the background. We found 770 significantly (p < 0.05) positively selected genes (PSGs) (**Fig. 4A and Supplementary Table 2**). The functional impact of amino acid changes within Jonathan’s PSGs was evaluated using the protein algorithms SIFT and PROVEAN, an approach previously employed in studying other long-lived species (**Supplementary Table 2**)^4,5,25^. In Jonathan’s genome, we identified PSGs involved in DNA repair (*POLE2*, *BARD1, RHNO1*, *XAB2*, *RAD23A*, *RAD51B*, *XRCC5*, *DDB2*, *RFC1*, *SIRT4, POLQ*, and *FANCL*), insulin regulation (*SOCS2, NOX4, APPL1, APPL2*), mitochondrial function (*ALDH2, HIGD1A, VPS13C, STOX1, GSTZ1, FSTP1*) and telomere function (*POT1, RFC1, LSM11*)^26^. Interestingly, *LSM11* is part of the U7 snRNP-telomerase holoenzyme complex and is implicated in the Aicardi-Goutieres syndrome. This rare human disorder affects white matter in the brain and causes premature aging in some individuals^27–29^.

Similarly, *HAS2* (*hyaluronic synthase 2*), a gene associated with soft tissue sarcoma and mucopolysaccharidosis in humans, is positively selected in Jonathan^30^. *HAS2* is also involved in excess hyaluronic secretion in the naked mole rat, and transgenic mice become cancer-resistant and have extended lifespans and healthspans when HAS2 is overexpressed^31,32^.

Gene ontology enrichment analysis was performed on the PSGs and revealed enrichment for components in development, metabolism, and immunity (**Extended Data Fig. 9)**. Pathways related to aging were identified including telomerase RNA localization, NAD biosynthesis, insulin regulation, and DNA repair. NAD levels have been shown to decrease with age in humans, and to our knowledge, this has not been studied in giant tortoises^33^.

Next, we used the STRING database to construct a functional association network of the PSGs and computed clusters to identify functional modules^34^. We found clusters related to DNA repair, mitochondrial function, insulin regulation, oxidative stress response, and heart muscle contraction prominently represented in Jonathan’s genome (**Extended Data Fig. 10**). The identification of a gene cluster related to cardiovascular function is noteworthy, as limited research has been conducted on the Aldabra tortoise’s cardiovascular system. Existing studies primarily focus on echocardiographic imaging, revealing a remarkably low resting heart rate of 21 beats per minute^35^. In contrast, most cardiovascular research on other long-lived species has centered on the naked mole rat, exploring its unique cardiac function and myofilament protein profile^36,37^.

## Non-synonymous single nucleotide variants (nsSNVs) shared among Lonesome George, Aldabras, and Jonathan, but absent in humans

Previous work on Lonesome George manually annotated 2,629 genes selected “a priori from a series of hypothesis-driven studies” on longevity^6^. The authors identified and confirmed 13 nsSNVs that affected known gene motifs or had human counterparts related to a known genetic disease from this set. Jonathan shares these same nsSNVs including genes involved in DNA repair (*ATM*, *XRCC5*, *XRCC6*, *TP53*), telomere function (*DCLRE1B*), glucose or insulin regulation (*MIF*, *IGF1R*, *IGF2R*, *GSK3A*), mitochondrial function (*ALDH2*), cell adhesion (*CDH1*), and neurological development (*CNDP1*, *XPNPEP1*, *PSEN1*)(**Extended Data Fig. 11A-G**)^26^.

## Unique Jonathan nsSNVs

Since Jonathan’s age far exceeds the typical age of giant tortoises, we compared his genome to that of Tank, Lonesome George, and humans. Employing the same gene set used in the Lonesome George study, we identified 287 genes harboring unique nsSNVs found exclusively in Jonathan (**Supplementary Table 3**).

We used four protein language models (Tranception, ESM1v, Progen2, and RITA), and one variant effect predictor (VESPA), to evaluate these nsSNVs^38–43^. These models were chosen for superior performance and sequence alignment independence compared to SIFT and PROVEAN. We computed the log-likelihood ratio of the mutated human sequence (with Jonathan’s amino acid inserted) versus the reference human sequence, and the log-likelihood ratio of the mutated Jonathan sequence (with human amino acid inserted) versus Jonathan’s sequence for each gene (**Supplementary Table 4**). This analysis identified 41 genes that were predicted by at least two models to have functionally significant nsSNVs (**Fig. 2B and Supplementary Table 5**). Of the 41 genes, 12 were identified in the aging databases LongevityMap and GenAge, of which two are part of the telomerase shelterin complex (*ACD* and *POT1*), two in the DNA repair pathway (*ERCC1* and *TP53BP1*), one in autophagy (*FOXO3*), and one a tumor suppressor (*CIC*) (**Supplementary Table 5**)^26,44–46^. Interestingly, a *POT1* variant was found in the immortal jellyfish (*Turritopis dohrnii*), a *FOXO3* variant is associated with longevity in Okinawans, and an *ERCC1* variant was found in the bowhead whale^2,5,47^.

**Figure 2.**
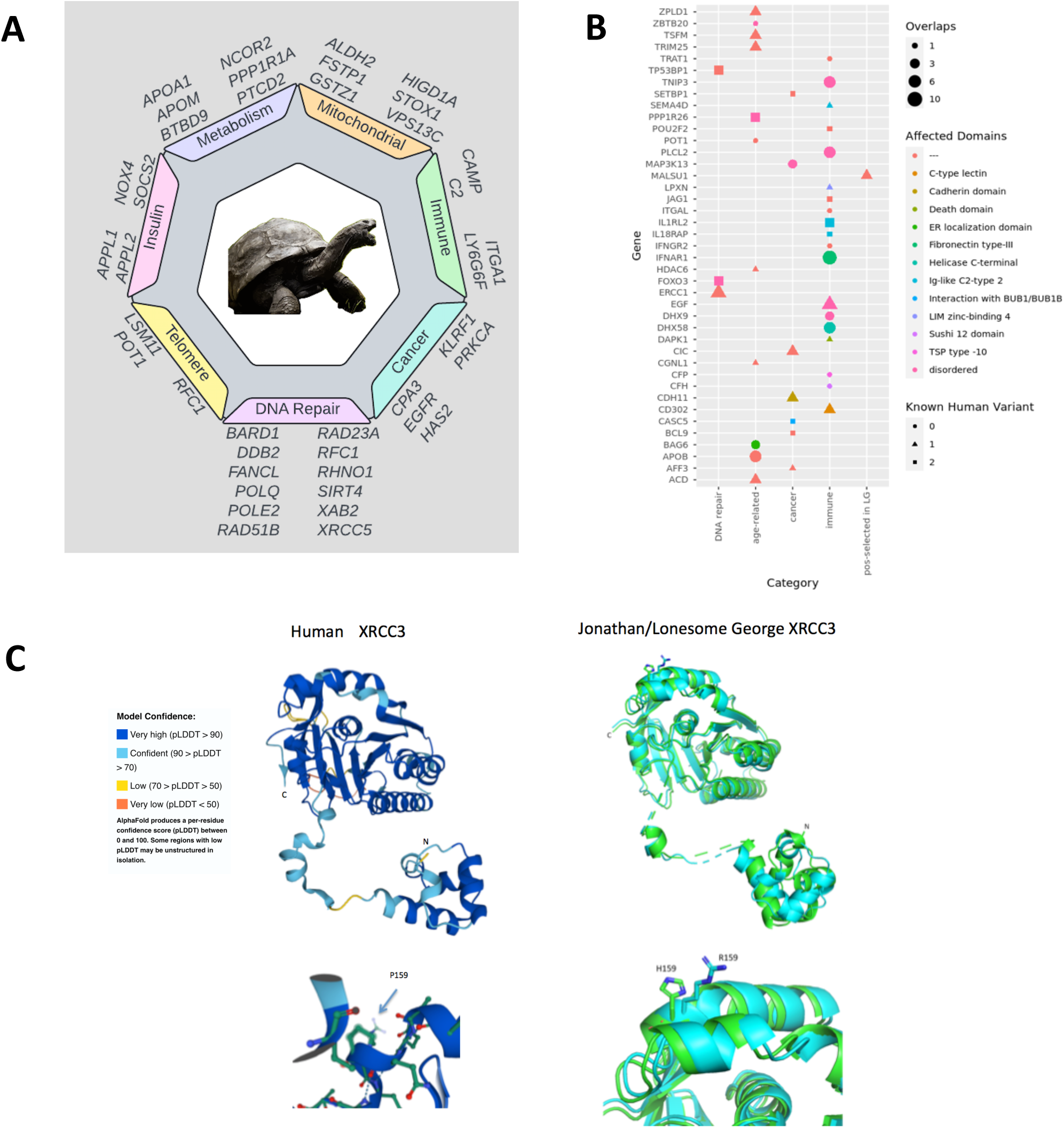
Evolutionary analysis of Jonathan’s genome. **A.** A subset of genes under positive selection in Jonathan and categories of function they fall into. **B.** Jonathan’s unique non-synonymous nucleotide sequence variants (nsNSVs) predicted to be significant by protein language models. **c**, Alphafold2 predictions; Left panel: Alphafold prediction found in UniProt of human *XRCC3* and model confidence; Right panel: AF2 predictions for Jonathan’s (cyan) and Lonesome George’s (green) *XRCC3*. The N and C-termini are labeled. Amino acids are labeled using single-letter codes: proline (P), histidine (H), and arginine (R).

Using AlphaFold2 (AF2), we generated protein structure predictions for all 287 genes of Jonathan with unique nsSNVs and their corresponding orthologs in Lonesome George (**Supplementary Table 3**)^48^. We primarily observed side-chain differences between the orthologs. For example, Jonathan’s gene, *XRCC3*, contains a hydrophilic arginine (P159R), and Lonesome George’s a hydrophilic histidine (P159H). In contrast, humans have a hydrophobic proline (**Fig. 2C**). Although in humans an XRCC3 variant P159R (rs199681385) is found in UniProtKB, this nsSNV is predicted to be benign and tolerated by Polyphen-2 and SIFT, respectively^49–51^.

## Gene expansion

Previous work identified 12 tortoise-specific gene families with increased copy number^6^. Jonathan has gene expansion in 10 of 12 genes (**Supplementary Table 6).**

*LAMTOR4*, a member of the Ragulator pentameric protein complex (*LAMTOR1-5*) that helps regulate *mTORC1,* a well-studied pathway in human longevity, is one of the expanded tortoise-specific genes^52^. Detailed analysis of *LAMTOR4* reveals that Lonesome George and all three Aldabra genomes (Jonathan, Tank, and Hermania) possess two copies of this gene (LAMTOR4JON1, LAMTOR4JON2) (**Extended Data Fig. 12A-C**). The second copy (LAMTOR4JON2) has five corresponding human variants (S41R, R54Q, P62L, H73N, and P96S), two of which have been predicted to be damaging by SIFT and PolyPhen-2 in UniProt (S41R. rs1798774135; P96S, rs371410641)^49^. Likewise, the bowhead whale has a duplication of *LAMTOR1*, with the extra copy containing several nsSNVs not found in the other gene^2^. Lastly, the killifish has positive selection for *LAMTOR5*^4^.

*VCP,* another tortoise-specific gene, was duplicated in both Lonesome George and Jonathan. *VCP* is a member of the AAA ATPase family of proteins involved in DNA damage response, where it is recruited to double-strand breaks (DSBs) sites and promotes the recruitment of TP53BP1^53^.

Expansion in non-tortoise-specific genes was identified in 16 families in both Jonathan and Lonesome George, including 4 tumor suppressor genes (*NF2*, *SMAD4*, *PML*, *P2RY8*), 9 immune genes (*APOBEC1*, *CHIA*, *CMAIL*, *CTSGL*, *GZMB*, *GZMH*, *PRF1*, *TLR2*, *TLR5*, *TLR8*, *TLR13*), two cancer genes (*MYCN*, *USP12*), and one DNA repair genes (*NEIL 1*)(**Supplementary Table 6**)^6,26^. On closer review of the perforin gene (*Prf1*), Jonathan had 10 copies while Lonesome George had 7. Strikingly, if this pore-forming protein is deleted (*Prf1^-/-^*) in mice, senescent cells accumulate with age^54^.

## Methylome analysis

Buccal DNA from Jonathan and a 5-year-old Aldabra hatchling was used for whole genome bisulfite sequencing (WGBS). Raw reads from Jonathan were 885M and from the hatchling 1,272M, resulting in 133 (Jonathan) and 192 (hatchling) raw Gb of paired-end sequence data. From this 73 (Jonathan) and 60 (hatchling) Gb were successfully mapped to our reference genome (Tank), providing 30-fold (Jonathan) and 25-fold (hatchling) sequencing depth. The bisulfite conversion rate was determined to be 99.4% (Jonathan) and 99.4% (hatchling). The WGBS data from Jonathan and the hatchling is illustrated in **Fig. 3A**.

**Figure 3.**
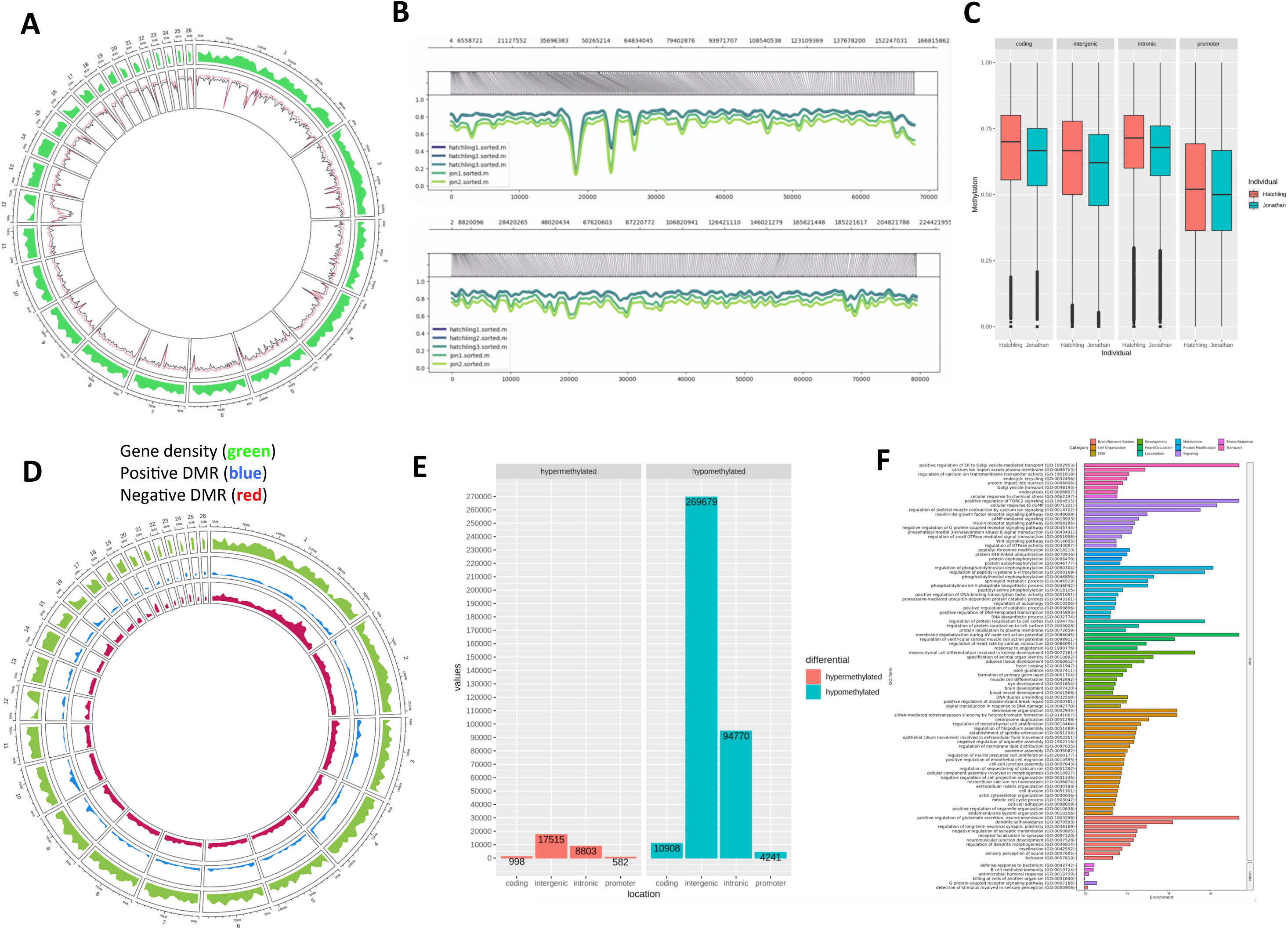
DNA methylome analysis, comparing Jonathan and hatchling. **A.** Circos plot of global DNA methylation of Jonathan (black) and hatchling (red) and gene density (green). **B.** Global DNA methylation of Jonathan and hatchling for contigs 1 and 8. **C.** Genomic regions of DNA methylation for Jonathan (green) and hatchling (red). **D.** Circos plot of DNA methylated regions (DMRs) in Jonathan with red (negative DMR) and blue (positive DMR), and green (gene density) across all contigs. **E.** Genomic regions for hypomethylation and hypermethylation in DMRs. **F.** GO analysis on hypermethylated DMR regions.

Comparing the two methylomes, we found a global decrease in DNA methylation in Jonathan, with the average methylation decreased for all covered CpGs (**Fig. 3A**). This decreased global methylation was appreciated across all chromosomes and genomic regions (**Fig. 3B, C**).

Given the discordance between the methylomes, we examined the WGBS data for differentially methylated regions (DMRs) (**see methods**). DMRs were identified across Jonathan’s and the hatchling’s genomes (**Fig. 3D**) In total, we identified 407,483 DMRs between Jonathan and the hatchling, of which 379,598 (93.2%) were hypomethylated in Jonathan. The hypomethylated DMRs were mostly confined to intergenic (71%) and intronic (25%) regions, with only 4% in either the promoter or exon (**Fig. 3E**). Similarly, the hypermethylated regions were located within the intergenic (63%) and intronic regions (31%) regions and 6% were located within the promoter or exon. The hypermethylated DMR regions encompassed 2,944 genes (**Supplementary Tabel 7**). GO analysis of these genes revealed significant involvement in insulin receptor recycling, positive regulation of TORC2 signaling, and many developmental pathways, especially cardiovascular development (**Fig.** 3F)(Supplementary Table 7).

In mammalian embryonic stem cells, age-related increases in DNA methylation are observed at specific genomic regions known as target sites of the Polycomb repressive complex 2 (PRC2)^19^. We identified several PRC2 target genes in Jonathan’s hypermethylated DMRs (**Supplementary Table 7**). Moreover, 159 genes had hypermethylated DMRs spanning their promoter and coding regions, these genes included developmental genes (*WNT5B*, *HOXA7*, and *IRX6),* cell cycle genes (*RGCC, EAPP*, *CDCA7L*), and genes involved in *cytochrome c oxidase* (*COX16*, *COX4I2*, *COA3*) (**Supplementary Table 7**)^26^.

Previous work has shown that information is encoded within the DNA methylome, and over time, stochastic changes in methylation can lead to information loss or what is defined as methylation entropy (ME)^11,55^. Additional studies revealed that ME increases with age, and increasing ME is associated with increasing gene expression variance (i.e. transcriptional noise, TN)^55–60^. We compared the ME in Jonathan’s and the hatchling’s methylome, and consistent with findings in humans, Jonathan had increased ME (**Fig. 4A, B**)^60^. The increased ME occurred in all genomic regions, except at the promoter regions, where Jonathan and the hatchling had equivalent and low ME (**Fig. 4C**).

**Figure 4.**
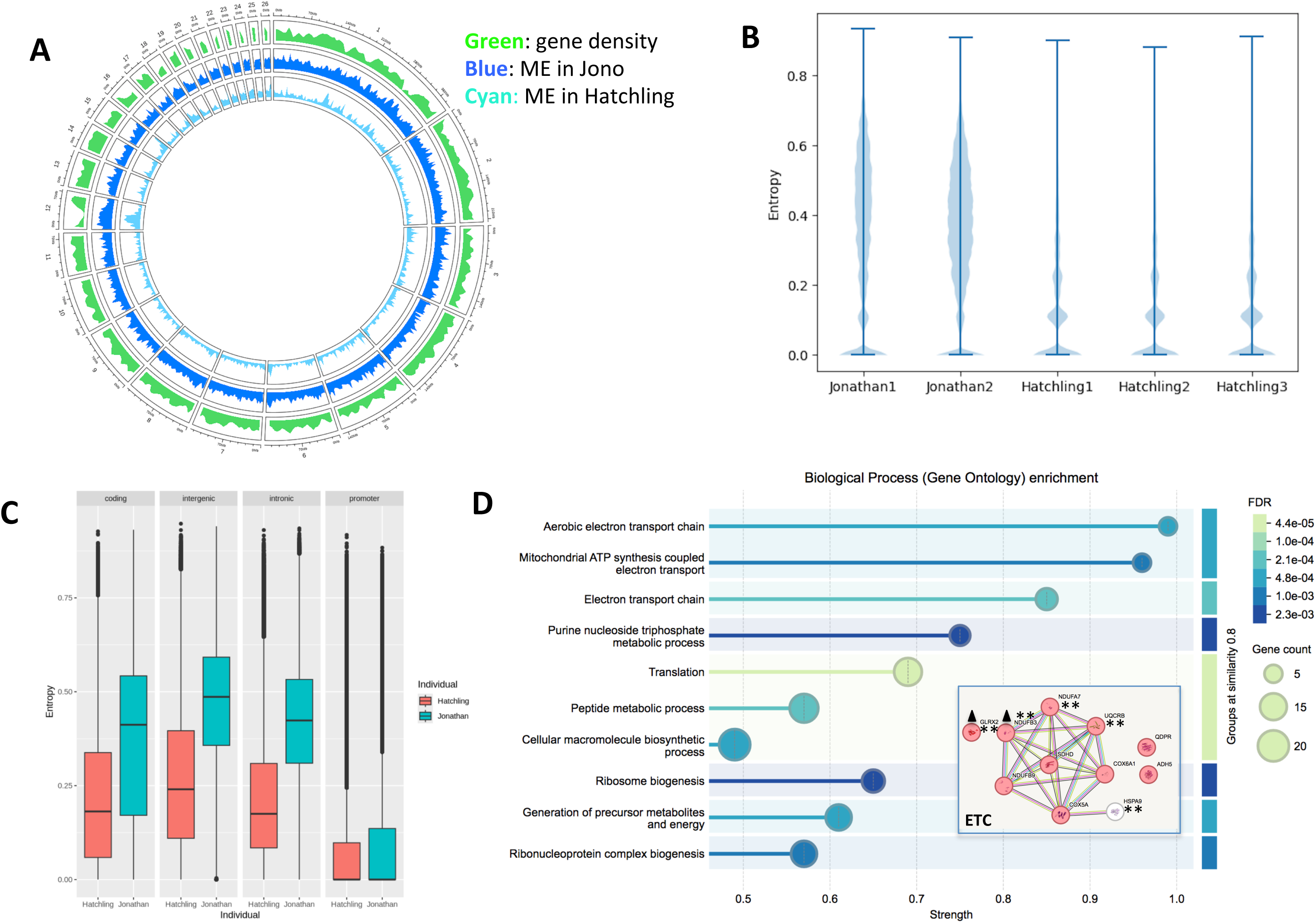
Methylation entropy (ME) in Jonathan and hatchling. **A.** Circos plot of methylation entropy, comparing Jonathan’s ME (blue) with the hatchling (cyan) and associated plot of gene density (green). **B.** ME density plot comparing ME variance between Jonathan’s samples and the hatchling. **C.** Genomic regions of ME. **D.** GO enrichment using the STRING database and the 169 genes within Jonathan’s low entropy regions (LERs). Inset, red circles, the 10 genes within Jonathan’s LERs involved in the electron transport chain (ETC), and white circle, the gene involved in mitochondrial integrity. **\*\*** Genes identified in the Open-Genes database for human longevity. The triangle represents genes involved in mediating reactive oxygen species (ROS).

Because ME increases with age, it has been postulated that lower entropy regions (LERs), are more likely to harbor genes of importance, as maintaining lower entropy would require energy expenditure^60^. We examined Jonathan’s methylome for lower entropy regions (LERs). LERs were defined as regions one standard deviation below the mean ME. 1,253,905 LERs meet this criterion in Jonathan and 1,067,310 LERs in the hatchling. Of Jonathan’s LERs, only 17 regions had lower entropy than the hatchling (**Supplementary Table 8**). We identified 11 genes within these LERs, two are involved in DNA repair (*HPF1* and *SETMAR*)^26^. Five genes are mitochondrially related, one (*UQCRFS1*) is involved in the electron transport chain (ETC) and two (*GLRX2*, *MAFF*) are involved in oxidative and cellular stress^26^. Lastly, one gene, *CEBPD*, is involved in immune and inflammatory responses^26^. To identify more genes within LERs we allowed LERs in Jonathan to be within 0.005 of the median ME of the hatchling. This identified an additional 169 genes with LERs in their promoters. We then cross-referenced this set of 169 genes with the “Open-Genes” human longevity database and identified 37 genes that have been associated with human longevity (**Supplementary Table 8)**^61^. Of these, the highest confidence was in the gene *RPSKB1*, a downstream effector of *mTORC1*. Of the remaining genes, five genes (*POLB*, *TAF9*, *RAN, PTTG1,* and *GLRX2*) are involved in DNA stability, two genes are involved in reactive oxygen species mediation (*GLRX2*, *NDUFB3*), and four genes are involved in mitochondrial integrity (*HSPA9*, *MTFP1*, *MRPL49*, *QRSL1*)^26,61^. Moreover, the gene *UPF1* emerged as an intriguing gene in our analysis. *UPF1* encodes for a nucleic acid-dependent ATPase and 5’-to-3’ helicase, and it is known to function in RNA quality control but it also interacts with telomeres and its depletion leads to telomere shortening^62^.

GO analysis of the 169 genes revealed enrichment for the electron transport chain (ETC), ribosome biogenesis, and cellular localization to the inner mitochondrial membrane (**Fig. 4D and Extended Data Fig. 13**). Twelve genes were associated with mitochondrial integrity of which 11 genes (*COQ3*, *QDPR*, *ADH5*, *GLRX2*, *SDHD*, *NDUFA7*, *UQCRB*, *NDUFB9*, *COX5A*, *NDUFB3*, and *COX6A1*) were specifically associated with the electron transport chain (ETC) (**Fig. 4d and Supplementary Table 8**)^26^. Three of these genes (*NDUFA7*, *NDUFB9*, *NDUFB3*) are subunits of Complex I and five genes (*GLRX2*, N*DUFA7*, *UQCRB*, *NDUFB3*, and *HSPA9*) are associated with human longevity (**Fig. 4D**)^26,61^.

## Discussion

Genome stability is a hallmark of long-lived species, and we found Jonathan to be no exception to this rule. Our genomic analysis highlights multiple unique variants involved in pathways such as DNA repair, telomere maintenance, autophagy, insulin regulation, mitochondrial function, and cancer. Moreover, we confirmed the expansion of the Ragulator subunit, *LAMTOR4*, in Jonathan^7^. *LAMTOR4* expansion is compelling, in that the extra copy contains two variants predicted to be damaging, and thus could exert a dominant negative effect, partially inhibiting mTORC1.

Here we also present the first whole-genome bisulfite sequencing (WGBS) data of a giant tortoise, revealing an age-related decline in global DNA methylation in Jonathan, an Aldabra tortoise estimated to be 192 years old. This observation aligns with previous findings in humans demonstrating decreased methylation in centenarians compared to newborns^63^. We also find Jonathan’s hypermethylated DMRs are enriched for developmental genes and targets of the PRC2 complex. These findings are consistent with established data showing that small regions of the genome are hypermethylated with age, and these areas occur in GpC-rich-regions that often encompass PRC2 target genes^19,64–67^. It has been proposed that the hypermethylation of PRC2 targets leads to loss of stem cell plasticity, resulting in age-related tissue decline^66^.

Jonathan’s methylome exhibited greater methylation entropy (ME) than the hatchling across most genomic regions, except for promoters, where ME was comparable. This pattern mirrors the human findings in long-lived individuals (LLIs), where ME increases with age, but is reduced in promoter regions of centenarians compared to elderly individuals^60^. Using methylation data, this group further identified lower entropy regions (LERs) in LLIs compared to elderly controls, particularly in active promoters of DNA repair genes. Other work has shown that ME in promoter regions leads to gene expression variability (i.e. transcriptional noise, TN)^59^. These studies imply that increasing ME, especially within promoter regions, increases transcriptional noise (TN), decreasing gene function. Thus, genes within actively maintained LERs, particularly in long-lived individuals, could be critical for longevity.

Given the potential importance of LERs in longevity, we identified LERs in Jonathan’s methylome; his LERs contain genes involved in DNA repair and to a lesser extent RNA metabolism. These results are consistent with findings in long-lived human LERs^60^. Remarkably, we also found enrichment in mitochondrial-related genes, especially the electron transport chain (ETC). We are not aware of such an enrichment in human LLIs. This implies that high-fidelity ETC gene transcription may be required for especially long-lived species.

Mitochondria are the source of both ATP and ROS generation, and mitochondrial dysfunction is one of the hallmarks of aging^68,69^. Previous work has shown that longer-lived species have decreased mitochondrial reactive oxygen species production (mitROSp) and decreased free radical leak (FRL) within the ETC compared to shorter-lived species^70–72^. Both Complex III (CxIII) and Complex I (CxI) are responsible for ROS production, but studies using selective ETC inhibitors localized the site for the drop in mitROSp and FRL in long-lived species to CxI^73^. Moreover, recent work in older human oocytes (34 years old), revealed they reduce mitochondrial ROS generation by suppressing CxI^74^. Intriguingly, three genes (*NDUFA7*, *NDUFB9*, *NDUFB3*) identified within Jonathan’s LERs are CxI subunits^26^. It is possible these genes could also affect other aspects of mitochondria integrity, such as the proton motive force (PMF), so these findings will need further validation^75^. Thus, we suspect Jonathan’s mitochondria are unique for mitROSp, FRL, or PMF, and maintaining the fidelity of the ETC is a key component of his extreme longevity.

In summary, aging is associated with distinct changes in the DNA methylome, including global DNA hypomethylation, hypermethylation at PRC2 target genes, and increased methylation entropy (**Fig.5A**)^11,57,76^. Building on the methylation entropy findings in long-lived humans and Jonathan, we propose a model for aging that integrates methylome changes with increasing entropy and loss of LERs^60^. We postulate that maintaining LERs that encompass genes involved in mitochondrial ATP production (e.g. ETC genes) and nuclear repair, is essential for the efficient function of both processes, likely by reducing transcriptional noise within these pathways (**Fig. 5B**). Efficient mitochondria, in turn, provide the necessary energy (ATP) to maintain efficient nuclear repair. Creating a positive feedback cycle, efficient nuclear repair and efficient energy production help maintain the low entropy state of the DNA regions. Consequently, our model suggests that mitochondrial decline may precede increased entropy in long-lived species. This model is supported by work showing long-lived species have more efficient mitochondria and the work in long-lived humans and the work here showing LERs in DNA repair and ETC genes^60,70–72^. However, further research is needed to validate and refine this model, including investigating the mechanisms that maintain LERs and how mitochondrial efficiency is regulated in long-lived species.

**Figure 5.**
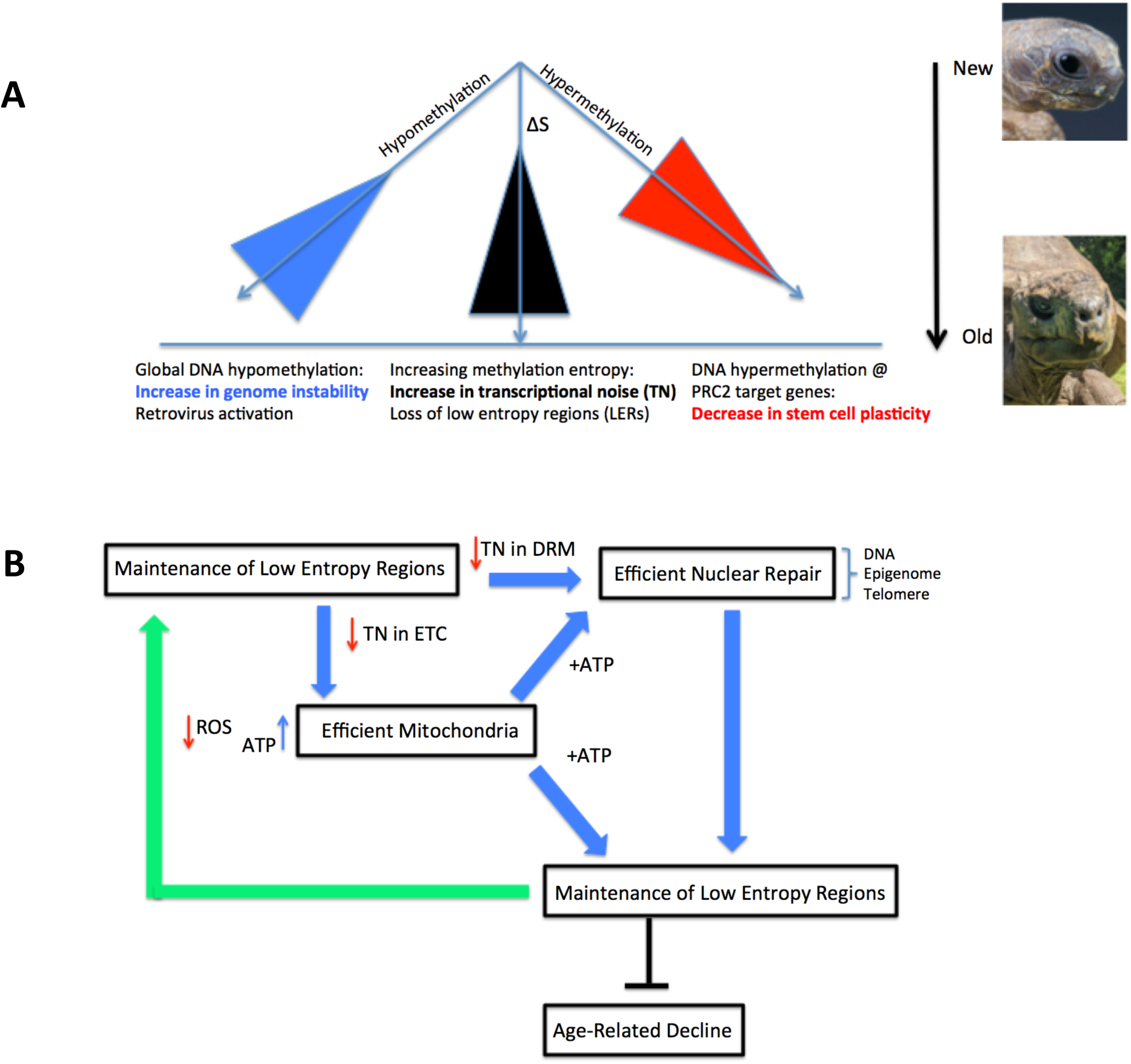
Aging methylome characteristics and aging model. **A.** Aging methylome: three concurrent events happen with age, global hypomethylation, resulting in increasing genomic instability (represented by an upright blue triangle, the tip represents a stable genome but genomic instability increases as you move toward the base); hypermethylation at PRC2 target gene sites leading to decreased stem cell plasticity (represented by the inverted red triangle, the base of the triangle represents high plasticity that decreases as you move down to the tip); and increasing methylation entropy, leading to loss of low entropy regions (LERs) and increasing transcriptional noise (TN) (represented by an upright black triangle, the tip of the triangle represents low transcriptional noise and the base represents transcriptional noise). Jonathan’s picture was provided by Jonathan Hollins of St. Helena. Hatchling picture credit: reptiles4all/Shutterstock. **B.** Mitochondrial-Entropy-Genome-Stability model for aging in extremely long-lived species. Efficient mitochondria provide ATP to maintain low entropy and efficient DNA/epigenetic repair. Low entropy regions reduce transcriptional noise (TN) in the electron transport chain (ETC) and DNA repair mechanisms (DRM). “Efficient Nuclear Repair” encompasses DNA/telomere repair and epigenome maintenance. Efficient mitochondria and nuclear repair help maintain low methylation entropy.

In this study, we were limited because blood could not be used for DNA isolation. This prohibited us from using: long-read sequencing technologies, isolation of RNA, use of RNA seq to verify Jonathan’s unique nsNSVs, and the relationship between Jonathan’s increasing methylation entropy with transcriptional variance. To partially overcome this limitation, we generated a reference genome from Tank and used state-of-the-art protein language models to screen for potentially impactful nsNSVs.

In conclusion, this study presents the first genomic and epigenomic characterization of Jonathan, the world’s oldest known living land animal, highlighting unique molecular features associated with his exceptional longevity. Drawing upon both previous findings in long-lived humans and our study of Jonathan, we propose an aging model that emphasizes the importance of maintaining low-entropy regions in genes essential for DNA repair and energy production^60^. This work should provide a valuable resource for future research into the mechanisms of aging and inform conservation strategies for Testudines by identifying cryptic diversity and inbred lineages in restricted populations.

## METHODS

### Buccal scrape

Scrapes were taken on a previous occasion and acquired only plant DNA. To avoid this, a succession of scrapes were taken using two different opportunities:

Early morning before the tortoises venture out to graze, giving a chance for overnight salivation and swallowing to cleanse exogenous organic debris from the buccal cavity.
Although the Island of St Helena is in the tropics, the maritime climate is subtropical and the tortoises are kept at 1,700 ft (500m) altitude. Days and nights can be cool, especially in winter. During periods of inclement weather Jonathan will cease grazing for several days and hunker down in heaped grass clippings, put under trees for this very purpose (fermentation in the grass creates warmth which the giant tortoises seek out).

Perhaps the hardest obstacle is physical resistance. Jonathan has clear taste preferences and responds rapidly to anything alien or unpalatable. He loathes kale, for example, and with the weekly hand feeding made this clear by using his forelegs to resist and remove it. The same applies to buccal scrapes. This was overcome, albeit with some difficulty, by pretending to feed Jonathan, whereupon through conditioning he would start biting the air, then by wedging the mouth open with the left hand protected in a leather welding glove while swabbing under the tongue with the right hand. It was also found possible to hold two swabs at the same time between different fingers. Jonathan would almost immediately dislodge the right hand with an upward sweep of his left foreleg, but in this manner it was possible to create an approximately 5-second window for acquiring samples.

### DNA Extraction from Jonathan’s buccal scrapes

DNA was extracted from the buccal scrapes using the Buccal-Prep Plus DNA Isolation Kit (Isohelix) kit with minor modifications. Briefly, 20µl PK solution was added to the tube and incubated in a 60°C bath for 1 hour. The liquid in the tube was transferred into a centrifuge using a sterile pipette tip. The tube was centrifuged for 2 minutes at 18,000 x g. The supernatant was recovered and added to the 400µl sample for a total volume of 500µl. The sample was vortexed briefly. Next, the tube was placed in a microcentrifuge and spun at 12,000 x g for 10 minutes. The supernatant was poured off and the tube re-spun. 50µl of TE solution was added to the tube. The pellet was resuspended in TE and left at room temperature for 10 minutes. The sample was then assessed by nanodrop.

### DNA Extraction from Tank’s Blood

High Molecular Weight genomic DNA (HMW gDNA) was extracted from blood with the MagAttract HMW DNA kit (Qiagen) with minor modifications. Briefly, 10µl of blood was incubated with 490µl buffer AL, and incubated overnight at room temperature with 20µl of proteinase K and 4ul RNase A, followed by incubation at 65°C for 10 minutes. The lysate was brought to room temperature, and 10µl MagAttract suspension G (beads) and 350 µl buffer MB were added to the lysate for DNA binding. Bead washings were done twice with 700µl each of buffer MW1, buffer PE, and 80% EtOH. DNA was eluted four times in 100µl of buffer AE each time.

### Jonathan’s Illumina Libraries and Sequencing

#### Jonathan and Hatchling’s Whole Genome Bisulfite Sequencing

A Quality Control analysis was performed on the DNA samples using a DNA Qubit assay to determine quantity. The samples were normalized, and libraries were prepared using the NEBNext® Enzymatic Methyl-seq kit (P/N: E7120L) following a modified version of the manufacturer’s protocol. The library quality was assessed using a Bioanalyzer and quantified using a qPCR-based method with the KAPA Library Quantification Kit (P/N: KK4873) and the QuantStudio 12K instrument.

Prepared libraries were pooled in equimolar ratios, and the resulting pool was subjected to cluster generation using the NovaSeq 6000 System, following the manufacturer’s protocols. 150 bp paired-end sequencing was performed on the NovaSeq 6000 platform targeting 400M reads per sample. Raw sequencing data (FASTQ files) obtained from the NovaSeq 6000 was subjected to quality control analysis, including read quality assessment. Real-Time Analysis Software (RTA) and NovaSeq Control Software (NCS) (1.8.0; Illumina) were used for base calling. MultiQC (v1.7; Illumina) was used for data quality assessments.

#### Whole Genome Sequencing

A Quality Control analysis was performed on the DNA sample using a DNA Qubit assay to determine quantity. The sample was normalized, and the library was prepared using the Twist Biosciences kit (P/N: 104207) following the manufacturer’s protocol. The quality was assessed post-library preparation using the Agilent Bioanalyzer and quantified using a qPCR-based method with the KAPA Library Quantification Kit (P/N: KK4873) and the QuantStudio 12K instrument.

Prepared library pools were pooled in equimolar ratios, and the resulting pool was subjected to cluster generation using the NovaSeq 6000 System, following the manufacturer’s protocols. 150 bp paired-end sequencing was performed on the NovaSeq 6000 platform targeting 400M reads per sample. Raw sequencing data (FASTQ files) obtained from the NovaSeq 6000 was subjected to quality control analysis, including read quality assessment. Real-Time Analysis Software (RTA) and NovaSeq Control Software (NCS) (v1.8.0; Illumina) were used for base calling. MultiQC (v1.7; Illumina) was used for data quality assessments.

#### Tank’s RNA Sequencing

A Quality Control analysis was performed on the RNA sample using the Agilent Bioanalyzer to assess quality and an RNA Qubit assay was used to determine quantity. Poly(A) RNA enrichment was conducted using NEBNext Poly(A) mRNA Magnetic Isolation Module (NEB), and the sequencing library was constructed by using the NEBNext Ultra II RNA Library Prep Kit (NEB) (P/N: E7765L, NEB, Ipswich, MA) following the manufacturer’s instructions. End repair, A-tailing, and adapter ligation were performed to generate the final cDNA library. The library quality was assessed using a Bioanalyzer and quantified using a qPCR-based method with the KAPA Library Quantification Kit (P/N: KK4873) and the QuantStudio 12K instrument.

Prepared libraries were pooled in equimolar ratios, and the resulting pool was subjected to cluster generation using the NovaSeq 6000 System, following the manufacturer’s protocols. 150 bp paired-end sequencing was performed on the NovaSeq 6000 platform targeting 50M reads per sample. Raw sequencing data (FASTQ files) obtained from the NovaSeq 6000 was subjected to quality control analysis, including read quality assessment. Real-Time Analysis Software (RTA) and NovaSeq Control Software (NCS) (1.8.0; Illumina) were used for base calling. MultiQC (v1.7; Illumina) was used for data quality assessments.

#### PacBio Libraries and Sequencing of Tank

HMW gDNA (8µg) was sheared on a Megaruptor 3 (Diagenode, NJ). Sheared DNAs were size selected for fragments 10kb to 20kb in a BluePippin (Sage Science, MA) using a 0.75% agarose gel Cassette. The PacBio HiFi library was prepared from 2µg of size-selected DNA with the SMRTBell Express Template Prep kit 2.0. Sequencing was performed on 3 SMRT cells on a Pacbio Sequel IIe (Pacific Biosciences, CA) using 30hs movie times. The circular consensus bam and fastq.gz files were generated on an instrument with the software SMRT Link V10.2 from Pacific Biosciences, which generates consensus sequences from a minimum of 3 passes around the template with a minimum predicted accuracy of 99%.

#### Omni-C Libraries and Sequencing of Tank

Proximity ligation Omni-C libraries were constructed with the Omni-C Library Preparation kit from Dovetail Genomics, CA. Ten microliters of blood were used to prepare two libraries. The libraries were constructed exactly as described in the manufacturer’s protocol. The individually barcoded libraries were quantitated with Qubit, run on a Fragment Analyzer (Agilent, CA) to determine library fragment lengths, and further quantitated by qPCR on a CFX Connect Real-Time qPCR system (Bio-Rad, CA) for accurate pooling of barcoded libraries and maximization of number of clusters in the flowcell. The libraries were sequenced on an SP lane with 2×150nt reads in a NovaSeq 6000 (Illumina, CA). FASTQ files were generated and demultiplexed with the bcl2fastq v2.20 Conversion Software (Illumina).

#### Genome assembly and annotation

PacBio HiFi sequencing yielded a total of 108 gigabases in approximately 6.8 million reads with the longest read reaching 64,815 bp. Illumina sequencing of the Omni-C reads resulted in a total of one billion reads in pairs of 150 nt. Hifi sequences were assembled using Hifiasm (version 0.16.1 r375) with default parameters^77^. This created an assembly of length 2.4 Gb in 193 contigs. The assembly has an N50 of 113 Mb and the largest contig is 377 Mb. The Omni-C reads were used to scaffold the assembly using SALSA v2.3^78^. SALSA was run with three iterations, enzyme set to DNASE, and “attempt to fix mis-assemblies” set to “yes.”

#### Assembly Quality Assessment

BUSCO version 5.3.2 (Benchmarking Universal Single-Copy Orthologs) was used to measure the quality of the assembly using the sauropsida dataset from OrthoDB (version 10) which contained 7480 orthologs^79,80^. Of those orthologs, 97.4% were complete, 1.4% were complete but duplicated, 0.4% were fragmented, and 2.2% were missing (**Fig. 2a**).

#### Jonathan’s Variation and Consensus Creation

The Illumina reads from Jonathan were aligned against the Tank reference assembly using BWA-MEM^81^. The alignment rate was 92.54%. Following GATK v4.2.6.1 best practices for non-model organism variant calling, PCR and optical duplicates were marked using the “MarkDuplicates” command in GATK^82^. Next variants were called using “HaplotypeCaller” and genotyped using “GenotypeGVCFs.” This resulted in 7,678,401 variants which were then filtered using “VariantFiltration” leaving 6,911,581 high-quality variants. These variants were then used to create a consensus genome using GATK’s “FastaAlternateReferenceMaker.”

#### Population Genetic Structure Analysis

We performed the population genetic structure analysis as described in Çilinger, *et al*^7^. Briefly, we subsampled Illumina reads from Tank and Jonathan to match the sequencing level of the previously published data. We aligned those reads against the “AldGig_1.0.” assembly using BWA using BWA-MEM version 0.7.17. We filtered the alignments for quality greater or equal to 20. Next, we used ANGSD version 0.941-24 to create a beagle file containing genotype likelihoods (baq 1, skipTriallelic 1 minQ 20 minInd 16 doMajorMinor 1 doMaf 1 GL 2 only_proper_pairs 1 uniqueOnly doGlf 2 SNP_pval 0.001)^83^. PCANGSD version 1.2 was used to generate PCA and admixture analysis at various eigenvalues^84^. The results were plotted using ggplot2.

#### Gene prediction for Tank and Jonathan

First, a custom repeat model was created for Tank’s genome using Dfam TE Tools (https://github.com/Dfam-consortium/TETools) using RepeatModeler (version 2.0.3)^85^. Next, both genomes were soft-masked using RepeatMasker (version 4.1.2-p1)^86^. RNA-Seq from Lonesome George a member of *Chelonoidis niger abingdonii*,*Gopherus evgoodei, and Tank* was aligned against the reference genomes using STAR^7,87,88^. The RNA-Seq for Lonesome George mapped at a rate of 81.57% for Tank and 81.71% for Jonathan. The mapping rates for *G. evgoodei* were 63.44% for Tank and 63.45% for Jonathan. The RNA-Seq alignments were used by BRAKER to generate a file of evidence^89^. Next, GenomeThreader via BRAKER was used to generate protein evidence from both Lonesome George and *G. evgoodei*^90^. The protein and RNA-Seq evidence were then used as input into Augustus, along with the ‘chicken’ model’, to generate gene predictions^91^.

#### Genome annotation and whole-genome alignment

The ERGO Annotation Pipeline was used to annotate gene functions using other Eukaryotic genomes (*Homo sapiens*, *Mus musculus*, *Chelonoidis abingodonii,* and others) as references^92,93^.

#### Identification of Positively Selected Genes

First, the longest transcript from each gene of the *H. sapiens* genome. Protein similarity between all coding sequences between all genomes was computed using DIAMOND (version 2.0.8) using the “--ultra-sensitive” option^94^. Next, using custom scripts coding sequences with corresponding, bidirectional best hits were selected. Incomplete transcripts or transcripts that included embedded stops were removed. The resulting groups of sequences were aligned with PRANK (v170427) using the codon model^95^. The alignments were then analyzed using codeml from the PAML package (version 4.10.6) using the nested models of M1 and M2a^96^. The M1 model tests for neutral evolution, while the M2A model tests for positive selection of the foreground branch. The foreground branch was set as Jonathan, the background contained *H. sapiens*, *H. glaber*, *A. carolinensis*, *P. sinensis*, *M. musculus*, *N. furzeri*, *D. rerio*, *C. picta bellii*, *G. agassizii*, and *B. mysticetus*. Next, data was imported into R (version 3.6.3)^97^. The likelihood ratio test was performed between the nested models with two degrees of freedom with the resulting values converted to p-values using R’s pchisq function. PSGs were tested for over-representation in GeneAge and LongevityMap using Fisher’s exact test (fisher.test) in R. Results were considered significant if P < 0.05.

#### Gene Ontology Term Enrichment Analysis of Positively Selected Genes

GO term enrichment analysis was performed using topGO with genes with a ω > 1 for each genome^98^. For each set, three statistical tests were run: Fisher’s exact test with the classic algorithm, Kolmogorov-Smirnov test with a classic algorithm, and Kolmogorv-Smirnov with the elimination algorithm using the complete human gene set as the background. The resulting tables were then ordered by the Kolmogorv-Smirnov test with the elimination algorithm p scores.

### DNA methylation

#### Differentially Methylated Regions

Generated hypomethylated regions using DNM tools ‘hmr’ command. Differentially methylated regions (DMRs) were calculated using DNM tools ‘dmr’ function.

Briefly, a proportion table was generated as described in the DNM tools documentation for DM detection using the ‘radmeth’ command^99^. Next, we created a design matrix which was used along with the proportion table to create a bed file of methylated read counts. Next, ‘radadjust’ was used to adjust the p-values of individual CpGs, then ‘radmerge’ was run with a p-value of 0.01 which joins neighboring differentially methylated sites with a p-value below 0.01. Custom scripts were used to detect if a DMR overlapped a gene or promoter region.

#### Methylation Entropy

Total genome methylation entropy was computed using Metheor’s ‘me’ command^100^. Low Entropy Regions (LERs) were identified by computing the mean of the entropy for every gene ‘region’ (between gene start and gene stop) for Jonathan and the hatchling. The region was considered an LER if the region had a non-zero mean within 0.005 of the hatchling’s mean for that region.

#### AlphaFold 2 structure prediction

AlphaFold structure prediction was carried out with the Phenix *PredictModel* tool which uses AlphaFold 2, and employs the mmseqs2 MSA server and scripts derived from ColabFold^48,101–104^. The *PredictModel* tool uses AlphaFold2 directly for sequences of up to 1500 residues. Longer sequences are split into overlapping chunks of length 600 with an overlap of 200. These chunks are run individually and then reassembled at the end. To improve the contacts between domains in different chunks, a contact map is created and domains that are in contact but in different chunks are re-predicted together. During assembly, the domain pairs are used to adjust the positions of domains, and also positions of domains are adjusted to minimize overlaps. The repositioned domains are then used as a template to morph the fully reassembled prediction. The result of this procedure is a full-length prediction from overlapping chunk predictions, with distortions (largely in low-confidence regions) that allow the assembled prediction to minimize overlaps and make domain-domain contacts predicted to be present between chunks.

#### Network analysis

The FASTA file with all protein sequences from Jonathan was uploaded to the STRING web interface as a custom proteome^34^. We subsequently retrieved the functional association network for the PSGs at a 0.4 confidence cutoff using Cytoscape stringApp and clustered it using the clusterMaker2 implementation of the MCL algorithm^105,106^. We finally highlighted genes belonging to the set of conserved PSGs, GeneAge, or LongevityMap using Omics Visualizer, and manually annotated selected clusters with descriptive labels^107^.

#### Structural analyses

The AlphaFold 2-predicted models were processed with the Phenix Process-predicted-model tool to remove the low-confidence regions (LDDT <0.7). Pairs of the processed models of the Lonesome George and Jonathan proteins are superposed and analyzed with the ProSMART program to identify backbone and side-chain differences between the two protein pairs^108^. Residues with their main-chain or side-chain deviations greater than 3 standard deviations of all the corresponding values of the analyzed residues are flagged for visual inspection, especially for those corresponding to key mutations of proteins related to Jonathan’s extended lifespan.

#### Gene family expansion

To detect gene family expansion, we created protein clusters utilizing protein similarities computed by DIAMOND as previously described using ‘grouping.py’ and ‘gen_table.py’ available on GitHub^94^. The clusters were created with a protein coverage cutoff of 20%, a match length cutoff of 20%, expected value threshold of 1E-100, with at least 50% identity, ensuring that proteins in the cluster were not coded by DNA sequences whose genomic locations overlapped. These clusters were then reduced to a table of non-overlapping genes indicating possible duplication. A specific subset of candidate gene family expansions of interest was then manually curated.

#### Reporting Summary

Further information on research design is available in the Nature Research Reporting Summary linked to this article.

## Data availability

The raw sequence is available at BioProject PRJNA1208048. Predicted protein sequences, Alphafold structures, Structural Analysis, and secondary Methylation data are available from https://github.com/bvaisvil/jg.

## Code availability

The code used during the analysis is available from https://github.com/bvaisvil/jg.

### Export Permit

#### EMDEP020/17

We received an export permit for Jonathan’s samples from the St. Helena government (Department of Environment & Natural Resources Directorate) on 4/8/2017.

### Institutional Animal Care and Use Committee (IACUC) Approval

We received approval for buccal scrap collections from Aldabras from the Vanderbilt University Medical Center and Vanderbilt University Institutional Animal Care and Use Committee (IACUC).

## Acknowledgments

We thank Terri Voland for her support throughout this project. We thank Professor Geoffrey Batchen (University of Oxford), for his evaluation of the Jonathan photograph. We thank Professor Lawrence Steinman (Stanford University) and Dr. Anamitra Bhattacharyya (AbbVie) for reviewing the manuscript. We are grateful to Rosemary Rees for providing her copy of Jonathan’s photograph and for her detailed family history. We appreciate the guidance from Rebecca Cairns-Wicks, Sean Burns, and the entire St. Helena Research Institute and the Joint Nature Conservation Committee during this project. We thank the people of St. Helena for the care they have given Jonathan over all these years and for allowing us to learn about his genome. We thank Dr. Stefan Green (Rush University and Kevin Kunstman (Rush University) for their assistance with HMW gDNA extraction. We thank, Ross Swofford (University of Massachusetts-Chan Medical School) for help with DNA extraction. We thank the men and women of the United States Space Force at the Patrick Space Force Base for transporting Jonathan’s DNA samples from St. Helena to Brevard County, Florida, USA. We thank Elinor Karlsson’s lab (Broad Institute), especially Diane Genereux for helping with the buccal scrapes, Jeremy Johnson for helping with shipments in the USA, and Dr. Karlsson for early suggestions. We thank the Ayuu Clinic, Chicago for providing blood vacutainers for sample collection. Lastly, we thank Alicia Clark for the carapace and scute drawings.

## Author contributions

S.W.C., B.V., D.P. S., A.J., V.K., Y.M., C.F., and S.H., conceived experiments and evaluated results. J.H., J.M.F., and M.L.T acquired Jonathan samples. S.P., and M.L.T., acquired hatchling and Tank samples. P.N., K.W., and B.R. evaluated the unique variants with protein language models. J.G. helped with the evaluation of previous Aldabra growth data. T.C.T and L-W.H., generated Alphafold predictions. L.J.J. analyzed data with the STRING database. S.W.C. supervised all aspects of study. S.W.C wrote the original manuscript and M.C. helped in figure production.

## Data and materials availability

The raw sequence is available at BioProject PRJNA1208048. Predicted protein sequences, Alphafold structures, Structural Analysis, and secondary Methylation data are available from https://github.com/bvaisvil/jg. The code used during the analysis is available from https://github.com/bvaisvil/jg.

## References

1. Kim, E. B. et al. Genome sequencing reveals insights into physiology and longevity of the naked mole rat. Nature 479, 223–227 (2011).

2. Keane, M. et al. Insights into the evolution of longevity from the bowhead whale genome. Cell Rep. 10, 112–122 (2015).

3. Seim, I. et al. Genome analysis reveals insights into physiology and longevity of the Brandt’s bat Myotis brandtii. Nat. Commun. 4, 2212 (2013).

4. Valenzano, D. R. et al. The African Turquoise Killifish Genome Provides Insights into Evolution and Genetic Architecture of Lifespan. Cell 163, 1539–1554 (2015).

5. Pascual-Torner, M. et al. Comparative genomics of mortal and immortal cnidarians unveils novel keys behind rejuvenation. Proc. Natl. Acad. Sci. U. S. A. 119, e2118763119 (2022).

6. Quesada, V. et al. Giant tortoise genomes provide insights into longevity and age-related disease. *Nat*. Ecol. Evol. 3, 87–95 (2019).

7. Çilingir, F. G. et al. Chromosome-level genome assembly for the Aldabra giant tortoise enables insights into the genetic health of a threatened population. GigaScience 11, giac090 (2022).

8. da Silva, R., Conde, D. A., Baudisch, A. & Colchero, F. Slow and negligible senescence among testudines challenges evolutionary theories of senescence. Science 376, 1466–1470 (2022).

9. Grubb, P. The growth, ecology and population structure of giant tortoises on aldabra. Phil Trans R Soc Lond B 260, 327–372 (1971).

10. Bourn, D & Coe, M. Features of tortoise mortality and decomposition on aldabra. Phil. Trans. R. Soc. Lond. B 286, 189–193 (1979).

11. Seale, K., Horvath, S., Teschendorff, A., Eynon, N. & Voisin, S. Making sense of the ageing methylome. Nat. Rev. Genet. 23, 585–605 (2022).

12. Crofts, S. J. C., Latorre-Crespo, E. & Chandra, T. DNA methylation rates scale with maximum lifespan across mammals. *Nat*. Aging 4, 27–32 (2024).

13. Haghani, A. et al. DNA methylation networks underlying mammalian traits. Science 381, eabq5693 (2023).

14. Li, C. Z. et al. Epigenetic predictors of species maximum life span and other life-history traits in mammals. Sci. Adv. 10, eadm7273 (2024).

15. Horvath, S., Zhang, J., Haghani, A., Lu, A. T. & Fei, Z. Fundamental equations linking methylation dynamics to maximum lifespan in mammals. Nat. Commun. 15, 8093 (2024).

16. Lowe, R. et al. DNA methylation clocks as a predictor for ageing and age estimation in naked mole-rats, Heterocephalus glaber. Aging 12, 4394–4406 (2020).

17. Kerepesi, C. et al. Epigenetic aging of the demographically non-aging naked mole-rat. Nat. Commun. 13, 355 (2022).

18. Horvath, S. et al. DNA methylation clocks tick in naked mole rats but queens age more slowly than nonbreeders. *Nat*. Aging 2, 46–59 (2022).

19. Lu, A. T. et al. Universal DNA methylation age across mammalian tissues. *Nat*. Aging 3, 1144–1166 (2023).

20. Kehlmaier, C. et al. Ancient DNA elucidates the lost world of western Indian Ocean giant tortoises and reveals a new extinct species from Madagascar. Sci. Adv. 9, eabq2574 (2023).

21. Turnbull, L. A. et al. Persistence of distinctive morphotypes in the native range of the CITES-listed Aldabra giant tortoise. Ecol. Evol. 5, 5499–5508 (2015).

22. Bourn, D. The size, structure and distribution of giant tortoise population of Aldabra. Philos Trans R Soc Lond B Biol Sci 282, 139–175 (1978).

23. Çilingir, F. G. et al. Low-coverage reduced representation sequencing reveals subtle within-island genetic structure in Aldabra giant tortoises. Ecol. Evol. 12, e8739 (2022).

24. Cheke, AS & Bourn, R. Unequal struggle - how humans displaced the domiance of tortoises in island ecosystems. in Western Indian Ocean Tortoises 31–120 (Siri Scientific, 2014).

25. Kolora, S. R. R. et al. Origins and evolution of extreme life span in Pacific Ocean rockfishes. Science 374, 842–847 (2021).

26. Stelzer, G. et al. The GeneCards Suite: From Gene Data Mining to Disease Genome Sequence Analyses. Curr. Protoc. Bioinforma. 54, 1.30.1–1.30.33 (2016).

27. Uggenti, C. et al. cGAS-mediated induction of type I interferon due to inborn errors of histone pre-mRNA processing. Nat. Genet. 52, 1364–1372 (2020).

28. Naesens, L. et al. Mutations in RNU7-1 Weaken Secondary RNA Structure, Induce MCP-1 and CXCL10 in CSF, and Result in Aicardi-Goutières Syndrome with Severe End-Organ Involvement. J. Clin. Immunol. 42, 962–974 (2022).

29. Lessel, D. et al. Atypical Aicardi-Goutieres syndrome: is the WRN locus a modifier? Am. J. Med. Genet. A. 164A, 2510–2513 (2014).

30. Seluanov, A. et al. Hypersensitivity to contact inhibition provides a clue to cancer resistance of naked mole-rat. Proc. Natl. Acad. Sci. U. S. A. 106, 19352–19357 (2009).

31. Tian, X. et al. High-molecular-mass hyaluronan mediates the cancer resistance of the naked mole rat. Nature 499, 346–349 (2013).

32. Zhang, Z. et al. Increased hyaluronan by naked mole-rat HAS2 improves healthspan in mice. Nature 621, 196 (2023).

33. Braidy, N. et al. Age related changes in NAD+ metabolism oxidative stress and Sirt1 activity in wistar rats. PloS One 6, e19194 (2011).

34. Szklarczyk, D. et al. The STRING database in 2023: protein-protein association networks and functional enrichment analyses for any sequenced genome of interest. Nucleic Acids Res. 51, D638–D646 (2023).

35. Campolo, M. et al. Echocardiographic evaluation of four giant Aldabra tortoises (Aldabrachelys gigantea). Vet. Rec. Open 6, e000274 (2019).

36. Can, E. et al. Naked mole-rats maintain cardiac function and body composition well into their fourth decade of life. GeroScience 44, 731–746 (2022).

37. Grimes, K. M. et al. The naked mole-rat exhibits an unusual cardiac myofilament protein profile providing new insights into heart function of this naturally subterranean rodent. Pflugers Arch. 469, 1603–1613 (2017).

38. Notin, P., et al. Tranception: protein fitness prediction with autoregressive transformers and inference-time retrieval. arxiv (2022) doi:10.48550/arXiv.2205.13760.

39. Meier, J. et al. Language models enable zero-shot prediction of the effects of mutations on protein function. bioRxiv 2021.07.09.450648 (2021) doi:10.1101/2021.07.09.450648.

40. Nijkamp, E., Ruffolo, J. A., Weinstein, E. N., Naik, N. & Madani, A. ProGen2: Exploring the boundaries of protein language models. Cell Syst. 14, 968–978.e3 (2023).

41. Hesslow, D., Zanichelli, N., Notin, P., Poli, I. & Marks, D. RITA: a Study on Scaling Up Generative Protein Sequence Models. arXiv (2022) doi:10.48550/arXiv.2205.05789.

42. Notin, P. et al. ProteinGym: Large-Scale Benchmarks for Protein Design and Fitness Prediction. BioRxiv Prepr. Serv. Biol. 2023.12.07.570727 (2023) doi:10.1101/2023.12.07.570727.

43. Marquet, C. et al. Embeddings from protein language models predict conservation and variant effects. Hum. Genet. 141, 1629–1647 (2022).

44. Tacutu, R. et al. Human Ageing Genomic Resources: new and updated databases. Nucleic Acids Res. 46, D1083–D1090 (2018).

45. Rice, C. et al. Structural and functional analysis of the human POT1-TPP1 telomeric complex. Nat. Commun. 8, 14928 (2017).

46. Bettegowda, C. et al. Mutations in CIC and FUBP1 contribute to human oligodendroglioma. Science 333, 1453–1455 (2011).

47. Torigoe, T. H. et al. Novel protective effect of the FOXO3 longevity genotype on mechanisms of cellular aging in Okinawans. Npj Aging 10, 18 (2024).

48. Jumper, J. et al. Highly accurate protein structure prediction with AlphaFold. Nature 596, 583–589 (2021).

49. UniProt Consortium. UniProt: the Universal Protein Knowledgebase in 2023. Nucleic Acids Res. 51, D523–D531 (2023).

50. Kircher, M. et al. A general framework for estimating the relative pathogenicity of human genetic variants. Nat. Genet. 46, 310–315 (2014).

51. Ng, P. C. & Henikoff, S. Predicting deleterious amino acid substitutions. Genome Res. 11, 863–874 (2001).

52. de Araujo, M. E. G. et al. Crystal structure of the human lysosomal mTORC1 scaffold complex and its impact on signaling. Science 358, 377–381 (2017).

53. Meerang, M. et al. The ubiquitin-selective segregase VCP/p97 orchestrates the response to DNA double-strand breaks. Nat. Cell Biol. 13, 1376–1382 (2011).

54. Ovadya, Y. et al. Impaired immune surveillance accelerates accumulation of senescent cells and aging. Nat. Commun. 9, 5435 (2018).

55. Jenkinson, G., Pujadas, E., Goutsias, J. & Feinberg, A. P. Potential energy landscapes identify the information-theoretic nature of the epigenome. Nat. Genet. 49, 719–729 (2017).

56. Landan, G. et al. Epigenetic polymorphism and the stochastic formation of differentially methylated regions in normal and cancerous tissues. Nat. Genet. 44, 1207–1214 (2012).

57. Hannum, G. et al. Genome-wide methylation profiles reveal quantitative views of human aging rates. Mol. Cell 49, 359–367 (2013).

58. Tsankov, A. M. et al. Loss of DNA methyltransferase activity in primed human ES cells triggers increased cell-cell variability and transcriptional repression. Dev. Camb. Engl. 146, dev174722 (2019).

59. Fang, Y. et al. DNA methylation entropy is associated with DNA sequence features and developmental epigenetic divergence. Nucleic Acids Res. 51, 2046–2065 (2023).

60. Wang, H.-T. et al. Methylation entropy landscape of Chinese long-lived individuals reveals lower epigenetic noise related to human healthy aging. Aging Cell 23, e14163 (2024).

61. Rafikova, E. et al. Open Genes-a new comprehensive database of human genes associated with aging and longevity. Nucleic Acids Res. 52, D950–D962 (2024).

62. Chawla, R. et al. Human UPF1 interacts with TPP1 and telomerase and sustains telomere leading-strand replication. EMBO J. 30, 4047–4058 (2011).

63. Heyn, H. et al. Distinct DNA methylomes of newborns and centenarians. Proc. Natl. Acad. Sci. U. S. A. 109, 10522–10527 (2012).

64. Teschendorff, A. E. et al. Age-dependent DNA methylation of genes that are suppressed in stem cells is a hallmark of cancer. Genome Res. 20, 440–446 (2010).

65. Day, K. et al. Differential DNA methylation with age displays both common and dynamic features across human tissues that are influenced by CpG landscape. Genome Biol. 14, R102 (2013).

66. Rakyan, V. K. et al. Human aging-associated DNA hypermethylation occurs preferentially at bivalent chromatin domains. Genome Res. 20, 434–439 (2010).

67. Bell, J. T. et al. DNA methylation patterns associate with genetic and gene expression variation in HapMap cell lines. Genome Biol. 12, R10 (2011).

68. López-Otín, C., Blasco, M. A., Partridge, L., Serrano, M. & Kroemer, G. The Hallmarks of Aging. Cell 153, 1194–1217 (2013).

69. López-Otín, C., Blasco, M. A., Partridge, L., Serrano, M. & Kroemer, G. Hallmarks of aging: An expanding universe. Cell 186, 243–278 (2023).

70. Barja, G., Cadenas, S., Rojas, C., Pérez-Campo, R. & López-Torres, M. Low mitochondrial free radical production per unit O2 consumption can explain the simultaneous presence of high longevity and high aerobic metabolic rate in birds. Free Radic. Res. 21, 317–327 (1994).

71. Gómez, J., Mota-Martorell, N., Jové, M., Pamplona, R. & Barja, G. Mitochondrial ROS production, oxidative stress and aging within and between species: Evidences and recent advances on this aging effector. Exp. Gerontol. 174, 112134 (2023).

72. Brunet-Rossinni, A. K. & Austad, S. N. Ageing studies on bats: a review. Biogerontology 5, 211–222 (2004).

73. Herrero, A. & Barja, G. Sites and mechanisms responsible for the low rate of free radical production of heart mitochondria in the long-lived pigeon. Mech. Ageing Dev. 98, 95–111 (1997).

74. Rodríguez-Nuevo, A. et al. Oocytes maintain ROS-free mitochondrial metabolism by suppressing complex I. Nature 607, 756–761 (2022).

75. Berry, B. J. & Kaeberlein, M. An energetics perspective on geroscience: mitochondrial protonmotive force and aging. GeroScience 43, 1591–1604 (2021).

76. Field, A. E. et al. DNA Methylation Clocks in Aging: Categories, Causes, and Consequences. Mol. Cell 71, 882–895 (2018).

77. Cheng, H., Concepcion, G. T., Feng, X., Zhang, H. & Li, H. Haplotype-resolved de novo assembly using phased assembly graphs with hifiasm. Nat. Methods 18, 170–175 (2021).

78. Ghurye, J., Pop, M., Koren, S., Bickhart, D. & Chin, C.-S. Scaffolding of long read assemblies using long range contact information. BMC Genomics 18, 527 (2017).

79. Manni, M., Berkeley, M. R., Seppey, M. & Zdobnov, E. M. BUSCO: Assessing Genomic Data Quality and Beyond. Curr. Protoc. 1, e323 (2021).

80. Zdobnov, E. M. et al. OrthoDB in 2020: evolutionary and functional annotations of orthologs. Nucleic Acids Res. 49, D389–D393 (2021).

81. Li, H. & Durbin, R. Fast and accurate short read alignment with Burrows-Wheeler transform. Bioinforma. Oxf. Engl. 25, 1754–1760 (2009).

82. Van der Auwera, G. A. et al. From FastQ data to high confidence variant calls: the Genome Analysis Toolkit best practices pipeline. Curr. Protoc. Bioinforma. 43, 11.10.1-11.10.33 (2013).

83. Korneliussen, T. S., Albrechtsen, A. & Nielsen, R. ANGSD: Analysis of Next Generation Sequencing Data. BMC Bioinformatics 15, 356 (2014).

84. Meisner, J. & Albrechtsen, A. Inferring Population Structure and Admixture Proportions in Low-Depth NGS Data. Genetics 210, 719–731 (2018).

85. Flynn, J. M. et al. RepeatModeler2 for automated genomic discovery of transposable element families. Proc. Natl. Acad. Sci. U. S. A. 117, 9451–9457 (2020).

86. Tarailo-Graovac, M. & Chen, N. Using RepeatMasker to identify repetitive elements in genomic sequences. Curr. Protoc. Bioinforma. Chapter 4, Unit 4.10 (2009).

87. Rhie, A. et al. Towards complete and error-free genome assemblies of all vertebrate species. Nature 592, 737–746 (2021).

88. Dobin, A. et al. STAR: ultrafast universal RNA-seq aligner. Bioinforma. Oxf. Engl. 29, 15–21 (2013).

89. Brůna, T., Hoff, K. J., Lomsadze, A., Stanke, M. & Borodovsky, M. BRAKER2: automatic eukaryotic genome annotation with GeneMark-EP+ and AUGUSTUS supported by a protein database. NAR Genomics Bioinforma. 3, lqaa108 (2021).

90. Gremme, G., Brendel, V., Sparks, M. & Kurtz, S. Engineering a software tool for gene structure prediction in higher organisms. Inf. Softw. Technol. 47, 965–978 (2008).

91. Stanke, M., Steinkamp, R., Waack, S. & Morgenstern, B. AUGUSTUS: a web server for gene finding in eukaryotes. Nucleic Acids Res. 32, W309–312 (2004).

92. Kapatral, V. et al. Genome sequence and analysis of the oral bacterium Fusobacterium nucleatum strain ATCC 25586. J. Bacteriol. 184, 2005–2018 (2002).

93. Overbeek, R. et al. The ERGO genome analysis and discovery system. Nucleic Acids Res. 31, 164–171 (2003).

94. Buchfink, B., Reuter, K. & Drost, H.-G. Sensitive protein alignments at tree-of-life scale using DIAMOND. Nat. Methods 18, 366–368 (2021).

95. Löytynoja, A. Phylogeny-aware alignment with PRANK. Methods Mol. Biol. Clifton NJ 1079, 155–170 (2014).

96. Yang, Z. PAML 4: phylogenetic analysis by maximum likelihood. Mol. Biol. Evol. 24, 1586–1591 (2007).

97. R Core Team. R: A language and environment for statistical computing. R Foundation for Statistical Computing (2020).

98. Alexa, A. & Rahnenfuhrer, J. topGO: Enrichment Analysis for Gene Ontology. (2022).

99. Dolzhenko, E. & Smith, A. D. Using beta-binomial regression for high-precision differential methylation analysis in multifactor whole-genome bisulfite sequencing experiments. BMC Bioinformatics 15, 215 (2014).

100. Lee, D., Koo, B., Yang, J. & Kim, S. Metheor: Ultrafast DNA methylation heterogeneity calculation from bisulfite read alignments. PLoS Comput. Biol. 19, e1010946 (2023).

101. Terwilliger, T. C. et al. AlphaFold predictions are valuable hypotheses and accelerate but do not replace experimental structure determination. Nat. Methods 21, 110–116 (2024).

102. Steinegger, M. & Söding, J. MMseqs2 enables sensitive protein sequence searching for the analysis of massive data sets. Nat. Biotechnol. 35, 1026–1028 (2017).

103. Mirdita, M., Steinegger, M. & Söding, J. MMseqs2 desktop and local web server app for fast, interactive sequence searches. Bioinforma. Oxf. Engl. 35, 2856–2858 (2019).

104. Mirdita, M. et al. ColabFold: making protein folding accessible to all. Nat. Methods 19, 679–682 (2022).

105. Doncheva, N. T. et al. Cytoscape stringApp 2.0: Analysis and Visualization of Heterogeneous Biological Networks. J. Proteome Res. 22, 637–646 (2023).

106. Utriainen, M. & Morris, J. H. clusterMaker2: a major update to clusterMaker, a multi-algorithm clustering app for Cytoscape. BMC Bioinformatics 24, 134 (2023).

107. Legeay, M., Doncheva, N. T., Morris, J. H. & Jensen, L. J. Visualize omics data on networks with Omics Visualizer, a Cytoscape App. F1000Research 9, 157 (2020).

108. Nicholls, R. A., Fischer, M., McNicholas, S. & Murshudov, G. N. Conformation-independent structural comparison of macromolecules with ProSMART. Acta Crystallogr. D Biol. Crystallogr. 70, 2487–2499 (2014).

