## Extended Data Figures for "Epigenomic insights into extreme longevity in the world’s oldest terrestrial animal, Jonathan"

### Slide 1
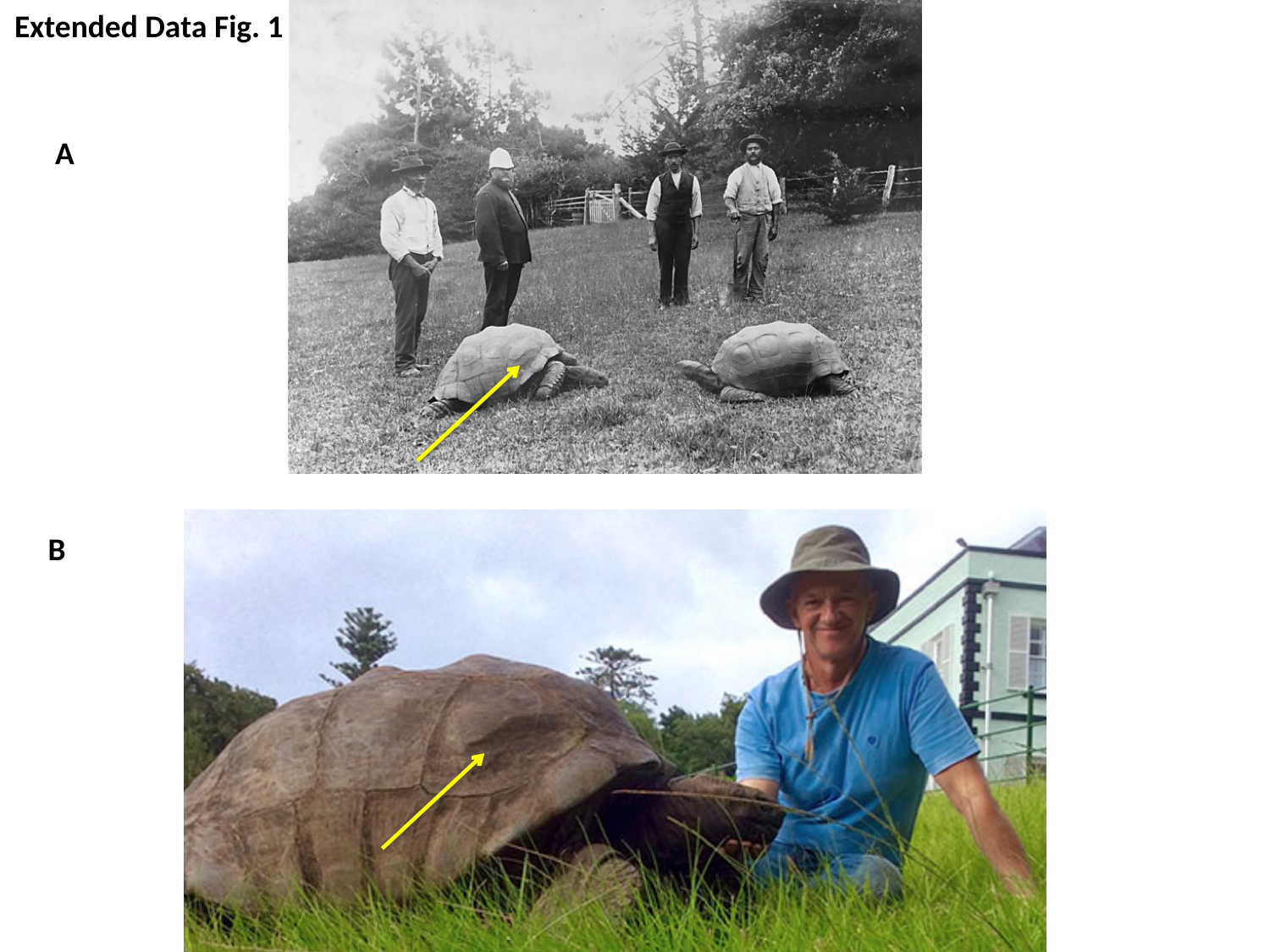

Extended Data Fig. 1
A
B

### Slide 2
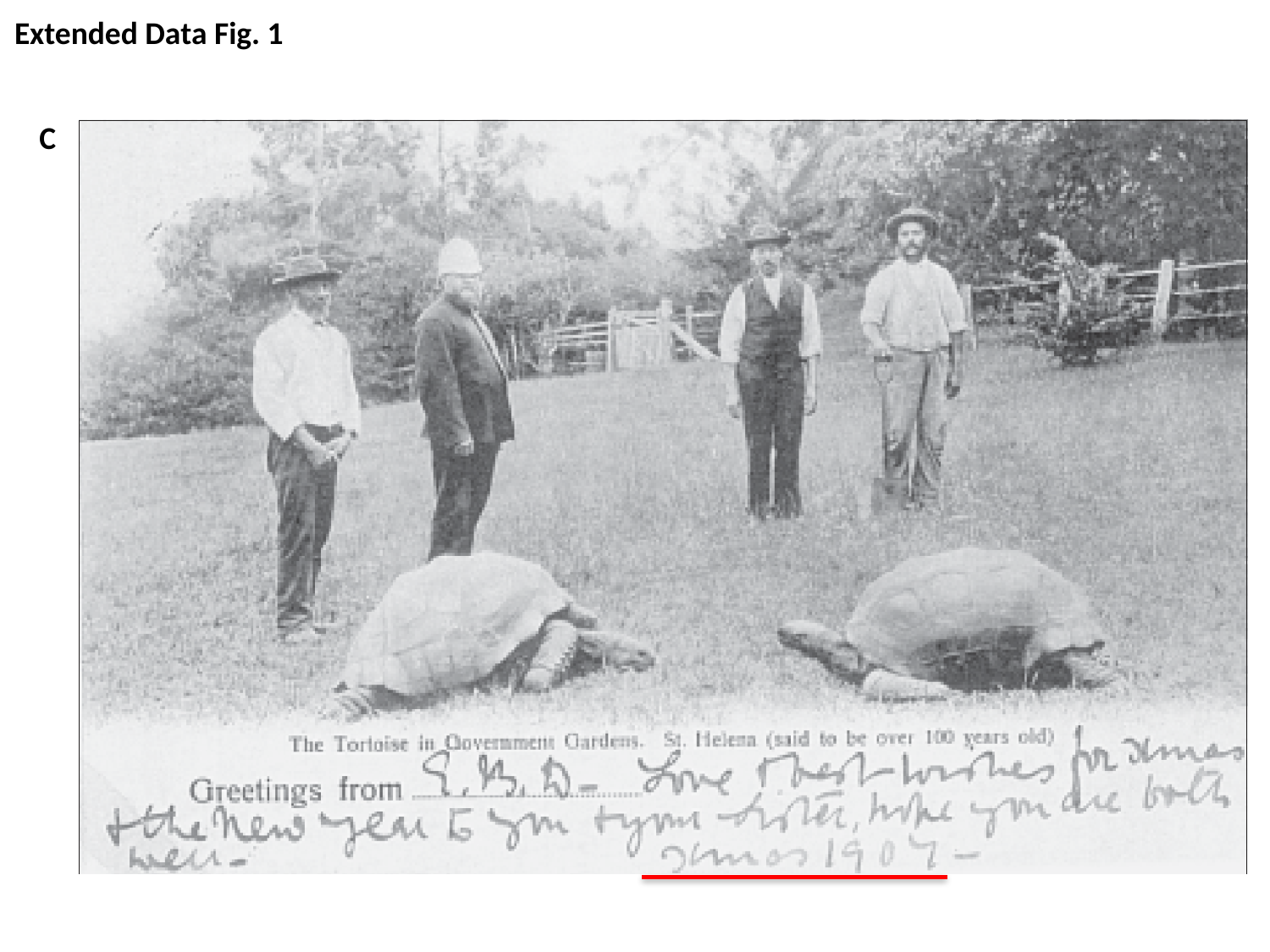

Extended Data Fig. 1
C

### Slide 3
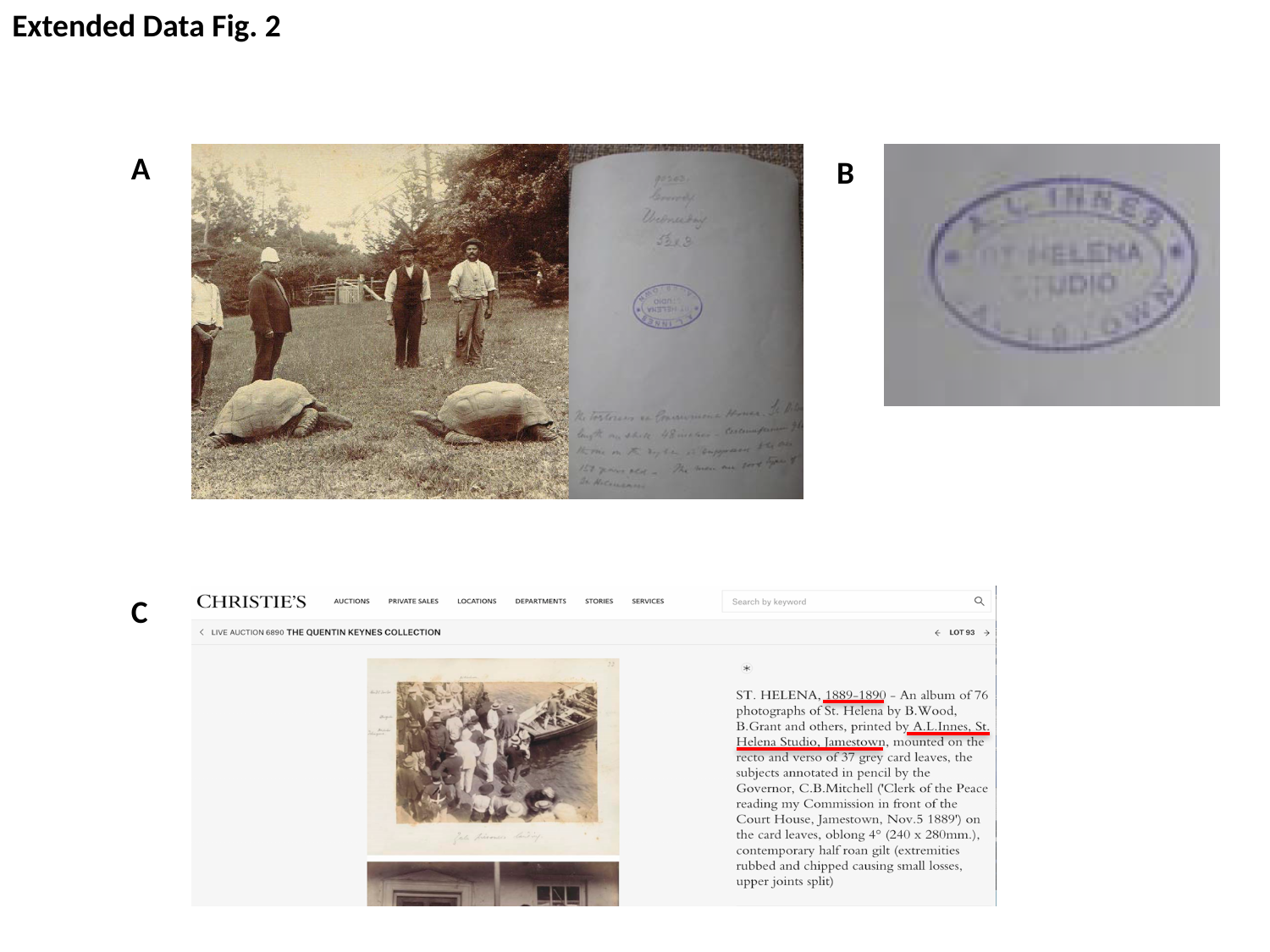

Extended Data Fig. 2
A
B
C

### Slide 4
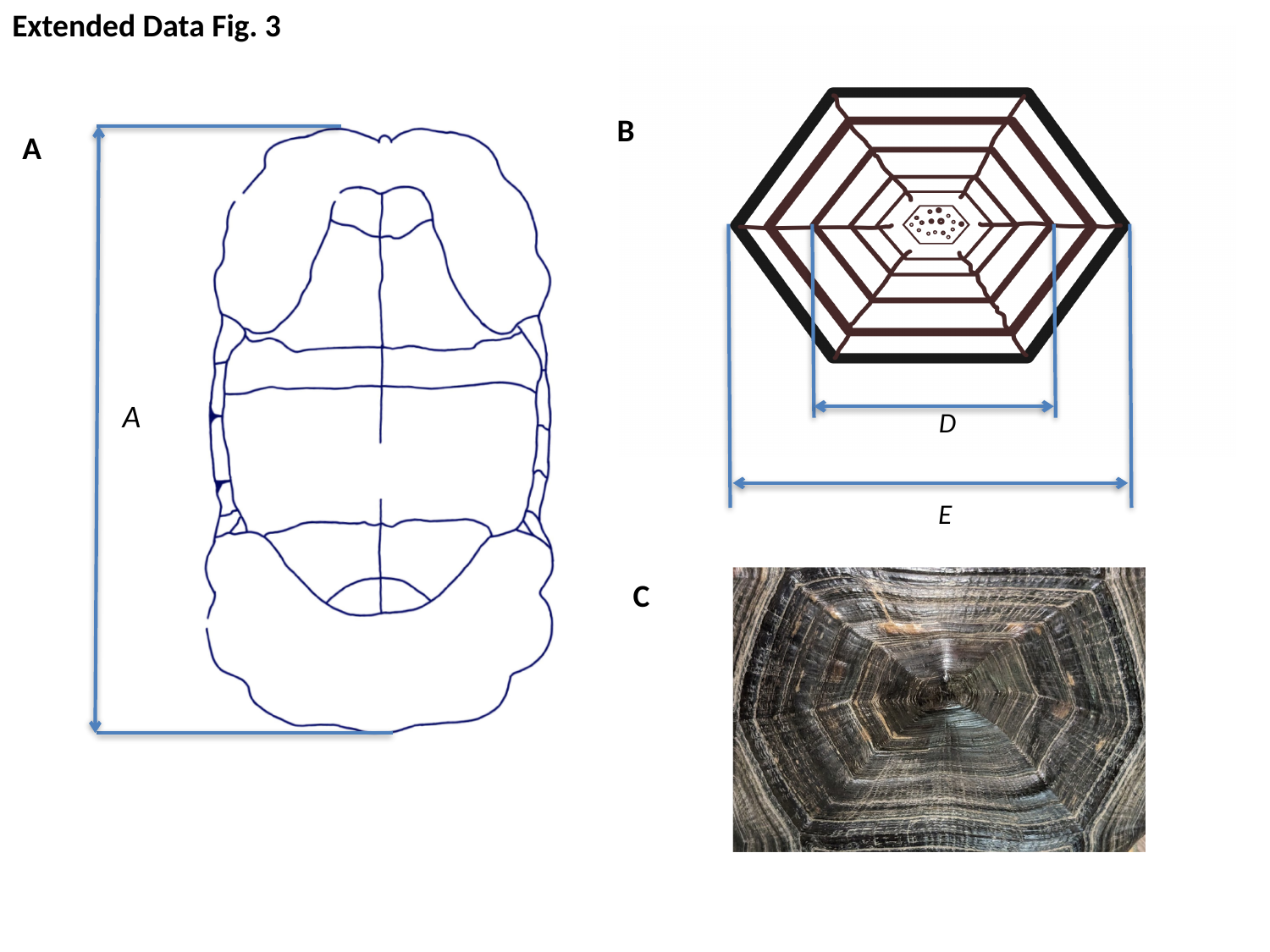

Extended Data Fig. 3
B
A
A
D
E
C

### Slide 5
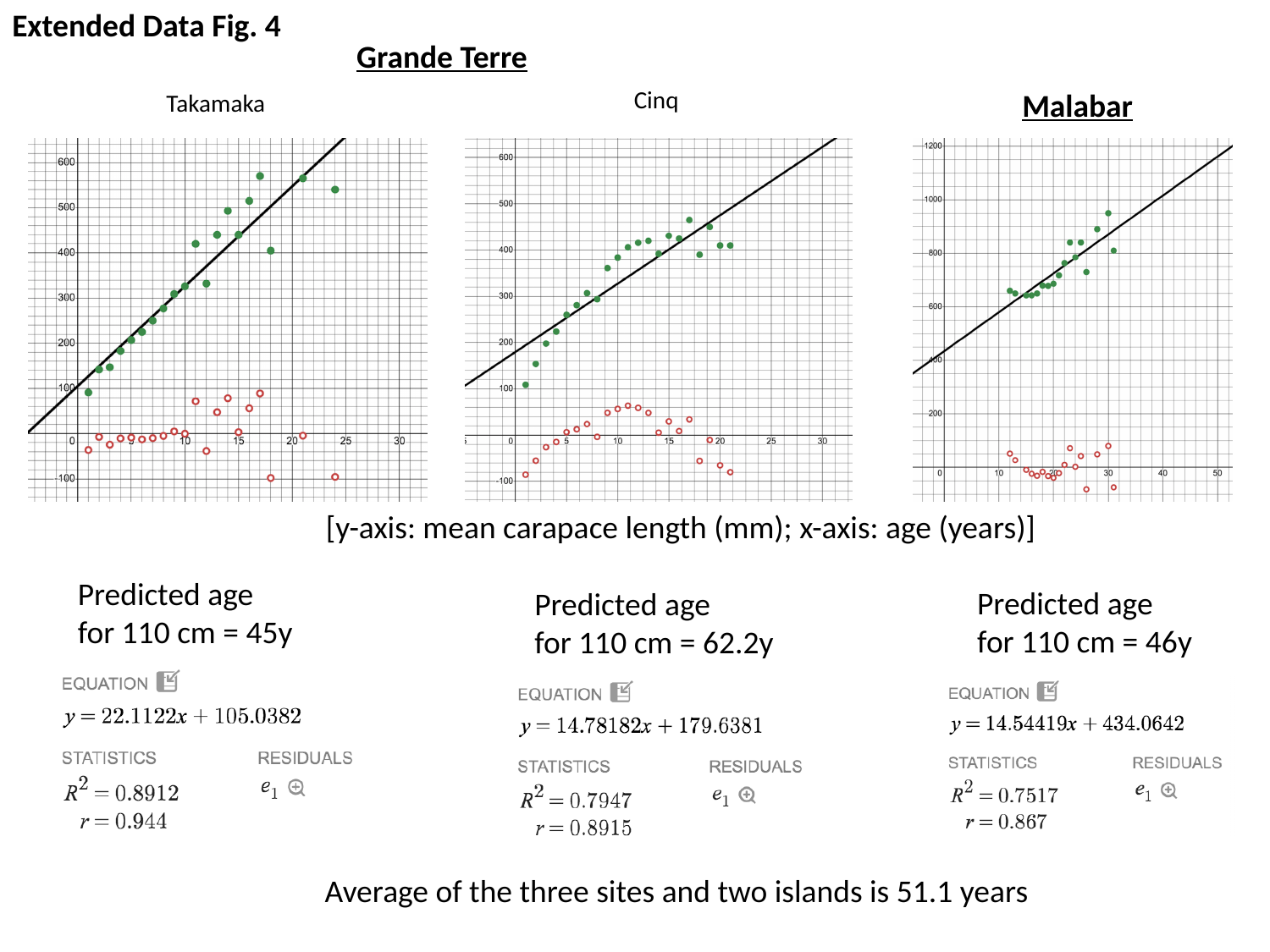

Extended Data Fig. 4
Grande Terre
Cinq
Malabar
Takamaka
[y-axis: mean carapace length (mm); x-axis: age (years)]
Predicted age
for 110 cm = 62.2y
 Predicted age
 for 110 cm = 45y
Predicted age
for 110 cm = 46y
Average of the three sites and two islands is 51.1 years

### Slide 6
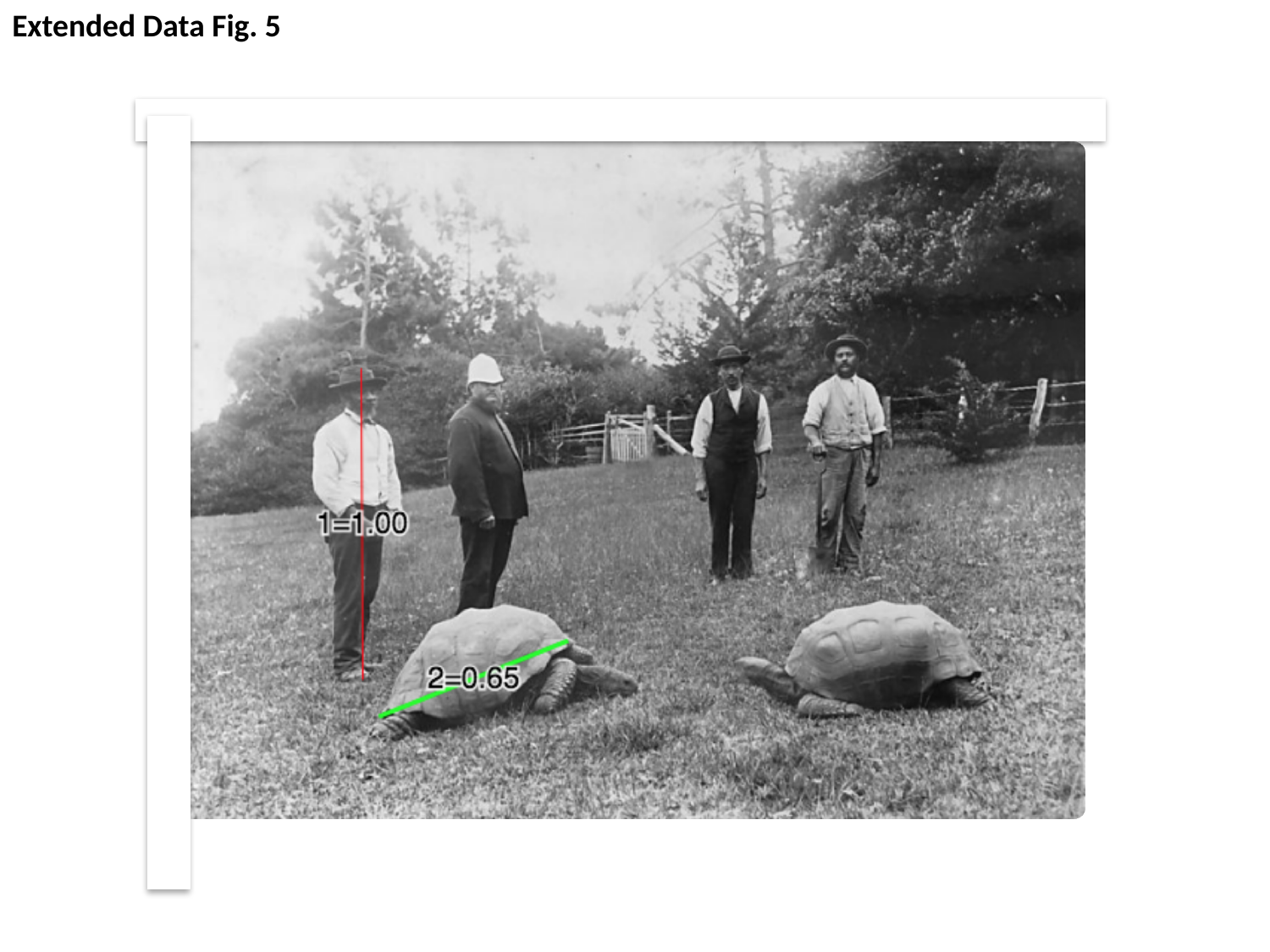

Extended Data Fig. 5

### Slide 7
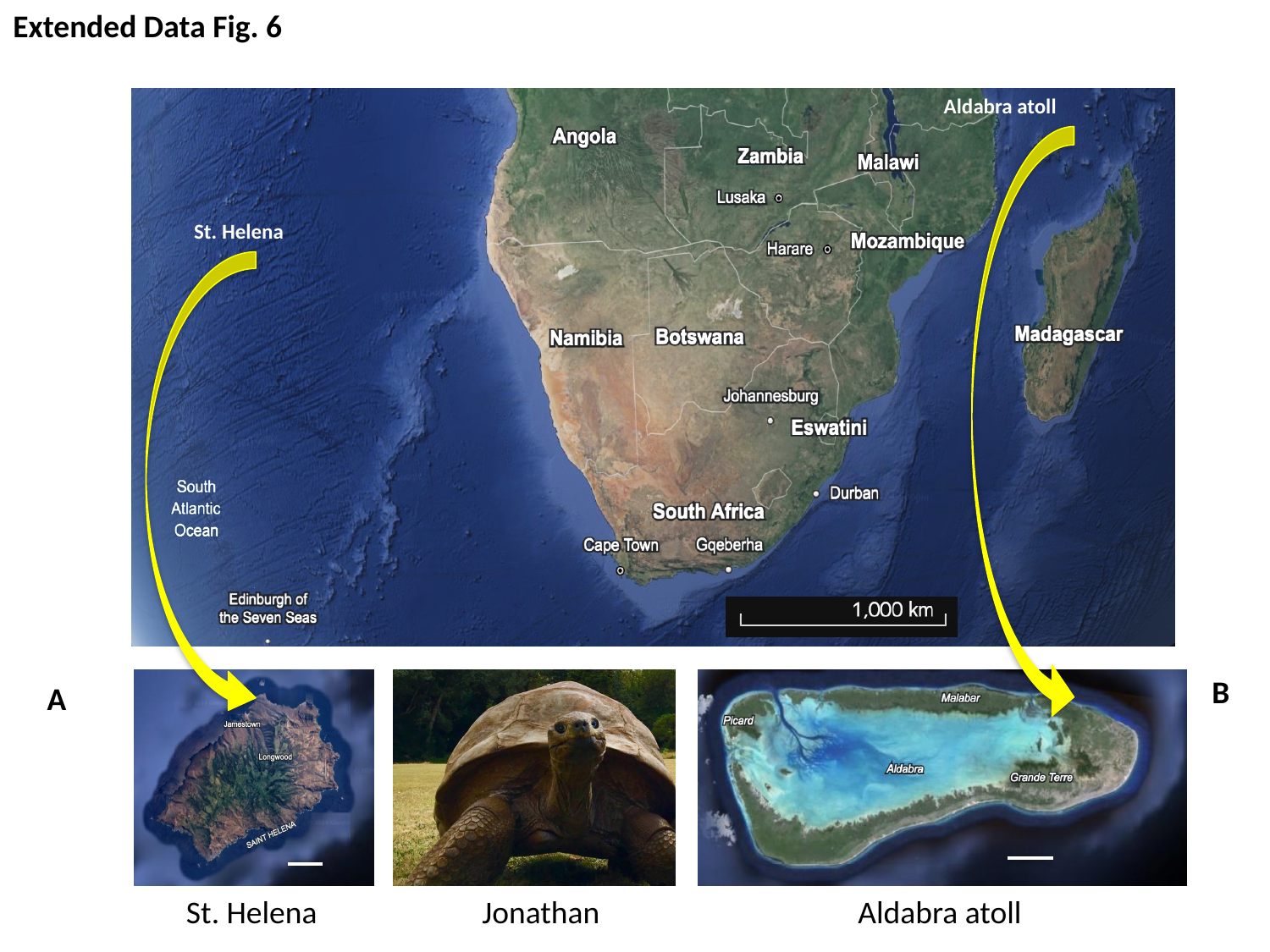

Extended Data Fig. 6
Aldabra atoll
St. Helena
B
A
St. Helena Jonathan Aldabra atoll

### Slide 8
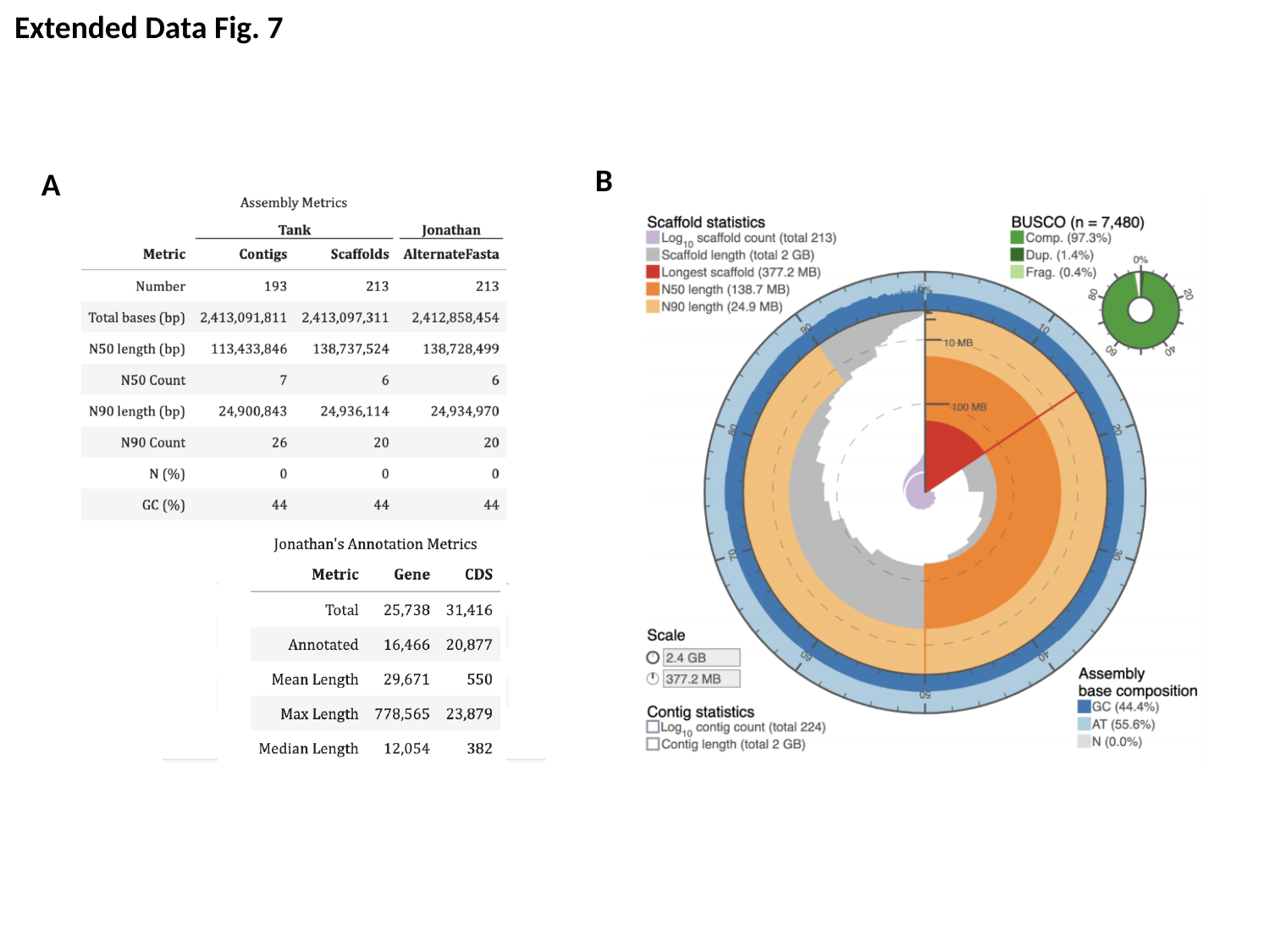

Extended Data Fig. 7
B
A

### Slide 9
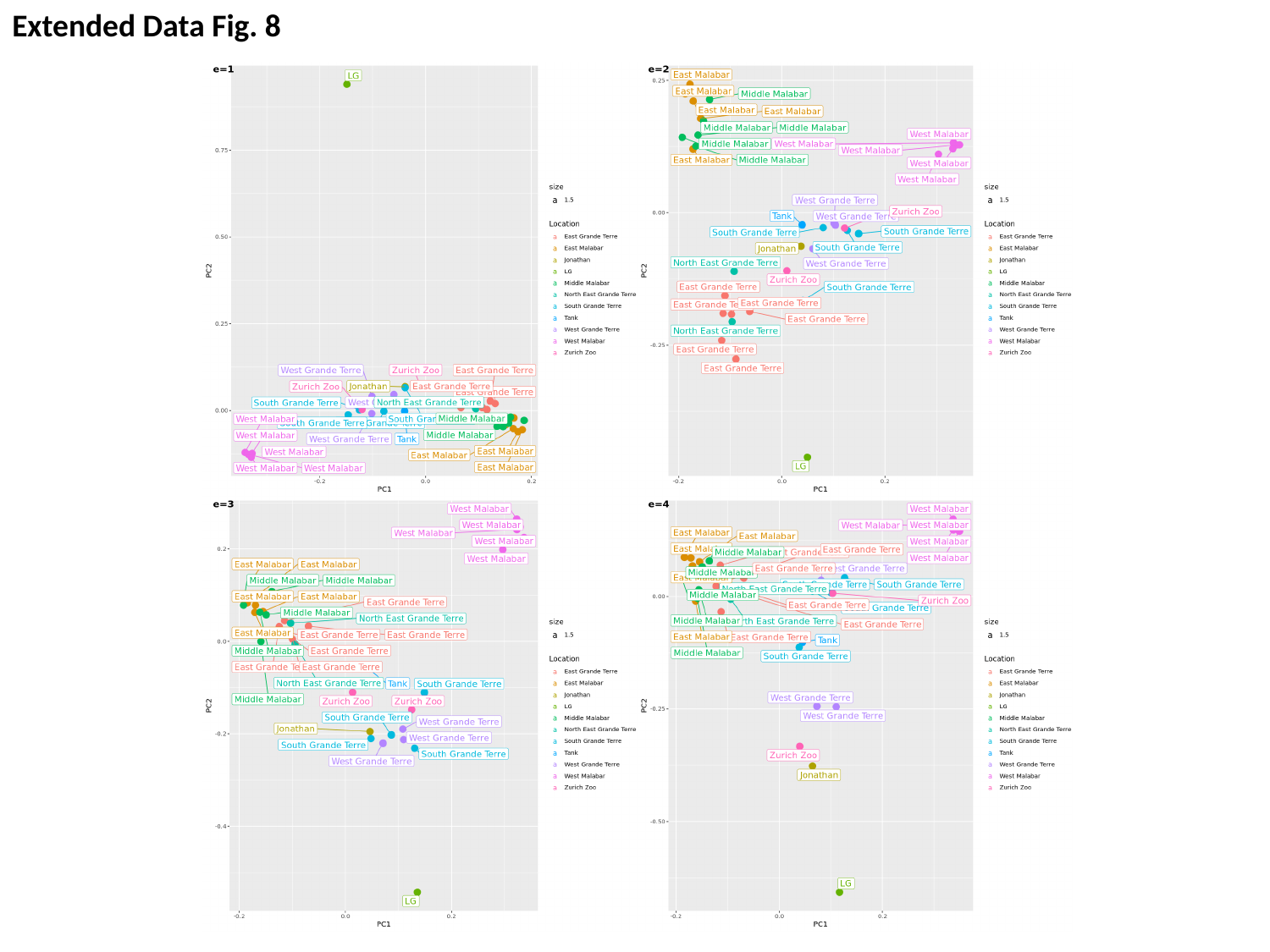

Extended Data Fig. 8

### Slide 10
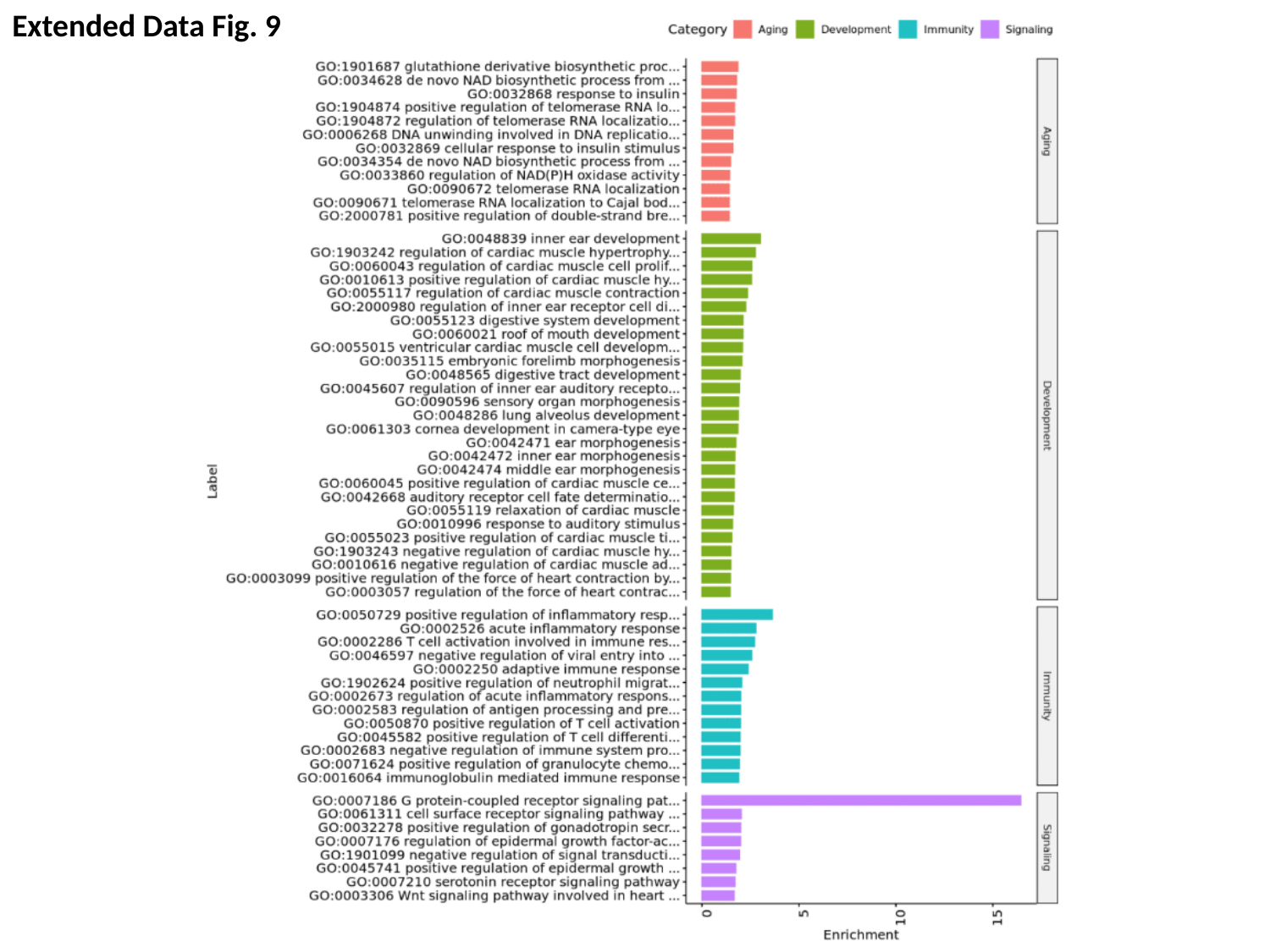

Extended Data Fig. 9

### Slide 11
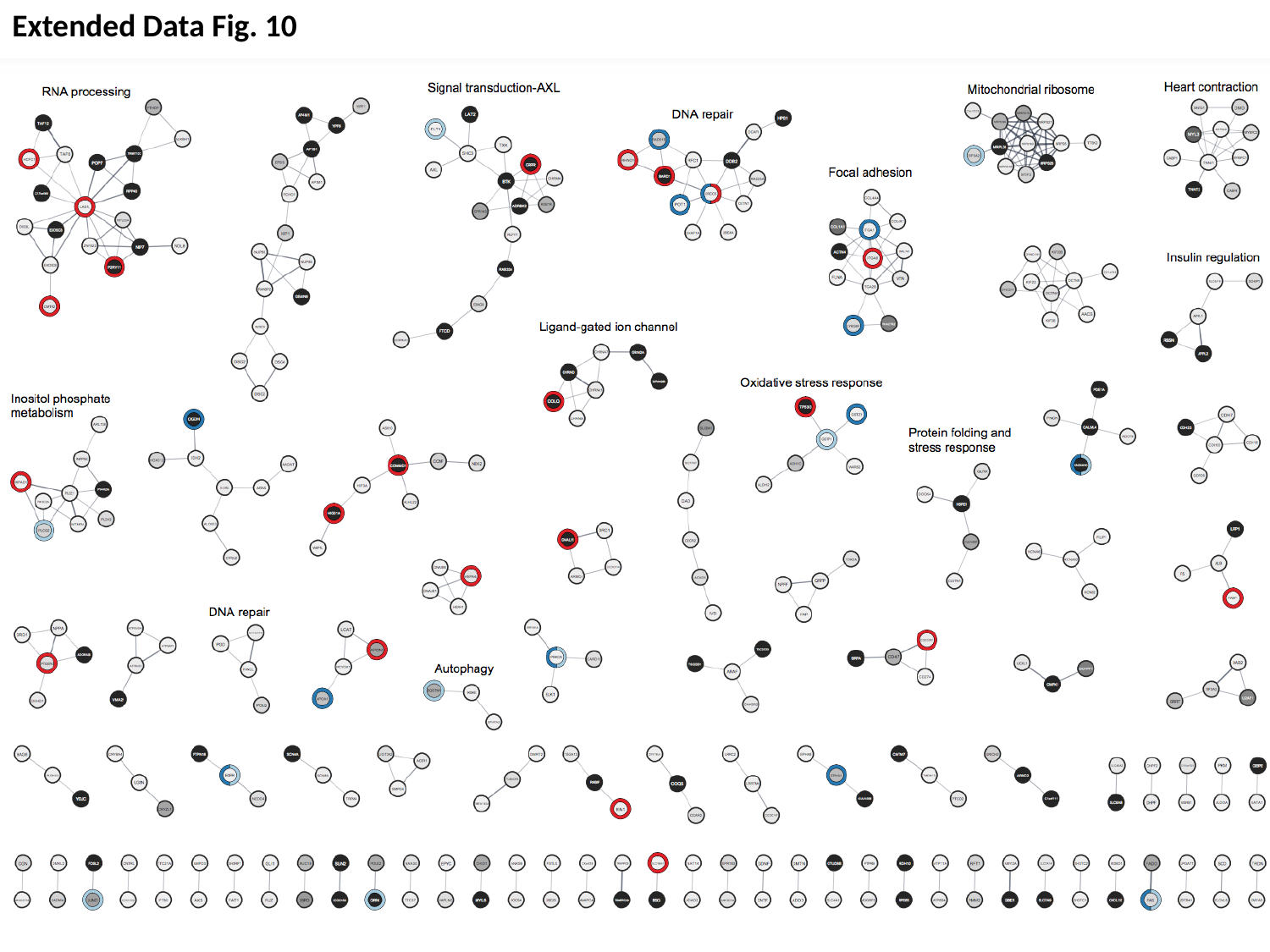

Extended Data Fig. 10

### Slide 12
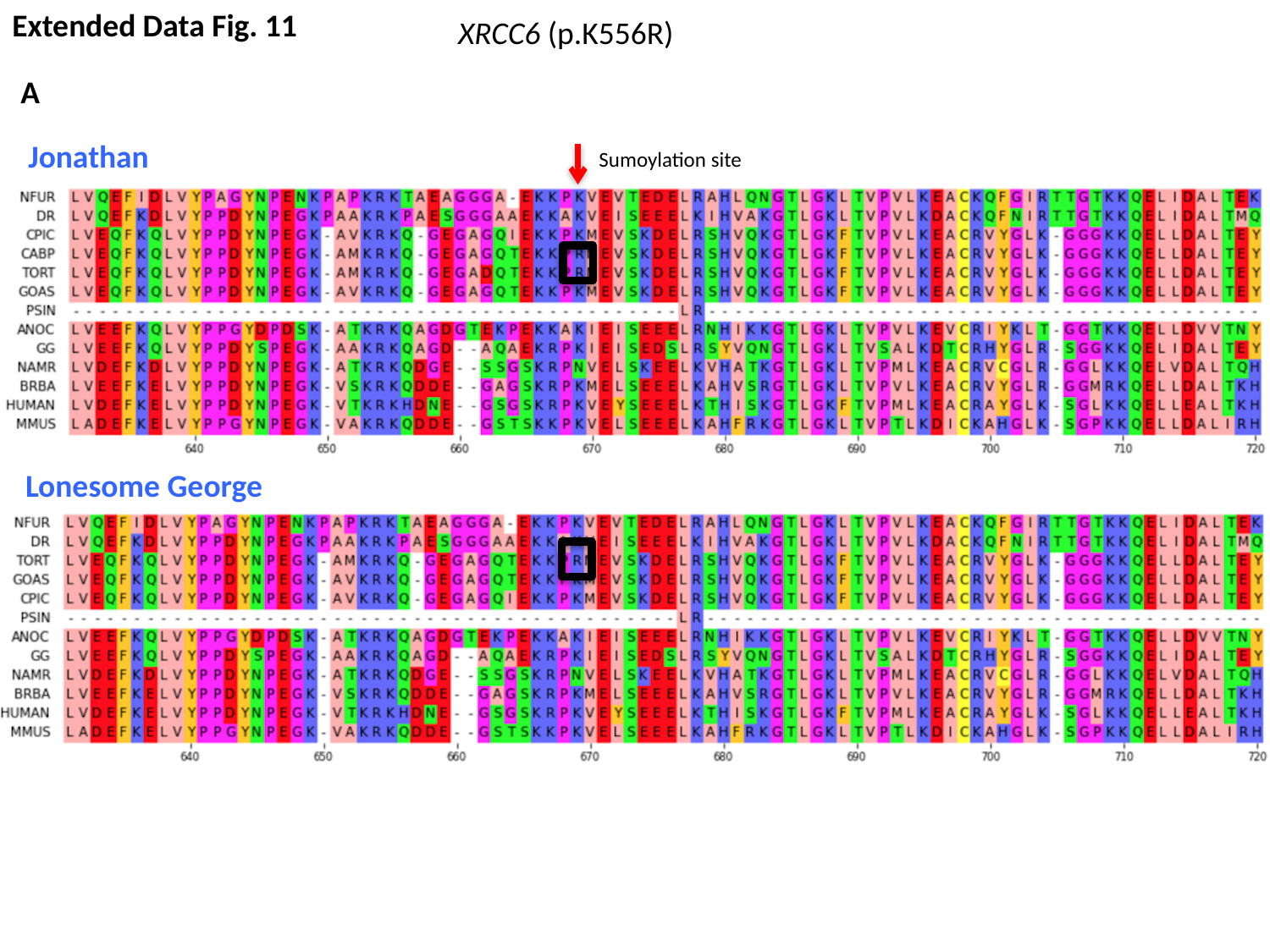

Extended Data Fig. 11
XRCC6 (p.K556R)
A
Jonathan
Sumoylation site
Lonesome George

### Slide 13
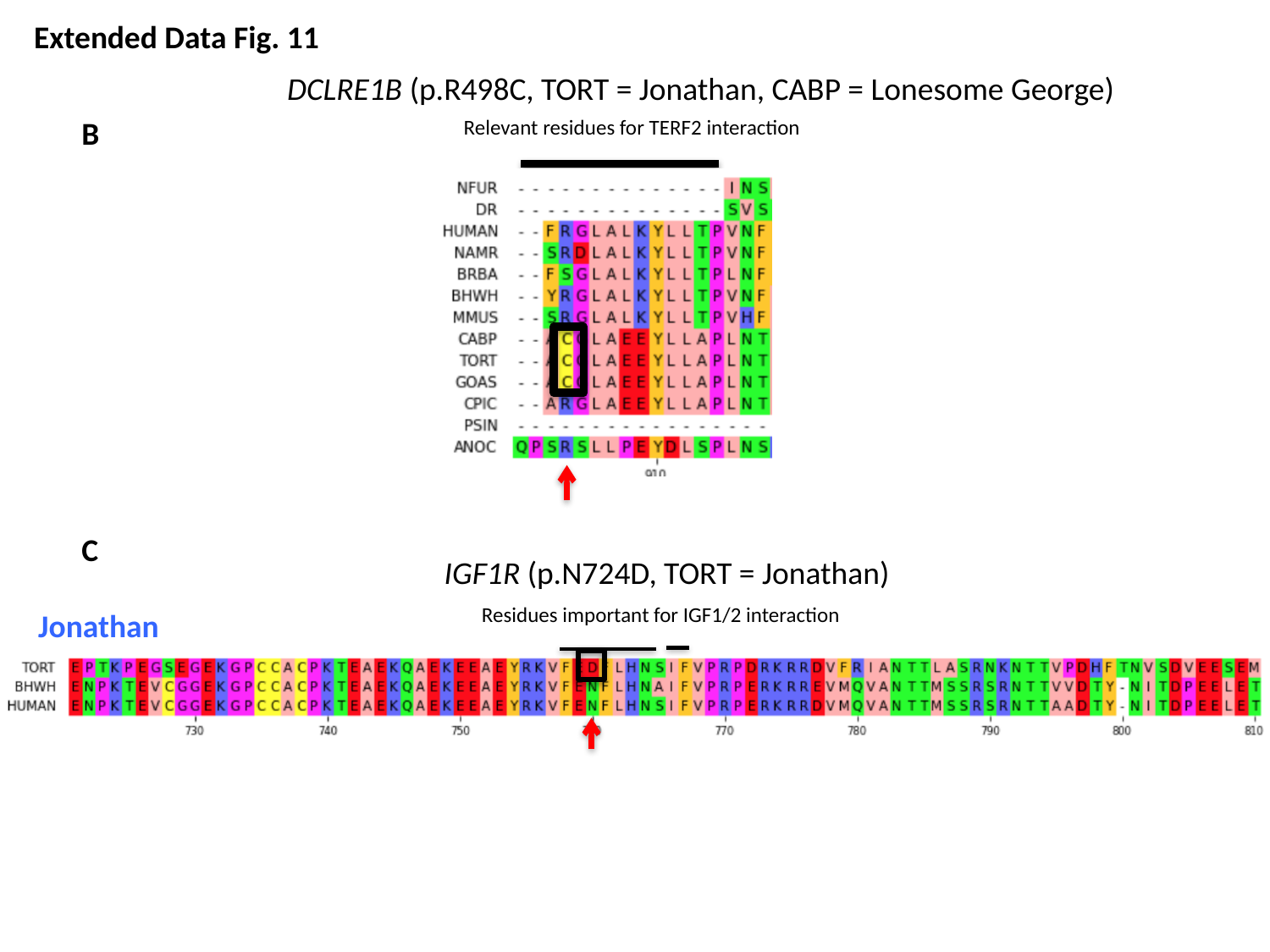

Extended Data Fig. 11
DCLRE1B (p.R498C, TORT = Jonathan, CABP = Lonesome George)
B
Relevant residues for TERF2 interaction
C
IGF1R (p.N724D, TORT = Jonathan)
Residues important for IGF1/2 interaction
Jonathan

### Slide 14
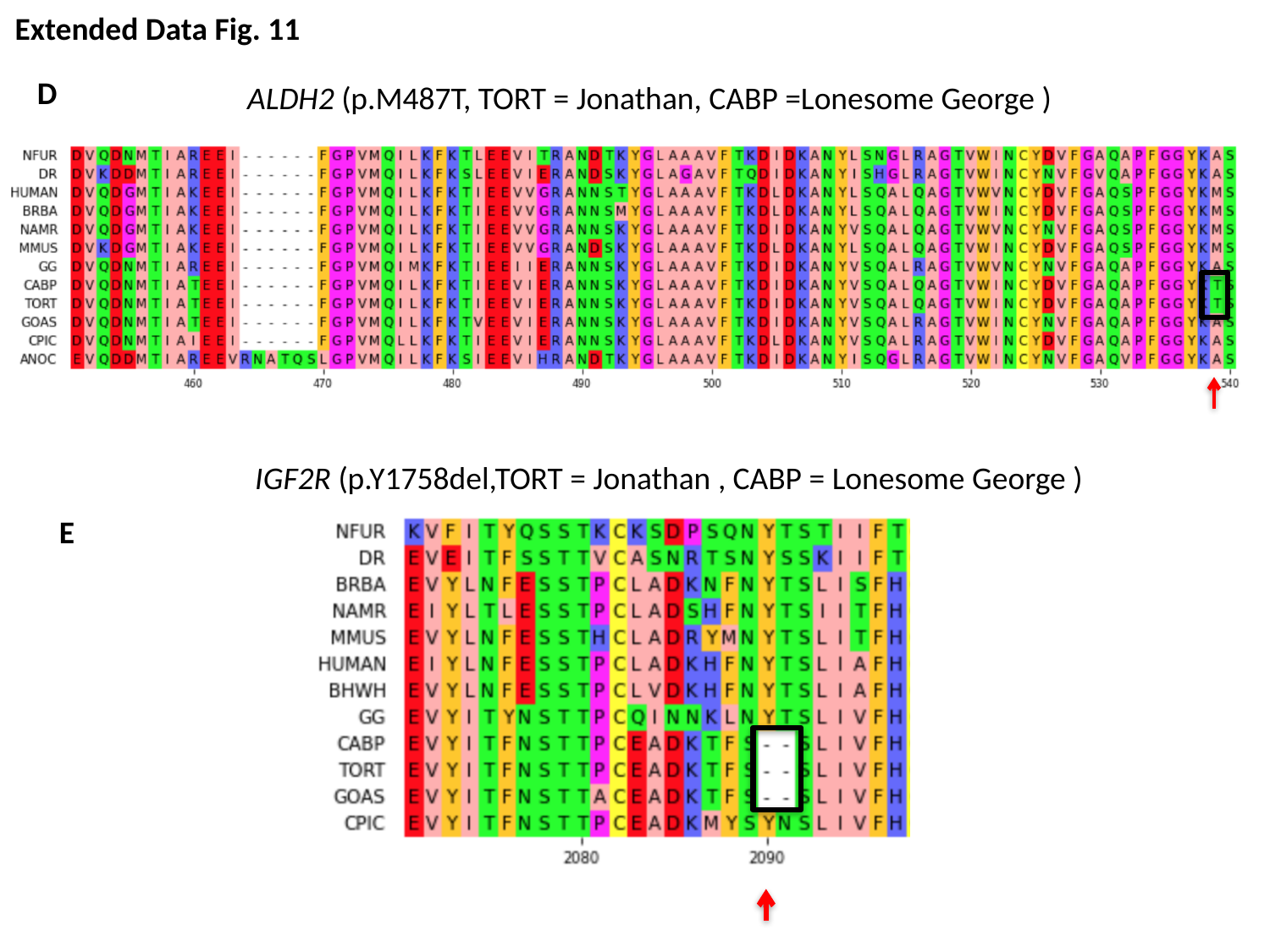

Extended Data Fig. 11
D
ALDH2 (p.M487T, TORT = Jonathan, CABP =Lonesome George )
IGF2R (p.Y1758del,TORT = Jonathan , CABP = Lonesome George )
E

### Slide 15
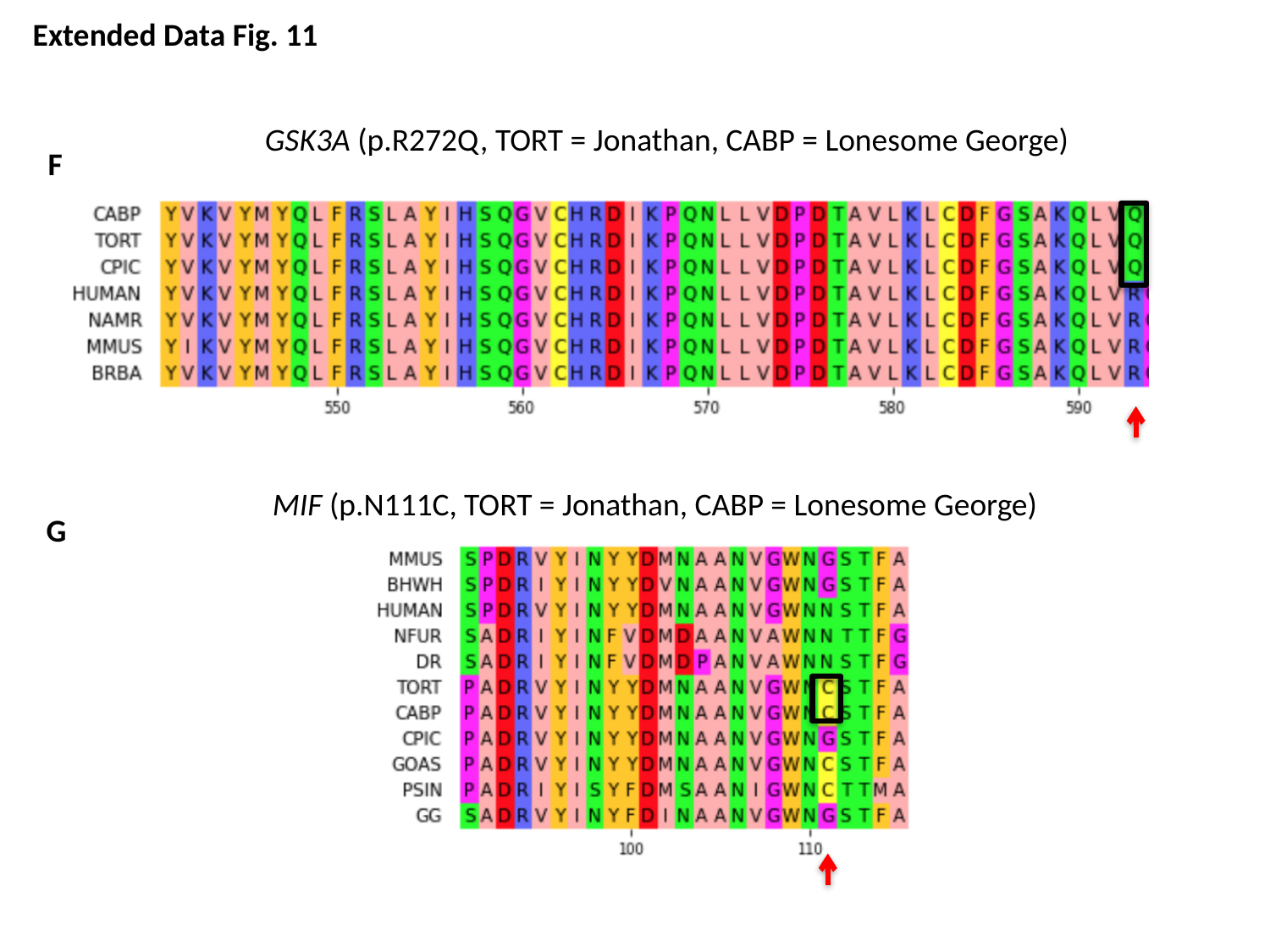

Extended Data Fig. 11
GSK3A (p.R272Q, TORT = Jonathan, CABP = Lonesome George)
F
MIF (p.N111C, TORT = Jonathan, CABP = Lonesome George)
G

### Slide 16
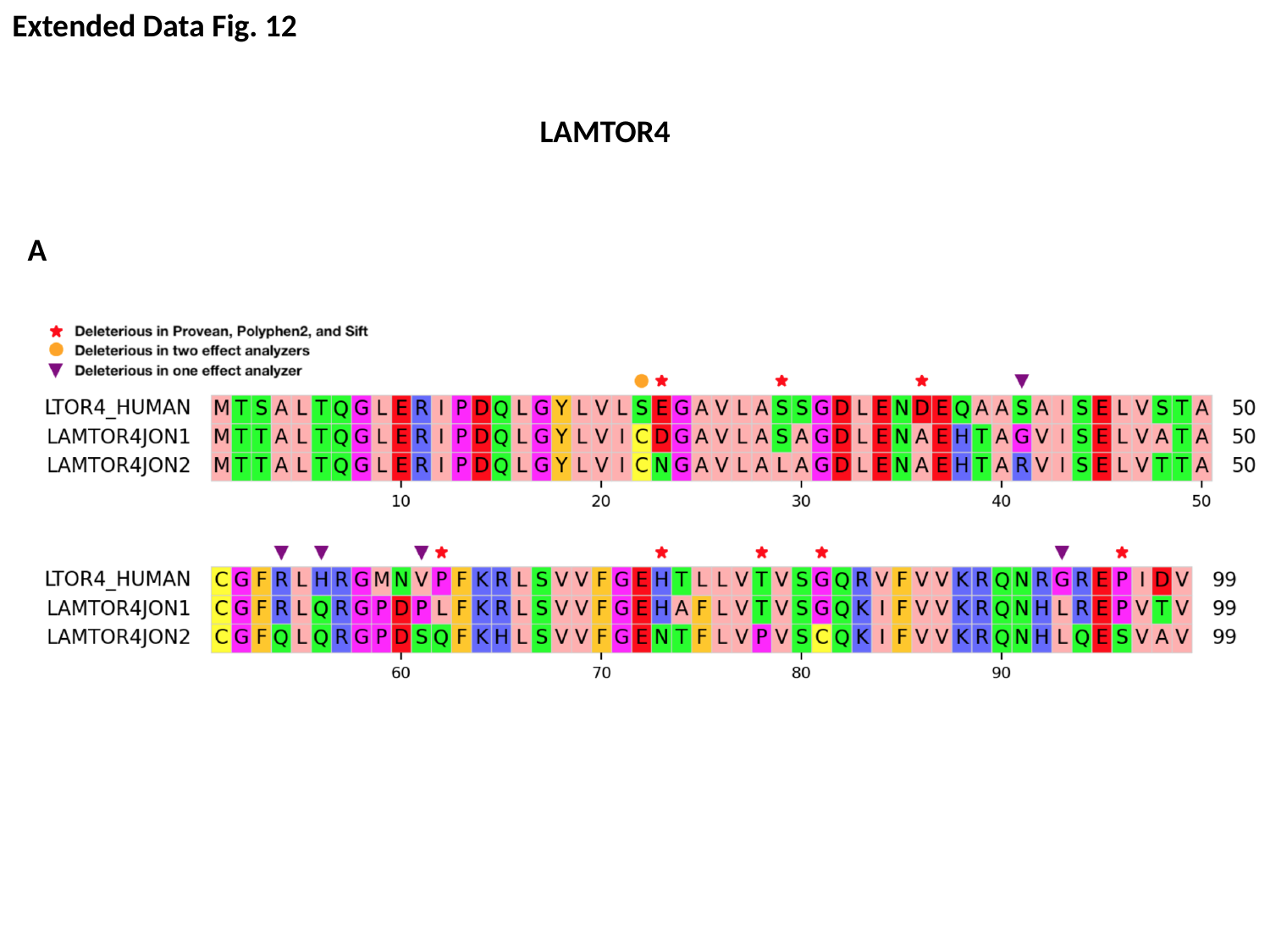

Extended Data Fig. 12
LAMTOR4
A

### Slide 17
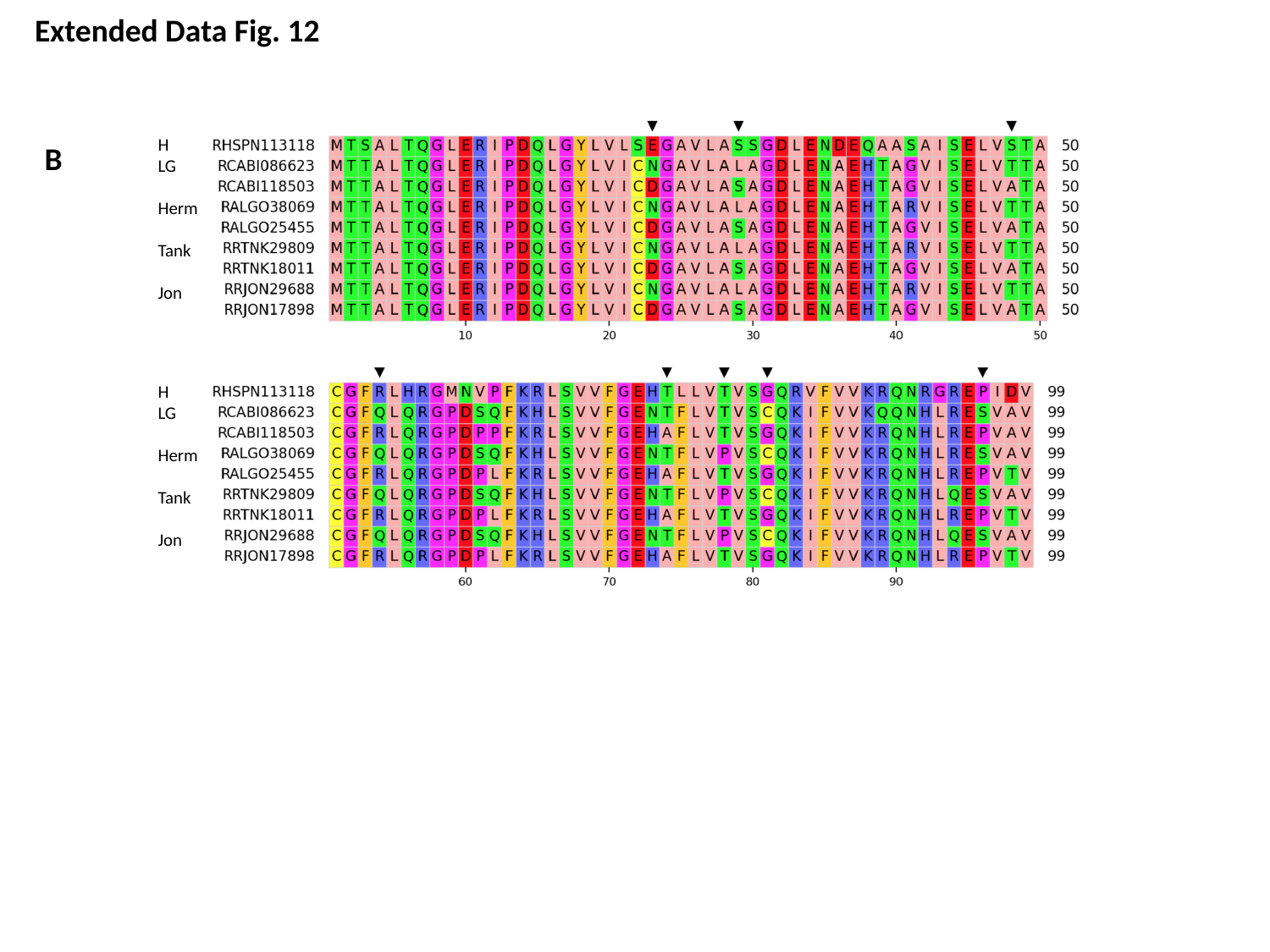

Extended Data Fig. 12
H
LG
Herm
Tank
Jon
H
LG
Herm
Tank
Jon
B

### Slide 18
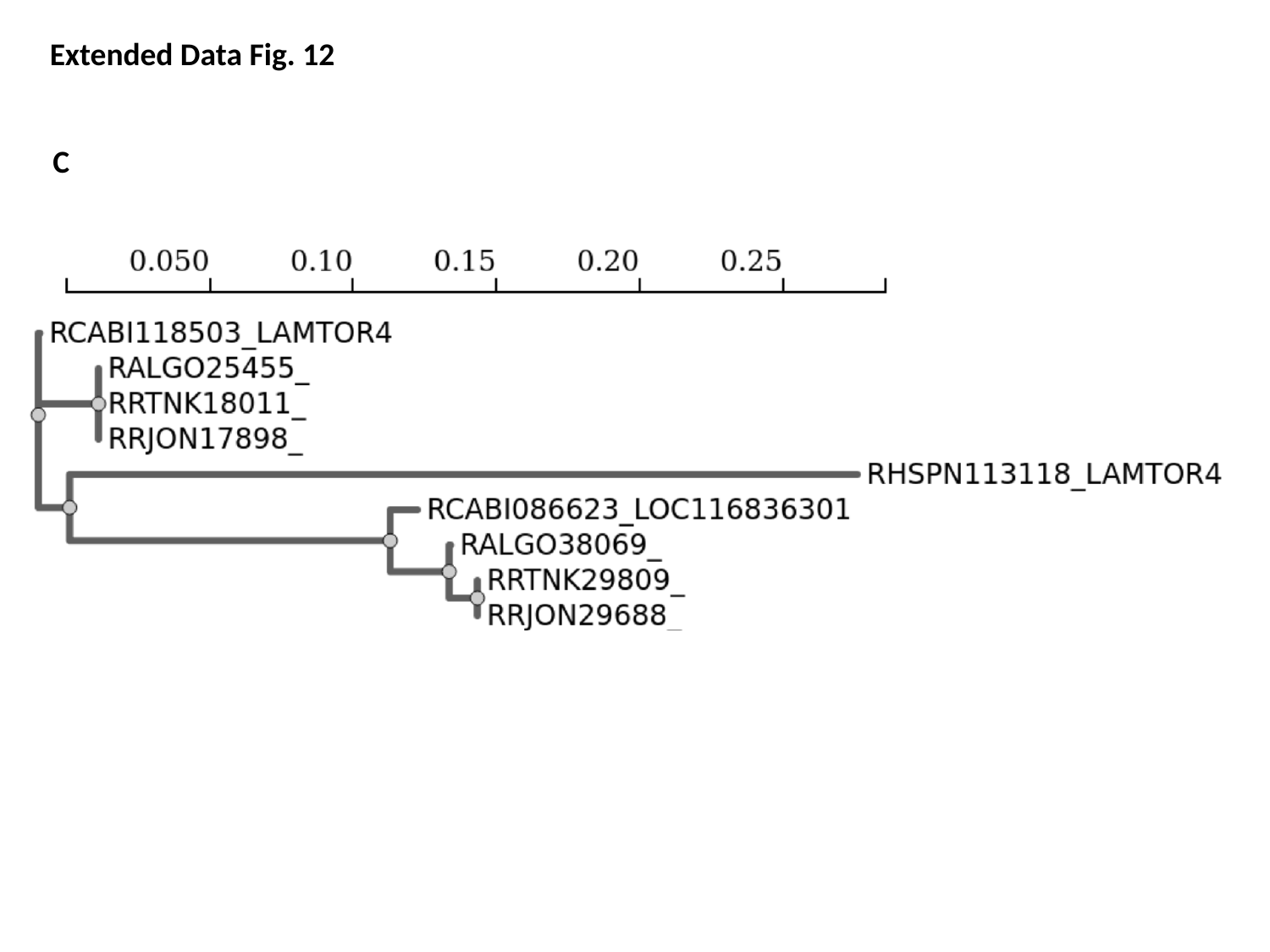

Extended Data Fig. 12
C

### Slide 19
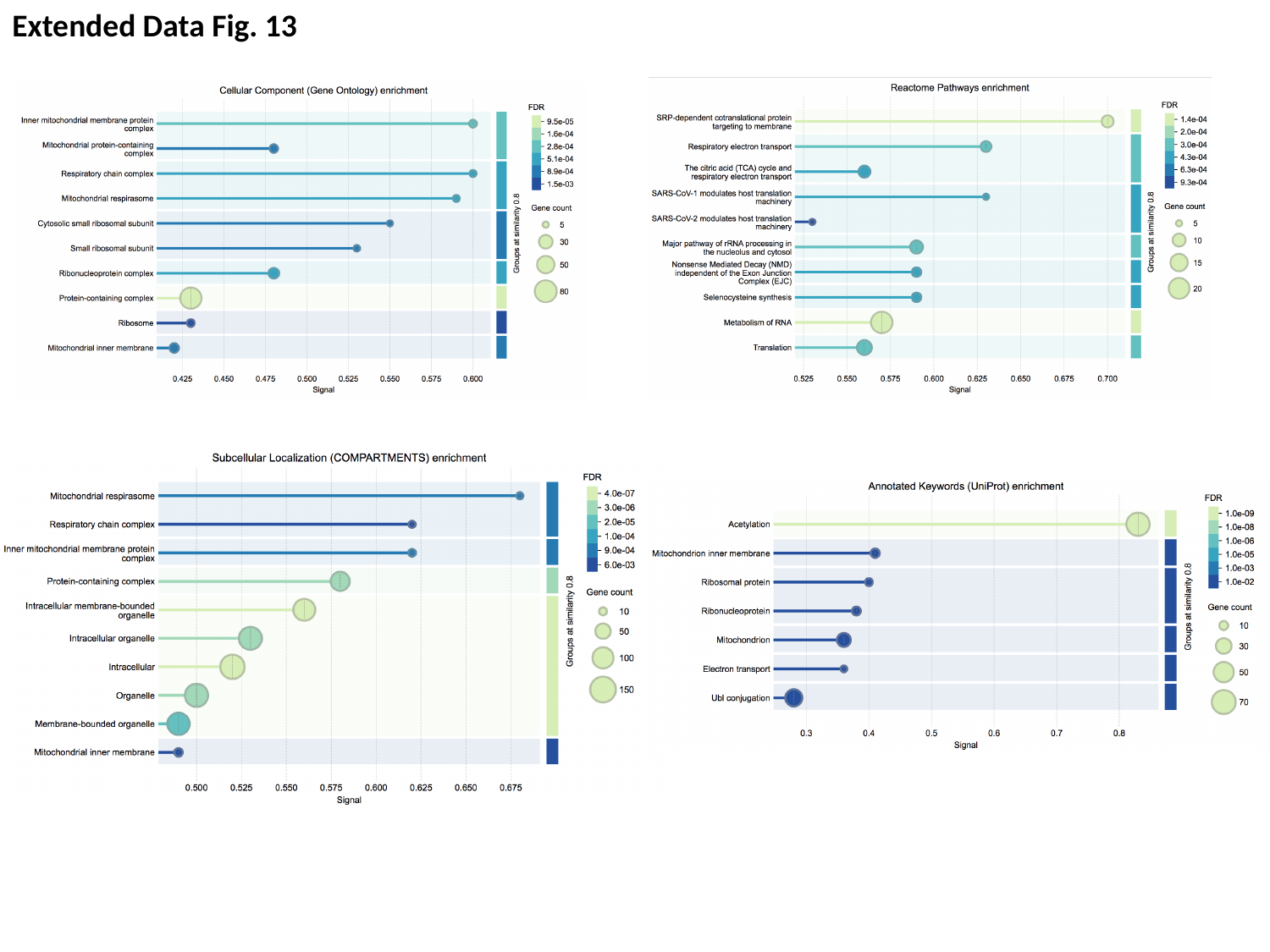

Extended Data Fig. 13
