## Supplement Tables for "Epigenomic insights into extreme longevity in the world’s oldest terrestrial animal, Jonathan"

**Supplementary Table S1**

**A** Assembly Metrics
**B** Repeat Analysis Jonathan
**C** Repeat Analysis Tank
**D** Mitochondrial Genome Jonathan

### **A** Assembly Metrics

| Metric | Jon | Tank |
|---|---|---|
| Size (bp) | ########## | ########## |
| N (%) | 0.00 | 0.00 |
| GC (%) | 44.40 | 44.41 |
| AT (%) | 55.60 | 55.59 |
| Scaffold Cou | 213.00 | 213.00 |
| Longest Scaff | ########## | ########## |
| scaffold N50 | ########## | ########## |
| scaffold N50 | 6.00 | 6.00 |
| scaffold N90 | ########## | ########## |
| scaffold N90 | 20.00 | 20.00 |
| contig count | 224.00 | 224.00 |
| contig N50 le | ########## | ########## |
| contig N50 co | 7.00 | 7.00 |
| contig N90 le | ########## | ########## |
| contig N90 co | 27.00 | 27.00 |
| gene count | 25738.00 | 25723.00 |
| CDS Count | 31416.00 | 31260.00 |
| Mean CDS Le | 550.00 | 549.00 |
| Median CDS | 382.00 | 380.00 |
| Max CDS Len | 23879.00 | 34300.00 |
| Annotated CI | 19993.00 | 20083.00 |
| Retroelemen | 1402438 | 1488176 |
| DNA transpo: | 2886784 | 2814310 |

### **B** Repeat Analysis Jonathan

| | | | | |
|---|---|---|---|---|
| sequences: | 213 | | | |
| total length: | 2412858454 | (2412852954 bp excl N/X-runs) | | |
| GC level: | 44.41% | | | |
| bases maske | 1102605401 bp ( 45.70 %) | | | |
| | | number of elements* | length occupied | percentage of sequence |
| **Retroelements** | | **1402438** | **536149276 b** | **22.22%** |
| | **SINEs:** | **281473** | **41550533 bp** | **1.72%** |
| | **Penelope:** | **0** | **0 bp** | **0.00%** |
| | **LINEs:** | **754296** | **289691774 b** | **12.01%** |
| | CRE/SLACS | 0 | 0 bp | 0.00% |

| | | | |
|---|---|---|---|
| L2/CR1/Rex | 585705 | 220820522 b | 9.15% |
| R1/LOA/Jock | 1037 | 1121809 bp | 0.05% |
| R2/R4/NeSL | 10741 | 9218617 bp | 0.38% |
| RTE/Bov-B | 16135 | 9685712 bp | 0.40% |
| L1/CIN4 | 13033 | 2738874 bp | 0.11% |
| **LTR element** | **366669** | **204906969 b** | **8.49%** |
| BEL/Pao | 0 | 0 bp | 0.00% |
| Ty1/Copia | 0 | 0 bp | 0.00% |
| Gypsy/DIRS1 | 171416 | 134991695 b | 5.59% |
| Retroviral | 233 | 117796 bp | 0.00% |
| **DNA transposons** | **2886784** | **485505338 b** | **20.12%** |
| hobo-Activat | 4464 | 1682109 bp | 0.07% |
| Tc1-IS630-Pc | 0 | 0 bp | 0.00% |
| En-Spm | 0 | 0 bp | 0.00% |
| MULE-MuDR | 0 | 0 bp | 0.00% |
| PiggyBac | 0 | 0 bp | 0.00% |
| Tourist/Harbi | 865 | 160846 bp | 0.01% |
| Other (Mirag | 0 | 0 bp | 0.00% |
| **Rolling-circles** | **4** | **551 bp** | **0.00%** |
| **Unclassified:** | **555375** | **80545699 bp** | **3.34%** |
| **Total interspersed repeats:** | | **1102200313** | **45.68%** |
| **Small RNA:** | **0** | **0 bp** | **0.00%** |
| **Satellites:** | **0** | **0 bp** | **0.00%** |
| **Simple repeats:** | **516** | **404537 bp** | **0.02%** |
| **Low complexity:** | **0** | **0 bp** | **0.00%** |

==================================================

### **C** Repeat Analysis Tank

file name: tank.fasta.mod
sequences: 213
total length: : (2413091811 bp excl N/X-runs)
GC level: 44.41%
bases masked: 1107087936 bp ( 45.88 %)

| | number of elements* | length occupied | percentage of sequence |
|---|---|---|---|
| **Retroelements** | **1488176** | **557549290 b** | **23.11%** |
| **SINEs:** | **223118** | **35270031 bp** | **1.46%** |

| | | | |
|---|---|---|---|
| **Penelope:** | **0** | **0 bp** | **0.00%** |
| **LINEs:** | **840751** | **301927609 b** | **12.51%** |
| CRE/SLACS | 0 | 0 bp | 0.00% |
| L2/CR1/Rex | 653935 | 229905605 b | 9.53% |
| R1/LOA/Jock | 4244 | 2286609 bp | 0.09% |
| R2/R4/NeSL | 5292 | 8072577 bp | 0.33% |
| RTE/Bov-B | 21285 | 9732375 bp | 0.40% |
| L1/CIN4 | 4177 | 1926307 bp | 0.08% |
| **LTR element** | **424307** | **220351650 b** | **9.13%** |
| BEL/Pao | 0 | 0 bp | 0.00% |
| Ty1/Copia | 0 | 0 bp | 0.00% |
| Gypsy/DIRS1 | 209449 | 142780980 b | 5.92% |
| Retroviral | 807 | 297658 bp | 0.01% |
| **DNA transposons** | **2814310** | **469662142 b** | **19.46%** |
| hobo-Activat | 10627 | 2898195 bp | 0.12% |
| Tc1-IS630-Pc | 138 | 14447 bp | 0.00% |
| En-Spm | 0 | 0 bp | 0.00% |
| MULE-MuDR | 0 | 0 bp | 0.00% |
| PiggyBac | 0 | 0 bp | 0.00% |
| Tourist/Harbi | 995 | 185550 bp | 0.01% |
| Other (Mirag | 0 | 0 bp | 0.00% |
| P-element, Transib) | | | |
| **Rolling-circles** | **1219** | **119228 bp** | **0.00%** |
| **Unclassified:** | **561923** | **79558530 bp** | **3.30%** |
| **Total interspersed repeats** | **1106769962** | **45.87%** | |
| **Small RNA:** | **0** | **0 bp** | **0.00%** |
| **Satellites:** | **0** | **0 bp** | **0.00%** |
| **Simple repeats:** | **352** | **198746 bp** | **0.01%** |
| **Low complexity:** | **0** | **0 bp** | **0.00%** |

==================================================

* most repeats fragmented by insertions or deletions
have been counted as one element

RepeatMasker version 4.1.5 , quick mode

run with rmblastn version 2.14.1+

**D** Mitochondrial Genome Jonathan

| Metric | Jon |
|---|---|
| Size (bp) | 16467 |
| N (%) | 0.00% |
| GC (%) | 40.13% |
| AT (%) | 59.87% |
| Scaffold Cou | 1 |
| Longest Scaf | 16467 |
| scaffold N50 | 16467 |
| scaffold N50 | 1 |
| scaffold N90 | 16467 |
| scaffold N90 | 1 |
| contig count | 1 |
| contig N50 le | 16467 |
| contig N50 c | 1 |
| contig N90 le | 16467 |
| contig N90 c | 1 |
| gene count | 13 |
| CDS Count | 13 |
| Mean CDS Le | 290.46 |
| Median CDS | 261 |
| Max CDS Len | 593 |
| Annotated C | 13 |

**Supplementary Table S2**

**A** Jonathan's positively selected genes (PSGs) with Provean an
**B** Jonathan's PSGs shared with Bowhead Whale PSGs
**C** Jonathan's PSGs shared with Brandit's bat PSGs
**D** Jonathan's PSGs shared with Killifish PSGs (highest confiden
**E** Jonathan's PSGs shared with NMR's PSGs
**F** Jonathan's PSGs shared with Rockfish's PSGs
**G** Jonathan's PSGs shared with LongevityMap significant huma
**H** Jonathan's PSGs shared with GeneAge 307 aging genes

**A** Jonathan's positively selected genes (PSGs) with Provean an

| n_all_relaxe | transcript.id | tortoise | lnL.m1 | w.m1 | w2.m1 | lnL.m2 |
|---|---|---|---|---|---|---|
| **gene** | RHSPN17901 | RJON | -26170.4 | 0.0781 | 1 | -26070.45 |
| **PTPRB** | RHSPN04656 | RJON | -7112.2902 | 0.04281 | 1 | -7025.9491 |
| **ASIC4** | RHSPN17579 | RJON | -7185.9907 | 0.05448 | 1 | -7122.045 |
| **ASIC1** | RHSPN13203 | RJON | -2415.8004 | 0.0159 | 1 | -2353.7644 |
| **TPM2** | RHSPN08505 | RJON | -39318.718 | 0.02947 | 1 | -39257.228 |
| **FBN2** | RHSPN05072 | RJON | -2556.1018 | 0.0808 | 1 | -2495.2161 |
| **COLQ** | RHSPN13203 | RJON | -2236.7364 | 0.01829 | 1 | -2179.2302 |
| **TPM2** | RHSPN03209 | RJON | -31871.623 | 0.13221 | 1 | -31820.691 |
| **THADA** | RHSPN19954 | RJON | -41907.802 | 0.00594 | 1 | -41862.174 |
| **DYNC1H1** | RHSPN18870 | RJON | -2784.6498 | 0.0708 | 1 | -2741.8107 |
| **KLF12** | RHSPN25092 | RJON | -4348.0295 | 0.04142 | 1 | -4306.0545 |
| **AXL** | RHSPN28493 | RJON | -17875.441 | 0.04641 | 1 | -17834.19 |
| **L1CAM** | RHSPN05355 | RJON | -1749.3698 | 0.02691 | 1 | -1708.635 |
| **MYL3** | RHSPN02316 | RJON | -2312.3523 | 0.03059 | 1 | -2274.1097 |
| **TNNT2** | RHSPN04371 | RJON | -14533.188 | 0.02544 | 1 | -14498.873 |
| **MYO1B** | RHSPN25073 | RJON | -13793.005 | 0.05433 | 1 | -13759.377 |
| **LTBP4** | RHSPN05006 | RJON | -14068.084 | 0.0228 | 1 | -14035.271 |
| **ATP2B2** | RHSPN06525 | RJON | -5195.0588 | 0.09077 | 1 | -5165.6986 |
| **GP5** | RHSPN06261 | RJON | -7724.9143 | 0.06111 | 1 | -7696.6276 |
| **TIPARP** | RHSPN10297 | RJON | -7625.3668 | 0.05161 | 1 | -7597.2717 |
| **PHACTR2** | RHSPN00676 | RJON | -2492.3496 | 0.02496 | 1 | -2465.2273 |
| **DNALI1** | RHSPN11338 | RJON | -10083.452 | 0.05236 | 1 | -10056.461 |
| **EPHB4** | RHSPN02811 | RJON | -11983.194 | 0.1633 | 1 | -11956.655 |
| **NLRP3** | RHSPN04854 | RJON | -7512.0172 | 0.07049 | 1 | -7486.3347 |
| **GPC1** | RHSPN26674 | RJON | -4382.9858 | 0.08944 | 1 | -4357.3165 |
| **CRYZL1** | RHSPN16699 | RJON | -33416.862 | 0.06032 | 1 | -33391.089 |
| **DYNC2H1** | RHSPN16368 | RJON | -2463.5235 | 0.04198 | 1 | -2437.8385 |
| **CABP4** | RHSPN09032 | RJON | -6643.2413 | 0.05556 | 1 | -6618.205 |
| **RASGEF1C** | RHSPN23483 | RJON | -959.60453 | 0.0932 | 1 | -934.81773 |
| **MXRA7** | RHSPN05241 | RJON | -19466.475 | 0.03405 | 1 | -19441.684 |
| **SCN5A** | RHSPN07357 | RJON | -2999.1885 | 0.02537 | 1 | -2976.1999 |
| **ELOVL6** | RHSPN05242 | RJON | -19646.04 | 0.03478 | 1 | -19623.098 |

| | | | | | | |
|---|---|---|---|---|---|---|
| **SCN5A** | RHSPN15514 | RJON | -4836.9183 | 0.09125 | 1 | -4814.1415 |
| **CHID1** | RHSPN07975 | RJON | -4476.7168 | 0.16366 | 1 | -4453.9475 |
| **UGT3A2** | RHSPN04499 | RJON | -7337.2226 | 0.05413 | 1 | -7314.5585 |
| **ADAM23** | RHSPN19414 | RJON | -4762.9512 | 0.05063 | 1 | -4740.4437 |
| **POLE2** | RHSPN07275 | RJON | -5490.9101 | 0.08563 | 1 | -5469.1187 |
| **ADH1C** | RHSPN18335 | RJON | -1992.2251 | 0.01796 | 1 | -1970.5574 |
| **CABP1** | RHSPN04035 | RJON | -4216.3411 | 0.08648 | 1 | -4195.0574 |
| **ARHGAP15** | RHSPN07960 | RJON | -2635.7701 | 0.21129 | 1 | -2614.7602 |
| **TTC23L** | RHSPN22612 | RJON | -2186.2233 | 0.01098 | 1 | -2165.2119 |
| **RPL23A** | RHSPN22370 | RJON | -11605.704 | 0.1 | 1 | -11584.82 |
| **ALOXE3** | RHSPN12623 | RJON | -7750.2022 | 0.03184 | 1 | -7729.546 |
| **GRHL2** | RHSPN02759 | RJON | -4660.2878 | 0.06582 | 1 | -4639.9344 |
| **PLD5** | RHSPN11094 | RJON | -2730.47 | 0.19597 | 1 | -2710.5497 |
| **LAT2** | RHSPN03338 | RJON | -1708.9571 | 0.06417 | 1 | -1689.1312 |
| **COMMD1** | RHSPN24078 | RJON | -7854.9764 | 0.02657 | 1 | -7835.1819 |
| **KIAA1468** | RHSPN05297 | RJON | -1436.9897 | 0.0422 | 1 | -1417.4568 |
| **HIGD1A** | RHSPN14989 | RJON | -21716.434 | 0.06235 | 1 | -21696.921 |
| **MYOF** | RHSPN06573 | RJON | -9138.5685 | 0.03112 | 1 | -9119.5502 |
| **DLG1** | RHSPN26858 | RJON | -6686.1323 | 0.05867 | 1 | -6667.4279 |
| **FTCD** | RHSPN08286 | RJON | -3006.6572 | 0.02457 | 1 | -2987.9777 |
| **EDIL3** | RHSPN00579 | RJON | -4320.6236 | 0.11375 | 1 | -4302.4216 |
| **COL16A1** | RHSPN15099 | RJON | -5357.34 | 0.09125 | 1 | -5339.2182 |
| **SCD** | RHSPN14721 | RJON | -667.07602 | 0.04298 | 1 | -649.04262 |
| **PBLD** | RHSPN28177 | RJON | -1479.335 | 0.04143 | 1 | -1461.5716 |
| **TSC22D3** | RHSPN28487 | RJON | -9323.8938 | 0.03563 | 1 | -9306.3135 |
| **ABCD1** | RHSPN21048 | RJON | -5091.4229 | 0.07114 | 1 | -5073.8634 |
| **DECR2** | RHSPN17078 | RJON | -22361.873 | 0.02805 | 1 | -22344.553 |
| **CACNA1C** | RHSPN19739 | RJON | -4017.438 | 0.15309 | 1 | -4000.2773 |
| **CIPC** | RHSPN09531 | RJON | -11321.961 | 0.01796 | 1 | -11304.952 |
| **GRM4** | RHSPN26796 | RJON | -2408.4763 | 0.01453 | 1 | -2391.5108 |
| **U2AF1** | RHSPN04659 | RJON | -11891.626 | 0.0419 | 1 | -11874.66 |
| **OBSL1** | RHSPN15973 | RJON | -3867.7987 | 0.0922 | 1 | -3850.9187 |
| **DDB2** | RHSPN26725 | RJON | -5023.6639 | 0.03067 | 1 | -5006.9402 |
| **ERG** | RHSPN12154 | RJON | -8797.3652 | 0.13368 | 1 | -8780.9069 |
| **NUGGC** | RHSPN20789 | RJON | -1459.339 | 0.01485 | 1 | -1442.9293 |
| **CRABP1** | RHSPN18809 | RJON | -6484.8475 | 0.1491 | 1 | -6468.4544 |
| **NEK5** | RHSPN03684 | RJON | -7681.8396 | 0.03941 | 1 | -7665.9306 |
| **MGAT4A** | RHSPN03880 | RJON | -4138.2559 | 0.08436 | 1 | -4122.59 |
| **MARCO** | RHSPN12310 | RJON | -2223.2034 | 0.03104 | 1 | -2208.078 |
| **SNTG1** | RHSPN21187 | RJON | -3699.8202 | 0.07449 | 1 | -3684.8397 |
| **ZNF263** | RHSPN02067 | RJON | -8393.8531 | 0.02831 | 1 | -8379.0723 |
| **DCAF6** | RHSPN17344 | RJON | -2914.1828 | 0.01844 | 1 | -2899.5443 |
| **SOX5** | RHSPN24280 | RJON | -2303.7974 | 0.01066 | 1 | -2289.2159 |
| **AES** | RHSPN25331 | RJON | -2374.1142 | 0.05622 | 1 | -2359.5434 |
| **FAM83E** | RHSPN03616 | RJON | -2523.4317 | 0.02956 | 1 | -2508.9378 |

| | | | | | | |
|---|---|---|---|---|---|---|
| **KCNIP3** | RHSPN18976 | RJON | -3234.9671 | 0.09686 | 1 | -3220.5788 |
| **UBAC2** | RHSPN08661 | RJON | -10397.279 | 0.12833 | 1 | -10383.056 |
| **PCDHB1** | RHSPN27657 | RJON | -7283.1202 | 0.09819 | 1 | -7268.9988 |
| **REPS2** | RHSPN18100 | RJON | -19277.523 | 0.05925 | 1 | -19263.44 |
| **MYBPC1** | RHSPN28498 | RJON | -18802.591 | 0.02143 | 1 | -18788.552 |
| **HCFC1** | RHSPN21187 | RJON | -3903.6185 | 0.0768 | 1 | -3889.6478 |
| **ZNF263** | RHSPN11350 | RJON | -6105.4346 | 0.06293 | 1 | -6091.5833 |
| **TRIM56** | RHSPN20585 | RJON | -4905.9367 | 0.09545 | 1 | -4892.1647 |
| **ANKDD1A** | RHSPN05818 | RJON | -5707.1841 | 0.23214 | 1 | -5693.6415 |
| **CD47** | RHSPN07090 | RJON | -8777.4236 | 0.19914 | 1 | -8763.8712 |
| **ALB** | RHSPN20875 | RJON | -19496.332 | 0.08833 | 1 | -19482.912 |
| **AKAP13** | RHSPN17273 | RJON | -8854.2612 | 0.01507 | 1 | -8840.9755 |
| **GRIN2B** | RHSPN02022 | RJON | -2182.2823 | 0.11185 | 1 | -2169.0153 |
| **C1orf111** | RHSPN05443 | RJON | -43200.138 | 0.07818 | 1 | -43186.9 |
| **BSN** | RHSPN09648 | RJON | -6709.0496 | 0.12721 | 1 | -6696.0044 |
| **PGC** | RHSPN26469 | RJON | -7316.4651 | 0.03291 | 1 | -7303.5129 |
| **YTHDF1** | RHSPN06288 | RJON | -2006.2454 | 0.00471 | 1 | -1993.3864 |
| **SCHIP1** | RHSPN18442 | RJON | -2890.6771 | 0.10762 | 1 | -2877.8298 |
| **GLT1D1** | RHSPN17982 | RJON | -7322.6574 | 0.10028 | 1 | -7309.8849 |
| **RASSF9** | RHSPN12679 | RJON | -30730.911 | 0.0211 | 1 | -30718.156 |
| **CSMD3** | RHSPN17741 | RJON | -2076.9695 | 0.01404 | 1 | -2064.4043 |
| **MYL6** | RHSPN00055 | RJON | -18140.088 | 0.12717 | 1 | -18127.632 |
| **TAS1R3** | RHSPN11591 | RJON | -3389.4019 | 0.21108 | 1 | -3377.0269 |
| **TSGA13** | RHSPN07949 | RJON | -20122.562 | 0.0795 | 1 | -20110.194 |
| **ADAMTS12** | RHSPN23292 | RJON | -22714.534 | 0.03123 | 1 | -22702.319 |
| **SCN4A** | RHSPN00210 | RJON | -6415.1391 | 0.0279 | 1 | -6402.9849 |
| **PGD** | RHSPN03299 | RJON | -20813.729 | 0.03912 | 1 | -20801.668 |
| **CCDC88A** | RHSPN17991 | RJON | -40639.652 | 0.07837 | 1 | -40627.761 |
| **CEP290** | RHSPN21530 | RJON | -5497.5873 | 0.03935 | 1 | -5485.713 |
| **DOC2A** | RHSPN03780 | RJON | -37732.992 | 0.0778 | 1 | -37721.139 |
| **RANBP2** | RHSPN19413 | RJON | -6019.2627 | 0.10777 | 1 | -6007.458 |
| **DNAAF2** | RHSPN28113 | RJON | -8104.2159 | 0.0307 | 1 | -8092.4902 |
| **BTK** | RHSPN02082 | RJON | -25308.398 | 0.11555 | 1 | -25296.722 |
| **F5** | RHSPN09763 | RJON | -6438.9469 | 0.17491 | 1 | -6427.3263 |
| **GLYATL3** | RHSPN09347 | RJON | -6351.6322 | 0.1276 | 1 | -6340.0918 |
| **TRIM27** | RHSPN15060 | RJON | -3706.6925 | 0.01265 | 1 | -3695.1664 |
| **PI4K2A** | RHSPN07569 | RJON | -10039.242 | 0.0439 | 1 | -10027.812 |
| **LRBA** | RHSPN09370 | RJON | -960.15777 | 0.17656 | 1 | -948.7636 |
| **MOG** | RHSPN15516 | RJON | -8979.9804 | 0.0794 | 1 | -8968.6402 |
| **MUC6** | RHSPN05777 | RJON | -2812.3837 | 0.1521 | 1 | -2801.1363 |
| **ADGRG7** | RHSPN22403 | RJON | -2879.0985 | 0.01285 | 1 | -2867.8629 |
| **NTN1** | RHSPN14165 | RJON | -2145.3704 | 0.01575 | 1 | -2134.2993 |
| **ZMYND19** | RHSPN12530 | RJON | -2479.405 | 0.05946 | 1 | -2468.5336 |
| **OTUD6B** | RHSPN22685 | RJON | -12902.582 | 0.03381 | 1 | -12891.741 |
| **MYO1D** | RHSPN10890 | RJON | -15033.981 | 0.02179 | 1 | -15023.225 |

| | | | | | | |
|---|---|---|---|---|---|---|
| **OGDH** | RHSPN02107 | RJON | -26485.705 | 0.06815 | 1 | -26474.984 |
| **PRRC2C** | RHSPN08196 | RJON | -4196.4254 | 0.13006 | 1 | -4185.8419 |
| **TMEM174** | RHSPN25364 | RJON | -12318.137 | 0.02266 | 1 | -12307.565 |
| **PPFIA3** | RHSPN23676 | RJON | -2747.9041 | 0.04401 | 1 | -2737.3427 |
| **CETN1** | RHSPN04308 | RJON | -6351.1335 | 0.03442 | 1 | -6340.5954 |
| **PDE1A** | RHSPN06135 | RJON | -7965.9354 | 0.14225 | 1 | -7955.5101 |
| **ESYT3** | RHSPN08640 | RJON | -3027.6655 | 0.0646 | 1 | -3017.2708 |
| **SRA1** | RHSPN02680 | RJON | -6043.374 | 0.01842 | 1 | -6033.0674 |
| **TTC13** | RHSPN21118 | RJON | -554.82011 | 0.02089 | 1 | -544.67861 |
| **MSRB1** | RHSPN11261 | RJON | -9612.0826 | 0.06155 | 1 | -9602.0373 |
| **ASNS** | RHSPN14998 | RJON | -27776.817 | 0.05771 | 1 | -27766.793 |
| **PLCE1** | RHSPN18801 | RJON | -3797.4581 | 0.07448 | 1 | -3787.4701 |
| **DHRS12** | RHSPN25655 | RJON | -2290.9472 | 0.10384 | 1 | -2280.9813 |
| **TMEM150B** | RHSPN20635 | RJON | -2647.0426 | 0.0763 | 1 | -2637.1279 |
| **CALML4** | RHSPN11111 | RJON | -11414.631 | 0.0411 | 1 | -11404.745 |
| **HIP1** | RHSPN21512 | RJON | -2741.1843 | 0.05875 | 1 | -2731.3259 |
| **KIF22** | RHSPN11328 | RJON | -6474.5136 | 0.0637 | 1 | -6464.6697 |
| **LRCH4** | RHSPN00904 | RJON | -3790.0004 | 0.0459 | 1 | -3780.1971 |
| **FOXD2** | RHSPN21148 | RJON | -1589.857 | 0.00443 | 1 | -1580.0904 |
| **ATP6V0C** | RHSPN12810 | RJON | -13427.469 | 0.03253 | 1 | -13417.7 |
| **TRAPPC9** | RHSPN23122 | RJON | -4247.1797 | 0.06451 | 1 | -4237.4305 |
| **COL1A1** | RHSPN25089 | RJON | -8236.5473 | 0.0721 | 1 | -8226.9463 |
| **CYP2A13** | RHSPN04765 | RJON | -5785.9721 | 0.06722 | 1 | -5776.3811 |
| **CHRND** | RHSPN01326 | RJON | -8580.3623 | 0.04999 | 1 | -8570.8328 |
| **AMY2A** | RHSPN01430 | RJON | -6077.6977 | 0.05976 | 1 | -6068.1845 |
| **SLC16A1** | RHSPN03047 | RJON | -11396.002 | 0.07543 | 1 | -11386.546 |
| **EMILIN1** | RHSPN08832 | RJON | -13984.779 | 0.08142 | 1 | -13975.413 |
| **ADAM19** | RHSPN04018 | RJON | -3928.6794 | 0.08817 | 1 | -3919.3414 |
| **LCT** | RHSPN27847 | RJON | -4669.29 | 0.06409 | 1 | -4659.9782 |
| **CCDC120** | RHSPN02240 | RJON | -10884.362 | 0.0875 | 1 | -10875.062 |
| **HMCN1** | RHSPN09753 | RJON | -12468.332 | 0.12572 | 1 | -12459.105 |
| **ADGRF5** | RHSPN25170 | RJON | -4734.8517 | 0.01306 | 1 | -4725.669 |
| **CADM4** | RHSPN22270 | RJON | -10258.033 | 0.02271 | 1 | -10248.88 |
| **KIF1C** | RHSPN05455 | RJON | -7480.1885 | 0.08919 | 1 | -7471.0932 |
| **TRAIP** | RHSPN21805 | RJON | -8492.0638 | 0.11101 | 1 | -8482.9958 |
| **CDH16** | RHSPN00361 | RJON | -2029.3105 | 0.06132 | 1 | -2020.2818 |
| **PLA2G2E** | RHSPN11076 | RJON | -6900.461 | 0.06677 | 1 | -6891.4857 |
| **TBL2** | RHSPN12253 | RJON | -1220.8251 | 0.04888 | 1 | -1211.9251 |
| **ZMAT4** | RHSPN20459 | RJON | -44305.406 | 0.04673 | 1 | -44296.508 |
| **DMXL2** | RHSPN00155 | RJON | -1081.8528 | 0.06012 | 1 | -1072.9996 |
| **HES3** | RHSPN18643 | RJON | -15267.683 | 0.06328 | 1 | -15258.836 |
| **STARD13** | RHSPN15004 | RJON | -8007.2027 | 0.12845 | 1 | -7998.4665 |
| **CYP2C18** | RHSPN12199 | RJON | -7416.0113 | 0.11446 | 1 | -7407.2986 |
| **TTI2** | RHSPN00884 | RJON | -3211.411 | 0.01624 | 1 | -3202.7394 |
| **MOB3C** | RHSPN09610 | RJON | -5633.8182 | 0.01937 | 1 | -5625.1565 |

| | | | | | | |
|---|---|---|---|---|---|---|
| **BTBD9** | RHSPN11264 | RJON | -16859.28 | 0.08685 | 1 | -16850.661 |
| **LMTK2** | RHSPN24403 | RJON | -11073.505 | 0.01347 | 1 | -11064.896 |
| **XAB2** | RHSPN09815 | RJON | -45019.492 | 0.0741 | 1 | -45010.885 |
| **DST** | RHSPN14748 | RJON | -1447.858 | 0.05788 | 1 | -1439.3288 |
| **COL13A1** | RHSPN15713 | RJON | -2953.9347 | 0.14954 | 1 | -2945.4252 |
| **ZBED5** | RHSPN05387 | RJON | -2603.0002 | 0.14109 | 1 | -2594.505 |
| **CAMP** | RHSPN01637 | RJON | -1774.7983 | 0.05109 | 1 | -1766.3045 |
| **MRPS21** | RHSPN16812 | RJON | -3673.4954 | 0.17721 | 1 | -3665.0484 |
| **APOA1** | RHSPN04376 | RJON | -1363.7693 | 0.04102 | 1 | -1355.3296 |
| **TMEFF2** | RHSPN20356 | RJON | -2896.2422 | 0.12654 | 1 | -2887.7981 |
| **ZSCAN29** | RHSPN18567 | RJON | -125.92295 | 0 | 1 | -117.50265 |
| **RNF17** | RHSPN06898 | RJON | -14736.742 | 0.06408 | 1 | -14728.34 |
| **RFC1** | RHSPN13493 | RJON | -6259.4903 | 0.07144 | 1 | -6251.0781 |
| **SHC3** | RHSPN22976 | RJON | -2091.7917 | 0.05829 | 1 | -2083.4006 |
| **GRN** | RHSPN00903 | RJON | -1770.2865 | 0.04762 | 1 | -1762.0111 |
| **CMPK1** | RHSPN15707 | RJON | -10014.908 | 0.03749 | 1 | -10006.663 |
| **AMPD3** | RHSPN24574 | RJON | -4541.5516 | 0.02755 | 1 | -4533.3276 |
| **RAD23A** | RHSPN02957 | RJON | -4363.1233 | 0.023 | 1 | -4354.917 |
| **FAM49A** | RHSPN18237 | RJON | -6378.7188 | 0.04103 | 1 | -6370.5282 |
| **ALDH2** | RHSPN02061 | RJON | -4500.1614 | 0.13007 | 1 | -4491.9918 |
| **RCSD1** | RHSPN27352 | RJON | -6737.0645 | 0.08945 | 1 | -6728.926 |
| **SUN2** | RHSPN08822 | RJON | -3154.6507 | 0.10824 | 1 | -3146.5106 |
| **TIMD4** | RHSPN09451 | RJON | -3664.8115 | 0.1607 | 1 | -3656.6777 |
| **LY6G6F** | RHSPN06347 | RJON | -10116.944 | 0.0329 | 1 | -10108.856 |
| **SLC7A14** | RHSPN23205 | RJON | -7194.9088 | 0.02783 | 1 | -7186.8225 |
| **MTMR4** | RHSPN05716 | RJON | -1009.3958 | 0.11946 | 1 | -1001.3361 |
| **GBE1** | RHSPN12517 | RJON | -4143.795 | 0.14318 | 1 | -4135.7501 |
| **CNBD1** | RHSPN11342 | RJON | -11006.319 | 0.02313 | 1 | -10998.323 |
| **SRRT** | RHSPN19643 | RJON | -21972.467 | 0.04279 | 1 | -21964.481 |
| **PCNX** | RHSPN21899 | RJON | -4081.7125 | 0.05369 | 1 | -4073.7308 |
| **EXOSC6** | RHSPN01667 | RJON | -771.39214 | 0.01299 | 1 | -763.41509 |
| **CDC42SE1** | RHSPN23199 | RJON | -15587.213 | 0.08609 | 1 | -15579.264 |
| **BZRAP1** | RHSPN22385 | RJON | -8811.8482 | 0.06721 | 1 | -8803.9094 |
| **ARHGEF15** | RHSPN02729 | RJON | -9732.3554 | 0.01575 | 1 | -9724.4349 |
| **ACTN2** | RHSPN27811 | RJON | -3725.0974 | 0.03509 | 1 | -3717.1736 |
| **ELK1** | RHSPN17491 | RJON | -10567.628 | 0.12955 | 1 | -10559.725 |
| **SCAF11** | RHSPN09833 | RJON | -9251.8726 | 0.10751 | 1 | -9244.0176 |
| **LGSN** | RHSPN16318 | RJON | -738.38252 | 0.02534 | 1 | -730.5344 |
| **EIF1AD** | RHSPN16067 | RJON | -493.6308 | 0.08339 | 1 | -485.81009 |
| **C11orf31** | RHSPN22280 | RJON | -6332.9847 | 0.07242 | 1 | -6325.1771 |
| **NUP88** | RHSPN03002 | RJON | -2361.0863 | 0.095 | 1 | -2353.3185 |
| **TP53I3** | RHSPN20493 | RJON | -8257.108 | 0.03819 | 1 | -8249.3391 |
| **NEDD4** | RHSPN20804 | RJON | -7709.1895 | 0.08148 | 1 | -7701.4185 |
| **ANKRD34C** | RHSPN15272 | RJON | -6138.0852 | 0.08652 | 1 | -6130.3245 |
| **PNLIPRP1** | RHSPN04000 | RJON | -65.097031 | 0 | 1 | -57.394496 |

| | | | | | | |
|---|---|---|---|---|---|---|
| **TMEM163** | RHSPN24900 | RJON | -6190.5414 | 0.06901 | 1 | -6182.8554 |
| **FFAR3** | RHSPN26504 | RJON | -4013.6362 | 0.01695 | 1 | -4005.9576 |
| **UCKL1** | RHSPN07787 | RJON | -58648.786 | 0.0636 | 1 | -58641.129 |
| **FAT1** | RHSPN06763 | RJON | -4636.4297 | 0.16256 | 1 | -4628.8209 |
| **CLNK** | RHSPN12671 | RJON | -1872.5654 | 0.04049 | 1 | -1865.0234 |
| **EBAG9** | RHSPN16617 | RJON | -5033.6626 | 0.07728 | 1 | -5026.1393 |
| **NOX4** | RHSPN09445 | RJON | -11448.433 | 0.04878 | 1 | -11440.906 |
| **BAG6** | RHSPN12437 | RJON | -8511.9692 | 0.04651 | 1 | -8504.4529 |
| **KCNB2** | RHSPN02243 | RJON | -8088.0427 | 0.1175 | 1 | -8080.532 |
| **C1orf27** | RHSPN17454 | RJON | -33934.729 | 0.09355 | 1 | -33927.238 |
| **LRRK2** | RHSPN06107 | RJON | -8671.9046 | 0.10237 | 1 | -8664.4638 |
| **KY** | RHSPN23207 | RJON | -3300.1704 | 0.01429 | 1 | -3292.7463 |
| **SEPT4** | RHSPN17555 | RJON | -5945.3809 | 0.09673 | 1 | -5937.9688 |
| **DNAJC22** | RHSPN01216 | RJON | -3492.7747 | 0.09503 | 1 | -3485.3726 |
| **GBP1** | RHSPN12766 | RJON | -9274.8806 | 0.02521 | 1 | -9267.5502 |
| **ASAP1** | RHSPN22283 | RJON | -8527.3692 | 0.04694 | 1 | -8520.0447 |
| **DHX33** | RHSPN02209 | RJON | -2224.0088 | 0.08451 | 1 | -2216.6921 |
| **RGS16** | RHSPN00550 | RJON | -8134.4706 | 0.0368 | 1 | -8127.1637 |
| **EPB41** | RHSPN08345 | RJON | -2104.0229 | 0.15511 | 1 | -2096.7932 |
| **GLRX** | RHSPN08339 | RJON | -27156.589 | 0.11455 | 1 | -27149.388 |
| **TTC37** | RHSPN00542 | RJON | -1759.0022 | 0.02424 | 1 | -1751.8053 |
| **TAF12** | RHSPN17780 | RJON | -36149.445 | 0.02138 | 1 | -36142.267 |
| **LRP1** | RHSPN17215 | RJON | -801.26211 | 0.01968 | 1 | -794.1232 |
| **KLRF1** | RHSPN23860 | RJON | -14690.99 | 0.1252 | 1 | -14683.88 |
| **DSG4** | RHSPN15081 | RJON | -7559.2905 | 0.04026 | 1 | -7552.1917 |
| **CNNM1** | RHSPN11521 | RJON | -10743.518 | 0.16163 | 1 | -10736.448 |
| **POT1** | RHSPN23530 | RJON | -7146.9766 | 0.05005 | 1 | -7139.9119 |
| **PGS1** | RHSPN04257 | RJON | -2529.6856 | 0.06128 | 1 | -2522.6247 |
| **HOXD12** | RHSPN07231 | RJON | -15194.222 | 0.0622 | 1 | -15187.182 |
| **HERC3** | RHSPN27809 | RJON | -150.87355 | 0.00412 | 1 | -143.85126 |
| **SYN1** | RHSPN20679 | RJON | -4995.215 | 0.01369 | 1 | -4988.2091 |
| **PARP6** | RHSPN10209 | RJON | -2236.0364 | 0.12701 | 1 | -2229.0532 |
| **AKAP7** | RHSPN24386 | RJON | -7693.4499 | 0.12982 | 1 | -7686.4868 |
| **ADGRE1** | RHSPN07901 | RJON | -7243.1259 | 0.06555 | 1 | -7236.171 |
| **ZNF622** | RHSPN04248 | RJON | -6632.552 | 0.03746 | 1 | -6625.6266 |
| **CHRNA1** | RHSPN26424 | RJON | -4928.8791 | 0.13202 | 1 | -4921.9606 |
| **FAM217B** | RHSPN05479 | RJON | -7230.245 | 0.07402 | 1 | -7223.3279 |
| **ZMYND10** | RHSPN07683 | RJON | -6008.4826 | 0.10029 | 1 | -6001.623 |
| **AADAT** | RHSPN04517 | RJON | -10342.772 | 0.11567 | 1 | -10335.976 |
| **PLEKHM3** | RHSPN24251 | RJON | -3990.8825 | 0.06712 | 1 | -3984.1019 |
| **SCAMP4** | RHSPN03945 | RJON | -20719.675 | 0.05281 | 1 | -20712.897 |
| **UGGT1** | RHSPN09467 | RJON | -11250.758 | 0.10544 | 1 | -11244.041 |
| **C2** | RHSPN23870 | RJON | -6491.5092 | 0.03125 | 1 | -6484.7962 |
| **GAREM** | RHSPN23468 | RJON | -8421.8104 | 0.05966 | 1 | -8415.1041 |
| **RNF157** | RHSPN07216 | RJON | -6702.3487 | 0.1459 | 1 | -6695.6968 |

| | | | | | | |
|---|---|---|---|---|---|---|
| **SPARCL1** | RHSPN08190 | RJON | -5979.0023 | 0.17634 | 1 | -5972.3753 |
| **PTCD2** | RHSPN28373 | RJON | -9676.196 | 0.11272 | 1 | -9669.5671 |
| **ADGRG4** | RHSPN26985 | RJON | -11773.31 | 0.05005 | 1 | -11766.688 |
| **KLHL22** | RHSPN05772 | RJON | -14906.747 | 0.07028 | 1 | -14900.148 |
| **FILIP1L** | RHSPN21546 | RJON | -1882.832 | 0.04683 | 1 | -1876.2556 |
| **DCTPP1** | RHSPN05924 | RJON | -2190.4218 | 0.04191 | 1 | -2183.8794 |
| **FSTL1** | RHSPN18339 | RJON | -2495.0542 | 0.03155 | 1 | -2488.5225 |
| **ACADS** | RHSPN06957 | RJON | -5917.7378 | 0.09924 | 1 | -5911.2118 |
| **TXK** | RHSPN23861 | RJON | -17312.195 | 0.12294 | 1 | -17305.672 |
| **DSG2** | RHSPN04698 | RJON | -16522.083 | 0.04169 | 1 | -16515.574 |
| **COL4A4** | RHSPN07088 | RJON | -20678.488 | 0.03391 | 1 | -20671.989 |
| **ANKRD17** | RHSPN21207 | RJON | -12544.841 | 0.04925 | 1 | -12538.364 |
| **ADCY9** | RHSPN25610 | RJON | -1571.6038 | 0.05449 | 1 | -1565.1324 |
| **TTYH1** | RHSPN22448 | RJON | -3429.8697 | 0.06314 | 1 | -3423.4366 |
| **ADORA2B** | RHSPN08229 | RJON | -1589.3388 | 0.10693 | 1 | -1582.9011 |
| **S100Z** | RHSPN05438 | RJON | -1523.7703 | 0.07672 | 1 | -1517.331 |
| **TCTA** | RHSPN16843 | RJON | -15324.374 | 0.03989 | 1 | -15317.94 |
| **UBE4A** | RHSPN24616 | RJON | -6769.3649 | 0.1206 | 1 | -6762.95 |
| **ADGRE3** | RHSPN00209 | RJON | -21682.951 | 0.02431 | 1 | -21676.534 |
| **KIF1B** | RHSPN27760 | RJON | -3691.1233 | 0.12776 | 1 | -3684.7098 |
| **CXorf38** | RHSPN07674 | RJON | -8617.7207 | 0.06837 | 1 | -8611.3142 |
| **SH3RF1** | RHSPN00203 | RJON | -14985.337 | 0.04874 | 1 | -14978.932 |
| **CLSTN1** | RHSPN05276 | RJON | -5023.6661 | 0.16347 | 1 | -5017.2699 |
| **ULK4** | RHSPN00154 | RJON | -4859.7823 | 0.06978 | 1 | -4853.3971 |
| **RNF207** | RHSPN13613 | RJON | -7904.3735 | 0.08242 | 1 | -7897.9942 |
| **TBC1D2** | RHSPN19606 | RJON | -2994.4907 | 0.06552 | 1 | -2988.1651 |
| **RAD51B** | RHSPN25660 | RJON | -2127.5672 | 0.01364 | 1 | -2121.2419 |
| **UBE2S** | RHSPN20344 | RJON | -12032.979 | 0.09286 | 1 | -12026.675 |
| **TTBK2** | RHSPN05169 | RJON | -2613.8227 | 0.14175 | 1 | -2607.5289 |
| **CMTM7** | RHSPN05334 | RJON | -9026.5723 | 0.05619 | 1 | -9020.3032 |
| **SLC6A20** | RHSPN26403 | RJON | -3170.1264 | 0.16381 | 1 | -3163.8679 |
| **ANKRD60** | RHSPN09019 | RJON | -8226.7307 | 0.04369 | 1 | -8220.487 |
| **RUFY1** | RHSPN00798 | RJON | -9620.2047 | 0.04095 | 1 | -9613.9743 |
| **KDM4A** | RHSPN24259 | RJON | -2175.9335 | 0.01978 | 1 | -2169.7145 |
| **SF3A2** | RHSPN24710 | RJON | -1850.4995 | 0.0201 | 1 | -1844.2849 |
| **JUND** | RHSPN22307 | RJON | -1800.5662 | 0.08331 | 1 | -1794.3642 |
| **C17orf49** | RHSPN24094 | RJON | -6564.2756 | 0.15506 | 1 | -6558.0752 |
| **SERPINB2** | RHSPN10062 | RJON | -1843.0488 | 0.16537 | 1 | -1836.8591 |
| **LOC1009966** | RHSPN19023 | RJON | -17504.65 | 0.0375 | 1 | -17498.459 |
| **COL4A1** | RHSPN23914 | RJON | -14925.104 | 0.05193 | 1 | -14918.936 |
| **FHOD3** | RHSPN13521 | RJON | -12499.463 | 0.12325 | 1 | -12493.318 |
| **NOL8** | RHSPN05740 | RJON | -4067.9079 | 0.06014 | 1 | -4061.7843 |
| **ARL13B** | RHSPN26755 | RJON | -11617.677 | 0.09792 | 1 | -11611.561 |
| **MX1** | RHSPN10662 | RJON | -3424.8378 | 0.04673 | 1 | -3418.7637 |
| **TSPAN13** | RHSPN16513 | RJON | -6552.9289 | 0.07867 | 1 | -6546.8645 |

| | | | | | | |
|---|---|---|---|---|---|---|
| **GDPD5** | RHSPN24193 | RJON | -5077.8037 | 0.12538 | 1 | -5071.7437 |
| **BSG** | RHSPN18663 | RJON | -1396.6712 | 0.02681 | 1 | -1390.6234 |
| **SERTM1** | RHSPN13796 | RJON | -9766.7381 | 0.15004 | 1 | -9760.7131 |
| **CNTRL** | RHSPN22000 | RJON | -11153.315 | 0.03913 | 1 | -11147.324 |
| **PLCG2** | RHSPN17791 | RJON | -13052.317 | 0.07094 | 1 | -13046.337 |
| **GLI1** | RHSPN09958 | RJON | -5401.6894 | 0.04995 | 1 | -5395.7203 |
| **RNGTT** | RHSPN11455 | RJON | -3210.0935 | 0.02661 | 1 | -3204.124 |
| **DOCK4** | RHSPN24628 | RJON | -4578.3453 | 0.13072 | 1 | -4572.3794 |
| **CASP14** | RHSPN16328 | RJON | -7285.4303 | 0.09226 | 1 | -7279.5006 |
| **RIN1** | RHSPN11726 | RJON | -14100.07 | 0.07717 | 1 | -14094.155 |
| **EPHA1** | RHSPN23470 | RJON | -4313.0406 | 0.10247 | 1 | -4307.1259 |
| **QRICH2** | RHSPN20781 | RJON | -4906.1802 | 0.15945 | 1 | -4900.2789 |
| **SH2D7** | RHSPN14953 | RJON | -5468.618 | 0.17048 | 1 | -5462.7777 |
| **FAS** | RHSPN09756 | RJON | -5979.9673 | 0.10974 | 1 | -5974.1834 |
| **CD2AP** | RHSPN18028 | RJON | -5441.565 | 0.0508 | 1 | -5435.7986 |
| **PLXNC1** | RHSPN02482 | RJON | -7736.1257 | 0.0547 | 1 | -7730.3884 |
| **NEK2** | RHSPN08680 | RJON | -12332.021 | 0.05012 | 1 | -12326.286 |
| **DIAPH1** | RHSPN23850 | RJON | -4966.535 | 0.05282 | 1 | -4960.8034 |
| **AQP4** | RHSPN04086 | RJON | -1951.2513 | 0.01938 | 1 | -1945.5353 |
| **ARL5A** | RHSPN15596 | RJON | -4987.6026 | 0.08367 | 1 | -4981.912 |
| **OR52R1** | RHSPN00647 | RJON | -13305.301 | 0.09787 | 1 | -13299.637 |
| **NCDN** | RHSPN24400 | RJON | -15503.188 | 0.02845 | 1 | -15497.537 |
| **PNPLA6** | RHSPN14775 | RJON | -30533.415 | 0.05096 | 1 | -30527.794 |
| **CDH23** | RHSPN18326 | RJON | -4963.5398 | 0.06194 | 1 | -4957.9182 |
| **SIRT4** | RHSPN16529 | RJON | -13021.094 | 0.0541 | 1 | -13015.48 |
| **C11orf30** | RHSPN13996 | RJON | -1263.294 | 0.10792 | 1 | -1257.7317 |
| **QRFP** | RHSPN04096 | RJON | -669.05678 | 0.00243 | 1 | -663.48611 |
| **RPRM** | RHSPN15954 | RJON | -6174.496 | 0.02469 | 1 | -6168.9381 |
| **CHRM4** | RHSPN17109 | RJON | -4377.9083 | 0.01722 | 1 | -4372.3523 |
| **KCNA6** | RHSPN28227 | RJON | -8688.1398 | 0.01472 | 1 | -8682.5928 |
| **PLS3** | RHSPN28484 | RJON | -3050.819 | 0.02987 | 1 | -3045.2712 |
| **PNCK** | RHSPN01691 | RJON | -11515.361 | 0.04888 | 1 | -11509.833 |
| **CGN** | RHSPN18332 | RJON | -4371.1892 | 0.05352 | 1 | -4365.6694 |
| **COQ5** | RHSPN00704 | RJON | -926.61129 | 0.10346 | 1 | -921.09076 |
| **MACF1** | RHSPN24205 | RJON | -5339.2896 | 0.03712 | 1 | -5333.7853 |
| **KISS1R** | RHSPN09163 | RJON | -1441.4681 | 0.04164 | 1 | -1435.9784 |
| **TMEM170B** | RHSPN00505 | RJON | -1922.1708 | 0.06152 | 1 | -1916.6753 |
| **TRNP1** | RHSPN04582 | RJON | -11443.336 | 0.10519 | 1 | -11437.845 |
| **XRCC5** | RHSPN05251 | RJON | -18901.441 | 0.11495 | 1 | -18895.947 |
| **TTC21A** | RHSPN03961 | RJON | -4613.7747 | 0.1104 | 1 | -4608.3199 |
| **PTPN18** | RHSPN11470 | RJON | -2806.7935 | 0.07435 | 1 | -2801.3605 |
| **MDFIC** | RHSPN01880 | RJON | -5392.3884 | 0.04499 | 1 | -5386.9601 |
| **HAPLN2** | RHSPN09606 | RJON | -2266.3968 | 0.14381 | 1 | -2260.9609 |
| **CCDC167** | RHSPN01709 | RJON | -3005.1234 | 0.23102 | 1 | -2999.6938 |
| **CRNN** | RHSPN24318 | RJON | -7618.9367 | 0.06642 | 1 | -7613.5056 |

| | | | | | | |
|---|---|---|---|---|---|---|
| **CHAF1A** | RHSPN05959 | RJON | -11356.354 | 0.03241 | 1 | -11350.919 |
| **SEMA5B** | RHSPN09029 | RJON | -4827.5006 | 0.07388 | 1 | -4822.0916 |
| **SQSTM1** | RHSPN08631 | RJON | -1198.957 | 0.02095 | 1 | -1193.5418 |
| **NRG2** | RHSPN19750 | RJON | -4060.885 | 0.08052 | 1 | -4055.4722 |
| **ALKBH1** | RHSPN14894 | RJON | -2764.5216 | 0.07235 | 1 | -2759.1114 |
| **SH2D4B** | RHSPN02648 | RJON | -11072.654 | 0.10439 | 1 | -11067.27 |
| **OBSCN** | RHSPN23645 | RJON | -2085.0482 | 0.12258 | 1 | -2079.6623 |
| **CD7** | RHSPN25477 | RJON | -6303.2159 | 0.13844 | 1 | -6297.8517 |
| **VSIG10L** | RHSPN11309 | RJON | -7441.9932 | 0.02819 | 1 | -7436.6426 |
| **TAF6** | RHSPN28478 | RJON | -4105.7906 | 0.02636 | 1 | -4100.4508 |
| **BGN** | RHSPN08737 | RJON | -771.33699 | 0.08917 | 1 | -766.0035 |
| **SPINK5** | RHSPN09512 | RJON | -7455.3163 | 0.05617 | 1 | -7449.9928 |
| **RGL2** | RHSPN28189 | RJON | -1586.1604 | 0.09989 | 1 | -1580.85 |
| **KCNE5** | RHSPN20215 | RJON | -3725.551 | 0.02844 | 1 | -3720.2481 |
| **CHRNA7** | RHSPN16733 | RJON | -8078.1935 | 0.07085 | 1 | -8072.8955 |
| **KDELC2** | RHSPN22972 | RJON | -13699.624 | 0.0624 | 1 | -13694.339 |
| **SLC4A1** | RHSPN00106 | RJON | -11958.773 | 0.04212 | 1 | -11953.511 |
| **PLCH2** | RHSPN19470 | RJON | -5342.0454 | 0.03427 | 1 | -5336.7986 |
| **SOCS4** | RHSPN24629 | RJON | -12397.566 | 0.06575 | 1 | -12392.317 |
| **ILVBL** | RHSPN01491 | RJON | -6697.5676 | 0.09768 | 1 | -6692.3184 |
| **WARS2** | RHSPN27017 | RJON | -2372.1303 | 0.07053 | 1 | -2366.894 |
| **YDJC** | RHSPN28119 | RJON | -2641.7438 | 0.12616 | 1 | -2636.5333 |
| **ARMCX3** | RHSPN13006 | RJON | -4380.8786 | 0.13147 | 1 | -4375.6879 |
| **CD274** | RHSPN20795 | RJON | -4570.8131 | 0.04318 | 1 | -4565.6399 |
| **CHRNB4** | RHSPN21941 | RJON | -27598.595 | 0.03377 | 1 | -27593.426 |
| **ZFHX3** | RHSPN26813 | RJON | -19732.666 | 0.05318 | 1 | -19727.509 |
| **TRAPPC10** | RHSPN10140 | RJON | -6098.8722 | 0.0648 | 1 | -6093.7184 |
| **MCM9** | RHSPN17633 | RJON | -5570.2374 | 0.05688 | 1 | -5565.0867 |
| **KRT6A** | RHSPN21145 | RJON | -11793.782 | 0.08574 | 1 | -11788.64 |
| **CCNF** | RHSPN11819 | RJON | -11196.16 | 0.03521 | 1 | -11191.018 |
| **CHPF2** | RHSPN03175 | RJON | -2649.2593 | 0.1263 | 1 | -2644.1211 |
| **GEMIN6** | RHSPN27808 | RJON | -7901.6096 | 0.0346 | 1 | -7896.4789 |
| **ARAF** | RHSPN09884 | RJON | -17143.995 | 0.04371 | 1 | -17138.877 |
| **FILIP1** | RHSPN01012 | RJON | -4548.5899 | 0.1006 | 1 | -4543.4747 |
| **TACSTD2** | RHSPN27963 | RJON | -218.63071 | 0 | 1 | -213.52886 |
| **LAS1L** | RHSPN11776 | RJON | -38900.486 | 0.08228 | 1 | -38895.409 |
| **SSPO** | RHSPN00537 | RJON | -8314.1117 | 0.0828 | 1 | -8309.0366 |
| **PHACTR4** | RHSPN12557 | RJON | -8247.4516 | 0.07182 | 1 | -8242.3732 |
| **DPY19L4** | RHSPN06160 | RJON | -9725.9536 | 0.06617 | 1 | -9720.9119 |
| **RASA2** | RHSPN22700 | RJON | -3072.4749 | 0.16143 | 1 | -3067.44 |
| **FNDC8** | RHSPN12081 | RJON | -4271.4714 | 0.05921 | 1 | -4266.4372 |
| **DMTN** | RHSPN11856 | RJON | -1885.6865 | 0.04978 | 1 | -1880.6544 |
| **HTR5A** | RHSPN06160 | RJON | -9738.0829 | 0.06657 | 1 | -9733.0574 |
| **RASA2** | RHSPN21533 | RJON | -4842.6767 | 0.02023 | 1 | -4837.659 |
| **ALDOA** | RHSPN09207 | RJON | -3357.707 | 0.12535 | 1 | -3352.6969 |

| | | | | | | |
|---|---|---|---|---|---|---|
| **NHLRC1** | RHSPN27691 | RJON | -10063.817 | 0.03726 | 1 | -10058.809 |
| **CNKSR2** | RHSPN20437 | RJON | -11737.032 | 0.08295 | 1 | -11732.025 |
| **ATP8B4** | RHSPN08052 | RJON | -8464.1308 | 0.09238 | 1 | -8459.119 |
| **ITGA1** | RHSPN23220 | RJON | -13892.402 | 0.02669 | 1 | -13887.394 |
| **SMG8** | RHSPN16393 | RJON | -2100.0725 | 0.13218 | 1 | -2095.0709 |
| **C11orf24** | RHSPN17080 | RJON | -22723.323 | 0.02758 | 1 | -22718.324 |
| **CACNA1C** | RHSPN25434 | RJON | -13850.633 | 0.04302 | 1 | -13845.634 |
| **MYBPC2** | RHSPN03182 | RJON | -6384.3328 | 0.03026 | 1 | -6379.3341 |
| **MAP4K3** | RHSPN28486 | RJON | -7710.1683 | 0.03305 | 1 | -7705.208 |
| **SLC6A8** | RHSPN17702 | RJON | -1363.8863 | 0.08991 | 1 | -1358.9363 |
| **PPP1R1A** | RHSPN01096 | RJON | -3718.9187 | 0.04035 | 1 | -3713.9782 |
| **RPE65** | RHSPN14421 | RJON | -5824.2433 | 0.04931 | 1 | -5819.3028 |
| **ARMC4** | RHSPN10826 | RJON | -594.13493 | 0.21261 | 1 | -589.21178 |
| **TARP** | RHSPN20409 | RJON | -7503.8692 | 0.07044 | 1 | -7498.9574 |
| **SLC24A5** | RHSPN03406 | RJON | -8896.272 | 0.113 | 1 | -8891.3603 |
| **PCYOX1** | RHSPN25811 | RJON | -3085.0211 | 0.25775 | 1 | -3080.1181 |
| **SIRPA** | RHSPN12168 | RJON | -19694.712 | 0.04605 | 1 | -19689.811 |
| **KIF13B** | RHSPN15609 | RJON | -4179.1678 | 0.1009 | 1 | -4174.2658 |
| **OR51I1** | RHSPN05533 | RJON | -34725.596 | 0.09707 | 1 | -34720.706 |
| **STAB1** | RHSPN13470 | RJON | -16635.397 | 0.06904 | 1 | -16630.509 |
| **ZCCHC6** | RHSPN18420 | RJON | -14921.812 | 0.06038 | 1 | -14916.93 |
| **NCOR2** | RHSPN20921 | RJON | -4797.2602 | 0.01446 | 1 | -4792.3757 |
| **IDH2** | RHSPN07276 | RJON | -3923.3953 | 0.16426 | 1 | -3918.5281 |
| **C4orf17** | RHSPN17631 | RJON | -7408.5459 | 0.10428 | 1 | -7403.6753 |
| **KRT84** | RHSPN22019 | RJON | -7744.6946 | 0.11013 | 1 | -7739.8606 |
| **ADAD2** | RHSPN17615 | RJON | -18519.499 | 0.0238 | 1 | -18514.668 |
| **SCN8A** | RHSPN05547 | RJON | -15889.361 | 0.1081 | 1 | -15884.538 |
| **ITIH3** | RHSPN15725 | RJON | -5811.7308 | 0.03686 | 1 | -5806.9096 |
| **RASSF10** | RHSPN00684 | RJON | -3753.271 | 0.03358 | 1 | -3748.4717 |
| **MANEAL** | RHSPN04805 | RJON | -4239.3394 | 0.0862 | 1 | -4234.5509 |
| **IQCA1** | RHSPN14346 | RJON | -3750.9487 | 0.14626 | 1 | -3746.1588 |
| **MALRD1** | RHSPN04569 | RJON | -4678.9235 | 0.13743 | 1 | -4674.1408 |
| **BARD1** | RHSPN14800 | RJON | -1931.9587 | 0.0677 | 1 | -1927.1756 |
| **MRPS16** | RHSPN13896 | RJON | -3020.6365 | 0.10034 | 1 | -3015.8569 |
| **TOR2A** | RHSPN19451 | RJON | -5347.1462 | 0.06689 | 1 | -5342.3811 |
| **PTGER2** | RHSPN03322 | RJON | -8662.1357 | 0.04893 | 1 | -8657.3788 |
| **PAPOLG** | RHSPN23329 | RJON | -972.23199 | 0.03478 | 1 | -967.48411 |
| **PITPNC1** | RHSPN21261 | RJON | -10441.404 | 0.06849 | 1 | -10436.663 |
| **GRIN2A** | RHSPN15076 | RJON | -11397.513 | 0.0809 | 1 | -11392.78 |
| **HPS1** | RHSPN24611 | RJON | -4248.6017 | 0.03306 | 1 | -4243.8686 |
| **DNAJB1** | RHSPN14546 | RJON | -1001.2939 | 0.02828 | 1 | -996.57234 |
| **CXCL12** | RHSPN16574 | RJON | -2013.343 | 0.11078 | 1 | -2008.6213 |
| **ANKRD42** | RHSPN10429 | RJON | -6065.9328 | 0.11876 | 1 | -6061.2205 |
| **PNLDC1** | RHSPN00414 | RJON | -12425.749 | 0.03148 | 1 | -12421.054 |
| **EPHA8** | RHSPN15742 | RJON | -515.10695 | 0.12483 | 1 | -510.41718 |

| | | | | | | |
|---|---|---|---|---|---|---|
| **INSC** | RHSPN22247 | RJON | -4424.1475 | 0.02274 | 1 | -4419.4566 |
| **ARRB2** | RHSPN25380 | RJON | -4400.4579 | 0.01337 | 1 | -4395.7739 |
| **SLC17A7** | RHSPN00680 | RJON | -2371.3998 | 0.11273 | 1 | -2366.7173 |
| **C1orf109** | RHSPN04066 | RJON | -2830.4103 | 0.08429 | 1 | -2825.7454 |
| **LYPD6B** | RHSPN18727 | RJON | -41.682446 | 0.01069 | 1 | -37.020357 |
| **TSC22D1** | RHSPN09202 | RJON | -2008.7677 | 0.02271 | 1 | -2004.1001 |
| **FAM8A1** | RHSPN08738 | RJON | -1533.4746 | 0.10779 | 1 | -1528.8077 |
| **SPINK6** | RHSPN18935 | RJON | -4311.8767 | 0.09876 | 1 | -4307.2248 |
| **GPC5** | RHSPN05586 | RJON | -8115.5394 | 0.03707 | 1 | -8110.9007 |
| **APPL1** | RHSPN09285 | RJON | -9154.6099 | 0.18791 | 1 | -9149.9795 |
| **BTN1A1** | RHSPN15695 | RJON | -12781.448 | 0.05048 | 1 | -12776.819 |
| **SCUBE2** | RHSPN07451 | RJON | -10040.81 | 0.05777 | 1 | -10036.188 |
| **HSPA4L** | RHSPN17615 | RJON | -18555.311 | 0.02392 | 1 | -18550.687 |
| **SCN8A** | RHSPN25441 | RJON | -4814.4416 | 0.02077 | 1 | -4809.8196 |
| **SYT3** | RHSPN10036 | RJON | -3287.203 | 0.1187 | 1 | -3282.583 |
| **C6orf203** | RHSPN06439 | RJON | -3344.3464 | 0.06964 | 1 | -3339.7349 |
| **FAM131A** | RHSPN01138 | RJON | -2351.2383 | 0.01766 | 1 | -2346.6273 |
| **AK5** | RHSPN11122 | RJON | -2495.8373 | 0.05784 | 1 | -2491.231 |
| **HSPB1** | RHSPN07640 | RJON | -8359.2566 | 0.05221 | 1 | -8354.6654 |
| **FSTL5** | RHSPN27242 | RJON | -8587.2886 | 0.05451 | 1 | -8582.7021 |
| **SLC5A1** | RHSPN09026 | RJON | -6712.2323 | 0.05484 | 1 | -6707.6596 |
| **CANX** | RHSPN23602 | RJON | -3814.4014 | 0.1103 | 1 | -3809.8259 |
| **C17orf70** | RHSPN16082 | RJON | -3916.6886 | 0.12824 | 1 | -3912.1124 |
| **CNTF** | RHSPN16231 | RJON | -2827.3483 | 0.02719 | 1 | -2822.7798 |
| **GPR137** | RHSPN19241 | RJON | -2131.7869 | 0.13001 | 1 | -2127.2259 |
| **IL25** | RHSPN08719 | RJON | -2662.9452 | 0.07361 | 1 | -2658.3847 |
| **GRXCR2** | RHSPN21733 | RJON | -6200.5425 | 0.07408 | 1 | -6195.9852 |
| **ADGRG5** | RHSPN05795 | RJON | -7066.7376 | 0.1218 | 1 | -7062.1836 |
| **TRMT10C** | RHSPN14829 | RJON | -3471.8329 | 0.01864 | 1 | -3467.2964 |
| **AP3M1** | RHSPN21414 | RJON | -5916.646 | 0.10229 | 1 | -5912.1147 |
| **SDR42E2** | RHSPN09858 | RJON | -4090.1186 | 0.01603 | 1 | -4085.5872 |
| **OGFRL1** | RHSPN24608 | RJON | -7614.7058 | 0.03785 | 1 | -7610.1719 |
| **PKN1** | RHSPN25323 | RJON | -7122.7702 | 0.1176 | 1 | -7118.242 |
| **CCDC114** | RHSPN12348 | RJON | -4180.0914 | 0.05984 | 1 | -4175.5789 |
| **IMPAD1** | RHSPN06272 | RJON | -6403.7756 | 0.06062 | 1 | -6399.2718 |
| **VEPH1** | RHSPN12837 | RJON | -2376.1533 | 0.10536 | 1 | -2371.6589 |
| **TSNARE1** | RHSPN25135 | RJON | -4901.1323 | 0.0273 | 1 | -4896.6591 |
| **ERF** | RHSPN22594 | RJON | -4812.6412 | 0.05687 | 1 | -4808.1635 |
| **VTN** | RHSPN12084 | RJON | -7540.2088 | 0.06659 | 1 | -7535.7356 |
| **FAM160B2** | RHSPN04863 | RJON | -6284.5299 | 0.12968 | 1 | -6280.0559 |
| **GPR35** | RHSPN04156 | RJON | -6976.013 | 0.08693 | 1 | -6971.5523 |
| **FAP** | RHSPN09944 | RJON | -1419.7371 | 0.0653 | 1 | -1415.2921 |
| **SMIM8** | RHSPN23733 | RJON | -6278.346 | 0.03666 | 1 | -6273.8986 |
| **PTPRM** | RHSPN11310 | RJON | -2870.9459 | 0.05522 | 1 | -2866.5013 |
| **CNPY4** | RHSPN13117 | RJON | -2986.52 | 0.27824 | 1 | -2982.0838 |

| | | | | | | |
|---|---|---|---|---|---|---|
| **IFNK** | RHSPN11643 | RJON | -8104.316 | 0.04536 | 1 | -8099.8944 |
| **ATP6V0A4** | RHSPN25002 | RJON | -4287.4188 | 0.0448 | 1 | -4282.9997 |
| **YIF1B** | RHSPN24441 | RJON | -7267.1734 | 0.15464 | 1 | -7262.7637 |
| **MUC16** | RHSPN00423 | RJON | -12944.011 | 0.08019 | 1 | -12939.604 |
| **LUZP1** | RHSPN02342 | RJON | -1358.0503 | 0.03899 | 1 | -1353.6617 |
| **RABIF** | RHSPN19743 | RJON | -4069.5081 | 0.06823 | 1 | -4065.1229 |
| **GSTZ1** | RHSPN15634 | RJON | -1767.5387 | 0.04736 | 1 | -1763.1572 |
| **APBB1** | RHSPN02438 | RJON | -6396.3867 | 0.0938 | 1 | -6392.0061 |
| **C4BPA** | RHSPN06537 | RJON | -8196.1363 | 0.03928 | 1 | -8191.756 |
| **ACAP2** | RHSPN18794 | RJON | -4373.6161 | 0.06314 | 1 | -4369.2379 |
| **FAM124A** | RHSPN25229 | RJON | -4317.9639 | 0.02375 | 1 | -4313.5893 |
| **CKM** | RHSPN12087 | RJON | -12670.976 | 0.12362 | 1 | -12666.603 |
| **HR** | RHSPN13251 | RJON | -1957.3576 | 0.06907 | 1 | -1952.9995 |
| **EXOSC3** | RHSPN02368 | RJON | -1563.8944 | 0.0585 | 1 | -1559.529 |
| **GOLT1A** | RHSPN27976 | RJON | -3544.7475 | 0.05896 | 1 | -3540.3906 |
| **YIPF6** | RHSPN05712 | RJON | -19280.251 | 0.03979 | 1 | -19275.892 |
| **ROBO1** | RHSPN03956 | RJON | -9740.6527 | 0.1012 | 1 | -9736.2969 |
| **SMPD4** | RHSPN21854 | RJON | -7212.8992 | 0.06848 | 1 | -7208.5556 |
| **LCAT** | RHSPN07663 | RJON | -4016.1622 | 0.07542 | 1 | -4011.8167 |
| **ANXA10** | RHSPN04872 | RJON | -14574.032 | 0.06934 | 1 | -14569.689 |
| **SNED1** | RHSPN02145 | RJON | -3618.4898 | 0.05862 | 1 | -3614.168 |
| **CACYBP** | RHSPN12261 | RJON | -8336.2784 | 0.04498 | 1 | -8331.9591 |
| **ANK1** | RHSPN19058 | RJON | -9945.2009 | 0.09619 | 1 | -9940.8912 |
| **F7** | RHSPN22753 | RJON | -1133.9841 | 0.08119 | 1 | -1129.6915 |
| **C17orf78** | RHSPN27133 | RJON | -2314.0585 | 0.05396 | 1 | -2309.7693 |
| **CRYBA4** | RHSPN24478 | RJON | -2146.0638 | 0.05903 | 1 | -2141.7891 |
| **P2RY11** | RHSPN08116 | RJON | -2767.5749 | 0.23183 | 1 | -2763.3145 |
| **LRRC70** | RHSPN03485 | RJON | -6893.9255 | 0.02292 | 1 | -6889.6641 |
| **LRRTM4** | RHSPN11815 | RJON | -6301.3898 | 0.09031 | 1 | -6297.1331 |
| **ASB10** | RHSPN07992 | RJON | -2732.41 | 0.08122 | 1 | -2728.152 |
| **GDNF** | RHSPN25408 | RJON | -5209.8263 | 0.06967 | 1 | -5205.5718 |
| **FUZ** | RHSPN01472 | RJON | -15289.944 | 0.05304 | 1 | -15285.699 |
| **IGSF3** | RHSPN26067 | RJON | -10091.355 | 0.01801 | 1 | -10087.111 |
| **KIF3B** | RHSPN13616 | RJON | -11399.865 | 0.08692 | 1 | -11395.62 |
| **ANKS6** | RHSPN00151 | RJON | -3449.0637 | 0.01271 | 1 | -3444.823 |
| **KCNAB2** | RHSPN17090 | RJON | -2632.2555 | 0.17468 | 1 | -2628.0209 |
| **RHNO1** | RHSPN03086 | RJON | -1187.9072 | 0.03771 | 1 | -1183.6774 |
| **FOSL2** | RHSPN06842 | RJON | -10771.546 | 0.05517 | 1 | -10767.317 |
| **ANAPC4** | RHSPN12771 | RJON | -11655.547 | 0.03654 | 1 | -11651.32 |
| **EFR3A** | RHSPN21924 | RJON | -9887.7949 | 0.06617 | 1 | -9883.5704 |
| **PHLPP2** | RHSPN27454 | RJON | -2404.2893 | 0.01658 | 1 | -2400.0817 |
| **SULT4A1** | RHSPN03829 | RJON | -9926.0546 | 0.07538 | 1 | -9921.8501 |
| **MERTK** | RHSPN07377 | RJON | -20594.596 | 0.04304 | 1 | -20590.391 |
| **ANK2** | RHSPN24519 | RJON | -3323.4844 | 0.02302 | 1 | -3319.2781 |
| **LPPR2** | RHSPN05601 | RJON | -1485.8362 | 0.23534 | 1 | -1481.6342 |

| | | | | | | |
|---|---|---|---|---|---|---|
| **LOC1053776** | RHSPN21519 | RJON | -2624.5939 | 0.05976 | 1 | -2620.393 |
| **ASPHD1** | RHSPN12886 | RJON | -5620.0731 | 0.03221 | 1 | -5615.874 |
| **NRBP2** | RHSPN18267 | RJON | -2539.4996 | 0.06762 | 1 | -2535.3076 |
| **SLC8B1** | RHSPN22977 | RJON | -7673.0342 | 0.08549 | 1 | -7668.8414 |
| **ITGA2B** | RHSPN19161 | RJON | -4743.4678 | 0.06077 | 1 | -4739.2776 |
| **OR4E2** | RHSPN02785 | RJON | -1990.9005 | 0.0302 | 1 | -1986.7246 |
| **DESI2** | RHSPN03235 | RJON | -1460.8441 | 0.1352 | 1 | -1456.6819 |
| **TMEM247** | RHSPN18179 | RJON | -2410.408 | 0.05082 | 1 | -2406.2496 |
| **USP30** | RHSPN04657 | RJON | -11017.202 | 0.03589 | 1 | -11013.05 |
| **CHPF** | RHSPN12844 | RJON | -3730.4391 | 0.06239 | 1 | -3726.2835 |
| **THEM6** | RHSPN17510 | RJON | -5202.5662 | 0.03418 | 1 | -5198.4149 |
| **VDR** | RHSPN24228 | RJON | -4500.5501 | 0.05237 | 1 | -4496.3972 |
| **DAZAP1** | RHSPN18157 | RJON | -7367.1118 | 0.12289 | 1 | -7362.974 |
| **MTERF2** | RHSPN20474 | RJON | -14357.315 | 0.07205 | 1 | -14353.195 |
| **FAM214A** | RHSPN16375 | RJON | -3003.9528 | 0.08343 | 1 | -2999.8373 |
| **GSTP1** | RHSPN20288 | RJON | -5836.9396 | 0.04895 | 1 | -5832.8384 |
| **IVD** | RHSPN24495 | RJON | -11223.19 | 0.05002 | 1 | -11219.093 |
| **SLC44A2** | RHSPN21398 | RJON | -57874.326 | 0.05135 | 1 | -57870.23 |
| **DNAH3** | RHSPN00931 | RJON | -8765.6161 | 0.08724 | 1 | -8761.5217 |
| **EPS15** | RHSPN27621 | RJON | -17945.112 | 0.08921 | 1 | -17941.023 |
| **TLR8** | RHSPN14964 | RJON | -27734.207 | 0.12963 | 1 | -27730.123 |
| **KIF20B** | RHSPN16307 | RJON | -3245.7031 | 0.11015 | 1 | -3241.6261 |
| **RNASEH2C** | RHSPN09633 | RJON | -1803.1471 | 0.05703 | 1 | -1799.0691 |
| **OARD1** | RHSPN07015 | RJON | -1491.6284 | 0.08048 | 1 | -1487.5524 |
| **HOPX** | RHSPN14921 | RJON | -7090.0407 | 0.02655 | 1 | -7085.9676 |
| **BMPR1A** | RHSPN24071 | RJON | -10639.87 | 0.03661 | 1 | -10635.799 |
| **CDH20** | RHSPN00562 | RJON | -7862.1866 | 0.05789 | 1 | -7858.1164 |
| **MATN1** | RHSPN18979 | RJON | -7378.6716 | 0.08086 | 1 | -7374.6035 |
| **GPR183** | RHSPN09961 | RJON | -3148.791 | 0.08544 | 1 | -3144.7288 |
| **PM20D2** | RHSPN08776 | RJON | -4755.9686 | 0.03106 | 1 | -4751.9056 |
| **DCTN4** | RHSPN16057 | RJON | -658.69926 | 0.18231 | 1 | -654.64868 |
| **PRG2** | RHSPN28515 | RJON | -2884.9547 | 0.1048 | 1 | -2880.9047 |
| **ATP6AP1** | RHSPN21923 | RJON | -5165.4816 | 0.13647 | 1 | -5161.4298 |
| **MARVELD3** | RHSPN15206 | RJON | -9843.4957 | 0.05037 | 1 | -9839.4422 |
| **ADD3** | RHSPN19026 | RJON | -3414.1104 | 0.09726 | 1 | -3410.0688 |
| **CARKD** | RHSPN11705 | RJON | -3025.1833 | 0.06219 | 1 | -3021.1404 |
| **PRSS2** | RHSPN07836 | RJON | -12228.011 | 0.05663 | 1 | -12223.971 |
| **SLC6A19** | RHSPN03611 | RJON | -5462.3765 | 0.07695 | 1 | -5458.3405 |
| **MRPS5** | RHSPN05459 | RJON | -6303.6552 | 0.0292 | 1 | -6299.6202 |
| **MON1A** | RHSPN26332 | RJON | -9396.3333 | 0.09145 | 1 | -9392.3002 |
| **SPATA2** | RHSPN01777 | RJON | -8501.3793 | 0.12024 | 1 | -8497.3717 |
| **PBXIP1** | RHSPN15159 | RJON | -1917.9469 | 0.12934 | 1 | -1913.958 |
| **AS3MT** | RHSPN09216 | RJON | -1333.2666 | 0.03488 | 1 | -1329.2765 |
| **ID4** | RHSPN19479 | RJON | -20253.331 | 0.10106 | 1 | -20249.35 |
| **KTN1** | RHSPN27544 | RJON | -4645.5994 | 0.06838 | 1 | -4641.6249 |

| | | | | | | |
|---|---|---|---|---|---|---|
| **SCO2** | RHSPN03694 | RJON | -3944.7661 | 0.11106 | 1 | -3940.7895 |
| **LYG2** | RHSPN14391 | RJON | -9736.802 | 0.09307 | 1 | -9732.8344 |
| **GPR158** | RHSPN10158 | RJON | -1181.5123 | 0.17787 | 1 | -1177.5496 |
| **TRDN** | RHSPN25365 | RJON | -11606.233 | 0.06899 | 1 | -11602.276 |
| **TRPM4** | RHSPN27732 | RJON | -37.122819 | 0.00297 | 1 | -33.172699 |
| **DMD** | RHSPN12442 | RJON | -925.05021 | 0.00996 | 1 | -921.10912 |
| **RDH10** | RHSPN09838 | RJON | -1996.8951 | 0.10385 | 1 | -1992.9666 |
| **EYS** | RHSPN27768 | RJON | -2843.3815 | 0.09676 | 1 | -2839.4584 |
| **GPR34** | RHSPN24691 | RJON | -6603.9256 | 0.04892 | 1 | -6600.0239 |
| **FCHO1** | RHSPN26372 | RJON | -7444.9538 | 0.07212 | 1 | -7441.0559 |
| **CYP24A1** | RHSPN24100 | RJON | -10775.115 | 0.02305 | 1 | -10771.22 |
| **CDH7** | RHSPN09707 | RJON | -2872.3298 | 0.06674 | 1 | -2868.4774 |
| **MRPS18A** | RHSPN11796 | RJON | -6278.2561 | 0.15649 | 1 | -6274.4104 |
| **GIMAP4** | RHSPN25058 | RJON | -1381.1116 | 0.024 | 1 | -1377.2757 |
| **TTC9B** | RHSPN04400 | RJON | -1666.0947 | 0.04378 | 1 | -1662.267 |
| **HSPE1** | RHSPN08867 | RJON | -4612.5954 | 0.07687 | 1 | -4608.7693 |
| **CCNG1** | RHSPN16425 | RJON | -2042.2492 | 0.13462 | 1 | -2038.4304 |
| **FADD** | RHSPN22173 | RJON | -5936.6276 | 0.04577 | 1 | -5932.8105 |
| **SLC43A2** | RHSPN15295 | RJON | -6084.32 | 0.13783 | 1 | -6080.5062 |
| **SFXN4** | RHSPN10537 | RJON | -6464.0034 | 0.09829 | 1 | -6460.1972 |
| **CYP2W1** | RHSPN09616 | RJON | -2728.1089 | 0.08734 | 1 | -2724.3058 |
| **SAYSD1** | RHSPN15167 | RJON | -5252.996 | 0.05721 | 1 | -5249.2074 |
| **INA** | RHSPN10488 | RJON | -8719.1996 | 0.02853 | 1 | -8715.4142 |
| **MLLT4** | RHSPN20879 | RJON | -12720.745 | 0.11366 | 1 | -12716.966 |
| **AGBL1** | RHSPN18005 | RJON | -3941.8457 | 0.07989 | 1 | -3938.0775 |
| **EPYC** | RHSPN18606 | RJON | -18604.854 | 0.11338 | 1 | -18601.091 |
| **FLT1** | RHSPN05932 | RJON | -28636.702 | 0.09265 | 1 | -28632.944 |
| **POLQ** | RHSPN20611 | RJON | -14129.693 | 0.05611 | 1 | -14125.938 |
| **DIS3L** | RHSPN18145 | RJON | -9850.0707 | 0.04875 | 1 | -9846.318 |
| **APPL2** | RHSPN24367 | RJON | -4670.5771 | 0.10624 | 1 | -4666.8318 |
| **ACER1** | RHSPN02346 | RJON | -3023.9506 | 0.08861 | 1 | -3020.2127 |
| **TMEM183A** | RHSPN18594 | RJON | -5510.0077 | 0.16968 | 1 | -5506.2717 |
| **MTIF3** | RHSPN15862 | RJON | -9558.043 | 0.11324 | 1 | -9554.3081 |
| **TCP11L1** | RHSPN12100 | RJON | -9364.2687 | 0.07287 | 1 | -9360.5393 |
| **CCAR2** | RHSPN26498 | RJON | -2820.46 | 0.08932 | 1 | -2816.7327 |
| **TPD52L2** | RHSPN09085 | RJON | -7007.3225 | 0.1262 | 1 | -7003.596 |
| **SERPINB6** | RHSPN04789 | RJON | -10925.83 | 0.10773 | 1 | -10922.128 |
| **UGT1A1** | RHSPN21252 | RJON | -3322.9635 | 0.06579 | 1 | -3319.262 |
| **PMM2** | RHSPN27649 | RJON | -3914.8925 | 0.05297 | 1 | -3911.1998 |
| **GRPR** | RHSPN11213 | RJON | -9042.811 | 0.03675 | 1 | -9039.1159 |
| **KRIT1** | RHSPN12184 | RJON | -891.42392 | 0.11203 | 1 | -887.74316 |
| **TEX15** | RHSPN23425 | RJON | -8417.9142 | 0.06363 | 1 | -8414.2329 |
| **NUP85** | RHSPN08744 | RJON | -18538.163 | 0.14572 | 1 | -18534.483 |
| **SH3TC2** | RHSPN20676 | RJON | -8191.1467 | 0.0348 | 1 | -8187.465 |
| **PKM** | RHSPN09673 | RJON | -1460.439 | 0.09762 | 1 | -1456.7546 |

| | | | | | | |
|---|---|---|---|---|---|---|
| **MEA1** | RHSPN25477 | RJON | -2815.0896 | 0.01542 | 1 | -2811.4136 |
| **IGLON5** | RHSPN01024 | RJON | -11607.498 | 0.11671 | 1 | -11603.826 |
| **CYP2J2** | RHSPN02420 | RJON | -9432.7417 | 0.08139 | 1 | -9429.0712 |
| **IKBKE** | RHSPN08931 | RJON | -7082.1875 | 0.03843 | 1 | -7078.5212 |
| **DRD1** | RHSPN24698 | RJON | -5707.7625 | 0.06682 | 1 | -5704.1023 |
| **ARRDC2** | RHSPN27834 | RJON | -2661.6641 | 0.03082 | 1 | -2658.0079 |
| **GATA1** | RHSPN17423 | RJON | -1622.1132 | 0.0956 | 1 | -1618.4629 |
| **METTL20** | RHSPN05063 | RJON | -1231.4229 | 0.02483 | 1 | -1227.7728 |
| **MRPS25** | RHSPN27606 | RJON | -5244.1037 | 0.10541 | 1 | -5240.4719 |
| **GPR143** | RHSPN01096 | RJON | -7984.845 | 0.08891 | 1 | -7981.2121 |
| **DEPDC1** | RHSPN07073 | RJON | -2609.6211 | 0.14943 | 1 | -2605.9933 |
| **IGJ** | RHSPN19144 | RJON | -15896.627 | 0.08837 | 1 | -15893.001 |
| **ARHGEF40** | RHSPN02845 | RJON | -3610.5979 | 0.0714 | 1 | -3606.981 |
| **FAM110C** | RHSPN19050 | RJON | -12558.264 | 0.0369 | 1 | -12554.651 |
| **ATP11A** | RHSPN20494 | RJON | -17198.587 | 0.07216 | 1 | -17194.974 |
| **RFX7** | RHSPN26781 | RJON | -6384.1254 | 0.0498 | 1 | -6380.5165 |
| **SLC37A1** | RHSPN18628 | RJON | -9850.2143 | 0.06058 | 1 | -9846.6064 |
| **HSPH1** | RHSPN01812 | RJON | -4327.0523 | 0.0624 | 1 | -4323.4494 |
| **SCAMP3** | RHSPN19459 | RJON | -7810.6053 | 0.04424 | 1 | -7807.0065 |
| **DDHD1** | RHSPN24997 | RJON | -18112.105 | 0.04843 | 1 | -18108.521 |
| **SIPA1L3** | RHSPN19490 | RJON | -1322.8597 | 0.01003 | 1 | -1319.2834 |
| **NAA30** | RHSPN25262 | RJON | -5349.9021 | 0.06926 | 1 | -5346.3243 |
| **HIF3A** | RHSPN20187 | RJON | -2278.0316 | 0.08866 | 1 | -2274.4586 |
| **FAM189A1** | RHSPN20947 | RJON | -3190.9207 | 0.02971 | 1 | -3187.3507 |
| **ST8SIA2** | RHSPN14701 | RJON | -32313.08 | 0.08079 | 1 | -32309.51 |
| **JMJD1C** | RHSPN21035 | RJON | -2442.4836 | 0.09801 | 1 | -2438.9166 |
| **HBA1** | RHSPN16327 | RJON | -5101.0366 | 0.01853 | 1 | -5097.4741 |
| **TMEM151A** | RHSPN21046 | RJON | -12096.211 | 0.10415 | 1 | -12092.661 |
| **TMEM8A** | RHSPN05771 | RJON | -3300.2188 | 0.11402 | 1 | -3296.667 |
| **CMSS1** | RHSPN14734 | RJON | -11005.117 | 0.18565 | 1 | -11001.575 |
| **STOX1** | RHSPN04763 | RJON | -8678.0461 | 0.0686 | 1 | -8674.5075 |
| **ALPI** | RHSPN20246 | RJON | -7634.9221 | 0.13688 | 1 | -7631.3957 |
| **ZNF770** | RHSPN07438 | RJON | -6613.0575 | 0.06481 | 1 | -6609.5315 |
| **SPATA5** | RHSPN26816 | RJON | -3917.4133 | 0.05157 | 1 | -3913.9062 |
| **C21orf33** | RHSPN26229 | RJON | -3734.8745 | 0.11596 | 1 | -3731.3769 |
| **FITM2** | RHSPN11878 | RJON | -3733.7521 | 0.05808 | 1 | -3730.2664 |
| **DNAJB6** | RHSPN06735 | RJON | -19674.258 | 0.12228 | 1 | -19670.785 |
| **SH3TC1** | RHSPN23596 | RJON | -19700.122 | 0.06374 | 1 | -19696.655 |
| **BAHCC1** | RHSPN25848 | RJON | -9998.0829 | 0.13044 | 1 | -9994.6212 |
| **SIGLEC1** | RHSPN18943 | RJON | -2311.4709 | 0.01349 | 1 | -2308.015 |
| **SOX21** | RHSPN05064 | RJON | -5303.8792 | 0.06665 | 1 | -5300.4342 |
| **RBSN** | RHSPN05861 | RJON | -3966.4129 | 0.16638 | 1 | -3962.9708 |
| **CD200R1** | RHSPN06198 | RJON | -5289.171 | 0.14447 | 1 | -5285.7287 |
| **CPA3** | RHSPN10962 | RJON | -18271.842 | 0.05624 | 1 | -18268.394 |
| **EGFR** | RHSPN12174 | RJON | -2589.5573 | 0.026 | 1 | -2586.111 |

| | | | | | | |
|---|---|---|---|---|---|---|
| **DCTN6** | RHSPN03561 | RJON | -2704.5228 | 0.12237 | 1 | -2701.0764 |
| **FABP1** | RHSPN09017 | RJON | -12756.249 | 0.05738 | 1 | -12752.81 |
| **ADAMTS2** | RHSPN19372 | RJON | -4156.5658 | 0.12346 | 1 | -4153.1283 |
| **MIPOL1** | RHSPN00980 | RJON | -4510.1202 | 0.06377 | 1 | -4506.689 |
| **YIPF1** | RHSPN28443 | RJON | -1424.7851 | 0.05924 | 1 | -1421.3653 |
| **VMA21** | RHSPN14798 | RJON | -3626.1046 | 0.14013 | 1 | -3622.6878 |
| **FAM149B1** | RHSPN19047 | RJON | -11006.305 | 0.03185 | 1 | -11002.892 |
| **TUBGCP3** | RHSPN21130 | RJON | -48835.951 | 0.08382 | 1 | -48832.536 |
| **PKD1** | RHSPN05883 | RJON | -10836.484 | 0.13665 | 1 | -10833.071 |
| **KIAA1407** | RHSPN12710 | RJON | -7300.8608 | 0.02116 | 1 | -7297.4632 |
| **HAS2** | RHSPN19856 | RJON | -7278.1329 | 0.14739 | 1 | -7274.7396 |
| **SERPINA10** | RHSPN17229 | RJON | -2444.6839 | 0.24593 | 1 | -2441.2946 |
| **OLR1** | RHSPN06714 | RJON | -1476.637 | 0.08413 | 1 | -1473.251 |
| **BLOC1S4** | RHSPN12745 | RJON | -13609.326 | 0.03296 | 1 | -13605.94 |
| **KIAA0196** | RHSPN01916 | RJON | -5693.6282 | 0.17196 | 1 | -5690.2446 |
| **CD1D** | RHSPN10585 | RJON | -5803.3851 | 0.09902 | 1 | -5800.0092 |
| **TNRC18** | RHSPN18426 | RJON | -1444.9958 | 0.03661 | 1 | -1441.6216 |
| **AACS** | RHSPN04750 | RJON | -6950.9526 | 0.04957 | 1 | -6947.5837 |
| **B3GNT7** | RHSPN20546 | RJON | -46162.081 | 0.07954 | 1 | -46158.719 |
| **VPS13C** | RHSPN09160 | RJON | -12439.943 | 0.08023 | 1 | -12436.591 |
| **NEDD9** | RHSPN09160 | RJON | -12435.723 | 0.08003 | 1 | -12432.377 |
| **NEDD9** | RHSPN22269 | RJON | -3061.5831 | 0.22488 | 1 | -3058.2409 |
| **INCA1** | RHSPN06672 | RJON | -8218.4566 | 0.07958 | 1 | -8215.1181 |
| **MFSD10** | RHSPN16866 | RJON | -1218.8839 | 0.08692 | 1 | -1215.5555 |
| **CCDC84** | RHSPN24850 | RJON | -4557.1812 | 0.0622 | 1 | -4553.8522 |
| **SLC7A9** | RHSPN26626 | RJON | -4168.1694 | 0.10546 | 1 | -4164.8459 |
| **RWDD2B** | RHSPN02029 | RJON | -4186.0874 | 0.10898 | 1 | -4182.7676 |
| **HSD17B7** | RHSPN27125 | RJON | -4765.9763 | 0.0322 | 1 | -4762.6604 |
| **ASPHD2** | RHSPN28181 | RJON | -2361.0089 | 0.0235 | 1 | -2357.7057 |
| **ATG4A** | RHSPN21407 | RJON | -2830.6075 | 0.22128 | 1 | -2827.3014 |
| **IGSF6** | RHSPN23621 | RJON | -4870.9568 | 0.0311 | 1 | -4867.6535 |
| **PCYT2** | RHSPN11336 | RJON | -1935.8957 | 0.04206 | 1 | -1932.6114 |
| **POP7** | RHSPN22701 | RJON | -9167.2213 | 0.04459 | 1 | -9163.9409 |
| **UNC45B** | RHSPN12346 | RJON | -4198.5412 | 0.08345 | 1 | -4195.2638 |
| **PENK** | RHSPN22847 | RJON | -4276.5938 | 0.09241 | 1 | -4273.3167 |
| **KRT222** | RHSPN04230 | RJON | -17934.251 | 0.07807 | 1 | -17930.976 |
| **ITGA6** | RHSPN21917 | RJON | -13344.884 | 0.09833 | 1 | -13341.612 |
| **CMTR2** | RHSPN12028 | RJON | -3958.0413 | 0.04521 | 1 | -3954.7698 |
| **ZDHHC2** | RHSPN02280 | RJON | -7645.8654 | 0.04103 | 1 | -7642.5957 |
| **ZBTB41** | RHSPN00267 | RJON | -21954.701 | 0.06621 | 1 | -21951.432 |
| **PRDM2** | RHSPN17672 | RJON | -1685.9908 | 0.14061 | 1 | -1682.7345 |
| **NPFF** | RHSPN21301 | RJON | -9510.3998 | 0.07566 | 1 | -9507.1525 |
| **MKL2** | RHSPN18025 | RJON | -2531.855 | 0.05561 | 1 | -2528.6075 |
| **SOCS2** | RHSPN15157 | RJON | -5091.8058 | 0.08123 | 1 | -5088.5578 |
| **WBP1L** | RHSPN12106 | RJON | -9425.1691 | 0.02049 | 1 | -9421.9256 |

| | | | | | | |
|---|---|---|---|---|---|---|
| **RHOBTB2** | RHSPN07217 | RJON | -4542.298 | 0.15236 | 1 | -4539.0641 |
| **DMP1** | RHSPN12975 | RJON | -5974.7953 | 0.083 | 1 | -5971.5644 |
| **DMRT2** | RHSPN20496 | RJON | -4124.207 | 0.08716 | 1 | -4120.9759 |
| **TEX9** | RHSPN03315 | RJON | -2667.3787 | 0.12404 | 1 | -2664.1462 |
| **FANCL** | RHSPN10056 | RJON | -1483.1273 | 0.10636 | 1 | -1479.9049 |
| **ARMC2** | RHSPN24677 | RJON | -2207.1682 | 0.05162 | 1 | -2203.9444 |
| **GTPBP3** | RHSPN08432 | RJON | -1770.5161 | 0.01648 | 1 | -1767.2922 |
| **PGGT1B** | RHSPN18177 | RJON | -2209.7786 | 0.08402 | 1 | -2206.5577 |
| **DAO** | RHSPN08254 | RJON | -19084.806 | 0.14837 | 1 | -19081.586 |
| **CMYA5** | RHSPN13723 | RJON | -4874.0918 | 0.03001 | 1 | -4870.8725 |
| **SNX30** | RHSPN06292 | RJON | -7940.7158 | 0.0595 | 1 | -7937.4985 |
| **IFT80** | RHSPN24392 | RJON | -17230.697 | 0.0808 | 1 | -17227.499 |
| **ARHGEF18** | RHSPN09111 | RJON | -4392.8991 | 0.12676 | 1 | -4389.7071 |
| **RPP40** | RHSPN23858 | RJON | -12810.188 | 0.11572 | 1 | -12807.006 |
| **DSC2** | RHSPN06349 | RJON | -1399.5286 | 0.01511 | 1 | -1396.351 |
| **EIF5A2** | RHSPN00667 | RJON | -1946.5398 | 0.14001 | 1 | -1943.3645 |
| **MRPS15** | RHSPN05553 | RJON | -8472.8149 | 0.07018 | 1 | -8469.6388 |
| **RFT1** | RHSPN28508 | RJON | -28554.676 | 0.02015 | 1 | -28551.507 |
| **FLNA** | RHSPN02571 | RJON | -22802.617 | 0.11902 | 1 | -22799.453 |
| **TLR5** | RHSPN11307 | RJON | -1302.306 | 0.09151 | 1 | -1299.145 |
| **AP4M1** | RHSPN25377 | RJON | -5698.6731 | 0.10005 | 1 | -5695.5174 |
| **CCDC155** | RHSPN05011 | RJON | -7023.8415 | 0.0234 | 1 | -7020.6906 |
| **SLC6A1** | RHSPN18087 | RJON | -11713.217 | 0.04374 | 1 | -11710.07 |
| **SCYL2** | RHSPN16148 | RJON | -6188.2654 | 0.01302 | 1 | -6185.125 |
| **DAGLA** | RHSPN25015 | RJON | -8364.178 | 0.01055 | 1 | -8361.0411 |
| **ACTN4** | RHSPN14102 | RJON | -6139.4132 | 0.07397 | 1 | -6136.2718 |
| **INPP5E** | RHSPN14315 | RJON | -2739.3296 | 0.16126 | 1 | -2736.1918 |
| **OLAH** | RHSPN25663 | RJON | -2410.3218 | 0.07688 | 1 | -2407.1899 |
| **NAT14** | RHSPN27306 | RJON | -7173.6427 | 0.06177 | 1 | -7170.514 |
| **CARD10** | RHSPN00952 | RJON | -801.72618 | 0.03011 | 1 | -798.59913 |
| **FAM159A** | RHSPN24001 | RJON | -3297.6214 | 0.11589 | 1 | -3294.4972 |
| **SKA1** | RHSPN27288 | RJON | -7017.9609 | 0.01327 | 1 | -7014.8383 |
| **EIF3D** | RHSPN05532 | RJON | -19289.296 | 0.07709 | 1 | -19286.172 |
| **NISCH** | RHSPN27117 | RJON | -8098.4392 | 0.01977 | 1 | -8095.3194 |
| **ADRBK2** | RHSPN28298 | RJON | -1716.0723 | 0.01965 | 1 | -1712.9521 |
| **RAB33A** | RHSPN15756 | RJON | -20971.791 | 0.06295 | 1 | -20968.673 |
| **PIK3C2A** | RHSPN08835 | RJON | -3002.6631 | 0.04905 | 1 | -2999.5517 |
| **LSM11.00** | RHSPN07842 | RJON | -1184.199 | 0.07419 | 1 | -1181.0953 |
| **MRPL36** | RHSPN24895 | RJON | -6451.4942 | 0.05532 | 1 | -6448.3998 |
| **LSR** | RHSPN21882 | RJON | -2330.2562 | 0.04254 | 1 | -2327.1633 |
| **NIP7** | RHSPN13842 | RJON | -1789.7299 | 0.00969 | 1 | -1786.6382 |
| **NEK6** | RHSPN19230 | RJON | -2601.0806 | 0.02295 | 1 | -2597.9955 |
| **CEBPE** | RHSPN07493 | RJON | -1162.9891 | 0.1699 | 1 | -1159.9079 |
| **NDUFC1** | RHSPN20580 | RJON | -6282.06 | 0.0716 | 1 | -6278.97 |
| **TRIP4** | RHSPN11737 | RJON | -4626.9562 | 0.05395 | 1 | -4623.8753 |

| | | | | | | |
|---|---|---|---|---|---|---|
| **OR6B1** | RHSPN18052 | RJON | -4165.432 | 0.14856 | 1 | -4162.3516 |
| **CCDC38** | RHSPN00130 | RJON | -6181.7066 | 0.06785 | 1 | -6178.6221 |
| **WRAP73** | RHSPN03867 | RJON | -6088.6861 | 0.03513 | 1 | -6085.6168 |
| **SLC35F5** | RHSPN05348 | RJON | -7104.4147 | 0.11726 | 1 | -7101.3457 |
| **LRRC2** | RHSPN02317 | RJON | -1617.5616 | 0.02597 | 1 | -1614.5033 |
| **TNNI1** | RHSPN03034 | RJON | -6534.2215 | 0.09542 | 1 | -6531.1662 |
| **DRC1** | RHSPN15066 | RJON | -1843.9258 | 0.02473 | 1 | -1840.8735 |
| **GOLGA7B** | RHSPN00239 | RJON | -1973.4662 | 0.13723 | 1 | -1970.4128 |
| **NPPA** | RHSPN09446 | RJON | -1591.6241 | 0.09259 | 1 | -1588.5741 |
| **APOM** | RHSPN20748 | RJON | -2333.1073 | 0.07874 | 1 | -2330.0629 |
| **MAN2C1** | RHSPN04402 | RJON | -6392.1195 | 0.09396 | 1 | -6389.0752 |
| **RFTN2** | RHSPN09396 | RJON | -3636.977 | 0.10416 | 1 | -3633.9313 |
| **MRPS18B** | RHSPN05489 | RJON | -6050.7563 | 0.09633 | 1 | -6047.7162 |
| **HEMK1** | RHSPN27163 | RJON | -10821.801 | 0.01807 | 1 | -10818.764 |
| **AP1B1** | RHSPN04348 | RJON | -1949.251 | 0.02051 | 1 | -1946.2196 |
| **ORMDL1** | RHSPN15158 | RJON | -10287.868 | 0.13792 | 1 | -10284.864 |
| **CYP17A1** | RHSPN02484 | RJON | -4410.2888 | 0.06137 | 1 | -4407.2868 |
| **LPGAT1** | RHSPN25014 | RJON | -8392.1739 | 0.07904 | 1 | -8389.1772 |
| **MAP4K1** | RHSPN23323 | RJON | -7425.0851 | 0.02468 | 1 | -7422.0884 |
| **PRKCA** | | | | | | |

Jonathan's PSGs shared with Bowhead Whale PSGs

| | | | |
|---|---|---|---|
| **B** | **ead Whale's** | **lection loca** | **nction of Gene** |
| **nathan's Ge** | **DMP1** | 710 | Dentin matrix acidic phosphoprotein |
| **DMP1** | **IGSF6** | 694 | Immunoglobulin superfamily member 6: |
| **IGSF6** | **DNALI1** | 23 | Dynein Axonemal Light Intermediate Chain 1 |
| **DNALI1** | | | |

Jonathan's PSGs shared with Brandit's bat PSGs

| | | | |
|---|---|---|---|
| **C** | **ndit's Bat G** | **lection loca** | **ene Function** |
| **nathan's Ge** | **ADAM19** | 151 | ADAM metallopeptidase domain 19 |
| **ADAM19** | **C17orf78** | 508 | chromosome 17 open reading frame 78 |
| **C17orf78** | **CD200R1** | 661 | CD200 receptor 1 |
| **CD200R1** | **CYNP4** | 483 | canopy 4 homolog (zebrafish) |
| **CYNP4** | **GTPBP3** | 715 | GTP binding protein 3 (mitochondrial) |
| **GTPBP3** | **P2RY11** | 510 | purinergic receptor P2Y, G-protein coupled, 11 |
| **P2RY11** | **PHACTR4** | 386 | phosphatase and actin regulator 4 |
| **PHACTR4** | **RHNO1** | 520 | RAD0-HUS Involved in microhomology-mediate |
| **RHNO1** | **SCN5A** | 32 | sodium channel, voltage-gated, type V, alpha sub |
| **SCN5A** | **SLC16A1** | 149 | solute carrier family 16, member 1 (monocarboxy |
| **SLC16A1** | **SLC44A2** | 546 | solute carrier family 44, member 2 |
| **SLC44A2** | **TPM2** | 6 | tropomyosin 2 (beta) |
| **TPM2** | | | |

Jonathan's PSGs shared with Killifish PSGs (highest confiden

| | | | |
|---|---|---|---|
| **D** | **illifish's Ger** | **lection loca** | **ene Function** |
| **nathan's Ge** | **BSN** | 93 | Bassoon presynaptic cytomatrix protein |
| **BSN** | **GRPR** | 613 | Gastrin-releasing peptide receptor |
| **GRPR** | **VPS13C** | 683 | Vacuolar Protein Sorting 13 Homolog C |
| **VPS13C** | **COLQ** | 8 | collagen-like molecule associated with acetylch |

| | | | |
|---|---|---|---|
| COLQ | LRRK2 | 224 | Serine/threonine-protein kinase;associated with |
| LRRK2 | DOMMD1 | 48 | Proposed scaffold protein;Can modulate activity |
| COMMD1 | MYO1B | 17 | Myosin IB |
| MYO1B | EYS | 581 | Eyes shut homolog; Associated with retinitis pig |
| EYS | MX1 | 302 | MX Dynamin Like GTPase 1; encodes a guanos |
| MX1 | OBSCN | 355 | obscurin gene spans more than 150 kb |
| OBSCN | XRCC5 | 342 | X-Ray Repair Cross Complementing 5 |
| XRCC5 | TLR5 | 728 | Toll Like Receptor 5 |
| TLR5 | NOX4 | 220 | NADPH Oxidase 4; catalyzes the reduction of m |
| NOX4 | RASA2 | 388 | member of the GAP1 family of GTPase-activatin |
| RASA2 | HSPA4 | 451 | Heat Shock Protein Family A Member 4 |
| HSPA4L | Jonathan's PSGs shared with NMR's PSGs | | |
| E | NMR's Gen | election loca | ene Function |
| nathan's Ge | STOX1 | 649 | regulates the levels of reactive oxidative species |
| STOX1 | INCA1 | 686 | Inhibitor Of CDK, Cyclin A1 Interacting Protein 1 |
| INCA1 | CASP14 | 312 | Caspase 14, Apoptosis-Related Cysteine Peptic |
| CASP14 | ANK2 | 527 | Ankyrin 2; link the integral membrane to the spe |
| ANK2 | CD1D | 679 | Antigen-Presenting Glycoprotein CD1d |
| CD1D | APOM | 763 | Apolipoprotein M; associated with HDL |
| APOM | CMTR2 | 701 | Cap Methyltransferase 2; Involved in 7-methylg |
| CMTR2 | SKA1 | 740 | Spindle And Kinetochore Associated Complex S |
| SKA1 | HIGD1A | 50 | Proposed subunit of cytochrome c oxidase |
| HIGD1A | THEM6 | 539 | Thioesterase Superfamily Member 6 |
| THEM6 | Jonathan's PSGs shared with Rockfish's PSGs | | |
| F | ockfish's Ge | election loca | ene function |
| nathan's Ge | HCFC1 | 84 | DNA damage; chromatin bindingl; cell cycle |
| HCFC1 | TP53I3 | 210 | Generation of reactive oxygen species (ROS); ind |
| TP53I3 | OBSCN | 355 | Myofibrillog Residual |
| OBSCN | CD7 | 356 | T-cell/B-cell interaction; hematopoietic cell linea |
| CD7 | LAS1L | 384 | rRNA processing; ribosomal biogenesis; endonu |
| LAS1L | IMPAD1 | 473 | 3'(2'),5'-bisphosphate nucleotidase activity; hydr |
| IMPAD1 | INA | 596 | Neuronal morphogenesis; structural constituent |
| INA | STOX1 | 649 | RNA polymerase II regulatory region sequence-s |
| STOX1 | NEK6 | 750 | Tumor cell senescence suppresion; mitotic cell c |
| NEK6 | BARD1 | 426 | protein which interacts with the N-terminal regio |
| BARD1 | FABP1 | 665 | fatty acid binding protein |
| FABP1 | PTGER2 | 429 | receptor for prostaglandin E2 |
| PTGER2 | RIN1 | 313 | Ras effector protein |
| RIN1 | ITGA6 | 700 | integrin alpha chain family of proteins |
| ITGA6 | Jonathan's PSGs shared with LongevityMap significant huma | | |
| G | election loca | ene Function | |
| nathan's Ge | 240 | member of the telombin family and encodes a nuclear protei | |
| POT1 | 255 | glycoprotein that functions as part of the classical pathway o | |
| C2 | 99 | Involved in dendrite development | |

| | | |
|---|---|---|
| CSMD3 | 317 | play a central role in the physiological regulation of progran |
| FAS | 397 | heterodimerizes with the beta 1 subunit to form a cell-surfac |
| ITGA1 | 411 | Involved in regulation of axonogenesis |
| KIF13B | 237 | expressed on nearly all natural killer (NK) cells and stimulat |
| KLRF1 | 192 | Belongs to the immunogloblin (Ig) superfamily |
| LY6G6F | 124 | subunit of the 2-oxoglutarate dehydrogenase complex |
| OGDH | 198 | component of the Notch signaling |
| PCNX | 285 | Essential for DNA repair by homologous recombination |
| RAD51B | 707 | interacts with the cytoplasmic domain of insulin-like growth |
| SOCS2 | 773 | Protein Kinase C Alpha; regulator of cardiac contractility anc |
| PRKCA | 342 | X-ray repair complementing defective repair in Chinese hams |
| XRCC5 | 169 | This domain is known to be involved in protein-protein intera |
| BTBD9 | 663 | This protein is a receptor for members of the epidermal grow |
| EGFR | 318 | a scaffolding molecule that regulates the actin cytoskeleton |
| CD2AP | 314 | belongs to the ephrin receptor subfamily of the protein-tyros |
| EPHA1 | 177 | A major protein component of high density lipoprotein (HDL |
| APOA1 | 61 | alpha-1 subunit of a voltage-dependent calcium channel |
| CACNA1C | 507 | coagulation factor VII which is a vitamin K-dependent factor |
| F7 | 490 | member of the glutathione S-transferase (GSTs) super-famil |
| GSTZ1 | | |

### Jonathan's PSGs shared with GeneAge 307 aging genes

| H | lection loca | ene Functic Evidence |
|---|---|---|
| nathan's Ge | 400 | calcium channel, voltage-dependent, P/Q type, alpha 1A sub |
| CACNA1C | 663 | epidermal growth factor receptor |
| EGFR | 724 | eukaryotic t Entry selected based on evidence directly linking the g |
| EIF5A2 | 317 | Fas cell surface death receptor |
| FAS | 600 | fms-related tyrosine kinase 1 |
| FLT1 | 183 | granulin |
| GRN | 544 | glutathione Entry slected based on evidence linking the geneprodu |
| GSTP1 | 294 | jun D proto- Entry selected based on evidence directly linking the g |
| JUND | 415 | nuclear rece Entry selected based on evidence directly linking the g |
| NCOR2 | 308 | phospholipase C, gamma 2 (phosphatidylinositol-specific) |
| PLCG2 | 773 | protein kinase C, alpha |
| PRKCA | 707 | suppressor Entry selected based on evidence linking the gene proc |
| SOCS2 | 351 | sequestosome 1 |
| SQSTM1 | 342 | X-ray repair Entry slected based on evidence linking the geneprodu |
| XRCC5 | | |

All 307 genes in GenAge were used to search Jonathan's PS

d SIFT analysis

ce, 249 genes)

n variants (274 genes)

d SIFT analysis

| bg.w | fg.w | bg.w1 | fg.w1 | LRT | P | Q |
|---|---|---|---|---|---|---|
| 0.08518 | 4.44769 | 1 | 4.44769 | 199.90106 | ########## | ########## |
| 0.02671 | 14.34999 | 1 | 14.34999 | 172.682296 | ########## | ########## |
| 0.05043 | 84.24533 | 1 | 84.24533 | 127.891374 | ########## | ########## |
| 0.00829 | 999 | 1 | 999 | 124.072148 | ########## | ########## |
| 0.02728 | 4.93175 | 1 | 4.93175 | 122.980006 | ########## | ########## |
| 0.06035 | 999 | 1 | 999 | 121.771338 | ########## | ########## |
| 0.00937 | 734.05354 | 1 | 734.05354 | 115.012472 | ########## | ########## |
| 0.12542 | 9.3611 | 1 | 9.3611 | 101.864096 | ########## | ########## |
| 0.00525 | 14.54884 | 1 | 14.54884 | 91.257506 | ########## | ########## |
| 0.05006 | 762.88403 | 1 | 762.88403 | 85.678264 | ########## | ########## |
| 0.03097 | 13.46195 | 1 | 13.46195 | 83.95 | ########## | ########## |
| 0.05021 | 104.70718 | 1 | 104.70718 | 82.501522 | ########## | ########## |
| 0.0181 | 727.4668 | 1 | 727.4668 | 81.469658 | ########## | ########## |
| 0.02634 | 999 | 1 | 999 | 76.485112 | ########## | ########## |
| 0.0258 | 8.50788 | 1 | 8.50788 | 68.628902 | ########## | ########## |
| 0.04556 | 2.67856 | 1 | 2.67856 | 67.257276 | ########## | ########## |
| 0.02164 | 387.3051 | 1 | 387.3051 | 65.626024 | ########## | ########## |
| 0.08701 | 568.02688 | 1 | 568.02688 | 58.720514 | ########## | ########## |
| 0.06368 | 1.6856 | 1 | 1.6856 | 56.5734 | ########## | ########## |
| 0.04946 | 602.13172 | 1 | 602.13172 | 56.190334 | ########## | ########## |
| 0.02139 | 999 | 1 | 999 | 54.244602 | ########## | ########## |
| 0.0461 | 2.86704 | 1 | 2.86704 | 53.981122 | ########## | ########## |
| 0.12945 | 5.14487 | 1 | 5.14487 | 53.076846 | ########## | ########## |
| 0.06765 | 999 | 1 | 999 | 51.36499 | ########## | ########## |
| 0.08377 | 411.21283 | 1 | 411.21283 | 51.33861 | ########## | ########## |
| 0.06001 | 256.87801 | 1 | 256.87801 | 51.544504 | ########## | ########## |
| 0.03659 | 161.65845 | 1 | 161.65845 | 51.369954 | ########## | ########## |
| 0.05182 | 119.66258 | 1 | 119.66258 | 50.072664 | ########## | ########## |
| 0.07026 | 229.17352 | 1 | 229.17352 | 49.573586 | ########## | ########## |
| 0.03241 | 6.01569 | 1 | 6.01569 | 49.581948 | ########## | ########## |
| 0.02114 | 1.00819 | 1 | 1.00819 | 45.977212 | ########## | ########## |
| 0.03326 | 6.39363 | 1 | 6.39363 | 45.88512 | ########## | ########## |

| | | | | | | |
|---|---|---|---|---|---|---|
| 0.08712 | 391.31412 | 1 | 391.31412 | 45.553518 | ########## | ########## |
| 0.15051 | 104.78968 | 1 | 104.78968 | 45.5385 | ########## | ########## |
| 0.05259 | 999 | 1 | 999 | 45.328308 | ########## | ########## |
| 0.04801 | 370.66271 | 1 | 370.66271 | 45.015174 | ########## | ########## |
| 0.07927 | 250.38087 | 1 | 250.38087 | 43.582752 | ########## | ########## |
| 0.01187 | 23.50307 | 1 | 23.50307 | 43.33554 | ########## | ########## |
| 0.08601 | 701.84583 | 1 | 701.84583 | 42.567402 | ########## | ########## |
| 0.21013 | 528.24302 | 1 | 528.24302 | 42.019894 | ########## | ########## |
| 0.00875 | 113.15562 | 1 | 113.15562 | 42.022858 | ########## | ########## |
| 0.09015 | 2.81722 | 1 | 2.81722 | 41.768652 | ########## | ########## |
| 0.03046 | 185.49314 | 1 | 185.49314 | 41.31231 | ########## | ########## |
| 0.0626 | 310.72109 | 1 | 310.72109 | 40.706786 | ########## | ########## |
| 0.19839 | 999 | 1 | 999 | 39.840556 | ########## | ########## |
| 0.05953 | 999 | 1 | 999 | 39.65172 | ########## | ########## |
| 0.02564 | 999 | 1 | 999 | 39.588948 | ########## | ########## |
| 0.03586 | 999 | 1 | 999 | 39.065914 | ########## | ########## |
| 0.06118 | 27.82926 | 1 | 27.82926 | 39.027058 | ########## | ########## |
| 0.03011 | 999 | 1 | 999 | 38.036616 | ########## | ########## |
| 0.05679 | 999 | 1 | 999 | 37.408776 | ########## | ########## |
| 0.02508 | 2.65044 | 1 | 2.65044 | 37.358836 | ########## | ########## |
| 0.11087 | 999 | 1 | 999 | 36.403866 | ########## | ########## |
| 0.07806 | 6.22094 | 1 | 6.22094 | 36.243576 | ########## | ########## |
| 0.01041 | 8.40745 | 1 | 8.40745 | 36.06679 | ########## | ########## |
| 0.03606 | 999 | 1 | 999 | 35.526736 | ########## | ########## |
| 0.0331 | 48.65651 | 1 | 48.65651 | 35.160732 | ########## | ########## |
| 0.06619 | 48.34104 | 1 | 48.34104 | 35.118894 | ########## | ########## |
| 0.02768 | 999 | 1 | 999 | 34.639422 | ########## | ########## |
| 0.15207 | 778.2451 | 1 | 778.2451 | 34.321406 | ########## | ########## |
| 0.01735 | 999 | 1 | 999 | 34.018504 | ########## | ########## |
| 0.01589 | 581.42899 | 1 | 581.42899 | 33.93093 | ########## | ########## |
| 0.04537 | 1.90738 | 1 | 1.90738 | 33.93093 | ########## | ########## |
| 0.09095 | 999 | 1 | 999 | 33.760024 | ########## | ########## |
| 0.02864 | 846.60991 | 1 | 846.60991 | 33.447242 | ########## | ########## |
| 0.1279 | 190.54149 | 1 | 190.54149 | 32.916622 | ########## | ########## |
| 0.01211 | 999 | 1 | 999 | 32.819252 | ########## | ########## |
| 0.14623 | 145.59849 | 1 | 145.59849 | 32.7863 | ########## | ########## |
| 0.03796 | 55.85838 | 1 | 55.85838 | 31.818134 | ########## | ########## |
| 0.0891 | 469.06485 | 1 | 469.06485 | 31.331834 | ########## | ########## |
| 0.02842 | 93.08541 | 1 | 93.08541 | 30.250682 | ########## | ########## |
| 0.0818 | 191.40384 | 1 | 191.40384 | 29.961108 | ########## | ########## |
| 0.02621 | 11.52178 | 1 | 11.52178 | 29.56162 | ########## | ########## |
| 0.01632 | 443.91418 | 1 | 443.91418 | 29.277056 | ########## | ########## |
| 0.00919 | 999 | 1 | 999 | 29.163018 | ########## | ########## |
| 0.04757 | 157.77001 | 1 | 157.77001 | 29.141566 | ########## | ########## |
| 0.02504 | 1.11925 | 1 | 1.11925 | 28.987856 | ########## | ########## |

| | | | | | | |
|---|---|---|---|---|---|---|
| 0.08993 | 120.31565 | 1 | 120.31565 | 28.776566 | ########## | ########## |
| 0.12034 | 5.8601 | 1 | 5.8601 | 28.445686 | ########## | 0.0001107 |
| 0.09412 | 41.51705 | 1 | 41.51705 | 28.242736 | ########## | 0.00012113 |
| 0.05811 | 11.20265 | 1 | 11.20265 | 28.166196 | ########## | 0.00012444 |
| 0.01937 | 3.19306 | 1 | 3.19306 | 28.077252 | ########## | 0.00012865 |
| 0.08471 | 137.69551 | 1 | 137.69551 | 27.94155 | ########## | 0.00013617 |
| 0.0533 | 2.03204 | 1 | 2.03204 | 27.702678 | ########## | 0.00015178 |
| 0.09166 | 72.92824 | 1 | 72.92824 | 27.544036 | ########## | 0.00016254 |
| 0.22614 | 363.46964 | 1 | 363.46964 | 27.085212 | ########## | 0.00020015 |
| 0.20231 | 21.36554 | 1 | 21.36554 | 27.104858 | ########## | 0.00020015 |
| 0.08705 | 42.90165 | 1 | 42.90165 | 26.84067 | ########## | 0.00022383 |
| 0.01453 | 999 | 1 | 999 | 26.571438 | ########## | 0.00025344 |
| 0.10969 | 999 | 1 | 999 | 26.533912 | ########## | 0.0002556 |
| 0.07653 | 1.17188 | 1 | 1.17188 | 26.47483 | ########## | 0.00026061 |
| 0.11958 | 3.0582 | 1 | 3.0582 | 26.090514 | ########## | 0.00031266 |
| 0.03185 | 340.288 | 1 | 340.288 | 25.90437 | ########## | 0.00033643 |
| 0.00422 | 112.16063 | 1 | 112.16063 | 25.718044 | ########## | 0.0003657 |
| 0.1119 | 999 | 1 | 999 | 25.694748 | ########## | 0.00036642 |
| 0.09787 | 219.90507 | 1 | 219.90507 | 25.544854 | ########## | 0.00039118 |
| 0.01981 | 1.63784 | 1 | 1.63784 | 25.509758 | ########## | 0.00039435 |
| 0.01316 | 999 | 1 | 999 | 25.130314 | ########## | 0.00047228 |
| 0.12473 | 19.83545 | 1 | 19.83545 | 24.912184 | ########## | 0.00052182 |
| 0.2001 | 42.37543 | 1 | 42.37543 | 24.74994 | ########## | 0.00055951 |
| 0.07814 | 9.99516 | 1 | 9.99516 | 24.736018 | ########## | 0.00055951 |
| 0.03107 | 999 | 1 | 999 | 24.429404 | ########## | 0.00064634 |
| 0.02614 | 5.76623 | 1 | 5.76623 | 24.308252 | ########## | 0.00068057 |
| 0.0389 | 999 | 1 | 999 | 24.123236 | ########## | 0.00073343 |
| 0.07766 | 19.87981 | 1 | 19.87981 | 23.783518 | ########## | 0.00085423 |
| 0.03542 | 999 | 1 | 999 | 23.748612 | ########## | 0.00086184 |
| 0.07635 | 2.82636 | 1 | 2.82636 | 23.706226 | ########## | 0.00086551 |
| 0.10596 | 221.7661 | 1 | 221.7661 | 23.60938 | ########## | 0.00090088 |
| 0.02988 | 999 | 1 | 999 | 23.451362 | ########## | 0.00095896 |
| 0.11456 | 22.04196 | 1 | 22.04196 | 23.350824 | ########## | 0.0010002 |
| 0.16549 | 5.71402 | 1 | 5.71402 | 23.24119 | ########## | 0.00104804 |
| 0.12631 | 23.01217 | 1 | 23.01217 | 23.080894 | ########## | 0.00112641 |
| 0.01165 | 999 | 1 | 999 | 23.052276 | ########## | 0.00113357 |
| 0.04219 | 17.1729 | 1 | 17.1729 | 22.859152 | ########## | 0.00123866 |
| 0.13611 | 999 | 1 | 999 | 22.788338 | ########## | 0.00127328 |
| 0.07921 | 232.72086 | 1 | 232.72086 | 22.68044 | ########## | 0.00133344 |
| 0.14727 | 53.59703 | 1 | 53.59703 | 22.494876 | ########## | 0.00145183 |
| 0.01357 | 3.49197 | 1 | 3.49197 | 22.471224 | ########## | 0.00145788 |
| 0.0127 | 1.56573 | 1 | 1.56573 | 22.142114 | ########## | 0.00169281 |
| 0.05839 | 999 | 1 | 999 | 21.742648 | ########## | 0.00203643 |
| 0.03325 | 158.01245 | 1 | 158.01245 | 21.68281 | ########## | 0.00208285 |
| 0.02154 | 999 | 1 | 999 | 21.511756 | ########## | 0.00225227 |

| | | | | | | |
|---|---|---|---|---|---|---|
| 0.06729 | 9.59874 | 1 | 9.59874 | 21.44247 | ########## | 0.00231476 |
| 0.11987 | 5.78814 | 1 | 5.78814 | 21.166986 | ########## | 0.00261865 |
| 0.0219 | 174.92395 | 1 | 174.92395 | 21.14473 | ########## | 0.00262918 |
| 0.03609 | 1.09725 | 1 | 1.09725 | 21.122724 | ########## | 0.00263954 |
| 0.03356 | 999 | 1 | 999 | 21.076204 | ########## | 0.00268277 |
| 0.13752 | 52.43343 | 1 | 52.43343 | 20.850678 | ########## | 0.00298215 |
| 0.05758 | 250.18412 | 1 | 250.18412 | 20.789414 | ########## | 0.0030537 |
| 0.01723 | 3.58907 | 1 | 3.58907 | 20.613266 | ########## | 0.00328948 |
| 0.00135 | 14.68875 | 1 | 14.68875 | 20.282998 | ########## | 0.00385389 |
| 0.0595 | 1.56049 | 1 | 1.56049 | 20.090676 | ########## | 0.00421442 |
| 0.05683 | 17.258 | 1 | 17.258 | 20.04667 | ########## | 0.00427945 |
| 0.07162 | 999 | 1 | 999 | 19.975942 | ########## | 0.00440414 |
| 0.10438 | 999 | 1 | 999 | 19.931858 | ########## | 0.00444344 |
| 0.07384 | 999 | 1 | 999 | 19.829362 | ########## | 0.00464672 |
| 0.04136 | 202.8023 | 1 | 202.8023 | 19.770626 | ########## | 0.00475434 |
| 0.05043 | 5.82076 | 1 | 5.82076 | 19.716706 | ########## | 0.00485295 |
| 0.0611 | 264.81177 | 1 | 264.81177 | 19.687876 | ########## | 0.00489205 |
| 0.04489 | 999 | 1 | 999 | 19.606462 | ########## | 0.00503121 |
| 0.00288 | 33.74244 | 1 | 33.74244 | 19.533202 | ########## | 0.00515409 |
| 0.03158 | 12.78686 | 1 | 12.78686 | 19.53844 | ########## | 0.00515409 |
| 0.06318 | 681.16907 | 1 | 681.16907 | 19.498514 | ########## | 0.00518 |
| 0.06814 | 46.7499 | 1 | 46.7499 | 19.201986 | ########## | 0.00593494 |
| 0.06606 | 999 | 1 | 999 | 19.18207 | ########## | 0.00595822 |
| 0.04841 | 13.80204 | 1 | 13.80204 | 19.058966 | ########## | 0.00626105 |
| 0.05745 | 2.02014 | 1 | 2.02014 | 19.026424 | ########## | 0.0063261 |
| 0.06645 | 1.65531 | 1 | 1.65531 | 18.911752 | ########## | 0.00666001 |
| 0.08075 | 204.84348 | 1 | 204.84348 | 18.73234 | ########## | 0.00724247 |
| 0.08129 | 1.18416 | 1 | 1.18416 | 18.675964 | ########## | 0.00740621 |
| 0.05933 | 3.07709 | 1 | 3.07709 | 18.623708 | ########## | 0.00755833 |
| 0.08483 | 1.27128 | 1 | 1.27128 | 18.599586 | ########## | 0.00760607 |
| 0.12419 | 29.83456 | 1 | 29.83456 | 18.455042 | ########## | 0.00812941 |
| 0.00756 | 169.5817 | 1 | 169.5817 | 18.365374 | 0.0001028 | 0.00845387 |
| 0.02209 | 60.37276 | 1 | 60.37276 | 18.305944 | 0.0001059 | 0.00865964 |
| 0.08815 | 454.52162 | 1 | 454.52162 | 18.190516 | 0.0001122 | 0.00912259 |
| 0.10282 | 1.1777 | 1 | 1.1777 | 18.135944 | 0.0001153 | 0.00932256 |
| 0.05265 | 20.97967 | 1 | 20.97967 | 18.057218 | 0.00011993 | 0.00964297 |
| 0.06492 | 34.44343 | 1 | 34.44343 | 17.950574 | 0.0001265 | 0.01011492 |
| 0.03938 | 999 | 1 | 999 | 17.800136 | 0.00013638 | 0.01081024 |
| 0.04616 | 9.49259 | 1 | 9.49259 | 17.79563 | 0.00013669 | 0.01081024 |
| 0.05521 | 999 | 1 | 999 | 17.706524 | 0.00014291 | 0.01118057 |
| 0.06231 | 10.82611 | 1 | 10.82611 | 17.693636 | 0.00014384 | 0.01119235 |
| 0.12118 | 7.58971 | 1 | 7.58971 | 17.472494 | 0.00016066 | 0.01243406 |
| 0.11491 | 999 | 1 | 999 | 17.425382 | 0.00016449 | 0.01266272 |
| 0.01495 | 50.67349 | 1 | 50.67349 | 17.343238 | 0.00017138 | 0.01312383 |
| 0.01755 | 24.4513 | 1 | 24.4513 | 17.323392 | 0.00017309 | 0.01318494 |

| | | | | | | |
|---|---|---|---|---|---|---|
| 0.0857 | 4.33886 | 1 | 4.33886 | 17.237028 | 0.00018073 | 0.01369469 |
| 0.01201 | 8.29517 | 1 | 8.29517 | 17.219164 | 0.00018235 | 0.01372196 |
| 0.07316 | 5.58057 | 1 | 5.58057 | 17.212216 | 0.00018298 | 0.01372196 |
| 0.04925 | 999 | 1 | 999 | 17.058302 | 0.00019762 | 0.01466766 |
| 0.1495 | 46.63364 | 1 | 46.63364 | 17.019046 | 0.00020154 | 0.01488208 |
| 0.12896 | 999 | 1 | 999 | 16.99035 | 0.00020445 | 0.01496616 |
| 0.04414 | 18.04858 | 1 | 18.04858 | 16.987474 | 0.00020475 | 0.01496616 |
| 0.17361 | 263.43007 | 1 | 263.43007 | 16.89391 | 0.00021455 | 0.01556143 |
| 0.03662 | 242.51883 | 1 | 242.51883 | 16.87939 | 0.00021612 | 0.01556143 |
| 0.11596 | 147.66695 | 1 | 147.66695 | 16.888166 | 0.00021517 | 0.01556143 |
| 0.02045 | 1.91685 | 1 | 1.91685 | 16.840606 | 0.00022035 | 0.01570983 |
| 0.06224 | 6.14335 | 1 | 6.14335 | 16.805412 | 0.00022426 | 0.01575587 |
| 0.06924 | 3.66108 | 1 | 3.66108 | 16.824478 | 0.00022213 | 0.01575587 |
| 0.05727 | 999 | 1 | 999 | 16.782034 | 0.0002269 | 0.01586411 |
| 0.04257 | 999 | 1 | 999 | 16.550866 | 0.0002547 | 0.01763751 |
| 0.03613 | 2.54666 | 1 | 2.54666 | 16.4898 | 0.00026259 | 0.01801198 |
| 0.02685 | 59.45748 | 1 | 59.45748 | 16.44788 | 0.00026816 | 0.01830674 |
| 0.02098 | 12.95607 | 1 | 12.95607 | 16.412692 | 0.00027292 | 0.0185442 |
| 0.03966 | 40.35684 | 1 | 40.35684 | 16.381042 | 0.00027727 | 0.01875196 |
| 0.12577 | 125.85753 | 1 | 125.85753 | 16.339154 | 0.00028314 | 0.01905978 |
| 0.08819 | 999 | 1 | 999 | 16.277034 | 0.00029207 | 0.01947986 |
| 0.10216 | 18.38683 | 1 | 18.38683 | 16.280126 | 0.00029162 | 0.01947986 |
| 0.15678 | 124.92394 | 1 | 124.92394 | 16.267716 | 0.00029343 | 0.01948105 |
| 0.03209 | 55.77957 | 1 | 55.77957 | 16.176292 | 0.00030716 | 0.02024563 |
| 0.02658 | 11.18623 | 1 | 11.18623 | 16.172458 | 0.00030775 | 0.02024563 |
| 0.11485 | 999 | 1 | 999 | 16.11936 | 0.00031603 | 0.02060303 |
| 0.14326 | 223.813 | 1 | 223.813 | 16.089864 | 0.00032072 | 0.02081537 |
| 0.02232 | 283.96069 | 1 | 283.96069 | 15.992918 | 0.00033665 | 0.02165499 |
| 0.04222 | 30.86904 | 1 | 30.86904 | 15.973362 | 0.00033996 | 0.02168936 |
| 0.04831 | 3.72585 | 1 | 3.72585 | 15.963256 | 0.00034168 | 0.02168936 |
| 0.01284 | 999 | 1 | 999 | 15.9541 | 0.00034325 | 0.02169373 |
| 0.08405 | 13.78853 | 1 | 13.78853 | 15.897988 | 0.00035302 | 0.02211782 |
| 0.05665 | 999 | 1 | 999 | 15.877434 | 0.00035666 | 0.02224998 |
| 0.01475 | 7.4627 | 1 | 7.4627 | 15.840982 | 0.00036322 | 0.02246556 |
| 0.03058 | 4.43771 | 1 | 4.43771 | 15.847558 | 0.00036203 | 0.02246556 |
| 0.12485 | 31.27486 | 1 | 31.27486 | 15.806206 | 0.00036959 | 0.02270165 |
| 0.10375 | 1.1637 | 1 | 1.1637 | 15.709878 | 0.00038783 | 0.02358441 |
| 0.02185 | 999 | 1 | 999 | 15.696234 | 0.00039049 | 0.02364649 |
| 0.03809 | 65.11404 | 1 | 65.11404 | 15.641426 | 0.00040134 | 0.0242022 |
| 0.06916 | 8.528 | 1 | 8.528 | 15.615168 | 0.00040664 | 0.02431938 |
| 0.09502 | 999 | 1 | 999 | 15.535632 | 0.00042314 | 0.02499613 |
| 0.03775 | 101.69482 | 1 | 101.69482 | 15.53775 | 0.00042269 | 0.02499613 |
| 0.07915 | 12.21638 | 1 | 12.21638 | 15.542014 | 0.00042179 | 0.02499613 |
| 0.07948 | 2.11868 | 1 | 2.11868 | 15.521326 | 0.00042617 | 0.02507323 |
| 0 | 999 | 1 | 999 | 15.40507 | 0.00045168 | 0.02635958 |

| | | | | | | |
|---|---|---|---|---|---|---|
| 0.06149 | 3.01169 | 1 | 3.01169 | 15.371976 | 0.00045922 | 0.02669175 |
| 0.018 | 48.43523 | 1 | 48.43523 | 15.357208 | 0.00046262 | 0.02678201 |
| 0.06294 | 1.24469 | 1 | 1.24469 | 15.313034 | 0.00047295 | 0.02727104 |
| 0.15825 | 376.34906 | 1 | 376.34906 | 15.217676 | 0.00049605 | 0.02848929 |
| 0.0374 | 122.12507 | 1 | 122.12507 | 15.084046 | 0.00053032 | 0.02986527 |
| 0.07575 | 671.29564 | 1 | 671.29564 | 15.046682 | 0.00054032 | 0.03019349 |
| 0.04315 | 5.30074 | 1 | 5.30074 | 15.054382 | 0.00053825 | 0.03019349 |
| 0.04521 | 25.48758 | 1 | 25.48758 | 15.03273 | 0.00054411 | 0.03028791 |
| 0.11295 | 1.19145 | 1 | 1.19145 | 15.021336 | 0.00054722 | 0.03034425 |
| 0.09218 | 4.81782 | 1 | 4.81782 | 14.983408 | 0.00055769 | 0.03080715 |
| 0.10035 | 55.95843 | 1 | 55.95843 | 14.881656 | 0.0005868 | 0.0322918 |
| 0.01351 | 16.23493 | 1 | 16.23493 | 14.848284 | 0.00059667 | 0.03271077 |
| 0.09276 | 11.73228 | 1 | 11.73228 | 14.824278 | 0.00060388 | 0.03298083 |
| 0.09187 | 27.50514 | 1 | 27.50514 | 14.804226 | 0.00060996 | 0.03318792 |
| 0.02475 | 71.22023 | 1 | 71.22023 | 14.660894 | 0.00065528 | 0.03552014 |
| 0.04585 | 67.47485 | 1 | 67.47485 | 14.6489 | 0.00065922 | 0.03560046 |
| 0.08096 | 14.62137 | 1 | 14.62137 | 14.633424 | 0.00066434 | 0.03561367 |
| 0.03497 | 1.1535 | 1 | 1.1535 | 14.613874 | 0.00067087 | 0.03582835 |
| 0.13566 | 100.00897 | 1 | 100.00897 | 14.459346 | 0.00072476 | 0.03856404 |
| 0.11397 | 23.33806 | 1 | 23.33806 | 14.402766 | 0.00074555 | 0.03952529 |
| 0.01811 | 998.99964 | 1 | 998.99964 | 14.393904 | 0.00074886 | 0.03955592 |
| 0.02144 | 999 | 1 | 999 | 14.357578 | 0.00076259 | 0.04013446 |
| 0.01354 | 153.45325 | 1 | 153.45325 | 14.277814 | 0.00079362 | 0.04146588 |
| 0.12229 | 8.36925 | 1 | 8.36925 | 14.221676 | 0.00081621 | 0.04249287 |
| 0.03833 | 1.04498 | 1 | 1.04498 | 14.197758 | 0.00082603 | 0.04271989 |
| 0.15681 | 1.16924 | 1 | 1.16924 | 14.1398 | 0.00085032 | 0.04364062 |
| 0.04921 | 190.48833 | 1 | 190.48833 | 14.129366 | 0.00085477 | 0.04371387 |
| 0.05924 | 155.07878 | 1 | 155.07878 | 14.121784 | 0.00085801 | 0.0437254 |
| 0.06023 | 3.87874 | 1 | 3.87874 | 14.081392 | 0.00087552 | 0.0444609 |
| 0 | 999 | 1 | 999 | 14.044586 | 0.00089178 | 0.04512835 |
| 0.0127 | 75.18916 | 1 | 75.18916 | 14.011732 | 0.00090655 | 0.04559768 |
| 0.11206 | 475.12984 | 1 | 475.12984 | 13.9664 | 0.00092733 | 0.04644035 |
| 0.12788 | 27.86225 | 1 | 27.86225 | 13.926368 | 0.00094608 | 0.04721589 |
| 0.06335 | 2.70551 | 1 | 2.70551 | 13.909754 | 0.00095397 | 0.04744614 |
| 0.03623 | 43.39786 | 1 | 43.39786 | 13.85064 | 0.00098259 | 0.04853585 |
| 0.12793 | 999 | 1 | 999 | 13.837054 | 0.00098929 | 0.04860722 |
| 0.0729 | 33.93082 | 1 | 33.93082 | 13.834096 | 0.00099075 | 0.04860722 |
| 0.0978 | 17.83787 | 1 | 17.83787 | 13.719006 | 0.00104944 | 0.05096805 |
| 0.11307 | 11.42763 | 1 | 11.42763 | 13.59122 | 0.00111868 | 0.05396863 |
| 0.0659 | 74.16876 | 1 | 74.16876 | 13.56115 | 0.00113562 | 0.05457672 |
| 0.05215 | 15.63482 | 1 | 15.63482 | 13.555522 | 0.00113882 | 0.05457672 |
| 0.1 | 4.9306 | 1 | 4.9306 | 13.435382 | 0.00120933 | 0.05757432 |
| 0.03138 | 140.67479 | 1 | 140.67479 | 13.426018 | 0.001215 | 0.05765486 |
| 0.05819 | 10.20114 | 1 | 10.20114 | 13.412622 | 0.00122317 | 0.05785265 |
| 0.1394 | 174.69388 | 1 | 174.69388 | 13.303668 | 0.00129165 | 0.06049859 |

| | | | | | | |
|---|---|---|---|---|---|---|
| 0.17174 | 53.85106 | 1 | 53.85106 | 13.254 | 0.00132413 | 0.06122724 |
| 0.10751 | 2.66061 | 1 | 2.66061 | 13.257728 | 0.00132166 | 0.06122724 |
| 0.04889 | 35.17336 | 1 | 35.17336 | 13.24313 | 0.00133135 | 0.06136486 |
| 0.06863 | 2.68054 | 1 | 2.68054 | 13.197826 | 0.00136185 | 0.06257149 |
| 0.0409 | 612.51545 | 1 | 612.51545 | 13.152686 | 0.00139293 | 0.06379726 |
| 0.03986 | 69.61348 | 1 | 69.61348 | 13.084832 | 0.001441 | 0.06579064 |
| 0.03002 | 105.3077 | 1 | 105.3077 | 13.063424 | 0.00145651 | 0.06628953 |
| 0.09585 | 16.38372 | 1 | 16.38372 | 13.051998 | 0.00146486 | 0.06641211 |
| 0.12095 | 4.40709 | 1 | 4.40709 | 13.04719 | 0.00146838 | 0.06641211 |
| 0.04175 | 13.94685 | 1 | 13.94685 | 13.018654 | 0.00148948 | 0.0671566 |
| 0.03267 | 1.92806 | 1 | 1.92806 | 12.998996 | 0.00150419 | 0.06760932 |
| 0.04817 | 26.03835 | 1 | 26.03835 | 12.954194 | 0.00153827 | 0.06892687 |
| 0.04278 | 999 | 1 | 999 | 12.942886 | 0.00154699 | 0.06910374 |
| 0.06358 | 999 | 1 | 999 | 12.866276 | 0.0016074 | 0.07092648 |
| 0.10159 | 120.13635 | 1 | 120.13635 | 12.875326 | 0.00160014 | 0.07092648 |
| 0.05573 | 25.94161 | 1 | 25.94161 | 12.87855 | 0.00159756 | 0.07092648 |
| 0.03905 | 1.35346 | 1 | 1.35346 | 12.868194 | 0.00160586 | 0.07092648 |
| 0.11543 | 8.65176 | 1 | 8.65176 | 12.829792 | 0.00163699 | 0.07168188 |
| 0.02368 | 4.55931 | 1 | 4.55931 | 12.833616 | 0.00163386 | 0.07168188 |
| 0.11631 | 1.86292 | 1 | 1.86292 | 12.826878 | 0.00163938 | 0.07168188 |
| 0.06603 | 17.91499 | 1 | 17.91499 | 12.812982 | 0.00165081 | 0.07183666 |
| 0.0481 | 13.61794 | 1 | 13.61794 | 12.810516 | 0.00165284 | 0.07183666 |
| 0.15529 | 12.48373 | 1 | 12.48373 | 12.79239 | 0.00166789 | 0.07227364 |
| 0.06838 | 107.7414 | 1 | 107.7414 | 12.7705 | 0.00168625 | 0.0728509 |
| 0.08048 | 31.60343 | 1 | 31.60343 | 12.758624 | 0.00169629 | 0.07306666 |
| 0.06202 | 210.51053 | 1 | 210.51053 | 12.65134 | 0.00178977 | 0.07644605 |
| 0.0123 | 32.86741 | 1 | 32.86741 | 12.65042 | 0.00179059 | 0.07644605 |
| 0.09147 | 17.01717 | 1 | 17.01717 | 12.607782 | 0.00182917 | 0.07786362 |
| 0.13702 | 999 | 1 | 999 | 12.587658 | 0.00184767 | 0.07819109 |
| 0.05502 | 18.82663 | 1 | 18.82663 | 12.538158 | 0.00189397 | 0.07991679 |
| 0.16434 | 124.79284 | 1 | 124.79284 | 12.517002 | 0.00191411 | 0.08029842 |
| 0.0426 | 16.23174 | 1 | 16.23174 | 12.487332 | 0.00194272 | 0.0810288 |
| 0.03933 | 62.56399 | 1 | 62.56399 | 12.460714 | 0.00196875 | 0.08141058 |
| 0.02028 | 125.36571 | 1 | 125.36571 | 12.438042 | 0.00199119 | 0.08210412 |
| 0.01827 | 433.26691 | 1 | 433.26691 | 12.429344 | 0.00199987 | 0.0822277 |
| 0.08044 | 999 | 1 | 999 | 12.403832 | 0.00202555 | 0.08294116 |
| 0.15197 | 12.69073 | 1 | 12.69073 | 12.400734 | 0.00202869 | 0.08294116 |
| 0.16114 | 998.99966 | 1 | 998.99966 | 12.379364 | 0.00205048 | 0.08312767 |
| 0.03673 | 4.71571 | 1 | 4.71571 | 12.381874 | 0.00204791 | 0.08312767 |
| 0.05125 | 40.1272 | 1 | 40.1272 | 12.335734 | 0.0020957 | 0.0844877 |
| 0.12114 | 2.89484 | 1 | 2.89484 | 12.290652 | 0.00214348 | 0.08617373 |
| 0.05791 | 19.16084 | 1 | 19.16084 | 12.247274 | 0.00219047 | 0.08781923 |
| 0.09678 | 9.853 | 1 | 9.853 | 12.233078 | 0.00220608 | 0.08820046 |
| 0.04136 | 3.15026 | 1 | 3.15026 | 12.148236 | 0.00230168 | 0.09126616 |
| 0.07688 | 11.07107 | 1 | 11.07107 | 12.128734 | 0.00232423 | 0.09165821 |

| | | | | | | |
|---|---|---|---|---|---|---|
| 0.12117 | 999 | 1 | 999 | 12.119958 | 0.00233445 | 0.09181112 |
| 0.02097 | 999 | 1 | 999 | 12.09553 | 0.00236314 | 0.09268751 |
| 0.14702 | 22.18291 | 1 | 22.18291 | 12.04998 | 0.00241758 | 0.09431152 |
| 0.03878 | 158.06266 | 1 | 158.06266 | 11.981706 | 0.00250153 | 0.09706336 |
| 0.06875 | 4.15093 | 1 | 4.15093 | 11.960546 | 0.00252814 | 0.09783346 |
| 0.04894 | 47.49991 | 1 | 47.49991 | 11.9382 | 0.00255654 | 0.09840644 |
| 0.02446 | 1.02005 | 1 | 1.02005 | 11.939026 | 0.00255549 | 0.09840644 |
| 0.13569 | 31.21568 | 1 | 31.21568 | 11.931814 | 0.00256472 | 0.0984593 |
| 0.08801 | 17.28367 | 1 | 17.28367 | 11.859288 | 0.00265943 | 0.10155649 |
| 0.07661 | 287.82063 | 1 | 287.82063 | 11.829138 | 0.00269982 | 0.10255785 |
| 0.10083 | 186.53099 | 1 | 186.53099 | 11.82949 | 0.00269935 | 0.10255785 |
| 0.14477 | 5.97835 | 1 | 5.97835 | 11.802672 | 0.00273579 | 0.10365196 |
| 0.16564 | 122.42617 | 1 | 122.42617 | 11.680618 | 0.00290794 | 0.10903283 |
| 0.10805 | 48.66124 | 1 | 48.66124 | 11.567988 | 0.0030764 | 0.11475461 |
| 0.04769 | 2.74213 | 1 | 2.74213 | 11.532896 | 0.00313086 | 0.11648565 |
| 0.05406 | 78.95414 | 1 | 78.95414 | 11.47446 | 0.00322369 | 0.1196195 |
| 0.04722 | 3.24346 | 1 | 3.24346 | 11.469544 | 0.00323162 | 0.1196195 |
| 0.0507 | 52.50813 | 1 | 52.50813 | 11.463176 | 0.00324192 | 0.11969485 |
| 0.01801 | 999 | 1 | 999 | 11.43195 | 0.00329294 | 0.12096116 |
| 0.08005 | 4.61619 | 1 | 4.61619 | 11.381238 | 0.0033775 | 0.12375337 |
| 0.09462 | 1.09733 | 1 | 1.09733 | 11.327656 | 0.00346921 | 0.12647329 |
| 0.02795 | 15.29564 | 1 | 15.29564 | 11.303356 | 0.00351162 | 0.12737761 |
| 0.05078 | 999 | 1 | 999 | 11.243268 | 0.00361872 | 0.13028773 |
| 0.06051 | 84.37335 | 1 | 84.37335 | 11.243196 | 0.00361885 | 0.13028773 |
| 0.05396 | 301.87179 | 1 | 301.87179 | 11.228702 | 0.00364517 | 0.13090971 |
| 0.09769 | 35.98653 | 1 | 35.98653 | 11.124528 | 0.00384007 | 0.13621905 |
| 0.00143 | 31.56391 | 1 | 31.56391 | 11.141338 | 0.00380793 | 0.13621905 |
| 0.02403 | 38.49306 | 1 | 38.49306 | 11.115836 | 0.0038568 | 0.1364118 |
| 0.0161 | 11.10172 | 1 | 11.10172 | 11.11192 | 0.00386436 | 0.1364118 |
| 0.01482 | 307.92794 | 1 | 307.92794 | 11.093996 | 0.00389914 | 0.13697166 |
| 0.02555 | 76.8992 | 1 | 76.8992 | 11.095588 | 0.00389604 | 0.13697166 |
| 0.048 | 123.18276 | 1 | 123.18276 | 11.056528 | 0.00397288 | 0.13888766 |
| 0.05251 | 999 | 1 | 999 | 11.039598 | 0.00400665 | 0.13939493 |
| 0.09101 | 15.86935 | 1 | 15.86935 | 11.041068 | 0.00400371 | 0.13939493 |
| 0.03588 | 346.72965 | 1 | 346.72965 | 11.008674 | 0.00406909 | 0.14122752 |
| 0.04201 | 999 | 1 | 999 | 10.979494 | 0.00412889 | 0.14157226 |
| 0.05694 | 29.82929 | 1 | 29.82929 | 10.991092 | 0.00410501 | 0.14157226 |
| 0.10338 | 10.16899 | 1 | 10.16899 | 10.98365 | 0.00412032 | 0.14157226 |
| 0.11297 | 5.78174 | 1 | 5.78174 | 10.98968 | 0.00410791 | 0.14157226 |
| 0.10702 | 999 | 1 | 999 | 10.9097 | 0.00427552 | 0.14584891 |
| 0.07363 | 999 | 1 | 999 | 10.866156 | 0.00436963 | 0.14743735 |
| 0.04264 | 102.20159 | 1 | 102.20159 | 10.856568 | 0.00439062 | 0.14743735 |
| 0.14481 | 64.16461 | 1 | 64.16461 | 10.87179 | 0.00435733 | 0.14743735 |
| 0.21308 | 9.99046 | 1 | 9.99046 | 10.859124 | 0.00438502 | 0.14743735 |
| 0.06432 | 2.12039 | 1 | 2.12039 | 10.862298 | 0.00437806 | 0.14743735 |

| | | | | | | |
|---|---|---|---|---|---|---|
| 0.03122 | 1.89792 | 1 | 1.89792 | 10.871066 | 0.00435891 | 0.14743735 |
| 0.07397 | 369.05866 | 1 | 369.05866 | 10.818028 | 0.00447605 | 0.14892389 |
| 0.01856 | 145.46287 | 1 | 145.46287 | 10.830392 | 0.00444847 | 0.14892389 |
| 0.07703 | 31.87883 | 1 | 31.87883 | 10.825606 | 0.00445912 | 0.14892389 |
| 0.06854 | 1.97193 | 1 | 1.97193 | 10.820258 | 0.00447106 | 0.14892389 |
| 0.10291 | 37.43064 | 1 | 37.43064 | 10.768198 | 0.00458897 | 0.15163518 |
| 0.10858 | 25.98265 | 1 | 25.98265 | 10.771748 | 0.00458083 | 0.15163518 |
| 0.13576 | 999 | 1 | 999 | 10.728444 | 0.0046811 | 0.1539763 |
| 0.0263 | 4.45852 | 1 | 4.45852 | 10.701302 | 0.00474506 | 0.15565432 |
| 0.02434 | 88.01468 | 1 | 88.01468 | 10.67968 | 0.00479664 | 0.15627123 |
| 0.08344 | 999 | 1 | 999 | 10.666982 | 0.00482719 | 0.15629509 |
| 0.0544 | 981.08137 | 1 | 981.08137 | 10.64705 | 0.00487554 | 0.15715735 |
| 0.09388 | 37.18015 | 1 | 37.18015 | 10.62086 | 0.0049398 | 0.15887501 |
| 0.0274 | 3.09003 | 1 | 3.09003 | 10.60567 | 0.00497746 | 0.15973131 |
| 0.06996 | 85.56464 | 1 | 85.56464 | 10.596034 | 0.0050015 | 0.16014765 |
| 0.06107 | 4.5318 | 1 | 4.5318 | 10.569778 | 0.00506759 | 0.16190573 |
| 0.04176 | 119.82626 | 1 | 119.82626 | 10.525156 | 0.00518193 | 0.16519394 |
| 0.03388 | 47.55668 | 1 | 47.55668 | 10.493636 | 0.00526424 | 0.16635236 |
| 0.06493 | 32.40253 | 1 | 32.40253 | 10.496926 | 0.00525559 | 0.16635236 |
| 0.0961 | 1.65658 | 1 | 1.65658 | 10.498356 | 0.00525183 | 0.16635236 |
| 0.0683 | 999 | 1 | 999 | 10.472462 | 0.00532027 | 0.16775661 |
| 0.11887 | 999 | 1 | 999 | 10.421074 | 0.00545874 | 0.17100512 |
| 0.12544 | 1.21278 | 1 | 1.21278 | 10.381398 | 0.00556811 | 0.17367954 |
| 0.04093 | 2.16918 | 1 | 2.16918 | 10.346342 | 0.00566657 | 0.17637052 |
| 0.03326 | 2.59027 | 1 | 2.59027 | 10.337542 | 0.00569156 | 0.17676811 |
| 0.05308 | 999 | 1 | 999 | 10.315098 | 0.00575579 | 0.17799903 |
| 0.06378 | 999 | 1 | 999 | 10.307548 | 0.00577756 | 0.17829128 |
| 0.05647 | 998.99738 | 1 | 998.99738 | 10.301418 | 0.00579529 | 0.17845808 |
| 0.08514 | 174.1572 | 1 | 174.1572 | 10.285272 | 0.00584227 | 0.17910263 |
| 0.03407 | 11.08324 | 1 | 11.08324 | 10.284886 | 0.0058434 | 0.17910263 |
| 0.12098 | 999 | 1 | 999 | 10.276324 | 0.00586847 | 0.17918631 |
| 0.0359 | 11.47639 | 1 | 11.47639 | 10.26133 | 0.00591263 | 0.17977616 |
| 0.04294 | 1.95686 | 1 | 1.95686 | 10.235864 | 0.00598839 | 0.18105318 |
| 0.09582 | 1.35576 | 1 | 1.35576 | 10.230248 | 0.00600523 | 0.18107029 |
| 0.14665 | 1.88613 | 1 | 1.88613 | 10.203706 | 0.00608546 | 0.18272792 |
| 0.08141 | 470.04981 | 1 | 470.04981 | 10.154672 | 0.0062365 | 0.18500706 |
| 0.08243 | 145.1962 | 1 | 145.1962 | 10.150158 | 0.00625059 | 0.18500706 |
| 0.07017 | 18.02526 | 1 | 18.02526 | 10.156778 | 0.00622994 | 0.18500706 |
| 0.06538 | 44.75437 | 1 | 44.75437 | 10.083386 | 0.0064628 | 0.18972833 |
| 0.16237 | 269.20165 | 1 | 269.20165 | 10.069762 | 0.00650697 | 0.19038372 |
| 0.05578 | 1.96634 | 1 | 1.96634 | 10.068392 | 0.00651143 | 0.19038372 |
| 0.0456 | 1.24035 | 1 | 1.24035 | 10.064234 | 0.00652498 | 0.19039531 |
| 0.06577 | 44.83125 | 1 | 44.83125 | 10.050996 | 0.00656831 | 0.19127408 |
| 0.01769 | 1.55011 | 1 | 1.55011 | 10.035406 | 0.00661971 | 0.1923838 |
| 0.12258 | 58.44581 | 1 | 58.44581 | 10.020244 | 0.00667009 | 0.19259822 |

| | | | | | | |
|---|---|---|---|---|---|---|
| 0.03694 | 52.51939 | 1 | 52.51939 | 10.016168 | 0.0066837 | 0.19259822 |
| 0.08171 | 15.00894 | 1 | 15.00894 | 10.013198 | 0.00669363 | 0.19259822 |
| 0.09044 | 11.60218 | 1 | 11.60218 | 10.023546 | 0.00665909 | 0.19259822 |
| 0.02598 | 1.19543 | 1 | 1.19543 | 10.017098 | 0.00668059 | 0.19259822 |
| 0.13281 | 999 | 1 | 999 | 10.003154 | 0.00672733 | 0.19259883 |
| 0.0278 | 999 | 1 | 999 | 9.99895 | 0.00674149 | 0.19259883 |
| 0.04191 | 115.5878 | 1 | 115.5878 | 9.997818 | 0.0067453 | 0.19259883 |
| 0.02857 | 2.66658 | 1 | 2.66658 | 9.99735 | 0.00674688 | 0.19259883 |
| 0.03253 | 999 | 1 | 999 | 9.920722 | 0.0070104 | 0.19894406 |
| 0.07728 | 7.84534 | 1 | 7.84534 | 9.900064 | 0.00708318 | 0.20022441 |
| 0.04074 | 998.99996 | 1 | 998.99996 | 9.880864 | 0.00715151 | 0.20136922 |
| 0.04897 | 31.02584 | 1 | 31.02584 | 9.880952 | 0.00715119 | 0.20136922 |
| 0.19672 | 999 | 1 | 999 | 9.846312 | 0.00727613 | 0.20448048 |
| 0.06907 | 31.26773 | 1 | 31.26773 | 9.823794 | 0.00735852 | 0.2060283 |
| 0.11125 | 4.58412 | 1 | 4.58412 | 9.823478 | 0.00735968 | 0.2060283 |
| 0.26057 | 999 | 1 | 999 | 9.806074 | 0.007424 | 0.20723359 |
| 0.0457 | 31.96498 | 1 | 31.96498 | 9.80024 | 0.00744569 | 0.20723359 |
| 0.09295 | 1.68181 | 1 | 1.68181 | 9.803916 | 0.00743202 | 0.20723359 |
| 0.0968 | 54.20788 | 1 | 54.20788 | 9.780952 | 0.00751784 | 0.20843744 |
| 0.06781 | 11.61107 | 1 | 11.61107 | 9.77715 | 0.00753215 | 0.20843744 |
| 0.05979 | 59.96101 | 1 | 59.96101 | 9.763196 | 0.00758488 | 0.20869966 |
| 0.01377 | 6.44381 | 1 | 6.44381 | 9.768892 | 0.00756331 | 0.20869966 |
| 0.16563 | 102.09619 | 1 | 102.09619 | 9.73454 | 0.00769434 | 0.21011361 |
| 0.1006 | 6.52718 | 1 | 6.52718 | 9.741078 | 0.00766923 | 0.21011361 |
| 0.10881 | 22.06501 | 1 | 22.06501 | 9.668072 | 0.00795435 | 0.21478236 |
| 0.0234 | 29.75159 | 1 | 29.75159 | 9.6619 | 0.00797894 | 0.215045 |
| 0.10736 | 59.52722 | 1 | 59.52722 | 9.646738 | 0.00803966 | 0.21627035 |
| 0.03616 | 32.23363 | 1 | 32.23363 | 9.642396 | 0.00805713 | 0.21627035 |
| 0.03196 | 1.0209 | 1 | 1.0209 | 9.598678 | 0.00823519 | 0.21990385 |
| 0.08583 | 188.17732 | 1 | 188.17732 | 9.57697 | 0.00832506 | 0.22148639 |
| 0.14172 | 33.89171 | 1 | 33.89171 | 9.579806 | 0.00831326 | 0.22148639 |
| 0.13529 | 999 | 1 | 999 | 9.565526 | 0.00837283 | 0.22194141 |
| 0.06201 | 20.61938 | 1 | 20.61938 | 9.566316 | 0.00836953 | 0.22194141 |
| 0.09768 | 88.44064 | 1 | 88.44064 | 9.559196 | 0.00839937 | 0.22223794 |
| 0.06776 | 98.36575 | 1 | 98.36575 | 9.530132 | 0.00852233 | 0.22426113 |
| 0.04759 | 5.20163 | 1 | 5.20163 | 9.513742 | 0.00859245 | 0.22569614 |
| 0.03309 | 3.00477 | 1 | 3.00477 | 9.49575 | 0.0086701 | 0.2273231 |
| 0.06827 | 998.99993 | 1 | 998.99993 | 9.483016 | 0.00872548 | 0.22836139 |
| 0.08041 | 984.42411 | 1 | 984.42411 | 9.467218 | 0.00879467 | 0.22904606 |
| 0.03155 | 3.45567 | 1 | 3.45567 | 9.466208 | 0.00879912 | 0.22904606 |
| 0.02394 | 999 | 1 | 999 | 9.443122 | 0.00890127 | 0.23046176 |
| 0.10268 | 1.37094 | 1 | 1.37094 | 9.443474 | 0.00889971 | 0.23046176 |
| 0.11683 | 105.74692 | 1 | 105.74692 | 9.42466 | 0.00898382 | 0.23218364 |
| 0.03028 | 2.25735 | 1 | 2.25735 | 9.390174 | 0.00914007 | 0.23413144 |
| 0.11329 | 999 | 1 | 999 | 9.379538 | 0.00918881 | 0.23454653 |

| | | | | | | |
|---|---|---|---|---|---|---|
| 0.02369 | 30.22265 | 1 | 30.22265 | 9.381884 | 0.00917804 | 0.23454653 |
| 0.0126 | 29.71757 | 1 | 29.71757 | 9.367978 | 0.00924207 | 0.23500905 |
| 0.10333 | 1.42878 | 1 | 1.42878 | 9.36507 | 0.00925552 | 0.23500905 |
| 0.08413 | 999 | 1 | 999 | 9.32972 | 0.00942057 | 0.23736493 |
| 0.00198 | 999 | 1 | 999 | 9.324178 | 0.00944671 | 0.23736493 |
| 0.02341 | 277.36599 | 1 | 277.36599 | 9.335132 | 0.00939511 | 0.23736493 |
| 0.09359 | 5.2776 | 1 | 5.2776 | 9.33378 | 0.00940146 | 0.23736493 |
| 0.09538 | 999 | 1 | 999 | 9.30382 | 0.00954336 | 0.23937782 |
| 0.03645 | 42.64501 | 1 | 42.64501 | 9.277252 | 0.00967098 | 0.241741 |
| 0.18457 | 3.0714 | 1 | 3.0714 | 9.26077 | 0.009751 | 0.2427604 |
| 0.04951 | 1.38731 | 1 | 1.38731 | 9.2585 | 0.00976208 | 0.2427604 |
| 0.05707 | 73.21467 | 1 | 73.21467 | 9.243704 | 0.00983457 | 0.24330883 |
| 0.02352 | 28.63995 | 1 | 28.63995 | 9.24715 | 0.00981764 | 0.24330883 |
| 0.0199 | 2.09261 | 1 | 2.09261 | 9.244088 | 0.00983268 | 0.24330883 |
| 0.1155 | 50.45716 | 1 | 50.45716 | 9.239968 | 0.00985295 | 0.24334778 |
| 0.06791 | 54.08963 | 1 | 54.08963 | 9.222982 | 0.00993699 | 0.24429957 |
| 0.01784 | 8.49681 | 1 | 8.49681 | 9.221948 | 0.00994213 | 0.24429957 |
| 0.05461 | 10.50841 | 1 | 10.50841 | 9.212692 | 0.00998825 | 0.24501682 |
| 0.05092 | 7.87642 | 1 | 7.87642 | 9.182376 | 0.0101408 | 0.24833816 |
| 0.05331 | 5.51297 | 1 | 5.51297 | 9.173036 | 0.01018827 | 0.24907916 |
| 0.05473 | 81.49694 | 1 | 81.49694 | 9.145402 | 0.01033002 | 0.25039249 |
| 0.10718 | 33.85903 | 1 | 33.85903 | 9.15101 | 0.0103011 | 0.25039249 |
| 0.12099 | 1.71208 | 1 | 1.71208 | 9.152342 | 0.01029424 | 0.25039249 |
| 0.0249 | 89.19836 | 1 | 89.19836 | 9.137032 | 0.01037334 | 0.25064004 |
| 0.12215 | 47.85573 | 1 | 47.85573 | 9.121992 | 0.01045164 | 0.25181948 |
| 0.07093 | 21.37225 | 1 | 21.37225 | 9.120976 | 0.01045695 | 0.25181948 |
| 0.07254 | 86.32096 | 1 | 86.32096 | 9.114636 | 0.01049016 | 0.25219938 |
| 0.12215 | 999 | 1 | 999 | 9.108166 | 0.01052415 | 0.25259697 |
| 0.01691 | 1.09688 | 1 | 1.09688 | 9.07288 | 0.01071147 | 0.25624319 |
| 0.1021 | 998.99988 | 1 | 998.99988 | 9.06263 | 0.01076651 | 0.25628895 |
| 0.01404 | 502.05989 | 1 | 502.05989 | 9.062656 | 0.01076637 | 0.25628895 |
| 0.03689 | 18.92263 | 1 | 18.92263 | 9.067774 | 0.01073885 | 0.25628895 |
| 0.11319 | 8.80167 | 1 | 8.80167 | 9.05641 | 0.01080004 | 0.2566651 |
| 0.05797 | 1.14154 | 1 | 1.14154 | 9.02497 | 0.01097116 | 0.25888357 |
| 0.05958 | 7.27379 | 1 | 7.27379 | 9.007506 | 0.01106738 | 0.25918807 |
| 0.10104 | 16.89072 | 1 | 16.89072 | 8.988744 | 0.01117169 | 0.26120829 |
| 0.02513 | 95.54675 | 1 | 95.54675 | 8.946312 | 0.01141125 | 0.26323133 |
| 0.05607 | 55.61127 | 1 | 55.61127 | 8.955352 | 0.01135978 | 0.26323133 |
| 0.06394 | 7.40269 | 1 | 7.40269 | 8.9465 | 0.01141017 | 0.26323133 |
| 0.1236 | 2.19798 | 1 | 2.19798 | 8.948054 | 0.01140131 | 0.26323133 |
| 0.08567 | 1.38161 | 1 | 1.38161 | 8.92139 | 0.01155433 | 0.26543781 |
| 0.05945 | 999 | 1 | 999 | 8.890018 | 0.011737 | 0.26763428 |
| 0.03588 | 220.8447 | 1 | 220.8447 | 8.894648 | 0.01170986 | 0.26763428 |
| 0.04952 | 4.06065 | 1 | 4.06065 | 8.889098 | 0.0117424 | 0.26763428 |
| 0.27671 | 999 | 1 | 999 | 8.872326 | 0.01184129 | 0.26904072 |

| | | | | | | |
|---|---|---|---|---|---|---|
| 0.04268 | 1.31193 | 1 | 1.31193 | 8.843224 | 0.01201485 | 0.27170454 |
| 0.04365 | 19.09303 | 1 | 19.09303 | 8.838072 | 0.01204584 | 0.27198038 |
| 0.15708 | 297.1907 | 1 | 297.1907 | 8.819566 | 0.01215782 | 0.27408111 |
| 0.07927 | 24.83921 | 1 | 24.83921 | 8.814584 | 0.01218814 | 0.27433738 |
| 0.03613 | 999 | 1 | 999 | 8.777352 | 0.01241716 | 0.27832873 |
| 0.06643 | 5.34974 | 1 | 5.34974 | 8.770326 | 0.01246086 | 0.27874186 |
| 0.04334 | 403.2326 | 1 | 403.2326 | 8.762958 | 0.01250685 | 0.27882631 |
| 0.09518 | 15.43528 | 1 | 15.43528 | 8.761072 | 0.01251865 | 0.27882631 |
| 0.03752 | 1.8986 | 1 | 1.8986 | 8.760468 | 0.01252243 | 0.27882631 |
| 0.06132 | 2.18978 | 1 | 2.18978 | 8.756402 | 0.01254791 | 0.27896456 |
| 0.02307 | 16.61265 | 1 | 16.61265 | 8.749202 | 0.01259317 | 0.27927866 |
| 0.11998 | 4.49258 | 1 | 4.49258 | 8.74563 | 0.01261568 | 0.27927866 |
| 0.0677 | 999 | 1 | 999 | 8.716122 | 0.01280319 | 0.28011281 |
| 0.05042 | 999 | 1 | 999 | 8.730804 | 0.01270954 | 0.28011281 |
| 0.05708 | 999 | 1 | 999 | 8.713884 | 0.01281752 | 0.28011281 |
| 0.03894 | 12.48879 | 1 | 12.48879 | 8.71692 | 0.01279808 | 0.28011281 |
| 0.10006 | 1.77896 | 1 | 1.77896 | 8.711656 | 0.01283181 | 0.28011281 |
| 0.06756 | 51.80915 | 1 | 51.80915 | 8.68722 | 0.01298955 | 0.28206636 |
| 0.07327 | 34.49757 | 1 | 34.49757 | 8.69114 | 0.01296412 | 0.28206636 |
| 0.06888 | 31.07323 | 1 | 31.07323 | 8.685726 | 0.01299926 | 0.28206636 |
| 0.05676 | 392.3463 | 1 | 392.3463 | 8.64368 | 0.01327543 | 0.28719785 |
| 0.04406 | 10.19391 | 1 | 10.19391 | 8.6387 | 0.01330853 | 0.28748414 |
| 0.09326 | 1.18828 | 1 | 1.18828 | 8.619412 | 0.0134375 | 0.28983745 |
| 0.04967 | 1.04967 | 1 | 1.04967 | 8.585228 | 0.01366915 | 0.29269331 |
| 0.04985 | 1.60786 | 1 | 1.60786 | 8.578328 | 0.01371639 | 0.29323083 |
| 0.05825 | 998.99961 | 1 | 998.99961 | 8.549284 | 0.01391703 | 0.29664385 |
| 0.2273 | 136.09224 | 1 | 136.09224 | 8.52098 | 0.01411538 | 0.29982545 |
| 0.02171 | 13.16118 | 1 | 13.16118 | 8.522912 | 0.01410176 | 0.29982545 |
| 0.08647 | 9.08339 | 1 | 9.08339 | 8.513274 | 0.01416988 | 0.29982545 |
| 0.07805 | 1.3402 | 1 | 1.3402 | 8.515958 | 0.01415087 | 0.29982545 |
| 0.0675 | 33.04021 | 1 | 33.04021 | 8.508878 | 0.01420106 | 0.30004653 |
| 0.05248 | 23.03245 | 1 | 23.03245 | 8.48968 | 0.01433803 | 0.30161794 |
| 0.01752 | 1.98012 | 1 | 1.98012 | 8.489644 | 0.01433829 | 0.30161794 |
| 0.08503 | 1.74505 | 1 | 1.74505 | 8.491132 | 0.01432762 | 0.30161794 |
| 0.01178 | 9.0429 | 1 | 9.0429 | 8.48137 | 0.01439773 | 0.30199752 |
| 0.16794 | 106.7392 | 1 | 106.7392 | 8.469234 | 0.01448536 | 0.30339591 |
| 0.03438 | 999 | 1 | 999 | 8.459734 | 0.01455433 | 0.30429767 |
| 0.05451 | 29.60092 | 1 | 29.60092 | 8.457518 | 0.01457046 | 0.30429767 |
| 0.03576 | 1.21403 | 1 | 1.21403 | 8.454444 | 0.01459287 | 0.30432659 |
| 0.06449 | 2.32804 | 1 | 2.32804 | 8.449112 | 0.01463183 | 0.30469996 |
| 0.01578 | 999 | 1 | 999 | 8.415242 | 0.01488173 | 0.30818508 |
| 0.07429 | 55.65606 | 1 | 55.65606 | 8.409174 | 0.01492695 | 0.30818508 |
| 0.04292 | 36.55123 | 1 | 36.55123 | 8.410258 | 0.01491886 | 0.30818508 |
| 0.0183 | 9.37678 | 1 | 9.37678 | 8.412626 | 0.01490121 | 0.30818508 |
| 0.22942 | 999 | 1 | 999 | 8.403904 | 0.01496633 | 0.30843234 |

| | | | | | | |
|---|---|---|---|---|---|---|
| 0.05169 | 1.18112 | 1 | 1.18112 | 8.401872 | 0.01498155 | 0.30843234 |
| 0.03115 | 81.45745 | 1 | 81.45745 | 8.398148 | 0.01500947 | 0.30856825 |
| 0.06714 | 82.06888 | 1 | 82.06888 | 8.383994 | 0.01511607 | 0.30987939 |
| 0.08376 | 14.45128 | 1 | 14.45128 | 8.385704 | 0.01510315 | 0.30987939 |
| 0.05886 | 37.80336 | 1 | 37.80336 | 8.380386 | 0.01514336 | 0.30999983 |
| 0.02836 | 999 | 1 | 999 | 8.351878 | 0.01536076 | 0.31356318 |
| 0.13719 | 68.53732 | 1 | 68.53732 | 8.324388 | 0.01557335 | 0.31597732 |
| 0.04855 | 999 | 1 | 999 | 8.31681 | 0.01563247 | 0.31608598 |
| 0.03524 | 32.74837 | 1 | 32.74837 | 8.30309 | 0.01574008 | 0.31608598 |
| 0.05837 | 2.57172 | 1 | 2.57172 | 8.311084 | 0.01567729 | 0.31608598 |
| 0.03349 | 2.07945 | 1 | 2.07945 | 8.302452 | 0.0157451 | 0.31608598 |
| 0.05176 | 1.23375 | 1 | 1.23375 | 8.305772 | 0.01571899 | 0.31608598 |
| 0.12073 | 2.61851 | 1 | 2.61851 | 8.275458 | 0.01595905 | 0.31946802 |
| 0.07196 | 2.16702 | 1 | 2.16702 | 8.24192 | 0.01622893 | 0.32397416 |
| 0.0782 | 3.68971 | 1 | 3.68971 | 8.231016 | 0.01631765 | 0.3252966 |
| 0.04739 | 4.60025 | 1 | 4.60025 | 8.202504 | 0.01655194 | 0.32906074 |
| 0.04987 | 998.99986 | 1 | 998.99986 | 8.194522 | 0.01661813 | 0.3298361 |
| 0.05118 | 22.7761 | 1 | 22.7761 | 8.19231 | 0.01663652 | 0.3298361 |
| 0.0873 | 237.11289 | 1 | 237.11289 | 8.188694 | 0.01666663 | 0.32998096 |
| 0.08927 | 164.37422 | 1 | 164.37422 | 8.177984 | 0.01675612 | 0.33129953 |
| 0.12941 | 204.75928 | 1 | 204.75928 | 8.168072 | 0.01683936 | 0.33203832 |
| 0.10647 | 782.98966 | 1 | 782.98966 | 8.153944 | 0.01695874 | 0.33259066 |
| 0.05357 | 97.63885 | 1 | 97.63885 | 8.156078 | 0.01694065 | 0.33259066 |
| 0.0773 | 44.96784 | 1 | 44.96784 | 8.151974 | 0.01697545 | 0.33259066 |
| 0.02624 | 116.79694 | 1 | 116.79694 | 8.146334 | 0.01702339 | 0.3329453 |
| 0.0353 | 10.99764 | 1 | 10.99764 | 8.143044 | 0.01705142 | 0.33304339 |
| 0.05701 | 15.40087 | 1 | 15.40087 | 8.140338 | 0.0170745 | 0.33304485 |
| 0.08172 | 135.19227 | 1 | 135.19227 | 8.136236 | 0.01710956 | 0.33327946 |
| 0.08266 | 999 | 1 | 999 | 8.124314 | 0.01721185 | 0.33437201 |
| 0.03082 | 86.22639 | 1 | 86.22639 | 8.125942 | 0.01719785 | 0.33437201 |
| 0.16151 | 999 | 1 | 999 | 8.101154 | 0.01741232 | 0.33663236 |
| 0.0978 | 7.22485 | 1 | 7.22485 | 8.10013 | 0.01742124 | 0.33663236 |
| 0.13381 | 2.84798 | 1 | 2.84798 | 8.103626 | 0.01739082 | 0.33663236 |
| 0.04925 | 1.00601 | 1 | 1.00601 | 8.10707 | 0.0173609 | 0.33663236 |
| 0.0961 | 366.86511 | 1 | 366.86511 | 8.083238 | 0.01756901 | 0.33768421 |
| 0.05995 | 59.47057 | 1 | 59.47057 | 8.085774 | 0.01754674 | 0.33768421 |
| 0.05647 | 2.90054 | 1 | 2.90054 | 8.07927 | 0.0176039 | 0.33776205 |
| 0.07562 | 1.98474 | 1 | 1.98474 | 8.072032 | 0.01766772 | 0.33814309 |
| 0.02798 | 1.56525 | 1 | 1.56525 | 8.069926 | 0.01768633 | 0.33814309 |
| 0.08979 | 2.34275 | 1 | 2.34275 | 8.066182 | 0.01771947 | 0.33832975 |
| 0.11835 | 10.24419 | 1 | 10.24419 | 8.015248 | 0.01817653 | 0.34568849 |
| 0.12235 | 999 | 1 | 999 | 7.977784 | 0.01852022 | 0.35084187 |
| 0.03192 | 999 | 1 | 999 | 7.980112 | 0.01849868 | 0.35084187 |
| 0.10009 | 13.30371 | 1 | 13.30371 | 7.962484 | 0.01866245 | 0.35307397 |
| 0.06705 | 23.2 | 1 | 23.2 | 7.949066 | 0.01878807 | 0.35406222 |

| | | | | | | |
|---|---|---|---|---|---|---|
| 0.10903 | 11.31978 | 1 | 11.31978 | 7.953106 | 0.01875016 | 0.35406222 |
| 0.09228 | 46.46394 | 1 | 46.46394 | 7.93517 | 0.01891907 | 0.35514353 |
| 0.15622 | 7.18315 | 1 | 7.18315 | 7.92547 | 0.01901105 | 0.35548695 |
| 0.06656 | 4.521 | 1 | 4.521 | 7.91544 | 0.01910663 | 0.35589475 |
| 4E-04 | 149.30087 | 1 | 149.30087 | 7.90024 | 0.01925239 | 0.35768917 |
| 0.00784 | 999 | 1 | 999 | 7.882196 | 0.01942687 | 0.35954619 |
| 0.09869 | 701.85358 | 1 | 701.85358 | 7.856868 | 0.01967446 | 0.36319954 |
| 0.09328 | 999 | 1 | 999 | 7.846276 | 0.01977893 | 0.36466302 |
| 0.04696 | 15.40228 | 1 | 15.40228 | 7.803522 | 0.0202063 | 0.37206836 |
| 0.07111 | 1.37232 | 1 | 1.37232 | 7.79581 | 0.02028436 | 0.37303123 |
| 0.02245 | 23.2636 | 1 | 23.2636 | 7.790784 | 0.0203354 | 0.37349526 |
| 0.06444 | 336.5355 | 1 | 336.5355 | 7.7048 | 0.02122873 | 0.38695637 |
| 0.15588 | 4.76126 | 1 | 4.76126 | 7.691426 | 0.02137116 | 0.38906261 |
| 0.02279 | 59.77303 | 1 | 59.77303 | 7.671798 | 0.02158193 | 0.39093271 |
| 0.04116 | 999 | 1 | 999 | 7.65534 | 0.02176026 | 0.39367027 |
| 0.07611 | 64.24503 | 1 | 64.24503 | 7.652112 | 0.02179541 | 0.3938139 |
| 0.13473 | 389.15448 | 1 | 389.15448 | 7.637692 | 0.02195312 | 0.3956756 |
| 0.04493 | 48.87482 | 1 | 48.87482 | 7.634338 | 0.02198997 | 0.39584675 |
| 0.13276 | 8.91218 | 1 | 8.91218 | 7.627518 | 0.02206508 | 0.39670547 |
| 0.09792 | 87.93434 | 1 | 87.93434 | 7.61241 | 0.02223239 | 0.39829522 |
| 0.08678 | 999 | 1 | 999 | 7.606318 | 0.02230021 | 0.39845802 |
| 0.0535 | 186.83632 | 1 | 186.83632 | 7.577324 | 0.02262586 | 0.40328079 |
| 0.02875 | 141.10117 | 1 | 141.10117 | 7.570966 | 0.0226979 | 0.40396906 |
| 0.1126 | 80.59487 | 1 | 80.59487 | 7.556368 | 0.02286418 | 0.40503452 |
| 0.0761 | 1.14201 | 1 | 1.14201 | 7.536484 | 0.02309262 | 0.40758482 |
| 0.11217 | 1.29748 | 1 | 1.29748 | 7.525332 | 0.02322175 | 0.40886663 |
| 0.09245 | 134.88589 | 1 | 134.88589 | 7.514842 | 0.02334387 | 0.41033407 |
| 0.05541 | 15.60633 | 1 | 15.60633 | 7.50945 | 0.02340689 | 0.41033407 |
| 0.04856 | 999 | 1 | 999 | 7.505538 | 0.02345272 | 0.4104367 |
| 0.1042 | 52.96634 | 1 | 52.96634 | 7.490538 | 0.02362927 | 0.41302712 |
| 0.08406 | 1.44464 | 1 | 1.44464 | 7.47579 | 0.02380416 | 0.41525247 |
| 0.16728 | 149.12216 | 1 | 149.12216 | 7.471958 | 0.02384981 | 0.41533647 |
| 0.11222 | 2.58027 | 1 | 2.58027 | 7.469748 | 0.02387618 | 0.41533647 |
| 0.07256 | 85.38615 | 1 | 85.38615 | 7.45875 | 0.02400784 | 0.41712534 |
| 0.08651 | 563.22898 | 1 | 563.22898 | 7.454576 | 0.02405799 | 0.41731765 |
| 0.12475 | 1.2326 | 1 | 1.2326 | 7.453032 | 0.02407657 | 0.41731765 |
| 0.10758 | 369.66427 | 1 | 369.66427 | 7.403788 | 0.02467674 | 0.42632898 |
| 0.06475 | 163.17836 | 1 | 163.17836 | 7.403132 | 0.02468484 | 0.42632898 |
| 0.0516 | 999 | 1 | 999 | 7.385378 | 0.02490494 | 0.42772371 |
| 0.03681 | 90.73775 | 1 | 90.73775 | 7.390094 | 0.02484629 | 0.42772371 |
| 0.09675 | 999 | 1 | 999 | 7.361516 | 0.02520386 | 0.42923779 |
| 0.06272 | 17.15179 | 1 | 17.15179 | 7.362778 | 0.02518796 | 0.42923779 |
| 0.14451 | 12.26638 | 1 | 12.26638 | 7.361096 | 0.02520916 | 0.42923779 |
| 0.03434 | 3.1192 | 1 | 3.1192 | 7.363372 | 0.02518048 | 0.42923779 |
| 0.09113 | 2.55246 | 1 | 2.55246 | 7.368728 | 0.02511314 | 0.42923779 |

| | | | | | | |
|---|---|---|---|---|---|---|
| 0.01414 | 999 | 1 | 999 | 7.351944 | 0.02532478 | 0.43069978 |
| 0.11593 | 16.96652 | 1 | 16.96652 | 7.343726 | 0.02542905 | 0.43196557 |
| 0.08009 | 14.79778 | 1 | 14.79778 | 7.34104 | 0.02546323 | 0.432039 |
| 0.03736 | 2.02899 | 1 | 2.02899 | 7.332508 | 0.02557208 | 0.43337795 |
| 0.06459 | 1.9627 | 1 | 1.9627 | 7.32043 | 0.02572698 | 0.43549309 |
| 0.0294 | 10.6854 | 1 | 10.6854 | 7.312234 | 0.02583263 | 0.43626091 |
| 0.09086 | 999 | 1 | 999 | 7.300576 | 0.02598364 | 0.43788463 |
| 0.02322 | 999 | 1 | 999 | 7.300142 | 0.02598928 | 0.43788463 |
| 0.10397 | 412.4779 | 1 | 412.4779 | 7.263628 | 0.02646813 | 0.44132858 |
| 0.08827 | 53.06507 | 1 | 53.06507 | 7.265752 | 0.02644003 | 0.44132858 |
| 0.14629 | 458.2094 | 1 | 458.2094 | 7.255644 | 0.026574 | 0.442584 |
| 0.08704 | 18.31709 | 1 | 18.31709 | 7.250954 | 0.02663639 | 0.4426839 |
| 0.06878 | 3.81084 | 1 | 3.81084 | 7.233774 | 0.02686618 | 0.44540004 |
| 0.03653 | 1.78126 | 1 | 1.78126 | 7.227004 | 0.02695728 | 0.44625722 |
| 0.07108 | 1.31561 | 1 | 1.31561 | 7.225352 | 0.02697955 | 0.44625722 |
| 0.04928 | 999 | 1 | 999 | 7.217766 | 0.02708208 | 0.44685172 |
| 0.05944 | 2.49612 | 1 | 2.49612 | 7.215844 | 0.02710812 | 0.44685172 |
| 0.06187 | 51.14567 | 1 | 51.14567 | 7.205856 | 0.02724384 | 0.44857797 |
| 0.0439 | 31.82055 | 1 | 31.82055 | 7.1976 | 0.02735653 | 0.44992167 |
| 0.04744 | 4.10056 | 1 | 4.10056 | 7.168356 | 0.02775948 | 0.45396938 |
| 0.00876 | 89.91043 | 1 | 89.91043 | 7.15256 | 0.02797959 | 0.4561956 |
| 0.06629 | 11.9067 | 1 | 11.9067 | 7.155596 | 0.02793715 | 0.4561956 |
| 0.08598 | 2.60585 | 1 | 2.60585 | 7.145986 | 0.02807171 | 0.45700995 |
| 0.02851 | 7.21356 | 1 | 7.21356 | 7.139892 | 0.02815737 | 0.4573962 |
| 0.08012 | 1.18008 | 1 | 1.18008 | 7.139802 | 0.02815864 | 0.4573962 |
| 0.09297 | 999 | 1 | 999 | 7.133988 | 0.02824062 | 0.45821352 |
| 0.0171 | 8.18352 | 1 | 8.18352 | 7.124834 | 0.02837017 | 0.45907458 |
| 0.1033 | 30.62523 | 1 | 30.62523 | 7.10014 | 0.02872263 | 0.46189179 |
| 0.10768 | 14.3419 | 1 | 14.3419 | 7.103514 | 0.02867421 | 0.46189179 |
| 0.184 | 69.75414 | 1 | 69.75414 | 7.084998 | 0.02894091 | 0.46437011 |
| 0.06755 | 29.62162 | 1 | 29.62162 | 7.077142 | 0.02905482 | 0.46516633 |
| 0.13831 | 140.69806 | 1 | 140.69806 | 7.052884 | 0.02940937 | 0.46946557 |
| 0.06245 | 2.26194 | 1 | 2.26194 | 7.052116 | 0.02942066 | 0.46946557 |
| 0.0501 | 5.14261 | 1 | 5.14261 | 7.014228 | 0.02998332 | 0.47529969 |
| 0.11637 | 74.43009 | 1 | 74.43009 | 6.995114 | 0.03027125 | 0.47777071 |
| 0.05586 | 1.00299 | 1 | 1.00299 | 6.971378 | 0.03063265 | 0.48189812 |
| 0.12147 | 3.84736 | 1 | 3.84736 | 6.94725 | 0.03100444 | 0.48616164 |
| 0.06321 | 38.11923 | 1 | 38.11923 | 6.93373 | 0.03121474 | 0.48892951 |
| 0.13081 | 16.98012 | 1 | 16.98012 | 6.923512 | 0.03137462 | 0.49037243 |
| 0.01363 | 61.63101 | 1 | 61.63101 | 6.91184 | 0.03155826 | 0.49271053 |
| 0.06515 | 999 | 1 | 999 | 6.890082 | 0.03190345 | 0.49528872 |
| 0.16469 | 26.09144 | 1 | 26.09144 | 6.884216 | 0.03199716 | 0.49528872 |
| 0.14299 | 11.35697 | 1 | 11.35697 | 6.884538 | 0.03199201 | 0.49528872 |
| 0.05563 | 7.24913 | 1 | 7.24913 | 6.895668 | 0.03181447 | 0.49528872 |
| 0.02533 | 2.6013 | 1 | 2.6013 | 6.892602 | 0.03186328 | 0.49528872 |

| | | | | | | |
|---|---|---|---|---|---|---|
| 0.11536 | 1.12413 | 1 | 1.12413 | 6.89271 | 0.03186156 | 0.49528872 |
| 0.05611 | 2.41221 | 1 | 2.41221 | 6.877894 | 0.03209847 | 0.49632598 |
| 0.12059 | 591.15706 | 1 | 591.15706 | 6.874866 | 0.0321471 | 0.49654749 |
| 0.0638 | 180.36336 | 1 | 180.36336 | 6.86243 | 0.03234761 | 0.49911197 |
| 0.0552 | 999 | 1 | 999 | 6.839792 | 0.03271584 | 0.50265001 |
| 0.13829 | 59.0645 | 1 | 59.0645 | 6.833548 | 0.03281814 | 0.50368704 |
| 0.03153 | 100.94859 | 1 | 100.94859 | 6.826886 | 0.03292764 | 0.5038486 |
| 0.0833 | 15.4431 | 1 | 15.4431 | 6.830108 | 0.03287463 | 0.5038486 |
| 0.13451 | 2.25555 | 1 | 2.25555 | 6.826554 | 0.0329331 | 0.5038486 |
| 0.02064 | 31.47869 | 1 | 31.47869 | 6.795126 | 0.0334547 | 0.51023612 |
| 0.14597 | 53.30882 | 1 | 53.30882 | 6.786582 | 0.03359792 | 0.51185553 |
| 0.24819 | 9.91818 | 1 | 9.91818 | 6.778648 | 0.03373147 | 0.51281049 |
| 0.08033 | 999 | 1 | 999 | 6.772004 | 0.03384371 | 0.51385171 |
| 0.03256 | 1.61678 | 1 | 1.61678 | 6.770394 | 0.03387097 | 0.51385171 |
| 0.16753 | 5.78322 | 1 | 5.78322 | 6.767274 | 0.03392385 | 0.51411504 |
| 0.09772 | 147.15356 | 1 | 147.15356 | 6.75189 | 0.0341858 | 0.51592393 |
| 0.03448 | 1.1009 | 1 | 1.1009 | 6.74842 | 0.03424516 | 0.51628148 |
| 0.04855 | 47.47679 | 1 | 47.47679 | 6.737936 | 0.03442515 | 0.51791594 |
| 0.079 | 1.1811 | 1 | 1.1811 | 6.724138 | 0.03466347 | 0.51988015 |
| 0.07954 | 3.42289 | 1 | 3.42289 | 6.70403 | 0.03501373 | 0.52404728 |
| 0.07935 | 3.42596 | 1 | 3.42596 | 6.691796 | 0.03522857 | 0.52671801 |
| 0.2257 | 999 | 1 | 999 | 6.684446 | 0.03535827 | 0.52781452 |
| 0.07765 | 1.67996 | 1 | 1.67996 | 6.677022 | 0.03548976 | 0.52789654 |
| 0.08014 | 999 | 1 | 999 | 6.656958 | 0.03584759 | 0.53212527 |
| 0.06209 | 999 | 1 | 999 | 6.658168 | 0.03582591 | 0.53212527 |
| 0.10214 | 1.38444 | 1 | 1.38444 | 6.646932 | 0.03602774 | 0.53370473 |
| 0.10722 | 269.67404 | 1 | 269.67404 | 6.639692 | 0.0361584 | 0.53454598 |
| 0.03083 | 4.54856 | 1 | 4.54856 | 6.63177 | 0.03630191 | 0.5361199 |
| 0.02258 | 318.99392 | 1 | 318.99392 | 6.60649 | 0.03676368 | 0.53963881 |
| 0.20708 | 27.49937 | 1 | 27.49937 | 6.612258 | 0.0366578 | 0.53963881 |
| 0.02996 | 2.20078 | 1 | 2.20078 | 6.606478 | 0.0367639 | 0.53963881 |
| 0.03769 | 999 | 1 | 999 | 6.568624 | 0.03746635 | 0.54772781 |
| 0.0446 | 44.06271 | 1 | 44.06271 | 6.560854 | 0.03761219 | 0.54875128 |
| 0.08397 | 36.93139 | 1 | 36.93139 | 6.554798 | 0.03772626 | 0.54947441 |
| 0.09041 | 1.22159 | 1 | 1.22159 | 6.554192 | 0.03773769 | 0.54947441 |
| 0.07759 | 43.60126 | 1 | 43.60126 | 6.549732 | 0.03782194 | 0.55014763 |
| 0.0979 | 58.90717 | 1 | 58.90717 | 6.54492 | 0.03791305 | 0.55086107 |
| 0.04477 | 38.65923 | 1 | 38.65923 | 6.543124 | 0.03794711 | 0.55086107 |
| 0.03992 | 25.69615 | 1 | 25.69615 | 6.5394 | 0.03801783 | 0.55129531 |
| 0.06558 | 13.64393 | 1 | 13.64393 | 6.53754 | 0.0380532 | 0.55129531 |
| 0.13702 | 28.38699 | 1 | 28.38699 | 6.512494 | 0.03853274 | 0.5560163 |
| 0.07565 | 85.77192 | 1 | 85.77192 | 6.494504 | 0.03888091 | 0.55825729 |
| 0.05456 | 76.15491 | 1 | 76.15491 | 6.494938 | 0.03887247 | 0.55825729 |
| 0.07951 | 1.04639 | 1 | 1.04639 | 6.495906 | 0.03885366 | 0.55825729 |
| 0.02028 | 54.40068 | 1 | 54.40068 | 6.48712 | 0.03902472 | 0.55865951 |

| | | | | | | |
|---|---|---|---|---|---|---|
| 0.15139 | 33.39183 | 1 | 33.39183 | 6.46772 | 0.0394051 | 0.56128757 |
| 0.08136 | 3.50633 | 1 | 3.50633 | 6.46197 | 0.03951855 | 0.56128757 |
| 0.08377 | 1.38892 | 1 | 1.38892 | 6.462196 | 0.03951409 | 0.56128757 |
| 0.11785 | 1.18417 | 1 | 1.18417 | 6.46505 | 0.03945774 | 0.56128757 |
| 0.10108 | 999 | 1 | 999 | 6.444682 | 0.03986163 | 0.56387552 |
| 0.04946 | 999 | 1 | 999 | 6.447642 | 0.03980268 | 0.56387552 |
| 0.01427 | 656.70242 | 1 | 656.70242 | 6.44786 | 0.03979834 | 0.56387552 |
| 0.08087 | 1.54811 | 1 | 1.54811 | 6.441736 | 0.03992039 | 0.56387552 |
| 0.14626 | 1.51581 | 1 | 1.51581 | 6.441028 | 0.03993453 | 0.56387552 |
| 0.02867 | 1.01923 | 1 | 1.01923 | 6.438638 | 0.03998228 | 0.56399951 |
| 0.05837 | 1.81182 | 1 | 1.81182 | 6.434414 | 0.04006681 | 0.5646416 |
| 0.08026 | 7.9126 | 1 | 7.9126 | 6.395944 | 0.04084495 | 0.57281881 |
| 0.12398 | 999 | 1 | 999 | 6.38408 | 0.04108797 | 0.57455665 |
| 0.11365 | 3.4648 | 1 | 3.4648 | 6.36313 | 0.04152062 | 0.57948698 |
| 0.01279 | 147.28122 | 1 | 147.28122 | 6.355116 | 0.04168733 | 0.58125313 |
| 0.13054 | 421.99908 | 1 | 421.99908 | 6.350672 | 0.04178006 | 0.58142583 |
| 0.06965 | 160.77136 | 1 | 160.77136 | 6.352164 | 0.04174891 | 0.58142583 |
| 0.01982 | 5.86371 | 1 | 5.86371 | 6.336836 | 0.0420701 | 0.5837781 |
| 0.11808 | 24.18113 | 1 | 24.18113 | 6.328374 | 0.04224848 | 0.58457188 |
| 0.0861 | 999 | 1 | 999 | 6.322052 | 0.04238223 | 0.58586254 |
| 0.09934 | 998.99999 | 1 | 998.99999 | 6.31143 | 0.04260793 | 0.58785939 |
| 0.02307 | 437.876 | 1 | 437.876 | 6.301866 | 0.04281216 | 0.59011472 |
| 0.04325 | 1.36043 | 1 | 1.36043 | 6.293656 | 0.04298827 | 0.59197833 |
| 0.01274 | 999 | 1 | 999 | 6.280886 | 0.04326363 | 0.59450696 |
| 0.01091 | 998.99999 | 1 | 998.99999 | 6.273746 | 0.04341836 | 0.59450696 |
| 0.07319 | 32.71833 | 1 | 32.71833 | 6.28285 | 0.04322116 | 0.59450696 |
| 0.16354 | 1.32964 | 1 | 1.32964 | 6.275572 | 0.04337873 | 0.59450696 |
| 0.07317 | 1.46863 | 1 | 1.46863 | 6.26374 | 0.04363612 | 0.59636033 |
| 0.06202 | 94.86965 | 1 | 94.86965 | 6.257422 | 0.04377419 | 0.59768282 |
| 0.03564 | 999 | 1 | 999 | 6.254102 | 0.04384691 | 0.59811154 |
| 0.11634 | 149.75915 | 1 | 149.75915 | 6.248454 | 0.04397091 | 0.59855973 |
| 0.01325 | 70.56504 | 1 | 70.56504 | 6.245078 | 0.0440452 | 0.59855973 |
| 0.07605 | 1.80784 | 1 | 1.80784 | 6.246782 | 0.04400769 | 0.59855973 |
| 0.01959 | 999 | 1 | 999 | 6.239604 | 0.04416591 | 0.59907521 |
| 0.01826 | 999 | 1 | 999 | 6.240434 | 0.04414759 | 0.59907521 |
| 0.06268 | 2.86433 | 1 | 2.86433 | 6.236228 | 0.04424053 | 0.59952542 |
| 0.0492 | 999 | 1 | 999 | 6.222738 | 0.04453994 | 0.6024547 |
| 0.0735 | 999 | 1 | 999 | 6.207402 | 0.04488278 | 0.60652523 |
| 0.05421 | 32.17338 | 1 | 32.17338 | 6.188774 | 0.04530277 | 0.61049074 |
| 0.04109 | 999 | 1 | 999 | 6.18587 | 0.0453686 | 0.61080909 |
| 0.00949 | 999 | 1 | 999 | 6.183482 | 0.0454228 | 0.61097049 |
| 0.01814 | 999 | 1 | 999 | 6.170364 | 0.04572171 | 0.611187 |
| 0.15965 | 999 | 1 | 999 | 6.162458 | 0.04590281 | 0.611187 |
| 0.07051 | 23.86246 | 1 | 23.86246 | 6.179888 | 0.0455045 | 0.611187 |
| 0.0517 | 11.18724 | 1 | 11.18724 | 6.161828 | 0.04591727 | 0.611187 |

| | | | | | | |
|---|---|---|---|---|---|---|
| 0.14354 | 7.74269 | 1 | 7.74269 | 6.160722 | 0.04594267 | 0.611187 |
| 0.06631 | 1.09662 | 1 | 1.09662 | 6.16917 | 0.04574902 | 0.611187 |
| 0.03365 | 12.54746 | 1 | 12.54746 | 6.138638 | 0.04645278 | 0.61587514 |
| 0.11552 | 1.1036 | 1 | 1.1036 | 6.13797 | 0.0464683 | 0.61587514 |
| 0.02476 | 8.34378 | 1 | 8.34378 | 6.116668 | 0.04696588 | 0.62133191 |
| 0.09432 | 23.0681 | 1 | 23.0681 | 6.110532 | 0.04711019 | 0.62267192 |
| 0.02364 | 999 | 1 | 999 | 6.104494 | 0.04725263 | 0.62295593 |
| 0.14858 | 24.94146 | 1 | 24.94146 | 6.106816 | 0.0471978 | 0.62295593 |
| 0.09177 | 420.81016 | 1 | 420.81016 | 6.10001 | 0.04735869 | 0.62367815 |
| 0.07539 | 999 | 1 | 999 | 6.088942 | 0.0476215 | 0.62497546 |
| 0.09255 | 23.76139 | 1 | 23.76139 | 6.088588 | 0.04762993 | 0.62497546 |
| 0.09955 | 4.02956 | 1 | 4.02956 | 6.091264 | 0.04756624 | 0.62497546 |
| 0.09688 | 128.30319 | 1 | 128.30319 | 6.080162 | 0.04783102 | 0.6256033 |
| 0.01794 | 999 | 1 | 999 | 6.073738 | 0.0479849 | 0.62622668 |
| 0.01957 | 999 | 1 | 999 | 6.062802 | 0.048248 | 0.62852677 |
| 0.13623 | 10.44395 | 1 | 10.44395 | 6.008726 | 0.04957032 | 0.64228402 |
| 0.05966 | 7.89078 | 1 | 7.89078 | 6.004002 | 0.04968754 | 0.64265221 |
| 0.07754 | 9.70807 | 1 | 9.70807 | 5.99341 | 0.04995139 | 0.64491207 |
| 0.0241 | 1.50967 | 1 | 1.50967 | 5.993488 | 0.04994944 | 0.64491207 |

d end-joining (MMEJ) DNA repair by promoting recruitment of polymerase theta (PO
ɔunit
lic acid transporter 1)

ce, 249 genes)

olinesterase

Parkinson disease-8
/ of cullin-RING E3 ubiquitin ligase (CRL) complexes

mentosa
sine triphosphate (GTP)-metabolizing protein that participates in the cellular antiviral [illegible]

olecular oxygen to various reactive oxygen species (ROS)
g proteins

and reactive nitrogen species and in mitochondrial homeostasis in the placenta

dase
ctrin-actin cytoskeleton

uanosine mRNA capping and cap2 mRNA methylation
Subunit 1

luced by P53 and involved in P53-mediated cell death

ge
clease activity
rolase activity
of cytoskeleton
pecific DNA binding; reactive oxidative species regulation
ycle progression
n of BRCA1; BARD1-BRCA1 heterodimer involved in DNA repair

### n variants (274 genes)

in involved in telomere maintenance
f the complement system

ımed cell death
e receptor for collagen and laminin

es their cytoxicity

factor-1 receptor (IGF1R) and is thought to be involved in the regulation of IGF1R me
ł Ca(2+) handling in myocytes
ter cells 5 (double-strand-break rejoining)
ıctions
vth factor family

sine kinase family
) in plasma

essential for hemostasis
y

unit

ene product to aging in a mammalian model organism

ct to a pathway or mechanism linked to aging
ene product to aging in a mammalian model organism
ene product to aging in a mammalian model organism

luct to the regulation or control of genes previously linked to aging

ct to a pathway or mechanism linked to aging

Gs

| Provean Tole | Provean Dele | SIFT Tolerate | SIFT Deleterious |
|---|---|---|---|
| 227 | 503 | 283 | 447 |
| 113 | 52 | 141 | 24 |
| 82 | 24 | 93 | 13 |
| 22 | 4 | 24 | 2 |
| 455 | 62 | 489 | 28 |
| 35 | 20 | 37 | 18 |
| 19 | 5 | 21 | 3 |
| 349 | 715 | 330 | 734 |
| 0 | 0 | 0 | 0 |
| 17 | 8 | 19 | 6 |
| 214 | 62 | 240 | 36 |
| 409 | 104 | 503 | 10 |
| 17 | 8 | 15 | 10 |
| 26 | 5 | 27 | 4 |
| 238 | 149 | 91 | 296 |
| 141 | 312 | 405 | 48 |
| 60 | 14 | 60 | 14 |
| 82 | 106 | 91 | 97 |
| 151 | 114 | 224 | 41 |
| 170 | 34 | 189 | 15 |
| 24 | 5 | 26 | 3 |
| 238 | 27 | 237 | 28 |
| 249 | 158 | 329 | 78 |
| 145 | 34 | 170 | 9 |
| 34 | 10 | 38 | 6 |
| 442 | 76 | 484 | 34 |
| 54 | 30 | 72 | 12 |
| 75 | 9 | 78 | 6 |
| 9 | 13 | 11 | 11 |
| 359 | 41 | 322 | 78 |
| 66 | 23 | 78 | 11 |
| 364 | 41 | 327 | 78 |

| | | | |
|---|---|---|---|
| 67 | 15 | 70 | 12 |
| 107 | 96 | 178 | 25 |
| 107 | 16 | 111 | 12 |
| 74 | 5 | 77 | 2 |
| 82 | 21 | 88 | 15 |
| 8 | 6 | 7 | 7 |
| 65 | 6 | 63 | 8 |
| 43 | 58 | 64 | 37 |
| 5 | 3 | 3 | 5 |
| 216 | 80 | 271 | 25 |
| 67 | 6 | 68 | 5 |
| 56 | 4 | 57 | 3 |
| 32 | 53 | 48 | 37 |
| 40 | 3 | 40 | 3 |
| 125 | 20 | 100 | 45 |
| 17 | 6 | 16 | 7 |
| 281 | 52 | 292 | 41 |
| 47 | 7 | 41 | 13 |
| 68 | 19 | 74 | 13 |
| 159 | 52 | 193 | 18 |
| 222 | 52 | 177 | 97 |
| 86 | 33 | 98 | 21 |
| 10 | 11 | 9 | 12 |
| 18 | 6 | 22 | 2 |
| 143 | 34 | 159 | 18 |
| 44 | 15 | 49 | 10 |
| 129 | 40 | 9 | 160 |
| 134 | 36 | 146 | 24 |
| 91 | 10 | 92 | 9 |
| 6 | 1 | 7 | 0 |
| 504 | 242 | 466 | 280 |
| 133 | 20 | 148 | 5 |
| 28 | 4 | 32 | 0 |
| 178 | 139 | 296 | 21 |
| 7 | 0 | 6 | 1 |
| 109 | 137 | 228 | 18 |
| 53 | 2 | 52 | 3 |
| 77 | 69 | 110 | 36 |
| 6 | 4 | 7 | 3 |
| 140 | 78 | 88 | 130 |
| 123 | 18 | 123 | 18 |
| 16 | 5 | 18 | 3 |
| 2 | 0 | 1 | 1 |
| 78 | 27 | 102 | 3 |
| 17 | 12 | 23 | 6 |

| | | | |
|---|---|---|---|
| 39 | 13 | 44 | 8 |
| 266 | 54 | 285 | 35 |
| 140 | 44 | 168 | 16 |
| 206 | 21 | 202 | 25 |
| 198 | 119 | 263 | 54 |
| 141 | 76 | 87 | 130 |
| 298 | 28 | 294 | 32 |
| 72 | 68 | 133 | 7 |
| 80 | 81 | 138 | 23 |
| 126 | 144 | 185 | 85 |
| 133 | 249 | 344 | 38 |
| 72 | 5 | 71 | 6 |
| 35 | 28 | 58 | 5 |
| 752 | 370 | 1021 | 101 |
| 104 | 23 | 124 | 3 |
| 59 | 0 | 57 | 2 |
| 51 | 2 | 47 | 6 |
| 42 | 18 | 52 | 8 |
| 123 | 11 | 127 | 7 |
| 154 | 29 | 160 | 23 |
| 13 | 2 | 15 | 0 |
| 188 | 261 | 219 | 230 |
| 42 | 85 | 99 | 28 |
| 352 | 117 | 426 | 43 |
| 196 | 27 | 194 | 29 |
| 49 | 10 | 53 | 6 |
| 215 | 28 | 218 | 25 |
| 568 | 91 | 623 | 36 |
| 76 | 12 | 76 | 12 |
| 359 | 356 | 628 | 87 |
| 137 | 196 | 315 | 18 |
| 61 | 9 | 67 | 3 |
| 228 | 475 | 597 | 106 |
| 102 | 55 | 142 | 15 |
| 187 | 23 | 204 | 6 |
| 43 | 4 | 44 | 3 |
| 50 | 9 | 52 | 7 |
| 40 | 34 | 59 | 15 |
| 128 | 177 | 260 | 45 |
| 133 | 156 | 236 | 53 |
| 130 | 17 | 141 | 6 |
| 13 | 3 | 13 | 3 |
| 49 | 7 | 53 | 3 |
| 79 | 9 | 80 | 8 |
| 59 | 10 | 55 | 14 |

| | | | |
|---|---|---|---|
| 293 | 211 | 430 | 74 |
| 67 | 21 | 78 | 10 |
| 138 | 28 | 139 | 27 |
| 27 | 24 | 24 | 27 |
| 0 | 0 | 0 | 0 |
| 222 | 94 | 298 | 18 |
| 55 | 14 | 61 | 8 |
| 50 | 16 | 51 | 15 |
| 0 | 0 | 0 | 0 |
| 88 | 8 | 94 | 2 |
| 484 | 53 | 494 | 43 |
| 64 | 9 | 66 | 7 |
| 35 | 39 | 67 | 7 |
| 24 | 22 | 41 | 5 |
| 169 | 27 | 184 | 12 |
| 130 | 16 | 136 | 10 |
| 178 | 61 | 224 | 15 |
| 108 | 4 | 101 | 11 |
| 9 | 1 | 10 | 0 |
| 94 | 10 | 92 | 12 |
| 29 | 7 | 29 | 7 |
| 100 | 46 | 136 | 10 |
| 136 | 19 | 149 | 6 |
| 91 | 8 | 93 | 6 |
| 100 | 5 | 103 | 2 |
| 342 | 34 | 353 | 23 |
| 227 | 42 | 251 | 18 |
| 213 | 50 | 225 | 38 |
| 124 | 48 | 159 | 13 |
| 1013 | 184 | 1098 | 99 |
| 180 | 218 | 349 | 49 |
| 69 | 10 | 75 | 4 |
| 183 | 17 | 184 | 16 |
| 132 | 28 | 151 | 9 |
| 269 | 202 | 205 | 266 |
| 36 | 12 | 43 | 5 |
| 123 | 5 | 125 | 3 |
| 24 | 0 | 23 | 1 |
| 378 | 44 | 383 | 39 |
| 26 | 18 | 42 | 2 |
| 174 | 35 | 176 | 33 |
| 92 | 102 | 122 | 72 |
| 133 | 42 | 159 | 16 |
| 33 | 3 | 34 | 2 |
| 54 | 4 | 56 | 2 |

| | | | |
|---|---|---|---|
| 325 | 163 | 459 | 29 |
| 107 | 13 | 114 | 6 |
| 744 | 651 | 183 | 1212 |
| 30 | 26 | 46 | 10 |
| 105 | 26 | 109 | 22 |
| 28 | 50 | 35 | 43 |
| 18 | 2 | 19 | 1 |
| 72 | 54 | 116 | 10 |
| 14 | 1 | 14 | 1 |
| 96 | 78 | 57 | 117 |
| 397 | 179 | 543 | 33 |
| 271 | 28 | 281 | 18 |
| 125 | 19 | 130 | 14 |
| 172 | 73 | 231 | 14 |
| 21 | 7 | 22 | 6 |
| 84 | 16 | 86 | 14 |
| 51 | 4 | 54 | 1 |
| 0 | 0 | 0 | 0 |
| 61 | 7 | 63 | 5 |
| 59 | 82 | 128 | 13 |
| 143 | 7 | 147 | 3 |
| 79 | 41 | 88 | 32 |
| 39 | 121 | 57 | 103 |
| 76 | 1 | 75 | 2 |
| 172 | 22 | 178 | 16 |
| 108 | 18 | 117 | 9 |
| 69 | 89 | 114 | 44 |
| 97 | 16 | 107 | 6 |
| 276 | 39 | 269 | 46 |
| 50 | 11 | 56 | 5 |
| 9 | 2 | 8 | 3 |
| 198 | 437 | 236 | 399 |
| 208 | 73 | 256 | 25 |
| 38 | 6 | 37 | 7 |
| 73 | 8 | 75 | 6 |
| 264 | 229 | 458 | 35 |
| 214 | 22 | 216 | 20 |
| 34 | 0 | 33 | 1 |
| 0 | 0 | 0 | 0 |
| 108 | 26 | 123 | 11 |
| 65 | 11 | 67 | 9 |
| 288 | 47 | 295 | 40 |
| 153 | 23 | 157 | 19 |
| 136 | 39 | 162 | 13 |
| 21 | 3 | 20 | 4 |

| | | | |
|---|---|---|---|
| 92 | 40 | 111 | 21 |
| 87 | 23 | 98 | 12 |
| 736 | 129 | 781 | 84 |
| 56 | 92 | 137 | 11 |
| 12 | 4 | 13 | 3 |
| 76 | 9 | 78 | 7 |
| 213 | 63 | 18 | 258 |
| 147 | 3 | 139 | 11 |
| 126 | 21 | 136 | 11 |
| 541 | 94 | 591 | 44 |
| 225 | 18 | 235 | 8 |
| 36 | 13 | 48 | 1 |
| 101 | 12 | 108 | 5 |
| 149 | 95 | 167 | 77 |
| 81 | 19 | 93 | 7 |
| 101 | 7 | 101 | 7 |
| 36 | 6 | 37 | 5 |
| 204 | 28 | 55 | 177 |
| 27 | 11 | 31 | 7 |
| 366 | 44 | 378 | 32 |
| 19 | 1 | 18 | 2 |
| 0 | 0 | 0 | 0 |
| 85 | 23 | 91 | 17 |
| 314 | 135 | 414 | 35 |
| 136 | 32 | 147 | 21 |
| 192 | 27 | 215 | 4 |
| 74 | 14 | 82 | 6 |
| 89 | 7 | 95 | 1 |
| 112 | 13 | 114 | 11 |
| 35 | 15 | 40 | 10 |
| 11 | 0 | 10 | 1 |
| 34 | 106 | 58 | 82 |
| 167 | 210 | 355 | 22 |
| 99 | 10 | 105 | 4 |
| 56 | 7 | 60 | 3 |
| 64 | 97 | 146 | 15 |
| 115 | 32 | 134 | 13 |
| 109 | 19 | 123 | 5 |
| 162 | 17 | 172 | 7 |
| 43 | 12 | 53 | 2 |
| 196 | 26 | 208 | 14 |
| 271 | 84 | 338 | 17 |
| 110 | 6 | 113 | 3 |
| 80 | 16 | 81 | 15 |
| 130 | 113 | 219 | 24 |

| | | | |
|---|---|---|---|
| 129 | 36 | 158 | 7 |
| 116 | 234 | 199 | 151 |
| 108 | 10 | 115 | 3 |
| 261 | 26 | 270 | 17 |
| 46 | 6 | 50 | 2 |
| 21 | 3 | 22 | 2 |
| 53 | 11 | 54 | 10 |
| 116 | 14 | 119 | 11 |
| 313 | 192 | 478 | 27 |
| 388 | 152 | 234 | 306 |
| 166 | 20 | 154 | 32 |
| 216 | 12 | 221 | 7 |
| 76 | 68 | 136 | 8 |
| 96 | 10 | 96 | 10 |
| 23 | 6 | 24 | 5 |
| 30 | 12 | 40 | 2 |
| 111 | 10 | 116 | 5 |
| 126 | 166 | 260 | 32 |
| 76 | 14 | 29 | 61 |
| 115 | 14 | 125 | 4 |
| 152 | 16 | 154 | 14 |
| 127 | 12 | 124 | 15 |
| 70 | 17 | 81 | 6 |
| 107 | 54 | 158 | 3 |
| 273 | 43 | 292 | 24 |
| 43 | 16 | 55 | 4 |
| 18 | 4 | 21 | 1 |
| 237 | 24 | 235 | 26 |
| 34 | 13 | 43 | 4 |
| 89 | 15 | 96 | 8 |
| 37 | 69 | 98 | 8 |
| 73 | 12 | 78 | 7 |
| 169 | 13 | 172 | 10 |
| 19 | 1 | 20 | 0 |
| 44 | 5 | 45 | 4 |
| 29 | 7 | 32 | 4 |
| 152 | 26 | 169 | 9 |
| 159 | 134 | 280 | 13 |
| 330 | 64 | 371 | 23 |
| 252 | 50 | 263 | 39 |
| 175 | 275 | 398 | 52 |
| 106 | 7 | 109 | 4 |
| 128 | 95 | 217 | 6 |
| 25 | 6 | 29 | 2 |
| 117 | 11 | 123 | 5 |

| | | | |
|---|---|---|---|
| 146 | 22 | 160 | 8 |
| 14 | 1 | 15 | 0 |
| 458 | 406 | 804 | 60 |
| 172 | 33 | 196 | 9 |
| 348 | 15 | 338 | 25 |
| 58 | 7 | 58 | 7 |
| 49 | 23 | 59 | 13 |
| 90 | 23 | 106 | 7 |
| 148 | 93 | 200 | 41 |
| 274 | 59 | 318 | 15 |
| 154 | 212 | 190 | 176 |
| 82 | 130 | 190 | 22 |
| 66 | 72 | 83 | 55 |
| 151 | 34 | 173 | 12 |
| 408 | 119 | 505 | 22 |
| 85 | 10 | 95 | 0 |
| 187 | 24 | 193 | 18 |
| 45 | 0 | 42 | 3 |
| 4 | 1 | 5 | 0 |
| 123 | 13 | 119 | 17 |
| 252 | 9 | 250 | 11 |
| 215 | 17 | 213 | 19 |
| 462 | 53 | 461 | 54 |
| 61 | 5 | 62 | 4 |
| 123 | 11 | 130 | 4 |
| 24 | 26 | 43 | 7 |
| 14 | 2 | 15 | 1 |
| 66 | 2 | 67 | 1 |
| 65 | 4 | 63 | 6 |
| 35 | 3 | 34 | 4 |
| 68 | 11 | 64 | 15 |
| 369 | 55 | 411 | 13 |
| 47 | 8 | 50 | 5 |
| 126 | 88 | 15 | 199 |
| 115 | 10 | 117 | 8 |
| 8 | 0 | 7 | 1 |
| 46 | 8 | 49 | 5 |
| 199 | 18 | 206 | 11 |
| 293 | 147 | 415 | 25 |
| 72 | 107 | 168 | 11 |
| 16 | 10 | 22 | 4 |
| 100 | 9 | 103 | 6 |
| 16 | 22 | 35 | 3 |
| 43 | 128 | 66 | 105 |
| 209 | 69 | 265 | 13 |

| | | | |
|---|---|---|---|
| 180 | 24 | 198 | 6 |
| 111 | 21 | 125 | 7 |
| 73 | 11 | 68 | 16 |
| 96 | 10 | 99 | 7 |
| 96 | 4 | 95 | 5 |
| 839 | 1933 | 416 | 2356 |
| 66 | 50 | 75 | 41 |
| 87 | 207 | 95 | 199 |
| 111 | 7 | 114 | 4 |
| 59 | 11 | 61 | 9 |
| 31 | 66 | 32 | 65 |
| 142 | 36 | 168 | 10 |
| 31 | 10 | 36 | 5 |
| 36 | 3 | 37 | 2 |
| 97 | 13 | 99 | 11 |
| 200 | 29 | 212 | 17 |
| 279 | 57 | 311 | 25 |
| 71 | 6 | 69 | 8 |
| 185 | 21 | 195 | 11 |
| 92 | 16 | 98 | 10 |
| 128 | 8 | 133 | 3 |
| 75 | 34 | 99 | 10 |
| 59 | 64 | 103 | 20 |
| 76 | 3 | 73 | 6 |
| 324 | 41 | 317 | 48 |
| 175 | 25 | 182 | 18 |
| 62 | 5 | 62 | 5 |
| 95 | 32 | 104 | 23 |
| 169 | 35 | 199 | 5 |
| 186 | 20 | 196 | 10 |
| 46 | 7 | 51 | 2 |
| 121 | 42 | 118 | 45 |
| 142 | 14 | 149 | 7 |
| 129 | 5 | 127 | 7 |
| 95 | 195 | 191 | 99 |
| 0 | 0 | 0 | 0 |
| 155 | 29 | 175 | 9 |
| 109 | 18 | 112 | 15 |
| 92 | 10 | 96 | 6 |
| 75 | 46 | 51 | 70 |
| 70 | 15 | 78 | 7 |
| 48 | 9 | 55 | 2 |
| 93 | 9 | 95 | 7 |
| 27 | 4 | 28 | 3 |
| 157 | 10 | 160 | 7 |

| | | | |
|---|---|---|---|
| 83 | 5 | 83 | 5 |
| 196 | 43 | 206 | 33 |
| 257 | 48 | 291 | 14 |
| 112 | 8 | 108 | 12 |
| 35 | 32 | 37 | 30 |
| 138 | 41 | 21 | 158 |
| 222 | 44 | 227 | 39 |
| 67 | 16 | 64 | 19 |
| 80 | 18 | 71 | 27 |
| 20 | 9 | 25 | 4 |
| 25 | 0 | 23 | 2 |
| 78 | 10 | 79 | 9 |
| 61 | 54 | 56 | 59 |
| 106 | 6 | 109 | 3 |
| 162 | 11 | 164 | 9 |
| 107 | 106 | 120 | 93 |
| 212 | 49 | 233 | 28 |
| 111 | 16 | 116 | 11 |
| 310 | 792 | 1067 | 35 |
| 348 | 88 | 417 | 19 |
| 217 | 331 | 64 | 484 |
| 29 | 4 | 24 | 9 |
| 62 | 101 | 103 | 60 |
| 152 | 45 | 176 | 21 |
| 44 | 165 | 84 | 125 |
| 114 | 20 | 95 | 39 |
| 242 | 38 | 261 | 19 |
| 99 | 9 | 104 | 4 |
| 64 | 7 | 68 | 3 |
| 230 | 18 | 238 | 10 |
| 90 | 81 | 155 | 16 |
| 206 | 94 | 279 | 21 |
| 33 | 5 | 35 | 3 |
| 54 | 3 | 54 | 3 |
| 99 | 13 | 108 | 4 |
| 96 | 17 | 96 | 17 |
| 43 | 29 | 59 | 13 |
| 220 | 21 | 226 | 15 |
| 133 | 96 | 214 | 15 |
| 53 | 4 | 53 | 4 |
| 18 | 8 | 24 | 2 |
| 106 | 50 | 147 | 9 |
| 139 | 35 | 163 | 11 |
| 150 | 13 | 153 | 10 |
| 111 | 13 | 119 | 5 |

| | | | |
|---|---|---|---|
| 39 | 10 | 40 | 9 |
| 57 | 7 | 59 | 5 |
| 55 | 22 | 62 | 15 |
| 35 | 6 | 37 | 4 |
| 261 | 19 | 267 | 13 |
| 41 | 3 | 39 | 5 |
| 16 | 26 | 39 | 3 |
| 41 | 7 | 45 | 3 |
| 52 | 4 | 52 | 4 |
| 105 | 133 | 180 | 58 |
| 189 | 36 | 205 | 20 |
| 138 | 12 | 135 | 15 |
| 112 | 20 | 94 | 38 |
| 109 | 13 | 111 | 11 |
| 78 | 18 | 90 | 6 |
| 63 | 1 | 62 | 2 |
| 193 | 35 | 208 | 20 |
| 53 | 6 | 56 | 3 |
| 54 | 6 | 54 | 6 |
| 113 | 8 | 112 | 9 |
| 71 | 12 | 77 | 6 |
| 171 | 152 | 296 | 27 |
| 80 | 17 | 83 | 14 |
| 76 | 12 | 74 | 14 |
| 23 | 55 | 65 | 13 |
| 92 | 6 | 93 | 5 |
| 104 | 124 | 213 | 15 |
| 150 | 13 | 155 | 8 |
| 18 | 6 | 20 | 4 |
| 71 | 33 | 95 | 9 |
| 98 | 18 | 112 | 4 |
| 179 | 46 | 199 | 26 |
| 129 | 218 | 135 | 212 |
| 49 | 3 | 52 | 0 |
| 137 | 12 | 140 | 9 |
| 64 | 26 | 65 | 25 |
| 65 | 6 | 66 | 5 |
| 95 | 25 | 115 | 5 |
| 141 | 77 | 203 | 15 |
| 79 | 78 | 133 | 24 |
| 0 | 0 | 0 | 0 |
| 13 | 3 | 16 | 0 |
| 47 | 8 | 45 | 10 |
| 67 | 7 | 66 | 8 |
| 38 | 50 | 85 | 3 |

| | | | |
|---|---|---|---|
| 153 | 10 | 153 | 10 |
| 46 | 4 | 49 | 1 |
| 1280 | 949 | 247 | 1982 |
| 374 | 20 | 383 | 11 |
| 25 | 3 | 28 | 0 |
| 40 | 10 | 43 | 7 |
| 139 | 74 | 169 | 44 |
| 116 | 142 | 156 | 102 |
| 69 | 16 | 69 | 16 |
| 146 | 35 | 174 | 7 |
| 31 | 3 | 33 | 1 |
| 299 | 268 | 490 | 77 |
| 50 | 6 | 52 | 4 |
| 26 | 10 | 30 | 6 |
| 50 | 0 | 49 | 1 |
| 109 | 19 | 92 | 36 |
| 168 | 32 | 178 | 22 |
| 87 | 12 | 92 | 7 |
| 48 | 3 | 48 | 3 |
| 237 | 144 | 353 | 28 |
| 48 | 7 | 51 | 4 |
| 199 | 39 | 173 | 65 |
| 107 | 86 | 188 | 5 |
| 36 | 80 | 104 | 12 |
| 38 | 13 | 42 | 9 |
| 59 | 77 | 129 | 7 |
| 215 | 17 | 216 | 16 |
| 41 | 3 | 43 | 1 |
| 132 | 41 | 163 | 10 |
| 45 | 5 | 48 | 2 |
| 84 | 80 | 149 | 15 |
| 239 | 17 | 246 | 10 |
| 30 | 1 | 31 | 0 |
| 199 | 33 | 222 | 10 |
| 5 | 0 | 4 | 1 |
| 36 | 52 | 64 | 24 |
| 13 | 3 | 13 | 3 |
| 125 | 15 | 132 | 8 |
| 73 | 8 | 76 | 5 |
| 230 | 37 | 246 | 21 |
| 12 | 2 | 14 | 0 |
| 135 | 31 | 154 | 12 |
| 421 | 231 | 580 | 72 |
| 70 | 21 | 77 | 14 |
| 44 | 11 | 27 | 28 |

| | | | |
|---|---|---|---|
| 94 | 12 | 89 | 17 |
| 69 | 15 | 74 | 10 |
| 98 | 39 | 130 | 7 |
| 280 | 181 | 443 | 18 |
| 87 | 8 | 91 | 4 |
| 11 | 2 | 11 | 2 |
| 28 | 53 | 60 | 21 |
| 90 | 14 | 98 | 6 |
| 252 | 16 | 254 | 14 |
| 52 | 9 | 59 | 2 |
| 60 | 16 | 70 | 6 |
| 21 | 2 | 19 | 4 |
| 107 | 18 | 115 | 10 |
| 270 | 48 | 307 | 11 |
| 41 | 18 | 55 | 4 |
| 49 | 9 | 50 | 8 |
| 115 | 13 | 122 | 6 |
| 711 | 95 | 762 | 44 |
| 156 | 34 | 178 | 12 |
| 393 | 48 | 423 | 18 |
| 283 | 451 | 684 | 50 |
| 26 | 49 | 70 | 5 |
| 27 | 1 | 24 | 4 |
| 29 | 1 | 22 | 8 |
| 45 | 1 | 42 | 4 |
| 86 | 5 | 89 | 2 |
| 85 | 17 | 94 | 8 |
| 82 | 6 | 88 | 0 |
| 91 | 16 | 103 | 4 |
| 37 | 2 | 38 | 1 |
| 36 | 36 | 49 | 23 |
| 147 | 44 | 177 | 14 |
| 89 | 69 | 104 | 54 |
| 80 | 6 | 80 | 6 |
| 47 | 2 | 47 | 2 |
| 47 | 9 | 50 | 6 |
| 130 | 5 | 127 | 8 |
| 102 | 10 | 101 | 11 |
| 105 | 7 | 107 | 5 |
| 155 | 8 | 160 | 3 |
| 106 | 217 | 207 | 116 |
| 76 | 18 | 88 | 6 |
| 21 | 2 | 22 | 1 |
| 346 | 52 | 355 | 43 |
| 98 | 11 | 101 | 8 |

| | | | |
|---|---|---|---|
| 72 | 18 | 87 | 3 |
| 169 | 18 | 170 | 17 |
| 59 | 24 | 68 | 15 |
| 220 | 204 | 379 | 45 |
| 183 | 12 | 180 | 15 |
| 21 | 0 | 21 | 0 |
| 95 | 55 | 147 | 3 |
| 122 | 10 | 125 | 7 |
| 139 | 128 | 252 | 15 |
| 108 | 19 | 122 | 5 |
| 38 | 8 | 41 | 5 |
| 61 | 7 | 63 | 5 |
| 96 | 59 | 141 | 14 |
| 47 | 7 | 38 | 16 |
| 8 | 2 | 9 | 1 |
| 36 | 6 | 39 | 3 |
| 44 | 12 | 52 | 4 |
| 77 | 4 | 77 | 4 |
| 100 | 59 | 102 | 57 |
| 105 | 111 | 203 | 13 |
| 28 | 20 | 44 | 4 |
| 99 | 39 | 125 | 13 |
| 156 | 13 | 87 | 82 |
| 224 | 83 | 287 | 20 |
| 65 | 9 | 71 | 3 |
| 313 | 45 | 337 | 21 |
| 351 | 310 | 622 | 39 |
| 154 | 37 | 176 | 15 |
| 77 | 7 | 79 | 5 |
| 66 | 58 | 115 | 9 |
| 54 | 9 | 61 | 2 |
| 64 | 64 | 98 | 30 |
| 139 | 24 | 151 | 12 |
| 281 | 21 | 289 | 13 |
| 27 | 10 | 32 | 5 |
| 92 | 47 | 122 | 17 |
| 147 | 47 | 172 | 22 |
| 26 | 5 | 27 | 4 |
| 61 | 5 | 59 | 7 |
| 49 | 3 | 47 | 5 |
| 518 | 704 | 669 | 553 |
| 114 | 20 | 125 | 9 |
| 496 | 39 | 496 | 39 |
| 51 | 6 | 52 | 5 |
| 39 | 10 | 46 | 3 |

| | | | |
|---|---|---|---|
| 40 | 43 | 77 | 6 |
| 87 | 79 | 150 | 16 |
| 179 | 44 | 213 | 10 |
| 67 | 3 | 66 | 4 |
| 104 | 15 | 111 | 8 |
| 70 | 8 | 68 | 10 |
| 65 | 13 | 70 | 8 |
| 26 | 6 | 30 | 2 |
| 92 | 26 | 112 | 6 |
| 219 | 24 | 225 | 18 |
| 49 | 8 | 55 | 2 |
| 469 | 86 | 481 | 74 |
| 79 | 27 | 96 | 10 |
| 0 | 0 | 0 | 0 |
| 256 | 12 | 262 | 6 |
| 66 | 11 | 67 | 10 |
| 129 | 17 | 131 | 15 |
| 63 | 17 | 71 | 9 |
| 115 | 14 | 112 | 17 |
| 290 | 85 | 350 | 25 |
| 39 | 7 | 44 | 2 |
| 132 | 68 | 186 | 14 |
| 65 | 36 | 95 | 6 |
| 29 | 7 | 33 | 3 |
| 442 | 58 | 473 | 27 |
| 33 | 19 | 47 | 5 |
| 126 | 20 | 119 | 27 |
| 208 | 85 | 284 | 9 |
| 84 | 6 | 83 | 7 |
| 228 | 227 | 436 | 19 |
| 0 | 0 | 0 | 0 |
| 258 | 8 | 260 | 6 |
| 147 | 29 | 162 | 14 |
| 35 | 12 | 43 | 4 |
| 84 | 11 | 91 | 4 |
| 15 | 8 | 19 | 4 |
| 253 | 222 | 458 | 17 |
| 348 | 368 | 647 | 69 |
| 320 | 270 | 250 | 340 |
| 10 | 1 | 10 | 1 |
| 130 | 14 | 129 | 15 |
| 68 | 87 | 97 | 58 |
| 137 | 19 | 151 | 5 |
| 224 | 18 | 231 | 11 |
| 19 | 5 | 19 | 5 |

| | | | |
|---|---|---|---|
| 24 | 12 | 28 | 8 |
| 187 | 15 | 196 | 6 |
| 57 | 14 | 67 | 4 |
| 54 | 4 | 55 | 3 |
| 11 | 1 | 10 | 2 |
| 132 | 115 | 228 | 19 |
| 71 | 10 | 79 | 2 |
| 688 | 1071 | 1681 | 78 |
| 165 | 228 | 371 | 22 |
| 24 | 3 | 25 | 2 |
| 124 | 70 | 175 | 19 |
| 58 | 69 | 59 | 68 |
| 59 | 9 | 60 | 8 |
| 67 | 15 | 82 | 0 |
| 71 | 110 | 95 | 86 |
| 642 | 228 | 0 | 870 |
| 95 | 5 | 98 | 2 |
| 98 | 32 | 124 | 6 |
| 641 | 145 | 742 | 44 |
| 208 | 27 | 221 | 14 |
| 208 | 27 | 221 | 14 |
| 37 | 73 | 38 | 72 |
| 105 | 23 | 121 | 7 |
| 73 | 42 | 105 | 10 |
| 87 | 10 | 93 | 4 |
| 86 | 7 | 88 | 5 |
| 78 | 30 | 91 | 17 |
| 44 | 3 | 44 | 3 |
| 64 | 10 | 71 | 3 |
| 56 | 73 | 124 | 5 |
| 51 | 11 | 51 | 11 |
| 23 | 10 | 30 | 3 |
| 129 | 20 | 133 | 16 |
| 67 | 5 | 69 | 3 |
| 54 | 8 | 60 | 2 |
| 247 | 29 | 253 | 23 |
| 184 | 12 | 188 | 8 |
| 26 | 5 | 24 | 7 |
| 139 | 24 | 149 | 14 |
| 367 | 23 | 367 | 23 |
| 19 | 20 | 37 | 2 |
| 221 | 38 | 244 | 15 |
| 53 | 6 | 57 | 2 |
| 61 | 20 | 76 | 5 |
| 37 | 1 | 34 | 4 |

| | | | |
|---|---|---|---|
| 99 | 96 | 126 | 69 |
| 101 | 16 | 110 | 7 |
| 102 | 30 | 117 | 15 |
| 81 | 14 | 85 | 10 |
| 236 | 44 | 268 | 12 |
| 122 | 29 | 147 | 4 |
| 23 | 1 | 24 | 0 |
| 91 | 18 | 102 | 7 |
| 253 | 636 | 488 | 401 |
| 44 | 1 | 43 | 2 |
| 88 | 5 | 87 | 6 |
| 362 | 79 | 407 | 34 |
| 100 | 13 | 109 | 4 |
| 221 | 175 | 357 | 39 |
| 3 | 0 | 2 | 1 |
| 37 | 66 | 93 | 10 |
| 129 | 10 | 135 | 4 |
| 296 | 70 | 334 | 32 |
| 319 | 81 | 376 | 24 |
| 125 | 21 | 136 | 10 |
| 82 | 169 | 96 | 155 |
| 44 | 4 | 43 | 5 |
| 103 | 13 | 101 | 15 |
| 96 | 4 | 93 | 7 |
| 43 | 3 | 43 | 3 |
| 142 | 52 | 187 | 7 |
| 43 | 37 | 72 | 8 |
| 80 | 11 | 87 | 4 |
| 320 | 35 | 322 | 33 |
| 10 | 9 | 14 | 5 |
| 63 | 4 | 65 | 2 |
| 23 | 2 | 23 | 2 |
| 325 | 127 | 418 | 34 |
| 47 | 3 | 43 | 7 |
| 19 | 2 | 21 | 0 |
| 309 | 64 | 348 | 25 |
| 81 | 11 | 85 | 7 |
| 36 | 15 | 45 | 6 |
| 120 | 47 | 160 | 7 |
| 22 | 0 | 20 | 2 |
| 30 | 1 | 30 | 1 |
| 58 | 7 | 58 | 7 |
| 16 | 23 | 31 | 8 |
| 138 | 20 | 149 | 9 |
| 82 | 5 | 84 | 3 |

| | | | |
|---|---|---|---|
| 107 | 153 | 138 | 122 |
| 61 | 17 | 72 | 6 |
| 41 | 7 | 45 | 3 |
| 105 | 14 | 110 | 9 |
| 18 | 4 | 20 | 2 |
| 86 | 114 | 194 | 6 |
| 10 | 4 | 9 | 5 |
| 50 | 17 | 63 | 4 |
| 78 | 13 | 74 | 17 |
| 223 | 84 | 272 | 35 |
| 96 | 17 | 108 | 5 |
| 80 | 17 | 89 | 8 |
| 55 | 48 | 96 | 7 |
| 33 | 3 | 34 | 2 |
| 5 | 0 | 5 | 0 |
| 170 | 79 | 233 | 16 |
| 70 | 7 | 74 | 3 |
| 204 | 49 | 237 | 16 |
| 44 | 4 | 45 | 3 |

LQ) to DNA damage sites during mitosis

response

diated cell signaling

**Supplementary Table S3**

**A** Gene set used to evaluate for nsSNVs
**B** Non-synonymous Variants (SNVs) Unique to Jonathan
**C** Key for abbreviations in tab "A"

### **A** Gene set used to evaluate for nsSNVs

| gene | ergo id | GG | CABP | NAMR | BRBA | ANOC |
|---|---|---|---|---|---|---|
| A2M | RHSPN17203 | RGG010277 | RCABP01030 | RNAMR035611 | | RANOC00813 |
| ABCC4 | RHSPN18945 | RGG019014 | RCABP01686 | RNAMR03689 | RBRBA03524 | RANOC00854 |
| ABCC8 | RHSPN157638 | | RCABP02877 | RNAMR01760 | RBRBA03653 | RANOC00025 |
| ABI1 | RHSPN14404 | RGG026605 | RCABP01711 | RNAMR02598 | RBRBA00165 | RANOC01809 |
| ABL1 | RHSPN13995 | RGG132491 | RCABP02739 | RNAMR01520 | RBRBA03852 | RANOC03273 |
| ABL2 | RHSPN02171 | RGG094257 | RCABP02604 | RNAMR01943 | RBRBA03254 | RANOC02418 |
| ABO | RHSPN140416 | | not predicted | | | |
| ABR | RHSPN22164 | RGG139978 | RCABP00155 | RNAMR02318 | RBRBA003493 | |
| ACD | RHSPN21842 | RGG109843 | RCABP03558 | RNAMR04173 | RBRBA02745 | RANOC02396 |
| ACE | RHSPN23272 | RGG158369 | RCABP03444 | RNAMR03467 | RBRBA00389 | RANOC01768 |
| ACKR3 | RHSPN04807 | RGG086794 | RCABP02953 | RNAMR03443 | RBRBA02133 | RANOC00045 |
| ACSL3 | RHSPN04678 | RGG100922 | RCABP01971 | RNAMR03053 | RBRBA036796 | |
| ACSL6 | RHSPN08520 | RGG120309 | RCABP04174 | RNAMR00274 | RBRBA01302 | RANOC00637 |
| ACTN3 | RHSPN163399 | | RCABP00297 | RNAMR01839 | RBRBA01140 | RANOC03137 |
| ACVR1 | RHSPN04113 | RGG092521 | RCABP00203 | RNAMR00202 | RBRBA00488 | RANOC02245 |
| ADA | RHSPN26238 | RGG142297 | RCABP04018 | RNAMR02427 | RBRBA02065 | RANOC01178 |
| ADAMTSL1 | RHSPN13069 | RGG173984 | RCABP02729 | RNAMR00450 | RBRBA01840 | RANOC00551 |
| ADAR | RHSPN01774 | RGG154128 | RCABP01997 | RNAMR0374 | RBRBA01560 | RANOC03441 |
| ADARB1 | RHSPN26842 | RGG087464 | RCABP03322 | RNAMR00862 | RBRBA03083 | RANOC01252 |
| ADARB2 | RHSPN14192 | RGG025992 | RCABP01342 | RNAMR02079 | RBRBA03278 | RANOC01717 |
| ADCY5 | RHSPN05963 | RGG091118 | RCABP02683 | RNAMR03968 | RBRBA02313 | RANOC00032 |
| ADGB | RHSPN10317 | RGG047594 | RCABP02034 | RNAMR02300 | RBRBA03708 | RANOC00258 |
| ADGRL4 | RHSPN01155 | RGG096568 | RCABP03608 | RNAMR00749 | RBRBA02170 | RANOC01910 |
| ADIPOQ | RHSPN06474 | RGG100519 | RCABP02808 | RNAMR00092 | RBRBA02729 | RANOC00834 |
| ADORA2B | RHSPN22448 | RGG140006 | RCABP00156 | RNAMR03684 | RBRBA019258 | |
| AFF1 | RHSPN072092 | | RCABP02970 | RNAMR0366 | RBRBA02437 | RANOC01520 |
| AFF3 | RHSPN03700 | RGG017593 | RCABP03834 | RNAMR01053 | RBRBA00350 | RANOC02745 |
| AFF4 | RHSPN08543 | RGG120466 | RCABP04178 | RNAMR00269 | RBRBA01306 | RANOC00460 |
| AGER | RHSPN094806 | | not predicted | RNAMR01298 | RBRBA005493 | |
| AGPAT2 | RHSPN14112 | RGG132965 | RCABP04022 | RNAMR01503 | RBRBA034667 | |
| AGT | RHSPN02675 | RGG046852 | RCABP03531 | RNAMR02077 | RBRBA00087 | RANOC00080 |
| AGTR1 | RHSPN06197 | RGG101595 | RCABP03516 | RNAMR0005 | RBRBA01032 | RANOC00903 |
| AGTR2 | RHSPN28229 | RGG056028 | RCABP00832 | RNAMR04128 | RBRBA03210 | RANOC02660 |
| AHSG | RHSPN06467 | RGG102707 | RCABP03301 | RNAMR00093 | RBRBA00359 | RANOC02464 |
| AICDA | RHSPN17191 | RGG010177 | RCABP01034 | RNAMR0356 | RBRBA02685 | RANOC00439 |
| AIF1 | RHSPN094384 | | RCABP00203 | RNAMR01310 | RBRBA000139 | |
| AIFM1 | RHSPN28297 | RGG055285 | RCABP00174 | RNAMR02614 | RBRBA02266 | RANOC02737 |

| | | | | | | |
|---|---|---|---|---|---|---|
| AIRE | RHSPN2682 | RGG099351 | RCABP03968 | RNAMR0085 | RBRBA03276 | RANOC0104 |
| AKAP10 | RHSPN2254 | RGG140151 | RCABP00172 | RNAMR0348 | RBRBA01924 | RANOC0263 |
| AKAP9 | RHSPN1120 | RGG027473 | RCABP02778 | RNAMR0284 | RBRBA02705 | RANOC0172 |
| AKT1 | RHSPN2001 | RGG077555 | RCABP04142 | RNAMR002957 | | RANOC0010 |
| AKT1S1 | RHSPN254146 | | RCABP04121 | RNAMR0329 | RBRBA0316 | RANOC0336 |
| AKT2 | RHSPN2506 | RGG168936 | RCABP02556 | RNAMR0402 | RBRBA01732 | RANOC0187 |
| AKT3 | RHSPN0277 | RGG046191 | RCABP03050 | RNAMR0196 | RBRBA01162 | RANOC0236 |
| ALDH2 | RHSPN1823 | RGG126749 | RCABP02995 | RNAMR0098 | RBRBA02720 | RANOC0234 |
| ALG1 | RHSPN2123 | RGG124316 | RCABP04315 | RNAMR0309 | RBRBA02290 | RANOC0296 |
| ALK | RHSPN0310 | RGG073458 | RCABP00686 | RNAMR0053 | RBRBA00756 | RANOC0029 |
| ALKBH2 | RHSPN1818 | RGG126929 | RCABP03442 | RNAMR0097 | RBRBA018521 | |
| ALKBH3 | RHSPN1591 | RGG072482 | RCABP02088 | RNAMR0180 | RBRBA01994 | RANOC0002 |
| AMER1 | RHSPN2795 | RGG057361 | RCABP04414 | RNAMR0383 | RBRBA03550 | RANOC0316 |
| ANG | RHSPN1913 | RGG062077 | RCABP037576 | | RBRBA01342 | RANOC0208 |
| ANGEL1 | RHSPN1973 | RGG075696 | RCABP03123 | RNAMR0032 | RBRBA00593 | RANOC0024 |
| ANGPT1 | RHSPN1266 | RGG039044 | RCABP00135 | RNAMR0164 | RBRBA01770 | RANOC0111 |
| ANGPTL8 | RHSPN245140 | | not predicted | RNAMR0316 | RBRBA000266 | |
| ANKRD17 | RHSPN0708 | RGG062328 | not predicted | RNAMR0041 | RBRBA034372 | |
| ANO9 | RHSPN1547 | RGG068803 | RCABP00532 | RNAMR0178 | RBRBA00475 | RANOC0015 |
| ANXA1 | RHSPN1338 | RGG174481 | RCABP00353 | RNAMR0286 | RBRBA01661 | RANOC0042 |
| ANXA3 | RHSPN071465 | | RCABP00979 | RNAMR0141 | RBRBA00226 | RANOC0145 |
| AOPEP | RHSPN1356 | RGG175136 | RCABP00619 | RNAMR0121 | RBRBA02393 | RANOC0073 |
| AP1G1 | RHSPN2192 | RGG112794 | RCABP03318 | RNAMR0090 | RBRBA03378 | RANOC0279 |
| APBB1IP | RHSPN1439 | RGG026648 | RCABP01713 | RNAMR0260 | RBRBA00165 | RANOC0166 |
| APBB2 | RHSPN0691 | RGG064754 | RCABP00867 | RNAMR0159 | RBRBA02885 | RANOC0163 |
| APC | RHSPN0842 | RGG175598 | RCABP04095 | RNAMR0078 | RBRBA01427 | RANOC0190 |
| APC2 | RHSPN2423 | RGG161767 | RCABP00024 | RNAMR0320 | RBRBA01460 | RANOC0331 |
| APEX1 | RHSPN1912 | RGG168061 | RCABP03862 | RNAMR0310 | RBRBA01385 | RANOC02085 |
| APEX2 | RHSPN279280 | | RCABP04264 | RNAMR0396 | RBRBA00689 | RANOC0007 |
| APLF | RHSPN0338 | RGG042879 | RCABP03527 | RNAMR0160 | RBRBA03818 | RANOC0036 |
| APOA1 | RHSPN1681 | RGG152224 | RCABP02598 | RNAMR0227 | RBRBA001493 | |
| APOA2 | RHSPN0199 | RGG154570 | RCABP03811 | RNAMR024750 | | |
| APOA4 | RHSPN1681 | RGG152213 | RCABP02598 | RNAMR0227 | RBRBA00150 | RANOC0319 |
| APOA5 | RHSPN1681 | RGG152210 | RCABP02601 | RNAMR0227 | RBRBA001502 | |
| APOB | RHSPN0298 | RGG053373 | RCABP01375 | RNAMR0051 | RBRBA02548 | RANOC0022 |
| APOBEC1 | RHSPN171750 | | RCABP01034 | RNAMR0355 | RBRBA02685 | RANOC0072 |
| APOBEC2 | RHSPN0963 | RGG157244 | RCABP00421 | RNAMR0127 | RBRBA03930 | RANOC0116 |
| APOBEC3A | RHSPN273542 | | not predicted | | | |
| APOBEC3B | RHSPN273549 | | not predicted | RNAMR012007 | | |
| APOBEC3C | RHSPN273555 | | not predicted | | | |
| APOBEC3D | RHSPN273560 | | not predicted | | RBRBA016072 | |
| APOBEC3F | RHSPN273569 | | not predicted | | RBRBA016072 | |
| APOBEC3G | RHSPN273575 | | not predicted | | | |
| APOBEC3H | RHSPN273594 | | not predicted | | RBRBA016072 | |
| APOBEC4 | RHSPN0222 | RGG094690 | not predicted | RNAMR0288 | RBRBA03397 | RANOC0199 |

| | | | | | | |
|---|---|---|---|---|---|---|
| APOC1 | RHSPN252176 | | not predicte | RNAMR039207 | | |
| APOC2 | RHSPN252189 | | not predicte | RNAMR0392 | RBRBA010177 | |
| APOC3 | RHSPN1681 | RGG152220 | RCABP0259 | RNAMR0227 | RBRBA000056 | |
| APOC4 | RHSPN252186 | | not predicte | RNAMR039209 | | |
| APOD | RHSPN0653 | RGG101690 | RCABP018304 | | RBRBA023272 | |
| APOE | RHSPN252166 | | not predicte | RNAMR0392 | RBRBA010178 | |
| APOF | RHSPN1775 | RGG167016 | RCABP0164 | RNAMR0274 | RBRBA027911 | |
| APOH | RHSPN2332 | RGG136101 | RCABP0008 | RNAMR0363 | RBRBA0039 | RANOC0226 |
| APOL1 | | | not predicted | | | |
| APOL2 | RHSPN272822 | | not predicte | RNAMR011795 | | |
| APOL3 | RHSPN272800 | | not predicted | | RBRBA03642 | RANOC0148 |
| APOL4 | RHSPN272810 | | not predicted | | | |
| APOL5 | RHSPN272726 | | not predicted | | RBRBA036454 | |
| APOL6 | RHSPN272720 | | RCABP00691 | RNAMR011822 | | |
| APOLD1 | RHSPN1726 | RGG005381 | RCABP0271 | RNAMR0120 | RBRBA014716 | |
| APOM | RHSPN094463 | | RCABP00202 | RNAMR0130 | RBRBA00554 | RANOC0049 |
| APOO | RHSPN2770 | RGG015837 | RCABP0322 | RNAMR0382 | RBRBA00027 | RANOC0086 |
| APP | RHSPN2661 | RGG013844 | RCABP0132 | RNAMR0083 | RBRBA0072 | RANOC0100 |
| APTX | RHSPN1313 | RGG177867 | RCABP0171 | RNAMR0044 | RBRBA0051 | RANOC0225 |
| AQP4 | RHSPN2385 | RGG036230 | RCABP02862 | RNAMR0192 | RBRBA0017 | RANOC0112 |
| AQP9 | RHSPN2051 | RGG106517 | RCABP00621 | RNAMR0218 | RBRBA036482 | |
| ARHGAP1 | RHSPN1596 | RGG073074 | RCABP0070 | RNAMR0180 | RBRBA0200 | RANOC0011 |
| ARHGAP26 | RHSPN0870 | RGG120531 | RCABP0417 | RNAMR0204 | RBRBA0336 | RANOC0046 |
| ARHGAP8 | RHSPN2746 | RGG009337 | RCABP03381 | RNAMR011520 | | |
| ARHGDIB | RHSPN1728 | RGG005236 | RCABP0156 | RNAMR0119 | RBRBA01061 | RANOC0141 |
| ARHGEF12 | RHSPN1690 | RGG151862 | RCABP001223 | | RBRBA0334 | RANOC0185 |
| ARID1A | RHSPN0050 | RGG148803 | RCABP0321 | RNAMR0167 | RBRBA0257 | RANOC0280 |
| ARID1B | RHSPN1039 | RGG048045 | RCABP0292 | RNAMR0230 | RBRBA0228 | RANOC0026 |
| ARID2 | RHSPN1748 | RGG003322 | RCABP0143 | RNAMR0292 | RBRBA0344 | RANOC0142 |
| ARNT | RHSPN0165 | RGG154678 | RCABP04064 | RNAMR0201 | RBRBA0154 | RANOC0317 |
| ARNTL | RHSPN1572 | RGG069664 | RCABP02214 | RNAMR0175 | RBRBA0329 | RANOC0216 |
| ARPP19 | RHSPN2047 | RGG106788 | RCABP0102 | RNAMR0362 | RBRBA010886 | |
| ARRB2 | RHSPN222478 | | RHSPN2224 | RNAMR0233 | RBRBA035441 | |
| ART1 | RHSPN1557 | RGG023245 | RCABP0024 | RNAMR0295 | RBRBA0151 | RANOC0331 |
| ASIC2 | RHSPN2268 | RGG158233 | RCABP0272 | RNAMR0366 | RBRBA0290 | RANOC0168 |
| ASPSCR1 | RHSPN2362 | RGG135473 | RCABP03754 | RNAMR0270 | RBRBA03702 | RANOC0062 |
| ASXL1 | RHSPN2606 | RGG143613 | RCABP03562 | RNAMR0066 | RBRBA03201 | RANOC0283 |
| ATCAY | RHSPN2430 | RGG160900 | RCABP0003 | RNAMR0319 | RBRBA004123 | |
| ATF1 | RHSPN1759 | RGG166318 | RCABP0173 | RNAMR0290 | RBRBA0179 | RANOC0058 |
| ATF2 | RHSPN0425 | RGG089002 | RCABP0156 | RNAMR0238 | RBRBA00251 | RANOC0260 |
| ATF3 | RHSPN024954 | | RCABP0341 | RNAMR0196 | RBRBA0125 | RANOC0008 |
| ATF4 | RHSPN2737 | RGG005935 | RCABP0114 | RNAMR0116 | RBRBA0208 | RANOC0141 |
| ATF5 | RHSPN254191 | | RCABP02772 | RNAMR0329 | RBRBA0317 | RANOC0224 |
| ATF6 | RHSPN0201 | RGG093430 | RCABP0339 | RNAMR0312 | RBRBA0328 | RANOC0113 |
| ATF6B | RHSPN094745 | | RCABP0364 | RNAMR0130 | RBRBA0055 | RANOC0070 |

| | | | | | | |
|---|---|---|---|---|---|---|
| ATF7 | RHSPN17673 | RGG166608 | RCABP01757 | RNAMR0388 | RBRBA01780 | RANOC00564 |
| ATG101 | RHSPN17625 | RGG166243 | RCABP01658 | RNAMR02899 | RBRBA01789 | RANOC00580 |
| ATG12 | RHSPN08439 | RGG178153 | RCABP01973 | RNAMR00228 | RBRBA00217 | RANOC01895 |
| ATG16L1 | RHSPN04775 | RGG099529 | RCABP03099 | RNAMR03418 | RBRBA03600 | RANOC00839 |
| ATG4C | RHSPN01040 | RGG098717 | RCABP02543 | RNAMR00763 | RBRBA02968 | RANOC01157 |
| ATG5 | RHSPN10030 | RGG050458 | RCABP01474 | RNAMR02153 | RBRBA03722 | RANOC00380 |
| ATG9A | RHSPN04639 | RGG090243 | RCABP02199 | RNAMR0306 | RBRBA03672 | RANOC00038 |
| ATIC | RHSPN04571 | RGG086539 | RCABP03430 | RNAMR01719 | RBRBA03510 | RANOC00045 |
| ATM | RHSPN16729 | RGG021743 | RCABP02113 | RNAMR02290 | RBRBA03973 | RANOC02458 |
| ATP1A1 | RHSPN01467 | RGG011308 | RCABP02943 | RNAMR01990 | RBRBA00561 | RANOC02670 |
| ATP2B3 | RHSPN284801 | | RCABP00628 | RNAMR04098 | RBRBA04037 | RANOC00580 |
| ATP4B | RHSPN19077 | RGG018203 | RCABP00954 | RNAMR02457 | RBRBA02916 | RANOC00859 |
| ATP5O | RHSPN26677 | RGG014254 | RCABP03866 | RNAMR00843 | RBRBA00667 | RANOC00877 |
| ATP6V0A2 | RHSPN18412 | RGG126003 | RCABP02997 | RNAMR01000 | RBRBA04052 | RANOC01858 |
| ATP6V1A | RHSPN05877 | RGG011096 | RCABP00575 | RNAMR00180 | RBRBA02341 | RANOC02777 |
| ATP7A | RHSPN28049 | RGG057579 | RCABP03978 | RNAMR03350 | RBRBA028410 | |
| ATR | RHSPN06171 | RGG101410 | RCABP03855 | RNAMR00048 | RBRBA01803 | RANOC00904 |
| ATRIP | RHSPN05396 | RGG114151 | RCABP00512 | RNAMR00003 | RBRBA01276 | RANOC00665 |
| ATRX | RHSPN28047 | RGG057562 | RCABP03979 | RNAMR03359 | RBRBA02841 | RANOC02697 |
| ATXN2 | RHSPN18234 | RGG126701 | RCABP03009 | RNAMR01019 | RBRBA02721 | RANOC02695 |
| ATXN7L3B | RHSPN179155 | | not predicted | RNAMR01113 | RBRBA037654 | |
| AXIN1 | RHSPN21044 | RGG123750 | RCABP01294 | RNAMR03970 | RBRBA01945 | RANOC03333 |
| AXIN2 | RHSPN23317 | RGG136114 | RCABP00086 | RNAMR03634 | RBRBA02656 | RANOC02264 |
| B2M | RHSPN20383 | RGG104922 | RCABP02156 | RNAMR02780 | RBRBA02894 | RANOC02811 |
| BAD | RHSPN16230 | RGG169170 | RCABP03346 | RNAMR0186 | RBRBA011237 | |
| BAG2 | RHSPN09821 | RGG051959 | RCABP01005 | RNAMR01228 | RBRBA01007 | RANOC00062 |
| BAG6 | RHSPN094450 | | RCABP00203 | RNAMR01309 | RBRBA00555 | RANOC00813 |
| BAK1 | RHSPN09522 | RGG157231 | RCABP00422 | RNAMR01287 | RBRBA02962 | RANOC01165 |
| BAMBI | RHSPN14428 | RGG026463 | RCABP03721 | RNAMR02780 | RBRBA02763 | RANOC02552 |
| BANF1 | RHSPN163196 | | RCABP03346 | RNAMR01844 | RBRBA01136 | RANOC01139 |
| BATF | RHSPN19722 | RGG075583 | RCABP03124 | RNAMR00327 | RBRBA01844 | RANOC00000 |
| BAX | RHSPN253519 | | RCABP03537 | RNAMR03308 | RBRBA03162 | RANOC02558 |
| BCL10 | RHSPN01190 | RGG096235 | RCABP02845 | RNAMR00748 | RBRBA02503 | RANOC01910 |
| BCL11A | RHSPN03320 | RGG041277 | RCABP03956 | RNAMR03331 | RBRBA03827 | RANOC00095 |
| BCL11B | RHSPN19890 | RGG076848 | RCABP00588 | RNAMR003088 | | RANOC00310 |
| BCL2 | RHSPN24084 | RGG033216 | RCABP02267 | RNAMR02458 | RBRBA03457 | RANOC03293 |
| BCL2A1 | RHSPN20810 | RGG107630 | RCABP04422 | RNAMR040875 | | |
| BCL2L1 | RHSPN26049 | RGG143480 | RCABP03922 | RNAMR00670 | RBRBA03203 | RANOC03099 |
| BCL2L10 | RHSPN204662 | | | RNAMR00190 | RBRBA007632 | |
| BCL2L11 | RHSPN03820 | RGG041565 | RCABP04305 | RNAMR01041 | RBRBA001977 | |
| BCL2L12 | RHSPN253994 | | | RNAMR03298 | RBRBA03168 | RANOC01800 |
| BCL2L13 | RHSPN26912 | RGG008030 | RCABP01993 | RNAMR00370 | RBRBA03497 | RANOC01477 |
| BCL2L14 | RHSPN17254 | RGG009612 | RCABP03221 | RNAMR01197 | RBRBA014733 | |
| BCL2L15 | RHSPN01439 | RGG156878 | RCABP03946 | RNAMR01987 | RBRBA005660 | |
| BCL2L2 | RHSPN192385 | | | RNAMR03024 | RBRBA035016 | |

| | | | | | | |
|---|---|---|---|---|---|---|
| BCL3 | RHSPN252118 | | RCABP01045 | RNAMR0391 | RBRBA01018 | RANOC0335 |
| BCL6 | RHSPN0648 | RGG102114 | RCABP00907 | RNAMR0009 | RBRBA02727 | RANOC0318 |
| BCL7A | RHSPN1836 | RGG126393 | RALGO09354 | RNAMR0099 | RBRBA02609 | RANOC0289 |
| BCL9 | RHSPN0158 | RGG013307 | RCABP01241 | RNAMR0199 | RBRBA01269 | RANOC0100 |
| BCOR | RHSPN2775 | RGG015314 | RCABP02568 | RNAMR0161 | RBRBA03334 | RANOC0098 |
| BCR | RHSPN2707 | RGG127634 | RCABP03064 | RNAMR0096 | RBRBA04054 | RANOC0319 |
| BDKRB2 | RHSPN198782 | | RCABP04206 | RNAMR0031 | RBRBA01693 | RANOC0000 |
| BDNF | RHSPN1583 | RGG069132 | RCABP04329 | RNAMR0176 | RBRBA02309 | RANOC0244 |
| BECN1 | RHSPN2292 | RGG159983 | RCABP00500 | RNAMR0253 | RBRBA02644 | RANOC0169 |
| BHLHE41 | RHSPN1736 | RGG008852 | RCABP04003 | RNAMR0118 | RBRBA02150 | RANOC0255 |
| BID | RHSPN2691 | RGG008044 | RCABP01993 | RNAMR0037 | RBRBA03494 | RANOC0160 |
| BIN2 | RHSPN1760 | RGG167380 | not predicted | RNAMR0292 | RBRBA01797 | RANOC0056 |
| BIRC2 | RHSPN1669 | RGG021988 | RCABP03605 | RNAMR0368 | RBRBA02202 | RANOC0088 |
| BIRC3 | RHSPN166899 | | | RNAMR0368 | RBRBA022021 | |
| BIRC5 | RHSPN2352 | RGG043350 | RCABP02579 | RNAMR0269 | RBRBA00040 | RANOC0060 |
| BLK | RHSPN1198 | RGG054417 | RCABP03662 | RNAMR0145 | RBRBA00525 | RANOC0290 |
| BLM | RHSPN2092 | RGG108895 | RCABP01928 | RNAMR0189 | RBRBA03026 | RANOC0064 |
| BLNK | RHSPN1503 | RGG083000 | RCABP04276 | RNAMR0046 | RBRBA03942 | RANOC0227 |
| BMP6 | RHSPN0913 | RGG032754 | RCABP04126 | RNAMR0138 | RBRBA04064 | RANOC0202 |
| BMPR1A | RHSPN1492 | RGG079509 | RCABP01213 | RNAMR0337 | RBRBA02052 | RANOC0195 |
| BNIP2 | RHSPN2053 | RGG106116 | RCABP02817 | RNAMR0218 | RBRBA00717 | RANOC0270 |
| BNIPL | RHSPN016666 | | RCABP04071 | RNAMR0201 | RBRBA015412 | |
| BOK | RHSPN0490 | RGG100447 | RCABP00269 | RNAMR0345 | RBRBA00254 | RANOC0083 |
| BPI | RHSPN2618 | RGG143746 | RCABP03563 | RNAMR0338 | RBRBA00337 | RANOC0292 |
| BRAF | RHSPN1166 | RGG007269 | RCABP01439 | RNAMR0173 | RBRBA02553 | RANOC0140 |
| BRCA1 | RHSPN2293 | RGG160047 | RCABP00497 | RNAMR0253 | RBRBA02642 | RANOC0177 |
| BRCA2 | RHSPN1863 | RGG020996 | RCABP02587 | RNAMR0208 | RBRBA03223 | RANOC0110 |
| BRD3 | RHSPN1406 | RGG132886 | RCABP04019 | RNAMR015095 | | RANOC0307 |
| BRD4 | RHSPN2463 | RGG163422 | RCABP00858 | RNAMR0385 | RBRBA02734 | RANOC0045 |
| BRE | RHSPN0308 | RGG045290 | RCABP02071 | RNAMR0053 | RBRBA02534 | RANOC0046 |
| BRIP1 | RHSPN2325 | RGG140522 | RCABP01087 | RNAMR0365 | RBRBA00381 | RANOC0318 |
| BSCL2 | RHSPN161804 | | RCABP00302 | RNAMR0404 | RBRBA00232 | RANOC0318 |
| BST1 | RHSPN0678 | RGG065596 | RCABP00605 | RNAMR0136 | RBRBA029793 | |
| BTBD19 | RHSPN0083 | RGG097304 | RCABP02094 | RNAMR0389 | RBRBA03388 | RANOC0127 |
| BTBD9 | RHSPN0961 | RGG045444 | not predicted | RNAMR0127 | RBRBA02398 | RANOC0008 |
| BTG3 | RHSPN265907 | | RCABP03246 | RNAMR0083 | RBRBA028782 | |
| BTK | RHSPN2811 | RGG055555 | RCABP04449 | RNAMR0386 | RBRBA01881 | RANOC0273 |
| BTLA | RHSPN058571 | | RCABP01270 | RNAMR0018 | RBRBA02342 | RANOC0101 |
| BTNL2 | RHSPN094868 | | not predicted | | RBRBA005847 | |
| BUB1B | RHSPN2028 | RGG068589 | RCABP03179 | RNAMR0276 | RBRBA00601 | RANOC0228 |
| BUB3 | RHSPN1534 | RGG085072 | RCABP02631 | RNAMR0048 | RBRBA03332 | RANOC0084 |
| C16orf87 | RHSPN2163 | RGG110913 | RCABP00979 | RNAMR039953 | | RANOC0187 |
| C19orf40 | RHSPN2485 | RGG111209 | RCABP00553 | RNAMR0248 | RBRBA00898 | RANOC0279 |
| C1QA | RHSPN0041 | RGG146794 | not predicted | RNAMR0166 | RBRBA00528 | RANOC0342 |
| C1QB | RHSPN0041 | RGG146787 | RCABP01839 | RNAMR0166 | RBRBA005274 | |

| | | | | | | |
|---|---|---|---|---|---|---|
| C1QBP | RHSPN22282 | RGG138654 | RCABP01781 | RNAMR0234 | RBRBA0355 | RANOC0335 |
| C1QC | RHSPN0041 | RGG146790 | RCABP01831 | RNAMR0166 | RBRBA005275 | |
| C1QTNF5 | RHSPN1688 | RGG151985 | RCABP0012 | RNAMR0228 | RBRBA0014 | RANOC0311 |
| C1R | RHSPN17162 | RGG010643 | RCABP00602 | RNAMR0356 | RBRBA023836 | |
| C2 | RHSPN09467 | RGG149223 | RCABP0020 | RNAMR0130 | RBRBA00551 | RANOC0072 |
| C20orf194 | RHSPN2583 | RGG067403 | RCABP03872 | RNAMR0068 | RBRBA02517 | RANOC0241 |
| C20orf78 | RHSPN259564 | | | | | |
| C3 | RHSPN2438 | RGG163052 | RCABP00147 | RNAMR0317 | RBRBA0470 | RANOC0057 |
| C3AR1 | RHSPN17182 | RGG008688 | RCABP0103 | RNAMR0355 | RBRBA00932 | RANOC0215 |
| C4BPA | RHSPN024389 | | RCABP02422 | RNAMR0281 | RBRBA039069 | |
| C4BPB | RHSPN024379 | | not predicted | RNAMR0281 | RBRBA039071 | |
| C5 | RHSPN13794 | RGG133208 | RCABP0402 | RNAMR0155 | RBRBA031179 | |
| C5AR1 | RHSPN252900 | | RCABP04321 | RNAMR0331 | RBRBA00282 | RANOC01879 |
| C5AR2 | RHSPN2529 | RGG168960 | not predicted | RNAMR0281 | RBRBA039071 | |
| C6 | RHSPN0801 | RGG171806 | RCABP0318 | RNAMR0146 | RBRBA0268 | RANOC00729 |
| C7 | RHSPN08012 | RGG171801 | RCABP0318 | RNAMR0146 | RBRBA02687 | RANOC00409 |
| C7orf50 | RHSPN10537 | RGG121399 | RCABP0241 | RNAMR006445 | | RANOC02829 |
| C8A | RHSPN0100 | RGG098242 | RCABP0323 | RNAMR0356 | RBRBA0187 | RANOC0126 |
| C8B | RHSPN0100 | RGG098245 | RCABP0323 | RNAMR0356 | RBRBA0187 | RANOC0115 |
| C8G | RHSPN1412 | RGG130994 | RCABP00564 | RNAMR0149 | RBRBA034631 | |
| C8orf4 | RHSPN1225 | RGG147893 | RCABP0066 | RNAMR0214 | RBRBA0009 | RANOC0268 |
| C9 | RHSPN080061 | | RCABP0052 | RNAMR0146 | RBRBA02244 | RANOC0040 |
| CACNA1C | RHSPN1708 | RGG007967 | RCABP02622 | RNAMR0037 | RBRBA01352 | RANOC01604 |
| CACNA1D | RHSPN0555 | RGG115026 | RCABP0332 | RNAMR0304 | RBRBA019356 | |
| CACNB3 | RHSPN17534 | RGG166197 | RCABP0165 | RNAMR0291 | RBRBA02871 | RANOC0059 |
| CACNB4 | RHSPN04087 | RGG092290 | RCABP0148 | RNAMR0360 | RBRBA0048 | RANOC0239 |
| CADM1 | RHSPN1680 | RGG152157 | RCABP0259 | RNAMR0227 | RBRBA0015 | RANOC0277 |
| CALCOCO2 | RHSPN23094 | RGG158911 | RCABP02181 | RNAMR0257 | RBRBA01637 | RANOC0179 |
| CALR | RHSPN2457 | RGG163370 | RCABP0021 | RNAMR0384 | RBRBA03427 | RANOC0075 |
| CAMK4 | RHSPN0841 | RGG175656 | RCABP00022 | RNAMR0078 | RBRBA0142 | RANOC0190 |
| CAMP | RHSPN05387 | RGG025126 | RCABP02914 | RNAMR0010 | RBRBA0152 | RANOC0250 |
| CAMP4 | | | | | | |
| CAMTA1 | RHSPN00172 | RGG144916 | RCABP03684 | RNAMR0297 | RBRBA0247 | RANOC0326 |
| CANT1 | RHSPN2354 | RGG136723 | RCABP0286 | RNAMR0269 | RBRBA0377 | RANOC0059 |
| CARD11 | RHSPN1057 | RGG121655 | RCABP0404 | RNAMR0065 | RBRBA0366 | RANOC0253 |
| CARD14 | RHSPN235621 | | RCABP0269 | RNAMR0269 | RBRBA0325 | RANOC0076 |
| CARD16 | RHSPN167091 | | not predicted | | | |
| CARD18 | RHSPN167111 | | not predicted | | | |
| CARD6 | RHSPN080115 | | not predicted | RNAMR0146 | RBRBA026877 | |
| CARD8 | RHSPN25317 | RGG128319 | not predicted | | | |
| CARD9 | RHSPN14097 | RGG133046 | RCABP0403 | RNAMR015007 | | |
| CARS | RHSPN15562 | RGG070665 | not predicted | RNAMR0182 | RBRBA04007 | RANOC0002 |
| CASC5 | RHSPN20294 | RGG068697 | RCABP01447 | RNAMR0278 | RBRBA0060 | RANOC0227 |
| CASP12 | | | | | | |
| CASP5 | | | | | | |

| | | | | | | |
|---|---|---|---|---|---|---|
| CASP8 | RHSPN04439 | RGG087945 | RCABP02899 | RNAMR03835 | RBRBA003285 | |
| CAT | RHSPN15876 | RGG072173 | RCABP04312 | RNAMR01815 | RBRBA01157 | RANOC02350 |
| CAV1 | RHSPN11478 | RGG002809 | RCABP04324 | RNAMR03134 | RBRBA02915 | RANOC01425 |
| CBFB | RHSPN21812 | RGG110252 | RCABP01669 | RNAMR00917 | RBRBA02739 | RANOC01445 |
| CBL | RHSPN16887 | RGG151999 | RCABP00123 | RNAMR02288 | RBRBA00140 | RANOC03118 |
| CBLB | RHSPN05807 | RGG012436 | RCABP03872 | RNAMR00188 | RBRBA02878 | RANOC00875 |
| CBLC | RHSPN252137 | | | RNAMR039197 | | |
| CCDC170 | RHSPN10359 | RGG047826 | RCABP01374 | RNAMR02303 | RBRBA03844 | RANOC02260 |
| CCDC6 | RHSPN14677 | RGG080216 | RCABP04147 | RNAMR02437 | RBRBA01242 | RANOC02227 |
| CCDC85A | RHSPN03306 | RGG041175 | RCABP00899 | RNAMR03333 | RBRBA03829 | RANOC00099 |
| CCDC88C | RHSPN19818 | RGG076338 | RCABP02055 | RNAMR00310 | RBRBA020609 | |
| CCL1 | RHSPN22693 | RGG139036 | not predicted | | RBRBA016480 | |
| CCL11 | RHSPN226923 | | not predicted | | RBRBA016480 | |
| CCL13 | RHSPN226929 | | not predicted | | RBRBA016480 | |
| CCL16 | RHSPN227264 | | not predicted | | RBRBA016480 | |
| CCL17 | RHSPN21729 | RGG109639 | RCABP00616 | RNAMR00927 | RBRBA023673 | |
| CCL2 | RHSPN226917 | | not predicted | | RBRBA016480 | |
| CCL20 | RHSPN04712 | RGG101179 | RCABP03898 | RNAMR03408 | RBRBA03543 | RANOC00900 |
| CCL3 | RHSPN227291 | | RCABP020068 | | | |
| CCL4 | RHSPN227294 | | RCABP02023 | RNAMR03660 | RBRBA000180 | |
| CCL5 | RHSPN22721 | RGG138067 | RCABP02006 | RNAMR03673 | RBRBA006001 | |
| CCL7 | RHSPN226920 | | not predicted | | | |
| CCL8 | RHSPN226926 | | not predicted | | RBRBA029081 | |
| CCNB1IP1 | RHSPN191171 | | RCABP02102 | RNAMR03108 | RBRBA01385 | RANOC02097 |
| CCND1 | RHSPN16417 | RGG071895 | RCABP00045 | RNAMR01820 | RBRBA01151 | RANOC00027 |
| CCND2 | RHSPN17102 | RGG009857 | RCABP00487 | RNAMR03543 | RBRBA028237 | |
| CCND3 | RHSPN09652 | RGG157456 | RCABP01676 | RNAMR01268 | RBRBA03990 | RANOC01167 |
| CCNE1 | RHSPN24832 | RGG111047 | RCABP03723 | RNAMR024870 | | RANOC02411 |
| CCNH | RHSPN08296 | RGG176964 | RCABP01394 | RNAMR00770 | RBRBA02100 | RANOC00412 |
| CCR2 | RHSPN053441 | | not predicted | RNAMR00113 | RBRBA028319 | |
| CCR3 | RHSPN053429 | | not predicted | RNAMR00113 | RBRBA028318 | |
| CCR4 | RHSPN05174 | RGG030298 | RCABP00134 | RNAMR02393 | RBRBA02838 | RANOC02474 |
| CCR6 | RHSPN10473 | RGG047185 | RCABP00133 | RNAMR023044 | | RANOC00270 |
| CCR7 | RHSPN22846 | RGG159562 | RCABP01960 | RNAMR02553 | RBRBA00819 | RANOC01787 |
| CD14 | RHSPN086443 | | not predicted | RNAMR00799 | RBRBA033645 | |
| CD164 | RHSPN10064 | RGG050171 | not predicted | RNAMR04003 | RBRBA03719 | RANOC00070 |
| CD180 | RHSPN08144 | RGG172590 | RCABP03421 | RNAMR02893 | RBRBA00294 | RANOC00510 |
| CD19 | RHSPN214983 | | RCABP00806 | RNAMR02213 | RBRBA01940 | RANOC02374 |
| CD1D | RHSPN01916 | RGG130374 | RCABP008803 | | RBRBA01099 | RANOC00688 |
| CD200 | RHSPN05854 | RGG012809 | RCABP01270 | RNAMR00184 | RBRBA02343 | RANOC01017 |
| CD207 | RHSPN034140 | | RCABP02501 | RNAMR01600 | RBRBA015957 | |
| CD209 | RHSPN244112 | | not predicted | RNAMR03324 | RBRBA039117 | |
| CD22 | RHSPN248994 | | RCABP02530 | RNAMR0249 | RBRBA00892 | RANOC02345 |
| CD226 | RHSPN24111 | RGG034974 | RCABP03259 | RNAMR02803 | RBRBA00777 | RANOC01130 |
| CD247 | RHSPN02059 | RGG013110 | RCABP00917 | RNAMR03133 | RBRBA03289 | RANOC00870 |

| | | | | | | |
|---|---|---|---|---|---|---|
| CD27 | RHSPN171240 | | not predicted | RNAMR0354[illegible] | RBRBA023918 | |
| CD274 | RHSPN1300[illegible] | RGG173485 | RCABP0010[illegible] | RNAMR0043[illegible] | RBRBA00731 | RANOC0074[illegible] |
| CD28 | RHSPN0448[illegible] | RGG088286 | RCABP0397[illegible] | RNAMR0174[illegible] | RBRBA02457 | RANOC0202[illegible] |
| CD300A | RHSPN233938 | | not predicted | RNAMR0266[illegible] | RBRBA008066 | |
| CD300E | RHSPN233980 | | RCABP016113 | | RBRBA038666 | |
| CD300LB | RHSPN233949 | | RCABP0161[illegible] | RNAMR0271[illegible] | RBRBA008065 | |
| CD300LF | RHSPN234014 | | RCABP01611 | RNAMR0418 | RBRBA0379[illegible] | RANOC0062[illegible] |
| CD302 | RHSPN041387 | | RCABP013382 | | RBRBA004920 | |
| CD34 | RHSPN0244[illegible] | RGG156585 | RCABP02422 | RNAMR0281[illegible] | RBRBA0386[illegible] | RANOC0224[illegible] |
| CD36 | RHSPN1115[illegible] | RGG001514 | RCABP0313[illegible] | RNAMR0058[illegible] | RBRBA00957 | RANOC0144[illegible] |
| CD37 | RHSPN253688 | | RCABP0353[illegible] | RNAMR0330[illegible] | RBRBA031645 | |
| CD38 | RHSPN0678[illegible] | RGG065590 | RCABP0060[illegible] | RNAMR0133[illegible] | RBRBA0297[illegible] | RANOC0138[illegible] |
| CD3E | RHSPN1684[illegible] | RGG152410 | RCABP00897 | RNAMR0227[illegible] | RBRBA0014[illegible] | RANOC0309[illegible] |
| CD3G | RHSPN168433 | | RCABP0089[illegible] | RNAMR0227[illegible] | RBRBA0014[illegible] | RANOC0309[illegible] |
| CD4 | RHSPN1714[illegible] | RGG010550 | RCABP00324 | RNAMR0355 | RBRBA0238[illegible] | RANOC0274[illegible] |
| CD40LG | RHSPN2837[illegible] | RGG056457 | RCABP0053[illegible] | RNAMR0284 | RBRBA0127[illegible] | RANOC0291[illegible] |
| CD44 | RHSPN1588[illegible] | RGG072290 | RCABP0340[illegible] | RNAMR0181[illegible] | RBRBA01161 | RANOC0033[illegible] |
| CD46 | RHSPN024446 | | not predicted | RNAMR0283[illegible] | RBRBA033447 | |
| CD53 | RHSPN013960 | | not predicted | RNAMR0283[illegible] | RBRBA033447 | |
| CD55 | RHSPN0244[illegible] | RGG156529 | RCABP02282 | RNAMR0283[illegible] | RBRBA0390[illegible] | RANOC0211[illegible] |
| CD58 | RHSPN014697 | | RCABP029416 | | RBRBA00561 | RANOC0267[illegible] |
| CD59 | RHSPN1586[illegible] | RGG072145 | RCABP01504 | RNAMR0187 | RBRBA01157 | RANOC0235[illegible] |
| CD5L | RHSPN019127 | | RCABP02764 | RNAMR024859 | | |
| CD6 | RHSPN1612[illegible] | RGG068123 | RCABP0209[illegible] | RNAMR0405[illegible] | RBRBA01314 | RANOC0214[illegible] |
| CD63 | RHSPN1772[illegible] | RGG167196 | not predicted | RNAMR0274[illegible] | RBRBA02782 | RANOC0194[illegible] |
| CD69 | RHSPN172138 | | RCABP01234 | RNAMR0356[illegible] | RBRBA020797 | |
| CD74 | RHSPN0876[illegible] | RGG119430 | RCABP00452 | RNAMR0205 | RBRBA0195[illegible] | RANOC0064[illegible] |
| CD79A | RHSPN251213 | | RCABP01882 | RNAMR0391[illegible] | RBRBA03717 | RANOC0342[illegible] |
| CD79B | RHSPN2329[illegible] | RGG158189 | RCABP0410[illegible] | RNAMR0347[illegible] | RBRBA0039[illegible] | RANOC0177[illegible] |
| CD80 | RHSPN059122 | | RCABP03384 | RNAMR0019[illegible] | RBRBA02597 | RANOC0100[illegible] |
| CD81 | RHSPN1555[illegible] | RGG070743 | RCABP0043[illegible] | RNAMR0181[illegible] | RBRBA0400[illegible] | RANOC0002[illegible] |
| CD83 | RHSPN0918[illegible] | RGG032332 | RCABP0147[illegible] | RNAMR0137[illegible] | RBRBA03014 | RANOC0122[illegible] |
| CD86 | RHSPN0594[illegible] | RGG010322 | not predicted | RNAMR0396[illegible] | RBRBA026038 | |
| CD8A | RHSPN0354[illegible] | RGG066880 | RCABP0086[illegible] | RNAMR0106[illegible] | RBRBA00357 | RANOC0176[illegible] |
| CD96 | RHSPN058387 | | RCABP033532 | | RBRBA0136[illegible] | RANOC0101[illegible] |
| CD97 | RHSPN246069 | | RCABP0127[illegible] | RNAMR0385[illegible] | RBRBA01342 | RANOC0062[illegible] |
| CDC37 | RHSPN2448[illegible] | RGG163026 | RCABP02264 | RNAMR0315[illegible] | RBRBA00467 | RANOC0041[illegible] |
| CDC42 | RHSPN0040[illegible] | RGG146830 | RCABP0405[illegible] | RNAMR0166[illegible] | RBRBA005292 | |
| CDC73 | RHSPN0226[illegible] | RGG093283 | RCABP0214[illegible] | RNAMR0288[illegible] | RBRBA03761 | RANOC0228[illegible] |
| CDH1 | RHSPN2187[illegible] | RGG112570 | RCABP0438[illegible] | RNAMR0090[illegible] | RBRBA0338[illegible] | RANOC0318[illegible] |
| CDH11 | RHSPN2178[illegible] | RGG111519 | RCABP0408[illegible] | RNAMR0092 | RBRBA0191[illegible] | RANOC0186[illegible] |
| CDH2 | RHSPN2385[illegible] | RGG036259 | RCABP0286[illegible] | RNAMR0192[illegible] | RBRBA0017[illegible] | RANOC0121[illegible] |
| CDH3 | RHSPN2187[illegible] | RGG112565 | | RNAMR0090[illegible] | RBRBA027514 | |
| CDH4 | RHSPN2643[illegible] | RGG142720 | RCABP0315[illegible] | RNAMR0070[illegible] | RBRBA03967 | RANOC0132[illegible] |
| CDK1 | RHSPN1468[illegible] | RGG080177 | RCABP015996 | | RBRBA01242 | RANOC0199[illegible] |

| | | | | | | |
|---|---|---|---|---|---|---|
| CDK12 | RHSPN22804 | RGG159363 | RCABP02169 | RNAMR02559 | RBRBA008265 | |
| CDK14 | RHSPN11200 | RGG027453 | RCABP02779 | RNAMR02847 | RBRBA02705 | RANOC01724 |
| CDK4 | RHSPN178112 | | RCABP01771 | RNAMR02720 | RBRBA03097 | RANOC00433 |
| CDK6 | RHSPN11223 | RGG027570 | RCABP01635 | RNAMR02843 | RBRBA02702 | RANOC01670 |
| CDK7 | RHSPN08153 | RGG172667 | RCABP03798 | RNAMR03889 | RBRBA00304 | RANOC00415 |
| CDK9 | RHSPN13900 | RGG131947 | RCABP02735 | RNAMR01540 | RBRBA03413 | RANOC03277 |
| CDKN1A | RHSPN09586 | RGG155916 | RCABP03390 | RNAMR01278 | RBRBA03448 | RANOC01163 |
| CDKN1B | RHSPN17262 | RGG009698 | RCABP03221 | RNAMR01190 | RBRBA01471 | RANOC02938 |
| CDKN2A | RHSPN130978 | | RCABP014445 | | RBRBA04042 | RANOC00429 |
| CDKN2B | RHSPN13100 | RGG178735 | RCABP01444 | RNAMR004457 | | |
| CDKN2C | RHSPN00924 | RGG097742 | RCABP00409 | RNAMR03534 | RBRBA01904 | RANOC01158 |
| CDKN2D | RHSPN244935 | | RCABP01192 | RNAMR03158 | RBRBA00466 | RANOC00438 |
| CDX4 | RHSPN28028 | RGG057445 | RCABP044161 | | | |
| CEBPA | RHSPN24856 | RGG111260 | RCABP00552 | RNAMR025005 | | RANOC03433 |
| CEBPB | RHSPN26340 | RGG144591 | RCABP03314 | RNAMR02414 | RBRBA01390 | RANOC01178 |
| CEBPG | RHSPN24858 | RGG111268 | RCABP00552 | RNAMR0366 | RBRBA00898 | RANOC02839 |
| CEL | RHSPN14036 | RGG132681 | RCABP00110 | RNAMR01569 | RBRBA02962 | RANOC01697 |
| CEP89 | RHSPN24851 | RGG111200 | RCABP00552 | RNAMR02523 | RBRBA00895 | RANOC02797 |
| CETN2 | RHSPN28464 | RGG057245 | RCABP03966 | RNAMR037133 | | |
| CETP | RHSPN21718 | RGG109554 | RCABP00615 | RNAMR009292 | | RANOC03289 |
| CFD | RHSPN24203 | RGG161597 | RCABP00017 | RNAMR032161 | | |
| CFH | RHSPN02272 | RGG093261 | RCABP01547 | RNAMR02622 | RBRBA03876 | RANOC02709 |
| CFHR5 | RHSPN022782 | | not predicted | | | RANOC02709 |
| CFP | RHSPN278113 | | RCABP04002 | RNAMR04130 | RBRBA02032 | RANOC00550 |
| CGNL1 | RHSPN20507 | RGG106556 | RCABP00620 | RNAMR02183 | RBRBA01498 | RANOC02920 |
| CHAF1A | RHSPN24318 | RGG161382 | RCABP00003 | RNAMR03183 | RBRBA00410 | RANOC03088 |
| CHCHD7 | RHSPN12344 | RGG036836 | RCABP02668 | RNAMR03758 | RBRBA02128 | RANOC02187 |
| CHEK1 | RHSPN16967 | RGG151194 | RCABP04255 | RNAMR04050 | RBRBA02016 | RANOC02465 |
| CHEK2 | RHSPN27143 | RGG127184 | RCABP00394 | RNAMR00940 | RBRBA018583 | |
| CHGA | RHSPN19837 | RGG076474 | RCABP00095 | RNAMR00313 | RBRBA02936 | RANOC00009 |
| CHIA | RHSPN01404 | RGG157477 | RCABP016666 | | RBRBA00573 | RANOC01287 |
| CHIC2 | RHSPN06986 | RGG064213 | RCABP00631 | RNAMR0158 | RBRBA02843 | RANOC01444 |
| CHID1 | RHSPN15514 | RGG071073 | RCABP01708 | RNAMR01773 | RBRBA00472 | RANOC00027 |
| CHIT1 | RHSPN023543 | | RCABP01675 | RNAMR028275 | | |
| CHN1 | RHSPN04249 | RGG089022 | RCABP01560 | RNAMR02379 | RBRBA002515 | |
| CHST11 | RHSPN18137 | RGG006877 | RCABP03564 | RNAMR01092 | RBRBA00792 | RANOC01408 |
| CHUK | RHSPN15093 | RGG080326 | RCABP02630 | RNAMR00470 | RBRBA03933 | RANOC02404 |
| CIC | RHSPN251378 | | RCABP00787 | RNAMR03909 | RBRBA03734 | RANOC03460 |
| CIITA | RHSPN21270 | RGG123345 | RCABP01807 | RNAMR03090 | RBRBA02139 | RANOC02947 |
| CISD2 | RHSPN07298 | RGG063573 | RCABP01416 | RNAMR03234 | RBRBA008711 | |
| CITED1 | RHSPN280184 | | RCABP02197 | RNAMR03872 | RBRBA039411 | |
| CLCF1 | RHSPN16359 | RGG068033 | RCABP02499 | RNAMR01834 | RBRBA01144 | RANOC00357 |
| CLEC2A | RHSPN172171 | | not predicted | | | |
| CLEC2B | RHSPN172156 | | not predictec | RNAMR02812 | RBRBA039071 | |
| CLEC2D | RHSPN17212 | RGG000130 | RCABP018628 | | | RANOC03247 |

| | | | | | | |
|---|---|---|---|---|---|---|
| CLEC2L | RHSPN116543 | | RCABP01638 | RNAMR0174 | RBRBA025538 | |
| CLIP1 | RHSPN18372 | RGG126498 | RCABP03018 | RNAMR0100 | RBRBA02613 | RANOC0269 |
| CLIP2 | RHSPN11096 | RGG138470 | RCABP01778 | RNAMR0061 | RBRBA02852 | RANOC0317 |
| CLIP3 | RHSPN249392 | | RCABP00809 | RNAMR0250 | RBRBA008851 | |
| CLIP4 | RHSPN03105 | RGG042354 | RCABP00685 | RNAMR0053 | RBRBA00756 | RANOC0030 |
| CLK2 | RHSPN01813 | RGG154248 | RCABP03993 | RNAMR0373 | RBRBA01565 | RANOC0285 |
| CLNK | RHSPN06763 | RGG065723 | RCABP00939 | RNAMR0136 | RBRBA013838 | |
| CLOCK | RHSPN06994 | RGG064157 | RCABP01378 | RNAMR0158 | RBRBA02844 | RANOC0156 |
| CLP1 | RHSPN16064 | RGG071770 | RCABP00772 | RNAMR0358 | RBRBA03850 | RANOC0003 |
| CLPS | RHSPN09566 | RGG155154 | RCABP03393 | RNAMR0128 | RBRBA03451 | RANOC0261 |
| CLRN2 | RHSPN06803 | RGG065483 | RCABP03025 | RNAMR0133 | RBRBA02977 | RANOC0116 |
| CLSTN2 | RHSPN06150 | RGG100780 | RCABP02689 | RNAMR001263 | | RANOC0091 |
| CLTC | RHSPN23224 | RGG140384 | RCABP01596 | RNAMR0364 | RBRBA00384 | RANOC0277 |
| CLTCL1 | RHSPN26947 | RGG125128 | RCABP037719 | | | RANOC0235 |
| CLU | RHSPN12146 | RGG053969 | RCABP028265 | | RBRBA01450 | RANOC0321 |
| CLYBL | RHSPN18982 | RGG018748 | RCABP04216 | RNAMR0247 | RBRBA00090 | RANOC0095 |
| CMA1 | RHSPN192946 | | not predicted | RNAMR0301 | RBRBA010125 | |
| CMAS | RHSPN17333 | RGG008697 | RCABP02799 | RNAMR0118 | RBRBA01835 | RANOC0146 |
| CMKLR1 | RHSPN18166 | RGG126842 | RCABP03443 | RNAMR0097 | RBRBA01854 | RANOC0235 |
| CMTM3 | RHSPN21795 | RGG111475 | RCABP03291 | RNAMR0092 | RBRBA02737 | RANOC0239 |
| CNBP | RHSPN06037 | RGG114535 | RCABP003542 | | RBRBA02095 | RANOC0048 |
| CNOT3 | RHSPN255929 | | RCABP02320 | RNAMR0328 | RBRBA03217 | RANOC0233 |
| CNR2 | RHSPN00438 | RGG150805 | not predicted | RNAMR0166 | RBRBA02585 | RANOC0282 |
| CNTRL | RHSPN13796 | RGG133212 | RCABP04019 | RNAMR0155 | RBRBA03117 | RANOC0326 |
| COCH | RHSPN19320 | RGG074881 | RCABP00057 | RNAMR0301 | RBRBA03351 | RANOC0054 |
| COL1A1 | RHSPN23122 | RGG158702 | RCABP02187 | RNAMR0257 | RBRBA01643 | RANOC0184 |
| COL2A1 | RHSPN175130 | | RCABP01761 | RNAMR0291 | RBRBA02874 | RANOC0288 |
| COL3A1 | RHSPN04336 | RGG085938 | RCABP03183 | RNAMR0383 | RBRBA01399 | RANOC0238 |
| COL4A3BP | RHSPN08215 | RGG172940 | RCABP03865 | RNAMR0411 | RBRBA00313 | RANOC0041 |
| COL5A1 | RHSPN14067 | RGG132930 | RCABP04030 | RNAMR0150 | RBRBA02144 | RANOC0189 |
| COL6A3 | RHSPN04812 | RGG086746 | RCABP02954 | RNAMR0344 | RBRBA00000 | RANOC0019 |
| COLEC12 | RHSPN23676 | RGG035834 | RCABP02264 | RNAMR0192 | RBRBA00786 | RANOC0121 |
| COQ7 | RHSPN21363 | RGG123058 | RCABP02965 | RNAMR0218 | RBRBA021761 | |
| COTL1 | RHSPN22025 | RGG111981 | RCABP00931 | RNAMR0088 | RBRBA00911 | RANOC0314 |
| COX18 | RHSPN07086 | RGG062320 | RCABP01910 | RNAMR0041 | RBRBA03437 | RANOC0184 |
| COX5B | RHSPN03669 | RGG148217 | RCABP020808 | | RBRBA00862 | RANOC0279 |
| COX6C | RHSPN12600 | RGG038574 | RCABP04160 | RNAMR0162 | RBRBA00674 | RANOC0264 |
| CPA2 | RHSPN11580 | RGG000602 | RCABP01234 | RNAMR0224 | RBRBA01753 | RANOC0312 |
| CPA3 | RHSPN061989 | | RCABP03516 | RNAMR0005 | RBRBA01032 | RANOC0083 |
| CPB2 | RHSPN18747 | RGG020106 | RCABP00218 | RNAMR0103 | RBRBA00753 | RANOC0005 |
| CR1 | RHSPN02442 | RGG156543 | RCABP022714 | | RBRBA039061 | |
| CR2 | RHSPN024415 | | RCABP02425 | RNAMR0283 | RBRBA039062 | |
| CREB1 | RHSPN04510 | RGG088136 | RCABP03800 | RNAMR0172 | RBRBA02454 | RANOC0254 |
| CREB3L1 | RHSPN15949 | RGG073173 | RCABP00708 | RNAMR0180 | RBRBA02000 | RANOC0011 |
| CREB3L2 | RHSPN11638 | RGG007320 | RCABP01440 | RNAMR0173 | RBRBA03535 | RANOC0140 |

| | | | | | | |
|---|---|---|---|---|---|---|
| CREBBP | RHSPN21206 | RGG123952 | RCABP01298 | RNAMR0308 | RBRBA02296 | RANOC02839 |
| CREBL2 | RHSPN17260 | RGG009673 | RCABP03221 | RNAMR01190 | RBRBA01472 | RANOC02937 |
| CRLF2 | RHSPN28555 | RGG017131 | RCABP00549 | RNAMR03338 | RBRBA012714 | |
| CRLS1 | RHSPN25881 | RGG043870 | RCABP01067 | RNAMR00690 | RBRBA02513 | RANOC00198 |
| CRTAM | RHSPN16914 | RGG151795 | RCABP03164 | RNAMR0370 | RBRBA00203 | RANOC01849 |
| CRTC1 | RHSPN24726 | RGG162257 | RCABP01401 | RNAMR0394 | RBRBA024061 | |
| CRTC3 | RHSPN209266 | | | RNAMR0189 | RBRBA030258 | |
| CRYAA | RHSPN26798 | RGG014950 | RCABP02065 | RNAMR00851 | RBRBA03982 | RANOC00870 |
| CRYGS | RHSPN06465 | RGG100058 | RCABP00473 | RNAMR0009 | RBRBA00361 | RANOC00838 |
| CSF1 | RHSPN01379 | RGG155726 | RCABP02553 | RNAMR0197 | RBRBA00578 | RANOC01269 |
| CSF1R | RHSPN08761 | RGG119473 | RCABP00453 | RNAMR02050 | RBRBA01957 | RANOC00644 |
| CSF3 | RHSPN22828 | RGG159495 | RCABP01956 | RNAMR0255 | RBRBA00823 | RANOC01784 |
| CSF3R | RHSPN00668 | RGG150040 | RCABP03958 | RNAMR01700 | RBRBA017043 | |
| CSMD3 | RHSPN12679 | RGG039125 | RCABP04147 | RNAMR0162 | RBRBA017688 | |
| CSNK1E | RHSPN27339 | RGG006108 | RCABP00230 | RNAMR0117 | RBRBA03703 | RANOC02751 |
| CTC1 | RHSPN22380 | RGG011164 | RCABP00574 | RNAMR0235 | RBRBA03067 | RANOC02777 |
| CTF1 | RHSPN215627 | | RCABP03815 | RNAMR0222 | RBRBA03355 | RANOC02080 |
| CTLA4 | RHSPN04482 | RGG088283 | RCABP03970 | RNAMR01720 | RBRBA02457 | RANOC03468 |
| CTNNA2 | RHSPN03495 | RGG067234 | RCABP00747 | RNAMR0107 | RBRBA03845 | RANOC01596 |
| CTNNA3 | RHSPN14709 | RGG079961 | RCABP02525 | RNAMR0122 | RBRBA02792 | RANOC02052 |
| CTNNB1 | RHSPN05275 | RGG030191 | RCABP03650 | RNAMR0012 | RBRBA00621 | RANOC02100 |
| CTNND2 | RHSPN07889 | RGG033680 | RCABP00160 | RNAMR01129 | RBRBA02039 | RANOC01228 |
| CTSG | RHSPN19295 | RGG160715 | RCABP02090 | RNAMR0301 | RBRBA01014 | RANOC03059 |
| CUEDC2 | RHSPN15147 | RGG082005 | RCABP00992 | RNAMR0325 | RBRBA029948 | |
| CUX1 | RHSPN11361 | RGG138772 | RCABP00971 | RNAMR006229 | | RANOC02965 |
| CX3CR1 | RHSPN05256 | RGG030292 | RCABP00134 | RNAMR0013 | RBRBA00624 | RANOC02474 |
| CXCL1 | RHSPN070947 | | RCABP03556 | RNAMR0000 | RBRBA025315 | |
| CXCL10 | RHSPN071212 | | RCABP04078 | RNAMR0141 | RBRBA001236 | |
| CXCL11 | RHSPN071216 | | RCABP04079 | RNAMR0142 | RBRBA00123 | RANOC01458 |
| CXCL12 | RHSPN14546 | RGG082457 | RCABP03948 | RNAMR04017 | RBRBA00507 | RANOC02401 |
| CXCL14 | RHSPN08579 | RGG120046 | RCABP01445 | RNAMR0023 | RBRBA01312 | RANOC02103 |
| CXCL16 | RHSPN222494 | | not predicted | RNAMR02367 | RBRBA035443 | |
| CXCL2 | RHSPN070975 | | not predicted | | | |
| CXCL5 | RHSPN070961 | | not predicted | RNAMR000031 | | |
| CXCL8 | RHSPN07093 | RGG062364 | RCABP02664 | RNAMR0000 | RBRBA02532 | RANOC03308 |
| CXCL9 | RHSPN071207 | | not predicted | RNAMR0141 | RBRBA001235 | |
| CXCR2 | RHSPN04598 | RGG090468 | RCABP03748 | RNAMR0306 | RBRBA03665 | RANOC00160 |
| CXCR3 | RHSPN280123 | | RCABP04300 | RNAMR03871 | RBRBA02718 | RANOC03161 |
| CXCR4 | RHSPN04020 | RGG091723 | RCABP00138 | RNAMR0022 | RBRBA00944 | RANOC00042 |
| CXCR5 | RHSPN16862 | RGG152487 | RCABP00894 | RNAMR022829 | | RANOC03247 |
| CYC1 | RHSPN129002 | | RCABP01362 | RNAMR00380 | RBRBA03340 | RANOC03113 |
| CYGB | RHSPN23479 | RGG135062 | RCABP02304 | RNAMR0268 | RBRBA03782 | RANOC00611 |
| CYP19A1 | RHSPN20455 | RGG107015 | RCABP010285 | | RBRBA01090 | RANOC02501 |
| CYP1B1 | RHSPN03165 | RGG045884 | RCABP00837 | RNAMR0054 | RBRBA011901 | |
| CYP27B1 | RHSPN178124 | | RCABP01753 | RNAMR0272 | RBRBA03097 | RANOC00433 |

| | | | | | | |
|---|---|---|---|---|---|---|
| CYP4X1 | RHSPN008919 | | not predicted | RNAMR03539 | RBRBA008759 | |
| CYSLTR2 | RHSPN18767 | RGG020273 | RCABP01636 | RNAMR01040 | RBRBA009922 | |
| DAB2IP | RHSPN13808 | RGG133262 | RCABP04028 | RNAMR01554 | RBRBA03116 | RANOC03354 |
| DAPK1 | RHSPN13476 | RGG175086 | RCABP00332 | RNAMR02854 | RBRBA01651 | RANOC00424 |
| DAPK3 | RHSPN24305 | RGG160881 | RCABP00001 | RNAMR03213 | RBRBA004121 | |
| DAXX | RHSPN09515 | RGG129761 | RCABP01904 | RNAMR01290 | RBRBA01617 | RANOC03303 |
| DCLRE1A | RHSPN15246 | RGG084289 | RCABP00713 | RNAMR00499 | RBRBA02494 | RANOC01993 |
| DCLRE1B | RHSPN014419 | | RCABP03941 | RNAMR01980 | RBRBA00566 | RANOC01171 |
| DCLRE1C | RHSPN14310 | RGG001358 | RCABP02946 | RNAMR02784 | RBRBA03076 | RANOC02911 |
| DCPS | RHSPN16983 | RGG151249 | RCABP04252 | RNAMR04183 | RBRBA02014 | RANOC02469 |
| DCTN1 | RHSPN03459 | RGG067684 | RCABP01868 | RNAMR01079 | RBRBA01820 | RANOC02930 |
| DDB1 | RHSPN16130 | RGG068161 | RCABP00827 | RNAMR04047 | RBRBA02195 | RANOC02140 |
| DDB2 | RHSPN15973 | RGG072971 | RCABP01349 | RNAMR01794 | RBRBA02005 | RANOC00019 |
| DDIT3 | RHSPN177967 | | RCABP04260 | RNAMR02731 | RBRBA03095 | RANOC02037 |
| DDX1 | RHSPN02953 | RGG053048 | RCABP01060 | RNAMR03600 | RBRBA03758 | RANOC00059 |
| DDX10 | RHSPN16736 | RGG021709 | RCABP01170 | RNAMR02261 | RBRBA03572 | RANOC01451 |
| DDX21 | RHSPN14735 | RGG080738 | RCABP02519 | RNAMR01227 | RBRBA02795 | RANOC02055 |
| DDX3X | RHSPN27764 | RGG015240 | RCABP00364 | RNAMR02758 | RBRBA03335 | RANOC00980 |
| DDX41 | RHSPN08982 | RGG118809 | RCABP02478 | RNAMR00233 | RBRBA01294 | RANOC00652 |
| DDX5 | RHSPN23304 | RGG135928 | RCABP03464 | RNAMR03631 | RBRBA00395 | RANOC02215 |
| DDX58 | RHSPN131306 | | RCABP01822 | RNAMR00444 | RBRBA005165 | |
| DDX6 | RHSPN16861 | RGG152472 | RCABP00894 | RNAMR02281 | RBRBA00143 | RANOC03247 |
| DDX60 | RHSPN07664 | RGG059003 | RCABP00330 | RNAMR01918 | RBRBA00671 | RANOC02059 |
| DEFA1 | RHSPN119332 | | not predicted | | | |
| DEFA1B | RHSPN119339 | | not predicted | | | |
| DEFA3 | RHSPN119346 | | not predicted | | | |
| DEFA4 | RHSPN119322 | | not predicted | | | |
| DEFA5 | RHSPN119353 | | not predicted | RNAMR03567 | RBRBA033211 | |
| DEFA6 | RHSPN119319 | | not predicted | | | |
| DEFB1 | RHSPN11931 | RGG054519 | not predicted | RNAMR035666 | | |
| DEFB103A | RHSPN119530 | | not predicted | RNAMR04149 | RBRBA033212 | |
| DEFB103B | RHSPN119411 | | not predicted | | | |
| DEFB104A | RHSPN119518 | | not predicted | RNAMR03567 | RBRBA033211 | |
| DEFB104B | RHSPN119429 | | not predicted | | | |
| DEFB105A | RHSPN119508 | | not predicted | RNAMR03567 | RBRBA033211 | |
| DEFB105B | RHSPN119436 | | not predicted | | | |
| DEFB106A | RHSPN119511 | | not predicted | RNAMR03567 | RBRBA033211 | |
| DEFB106B | RHSPN119432 | | not predicted | | | |
| DEFB107A | RHSPN119504 | | not predicted | RNAMR03567 | RBRBA033211 | |
| DEFB107B | RHSPN119439 | | not predicted | | | |
| DEFB108B | RHSPN164397 | | not predicted | | | |
| DEFB108P1 | RHSPN119539 | | not predicted | | | |
| DEFB108P2 | RHSPN119403 | | not predicted | | | |
| DEFB109P1 | | | | | | |
| DEFB109P1B | | | | | | |

| | | | | | | |
|---|---|---|---|---|---|---|
| DEFB110 | RHSPN097739 | | not predictec | RNAMR0356 | RBRBA033211 | |
| DEFB112 | RHSPN097742 | | not predictec | RNAMR0356 | RBRBA033211 | |
| DEFB113 | RHSPN097734 | | not predictec | RNAMR0356 | RBRBA033211 | |
| DEFB114 | RHSPN097731 | | not predicted | | | |
| DEFB115 | RHSPN260373 | | not predictec | RNAMR0356 | RBRBA033211 | |
| DEFB116 | RHSPN260380 | | not predictec | RNAMR0356 | RBRBA033211 | |
| DEFB118 | RHSPN260387 | | not predictec | RNAMR0356 | RBRBA033211 | |
| DEFB119 | RHSPN260398 | | not predictec | RNAMR0356 | RBRBA033211 | |
| DEFB121 | RHSPN260411 | | not predictec | RNAMR0356 | RBRBA033211 | |
| DEFB123 | RHSPN260417 | | not predictec | RNAMR0356 | RBRBA033211 | |
| DEFB124 | RHSPN260420 | | not predictec | RNAMR0356 | RBRBA033211 | |
| DEFB125 | RHSPN257689 | | not predictec | RNAMR0356 | RBRBA033211 | |
| DEFB126 | RHSPN257692 | | not predictec | RNAMR0356 | RBRBA033211 | |
| DEFB127 | RHSPN257695 | | not predictec | RNAMR0356 | RBRBA033211 | |
| DEFB128 | RHSPN257698 | | not predictec | RNAMR0356 | RBRBA033211 | |
| DEFB129 | RHSPN257701 | | not predictec | RNAMR0356 | RBRBA033211 | |
| DEFB130 | RHSPN120018 | | not predictec | RNAMR0356 | RBRBA033211 | |
| DEFB131 | RHSPN067534 | | not predictec | RNAMR0356 | RBRBA033211 | |
| DEFB132 | RHSPN257704 | | not predictec | RNAMR0356 | RBRBA033211 | |
| DEFB133 | RHSPN097728 | | not predictec | RNAMR0356 | RBRBA033211 | |
| DEFB134 | RHSPN119946 | | not predictec | RNAMR0356 | RBRBA033211 | |
| DEFB135 | RHSPN119943 | | not predictec | RNAMR0356 | RBRBA033211 | |
| DEFB136 | RHSPN119940 | | not predictec | RNAMR0356 | RBRBA033211 | |
| DEFB4A | RHSPN1195 | RGG054497 | not predictec | RNAMR0356 | RBRBA033211 | |
| DEFB4B | RHSPN119406 | | not predicted | | | |
| DEK | RHSPN0921 | RGG032257 | RCABP008775 | | RBRBA0230 | RANOC0122 |
| DGAT1 | RHSPN129093 | | RCABP0171 | RNAMR0038 | RBRBA0123 | RANOC0010 |
| DHX15 | RHSPN0683 | RGG065322 | RCABP0254 | RNAMR0133 | RBRBA0291 | RANOC0165 |
| DHX29 | RHSPN0806 | RGG172141 | RCABP0416 | RNAMR0146 | RBRBA0244 | RANOC0050 |
| DHX33 | RHSPN2228 | RGG138651 | RCABP01781 | RNAMR0234 | RBRBA0355 | RANOC0263 |
| DHX36 | RHSPN0624 | RGG104040 | RCABP0181 | RNAMR0005 | RBRBA0247 | RANOC0090 |
| DHX58 | RHSPN2289 | RGG159762 | RCABP0196 | RNAMR0254 | RBRBA02651 | RANOC0140 |
| DHX9 | RHSPN022140 | | RCABP0110 | RNAMR0289 | RBRBA0324 | RANOC0199 |
| DICER1 | RHSPN1986 | RGG076651 | RCABP0443 | RNAMR0031 | RBRBA0205 | RANOC0000 |
| DKC1 | RHSPN2853 | RGG055638 | RCABP0046 | RNAMR0412 | RBRBA0388 | RANOC0198 |
| DLG1 | RHSPN0657 | RGG101660 | RCABP0182 | RNAMR0008 | RBRBA023257 | |
| DLGAP5 | RHSPN1947 | RGG078473 | RCABP0428 | RNAMR0015 | RBRBA0339 | RANOC0248 |
| DLL1 | RHSPN1051 | RGG046951 | RCABP0171 | RNAMR0349 | RBRBA0018 | RANOC0008 |
| DMC1 | RHSPN2734 | RGG006082 | RCABP00231 | RNAMR0117 | RBRBA0363 | RANOC0141 |
| DNAJA3 | RHSPN2121 | RGG124008 | RCABP0130 | RNAMR030868 | | RANOC0345 |
| DNM2 | RHSPN244991 | | RCABP0118 | RNAMR0316 | RBRBA0046 | RANOC0043 |
| DNMT3A | RHSPN0301 | RGG053761 | RCABP0355 | RNAMR0052 | RBRBA0254 | RANOC0294 |
| DOCK1 | RHSPN1539 | RGG085257 | RCABP0309 | RNAMR0048 | RBRBA0353 | RANOC0084 |
| DOCK2 | RHSPN0888 | RGG117991 | RCABP0328 | RNAMR0025 | RBRBA0061 | RANOC0063 |
| DOCK8 | RHSPN1296 | RGG173171 | RCABP03111 | RNAMR0043 | RBRBA0248 | RANOC0054 |

| | | | | | | |
|---|---|---|---|---|---|---|
| DOT1L | RHSPN2425 | RGG161121 | RCABP01134 | RNAMR0319 | RBRBA00421 | RANOC0344 |
| DRD2 | RHSPN1678 | RGG152567 | RCABP00572 | RNAMR0227 | RBRBA0015 | RANOC0255 |
| DRD4 | RHSPN1549 | RGG068572 | RCABP00344 | RNAMR017793 | | RANOC0014 |
| DUSP10 | RHSPN0255 | RGG044186 | RCABP0229 | RNAMR0334 | RBRBA0142 | RANOC0008 |
| DUSP3 | RHSPN2294 | RGG158844 | RCABP0049 | RNAMR0253 | RBRBA0263 | RANOC0169 |
| DUT | RHSPN2041 | RGG107158 | RCABP00371 | RNAMR0015 | RBRBA0288 | RANOC0312 |
| DYNLRB1 | RHSPN2610 | RGG141720 | RCABP01231 | RNAMR0339 | RBRBA0161 | RANOC0138 |
| DZIP3 | RHSPN058258 | | not predicte | RNAMR0018 | RBRBA001685 | |
| EBF1 | RHSPN0884 | RGG119068 | RCABP0299 | RNAMR0025 | RBRBA01971 | RANOC0200 |
| EBF3 | RHSPN1540 | RGG085418 | RCABP0018 | RNAMR004826 | | RANOC0093 |
| EBI3 | RHSPN243136 | | RCABP00002 | RNAMR0318 | RBRBA0040 | RANOC0341 |
| ECHS1 | RHSPN1544 | RGG080365 | RCABP0262 | RNAMR0047 | RBRBA031816 | |
| EDA | RHSPN2798 | RGG055035 | RCABP0268 | RNAMR0378 | RBRBA000888 | |
| EDN1 | RHSPN0916 | RGG032580 | RCABP02347 | RNAMR0137 | RBRBA0301 | RANOC0202 |
| EEF1A1 | RHSPN0987 | RGG051558 | RCABP0388 | RNAMR0300 | RBRBA0075 | RANOC0253 |
| EEF1A2 | RHSPN2648 | RGG143050 | RCABP0392 | RNAMR0071 | RBRBA0396 | RANOC0118 |
| EEF2 | RHSPN2430 | RGG160873 | RCABP00001 | RNAMR0318 | RBRBA00411 | RANOC0001 |
| EFEMP1 | RHSPN0330 | RGG041145 | RCABP01202 | RNAMR0333 | RBRBA0382 | RANOC0009 |
| EGF | RHSPN0735 | RGG063212 | RCABP01567 | RNAMR0336 | RBRBA0222 | RANOC0159 |
| EGFR | RHSPN1096 | RGG031471 | RCABP0236 | RNAMR0059 | RBRBA0035 | RANOC0175 |
| EGLN2 | RHSPN250859 | | not predicte | RNAMR0402 | RBRBA0172 | RANOC0188 |
| EGR1 | RHSPN0861 | RGG120717 | RCABP01704 | RNAMR0079 | RBRBA03357 | RANOC0065 |
| EIF2AK1 | RHSPN1059 | RGG121093 | RCABP0204 | RNAMR0065 | RBRBA0314 | RANOC0283 |
| EIF2AK2 | RHSPN0315 | RGG045892 | RCABP0046 | RNAMR0054 | RBRBA01191 | RANOC0028 |
| EIF2AK3 | RHSPN0356 | RGG067198 | RCABP0151 | RNAMR0106 | RBRBA0032 | RANOC0168 |
| EIF2AK4 | RHSPN2027 | RGG074473 | RCABP03181 | RNAMR0276 | RBRBA0060 | RANOC0227 |
| EIF2S1 | RHSPN1959 | RGG074359 | RCABP0067 | RNAMR0034 | RBRBA0294 | RANOC0229 |
| EIF2S2 | RHSPN2609 | RGG141780 | RCABP01232 | RNAMR0340 | RBRBA0319 | RANOC0131 |
| EIF2S3 | RHSPN2770 | RGG015796 | RCABP0322 | RNAMR0382 | RBRBA00801 | RANOC0086 |
| EIF3E | RHSPN1266 | RGG039052 | RCABP0297 | RNAMR0163 | RBRBA0177 | RANOC0111 |
| EIF4A2 | RHSPN0647 | RGG102723 | RCABP0329 | RNAMR0009 | RBRBA0272 | RANOC0246 |
| EIF4E | RHSPN0726 | RGG063357 | RCABP0229 | RNAMR0142 | RBRBA0295 | RANOC0242 |
| EIF4E3 | RHSPN0568 | RGG116849 | RCABP0258 | RNAMR0202 | RBRBA03887 | RANOC0049 |
| EIF4EBP1 | RHSPN1221 | RGG147742 | RCABP03294 | RNAMR0214 | RBRBA0009 | RANOC0266 |
| EIF5A2 | RHSPN0634 | RGG103170 | RCABP0148 | RNAMR0006 | RBRBA04087 | RANOC0082 |
| ELF1 | RHSPN1869 | RGG019751 | RCABP0087 | RNAMR0103 | RBRBA0304 | RANOC0094 |
| ELF4 | RHSPN282965 | | RCABP0017 | RNAMR0261 | RBRBA022662 | |
| ELK4 | RHSPN0240 | RGG156267 | RCABP0227 | RNAMR0281 | RBRBA0391 | RANOC0211 |
| ELL | RHSPN2471 | RGG162310 | RCABP0140 | RNAMR0394 | RBRBA024082 | |
| ELL2 | RHSPN0834 | RGG176678 | RCABP0282 | RNAMR0078 | RBRBA0210 | RANOC0041 |
| ELMO1 | RHSPN1081 | RGG030698 | RCABP0164 | RNAMR0063 | RBRBA0162 | RANOC0168 |
| ELMO2 | RHSPN2628 | RGG143952 | RCABP02551 | RNAMR0241 | RBRBA008555 | |
| ELN | RHSPN1109 | RGG138248 | RCABP04247 | RNAMR0066 | RBRBA0285 | RANOC0323 |
| ELOVL2 | RHSPN0915 | RGG032639 | RCABP0177 | RNAMR0137 | RBRBA03017 | RANOC0202 |
| ELTD1 | RHSPN0115 | RGG096568 | RCABP0360 | RNAMR0074 | RBRBA0217 | RANOC0191 |

| | | | | | | |
|---|---|---|---|---|---|---|
| EMC2 | RHSPN12665 | RGG039058 | RCABP02979 | RNAMR01640 | RBRBA01770 | RANOC01114 |
| EME1 | RHSPN23127 | RGG137060 | RCABP01611 | RNAMR02577 | RBRBA01643 | RANOC00627 |
| EME2 | RHSPN21111 | RGG124236 | | RNAMR03973 | RBRBA014406 | |
| EML4 | RHSPN03196 | RGG044854 | RCABP00577 | RNAMR00543 | RBRBA01197 | RANOC02957 |
| EMP2 | RHSPN21264 | RGG123375 | RCABP01807 | RNAMR03093 | RBRBA02141 | RANOC02947 |
| ENDOU | RHSPN17503 | RGG166438 | RCABP04311 | RNAMR0292 | RBRBA02875 | RANOC02223 |
| ENDOV | RHSPN23571 | RGG136553 | RCABP02697 | RNAMR02698 | RBRBA03259 | RANOC00597 |
| ENO1 | RHSPN00188 | RGG145864 | RCABP00745 | RNAMR02974 | RBRBA024716 | |
| ENOX1 | RHSPN18718 | RGG019944 | RCABP01311 | RNAMR01034 | RBRBA00080 | RANOC00945 |
| ENPP1 | RHSPN10214 | RGG049039 | RCABP03021 | RNAMR01210 | RBRBA00446 | RANOC00253 |
| ENPP2 | RHSPN12702 | RGG039293 | RCABP00899 | RNAMR01630 | RBRBA03998 | RANOC02297 |
| EOMES | RHSPN05148 | RGG029523 | RCABP00206 | RNAMR02389 | RBRBA02964 | RANOC00240 |
| EP300 | RHSPN27391 | RGG005819 | RCABP00190 | RNAMR01163 | RBRBA02082 | RANOC01494 |
| EPB41L4B | RHSPN13690 | RGG034255 | RCABP03761 | RNAMR01424 | RBRBA00543 | RANOC01828 |
| EPHB6 | RHSPN117128 | | not predicted | not predicted | not predicted | not predicted |
| EPRS | RHSPN02540 | RGG044274 | RCABP00316 | RNAMR03347 | RBRBA03206 | RANOC00087 |
| EPS15 | RHSPN00931 | RGG097800 | RCABP00410 | RNAMR03533 | RBRBA02237 | RANOC01158 |
| EPS8 | RHSPN17290 | RGG008263 | RCABP04296 | RNAMR01192 | RBRBA01467 | RANOC02518 |
| EQTN | RHSPN131168 | | not predicted | | RBRBA014818 | |
| ERBB2 | RHSPN22812 | RGG159408 | RCABP02167 | RNAMR02557 | RBRBA008254 | |
| ERBB2IP | RHSPN08135 | RGG172522 | RCABP01251 | RNAMR02894 | RBRBA00300 | RANOC00509 |
| ERBB3 | RHSPN17738 | RGG166918 | RCABP01660 | RNAMR02743 | RBRBA02787 | RANOC01950 |
| ERBB4 | RHSPN04551 | RGG086431 | RCABP03188 | RNAMR01720 | RBRBA03508 | RANOC00184 |
| ERC1 | RHSPN17066 | RGG007838 | RCABP02657 | RNAMR00373 | RBRBA01354 | RANOC01604 |
| ERCC1 | RHSPN252334 | | RCABP02776 | RNAMR03924 | RBRBA01015 | RANOC03160 |
| ERCC2 | RHSPN252308 | | RCABP02773 | RNAMR03924 | RBRBA01015 | RANOC03486 |
| ERCC3 | RHSPN03929 | RGG090642 | RCABP01742 | RNAMR00203 | RBRBA03030 | RANOC02542 |
| ERCC4 | RHSPN21298 | RGG120886 | RCABP01122 | RNAMR02194 | RBRBA02171 | RANOC03007 |
| ERCC5 | RHSPN19004 | RGG018637 | RCABP00313 | RNAMR02472 | RBRBA00324 | RANOC01075 |
| ERCC6 | RHSPN14631 | RGG082440 | RCABP03947 | RNAMR04059 | RBRBA00071 | RANOC02404 |
| ERCC8 | RHSPN08107 | RGG172332 | RCABP03058 | RNAMR02898 | RBRBA00294 | RANOC00507 |
| EREG | RHSPN07102 | RGG061247 | RCABP03556 | RNAMR00124 | RBRBA02530 | RANOC01760 |
| ERG | RHSPN26725 | RGG014535 | RCABP01706 | RNAMR00847 | RBRBA00741 | RANOC00994 |
| ERGIC1 | RHSPN08914 | RGG118520 | RCABP03081 | RNAMR00248 | RBRBA03142 | RANOC00633 |
| ERN1 | RHSPN23296 | RGG135846 | RCABP03461 | RNAMR0347 | RBRBA00394 | RANOC02483 |
| ERN2 | RHSPN214297 | | | RNAMR02204 | RBRBA035143 | |
| ESR1 | RHSPN10361 | RGG047830 | RCABP00530 | RNAMR02294 | RBRBA03844 | RANOC02259 |
| ESRRG | RHSPN02525 | RGG044362 | RCABP01715 | RNAMR03340 | RBRBA03208 | RANOC00300 |
| ETNK1 | RHSPN17340 | RGG008619 | RCABP02801 | RNAMR01183 | RBRBA01832 | RANOC01407 |
| ETS1 | RHSPN16993 | RGG151311 | RCABP04433 | RNAMR02869 | RBRBA00866 | RANOC01947 |
| ETV1 | RHSPN10646 | RGG027992 | RCABP01520 | RNAMR0057 | RBRBA02217 | RANOC01737 |
| ETV4 | RHSPN22938 | RGG158857 | RCABP00495 | RNAMR02530 | RBRBA02639 | RANOC01797 |
| ETV5 | RHSPN06464 | RGG100462 | RCABP00475 | RNAMR00094 | RBRBA00361 | RANOC00838 |
| ETV6 | RHSPN17252 | RGG009592 | RCABP03221 | RNAMR01197 | RBRBA01473 | RANOC01537 |
| EWSR1 | RHSPN27159 | RGG128684 | RCABP03766 | RNAMR00948 | RBRBA01860 | RANOC02688 |

| | | | | | | |
|---|---|---|---|---|---|---|
| EXO1 | RHSPN02755 | RGG046282 | RCABP04350 | RNAMR01967 | RBRBA03528 | RANOC00079 |
| EXOSC3 | RHSPN13251 | RGG079716 | RCABP01723 | RNAMR04063 | RBRBA03726 | RANOC02138 |
| EXT1 | RHSPN12693 | RGG039206 | RCABP04256 | RNAMR01629 | RBRBA03997 | RANOC02290 |
| EXT2 | RHSPN15918 | RGG072496 | RCABP02089 | RNAMR01808 | RBRBA01994 | RANOC00019 |
| EZH2 | RHSPN11758 | RGG031606 | RCABP01341 | RNAMR02653 | RBRBA00942 | RANOC01824 |
| EZR | RHSPN10417 | RGG048186 | RCABP00244 | RNAMR0406 | RBRBA02286 | RANOC00077 |
| F10 | RHSPN19059 | RGG018277 | RCABP00956 | RNAMR02460 | RBRBA03806 | RANOC01070 |
| F11 | RHSPN07784 | RGG063707 | | RNAMR019147 | | |
| F2R | RHSPN08227 | RGG172831 | RCABP00067 | RNAMR03624 | RBRBA00315 | RANOC00413 |
| F2RL1 | RHSPN08228 | RGG172823 | RCABP00067 | RNAMR03624 | RBRBA00315 | RANOC00731 |
| F7 | RHSPN19058 | RGG018282 | RCABP00956 | RNAMR02460 | RBRBA03807 | RANOC01070 |
| FADD | RHSPN16425 | RGG071935 | RCABP00836 | RNAMR018257 | | RANOC02359 |
| FADS1 | RHSPN16152 | RGG071589 | RCABP02103 | RNAMR04030 | RBRBA02199 | RANOC02140 |
| FAM131B | RHSPN11723 | RGG010694 | RCABP03705 | RNAMR02657 | RBRBA00938 | RANOC02631 |
| FAM13A | RHSPN07234 | RGG060378 | RCABP00412 | RNAMR01410 | RBRBA00934 | RANOC01515 |
| FAM19A5 | RHSPN27504 | RGG002187 | RCABP03390 | RNAMR011474 | | RANOC01531 |
| FAM46C | RHSPN01482 | RGG011086 | RCABP00576 | RNAMR0199 | RBRBA005593 | |
| FAN1 | RHSPN20206 | RGG106245 | RCABP00794 | RNAMR01874 | RBRBA02043 | RANOC02704 |
| FANCA | RHSPN22123 | RGG112457 | RCABP04380 | RNAMR00873 | RBRBA00734 | RANOC03248 |
| FANCB | RHSPN27636 | RGG016407 | RCABP02583 | RNAMR04092 | RBRBA02157 | RANOC00975 |
| FANCC | RHSPN13566 | RGG175170 | RCABP00619 | RNAMR01218 | RBRBA02393 | RANOC00424 |
| FANCD2 | RHSPN04996 | RGG113870 | RCABP01636 | RNAMR00143 | RBRBA01908 | RANOC00650 |
| FANCE | RHSPN09558 | RGG157012 | RCABP01969 | RNAMR0128 | RBRBA03454 | RANOC01289 |
| FANCF | RHSPN15810 | RGG069038 | RCABP00723 | RNAMR04178 | RBRBA01817 | RANOC00130 |
| FANCG | RHSPN13184 | RGG171007 | RCABP02987 | RNAMR03998 | RBRBA03750 | RANOC03122 |
| FANCI | RHSPN20902 | RGG107722 | RCABP03539 | RNAMR01883 | RBRBA04079 | RANOC02731 |
| FANCL | RHSPN03315 | RGG041241 | RCABP02290 | RNAMR03332 | RBRBA03828 | RANOC00090 |
| FANCM | RHSPN19400 | RGG078975 | RCABP04222 | RNAMR0017 | RBRBA02799 | RANOC02308 |
| FAS | RHSPN14953 | RGG082529 | RCABP04278 | RNAMR02448 | RBRBA02682 | RANOC02202 |
| FASLG | RHSPN02122 | RGG093637 | RCABP00082 | RNAMR0195 | RBRBA01584 | RANOC02417 |
| FAU | RHSPN162797 | | RCABP03347 | RNAMR0185 | RBRBA01129 | RANOC03340 |
| FBXO11 | RHSPN03253 | RGG042435 | RCABP04436 | RNAMR0055 | RBRBA01204 | RANOC00098 |
| FBXO31 | RHSPN22056 | RGG112162 | RCABP02388 | RNAMR00880 | RBRBA01661 | RANOC03204 |
| FBXO9 | RHSPN09793 | RGG052253 | RCABP03709 | RNAMR03488 | RBRBA02973 | RANOC00220 |
| FBXW7 | RHSPN07581 | RGG059913 | RCABP04366 | RNAMR03520 | RBRBA02090 | RANOC01580 |
| FCER1A | RHSPN019345 | | not predicted | RNAMR02484 | RBRBA002753 | |
| FCER1G | RHSPN01994 | RGG154594 | RCABP03813 | RNAMR02474 | RBRBA032828 | |
| FCER2 | RHSPN244084 | | not predicted | RNAMR03144 | RBRBA01216 | RANOC00602 |
| FCGR1A | RHSPN016149 | | not predicted | RNAMR02022 | RBRBA015517 | |
| FCGR1B | RHSPN015175 | | not predicted | | | |
| FCGR2A | RHSPN020066 | | not predicted | RNAMR03122 | RBRBA032836 | |
| FCGR2B | RHSPN020119 | | | RNAMR03122 | RBRBA032839 | |
| FCGR3A | RHSPN020091 | | not predicted | RNAMR03122 | RBRBA032836 | |
| FCGR3B | RHSPN020108 | | not predicted | | | |
| FCN1 | RHSPN140718 | | RCABP03818 | RNAMR01510 | RBRBA040179 | |

| | | | | | | |
|---|---|---|---|---|---|---|
| FCN2 | RHSPN1407 | RGG130901 | RCABP032055 | | | RANOC0003 |
| FCN3 | RHSPN005137 | | not predicte | RNAMR0168 | RBRBA025733 | |
| FCRL4 | RHSPN0190 | RGG154440 | | RNAMR025234 | | |
| FCRL5 | RHSPN019037 | | not predicte | RNAMR025235 | | |
| FEN1 | RHSPN1615 | RGG068334 | RCABP0125 | RNAMR0403 | RBRBA0219 | RANOC0215 |
| FER | RHSPN0839 | RGG175699 | RCABP03582 | RNAMR0078 | RBRBA01424 | RANOC0189 |
| FEV | RHSPN0462 | RGG090312 | RCABP03181 | RNAMR0306 | RBRBA03671 | RANOC0003 |
| FGA | RHSPN0760 | RGG058571 | RCABP0106 | RNAMR0352 | RBRBA0063 | RANOC0145 |
| FGB | RHSPN0760 | RGG058567 | RCABP0106 | RNAMR0352 | RBRBA0063 | RANOC0145 |
| FGF19 | RHSPN1641 | RGG071905 | RCABP0307 | RNAMR0187 | RBRBA01152 | RANOC0002 |
| FGF21 | RHSPN253435 | | RCABP03532 | RNAMR0330 | RBRBA0316 | RANOC0256 |
| FGFR1 | RHSPN1223 | RGG147810 | RCABP00662 | RNAMR0214 | RBRBA000964 | |
| FGFR1OP | RHSPN1047 | RGG047195 | RCABP00132 | RNAMR0230 | RBRBA00192 | RANOC0027 |
| FGFR2 | RHSPN1531 | RGG084908 | RCABP00854 | RNAMR0049 | RBRBA03244 | RANOC0094 |
| FGFR3 | RHSPN0664 | RGG066487 | RCABP04061 | RNAMR013525 | | |
| FGFR4 | RHSPN0896 | RGG118852 | RCABP0181 | RNAMR0024 | RBRBA0129 | RANOC0047 |
| FH | RHSPN0274 | RGG034347 | RCABP03847 | RNAMR0195 | RBRBA0376 | RANOC0175 |
| FHIT | RHSPN0562 | RGG116263 | RCABP0131 | RNAMR0305 | RBRBA00094 | RANOC0068 |
| FIP1L1 | RHSPN0698 | RGG064241 | RCABP00541 | RNAMR0158 | RBRBA02842 | RANOC0144 |
| FLCN | RHSPN2247 | RGG121976 | RCABP03591 | RNAMR0347 | RBRBA01917 | RANOC0282 |
| FLI1 | RHSPN1699 | RGG151325 | RCABP0443 | RNAMR0286 | RBRBA03607 | RANOC0194 |
| FLOT1 | RHSPN094088 | | RCABP02884 | RNAMR0131 | RBRBA01001 | RANOC0297 |
| FLT3 | RHSPN1860 | RGG021194 | RCABP03827 | RNAMR0209 | RBRBA0285 | RANOC0102 |
| FLT4 | RHSPN0903 | RGG119491 | RCABP00454 | RNAMR0028 | RBRBA0122 | RANOC0064 |
| FOS | RHSPN1971 | RGG075573 | RCABP0312 | RNAMR0032 | RBRBA03694 | RANOC0025 |
| FOSB | RHSPN252346 | | RCABP02767 | RNAMR0392 | RBRBA0101 | RANOC0331 |
| FOSL1 | RHSPN163146 | | RCABP0333 | RNAMR0184 | RBRBA0113 | RANOC0327 |
| FOSL2 | RHSPN0308 | RGG045280 | RCABP02072 | RNAMR0053 | RBRBA02534 | RANOC0063 |
| FOXA1 | RHSPN1937 | RGG075282 | RCABP0204 | RNAMR0411 | RBRBA0122 | RANOC0010 |
| FOXA2 | RHSPN2598 | RGG041661 | RCABP0212 | RNAMR0070 | RBRBA0134 | RANOC0136 |
| FOXJ1 | RHSPN2346 | RGG135166 | RCABP0230 | RNAMR026830 | | RANOC0045 |
| FOXJ1L | | | | | | |
| FOXL2 | RHSPN0614 | RGG100736 | RCABP0307 | RNAMR0004 | RBRBA01034 | RANOC0105 |
| FOXO1 | RHSPN1869 | RGG020651 | RCABP02514 | RNAMR0102 | RBRBA00934 | RANOC0005 |
| FOXO1B | | | | | | |
| FOXO3 | RHSPN1005 | RGG050304 | RCABP0054 | RNAMR0215 | RBRBA0372 | RANOC0007 |
| FOXO4 | RHSPN2799 | RGG055803 | | RNAMR0386 | RBRBA0271 | RANOC0309 |
| FOXO6 | RHSPN007565 | | | RNAMR0378 | RBRBA013693 | |
| FOXP1 | RHSPN0568 | RGG116796 | RCABP0050 | RNAMR0202 | RBRBA0388 | RANOC0067 |
| FOXP3 | RHSPN278569 | | not predicte | RNAMR0380 | RBRBA011668 | |
| FRRS1L | RHSPN1368 | RGG034239 | RCABP00727 | RNAMR0145 | RBRBA0054 | RANOC0176 |
| FSHR | RHSPN0326 | RGG042165 | RCABP01447 | RNAMR0055 | RBRBA0120 | RANOC0009 |
| FSTL3 | RHSPN2419 | RGG161534 | RCABP0000 | RNAMR0320 | RBRBA014542 | |
| FUBP1 | RHSPN0114 | RGG096660 | RCABP0317 | RNAMR0075 | RBRBA0216 | RANOC0192 |
| FUS | RHSPN215772 | | RCABP01604 | RNAMR0223 | RBRBA0348 | RANOC0208 |

| | | | | | | |
|---|---|---|---|---|---|---|
| FYB | RHSPN0800 | RGG171724 | RCABP0052 | RNAMR0147 | RBRBA0224 | RANOC0050 |
| FYN | RHSPN1009 | RGG049933 | RCABP00487 | RNAMR0406 | RBRBA0371 | RANOC0007 |
| FZD5 | RHSPN0451 | RGG088110 | RCABP03804 | RNAMR0172 | RBRBA0245 | RANOC0254 |
| G0S2 | RHSPN0246 | RGG156634 | RCABP0242 | RNAMR028356 | | RANOC0116 |
| GABRR3 | RHSPN0575 | RGG013027 | RCABP0035 | RNAMR024032 | | RANOC0100 |
| GADD45A | RHSPN0109 | RGG098994 | RCABP0442 | RNAMR0075 | RBRBA0162 | RANOC0248 |
| GADD45B | RHSPN2426 | RGG161854 | RCABP00027 | RNAMR0318 | RBRBA004199 | |
| GADD45G | RHSPN1350 | RGG175390 | RCABP0134 | RNAMR0174 | RBRBA0361 | RANOC0284 |
| GAPDH | RHSPN1712 | RGG010403 | RCABP0386 | RNAMR0354 | RBRBA0239 | RANOC0261 |
| GAPT | RHSPN080952 | | not predicte | RNAMR014770 | | |
| GAS6 | RHSPN1908 | RGG018179 | RCABP00954 | RNAMR0245 | RBRBA0291 | RANOC0085 |
| GAS7 | RHSPN2241 | RGG134263 | RCABP0071 | RNAMR0236 | RBRBA0307 | RANOC0060 |
| GATA1 | RHSPN278341 | | RCABP0105 | RNAMR0380 | RBRBA0162 | RANOC0043 |
| GATA2 | RHSPN0602 | RGG115453 | RCABP03071 | RNAMR0412 | RBRBA0149 | RANOC0072 |
| GATA3 | RHSPN1425 | RGG001084 | RCABP0055 | RNAMR0206 | RBRBA03924 | RANOC0326 |
| GATA4 | RHSPN1198 | RGG054432 | RCABP0366 | RNAMR0145 | RBRBA0052 | RANOC0304 |
| GBA3 | | | | | | |
| GBE1 | RHSPN0571 | RGG013485 | RCABP01021 | RNAMR0366 | RBRBA0084 | RANOC0100 |
| GBF1 | RHSPN1514 | RGG081930 | RCABP0099 | RNAMR0324 | RBRBA0299 | RANOC0193 |
| GBX1 | RHSPN1181 | RGG024216 | RCABP0161 | RNAMR0266 | RBRBA032340 | |
| GCH1 | RHSPN1946 | RGG078507 | RCABP0428 | RNAMR001608 | | RANOC0047 |
| GCLC | RHSPN0979 | RGG052228 | RCABP03707 | RNAMR0348 | RBRBA0297 | RANOC0021 |
| GCLM | RHSPN0126 | RGG095645 | RCABP00524 | RNAMR0073 | RBRBA03091 | RANOC0114 |
| GCNT3 | RHSPN2053 | RGG106156 | RCABP0281 | RNAMR0218 | RBRBA0071 | RANOC0270 |
| GDPD1 | RHSPN2322 | RGG140339 | RCABP0159 | RNAMR0364 | RBRBA00384 | RANOC0330 |
| GEM | RHSPN1255 | RGG038253 | RCABP01801 | RNAMR0162 | RBRBA03114 | RANOC0111 |
| GEN1 | RHSPN0296 | RGG053172 | RCABP0060 | RNAMR0051 | RBRBA0375 | RANOC0021 |
| GFI1 | RHSPN0125 | RGG095805 | RCABP0052 | RNAMR0073 | RBRBA03087 | RANOC0114 |
| GH1 | RHSPN2329 | RGG158186 | RCABP0410 | RNAMR0347 | RBRBA0039 | RANOC0169 |
| GH2 | RHSPN232878 | | | | | |
| GHRH | RHSPN2617 | RGG142163 | RCABP04011 | RNAMR0415 | RBRBA0168 | RANOC0238 |
| GHRHR | RHSPN1077 | RGG024669 | RCABP0376 | RNAMR0223 | RBRBA0095 | RANOC0170 |
| GHRL | RHSPN0500 | RGG117190 | RCABP0153 | RNAMR0014 | RBRBA01907 | RANOC0230 |
| GHSR | RHSPN0635 | RGG103116 | RCABP01481 | RNAMR0007 | RBRBA00684 | RANOC0082 |
| GJD2 | RHSPN2024 | RGG074736 | RCABP0070 | RNAMR0275 | RBRBA03987 | RANOC0001 |
| GLIS1 | RHSPN0097 | RGG098052 | RCABP0106 | RNAMR0407 | RBRBA0304 | RANOC0137 |
| GLL13L | | | | | | |
| GLL14L | | | | | | |
| GLUD1 | RHSPN1492 | RGG079534 | RCABP0121 | RNAMR0337 | RBRBA0205 | RANOC0195 |
| GMPS | RHSPN0625 | RGG103957 | RCABP0181 | RNAMR0006 | RBRBA02477 | RANOC0089 |
| GNA11 | RHSPN2428 | RGG160707 | RCABP00011 | RNAMR0319 | RBRBA004144 | |
| GNAQ | RHSPN1341 | RGG174638 | RCABP0209 | RNAMR0286 | RBRBA0165 | RANOC0042 |
| GNAS | RHSPN2641 | RGG144044 | RCABP01381 | RNAMR0070 | RBRBA03324 | RANOC0278 |
| GNLY | RHSPN035323 | | not predicted | | RBRBA039716 | |
| GOLGA5 | RHSPN1983 | RGG076465 | RCABP0009 | RNAMR0031 | RBRBA0293 | RANOC0031 |

| | | | | | | |
|---|---|---|---|---|---|---|
| GOPC | RHSPN1013 | RGG049635 | RCABP0197 | RNAMR0262 | RBRBA00431 | RANOC0006 |
| GPC3 | RHSPN2832 | RGG056183 | RCABP0116 | RNAMR0261 | RBRBA022757 | |
| GPC6 | RHSPN1893 | RGG019051 | RCABP0168 | RNAMR0368 | RBRBA0352 | RANOC0085 |
| GPHN | RHSPN1958 | RGG074441 | RCABP0079 | RNAMR0034 | RBRBA0294 | RANOC0230 |
| GPLD1 | RHSPN0923 | RGG032072 | RCABP0290 | RNAMR0137 | RBRBA00981 | RANOC0134 |
| GPR183 | RHSPN1897 | RGG018776 | RCABP0421 | RNAMR0247 | RBRBA0369 | RANOC0085 |
| GPR65 | RHSPN1978 | RGG076113 | RCABP0063 | RNAMR0032 | RBRBA0205 | RANOC0021 |
| GPRC5B | RHSPN2137 | RGG124847 | RCABP02967 | RNAMR021910 | | RANOC0321 |
| GPX1 | RHSPN0543 | RGG113882 | RCABP03867 | RNAMR000136 | | RANOC0065 |
| GPX2 | RHSPN1957 | RGG068406 | RCABP0077 | RNAMR0034 | RBRBA0237 | RANOC0017 |
| GPX4 | RHSPN2421 | RGG161664 | RCABP0001 | RNAMR0320 | RBRBA0145 | RANOC0347 |
| GPX5 | RHSPN093398 | | | | RBRBA040638 | |
| GPX6 | RHSPN093394 | | | RNAMR013988 | | |
| GPX7 | RHSPN0095 | RGG097921 | RCABP0411 | RNAMR0353 | RBRBA0222 | RANOC0116 |
| GPX8 | RHSPN0806 | RGG172114 | RCABP0416 | RNAMR0146 | RBRBA0244 | RANOC0040 |
| GRAMD1B | RHSPN1692 | RGG151749 | RCABP0316 | RNAMR0369 | RBRBA0106 | RANOC0184 |
| GRIA1 | RHSPN0880 | RGG119291 | RCABP0044 | RNAMR0026 | RBRBA0196 | RANOC0221 |
| GRIN2A | RHSPN2126 | RGG123383 | RCABP00251 | RNAMR0309 | RBRBA021416 | |
| GRIN2B | RHSPN1727 | RGG005428 | RCABP0271 | RNAMR0119 | RBRBA0147 | RANOC0264 |
| GRN | RHSPN229767 | | RCABP02591 | RNAMR0253 | RBRBA0107 | RANOC0177 |
| GSG1L | RHSPN2146 | RGG123634 | RCABP0247 | RNAMR0221 | RBRBA01941 | RANOC0315 |
| GSK3A | RHSPN251348 | | RCABP00787 | RNAMR0391 | RBRBA037342 | |
| GSR | RHSPN1218 | RGG060254 | RCABP0264 | RNAMR0215 | RBRBA0110 | RANOC0225 |
| GSTA1 | RHSPN0978 | RGG052303 | RCABP03707 | RNAMR035101 | | RANOC0006 |
| GSTA2 | RHSPN0978 | RGG052343 | RCABP022065 | | | RANOC0006 |
| GSTA3 | RHSPN097897 | | RCABP037071 | | RBRBA029434 | |
| GSTA4 | RHSPN097908 | | not predicte | RNAMR0418 | RBRBA0268 | RANOC0006 |
| GSTP1 | RHSPN163754 | | RCABP00782 | RNAMR0183 | RBRBA0114 | RANOC0004 |
| GSTT1 | RHSPN2874 | RGG127475 | RCABP03067 | RNAMR0096 | RBRBA0405 | RANOC0332 |
| GSTZ1 | RHSPN1974 | RGG075801 | RCABP03121 | RNAMR0032 | RBRBA0059 | RANOC0000 |
| GTF2H1 | RHSPN1577 | RGG070530 | RCABP0384 | RNAMR0176 | RBRBA0365 | RANOC0002 |
| GTF2H2 | RHSPN0818 | RGG178177 | RCABP0421 | RNAMR0389 | RBRBA0030 | RANOC0041 |
| GTF2H3 | RHSPN1841 | RGG126019 | RCABP0301 | RNAMR0100 | RBRBA0405 | RANOC0185 |
| GTF2H4 | RHSPN0941 | RGG130510 | RCABP01857 | RNAMR013158 | | RANOC0049 |
| GTF2H5 | RHSPN1040 | RGG048138 | RCABP002453 | | RBRBA0228 | RANOC0026 |
| GTPBP10 | RHSPN1119 | RGG027440 | RCABP02782 | RNAMR0284 | RBRBA0270 | RANOC0166 |
| GULOP | RHSPN121457 | | | | | |
| GZMA | RHSPN0806 | RGG172107 | RCABP0416 | RNAMR0146 | RBRBA0244 | RANOC0040 |
| GZMB | RHSPN192965 | | RCABP0204 | RNAMR0302 | RBRBA002059 | |
| GZMH | RHSPN192960 | | RCABP020902 | | | |
| GZMK | RHSPN080635 | | not predicte | RNAMR0147 | RBRBA024455 | |
| GZMM | RHSPN2419 | RGG161773 | RCABP000242 | | RBRBA014535 | |
| H2AFX | RHSPN1687 | RGG151022 | RCABP0080 | RNAMR022855 | | |
| H3F3A | RHSPN2345 | RGG043931 | RCABP0160 | RNAMR0267 | RBRBA0339 | RANOC0009 |
| H3F3B | RHSPN2345 | RGG043931 | RCABP0160 | RNAMR0267 | RBRBA0339 | RANOC0009 |

| | | | | | | |
|---|---|---|---|---|---|---|
| H6PD | RHSPN0019 | RCABP00743 | RCABP00743 | RNAMR0297 | RBRBA02351 | RANOC0304 |
| HAMP | RHSPN248980 | | RCABP00837 | RNAMR0249 | RBRBA008936 | |
| HAS2 | RHSPN1270 | RGG039379 | RCABP00901 | RNAMR0041 | RBRBA0400 | RANOC0229 |
| HAVCR2 | RHSPN088267 | | RCABP02992 | RNAMR0028 | RBRBA019691 | |
| HBEGF | RHSPN0863 | RGG117371 | RCABP03108 | RNAMR0079 | RBRBA03198 | RANOC0124 |
| HCK | RHSPN2606 | RGG143573 | RCABP03914 | RNAMR0066 | RBRBA03201 | RANOC0283 |
| HDAC1 | RHSPN0059 | RGG150469 | RCABP01855 | RNAMR0169 | RBRBA03008 | RANOC0304 |
| HDAC10 | RHSPN2752 | RGG002369 | RCABP04195 | RNAMR0114 | RBRBA02957 | RANOC0165 |
| HDAC11 | RHSPN0504 | RGG114738 | RCABP00383 | RNAMR0260 | RBRBA04079 | RANOC0068 |
| HDAC2 | RHSPN1010 | RGG049816 | RCABP00119 | RNAMR0351 | RBRBA03711 | RANOC0006 |
| HDAC3 | RHSPN0868 | RGG117678 | RCABP00480 | RNAMR0203 | RBRBA03372 | RANOC0114 |
| HDAC4 | RHSPN0484 | RGG087031 | RCABP04361 | RNAMR0344 | RBRBA00269 | RANOC0019 |
| HDAC5 | RHSPN229587 | | RCABP02188 | RNAMR0253 | RBRBA01076 | RANOC0179 |
| HDAC6 | RHSPN278348 | | RCABP01050 | RNAMR0380 | RBRBA02960 | RANOC0072 |
| HDAC7 | RHSPN1750 | RGG166418 | RCABP01512 | RNAMR0292 | RBRBA02875 | RANOC0222 |
| HDAC8 | RHSPN2802 | RGG055460 | RCABP02196 | RNAMR0387 | RBRBA02522 | RANOC0303 |
| HDAC9 | RHSPN1066 | RGG028210 | RCABP03379 | RNAMR0056 | RBRBA02213 | RANOC0181 |
| HECW1 | RHSPN1085 | RGG031382 | RCABP02361 | RNAMR0063 | RBRBA01634 | RANOC0175 |
| HECW2 | RHSPN0438 | RGG087619 | RCABP03276 | RNAMR0350 | RBRBA02369 | RANOC0233 |
| HELLS | RHSPN1500 | RGG081856 | RCABP00236 | RNAMR0046 | RBRBA037577 | |
| HELQ | RHSPN0718 | RGG061782 | RCABP02988 | RNAMR0383 | RBRBA00220 | RANOC0151 |
| HERPUD1 | RHSPN2171 | RGG109551 | RCABP00615 | RNAMR0092 | RBRBA023685 | |
| HEY1 | RHSPN1247 | RGG037696 | RCABP03648 | RNAMR0081 | RBRBA02986 | RANOC0112 |
| HFE | RHSPN092630 | | not predicte | RNAMR0136 | RBRBA009732 | |
| HIF1A | RHSPN1954 | RGG077938 | RCABP01979 | RNAMR0035 | RBRBA02432 | RANOC0026 |
| HIP1 | RHSPN1111 | RGG138050 | RCABP02023 | RNAMR0066 | RBRBA028483 | |
| HIST1H3B | RHSPN0925 | RGG005366 | RCABP01572 | RNAMR0136 | RBRBA01551 | RANOC0141 |
| HIST1H4I | RHSPN0925 | RGG005321 | RCABP01046 | RNAMR0119 | RBRBA00408 | RANOC0222 |
| HLA-A | RHSPN0937 | RGG130274 | | RNAMR0131 | RBRBA03659 | RANOC0325 |
| HLA-DQA1 | RHSPN094892 | | not predicte | RNAMR0129 | RBRBA016353 | |
| HLA-DQB1 | RHSPN094897 | | not predicte | RNAMR0129 | RBRBA031865 | |
| HLA-DRB1 | RHSPN0948 | RGG130225 | RCABP00701 | RNAMR012969 | | |
| HLF | RHSPN2316 | RGG135572 | RCABP03463 | RNAMR0258 | RBRBA02242 | RANOC0218 |
| HLTF | RHSPN062010 | | RCABP03514 | RNAMR0005 | RBRBA01031 | RANOC0090 |
| HMOX1 | RHSPN2726 | RGG006634 | RCABP00690 | RNAMR0108 | RBRBA03867 | RANOC0160 |
| HNF1A | RHSPN1834 | RGG127817 | RCABP00583 | RNAMR0097 | RBRBA010679 | |
| HNRNPA2B1 | RHSPN1072 | RGG028823 | RCABP00595 | RNAMR0056 | RBRBA01725 | RANOC0181 |
| HNRNPD | RHSPN0716 | RGG061750 | RCABP02666 | RNAMR0140 | RBRBA00224 | RANOC0151 |
| HOOK3 | RHSPN1228 | RGG176241 | RCABP01178 | RNAMR0213 | RBRBA00103 | RANOC0252 |
| HOXA1 | RHSPN1073 | RGG028883 | RCABP00594 | RNAMR0056 | RBRBA00964 | RANOC0230 |
| HOXA10 | RHSPN1073 | RGG028966 | RCABP00601 | RNAMR005612 | | RANOC0231 |
| HOXA11 | RHSPN1073 | RGG028972 | RCABP00601 | RNAMR005609 | | RANOC0231 |
| HOXA13 | RHSPN1074 | RGG028979 | RCABP00601 | RNAMR0059 | RBRBA009636 | |
| HOXA2 | RHSPN1073 | RGG028886 | RCABP00594 | RNAMR0056 | RBRBA00964 | RANOC0230 |
| HOXA3 | RHSPN1073 | RGG028904 | RCABP00593 | RNAMR0056 | RBRBA00964 | RANOC0230 |

| | | | | | | |
|---|---|---|---|---|---|---|
| HOXA4 | RHSPN10736 | RGG028918 | RCABP00593 | RNAMR005622 | | RANOC02309 |
| HOXA5 | RHSPN10737 | RGG028928 | RCABP00593 | RNAMR00562 | RBRBA00964 | RANOC02309 |
| HOXA6 | RHSPN10737 | RGG028932 | RCABP00594 | RNAMR00562 | RBRBA00964 | RANOC02311 |
| HOXA7 | RHSPN10737 | RGG028940 | RCABP00593 | RNAMR0056 | RBRBA00964 | RANOC02309 |
| HOXA9 | RHSPN10738 | RGG028948 | RCABP00601 | RNAMR0056 | RBRBA00964 | RANOC02309 |
| HOXB1 | RHSPN23077 | RGG159056 | RCABP02165 | RNAMR02569 | RBRBA00834 | RANOC01697 |
| HOXB13 | RHSPN23092 | RGG158919 | RCABP02165 | RNAMR02570 | RBRBA016155 | |
| HOXB2 | RHSPN23078 | RGG159049 | RCABP02165 | RNAMR025693 | | RANOC01697 |
| HOXB3 | RHSPN23081 | RGG159009 | RCABP02180 | RNAMR02569 | RBRBA01637 | RANOC01794 |
| HOXB4 | RHSPN23084 | RGG158977 | RCABP02180 | RNAMR02569 | RBRBA01637 | RANOC01697 |
| HOXB5 | RHSPN23085 | RGG158956 | RCABP02180 | RNAMR02570 | RBRBA01636 | RANOC01794 |
| HOXB6 | RHSPN23086 | RGG158953 | RCABP02180 | RNAMR02570 | RBRBA01637 | RANOC01695 |
| HOXB7 | RHSPN23087 | RGG158940 | RCABP02180 | RNAMR025705 | | RANOC01695 |
| HOXB8 | RHSPN23088 | RGG158937 | RCABP02180 | RNAMR033966 | | RANOC01695 |
| HOXB9 | RHSPN23088 | RGG158934 | RCABP02181 | RNAMR02570 | RBRBA01611 | RANOC01695 |
| HOXC10 | RHSPN17680 | RGG166581 | RCABP01752 | RNAMR03884 | RBRBA01779 | RANOC00749 |
| HOXC11 | RHSPN17679 | RGG166584 | RCABP01752 | RNAMR03884 | RBRBA01779 | RANOC00563 |
| HOXC12 | RHSPN17679 | RGG166590 | RCABP01752 | RNAMR03884 | RBRBA01779 | RANOC00432 |
| HOXC13 | RHSPN17678 | RGG166593 | RCABP01752 | RNAMR03884 | RBRBA01780 | RANOC00432 |
| HOXC4 | RHSPN176836 | | RCABP01756 | RNAMR03884 | RBRBA01778 | RANOC00432 |
| HOXC5 | RHSPN176829 | | RCABP01757 | RNAMR03884 | RBRBA01779 | RANOC00562 |
| HOXC6 | RHSPN176826 | | RCABP01756 | RNAMR03884 | RBRBA01779 | RANOC00432 |
| HOXC8 | RHSPN17681 | RGG166570 | RCABP01752 | RNAMR03884 | RBRBA01779 | RANOC00432 |
| HOXC9 | RHSPN17681 | RGG166575 | RCABP01752 | RNAMR03884 | RBRBA01779 | RANOC00434 |
| HOXD1 | RHSPN042642 | | RCABP01558 | RNAMR0238 | RBRBA002537 | |
| HOXD10 | RHSPN04258 | RGG088914 | RCABP01559 | RNAMR02380 | RBRBA00253 | RANOC02560 |
| HOXD11 | RHSPN04257 | RGG088922 | RCABP015593 | | | RANOC02560 |
| HOXD12 | RHSPN04257 | RGG088925 | RCABP01559 | RNAMR02380 | RBRBA00253 | RANOC02565 |
| HOXD13 | RHSPN04257 | RGG088928 | RCABP01559 | RNAMR02380 | RBRBA00253 | RANOC02560 |
| HOXD3 | RHSPN04261 | RGG088862 | RCABP01558 | RNAMR0238 | RBRBA00253 | RANOC02565 |
| HOXD4 | RHSPN04260 | RGG088881 | RCABP01558 | RNAMR0238 | RBRBA00253 | RANOC02565 |
| HOXD8 | RHSPN04259 | RGG088896 | RCABP01559 | RNAMR0238 | RBRBA00253 | RANOC02565 |
| HOXD9 | RHSPN04258 | RGG088902 | RCABP01559 | RNAMR023810 | | |
| HP | RHSPN219329 | | RCABP03317 | RNAMR00902 | RBRBA033802 | |
| HPCAL1 | RHSPN02923 | RGG052866 | RCABP00985 | RNAMR03604 | RBRBA01528 | RANOC00059 |
| HRAS | RHSPN15482 | RGG071259 | RCABP00559 | RNAMR0178 | RBRBA00475 | RANOC00149 |
| HRH1 | RHSPN05014 | RGG114312 | RCABP00533 | RNAMR04090 | RBRBA01906 | RANOC00668 |
| HRH4 | RHSPN238363 | | not predicted | | RBRBA039716 | |
| HS3ST3B1 | RHSPN22434 | RGG134639 | RCABP02295 | RNAMR03682 | RBRBA00215 | RANOC00454 |
| HSBP1 | RHSPN22007 | RGG111955 | RCABP00930 | RNAMR00884 | RBRBA009164 | |
| HSD17B10 | RHSPN27909 | RGG179880 | RCABP04267 | RNAMR03957 | RBRBA00439 | RANOC00583 |
| HSF1 | RHSPN12908 | RGG040677 | RCABP03269 | RNAMR00382 | RBRBA01233 | RANOC03085 |
| HSF2 | RHSPN10152 | RGG049465 | RCABP03557 | RNAMR02624 | RBRBA00434 | RANOC00235 |
| HSF4 | RHSPN21815 | RGG110278 | RCABP01668 | RNAMR00910 | RBRBA027404 | |
| HSF5 | RHSPN23203 | RGG138139 | RCABP04247 | RNAMR03640 | RBRBA00351 | RANOC03224 |

| | | | | | | |
|---|---|---|---|---|---|---|
| HSP90AA1 | RHSPN1995 | RGG077066 | RCABP0091 | RNAMR0030 | RBRBA02941 | RANOC0213 |
| HSP90AB1 | RHSPN0972 | RGG045532 | RCABP0431 | RNAMR0125 | RBRBA0069 | RANOC0008 |
| HSP90B1 | RHSPN1812 | RGG006951 | RCABP0136 | RNAMR0109 | RBRBA01261 | RANOC0140 |
| HSPA12A | RHSPN1527 | RGG084508 | RCABP0361 | RNAMR0049 | RBRBA0318 | RANOC0200 |
| HSPA12B | RHSPN258498 | | RCABP0356 | RNAMR0068 | RBRBA0251 | RANOC0307 |
| HSPA13 | RHSPN2657 | RGG013579 | RCABP0177 | RNAMR0083 | RBRBA0102 | RANOC0212 |
| HSPA14 | RHSPN1430 | RGG001350 | RCABP0294 | RNAMR0278 | RBRBA0307 | RANOC0291 |
| HSPA1B | RHSPN094610 | | | RNAMR0130 | RBRBA005519 | |
| HSPA1L | RHSPN094600 | | | RNAMR0130 | RBRBA005521 | |
| HSPA2 | RHSPN1957 | RGG077735 | RCABP0064 | RNAMR0035 | RBRBA0242 | RANOC0000 |
| HSPA4 | RHSPN0854 | RGG120479 | RCABP0417 | RNAMR0026 | RBRBA0130 | RANOC0046 |
| HSPA5 | RHSPN1385 | RGG133696 | RCABP0404 | RNAMR0154 | RBRBA03421 | RANOC0339 |
| HSPA6 | RHSPN020077 | | | RNAMR041845 | | |
| HSPA8 | RHSPN1691 | RGG151775 | RCABP03161 | RNAMR0369 | RBRBA0362 | RANOC0184 |
| HSPA9 | RHSPN0861 | RGG117865 | RCABP0413 | RNAMR0079 | RBRBA0335 | RANOC0005 |
| HSPB1 | RHSPN1112 | RGG138879 | RCABP0097 | RNAMR0062 | RBRBA031129 | |
| HSPB11 | RHSPN0098 | RGG098071 | RCABP0397 | RNAMR0407 | RBRBA0202 | RANOC0116 |
| HSPB2 | RHSPN1676 | RGG152684 | RCABP0187 | RNAMR0226 | RBRBA0015 | RANOC0286 |
| HSPB3 | RHSPN0806 | RGG172077 | RCABP0162 | RNAMR014624 | | RANOC0040 |
| HSPB6 | RHSPN249237 | | | RNAMR0250 | RBRBA008875 | |
| HSPB8 | RHSPN1830 | RGG128197 | RCABP0134 | RNAMR0097 | RBRBA0107 | RANOC0252 |
| HSPB9 | RHSPN228919 | | | RNAMR0259 | RBRBA02651 | RANOC0284 |
| HSPD1 | RHSPN0439 | RGG087688 | RCABP0327 | RNAMR0377 | RBRBA0237 | RANOC0271 |
| HSPH1 | RHSPN1862 | RGG021048 | RCABP0258 | RNAMR0208 | RBRBA0285 | RANOC0087 |
| HTRA2 | RHSPN0347 | RGG067853 | RCABP0205 | RNAMR0107 | RBRBA0182 | RANOC0316 |
| HTT | RHSPN0667 | RGG066190 | RCABP0124 | RNAMR0134 | RBRBA04001 | RANOC0153 |
| HUS1 | RHSPN1091 | RGG031702 | RCABP0266 | RNAMR0059 | RBRBA0359 | RANOC0167 |
| HYOU1 | RHSPN1687 | RGG152514 | RCABP01681 | RNAMR0228 | RBRBA00141 | RANOC0328 |
| ICAM1 | RHSPN244821 | | RCABP0233 | RNAMR0315 | RBRBA004690 | |
| ICOS | RHSPN0448 | RGG088278 | RCABP0396 | RNAMR0174 | RBRBA024571 | |
| IDH1 | RHSPN0452 | RGG088390 | RCABP0380 | RNAMR0172 | RBRBA0245 | RANOC0114 |
| IDH2 | RHSPN2092 | RGG108965 | RALGO16621 | RNAMR0188 | RBRBA0390 | RANOC0327 |
| IDO1 | RHSPN122503 | | RCABP0066 | RNAMR0214 | RBRBA000985 | |
| IFI16 | RHSPN019312 | | not predicted | | RBRBA039716 | |
| IFI27 | RHSPN1985 | RGG150219 | not predicted | | RBRBA039716 | |
| IFI30 | RHSPN2470 | RGG162376 | RCABP0140 | RNAMR0394 | RBRBA02411 | RANOC0337 |
| IFI35 | RHSPN2292 | RGG160025 | RCABP0049 | RNAMR0253 | RBRBA0264 | RANOC0177 |
| IFI6 | RHSPN0052 | RGG034703 | RCABP032188 | | RBRBA025717 | |
| IFIH1 | RHSPN0415 | RGG089948 | RCABP0327 | RNAMR0019 | RBRBA0049 | RANOC0257 |
| IFIT1 | RHSPN149605 | | not predicted | | RBRBA039716 | |
| IFIT2 | RHSPN149576 | | not predicted | | RBRBA039716 | |
| IFIT3 | RHSPN149580 | | not predicted | | RBRBA039716 | |
| IFIT5 | RHSPN1496 | RGG082538 | RCABP0428 | RNAMR0045 | RBRBA0268 | RANOC0272 |
| IFITM1 | RHSPN154700 | | RCABP020770 | | | |
| IFITM2 | RHSPN154697 | | not predicted | | | |

| | | | | | | |
|---|---|---|---|---|---|---|
| IFITM3 | RHSPN154704 | | not predicted | | RBRBA039716 | |
| IFNA1 | RHSPN130940 | | not predicted | | RBRBA039716 | |
| IFNA10 | RHSPN130902 | | not predicted | | | |
| IFNA13 | RHSPN130928 | | not predicted | | RBRBA039716 | |
| IFNA14 | RHSPN130913 | | not predicted | | | |
| IFNA16 | RHSPN130906 | | not predicted | | | |
| IFNA17 | RHSPN130909 | | not predicted | | | |
| IFNA2 | RHSPN130931 | | not predicted | | | |
| IFNA21 | RHSPN130891 | | not predicted | | | |
| IFNA4 | RHSPN130895 | | not predicted | | RBRBA039716 | |
| IFNA5 | RHSPN130918 | | not predicted | | | |
| IFNA6 | RHSPN130925 | | not predicted | | | |
| IFNA7 | RHSPN130899 | | not predicted | | RBRBA016480 | |
| IFNA8 | RHSPN130936 | | not predicted | | | |
| IFNAR1 | RHSPN2666 | RGG014126 | RCABP01971 | RNAMR0084 | RBRBA0066 | RANOC0298 |
| IFNAR2 | RHSPN2666 | RGG014107 | RCABP00581 | RNAMR0084 | RBRBA0066 | RANOC0298 |
| IFNB1 | RHSPN130885 | | RCABP0273 | RNAMR0044 | RBRBA000514 | |
| IFNE | RHSPN130946 | | RCABP027323 | | RBRBA038899 | |
| IFNG | RHSPN1786 | RGG003734 | RCABP0098 | RNAMR0271 | RBRBA001185 | |
| IFNGR1 | RHSPN1026 | RGG048579 | RCABP0043 | RNAMR0375 | RBRBA0045 | RANOC0196 |
| IFNGR2 | RHSPN2666 | RGG014132 | RCABP01971 | RNAMR0084 | RBRBA0066 | RANOC0099 |
| IFNK | RHSPN1311 | RGG174117 | RCABP0113 | RNAMR0045 | RBRBA0003 | RANOC0055 |
| IFNL1 | RHSPN250349 | | not predicted | | | |
| IFNL2 | RHSPN250345 | | not predicted | | | |
| IFNL3 | RHSPN2503 | RGG086581 | RCABP03431 | RNAMR0416 | RBRBA040567 | |
| IFNLR1 | RHSPN0044 | RGG150848 | RCABP02011 | RNAMR0171 | RBRBA0258 | RANOC0287 |
| IFNW1 | RHSPN130888 | | not predicted | | RBRBA040395 | |
| IGF1 | RHSPN1811 | RGG006991 | RCABP0285 | RNAMR0109 | RBRBA0331 | RANOC0148 |
| IGF1R | RHSPN2097 | RGG108077 | RCABP0142 | RNAMR0188 | RBRBA0234 | RANOC0261 |
| IGF2R | RHSPN1043 | RGG047400 | RCABP0233 | RNAMR0350 | RBRBA0259 | RANOC0026 |
| IGFBP3 | RHSPN109076 | | RCABP0266 | RNAMR006648 | | RANOC0167 |
| IGFBP4 | RHSPN2284 | RGG159553 | RCABP0196 | RNAMR0255 | RBRBA0081 | RANOC0179 |
| IK | RHSPN0864 | RGG117510 | RCABP0310 | RNAMR0080 | RBRBA0336 | RANOC0114 |
| IKBKAP | RHSPN1368 | RGG177861 | RCABP0171 | RNAMR0142 | RBRBA0054 | RANOC0225 |
| IKBKB | RHSPN1227 | RGG148006 | RCABP0428 | RNAMR0213 | RBRBA0010 | RANOC0289 |
| IKBKE | RHSPN0242 | RGG156434 | RCABP0227 | RNAMR0281 | RBRBA03911 | RANOC0212 |
| IKBKG | RHSPN2852 | RGG179954 | RCABP0096 | RNAMR0410 | RBRBA0402 | RANOC0058 |
| IKZF1 | RHSPN1093 | RGG033997 | RCABP0297 | RNAMR005967 | | RANOC0175 |
| IL10 | RHSPN0242 | RGG156466 | RCABP0228 | RNAMR0283 | RBRBA0243 | RANOC0211 |
| IL12A | RHSPN0629 | RGG103674 | RCABP0378 | RNAMR0006 | RBRBA011173 | |
| IL12B | RHSPN0884 | RGG118306 | RCABP0244 | RNAMR0025 | RBRBA0197 | RANOC0063 |
| IL12RB1 | RHSPN2470 | RGG162401 | RCABP0140 | RNAMR0418 | RBRBA0241 | RANOC0337 |
| IL12RB2 | RHSPN0108 | RGG098972 | RCABP0442 | RNAMR0076 | RBRBA0162 | RANOC0329 |
| IL13 | RHSPN085321 | | not predicte | RNAMR0027 | RBRBA013014 | |
| IL13RA1 | RHSPN2823 | RGG056094 | RCABP0083 | RNAMR0382 | RBRBA005047 | |

| | | | | | | |
|---|---|---|---|---|---|---|
| IL15 | RHSPN07512 | RGG059415 | RCABP04159 | RNAMR03490 | RBRBA03773 | RANOC01449 |
| IL16 | RHSPN20818 | RGG107542 | RCABP04423 | RNAMR01897 | RBRBA009466 | |
| IL17A | RHSPN09782 | RGG054555 | not predicted | RNAMR0351 | RBRBA03899 | RANOC00195 |
| IL17B | RHSPN08750 | RGG118416 | RCABP02447 | RNAMR02048 | RBRBA01955 | RANOC00632 |
| IL17C | RHSPN22080 | RGG112265 | RCABP03728 | RNAMR0093 | RBRBA00904 | RANOC03019 |
| IL17F | RHSPN09782 | RGG054552 | RCABP02653 | RNAMR03513 | RBRBA038989 | |
| IL17RD | RHSPN05584 | RGG115164 | RCABP02843 | RNAMR03039 | RBRBA01671 | RANOC0067 |
| IL17RE | RHSPN04987 | RGG115990 | RCABP02041 | RNAMR0014 | RBRBA019095 | |
| IL18 | RHSPN16774 | RGG152636 | RCABP01873 | RNAMR02267 | RBRBA00126 | RANOC03209 |
| IL18R1 | RHSPN03737 | RGG017792 | RCABP02405 | RNAMR01057 | RBRBA00404 | RANOC02185 |
| IL18RAP | RHSPN03738 | RGG017804 | RCABP02401 | RNAMR01088 | RBRBA00404 | RANOC02180 |
| IL19 | RHSPN024275 | | not predicted | RNAMR02814 | RBRBA024277 | |
| IL1A | RHSPN038424 | | not predicted | RNAMR01063 | RBRBA034543 | |
| IL1B | RHSPN03842 | RGG148452 | RCABP02083 | RNAMR01063 | RBRBA034537 | |
| IL1R1 | RHSPN03730 | RGG017757 | RCABP02404 | RNAMR01050 | RBRBA01756 | RANOC02187 |
| IL1R2 | RHSPN03727 | RGG017751 | RCABP02406 | RNAMR01050 | RBRBA01756 | RANOC02185 |
| IL1RAP | RHSPN06501 | RGG101992 | RCABP00910 | RNAMR00089 | RBRBA03322 | RANOC02620 |
| IL1RL1 | RHSPN03736 | RGG017781 | RCABP02405 | RNAMR01050 | RBRBA000550 | |
| IL1RL2 | RHSPN037336 | | RCABP02405 | RNAMR01088 | RBRBA01755 | RANOC02187 |
| IL2 | RHSPN074353 | | | RNAMR02103 | RBRBA012600 | |
| IL20 | RHSPN02428 | RGG156469 | RCABP02271 | RNAMR02830 | RBRBA039094 | |
| IL20RA | RHSPN10258 | RGG048594 | RCABP00434 | RNAMR03759 | RBRBA004551 | |
| IL20RB | RHSPN06119 | RGG123619 | RCABP02470 | RNAMR00123 | RBRBA01037 | RANOC02313 |
| IL21 | RHSPN07435 | RGG062610 | not predicted | RNAMR01063 | RBRBA034543 | |
| IL21R | RHSPN21457 | RGG122814 | RCABP01980 | RNAMR02230 | RBRBA01941 | RANOC02963 |
| IL22 | RHSPN17870 | RGG003740 | RCABP02639 | RNAMR02717 | RBRBA00118 | RANOC02248 |
| IL23A | RHSPN17751 | RGG166998 | not predicted | RNAMR02738 | RBRBA02790 | RANOC00593 |
| IL23R | RHSPN01085 | RGG098959 | RCABP04429 | RNAMR00758 | RBRBA01627 | RANOC0267 |
| IL24 | RHSPN024295 | | not predicted | RNAMR01063 | RBRBA034543 | |
| IL25 | RHSPN192416 | | not predicted | RNAMR03029 | RBRBA03501 | RANOC03029 |
| IL26 | RHSPN17870 | RGG003737 | RCABP00984 | RNAMR02747 | RBRBA001186 | |
| IL27 | RHSPN214718 | | not predicted | RNAMR02213 | RBRBA01932 | RANOC02375 |
| IL27RA | RHSPN245991 | | RCABP01184 | RNAMR03860 | RBRBA00205 | RANOC00569 |
| IL2RB | RHSPN27300 | RGG006381 | RCABP01077 | RNAMR01177 | RBRBA03640 | RANOC01490 |
| IL2RG | RHSPN27998 | RGG055792 | RCABP02487 | RNAMR04188 | RBRBA02715 | RANOC02951 |
| IL31 | RHSPN183683 | | not predicted | RNAMR01063 | RBRBA034543 | |
| IL31RA | RHSPN08078 | RGG172174 | RCABP04163 | RNAMR01477 | RBRBA02446 | RANOC0050 |
| IL32 | RHSPN211766 | | not predicted | RNAMR01063 | RBRBA034543 | |
| IL33 | RHSPN130155 | | not predicted | RNAMR01063 | RBRBA034543 | |
| IL34 | RHSPN21907 | RGG110073 | RCABP02534 | RNAMR00899 | RBRBA009272 | |
| IL36A | RHSPN038465 | | not predicted | RNAMR01063 | RBRBA034543 | |
| IL36B | RHSPN038469 | | not predicted | RNAMR01063 | RBRBA034543 | |
| IL36G | RHSPN038456 | | not predicted | RNAMR01063 | RBRBA034543 | |
| IL36RN | RHSPN038477 | | RCABP02078 | RNAMR01063 | RBRBA00500 | RANOC02170 |
| IL37 | RHSPN038442 | | not predicted | RNAMR01063 | RBRBA034543 | |

| | | | | | | |
|---|---|---|---|---|---|---|
| IL4 | RHSPN085325 | | not predicted | RNAMR01063 | RBRBA034543 | |
| IL4R | RHSPN21454 | RGG122805 | RCABP01981 | RNAMR02208 | RBRBA03490 | RANOC02963 |
| IL5 | RHSPN085310 | | not predicted | RNAMR01063 | RBRBA034543 | |
| IL5RA | RHSPN04934 | RGG116984 | RCABP01528 | RNAMR02032 | RBRBA034682 | |
| IL6 | RHSPN10693 | RGG028362 | RCABP02163 | RNAMR00592 | RBRBA01975 | RANOC01730 |
| IL6ST | RHSPN08080 | RGG172184 | RCABP04161 | RNAMR01477 | RBRBA02447 | RANOC00507 |
| IL7R | RHSPN07971 | RGG171539 | RCABP01144 | RNAMR01472 | RBRBA02249 | RANOC02182 |
| IL9 | RHSPN085848 | | not predicted | RNAMR01063 | RBRBA034543 | |
| ILF2 | RHSPN01737 | RGG153909 | RCABP02377 | RNAMR03749 | RBRBA01551 | RANOC03347 |
| ILF3 | RHSPN244964 | | RCABP01191 | RNAMR03160 | RBRBA00465 | RANOC00575 |
| INPP5D | RHSPN04773 | RGG099508 | RCABP03098 | RNAMR03410 | RBRBA03601 | RANOC00842 |
| INSR | RHSPN24391 | RGG162453 | RCABP01409 | RNAMR03139 | RBRBA012149 | |
| IPO7 | RHSPN15698 | RGG070056 | RCABP01882 | RNAMR01749 | RBRBA02422 | RANOC00024 |
| IRAK1 | RHSPN285005 | | RCABP00929 | RNAMR04104 | RBRBA04033 | RANOC00581 |
| IRAK1BP1 | RHSPN09894 | RGG051438 | RCABP02492 | RNAMR03005 | RBRBA00763 | RANOC00074 |
| IRAK2 | RHSPN04999 | RGG117184 | RCABP01533 | RNAMR00190 | RBRBA01908 | RANOC02303 |
| IRAK3 | RHSPN178603 | | not predicted | RNAMR01063 | RBRBA034543 | |
| IRAK4 | RHSPN17478 | RGG003263 | RCABP03353 | RNAMR02924 | RBRBA03444 | RANOC01428 |
| IRF1 | RHSPN08529 | RGG120373 | RCABP04173 | RNAMR00272 | RBRBA01303 | RANOC00630 |
| IRF2 | RHSPN07740 | RGG060899 | RCABP04191 | RNAMR01909 | RBRBA03729 | RANOC01643 |
| IRF2BP1 | RHSPN252522 | | RCABP02768 | RNAMR03320 | RBRBA01020 | RANOC03228 |
| IRF3 | RHSPN253963 | | RCABP04124 | RNAMR03294 | RBRBA03167 | RANOC01700 |
| IRF4 | RHSPN09066 | RGG033030 | RCABP02805 | RNAMR01389 | RBRBA01832 | RANOC02207 |
| IRF5 | RHSPN11562 | RGG000423 | RCABP01578 | RNAMR02240 | RBRBA017496 | |
| IRF6 | RHSPN02464 | RGG156704 | RCABP02425 | RNAMR02810 | RBRBA00372 | RANOC01167 |
| IRF7 | RHSPN15492 | RGG071176 | RCABP00561 | RNAMR017801 | | RANOC00148 |
| IRF8 | RHSPN22043 | RGG112071 | RCABP02391 | RNAMR00883 | RBRBA00909 | RANOC02743 |
| IRF9 | RHSPN192734 | | RCABP03374 | RNAMR03020 | RBRBA005208 | |
| IRG1 | RHSPN18886 | RGG019380 | RCABP02938 | RNAMR01133 | RBRBA02122 | RANOC00855 |
| IRGM | RHSPN087778 | | not predicted | RNAMR01063 | RBRBA034543 | |
| IRS1 | RHSPN04692 | RGG101046 | RCABP01950 | RNAMR03405 | RBRBA03681 | RANOC00833 |
| IRS2 | RHSPN19021 | RGG018360 | RCABP00382 | RNAMR02464 | RBRBA038104 | |
| ISG20 | RHSPN208933 | | RCABP03542 | RNAMR018841 | | |
| ITCH | RHSPN26104 | RGG141754 | RCABP01231 | RNAMR03399 | RBRBA03192 | RANOC01310 |
| ITGA1 | RHSPN08052 | RGG172018 | RCABP02853 | RNAMR01463 | RBRBA01824 | RANOC00728 |
| ITGA4 | RHSPN04301 | RGG088481 | RCABP01495 | RNAMR03424 | RBRBA00260 | RANOC02743 |
| ITGAL | RHSPN215472 | | RCABP01584 | RNAMR02220 | RBRBA03484 | RANOC02850 |
| ITGB1 | RHSPN14469 | RGG026219 | RCABP02867 | RNAMR02790 | RBRBA03953 | RANOC01712 |
| ITGB2 | RHSPN26837 | RGG087431 | RCABP01845 | RNAMR0086 | RBRBA03083 | RANOC02997 |
| ITGB3 | RHSPN23054 | RGG158253 | RCABP03448 | RNAMR0347 | RBRBA00387 | RANOC01840 |
| ITGB7 | RHSPN176532 | | RCABP01764 | RNAMR03878 | RBRBA01787 | RANOC00434 |
| ITK | RHSPN08827 | RGG119131 | RCABP02994 | RNAMR00259 | RBRBA01969 | RANOC01998 |
| ITM2B | RHSPN18762 | RGG020216 | RCABP01884 | RNAMR01029 | RBRBA00993 | RANOC00208 |
| JAG1 | RHSPN25902 | RGG043066 | RCABP02963 | RNAMR00693 | RBRBA02510 | RANOC00055 |
| JAK1 | RHSPN01057 | RGG098791 | RCABP01116 | RNAMR00762 | RBRBA01625 | RANOC01259 |

| | | | | | | |
|---|---|---|---|---|---|---|
| JAK2 | RHSPN13002 | RGG173456 | RCABP00099 | RNAMR0043 | RBRBA00743 | RANOC0042 |
| JAK3 | RHSPN24693 | RGG161888 | RCABP00028 | RNAMR0393 | RBRBA024131 | |
| JAM3 | RHSPN17032 | RGG151567 | RCABP03166 | RNAMR0287 | RBRBA03272 | RANOC0264 |
| JAZF1 | RHSPN10745 | RGG029010 | RCABP00592 | RNAMR0056 | RBRBA00952 | RANOC0231 |
| JUN | RHSPN0101 | RGG098378 | RCABP03232 | RNAMR0356 | RBRBA01873 | RANOC0126 |
| JUNB | RHSPN245639 | | RCABP04085 | RNAMR0384 | RBRBA034270 | |
| JUND | RHSPN2471 | RGG162347 | | | | |
| KAT6A | RHSPN14833 | RGG081507 | | RNAMR0124 | RBRBA01133 | RANOC0192 |
| KAT6B | RHSPN14833 | RGG081507 | | RNAMR0124 | RBRBA01133 | RANOC0192 |
| KCNJ5 | RHSPN1700 | RGG151344 | RCABP04433 | RNAMR0287 | RBRBA03606 | RANOC0194 |
| KCNK18 | RHSPN1528 | RGG084542 | RCABP01220 | RNAMR0049 | RBRBA03185 | RANOC0200 |
| KCNMA1 | RHSPN14849 | RGG081396 | RCABP04054 | RNAMR0124 | RBRBA01180 | RANOC0194 |
| KCNQ4 | RHSPN0074 | RGG150399 | RCABP03192 | RNAMR0378 | RBRBA01367 | RANOC0243 |
| KCNT2 | RHSPN0227 | RGG093269 | RCABP02102 | RNAMR0262 | RBRBA03761 | RANOC0272 |
| KCTD1 | RHSPN23847 | RGG036213 | RCABP02863 | RNAMR0192 | RBRBA00180 | RANOC0121 |
| KDM5A | RHSPN17053 | RGG007747 | RCABP02655 | RNAMR0036 | RBRBA01357 | RANOC0140 |
| KDM5C | RHSPN279019 | | | RNAMR0395 | RBRBA01717 | RANOC0039 |
| KDM6A | RHSPN27778 | RGG015104 | RCABP03042 | RNAMR0376 | RBRBA021179 | |
| KDR | RHSPN0699 | RGG064193 | RCABP01378 | RNAMR0159 | RBRBA02844 | RANOC0155 |
| KDSR | RHSPN24086 | RGG033211 | RCABP02267 | RNAMR0245 | RBRBA03457 | RANOC0220 |
| KIAA1549 | RHSPN11645 | RGG007134 | RCABP01329 | RNAMR0174 | RBRBA02551 | RANOC0148 |
| KIF13B | RHSPN12168 | RGG054299 | | RNAMR0145 | RBRBA025074 | |
| KIF5B | RHSPN14458 | RGG026252 | RCABP02865 | RNAMR0278 | RBRBA02759 | RANOC0166 |
| KIR2DS1 | RHSPN256297 | | not predicted | | | |
| KIR2DS2 | | | | | | |
| KIR2DS4 | RHSPN256297 | | not predicted | | | |
| KIR2DS5 | RHSPN256273 | | not predicted | | | |
| KIR3DL2 | RHSPN256304 | | not predicted | | | |
| KIR3DS1 | | | | | | |
| KIT | RHSPN069897 | | RCABP00632 | RNAMR0159 | RBRBA02844 | RANOC0156 |
| KL | RHSPN18642 | RGG020947 | RCABP02629 | RNAMR0210 | RBRBA03625 | RANOC0323 |
| KLF4 | RHSPN13677 | RGG176479 | RCABP00405 | RNAMR0142 | RBRBA00544 | RANOC0207 |
| KLF6 | RHSPN14209 | RGG026038 | RCABP00392 | RNAMR0259 | RBRBA01828 | RANOC0166 |
| KLHL6 | RHSPN06417 | RGG099773 | RCABP03779 | RNAMR0010 | RBRBA02188 | RANOC0084 |
| KLKB1 | RHSPN077831 | | RCABP02875 | RNAMR0191 | RBRBA03859 | RANOC0144 |
| KLRB1 | RHSPN17209 | RGG130201 | not predicted | | | RANOC0272 |
| KLRC2 | RHSPN172356 | | not predicted | RNAMR0141 | RBRBA001235 | |
| KLRD1 | RHSPN17232 | RGG011005 | RCABP03061 | RNAMR0120 | RBRBA012549 | |
| KLRF1 | RHSPN172151 | | RCABP018597 | | RBRBA02079 | RANOC0272 |
| KLRF2 | RHSPN172161 | | RCABP018598 | | RBRBA020795 | |
| KLRG1 | RHSPN172017 | | RCABP01036 | RNAMR035612 | | |
| KLRG2 | RHSPN116535 | | not predicted | RNAMR0174 | RBRBA025537 | |
| KLRK1 | RHSPN172344 | | not predicted | | | |
| KMT2A | RHSPN16845 | RGG152068 | RCABP00892 | RNAMR0228 | RBRBA00145 | RANOC0309 |
| KMT2C | RHSPN1184 | RGG025631 | RCABP04052 | RNAMR0264 | RBRBA03229 | RANOC0180 |

| | | | | | | |
|---|---|---|---|---|---|---|
| KMT2D | RHSPN1754[illegible] | RGG166844 | RCABP0166[illegible] | RNAMR0291[illegible] | RBRBA0286[illegible] | RANOC0195[illegible] |
| KRAS | RHSPN1735[illegible] | RGG008809 | RCABP02031 | RNAMR0118[illegible] | RBRBA0214[illegible] | RANOC0326[illegible] |
| KTN1 | RHSPN1947[illegible] | RGG078382 | RCABP0428[illegible] | RNAMR0371[illegible] | RBRBA0339[illegible] | RANOC0274[illegible] |
| LAG3 | RHSPN1714[illegible] | RGG010546 | RCABP0032[illegible] | RNAMR0355 | RBRBA0238[illegible] | RANOC0274[illegible] |
| LAIR1 | RHSPN256098 | | not predicted | RNAMR0332[illegible] | RBRBA039117 | |
| LAMP1 | RHSPN1906[illegible] | RGG018254 | RCABP0095[illegible] | RNAMR0245[illegible] | RBRBA0380[illegible] | RANOC0085[illegible] |
| LAMTOR1 | RHSPN1645[illegible] | RGG023305 | RCABP0024[illegible] | RNAMR0293[illegible] | RBRBA0002[illegible] | RANOC0311[illegible] |
| LAMTOR2 | RHSPN0184[illegible] | RGG152884 | RCABP0224[illegible] | RNAMR0372[illegible] | RBRBA015714 | |
| LAMTOR4 | RHSPN113117 | | RCABP0396[illegible] | RNAMR0063[illegible] | RBRBA007209 | |
| LASP1 | RHSPN2279[illegible] | RGG159298 | RCABP02171 | RNAMR0256 | RBRBA0082[illegible] | RANOC0170[illegible] |
| LAT | RHSPN215011 | | not predicted | RNAMR0221 | RBRBA01941 | RANOC0237[illegible] |
| LAT2 | RHSPN1109[illegible] | RGG138531 | RCABP0177[illegible] | RNAMR0061[illegible] | RBRBA028515 | |
| LATS1 | RHSPN1033[illegible] | RGG047725 | RCABP0441[illegible] | RNAMR0229[illegible] | RBRBA03841 | RANOC0226[illegible] |
| LATS2 | RHSPN1853[illegible] | RGG021550 | RCABP0091[illegible] | RNAMR0209[illegible] | RBRBA03221 | RANOC0088[illegible] |
| LBP | RHSPN261861 | | | RNAMR0338[illegible] | RBRBA003374 | |
| LCK | RHSPN0059[illegible] | RGG150478 | RCABP01854 | RNAMR0169[illegible] | RBRBA030080 | |
| LCP1 | RHSPN1874[illegible] | RGG020116 | RCABP0021[illegible] | RNAMR0103[illegible] | RBRBA007528 | |
| LCP2 | RHSPN0889[illegible] | RGG117981 | RCABP03282 | RNAMR0025[illegible] | RBRBA0061[illegible] | RANOC0063[illegible] |
| LEP | RHSPN115374 | | RCABP00252 | RNAMR0224[illegible] | RBRBA0174[illegible] | RANOC0311[illegible] |
| LEPR | RHSPN0106[illegible] | RGG098854 | RCABP0442[illegible] | RNAMR0076 | RBRBA0162[illegible] | RANOC0125[illegible] |
| LFNG | RHSPN1056[illegible] | RGG121613 | RCABP0241[illegible] | RNAMR006526 | | RANOC0253[illegible] |
| LGALS3 | RHSPN1947[illegible] | RGG078477 | RCABP0428[illegible] | RNAMR0015[illegible] | RBRBA0333[illegible] | RANOC0248[illegible] |
| LGR4 | RHSPN1582[illegible] | RGG069111 | RCABP0432[illegible] | RNAMR0176[illegible] | RBRBA0230[illegible] | RANOC0244[illegible] |
| LHFP | RHSPN1868[illegible] | RGG020666 | RCABP02514 | RNAMR0102 | RBRBA0093[illegible] | RANOC0005[illegible] |
| LIFR | RHSPN0799[illegible] | RGG171678 | RCABP0373[illegible] | RNAMR0146[illegible] | RBRBA0224[illegible] | RANOC0218[illegible] |
| LIG1 | RHSPN2531[illegible] | RGG163479 | RCABP02772 | RNAMR0331[illegible] | RBRBA03157 | RANOC0223[illegible] |
| LIG3 | RHSPN2269[illegible] | RGG139013 | RCABP02022 | RNAMR0367[illegible] | RBRBA005964 | |
| LIG4 | RHSPN1901[illegible] | RGG018570 | RCABP0243[illegible] | RNAMR0246[illegible] | RBRBA0381[illegible] | RANOC0005[illegible] |
| LILRA2 | RHSPN256178 | | not predicted | | | |
| LILRA4 | RHSPN256083 | | not predicted | | | |
| LILRA5 | RHSPN256075 | | not predicted | | RBRBA004633 | |
| LINGO2 | RHSPN1312[illegible] | RGG177807 | RCABP0203[illegible] | RNAMR0044[illegible] | RBRBA0263[illegible] | RANOC0074[illegible] |
| LMNA | RHSPN018493 | | RCABP0224[illegible] | RNAMR0372[illegible] | RBRBA01571 | RANOC0329[illegible] |
| LMNB1 | RHSPN0849[illegible] | RGG176540 | RCABP0281[illegible] | RNAMR0205[illegible] | RBRBA0266[illegible] | RANOC0041[illegible] |
| LMNB2 | RHSPN2426[illegible] | RGG160330 | RCABP0009[illegible] | RNAMR0321[illegible] | RBRBA004200 | |
| LMNTD1 | RHSPN173620 | | RCABP0400[illegible] | RNAMR0118[illegible] | RBRBA0214[illegible] | RANOC0255[illegible] |
| LMO1 | RHSPN1567[illegible] | RGG070202 | RCABP0162[illegible] | RNAMR0174[illegible] | RBRBA0241[illegible] | RANOC0238[illegible] |
| LMO2 | RHSPN1587[illegible] | RGG072152 | RCABP0431[illegible] | RNAMR0181[illegible] | RBRBA0272[illegible] | RANOC0235[illegible] |
| LMO4 | RHSPN0120[illegible] | RGG096151 | RCABP0336[illegible] | RNAMR0074[illegible] | RBRBA0392[illegible] | RANOC0114[illegible] |
| LMX1B | RHSPN1387[illegible] | RGG133826 | RCABP0418[illegible] | RNAMR0154[illegible] | RBRBA034181 | |
| LPA | RHSPN104362 | | | | | |
| LPP | RHSPN0648[illegible] | RGG102091 | RCABP0090[illegible] | RNAMR0009[illegible] | RBRBA027262 | |
| LPXN | RHSPN1608[illegible] | RGG071687 | RCABP00774 | RNAMR0358 | RBRBA0181[illegible] | RANOC0016[illegible] |
| LRAS | | | | | | |
| LRIG3 | RHSPN1782[illegible] | RGG003430 | RCABP0026[illegible] | RNAMR0272[illegible] | RBRBA0310[illegible] | RANOC0142[illegible] |

| | | | | | | |
|---|---|---|---|---|---|---|
| LRP1B | RHSPN0402 | RGG091809 | RCABP03697 | RNAMR0022 | RBRBA0277 | RANOC0036 |
| LSM14A | RHSPN2486 | RGG111317 | RCABP03951 | RNAMR0249 | RBRBA0089 | RANOC0324 |
| LSP1 | RHSPN1553 | RGG070877 | RCABP0044 | RNAMR0181 | RBRBA0401 | RANOC0014 |
| LTB4R | RHSPN1928 | RGG130368 | RCABP0337 | RNAMR0301 | RBRBA0051 | RANOC0208 |
| LTBR | RHSPN171228 | | not predicted | | RBRBA004633 | |
| LY6G6F | RHSPN094516 | | not predicted | RNAMR0130 | RBRBA0055 | RANOC0294 |
| LY75 | RHSPN0413 | RGG090071 | RCABP01337 | RNAMR0019 | RBRBA0049 | RANOC0333 |
| LY86 | RHSPN0912 | RGG032803 | RCABP0052 | RNAMR0138 | RBRBA0028 | RANOC0243 |
| LY9 | RHSPN0197 | RGG153312 | RCABP022357 | | | |
| LY96 | RHSPN1245 | RGG037525 | RCABP0158 | RNAMR0081 | RBRBA0141 | RANOC0121 |
| LYL1 | RHSPN245786 | | RCABP0021 | RNAMR0384 | RBRBA0342 | RANOC0057 |
| LYN | RHSPN1234 | RGG036762 | RCABP0266 | RNAMR0375 | RBRBA0212 | RANOC0219 |
| LYPD3 | RHSPN251629 | | RCABP0271 | RNAMR0391 | RBRBA0407 | RANOC0188 |
| LYST | RHSPN0272 | RGG046541 | RCABP0272 | RNAMR0208 | RBRBA0340 | RANOC0040 |
| LYZ | RHSPN1788 | RGG003820 | RCABP0263 | RNAMR0410 | RBRBA0012 | RANOC0043 |
| MAD2L2 | RHSPN0023 | RGG146709 | RCABP0073 | RNAMR0296 | RBRBA023574 | |
| MADCAM1 | RHSPN2419 | RGG161488 | RCABP0001 | RNAMR0321 | RBRBA014531 | |
| MAF | RHSPN2198 | RGG111703 | RCABP04081 | RNAMR0089 | RBRBA0092 | RANOC0187 |
| MAFB | RHSPN2619 | RGG142082 | RCABP0401 | RNAMR0400 | RBRBA02071 | RANOC0238 |
| MALSU1 | RHSPN1070 | RGG028592 | RCABP0060 | RNAMR0056 | RBRBA0220 | RANOC0173 |
| MAML2 | RHSPN1666 | RGG022190 | RCABP03604 | RNAMR0041 | RBRBA0013 | RANOC0088 |
| MAP2K1 | RHSPN2061 | RGG108474 | RCABP03897 | RNAMR0216 | RBRBA008436 | |
| MAP2K2 | RHSPN2430 | RGG161458 | RCABP0000 | RNAMR0318 | RBRBA00411 | RANOC0341 |
| MAP2K3 | RHSPN2256 | RGG121877 | RCABP0141 | RNAMR0348 | RBRBA0191 | RANOC0262 |
| MAP2K4 | RHSPN2242 | RGG134352 | RCABP0034 | RNAMR0236 | RBRBA02607 | RANOC0060 |
| MAP2K6 | RHSPN2336 | RGG136223 | RCABP02697 | RNAMR0363 | RBRBA0081 | RANOC0044 |
| MAP2K7 | RHSPN244185 | | RCABP0128 | RNAMR0314 | RBRBA0198 | RANOC0046 |
| MAP3K1 | RHSPN0808 | RGG172205 | RCABP00731 | RNAMR0146 | RBRBA024473 | |
| MAP3K13 | RHSPN0645 | RGG099945 | RCABP0367 | RNAMR0009 | RBRBA0036 | RANOC0091 |
| MAP3K14 | RHSPN2301 | RGG158039 | RCABP0267 | RNAMR0252 | RBRBA0108 | RANOC0169 |
| MAP3K5 | RHSPN1025 | RGG048611 | RCABP0043 | RNAMR0375 | RBRBA0045 | RANOC0197 |
| MAP3K7 | RHSPN0997 | RGG050878 | RCABP00491 | RNAMR0262 | RBRBA02187 | RANOC0038 |
| MAP3K8 | RHSPN1444 | RGG026454 | RCABP0371 | RNAMR0278 | RBRBA0276 | RANOC0171 |
| MAP4K2 | RHSPN162557 | | RCABP0029 | RNAMR0185 | RBRBA011270 | |
| MAPK1 | RHSPN2702 | RGG125231 | RCABP0009 | RNAMR0094 | RBRBA0262 | RANOC0234 |
| MAPK10 | RHSPN0719 | RGG061531 | RCABP0296 | RNAMR0365 | RBRBA0243 | RANOC0151 |
| MAPK11 | RHSPN2752 | RGG002392 | RCABP0419 | RNAMR011429 | | RANOC0142 |
| MAPK14 | RHSPN0957 | RGG155195 | RCABP0338 | RNAMR0127 | RBRBA0345 | RANOC0261 |
| MAPK3 | RHSPN215382 | | | RNAMR0223 | RBRBA034778 | |
| MAPK8 | RHSPN1461 | RGG082357 | RCABP02717 | RNAMR0406 | RBRBA0074 | RANOC0240 |
| MAPK9 | RHSPN0903 | RGG119700 | RCABP0032 | RNAMR0028 | RBRBA0121 | RANOC0047 |
| MAPKAPK2 | RHSPN0242 | RGG156463 | RCABP0227 | RNAMR0283 | RBRBA0391 | RANOC0211 |
| MAPKAPK3 | RHSPN0549 | RGG113498 | RCABP0347 | RNAMR0002 | RBRBA0347 | RANOC0048 |
| MARCH1 | RHSPN0764 | RGG058885 | RCABP0033 | RNAMR0143 | RBRBA0063 | RANOC0157 |
| MARCH8 | RHSPN1455 | RGG082248 | RCABP0271 | RNAMR0412 | RBRBA0011 | RANOC0124 |

| | | | | | | |
|---|---|---|---|---|---|---|
| MARCO | RHSPN03880 | RGG091422 | RCABP02413 | RNAMR00217 | RBRBA03987 | RANOC02825 |
| MASP2 | RHSPN00217 | RGG146088 | RCABP00739 | RNAMR03000 | RBRBA023555 | |
| MAVS | RHSPN25855 | RGG067311 | RCABP03905 | RNAMR00683 | RBRBA025164 | |
| MAX | RHSPN19579 | RGG068395 | RCABP00775 | RNAMR00349 | RBRBA02428 | RANOC00170 |
| MB | RHSPN27271 | RGG006617 | RCABP00690 | RNAMR01083 | RBRBA03866 | RANOC01410 |
| MB21D1 | RHSPN09874 | RGG051572 | RCABP03714 | RNAMR03009 | RBRBA01801 | RANOC02533 |
| MBD4 | RHSPN06043 | RGG117211 | RCABP01535 | RNAMR03652 | RBRBA01902 | RANOC02303 |
| MBL2 | RHSPN14657 | RGG079754 | not predicted | RNAMR02440 | RBRBA026777 | |
| MDC1 | RHSPN094060 | | RCABP02883 | RNAMR01317 | RBRBA01000 | RANOC02723 |
| MDM2 | RHSPN17877 | RGG003790 | RCABP02638 | RNAMR04109 | RBRBA00119 | RANOC02249 |
| MDM2L | | | | | | |
| MDM4 | RHSPN02378 | RGG156112 | RCABP01671 | RNAMR02823 | RBRBA03921 | RANOC01160 |
| MDS2 | | | | | | |
| MECOM | RHSPN06326 | RGG103303 | RCABP00858 | RNAMR00067 | RBRBA04081 | RANOC00893 |
| MED1 | RHSPN22800 | RGG159353 | RCABP02169 | RNAMR02559 | RBRBA00826 | RANOC01693 |
| MED12 | RHSPN27999 | RGG055786 | RCABP02487 | RNAMR03869 | RBRBA02715 | RANOC02950 |
| MED19 | RHSPN16065 | RGG071751 | RCABP00772 | RNAMR03583 | RBRBA03849 | RANOC00030 |
| MEF2C | RHSPN08305 | RGG176860 | RCABP01593 | RNAMR00777 | RBRBA02101 | RANOC00733 |
| MEFV | RHSPN211861 | | not predicted | RNAMR03080 | RBRBA023004 | |
| MEN1 | RHSPN162573 | | RCABP00292 | RNAMR01853 | RBRBA01127 | RANOC03331 |
| MEP1A | RHSPN09751 | RGG054767 | RCABP00936 | RNAMR0125 | RBRBA00692 | RANOC00214 |
| MET | RHSPN11479 | RGG002805 | RCABP02085 | RNAMR03134 | RBRBA02919 | RANOC01524 |
| METTL1 | RHSPN178129 | | RCABP01770 | RNAMR02723 | RBRBA03098 | RANOC00433 |
| METTL21B | RHSPN178135 | | RCABP01753 | RNAMR02723 | RBRBA03098 | RANOC00433 |
| MFNG | RHSPN27306 | RGG006308 | RCABP01075 | RNAMR01170 | RBRBA03639 | RANOC01411 |
| MGAT5 | RHSPN03998 | RGG091578 | RCABP00141 | RNAMR00220 | RBRBA01738 | RANOC00042 |
| MGMT | RHSPN15404 | RGG085357 | RCABP00183 | RNAMR00483 | RBRBA03177 | RANOC00937 |
| MGP | RHSPN17282 | RGG005244 | RCABP01569 | RNAMR01193 | RBRBA01468 | RANOC01417 |
| MID2 | RHSPN28178 | RGG055363 | RCABP00590 | RNAMR02754 | RBRBA012580 | |
| MIF | RHSPN27086 | RGG127464 | RCABP030691 | | | |
| MINPP1 | RHSPN14931 | RGG080445 | RCABP04278 | RNAMR02441 | RBRBA02679 | RANOC02204 |
| MKL1 | RHSPN27384 | RGG005853 | RCABP00191 | RNAMR01163 | RBRBA02083 | RANOC01494 |
| MLH1 | RHSPN05204 | RGG030592 | RCABP02285 | RNAMR02397 | RBRBA02834 | RANOC01678 |
| MLH3 | RHSPN19714 | RGG075541 | RCABP03135 | RNAMR00323 | RBRBA03692 | RANOC00010 |
| MLLT1 | RHSPN24366 | RGG160980 | RCABP00016 | RNAMR03173 | RBRBA038027 | |
| MLLT10 | RHSPN14360 | RGG026906 | RCABP00757 | RNAMR0260 | RBRBA01347 | RANOC01817 |
| MLLT11 | RHSPN01667 | RGG154743 | RCABP04071 | RNAMR02011 | RBRBA032880 | |
| MLLT3 | RHSPN13083 | RGG174095 | RCABP02729 | RNAMR00450 | RBRBA03214 | RANOC00557 |
| MLLT4 | RHSPN10488 | RGG047143 | RCABP00130 | RNAMR0230 | RBRBA00191 | RANOC00275 |
| MLLT6 | RHSPN22782 | RGG159240 | RCABP02172 | RNAMR02563 | RBRBA00829 | RANOC01790 |
| MMP20 | RHSPN166923 | | not predicted | RNAMR03680 | RBRBA02202 | RANOC01032 |
| MMS19 | RHSPN15056 | RGG083236 | RCABP00872 | RNAMR00469 | RBRBA03938 | RANOC03124 |
| MN1 | RHSPN27137 | RGG127158 | RCABP04242 | RNAMR00940 | RBRBA018576 | |
| MNAT1 | RHSPN19537 | RGG077971 | RCABP01980 | RNAMR00354 | RBRBA02433 | RANOC02265 |
| MPG | RHSPN21029 | RGG123706 | RCABP02468 | RNAMR03974 | RBRBA01942 | RANOC03390 |

| | | | | | | |
|---|---|---|---|---|---|---|
| MPL | RHSPN00787 | RGG096927 | RCABP00253 | RNAMR0379 | RBRBA01375 | RANOC0127 |
| MPLKIP | RHSPN1084 | RGG031279 | RCABP02360 | RNAMR0063 | RBRBA02633 | RANOC0174 |
| MR1 | RHSPN02197 | RGG130294 | RCABP00701 | RNAMR0194 | RBRBA03250 | RANOC0080 |
| MRAS | RHSPN0613 | RGG090786 | RCABP02574 | RNAMR0004 | RBRBA01035 | RANOC0003 |
| MRC2 | RHSPN23262 | RGG158321 | RCABP03445 | RNAMR0346 | RBRBA00388 | RANOC0168 |
| MRE11A | RHSPN1665 | RGG022273 | RCABP00505 | RNAMR0041 | RBRBA03615 | RANOC0103 |
| MS4A1 | RHSPN161132 | | RCABP00405 | RNAMR0403 | RBRBA006280 | |
| MSH2 | RHSPN03248 | RGG042408 | RCABP04409 | RNAMR0055 | RBRBA01204 | RANOC0009 |
| MSH3 | RHSPN08266 | RGG177137 | RCABP00251 | RNAMR0077 | RBRBA00320 | RANOC0051 |
| MSH5 | RHSPN09456 | RGG049065 | RCABP03021 | RNAMR0130 | RBRBA00552 | RANOC0025 |
| MSH6 | RHSPN0325 | RGG042428 | RCABP04436 | RNAMR0055 | RBRBA01204 | RANOC0009 |
| MSI2 | RHSPN2318 | RGG140876 | RCABP01095 | RNAMR0258 | RBRBA02240 | RANOC0274 |
| MSN | RHSPN2796 | RGG054909 | RCABP03882 | RNAMR0383 | RBRBA02713 | RANOC0276 |
| MSRA | RHSPN1197 | RGG054321 | RCABP01014 | RNAMR0145 | RBRBA02507 | RANOC0112 |
| MSRB1 | RHSPN21118 | RGG122400 | RCABP03589 | RNAMR0418 | RBRBA01442 | RANOC0227 |
| MSTN | RHSPN0435 | RGG085824 | RCABP03183 | RNAMR035060 | | RANOC0046 |
| MT1A | RHSPN217103 | | not predicted | RNAMR041583 | | |
| MT1E | RHSPN217091 | | | | | |
| MT2A | RHSPN217084 | | | | | |
| MTCP1 | RHSPN28538 | RGG055698 | RCABP00462 | RNAMR0412 | RBRBA03881 | RANOC0287 |
| MTFMT | RHSPN20587 | RGG108338 | RCABP03895 | RNAMR0217 | RBRBA00710 | RANOC0263 |
| MTOR | RHSPN0022 | RGG146117 | RCABP00738 | RNAMR0296 | RBRBA02356 | RANOC0296 |
| MTR | RHSPN0273 | RGG046478 | RCABP01342 | RNAMR0206 | RBRBA03399 | RANOC0008 |
| MTTP | RHSPN07278 | RGG063404 | RCABP03619 | RNAMR0323 | RBRBA00202 | RANOC0247 |
| MUL1 | RHSPN0037 | RGG146431 | RCABP01111 | RNAMR0165 | RBRBA00533 | RANOC0219 |
| MUS81 | RHSPN163116 | | RCABP03337 | RNAMR0184 | RBRBA011349 | |
| MUSK | RHSPN1370 | RGG177500 | RCABP03793 | RNAMR0142 | RBRBA00540 | RANOC0275 |
| MUTYH | RHSPN00847 | RGG097349 | RCABP03360 | RNAMR0389 | RBRBA03389 | RANOC0127 |
| MVK | RHSPN1819 | RGG126995 | RCABP03443 | RNAMR0097 | RBRBA010663 | |
| MYB | RHSPN10238 | RGG048795 | RCABP01505 | RNAMR0375 | RBRBA00451 | RANOC0196 |
| MYC | RHSPN1275 | RGG039691 | RCABP01453 | RNAMR0040 | RBRBA03952 | RANOC0111 |
| MYCL | RHSPN00717 | RGG150735 | RCABP02014 | RNAMR0171 | RBRBA01713 | RANOC0279 |
| MYCN | RHSPN0295 | RGG053058 | RCABP01060 | RNAMR0360 | RBRBA03758 | RANOC0021 |
| MYD88 | RHSPN0522 | RGG025299 | RCABP01515 | RNAMR0013 | RBRBA03555 | RANOC0165 |
| MYH11 | RHSPN2134 | RGG122958 | RCABP02357 | RNAMR0218 | RBRBA021747 | |
| MYH9 | RHSPN2728 | RGG006540 | RCABP01309 | RNAMR0117 | RBRBA03642 | RANOC0141 |
| MYLK4 | RHSPN0907 | RGG033120 | RCABP02269 | RNAMR0138 | RBRBA01830 | RANOC0244 |
| MYO5A | RHSPN2047 | RGG106817 | RCABP01025 | RNAMR0014 | RBRBA010890 | |
| MYO5AB | | | | | | |
| MYO5B | RHSPN239912 | | RCABP04087 | RNAMR0212 | RBRBA01395 | RANOC0076 |
| MYO9B | RHSPN2466 | RGG162483 | RCABP01410 | RNAMR0393 | RBRBA001305 | |
| MYT1L | RHSPN028680 | | RCABP02150 | RNAMR0344 | RBRBA01531 | RANOC0037 |
| NAB1 | RHSPN0436 | RGG085743 | RCABP03645 | RNAMR0350 | RBRBA02230 | RANOC0046 |
| NAB2 | RHSPN177772 | | RCABP01644 | RNAMR0273 | RBRBA03092 | RANOC0203 |
| NABP2 | RHSPN17746 | RGG166957 | RCABP01649 | RNAMR0274 | RBRBA02786 | RANOC0076 |

| | | | | | | |
|---|---|---|---|---|---|---|
| NACA | RHSPN17768 | RGG167066 | RCABP01645 | RNAMR02730 | RBRBA02183 | RANOC02757 |
| NAIP | RHSPN081788 | | not predicted | RNAMR03890 | RBRBA00306 | RANOC00515 |
| NAV2 | RHSPN15797 | RGG068919 | RCABP02460 | RNAMR01764 | RBRBA01814 | RANOC00027 |
| NBEA | RHSPN18649 | RGG020894 | RCABP02342 | RNAMR02084 | RBRBA03625 | RANOC01022 |
| NBN | RHSPN12523 | RGG038046 | RCABP02528 | RNAMR00827 | RBRBA03265 | RANOC01198 |
| NCAM1 | RHSPN16780 | RGG152592 | RCABP00570 | RNAMR02269 | RBRBA00155 | RANOC02557 |
| NCAM2 | RHSPN26601 | RGG013772 | RCABP00278 | RNAMR00833 | RBRBA01993 | RANOC01003 |
| NCF1 | RHSPN11101 | RGG138401 | RCABP01776 | RNAMR00611 | RBRBA02853 | RANOC03469 |
| NCF2 | RHSPN02222 | RGG094684 | RCABP00517 | RNAMR02889 | RBRBA03397 | RANOC01997 |
| NCF4 | RHSPN27291 | RGG006470 | RCABP01078 | RNAMR01174 | RBRBA03641 | RANOC01417 |
| NCKAP1L | RHSPN17700 | RGG166473 | RCABP01258 | RNAMR03887 | RBRBA01776 | RANOC02227 |
| NCKIPSD | RHSPN05407 | RGG115374 | RCABP03070 | RNAMR00008 | RBRBA01277 | RANOC00485 |
| NCOA1 | RHSPN03007 | RGG054083 | RCABP02099 | RNAMR00554 | RBRBA02545 | RANOC02517 |
| NCOA2 | RHSPN12425 | RGG037369 | RCABP02450 | RNAMR00810 | RBRBA01412 | RANOC01347 |
| NCOA4 | RHSPN14569 | RGG079600 | RCABP00347 | RNAMR04057 | RBRBA03622 | RANOC03459 |
| NCOR1 | RHSPN22453 | RGG140037 | RCABP00157 | RNAMR03680 | RBRBA01926 | RANOC03343 |
| NCOR2 | RHSPN18420 | RGG125959 | RCABP03008 | RNAMR010075 | | RANOC01857 |
| NCR2 | RHSPN09639 | RGG157325 | not predicted | RNAMR03890 | RBRBA00306 | RANOC00515 |
| NCR3 | RHSPN094372 | | RCABP02574 | RNAMR01310 | RBRBA005553 | |
| NDFIP1 | RHSPN08696 | RGG120493 | RCABP04174 | RNAMR02040 | RBRBA03374 | RANOC00635 |
| NDRG1 | RHSPN12788 | RGG040005 | RCABP00059 | RNAMR00401 | RBRBA02269 | RANOC01184 |
| NDRG2 | RHSPN191395 | | RCABP01587 | RNAMR03110 | RBRBA00111 | RANOC02229 |
| NDRG3 | RHSPN26164 | RGG141508 | RCABP01986 | RNAMR03390 | RBRBA01684 | RANOC01173 |
| NDRG4 | RHSPN21763 | RGG109307 | RCABP00165 | RNAMR00923 | RBRBA023641 | |
| NDUFA11 | RHSPN24358 | RGG160388 | RCABP00062 | RNAMR03174 | RBRBA037319 | |
| NDUFA3 | RHSPN255901 | | RCABP02323 | RNAMR03280 | RBRBA03580 | RANOC02333 |
| NDUFS1 | RHSPN04492 | RGG088238 | RCABP01081 | RNAMR01744 | RBRBA02455 | RANOC02020 |
| NEDD4 | RHSPN20493 | RGG106702 | RCABP00937 | RNAMR00144 | RBRBA032379 | |
| NEIL1 | RHSPN20744 | RGG105516 | RCABP02928 | RNAMR0381 | RBRBA035741 | |
| NEIL2 | RHSPN11988 | RGG054443 | RCABP03663 | RNAMR01454 | RBRBA00525 | RANOC03047 |
| NEIL3 | RHSPN07716 | RGG061043 | RCABP01470 | RNAMR01905 | RBRBA02730 | RANOC01577 |
| NETO1 | RHSPN24122 | RGG034916 | RCABP01356 | RNAMR02804 | RBRBA00778 | RANOC01130 |
| NF1 | RHSPN22662 | RGG140915 | RCABP01096 | RNAMR03674 | RBRBA02905 | RANOC02890 |
| NF2 | RHSPN27170 | RGG128634 | RCABP03766 | RNAMR00957 | RBRBA01862 | RANOC02680 |
| NF2L | | | | | | |
| NFAM1 | RHSPN274279 | | RCABP03235 | RNAMR01150 | RBRBA00655 | RANOC02967 |
| NFAT5 | RHSPN21886 | RGG112634 | RCABP03189 | RNAMR00904 | RBRBA03381 | RANOC02993 |
| NFATC1 | RHSPN24163 | RGG032014 | RCABP00197 | RNAMR02800 | RBRBA00781 | RANOC01227 |
| NFATC2 | RHSPN26357 | RGG144441 | RCABP03313 | RNAMR02413 | RBRBA01388 | RANOC02114 |
| NFATC3 | RHSPN21861 | RGG110426 | RCABP01237 | RNAMR00909 | RBRBA02749 | RANOC01877 |
| NFATC4 | RHSPN192889 | | RCABP00686 | RNAMR03014 | RBRBA00517 | RANOC02079 |
| NFE2L1 | RHSPN23071 | RGG159106 | RCABP02179 | RNAMR02567 | RBRBA00833 | RANOC01695 |
| NFE2L2 | RHSPN04272 | RGG088687 | RCABP04115 | RNAMR02383 | RBRBA00254 | RANOC02672 |
| NFIB | RHSPN13044 | RGG173812 | RCABP00084 | RNAMR00440 | RBRBA01842 | RANOC00528 |
| NFIL3 | RHSPN13513 | RGG175456 | RCABP01707 | RNAMR03611 | RBRBA03619 | RANOC00418 |

| | | | | | | |
|---|---|---|---|---|---|---|
| NFKB1 | RHSPN07291 | RGG063509 | RCABP01416 | RNAMR03233 | RBRBA008717 | |
| NFKB2 | RHSPN15144 | RGG081956 | RCABP00992 | RNAMR03249 | RBRBA02993 | RANOC01929 |
| NFKBIA | RHSPN19355 | RGG075132 | RCABP03602 | RNAMR03363 | RBRBA01214 | RANOC00427 |
| NFKBIB | RHSPN25025 | RGG168785 | RCABP00815 | RNAMR02517 | RBRBA04013 | RANOC03177 |
| NFKBID | RHSPN24932 | RGG180174 | RCABP04192 | RNAMR02504 | RBRBA00886 | RANOC02319 |
| NFKBIE | RHSPN09727 | RGG045517 | RCABP04317 | RNAMR01253 | RBRBA00696 | RANOC00279 |
| NFKBIZ | RHSPN05800 | RGG012392 | RCABP03359 | RNAMR02408 | RBRBA01618 | RANOC01015 |
| NGB | RHSPN19741 | RGG075779 | RCABP03120 | RNAMR00324 | RBRBA00014 | RANOC00010 |
| NHEJ1 | RHSPN04635 | RGG090274 | RCABP02200 | RNAMR03063 | RBRBA03671 | RANOC00034 |
| NIN | RHSPN19435 | RGG078765 | RCABP00079 | RNAMR00160 | RBRBA01610 | RANOC03145 |
| NKIRAS1 | RHSPN05120 | RGG029333 | RCABP01257 | RNAMR02383 | RBRBA01055 | RANOC02567 |
| NKIRAS2 | RHSPN22885 | RGG159745 | RCABP01963 | RNAMR02543 | RBRBA02652 | RANOC01680 |
| NKX2-1 | RHSPN19365 | RGG075207 | RCABP02050 | RNAMR03362 | RBRBA01224 | RANOC00102 |
| NLN | RHSPN08133 | RGG172481 | RCABP00579 | RNAMR02894 | RBRBA03000 | RANOC00509 |
| NLRC3 | RHSPN21199 | RGG124019 | RCABP01301 | RNAMR03083 | RBRBA02297 | RANOC02853 |
| NLRC4 | RHSPN031301 | | not predicted | RNAMR00537 | RBRBA04022 | RANOC00283 |
| NLRC5 | RHSPN21720 | RGG109563 | RCABP00615 | RNAMR00929 | RBRBA02368 | RANOC03408 |
| NLRP1 | RHSPN222884 | | not predicted | | RBRBA035506 | |
| NLRP10 | RHSPN156647 | | | RNAMR04177 | RBRBA024182 | |
| NLRP11 | RHSPN256769 | | | | RBRBA026904 | |
| NLRP12 | RHSPN255797 | | RCABP008481 | | RBRBA00978 | RANOC00152 |
| NLRP13 | RHSPN256786 | | | | RBRBA026902 | |
| NLRP14 | RHSPN156518 | | | RNAMR02959 | RBRBA024161 | |
| NLRP2 | RHSPN256376 | | | | RBRBA020345 | |
| NLRP3 | RHSPN02812 | RGG068511 | | RNAMR04189 | RBRBA000686 | |
| NLRP4 | RHSPN256777 | | | RNAMR03324 | RBRBA026903 | |
| NLRP5 | RHSPN256795 | | | | RBRBA026899 | |
| NLRP6 | RHSPN154670 | | | RNAMR017899 | | |
| NLRP7 | RHSPN256361 | | | | RBRBA011005 | |
| NLRP8 | RHSPN256790 | | | RNAMR03324 | RBRBA026900 | |
| NLRP9 | RHSPN256749 | | | | RBRBA035314 | |
| NLRPs | | | | | | |
| NLRX1 | RHSPN16879 | RGG152016 | RCABP00124 | RNAMR02287 | RBRBA00140 | RANOC03062 |
| NMI | RHSPN04074 | RGG092157 | RCABP01239 | RNAMR03607 | RBRBA00479 | RANOC02420 |
| NNMT | RHSPN167958 | | RCABP02600 | RNAMR0227 | RBRBA00151 | RANOC02867 |
| NOD1 | RHSPN10769 | RGG029789 | RCABP01522 | RNAMR00558 | RBRBA00960 | RANOC01675 |
| NOD2 | RHSPN216636 | | not predicted | RNAMR01404 | RBRBA032621 | |
| NOG | RHSPN23176 | RGG135629 | RCABP03462 | RNAMR02587 | RBRBA02232 | RANOC01950 |
| NONO | RHSPN28004 | RGG055748 | RCABP02490 | RNAMR0387 | RBRBA02717 | RANOC02950 |
| NOS1 | RHSPN18290 | RGG128733 | RCABP02952 | RNAMR00980 | RBRBA01073 | RANOC01855 |
| NOS2 | RHSPN22586 | RGG140942 | RCABP01086 | RNAMR02303 | RBRBA02813 | RANOC03069 |
| NOTCH1 | RHSPN14107 | RGG132984 | RCABP04027 | RNAMR015029 | | RANOC03327 |
| NOTCH2 | RHSPN01505 | RGG093558 | RCABP00999 | RNAMR01994 | RBRBA030299 | |
| NPAT | RHSPN16728 | RGG021762 | RCABP02113 | RNAMR02260 | RBRBA03973 | RANOC03017 |
| NPM1 | RHSPN08901 | RGG117928 | RCABP04140 | RNAMR03633 | RBRBA03140 | RANOC00050 |

| | | | | | | |
|---|---|---|---|---|---|---|
| NPR2 | RHSPN13209 | RGG171162 | RCABP03906 | RNAMR03619 | RBRBA037455 | |
| NPY | RHSPN10710 | RGG028651 | RCABP00598 | RNAMR00563 | RBRBA02190 | RANOC01667 |
| NR3C1 | RHSPN08709 | RGG120540 | RCABP04179 | RNAMR02042 | RBRBA03375 | RANOC00465 |
| NRAS | RHSPN01451 | RGG156949 | | RNAMR01987 | RBRBA00562 | RANOC01170 |
| NRDE2 | RHSPN19808 | RGG076267 | RCABP01549 | RNAMR00318 | RBRBA02059 | RANOC00195 |
| NRG1 | RHSPN12194 | RGG176305 | RCABP01179 | RNAMR02150 | RBRBA01102 | RANOC00560 |
| NRROS | RHSPN06562 | RGG100171 | RCABP00359 | RNAMR0008 | RBRBA02323 | RANOC00914 |
| NRXN3 | RHSPN19758 | RGG075969 | RCABP03116 | RNAMR0032 | RBRBA00589 | RANOC00314 |
| NSD1 | RHSPN08967 | RGG118844 | RCABP02481 | RNAMR00239 | RBRBA01296 | RANOC00789 |
| NT5C2 | RHSPN15164 | RGG083728 | RCABP01701 | RNAMR03254 | RBRBA02991 | RANOC00845 |
| NT5DC1 | RHSPN10113 | RGG049751 | not predicted | RNAMR03513 | RBRBA03839 | RANOC00069 |
| NT5DC4 | RHSPN038409 | | | RNAMR010839 | | |
| NTF3 | RHSPN17112 | RGG009903 | RCABP03579 | RNAMR03544 | RBRBA03630 | RANOC01535 |
| NTHL1 | RHSPN21125 | RGG122461 | RCABP03584 | RNAMR03972 | RBRBA01806 | RANOC03254 |
| NTRK1 | RHSPN01892 | RGG153211 | RCABP02251 | RNAMR037320 | | RANOC02989 |
| NTRK3 | RHSPN20882 | RGG107807 | RCABP03540 | RNAMR01882 | RBRBA023485 | |
| NUDT1 | RHSPN10559 | RGG121559 | RCABP02419 | RNAMR00640 | RBRBA03654 | RANOC0253 |
| NUMA1 | RHSPN16447 | RGG023296 | RCABP00238 | RNAMR02939 | RBRBA00174 | RANOC03110 |
| NUP214 | RHSPN14000 | RGG132514 | RCABP02741 | RNAMR01519 | RBRBA03496 | RANOC03482 |
| NUTM1 | RHSPN202391 | | RCABP02090 | RNAMR03114 | RBRBA039842 | |
| NUTM2A | | | | | | |
| NUTM2B | RHSPN148746 | | | | | |
| NUTM2D | | | | | | |
| NUTM2E | | | | | | |
| NUTM2F | | | | | | |
| NUTM2G | RHSPN135946 | | | | RBRBA036184 | |
| OASL | RHSPN18343 | RGG114102 | RCABP02885 | RNAMR00974 | RBRBA010676 | |
| ODF1 | RHSPN12631 | RGG038775 | RCABP04372 | RNAMR01633 | RBRBA03132 | RANOC01115 |
| OGDH | RHSPN10890 | RGG148461 | RCABP00655 | RNAMR0060 | RBRBA03595 | RANOC02170 |
| OGG1 | RHSPN04980 | RGG115957 | RCABP02853 | RNAMR00140 | RBRBA01911 | RANOC00493 |
| OLIG2 | RHSPN26664 | RGG014099 | RCABP00581 | RNAMR008407 | | RANOC02982 |
| OMD | RHSPN13525 | RGG114196 | RCABP01679 | RNAMR03613 | RBRBA03438 | RANOC00487 |
| OPRD1 | RHSPN00546 | RGG149407 | RCABP03207 | RNAMR01687 | RBRBA02566 | RANOC03180 |
| OPRK1 | RHSPN12323 | RGG036644 | RCABP00673 | RNAMR00362 | RBRBA00213 | RANOC02188 |
| OPRM1 | RHSPN10377 | RGG047903 | RCABP043261 | | RBRBA02281 | RANOC00079 |
| OR56A1 | RHSPN156229 | | not predicted | | RBRBA040457 | |
| ORC5 | RHSPN11397 | RGG001762 | RCABP03036 | RNAMR0058 | RBRBA00952 | RANOC01549 |
| OSGIN1 | RHSPN22010 | RGG110329 | RCABP01667 | RNAMR00888 | RBRBA00916 | RANOC01872 |
| OTOL1 | RHSPN06304 | RGG103480 | RCABP00102 | RNAMR00127 | RBRBA01119 | RANOC01040 |
| OTUD6A | RHSPN279853 | | not predicted | | RBRBA000894 | |
| OTUD7B | RHSPN01625 | RGG152966 | RCABP02249 | RNAMR02004 | RBRBA015491 | |
| OTULIN | RHSPN07895 | RGG033587 | RCABP02028 | RNAMR01129 | RBRBA02039 | RANOC01347 |
| OTX2 | RHSPN19488 | RGG078317 | RCABP00974 | RNAMR03713 | RBRBA03292 | RANOC01878 |
| P2RX7 | RHSPN18346 | RGG126248 | RCABP03012 | RNAMR00993 | RBRBA00053 | RANOC01860 |
| P2RY8 | RHSPN28566 | RGG017082 | RCABP00199 | RNAMR03344 | RBRBA01097 | RANOC00968 |

| | | | | | | |
|---|---|---|---|---|---|---|
| PAFAH1B2 | RHSPN16817 | RGG152245 | RCABP02597 | RNAMR02274 | RBRBA001498 | |
| PALB2 | RHSPN21427 | RGG122705 | RCABP03906 | RNAMR02204 | RBRBA03515 | RANOC03170 |
| PAPD5 | RHSPN21654 | RGG110745 | RCABP01518 | RNAMR01403 | RBRBA03262 | RANOC02117 |
| PARK2 | RHSPN10442 | RGG047324 | RCABP02338 | RNAMR03500 | RBRBA01407 | RANOC00370 |
| PARP1 | RHSPN02616 | RGG043890 | RCABP03934 | RNAMR02833 | RBRBA03241 | RANOC00091 |
| PARP2 | RHSPN191183 | | RCABP03860 | RNAMR03100 | RBRBA01385 | RANOC02084 |
| PARP3 | RHSPN05511 | RGG114066 | RCABP00224 | RNAMR00028 | RBRBA03470 | RANOC00477 |
| PARVG | RHSPN27458 | RGG009276 | RCABP01330 | RNAMR01152 | RBRBA00658 | RANOC01632 |
| PATZ1 | RHSPN27221 | RGG127781 | RCABP00585 | RNAMR00958 | RBRBA018706 | |
| PAWR | RHSPN17950 | RGG004182 | RCABP02854 | RNAMR01109 | RBRBA03771 | RANOC01421 |
| PAX3 | RHSPN04672 | RGG100891 | RCABP03275 | RNAMR03053 | RBRBA03679 | RANOC00833 |
| PAX4 | RHSPN115336 | | RCABP04250 | RNAMR02250 | RBRBA031147 | |
| PAX5 | RHSPN13237 | RGG179012 | RCABP01037 | RNAMR04063 | RBRBA03740 | RANOC01978 |
| PAX7 | RHSPN00336 | RGG146296 | RCABP01832 | RNAMR01650 | RBRBA036104 | |
| PAX8 | RHSPN038530 | | RCABP02084 | RNAMR01064 | RBRBA010870 | |
| PAXIP1 | RHSPN11855 | RGG025684 | RCABP00755 | RNAMR02643 | RBRBA03860 | RANOC01661 |
| PBRM1 | RHSPN05541 | RGG113094 | RCABP03478 | RNAMR03053 | RBRBA01402 | RANOC00793 |
| PBX1 | RHSPN02035 | RGG093915 | RCABP03505 | RNAMR03124 | RBRBA03286 | RANOC02662 |
| PCK1 | RHSPN26397 | RGG144153 | RCABP01385 | RNAMR00703 | RBRBA03309 | RANOC02062 |
| PCK2 | RHSPN192687 | | not predicted | RNAMR03022 | RBRBA00522 | RANOC03229 |
| PCM1 | RHSPN12042 | RGG063775 | RCABP02788 | RNAMR01910 | RBRBA01112 | RANOC01642 |
| PCMT1 | RHSPN10337 | RGG047736 | RCABP04412 | RNAMR02294 | RBRBA03647 | RANOC02261 |
| PCNA | RHSPN25870 | RGG147268 | RCABP00180 | RNAMR00688 | RBRBA02510 | RANOC03348 |
| PCNXL2 | RHSPN02701 | RGG046705 | RCABP02118 | RNAMR02073 | RBRBA00084 | RANOC00271 |
| PCP4 | RHSPN26746 | RGG014632 | RCABP01944 | RNAMR00858 | RBRBA03977 | RANOC00871 |
| PCSK1 | RHSPN08348 | RGG176673 | RCABP02828 | RNAMR0078 | RBRBA02108 | RANOC00410 |
| PDCD1 | RHSPN04916 | RGG102133 | RCABP00912 | RNAMR03459 | RBRBA002623 | |
| PDCD1LG2 | RHSPN130075 | | | RNAMR00450 | RBRBA007449 | |
| PDE4B | RHSPN01066 | RGG098863 | RCABP04426 | RNAMR0076 | RBRBA01625 | RANOC02669 |
| PDE4D | RHSPN08100 | RGG172286 | RCABP00278 | RNAMR01459 | RBRBA00293 | RANOC00508 |
| PDE4DIP | RHSPN01603 | RGG093574 | RCABP04147 | RNAMR01993 | RBRBA03100 | RANOC01235 |
| PDGFA | RHSPN10529 | RGG121266 | RCABP04342 | RNAMR00641 | RBRBA006777 | |
| PDGFB | RHSPN27362 | RGG005993 | RCABP02074 | RNAMR01168 | RBRBA01875 | RANOC01493 |
| PDGFRA | RHSPN06988 | RGG064204 | RCABP00631 | RNAMR01580 | RBRBA02843 | RANOC01443 |
| PDGFRB | RHSPN08762 | RGG119469 | RCABP00453 | RNAMR02050 | RBRBA01958 | RANOC00643 |
| PDPK1 | RHSPN21151 | RGG122823 | RCABP02003 | RNAMR03073 | RBRBA01811 | RANOC02962 |
| PELI1 | RHSPN03357 | RGG042679 | RCABP00839 | RNAMR01607 | RBRBA03822 | RANOC00040 |
| PER1 | RHSPN223736 | | RCABP03472 | RNAMR02350 | RBRBA03066 | RANOC02652 |
| PER2 | RHSPN04837 | RGG100671 | RCABP00445 | RNAMR03448 | RBRBA01062 | RANOC00911 |
| PEX5 | RHSPN17168 | RGG010657 | RCABP00602 | RNAMR03550 | RBRBA00838 | RANOC02068 |
| PGLYRP2 | RHSPN24638 | RGG163144 | RCABP00748 | RNAMR03859 | RBRBA02736 | RANOC00720 |
| PGPEP1 | RHSPN24713 | RGG162341 | RCABP01402 | RNAMR0394 | RBRBA024080 | |
| PHB | RHSPN23105 | RGG158867 | RCABP02184 | RNAMR02574 | RBRBA01638 | RANOC01690 |
| PHF6 | RHSPN28327 | RGG056221 | RCABP00534 | RNAMR02617 | RBRBA02276 | RANOC03093 |
| PHOX2B | RHSPN06929 | RGG064668 | RCABP03796 | RNAMR01589 | RBRBA028858 | |

| | | | | | | |
|---|---|---|---|---|---|---|
| PHPT1 | RHSPN1412 | RGG131036 | RCABP03511 | RNAMR0149 | RBRBA03462 | RANOC0336 |
| PHYHIP | RHSPN120914 | | RCABP0303 | RNAMR0143 | RBRBA0117 | RANOC0272 |
| PICALM | RHSPN1659 | RGG022845 | RCABP01061 | RNAMR0343 | RBRBA01481 | RANOC0103 |
| PIGC | RHSPN0211 | RGG093652 | RCABP00082 | RNAMR0195 | RBRBA0158 | RANOC0320 |
| PIGR | RHSPN0243 | RGG156473 | RCABP0227 | RNAMR0281 | RBRBA0390 | RANOC0212 |
| PIK3AP1 | RHSPN1504 | RGG081897 | RCABP0098 | RNAMR0046 | RBRBA0394 | RANOC0192 |
| PIK3C2G | RHSPN1730 | RGG008383 | RCABP0259 | RNAMR0119 | RBRBA014661 | |
| PIK3CA | RHSPN0638 | RGG102945 | RCABP00151 | RNAMR0007 | RBRBA0360 | RANOC0082 |
| PIK3R1 | RHSPN0814 | RGG172599 | RCABP0342 | RNAMR0289 | RBRBA0030 | RANOC0041 |
| PIK3R6 | RHSPN2239 | RGG134549 | RCABP03102 | RNAMR0236 | RBRBA0307 | RANOC0076 |
| PIM1 | RHSPN0959 | RGG155222 | RCABP0339 | RNAMR0127 | RBRBA0239 | RANOC0261 |
| PIM3 | RHSPN2751 | RGG002273 | RCABP01392 | RNAMR011460 | | RANOC0232 |
| PITPNM3 | RHSPN2229 | RGG141240 | RCABP03607 | RNAMR0233 | RBRBA004241 | |
| PKN1 | RHSPN246087 | | RCABP0127 | RNAMR0385 | RBRBA01341 | RANOC0060 |
| PKN2 | RHSPN0121 | RGG096133 | RCABP02152 | RNAMR0074 | RBRBA03924 | RANOC0137 |
| PKN3 | RHSPN139498 | | not predicte | RNAMR0152 | RBRBA034058 | |
| PKNOX2 | RHSPN1696 | RGG151072 | RCABP00407 | RNAMR0405 | RBRBA02017 | RANOC0258 |
| PLA2G1B | RHSPN1832 | RGG128058 | RCABP0184 | RNAMR0097 | RBRBA0107 | RANOC0252 |
| PLAG1 | RHSPN1234 | RGG036806 | RCABP0266 | RNAMR0375 | RBRBA0212 | RANOC0218 |
| PLAT | RHSPN1226 | RGG147995 | RCABP0065 | RNAMR0213 | RBRBA0010 | RANOC0289 |
| PLAU | RHSPN1482 | RGG081553 | RCABP0262 | RNAMR0124 | RBRBA01182 | RANOC0193 |
| PLCL2 | RHSPN0508 | RGG029157 | RCABP01504 | RNAMR0303 | RBRBA01051 | RANOC0173 |
| PLEKHA1 | RHSPN1533 | RGG084984 | RCABP00852 | RNAMR0048 | RBRBA03242 | RANOC0085 |
| PLG | RHSPN1043 | RGG047380 | RCABP0233 | RNAMR0350 | RBRBA02592 | RANOC0026 |
| PLXNA4 | RHSPN1159 | RGG000873 | RCABP0342 | RNAMR0224 | RBRBA0019 | RANOC0139 |
| PML | RHSPN2070 | RGG105494 | RCABP00492 | RNAMR0215 | RBRBA035813 | |
| PMS1 | RHSPN0435 | RGG085836 | RCABP031860 | | RBRBA0140 | RANOC0046 |
| PMS2 | RHSPN1059 | RGG121113 | RCABP0204 | RNAMR0065 | RBRBA0314 | RANOC0283 |
| PNKP | RHSPN254139 | | RCABP04122 | RNAMR0329 | RBRBA0316 | RANOC0169 |
| POLA1 | RHSPN2771 | RGG015732 | RCABP0047 | RNAMR0382 | RBRBA0080 | RANOC0098 |
| POLA2 | RHSPN1628 | RGG169213 | RCABP0333 | RNAMR0185 | RBRBA0113 | RANOC0345 |
| POLB | RHSPN1227 | RGG148047 | RCABP0428 | RNAMR0213 | RBRBA00101 | RANOC0295 |
| POLD1 | RHSPN2543 | RGG180364 | RCABP04124 | RNAMR0329 | RBRBA03171 | RANOC0170 |
| POLD2 | RHSPN1087 | RGG148536 | RCABP0037 | RNAMR0060 | RBRBA0359 | RANOC0216 |
| POLD3 | RHSPN1649 | RGG023425 | RCABP0039 | RNAMR0293 | RBRBA0151 | RANOC0281 |
| POLD4 | RHSPN1635 | RGG068003 | RCABP0249 | RNAMR0183 | RBRBA01144 | RANOC0003 |
| POLE | RHSPN1848 | RGG128314 | RCABP0409 | RNAMR0101 | RBRBA02337 | RANOC0267 |
| POLE2 | RHSPN1941 | RGG078661 | RCABP0413 | RNAMR0016 | RBRBA0193 | RANOC0291 |
| POLE3 | RHSPN1373 | RGG131285 | RCABP0056 | RNAMR0156 | RBRBA016052 | |
| POLE4 | RHSPN0347 | RGG160458 | RCABP0134 | RNAMR0107 | RBRBA000598 | |
| POLG | RHSPN2090 | RGG107715 | RCABP0353 | RNAMR0188 | RBRBA0408 | RANOC0273 |
| POLG2 | RHSPN2330 | RGG135923 | RCABP0345 | RNAMR0363 | RBRBA0039 | RANOC0221 |
| POLH | RHSPN0970 | RGG045658 | RCABP0429 | RNAMR0125 | RBRBA0069 | RANOC0008 |
| POLI | RHSPN240133 | | RCABP00414 | RNAMR0212 | RBRBA0126 | RANOC0204 |
| POLK | RHSPN0821 | RGG172931 | RCABP0386 | RNAMR0389 | RBRBA0031 | RANOC0051 |

| | | | | | | |
|---|---|---|---|---|---|---|
| POLL | RHSPN15121 | RGG083386 | RCABP02563 | RNAMR03241 | RBRBA03000 | RANOC02175 |
| POLM | RHSPN108730 | | RCABP02077 | RNAMR00603 | RBRBA035973 | |
| POLN | RHSPN06655 | RGG066403 | RCABP00785 | RNAMR01349 | RBRBA040701 | |
| POLQ | RHSPN05932 | RGG011182 | RCABP00573 | RNAMR00192 | RBRBA02602 | RANOC02483 |
| POLR2L | RHSPN15509 | RGG071121 | RCABP00341 | RNAMR01773 | RBRBA039856 | |
| POLR3B | RHSPN18152 | RGG006792 | RCABP04224 | RNAMR0109 | RBRBA00790 | RANOC01409 |
| POLR3D | RHSPN120922 | | RCABP03033 | RNAMR01439 | RBRBA01175 | RANOC02727 |
| POMT2 | RHSPN19742 | RGG075788 | RCABP03119 | RNAMR00324 | RBRBA00591 | RANOC00009 |
| PON1 | RHSPN112472 | | not predicted | RNAMR02879 | RBRBA026987 | |
| POT1 | RHSPN11521 | RGG002499 | RCABP03038 | RNAMR0225 | RBRBA01838 | RANOC01528 |
| POU1F1 | RHSPN05729 | RGG013412 | RCABP00555 | RNAMR02399 | RBRBA02974 | RANOC01094 |
| POU2AF1 | RHSPN16749 | RGG152053 | RCABP00125 | RNAMR02261 | RBRBA00169 | RANOC02862 |
| POU2F2 | RHSPN251302 | | RCABP01840 | RNAMR03910 | RBRBA03733 | RANOC03181 |
| POU5F1 | RHSPN094215 | | RCABP01164 | RNAMR01313 | RBRBA00556 | RANOC02978 |
| PPARG | RHSPN05026 | RGG114375 | RCABP00483 | RNAMR04117 | RBRBA01904 | RANOC00799 |
| PPARGC1A | RHSPN06831 | RGG065327 | RCABP02547 | RNAMR01330 | RBRBA02914 | RANOC01532 |
| PPFIBP1 | RHSPN17383 | RGG009020 | RCABP00783 | RNAMR04074 | RBRBA02153 | RANOC01635 |
| PPM1D | RHSPN23241 | RGG140607 | RCABP01089 | RNAMR03640 | RBRBA00379 | RANOC02638 |
| PPP1R1A | RHSPN17702 | RGG166461 | RCABP01259 | RNAMR03887 | RBRBA01776 | RANOC02225 |
| PPP1R26 | RHSPN14075 | RGG132946 | RCABP04022 | RNAMR01510 | RBRBA02022 | RANOC03204 |
| PPP2R1A | RHSPN255081 | | RCABP03250 | RNAMR0328 | RBRBA03320 | RANOC02341 |
| PPP2R1B | RHSPN167593 | | | RNAMR02264 | RBRBA001526 | |
| PPP2R2B | RHSPN08729 | RGG120623 | RCABP03084 | RNAMR02044 | RBRBA04014 | RANOC02142 |
| PPP2R2C | RHSPN06710 | RGG065905 | RCABP02227 | RNAMR013421 | | RANOC01534 |
| PPP2R3C | RHSPN19350 | RGG075114 | RCABP03603 | RNAMR03363 | RBRBA01223 | RANOC00420 |
| PPP3CB | RHSPN14807 | RGG081707 | RCABP04135 | RNAMR01238 | RBRBA01185 | RANOC01930 |
| PPP6C | RHSPN13854 | RGG133689 | RCABP04040 | RNAMR01548 | RBRBA03422 | RANOC03470 |
| PRCC | RHSPN018890 | | RCABP02238 | RNAMR0373 | RBRBA01575 | RANOC02988 |
| PRDM1 | RHSPN10028 | RGG050495 | RCABP01474 | RNAMR02153 | RBRBA03723 | RANOC00239 |
| PRDM16 | RHSPN00123 | RGG145184 | RCABP03684 | RNAMR029859 | | RANOC02982 |
| PRDM1L | | | | | | |
| PREX1 | RHSPN26317 | RGG142560 | RCABP04011 | RNAMR02433 | RBRBA02095 | RANOC01179 |
| PRF1 | RHSPN14758 | RGG130843 | RCABP03702 | RNAMR0123 | RBRBA03568 | RANOC02087 |
| PRF1G | | | | | | |
| PRF1I | | | | | | |
| PRF1K | | | | | | |
| PRG4 | RHSPN02242 | RGG094977 | RCABP01462 | RNAMR0289 | RBRBA01688 | RANOC01990 |
| PRKCA | RHSPN23323 | RGG136086 | RCABP00085 | RNAMR03633 | RBRBA00399 | RANOC02263 |
| PRKCB | RHSPN21431 | RGG122625 | RCABP00043 | RNAMR022050 | | |
| PRKCD | RHSPN05554 | RGG113327 | RCABP00513 | RNAMR03042 | RBRBA01934 | RANOC0048 |
| PRKCE | RHSPN03228 | RGG045160 | RCABP03617 | RNAMR00549 | RBRBA01201 | RANOC02360 |
| PRKCG | RHSPN255832 | | RCABP02326 | RNAMR032808 | | RANOC02494 |
| PRKCH | RHSPN19542 | RGG077950 | RCABP01979 | RNAMR00354 | RBRBA02433 | RANOC00002 |
| PRKCQ | RHSPN14244 | RGG000985 | RCABP00572 | RNAMR02062 | RBRBA039229 | |
| PRKCZ | RHSPN00091 | RGG145326 | RCABP01472 | RNAMR02984 | RBRBA028423 | |

| | | | | | | |
|---|---|---|---|---|---|---|
| PRKD2 | RHSPN252715 | | RCABP02768 | RNAMR03317 | RBRBA002803 | |
| PRKDC | RHSPN12300 | RGG036491 | RCABP02956 | RNAMR00364 | RBRBA02775 | RANOC02010 |
| PRKRA | RHSPN042814 | | RCABP04116 | RNAMR03422 | RBRBA00257 | RANOC02462 |
| PRNP | RHSPN25863 | RGG147304 | RCABP00181 | RNAMR00687 | RBRBA02515 | RANOC02660 |
| PROC | RHSPN03931 | RGG099587 | RCABP03099 | RNAMR00200 | RBRBA03156 | RANOC00839 |
| PROM1 | RHSPN06793 | RGG065564 | RCABP00604 | RNAMR01333 | RBRBA02978 | RANOC01387 |
| PROP1 | RHSPN089959 | | not predicted | RNAMR0028 | RBRBA00876 | RANOC02226 |
| PROS1 | RHSPN05738 | RGG012891 | RCABP02508 | RNAMR02409 | RBRBA03188 | RANOC01017 |
| PRPF19 | RHSPN16121 | RGG068112 | RCABP02096 | RNAMR04033 | RBRBA01315 | RANOC02149 |
| PRPS1 | RHSPN28175 | RGG055501 | RCABP04445 | RNAMR02754 | RBRBA012601 | |
| PRR5-ARHGA | RHSPN274634 | | RCABP03381 | RNAMR0115 | RBRBA04021 | RANOC01437 |
| PRR5L | RHSPN15896 | RGG072361 | RCABP03410 | RNAMR0181 | RBRBA02253 | RANOC00120 |
| PRUNE2 | RHSPN13409 | RGG174595 | RCABP00697 | RNAMR02860 | RBRBA01658 | RANOC00739 |
| PSEN1 | RHSPN19667 | RGG073880 | RCABP01274 | RNAMR00333 | RBRBA01337 | RANOC00117 |
| PSIP1 | RHSPN13056 | RGG173902 | RCABP02205 | RNAMR00442 | RBRBA01841 | RANOC00419 |
| PSMC1 | RHSPN19807 | RGG076263 | RCABP01549 | RNAMR00358 | RBRBA02059 | RANOC00009 |
| PSMC2 | RHSPN11394 | RGG001724 | RCABP00657 | RNAMR0058 | RBRBA00953 | RANOC01447 |
| PSMC3 | RHSPN15996 | RGG072780 | RCABP02359 | RNAMR01794 | RBRBA02009 | RANOC00018 |
| PSMC4 | RHSPN250519 | | RCABP04192 | RNAMR04023 | RBRBA01732 | RANOC02844 |
| PSMC5 | RHSPN23286 | RGG158221 | RCABP01625 | RNAMR03470 | RBRBA00392 | RANOC01689 |
| PSMC6 | RHSPN19454 | RGG078875 | RCABP03882 | RNAMR00163 | RBRBA01608 | RANOC02868 |
| PSMD10 | RHSPN28181 | RGG057808 | RCABP01890 | RNAMR02753 | RBRBA01258 | RANOC01987 |
| PSMD11 | RHSPN22684 | RGG158069 | RCABP02590 | RNAMR03667 | RBRBA02908 | RANOC01680 |
| PSMD12 | RHSPN23329 | RGG136036 | RCABP02454 | RNAMR03633 | RBRBA00398 | RANOC02219 |
| PSMD13 | RHSPN15466 | RGG068852 | RCABP02944 | RNAMR017860 | | RANOC00028 |
| PSMD2 | RHSPN06436 | RGG102571 | RCABP03298 | RNAMR00099 | RBRBA02192 | RANOC02194 |
| PSMD4 | RHSPN01680 | RGG154791 | not predicted | RNAMR02010 | RBRBA03298 | RANOC03018 |
| PSMD5 | RHSPN13789 | RGG133169 | RCABP04030 | RNAMR01550 | RBRBA03118 | RANOC03398 |
| PTAFR | RHSPN00532 | RGG149174 | RCABP03489 | RNAMR01684 | RBRBA02569 | RANOC03079 |
| PTCD2 | RHSPN08190 | RGG173114 | RCABP04215 | RNAMR03894 | RBRBA00308 | RANOC00417 |
| PTCH1 | RHSPN13570 | RGG175179 | RCABP00619 | RNAMR01218 | RBRBA02394 | RANOC00739 |
| PTEN | RHSPN14936 | RGG080404 | RCABP04281 | RNAMR02443 | RBRBA02679 | RANOC02203 |
| PTGER4 | RHSPN08009 | RGG171779 | RCABP01261 | RNAMR01468 | RBRBA02686 | RANOC00500 |
| PTGS1 | RHSPN13819 | RGG133356 | RCABP04036 | RNAMR01551 | RBRBA03115 | RANOC03346 |
| PTGS2 | RHSPN02246 | RGG094946 | RCABP01461 | RNAMR02880 | RBRBA01687 | RANOC01990 |
| PTH | RHSPN15731 | RGG069628 | RCABP02211 | RNAMR01753 | RBRBA03294 | RANOC02160 |
| PTK2 | RHSPN12819 | RGG040182 | RCABP03240 | RNAMR00399 | RBRBA02932 | RANOC01110 |
| PTK2B | RHSPN12142 | RGG053876 | RCABP02101 | RNAMR01448 | RBRBA014516 | |
| PTPN1 | RHSPN26342 | RGG144577 | RCABP03315 | RNAMR02414 | RBRBA01389 | RANOC02113 |
| PTPN11 | RHSPN18251 | RGG126803 | RCABP03013 | RNAMR00980 | RBRBA00175 | RANOC02350 |
| PTPN2 | RHSPN23776 | RGG035191 | RCABP00209 | RNAMR01937 | RBRBA01488 | RANOC01344 |
| PTPN6 | RHSPN17157 | RGG010621 | RCABP00603 | RNAMR03554 | RBRBA023839 | |
| PTPRB | RHSPN17901 | RGG003902 | RCABP03294 | RNAMR01114 | RBRBA01378 | RANOC01509 |
| PTPRC | RHSPN02291 | RGG093139 | RCABP01713 | RNAMR0262 | RBRBA03878 | RANOC02032 |
| PTPRJ | RHSPN16014 | RGG070603 | RCABP03845 | RNAMR01790 | RBRBA02011 | RANOC00142 |

| | | | | | | |
|---|---|---|---|---|---|---|
| PTPRK | RHSPN10190 | RGG049226 | RCABP00939 | RNAMR01200 | RBRBA00440 | RANOC02569 |
| PTX3 | RHSPN06273 | RGG103875 | RCABP03450 | RNAMR0006 | RBRBA01115 | RANOC00828 |
| PUM1 | RHSPN00566 | RGG148968 | RCABP03151 | RNAMR01689 | RBRBA02562 | RANOC02917 |
| PUM2 | RHSPN02977 | RGG053274 | RCABP00622 | RNAMR00513 | RBRBA02549 | RANOC00375 |
| PVRL1 | RHSPN16893 | RGG151942 | RCABP00122 | RNAMR02289 | RBRBA00139 | RANOC01848 |
| PWWP2A | RHSPN08851 | RGG118278 | RCABP02442 | RNAMR00250 | RBRBA01271 | RANOC02360 |
| PYCARD | RHSPN215783 | | RCABP01605 | RNAMR02233 | RBRBA03489 | RANOC02090 |
| RAB17 | RHSPN04817 | RGG086701 | RCABP005466 | | RBRBA01036 | RANOC00194 |
| RAB27A | RHSPN20484 | RGG106732 | RCABP02649 | RNAMR00140 | RBRBA035227 | |
| RABEP1 | RHSPN22279 | RGG138680 | RCABP01781 | RNAMR02343 | RBRBA03548 | RANOC02964 |
| RABGEF1 | RHSPN11044 | RGG139235 | RCABP00653 | RNAMR00610 | RBRBA02181 | RANOC01852 |
| RAC1 | RHSPN10601 | RGG123259 | RCABP03145 | RNAMR00654 | RBRBA03150 | RANOC02839 |
| RAC2 | RHSPN27303 | RGG006342 | RCABP01076 | RNAMR01170 | RBRBA03640 | RANOC01417 |
| RAD1 | RHSPN07962 | RGG171409 | RCABP02608 | RNAMR01473 | RBRBA02245 | RANOC02182 |
| RAD17 | RHSPN08160 | RGG177191 | RCABP04210 | RNAMR03894 | RBRBA00305 | RANOC00510 |
| RAD21 | RHSPN12687 | RGG039186 | RCABP03823 | RNAMR01629 | RBRBA00278 | RANOC01115 |
| RAD23A | RHSPN24574 | RGG163366 | RCABP00211 | RNAMR03847 | RBRBA03428 | RANOC00437 |
| RAD23B | RHSPN13676 | RGG176471 | | RNAMR01420 | RBRBA00544 | RANOC02068 |
| RAD50 | RHSPN08531 | RGG120385 | RCABP04172 | RNAMR0027 | RBRBA01304 | RANOC00460 |
| RAD51 | RHSPN20296 | RGG068703 | RCABP01447 | RNAMR02763 | RBRBA00603 | RANOC02279 |
| RAD51B | RHSPN19606 | RGG074221 | RCABP00679 | RNAMR00344 | RBRBA02951 | RANOC02298 |
| RAD51C | RHSPN23212 | RGG140293 | RCABP01599 | RNAMR0364 | RBRBA00352 | RANOC03460 |
| RAD51D | RHSPN22700 | RGG138988 | RCABP02022 | RNAMR0367 | RBRBA005972 | |
| RAD52 | RHSPN17062 | RGG007807 | RCABP02658 | RNAMR00368 | RBRBA01354 | RANOC01475 |
| RAD54B | RHSPN12553 | RGG038262 | RCABP01801 | RNAMR01620 | RBRBA03121 | RANOC01330 |
| RAD54L | RHSPN00873 | RGG097475 | RCABP03406 | RNAMR0390 | RBRBA01715 | RANOC01277 |
| RAD9A | RHSPN16360 | RGG068036 | RCABP02498 | RNAMR01833 | RBRBA01144 | RANOC00169 |
| RAE1 | RHSPN26390 | RGG144188 | RCABP01384 | RNAMR00703 | RBRBA03309 | RANOC02062 |
| RAF1 | RHSPN05032 | RGG114437 | RCABP00482 | RNAMR0365 | RBRBA01903 | RANOC00483 |
| RAG1 | RHSPN15898 | RGG072383 | RCABP03409 | RNAMR0181 | RBRBA02254 | RANOC00027 |
| RALGDS | RHSPN14037 | RGG132715 | RCABP00109 | RNAMR0151 | RBRBA02306 | RANOC03290 |
| RALGSL | | | | | | |
| RANBP17 | RHSPN088981 | | RCABP04140 | RNAMR00249 | RBRBA03140 | RANOC00367 |
| RAPGEF3 | RHSPN17505 | RGG166431 | RCABP01511 | RNAMR02920 | RBRBA02875 | RANOC02224 |
| RARA | RHSPN22842 | RGG159540 | RCABP01956 | RNAMR02553 | RBRBA00820 | RANOC01786 |
| RASGRP1 | RHSPN20263 | RGG074557 | RCABP01888 | RNAMR0276 | RBRBA03137 | RANOC00013 |
| RB1 | RHSPN18763 | RGG020224 | RCABP01884 | RNAMR01029 | RBRBA00993 | RANOC00059 |
| RBBP8 | RHSPN23816 | RGG035957 | RCABP02573 | RNAMR01924 | RBRBA00783 | RANOC01349 |
| RBM15 | RHSPN01388 | RGG155789 | RCABP03393 | RNAMR01977 | RBRBA00577 | RANOC01372 |
| RBPJ | RHSPN06849 | RGG065229 | RCABP03548 | RNAMR01324 | RBRBA02913 | RANOC02549 |
| RBPMS | RHSPN12177 | RGG060283 | RCABP04204 | RNAMR033736 | | RANOC02259 |
| RC3H1 | RHSPN02140 | RGG094586 | RCABP03502 | RNAMR01960 | RBRBA01586 | RANOC01233 |
| RC3H2 | RHSPN13827 | RGG133368 | RCABP04039 | RNAMR0157 | RBRBA033389 | |
| RDM1 | RHSPN22724 | RGG158163 | RCABP04103 | RNAMR025525 | | RANOC01773 |
| RECQL | RHSPN17327 | RGG008527 | RCABP00484 | RNAMR01188 | RBRBA01836 | RANOC03409 |

| | | | | | | |
|---|---|---|---|---|---|---|
| RECQL4 | RHSPN12929 | RGG040724 | RCABP02139 | RNAMR00379 | RBRBA01235 | RANOC02985 |
| RECQL5 | RHSPN23444 | RGG135388 | RCABP02311 | RNAMR02674 | RBRBA03787 | RANOC00610 |
| REL | RHSPN03323 | RGG041303 | RCABP03955 | RNAMR0333 | RBRBA03827 | RANOC02864 |
| RELA | RHSPN16305 | RGG169181 | RCABP03335 | RNAMR01847 | RBRBA011335 | |
| RELB | RHSPN252203 | | RCABP02774 | RNAMR0392 | RBRBA01017 | RANOC01873 |
| RELT | RHSPN16476 | RGG023651 | RCABP02627 | RNAMR02933 | RBRBA01513 | RANOC02520 |
| RET | RHSPN14530 | RGG079785 | RCABP01726 | RNAMR04013 | RBRBA00804 | RANOC02147 |
| REV1 | RHSPN03697 | RGG017573 | RCABP03835 | RNAMR01053 | RBRBA00350 | RANOC02745 |
| REV3L | RHSPN10089 | RGG050003 | RCABP02215 | RNAMR04064 | RBRBA03709 | RANOC00243 |
| RFC1 | RHSPN06898 | RGG064912 | RCABP00219 | RNAMR03637 | RBRBA02880 | RANOC01628 |
| RFX1 | RHSPN245980 | | RCABP01184 | RNAMR03853 | RBRBA03432 | RANOC00569 |
| RGCC | RHSPN18704 | RGG019846 | RCABP00875 | RNAMR01033 | RBRBA03045 | RANOC00852 |
| RGS1 | RHSPN02255 | RGG093351 | RCABP01574 | RNAMR02886 | RBRBA03230 | RANOC02288 |
| RHOA | RHSPN054379 | | RCABP038677 | | RBRBA01279 | RANOC00470 |
| RHOH | RHSPN06911 | RGG064832 | RCABP01324 | RNAMR0159 | RBRBA02883 | RANOC01550 |
| RICTOR | RHSPN08002 | RGG171706 | RCABP00525 | RNAMR01474 | RBRBA02244 | RANOC02179 |
| RIF1 | RHSPN04077 | RGG092171 | RCABP01238 | RNAMR03610 | RBRBA00482 | RANOC02427 |
| RIMBP2 | RHSPN18450 | RGG125808 | RCABP02160 | RNAMR010093 | | RANOC01862 |
| RIPK1 | RHSPN09091 | RGG033004 | RCABP02803 | RNAMR04170 | RBRBA01844 | RANOC02208 |
| RIPK2 | RHSPN12521 | RGG038008 | RCABP01895 | RNAMR00824 | RBRBA03266 | RANOC01198 |
| RMI2 | RHSPN21281 | RGG123293 | RCABP01118 | RNAMR030977 | | RANOC00597 |
| RNASE1 | RHSPN191328 | | RCABP01689 | RNAMR03107 | RBRBA024582 | |
| RNF135 | RHSPN226588 | | RCABP03466 | RNAMR03670 | RBRBA029079 | |
| RNF145 | RHSPN08842 | RGG119050 | RCABP02990 | RNAMR00257 | RBRBA01971 | RANOC01999 |
| RNF168 | RHSPN06558 | RGG100191 | RCABP00360 | RNAMR00080 | RBRBA02322 | RANOC00837 |
| RNF213 | RHSPN23567 | RGG136561 | RCABP02700 | RNAMR02710 | RBRBA03259 | RANOC00763 |
| RNF4 | RHSPN06662 | RGG066339 | RCABP01248 | RNAMR01344 | RBRBA04004 | RANOC01124 |
| RNF43 | RHSPN23203 | RGG138133 | RCABP04247 | RNAMR03640 | RBRBA00351 | RANOC03224 |
| RNF8 | RHSPN09605 | RGG045478 | RCABP04317 | RNAMR01270 | RBRBA02397 | RANOC00280 |
| RORA | RHSPN20540 | RGG106061 | RCABP02815 | RNAMR02180 | RBRBA00716 | RANOC02705 |
| ROS1 | RHSPN10129 | RGG049647 | RCABP02056 | RNAMR02627 | RBRBA004312 | |
| RPA1 | RHSPN22181 | RGG139414 | RCABP01899 | RNAMR02324 | RBRBA00343 | RANOC02630 |
| RPA2 | RHSPN00526 | RGG149109 | RCABP03491 | RNAMR01683 | RBRBA02568 | RANOC03078 |
| RPA3 | RHSPN10620 | RGG027847 | RCABP01161 | RNAMR02883 | RBRBA02694 | RANOC01665 |
| RPA4 | RHSPN280924 | | | | RBRBA016240 | |
| RPL10 | RHSPN285095 | | RCABP02511 | RNAMR04103 | RBRBA02798 | RANOC00447 |
| RPL11 | RHSPN00434 | RGG150753 | RCABP02013 | RNAMR01663 | RBRBA02587 | RANOC02825 |
| RPL22 | RHSPN00153 | RGG145028 | RCABP03683 | RNAMR0298 | RBRBA02467 | RANOC03138 |
| RPL5 | RHSPN01254 | RGG095753 | RCABP00523 | RNAMR00730 | RBRBA03089 | RANOC01147 |
| RPN1 | RHSPN06027 | RGG115443 | RCABP03071 | RNAMR04120 | RBRBA01493 | RANOC00487 |
| RPS25 | RHSPN16867 | RGG152503 | RCABP01682 | RNAMR02283 | RBRBA00142 | RANOC03280 |
| RPS6KA3 | RHSPN27689 | RGG015973 | RCABP00970 | RNAMR03223 | RBRBA02024 | RANOC00980 |
| RPS8 | RHSPN00833 | RGG097271 | RCABP02351 | RNAMR03890 | RBRBA03383 | RANOC01277 |
| RPTOR | RHSPN23575 | RGG136543 | RCABP02706 | RNAMR02694 | RBRBA03259 | RANOC00763 |
| RRM2B | RHSPN12628 | RGG038735 | RCABP04369 | RNAMR01633 | RBRBA03132 | RANOC01190 |

| | | | | | | |
|---|---|---|---|---|---|---|
| RSAD2 | RHSPN0289[illegible] | RGG052651 | RCABP0405[illegible] | RNAMR0343[illegible] | RBRBA0153[illegible] | RANOC0005[illegible] |
| RSPO2 | RHSPN1266[illegible] | RGG039049 | RCABP0298[illegible] | RNAMR0163[illegible] | RBRBA0174[illegible] | RANOC0118[illegible] |
| RSPO3 | RHSPN1017[illegible] | RGG049342 | RCABP0158[illegible] | RNAMR0120[illegible] | RBRBA0043[illegible] | RANOC0303[illegible] |
| RTEL1 | RHSPN2648[illegible] | RGG143270 | RCABP0393[illegible] | RNAMR0072[illegible] | RBRBA0395[illegible] | RANOC0265[illegible] |
| RYR3 | RHSPN2023[illegible] | RGG074523 | RCABP0379[illegible] | RNAMR0278[illegible] | RBRBA0398[illegible] | RANOC0032[illegible] |
| S1PR4 | RHSPN2428[illegible] | RGG160718 | RCABP0001[illegible] | RNAMR0319[illegible] | RBRBA0041[illegible] | RANOC0305[illegible] |
| SAMHD1 | RHSPN2616[illegible] | RGG141468 | RCABP0198[illegible] | RNAMR0338[illegible] | RBRBA0168[illegible] | RANOC0117[illegible] |
| SAMSN1 | RHSPN2657[illegible] | RGG013587 | RCABP0177[illegible] | RNAMR0083[illegible] | RBRBA0102[illegible] | RANOC0212[illegible] |
| SARM1 | RHSPN2259[illegible] | RGG139547 | RCABP0058[illegible] | RNAMR0230[illegible] | RBRBA0281[illegible] | RANOC0277[illegible] |
| SASH3 | RHSPN2829[illegible] | RGG055311 | RCABP0017[illegible] | RNAMR0261[illegible] | RBRBA0226[illegible] | RANOC0273[illegible] |
| SBDS | RHSPN1104[illegible] | RGG138283 | RCABP0064[illegible] | RNAMR0061[illegible] | RBRBA021816 | |
| SBNO2 | RHSPN2421[illegible] | RGG161675 | RCABP0001[illegible] | RNAMR0320[illegible] | RBRBA014573 | |
| SDC4 | RHSPN2625[illegible] | RGG142234 | RCABP0401[illegible] | RNAMR0242[illegible] | RBRBA0219[illegible] | RANOC0131[illegible] |
| SDHAF2 | RHSPN1614[illegible] | RGG068198 | RCABP0082[illegible] | RNAMR0403[illegible] | RBRBA021985 | |
| SDHB | RHSPN0032[illegible] | RGG144760 | RCABP0183[illegible] | RNAMR0164[illegible] | RBRBA0022[illegible] | RANOC0336[illegible] |
| SDHC | RHSPN0200[illegible] | RGG154540 | RCABP0242[illegible] | RNAMR0247[illegible] | RBRBA0327[illegible] | RANOC0321[illegible] |
| SDHD | RHSPN1677[illegible] | RGG152645 | RCABP0187[illegible] | RNAMR0226[illegible] | RBRBA001542 | |
| SEMA3A | RHSPN1116[illegible] | RGG001432 | RCABP0308[illegible] | RNAMR0058[illegible] | RBRBA0322[illegible] | RANOC0143[illegible] |
| SEMA3C | RHSPN1115[illegible] | RGG001501 | RCABP0313[illegible] | RNAMR0058[illegible] | RBRBA0322[illegible] | RANOC0144[illegible] |
| SEMA4A | RHSPN018523 | | RCABP0224[illegible] | RNAMR0372[illegible] | RBRBA015720 | |
| SEMA4D | RHSPN1350[illegible] | RGG175382 | RCABP0046[illegible] | RNAMR0174[illegible] | RBRBA0361[illegible] | RANOC0073[illegible] |
| SEMA6A | RHSPN0844[illegible] | RGG178095 | RCABP0197[illegible] | RNAMR0022[illegible] | RBRBA0021[illegible] | RANOC0189[illegible] |
| SEMA7A | RHSPN2071[illegible] | RGG105423 | RCABP0293[illegible] | RNAMR0215[illegible] | RBRBA0358[illegible] | RANOC0336[illegible] |
| SERINC3 | RHSPN2623[illegible] | RGG142326 | RCABP0400[illegible] | RNAMR0242[illegible] | RBRBA0206[illegible] | RANOC0117[illegible] |
| SERINC5 | RHSPN0825[illegible] | RGG172676 | RCABP0379[illegible] | RNAMR0077[illegible] | RBRBA0031[illegible] | RANOC0041[illegible] |
| SERPINE1 | RHSPN113505 | | RCABP0396[illegible] | RNAMR0062[illegible] | RBRBA0310[illegible] | RANOC0346[illegible] |
| SERPING1 | RHSPN1606[illegible] | RGG071779 | RCABP0077[illegible] | RNAMR0358[illegible] | RBRBA0385[illegible] | RANOC0016[illegible] |
| SERPINH1 | RHSPN1651[illegible] | RGG023702 | RCABP0262[illegible] | RNAMR0293[illegible] | RBRBA0150[illegible] | RANOC0240[illegible] |
| SET | RHSPN1394[illegible] | RGG132135 | RCABP0274[illegible] | RNAMR0152[illegible] | RBRBA034059 | |
| SETBP1 | RHSPN2394[illegible] | RGG170541 | RCABP0407[illegible] | RNAMR0211[illegible] | RBRBA0133[illegible] | RANOC0010[illegible] |
| SETD2 | RHSPN0536[illegible] | RGG025078 | RCABP0430[illegible] | RNAMR0013[illegible] | RBRBA0152[illegible] | RANOC0318[illegible] |
| SETMAR | RHSPN049423 | | RCABP0152[illegible] | RNAMR0203[illegible] | RBRBA0346[illegible] | RANOC0049[illegible] |
| SF3B1 | RHSPN0439[illegible] | RGG087675 | RCABP0327[illegible] | RNAMR0377[illegible] | RBRBA0237[illegible] | RANOC0271[illegible] |
| SFTPD | RHSPN148770 | | RCABP0172[illegible] | RNAMR0338[illegible] | RBRBA0214[illegible] | RANOC0214[illegible] |
| SGK1 | RHSPN1023[illegible] | RGG048823 | RCABP0161[illegible] | RNAMR0408[illegible] | RBRBA0045[illegible] | RANOC0196[illegible] |
| SGK2 | RHSPN2622[illegible] | RGG142004 | RCABP0422[illegible] | RNAMR0242[illegible] | RBRBA0206[illegible] | RANOC0258[illegible] |
| SGPP2 | RHSPN0467[illegible] | RGG100905 | RCABP0327[illegible] | RNAMR0305[illegible] | RBRBA0367[illegible] | RANOC0083[illegible] |
| SGSH | RHSPN2356[illegible] | RGG136589 | RCABP0270[illegible] | RNAMR0269[illegible] | RBRBA0325[illegible] | RANOC0045[illegible] |
| SH2B2 | RHSPN1136[illegible] | RGG138787 | RCABP0097[illegible] | RNAMR0062[illegible] | RBRBA0310[illegible] | RANOC0296[illegible] |
| SH2B3 | RHSPN1823[illegible] | RGG126631 | RCABP0299[illegible] | RNAMR0098[illegible] | RBRBA0272[illegible] | RANOC0269[illegible] |
| SH2D1A | RHSPN2827[illegible] | RGG057927 | RCABP0366[illegible] | RNAMR0261[illegible] | RBRBA022634 | |
| SH2D4A | RHSPN1205[illegible] | RGG060010 | RCABP0264[illegible] | RNAMR0143[illegible] | RBRBA001063 | |
| SH3GL1 | RHSPN2431[illegible] | RGG161374 | RCABP0000[illegible] | RNAMR0318[illegible] | RBRBA004103 | |
| SHFM1 | RHSPN1125[illegible] | RGG027782 | RCABP039044 | | RBRBA009521 | |
| SHPRH | RHSPN1031[illegible] | RGG047572 | RCABP0025[illegible] | RNAMR0121[illegible] | RBRBA0370[illegible] | RANOC0039[illegible] |

| | | | | | | |
|---|---|---|---|---|---|---|
| SIGIRR | RHSPN1547 | RGG068812 | RCABP0053 | RNAMR0178 | RBRBA0047 | RANOC0015 |
| SIKE1 | RHSPN0145 | RGG156961 | RCABP0394 | RNAMR0198 | RBRBA0056 | RANOC0129 |
| SIN3A | RHSPN2075 | RGG105531 | RCABP0292 | RNAMR0381 | RBRBA0357 | RANOC0280 |
| SIRT1 | RHSPN1471 | RGG079941 | RCABP0252 | RNAMR0132 | RBRBA0279 | RANOC0205 |
| SIRT2 | RHSPN2502 | RGG168790 | RCABP0081 | RNAMR0251 | RBRBA0401 | RANOC0317 |
| SIRT3 | RHSPN1546 | RGG068858 | RCABP0294 | RNAMR0178 | RBRBA0131 | RANOC0002 |
| SIRT4 | RHSPN1832 | RGG128088 | RCABP0184 | RNAMR0097 | RBRBA0107 | RANOC0252 |
| SIRT5 | RHSPN0918 | RGG032301 | RCABP0087 | RNAMR0137 | RBRBA0301 | RANOC0113 |
| SIRT6 | RHSPN2431 | RGG161448 | RCABP0000 | RNAMR0318 | RBRBA0041 | RANOC0348 |
| SIRT7 | RHSPN2362 | RGG136878 | RCABP0375 | RNAMR0270 | RBRBA0370 | RANOC0062 |
| SIVA1 | RHSPN2001 | RGG077548 | RCABP0414 | RNAMR0029 | RBRBA0008 | RANOC0010 |
| SKAP1 | RHSPN2307 | RGG159074 | RCABP0218 | RNAMR0256 | RBRBA0083 | RANOC0179 |
| SLAMF1 | RHSPN0196 | RGG153309 | RCABP0223 | RNAMR0247 | RBRBA002705 | |
| SLC11A1 | RHSPN0460 | RGG090431 | not predicte | RNAMR0306 | RBRBA0366 | RANOC0015 |
| SLC11A2 | RHSPN1759 | RGG167432 | RCABP0176 | RNAMR0290 | RBRBA0179 | RANOC0055 |
| SLC13A1 | RHSPN1150 | RGG002574 | RCABP0414 | RNAMR0225 | RBRBA0307 | RANOC0152 |
| SLC13A2 | RHSPN2259 | RGG139555 | RCABP0211 | RNAMR0230 | RBRBA0281 | RANOC0277 |
| SLC13A3 | RHSPN2629 | RGG142411 | RCABP0401 | RNAMR0241 | RBRBA0085 | RANOC0139 |
| SLC13A4 | RHSPN1162 | RGG007403 | RCABP0260 | RNAMR0173 | RBRBA0065 | RANOC0295 |
| SLC13A5 | RHSPN2230 | RGG141172 | RCABP0065 | RNAMR0233 | RBRBA004249 | |
| SLC25A21 | RHSPN1936 | RGG075237 | RCABP020503 | | RBRBA0122 | RANOC0010 |
| SLC26A6 | RHSPN0540 | RGG115397 | RCABP0307 | RNAMR000073 | | RANOC0079 |
| SLC34A2 | RHSPN0684 | RGG065249 | RCABP0355 | RNAMR0132 | RBRBA0291 | RANOC0254 |
| SLC39A10 | RHSPN0438 | RGG087583 | RCABP0327 | RNAMR0350 | RBRBA0017 | RANOC0233 |
| SLC45A3 | RHSPN0240 | RGG156318 | RCABP0227 | RNAMR0281 | RBRBA0391 | RANOC0211 |
| SLC9A3R2 | RHSPN2112 | RGG122458 | RCABP0358 | RNAMR0397 | RBRBA018059 | |
| SLX1A | RHSPN2153 | RGG099212 | RCABP0157 | RNAMR0221 | RBRBA0347 | RANOC0232 |
| SLX4 | RHSPN2120 | RGG123933 | RCABP0129 | RNAMR0308 | RBRBA0229 | RANOC0283 |
| SMAD3 | RHSPN2062 | RGG108514 | RCABP0373 | RNAMR0216 | RBRBA008453 | |
| SMAD4 | RHSPN2400 | RGG169320 | RCABP0094 | RNAMR0212 | RBRBA0139 | RANOC0277 |
| SMARCA2 | RHSPN1297 | RGG173297 | RCABP0311 | RNAMR0043 | RBRBA0248 | RANOC0054 |
| SMARCA4 | RHSPN2450 | RGG168942 | RCABP0118 | RNAMR0316 | RBRBA007513 | |
| SMARCD1 | RHSPN1758 | RGG166386 | RCABP0136 | RNAMR0290 | RBRBA0286 | RANOC0058 |
| SMARCE1 | RHSPN2284 | RGG159572 | RCABP0195 | RNAMR0259 | RBRBA0081 | RANOC0168 |
| SMO | RHSPN1156 | RGG000441 | RCABP0123 | RNAMR0224 | RBRBA0175 | RANOC0300 |
| SMUG1 | RHSPN1769 | RGG166565 | RCABP0177 | RNAMR0388 | RBRBA0177 | RANOC0056 |
| SNAP23 | RHSPN2033 | RGG073638 | RCABP0128 | RNAMR0277 | RBRBA0061 | RANOC0001 |
| SNCA | RHSPN0724 | RGG060423 | RCABP0134 | RNAMR014178 | | RANOC0142 |
| SNRPC | RHSPN0954 | RGG157138 | RCABP0228 | RNAMR0128 | RBRBA0345 | RANOC0116 |
| SNX4 | RHSPN0598 | RGG091270 | RCABP0240 | RNAMR0013 | RBRBA0231 | RANOC0003 |
| SOCS1 | RHSPN2127 | RGG123298 | RCABP0111 | RNAMR0309 | RBRBA0213 | RANOC0341 |
| SOCS2 | RHSPN1802 | RGG004681 | RCABP0069 | RNAMR0110 | RBRBA0282 | RANOC0141 |
| SOCS3 | RHSPN2352 | RGG136768 | RCABP0286 | RNAMR0269 | RBRBA0377 | RANOC0060 |
| SOCS5 | RHSPN0324 | RGG045221 | RCABP0130 | RNAMR0055 | RBRBA0120 | RANOC0064 |
| SOCS6 | RHSPN2411 | RGG034939 | RCABP0290 | RNAMR0280 | RBRBA0077 | RANOC0113 |

| | | | | | | |
|---|---|---|---|---|---|---|
| SORCS1 | RHSPN15195 | RGG083913 | RCABP00581 | RNAMR0326 | RBRBA029868 | |
| SORCS2 | RHSPN0672 | RGG065998 | RCABP02033 | RNAMR0134 | RBRBA04066 | RANOC0124 |
| SPECC1 | RHSPN2255 | RGG139996 | RCABP00155 | RNAMR0368 | RBRBA019246 | |
| SPEN | RHSPN0029 | RGG146175 | RCABP01836 | RNAMR0164 | RBRBA0404 | RANOC0296 |
| SPG21 | RHSPN2058 | RGG108301 | RCABP03891 | RNAMR0217 | RBRBA0071 | RANOC0263 |
| SPNS2 | RHSPN2224 | RGG138551 | RCABP0177 | RNAMR0233 | RBRBA017644 | |
| SPON2 | RHSPN0662 | RGG066649 | RCABP0284 | RNAMR0135 | RBRBA04071 | RANOC0273 |
| SPOP | RHSPN2310 | RGG158790 | RCABP02185 | RNAMR0257 | RBRBA0164 | RANOC0169 |
| SPPL2A | RHSPN2045 | RGG107317 | RCABP02106 | RNAMR0015 | RBRBA01092 | RANOC0308 |
| SPRTN | RHSPN0268 | RGG046779 | RCABP0148 | RNAMR0207 | RBRBA00085 | RANOC0027 |
| SQSTM1 | RHSPN0902 | RGG119519 | RCABP00455 | RNAMR0028 | RBRBA01218 | RANOC0072 |
| SRC | RHSPN2617 | RGG142189 | RCABP04014 | RNAMR0338 | RBRBA01681 | RANOC0238 |
| SRGAP3 | RHSPN0496 | RGG117159 | RCABP01532 | RNAMR0013 | RBRBA0222 | RANOC0230 |
| SRSF2 | RHSPN2349 | RGG135011 | RCABP0230 | RNAMR0268 | RBRBA01801 | RANOC0045 |
| SRSF3 | RHSPN0958 | RGG155908 | RCABP03388 | RNAMR0127 | RBRBA03448 | RANOC0116 |
| SS18 | RHSPN2384 | RGG036188 | RCABP0286 | RNAMR0192 | RBRBA0018 | RANOC0112 |
| SS18L1 | RHSPN2643 | RGG142765 | RCABP03157 | RNAMR0070 | RBRBA03966 | RANOC0118 |
| SSPN | RHSPN1736 | RGG008854 | RCABP04004 | RNAMR0118 | RBRBA0215 | RANOC0143 |
| SSX1 | RHSPN278197 | | | RNAMR0380 | RBRBA012709 | |
| SSX2 | | | | | | |
| SSX4 | | | | | | |
| ST3GAL3 | RHSPN0080 | RGG097098 | RCABP01686 | RNAMR0379 | RBRBA01446 | RANOC0127 |
| ST6GAL1 | RHSPN064767 | | RCABP02807 | RNAMR0009 | RBRBA02728 | RANOC0091 |
| STAG2 | RHSPN2827 | RGG057950 | RCABP03665 | RNAMR0261 | RBRBA02262 | RANOC0266 |
| STAP1 | RHSPN0704 | RGG062413 | RCABP01983 | RNAMR0159 | RBRBA017442 | |
| STAT1 | RHSPN0436 | RGG087480 | RCABP03647 | RNAMR0350 | RBRBA02232 | RANOC0046 |
| STAT3 | RHSPN2289 | RGG159838 | RCABP00503 | RNAMR0254 | RBRBA0264 | RANOC0183 |
| STAT5B | RHSPN2289 | RGG159817 | RCABP00503 | RNAMR0254 | RBRBA0265 | RANOC0178 |
| STAT6 | RHSPN1777 | RGG167132 | RCABP0165 | RNAMR0273 | RBRBA0309 | RANOC0204 |
| STK11 | RHSPN2421 | RGG161688 | RCABP0002 | RNAMR0320 | RBRBA01458 | RANOC0319 |
| STK24 | RHSPN0080 | RGG097098 | RCABP01686 | RNAMR0379 | RBRBA01446 | RANOC0127 |
| STK3 | RHSPN1258 | RGG038543 | | RNAMR0162 | RBRBA03127 | RANOC0111 |
| STK4 | RHSPN2624 | RGG142262 | RCABP04015 | RNAMR024265 | | RANOC0117 |
| STOML2 | RHSPN1318 | RGG171017 | RCABP02987 | RNAMR0399 | RBRBA0374 | RANOC0312 |
| STRN | RHSPN0315 | RGG045919 | RCABP00468 | RNAMR0053 | RBRBA04028 | RANOC0028 |
| STUB1 | RHSPN2106 | RGG124180 | RCABP02471 | RNAMR0398 | RBRBA01948 | RANOC0325 |
| STX7 | RHSPN1021 | RGG049017 | RCABP01826 | RNAMR0121 | RBRBA00447 | RANOC0197 |
| STXBP1 | RHSPN1388 | RGG131665 | RCABP02752 | RNAMR0154 | RBRBA034146 | |
| STXBP2 | RHSPN244055 | | RCABP01825 | RNAMR0314 | RBRBA01216 | RANOC0060 |
| STXBP3 | RHSPN0134 | RGG092975 | RCABP03287 | RNAMR0196 | RBRBA00583 | RANOC0203 |
| SUCLA2 | RHSPN1875 | RGG020193 | RCABP01884 | RNAMR0102 | RBRBA00754 | RANOC0005 |
| SUFU | RHSPN1515 | RGG083595 | RCABP02562 | RNAMR0325 | RBRBA02994 | RANOC0084 |
| SULT1A1 | RHSPN214761 | | RCABP02644 | RNAMR0221 | RBRBA02205 | RANOC0237 |
| SUMF1 | RHSPN0494 | RGG117058 | RCABP01528 | RNAMR0203 | RBRBA03468 | RANOC0049 |
| SUSD4 | RHSPN0257 | RGG044120 | RCABP01308 | RNAMR0335 | RBRBA01431 | RANOC0009 |

| | | | | | | |
|---|---|---|---|---|---|---|
| SUV39H1 | RHSPN278326 | | RCABP04268 | RNAMR0380 | RBRBA01622 | RANOC0043 |
| SUZ12 | RHSPN22670 | RGG135723 | RCABP03466 | RNAMR0367 | RBRBA02907 | RANOC0195 |
| SVEP1 | RHSPN13700 | RGG177495 | RCABP03792 | RNAMR0142 | RBRBA00540 | RANOC0278 |
| SWAP70 | RHSPN15702 | RGG070017 | RCABP01883 | RNAMR0178 | RBRBA02424 | RANOC0002 |
| SYK | RHSPN1350 | RGG175421 | RCABP01740 | RNAMR0361 | RBRBA03618 | RANOC0052 |
| SYNCRIP | RHSPN0993 | RGG051140 | RCABP03424 | RNAMR0263 | RBRBA00769 | RANOC0006 |
| SYNE1 | RHSPN1036 | RGG047833 | RCABP00529 | RNAMR0229 | RBRBA03845 | RANOC0226 |
| SYT13 | RHSPN1593 | RGG072565 | RCABP00058 | RNAMR0180 | RBRBA01996 | RANOC0012 |
| TAB2 | RHSPN1033 | RGG047664 | RCABP04413 | RNAMR0229 | RBRBA03839 | RANOC0226 |
| TAC1 | RHSPN1126 | RGG027799 | RCABP01004 | RNAMR0288 | RBRBA01608 | RANOC0167 |
| TAF15 | RHSPN22718 | RGG138941 | RCABP02021 | RNAMR0367 | RBRBA005998 | |
| TAL1 | RHSPN0089 | RGG097562 | RCABP03403 | RNAMR0353 | RBRBA03856 | RANOC0221 |
| TANK | RHSPN0414 | RGG090009 | RCABP03271 | RNAMR0019 | RBRBA00493 | RANOC0259 |
| TAPBP | RHSPN0951 | RGG130230 | RCABP01907 | RNAMR0129 | RBRBA016175 | |
| TAZ | RHSPN285133 | | RCABP02481 | RNAMR0410 | RBRBA04031 | RANOC0044 |
| TBK1 | RHSPN1784 | RGG003560 | RCABP00720 | RNAMR0272 | RBRBA02260 | RANOC0142 |
| TBL1XR1 | RHSPN0637 | RGG102995 | RCABP00149 | RNAMR0007 | RBRBA03609 | RANOC0089 |
| TBXAS1 | RHSPN1165 | RGG007092 | RCABP01329 | RNAMR0173 | RBRBA02554 | RANOC0148 |
| TCEA1 | RHSPN1233 | RGG036664 | RCABP00672 | RNAMR0036 | RBRBA00207 | RANOC0219 |
| TCF12 | RHSPN2050 | RGG106613 | RCABP00620 | RNAMR0218 | RBRBA01498 | RANOC0292 |
| TCF3 | RHSPN2424 | RGG161292 | RCABP02857 | RNAMR0320 | RBRBA014622 | |
| TCF7 | RHSPN085540 | | RCABP01466 | RNAMR0026 | RBRBA013080 | |
| TCF7L2 | RHSPN1523 | RGG084194 | RCABP02369 | RNAMR0050 | RBRBA02496 | RANOC0199 |
| TDG | RHSPN1813 | RGG006941 | RCABP01368 | RNAMR0109 | RBRBA00793 | RANOC0140 |
| TDO2 | RHSPN0761 | RGG058661 | RCABP01069 | RNAMR0346 | RBRBA00669 | RANOC0145 |
| TDP1 | RHSPN1980 | RGG076252 | RCABP01550 | RNAMR0035 | RBRBA02059 | RANOC0000 |
| TDP2 | RHSPN0923 | RGG034768 | RCABP02907 | RNAMR0137 | RBRBA01762 | RANOC0113 |
| TEC | RHSPN0695 | RGG064486 | RCABP04152 | RNAMR0158 | RBRBA00063 | RANOC0238 |
| TEK | RHSPN1311 | RGG177759 | RCABP01503 | RNAMR0044 | RBRBA03865 | RANOC0054 |
| TENM4 | RHSPN1656 | RGG022915 | RCABP00765 | RNAMR0357 | RBRBA03509 | RANOC0251 |
| TEP1 | RHSPN1911 | RGG168057 | RCABP03860 | RNAMR0311 | RBRBA01385 | RANOC0208 |
| TERF1 | RHSPN1243 | RGG037456 | RCABP02453 | RNAMR0081 | RBRBA01416 | RANOC0120 |
| TERF2 | RHSPN2188 | RGG112626 | RCABP04380 | RNAMR0090 | RBRBA03383 | RANOC0299 |
| TERT | RHSPN0783 | RGG034392 | RCABP03954 | RNAMR0112 | RBRBA00288 | RANOC0122 |
| TESPA1 | RHSPN1770 | RGG166458 | RCABP04311 | RNAMR0413 | RBRBA01100 | RANOC0222 |
| TET1 | RHSPN1473 | RGG080700 | RCABP02521 | RNAMR0122 | RBRBA02796 | RANOC0205 |
| TET2 | RHSPN0731 | RGG060726 | RCABP02433 | RNAMR0323 | RBRBA02229 | RANOC0260 |
| TFE3 | RHSPN278461 | | RCABP04269 | RNAMR0380 | RBRBA01164 | RANOC0248 |
| TFEB | RHSPN096467 | | RCABP00419 | RNAMR0127 | RBRBA03991 | RANOC0116 |
| TFG | RHSPN0577 | RGG012146 | RCABP00924 | RNAMR0240 | RBRBA03247 | RANOC0101 |
| TFPT | RHSPN255906 | | RCABP02322 | RNAMR0328 | RBRBA03217 | RANOC0233 |
| TGFA | RHSPN0340 | RGG148376 | RCABP00389 | RNAMR0161 | RBRBA015963 | |
| TGFB1 | RHSPN250973 | | RCABP02559 | RNAMR0390 | RBRBA01726 | RANOC0188 |
| TGFB2 | RHSPN0253 | RGG044297 | RCABP01717 | RNAMR0334 | RBRBA03207 | RANOC0008 |
| TGFB3 | RHSPN1972 | RGG075617 | RCABP03129 | RNAMR0032 | RBRBA03695 | RANOC0001 |

| | | | | | | |
|---|---|---|---|---|---|---|
| TH | RHSPN15545 | RGG070758 | RCABP00444 | RNAMR018194 | | RANOC00029 |
| THBS1 | RHSPN20271 | RGG074481 | RCABP03177 | RNAMR0276 | RBRBA005996 | |
| THEMIS | RHSPN10188 | RGG049235 | RCABP00938 | RNAMR01200 | RBRBA00439 | RANOC02569 |
| THOC1 | RHSPN23675 | RGG035844 | RCABP02264 | RNAMR01927 | RBRBA00786 | RANOC01127 |
| THRAP3 | RHSPN00661 | RGG150081 | RCABP03960 | RNAMR01704 | RBRBA01702 | RANOC02794 |
| THSD7B | RHSPN04021 | RGG091749 | RCABP02692 | RNAMR00222 | RBRBA00359 | RANOC00179 |
| THY1 | RHSPN16891 | RGG151953 | RCABP00122 | RNAMR02289 | RBRBA00139 | RANOC03345 |
| TICAM1 | RHSPN24330 | RGG162734 | RCABP00004 | RNAMR03179 | RBRBA00408 | RANOC03013 |
| TICAM2 | RHSPN084370 | | RCABP019738 | | RBRBA00217 | RANOC01907 |
| TINAGL1 | RHSPN00575 | RGG150405 | RCABP03193 | RNAMR0169 | RBRBA03003 | RANOC02431 |
| TIRAP | RHSPN16981 | RGG151238 | RCABP04255 | RNAMR02868 | RBRBA02014 | RANOC02468 |
| TKFC | RHSPN16132 | RGG068164 | RCABP00825 | RNAMR0404 | RBRBA02196 | RANOC02150 |
| TLK2 | RHSPN23259 | RGG158295 | RCABP03445 | RNAMR03463 | RBRBA00388 | RANOC01777 |
| TLR1 | RHSPN06889 | RGG065018 | RCABP00077 | RNAMR03377 | RBRBA02879 | RANOC02572 |
| TLR10 | RHSPN068870 | | not predicted | | RBRBA016480 | |
| TLR13 | | | | | | |
| TLR14 | | | | | | |
| TLR15 | | | | | | |
| TLR2 | RHSPN07598 | RGG058531 | RCABP01069 | RNAMR03528 | RBRBA02094 | RANOC01452 |
| TLR21 | | | | | | |
| TLR3 | RHSPN07778 | RGG063671 | RCABP02875 | RNAMR01914 | RBRBA00974 | RANOC01445 |
| TLR4 | RHSPN13778 | RGG131779 | RCABP02747 | RNAMR01557 | RBRBA03343 | RANOC02735 |
| TLR5 | RHSPN02571 | RGG044133 | RCABP01866 | RNAMR03350 | RBRBA01431 | RANOC00090 |
| TLR6 | RHSPN068903 | | not predicted | RNAMR03377 | RBRBA028796 | |
| TLR7 | RHSPN27620 | RGG016573 | RCABP01369 | RNAMR03710 | RBRBA03344 | RANOC00864 |
| TLR8 | RHSPN276215 | | RCABP02584 | RNAMR03703 | RBRBA033442 | |
| TLR9 | RHSPN055228 | | not predicted | RNAMR00123 | RBRBA034694 | |
| TLX1 | RHSPN15115 | RGG083493 | RHSPN15115 | RNAMR03243 | RBRBA03000 | RANOC02179 |
| TLX3 | RHSPN08899 | RGG117931 | RCABP04140 | RNAMR002493 | | RANOC00052 |
| TMEM151B | RHSPN09727 | RGG045514 | RCABP04318 | RNAMR01253 | RBRBA00696 | RANOC00383 |
| TMEM173 | RHSPN08626 | RGG117478 | RCABP00942 | RNAMR00794 | RBRBA03361 | RANOC01140 |
| TMEM2 | RHSPN13373 | RGG174421 | RCABP02784 | RNAMR00424 | RBRBA01662 | RANOC00422 |
| TMEM38A | RHSPN24661 | RGG162529 | RCABP02255 | RNAMR03933 | RBRBA001261 | |
| TMPRSS6 | RHSPN27300 | RGG006388 | RCABP01077 | RNAMR01177 | RBRBA03640 | RANOC03409 |
| TNF | RHSPN094336 | | RCABP00880 | RNAMR0131 | RBRBA00555 | RANOC00497 |
| TNFAIP1 | RHSPN22592 | RGG140988 | RCABP01099 | RNAMR02300 | RBRBA02812 | RANOC03322 |
| TNFAIP8L2 | RHSPN016722 | | RCABP01265 | RNAMR0201 | RBRBA03296 | RANOC03018 |
| TNFRSF11A | RHSPN24081 | RGG033257 | RCABP01893 | RNAMR02454 | RBRBA03458 | RANOC02285 |
| TNFRSF11B | RHSPN12697 | RGG039235 | RCABP00901 | RNAMR01630 | RBRBA03997 | RANOC02290 |
| TNFRSF12A | RHSPN211701 | | not predicted | RNAMR03078 | RBRBA00986 | RANOC02954 |
| TNFRSF13B | RHSPN22473 | RGG121957 | RCABP03590 | RNAMR03487 | RBRBA01917 | RANOC02625 |
| TNFRSF14 | RHSPN001111 | | RCABP03686 | RNAMR0300 | RBRBA00558 | RANOC03168 |
| TNFRSF17 | RHSPN212892 | | RCABP00063 | RNAMR02194 | RBRBA021368 | |
| TNFRSF18 | RHSPN00034 | RGG145672 | not predicted | RNAMR0300 | RBRBA016746 | |
| TNFRSF1A | RHSPN17120 | RGG011172 | RCABP00574 | RNAMR03544 | RBRBA02392 | RANOC02779 |

| | | | | | | |
|---|---|---|---|---|---|---|
| TNFRSF1B | RHSPN00247 | RGG146606 | RCABP00734 | RNAMR0296 | RBRBA023601 | |
| TNFRSF21 | RHSPN09755 | RGG054783 | RCABP00933 | RNAMR0125 | RBRBA00691 | RANOC0021 |
| TNFRSF4 | RHSPN00035 | RGG145666 | RCABP01150 | RNAMR0300 | RBRBA020743 | |
| TNFRSF6B | RHSPN26489 | RGG143259 | RCABP039344 | | RBRBA03957 | RANOC0265 |
| TNFRSF8 | RHSPN00245 | RGG146615 | RCABP00734 | RNAMR0300 | RBRBA023599 | |
| TNFRSF9 | RHSPN00181 | RGG144848 | RCABP03694 | RNAMR0300 | RBRBA024710 | |
| TNFSF10 | RHSPN06360 | RGG103109 | RCABP01482 | RNAMR0007 | RBRBA00683 | RANOC0104 |
| TNFSF11 | RHSPN18712 | RGG019893 | RCABP04123 | RNAMR0103 | RBRBA00557 | RANOC0085 |
| TNFSF13B | RHSPN19017 | RGG018557 | RCABP02440 | RNAMR0247 | RBRBA03810 | RANOC0005 |
| TNFSF14 | RHSPN243804 | | RCABP00217 | RNAMR0317 | RBRBA00764 | RANOC0056 |
| TNFSF15 | RHSPN13765 | RGG131706 | RCABP02760 | RNAMR0155 | RBRBA03342 | RANOC0282 |
| TNFSF8 | RHSPN13765 | RGG131710 | RCABP02760 | RNAMR0155 | RBRBA03342 | RANOC0282 |
| TNFSF9 | RHSPN243787 | | not predicted | RNAMR0321 | RBRBA038050 | |
| TNIP1 | RHSPN08782 | RGG119376 | RCABP00450 | RNAMR0027 | RBRBA01962 | RANOC0064 |
| TNIP2 | RHSPN06665 | RGG066323 | RCABP01247 | RNAMR0134 | RBRBA040690 | |
| TNIP3 | RHSPN07420 | RGG062711 | RCABP03574 | RNAMR0210 | RBRBA01624 | RANOC0158 |
| TOLLIP | RHSPN15519 | RGG071006 | RCABP00201 | RNAMR0177 | RBRBA00471 | RANOC0014 |
| TOMM40 | RHSPN252159 | | RCABP04299 | RNAMR0392 | RBRBA01017 | RANOC0312 |
| TOP1 | RHSPN26200 | RGG142087 | RCABP04010 | RNAMR0400 | RBRBA02068 | RANOC0238 |
| TOP2A | RHSPN22843 | RGG159547 | RCABP01958 | RNAMR0259 | RBRBA00819 | RANOC0168 |
| TOP3B | RHSPN27025 | RGG125216 | RCABP00090 | RNAMR0094 | RBRBA02619 | RANOC0234 |
| TOPBP1 | RHSPN06093 | RGG029997 | RCABP01782 | RNAMR0003 | RBRBA01042 | RANOC0210 |
| TOX | RHSPN12356 | RGG036940 | RCABP03148 | RNAMR0082 | RBRBA02125 | RANOC0120 |
| TP53 | RHSPN223492 | | RCABP02458 | RNAMR0235 | RBRBA030625 | |
| TP53BP1 | RHSPN20358 | RGG109192 | RCABP04130 | RNAMR0277 | RBRBA02900 | RANOC0333 |
| TP53BP1 | RHSPN20358 | RGG109192 | RCABP04130 | RNAMR0277 | RBRBA02900 | RANOC0333 |
| TP63 | RHSPN06496 | RGG102052 | RCABP00909 | RNAMR0008 | RBRBA033216 | |
| TP73 | RHSPN00131 | RGG145124 | RCABP03681 | RNAMR029841 | | RANOC0325 |
| TPM3 | RHSPN01755 | RGG154022 | RCABP02379 | RNAMR0374 | RBRBA01555 | RANOC0328 |
| TPM4 | RHSPN24650 | RGG162621 | RCABP00928 | RNAMR0393 | RBRBA001279 | |
| TPP2 | RHSPN18998 | RGG018668 | RCABP00313 | RNAMR0246 | RBRBA00326 | RANOC0085 |
| TPT1 | RHSPN18735 | RGG020053 | RCABP04273 | RNAMR0287 | RBRBA00994 | RANOC0264 |
| TRADD | RHSPN21814 | RGG110269 | RCABP01669 | RNAMR0091 | RBRBA02738 | RANOC0156 |
| TRAF1 | RHSPN13793 | RGG133195 | RCABP04021 | RNAMR0155 | RBRBA031181 | |
| TRAF2 | RHSPN14122 | RGG131014 | RCABP02421 | RNAMR0149 | RBRBA03462 | RANOC0317 |
| TRAF3 | RHSPN19974 | RGG077180 | RCABP04395 | RNAMR0030 | RBRBA03762 | RANOC0213 |
| TRAF3IP2 | RHSPN10092 | RGG049994 | RCABP00487 | RNAMR0406 | RBRBA03714 | RANOC0024 |
| TRAF4 | RHSPN226173 | | RCABP019364 | | | |
| TRAF5 | RHSPN02478 | RGG044665 | RCABP03615 | RNAMR0280 | RBRBA01253 | RANOC0030 |
| TRAF6 | RHSPN15897 | RGG072374 | RCABP03408 | RNAMR0181 | RBRBA02254 | RANOC0002 |
| TRAF7 | RHSPN21135 | RGG122492 | RCABP01010 | RNAMR0397 | RBRBA01807 | RANOC0332 |
| TRAFD1 | RHSPN18248 | RGG126785 | RCABP03002 | RNAMR0098 | RBRBA02719 | RANOC0235 |
| TRAIP | RHSPN05455 | RGG113763 | RCABP01428 | RNAMR0001 | RBRBA01282 | RANOC0065 |
| TRAP1 | RHSPN21204 | RGG123938 | RCABP01300 | RNAMR0308 | RBRBA02296 | RANOC0283 |
| TRAT1 | RHSPN05827 | RGG012627 | RCABP03820 | RNAMR0019 | RBRBA00161 | RANOC0109 |

| | | | | | | |
|---|---|---|---|---|---|---|
| TREM1 | RHSPN096381 | | not predicted | | RBRBA039930 | |
| TREM2 | RHSPN0963 | RGG157332 | RCABP0042 | RNAMR0127 | RBRBA03994 | RANOC0116 |
| TREML1 | RHSPN096350 | | RCABP00421 | RNAMR0127 | RBRBA039942 | |
| TREML2 | RHSPN096365 | | RCABP0042 | RNAMR0127 | RBRBA039938 | |
| TREX1 | RHSPN053969 | | | RNAMR000050 | | RANOC0066 |
| TREX2 | RHSPN284769 | | RCABP0281 | RNAMR0409 | RBRBA04034 | RANOC0044 |
| TRIL | RHSPN1074 | RGG029022 | RCABP02504 | RNAMR0056 | RBRBA00962 | RANOC0230 |
| TRIM13 | RHSPN1878 | RGG020388 | RCABP0061 | RNAMR0102 | RBRBA0098 | RANOC0005 |
| TRIM14 | RHSPN1361 | RGG175945 | RCABP0234 | RNAMR0310 | RBRBA01981 | RANOC0197 |
| TRIM24 | RHSPN1164 | RGG007160 | RCABP0133 | RNAMR0173 | RBRBA02551 | RANOC0140 |
| TRIM25 | RHSPN2318 | RGG135644 | RCABP03467 | RNAMR0258 | RBRBA02241 | RANOC0195 |
| TRIM28 | RHSPN2576 | RGG164254 | RCABP02317 | RNAMR0326 | RBRBA002072 | |
| TRIM32 | RHSPN1377 | RGG131771 | RCABP0273 | RNAMR0155 | RBRBA0334 | RANOC0288 |
| TRIM33 | RHSPN0144 | RGG156915 | RCABP0394 | RNAMR0198 | RBRBA00564 | RANOC0131 |
| TRIM8 | RHSPN1515 | RGG083609 | RCABP02567 | RNAMR0325 | RBRBA02994 | RANOC0092 |
| TRIP11 | RHSPN1982 | RGG076375 | RCABP0009 | RNAMR0031 | RBRBA02062 | RANOC0018 |
| TRRAP | RHSPN1126 | RGG121221 | RCABP04342 | RNAMR0065 | RBRBA0130 | RANOC0262 |
| TSC1 | RHSPN1403 | RGG132670 | RCABP0011 | RNAMR0151 | RBRBA03492 | RANOC0327 |
| TSC2 | RHSPN2112 | RGG122471 | RCABP0359 | RNAMR0397 | RBRBA0180 | RANOC0325 |
| TSFM | RHSPN178143 | | RCABP0177 | RNAMR0272 | RBRBA0309 | RANOC0043 |
| TSHR | RHSPN1976 | RGG076045 | RCABP0273 | RNAMR0035 | RBRBA0058 | RANOC0000 |
| TSPAN6 | RHSPN2809 | RGG056649 | RCABP0444 | RNAMR0386 | RBRBA018843 | |
| TTC6 | RHSPN1937 | RGG075309 | RCABP0204 | RNAMR0411 | RBRBA0277 | RANOC0032 |
| TTL | RHSPN0383 | RGG043523 | RCABP04305 | RNAMR0106 | RBRBA0386 | RANOC0272 |
| TTR | RHSPN2386 | RGG036309 | RCABP01627 | RNAMR0192 | RBRBA00177 | RANOC0201 |
| TUBB | RHSPN094078 | | RCABP02884 | RNAMR0131 | RBRBA0100 | RANOC0272 |
| TUBE1 | RHSPN1009 | RGG049874 | RCABP021622 | | RBRBA0371 | RANOC0007 |
| TUBG1 | RHSPN2290 | RGG159899 | RCABP00501 | RNAMR0254 | RBRBA02647 | RANOC0176 |
| TUSC3 | RHSPN1202 | RGG063949 | RCABP0169 | RNAMR0337 | RBRBA0110 | RANOC0144 |
| TXK | RHSPN0695 | RGG064502 | RCABP0415 | RNAMR0158 | RBRBA000634 | |
| TXN | RHSPN1369 | RGG177490 | RCABP03794 | RNAMR0142 | RBRBA0362 | RANOC0278 |
| TYRO3 | RHSPN2031 | RGG073482 | RCABP02397 | RNAMR0277 | RBRBA0060 | RANOC0011 |
| U2AF1 | RHSPN2679 | RGG014942 | RCABP0206 | RNAMR0085 | RBRBA0082 | RANOC0087 |
| UBASH3A | RHSPN2677 | RGG014815 | RCABP0037 | RNAMR0085 | RBRBA0398 | RANOC0099 |
| UBE2A | RHSPN2824 | RGG058086 | RCABP03984 | RNAMR038224 | | |
| UBE2B | RHSPN0856 | RGG120150 | RCABP0146 | RNAMR0026 | RBRBA0130 | RANOC0046 |
| UBE2J1 | RHSPN0996 | RGG050975 | RCABP01114 | RNAMR0262 | RBRBA02184 | RANOC0023 |
| UBE2K | RHSPN0690 | RGG064865 | RCABP0022 | RNAMR0159 | RBRBA02881 | RANOC0155 |
| UBE2N | RHSPN180214 | | | RNAMR021544 | | |
| UBE2V2 | RHSPN1230 | RGG036520 | RCABP0071 | RNAMR0036 | RBRBA0277 | RANOC0200 |
| UBR5 | RHSPN1263 | RGG038753 | RCABP04371 | RNAMR0163 | RBRBA03132 | RANOC0133 |
| UCHL3 | RHSPN1887 | RGG019432 | RCABP0293 | RNAMR0113 | RBRBA0212 | RANOC0094 |
| UCP1 | RHSPN075087 | | not predicte | RNAMR0414 | RBRBA029206 | |
| UCP3 | RHSPN1648 | RGG023504 | RCABP01254 | RNAMR0293 | RBRBA01511 | RANOC0321 |
| ULK1 | RHSPN1846 | RGG125735 | RCABP03494 | RNAMR010111 | | RANOC0257 |

| | | | | | | |
|---|---|---|---|---|---|---|
| UNC13D | RHSPN2345 | RGG135317 | RCABP0231 | RNAMR0268 | RBRBA0378 | RANOC0061 |
| UNC93B1 | RHSPN1638 | RGG068385 | RCABP0077 | RNAMR0183 | RBRBA0114 | RANOC0004 |
| UNG | RHSPN1818 | RGG126934 | RCABP03441 | RNAMR0097 | RBRBA018518 | |
| UQCRB | RHSPN1257 | RGG038426 | RCABP0370 | RNAMR0162 | RBRBA0312 | RANOC0111 |
| USP12 | RHSPN1858 | RGG021279 | RCABP0382 | RNAMR0209 | RBRBA0122 | RANOC0242 |
| UVSSA | RHSPN0663 | RGG066544 | RCABP0011 | RNAMR0135 | RBRBA0407 | RANOC0273 |
| VAMP7 | RHSPN2868 | RGG057169 | RCABP0246 | RNAMR0413 | RBRBA0388 | RANOC0198 |
| VAMP8 | RHSPN035242 | | RCABP0207 | RNAMR0107 | RBRBA0352 | RANOC0341 |
| VASH1 | RHSPN1973 | RGG075688 | RCABP0312 | RNAMR0032 | RBRBA0059 | RANOC0001 |
| VAV1 | RHSPN243851 | | RCABP0014 | RNAMR0317 | RBRBA035512 | |
| VAV3 | RHSPN0133 | RGG092902 | RCABP0328 | RNAMR0196 | RBRBA0058 | RANOC0114 |
| VCAM1 | RHSPN0131 | RGG095242 | RCABP0340 | RNAMR0352 | RBRBA0303 | RANOC0124 |
| VCP | RHSPN1318 | RGG171004 | RCABP0298 | RNAMR0399 | RBRBA0375 | RANOC0312 |
| VEGFA | RHSPN0971 | RGG045620 | RCABP0429 | RNAMR0132 | RBRBA0069 | RANOC0027 |
| VEGFC | RHSPN0771 | RGG061231 | RCABP0147 | RNAMR0190 | RBRBA0273 | RANOC0144 |
| VHL | RHSPN0499 | RGG117180 | RCABP0153 | RNAMR0019 | RBRBA0190 | RANOC0230 |
| VIMP | RHSPN2101 | RGG108257 | RCABP0388 | RNAMR0187 | RBRBA0251 | RANOC0319 |
| VIP | RHSPN1037 | RGG047848 | RCABP0052 | RNAMR0229 | RBRBA0379 | RANOC0007 |
| VIPR1 | RHSPN0528 | RGG024730 | RCABP0011 | RNAMR0012 | RBRBA0062 | RANOC0165 |
| VPS13D | RHSPN0024 | RGG146598 | RCABP0073 | RNAMR0296 | RBRBA0236 | RANOC0328 |
| VPS35 | RHSPN2163 | RGG110934 | RCABP0097 | RNAMR0399 | RBRBA0326 | RANOC0187 |
| VSIR | RHSPN1478 | RGG081109 | RCABP0187 | RNAMR0123 | RBRBA0356 | RANOC0093 |
| VTI1A | RHSPN1522 | RGG084155 | RCABP0236 | RNAMR0050 | RBRBA0249 | RANOC0199 |
| VWA5A | RHSPN1693 | RGG157599 | RCABP0158 | RNAMR0369 | RBRBA0327 | RANOC0223 |
| WAS | RHSPN278322 | | RCABP0427 | RNAMR0380 | RBRBA0162 | RANOC0043 |
| WASL | RHSPN1151 | RGG002535 | RCABP0414 | RNAMR0225 | RBRBA0353 | RANOC0143 |
| WDR27 | RHSPN1050 | RGG046993 | RCABP0063 | RNAMR0349 | RBRBA0019 | RANOC0027 |
| WDR72 | RHSPN2048 | RGG106753 | RCABP02651 | RNAMR0014 | RBRBA0352 | RANOC0334 |
| WHSC1L1 | RHSPN1222 | RGG147789 | RCABP0066 | RNAMR0214 | RBRBA0009 | RANOC0268 |
| WIF1 | RHSPN1785 | RGG003608 | RCABP00721 | RNAMR0271 | RBRBA0226 | RANOC0151 |
| WNT11 | RHSPN1652 | RGG023199 | RCABP0352 | RNAMR0292 | RBRBA0150 | RANOC0241 |
| WNT4 | RHSPN0041 | RGG146824 | RCABP0405 | RNAMR0166 | RBRBA0052 | RANOC0338 |
| WRAP53 | RHSPN2235 | RGG179760 | RCABP03471 | RNAMR0235 | RBRBA0306 | RANOC0337 |
| WRN | RHSPN1218 | RGG060267 | RCABP0420 | RNAMR0215 | RBRBA0110 | RANOC0225 |
| WT1 | RHSPN1585 | RGG069314 | RCABP0290 | RNAMR0177 | RBRBA0062 | RANOC0033 |
| WWOX | RHSPN2198 | RGG111685 | RCABP04081 | RNAMR0089 | RBRBA0092 | RANOC0273 |
| WWTR1 | RHSPN0620 | RGG104300 | RCABP0351 | RNAMR0005 | RBRBA0102 | RANOC0105 |
| XAB2 | RHSPN244032 | | RCABP0085 | RNAMR0314 | RBRBA0121 | RANOC0045 |
| XBP1 | RHSPN2714 | RGG127219 | RCABP0039 | RNAMR0108 | RBRBA018584 | |
| XDH | RHSPN0312 | RGG041910 | RCABP0279 | RNAMR0053 | RBRBA0402 | RANOC0009 |
| XIAP | RHSPN2827 | RGG057972 | RCABP0204 | RNAMR0261 | RBRBA0226 | RANOC0266 |
| XKR6 | RHSPN1197 | RGG054373 | RCABP0101 | RNAMR0145 | RBRBA0052 | RANOC0290 |
| XPA | RHSPN1360 | RGG177925 | RCABP0395 | RNAMR0310 | RBRBA0198 | RANOC0289 |
| XPC | RHSPN0504 | RGG115736 | RCABP0153 | RNAMR0365 | RBRBA0189 | RANOC0048 |
| XPO1 | RHSPN0333 | RGG041386 | RCABP0209 | RNAMR0333 | RBRBA0382 | RANOC0286 |

| | | | | | | |
|---|---|---|---|---|---|---|
| XRCC1 | RHSPN25166 | RGG023288 | RCABP02711 | RNAMR0391 | RBRBA04078 | RANOC0187 |
| XRCC2 | RHSPN11842 | RGG025646 | RCABP04053 | RNAMR02644 | RBRBA03226 | RANOC0166 |
| XRCC3 | RHSPN19991 | RGG077395 | RCABP04394 | RNAMR00357 | RBRBA00081 | RANOC00003 |
| XRCC4 | RHSPN08281 | RGG177043 | RCABP03655 | RNAMR00774 | RBRBA03696 | RANOC00412 |
| XRCC5 | RHSPN04582 | RGG090592 | RCABP03746 | RNAMR0306 | RBRBA03662 | RANOC02543 |
| XRCC6 | RHSPN27404 | RGG005752 | RCABP00185 | RNAMR0116 | RBRBA02080 | RANOC01413 |
| YAP1 | RHSPN16687 | RGG022011 | RCABP03605 | RNAMR03688 | RBRBA02201 | RANOC01032 |
| YBX2 | RHSPN223253 | | RCABP02457 | RNAMR023478 | | RANOC03454 |
| YTHDF2 | RHSPN00545 | RGG149437 | RCABP03207 | RNAMR01687 | RBRBA02566 | RANOC03227 |
| YWHAE | RHSPN22168 | RGG139290 | RCABP01897 | RNAMR0232 | RBRBA003472 | |
| YWHAG | RHSPN11122 | RGG138876 | RCABP00973 | RNAMR0062 | RBRBA03112 | RANOC03323 |
| ZAP70 | RHSPN03671 | RGG161027 | RCABP00003 | RNAMR010496 | | |
| ZBTB16 | RHSPN16794 | RGG152125 | RCABP02600 | RNAMR0227 | RBRBA001514 | |
| ZBTB20 | RHSPN05890 | RGG011544 | RCABP00863 | RNAMR0017 | RBRBA02594 | RANOC0101 |
| ZFAND2A | RHSPN105427 | | RCABP02415 | RNAMR00644 | RBRBA006803 | |
| ZFAND2B | RHSPN04638 | RGG090254 | RCABP02198 | RNAMR0306 | RBRBA03672 | RANOC0017 |
| ZFYVE1 | RHSPN19665 | RGG073903 | RCABP01274 | RNAMR0033 | RBRBA01336 | RANOC00014 |
| ZFYVE28 | RHSPN06659 | RGG066364 | RCABP01248 | RNAMR0134 | RBRBA04069 | RANOC0065 |
| ZMPSTE24 | RHSPN00725 | RGG149216 | RCABP00827 | RNAMR01714 | RBRBA017143 | |
| ZMYM2 | RHSPN18522 | RGG021615 | RCABP01392 | RNAMR0209 | RBRBA03226 | RANOC0102 |
| ZNF278 | | | | | | |
| ZNF331 | RHSPN255564 | | | RNAMR032816 | | |
| ZNF384 | RHSPN17141 | RGG010481 | RCABP00323 | RNAMR0355 | RBRBA02389 | RANOC02614 |
| ZPLD1 | RHSPN05801 | RGG012403 | RCABP03359 | RNAMR0240 | RBRBA03567 | RANOC0087 |

### B Non-synonymous Variants (SNVs) Unique to Jonathan

| gene | Human Pos | Jonathan Pos |
|---|---|---|
| **ABL1** | V753M | M740V |
| **ACD** | S429P | P470S |
| **ACSL6** | V519L | L526V |
| **ADAMTSL1** | A1003V | V874A |
| **ADAR** | G488D | D397G |
| **ADGB** | A1166V | V587A |
| **AFF3** | R978K | K490R |
| **AGPAT2** | P232D | D236P |
| **AIF1** | S168G | G102S |
| **AKT1S1** | P102T | T83P |
| **ALK** | G2I,A3S,I4H,L80T,K81R,L422M | I2G,S3A,H4I,T37L,R38K,M75L |
| **ALKBH3** | A205V,D280G | V208A,G283D |
| **ANG** | L8M | M39L |
| **AOPEP** | E61K | K75E |
| **APBB1IP** | T349I | I318T |

| | | |
|---|---|---|
| **APC2** | V196M,A1545Q | M195V,Q1698A |
| **APEX2** | R132C,A133T | C118R,T119A |
| **APOA4** | A268N | N355A |
| **APOA5** | A160V | V191A |
| **APOB** | R2012Q,A4286S | Q2004R,S4283A |
| **APOBEC2** | L24S | S21L |
| **APOM** | V70I | I70V |
| **AQP9** | V62A | A90V |
| **ARID1B** | T826A | A894T |
| **ART1** | P126L,S300N | L157P,N330S |
| **ATF6** | R403S,T606K | S391R,K577T |
| **ATF6B** | L381A | A340L |
| **ATF7** | M1A | A13M |
| **ATG16L1** | G583E | E582G |
| **ATG4C** | V245I | I245V |
| **ATG9A** | A2M,P360L,L635V | M2A,L360P,V633L |
| **ATM** | E1245Q,K1451V,D1452K,V1532S,K2456R | Q326E,V495K,K496D,S576V,R1508K |
| **ATP7A** | R512G,D828G,Q1149N | G576R,G891D,N1212Q |
| **ATRX** | R1131Q | Q1136R |
| **BAG6** | V1055L | L1075V |
| **BCL11B** | L127R | R65L |
| **BCL9** | M1074V | V1073M |
| **BMP6** | D402G,K482E,L483V,N484M,A485R,S487A | G206D,E208K,V209L,M210N,R211A,A213S |

| | | |
|---|---|---|
| BNIPL | [illegible] E51D,A69N, T71E,P72A,S73D,T74I,L77V,C78T,R81Q,P82K,M83V,R86K,S88M,L92I,S95T,T97N,S119G,D120E,L121V,E122D,I123L,E125D,E127D,S132N,G138S, H139N,E146D,R149K,A150P,E151K, G152T,L153T,G154E,T155G,T158P, E160Y,R161T,L162A,G163A,G172R, E173D,H177R,V180M,M183I,R186Q, E187D,Q188H,T193K,V194A,I216V,L217V,S220V, | [illegible] D80E,N82A, E84T,A85P, D86S,I87T,V90L,T91C,Q94R,K103P, V104M,K107R,M109S,I113L,T116S, N130T,G133S,E134D,V135L,D136E, L137I,D139E,D141E,N146S,S147G, N148H,D155E,K158R,P159A,K160E, T161G,T162L,E163G,G168T,P171T,Y173E,T174R,A175L,A176G,R179G,D180E,R184H, M187V,I190M,Q193R,D194E,H195Q,K200T,A201V,V223I,V2 |
| BRCA1 | M1083R | R1105M |
| BRD4 | V997I | I956V |
| BSCL2 | R350Q | Q288R |
| C5 | D665G | G662D |
| C5AR1 | F182N | N184F |
| CASC5 | H415D,N2321C | D408H,C2403N |
| CBLB | L590P,G748S | P590L,S749G |
| CCL5 | A23P | P22A |
| CD1D | G46T,K232Q | T122G,Q312K |
| CD200 | S157R | R79S |

| | | |
|---|---|---|
| **CD207** | M1S,A7D,K38E,P40S,T50C,N77E,G83E,R84N,N87D,Q197S,T231A,A246I,D263T,S265C,R279E,N291T | S2M,D16A,E34K,S36P,C46T,E73N,E84G,N85R,D88N,S150Q,A184T,I199A,T216D,C218S,E232R,T246N |
| **CD22** | S123I,H133G,H213D,H240C,L352R | I264S,G281H,D356H,C383H,R542L |
| **CD302** | M110V | V113M |
| **CD38** | R47M | M40R |
| **CD5L** | E84K | K117E |
| **CD97** | F51T,T59M,S72R,K107T,V296H,G348N,A391H,R440H,I710M | T54F,M63T,R76S,T111K,H230V,N262G,H308A,H358R,M609I |
| **CDH11** | M382V | V382M |
| **CEL** | L542M | M483L |
| **CEP89** | Q688H | H698Q |
| **CFH** | N718D,S720D,S722R,T724K,H728S,S730K,T732E,H735D,V745T | D250N,D252S,R254S,K256T,S260H,K262S,E264T,D267H,T277V |
| **CFP** | K44E | E38K |
| **CGNL1** | Q27D,C234D | D27Q,D235C |
| **CHIT1** | D213K | K198D |
| **CIC** | S1258A,L2144P | A1146S,P1995L |
| **CLIP4** | P511A | A457P |
| **CMAS** | P43A | A21P |
| **CNOT3** | A442I | I418A |
| **COLEC12** | P493L,E672A | L484P,A666E |
| **CPA2** | N194E | E194N |
| **CRLS1** | V298L | L234V |
| **CSMD3** | M1A,V1966M | A96M,M1405V |

| | | |
|---|---|---|
| **CTSG** | Q2L,L5I,L8A,T14S,E17V,Q53E,S56H,A58D,Q60S,R69L,N92D,Q94T,R95K,R96Q,N98R,H102V,T104R,I109Y,P112Q,Q113D,R117K,T118S,I119H,Q127K,S129E,R130H,R131N,R133L,R136E,N137Y,N139S,A142P,R145M,E148Q,G149R,L150M,L155V,R163Q,M166A,R167H,R168N,G169N,T170L,Q177E,L178V,R179V,Q181M,R182Q,R184A,Q185E,G191R,S192N,D194N,R | L2Q,I5L,A9L,S15T,V18E,E40Q,H43S,D45A,S47Q,L56R,D79N,T81Q,K82R,Q83R,R85N,V89H,R91T,Y96I,Q99P,D100Q,K104R,S105T,H106I,K114Q,E116S,H117R,N118R,L120R,E123R,Y124N,S126N,P129A,M132R,Q135E,R136G,M137L,V142L,Q150R,A153M,H154R,N155R,N156G,L157T,E166Q,V167L,V168R,M170Q,Q171R,A173R,E174Q,R184G,N185S,N187D,P197R |
| **CYC1** | P38S | S27P |
| **CYP19A1** | V158M,K461H | M158V,H461K |
| **CYP27B1** | V397L | L394V |
| **DAPK1** | P1355S | S1355P |
| **DAXX** | S188L | L156S |
| **DHX33** | E4G | G4E |
| **DHX58** | T395S | S419T |
| **DHX9** | T108A | A108T |
| **DKC1** | E210G | G209E |
| **DMC1** | E197K,H198T,Q199P,M200F,E201P,E361D,E362H | K199E,T207H,P208Q,F209M,P210E,D230E,H231E |
| **ECHS1** | V47I | I50V |
| **EGF** | R1082S | S1136R |
| **EIF2AK1** | A18T,A20P,P57H,N348Y,G624R | T21A,P23A,H59P,Y347N,R635G |
| **EIF2AK4** | Y60H | H58Y |

| | | |
|---|---|---|
| EME1 | T188V,L193A,E304K,R391Q,G413S,T469S | V205T,A210L,K323E,Q398R,S427G,S483T |
| ENDOV | P40C,P184L | C40P,L184P |
| ENOX1 | C443S | S354C |
| ERBB2 | T166M | M162T |
| ERBB2IP | D724Q | Q720D |
| ERCC1 | L69S | S93L |
| ERCC5 | Q231E,D469K,L503S,G946E,G1080K | E233Q,K435D,S464L,E913G,K1043G |
| EXOSC3 | Q39Y | Y73Q |
| F10 | E29A | A32E |
| F7 | V24I,K70Q,A220T,P310S | I26V,Q72K,T186A,S274P |
| FANCA | G17S,A21N,L704F | S38G,N44A,F723L |
| FANCM | K1355H,Q2011N | H1176K,N1827Q |
| FBXO31 | G396D,V450M | D424G,M465V |
| FGA | P352E | E251P |
| FGF19 | R63S | S64R |
| FLT3 | I21A,V56I,D779N | A19I,I54V,N787D |
| FOSB | G111T | T120G |
| FOSL2 | S83N,P92D,G101D,P121S | N82S,D92P,D101G,S121P |
| FOXO1 | Q6K | K6Q |
| FOXO3 | A98G | G38A |
| GCH1 | P28Q | Q33P |
| GCNT3 | R6Y | Y7R |
| GHSR | R336S | S328R |
| GOPC | Q109K | K112Q |
| GRN | P430L | L469P |
| GSR | S120L | L60S |
| GZMB | K92Y,Q149S,Q214Y | Y93K,S150Q,Y219Q |
| GZMH | P3L,L7F,L8I,L11P,T13L,G17Q | L74P,F78L,I80L,P83L,L85T,Q89G |
| HAVCR2 | P138Q | Q179P |
| HDAC6 | N60R | R16N |
| HELQ | S517D | D487S |

| | | |
|---|---|---|
| **HIF1A** | S616R | R584S |
| **HIP1** | D344G | G319D |
| **HLA-DRB1** | L6G,G9S,S10C,C11W,M12A,M20T,R33P,P34R,R35G,L37V,W38Y,P40T,K41N,H45Y,S66H,V67L,F76W,P85A,D86T,Y89L,I96S,E98Q,Q99Y,A100K,G113Q,V114E,V115T,Q121S,T129K,Y131S,L138E,H141P,S149T,Q165L,T210Q,S211G | G8L,S11G,C12S,W13C,A14M,T22M,P35R,R36P,G37R,V39L,Y40W,T42P,N43K,Y47H,H68S,L69V,W78F,A87P,T88D,L91Y,S98I,Q100E,Y101Q,K102A,Q115G,E116V,T117V,S123Q,K131T,S133Y,E140L,P143H,T151S,L167Q,Q212T,G213S |
| **HLTF** | R36H,L328F | H31R,F329L |
| **HOXD12** | A215C | C211A |
| **HRH1** | V42I | I27V |
| **HSF4** | P270S,E342S | S270P,S340E |
| **HSF5** | P178S | S182P |
| **HSPA13** | Q270E | E270Q |
| **HSPA2** | E559G | G559E |
| **IDH1** | G204S | S267G |
| **IFNAR1** | T68E | E70T |
| **IFNGR2** | L242K,S246G,L314Q,D343G | K200L,G203S,Q269L,G297D |
| **IFNL3** | Q52W,Q55K,A165T | W15Q,K18Q,T125A |
| **IGF2R** | K1862S | S1849K |
| **IL17C** | L70V | V87L |
| **IL18RAP** | H55Y,Q66R,G117E,V300F | Y55H,R66Q,E102G,F296V |
| **IL1RL1** | I332T,T391K,T436M,V537R | T356I,K413T,M457T,R553V |
| **IL1RL2** | G127S | S89G |
| **IL31RA** | N536Y | Y675N |
| **IRG1** | K150S | S164K |

| | | |
|---|---|---|
| **ITCH** | V24I,N159R | I37V,R126N |
| **ITGAL** | M1125L | L1193M |
| **ITGB3** | A50T,T221A,M594V | T96A,A266T,V639M |
| **ITGB7** | Q102R,R129A,V675M | R116Q,A143R,M683V |
| **JAG1** | G1100T | T1095G |
| **KCNT2** | I974M | M988I |
| **KIAA1549** | D155N,P196S | N105D,S147P |
| **KLRF1** | P202L | L215P |
| **KMT2A** | S570R | R513S |
| **KMT2D** | P1165L | L827P |
| **LAMP1** | T39E | E35T |
| **LATS1** | S1023A | A1031S |
| **LATS2** | A325S | S350A |
| **LEP** | L147V | V148L |
| **LIG3** | A197M | M193A |
| **LINGO2** | I470V,V591I | V470I,I591V |
| **LMNB1** | T96M | M94T |
| **LMX1B** | M196I | I78M |
| **LPXN** | P383T | T386P |
| **LRP1B** | T2167M,P2946S,E3112Q | M1837T,S2616P,Q2782E |
| **LY75** | P153S,K488N,Q959E,T1354H,K1519Q | S150P,N486K,E957Q,H1354T,Q1519K |
| **LY9** | T120Q,I122F,A133I,E141T,K158P,N171G,N181I,Q287D,P355F | Q20T,F22I,I34A,T42E,P60K,G73N,I83N,D184Q,F313P |
| **MADCAM1** | E263S,R292S,P355A | S254E,S353R,A474P |
| **MALSU1** | V28S | S17V |
| **MAP3K13** | I745L | L748I |
| **MAP4K2** | N349S,V723I | S352N,I749V |
| **MAPK1** | Y358D | D366Y |
| **MARCH8** | T538I | I534T |
| **MARCO** | L74M | M46L |
| **MASP2** | S112A,S185E | A87S,E160S |
| **MBD4** | S122R | R107S |
| **MET** | S122Q | Q102S |

| | | |
|---|---|---|
| **METTL1** | Q117H | H108Q |
| **METTL21B** | H194V,Q198P | V215H,P219Q |
| **MID2** | V59L | L39V |
| **MINPP1** | N424H | H407N |
| **MMS19** | T957N | N948T |
| **MR1** | M5L,I7L,L8R,P10L,V17L,K18R,D21R,T24L,L27R,Y29I,R31H,E43W,F44Y,I45S,S46R,V47I,P54V,T57N,D59T,T62S,R63Q,K65V,A69S,P70V,A77D,H80F,R83F,T85N,Q86F,L87W,G90R,V97A,H105L,T113I,Y114I,R116M,M117I,I118T,E121D,L123Q,G126N,S127T,T129W,Y134D,A135S,I143T,N145H,D147E,L149M,N156I,V157G,H159Q,Q163R,A164R,A167T,N168E,Q169I,H1 | L5M,L7I,R6L,L8P,L17V,R18K,R21D,L24T,R27L,I29Y,H31R,W43E,Y44F,S45I,R46S,I47V,V54P,N57T,T59D,S62T,Q63R,V65K,S69A,V70P,D77A,F80H,F83R,N85T,F86Q,W87L,R90G,A97V,L105H,I114T,I115Y,M117R,I118M,T119I,D122E,Q124L,N127G,T128S,W130T,D135Y,S136A,T144I,H146N,E148D,M150L,I157N,G158V,Q160H,R164Q,R165A,T168A,E169N,I170Q,G1 |
| **MRC2** | K241R | R15K |
| **MRE11A** | N617S,D622G | S620N,G625D |
| **MSH6** | S602A,V1056G | A611S,G1065V |
| **MSRA** | A96S,E177G | S129A,G210E |
| **MTFMT** | G16V,S317L | V16G,L307S |
| **MUTYH** | T256L | L221T |
| **NAB2** | G494R | R437G |
| **NAV2** | S1179C | C1122S |
| **NCAM2** | V163I,R307E | I135V,E279R |
| **NCOA4** | S511R | R499S |

| | | |
|---|---|---|
| **NCOR2** | T946S,H1853L,A2237I | S899T,L1821H,I2205A |
| **NDRG1** | E38D | D75E |
| **NEDD4** | K261Q,K347T,F753A,S811A | Q259K,T346K,A757F,A815S |
| **NFATC1** | V928A | A935V |
| **NFKB1** | T212A | A208T |
| **NFKBIZ** | C177H,I353V | H160C,V334I |
| **NOTCH1** | N2396T | T2395N |
| **NRG1** | P49H | H42P |
| **NSD1** | A480T,L2513F | T406A,F2368L |
| **NTRK3** | F720Y | Y639F |
| **NUTM1** | S165T | T79S |
| **OASL** | L119V | V108L |
| **PARVG** | L339P,N341Q | P303L,Q306N |
| **PER1** | P355S,I1085G,R1112L | S402P,G1075I,L1103R |
| **PIK3AP1** | R706I,L709M | I705R,M708L |
| **PIK3C2G** | N426M | M1N |
| **PKN2** | E613G | G566E |
| **PLCL2** | S86G | G77S |
| **PLEKHA1** | P294S | S296P |
| **POLE** | V146M,R1231H | M145V,H1230R |
| **POLE2** | L107V | V107L |
| **POLM** | T21A,P146Q,L331T,L359M | A18T,Q140P,T320L,M348L |
| **POLQ** | S244I,D2414K | I186S,K2402D |
| **POT1** | K440E | E634K |
| **POU2F2** | G528A | A555G |
| **PPFIBP1** | Q249H,V288M | H283Q,M322V |
| **PPM1D** | V440I | I437V |
| **PPP1R1A** | E49K | K17E |
| **PPP1R26** | D503G | G499D |
| **PRKCG** | T438N,P602L | N427T,L591P |

| | | |
|---|---|---|
| **PRKDC** | V291R,T516S,G599S,S621V,V749I,Y779H,H783Y,D791Y,T809P | R15V,S244T,S326G,V348S,I476V,H506Y,Y510H,Y518D,P536T |
| **PROM1** | I169M,V371L,V485S | M1I,L92V,S155V |
| **PROS1** | G5S,S428N | S5G,N398S |
| **PRUNE2** | E78Q,P203L,P1564A | Q38E,L163P,A1554P |
| **PSMD5** | S266A | A266S |
| **PTCD2** | A316C | C318A |
| **PTGS1** | V581L | L600V |
| **PTPRB** | M489V,V613H,E646A,E663T,M750N,A885P,V889R,H1182V,L1201T,L1442A,R1454Q,Y1538W,Q2100G | V50M,H178V,A213E,T228E,N318M,P365A,R369V,V491H,T514L,A576L,Q588R,W668Y,G1192Q |
| **PUM2** | N242S | S372N |
| **RABGEF1** | V26L | L14V |
| **RAD50** | E384K | K384E |
| **RALGDS** | D41G | G42D |
| **RAPGEF3** | V906T | T835V |
| **RB1** | H69V,P253A | V54H,A245P |
| **RBPMS** | N17G | G16N |
| **REV1** | Q637R | R657Q |
| **RIF1** | S1277L,Q2111V | L1303S,V1979Q |
| **RNF213** | V1824M,V3576N | M1803V,N3526V |
| **RNF8** | P169H | H168P |
| **ROS1** | Y1312C,Q1368N,Y1488A,L1612T,K2100R | C1274Y,N1331Q,A1450Y,T1580L,R2072K |
| **RRM2B** | D414G | G344D |
| **SEMA4A** | A419T,G757R | T410A,R771G |
| **SEMA4D** | G602H | H551G |
| **SETBP1** | L556I | I393L |
| **SETD2** | A1408I,D1444E | I1400A,E1436D |
| **SH2B2** | S639N | N644S |
| **SH2D4A** | R334Q | Q337R |

| | | |
|---|---|---|
| SHPRH | N637R,N647G | R670N,G731N |
| SIRT1 | G19V | V19G |
| SMUG1 | R66K | K62R |
| SORCS1 | G807R | R485G |
| SPPL2A | T78S | S62T |
| SSPN | E28K | K37E |
| ST6GAL1 | D146G | G160D |
| STK11 | K108R | R108K |
| STRN | D226H | H236D |
| SVEP1 | V494L,T625I,G631N,S832L,S3235V | L261V,I392T,N398G,L599S,V3008S |
| SYNE1 | A2020G,P2769I,S3321G,S3915K,E4957K,A5702V,L5842F,A6283H | G677A,I1426P,G1978S,K2572S,K3614E,V4361A,F4501L,H4935A |
| SYT13 | A159I | I160A |
| TAPBP | R57K | K70R |
| TDG | S376V | V399S |
| TEP1 | A1976V,S2001L | V1966A,L1991S |
| TERF2 | R322F | F360R |
| TERT | R19S,L593Y | S19R,Y197L |
| TET1 | L1329H | H1331L |
| THEMIS | D630N | N555D |
| THRAP3 | H693L | L706H |
| TINAGL1 | H467W | W465H |
| TLR4 | G420C | C819G |
| TMEM173 | E249D | D256E |
| TNFSF8 | P69H | H69P |
| TNIP3 | G114T | T88G |
| TOP2A | P681S | S685P |
| TOPBP1 | T548I | I538T |
| TP53BP1 | M721V | V678M |
| TPP2 | A636L | L635A |
| TRAT1 | K92D | D115K |
| TREML1 | L5P,G15E,F54V,R110L | P5L,E15G,V54F,L114R |
| TRIM24 | Y540S | S539Y |
| TRIM25 | K332T | T335K |
| TRIM33 | G877E | E808G |
| TRIM8 | V225L | L225V |
| TSFM | L143A,I219T | A157L,T211I |

| | | |
|---|---|---|
| **VAV3** | Q634Y | Y617Q |
| **VWA5A** | K98R | R81K |
| **WDR27** | E651K | K630E |
| **XDH** | T510M | M533T |
| **XRCC3** | P159R | R159P |
| **XRCC6** | Q68W | W68Q |
| **ZBTB20** | N437D | D455N |
| **ZFYVE28** | G592K | K556G |
| **ZPLD1** | T38I | I11T |

### C Key for abbreviations in tab "A"

| Abbreviation | Description |
|---|---|
| ANOC | Anolis carolinensis (green anole) |
| ALGO | Cillinger's Aldabra (Aldabra tortoise) |
| BHWH | Bowhead Whale |
| BRBA | Brandt's Bat |
| CPIC | Chrysemys picta bellii (Western painted turtle) |
| CABP | Lonesome George (Galpagos tortoise) |
| DTNK | Tank (Aldabra tortoise) |
| DJON | Jonathan (Aldabra tortoise) |
| DR | Danio Rerio (zebrafish) |
| GOAS | Gopherus agassizii (desert tortoise) |
| GG | Galus Galus (chicken) |
| MMUS | mouse |
| NAMR | Naked Mole Rat |
| NFUR | Nothobranchius furzeri (turquoise killifish) |
| PSIN | Pelodiscus sinensis (Chinese softshell) |
| TORT | Reference tortoise genome in a particular alignment |
| JON | Jonathan (Aldabra tortoise) |
| LG | Lonesome George (Galpagos tortoise) |

| PSIN | MMUS | NFUR | DR | CPIC | GOAS | ALGO |
|---|---|---|---|---|---|---|
| RPSIN00819 | RMMUS101729 | | | RCPIC05126 | RGOAS0243 | RALGO6097 |
| RPSIN02379 | RMMUS2262 | RNFUR0011 | RDR050214 | RCPIC04684 | RGOAS0108 | RALGO6953 |
| RPSIN03211 | RMMUS1154 | RNFUR0420 | RDR046621 | RCPIC03089 | RGOAS0238 | RALGO3993 |
| RPSIN00674 | RMMUS0200 | RNFUR0170 | RDR003368 | RCPIC00275 | RGOAS0260 | RALGO32761 |
| RPSIN02405 | RMMUS0228 | RNFUR0405 | RDR010816 | RCPIC02025 | RGOAS0271 | RALGO1743 |
| RPSIN02219 | RMMUS0134 | RNFUR0499 | RDR018203 | RCPIC01245 | RGOAS0163 | RALGO43151 |
| | | | | | | not predicte |
| | RMMUS1823 | RNFUR0146 | RDR010392 | RCPIC02857 | RGOAS0338 | RALGO24325 |
| 69 | RMMUS139568 | | | RCPIC056821 | | |
| RPSIN02063 | RMMUS1887 | RNFUR0278 | RDR025580 | RCPIC02691 | RGOAS0166 | RALGO11681 |
| RPSIN02280 | RMMUS008361 | | RDR012311 | RCPIC02097 | RGOAS0191 | RALGO0701 |
| RPSIN03244 | RMMUS0068 | RNFUR0219 | RDR002626 | RCPIC01736 | RGOAS0210 | RALGO3343 |
| RPSIN00776 | RMMUS1774 | RNFUR0490 | RDR041169 | RCPIC00946 | RGOAS0275 | RALGO5366 |
| RPSIN01055 | RMMUS2678 | RNFUR0447 | RDR014933 | RCPIC00966 | RGOAS0099 | RALGO1232 |
| RPSIN02050 | RMMUS0255 | RNFUR0510 | RDR002775 | RCPIC034935 | | RALGO39312 |
| RPSIN02903 | RMMUS0383 | RNFUR0440 | RDR043251 | RCPIC02978 | RGOAS0114 | RALGO2212 |
| RPSIN02617 | RMMUS0609 | RNFUR023702 | | RCPIC01124 | RGOAS0184 | RALGO55604 |
| RPSIN00003 | RMMUS0476 | RNFUR0267 | RDR031182 | RCPIC04554 | RGOAS0117 | RALGO41711 |
| RPSIN00423 | RMMUS1649 | RNFUR0213 | RDR020426 | RCPIC03760 | RGOAS0167 | RALGO1147 |
| RPSIN00231 | RMMUS2041 | RNFUR0482 | RDR046320 | RCPIC003784 | | RALGO27665 |
| RPSIN03634 | RMMUS2423 | RNFUR0149 | RDR019073 | RCPIC045188 | | RALGO46563 |
| RPSIN00153 | RMMUS159605 | | RDR038992 | RCPIC00554 | RGOAS0070 | RALGO56937 |
| RPSIN03448 | RMMUS0541 | RNFUR0515 | RDR022821 | RCPIC03121 | RGOAS0018 | RALGO2774 |
| RPSIN01983 | RMMUS240955 | | RDR002433 | RCPIC01734 | RGOAS0278 | RALGO37055 |
| | RMMUS179160 | | RDR009576 | RCPIC02873 | RGOAS0338 | RALGO24172 |
| RPSIN00681 | RMMUS081227 | | | RCPIC025250 | | RALGO2419 |
| RPSIN00450 | RMMUS002595 | | | RCPIC04004 | RGOAS0327 | RALGO68674 |
| RPSIN02431 | RMMUS1771 | RNFUR0203 | RDR040564 | RCPIC01328 | RGOAS0259 | RALGO5327 |
| | RMMUS252893 | | | | | |
| RPSIN01392 | RMMUS021444 | | | RCPIC02036 | RGOAS0281 | RALGO2125 |
| RPSIN01554 | RMMUS1423 | RNFUR0536 | RDR026106 | RCPIC006319 | | RALGO59654 |
| RPSIN00762 | RMMUS2063 | RNFUR0429 | RDR003805 | RCPIC01837 | RGOAS0084 | RALGO28844 |
| RPSIN03316 | RMMUS276812 | | RDR009489 | RCPIC01285 | RGOAS0287 | RALGO55048 |
| RPSIN03733 | RMMUS2408 | RNFUR0399 | RDR002506 | RCPIC01753 | RGOAS0075 | RALGO1199 |
| RPSIN01278 | RMMUS1018 | RNFUR0033 | RDR030775 | RCPIC05122 | RGOAS0145 | RALGO60857 |
| | RMMUS253478 | | | RCPIC055768 | | RALGO1758 |
| RPSIN01035 | RMMUS2781 | RNFUR0469 | RDR010185 | RCPIC01919 | RGOAS0113 | not predicte |

| | | | | | | |
|---|---|---|---|---|---|---|
| RPSIN00396 | RMMUS1652 | RNFUR0299 | RDR002525 | RCPIC03077 | RGOAS0196 | RALGO4301 |
| RPSIN03753 | RMMUS1791 | RNFUR0193 | RDR010625 | RCPIC02835 | RGOAS0142 | RALGO23446 |
| RPSIN00570 | RMMUS0728 | RNFUR0476 | RDR037573 | RCPIC00486 | RGOAS0334 | RALGO39414 |
| RPSIN02927 | RMMUS2027 | RNFUR0474 | RDR033003 | RCPIC02937 | RGOAS0074 | RALGO60290 |
| 18 | RMMUS1144 | RNFUR0077 | RDR006879 | RCPIC05502 | RGOAS0071 | RALGO22617 |
| 69 | RMMUS1107 | RNFUR0438 | RDR034825 | RCPIC03921 | RGOAS0289 | RALGO01686 |
| RPSIN03299 | RMMUS0160 | RNFUR0024 | RDR025963 | RCPIC00612 | RGOAS0275 | RALGO60969 |
| RPSIN00560 | RMMUS0845 | RNFUR0070 | RDR010299 | RCPIC01587 | RGOAS0056 | RALGO07708 |
| RPSIN01905 | RMMUS2383 | RNFUR0286 | RDR005081 | RCPIC03692 | RGOAS0285 | RALGO28724 |
| RPSIN00732 | RMMUS2579 | RNFUR0430 | RDR034357 | RCPIC00701 | RGOAS0295 | RALGO63406 |
| RPSIN02186 | RMMUS0833 | RNFUR0303 | RDR009978 | RCPIC01585 | RGOAS0319 | RALGO05905 |
| RPSIN01892 | RMMUS0302 | RNFUR0265 | RDR046394 | RCPIC00802 | RGOAS0164 | RALGO49421 |
| RPSIN03438 | RMMUS2815 | RNFUR0469 | RDR010078 | RCPIC01217 | RGOAS0282 | RALGO50456 |
| RPSIN00332 | RMMUS2202 | RNFUR0095 | RDR053098 | RCPIC05313 | RGOAS0075 | RALGO42098 |
| RPSIN00996 | RMMUS1994 | RNFUR0278 | RDR034390 | RCPIC03029 | RGOAS0263 | RALGO56922 |
| RPSIN00379 | RMMUS2291 | RNFUR0355 | RDR036457 | RCPIC04754 | RGOAS0065 | RALGO69070 |
| | RMMUS144794 | | | | | RALGO10504 |
| RPSIN02015 | RMMUS079748 | | | RCPIC02527 | RGOAS0244 | RALGO16444 |
| RPSIN02829 | RMMUS1271 | RNFUR0520 | RDR015056 | RCPIC038306 | | RALGO33355 |
| RPSIN03652 | RMMUS2706 | RNFUR0408 | RDR010141 | RCPIC01128 | RGOAS0027 | RALGO54061 |
| RPSIN03115 | RMMUS080581 | | RDR020619 | RCPIC02512 | RGOAS0186 | RALGO19168 |
| RPSIN03391 | RMMUS2101 | RNFUR0471 | RDR017066 | RCPIC02128 | RGOAS0296 | RALGO51832 |
| RPSIN01465 | RMMUS1402 | RNFUR0398 | RDR015804 | RCPIC02664 | RGOAS0322 | RALGO36308 |
| RPSIN00678 | RMMUS0200 | RNFUR0433 | RDR045333 | RCPIC002756 | | RALGO32848 |
| RPSIN02559 | RMMUS0780 | RNFUR0012 | RDR001092 | RCPIC04295 | RGOAS0137 | RALGO54562 |
| RPSIN03199 | RMMUS262306 | | RDR020843 | RCPIC03920 | RGOAS0334 | RALGO47515 |
| RPSIN01959 | RMMUS1659 | RNFUR0511 | RDR023871 | RCPIC03791 | RGOAS0056 | RALGO39819 |
| 53 | RMMUS2201 | RNFUR0061 | RDR007359 | RCPIC02180 | RGOAS0189 | RALGO41924 |
| 62 | RMMUS2850 | RNFUR0380 | RDR017360 | RCPIC05179 | RGOAS0141 | RALGO24031 |
| RPSIN00517 | RMMUS0980 | RNFUR0035 | RDR001282 | RCPIC05444 | RGOAS0098 | RALGO65842 |
| RPSIN01509 | RMMUS147948 | | | RCPIC01446 | RGOAS0127 | RALGO36815 |
| RPSIN00665 | RMMUS015095 | | | RCPIC04588 | RGOAS0068 | RALGO45053 |
| RPSIN01509 | RMMUS1479 | RNFUR0032 | RDR030892 | RCPIC01446 | RGOAS0127 | RALGO48571 |
| RPSIN01509 | RMMUS147964 | | | RCPIC014501 | | RALGO36782 |
| RPSIN02859 | RMMUS1928 | RNFUR0391 | RDR034372 | RCPIC00601 | RGOAS0285 | RALGO23491 |
| RPSIN01278 | RMMUS101892 | | | RCPIC05121 | RGOAS0145 | RALGO60854 |
| RPSIN01195 | RMMUS2559 | RNFUR0285 | RDR043494 | RCPIC00830 | RGOAS0124 | RALGO21450 |
| | | | | | | not predicte |
| | | | | | | not predicte |
| | | | | | | not predicte |
| | | | | | | not predicte |
| | | | | | | not predicte |
| | | | | | | not predicte |
| | | | | | | not predicte |
| RPSIN02219 | RMMUS012918 | | | RCPIC012113 | | not predicte |

| | | | | | | |
|---|---|---|---|---|---|---|
| | RMMUS109109 | | RDR031088 | | | not predicted |
| | RMMUS109100 | | | | | not predicted |
| | RMMUS147955 | | | RCPIC01459 | RGOAS01273 | RALGO36803 |
| | RMMUS109105 | | | | | not predicted |
| RPSIN026004 | RMMUS2415 | RNFUR02337 | RDR045114 | RCPIC04508 | RGOAS02237 | RALGO01011 |
| | RMMUS109117 | | | | | not predicted |
| RPSIN027777 | RMMUS1715 | RNFUR02068 | RDR013999 | RCPIC04789 | RGOAS00920 | RALGO15029 |
| RPSIN000543 | RMMUS1892 | RNFUR009337 | | RCPIC04024 | RGOAS03226 | RALGO20970 |
| | | | | | | not predicted |
| | RMMUS232384 | | | | | not predicted |
| 35 | | | RDR015044 | | | not predicted |
| | | | | | | not predicted |
| | | | | | | not predicted |
| RPSIN036752 | RMMUS2321 | RNFUR049172 | | RCPIC04660 | RGOAS03428 | RALGO54643 |
| RPSIN000334 | RMMUS103804 | | RDR002386 | RCPIC03295 | RGOAS03133 | RALGO52700 |
| 63 | RMMUS253350 | | | RCPIC05576 | RGOAS00204 | RALGO02275 |
| RPSIN003137 | RMMUS2814 | RNFUR04346 | RDR009574 | RCPIC03144 | RGOAS00248 | RALGO67663 |
| RPSIN024705 | RMMUS2451 | RNFUR04924 | RDR000538 | RCPIC02375 | RGOAS02118 | RALGO66058 |
| RPSIN006183 | RMMUS0567 | RNFUR00348 | RDR000726 | RCPIC03583 | RGOAS01654 | RALGO60041 |
| RPSIN022470 | RMMUS2609 | RNFUR01401 | RDR037748 | RCPIC02162 | RGOAS00126 | RALGO67192 |
| RPSIN003085 | RMMUS1518 | RNFUR04980 | RDR014638 | RCPIC01804 | RGOAS02462 | RALGO43148 |
| RPSIN009311 | RMMUS0297 | RNFUR02411 | RDR014332 | RCPIC00833 | RGOAS01333 | not predicted |
| RPSIN024289 | RMMUS2633 | RNFUR053183 | | RCPIC01244 | RGOAS01095 | RALGO53075 |
| | RMMUS234268 | | | | | RALGO59390 |
| RPSIN020771 | RMMUS104060 | | RDR005139 | RCPIC03291 | RGOAS00298 | RALGO52469 |
| RPSIN030811 | RMMUS1470 | RNFUR04395 | RDR011125 | RCPIC01457 | RGOAS03076 | RALGO33553 |
| RPSIN038245 | RMMUS0678 | RNFUR03789 | RDR032505 | RCPIC04924 | RGOAS01266 | RALGO12764 |
| RPSIN006417 | RMMUS246953 | | RDR038250 | RCPIC00657 | RGOAS03198 | RALGO57941 |
| RPSIN021540 | RMMUS2355 | RNFUR00008 | RDR007137 | RCPIC00051 | RGOAS02749 | RALGO43792 |
| RPSIN028832 | RMMUS0489 | RNFUR043903 | | RCPIC04592 | RGOAS01397 | RALGO44003 |
| RPSIN004810 | RMMUS1229 | RNFUR04464 | RDR046705 | RCPIC03382 | RGOAS03245 | RALGO43236 |
| RPSIN010927 | RMMUS152229 | | RDR047153 | RCPIC017970 | | RALGO44303 |
| RPSIN008130 | RMMUS180892 | | RDR014922 | | | RALGO26125 |
| RPSIN035552 | RMMUS1210 | RNFUR022741 | | RCPIC05584 | RGOAS02119 | RALGO71689 |
| RPSIN000374 | RMMUS1834 | RNFUR02700 | RDR005563 | | RGOAS01516 | RALGO13880 |
| RPSIN011093 | RMMUS1919 | RNFUR00299 | RDR022407 | RCPIC04084 | RGOAS02365 | RALGO39702 |
| RPSIN010037 | RMMUS0364 | RNFUR03630 | RDR044043 | RCPIC04416 | RGOAS02684 | RALGO03352 |
| RPSIN033528 | RMMUS1665 | RNFUR01001 | RDR043006 | RCPIC04150 | RGOAS00086 | RALGO37364 |
| RPSIN027701 | RMMUS2365 | RNFUR045124 | | RCPIC02739 | RGOAS01194 | RALGO13109 |
| RPSIN012167 | RMMUS0272 | RNFUR05021 | RDR019643 | RCPIC03509 | RGOAS00684 | RALGO33262 |
| RPSIN021733 | RMMUS0175 | RNFUR01612 | RDR038364 | RCPIC03396 | RGOAS00752 | RALGO64426 |
| RPSIN001801 | RMMUS2333 | RNFUR00616 | RDR013126 | RCPIC04650 | RGOAS01540 | RALGO53959 |
| RPSIN030014 | RMMUS1144 | RNFUR01558 | RDR030680 | RCPIC05502 | RGOAS00077 | RALGO10330 |
| RPSIN035689 | RMMUS0149 | RNFUR05412 | RDR038122 | RCPIC03639 | RGOAS02984 | RALGO46527 |
| 26 | RMMUS2529 | RNFUR03097 | RDR048034 | RCPIC05547 | RGOAS00131 | RALGO01927 |

| | | | | | | |
|---|---|---|---|---|---|---|
| RPSIN02598 | RMMUS2374 | RNFUR0514 | RDR043348 | RCPIC04336 | RGOAS0273 | RALGO1879 |
| RPSIN02764 | RMMUS2369 | RNFUR0182 | RDR020716 | RCPIC02740 | RGOAS0288 | RALGO1362 |
| RPSIN03195 | RMMUS2638 | RNFUR0074 | RDR031059 | RCPIC02109 | RGOAS0013 | RALGO4838 |
| RPSIN02746 | RMMUS0080 | RNFUR0038 | RDR028969 | RCPIC01805 | RGOAS0016 | RALGO39282 |
| RPSIN02188 | RMMUS0620 | RNFUR0547 | RDR012083 | RCPIC01273 | RGOAS0005 | RALGO19952 |
| RPSIN01083 | RMMUS1621 | RNFUR0431 | RDR030779 | RCPIC02791 | RGOAS0054 | RALGO50462 |
| RPSIN00489 | RMMUS0063 | RNFUR0547 | RDR012307 | RCPIC04514 | RGOAS0066 | RALGO4958 |
| RPSIN02289 | RMMUS0056 | RNFUR0075 | RDR000227 | RCPIC032535 | | RALGO13841 |
| RPSIN01747 | RMMUS1488 | RNFUR0327 | RDR029484 | RCPIC00229 | RGOAS0068 | RALGO7094 |
| RPSIN02988 | RMMUS0499 | RNFUR0040 | RDR000546 | RCPIC00032 | RGOAS0292 | RALGO63781 |
| 02 | RMMUS2801 | RNFUR0198 | RDR043464 | RCPIC036524 | | RALGO24954 |
| RPSIN02139 | RMMUS129877 | | | RCPIC05168 | RGOAS0023 | RALGO6902 |
| RPSIN02093 | RMMUS2459 | RNFUR0250 | RDR018753 | RCPIC02378 | RGOAS0068 | RALGO6641 |
| RPSIN01242 | RMMUS0856 | RNFUR0414 | RDR009819 | RCPIC01660 | RGOAS0180 | RALGO11341 |
| RPSIN02991 | RMMUS2432 | RNFUR0470 | RDR010231 | RCPIC00034 | RGOAS0263 | RALGO63562 |
| RPSIN03438 | RMMUS2827 | RNFUR0237 | RDR027582 | RCPIC01214 | RGOAS0173 | RALGO5110 |
| RPSIN01826 | RMMUS154043 | | RDR004426 | RCPIC01837 | RGOAS0224 | RALGO3044 |
| RPSIN02304 | RMMUS1562 | RNFUR0172 | RDR042365 | RCPIC03741 | RGOAS0130 | RALGO2793 |
| RPSIN03429 | RMMUS2827 | RNFUR0054 | RDR056433 | RCPIC01213 | RGOAS0227 | RALGO51073 |
| RPSIN00561 | RMMUS0845 | RNFUR0250 | RDR011256 | RCPIC01619 | RGOAS0057 | RALGO07843 |
| | RMMUS169738 | | | | | not predicte |
| RPSIN01902 | RMMUS2506 | RNFUR0006 | RDR005946 | RCPIC01997 | RGOAS0306 | RALGO3362 |
| RPSIN00054 | RMMUS1892 | RNFUR0472 | RDR005032 | RCPIC04061 | RGOAS0322 | RALGO36512 |
| 13 | RMMUS0327 | RNFUR0095 | RDR017436 | RCPIC01387 | RGOAS0025 | RALGO3602 |
| RPSIN02576 | RMMUS2687 | RNFUR0261 | RDR014796 | RCPIC00999 | RGOAS0245 | RALGO1511 |
| RPSIN00109 | RMMUS0017 | RNFUR0074 | RDR026170 | RCPIC00609 | RGOAS0157 | RALGO3948 |
| 37 | RMMUS2533 | RNFUR0396 | RDR037299 | RCPIC055773 | | RALGO02281 |
| RPSIN01197 | RMMUS250860 | | | RCPIC02923 | RGOAS0250 | RALGO21501 |
| RPSIN01932 | RMMUS2599 | RNFUR0208 | RDR024316 | RCPIC00277 | RGOAS0229 | RALGO3166 |
| RPSIN02577 | RMMUS2679 | RNFUR0250 | RDR040141 | RCPIC00951 | RGOAS0297 | RALGO1535 |
| RPSIN00985 | RMMUS1992 | RNFUR0016 | RDR038756 | RCPIC030167 | | RALGO56781 |
| RPSIN02848 | RMMUS1149 | RNFUR0077 | RDR005033 | RCPIC053665 | | RALGO2218 |
| RPSIN01322 | RMMUS0537 | RNFUR0405 | RDR003971 | RCPIC03951 | RGOAS0221 | RALGO29261 |
| RPSIN01578 | RMMUS1745 | RNFUR0139 | RDR025761 | RCPIC02702 | RGOAS0189 | RALGO65701 |
| RPSIN01794 | RMMUS2014 | RNFUR0428 | RDR033048 | RCPIC03020 | RGOAS0328 | RALGO59132 |
| RPSIN01676 | RMMUS0096 | RNFUR0164 | RDR002992 | RCPIC03437 | RGOAS0228 | RALGO64438 |
| RPSIN03839 | RMMUS153445 | | | RCPIC02563 | RGOAS0328 | RALGO46614 |
| RPSIN02612 | RMMUS0363 | RNFUR0457 | RDR043606 | RCPIC04423 | RGOAS0268 | RALGO0402 |
| | RMMUS152292 | | | RCPIC017966 | | RALGO44399 |
| RPSIN00866 | RMMUS033617 | | | RCPIC016670 | | RALGO6694 |
| 08 | RMMUS1146 | RNFUR0124 | RDR006588 | RCPIC052646 | | RALGO22572 |
| RPSIN03334 | RMMUS1015 | RNFUR0101 | RDR034545 | RCPIC00020 | RGOAS0001 | RALGO57434 |
| RPSIN00703 | RMMUS1037 | RNFUR0121 | RDR034942 | RCPIC00054 | RGOAS0170 | RALGO60017 |
| RPSIN01132 | RMMUS050243 | | | RCPIC01575 | RGOAS0106 | RALGO1292 |
| | RMMUS221534 | | | | | |

| | | | | | | |
|---|---|---|---|---|---|---|
| 61 | RMMUS1091 | RNFUR0072 | RDR032666 | RCPIC039941 | | RALGO48544 |
| RPSIN01587( | RMMUS2410 | RNFUR0105: | RDR012625 | RCPIC04748 | RGOAS0014: | RALGO07003 |
| RPSIN01736( | RMMUS0851 | RNFUR0138: | RDR010266 | RCPIC01589 | RGOAS0163: | RCABP03001 |
| RPSIN01858{ | RMMUS0495 | RNFUR0167 | RDR000062 | RCPIC00092 | RGOAS0198 | RALGO65371 |
| RPSIN03124 | RMMUS2762 | RNFUR0366! | RDR042351 | RCPIC03176 | RGOAS0038 | RALGO67303 |
| RPSIN02024( | RMMUS1644 | RNFUR0291: | RDR040263 | RCPIC04161 | RGOAS0335 | RALGO23542 |
| RPSIN03089{ | RMMUS2012 | RNFUR0520( | RDR033561 | RCPIC03021 | RGOAS0011 | RALGO58844 |
| RPSIN0017 | RMMUS0310 | RNFUR0232( | RDR014682 | RCPIC008207 | | RALGO45303 |
| RPSIN02063( | RMMUS1874 | RNFUR0158{ | RDR024383 | RCPIC04763 | RGOAS0095 | RALGO01215 |
| RPSIN02317 | RMMUS1050 | RNFUR0521 | RDR034513 | RCPIC00046 | RGOAS0178{ | RALGO5882C |
| RPSIN0333 | RMMUS101566 | | | RCPIC00084 | RGOAS0002 | not predictec |
| RPSIN02621( | RMMUS2367 | RNFUR0280 | RDR012582 | RCPIC00345 | RGOAS0268 | not predictec |
| 53 | RMMUS1430 | RNFUR0532( | RDR040390 | | RGOAS018409 | |
| RPSIN0008 | RMMUS143103 | | | RCPIC001096 | | RALGO71035 |
| 09 | RMMUS1910 | RNFUR0326 | RDR024324 | RCPIC00563 | RGOAS0070 | RALGO6655C |
| RPSIN0074 | RMMUS222990 | | RDR037896 | RCPIC00634 | RGOAS0279 | RALGO12878 |
| RPSIN02886{ | RMMUS1183 | RNFUR0148( | RDR035495 | RCPIC02587 | RGOAS0068( | RALGO50578 |
| RPSIN00400 | RMMUS272591 | | RDR026210 | RCPIC02641 | RGOAS0283( | RALGO31762 |
| 60 | RMMUS2070 | RNFUR009993 | | RCPIC03451 | RGOAS0205( | RALGO64852 |
| RPSIN00299{ | RMMUS2185 | RNFUR0448{ | RDR026470 | RCPIC02139 | RGOAS0235 | RALGO15641 |
| RPSIN03520( | RMMUS1516 | RNFUR0284( | RDR047291 | RCPIC018452 | | RALGO41308 |
| RPSIN01202 | RMMUS048899 | | | RCPIC04614 | RGOAS0331{ | RALGO44063 |
| RPSIN00062( | RMMUS0091 | RNFUR0286! | RDR012930 | RCPIC01779 | RGOAS0052 | RALGO42434 |
| RPSIN03727 | RMMUS0378 | RNFUR040813 | | RCPIC04412 | RGOAS0268( | RALGO02701 |
| RPSIN03052 | RMMUS0928 | RNFUR0063 | RDR007463 | RCPIC00041 | RGOAS0331 | RALGO56208 |
| RPSIN02055 | RMMUS1875 | RNFUR015815 | | RCPIC04766 | RGOAS0244 | RALGO00693 |
| RPSIN02005 | RMMUS0899 | RNFUR0426: | RDR029128 | RCPIC04480 | RGOAS0301 | RALGO70687 |
| RPSIN01794( | RMMUS0216 | RNFUR0453: | RDR010632 | RCPIC02021 | RGOAS0225( | RALGO19874 |
| RPSIN01280! | RMMUS2518 | RNFUR0076( | RDR004830 | RCPIC04511 | RGOAS0144: | RALGO22704 |
| RPSIN0072 | RMMUS0754 | RNFUR0374 | RDR032779 | RCPIC00642 | RGOAS0090: | RALGO63268 |
| RPSIN03767 | RMMUS1842 | RNFUR0140! | RDR030225 | RCPIC02865 | RGOAS0042 | RALGO21522 |
| RPSIN01052( | RMMUS2692 | RNFUR0525: | RDR039758 | RCPIC00937 | RGOAS021368 | |
| RPSIN03536( | RMMUS076951 | | | RCPIC02299 | RGOAS0160 | RALGO57181 |
| 68 | RMMUS064402 | | | RCPIC05023 | RGOAS0256 | RALGO04566 |
| RPSIN02159 | RMMUS2515 | RNFUR0149{ | RDR025610 | RCPIC00682 | RGOAS0107 | RALGO6272C |
| | RMMUS245009 | | RDR051754 | | | not predictec |
| RPSIN03599 | RMMUS2836 | RNFUR0172 | RDR053113 | RCPIC03681 | RGOAS0230{ | RALGO45825 |
| 14 | RMMUS243413 | | | RCPIC00090 | RGOAS0009( | RALGO64936 |
| | RMMUS252751 | | | | | RALGO10369 |
| RPSIN02444 | RMMUS0318 | RNFUR0493 | RDR034377 | RCPIC027644 | | RALGO53342 |
| RPSIN03420 | RMMUS1260 | RNFUR0160 | RDR024396 | RCPIC02283 | RGOAS0194( | RALGO38066 |
| RPSIN01440 | RMMUS1373 | RNFUR0174: | RDR014221 | RCPIC01504 | RGOAS0156 | RALGO20904 |
| RPSIN00631( | RMMUS1128 | RNFUR0374: | RDR046552 | RCPIC01530 | RGOAS0120 | RALGO25146 |
| RPSIN01514 | RMMUS068584 | | | | RGOAS0227( | RALGO21069 |
| RPSIN03161 | RMMUS068576 | | | RCPIC01683 | RGOAS0227 | RALGO21054 |

| | | | | | | |
|---|---|---|---|---|---|---|
| RPSIN034162 | RMMUS1812 | RNFUR05388 | RDR009524 | RCPIC02832 | RGOAS01684 | RALGO10402 |
| | RMMUS0685 | RNFUR02954 | RDR039660 | RCPIC01683 | RGOAS02277 | RALGO21066 |
| RPSIN001387 | RMMUS1471 | RNFUR04587 | RDR028994 | RCPIC01452 | RGOAS02918 | RALGO34168 |
| RPSIN026641 | RMMUS102166 | | | RCPIC02223 | RGOAS02708 | RALGO63087 |
| RPSIN005362 | RMMUS253089 | | | RCPIC055461 | | RALGO02104 |
| RPSIN038326 | RMMUS034091 | | RDR025899 | RCPIC02359 | RGOAS02019 | RALGO60554 |
| RPSIN019290 | RMMUS2569 | RNFUR04059 | RDR000528 | RCPIC04800 | RGOAS00964 | RALGO07114 |
| RPSIN012762 | RMMUS101987 | | | RCPIC00069 | RGOAS00027 | RALGO60815 |
| | RMMUS010751 | | | RCPIC00867 | RGOAS01721 | RALGO16837 |
| | | | | | | not predicted |
| RPSIN013884 | RMMUS0238 | RNFUR04640 | RDR010340 | RCPIC02063 | RGOAS00572 | RALGO22986 |
| 93 | RMMUS1082 | RNFUR00811 | RDR021460 | RCPIC03923 | RGOAS00812 | RALGO00864 |
| | | | | | | not predicted |
| RPSIN038674 | RMMUS2269 | RNFUR00641 | RDR040391 | RCPIC04777 | RGOAS01158 | RALGO29006 |
| RPSIN038673 | RMMUS2269 | RNFUR03269 | RDR040423 | RCPIC04776 | RGOAS01158 | RALGO28967 |
| RPSIN003606 | RMMUS0882 | RNFUR04604 | RDR046288 | RCPIC01854 | RGOAS01984 | RALGO14696 |
| RPSIN030600 | RMMUS0626 | RNFUR01173 | RDR003509 | RCPIC01305 | RGOAS00297 | RALGO17056 |
| RPSIN030601 | RMMUS0626 | RNFUR01162 | RDR002692 | RCPIC013056 | | not predicted |
| RPSIN032334 | RMMUS0209 | RNFUR02401 | RDR045339 | RCPIC04637 | RGOAS01184 | not predicted |
| RPSIN004103 | RMMUS1309 | RNFUR02880 | RDR009491 | RCPIC05415 | RGOAS01987 | RALGO63949 |
| RPSIN038664 | RMMUS2270 | RNFUR01748 | RDR010063 | RCPIC04777 | RGOAS02886 | not predicted |
| RPSIN033337 | RMMUS1011 | RNFUR01860 | RDR007397 | RCPIC00168 | RGOAS03130 | RALGO57374 |
| RPSIN025918 | RMMUS2177 | RNFUR03037 | RDR023480 | RCPIC01035 | RGOAS02205 | RALGO17511 |
| 03 | RMMUS2361 | RNFUR04489 | RDR043890 | RCPIC02740 | RGOAS00907 | RALGO13853 |
| RPSIN020503 | RMMUS0252 | RNFUR01317 | RDR018965 | RCPIC03492 | RGOAS01678 | RALGO40698 |
| RPSIN015108 | RMMUS1480 | RNFUR01667 | RDR040681 | RCPIC01443 | RGOAS03057 | RALGO36125 |
| RPSIN013433 | RMMUS185769 | | | RCPIC04902 | RGOAS00298 | RALGO08416 |
| RPSIN002311 | RMMUS1371 | RNFUR05068 | RDR041949 | RCPIC04863 | RGOAS00088 | RALGO08847 |
| RPSIN031945 | RMMUS2621 | RNFUR01184 | RDR009511 | RCPIC03905 | RGOAS01803 | RALGO39717 |
| RPSIN005596 | RMMUS156447 | | | RCPIC00420 | RGOAS00093 | RALGO08970 |
| RPSIN000714 | RMMUS0712 | RNFUR04862 | RDR044042 | RCPIC03146 | RGOAS02125 | RALGO00534 |
| RPSIN012583 | RMMUS1911 | RNFUR04951 | RDR005230 | RCPIC05350 | RGOAS03321 | RALGO39261 |
| RPSIN003564 | RMMUS0885 | RNFUR00630 | RDR006117 | RCPIC018990 | | RALGO12758 |
| 36 | RMMUS1914 | RNFUR04707 | RDR013879 | RCPIC04120 | RGOAS011762 | |
| | | | | | | not predicted |
| | | | | | | not predicted |
| | RMMUS226953 | | | | | not predicted |
| | | | | | | not predicted |
| RPSIN013901 | RMMUS021295 | | RDR009775 | RCPIC02050 | RGOAS00140 | RALGO21714 |
| RPSIN032511 | RMMUS128191 | | RDR014560 | RCPIC03081 | RGOAS03413 | RALGO38342 |
| RPSIN024463 | RMMUS031930 | | RDR048648 | RCPIC00912 | RGOAS01635 | RALGO53072 |

| | | | | | | |
|---|---|---|---|---|---|---|
| RPSIN03084 | RMMUS0042 | RNFUR0107 | RDR012886 | RCPIC01422 | RGOAS0325 | RALGO22911 |
| RPSIN01886 | RMMUS0306 | RNFUR0362 | RDR046690 | RCPIC00813 | RGOAS0304 | RALGO47956 |
| 58 | RMMUS0909 | RNFUR0131 | RDR046787 | RCPIC002618 | | RALGO3997 |
| RPSIN00484 | RMMUS1392 | RNFUR0297 | RDR034829 | RCPIC01518 | RGOAS0063 | RALGO11458 |
| RPSIN00145 | RMMUS1471 | RNFUR0452 | RDR028938 | RCPIC01452 | RGOAS0292 | RALGO3420 |
| RPSIN00869 | RMMUS243885 | | RDR021142 | RCPIC00008 | RGOAS0168 | RALGO64660 |
| | RMMUS109150 | | | | | |
| RPSIN00636 | RMMUS159110 | | RDR033285 | RCPIC00689 | RGOAS0166 | RALGO5739 |
| RPSIN01675 | RMMUS1641 | RNFUR0513 | RDR024368 | RCPIC02144 | RGOAS0034 | RALGO1903 |
| RPSIN01573 | RMMUS1746 | RNFUR0440 | RDR012125 | RCPIC04441 | RGOAS0292 | RALGO65638 |
| RPSIN03342 | RMMUS200549 | | RDR033729 | RCPIC03031 | RGOAS0106 | RALGO5829 |
| | RMMUS183506 | | | | | not predicte |
| | RMMUS183506 | | | | | not predicte |
| | RMMUS183506 | | | | | not predicte |
| | RMMUS183506 | | | | | not predicte |
| | RMMUS138354 | | | | RGOAS0060 | RALGO0958 |
| | RMMUS183506 | | | | | not predicte |
| RPSIN01270 | RMMUS0072 | RNFUR0365 | RDR045197 | RCPIC01943 | RGOAS0142 | RALGO31687 |
| | | RNFUR019663 | | | | RALGO05001 |
| | RMMUS183834 | | | RCPIC02834 | RGOAS0144 | RALGO04947 |
| RPSIN0075 | RMMUS183812 | | | RCPIC02911 | RGOAS0144 | RALGO04974 |
| | | | | | | not predicte |
| | | | | | | not predicte |
| RPSIN01002 | RMMUS2201 | RNFUR0012 | RDR016636 | RCPIC05118 | RGOAS0006 | RALGO09672 |
| RPSIN00174 | RMMUS1286 | RNFUR0229 | RDR014775 | RCPIC00792 | RGOAS0140 | RALGO47136 |
| RPSIN01698 | RMMUS1028 | RNFUR0512 | RDR046549 | RCPIC00153 | RGOAS0150 | RALGO60617 |
| RPSIN02668 | RMMUS255630 | | | RCPIC008289 | | RALGO20651 |
| RPSIN01441 | RMMUS1130 | RNFUR0359 | RDR050989 | RCPIC01505 | RGOAS0099 | RALGO2266 |
| RPSIN02921 | RMMUS2120 | RNFUR0438 | RDR009490 | RCPIC02397 | RGOAS0001 | RALGO4247 |
| | RMMUS158844 | | | | | not predicte |
| | RMMUS158834 | | | | | not predicte |
| RPSIN01518 | RMMUS157531 | | | RCPIC00248 | RGOAS0149 | RALGO5167 |
| RPSIN02331 | RMMUS2474 | RNFUR0074 | RDR038106 | RCPIC00700 | RGOAS0022 | RALGO59057 |
| RPSIN00543 | RMMUS1867 | RNFUR0109 | RDR012339 | RCPIC04895 | RGOAS0283 | RALGO03684 |
| | RMMUS262854 | | | | | not predicte |
| 04 | RMMUS1618 | RNFUR0024 | RDR037935 | RCPIC02801 | RGOAS0170 | RALGO51082 |
| RPSIN01176 | RMMUS213334 | | RDR010012 | RCPIC042381 | | RALGO3677 |
| 41 | RMMUS124524 | | | RCPIC055300 | | RALGO0019 |
| RPSIN00612 | RMMUS046594 | | | RCPIC05577 | RGOAS0112 | RALGO01437 |
| RPSIN00870 | RMMUS2434 | RNFUR0531 | RDR000817 | RCPIC00010 | RGOAS0008 | RALGO64927 |
| | RMMUS097269 | | | RCPIC055367 | | RALGO61251 |
| | RMMUS105993 | | | | RGOAS0028 | not predicte |
| RPSIN00014 | RMMUS112170 | | RDR002090 | RCPIC05650 | RGOAS0038 | RALGO4710 |
| RPSIN01476 | RMMUS2671 | RNFUR0282 | RDR045542 | RCPIC03435 | RGOAS0282 | RALGO65908 |
| RPSIN01870 | RMMUS0145 | RNFUR0344 | RDR018944 | RCPIC00012 | RGOAS0206 | RALGO6526 |

| | | | | | | |
|---|---|---|---|---|---|---|
| | RMMUS102595 | | RDR032623 | | | not predicted |
| RPSIN006592 | RMMUS2714 | RNFUR05433 | RDR040582 | RCPIC01118 | RGOAS02146 | RALGO52607 |
| RPSIN028486 | RMMUS004721 | | | RCPIC01420 | RGOAS01791 | RALGO21024 |
| | RMMUS189937 | | | | RGOAS03254 | not predicted |
| | RMMUS189992 | | | RCPIC040347 | | not predicted |
| RPSIN031815 | RMMUS1899 | RNFUR014724 | | | RGOAS00143 | RALGO32512 |
| 05 | RMMUS190005 | | | RCPIC04068 | RGOAS03254 | RALGO32461 |
| RPSIN002010 | RMMUS0259 | RNFUR04904 | RDR020453 | RCPIC03519 | RGOAS01070 | RALGO38579 |
| RPSIN035716 | RMMUS017899 | | | RCPIC00832 | RGOAS01720 | RALGO16696 |
| RPSIN009121 | RMMUS0736 | RNFUR02252 | RDR006930 | RCPIC03212 | RGOAS01035 | RALGO25620 |
| | RMMUS114851 | | | | RGOAS00321 | RALGO22152 |
| RPSIN035383 | RMMUS0769 | RNFUR03250 | RDR046170 | RCPIC02300 | RGOAS01600 | RALGO57175 |
| RPSIN014099 | RMMUS147664 | | | RCPIC01460 | RGOAS02647 | RALGO37346 |
| RPSIN014120 | RMMUS147655 | | | RCPIC01447 | RGOAS02647 | not predicted |
| RPSIN018502 | RMMUS102325 | | | RCPIC02221 | RGOAS02818 | RALGO63138 |
| RPSIN006980 | RMMUS279046 | | | RCPIC01290 | RGOAS02715 | RALGO54478 |
| RPSIN018931 | RMMUS030546 | | RDR047435 | RCPIC00848 | RGOAS02568 | RALGO48205 |
| | RMMUS0179 | RNFUR016423 | | | | not predicted |
| | RMMUS0179 | RNFUR016423 | | | | not predicted |
| RPSIN035721 | RMMUS010736 | | | RCPIC00868 | RGOAS01322 | RALGO16987 |
| 70 | | RNFUR01866 | RDR020223 | RCPIC00105 | RGOAS01754 | RALGO63766 |
| RPSIN022667 | RMMUS0306 | RNFUR046524 | | RCPIC00814 | RGOAS02410 | RALGO47677 |
| | RMMUS046633 | | | | | not predicted |
| RPSIN000261 | RMMUS2697 | RNFUR036153 | | RCPIC02698 | RGOAS01670 | RALGO31116 |
| RPSIN036307 | RMMUS1719 | RNFUR04228 | RDR043639 | RCPIC027415 | | RALGO14294 |
| RPSIN019115 | RMMUS103159 | | | RCPIC044402 | | RALGO04473 |
| RPSIN037083 | RMMUS2645 | RNFUR03359 | RDR027786 | RCPIC01263 | RGOAS00357 | RALGO55313 |
| 25 | RMMUS1099 | RNFUR00683 | RDR031120 | RCPIC05696 | RGOAS01702 | RALGO48451 |
| RPSIN014705 | RMMUS1888 | RNFUR04366 | RDR052205 | RCPIC04964 | RGOAS00020 | not predicted |
| 72 | RMMUS242818 | | | RCPIC00074 | RGOAS01072 | RALGO65359 |
| RPSIN032521 | RMMUS1280 | RNFUR00912 | RDR014791 | RCPIC01766 | RGOAS01476 | RALGO37823 |
| RPSIN015312 | RMMUS207455 | | | RCPIC03446 | RGOAS01788 | RALGO65314 |
| | RMMUS242516 | | | | | not predicted |
| RPSIN001973 | RMMUS096352 | | | RCPIC05372 | RGOAS02579 | RALGO59402 |
| RPSIN008743 | RMMUS243581 | | | RCPIC00088 | RGOAS02108 | RALGO64855 |
| 30 | RMMUS1366 | RNFUR02615 | RDR055635 | RCPIC04913 | RGOAS03376 | RALGO04422 |
| RPSIN035141 | RMMUS1443 | RNFUR01483 | RDR000647 | RCPIC05427 | RGOAS02386 | RALGO11476 |
| RPSIN005521 | RMMUS0686 | RNFUR04341 | RDR022763 | RCPIC05002 | RGOAS01270 | RALGO22038 |
| RPSIN025116 | RMMUS0124 | RNFUR03101 | RDR002712 | RCPIC03944 | RGOAS02263 | RALGO39171 |
| RPSIN015903 | RMMUS1398 | RNFUR04336 | RDR014818 | | RGOAS01203 | RALGO34847 |
| RPSIN026841 | RMMUS1388 | RNFUR01977 | RDR014781 | RCPIC01516 | RGOAS01260 | RALGO28010 |
| RPSIN022490 | RMMUS2610 | RNFUR01312 | RDR038264 | RCPIC02162 | RGOAS03325 | RALGO67210 |
| | RMMUS139854 | | | RCPIC015432 | | |
| RPSIN021151 | RMMUS0406 | RNFUR01904 | RDR022706 | RCPIC02965 | RGOAS00718 | RALGO11852 |
| RPSIN004558 | RMMUS1639 | RNFUR01370 | RDR033165 | RCPIC05540 | RGOAS00361 | RALGO19712 |

| | | | | | | | |
|---|---|---|---|---|---|---|---|
| | RPSIN021184 | RMMUS1865 | RNFUR00045 | RDR036668 | RCPIC04880 | RGOAS01843 | RALGO05353 |
| | RPSIN005705 | RMMUS072872 | | RDR032313 | RCPIC00353 | RGOAS01688 | RALGO38804 |
| 30 | | RMMUS1709 | RNFUR05071 | RDR022505 | RCPIC00472 | RGOAS02573 | RALGO16549 |
| | RPSIN005660 | RMMUS0727 | RNFUR05339 | RDR036868 | RCPIC00485 | RGOAS01901 | RALGO39828 |
| | RPSIN011704 | RMMUS2132 | RNFUR04613 | RDR010285 | RCPIC04262 | RGOAS00396 | RALGO37238 |
| | RPSIN011549 | RMMUS0232 | RNFUR00305 | RDR040155 | RCPIC02029 | RGOAS01575 | RALGO15152 |
| | RPSIN020016 | RMMUS2513 | RNFUR01269 | RDR041859 | RCPIC00844 | RGOAS03229 | RALGO24401 |
| | RPSIN007044 | RMMUS1037 | RNFUR04288 | RDR046790 | RCPIC00026 | RGOAS01428 | RALGO60134 |
| | RPSIN024860 | | | RDR000801 | RCPIC01105 | RGOAS02964 | RALGO56931 |
| | | RMMUS0613 | RNFUR005124 | | RCPIC011058 | | RALGO56931 |
| | RPSIN021435 | RMMUS063562 | | RDR017183 | RCPIC05007 | RGOAS01544 | RALGO12959 |
| 39 | | RMMUS1444 | RNFUR04897 | RDR012772 | | RGOAS002107 | |
| | RPSIN034358 | RMMUS2824 | RNFUR00357 | RDR027773 | RCPIC01309 | RGOAS02253 | RALGO50602 |
| | RPSIN006321 | RMMUS1127 | RNFUR00921 | RDR014823 | RCPIC01507 | RGOAS03062 | RALGO25647 |
| | RPSIN025495 | RMMUS039368 | | | RCPIC02994 | RGOAS03274 | RALGO18937 |
| | RPSIN006322 | RMMUS1127 | RNFUR00921 | RDR014824 | RCPIC01507 | RGOAS03062 | RALGO25705 |
| | RPSIN035160 | RMMUS0218 | RNFUR00932 | RDR041266 | RCPIC02038 | RGOAS027607 | |
| | RPSIN006315 | RMMUS1128 | RNFUR02907 | RDR015813 | RCPIC01506 | RGOAS01209 | RALGO25104 |
| | RPSIN001575 | RMMUS279861 | | RDR027992 | RCPIC03067 | RGOAS01969 | RALGO43031 |
| | RPSIN006044 | | RNFUR04502 | RDR034448 | RCPIC01526 | RGOAS03053 | RALGO10138 |
| | | RMMUS1656 | RNFUR03407 | RDR025716 | RCPIC05618 | RGOAS01814 | RALGO38951 |
| 96 | | RMMUS012425 | | RDR054594 | RCPIC03956 | RGOAS03301 | RALGO38045 |
| 97 | | | | | | | not predicted |
| | RPSIN030975 | RMMUS2767 | RNFUR02315 | RDR017022 | RCPIC04231 | RGOAS01289 | RALGO23244 |
| | RPSIN003089 | RMMUS1518 | RNFUR02160 | RDR015123 | RCPIC01809 | RGOAS00913 | RALGO43260 |
| | RPSIN000752 | RMMUS2564 | RNFUR01991 | RDR054584 | RCPIC04170 | RGOAS01812 | RALGO37595 |
| | RPSIN021082 | RMMUS0550 | RNFUR02156 | RDR014572 | RCPIC02662 | RGOAS02118 | RALGO67606 |
| | RPSIN008557 | RMMUS1460 | RNFUR04039 | RDR041028 | RCPIC01471 | RGOAS03214 | RALGO28904 |
| | RPSIN012371 | RMMUS0828 | RNFUR04528 | RDR010181 | RCPIC016002 | | RALGO00453 |
| | RPSIN030908 | RMMUS2007 | RNFUR00459 | RDR038625 | RCPIC03012 | RGOAS00421 | RALGO58481 |
| | RPSIN026688 | RMMUS0505 | RNFUR04440 | RDR043644 | RCPIC00828 | RGOAS03375 | RALGO20540 |
| | RPSIN004385 | RMMUS0787 | RNFUR03778 | RDR000651 | RCPIC04275 | RGOAS02810 | RALGO52940 |
| | RPSIN006234 | RMMUS1275 | RNFUR01401 | RDR046744 | RCPIC03827 | RGOAS03299 | RALGO36137 |
| | RPSIN030141 | RMMUS011613 | | | RCPIC00828 | RGOAS03372 | RALGO20447 |
| | RPSIN012101 | RMMUS0271 | RNFUR01240 | RDR019283 | | RGOAS02315 | RALGO33325 |
| | RPSIN036703 | RMMUS1672 | RNFUR03758 | RDR007472 | RCPIC04680 | RGOAS00245 | RALGO55499 |
| | RPSIN021375 | RMMUS2730 | RNFUR05166 | RDR026143 | RCPIC02466 | RGOAS02478 | RALGO01218 |
| | RPSIN023451 | RMMUS1102 | RNFUR00576 | RDR031399 | RCPIC05209 | RGOAS01526 | RALGO48355 |
| | RPSIN013704 | RMMUS2386 | RNFUR00173 | RDR005266 | RCPIC03700 | RGOAS03170 | RALGO24741 |
| | RPSIN022116 | RMMUS0529 | RNFUR00317 | RDR000615 | RCPIC03334 | RGOAS00031 | RALGO32323 |
| | | RMMUS2822 | RNFUR01816 | RDR014555 | RCPIC03064 | RGOAS03401 | RALGO43822 |
| | RPSIN003731 | RMMUS2676 | RNFUR00766 | RDR028691 | RCPIC02929 | RGOAS02848 | RALGO29243 |
| | | | | | | | not predicted |
| | | | | | | | not predicted |
| 78 | | RMMUS103146 | | | RCPIC05698 | RGOAS00294 | RALGO05154 |

| | | | | | | |
|---|---|---|---|---|---|---|
| | RMMUS092674 | | | RCPIC02229 | RGOAS0174 | RALGO62242 |
| RPSIN01735 | RMMUS0853 | RNFUR0518 | RDR011163 | RCPIC01606 | RGOAS003015 | |
| RPSIN03418 | RMMUS0866 | RNFUR0471 | RDR011945 | RCPIC02853 | RGOAS0322 | RALGO1184 |
| RPSIN02800 | RMMUS1118 | RNFUR0338 | RDR030657 | | RGOAS0200 | RALGO0613 |
| RPSIN00728 | RMMUS2579 | RNFUR030383 | | RCPIC00642 | RGOAS0295 | RALGO6339 |
| RPSIN02292 | RMMUS0473 | RNFUR0370 | RDR036109 | RCPIC05308 | RGOAS0237 | RALGO4141 |
| RPSIN02382 | RMMUS0767 | RNFUR0039 | RDR028359 | RCPIC022959 | | RALGO57467 |
| RPSIN00439 | RMMUS0788 | RNFUR0299 | RDR038260 | RCPIC04285 | RGOAS0116 | RALGO5273 |
| RPSIN00102 | RMMUS0281 | RNFUR0077 | RDR027760 | RCPIC01577 | RGOAS0166 | RALGO11257 |
| RPSIN00724 | RMMUS251200 | | | RCPIC036748 | | RALGO2368 |
| RPSIN03537 | RMMUS0771 | RNFUR0423 | RDR000292 | RCPIC02271 | RGOAS001696 | |
| RPSIN00656 | RMMUS1543 | RNFUR0420 | RDR003692 | RCPIC017360 | | RALGO35420 |
| RPSIN03767 | RMMUS1843 | RNFUR0390 | RDR029558 | RCPIC02844 | RGOAS0316 | RALGO22092 |
| RPSIN032375 | | RNFUR0352 | RDR029194 | RCPIC04167 | RGOAS0202 | RALGO21027 |
| RPSIN02948 | RMMUS2233 | RNFUR0049 | RDR038355 | RCPIC03664 | RGOAS0003 | RALGO06304 |
| RPSIN01622 | RMMUS2266 | RNFUR0348 | RDR018856 | RCPIC04683 | RGOAS0229 | RALGO69373 |
| | RMMUS222014 | | | | | not predicte |
| RPSIN01456 | RMMUS1045 | RNFUR0023 | RDR007061 | RCPIC00045 | RGOAS0271 | RALGO5862 |
| RPSIN01776 | RMMUS0832 | RNFUR0299 | RDR031440 | RCPIC01586 | RGOAS0330 | RALGO06634 |
| RPSIN02684 | RMMUS138974 | | RDR014236 | RCPIC01524 | RGOAS0261 | RALGO27317 |
| RPSIN00970 | RMMUS0981 | RNFUR0412 | RDR029559 | RCPIC00969 | RGOAS0132 | RALGO22092 |
| RPSIN01402 | RMMUS1058 | RNFUR0414 | RDR031653 | RCPIC05518 | RGOAS0161 | RALGO20774 |
| 58 | RMMUS0683 | RNFUR043107 | | RCPIC04935 | RGOAS0060 | RALGO1087 |
| RPSIN01391 | RMMUS0238 | RNFUR0464 | RDR011146 | RCPIC02049 | RGOAS0056 | RALGO2305 |
| RPSIN00816 | RMMUS1959 | RNFUR0283 | RDR032730 | RCPIC02757 | RGOAS0219 | RALGO55544 |
| 09 | RMMUS1855 | RNFUR0537 | RDR005064 | RCPIC04887 | RGOAS0294 | RALGO09984 |
| 14 | RMMUS2358 | RNFUR0190 | RDR022638 | RCPIC043351 | | RALGO18162 |
| RPSIN00691 | RMMUS003271 | | | RCPIC035026 | | RALGO28643 |
| RPSIN01860 | RMMUS2127 | RNFUR0394 | RDR009532 | RCPIC02404 | RGOAS0028 | RALGO3890 |
| RPSIN01391 | RMMUS0217 | RNFUR0359 | RDR041240 | RCPIC02037 | RGOAS0112 | RALGO20546 |
| RPSIN02287 | RMMUS008418 | | RDR020329 | RCPIC02102 | RGOAS0286 | RALGO07702 |
| RPSIN03809 | RMMUS2604 | RNFUR0121 | RDR003206 | RCPIC02164 | RGOAS0280 | RALGO66835 |
| RPSIN01713 | RMMUS1234 | RNFUR0191 | RDR012305 | RCPIC01798 | RGOAS0175 | RALGO1912 |
| RPSIN03246 | RMMUS1412 | RNFUR0503 | RDR034855 | RCPIC01514 | RGOAS0178 | RALGO31561 |
| 59 | RMMUS0797 | RNFUR0053 | RDR027661 | RCPIC02511 | RGOAS0244 | RALGO1657 |
| RPSIN03206 | RMMUS0022 | RNFUR0194 | RDR017490 | RCPIC03192 | RGOAS0139 | RALGO61994 |
| RPSIN01613 | RMMUS2284 | RNFUR0200 | RDR012430 | RCPIC05027 | RGOAS011907 | |
| RPSIN01325 | RMMUS091942 | | RDR046376 | RCPIC04356 | RGOAS0068 | RALGO02653 |
| RPSIN00762 | RMMUS042392 | | | RCPIC017861 | | RALGO28811 |
| RPSIN01023 | RMMUS2244 | RNFUR0455 | RDR000122 | RCPIC04476 | RGOAS029474 | |
| RPSIN035772 | | | | | | not predicte |
| | RMMUS017954 | | | | | RALGO1687 |
| RPSIN02496 | RMMUS0050 | RNFUR0248 | RDR000640 | RCPIC03260 | RGOAS0242 | RALGO1883 |
| RPSIN00929 | RMMUS0298 | RNFUR0498 | RDR014659 | RCPIC00807 | RGOAS0081 | RALGO50024 |
| RPSIN03048 | RMMUS0924 | RNFUR0188 | RDR007141 | RCPIC00040 | RGOAS0260 | RALGO5634 |

| | | | | | | |
|---|---|---|---|---|---|---|
| RPSIN026914 | RMMUS2379 | RNFUR00260 | RDR042225 | RCPIC01985 | RGOAS00913 | RALGO32953 |
| RPSIN007056 | RMMUS1037 | RNFUR02796 | RDR006941 | RCPIC00026 | RGOAS02882 | RALGO60098 |
| RPSIN033850 | RMMUS082473 | | | RCPIC04002 | RGOAS01638 | RALGO68385 |
| RPSIN034088 | RMMUS0345 | RNFUR02191 | RDR033183 | RCPIC00565 | RGOAS01002 | RALGO66436 |
| RPSIN031396 | RMMUS146825 | | RDR021415 | RCPIC014891 | | not predicted |
| RPSIN029150 | RMMUS1347 | RNFUR03472 | RDR003772 | RCPIC03775 | RGOAS02892 | RALGO40653 |
| | RMMUS118342 | | RDR015210 | | | |
| RPSIN036561 | RMMUS2517 | RNFUR02377 | RDR000563 | RCPIC03169 | RGOAS00379 | RALGO67114 |
| RPSIN001357 | RMMUS2408 | RNFUR04782 | RDR042357 | RCPIC01927 | RGOAS01912 | RALGO40746 |
| RPSIN029265 | RMMUS0507 | RNFUR05352 | RDR017783 | RCPIC00843 | RGOAS02808 | RALGO25482 |
| RPSIN037018 | RMMUS2646 | RNFUR02883 | RDR027789 | RCPIC01219 | RGOAS00936 | RALGO55217 |
| RPSIN009449 | RMMUS186652 | | | RCPIC04878 | RGOAS00755 | RALGO04518 |
| RPSIN031652 | RMMUS0660 | RNFUR02736 | RDR030976 | RCPIC02000 | RGOAS03280 | RALGO03786 |
| RPSIN019118 | RMMUS229384 | | RDR037326 | RCPIC047581 | | RALGO69169 |
| RPSIN011465 | RMMUS2330 | RNFUR02346 | RDR024429 | RCPIC04648 | RGOAS02740 | RALGO54208 |
| RPSIN029975 | RMMUS180094 | | RDR043393 | RCPIC00033 | RGOAS02624 | RALGO63607 |
| RPSIN031839 | RMMUS125231 | | | RCPIC04840 | RGOAS02203 | RALGO03708 |
| RPSIN005366 | RMMUS004741 | | RDR019433 | RCPIC01423 | RGOAS01791 | RALGO20985 |
| 64 | RMMUS0967 | RNFUR00284 | RDR000959 | RCPIC001417 | | RALGO60146 |
| 23 | RMMUS163654 | | | RCPIC026574 | | RALGO22803 |
| RPSIN028490 | RMMUS1582 | RNFUR00857 | RDR031161 | RCPIC00250 | RGOAS01962 | RALGO51262 |
| RPSIN013045 | RMMUS2280 | RNFUR01093 | RDR045435 | RCPIC03325 | RGOAS03022 | RALGO59981 |
| 93 | RMMUS222049 | | RDR043074 | RCPIC05397 | RGOAS02610 | RALGO34850 |
| RPSIN008007 | RMMUS2737 | RNFUR01448 | RDR025711 | RCPIC00920 | RGOAS02669 | RALGO15272 |
| RPSIN005341 | RMMUS087269 | | RDR021185 | RCPIC028540 | | RALGO09231 |
| RPSIN015193 | RMMUS158137 | | | RCPIC05629 | RGOAS00092 | RALGO51649 |
| RPSIN020217 | | | | RCPIC055115 | | RALGO16004 |
| RPSIN031172 | RMMUS0801 | RNFUR015408 | | RCPIC02509 | RGOAS02511 | RALGO17874 |
| RPSIN031186 | | | RDR010328 | RCPIC02511 | RGOAS00447 | RALGO17841 |
| RPSIN010437 | RMMUS1008 | RNFUR02388 | RDR026100 | RCPIC02466 | RGOAS00325 | RALGO11371 |
| RPSIN007741 | RMMUS2092 | RNFUR04119 | RDR027788 | RCPIC01254 | RGOAS02755 | RALGO54301 |
| | RMMUS180908 | | | | | not predicted |
| | RMMUS079906 | | RDR000755 | | | not predicted |
| | | | | | | not predicted |
| RPSIN020178 | | | RDR014556 | RCPIC02511 | RGOAS02476 | RALGO16184 |
| | RMMUS080099 | | | | | not predicted |
| RPSIN035442 | RMMUS005973 | | | RCPIC04515 | RGOAS01627 | RALGO49115 |
| RPSIN024552 | RMMUS2821 | RNFUR00665 | RDR053294 | RCPIC039931 | | RALGO48610 |
| RPSIN008980 | RMMUS0106 | RNFUR05112 | RDR012537 | RCPIC00014 | RGOAS00624 | RALGO44024 |
| RPSIN014138 | RMMUS147387 | | RDR035830 | RCPIC01470 | RGOAS02656 | RALGO37544 |
| 39 | RMMUS2317 | RNFUR01046 | RDR012226 | RCPIC03623 | RGOAS02483 | RALGO70093 |
| RPSIN034655 | RMMUS1907 | RNFUR02553 | RDR012201 | RCPIC04052 | RGOAS00294 | RALGO30854 |
| RPSIN032789 | RMMUS1489 | RNFUR05125 | RDR046706 | RCPIC01729 | RGOAS01746 | RALGO44666 |
| RPSIN027846 | RMMUS2585 | RNFUR00212 | RDR025766 | RCPIC00640 | RGOAS01378 | RALGO61934 |
| 31 | RMMUS1709 | RNFUR05077 | RDR022655 | RCPIC00304 | RGOAS02699 | RALGO16585 |

| | | | | | | |
|---|---|---|---|---|---|---|
| | RMMUS063907 | | | | | not predicted |
| RPSIN023514 | RMMUS224294 | | | RCPIC04489 | RGOAS00766 | RALGO70438 |
| RPSIN013905 | RMMUS0239 | RNFUR02842 | RDR017927 | RCPIC02032 | RGOAS00563 | RALGO23467 |
| RPSIN033903 | RMMUS2098 | RNFUR01515 | RDR009706 | RCPIC02114 | RGOAS00441 | RALGO51910 |
| RPSIN033530 | RMMUS1665 | RNFUR01298 | RDR004398 | RCPIC04170 | RGOAS00083 | RALGO37412 |
| RPSIN030966 | RMMUS2525 | RNFUR04038 | RDR036130 | RCPIC051933 | | RALGO00180 |
| RPSIN032632 | RMMUS2746 | RNFUR02302 | RDR024020 | RCPIC02267 | RGOAS01256 | RALGO41476 |
| RPSIN011257 | RMMUS0502 | RNFUR04944 | RDR016950 | RCPIC01575 | RGOAS01560 | RALGO12890 |
| RPSIN009098 | RMMUS0182 | RNFUR05125 | RDR049535 | RCPIC03213 | RGOAS02083 | RALGO20336 |
| RPSIN001489 | RMMUS1458 | RNFUR00028 | RDR021060 | RCPIC01498 | RGOAS00708 | RALGO29000 |
| RPSIN023521 | RMMUS0971 | RNFUR02222 | RDR027187 | RCPIC04980 | RGOAS00495 | RALGO61389 |
| RPSIN000224 | RMMUS2696 | RNFUR02513 | RDR034872 | RCPIC02922 | RGOAS01774 | RALGO31350 |
| RPSIN033222 | RMMUS0296 | RNFUR01457 | RDR035034 | RCPIC00810 | RGOAS01932 | RALGO50231 |
| RPSIN000034 | RMMUS1711 | RNFUR03825 | RDR012298 | RCPIC00303 | RGOAS01426 | RALGO16190 |
| RPSIN036193 | RMMUS193264 | | RDR038184 | RCPIC00590 | RGOAS02716 | RALGO27995 |
| RPSIN009240 | RMMUS1488 | RNFUR04711 | RDR040242 | RCPIC00230 | RGOAS03160 | RALGO70936 |
| RPSIN019991 | RMMUS1634 | RNFUR00884 | RDR025866 | RCPIC02616 | RGOAS02886 | RALGO24346 |
| RPSIN031257 | RMMUS276325 | | | RCPIC03164 | RGOAS00477 | RALGO67252 |
| RPSIN024000 | RMMUS2091 | RNFUR00259 | RDR047992 | RCPIC04827 | RGOAS02376 | RALGO56376 |
| RPSIN000561 | RMMUS1889 | RNFUR04903 | RDR005178 | RCPIC04022 | RGOAS02681 | RALGO35486 |
| RPSIN017075 | RMMUS056759 | | | RCPIC01107 | RGOAS02332 | RALGO57683 |
| RPSIN014103 | RMMUS1474 | RNFUR00997 | RDR035916 | RCPIC01448 | RGOAS02655 | RALGO37526 |
| RPSIN002062 | RMMUS133819 | | | RCPIC04272 | RGOAS01095 | RALGO47923 |
| | | | | | | not predicted |
| | | | | | | not predicted |
| | | | | | | not predicted |
| | RMMUS130461 | | | | | not predicted |
| | RMMUS130501 | | | | | not predicted |
| | | | | | | not predicted |
| | RMMUS130593 | | | | RGOAS01187 | not predicted |
| | RMMUS130305 | | | | | not predicted |
| | | | | | | not predicted |
| | RMMUS130501 | | | | | not predicted |
| | | | | | | not predicted |
| | RMMUS130501 | | | | | not predicted |
| | | | | | | not predicted |
| | RMMUS130501 | | | | | not predicted |
| | | | | | | not predicted |
| | RMMUS130501 | | | | | not predicted |
| | | | | | | not predicted |
| | | | | | | not predicted |
| | | | | | | not predicted |
| | | | | | | not predicted |

| | | | | | | |
|---|---|---|---|---|---|---|
| | RMMUS130501 | | | | | not predicted |
| | RMMUS130501 | | | | | not predicted |
| | RMMUS130501 | | | | | not predicted |
| | | | | | | not predicted |
| | RMMUS130501 | | | | | not predicted |
| | RMMUS130501 | | | | | not predicted |
| | RMMUS130501 | | | | | not predicted |
| | RMMUS130501 | | | | | not predicted |
| | RMMUS130501 | | | | | not predicted |
| | RMMUS130501 | | | | | not predicted |
| | RMMUS130501 | | | | | not predicted |
| | RMMUS130501 | | | | | not predicted |
| | RMMUS130501 | | | | | not predicted |
| | RMMUS130501 | | | | | not predicted |
| | RMMUS130501 | | | | | not predicted |
| | RMMUS130501 | | | | | not predicted |
| | RMMUS130501 | | | | | not predicted |
| | RMMUS130501 | | | | | not predicted |
| | RMMUS130501 | | | | | not predicted |
| | RMMUS130501 | | | | | not predicted |
| | RMMUS130501 | | | | | not predicted |
| | RMMUS130501 | | | | | not predicted |
| | RMMUS130501 | | | | | not predicted |
| | RMMUS130501 | | | | | not predicted |
| | | | | | | not predicted |
| RPSIN015317 | RMMUS2077 | RNFUR01636 | RDR030858 | RCPIC03439 | RGOAS00376 | RALGO65356 |
| RPSIN023692 | RMMUS2318 | RNFUR04656 | RDR036486 | RCPIC03630 | RGOAS01751 | RALGO70027 |
| RPSIN036943 | RMMUS0773 | RNFUR01722 | RDR014559 | RCPIC02279 | RGOAS01252 | RALGO56478 |
| RPSIN038713 | RMMUS2141 | RNFUR00739 | RDR022316 | RCPIC02777 | RGOAS02576 | RALGO32020 |
| RPSIN034160 | RMMUS1812 | RNFUR05390 | RDR009731 | RCPIC02853 | RGOAS01685 | RALGO10420 |
| RPSIN017566 | RMMUS0451 | RNFUR00081 | RDR035113 | RCPIC01757 | RGOAS00032 | RALGO27488 |
| RPSIN027390 | RMMUS1872 | RNFUR01225 | RDR004919 | RCPIC04875 | RGOAS03050 | not predicted |
| RPSIN022145 | RMMUS0129 | RNFUR02527 | RDR002667 | RCPIC01209 | RGOAS03234 | RALGO42002 |
| RPSIN030891 | RMMUS201088 | | RDR056555 | RCPIC03024 | RGOAS02104 | RALGO58700 |
| RPSIN001599 | RMMUS2806 | RNFUR01241 | RDR039749 | RCPIC03065 | RGOAS01385 | RALGO43350 |
| RPSIN026016 | RMMUS2416 | RNFUR02634 | RDR003290 | RCPIC04507 | RGOAS02532 | RALGO00213 |
| RPSIN013026 | RMMUS2196 | RNFUR02989 | RDR024704 | RCPIC04133 | RGOAS01534 | RALGO62527 |
| RPSIN015594 | RMMUS2483 | RNFUR00663 | RDR026062 | RCPIC00619 | RGOAS00867 | RALGO59447 |
| RPSIN011468 | RMMUS2331 | RNFUR04124 | RDR024027 | RCPIC046485 | | RALGO54163 |
| RPSIN006055 | RMMUS2380 | RNFUR00509 | RDR005655 | RCPIC01984 | RGOAS02154 | RALGO32560 |
| RPSIN008057 | RMMUS1446 | RNFUR04339 | RDR006289 | RCPIC04850 | RGOAS02152 | RALGO10864 |
| 31 | RMMUS1924 | RNFUR03692 | RDR033045 | RCPIC03671 | RGOAS02409 | RALGO02752 |
| 90 | RMMUS1263 | RNFUR04827 | RDR024133 | RCPIC026275 | | RALGO36344 |
| RPSIN007398 | RMMUS1754 | RNFUR003434 | | RCPIC01319 | RGOAS00933 | RALGO49778 |
| RPSIN036546 | RMMUS2710 | RNFUR04482 | RDR017501 | RCPIC01115 | RGOAS01064 | RALGO53366 |

| | | | | | | |
|---|---|---|---|---|---|---|
| RPSIN024942 | RMMUS1662 | RNFUR02655 | RDR042566 | RCPIC04186 | RGOAS00391 | RALGO36521 |
| RPSIN001416 | RMMUS1483 | RNFUR03107 | RDR029315 | RCPIC01441 | RGOAS00454 | RALGO35537 |
| RPSIN019422 | RMMUS1273 | RNFUR04340 | RDR014299 | RCPIC03819 | RGOAS01465 | RALGO35216 |
| RPSIN007167 | RMMUS0169 | RNFUR00011 | RDR038521 | RCPIC00956 | RGOAS01104 | RALGO64051 |
| RPSIN020561 | RMMUS1877 | RNFUR02708 | RDR024450 | RCPIC04770 | RGOAS03216 | RALGO00204 |
| RPSIN032777 | RMMUS0331 | RNFUR02654 | RDR035616 | RCPIC04267 | RGOAS01928 | RALGO45140 |
| RPSIN028350 | RMMUS0369 | RNFUR01271 | RDR022782 | RCPIC05535 | RGOAS03191 | RALGO27896 |
| | RMMUS243676 | | | | | not predicted |
| 11 | RMMUS1758 | RNFUR04707 | RDR028483 | RCPIC049842 | | RALGO56064 |
| RPSIN034941 | RMMUS1265 | RNFUR01603 | RDR024970 | RCPIC02623 | RGOAS00130 | RALGO35006 |
| RPSIN033552 | RMMUS2563 | RNFUR03567 | RDR002538 | RCPIC04150 | RGOAS01863 | RALGO37532 |
| RPSIN010436 | RMMUS1268 | RNFUR00581 | RDR025639 | RCPIC02477 | RGOAS01134 | RALGO00342 |
| RPSIN004677 | RMMUS281794 | | RDR009813 | RCPIC01834 | RGOAS01696 | RALGO49679 |
| RPSIN037727 | RMMUS2073 | RNFUR04909 | RDR036442 | RCPIC03441 | RGOAS00119 | RALGO65080 |
| RPSIN026064 | RMMUS1527 | RNFUR03945 | RDR026137 | RCPIC02663 | RGOAS01314 | RALGO43494 |
| RPSIN029327 | RMMUS041129 | | RDR054804 | RCPIC02970 | RGOAS01047 | RALGO08329 |
| RPSIN033541 | RMMUS1665 | RNFUR03574 | RDR002867 | RCPIC04170 | RGOAS00080 | RALGO37427 |
| RPSIN015728 | RMMUS1747 | RNFUR00710 | RDR026586 | RCPIC04441 | RGOAS03146 | RALGO65632 |
| RPSIN011859 | RMMUS0524 | RNFUR00982 | RDR027899 | RCPIC03343 | RGOAS00498 | RALGO29945 |
| RPSIN018457 | RMMUS1738 | RNFUR04210 | RDR002821 | RCPIC002699 | | RALGO54628 |
| 58 | RMMUS1105 | RNFUR02330 | RDR030648 | | | not predicted |
| RPSIN026474 | RMMUS2625 | RNFUR02740 | RDR027775 | RCPIC04825 | RGOAS03262 | RALGO56538 |
| RPSIN003709 | RMMUS0889 | RNFUR03011 | RDR052236 | RCPIC01792 | RGOAS00317 | RALGO17368 |
| RPSIN027840 | RMMUS2584 | RNFUR04046 | RDR025843 | RCPIC00595 | RGOAS00241 | RALGO61862 |
| RPSIN001974 | RMMUS0962 | RNFUR03602 | RDR027193 | RCPIC02295 | RGOAS02985 | RALGO59504 |
| RPSIN024443 | RMMUS031770 | | RDR048188 | RCPIC04273 | RGOAS01211 | RALGO53396 |
| RPSIN004301 | RMMUS1981 | RNFUR00134 | RDR038282 | RCPIC00825 | RGOAS01230 | RALGO50839 |
| RPSIN028345 | RMMUS0368 | RNFUR03011 | RDR012682 | RCPIC02962 | RGOAS02065 | RALGO28073 |
| RPSIN003148 | RMMUS2813 | RNFUR04805 | RDR045360 | RCPIC03138 | RGOAS00247 | RALGO67648 |
| RPSIN003793 | RMMUS2292 | RNFUR03518 | RDR036953 | RCPIC04762 | RGOAS01939 | RALGO69085 |
| RPSIN037333 | RMMUS2409 | RNFUR02282 | RDR002964 | RCPIC01836 | RGOAS02497 | RALGO12083 |
| RPSIN026360 | RMMUS0532 | RNFUR04995 | RDR053124 | RCPIC03332 | RGOAS02696 | RALGO33547 |
| RPSIN012876 | RMMUS0993 | RNFUR01601 | RDR043277 | RCPIC00969 | RGOAS02933 | RALGO30590 |
| 55 | RMMUS1315 | RNFUR049543 | | | RGOAS03169 | RALGO63778 |
| RPSIN016704 | RMMUS042717 | | | RCPIC01783 | RGOAS03233 | RALGO18886 |
| RPSIN018767 | RMMUS224940 | | RDR027774 | RCPIC04461 | RGOAS00413 | RALGO70294 |
| | RMMUS2781 | RNFUR020337 | | RCPIC01883 | RGOAS01131 | RALGO44648 |
| RPSIN038301 | RMMUS0110 | RNFUR03510 | RDR022703 | RCPIC00822 | RGOAS01453 | RALGO18465 |
| RPSIN029156 | RMMUS1348 | RNFUR04090 | RDR041983 | RCPIC03776 | RGOAS00428 | RALGO40722 |
| RPSIN011116 | RMMUS2114 | RNFUR01803 | RDR039843 | RCPIC02406 | RGOAS01082 | RALGO44342 |
| RPSIN018312 | RMMUS2050 | RNFUR03760 | RDR036576 | RCPIC034046 | | RALGO53423 |
| RPSIN001220 | RMMUS0388 | RNFUR026986 | | RCPIC04410 | RGOAS02537 | RALGO00273 |
| RPSIN008512 | RMMUS086727 | | RDR029080 | RCPIC051644 | | RALGO01762 |
| RPSIN037830 | RMMUS207237 | | RDR045507 | | RGOAS03073 | RALGO65032 |
| RPSIN034488 | RMMUS0541 | RNFUR05151 | RDR022821 | RCPIC03121 | RGOAS00182 | RALGO27740 |

| | | | | | | |
|---|---|---|---|---|---|---|
| RPSIN003794 | RMMUS2292 | RNFUR00621 | RDR031335 | RCPIC04755 | RGOAS02610 | RALGO69100 |
| RPSIN019745 | RMMUS1854 | RNFUR03081 | RDR024598 | RCPIC04104 | RGOAS01801 | RALGO32644 |
| RPSIN026243 | RMMUS250033 | | | RCPIC014171 | | |
| RPSIN031293 | RMMUS2589 | RNFUR00021 | RDR026922 | RCPIC02242 | RGOAS01998 | RALGO64693 |
| RPSIN038205 | RMMUS2385 | RNFUR01957 | RDR006236 | RCPIC03690 | RGOAS03094 | RALGO25014 |
| RPSIN032191 | RMMUS2357 | RNFUR04457 | RDR057075 | RCPIC05719 | RGOAS00848 | RALGO20922 |
| RPSIN011988 | RMMUS1914 | RNFUR039230 | | RCPIC04026 | RGOAS02880 | RALGO38696 |
| RPSIN022394 | RMMUS0710 | RNFUR04758 | RDR043957 | RCPIC01690 | RGOAS01456 | RALGO12776 |
| RPSIN018800 | RMMUS224784 | | RDR019723 | RCPIC05074 | RGOAS00170 | RALGO70336 |
| RPSIN022649 | RMMUS160703 | | RDR038570 | RCPIC00650 | RGOAS02742 | RALGO55063 |
| RPSIN027124 | RMMUS2296 | RNFUR04231 | RDR031775 | RCPIC03565 | RGOAS00317 | RALGO69289 |
| RPSIN031719 | RMMUS1578 | RNFUR03321 | RDR036455 | RCPIC00295 | RGOAS03230 | RALGO49010 |
| RPSIN018214 | RMMUS2334 | RNFUR00740 | RDR024892 | RCPIC04661 | RGOAS01384 | RALGO53699 |
| RPSIN003849 | RMMUS059141 | | RDR031877 | RCPIC00394 | RGOAS016078 | |
| [illegible] | RMMUS093240 | | | | | not predicted |
| RPSIN007160 | RMMUS0170 | RNFUR03611 | RDR033005 | RCPIC00983 | RGOAS02298 | RALGO64309 |
| RPSIN021506 | RMMUS0635 | RNFUR05083 | RDR017526 | RCPIC05012 | RGOAS02645 | RALGO13136 |
| RPSIN026511 | RMMUS1041 | RNFUR02802 | RDR007828 | RCPIC00084 | RGOAS01451 | RALGO57818 |
| | RMMUS061629 | | | | | not predicted |
| | RMMUS1865 | RNFUR04798 | RDR024356 | RCPIC04891 | RGOAS02318 | RALGO04938 |
| RPSIN011734 | RMMUS213452 | | RDR021293 | RCPIC04239 | RGOAS01357 | RALGO36218 |
| RPSIN027738 | RMMUS1717 | RNFUR01933 | RDR012834 | RCPIC04793 | RGOAS02650 | RALGO14582 |
| 46 | RMMUS0055 | RNFUR00706 | RDR001785 | RCPIC03252 | RGOAS00650 | RALGO16621 |
| RPSIN033311 | RMMUS1012 | RNFUR01273 | RDR008760 | RCPIC00018 | RGOAS02214 | RALGO57212 |
| RPSIN021513 | RMMUS1089 | RNFUR05440 | RDR018948 | RCPIC04872 | RGOAS02102 | RALGO09702 |
| 63 | RMMUS1089 | RNFUR01874 | RDR029363 | RCPIC05493 | RGOAS01134 | RALGO09045 |
| RPSIN019274 | RMMUS2620 | RNFUR02150 | RDR012657 | RCPIC04524 | RGOAS02371 | RALGO48713 |
| RPSIN015521 | RMMUS2388 | RNFUR04375 | RDR024342 | RCPIC03700 | RGOAS00815 | RALGO23100 |
| RPSIN027494 | RMMUS0032 | RNFUR01940 | RDR000814 | | RGOAS02735 | RALGO69262 |
| RPSIN010467 | RMMUS2182 | RNFUR00614 | RDR026882 | RCPIC02469 | RGOAS02872 | RALGO11840 |
| RPSIN031921 | RMMUS2137 | RNFUR04312 | RDR016793 | RCPIC04231 | RGOAS00164 | RALGO34213 |
| RPSIN020147 | RMMUS0799 | RNFUR039551 | | RCPIC02503 | RGOAS01101 | RALGO15752 |
| RPSIN002186 | RMMUS246526 | | | RCPIC03167 | RGOAS02808 | RALGO66721 |
| RPSIN036903 | RMMUS2506 | RNFUR00313 | RDR027560 | RCPIC01228 | RGOAS02407 | RALGO51967 |
| RPSIN019855 | RMMUS1889 | RNFUR04598 | RDR000769 | RCPIC04022 | RGOAS007276 | |
| | RMMUS124145 | | | | | |
| RPSIN006365 | RMMUS159131 | | RDR054245 | RCPIC00539 | RGOAS03322 | RALGO57413 |
| RPSIN035869 | RMMUS0172 | RNFUR01728 | RDR033422 | RCPIC01073 | RGOAS02858 | RALGO64174 |
| RPSIN014549 | RMMUS1046 | RNFUR00914 | RDR009346 | RCPIC00044 | RGOAS00397 | RALGO58565 |
| RPSIN001338 | RMMUS1456 | RNFUR03118 | RDR034467 | RCPIC01461 | RGOAS02768 | RALGO29819 |
| RPSIN035392 | RMMUS1953 | RNFUR02504 | RDR029649 | RCPIC00481 | RGOAS03178 | RALGO43070 |
| RPSIN020609 | RMMUS1876 | RNFUR03964 | RDR024303 | RCPIC04762 | RGOAS03215 | RALGO00447 |
| RPSIN005379 | RMMUS2407 | RNFUR04361 | RDR056100 | RCPIC01801 | RGOAS01135 | RALGO40422 |
| RPSIN007036 | RMMUS1036 | RNFUR040488 | | RCPIC00110 | RGOAS00693 | RALGO59990 |
| RPSIN000304 | RMMUS1729 | RNFUR04488 | RDR021875 | RCPIC02456 | RGOAS01863 | RALGO29621 |

| | | | | | | |
|---|---|---|---|---|---|---|
| RPSIN00948 | RMMUS0159 | RNFUR0043 | RDR056559 | RCPIC00613 | RGOAS0093 | RALGO6077 |
| RPSIN02611 | RMMUS0581 | RNFUR0041 | RDR048494 | RCPIC02138 | RGOAS0160 | RALGO1190 |
| RPSIN02713 | RMMUS2295 | RNFUR0406 | RDR036030 | RCPIC04756 | RGOAS0279 | RALGO6923 |
| RPSIN01888 | RMMUS0302 | RNFUR0404 | RDR014285 | RCPIC00850 | RGOAS0192 | RALGO4946 |
| RPSIN01255 | RMMUS0936 | RNFUR0198 | RDR045139 | RCPIC00310 | RGOAS0194 | RALGO5532 |
| RPSIN00638 | RMMUS2472 | RNFUR0381 | RDR037904 | RCPIC02715 | RGOAS0170 | RALGO5815 |
| RPSIN03798 | RMMUS1297 | RNFUR0345 | RDR000658 | RCPIC03656 | RGOAS0096 | RALGO6905 |
| RPSIN02417 | RMMUS132697 | | | | | |
| RPSIN01168 | RMMUS2126 | RNFUR0310 | RDR009810 | RCPIC02404 | RGOAS0252 | RALGO3850 |
| RPSIN01171 | RMMUS2126 | RNFUR0233 | RDR039817 | RCPIC02405 | RGOAS0024 | RALGO3846 |
| RPSIN03798 | RMMUS1297 | RNFUR0345 | RDR000246 | RCPIC03656 | RGOAS009685 | |
| RPSIN02751 | RMMUS1284 | RNFUR0316 | RDR015064 | RCPIC00792 | RGOAS0266 | RALGO4726 |
| RPSIN00212 | RMMUS269453 | | | RCPIC04222 | RGOAS0005 | RALGO3204 |
| RPSIN02272 | RMMUS0933 | RNFUR0323 | RDR031823 | RCPIC02216 | RGOAS0253 | RALGO6301 |
| RPSIN00120 | RMMUS0953 | RNFUR0094 | RDR000974 | RCPIC02512 | RGOAS0252 | RALGO2514 |
| RPSIN01094 | RMMUS2345 | RNFUR0040 | RDR007793 | RCPIC00262 | RGOAS0070 | RALGO3212 |
| RPSIN02991 | RMMUS0498 | RNFUR0172 | RDR019232 | RCPIC00035 | RGOAS0209 | RALGO6354 |
| RPSIN03521 | RMMUS1169 | RNFUR0479 | RDR034575 | RCPIC018470 | | RALGO4168 |
| RPSIN03857 | RMMUS142035 | | RDR047704 | RCPIC05077 | RGOAS0036 | RALGO3460 |
| RPSIN01664 | RMMUS2864 | RNFUR0192 | RDR020005 | RCPIC03858 | RGOAS0027 | RALGO6797 |
| RPSIN03390 | RMMUS2101 | RNFUR0471 | RDR017769 | RCPIC02124 | RGOAS0112 | RALGO5181 |
| RPSIN03205 | RMMUS100272 | | RDR012650 | RCPIC01059 | RGOAS0214 | RALGO1849 |
| RPSIN00727 | RMMUS2511 | RNFUR0512 | RDR016912 | RCPIC03674 | RGOAS0133 | RALGO2337 |
| 03 | RMMUS1160 | RNFUR0517 | RDR046500 | RCPIC02277 | RGOAS0114 | RALGO4618 |
| RPSIN02971 | RMMUS0575 | RNFUR0490 | RDR055772 | RCPIC03928 | RGOAS0103 | RALGO0388 |
| RPSIN03838 | RMMUS1179 | RNFUR0523 | RDR047460 | RCPIC02598 | RGOAS0325 | RALGO4688 |
| RPSIN01577 | RMMUS1746 | RNFUR0287 | RDR026213 | RCPIC02703 | RGOAS0063 | RALGO6565 |
| RPSIN01381 | RMMUS1968 | RNFUR0179 | RDR032932 | RCPIC02914 | RGOAS0180 | RALGO6092 |
| RPSIN01483 | RMMUS2717 | RNFUR0484 | RDR034202 | RCPIC02612 | RGOAS0272 | RALGO2981 |
| RPSIN02467 | RMMUS0140 | RNFUR0416 | RDR037928 | RCPIC00061 | RGOAS0296 | RALGO4538 |
| RPSIN03482 | RMMUS2683 | RNFUR0234 | RDR038351 | RCPIC00961 | RGOAS0342 | RALGO1374 |
| RPSIN02843 | RMMUS2594 | RNFUR0117 | RDR025589 | RCPIC05449 | RGOAS0154 | RALGO6555 |
| RPSIN03590 | RMMUS1415 | RNFUR0522 | RDR055366 | RCPIC01512 | RGOAS0162 | RALGO3290 |
| RPSIN03643 | RMMUS1526 | RNFUR0405 | RDR026122 | RCPIC00610 | RGOAS0295 | RALGO3806 |
| RPSIN00185 | RMMUS0463 | RNFUR0407 | RDR001136 | RCPIC01084 | RGOAS0090 | RALGO4444 |
| | RMMUS015554 | | | | | not predicte |
| RPSIN00668 | RMMUS0151 | RNFUR0488 | RDR014513 | RCPIC04587 | RGOAS0059 | RALGO4498 |
| 20 | RMMUS128992 | | | | | not predicte |
| | RMMUS0493 | RNFUR009508 | | | | not predicte |
| | | | | | | not predicte |
| | RMMUS014996 | | | | | not predicte |
| | RMMUS014968 | | | | | |
| | RMMUS014996 | | | | | not predicte |
| | | | | | | not predicte |
| | RMMUS021750 | | | | RGOAS0240 | RALGO0871 |

| | | | | | | |
|---|---|---|---|---|---|---|
| RPSIN007467 | | | | RCPIC032807 | | RALGO08655 |
| | | | | | | not predicte |
| | | | | | | not predicte |
| RPSIN00018 | RMMUS2694 | RNFUR0095 | RDR045266 | RCPIC02921 | RGOAS0136 | RALGO31804 |
| RPSIN02479 | RMMUS2571 | RNFUR0412 | RDR012047 | RCPIC03906 | RGOAS0037 | RALGO46623 |
| RPSIN03009 | RMMUS0062 | RNFUR0490 | RDR019123 | RCPIC05532 | RGOAS0050 | RALGO49457 |
| RPSIN00184 | RMMUS0461 | RNFUR0073 | RDR000410 | RCPIC04261 | RGOAS0309 | not predicte |
| RPSIN00184 | RMMUS0461 | RNFUR0073 | RDR000683 | RCPIC04292 | RGOAS0309 | RALGO4499 |
| RPSIN00174 | RMMUS1285 | RNFUR0381 | RDR014687 | RCPIC00792 | RGOAS0140 | RALGO4716 |
| 42 | RMMUS115086 | | | RCPIC05558 | RGOAS0017 | not predicte |
| RPSIN03138 | RMMUS1311 | RNFUR0353 | RDR017306 | RCPIC03883 | RGOAS0247 | RALGO6385 |
| RPSIN02330 | RMMUS2473 | RNFUR0074 | RDR039085 | RCPIC00616 | RGOAS0022 | RALGO5904 |
| RPSIN00170 | RMMUS1256 | RNFUR0369 | RDR026556 | RCPIC02318 | RGOAS0130 | RALGO3863 |
| RPSIN03229 | RMMUS0757 | RNFUR0432 | RDR027089 | RCPIC02313 | RGOAS0194 | RALGO5880 |
| 78 | RMMUS208937 | | | RCPIC04829 | RGOAS0238 | RALGO56331 |
| RPSIN03145 | RMMUS0158 | RNFUR0535 | RDR024388 | RCPIC00285 | RGOAS0004 | RALGO57455 |
| RPSIN02191 | RMMUS215275 | | RDR012626 | RCPIC00945 | RGOAS0174 | RALGO35135 |
| RPSIN00442 | RMMUS0786 | RNFUR0378 | RDR001398 | RCPIC04254 | RGOAS0117 | RALGO53015 |
| RPSIN02167 | RMMUS1784 | RNFUR0316 | RDR030847 | RCPIC01769 | RGOAS0134 | RALGO09723 |
| RPSIN00134 | RMMUS1456 | RNFUR0097 | RDR034801 | RCPIC01456 | RGOAS0154 | RALGO2987 |
| 70 | RMMUS2538 | RNFUR0045 | RDR036752 | RCPIC03609 | RGOAS0081 | RALGO0357 |
| RPSIN03796 | RMMUS089561 | | RDR046213 | RCPIC04467 | RGOAS0085 | RALGO7075 |
| RPSIN03701 | RMMUS1764 | RNFUR0028 | RDR027765 | RCPIC01263 | RGOAS0318 | RALGO55157 |
| RPSIN00985 | RMMUS1992 | RNFUR0020 | RDR038379 | RCPIC03016 | RGOAS0004 | RALGO56763 |
| RPSIN01729 | RMMUS1089 | RNFUR0204 | RDR035190 | RCPIC05624 | RGOAS0099 | RALGO08598 |
| RPSIN02583 | RMMUS2680 | RNFUR0033 | RDR027691 | RCPIC00949 | RGOAS0078 | RALGO15893 |
| RPSIN00721 | RMMUS0754 | RNFUR0374 | RDR032891 | RCPIC00569 | RGOAS0090 | RALGO63313 |
| RPSIN00980 | RMMUS1965 | RNFUR0129 | RDR033061 | RCPIC03012 | RGOAS0253 | RALGO56394 |
| 19 | RMMUS0356 | RNFUR0484 | RDR033053 | RCPIC007004 | | RALGO66730 |
| RPSIN03464 | RMMUS1906 | RNFUR0471 | RDR004722 | RCPIC04039 | RGOAS0111 | RALGO31083 |
| RPSIN01961 | RMMUS1544 | RNFUR0471 | RDR029034 | RCPIC01735 | RGOAS0087 | RALGO3620 |
| RPSIN01951 | RMMUS0442 | RNFUR0218 | RDR029058 | RCPIC04465 | RGOAS0312 | RALGO7055 |
| RPSIN01718 | RMMUS1619 | RNFUR0022 | RDR032758 | RCPIC02789 | RGOAS0136 | RALGO19207 |
| 06 | RMMUS2820 | RNFUR0027 | RDR028308 | RCPIC01292 | RGOAS0246 | RALGO55421 |
| | RMMUS065285 | | | | | |
| RPSIN01287 | RMMUS0993 | RNFUR0159 | RDR013739 | RCPIC01051 | RGOAS0339 | RALGO3083 |
| | RMMUS275512 | | | | | not predicte |
| RPSIN00384 | RMMUS0591 | RNFUR0254 | RDR031442 | RCPIC00284 | RGOAS0315 | RALGO5717 |
| RPSIN01570 | RMMUS2595 | RNFUR0378 | RDR023928 | RCPIC04393 | RGOAS0099 | RALGO65491 |
| | RMMUS1655 | RNFUR0508 | RDR041648 | RCPIC03781 | RGOAS0062 | RALGO38807 |
| RPSIN02569 | RMMUS0542 | RNFUR0233 | RDR003544 | RCPIC03132 | RGOAS0246 | RALGO27464 |
| RPSIN00072 | RMMUS1254 | RNFUR0427 | RDR049388 | RCPIC05406 | RGOAS0216 | RALGO0644 |

| | | | | | | |
|---|---|---|---|---|---|---|
| RPSIN038754 | RMMUS2270 | RNFUR03594 | RDR040249 | RCPIC04776 | RGOAS03021 | not predicted |
| RPSIN017210 | RMMUS1615 | RNFUR04358 | RDR033044 | RCPIC02824 | RGOAS01547 | RALGO51517 |
| RPSIN024987 | RMMUS005067 | | RDR019153 | RCPIC03256 | RGOAS01059 | RALGO18715 |
| RPSIN011216 | RMMUS017831 | | | RCPIC01599 | RGOAS00341 | RALGO15572 |
| RPSIN018560 | RMMUS2444 | RNFUR05048 | RDR020834 | RCPIC00091 | RGOAS01320 | RALGO65224 |
| RPSIN019653 | RMMUS0956 | RNFUR02522 | RDR012091 | RCPIC01239 | RGOAS01403 | RALGO22587 |
| | RMMUS1664 | RNFUR05117 | RDR022809 | RCPIC03772 | RGOAS01712 | RALGO39999 |
| RPSIN028622 | RMMUS2084 | RNFUR02169 | RDR017409 | RCPIC02119 | RGOAS03156 | RALGO51007 |
| RPSIN018461 | RMMUS1025 | RNFUR00505 | RDR030989 | RCPIC02220 | RGOAS00561 | RALGO63220 |
| | RMMUS213887 | | | | | |
| RPSIN021394 | RMMUS1299 | RNFUR04369 | RDR000601 | RCPIC03658 | RGOAS00229 | RALGO69010 |
| RPSIN016891 | RMMUS1798 | RNFUR04743 | RDR004924 | RCPIC04045 | RGOAS03058 | RALGO25437 |
| 99 | RMMUS275762 | | | RCPIC051818 | | RALGO24262 |
| RPSIN027331 | RMMUS0981 | RNFUR04508 | RDR022710 | RCPIC01038 | RGOAS01514 | RALGO21786 |
| RPSIN026972 | RMMUS0187 | RNFUR00984 | RDR007382 | RCPIC03220 | RGOAS01896 | not predicted |
| RPSIN007489 | RMMUS2229 | RNFUR00824 | RDR038267 | RCPIC00790 | RGOAS00329 | not predicted |
| RPSIN035343 | RMMUS2447 | RNFUR02260 | RDR029851 | RCPIC05081 | RGOAS02238 | RALGO65578 |
| RPSIN008011 | RMMUS2736 | RNFUR00389 | RDR026608 | RCPIC00979 | RGOAS01517 | RALGO14750 |
| RPSIN022447 | RMMUS0742 | RNFUR02703 | RDR045299 | RCPIC02681 | RGOAS00415 | RALGO00669 |
| RPSIN013018 | RMMUS2196 | RNFUR00959 | RDR032961 | RCPIC04126 | RGOAS01292 | RALGO62584 |
| RPSIN036473 | RMMUS1525 | RNFUR04175 | RDR026112 | RCPIC00610 | RGOAS00531 | RALGO38153 |
| RPSIN005238 | RMMUS0518 | RNFUR05307 | RDR017343 | RCPIC03946 | RGOAS02062 | RALGO31726 |
| RPSIN035209 | RMMUS151643 | | RDR046882 | RCPIC01960 | RGOAS03288 | RALGO41341 |
| RPSIN037601 | RMMUS1844 | RNFUR01833 | RDR028925 | RCPIC02844 | RGOAS03100 | RALGO22350 |
| RPSIN001118 | RMMUS055499 | | | RCPIC050250 | | RALGO68554 |
| RPSIN036181 | RMMUS1931 | RNFUR00245 | RDR007838 | RCPIC00589 | RGOAS02958 | RALGO26566 |
| RPSIN005258 | RMMUS0819 | RNFUR01720 | RDR050399 | RCPIC039525 | | RALGO31167 |
| RPSIN014703 | RMMUS1888 | RNFUR03441 | RDR004621 | RCPIC04969 | RGOAS00030 | RALGO17646 |
| RPSIN015444 | RMMUS0376 | RNFUR04141 | RDR022517 | RCPIC02958 | RGOAS01791 | not predicted |
| RPSIN013641 | RMMUS0949 | RNFUR05420 | RDR043411 | RCPIC003603 | | RALGO15275 |
| RPSIN029780 | RMMUS100329 | | | RCPIC00976 | RGOAS02967 | not predicted |
| RPSIN032189 | RMMUS0425 | RNFUR04955 | RDR002627 | RCPIC01784 | RGOAS00547 | RALGO17889 |
| RPSIN017307 | RMMUS0314 | RNFUR05025 | RDR038299 | RCPIC02759 | RGOAS02072 | RALGO54688 |
| RPSIN030681 | RMMUS0630 | RNFUR03245 | RDR018242 | RCPIC01275 | RGOAS03022 | RALGO15008 |
| RPSIN003022 | RMMUS2185 | RNFUR02911 | RDR026198 | RCPIC02139 | RGOAS02351 | RALGO15416 |
| RPSIN017585 | RMMUS0452 | RNFUR02978 | RDR034871 | RCPIC01826 | RGOAS02498 | RALGO26966 |
| | RMMUS1667 | RNFUR04278 | RDR002316 | | RGOAS02508 | RALGO38507 |
| RPSIN016233 | RMMUS2704 | RNFUR05300 | RDR009926 | RCPIC011276 | | RALGO54694 |
| 21 | RMMUS0401 | RNFUR03036 | RDR013153 | RCPIC02998 | RGOAS03321 | RALGO13733 |
| | | | | | | not predicted |
| RPSIN030907 | RMMUS2007 | RNFUR01414 | RDR026231 | RCPIC03012 | RGOAS00421 | RALGO58466 |

| | | | | | | |
|---|---|---|---|---|---|---|
| RPSIN017057 | RMMUS1624 | RNFUR03023 | RDR038806 | RCPIC00646 | RGOAS02841 | RALGO52435 |
| RPSIN033160 | RMMUS278519 | | RDR027740 | RCPIC01287 | RGOAS00545 | RALGO54715 |
| RPSIN004654 | RMMUS2261 | RNFUR02141 | RDR001680 | RCPIC046853 | | not predicted |
| RPSIN004303 | RMMUS1980 | RNFUR00177 | RDR032976 | RCPIC00899 | RGOAS01233 | RALGO50590 |
| RPSIN031308 | RMMUS206018 | | | RCPIC03431 | RGOAS01241 | RALGO65530 |
| RPSIN006920 | RMMUS2266 | RNFUR03492 | RDR018942 | RCPIC04683 | RGOAS02296 | RALGO69406 |
| RPSIN002533 | RMMUS200175 | | | RCPIC03039 | RGOAS03252 | RALGO57914 |
| RPSIN017127 | RMMUS1235 | RNFUR00013 | RDR004962 | RCPIC01795 | RGOAS01768 | RALGO18757 |
| RPSIN009726 | RMMUS1559 | RNFUR04215 | RDR022387 | RCPIC05087 | RGOAS02798 | RALGO18859 |
| RPSIN001033 | RMMUS1978 | RNFUR04083 | RDR033051 | RCPIC01612 | RGOAS01404 | RALGO10426 |
| RPSIN019570 | RMMUS1657 | RNFUR01917 | RDR002383 | RCPIC037806 | | RALGO39342 |
| | RMMUS205093 | | | | | |
| | RMMUS205104 | | | | | |
| RPSIN021417 | RMMUS0632 | RNFUR01095 | RDR043326 | RCPIC01196 | RGOAS02638 | RALGO13928 |
| RPSIN038711 | RMMUS2141 | RNFUR03957 | RDR017364 | RCPIC02780 | RGOAS02432 | RALGO31927 |
| RPSIN008457 | RMMUS1467 | RNFUR00827 | RDR022195 | RCPIC01450 | RGOAS03284 | RALGO32662 |
| RPSIN037039 | RMMUS1777 | RNFUR04659 | RDR028216 | RCPIC01236 | RGOAS03332 | RALGO55649 |
| RPSIN038207 | RMMUS2385 | RNFUR03793 | RDR006554 | RCPIC036909 | | RALGO25368 |
| RPSIN023356 | RMMUS1039 | RNFUR02498 | RDR000306 | RCPIC03294 | RGOAS01503 | RALGO52850 |
| 50 | RMMUS1879 | RNFUR05285 | RDR005493 | RCPIC04973 | RGOAS02448 | RALGO17374 |
| RPSIN018990 | RMMUS1244 | RNFUR04408 | RDR004676 | RCPIC036946 | | RALGO34258 |
| | RMMUS110194 | | | RCPIC052091 | | RALGO48373 |
| RPSIN020131 | RMMUS1318 | RNFUR00334 | RDR028568 | RCPIC02518 | RGOAS01339 | RALGO06688 |
| RPSIN036444 | RMMUS152670 | | RDR026845 | RCPIC05271 | RGOAS023448 | |
| 05 | | | | | | RALGO38015 |
| | RMMUS001186 | | | | RGOAS02344 | RALGO38024 |
| 00 | | RNFUR036724 | | | | |
| RPSIN001059 | RMMUS267485 | | RDR027792 | RCPIC03578 | RGOAS02348 | RALGO09831 |
| RPSIN015077 | RMMUS1645 | RNFUR04556 | RDR017228 | RCPIC04189 | RGOAS01178 | RALGO22527 |
| RPSIN032407 | RMMUS1995 | RNFUR04723 | RDR032815 | RCPIC03036 | RGOAS01466 | RALGO56988 |
| RPSIN030664 | RMMUS1155 | RNFUR05305 | RDR046407 | RCPIC03087 | RGOAS03335 | RALGO38810 |
| RPSIN018633 | RMMUS2131 | RNFUR03416 | RDR010243 | RCPIC02401 | RGOAS01390 | RALGO40440 |
| RPSIN012422 | RMMUS0856 | RNFUR05205 | RDR009881 | RCPIC01593 | RGOAS01815 | RALGO11296 |
| RPSIN000157 | RMMUS2537 | RNFUR03137 | RDR036492 | RCPIC05671 | RGOAS00519 | RALGO06655 |
| RPSIN006382 | RMMUS2470 | RNFUR00264 | RDR038153 | RCPIC00616 | RGOAS01182 | RALGO58076 |
| RPSIN005642 | RMMUS0729 | RNFUR02265 | RDR030724 | RCPIC00258 | RGOAS02744 | RALGO38678 |
| RPSIN038728 | RMMUS2141 | RNFUR011826 | | RCPIC02777 | RGOAS02437 | RALGO31882 |
| | RMMUS222104 | | | | RGOAS03217 | RALGO34505 |
| RPSIN028343 | | | RDR019039 | | | RALGO34568 |
| | RMMUS214173 | | RDR041299 | | | not predicted |
| | RMMUS165536 | | | | RGOAS00560 | RALGO39843 |
| | RMMUS147269 | | RDR010225 | RCPIC014495 | | RALGO37715 |
| RPSIN010310 | RMMUS0164 | RNFUR03439 | RDR005199 | RCPIC00572 | RGOAS00854 | RALGO02836 |
| RPSIN010310 | RMMUS0164 | RNFUR03439 | RDR005199 | RCPIC00572 | RGOAS00854 | RALGO02836 |

| | | | | | | |
|---|---|---|---|---|---|---|
| RPSIN02239 | RMMUS0710 | RNFUR0384 | RDR014045 | RCPIC01701 | RGOAS0145 | RALGO1295 |
| | RMMUS112211 | | | RCPIC05616 | RGOAS0010 | not predicte |
| RPSIN03607 | RMMUS2297 | RNFUR0083 | RDR031198 | RCPIC03553 | RGOAS0044 | RALGO69331 |
| RPSIN03713 | RMMUS1760 | RNFUR027436 | | | RGOAS0007 | RALGO5589 |
| RPSIN00028 | RMMUS2627 | RNFUR0359 | RDR027653 | RCPIC03981 | RGOAS0035 | RALGO4837 |
| 46 | RMMUS0364 | RNFUR0294 | RDR044114 | RCPIC04413 | RGOAS0268 | RALGO03549 |
| RPSIN00963 | RMMUS066696 | | | RCPIC04934 | RGOAS0312 | RALGO09111 |
| RPSIN01633 | RMMUS2347 | RNFUR0466 | RDR046737 | RCPIC00368 | RGOAS0105 | RALGO35402 |
| RPSIN02070 | RMMUS098676 | | RDR022338 | RCPIC00959 | RGOAS0100 | RALGO10345 |
| RPSIN01724 | RMMUS1614 | RNFUR0447 | RDR036824 | RCPIC02786 | RGOAS0052 | RALGO5185 |
| RPSIN00418 | RMMUS2631 | RNFUR0226 | RDR027880 | RCPIC03984 | RGOAS0246 | RALGO47572 |
| RPSIN02280 | RMMUS0087 | RNFUR0149 | RDR019617 | RCPIC03254 | RGOAS0180 | RALGO02035 |
| RPSIN01066 | RMMUS1878 | RNFUR0493 | RDR005824 | RCPIC02690 | RGOAS0289 | RALGO10393 |
| 11 | RMMUS2757 | RNFUR0491 | RDR017907 | RCPIC05186 | RGOAS0126 | RALGO24244 |
| RPSIN03219 | RMMUS2358 | RNFUR052673 | | RCPIC05718 | RGOAS0216 | RALGO2118 |
| RPSIN01915 | RMMUS2823 | RNFUR0180 | RDR014982 | RCPIC03063 | RGOAS0340 | RALGO43801 |
| RPSIN03725 | RMMUS195148 | | | RCPIC004783 | | RALGO44102 |
| RPSIN01844 | RMMUS2047 | RNFUR0211 | RDR003739 | RCPIC00354 | RGOAS0197 | RALGO54472 |
| RPSIN01008 | RMMUS0037 | RNFUR0309 | RDR000815 | RCPIC01426 | RGOAS0169 | RALGO2599 |
| RPSIN03395 | RMMUS2723 | RNFUR0031 | RDR025744 | RCPIC00924 | RGOAS0123 | RALGO18063 |
| RPSIN00096 | RMMUS0810 | RNFUR0080 | RDR039994 | RCPIC02513 | RGOAS0191 | RALGO2181 |
| RPSIN00604 | RMMUS1382 | RNFUR0450 | RDR035079 | RCPIC01517 | RGOAS0305 | RALGO1018 |
| RPSIN02814 | RMMUS0417 | RNFUR0303 | RDR036555 | RCPIC00312 | RGOAS0005 | RALGO6822 |
| | RMMUS205740 | | | | | not predicte |
| RPSIN00081 | RMMUS1976 | RNFUR0359 | RDR038220 | RCPIC041282 | | RALGO6173 |
| RPSIN00754 | RMMUS0869 | RNFUR0185 | RDR029890 | RCPIC02833 | RGOAS0279 | RALGO0462 |
| RPSIN02079 | RMMUS2057 | RNFUR0050 | RDR015487 | RCPIC00572 | RGOAS0089 | RALGO2298 |
| RPSIN02078 | RMMUS2053 | RNFUR0050 | RDR009601 | RCPIC03295 | RGOAS0023 | RALGO2293 |
| 65 | RMMUS252581 | | | | | |
| | RMMUS252734 | | | | | not predicte |
| | RMMUS252731 | | | | | not predicte |
| RPSIN02867 | RMMUS2527 | RNFUR038047 | | | | RALGO01017 |
| RPSIN01979 | RMMUS1849 | RNFUR0007 | RDR004733 | RCPIC04020 | RGOAS0276 | RALGO34174 |
| RPSIN00767 | RMMUS0423 | RNFUR0546 | RDR003635 | RCPIC01808 | RGOAS0248 | RALGO28781 |
| RPSIN03667 | RMMUS135806 | | | RCPIC04644 | RGOAS0211 | RALGO5470 |
| RPSIN00587 | RMMUS083592 | | | RCPIC04163 | RGOAS0260 | RALGO24597 |
| RPSIN03722 | RMMUS094547 | | | RCPIC00399 | RGOAS0078 | RALGO45822 |
| RPSIN00097 | RMMUS0808 | RNFUR0188 | RDR020806 | RCPIC02508 | RGOAS0040 | RALGO21231 |
| RPSIN00127 | RMMUS1313 | RNFUR023380 | | RCPIC03155 | RGOAS0277 | RALGO5918 |
| RPSIN03719 | RMMUS0946 | RNFUR0123 | RDR036448 | RCPIC00288 | RGOAS0326 | RALGO46101 |
| RPSIN03726 | RMMUS0946 | RNFUR0125 | RDR031499 | RCPIC002892 | | RALGO4614 |
| RPSIN03719 | RMMUS0946 | RNFUR0125 | RDR031164 | RCPIC00289 | RGOAS0327 | RALGO46161 |
| RPSIN03719 | RMMUS0946 | RNFUR0125 | RDR031169 | RCPIC00289 | RGOAS0327 | RALGO46164 |
| RPSIN03718 | RMMUS0946 | RNFUR0124 | RDR031163 | RCPIC00288 | RGOAS0326 | RALGO46104 |
| RPSIN03721 | RMMUS0946 | RNFUR012472 | | RCPIC00288 | RGOAS0326 | RALGO4611 |

| | | | | | | |
|---|---|---|---|---|---|---|
| RPSIN037203 | RMMUS094635 | | | RCPIC00288 | RGOAS03268 | RALGO46125 |
| RPSIN037194 | RMMUS0946 | RNFUR010822 | | RCPIC00288 | RGOAS03269 | RALGO46131 |
| RPSIN037193 | RMMUS094643 | | | RCPIC00288 | RGOAS03269 | RALGO46134 |
| RPSIN037191 | RMMUS0946 | RNFUR012487 | | RCPIC00289 | RGOAS03269 | RALGO46137 |
| RPSIN037241 | RMMUS0946 | RNFUR012494 | | RCPIC00289 | RGOAS03270 | RALGO46140 |
| RPSIN013442 | RMMUS1858 | RNFUR02477 | RDR005007 | RCPIC04884 | RGOAS02805 | RALGO08074 |
| | RMMUS1857 | RNFUR03420 | RDR004693 | RCPIC04888 | RGOAS02308 | RALGO08311 |
| RPSIN013440 | RMMUS1858 | RNFUR04947 | RDR005008 | RCPIC04884 | RGOAS02805 | RALGO08104 |
| RPSIN013459 | RMMUS1858 | RNFUR04947 | RDR005009 | RCPIC04900 | RGOAS02804 | RALGO08119 |
| RPSIN013460 | RMMUS1858 | RNFUR04949 | RDR005010 | RCPIC04884 | RGOAS02804 | RALGO08152 |
| RPSIN013438 | RMMUS1858 | RNFUR02475 | RDR005005 | RCPIC04884 | RGOAS02804 | RALGO08170 |
| RPSIN013439 | RMMUS1858 | RNFUR02471 | RDR005011 | RCPIC04884 | RGOAS02804 | RALGO08179 |
| RPSIN013437 | RMMUS185822 | | RDR004850 | RCPIC04884 | RGOAS02804 | RALGO08191 |
| RPSIN013435 | RMMUS1858 | RNFUR04949 | RDR005686 | RCPIC04885 | RGOAS02306 | RALGO08206 |
| RPSIN013434 | RMMUS185807 | | RDR005013 | RCPIC04894 | RGOAS02307 | RALGO08218 |
| RPSIN025977 | RMMUS237495 | | | RCPIC043365 | | RALGO19387 |
| RPSIN025978 | RMMUS2374 | RNFUR03695 | RDR043587 | RCPIC04336 | RGOAS00804 | RALGO19360 |
| RPSIN025989 | RMMUS2374 | RNFUR03715 | RDR043443 | RCPIC04336 | RGOAS00804 | RALGO19330 |
| RPSIN025979 | RMMUS2374 | RNFUR03714 | RDR043600 | RCPIC04336 | RGOAS00805 | RALGO19321 |
| RPSIN025974 | RMMUS2375 | RNFUR03721 | RDR043582 | RCPIC04351 | RGOAS00802 | RALGO19468 |
| RPSIN025973 | RMMUS2375 | RNFUR03720 | RDR043584 | RCPIC04336 | RGOAS00802 | RALGO19438 |
| RPSIN025981 | RMMUS2375 | RNFUR03722 | RDR043583 | RCPIC04336 | RGOAS00803 | RALGO19432 |
| RPSIN025972 | RMMUS2375 | RNFUR03718 | RDR043285 | RCPIC04336 | RGOAS00803 | RALGO19408 |
| RPSIN025971 | RMMUS2375 | RNFUR03718 | RDR043598 | RCPIC04338 | RGOAS00803 | RALGO32914 |
| RPSIN012091 | RMMUS027393 | | | RCPIC03500 | RGOAS00301 | RALGO32887 |
| RPSIN012096 | RMMUS0273 | RNFUR05012 | RDR019156 | RCPIC03497 | RGOAS00303 | RALGO32917 |
| RPSIN012094 | RMMUS0273 | RNFUR050125 | | RCPIC03497 | RGOAS00304 | RALGO32920 |
| RPSIN012097 | RMMUS0273 | RNFUR05012 | RDR019614 | RCPIC03497 | RGOAS00304 | RALGO32923 |
| RPSIN012098 | RMMUS027349 | | | RCPIC03497 | RGOAS00305 | RALGO32926 |
| RPSIN012093 | RMMUS0273 | RNFUR05011 | RDR019151 | RCPIC03497 | RGOAS00301 | RALGO32896 |
| RPSIN012092 | RMMUS0273 | RNFUR05010 | RDR019008 | RCPIC03497 | RGOAS00301 | RALGO19468 |
| RPSIN012149 | RMMUS027372 | | | RCPIC03497 | RGOAS00302 | RALGO32908 |
| RPSIN012224 | RMMUS027364 | | | RCPIC03497 | RGOAS00303 | RALGO32914 |
| | RMMUS140143 | | | RCPIC05290 | RGOAS00190 | not predicted |
| RPSIN027123 | RMMUS1935 | RNFUR04778 | RDR036525 | RCPIC03613 | RGOAS02509 | RALGO29741 |
| RPSIN019416 | RMMUS1272 | RNFUR03768 | RDR046449 | RCPIC04521 | RGOAS02554 | RALGO34724 |
| RPSIN009722 | RMMUS1004 | RNFUR01069 | RDR022467 | RCPIC00972 | RGOAS01465 | RALGO06187 |
| | | | | | | not predicted |
| 44 | RMMUS1795 | RNFUR05447 | RDR024087 | RCPIC04044 | RGOAS00186 | RALGO28655 |
| | RMMUS141132 | | RDR034880 | RCPIC01514 | RGOAS02197 | not predicted |
| RPSIN000330 | RMMUS2853 | RNFUR04330 | RDR043295 | RCPIC05178 | RGOAS00721 | RALGO23956 |
| 50 | RMMUS2318 | RNFUR04049 | RDR037027 | RCPIC03624 | RGOAS01584 | RALGO70039 |
| RPSIN029540 | RMMUS1626 | RNFUR00611 | RDR038287 | RCPIC00541 | RGOAS00510 | RALGO53504 |
| RPSIN014152 | RMMUS139265 | | RDR034779 | RCPIC01535 | RGOAS006254 | |
| 45 | RMMUS1845 | RNFUR02706 | RDR009681 | RCPIC05160 | RGOAS02025 | RALGO02821 |

| | | | | | | |
|---|---|---|---|---|---|---|
| RPSIN012834 | RMMUS2021 | RNFUR02482 | RDR037982 | RCPIC02936 | RGOAS01657 | RALGO59723 |
| 40 | RMMUS2550 | RNFUR00092 | RDR038268 | RCPIC00599 | RGOAS01778 | RALGO62470 |
| RPSIN036717 | RMMUS1676 | RNFUR03774 | RDR007408 | RCPIC02120 | RGOAS00656 | RALGO55613 |
| RPSIN014196 | RMMUS2749 | RNFUR02585 | RDR033820 | RCPIC02287 | RGOAS00577 | RALGO40578 |
| 61 | RMMUS034179 | | RDR014400 | RCPIC02364 | RGOAS03283 | RALGO61065 |
| RPSIN035321 | RMMUS2448 | RNFUR02149 | RDR029099 | RCPIC02375 | RGOAS01295 | RALGO65704 |
| RPSIN009097 | RMMUS0182 | RNFUR00548 | RDR049534 | RCPIC03221 | RGOAS01123 | RALGO20294 |
| | RMMUS253145 | | | | | |
| | RMMUS253148 | | RDR017001 | | | |
| RPSIN031414 | RMMUS197828 | | | RCPIC04124 | RGOAS00936 | RALGO61473 |
| RPSIN024280 | RMMUS1771 | RNFUR00700 | RDR040187 | RCPIC01243 | RGOAS02509 | RALGO53261 |
| RPSIN005448 | RMMUS0237 | RNFUR04514 | RDR010291 | RCPIC02018 | RGOAS01241 | RALGO25954 |
| RPSIN008452 | RMMUS1467 | RNFUR02857 | RDR020803 | RCPIC01450 | RGOAS01897 | RALGO32821 |
| RPSIN007426 | RMMUS2625 | RNFUR02996 | RDR027890 | RCPIC01259 | RGOAS02905 | RALGO48950 |
| RPSIN009224 | RMMUS0870 | RNFUR01612 | RDR009474 | RCPIC02831 | RGOAS02720 | RALGO08347 |
| RPSIN030579 | RMMUS0629 | RNFUR04108 | RDR009640 | RCPIC01198 | RGOAS01875 | RALGO15467 |
| RPSIN001393 | RMMUS148586 | | RDR009519 | RCPIC01442 | RGOAS02663 | RALGO34715 |
| RPSIN038709 | RMMUS214215 | | RDR009699 | RCPIC02773 | RGOAS01031 | RALGO31615 |
| RPSIN010576 | RMMUS111980 | | | | | |
| RPSIN031534 | RMMUS0839 | RNFUR03189 | RDR009742 | RCPIC04565 | RGOAS02189 | RALGO25882 |
| 15 | RMMUS187251 | | | RCPIC036260 | | RALGO69859 |
| RPSIN010139 | RMMUS0038 | RNFUR01023 | RDR019186 | RCPIC01419 | RGOAS02180 | RALGO25332 |
| RPSIN020055 | RMMUS089874 | | | RCPIC04497 | RGOAS02794 | RALGO70702 |
| RPSIN012576 | RMMUS0970 | RNFUR03584 | RDR021977 | | | RALGO61627 |
| RPSIN023883 | RMMUS076002 | | RDR000934 | RCPIC02282 | RGOAS03339 | RALGO58445 |
| RPSIN034534 | RMMUS1735 | RNFUR02549 | RDR048104 | RCPIC00248 | RGOAS00303 | RALGO55878 |
| RPSIN018073 | RMMUS1472 | RNFUR02981 | RDR021069 | RCPIC01460 | RGOAS00048 | RALGO37655 |
| | RMMUS144331 | | | | | RALGO11663 |
| RPSIN005365 | RMMUS004758 | | | RCPIC01419 | RGOAS01790 | RALGO20958 |
| RPSIN005364 | RMMUS0051 | RNFUR03439 | RDR019473 | RCPIC03260 | RGOAS01050 | RALGO18342 |
| 96 | RMMUS1181 | RNFUR017370 | | RCPIC02557 | RGOAS00786 | RALGO50725 |
| RPSIN004108 | RMMUS131019 | | | RCPIC054144 | | RALGO63931 |
| | | | | | | not predicted |
| | | | | | | not predicted |
| RPSIN021976 | RMMUS1349 | RNFUR00090 | RDR002298 | RCPIC03787 | RGOAS00146 | RALGO40882 |
| RPSIN020625 | RMMUS1875 | RNFUR04586 | RDR006862 | RCPIC04768 | RGOAS00946 | RALGO00975 |
| | | | | | RGOAS00190 | not predicted |
| RPSIN006204 | RMMUS0261 | RNFUR04301 | RDR019105 | RCPIC03511 | RGOAS00825 | RALGO02344 |
| | | | | | | not predicted |
| | | | | | | not predicted |
| | | | | | | not predicted |
| RPSIN014848 | | | RDR024217 | RCPIC02612 | RGOAS01881 | RALGO29867 |
| RPSIN028255 | | | | | | RALGO33214 |
| | | | | | | not predicted |

| | | | | | | |
|---|---|---|---|---|---|---|
| | | | | | | not predicted |
| | RMMUS061200 | | RDR004827 | | | not predicted |
| | | | | | | not predicted |
| | | | | | | not predicted |
| | | | | | | not predicted |
| | | | | | | not predicted |
| | | | | | | not predicted |
| | | | | | | not predicted |
| | | | | | | not predicted |
| | | | | | | not predicted |
| | | | | | | not predicted |
| | | | | | | not predicted |
| | RMMUS183506 | | | | | not predicted |
| | | | | | | not predicted |
| RPSIN020945 | RMMUS245886 | | RDR020180 | RCPIC02372 | RGOAS02347 | RALGO66370 |
| RPSIN020947 | RMMUS245865 | | | RCPIC02372 | RGOAS03167 | RALGO66364 |
| | RMMUS061178 | | | RCPIC011520 | | RALGO56889 |
| RPSIN026179 | RMMUS061289 | | | RCPIC01151 | RGOAS01505 | RALGO55283 |
| RPSIN014934 | RMMUS170274 | | | RCPIC03366 | RGOAS02470 | not predicted |
| RPSIN003382 | RMMUS160179 | | | RCPIC00652 | RGOAS03024 | RALGO55866 |
| RPSIN021017 | RMMUS245896 | | | RCPIC02372 | RGOAS01413 | RALGO66376 |
| RPSIN026182 | RMMUS056593 | | | | RGOAS02071 | RALGO55229 |
| | | | | | | not predicted |
| | | | | | | not predicted |
| RPSIN022901 | RMMUS111098 | | | RCPIC03263 | RGOAS01474 | RALGO13316 |
| RPSIN026782 | RMMUS068324 | | | RCPIC04935 | RGOAS00171 | RALGO11074 |
| RPSIN029661 | | | | | | not predicted |
| RPSIN036660 | RMMUS1677 | RNFUR04873 | RDR007395 | RCPIC00060 | RGOAS03330 | RALGO55731 |
| RPSIN025174 | RMMUS1174 | RNFUR01799 | RDR034816 | RCPIC02552 | RGOAS03413 | RALGO48821 |
| RPSIN006479 | RMMUS2477 | RNFUR01072 | RDR037920 | RCPIC04047 | RGOAS02838 | RALGO58253 |
| RPSIN030379 | RMMUS1734 | RNFUR01797 | RDR038386 | RCPIC00310 | RGOAS01050 | RALGO55677 |
| RPSIN005444 | RMMUS1867 | RNFUR010431 | | RCPIC04877 | RGOAS00916 | RALGO03897 |
| RPSIN025936 | RMMUS262867 | | RDR040081 | RCPIC03983 | RGOAS00005 | RALGO48190 |
| RPSIN030387 | RMMUS0590 | RNFUR00750 | RDR028863 | RCPIC03583 | RGOAS02516 | RALGO60029 |
| RPSIN004111 | RMMUS1308 | RNFUR03342 | RDR017071 | RCPIC05105 | RGOAS01147 | RALGO64030 |
| RPSIN035720 | RMMUS0108 | RNFUR043041 | | RCPIC00841 | RGOAS01207 | RALGO17838 |
| RPSIN026277 | RMMUS2805 | RNFUR02095 | RDR043315 | RCPIC03650 | RGOAS00972 | RALGO25239 |
| RPSIN034570 | RMMUS1736 | RNFUR05453 | RDR026351 | RCPIC00247 | RGOAS01430 | RALGO56136 |
| RPSIN035766 | RMMUS0108 | RNFUR04329 | RDR022434 | RCPIC00824 | RGOAS01024 | RALGO17571 |
| RPSIN017524 | RMMUS045569 | | | RCPIC01972 | RGOAS00854 | RALGO24852 |
| RPSIN036859 | RMMUS1758 | RNFUR02181 | RDR027464 | RCPIC012267 | | not predicted |
| RPSIN022020 | RMMUS1350 | RNFUR034655 | | RCPIC037804 | | RALGO40924 |
| RPSIN019640 | RMMUS0956 | RNFUR02242 | RDR013588 | RCPIC01239 | RGOAS01740 | RALGO22416 |
| | RMMUS177278 | | | | | not predicted |
| RPSIN033159 | RMMUS277378 | | | RCPIC01383 | RGOAS00019 | RALGO54901 |

| | | | | | | |
|---|---|---|---|---|---|---|
| RPSIN015028 | RMMUS136409 | | | RCPIC01091 | RGOAS03141 | RALGO42284 |
| RPSIN038367 | RMMUS118749 | | RDR015113 | RCPIC02581 | RGOAS01579 | RALGO46332 |
| RPSIN035470 | RMMUS001134 | | | | | not predicted |
| RPSIN036937 | RMMUS264799 | | | RCPIC01227 | RGOAS02537 | RALGO51763 |
| RPSIN016561 | RMMUS1417 | RNFUR04816 | RDR034486 | RCPIC01574 | RGOAS02461 | RALGO33766 |
| RPSIN035473 | RMMUS0011 | RNFUR01120 | RDR037827 | RCPIC05062 | RGOAS01919 | RALGO14687 |
| RPSIN027291 | RMMUS2174 | RNFUR01443 | RDR022690 | RCPIC04370 | RGOAS01245 | RALGO20021 |
| | RMMUS1002 | RNFUR053030 | | RCPIC04816 | RGOAS00867 | RALGO36296 |
| RPSIN001477 | RMMUS148511 | | | RCPIC01442 | RGOAS00860 | RALGO34844 |
| RPSIN023154 | RMMUS002925 | | | RCPIC001331 | | RALGO68782 |
| RPSIN023158 | RMMUS002939 | | | RCPIC00135 | RGOAS02494 | not predicted |
| | RMMUS010826 | | | | | not predicted |
| | RMMUS033783 | | | | | not predicted |
| RPSIN012519 | RMMUS0337 | RNFUR04370 | RDR021077 | RCPIC031935 | | RALGO61792 |
| RPSIN030255 | RMMUS002874 | | RDR020462 | RCPIC03596 | RGOAS02775 | RALGO68773 |
| RPSIN030264 | RMMUS0028 | RNFUR019628 | | RCPIC035996 | | RALGO68767 |
| RPSIN035499 | RMMUS2411 | RNFUR001859 | | RCPIC04745 | RGOAS02448 | RALGO05172 |
| RPSIN023153 | RMMUS002909 | | RDR020473 | RCPIC000942 | | not predicted |
| RPSIN030269 | RMMUS002893 | | | RCPIC03596 | RGOAS02775 | RALGO68776 |
| | RMMUS043557 | | | | | |
| | RMMUS010818 | | RDR022532 | RCPIC00858 | RGOAS01613 | RALGO17520 |
| RPSIN003383 | RMMUS1601 | RNFUR00106 | RDR038032 | RCPIC007251 | | RALGO55857 |
| RPSIN019018 | RMMUS1546 | RNFUR04327 | RDR003107 | RCPIC03708 | RGOAS00937 | RALGO34348 |
| | RMMUS033783 | | | | | not predicted |
| RPSIN017340 | RMMUS1244 | RNFUR00980 | RDR004832 | RCPIC01762 | RGOAS018214 | |
| RPSIN014965 | RMMUS170235 | | | RCPIC03366 | RGOAS01282 | RALGO46338 |
| 36 | RMMUS171525 | | | RCPIC04791 | RGOAS00919 | RALGO14978 |
| RPSIN019655 | RMMUS095674 | | | RCPIC01301 | RGOAS01738 | RALGO22356 |
| | RMMUS033783 | | | | | not predicted |
| 93 | RMMUS221563 | | | RCPIC049969 | | not predicted |
| RPSIN014966 | | | | RCPIC03369 | RGOAS01281 | RALGO46335 |
| 51 | RMMUS124696 | | | | | RALGO00786 |
| 92 | RMMUS136750 | | | RCPIC04854 | RGOAS00489 | RALGO10291 |
| RPSIN011481 | RMMUS232543 | | RDR005697 | RCPIC04646 | RGOAS01225 | RALGO54451 |
| RPSIN014624 | RMMUS2820 | RNFUR01714 | RDR020823 | RCPIC01284 | RGOAS00748 | RALGO55448 |
| | RMMUS033783 | | | | | not predicted |
| RPSIN038718 | RMMUS214053 | | | RCPIC02784 | RGOAS00746 | RALGO32368 |
| | RMMUS033783 | | | | | not predicted |
| | RMMUS033783 | | | | | not predicted |
| RPSIN004879 | RMMUS1404 | RNFUR04804 | RDR034718 | RCPIC03402 | RGOAS01423 | RALGO05151 |
| | RMMUS033783 | | | | | not predicted |
| | RMMUS033783 | | | | | not predicted |
| | RMMUS033783 | | | | | not predicted |
| RPSIN012529 | RMMUS020209 | | | RCPIC03204 | RGOAS02609 | RALGO61844 |
| | RMMUS033783 | | | | | not predicted |

| | | | | | | |
|---|---|---|---|---|---|---|
| | RMMUS033783 | | | | | not predicted |
| RPSIN017331 | RMMUS1244 | RNFUR006264 | | RCPIC01822 | RGOAS01821 | not predicted |
| | RMMUS033783 | | | | | not predicted |
| RPSIN029805 | RMMUS099707 | | | RCPIC00935 | RGOAS02702 | RALGO28667 |
| RPSIN037168 | RMMUS074910 | | | RCPIC05065 | RGOAS01323 | RALGO45023 |
| RPSIN038723 | RMMUS2140 | RNFUR00366 | RDR021244 | RCPIC02778 | RGOAS00747 | RALGO32446 |
| RPSIN020698 | RMMUS2273 | RNFUR00279 | RDR039889 | RCPIC03390 | RGOAS03379 | RALGO27506 |
| | RMMUS033783 | | | | | not predicted |
| RPSIN001835 | RMMUS0479 | RNFUR04822 | RDR036585 | RCPIC04551 | RGOAS01540 | RALGO41948 |
| RPSIN009427 | RMMUS1445 | RNFUR00645 | RDR005396 | RCPIC04862 | RGOAS00182 | RALGO11044 |
| RPSIN027476 | RMMUS0080 | RNFUR01079 | RDR057057 | RCPIC01842 | RGOAS00167 | RALGO39315 |
| RPSIN021999 | RMMUS1287 | RNFUR05342 | RDR002616 | RCPIC03787 | RGOAS02802 | RALGO40996 |
| RPSIN025044 | RMMUS1226 | RNFUR00926 | RDR014913 | RCPIC03079 | RGOAS02625 | RALGO41663 |
| 19 | RMMUS2803 | RNFUR04782 | RDR017587 | RCPIC053575 | | RALGO24816 |
| RPSIN028708 | RMMUS1529 | RNFUR00700 | RDR043336 | RCPIC00607 | RGOAS00843 | RALGO44621 |
| RPSIN029779 | RMMUS100303 | | | RCPIC00932 | RGOAS02967 | RALGO26593 |
| | RMMUS033783 | | | | | not predicted |
| RPSIN021548 | RMMUS2354 | RNFUR05324 | RDR046748 | RCPIC00080 | RGOAS02024 | RALGO43368 |
| RPSIN024269 | RMMUS1773 | RNFUR04859 | RDR040151 | RCPIC01259 | RGOAS02781 | RALGO53477 |
| RPSIN026393 | RMMUS1330 | RNFUR00006 | RDR027745 | RCPIC04248 | RGOAS01995 | not predicted |
| 38 | RMMUS1087 | RNFUR02070 | RDR034531 | RCPIC054547 | | RALGO07849 |
| 09 | RMMUS1146 | RNFUR02598 | RDR024166 | RCPIC055028 | | RALGO22566 |
| RPSIN037761 | RMMUS2063 | RNFUR01635 | RDR002562 | RCPIC03445 | RGOAS03090 | RALGO64609 |
| | RMMUS0917 | RNFUR04980 | RDR007311 | RCPIC04356 | RGOAS00778 | RALGO00252 |
| RPSIN011290 | RMMUS0178 | RNFUR04527 | RDR042000 | RCPIC01598 | RGOAS00334 | RALGO15476 |
| RPSIN019431 | RMMUS1273 | RNFUR03757 | RDR046762 | RCPIC03818 | RGOAS01755 | RALGO35114 |
| RPSIN035890 | RMMUS1414 | RNFUR00447 | RDR034424 | RCPIC01513 | RGOAS00696 | RALGO32527 |
| | RMMUS2218 | RNFUR04259 | RDR024397 | RCPIC04993 | RGOAS01076 | not predicted |
| RPSIN008260 | RMMUS2256 | RNFUR03046 | RDR019013 | RCPIC03720 | RGOAS00699 | RALGO69904 |
| | RMMUS033783 | | | | | not predicted |
| RPSIN012713 | RMMUS007063 | | RDR030647 | RCPIC01865 | RGOAS01311 | RALGO32206 |
| RPSIN034318 | RMMUS1294 | RNFUR01376 | RDR020548 | RCPIC03654 | RGOAS01124 | RALGO69160 |
| | RMMUS117913 | | | RCPIC02554 | RGOAS01698 | RALGO47022 |
| RPSIN028349 | RMMUS0369 | RNFUR01236 | RDR012445 | RCPIC02982 | RGOAS03190 | RALGO27917 |
| RPSIN038701 | RMMUS2142 | RNFUR00787 | RDR051551 | RCPIC02779 | RGOAS00449 | RALGO31101 |
| RPSIN015115 | RMMUS027867 | | RDR019833 | RCPIC03500 | RGOAS03212 | RALGO30924 |
| RPSIN000655 | RMMUS1250 | RNFUR03656 | RDR001173 | | RGOAS00104 | RALGO08559 |
| RPSIN013842 | RMMUS1427 | RNFUR04108 | RDR045906 | RCPIC00340 | RGOAS02541 | RALGO30161 |
| RPSIN004250 | RMMUS165025 | | RDR020503 | RCPIC03756 | RGOAS00762 | RALGO10876 |
| RPSIN020634 | RMMUS1885 | RNFUR01650 | RDR024096 | RCPIC04771 | RGOAS02539 | RALGO12737 |
| 47 | RMMUS237201 | | | RCPIC043470 | | RALGO17802 |
| RPSIN037124 | RMMUS1760 | RNFUR02911 | RDR053044 | RCPIC012351 | | RALGO55917 |
| RPSIN016905 | RMMUS2243 | RNFUR01548 | RDR012574 | RCPIC04472 | RGOAS01511 | RALGO70426 |
| RPSIN034135 | RMMUS0348 | RNFUR02965 | RDR026091 | RCPIC00559 | RGOAS02922 | RALGO66202 |
| RPSIN019659 | RMMUS0621 | RNFUR05479 | RDR012516 | RCPIC01238 | RGOAS02399 | RALGO21099 |

| | | | | | | |
|---|---|---|---|---|---|---|
| RPSIN006587 | RMMUS2713 | RNFUR05168 | RDR010095 | RCPIC01117 | RGOAS02145 | RALGO52685 |
| | RMMUS135463 | | | RCPIC037725 | | RALGO40062 |
| RPSIN032877 | RMMUS1453 | RNFUR02631 | RDR039841 | RCPIC01458 | RGOAS01715 | RALGO32065 |
| RPSIN030358 | RMMUS0947 | RNFUR01265 | RDR032259 | RCPIC00289 | RGOAS01917 | RALGO46233 |
| RPSIN030607 | RMMUS0616 | RNFUR04166 | RDR038321 | RCPIC01196 | RGOAS02978 | RALGO18027 |
| RPSIN002285 | RMMUS1372 | RNFUR01377 | RDR000721 | RCPIC04852 | RGOAS00181 | RALGO09195 |
| | RMMUS1349 | RNFUR04671 | RDR002585 | RCPIC05456 | RGOAS02275 | RALGO40837 |
| RPSIN021363 | RMMUS2167 | RNFUR04641 | RDR026722 | RCPIC02463 | RGOAS01821 | RALGO02923 |
| RPSIN021363 | RMMUS2167 | RNFUR04641 | RDR026722 | RCPIC02463 | RGOAS01821 | RALGO02923 |
| RPSIN001343 | RMMUS1456 | RNFUR00983 | RDR035944 | RCPIC01455 | RGOAS01548 | RALGO29966 |
| RPSIN001630 | RMMUS2750 | RNFUR00402 | RDR032977 | RCPIC02287 | RGOAS02390 | RALGO40416 |
| RPSIN005482 | RMMUS2169 | RNFUR05058 | RDR026374 | RCPIC02480 | RGOAS03079 | RALGO05809 |
| RPSIN020857 | RMMUS0653 | RNFUR041345 | | RCPIC01995 | RGOAS01544 | RALGO07789 |
| RPSIN014262 | RMMUS012459 | | | RCPIC03978 | RGOAS03303 | RALGO38087 |
| RPSIN022468 | RMMUS2609 | RNFUR01402 | RDR038906 | RCPIC02167 | RGOAS00275 | RALGO67189 |
| RPSIN033336 | RMMUS1014 | RNFUR00559 | RDR048142 | RCPIC00083 | RGOAS00659 | RALGO57130 |
| 46 | RMMUS2854 | RNFUR04485 | RDR018186 | RCPIC051854 | | RALGO23881 |
| RPSIN036598 | RMMUS2765 | RNFUR00087 | RDR019526 | RCPIC03166 | RGOAS02187 | RALGO67207 |
| RPSIN004448 | RMMUS0787 | RNFUR02945 | RDR037830 | RCPIC04255 | RGOAS02931 | RALGO52826 |
| RPSIN016760 | RMMUS0097 | RNFUR01621 | RDR002896 | RCPIC03437 | RGOAS02389 | RALGO64453 |
| RPSIN036816 | RMMUS0925 | RNFUR01414 | RDR056614 | RCPIC000426 | | RALGO55965 |
| RPSIN033508 | RMMUS2231 | RNFUR02895 | RDR034398 | RCPIC007893 | | RALGO10948 |
| RPSIN016544 | RMMUS2602 | RNFUR04199 | RDR024490 | RCPIC00278 | RGOAS02180 | RALGO30536 |
| | | | | | | not predicted |
| | | | | | | not predicted |
| | | | | | | not predicted |
| | | | | | | not predicted |
| RPSIN004405 | RMMUS0787 | RNFUR02895 | RDR038263 | RCPIC04275 | RGOAS02931 | RALGO52871 |
| RPSIN031599 | RMMUS0900 | RNFUR02520 | RDR022165 | RCPIC044740 | | RALGO70675 |
| RPSIN001236 | RMMUS0590 | RNFUR00167 | RDR039888 | RCPIC035836 | | RALGO59894 |
| RPSIN002316 | RMMUS2041 | RNFUR04446 | RDR045349 | RCPIC00280 | RGOAS01724 | RALGO28559 |
| RPSIN019842 | RMMUS2402 | RNFUR01864 | RDR002291 | RCPIC01732 | RGOAS03065 | RALGO37829 |
| 56 | RMMUS132704 | | | RCPIC042908 | | |
| 19 | RMMUS103065 | | | RCPIC052216 | | not predicted |
| | RMMUS080099 | | | | | not predicted |
| | RMMUS103257 | | | RCPIC05121 | RGOAS02242 | RALGO62104 |
| RPSIN000613 | | | | RCPIC05376 | RGOAS02091 | RALGO04722 |
| | | | | RCPIC052886 | | RALGO05314 |
| RPSIN003171 | RMMUS101822 | | | RCPIC05126 | RGOAS02433 | RALGO60963 |
| | RMMUS092671 | | | | | not predicted |
| | | | | | | not predicted |
| RPSIN014115 | RMMUS1475 | RNFUR04591 | RDR029124 | RCPIC01447 | RGOAS02648 | RALGO37379 |
| RPSIN025358 | RMMUS0744 | RNFUR01660 | RDR045493 | RCPIC00344 | RGOAS00152 | RALGO20910 |

| | | | | | | |
|---|---|---|---|---|---|---|
| 31 | RMMUS236197 | | RDR044474 | RCPIC027458 | | RALGO13964 |
| RPSIN023234 | RMMUS1049 | RNFUR05214 | RDR047276 | RCPIC00161 | RGOAS01343 | RALGO58727 |
| RPSIN013028 | RMMUS2198 | RNFUR04517 | RDR033470 | RCPIC04133 | RGOAS00636 | RALGO62476 |
| 22 | RMMUS1023 | RNFUR005746 | | RCPIC02228 | RGOAS02818 | RALGO63144 |
| | RMMUS105993 | | | | RGOAS00285 | not predicted |
| RPSIN037978 | RMMUS1297 | RNFUR04348 | RDR000592 | RCPIC03656 | RGOAS00843 | RALGO69046 |
| RPSIN002154 | RMMUS1209 | RNFUR00838 | RDR029204 | RCPIC03864 | RGOAS00187 | RALGO71647 |
| RPSIN004940 | RMMUS0471 | RNFUR00974 | RDR030745 | RCPIC04578 | RGOAS02671 | RALGO43193 |
| RPSIN022620 | RMMUS0879 | RNFUR04092 | RDR007089 | RCPIC05476 | RGOAS03083 | RALGO25455 |
| RPSIN021190 | RMMUS1863 | RNFUR00611 | RDR005179 | RCPIC04900 | RGOAS00504 | RALGO05863 |
| 47 | RMMUS124498 | | | | | RALGO00417 |
| RPSIN034192 | RMMUS086700 | | | RCPIC02853 | RGOAS03220 | RALGO11756 |
| RPSIN006356 | RMMUS1594 | RNFUR01197 | RDR037814 | RCPIC00729 | RGOAS02589 | RALGO57248 |
| RPSIN037853 | RMMUS2223 | RNFUR01774 | RDR019024 | RCPIC04463 | RGOAS00692 | RALGO70864 |
| | RMMUS037850 | | RDR019311 | | | |
| RPSIN009672 | RMMUS0667 | RNFUR03296 | RDR032702 | RCPIC04934 | RGOAS03124 | RALGO09135 |
| RPSIN010239 | RMMUS2244 | RNFUR01560 | RDR019164 | RCPIC04459 | RGOAS02946 | RALGO70396 |
| RPSIN007430 | RMMUS1754 | RNFUR01151 | RDR040188 | RCPIC01316 | RGOAS00944 | RALGO49637 |
| 36 | RMMUS091514 | | | RCPIC03550 | RGOAS00191 | RALGO13139 |
| RPSIN019645 | RMMUS0622 | RNFUR01153 | RDR012459 | RCPIC01238 | RGOAS01485 | RALGO21375 |
| RPSIN003556 | RMMUS0884 | RNFUR00133 | RDR004999 | RCPIC01772 | RGOAS02861 | RALGO13076 |
| RPSIN013014 | RMMUS2196 | RNFUR02520 | RDR033205 | RCPIC04140 | RGOAS01535 | not predicted |
| RPSIN001770 | RMMUS0311 | RNFUR03746 | RDR016393 | RCPIC00821 | RGOAS00836 | RALGO45354 |
| RPSIN019491 | RMMUS0442 | RNFUR02034 | RDR020631 | RCPIC044650 | | RALGO70570 |
| RPSIN038659 | RMMUS2271 | RNFUR02763 | RDR009499 | RCPIC04775 | RGOAS00500 | RALGO28370 |
| RPSIN030015 | RMMUS1077 | RNFUR04359 | RDR002569 | RCPIC05503 | RGOAS00084 | RALGO10300 |
| RPSIN012314 | RMMUS1835 | RNFUR04852 | RDR009567 | RCPIC02877 | RGOAS00394 | RALGO07144 |
| RPSIN007149 | RMMUS1293 | RNFUR00461 | RDR000262 | RCPIC04803 | RGOAS00764 | RALGO69187 |
| | | | | | | not predicted |
| | | | | | | not predicted |
| | RMMUS106071 | | | | | not predicted |
| RPSIN020220 | RMMUS0566 | RNFUR04144 | RDR028792 | RCPIC01107 | RGOAS01546 | RALGO57332 |
| RPSIN004995 | RMMUS0471 | RNFUR00968 | RDR031195 | RCPIC04595 | RGOAS02669 | RALGO43245 |
| RPSIN026335 | RMMUS2642 | RNFUR02774 | RDR020978 | RCPIC02120 | RGOAS03425 | RALGO49901 |
| RPSIN020363 | RMMUS166412 | | RDR041948 | RCPIC05215 | RGOAS00658 | RALGO33331 |
| RPSIN023275 | RMMUS105001 | | | RCPIC00087 | RGOAS01146 | RALGO58742 |
| RPSIN021341 | RMMUS1224 | RNFUR04445 | RDR014829 | RCPIC03086 | RGOAS02301 | RALGO41116 |
| RPSIN022671 | RMMUS0306 | RNFUR03003 | RDR034796 | RCPIC00814 | RGOAS01754 | RALGO47743 |
| RPSIN013170 | RMMUS0535 | RNFUR01700 | RDR012683 | RCPIC03936 | RGOAS01367 | RALGO29834 |
| RPSIN027115 | RMMUS0234 | RNFUR03665 | RDR016875 | RCPIC02020 | RGOAS01021 | RALGO26834 |
| RPSIN015869 | RMMUS2410 | RNFUR01046 | RDR012646 | RCPIC04745 | RGOAS02905 | RALGO06262 |
| 41 | RMMUS270145 | | | RCPIC01578 | RGOAS02811 | RALGO10906 |
| RPSIN021523 | RMMUS1708 | RNFUR039430 | | RCPIC00114 | RGOAS00665 | RALGO44336 |

| | | | | | | |
|---|---|---|---|---|---|---|
| RPSIN009018 | RMMUS0246 | RNFUR04980 | RDR020405 | RCPIC03547 | RGOAS00998 | RALGO15575 |
| RPSIN026858 | RMMUS1126 | RNFUR05018 | RDR047470 | RCPIC01508 | RGOAS02043 | RALGO26488 |
| RPSIN032603 | RMMUS127806 | | | RCPIC03820 | RGOAS02409 | RALGO37415 |
| RPSIN021813 | RMMUS2219 | RNFUR04021 | RDR014686 | RCPIC02254 | RGOAS01275 | RALGO34285 |
| | RMMUS106071 | | | | | not predicted |
| 41 | RMMUS2532 | RNFUR004768 | | RCPIC054436 | | RALGO03166 |
| RPSIN002009 | RMMUS025918 | | RDR023197 | RCPIC03504 | RGOAS01069 | RALGO38540 |
| RPSIN037748 | RMMUS206920 | | RDR045709 | RCPIC03443 | RGOAS03109 | RALGO64801 |
| RPSIN013054 | RMMUS015240 | | | RCPIC046076 | | not predicted |
| RPSIN028114 | RMMUS000972 | | | RCPIC00325 | RGOAS03432 | RALGO68118 |
| 04 | RMMUS1370 | RNFUR02600 | RDR013164 | | | RALGO08479 |
| RPSIN021102 | RMMUS0549 | RNFUR01139 | RDR003175 | RCPIC02669 | RGOAS02120 | RALGO67588 |
| RPSIN025731 | RMMUS109887 | | | RCPIC05407 | RGOAS00055 | RALGO48154 |
| RPSIN015637 | RMMUS2045 | RNFUR01981 | RDR026902 | RCPIC00741 | RGOAS00671 | RALGO60341 |
| RPSIN014955 | RMMUS1700 | RNFUR007080 | | RCPIC03368 | RGOAS01230 | RALGO46536 |
| RPSIN029670 | RMMUS0705 | RNFUR01792 | RDR016842 | RCPIC01686 | RGOAS00208 | RALGO14726 |
| | RMMUS165515 | | | RCPIC03797 | RGOAS00628 | RALGO38699 |
| RPSIN016160 | RMMUS1409 | RNFUR00474 | RDR046662 | RCPIC01516 | RGOAS03219 | RALGO29768 |
| RPSIN015373 | RMMUS0379 | RNFUR02829 | RDR043578 | RCPIC02959 | RGOAS00190 | RALGO24040 |
| RPSIN037242 | RMMUS0942 | RNFUR03602 | RDR031139 | RCPIC002755 | | RALGO45179 |
| RPSIN024621 | RMMUS1433 | RNFUR01422 | RDR029504 | RCPIC02440 | RGOAS02552 | RALGO71077 |
| RPSIN025141 | RMMUS1507 | RNFUR01824 | RDR034923 | RCPIC02583 | RGOAS00511 | RALGO49712 |
| RPSIN033538 | RMMUS1664 | RNFUR04105 | RDR041655 | RCPIC04202 | RGOAS00153 | RALGO37484 |
| RPSIN003585 | RMMUS178762 | | | RCPIC01769 | RGOAS01179 | RALGO10414 |
| RPSIN027977 | RMMUS1796 | RNFUR03884 | RDR005167 | RCPIC04046 | RGOAS02992 | RALGO26140 |
| RPSIN025120 | RMMUS1894 | RNFUR02730 | RDR024222 | RCPIC04025 | RGOAS03343 | RALGO36959 |
| RPSIN028801 | RMMUS1291 | RNFUR05134 | RDR000408 | RCPIC03110 | RGOAS02760 | RALGO03588 |
| RPSIN038737 | RMMUS213973 | | RDR021256 | RCPIC02780 | RGOAS01349 | RALGO32776 |
| RPSIN017878 | RMMUS2407 | RNFUR05427 | RDR020540 | RCPIC01802 | RGOAS01065 | RALGO40251 |
| RPSIN002237 | RMMUS1882 | RNFUR00778 | RDR005324 | RCPIC04965 | RGOAS00059 | RALGO16645 |
| RPSIN003402 | RMMUS1602 | RNFUR00073 | RDR038167 | RCPIC006524 | | RALGO55815 |
| RPSIN030471 | RMMUS0562 | RNFUR03767 | RDR038920 | RCPIC02797 | RGOAS02770 | RALGO47404 |
| RPSIN000677 | RMMUS2599 | RNFUR00043 | RDR025412 | RCPIC00277 | RGOAS01156 | RALGO31308 |
| RPSIN028241 | RMMUS268527 | | RDR014399 | RCPIC00964 | RGOAS02811 | not predicted |
| RPSIN032614 | RMMUS2394 | RNFUR01220 | RDR010144 | RCPIC04144 | RGOAS02673 | RALGO20153 |
| RPSIN006824 | RMMUS0811 | RNFUR00521 | RDR040502 | RCPIC02506 | RGOAS00520 | RALGO23617 |
| RPSIN016331 | RMMUS234781 | | RDR006919 | RCPIC00264 | RGOAS01055 | RALGO35474 |
| 70 | RMMUS2512 | RNFUR02027 | RDR022661 | RCPIC03676 | RGOAS01095 | RALGO23824 |
| | RMMUS124738 | | | RCPIC05609 | RGOAS00194 | RALGO00927 |
| RPSIN010417 | RMMUS218347 | | | RCPIC02465 | RGOAS03024 | RALGO12698 |
| RPSIN036998 | RMMUS176454 | | | RCPIC01262 | RGOAS01506 | RALGO54940 |
| RPSIN035708 | RMMUS0108 | RNFUR04313 | RDR022785 | RCPIC00823 | RGOAS01205 | RALGO17580 |
| RPSIN023079 | RMMUS155453 | | RDR022515 | RCPIC03748 | RGOAS00399 | RALGO25263 |
| RPSIN002060 | RMMUS134009 | | RDR001870 | RCPIC04257 | RGOAS03270 | RALGO46797 |
| RPSIN010434 | RMMUS1007 | RNFUR03964 | RDR026649 | RCPIC02470 | RGOAS00645 | RALGO13793 |

| | | | | | | |
|---|---|---|---|---|---|---|
| 50 | RMMUS010287 | | RDR019638 | RCPIC001262 | | RALGO45642 |
| RPSIN02232( | RMMUS070707 | | RDR043461 | RCPIC016874 | | RALGO14189 |
| RPSIN038328 | RMMUS0342 | RNFUR02343 | RDR025798 | RCPIC02313 | RGOAS0158( | RALGO60410 |
| RPSIN001008 | RMMUS1979 | RNFUR0406( | RDR038266 | RCPIC01578 | RGOAS01402 | RALGO10513 |
| RPSIN03667 | RMMUS2321 | RNFUR04026 | RDR000626 | RCPIC04644 | RGOAS02197 | RALGO54655 |
| RPSIN02606( | RMMUS1527 | RNFUR0150( | RDR027364 | RCPIC02664 | RGOAS0333( | RALGO43473 |
| RPSIN02978( | RMMUS1006 | RNFUR04823 | RDR018294 | RCPIC00931 | RGOAS02382 | RALGO26500 |
| | RMMUS271522 | | RDR002549 | | | not predicted |
| 36 | RMMUS2538 | RNFUR00481 | RDR032104 | RCPIC03609 | RGOAS0083( | RALGO03513 |
| RPSIN01497( | RMMUS1701 | RNFUR05229 | RDR007388 | RCPIC03379 | RGOAS0224( | RALGO46434 |
| RPSIN03011 | RMMUS0112 | RNFUR03363 | RDR022801 | RCPIC00839 | RGOAS03377 | RALGO19597 |
| RPSIN02836( | RMMUS0428 | RNFUR025088 | | RCPIC01781 | RGOAS01354 | RALGO19763 |
| RPSIN02118 | RMMUS1864 | RNFUR05247 | RDR025545 | RCPIC04880 | RGOAS01844 | RALGO05419 |
| 09 | RMMUS2820 | RNFUR0006( | RDR028251 | RCPIC01284 | RGOAS00783 | RALGO55454 |
| RPSIN00103 | RMMUS0281 | RNFUR00767 | RDR000039 | RCPIC01577 | RGOAS01662 | RALGO11170 |
| RPSIN02921 | RMMUS2119 | RNFUR04355 | RDR011079 | RCPIC02419 | RGOAS00899 | RALGO42779 |
| | RMMUS237786 | | | | | not predicted |
| RPSIN02823 | RMMUS2685 | RNFUR0120( | RDR014772 | RCPIC00964 | RGOAS02811 | RALGO12743 |
| RPSIN01374 | RMMUS2547 | RNFUR0210( | RDR038146 | RCPIC00626 | RGOAS00749 | RALGO18619 |
| RPSIN01628 | RMMUS0909 | RNFUR01313 | RDR046418 | RCPIC00262 | RGOAS01982 | RALGO39915 |
| RPSIN00041( | RMMUS1709 | RNFUR05076 | RDR022381 | RCPIC00303 | RGOAS02696 | RALGO16591 |
| RPSIN00041 | RMMUS1709 | RNFUR044545 | | RCPIC00304 | RGOAS02696 | RALGO16615 |
| RPSIN01144 | RMMUS232628 | | | RCPIC04646 | RGOAS0133( | RALGO54367 |
| RPSIN00899 | RMMUS0105 | RNFUR0035( | RDR041729 | RCPIC00015 | RGOAS00207 | RALGO44372 |
| RPSIN03495 | RMMUS1264 | RNFUR01484 | RDR032999 | RCPIC02628 | RGOAS00125 | RALGO35120 |
| RPSIN02077 | RMMUS104040 | | | RCPIC03291 | RGOAS00296 | RALGO52496 |
| RPSIN02033 | RMMUS2843 | RNFUR00854 | RDR028267 | | | RALGO44093 |
| RPSIN02649 | RMMUS1645 | RNFUR01948 | RDR027583 | RCPIC04158 | RGOAS01176 | RALGO22449 |
| RPSIN014808 | RMMUS2716 | RNFUR04715 | RDR024390 | RCPIC02610 | RGOAS02863 | RALGO29381 |
| RPSIN01824 | RMMUS2334 | RNFUR0405( | RDR006546 | RCPIC04670 | RGOAS01155 | RALGO53801 |
| RPSIN01828 | RMMUS1570 | RNFUR05401 | RDR026164 | RCPIC03412 | RGOAS03368 | RALGO52135 |
| RPSIN00984 | RMMUS1991 | RNFUR04779 | RDR044790 | RCPIC03061 | RGOAS02584 | RALGO56742 |
| RPSIN02037 | RMMUS2568 | RNFUR03464 | RDR017689 | RCPIC04182 | RGOAS03374 | RALGO37079 |
| RPSIN01213 | RMMUS0196 | RNFUR0226( | RDR045931 | RCPIC00313 | RGOAS01087 | RALGO34910 |
| RPSIN01202 | RMMUS0488 | RNFUR03493 | RDR036342 | RCPIC04583 | RGOAS03317 | RALGO44108 |
| RPSIN02616 | RMMUS0611 | RNFUR04072 | RDR015530 | RCPIC01125 | RGOAS02737 | RALGO55331 |
| RPSIN01685( | RMMUS2479 | RNFUR00371 | RDR039499 | RCPIC00617 | RGOAS00988 | RALGO59123 |
| RPSIN02119 | RMMUS1862 | RNFUR010263 | | RCPIC04891 | RGOAS03004 | RALGO06178 |
| 20 | RMMUS1430 | RNFUR01091 | RDR030151 | | | not predicted |
| RPSIN029728 | RMMUS2727 | RNFUR03755 | RDR024280 | RCPIC05240 | RGOAS02458 | RALGO08787 |
| RPSIN01745 | RMMUS0828 | RNFUR04801 | RDR010342 | RCPIC01660 | RGOAS01878 | RALGO01458 |
| RPSIN00080 | RMMUS1975 | RNFUR03991 | RDR025654 | RCPIC05047 | RGOAS03086 | RALGO61823 |
| RPSIN01903 | RMMUS1751 | RNFUR01709 | RDR004766 | RCPIC03711 | RGOAS00596 | RALGO33946 |

| | | | | | | |
|---|---|---|---|---|---|---|
| 18 | RMMUS0649 | RNFUR05452 | RDR055913 | RCPIC05070 | RGOAS00825 | RALGO00507 |
| 94 | RMMUS204800 | | RDR002670 | RCPIC04957 | RGOAS02008 | RALGO53965 |
| RPSIN022904 | RMMUS0132 | RNFUR04035 | RDR036439 | RCPIC05708 | RGOAS03404 | RALGO01113 |
| RPSIN036305 | RMMUS1545 | RNFUR04069 | RDR020371 | RCPIC04516 | RGOAS02582 | RALGO47791 |
| RPSIN020633 | RMMUS1886 | RNFUR00666 | RDR049261 | RCPIC04771 | RGOAS00096 | RALGO12449 |
| RPSIN031895 | RMMUS1435 | RNFUR03099 | RDR029233 | RCPIC02442 | RGOAS00050 | RALGO71116 |
| RPSIN000254 | RMMUS269833 | | | | | not predicted |
| RPSIN020314 | RMMUS2594 | RNFUR03376 | RDR024437 | RCPIC02244 | RGOAS02465 | RALGO65197 |
| RPSIN018698 | RMMUS2122 | RNFUR02839 | RDR011087 | RCPIC02400 | RGOAS01089 | RALGO40629 |
| RPSIN011074 | RMMUS2532 | RNFUR03203 | RDR029723 | RCPIC00546 | RGOAS01521 | RALGO54946 |
| RPSIN028431 | RMMUS2594 | RNFUR01342 | RDR026103 | RCPIC04390 | RGOAS01548 | RALGO65566 |
| RPSIN037575 | RMMUS1848 | RNFUR04754 | RDR029680 | RCPIC02864 | RGOAS02102 | RALGO18664 |
| RPSIN034373 | RMMUS2816 | RNFUR04454 | RDR009435 | RCPIC01731 | RGOAS01388 | RALGO50273 |
| RPSIN002463 | RMMUS2231 | RNFUR01233 | RDR038047 | RCPIC00580 | RGOAS02161 | RALGO11624 |
| RPSIN025312 | RMMUS2499 | RNFUR03866 | RDR005044 | RCPIC01970 | RGOAS03165 | RALGO05016 |
| RPSIN004196 | RMMUS003624 | | RDR019146 | RCPIC04389 | RGOAS03219 | RALGO28373 |
| | | | | | | not predicted |
| RPSIN001589 | RMMUS280706 | | | RCPIC03074 | RGOAS01388 | RALGO43437 |
| RPSIN025155 | RMMUS1510 | RNFUR02536 | RDR014445 | RCPIC02575 | RGOAS00647 | RALGO49301 |
| RPSIN022312 | RMMUS0706 | RNFUR03248 | RDR016948 | RCPIC01699 | RGOAS00192 | RALGO14342 |
| RPSIN025995 | RMMUS2044 | RNFUR04058 | RDR024325 | RCPIC00581 | RGOAS01983 | RALGO60443 |
| RPSIN031030 | RMMUS0531 | RNFUR00373 | RDR000697 | RCPIC03333 | RGOAS02795 | RALGO33301 |
| RPSIN029668 | RMMUS0690 | RNFUR01462 | RDR043420 | RCPIC01694 | RGOAS01994 | RALGO16972 |
| RPSIN025930 | RMMUS268063 | | RDR014987 | RCPIC00949 | RGOAS00326 | RALGO15842 |
| RPSIN010223 | RMMUS0592 | RNFUR04212 | RDR020597 | RCPIC01140 | RGOAS00701 | RALGO57785 |
| RPSIN009341 | RMMUS0643 | RNFUR03294 | RDR018677 | RCPIC05014 | RGOAS03044 | RALGO05350 |
| RPSIN021867 | RMMUS0834 | RNFUR05457 | RDR009464 | RCPIC01618 | RGOAS00787 | RALGO05503 |
| RPSIN018106 | RMMUS1603 | RNFUR05097 | RDR043546 | RCPIC00548 | RGOAS00235 | RALGO55646 |
| RPSIN017688 | RMMUS2301 | RNFUR01405 | RDR045282 | RCPIC03556 | RGOAS02782 | RALGO69523 |
| RPSIN038218 | RMMUS0655 | RNFUR04438 | RDR036155 | RCPIC04929 | RGOAS00060 | RALGO10552 |
| RPSIN036189 | RMMUS1932 | RNFUR02574 | RDR038392 | RCPIC00661 | RGOAS01563 | RALGO27782 |
| RPSIN013583 | RMMUS1580 | RNFUR04516 | RDR045364 | RCPIC00297 | RGOAS01082 | RALGO17296 |
| RPSIN012287 | RMMUS239139 | | RDR013698 | RCPIC03700 | RGOAS020415 | |
| RPSIN031063 | RMMUS2323 | RNFUR00316 | RDR005387 | RCPIC05166 | RGOAS01751 | RALGO54565 |
| RPSIN007141 | RMMUS2064 | RNFUR04951 | RDR003963 | RCPIC03452 | RGOAS02889 | RALGO64513 |
| RPSIN010933 | RMMUS152245 | | RDR035376 | RCPIC01969 | RGOAS03193 | RALGO44312 |
| RPSIN035090 | RMMUS266108 | | RDR040680 | RCPIC011467 | | RALGO18889 |
| RPSIN021985 | RMMUS1351 | RNFUR05491 | RDR042446 | RCPIC03809 | RGOAS02794 | RALGO41044 |
| RPSIN027513 | RMMUS1946 | RNFUR04119 | RDR038698 | RCPIC00670 | RGOAS02459 | RALGO33190 |
| RPSIN004214 | RMMUS0035 | RNFUR01826 | RDR018990 | RCPIC04395 | RGOAS02349 | RALGO28247 |
| 75 | RMMUS1712 | RNFUR04415 | RDR022571 | RCPIC04788 | RGOAS00345 | RALGO15503 |
| RPSIN027663 | RMMUS171625 | | RDR023084 | RCPIC04797 | RGOAS00839 | RALGO14762 |

| | | | | | | |
|---|---|---|---|---|---|---|
| RPSIN02775 | RMMUS1714 | RNFUR0309 | RDR043499 | RCPIC04796 | RGOAS0138 | RALGO15254 |
| 56 | RMMUS213141 | | | | | not predicte |
| RPSIN00179 | RMMUS1158 | RNFUR0110 | RDR016266 | RCPIC00816 | RGOAS0169 | RALGO4865 |
| RPSIN03159 | RMMUS0446 | RNFUR0185 | RDR021626 | RCPIC044698 | | RALGO70651 |
| RPSIN02293 | RMMUS0557 | RNFUR0227 | RDR030797 | RCPIC00477 | RGOAS0179 | RALGO6845 |
| RPSIN00144 | RMMUS1483 | RNFUR0175 | RDR040379 | RCPIC01473 | RGOAS0161 | RALGO3533 |
| RPSIN02470 | RMMUS2450 | RNFUR0121 | RDR019176 | RCPIC02373 | RGOAS002719 | |
| RPSIN03414 | RMMUS0864 | RNFUR0186 | RDR009561 | RCPIC02852 | RGOAS0185 | RALGO12152 |
| RPSIN02214 | RMMUS0129 | RNFUR0359 | RDR014536 | RCPIC01209 | RGOAS0080 | RALGO41792 |
| RPSIN03105 | RMMUS2324 | RNFUR0112 | RDR054676 | RCPIC04645 | RGOAS0005 | RALGO54502 |
| RPSIN02772 | RMMUS2376 | RNFUR016845 | | RCPIC031074 | | RALGO20267 |
| RPSIN02734 | RMMUS1561 | RNFUR015278 | | RCPIC00926 | RGOAS0291 | RALGO21141 |
| RPSIN00860 | RMMUS1925 | RNFUR0044 | RDR039467 | RCPIC00620 | RGOAS0315 | RALGO0870 |
| RPSIN02807 | RMMUS0006 | RNFUR0217 | RDR045759 | RCPIC00334 | RGOAS0035 | RALGO68004 |
| RPSIN00298 | RMMUS2181 | RNFUR0328 | RDR026177 | RCPIC02140 | RGOAS0248 | RALGO15098 |
| RPSIN03766 | RMMUS1792 | RNFUR0138 | RDR010522 | RCPIC02873 | RGOAS0339 | RALGO24073 |
| RPSIN03220 | RMMUS0857 | RNFUR0522 | RDR018192 | RCPIC01660 | RGOAS0211 | RALGO11768 |
| 56 | RMMUS213141 | | | | | not predicte |
| | | | | RCPIC05690 | RGOAS0102 | RALGO02401 |
| RPSIN02430 | RMMUS2632 | RNFUR0530 | RDR039622 | RCPIC01244 | RGOAS0241 | RALGO53234 |
| RPSIN01764 | RMMUS2305 | RNFUR0408 | RDR036591 | RCPIC03561 | RGOAS0103 | RALGO69622 |
| 93 | RMMUS2204 | RNFUR0282 | RDR014287 | RCPIC02182 | RGOAS0201 | RALGO30695 |
| 33 | RMMUS0375 | RNFUR0246 | RDR044185 | RCPIC02185 | RGOAS0308 | RALGO29750 |
| RPSIN02370 | RMMUS1386 | RNFUR0312 | RDR046974 | RCPIC01522 | RGOAS0295 | RALGO07621 |
| RPSIN02200 | RMMUS2567 | RNFUR0482 | RDR022761 | RCPIC05335 | RGOAS0309 | RALGO3295 |
| RPSIN01402 | RMMUS1057 | RNFUR0299 | RDR030744 | | RGOAS0041 | RALGO42071 |
| RPSIN02498 | RMMUS0048 | RNFUR0497 | RDR012155 | RCPIC03258 | RGOAS0252 | RALGO19621 |
| RPSIN00309 | RMMUS1520 | RNFUR020513 | | RCPIC01846 | RGOAS0127 | RALGO43654 |
| RPSIN02123 | RMMUS1494 | RNFUR0223 | RDR046751 | RCPIC05346 | RGOAS0121 | RALGO37742 |
| RPSIN00748 | RMMUS222948 | | | RCPIC005814 | | RALGO13112 |
| RPSIN02640 | RMMUS1333 | RNFUR0116 | RDR028073 | RCPIC04273 | RGOAS0342 | RALGO49484 |
| RPSIN01474 | RMMUS2670 | RNFUR0306 | RDR045547 | RCPIC03464 | RGOAS0292 | RALGO65824 |
| RPSIN03749 | RMMUS183130 | | RDR030367 | RCPIC02861 | RGOAS0299 | RALGO18372 |
| RPSIN02353 | RMMUS1728 | RNFUR0464 | RDR009895 | RCPIC02456 | RGOAS0002 | RALGO1481 |
| RPSIN01349 | RMMUS233904 | | | RCPIC032975 | | RALGO53183 |
| RPSIN01590 | RMMUS1399 | RNFUR0412 | RDR015763 | RCPIC01523 | RGOAS0099 | RALGO35075 |
| RPSIN02472 | RMMUS2666 | RNFUR0080 | RDR036159 | RCPIC03433 | RGOAS0120 | RALGO6562 |
| RPSIN02550 | RMMUS0395 | RNFUR0512 | RDR043838 | RCPIC02954 | RGOAS0193 | RALGO1803 |
| RPSIN00976 | RMMUS1397 | RNFUR0064 | RDR050938 | RCPIC01498 | RGOAS0051 | RALGO13064 |
| 99 | RMMUS221981 | | | RCPIC02253 | RGOAS0048 | RALGO3433 |
| RPSIN01344 | RMMUS1859 | RNFUR0249 | RDR024212 | RCPIC04884 | RGOAS0262 | RALGO07543 |
| RPSIN01206 | RMMUS0275 | RNFUR0501 | RDR019613 | RCPIC034981 | | RALGO32551 |
| RPSIN00999 | RMMUS0606 | RNFUR024584 | | RCPIC01111 | RGOAS0285 | RALGO56061 |
| RPSIN02862 | RMMUS2085 | RNFUR0077 | RDR020595 | RCPIC02113 | RGOAS0259 | RALGO5062 |

| | | | | | | |
|---|---|---|---|---|---|---|
| RPSIN02211 | RMMUS0529 | RNFUR002057 | | RCPIC03334 | RGOAS0002 | RALGO3244 |
| 93 | RMMUS273665 | | RDR026433 | RCPIC00920 | RGOAS0266 | RALGO15071 |
| RPSIN00988 | RMMUS1963 | RNFUR0379 | RDR033100 | RCPIC02759 | RGOAS0267 | RALGO5607 |
| RPSIN02379 | RMMUS111189 | | | RCPIC04988 | RGOAS0117 | RALGO0441 |
| 97 | RMMUS1118 | RNFUR0061 | RDR032467 | RCPIC016710 | | RALGO4587 |
| RPSIN02158 | RMMUS2550 | RNFUR0007 | RDR038037 | RCPIC00599 | RGOAS0177 | RALGO62482 |
| RPSIN00868 | RMMUS2439 | RNFUR0086 | RDR001761 | RCPIC00007 | RGOAS0020 | RALGO64552 |
| RPSIN00987 | RMMUS1995 | RNFUR0463 | RDR033081 | RCPIC03018 | RGOAS025179 | |
| RPSIN00754 | RMMUS0062 | RNFUR0130 | RDR012717 | RCPIC04537 | RGOAS007540 | |
| RPSIN00807 | RMMUS1971 | RNFUR0321 | RDR026887 | RCPIC03326 | RGOAS0088 | RALGO62981 |
| RPSIN03173 | RMMUS214693 | | RDR036263 | RCPIC00293 | RGOAS0017 | RALGO48613 |
| RPSIN02737 | RMMUS1871 | RNFUR0501 | RDR004518 | RCPIC04896 | RGOAS0304 | RALGO0275 |
| RPSIN00980 | RMMUS1964 | RNFUR0440 | RDR033079 | RCPIC044640 | | RALGO5622 |
| RPSIN00894 | RMMUS213465 | | | RCPIC042370 | | not predicte |
| RPSIN00605 | RMMUS2378 | RNFUR0117 | RDR045856 | RCPIC01984 | RGOAS0175 | RALGO32494 |
| 33 | RMMUS258171 | | | | | not predicte |
| RPSIN01425 | RMMUS138272 | | RDR034996 | RCPIC01574 | RGOAS0305 | RALGO0977 |
| | RMMUS181243 | | | | | not predicte |
| | RMMUS122465 | | | | | |
| RPSIN02828 | RMMUS1056 | RNFUR022731 | | RCPIC050815 | | RALGO32806 |
| | RMMUS122216 | | | | | |
| | RMMUS106629 | | | | | |
| | RMMUS178342 | | RDR026311 | | RGOAS014161 | |
| | RMMUS106684 | | | | | |
| | RMMUS109577 | | | | | |
| | RMMUS127068 | | | | | |
| | RMMUS109218 | | | | | |
| RPSIN00138 | RMMUS1472 | RNFUR0271 | RDR030339 | RCPIC01456 | RGOAS0292 | RALGO34282 |
| RPSIN02047 | RMMUS0250 | RNFUR032551 | | RCPIC03544 | RGOAS0146 | RALGO40894 |
| RPSIN03080 | RMMUS148242 | | RDR022562 | RCPIC01475 | RGOAS0242 | RALGO35954 |
| RPSIN02040 | RMMUS094905 | | RDR031421 | RCPIC00254 | RGOAS0074 | RALGO50171 |
| | RMMUS1377 | RNFUR0115 | RDR014733 | | | not predicte |
| RPSIN01980 | RMMUS1848 | RNFUR0429 | RDR052426 | RCPIC04050 | RGOAS0158 | RALGO34589 |
| RPSIN01176 | RMMUS282139 | | RDR011398 | RCPIC01203 | RGOAS0336 | RALGO5553 |
| RPSIN02402 | RMMUS0840 | RNFUR0242 | RDR010115 | RCPIC04568 | RGOAS0089 | RALGO2681 |
| RPSIN03769 | RMMUS183062 | | RDR029135 | RCPIC02864 | RGOAS0036 | RALGO18114 |
| RPSIN01390 | RMMUS0214 | RNFUR0080 | RDR010688 | RCPIC02051 | RGOAS0015 | RALGO21504 |
| RPSIN01561 | RMMUS049644 | | RDR017047 | RCPIC00082 | RGOAS0149 | RALGO45735 |
| RPSIN01747 | RMMUS1488 | RNFUR0327 | RDR030658 | RCPIC00023 | RGOAS0068 | RALGO70951 |
| RPSIN00740 | RMMUS1753 | RNFUR0541 | RDR051650 | RCPIC01289 | RGOAS0260 | RALGO49211 |

| | | | | | | |
|---|---|---|---|---|---|---|
| | RMMUS0577 | RNFUR0013 | RDR011104 | | RGOAS0104 | RALGO0144 |
| RPSIN03717 | RMMUS0943 | RNFUR0421 | RDR036422 | RCPIC00274 | RGOAS0093 | RALGO4536 |
| RPSIN02429 | RMMUS2634 | RNFUR0068 | RDR027536 | RCPIC01244 | RGOAS0174 | RALGO5303 |
| 5 | RMMUS050098 | | RDR017278 | RCPIC01575 | RGOAS0225 | RALGO12464 |
| RPSIN0160 | RMMUS2004 | RNFUR0318 | RDR033334 | RCPIC03009 | RGOAS0036 | RALGO5819 |
| RPSIN0012 | RMMUS1317 | RNFUR0079 | RDR021441 | RCPIC03155 | RGOAS0216 | RALGO59261 |
| RPSIN01939 | RMMUS2417 | RNFUR0504 | RDR045029 | RCPIC01924 | RGOAS0270 | RALGO42107 |
| RPSIN0077 | RMMUS1999 | RNFUR0053 | RDR034002 | | RGOAS0269 | not predicte |
| RPSIN0152 | RMMUS2089 | RNFUR0278 | RDR028846 | RCPIC04833 | RGOAS0221 | RALGO5634 |
| RPSIN0140 | RMMUS2739 | RNFUR0030 | RDR026052 | RCPIC02279 | RGOAS0320 | RALGO04137 |
| 8 | RMMUS1613 | RNFUR0543 | RDR038115 | | RGOAS0312 | not predicte |
| RPSIN0238 | RMMUS1026 | RNFUR0454 | RDR007309 | RCPIC05082 | RGOAS0171 | RALGO6051 |
| RPSIN0252 | RMMUS2498 | RNFUR0411 | RDR000765 | RCPIC01892 | RGOAS0277 | RALGO04461 |
| RPSIN0050 | RMMUS0467 | RNFUR0104 | RDR031091 | RCPIC045913 | | RALGO42842 |
| RPSIN0384 | RMMUS1178 | RNFUR0314 | RDR046643 | RCPIC02553 | RGOAS0106 | RALGO47148 |
| RPSIN0035 | RMMUS0883 | RNFUR0011 | RDR005220 | RCPIC01772 | RGOAS0040 | RALGO13343 |
| RPSIN0021 | RMMUS1209 | RNFUR0075 | RDR030051 | RCPIC03863 | RGOAS0067 | RALGO7165 |
| RPSIN0240 | RMMUS0228 | RNFUR0011 | RDR011349 | RCPIC02040 | RGOAS0152 | RALGO17604 |
| | RMMUS031355 | | | RCPIC05190 | RGOAS0021 | RALGO3498 |
| | RMMUS208258 | | | | | |
| | RMMUS083580 | | | | RGOAS0169 | RALGO21927 |
| RPSIN0161 | RMMUS228780 | | | RCPIC05042 | RGOAS0215 | RALGO68911 |
| RPSIN0125 | RMMUS1732 | RNFUR0107 | RDR018090 | RCPIC03325 | RGOAS0260 | RALGO6176 |
| RPSIN0160 | RMMUS1001 | RNFUR0538 | RDR017089 | RCPIC04814 | RGOAS0030 | RALGO3638 |
| RPSIN0209 | RMMUS2458 | RNFUR0239 | RDR019184 | RCPIC02372 | RGOAS0316 | RALGO66346 |
| RPSIN0230 | RMMUS208088 | | RDR042738 | RCPIC00473 | RGOAS0124 | RALGO2810 |
| RPSIN0024 | RMMUS0674 | RNFUR0514 | RDR036429 | RCPIC02722 | RGOAS0314 | RALGO1392 |
| RPSIN0027 | RMMUS0001 | RNFUR0136 | RDR002812 | RCPIC02663 | RGOAS0044 | RALGO67534 |
| RPSIN0064 | RMMUS1592 | RNFUR0008 | RDR026085 | RCPIC00615 | RGOAS0222 | RALGO57707 |
| | | | | | | not predicte |
| RPSIN0361 | RMMUS0739 | RNFUR0130 | RDR007480 | RCPIC03218 | RGOAS0075 | RALGO28217 |
| RPSIN0312 | RMMUS1411 | RNFUR031089 | | RCPIC01519 | RGOAS0005 | RALGO12131 |
| RPSIN0175 | RMMUS0456 | RNFUR0095 | RDR002499 | RCPIC01972 | RGOAS0017 | RALGO2387 |
| | RMMUS281830 | | | | | not predicte |
| RPSIN0322 | RMMUS0492 | RNFUR0103 | RDR031670 | RCPIC04581 | RGOAS0331 | RALGO43705 |
| RPSIN0090 | RMMUS2279 | RNFUR0138 | RDR045401 | RCPIC03320 | RGOAS0179 | RALGO60497 |
| RPSIN0130 | RMMUS2199 | RNFUR0452 | RDR033060 | RCPIC00038 | RGOAS0012 | RALGO62404 |
| RPSIN0329 | RMMUS0849 | RNFUR0287 | RDR017328 | RCPIC01591 | RGOAS0329 | RALGO10162 |
| RPSIN033845 | | RNFUR0323 | RDR020537 | RCPIC05147 | RGOAS0130 | RALGO6835 |

| | | | | | | |
|---|---|---|---|---|---|---|
| RPSIN01509 | RMMUS1479 | RNFUR0353 | RDR009657 | RCPIC01446 | RGOAS0232 | RALGO36887 |
| RPSIN01138 | RMMUS1241 | RNFUR0015 | RDR048634 | RCPIC01821 | RGOAS0028 | RALGO01734 |
| RPSIN01449 | RMMUS1376 | RNFUR0093 | RDR014703 | RCPIC01501 | RGOAS0039 | RALGO18505 |
| RPSIN00648 | RMMUS2476 | RNFUR0328 | RDR025696 | RCPIC02713 | RGOAS0332 | RALGO58382 |
| RPSIN01035 | RMMUS0164 | RNFUR0098 | RDR037966 | RCPIC00571 | RGOAS0120 | RALGO63664 |
| 46 | RMMUS2201 | RNFUR0489 | RDR002671 | RCPIC05706 | RGOAS0029 | RALGO41882 |
| 72 | RMMUS1553 | RNFUR0424 | RDR012614 | RCPIC03737 | RGOAS0287 | RALGO23989 |
| RPSIN02321 | RMMUS2341 | RNFUR0468 | RDR046554 | RCPIC000503 | | not predicted |
| RPSIN00586 | RMMUS1723 | RNFUR0467 | RDR013373 | RCPIC04162 | RGOAS0035 | RALGO24413 |
| RPSIN01092 | RMMUS1694 | RNFUR0385 | RDR006960 | RCPIC000697 | | RALGO48881 |
| RPSIN03243 | RMMUS006767 | | RDR003886 | RCPIC017365 | | |
| | RMMUS091457 | | | RCPIC035517 | | not predicted |
| RPSIN01610 | RMMUS0579 | RNFUR0168 | RDR001127 | RCPIC01191 | RGOAS0010 | not predicted |
| | RMMUS0692 | RNFUR0445 | RDR023323 | | RGOAS032066 | |
| | RMMUS0202 | RNFUR0418 | RDR010330 | RCPIC03209 | RGOAS029994 | |
| RPSIN02534 | RMMUS0746 | RNFUR0221 | RDR002293 | RCPIC00300 | RGOAS0296 | RALGO23064 |
| RPSIN02312 | RMMUS2178 | RNFUR0447 | RDR023168 | RCPIC03744 | RGOAS0179 | RALGO26930 |
| RPSIN03849 | RMMUS0147 | RNFUR0179 | RDR012313 | RCPIC00103 | RGOAS0293 | RALGO44657 |
| RPSIN02007 | RMMUS0399 | RNFUR0173 | RDR012718 | RCPIC02969 | RGOAS0013 | RALGO14951 |
| 96 | RMMUS221751 | | | RCPIC049941 | | not predicted |
| RPSIN02417 | RMMUS132482 | | RDR002119 | RCPIC04289 | RGOAS0329 | RALGO51529 |
| RPSIN00643 | RMMUS1593 | RNFUR0120 | RDR038530 | RCPIC00555 | RGOAS0259 | RALGO57260 |
| RPSIN00083 | RMMUS0344 | RNFUR0312 | RDR020970 | RCPIC03878 | RGOAS012032 | |
| RPSIN01564 | RMMUS1425 | RNFUR0009 | RDR027382 | RCPIC00702 | RGOAS0291 | RALGO60128 |
| RPSIN01745 | RMMUS2466 | RNFUR0355 | RDR030253 | RCPIC03168 | RGOAS0289 | not predicted |
| RPSIN01113 | RMMUS2114 | RNFUR0174 | RDR040846 | RCPIC05221 | RGOAS0108 | RALGO44405 |
| RPSIN00296 | RMMUS009188 | | | RCPIC04754 | RGOAS0305 | RALGO07423 |
| | RMMUS271431 | | | | | |
| RPSIN01965 | RMMUS0623 | RNFUR0152 | RDR013950 | RCPIC01238 | RGOAS0279 | RALGO21732 |
| RPSIN00333 | RMMUS2138 | RNFUR0413 | RDR018509 | RCPIC027858 | | RALGO33469 |
| RPSIN02848 | RMMUS0496 | RNFUR0491 | RDR039001 | RCPIC00081 | RGOAS0009 | RALGO45648 |
| RPSIN00360 | RMMUS0881 | RNFUR0003 | RDR041972 | RCPIC01775 | RGOAS0285 | RALGO15452 |
| RPSIN00180 | RMMUS233248 | | RDR055585 | RCPIC04649 | RGOAS0050 | RALGO54058 |
| RPSIN00438 | RMMUS0787 | RNFUR0257 | RDR038275 | RCPIC04254 | RGOAS0293 | RALGO52895 |
| RPSIN03707 | RMMUS2646 | RNFUR0284 | RDR027704 | RCPIC01335 | RGOAS0036 | RALGO55223 |
| RPSIN01734 | RMMUS2495 | RNFUR0448 | RDR045355 | RCPIC01762 | RGOAS0278 | not predicted |
| RPSIN00514 | RMMUS1741 | RNFUR0523 | RDR032872 | RCPIC02699 | RGOAS0013 | RALGO65974 |
| 24 | RMMUS1801 | RNFUR0021 | RDR014945 | RCPIC05674 | RGOAS0150 | RALGO27695 |
| RPSIN02106 | RMMUS0086 | RNFUR0130 | RDR002810 | RCPIC01735 | RGOAS0087 | RALGO36542 |
| RPSIN02274 | RMMUS1021 | RNFUR0028 | RDR031777 | RCPIC05516 | RGOAS0270 | RALGO63072 |
| RPSIN00348 | RMMUS2521 | RNFUR0313 | RDR012242 | RCPIC04506 | RGOAS0016 | RALGO22035 |
| RPSIN02912 | RMMUS1348 | RNFUR0464 | RDR002362 | RCPIC03776 | RGOAS0227 | RALGO40807 |
| RPSIN01342 | RMMUS1856 | RNFUR0060 | RDR005143 | RCPIC04886 | RGOAS0280 | RALGO08916 |
| RPSIN03314 | RMMUS2785 | RNFUR0022 | RDR027838 | RCPIC01286 | RGOAS0029 | RALGO54646 |
| RPSIN00238 | RMMUS0781 | RNFUR0052 | RDR027508 | RCPIC04263 | RGOAS0302 | RALGO54418 |

| | | | | | | |
|---|---|---|---|---|---|---|
| RPSIN032329 | RMMUS0209 | RNFUR00916 | RDR039643 | RCPIC04633 | RGOAS00114 | RALGO03657 |
| 75 | RMMUS224024 | | | RCPIC03873 | RGOAS03044 | RALGO63397 |
| RPSIN010723 | RMMUS1193 | RNFUR00600 | RDR021895 | RCPIC02431 | RGOAS02560 | RALGO51613 |
| RPSIN024692 | RMMUS0140 | RNFUR04625 | RDR038427 | RCPIC00061 | RGOAS03040 | RALGO45312 |
| RPSIN035763 | RMMUS010807 | | RDR003263 | RCPIC00860 | RGOAS01614 | RALGO17451 |
| RPSIN008008 | RMMUS272625 | | RDR026717 | RCPIC00971 | RGOAS02978 | RALGO14405 |
| RPSIN014587 | RMMUS1042 | RNFUR03905 | RDR035538 | RCPIC00164 | RGOAS00415 | RALGO58046 |
| RPSIN037353 | RMMUS0430 | RNFUR04162 | RDR023305 | RCPIC01750 | RGOAS01108 | RALGO14378 |
| RPSIN011707 | RMMUS2132 | RNFUR01278 | RDR009993 | RCPIC04238 | RGOAS03327 | RALGO37007 |
| RPSIN003229 | RMMUS1799 | RNFUR02889 | RDR013295 | RCPIC04051 | RGOAS01367 | RALGO27941 |
| RPSIN033711 | RMMUS2514 | RNFUR02419 | RDR016955 | | RGOAS00073 | RALGO24130 |
| RPSIN003754 | RMMUS2346 | RNFUR00420 | RDR046484 | RCPIC00263 | RGOAS01916 | RALGO34114 |
| RPSIN026230 | RMMUS1813 | RNFUR00542 | RDR029666 | RCPIC02912 | RGOAS00556 | RALGO16085 |
| 17 | RMMUS1365 | RNFUR01560 | RDR000950 | RCPIC057237 | | RALGO04659 |
| RPSIN035830 | RMMUS0534 | RNFUR02643 | RDR012448 | RCPIC03937 | RGOAS01650 | RALGO30149 |
| | RMMUS022288 | | | | | not predicted |
| RPSIN015980 | RMMUS1460 | RNFUR01685 | RDR020979 | RCPIC01445 | RGOAS01457 | RALGO28061 |
| RPSIN025034 | RMMUS0837 | RNFUR03982 | RDR017035 | RCPIC04166 | RGOAS02037 | not predicted |
| RPSIN021079 | RMMUS0549 | RNFUR021590 | | RCPIC02667 | RGOAS02118 | RALGO67600 |
| RPSIN004114 | RMMUS1308 | RNFUR03513 | RDR017520 | RCPIC05105 | RGOAS02458 | RALGO64027 |
| RPSIN021368 | RMMUS216733 | | RDR025112 | RCPIC02463 | RGOAS01750 | RALGO01602 |
| RPSIN030369 | RMMUS2560 | RNFUR01162 | RDR031424 | RCPIC00291 | RGOAS01069 | RALGO47073 |
| RPSIN001669 | RMMUS1259 | RNFUR01782 | RDR024570 | RCPIC02284 | RGOAS02769 | RALGO38348 |
| RPSIN006397 | RMMUS2477 | RNFUR00367 | RDR038373 | RCPIC040583 | | RALGO58286 |
| RPSIN013311 | RMMUS0921 | RNFUR03492 | RDR006951 | RCPIC04573 | RGOAS00567 | RALGO06886 |
| RPSIN014309 | RMMUS149841 | | | RCPIC01401 | RGOAS01142 | RALGO29495 |
| RPSIN004197 | RMMUS0036 | RNFUR03034 | RDR019251 | RCPIC04389 | RGOAS03220 | RALGO28406 |
| 37 | RMMUS0889 | RNFUR02993 | RDR052242 | RCPIC03375 | RGOAS02037 | RALGO17344 |
| 99 | RMMUS1144 | RNFUR02028 | RDR029064 | RCPIC05503 | RGOAS004301 | |
| RPSIN003155 | RMMUS2813 | RNFUR03981 | RDR045246 | RCPIC03144 | RGOAS00874 | RALGO67621 |
| RPSIN034878 | RMMUS2683 | RNFUR00882 | RDR017333 | RCPIC00960 | RGOAS01758 | RALGO13865 |
| RPSIN004092 | RMMUS1307 | RNFUR03287 | RDR016775 | RCPIC05105 | RGOAS01148 | RALGO64042 |
| 17 | RMMUS1142 | RNFUR00448 | RDR004674 | RCPIC05717 | RGOAS00406 | RALGO22701 |
| RPSIN020842 | RMMUS1731 | RNFUR03877 | RDR017447 | RCPIC03196 | RGOAS01709 | RALGO62197 |
| RPSIN023005 | RMMUS1203 | RNFUR04323 | RDR039778 | RCPIC05263 | RGOAS00258 | RALGO71566 |
| RPSIN004760 | RMMUS2676 | RNFUR03540 | RDR000425 | RCPIC02919 | RGOAS01948 | RALGO29138 |
| RPSIN015969 | RMMUS082673 | | RDR009882 | RCPIC05368 | RGOAS02728 | RALGO28853 |
| RPSIN013809 | RMMUS1969 | RNFUR05208 | RDR026095 | RCPIC02915 | RGOAS02857 | RALGO61398 |
| RPSIN028527 | RMMUS0597 | RNFUR00653 | RDR010216 | RCPIC04634 | RGOAS00581 | RALGO04308 |
| RPSIN018132 | RMMUS0969 | RNFUR04282 | RDR022701 | RCPIC04183 | RGOAS02430 | RALGO34559 |
| RPSIN038418 | RMMUS1180 | RNFUR03008 | RDR046842 | RCPIC02555 | RGOAS03261 | RALGO46863 |
| RPSIN000600 | RMMUS1889 | RNFUR04897 | RDR012764 | RCPIC04098 | RGOAS02681 | RALGO35483 |
| RPSIN027868 | RMMUS2552 | RNFUR03541 | RDR025754 | RCPIC00641 | RGOAS01284 | RALGO62356 |
| RPSIN035306 | RMMUS2658 | RNFUR01894 | RDR011225 | RCPIC01135 | RGOAS00098 | RALGO21546 |
| RPSIN018606 | RMMUS2126 | RNFUR03937 | RDR012041 | RCPIC02414 | RGOAS00290 | RALGO38879 |

| | | | | | | |
|---|---|---|---|---|---|---|
| RPSIN02840 | RMMUS2734 | RNFUR0209 | RDR000707 | RCPIC05051 | RGOAS0034 | RALGO06931 |
| | RMMUS1731 | RNFUR0237 | RDR021019 | RCPIC03202 | RGOAS0165 | RALGO61949 |
| RPSIN03230 | RMMUS0757 | RNFUR0104 | RDR050744 | RCPIC023526 | | RALGO5865 |
| RPSIN02994 | RMMUS242634 | | RDR040706 | RCPIC00101 | RGOAS0172 | RALGO6361 |
| RPSIN00624 | RMMUS1275 | RNFUR0402 | RDR014623 | RCPIC03824 | RGOAS0226 | RALGO35702 |
| RPSIN03669 | RMMUS1673 | RNFUR0486 | RDR035743 | RCPIC04643 | RGOAS0184 | RALGO5523 |
| RPSIN03346 | RMMUS2240 | RNFUR0487 | RDR048219 | RCPIC03873 | RGOAS0304 | RALGO63388 |
| RPSIN03240 | RMMUS199591 | | RDR032841 | RCPIC03018 | RGOAS0146 | RALGO56976 |
| | RMMUS090474 | | | | | not predicte |
| RPSIN01632 | RMMUS0913 | RNFUR0116 | RDR047517 | RCPIC00265 | RGOAS0101 | RALGO36842 |
| RPSIN00084 | RMMUS2446 | RNFUR0223 | RDR056067 | RCPIC00946 | RGOAS0041 | RALGO65416 |
| RPSIN00139 | RMMUS1487 | RNFUR0155 | RDR030287 | RCPIC01444 | RGOAS0293 | RALGO34357 |
| 17 | RMMUS1101 | RNFUR0057 | RDR037619 | RCPIC05208 | RGOAS0005 | RALGO48406 |
| RPSIN01791 | RMMUS253664 | | | RCPIC055836 | | RALGO0338 |
| RPSIN00971 | RMMUS1005 | RNFUR0099 | RDR022720 | RCPIC00968 | RGOAS0113 | RALGO07804 |
| RPSIN03694 | RMMUS0772 | RNFUR0505 | RDR015103 | RCPIC02278 | RGOAS0084 | RALGO56541 |
| RPSIN02318 | RMMUS1052 | RNFUR0486 | RDR035645 | RCPIC00047 | RGOAS0009 | RALGO58994 |
| RPSIN03762 | RMMUS1841 | RNFUR0140 | RDR009463 | RCPIC02845 | RGOAS0041 | not predicte |
| RPSIN02767 | RMMUS2377 | RNFUR016152 | | RCPIC03107 | RGOAS0045 | RALGO20621 |
| RPSIN01387 | RMMUS0217 | RNFUR0356 | RDR040516 | RCPIC02053 | RGOAS0283 | RALGO2115 |
| 11 | RMMUS248722 | | | RCPIC05615 | RGOAS0076 | RALGO47401 |
| RPSIN00622 | RMMUS1486 | RNFUR0418 | RDR028928 | | | |
| RPSIN02430 | RMMUS263561 | | RDR039740 | RCPIC01326 | RGOAS0203 | RALGO52517 |
| RPSIN02386 | RMMUS0764 | RNFUR006037 | | RCPIC022748 | | RALGO57851 |
| RPSIN00979 | RMMUS1963 | RNFUR0328 | RDR033118 | RCPIC02756 | RGOAS0186 | RALGO56034 |
| 4 | RMMUS216515 | | | RCPIC00923 | RGOAS0146 | RALGO1744 |
| RPSIN00545 | RMMUS0246 | RNFUR0038 | RDR010222 | RCPIC02019 | RGOAS0124 | RALGO25912 |
| RPSIN00498 | RMMUS0467 | RNFUR0479 | RDR036302 | RCPIC04597 | RGOAS0303 | RALGO4289 |
| RPSIN01080 | RMMUS1621 | RNFUR0431 | RDR031219 | RCPIC02807 | RGOAS0054 | RALGO5043 |
| RPSIN01922 | RMMUS0718 | RNFUR0522 | RDR017511 | RCPIC01974 | RGOAS0114 | RALGO0558 |
| RPSIN02908 | RMMUS0391 | RNFUR0124 | RDR018712 | RCPIC03598 | RGOAS0158 | RALGO2003 |
| RPSIN03181 | RMMUS1631 | RNFUR0010 | RDR004956 | RCPIC04839 | RGOAS0208 | RALGO04191 |
| RPSIN00094 | RMMUS0126 | RNFUR0231 | RDR038243 | RCPIC01299 | RGOAS0315 | RALGO4111 |
| RPSIN00059 | RMMUS1892 | RNFUR0095 | RDR005989 | RCPIC04061 | RGOAS0078 | RALGO36044 |
| RPSIN03109 | RMMUS1241 | RNFUR0021 | RDR005129 | RCPIC017636 | | RALGO02518 |
| RPSIN02304 | RMMUS2177 | RNFUR0278 | RDR012713 | RCPIC037409 | | RALGO27761 |
| RPSIN02029 | RMMUS2592 | RNFUR0303 | RDR026274 | RCPIC022395 | | RALGO65068 |
| 47 | RMMUS105701 | | | RCPIC045719 | | RALGO2097 |
| RPSIN00080 | RMMUS1976 | RNFUR0373 | RDR025858 | RCPIC04123 | RGOAS0215 | RALGO61762 |
| RPSIN02694 | RMMUS0191 | RNFUR0110 | RDR007168 | RCPIC03211 | RGOAS0087 | RALGO09561 |
| RPSIN01180 | RMMUS0720 | RNFUR0178 | RDR016879 | RCPIC01693 | RGOAS0217 | RALGO07978 |

| | | | | | | |
|---|---|---|---|---|---|---|
| | RMMUS1083 | RNFUR02734 | RDR030263 | RCPIC05413 | RGOAS01361 | RALGO06991 |
| RPSIN002690 | RMMUS239225 | | RDR015670 | RCPIC02659 | RGOAS03056 | RALGO67417 |
| RPSIN012214 | RMMUS0276 | RNFUR04417 | RDR018930 | RCPIC05695 | RGOAS01307 | RALGO32026 |
| RPSIN000823 | RMMUS034386 | | | RCPIC03877 | RGOAS01822 | not predicted |
| RPSIN027461 | RMMUS2620 | RNFUR04192 | RDR002864 | RCPIC01754 | RGOAS00147 | RALGO39132 |
| RPSIN035407 | RMMUS0770 | RNFUR00438 | RDR028341 | RCPIC02270 | RGOAS02005 | RALGO57160 |
| 55 | RMMUS1767 | RNFUR05453 | RDR025960 | | | not predicted |
| RPSIN008711 | RMMUS2445 | RNFUR04389 | RDR000197 | RCPIC00010 | RGOAS01273 | RALGO65035 |
| RPSIN000200 | RMMUS2697 | RNFUR04965 | RDR027891 | RCPIC02687 | RGOAS01669 | RALGO31077 |
| RPSIN036003 | RMMUS2843 | RNFUR00480 | RDR027594 | RCPIC03681 | RGOAS02849 | RALGO45693 |
| RPSIN023263 | RMMUS234264 | | | RCPIC00053 | RGOAS02825 | RALGO59390 |
| RPSIN018910 | RMMUS030454 | | RDR047775 | RCPIC00812 | RGOAS00647 | RALGO48493 |
| RPSIN016230 | RMMUS2704 | RNFUR00833 | RDR009783 | RCPIC01131 | RGOAS02330 | RALGO54538 |
| RPSIN024395 | RMMUS1988 | RNFUR01356 | RDR033539 | RCPIC00794 | RGOAS03240 | RALGO51862 |
| RPSIN009985 | RMMUS0608 | RNFUR01094 | RDR000881 | RCPIC01110 | RGOAS01938 | RALGO55884 |
| RPSIN016056 | RMMUS2004 | RNFUR03748 | RDR033114 | RCPIC03008 | RGOAS00362 | RALGO58193 |
| RPSIN036114 | RMMUS0739 | RNFUR00258 | RDR007444 | RCPIC03224 | RGOAS01627 | RALGO28022 |
| RPSIN004322 | RMMUS0294 | RNFUR01256 | RDR015319 | RCPIC00811 | RGOAS00678 | RALGO50357 |
| RPSIN011784 | RMMUS1109 | RNFUR00880 | RDR031228 | RCPIC044063 | | RALGO45777 |
| RPSIN032399 | RMMUS1888 | RNFUR04045 | RDR023960 | RCPIC03061 | RGOAS01196 | RALGO14525 |
| RPSIN008086 | RMMUS2194 | RNFUR00860 | RDR037751 | RCPIC04125 | RGOAS00451 | RALGO62783 |
| RPSIN034350 | RMMUS2844 | RNFUR03154 | RDR027900 | RCPIC01279 | RGOAS00251 | RALGO51547 |
| RPSIN002242 | RMMUS1833 | RNFUR02812 | RDR004896 | RCPIC04965 | RGOAS02477 | RALGO16996 |
| RPSIN000573 | RMMUS1891 | RNFUR01149 | RDR012656 | RCPIC04024 | RGOAS02914 | RALGO35765 |
| RPSIN002101 | RMMUS1270 | RNFUR02195 | RDR046773 | RCPIC04223 | RGOAS00537 | RALGO32452 |
| RPSIN037321 | RMMUS2404 | RNFUR02907 | RDR048010 | RCPIC04727 | RGOAS02958 | RALGO11017 |
| RPSIN005015 | RMMUS0486 | RNFUR02187 | RDR035955 | RCPIC04607 | RGOAS00641 | RALGO44450 |
| RPSIN013930 | RMMUS0237 | RNFUR00885 | RDR039702 | RCPIC02034 | RGOAS00576 | RALGO22842 |
| RPSIN032503 | RMMUS0675 | RNFUR01099 | RDR033238 | RCPIC03733 | RGOAS00460 | RALGO23401 |
| RPSIN018626 | RMMUS2130 | RNFUR01100 | RDR049913 | RCPIC02401 | RGOAS02853 | RALGO40212 |
| RPSIN025383 | RMMUS2102 | RNFUR04721 | RDR017203 | RCPIC02114 | RGOAS01126 | RALGO51799 |
| RPSIN014836 | RMMUS2716 | RNFUR04841 | RDR033650 | RCPIC02645 | RGOAS02864 | RALGO29468 |
| RPSIN038665 | RMMUS2269 | RNFUR02344 | RDR016893 | RCPIC04774 | RGOAS00962 | RALGO28877 |
| 55 | RMMUS0240 | RNFUR00752 | RDR010128 | RCPIC02049 | RGOAS02336 | RALGO24067 |
| RPSIN000942 | RMMUS0126 | RNFUR01712 | RDR037842 | RCPIC05617 | RGOAS01154 | RALGO41026 |
| 09 | RMMUS1230 | RNFUR04481 | RDR014967 | RCPIC03383 | RGOAS03245 | RALGO43263 |
| RPSIN017634 | RMMUS2308 | RNFUR04464 | RDR031574 | RCPIC03562 | RGOAS00181 | RALGO69697 |
| RPSIN008634 | RMMUS2233 | RNFUR03632 | RDR033346 | RCPIC00637 | RGOAS02450 | RALGO07681 |
| RPSIN025482 | RMMUS0394 | RNFUR033513 | | RCPIC02955 | RGOAS03275 | RALGO18676 |
| RPSIN015237 | RMMUS0844 | RNFUR01324 | RDR021003 | RCPIC01586 | RGOAS00752 | RALGO07246 |
| RPSIN009549 | RMMUS2654 | RNFUR02789 | RDR036561 | RCPIC00446 | RGOAS02043 | RALGO66133 |
| RPSIN018464 | RMMUS1022 | RNFUR00056 | RDR031235 | RCPIC02222 | RGOAS00638 | RALGO63108 |
| RPSIN014945 | RMMUS1698 | RNFUR03714 | RDR007299 | RCPIC03369 | RGOAS00075 | RALGO46857 |
| RPSIN004560 | RMMUS0122 | RNFUR03736 | RDR043080 | RCPIC03962 | RGOAS01651 | RALGO37163 |
| RPSIN030570 | RMMUS029224 | | RDR014613 | RCPIC03106 | RGOAS00044 | RALGO38519 |

| | | | | | | |
|---|---|---|---|---|---|---|
| RPSIN038630 | RMMUS1610 | RNFUR00906 | RDR039389 | RCPIC00544 | RGOAS01374 | RALGO54448 |
| RPSIN017542 | RMMUS0454 | RNFUR02879 | RDR004416 | RCPIC01757 | RGOAS02167 | RALGO26329 |
| RPSIN032085 | RMMUS0670 | RNFUR04710 | RDR036377 | RCPIC05119 | RGOAS00148 | RALGO17191 |
| RPSIN036173 | RMMUS1929 | RNFUR034791 | | RCPIC00624 | RGOAS02669 | RALGO24525 |
| RPSIN009214 | RMMUS1471 | RNFUR01041 | RDR034734 | RCPIC01456 | RGOAS03353 | RALGO33913 |
| RPSIN035631 | RMMUS1758 | RNFUR02703 | RDR041511 | RCPIC01226 | RGOAS00034 | RALGO51472 |
| RPSIN014323 | RMMUS1254 | RNFUR00906 | RDR031177 | RCPIC05176 | RGOAS01575 | RALGO06556 |
| RPSIN022869 | RMMUS0084 | RNFUR031968 | | RCPIC02096 | RGOAS01712 | RALGO12716 |
| RPSIN003454 | RMMUS1521 | RNFUR02004 | RDR034499 | RCPIC017777 | | RALGO43789 |
| RPSIN034169 | RMMUS1811 | RNFUR01020 | RDR010800 | RCPIC02830 | RGOAS01682 | RALGO10288 |
| RPSIN003912 | RMMUS0862 | RNFUR02676 | RDR021402 | RCPIC02886 | RGOAS03065 | RALGO15302 |
| 95 | RMMUS0888 | RNFUR05032 | RDR004675 | RCPIC018135 | | RALGO17568 |
| RPSIN011438 | RMMUS2325 | RNFUR00039 | RDR004484 | RCPIC04646 | RGOAS02220 | RALGO54433 |
| RPSIN008321 | RMMUS2273 | RNFUR00217 | RDR029361 | RCPIC05109 | RGOAS01470 | RALGO27137 |
| RPSIN018638 | RMMUS213186 | | RDR009399 | RCPIC02400 | RGOAS03302 | RALGO40485 |
| RPSIN027134 | RMMUS2294 | RNFUR04070 | RDR036153 | RCPIC04756 | RGOAS01800 | RALGO69226 |
| RPSIN002295 | RMMUS1371 | RNFUR03281 | RDR000051 | RCPIC04852 | RGOAS00093 | RALGO08814 |
| RPSIN001238 | RMMUS0590 | RNFUR00170 | RDR040114 | RCPIC03582 | RGOAS01291 | RALGO59873 |
| RPSIN024272 | RMMUS1772 | RNFUR01366 | RDR019588 | RCPIC01280 | RGOAS03152 | RALGO53432 |
| RPSIN024501 | RMMUS0319 | RNFUR03855 | RDR038425 | RCPIC00799 | RGOAS01634 | RALGO53066 |
| RPSIN015603 | RMMUS198269 | | RDR038430 | RCPIC00857 | RGOAS02779 | RALGO51013 |
| RPSIN037529 | RMMUS1844 | RNFUR04779 | RDR020616 | RCPIC02864 | RGOAS02120 | RALGO22614 |
| RPSIN012324 | RMMUS1836 | RNFUR04845 | RDR010264 | RCPIC02852 | RGOAS00396 | RALGO07237 |
| RPSIN033407 | RMMUS1013 | RNFUR04353 | RDR046962 | RCPIC00018 | RGOAS00434 | RALGO57190 |
| RPSIN001115 | RMMUS0554 | RNFUR02807 | RDR032646 | RCPIC05031 | RGOAS02397 | RALGO68560 |
| RPSIN001731 | RMMUS0641 | RNFUR03482 | RDR012640 | RCPIC03655 | RGOAS01768 | RALGO07555 |
| RPSIN003728 | RMMUS2676 | RNFUR00726 | RDR052875 | RCPIC02920 | RGOAS01203 | RALGO29267 |
| RPSIN020083 | RMMUS0399 | RNFUR01719 | RDR012635 | RCPIC02969 | RGOAS00137 | RALGO15221 |
| RPSIN009738 | RMMUS1005 | RNFUR05082 | RDR022619 | RCPIC01006 | RGOAS02941 | RALGO08023 |
| RPSIN018874 | RMMUS0304 | RNFUR03704 | RDR046701 | RCPIC00815 | RGOAS00876 | RALGO48520 |
| RPSIN035157 | RMMUS0218 | RNFUR02941 | RDR017094 | RCPIC02023 | RGOAS02759 | RALGO18841 |
| RPSIN007441 | RMMUS175329 | | | RCPIC01193 | RGOAS02124 | RALGO49253 |
| RPSIN032195 | RMMUS2357 | RNFUR02343 | RDR044333 | | RGOAS00253 | RALGO20997 |
| 63 | RMMUS1867 | RNFUR02382 | RDR024302 | | RGOAS02505 | RALGO04059 |
| RPSIN029651 | RMMUS0317 | RNFUR035492 | | RCPIC02765 | RGOAS01234 | RALGO54019 |
| RPSIN016907 | RMMUS2243 | RNFUR02341 | RDR039815 | RCPIC04472 | RGOAS01511 | RALGO70429 |
| RPSIN038113 | RMMUS2605 | RNFUR05049 | RDR045060 | RCPIC02175 | RGOAS01121 | RALGO66982 |
| RPSIN029272 | RMMUS0507 | RNFUR04875 | RDR018556 | RCPIC00832 | RGOAS01515 | RALGO25074 |
| RPSIN036953 | RMMUS0774 | RNFUR00183 | RDR014481 | RCPIC02288 | RGOAS02420 | RALGO56292 |
| RPSIN020200 | RMMUS1319 | RNFUR000607 | | RCPIC025342 | | RALGO06079 |
| RPSIN038482 | RMMUS0138 | RNFUR04179 | RDR002547 | RCPIC01249 | RGOAS02323 | RALGO44186 |
| RPSIN015476 | RMMUS0242 | RNFUR05381 | RDR021785 | RCPIC02088 | RGOAS02335 | RALGO24115 |
| RPSIN014704 | RMMUS1876 | RNFUR03458 | RDR004573 | RCPIC04973 | RGOAS00032 | RALGO17601 |
| RPSIN014554 | RMMUS1045 | RNFUR01997 | RDR007109 | RCPIC00044 | RGOAS03364 | RALGO58370 |

| | | | | | | |
|---|---|---|---|---|---|---|
| RPSIN027619 | RMMUS2320 | RNFUR04673 | RDR036623 | RCPIC03627 | RGOAS02852 | RALGO69928 |
| RPSIN034669 | RMMUS1903 | RNFUR02015 | RDR005617 | RCPIC04054 | RGOAS00790 | RALGO31473 |
| RPSIN015749 | RMMUS1745 | RNFUR00188 | RDR025605 | RCPIC02702 | RGOAS02016 | RALGO65722 |
| RPSIN025790 | RMMUS2681 | RNFUR00668 | RDR014201 | RCPIC00950 | RGOAS00958 | RALGO15746 |
| RPSIN021509 | RMMUS1090 | RNFUR02150 | RDR030283 | RCPIC04874 | RGOAS00695 | RALGO09819 |
| RPSIN027253 | RMMUS1205 | RNFUR00470 | RDR034906 | RCPIC00075 | RGOAS00857 | RALGO71482 |
| RPSIN005578 | RMMUS1009 | RNFUR05433 | RDR026101 | RCPIC02464 | RGOAS00379 | RALGO10828 |
| RPSIN004505 | RMMUS0025 | RNFUR04335 | RDR056080 | RCPIC04011 | RGOAS02906 | RALGO68671 |
| RPSIN017288 | RMMUS161569 | | RDR038466 | RCPIC02786 | RGOAS02723 | RALGO51442 |
| RPSIN025628 | RMMUS0778 | RNFUR01743 | RDR000243 | RCPIC046296 | | RALGO54799 |
| RPSIN004787 | RMMUS1367 | RNFUR01976 | RDR005440 | RCPIC04854 | RGOAS00491 | RALGO10225 |
| RPSIN018812 | RMMUS224902 | | RDR018722 | RCPIC04462 | RGOAS00010 | RALGO70306 |
| RPSIN025104 | RMMUS0125 | RNFUR02134 | RDR056804 | RCPIC03943 | RGOAS00922 | RALGO39471 |
| RPSIN009748 | RMMUS1559 | RNFUR01458 | RDR043734 | RCPIC02396 | RGOAS02797 | RALGO23431 |
| RPSIN025561 | RMMUS0779 | RNFUR02355 | RDR048819 | RCPIC04623 | RGOAS01126 | RALGO54673 |
| RPSIN038661 | RMMUS2270 | RNFUR03797 | RDR040696 | RCPIC04775 | RGOAS03020 | RALGO28502 |
| RPSIN020496 | RMMUS025085 | | RDR019459 | RCPIC03505 | RGOAS01589 | RALGO40873 |
| RPSIN027165 | RMMUS0860 | RNFUR05308 | RDR018082 | RCPIC01622 | RGOAS00356 | RALGO14474 |
| RPSIN037788 | RMMUS2066 | RNFUR04957 | RDR002414 | RCPIC03444 | RGOAS00788 | RALGO64630 |
| RPSIN005464 | RMMUS0557 | RNFUR03177 | RDR002819 | RCPIC00459 | RGOAS01787 | RALGO68442 |
| 75 | RMMUS2386 | RNFUR00015 | RDR006876 | RCPIC03698 | RGOAS03369 | RALGO24307 |
| | RMMUS220259 | | | | | RALGO42359 |
| RPSIN019811 | RMMUS183262 | | | RCPIC04021 | RGOAS00714 | not predicted |
| RPSIN037090 | RMMUS1758 | RNFUR01902 | RDR028077 | RCPIC04984 | RGOAS00226 | RALGO56112 |
| RPSIN019399 | RMMUS2417 | RNFUR00737 | RDR029330 | RCPIC01786 | RGOAS02704 | RALGO42164 |
| RPSIN011992 | RMMUS1914 | RNFUR04433 | RDR006692 | RCPIC04091 | RGOAS02878 | RALGO38720 |
| RPSIN032277 | RMMUS0758 | RNFUR00534 | RDR028683 | RCPIC05127 | RGOAS00796 | RALGO58568 |
| RPSIN021872 | RMMUS1845 | RNFUR02707 | RDR011684 | RCPIC05690 | RGOAS01015 | RALGO02887 |
| 00 | RMMUS2515 | RNFUR00063 | RDR038388 | RCPIC00599 | RGOAS030335 | |
| RPSIN035242 | RMMUS1515 | RNFUR02853 | RDR046573 | RCPIC01960 | RGOAS02278 | RALGO41104 |
| RPSIN017045 | RMMUS162456 | | RDR034367 | RCPIC00715 | RGOAS01097 | RALGO52339 |
| RPSIN027822 | RMMUS1820 | RNFUR00495 | RDR029326 | RCPIC02840 | RGOAS01707 | RALGO26542 |
| RPSIN032497 | RMMUS0676 | RNFUR00233 | RDR031187 | RCPIC03724 | RGOAS00650 | RALGO23614 |
| RPSIN005762 | RMMUS0906 | RNFUR01947 | RDR036504 | RCPIC00308 | RGOAS01510 | RALGO41245 |
| RPSIN016065 | RMMUS1968 | RNFUR01821 | RDR043659 | RCPIC03651 | RGOAS00625 | RALGO25086 |
| RPSIN026767 | RMMUS0684 | RNFUR04629 | RDR035956 | RCPIC04928 | RGOAS00071 | RALGO10744 |
| RPSIN019198 | RMMUS0715 | RNFUR01508 | RDR022458 | RCPIC01982 | RGOAS00492 | RALGO02872 |
| RPSIN005251 | RMMUS0820 | RNFUR01770 | RDR050390 | RCPIC03938 | RGOAS02017 | RALGO31278 |
| RPSIN027337 | RMMUS0981 | RNFUR04429 | RDR022734 | RCPIC00927 | RGOAS01513 | RALGO21630 |
| RPSIN018071 | RMMUS1473 | RNFUR04154 | RDR021040 | RCPIC01449 | RGOAS00030 | RALGO37619 |
| RPSIN007930 | RMMUS2858 | RNFUR04654 | RDR045768 | RCPIC03136 | RGOAS03124 | RALGO67753 |
| RPSIN006020 | RMMUS0644 | RNFUR01969 | RDR002977 | RCPIC01245 | RGOAS02570 | RALGO04443 |
| RPSIN011985 | RMMUS1914 | RNFUR04459 | RDR013523 | RCPIC04026 | RGOAS00222 | RALGO38567 |
| RPSIN016143 | RMMUS228741 | | | RCPIC05254 | RGOAS01972 | RALGO68905 |

| | | | | | | |
|---|---|---|---|---|---|---|
| RPSIN004081 | RMMUS1943 | RNFUR02213 | RDR032806 | RCPIC00567 | RGOAS03295 | RALGO31239 |
| RPSIN003797 | RMMUS2291 | RNFUR00631 | RDR031078 | RCPIC04762 | RGOAS01939 | RALGO69076 |
| RPSIN038592 | RMMUS1611 | RNFUR00905 | RDR030798 | RCPIC00543 | RGOAS01674 | RALGO54268 |
| RPSIN029301 | RMMUS0412 | RNFUR01515 | RDR022399 | RCPIC05562 | RGOAS02495 | RALGO05728 |
| RPSIN029629 | RMMUS031412 | | RDR038988 | RCPIC04282 | RGOAS01621 | not predicted |
| RPSIN010510 | RMMUS1667 | RNFUR05491 | RDR041844 | | | RALGO38471 |
| RPSIN004050 | RMMUS0376 | RNFUR02108 | RDR044131 | RCPIC02184 | RGOAS01247 | RALGO29333 |
| RPSIN035352 | RMMUS2448 | RNFUR02145 | RDR021676 | RCPIC02382 | RGOAS01509 | RALGO65707 |
| RPSIN010278 | RMMUS1829 | RNFUR00273 | RDR029171 | RCPIC02840 | RGOAS03396 | RALGO26203 |
| 32 | RMMUS2781 | RNFUR00039 | RDR028865 | RCPIC01920 | RGOAS01129 | RALGO44561 |
| RPSIN003915 | RMMUS0863 | RNFUR02350 | RDR029369 | RCPIC02828 | RGOAS02919 | RALGO14798 |
| RPSIN019576 | RMMUS1657 | RNFUR01163 | RDR042076 | RCPIC03796 | RGOAS00981 | RALGO39354 |
| RPSIN015391 | RMMUS0385 | RNFUR012885 | | RCPIC02957 | RGOAS01424 | RALGO22875 |
| RPSIN000188 | RMMUS2695 | RNFUR04033 | RDR014477 | RCPIC02922 | RGOAS00828 | RALGO31443 |
| RPSIN006549 | RMMUS0694 | RNFUR01430 | RDR041788 | RCPIC05435 | RGOAS00556 | RALGO19216 |
| RPSIN006648 | RMMUS0150 | RNFUR04561 | RDR002275 | RCPIC04605 | RGOAS02908 | RALGO45143 |
| RPSIN001429 | RMMUS1485 | RNFUR01505 | RDR028924 | | RGOAS00860 | RALGO34832 |
| RPSIN009059 | RMMUS0734 | RNFUR04493 | RDR034795 | RCPIC03214 | RGOAS03157 | RALGO22086 |
| RPSIN009120 | RMMUS0736 | RNFUR02288 | RDR008229 | RCPIC03212 | RGOAS03031 | RALGO25470 |
| RPSIN004943 | RMMUS047101 | | | RCPIC04578 | RGOAS02657 | RALGO43269 |
| RPSIN028634 | RMMUS2084 | RNFUR00202 | RDR041200 | RCPIC02118 | RGOAS01685 | RALGO51046 |
| RPSIN031963 | RMMUS2638 | RNFUR027384 | | RCPIC02110 | RGOAS01434 | RALGO48502 |
| RPSIN023640 | RMMUS1497 | RNFUR04155 | RDR046684 | RCPIC04696 | RGOAS01195 | RALGO34841 |
| RPSIN029035 | RMMUS038325 | | | RCPIC03596 | RGOAS02831 | RALGO21969 |
| RPSIN011702 | RMMUS212318 | | RDR010311 | RCPIC04233 | RGOAS01497 | RALGO37271 |
| RPSIN010210 | RMMUS087398 | | RDR014549 | RCPIC054544 | | RALGO25605 |
| RPSIN001022 | RMMUS0281 | RNFUR00153 | RDR017079 | RCPIC01654 | RGOAS01661 | RALGO11308 |
| RPSIN027245 | RMMUS1201 | RNFUR01351 | RDR029224 | RCPIC00027 | RGOAS01577 | RALGO71458 |
| RPSIN011623 | RMMUS0222 | RNFUR01210 | RDR040359 | RCPIC02028 | RGOAS02729 | RALGO15809 |
| RPSIN035075 | RMMUS2665 | RNFUR04618 | RDR010662 | RCPIC01147 | RGOAS00633 | RALGO14663 |
| RPSIN005593 | RMMUS1567 | RNFUR03274 | RDR032442 | RCPIC00283 | RGOAS02657 | RALGO09366 |
| RPSIN029763 | RMMUS0997 | RNFUR01500 | RDR022564 | | RGOAS00286 | RALGO28436 |
| RPSIN010120 | RMMUS0038 | RNFUR01042 | RDR020498 | RCPIC01418 | RGOAS02227 | RALGO25410 |
| RPSIN026103 | RMMUS218879 | | RDR004054 | RCPIC02139 | RGOAS01179 | RALGO11651 |
| RPSIN034447 | RMMUS1603 | RNFUR05234 | RDR044028 | RCPIC00693 | RGOAS02854 | RALGO55505 |
| RPSIN021801 | RMMUS038183 | | RDR043449 | RCPIC02960 | RGOAS00358 | RALGO27038 |
| RPSIN032448 | RMMUS0068 | RNFUR03827 | RDR029497 | RCPIC017366 | | RALGO33568 |
| RPSIN019119 | RMMUS1914 | RNFUR00316 | RDR041849 | RCPIC04033 | RGOAS01176 | RALGO38753 |
| RPSIN005351 | RMMUS0872 | RNFUR04315 | RDR029813 | RCPIC02881 | RGOAS02334 | RALGO08961 |
| RPSIN005611 | RMMUS0846 | RNFUR02707 | RDR011277 | RCPIC01587 | RGOAS00571 | RALGO07948 |
| RPSIN006923 | RMMUS2779 | RNFUR00968 | RDR028823 | RCPIC01282 | RGOAS01063 | RALGO51997 |
| RPSIN020124 | RMMUS134158 | | RDR021334 | RCPIC02518 | RGOAS01326 | RALGO06748 |
| RPSIN033535 | RMMUS256396 | | RDR018334 | RCPIC04170 | RGOAS01813 | RALGO37586 |
| RPSIN005677 | RMMUS0905 | RNFUR04768 | RDR036481 | RCPIC00307 | RGOAS00611 | RALGO40942 |
| RPSIN001530 | RMMUS1596 | RNFUR04150 | RDR033284 | RCPIC00654 | RGOAS03057 | RALGO56847 |

| | | | | | | |
|---|---|---|---|---|---|---|
| RPSIN028252 | RMMUS1271 | RNFUR05202 | RDR046558 | RCPIC03833 | RGOAS01859 | not predicted |
| RPSIN011245 | RMMUS0500 | RNFUR01540 | RDR044359 | RCPIC01576 | RGOAS02256 | RALGO12422 |
| RPSIN021229 | RMMUS1493 | RNFUR01596 | RDR046557 | RCPIC04385 | RGOAS02928 | RALGO38447 |
| RPSIN019988 | RMMUS1636 | RNFUR05425 | RDR026568 | RCPIC02614 | RGOAS00340 | RALGO23620 |
| RPSIN023794 | RMMUS1111 | RNFUR04315 | RDR029321 | RCPIC04988 | RGOAS01170 | RALGO04395 |
| RPSIN002134 | RMMUS1270 | RNFUR02187 | RDR015717 | RCPIC04222 | RGOAS00537 | RALGO32440 |
| RPSIN025029 | RMMUS0837 | RNFUR04672 | RDR012138 | RCPIC04165 | RGOAS02056 | RALGO25173 |
| RPSIN015295 | RMMUS2074 | RNFUR01780 | RDR037733 | RCPIC03438 | RGOAS00101 | RALGO65335 |
| RPSIN033554 | RMMUS1668 | RNFUR03519 | RDR002236 | RCPIC04150 | RGOAS00733 | RALGO37508 |
| RPSIN011106 | RMMUS1918 | RNFUR00312 | RDR042072 | RCPIC040310 | | RALGO39645 |
| RPSIN029278 | RMMUS2027 | RNFUR04079 | RDR033038 | RCPIC02942 | RGOAS03020 | RALGO60287 |
| RPSIN013456 | RMMUS1858 | RNFUR02479 | RDR024080 | RCPIC04884 | RGOAS02288 | RALGO07651 |
| RPSIN026900 | RMMUS015280 | | | RCPIC04597 | RGOAS00014 | RALGO42596 |
| 92 | RMMUS006044 | | | RCPIC04515 | RGOAS01824 | RALGO49235 |
| RPSIN026215 | RMMUS2366 | RNFUR01787 | RDR022465 | RCPIC00381 | RGOAS02691 | RALGO16693 |
| RPSIN016307 | RMMUS0912 | RNFUR02779 | RDR046726 | RCPIC00412 | RGOAS01553 | RALGO37421 |
| 43 | RMMUS1829 | RNFUR01539 | RDR040180 | RCPIC02840 | RGOAS016462 | |
| RPSIN029026 | RMMUS0389 | RNFUR04825 | RDR017427 | RCPIC03595 | RGOAS02836 | RALGO21411 |
| RPSIN005542 | RMMUS0923 | RNFUR04824 | RDR007598 | RCPIC00137 | RGOAS01880 | RALGO56637 |
| RPSIN029729 | RMMUS1814 | RNFUR00516 | RDR039943 | RCPIC02851 | RGOAS02335 | RALGO15914 |
| RPSIN009805 | RMMUS1964 | RNFUR04406 | RDR032858 | RCPIC04464 | RGOAS01946 | RALGO56274 |
| RPSIN027336 | RMMUS1562 | RNFUR04496 | RDR017494 | RCPIC01008 | RGOAS02912 | not predicted |
| RPSIN036949 | RMMUS0774 | RNFUR03899 | RDR043625 | RCPIC02279 | RGOAS02737 | RALGO56349 |
| RPSIN010082 | RMMUS0033 | RNFUR01340 | RDR019238 | RCPIC01430 | RGOAS00929 | RALGO26386 |
| RPSIN038287 | RMMUS0109 | RNFUR05122 | RDR046783 | RCPIC00870 | RGOAS02618 | RALGO18408 |
| RPSIN025270 | RMMUS2498 | RNFUR02222 | RDR006499 | RCPIC01764 | RGOAS02770 | RALGO04488 |
| 30 | RMMUS1247 | RNFUR04097 | RDR014715 | | | |
| RPSIN026913 | RMMUS237911 | | RDR054653 | RCPIC01985 | RGOAS00911 | RALGO33109 |
| | RMMUS1506 | RNFUR01719 | RDR014795 | RCPIC02561 | RGOAS03025 | RALGO49874 |
| RPSIN035035 | RMMUS2659 | RNFUR03729 | RDR020819 | RCPIC01122 | RGOAS01027 | RALGO19721 |
| RPSIN026077 | RMMUS2711 | RNFUR01579 | RDR050080 | RCPIC01116 | RGOAS03225 | RALGO53081 |
| RPSIN008062 | RMMUS1447 | RNFUR04390 | RDR005048 | RCPIC04857 | RGOAS02131 | RALGO10615 |
| RPSIN027710 | RMMUS2365 | RNFUR02553 | RDR041974 | RCPIC02742 | RGOAS00809 | RALGO12830 |
| RPSIN005428 | RMMUS1867 | RNFUR01050 | RDR005503 | RCPIC04877 | RGOAS02839 | RALGO03606 |
| RPSIN013310 | RMMUS0917 | RNFUR03355 | RDR007378 | RCPIC04355 | RGOAS00529 | RALGO01236 |
| 26 | RMMUS2375 | RNFUR04903 | RDR023392 | RCPIC04336 | RGOAS008568 | |
| RPSIN033268 | RMMUS0322 | RNFUR054439 | | RCPIC00796 | RGOAS00341 | RALGO52108 |
| RPSIN028461 | RMMUS095397 | | | RCPIC02504 | RGOAS00176 | RALGO25738 |
| RPSIN007235 | RMMUS2510 | RNFUR02467 | RDR012546 | RCPIC02924 | RGOAS00344 | RALGO22815 |
| RPSIN036269 | RMMUS2420 | RNFUR03826 | RDR018809 | RCPIC04519 | RGOAS00717 | RALGO46053 |
| RPSIN024207 | RMMUS2386 | RNFUR00181 | RDR004521 | RCPIC036983 | | RALGO24364 |
| RPSIN022101 | RMMUS1686 | RNFUR02824 | RDR007211 | RCPIC03304 | RGOAS03263 | RALGO51238 |
| RPSIN014266 | RMMUS1910 | RNFUR01316 | RDR024431 | RCPIC04029 | RGOAS00240 | RALGO39459 |
| RPSIN020304 | RMMUS2593 | RNFUR04397 | RDR026345 | RCPIC02240 | RGOAS01988 | RALGO65131 |
| RPSIN014759 | RMMUS2671 | RNFUR03052 | RDR046324 | RCPIC03435 | RGOAS02143 | RALGO65899 |

| | | | | | | |
|---|---|---|---|---|---|---|
| RPSIN032712 | RMMUS2741 | RNFUR05180 | RDR001974 | RCPIC023198 | | RALGO00747 |
| RPSIN023870 | RMMUS0763 | RNFUR00703 | RDR015084 | RCPIC02349 | RGOAS00989 | RALGO58043 |
| RPSIN037552 | RMMUS179139 | | RDR011136 | RCPIC02857 | RGOAS033871 | |
| RPSIN000434 | RMMUS0696 | RNFUR01790 | RDR043834 | RCPIC01705 | RGOAS02679 | RALGO19940 |
| RPSIN025161 | RMMUS1510 | RNFUR04297 | RDR014981 | RCPIC02571 | RGOAS01086 | RALGO49226 |
| RPSIN034185 | RMMUS1815 | RNFUR05335 | RDR009670 | RCPIC02853 | RGOAS01019 | RALGO11491 |
| RPSIN001987 | RMMUS0756 | RNFUR00841 | RDR027769 | RCPIC02264 | RGOAS02472 | RALGO59204 |
| RPSIN013423 | RMMUS1856 | RNFUR04696 | RDR005138 | RCPIC04889 | RGOAS00063 | RALGO09258 |
| RPSIN032801 | RMMUS033354 | | | RCPIC05155 | RGOAS02937 | RALGO45531 |
| RPSIN015649 | RMMUS1424 | RNFUR00699 | RDR038832 | RCPIC00584 | RGOAS01599 | RALGO59927 |
| RPSIN037010 | RMMUS1765 | RNFUR00534 | RDR027744 | RCPIC01262 | RGOAS02485 | RALGO55108 |
| RPSIN015385 | RMMUS0377 | RNFUR03259 | RDR043269 | RCPIC02958 | RGOAS01909 | RALGO23061 |
| RPSIN029774 | RMMUS0999 | RNFUR04280 | RDR014169 | RCPIC00932 | RGOAS00384 | RALGO26795 |
| RPSIN034680 | RMMUS1907 | RNFUR01181 | RDR024432 | RCPIC04040 | RGOAS03231 | RALGO30749 |
| RPSIN020017 | RMMUS2513 | RNFUR01268 | RDR012080 | RCPIC00793 | RGOAS03229 | RALGO24431 |
| RPSIN022464 | RMMUS2609 | RNFUR01409 | RDR002837 | RCPIC02166 | RGOAS00185 | RALGO67162 |
| RPSIN021115 | RMMUS0406 | RNFUR032014 | | RCPIC02965 | RGOAS00461 | RALGO11615 |
| RPSIN023175 | RMMUS1050 | RNFUR05214 | RDR053871 | RCPIC00087 | RGOAS01788 | RALGO58826 |
| | RMMUS275925 | | | | | |

| | | | | | | |
|---|---|---|---|---|---|---|
| RPSIN031143 | RMMUS0647 | RNFUR03450 | RDR003141 | RCPIC01380 | RGOAS00977 | RALGO02410 |
| RPSIN019829 | RMMUS240972 | | RDR039649 | | RGOAS02191 | RALGO37004 |
| RPSIN006924 | RMMUS2779 | RNFUR02517 | RDR028856 | RCPIC01253 | RGOAS02244 | RALGO52009 |
| RPSIN023581 | RMMUS0793 | RNFUR006105 | | | RGOAS02140 | RALGO07819 |
| RPSIN010126 | RMMUS0035 | RNFUR01458 | RDR041955 | RCPIC03526 | RGOAS00057 | RALGO28151 |
| RPSIN003923 | RMMUS1873 | RNFUR00529 | RDR005412 | RCPIC047653 | | RALGO01984 |
| RPSIN003932 | RMMUS1872 | RNFUR00515 | RDR005055 | RCPIC04770 | RGOAS00763 | RALGO02113 |
| 29 | RMMUS171288 | | | RCPIC04795 | RGOAS00347 | RALGO15509 |
| RPSIN019569 | RMMUS1658 | RNFUR04943 | RDR002355 | RCPIC03782 | RGOAS03367 | RALGO39435 |
| RPSIN031143 | RMMUS0647 | RNFUR03450 | RDR003141 | RCPIC01380 | RGOAS00977 | RALGO02410 |
| 35 | RMMUS2283 | RNFUR02745 | RDR036484 | | RGOAS00130 | RALGO68770 |
| RPSIN029042 | RMMUS038423 | | | RCPIC02972 | RGOAS01727 | RALGO22563 |
| RPSIN029706 | RMMUS0575 | RNFUR03988 | RDR021047 | RCPIC039283 | | RALGO03813 |
| RPSIN027875 | RMMUS2583 | RNFUR03869 | RDR022521 | RCPIC00594 | RGOAS02982 | RALGO61817 |
| RPSIN019037 | RMMUS2504 | RNFUR02761 | RDR045312 | RCPIC03708 | RGOAS00938 | RALGO34393 |
| RPSIN022648 | RMMUS1606 | RNFUR004612 | | RCPIC00650 | RGOAS00645 | RALGO55169 |
| RPSIN035661 | RMMUS023281 | | RDR040368 | RCPIC03275 | RGOAS02286 | RALGO10807 |
| 28 | RMMUS1289 | RNFUR00777 | RDR005211 | RCPIC03109 | RGOAS02176 | RALGO02614 |
| RPSIN029512 | RMMUS0511 | RNFUR04185 | RDR043738 | RCPIC03932 | RGOAS02740 | RALGO36365 |
| RPSIN016904 | RMMUS2243 | RNFUR01511 | RDR012692 | RCPIC04459 | RGOAS00204 | RALGO70417 |
| RPSIN014015 | RMMUS2738 | RNFUR00809 | RDR026328 | RCPIC05053 | RGOAS02581 | RALGO05311 |
| RPSIN012661 | RMMUS124715 | | | RCPIC025327 | | RALGO00870 |
| RPSIN029764 | RMMUS0997 | RNFUR04796 | RDR012975 | RCPIC00934 | RGOAS00286 | RALGO28403 |
| RPSIN007185 | RMMUS0167 | RNFUR04090 | RDR033024 | RCPIC00573 | RGOAS00064 | RALGO63955 |

| | | | | | | |
|---|---|---|---|---|---|---|
| RPSIN02721 | RMMUS2757 | RNFUR0350 | RDR017122 | RCPIC052356 | | RALGO20312 |
| RPSIN01980 | RMMUS1832 | RNFUR0280 | RDR055864 | RCPIC04032 | RGOAS0054 | RALGO35123 |
| RPSIN01135 | RMMUS0592 | RNFUR0111 | RDR016322 | RCPIC01140 | RGOAS0295 | RALGO57797 |
| RPSIN02533 | RMMUS1227 | RNFUR0096 | RDR034560 | RCPIC03078 | RGOAS0200 | RALGO4175 |
| RPSIN02864 | RMMUS2085 | RNFUR0076 | RDR021268 | RCPIC02119 | RGOAS0183 | RALGO50737 |
| RPSIN03042 | RMMUS1533 | RNFUR0194 | RDR038435 | RCPIC02812 | RGOAS0168 | RALGO46272 |
| RPSIN00647 | RMMUS1591 | RNFUR0372 | RDR033580 | RCPIC00655 | RGOAS0331 | RALGO5744 |
| RPSIN01888 | RMMUS030078 | | RDR014321 | RCPIC00806 | RGOAS0132 | RALGO49784 |
| RPSIN00640 | RMMUS1594 | RNFUR0389 | RDR038222 | RCPIC00554 | RGOAS0319 | RALGO57224 |
| RPSIN00568 | RMMUS0906 | RNFUR0496 | RDR036370 | RCPIC00307 | RGOAS0184 | RALGO41131 |
| RPSIN00415 | RMMUS1837 | RNFUR017647 | | RCPIC028547 | | RALGO07663 |
| RPSIN02144 | RMMUS0638 | RNFUR0417 | RDR042055 | RCPIC05012 | RGOAS0278 | RALGO09237 |
| RPSIN00619 | RMMUS0260 | RNFUR0487 | RDR018887 | RCPIC03504 | RGOAS0091 | RALGO3824 |
| RPSIN03096 | RMMUS2525 | RNFUR0403 | RDR036770 | RCPIC05191 | RGOAS0192 | RALGO0022 |
| RPSIN01606 | RMMUS2804 | RNFUR0349 | RDR043224 | RCPIC03648 | RGOAS0072 | RALGO25101 |
| RPSIN02646 | RMMUS1705 | RNFUR0278 | RDR007066 | RCPIC00059 | RGOAS0300 | RALGO45453 |
| RPSIN03830 | RMMUS0424 | RNFUR0191 | RDR023087 | RCPIC01835 | RGOAS0073 | RALGO15617 |
| RPSIN03684 | RMMUS0927 | RNFUR0363 | RDR034899 | RCPIC00098 | RGOAS0012 | RALGO55845 |
| RPSIN00274 | RMMUS0001 | RNFUR0262 | RDR002849 | RCPIC02663 | RGOAS0177 | RALGO67543 |
| RPSIN00309 | RMMUS1519 | RNFUR0213 | RDR015219 | RCPIC01965 | RGOAS0091 | RALGO43311 |
| RPSIN02490 | RMMUS1661 | RNFUR0417 | RDR042325 | RCPIC04172 | RGOAS016775 | |
| | RMMUS177127 | | RDR039683 | | | not predicte |
| RPSIN03263 | RMMUS2745 | RNFUR0491 | RDR024294 | RCPIC022680 | | RALGO4172 |
| RPSIN03671 | RMMUS1671 | RNFUR0255 | RDR046937 | RCPIC02120 | RGOAS003651 | |
| RPSIN01534 | RMMUS0460 | RNFUR0104 | RDR027640 | RCPIC04260 | RGOAS0104 | RALGO4530 |
| RPSIN01602 | RMMUS2003 | RNFUR0342 | RDR034397 | RCPIC03030 | RGOAS0150 | RALGO5811 |
| RPSIN03133 | RMMUS2060 | RNFUR0326 | RDR036188 | RCPIC03431 | RGOAS0032 | RALGO6553 |
| RPSIN00436 | RMMUS0784 | RNFUR0029 | RDR043456 | RCPIC04252 | RGOAS0182 | RALGO53489 |
| RPSIN02260 | RMMUS061623 | | RDR010111 | RCPIC01140 | RGOAS0106 | RALGO57257 |
| RPSIN02867 | RMMUS1196 | RNFUR0147 | RDR029581 | RCPIC02443 | RGOAS0019 | RALGO71317 |
| 48 | RMMUS2201 | RNFUR0490 | RDR003689 | RCPIC02215 | RGOAS0104 | RALGO4188 |
| RPSIN02808 | RMMUS0008 | RNFUR0276 | RDR046192 | RCPIC00332 | RGOAS0062 | RALGO68088 |
| RPSIN01588 | RMMUS139955 | | RDR008501 | RCPIC01523 | RGOAS0002 | RALGO35024 |
| RPSIN03553 | RMMUS211134 | | RDR036237 | RCPIC00273 | RGOAS0287 | not predicte |
| RPSIN02204 | RMMUS172190 | | RDR012797 | RCPIC03111 | RGOAS0157 | RALGO2069 |
| RPSIN01996 | RMMUS163486 | | RDR052664 | RCPIC02615 | RGOAS0234 | RALGO2411 |
| RPSIN01188 | RMMUS0528 | RNFUR0050 | RDR000883 | RCPIC03335 | RGOAS0012 | RALGO3150 |
| 53 | RMMUS275595 | | RDR017303 | RCPIC05179 | RGOAS0018 | RALGO2381 |
| RPSIN01194 | RMMUS2556 | RNFUR0425 | RDR022633 | RCPIC00829 | RGOAS0273 | RALGO20952 |
| RPSIN02701 | RMMUS2441 | RNFUR0368 | RDR019136 | RCPIC00005 | RGOAS0163 | RALGO6443 |
| RPSIN01402 | RMMUS1057 | RNFUR0300 | RDR031025 | RCPIC05518 | RGOAS0041 | RALGO20753 |
| RPSIN02504 | RMMUS0977 | RNFUR0473 | RDR012963 | RCPIC03879 | RGOAS0338 | RALGO6263 |
| 05 | RMMUS1103 | RNFUR023890 | | RCPIC03587 | RGOAS0241 | RALGO0262 |
| RPSIN00715 | RMMUS017102 | | RDR031210 | RCPIC00958 | RGOAS0163 | RALGO64264 |
| RPSIN00986 | RMMUS1993 | RNFUR0272 | RDR033095 | RCPIC03017 | RGOAS0153 | RALGO5682 |

| | | | | | | |
|---|---|---|---|---|---|---|
| RPSIN03252 | RMMUS1279 | RNFUR0387 | RDR046694 | RCPIC03823 | RGOAS0148 | RALGO3768 |
| RPSIN01820 | RMMUS0317 | RNFUR0056 | RDR039567 | RCPIC04265 | RGOAS0048 | RALGO5351 |
| RPSIN03858 | RMMUS161043 | | RDR038251 | RCPIC00544 | RGOAS0046 | RALGO5441 |
| RPSIN03811 | RMMUS2604 | RNFUR0228 | RDR002902 | RCPIC02168 | RGOAS0280 | RALGO6684 |
| RPSIN03165 | RMMUS0661 | RNFUR0006 | RDR036602 | RCPIC01979 | RGOAS0328 | RALGO0406 |
| RPSIN00897 | RMMUS0107 | RNFUR0510 | RDR019684 | | RGOAS0062 | RALGO43810 |
| RPSIN00138 | RMMUS147123 | | | RCPIC01451 | RGOAS0109 | RALGO3413 |
| RPSIN00048 | RMMUS2565 | RNFUR0486 | RDR041678 | RCPIC04148 | RGOAS0010 | RALGO3788 |
| RPSIN03198 | RMMUS263796 | | | RCPIC02109 | RGOAS0012 | RALGO4833 |
| RPSIN02085 | RMMUS0669 | RNFUR0411 | RDR036135 | RCPIC01981 | RGOAS0154 | RALGO0791 |
| RPSIN00142 | RMMUS1458 | RNFUR0098 | RDR034981 | RCPIC01463 | RGOAS0070 | RALGO2899 |
| RPSIN00018 | RMMUS2696 | RNFUR0242 | RDR014426 | RCPIC02928 | RGOAS0177 | RALGO3137 |
| RPSIN02061 | RMMUS1886 | RNFUR0074 | RDR049260 | RCPIC04771 | RGOAS0009 | not predicte |
| RPSIN02557 | RMMUS0777 | RNFUR0508 | RDR027678 | RCPIC04625 | RGOAS0135 | RALGO5486 |
| | RMMUS183506 | | | | | not predicte |
| RPSIN00184 | RMMUS046196 | | RDR000686 | RCPIC04279 | RGOAS0133 | RALGO4481 |
| RPSIN02418 | RMMUS1327 | RNFUR0342 | RDR000091 | RCPIC04277 | RGOAS0219 | RALGO5092 |
| RPSIN01426 | RMMUS060174 | | RDR026208 | RCPIC02037 | RGOAS0071 | RALGO1297 |
| RPSIN00720 | RMMUS0167 | RNFUR0216 | RDR038148 | RCPIC00644 | RGOAS0063 | RALGO6397 |
| | RMMUS077769 | | | | | not predicte |
| RPSIN01667 | RMMUS2866 | RNFUR0200 | RDR020465 | RCPIC03852 | RGOAS0249 | RALGO6804 |
| RPSIN01663 | RMMUS286589 | | RDR019353 | RCPIC03848 | RGOAS0249 | RALGO6803 |
| RPSIN01239 | RMMUS155210 | | RDR017152 | | | not predicte |
| RPSIN01522 | RMMUS2733 | RNFUR0033 | RDR026094 | RCPIC05057 | RGOAS0192 | RALGO06271 |
| RPSIN00745 | RMMUS1753 | RNFUR0058 | RDR051649 | RCPIC01193 | RGOAS0212 | RALGO4923 |
| RPSIN02156 | RMMUS2550 | RNFUR0006 | RDR038114 | RCPIC00600 | RGOAS0178 | RALGO62491 |
| RPSIN00028 | RMMUS2626 | RNFUR0020 | RDR027726 | RCPIC01317 | RGOAS0327 | RALGO4869 |
| RPSIN03653 | RMMUS2707 | RNFUR0189 | RDR010775 | RCPIC01129 | RGOAS0220 | RALGO5386 |
| RPSIN01619 | RMMUS1357 | RNFUR0359 | RDR022769 | RCPIC03778 | RGOAS0178 | RALGO4115 |
| RPSIN03106 | RMMUS2325 | RNFUR0110 | RDR013781 | RCPIC04676 | RGOAS0122 | RALGO54460 |
| RPSIN02360 | RMMUS2534 | RNFUR039645 | | RCPIC05567 | RGOAS0066 | RALGO0248 |
| RPSIN03759 | RMMUS182992 | | RDR029325 | RCPIC02861 | RGOAS0035 | RALGO17901 |
| RPSIN00496 | RMMUS048753 | | | RCPIC04584 | RGOAS0137 | RALGO4450 |
| RPSIN01677 | RMMUS0096 | RNFUR0340 | RDR045530 | RCPIC03437 | RGOAS0314 | RALGO6437 |
| RPSIN02712 | RMMUS2295 | RNFUR012882 | | RCPIC03561 | RGOAS0258 | RALGO6925 |
| RPSIN01551 | RMMUS249284 | | | RCPIC05729 | RGOAS0054 | RALGO01491 |
| RPSIN02169 | RMMUS178780 | | | RCPIC01776 | RGOAS0266 | RALGO0998 |
| RPSIN01917 | RMMUS071884 | | RDR018658 | | RGOAS0272 | RALGO0666 |
| RPSIN02804 | RMMUS238785 | | | RCPIC03718 | RGOAS0057 | RALGO2146 |
| | RMMUS072451 | | | | | not predicte |
| RPSIN02990 | RMMUS1026 | RNFUR0271 | RDR031327 | RCPIC00201 | RGOAS0262 | RALGO6361 |

| | | | | | | |
|---|---|---|---|---|---|---|
| RPSIN023489 | RMMUS0701 | RNFUR040438 | | RCPIC01685 | RGOAS02082 | RALGO15209 |
| RPSIN013748 | RMMUS2546 | RNFUR04216 | RDR037927 | RCPIC00603 | RGOAS02500 | RALGO18832 |
| RPSIN022397 | RMMUS072442 | | | RCPIC01709 | RGOAS00219 | not predicted |
| RPSIN029302 | | | | RCPIC05561 | RGOAS02499 | RALGO05911 |
| RPSIN023494 | RMMUS070148 | | | RCPIC01705 | RGOAS02082 | RALGO15146 |
| | RMMUS0711 | RNFUR048771 | | RCPIC03145 | RGOAS02127 | RALGO00387 |
| RPSIN032169 | RMMUS0425 | RNFUR04728 | RDR045268 | RCPIC01809 | RGOAS00548 | RALGO17853 |
| RPSIN018777 | RMMUS224829 | | | RCPIC05075 | RGOAS01058 | RALGO70321 |
| RPSIN007150 | RMMUS1293 | RNFUR01383 | RDR019912 | RCPIC04803 | RGOAS02128 | RALGO69175 |
| RPSIN019309 | RMMUS2569 | RNFUR02911 | RDR004926 | RCPIC04799 | RGOAS01224 | RALGO07585 |
| RPSIN035653 | RMMUS060098 | | | RCPIC03279 | RGOAS02326 | RALGO11314 |
| RPSIN035656 | RMMUS060106 | | | RCPIC03274 | RGOAS00474 | RALGO11446 |
| | RMMUS256910 | | | | | not predicted |
| RPSIN037059 | RMMUS1775 | RNFUR02672 | RDR027614 | RCPIC01218 | RGOAS03207 | RALGO55379 |
| RPSIN032274 | RMMUS0758 | RNFUR03669 | RDR002165 | RCPIC05128 | RGOAS02981 | RALGO58529 |
| RPSIN020410 | RMMUS095511 | | | RCPIC043673 | | RALGO36581 |
| RPSIN006305 | RMMUS1276 | RNFUR02877 | RDR014943 | RCPIC03822 | RGOAS00092 | RALGO36854 |
| 72 | RMMUS1091 | RNFUR00724 | RDR031225 | RCPIC03993 | RGOAS00196 | RALGO48568 |
| RPSIN015374 | RMMUS0380 | RNFUR03242 | RDR022469 | RCPIC02973 | RGOAS02301 | RALGO23638 |
| RPSIN005426 | RMMUS1867 | RNFUR01828 | RDR024221 | RCPIC04877 | RGOAS02473 | RALGO03987 |
| RPSIN000512 | RMMUS2393 | RNFUR03852 | RDR010381 | RCPIC04144 | RGOAS02672 | RALGO20345 |
| RPSIN030233 | RMMUS1548 | RNFUR02764 | RDR045505 | RCPIC05074 | RGOAS01440 | RALGO50809 |
| RPSIN004612 | RMMUS0551 | RNFUR05014 | RDR003151 | RCPIC02660 | RGOAS00529 | RALGO67696 |
| RPSIN002657 | RMMUS180357 | | RDR010105 | RCPIC05641 | RGOAS00570 | RALGO27242 |
| RPSIN008922 | RMMUS0324 | RNFUR01104 | RDR015826 | RCPIC02584 | RGOAS00055 | RALGO51022 |
| RPSIN008922 | RMMUS0324 | RNFUR01104 | RDR015826 | RCPIC02584 | RGOAS00055 | RALGO51022 |
| RPSIN015962 | RMMUS2411 | RNFUR03436 | RDR012538 | RCPIC04745 | RGOAS03349 | RALGO05611 |
| RPSIN019186 | RMMUS0717 | RNFUR05162 | RDR017319 | RCPIC01983 | RGOAS02946 | RALGO05064 |
| RPSIN029744 | RMMUS047791 | | RDR014779 | RCPIC04561 | RGOAS01315 | RALGO41810 |
| RPSIN000482 | | RNFUR01198 | RDR003889 | RCPIC03779 | RGOAS02522 | RALGO41269 |
| RPSIN027491 | RMMUS0031 | RNFUR01735 | RDR001058 | RCPIC04804 | RGOAS01658 | RALGO69286 |
| RPSIN010234 | RMMUS2245 | RNFUR02517 | RDR000578 | RCPIC04459 | RGOAS00020 | RALGO70369 |
| RPSIN014147 | RMMUS1392 | RNFUR01933 | RDR014800 | RCPIC01527 | RGOAS02098 | RALGO11672 |
| RPSIN013949 | RMMUS023798 | | | RCPIC02034 | RGOAS00574 | RALGO03223 |
| RPSIN032332 | RMMUS0209 | RNFUR01141 | RDR010339 | RCPIC04635 | RGOAS01169 | RALGO03942 |
| RPSIN012824 | RMMUS2023 | RNFUR01265 | RDR032725 | RCPIC02944 | RGOAS01131 | RALGO59888 |
| RPSIN017208 | RMMUS1615 | RNFUR03477 | RDR023804 | RCPIC02786 | RGOAS01548 | RALGO51508 |
| | | | | | | RALGO25759 |
| RPSIN027574 | RMMUS017654 | | RDR043128 | RCPIC02243 | RGOAS02367 | not predicted |
| RPSIN018873 | RMMUS0304 | RNFUR05386 | RDR014406 | RCPIC00812 | RGOAS00647 | RALGO48505 |
| RPSIN025282 | RMMUS2497 | RNFUR03072 | RDR004752 | RCPIC01764 | RGOAS03148 | RALGO03732 |
| RPSIN015224 | RMMUS0844 | RNFUR05086 | RDR009683 | RCPIC01639 | RGOAS00559 | RALGO07468 |
| RPSIN032058 | RMMUS1557 | RNFUR02821 | RDR022790 | RCPIC00939 | RGOAS01561 | not predicted |
| RPSIN026932 | RMMUS2379 | RNFUR00116 | RDR004833 | RCPIC01988 | RGOAS00912 | RALGO33061 |
| RPSIN008719 | RMMUS243650 | | | RCPIC00214 | RGOAS00506 | RALGO64777 |

| | | | | | | |
|---|---|---|---|---|---|---|
| | RMMUS255790 | | | | | not predicted |
| RPSIN011949 | RMMUS255844 | | | RCPIC05403 | RGOAS01250 | RALGO21348 |
| RPSIN011951 | RMMUS255851 | | | RCPIC008301 | | RALGO21378 |
| | RMMUS255827 | | | RCPIC05253 | RGOAS02041 | RALGO21315 |
| 56 | RMMUS156285 | | | | | |
| RPSIN027216 | RMMUS280100 | | | RCPIC051794 | | RALGO23857 |
| RPSIN030375 | RMMUS0947 | RNFUR01267 | RDR037113 | RCPIC00290 | RGOAS00824 | RALGO46419 |
| RPSIN014670 | RMMUS2227 | RNFUR03726 | RDR019494 | RCPIC04457 | RGOAS01829 | RALGO70477 |
| RPSIN032942 | RMMUS058299 | | | RCPIC03155 | RGOAS01292 | RALGO58559 |
| RPSIN036742 | RMMUS092510 | | | RCPIC00196 | RGOAS01063 | RALGO56028 |
| RPSIN019913 | RMMUS184869 | | RDR013461 | RCPIC04021 | RGOAS01586 | RALGO34682 |
| RPSIN030006 | RMMUS107728 | | | RCPIC05274 | RGOAS02529 | RALGO19159 |
| RPSIN014271 | RMMUS0601 | RNFUR04708 | RDR048392 | RCPIC03275 | RGOAS00153 | RALGO12479 |
| RPSIN011258 | RMMUS0501 | RNFUR04156 | RDR016759 | RCPIC01610 | RGOAS02163 | RALGO12599 |
| RPSIN013996 | RMMUS2738 | RNFUR01197 | RDR025990 | RCPIC05118 | RGOAS02575 | RALGO05061 |
| RPSIN030920 | RMMUS2006 | RNFUR01345 | RDR026951 | RCPIC03010 | RGOAS02644 | RALGO58409 |
| RPSIN003597 | RMMUS0891 | RNFUR03025 | RDR024881 | RCPIC01903 | RGOAS02773 | RALGO16600 |
| RPSIN035166 | RMMUS0219 | RNFUR00951 | RDR039992 | RCPIC02039 | RGOAS02761 | RALGO18712 |
| RPSIN025268 | RMMUS2498 | RNFUR04120 | RDR001676 | RCPIC01820 | RGOAS02771 | RALGO04404 |
| RPSIN000411 | RMMUS1709 | RNFUR04460 | RDR057074 | RCPIC00382 | RGOAS02696 | RALGO16618 |
| RPSIN007693 | RMMUS2000 | RNFUR00526 | RDR038159 | RCPIC03028 | RGOAS02597 | RALGO57359 |
| | RMMUS283492 | | | RCPIC01833 | RGOAS01877 | RALGO46059 |
| RPSIN009946 | RMMUS196551 | | RDR034098 | RCPIC03012 | RGOAS02531 | RALGO56400 |
| RPSIN034005 | RMMUS0337 | RNFUR05453 | RDR026388 | RCPIC01662 | RGOAS01313 | RALGO66889 |
| RPSIN004582 | RMMUS2611 | RNFUR00297 | RDR037762 | RCPIC02161 | RGOAS02572 | not predicted |
| 20 | RMMUS2538 | RNFUR00469 | RDR031216 | RCPIC03609 | RGOAS00813 | RALGO03561 |
| RPSIN017289 | RMMUS1614 | RNFUR04642 | RDR038366 | RCPIC02786 | RGOAS01529 | RALGO51586 |
| RPSIN020590 | RMMUS1874 | RNFUR00692 | RDR005126 | RCPIC04764 | RGOAS02132 | RALGO01572 |
| RPSIN024253 | RMMUS132270 | | RDR001609 | RCPIC04251 | RGOAS00877 | RALGO51850 |
| RPSIN004481 | RMMUS078392 | | | RCPIC043041 | | RALGO53507 |
| RPSIN011343 | RMMUS0592 | RNFUR01062 | RDR000604 | RCPIC01141 | RGOAS00018 | RALGO57833 |
| RPSIN024485 | RMMUS0321 | RNFUR01316 | RDR033070 | RCPIC00798 | RGOAS03139 | RALGO52559 |
| RPSIN032486 | RMMUS2517 | RNFUR00507 | RDR019183 | RCPIC03172 | RGOAS02555 | RALGO67108 |
| 12 | RMMUS251642 | | | RCPIC00061 | RGOAS01518 | RALGO67030 |
| RPSIN006947 | RMMUS2774 | RNFUR02037 | RDR027887 | RCPIC012317 | | RALGO52454 |
| RPSIN007751 | RMMUS176991 | | | RCPIC01255 | RGOAS01679 | RALGO54046 |
| RPSIN030451 | RMMUS0563 | RNFUR03797 | RDR038438 | RCPIC02795 | RGOAS00642 | RALGO47079 |
| RPSIN025562 | RMMUS0778 | RNFUR01903 | RDR000084 | RCPIC04623 | RGOAS01721 | RALGO54733 |
| RPSIN002698 | RMMUS2392 | RNFUR03348 | RDR045381 | RCPIC02659 | RGOAS03080 | RALGO67435 |
| RPSIN016126 | RMMUS2287 | RNFUR02073 | RDR031032 | RCPIC05045 | RGOAS00651 | RALGO68908 |
| RPSIN009775 | RMMUS2254 | RNFUR03008 | RDR018845 | RCPIC03147 | RGOAS01990 | RALGO69961 |
| | RMMUS136476 | | | | | not predicted |
| RPSIN022967 | RMMUS120437 | | | RCPIC00130 | RGOAS00577 | RALGO71530 |
| RPSIN001329 | RMMUS0828 | RNFUR02143 | RDR040262 | RCPIC01601 | RGOAS00303 | RALGO16313 |

| | | | | | | |
|---|---|---|---|---|---|---|
| RPSIN03462[illegible] | RMMUS190480 | | RDR024948 | RCPIC04053 | RGOAS0085[illegible] | RALGO31272 |
| RPSIN00105[illegible] | RMMUS2674 | RNFUR0090[illegible] | RDR000439 | RCPIC01578 | RGOAS0140[illegible] | RALGO10534 |
| RPSIN02185[illegible] | RMMUS0833 | RNFUR0303[illegible] | RDR010219 | RCPIC01608 | RGOAS0319[illegible] | RALGO05899 |
| RPSIN02688[illegible] | RMMUS2105 | RNFUR0368[illegible] | RDR036065 | RCPIC05026 | RGOAS0017[illegible] | RALGO68680 |
| RPSIN03428[illegible] | RMMUS0894 | RNFUR0518[illegible] | RDR045161 | RCPIC04473 | RGOAS0321[illegible] | RALGO70780 |
| RPSIN01681[illegible] | RMMUS0756 | RNFUR0296[illegible] | RDR027758 | RCPIC02263 | RGOAS0329[illegible] | RALGO59081 |
| RPSIN00157[illegible] | RMMUS2868 | RNFUR0052[illegible] | RDR027851 | RCPIC03071 | RGOAS0100[illegible] | RALGO43106 |
| RPSIN02715[illegible] | RMMUS0965 | RNFUR0413[illegible] | RDR010167 | RCPIC03198 | RGOAS0203[illegible] | RALGO61922 |
| RPSIN00991[illegible] | RMMUS1994 | RNFUR0278[illegible] | RDR033304 | RCPIC03036 | RGOAS0263[illegible] | RALGO56910 |
| RPSIN02833[illegible] | RMMUS2569 | RNFUR002767 | | RCPIC01682 | RGOAS0128[illegible] | RALGO06790 |
| RPSIN01258[illegible] | RMMUS0511 | RNFUR0426[illegible] | RDR004298 | RCPIC03956 | RGOAS0291[illegible] | RALGO36062 |
| RPSIN01259[illegible] | RMMUS0513 | RNFUR0133[illegible] | RDR041772 | RCPIC03952 | RGOAS0288[illegible] | RALGO33877 |
| RPSIN02971[illegible] | RMMUS0574 | RNFUR0488[illegible] | RDR010238 | RCPIC03928 | RGOAS0142[illegible] | RALGO03936 |
| RPSIN02786[illegible] | RMMUS2551 | RNFUR0098[illegible] | RDR031175 | RCPIC00641 | RGOAS0014[illegible] | RALGO62401 |
| RPSIN02819[illegible] | RMMUS1333 | RNFUR0512[illegible] | RDR000924 | RCPIC04268 | RGOAS0335[illegible] | RALGO49394 |
| RPSIN02981[illegible] | RMMUS1002 | RNFUR0197[illegible] | RDR012293 | RCPIC00932 | RGOAS0296[illegible] | RALGO26629 |
| RPSIN02516[illegible] | RMMUS1171 | RNFUR0156[illegible] | RDR014418 | RCPIC02570 | RGOAS0116[illegible] | RALGO49118 |
| RPSIN00637[illegible] | RMMUS1592 | RNFUR0114[illegible] | RDR025844 | RCPIC00615 | RGOAS0300[illegible] | RALGO57527 |
| RPSIN01364[illegible] | RMMUS1583 | RNFUR0456[illegible] | RDR045570 | RCPIC00267 | RGOAS0048[illegible] | RALGO13913 |
| RPSIN02348[illegible] | RMMUS0701 | RNFUR0423[illegible] | RDR044774 | RCPIC01697 | RGOAS0326[illegible] | RALGO15269 |
| RPSIN01440[illegible] | RMMUS1373 | RNFUR0202[illegible] | RDR014369 | RCPIC01504 | RGOAS0156[illegible] | RALGO21033 |
| RPSIN01486[illegible] | RMMUS1630 | RNFUR0537[illegible] | RDR025790 | RCPIC02608 | RGOAS0204[illegible] | not predicted |
| RPSIN03265[illegible] | RMMUS2744 | RNFUR0486[illegible] | RDR024928 | RCPIC02311 | RGOAS0308[illegible] | RALGO41825 |
| 30 | RMMUS1464 | RNFUR0461[illegible] | RDR047138 | RCPIC03690 | RGOAS0103[illegible] | RALGO30743 |
| 45 | RMMUS2757 | RNFUR0442[illegible] | RDR017349 | RCPIC05236 | RGOAS011716 | |
| RPSIN01644[illegible] | RMMUS0913 | RNFUR0120[illegible] | RDR046541 | RCPIC00322 | RGOAS0071[illegible] | RALGO37226 |
| RPSIN01553[illegible] | RMMUS2480 | RNFUR0086[illegible] | RDR026752 | RCPIC006850 | | not predicted |
| RPSIN00346[illegible] | RMMUS152175 | | | RCPIC01877 | RGOAS0338[illegible] | RALGO44009 |
| RPSIN03135[illegible] | RMMUS1312 | RNFUR0325[illegible] | RDR021204 | RCPIC03882 | RGOAS0317[illegible] | RALGO63829 |
| RPSIN02643[illegible] | RMMUS170537 | | RDR007384 | RCPIC00059 | RGOAS0303[illegible] | RALGO45615 |
| RPSIN03074[illegible] | RMMUS1201 | RNFUR0179[illegible] | RDR020959 | RCPIC00028 | RGOAS0172[illegible] | RALGO71422 |
| RPSIN00553[illegible] | RMMUS0686 | RNFUR0300[illegible] | RDR022466 | RCPIC05003 | RGOAS0040[illegible] | RALGO21873 |
| RPSIN00263[illegible] | RMMUS1803 | RNFUR0030[illegible] | RDR014666 | RCPIC04725 | RGOAS0073[illegible] | RALGO27377 |
| RPSIN02013[illegible] | RMMUS131813 | | RDR022011 | RCPIC02518 | RGOAS0295[illegible] | RALGO06259 |
| RPSIN02476[illegible] | RMMUS0308 | RNFUR0419[illegible] | RDR046693 | RCPIC03169 | RGOAS0002[illegible] | RALGO44246 |
| RPSIN01617[illegible] | RMMUS1408 | RNFUR0406[illegible] | RDR046770 | RCPIC01517 | RGOAS0276[illegible] | RALGO29345 |
| RPSIN00763[illegible] | RMMUS044708 | | RDR042247 | RCPIC01800 | RGOAS0248[illegible] | RALGO28613 |
| RPSIN01280[illegible] | RMMUS1288 | RNFUR0094[illegible] | RDR004692 | RCPIC04847 | RGOAS0278[illegible] | RALGO23067 |
| | RMMUS1730 | RNFUR0321[illegible] | RDR010124 | RCPIC01614 | RGOAS0260[illegible] | RALGO00312 |
| RPSIN01033[illegible] | RMMUS2580 | RNFUR0353[illegible] | RDR034350 | RCPIC00643 | RGOAS0329[illegible] | RALGO63601 |
| RPSIN00692[illegible] | RMMUS2778 | RNFUR0363[illegible] | RDR027814 | RCPIC01253 | RGOAS0320[illegible] | RALGO52054 |
| RPSIN01896[illegible] | RMMUS2230 | RNFUR0529[illegible] | RDR053481 | RCPIC00580 | RGOAS0143[illegible] | RALGO12146 |
| RPSIN00617[illegible] | RMMUS0582 | RNFUR0226[illegible] | RDR000853 | RCPIC04955 | RGOAS0249[illegible] | RALGO60113 |
| RPSIN02780[illegible] | RMMUS0987 | RNFUR0234[illegible] | RDR018374 | RCPIC00929 | RGOAS0340[illegible] | RALGO24804 |
| RPSIN01578[illegible] | RMMUS1744 | RNFUR0019[illegible] | RDR033652 | RCPIC02701 | RGOAS0223[illegible] | RALGO65791 |

| | | | | | | |
|---|---|---|---|---|---|---|
| RPSIN02573 | RMMUS1098 | RNFUR0272 | RDR030751 | RCPIC05409 | RGOAS0142 | RALGO48187 |
| RPSIN02533 | RMMUS0745 | RNFUR0196 | RDR046319 | RCPIC00300 | RGOAS0219 | RALGO21210 |
| RPSIN00880 | RMMUS2025 | RNFUR0083 | RDR037792 | RCPIC02943 | RGOAS0230 | RALGO60065 |
| RPSIN01865 | RMMUS2121 | RNFUR0061 | RDR021037 | RCPIC02398 | RGOAS0090 | RALGO41203 |
| RPSIN01926 | RMMUS0057 | RNFUR0539 | RDR018810 | RCPIC04522 | RGOAS0079 | RALGO48767 |
| RPSIN01823 | RMMUS2336 | RNFUR0402 | RDR024349 | RCPIC04654 | RGOAS0223 | RALGO53543 |
| RPSIN00087 | RMMUS1431 | RNFUR0141 | RDR034726 | RCPIC00227 | RGOAS0184 | RALGO71038 |
| RPSIN00268 | RMMUS180649 | | | RCPIC05644 | RGOAS0205 | RALGO27161 |
| RPSIN00246 | RMMUS0674 | RNFUR0112 | RDR033123 | RCPIC02722 | RGOAS0055 | RALGO14372 |
| RPSIN02227 | RMMUS1821 | RNFUR0294 | RDR029391 | RCPIC02911 | RGOAS0045 | RALGO27035 |
| RPSIN00921 | RMMUS0870 | RNFUR0008 | RDR030650 | RCPIC02831 | RGOAS0272 | RALGO08410 |
| RPSIN02038 | RMMUS0022 | RNFUR0358 | RDR016865 | RCPIC041515 | | RALGO36935 |
| RPSIN03083 | RMMUS1482 | RNFUR0373 | RDR040666 | RCPIC01483 | RGOAS0178 | RALGO35771 |
| RPSIN00283 | RMMUS2430 | RNFUR0104 | RDR011623 | RCPIC00106 | RGOAS0097 | RALGO64117 |
| | RMMUS088268 | | | | RGOAS0028 | RALGO14522 |
| RPSIN00353 | RMMUS0062 | RNFUR0007 | RDR006216 | RCPIC045501 | | RALGO49562 |
| RPSIN02436 | RMMUS1987 | RNFUR0047 | RDR038455 | RCPIC00795 | RGOAS0323 | RALGO51847 |
| RPSIN03231 | RMMUS0758 | RNFUR0405 | RDR033466 | RCPIC02294 | RGOAS0254 | RALGO58613 |
| RPSIN00414 | RMMUS0654 | RNFUR0185 | RDR053662 | RCPIC02726 | RGOAS0299 | RALGO22644 |
| RPSIN03784 | RMMUS2222 | RNFUR0182 | RDR019329 | RCPIC00095 | RGOAS0259 | RALGO70894 |
| RPSIN01851 | RMMUS1024 | RNFUR0435 | RDR036387 | RCPIC02224 | RGOAS027768 | |
| RPSIN00868 | RMMUS2439 | RNFUR0280 | RDR020911 | RCPIC00007 | RGOAS0256 | RALGO64564 |

| BHWH | RTNK | RJON | Notes |
|---|---|---|---|
| RBHWH0055 | RRTNK77294 | RRJON77087 | |
| RBHWH0196 | RRTNK83962 | RRJON83711 | |
| RBHWH0036 | RRTNK50525 | RRJON50302 | |
| RBHWH0074 | RRTNK64300 | RRJON64061 | |
| RBHWH0168 | RRTNK26840 | RRJON26726 | |
| | RRTNK52859 | RRJON52623 | Manually edited ALGO, TANK, JON, LG |
| | not predicted | not predicted | |
| | RRTNK33815 | RRJON33690 | |
| RBHWH0052 | RRTNK07549 | RRJON07465 | |
| RBHWH0175 | RRTNK21194 | RRJON21097 | |
| RBHWH0124 | RRTNK59306 | RRJON59055 | |
| RBHWH0136 | RRTNK56545 | RRJON56294 | |
| RBHWH0083 | RRTNK65580 | RRJON65329 | Manually edited ALGO, TANK, JON |
| RBHWH0077 | RRTNK65818 | RRJON65560 | |
| RBHWH0106 | RRTNK38962 | RRJON38800 | |
| RBHWH0087 | RRTNK31053 | RRJON30948 | |
| | RRTNK39364 | RRJON39176 | |
| RBHWH0096 | RRTNK04114 | RRJON04069 | |
| RBHWH0184 | RRTNK58039 | RRJON57776 | |
| | RRTNK68191 | RRJON67948 | |
| | RRTNK23084 | RRJON22976 | |
| RBHWH0173 | RRTNK70960 | RRJON70770 | |
| RBHWH0186 | RRTNK36127 | RRJON35997 | |
| RBHWH0110 | RRTNK54105 | RRJON53850 | |
| RBHWH0106 | RRTNK33606 | RRJON33487 | |
| RBHWH0116 | RRTNK73654 | RRJON73418 | Manually edited ALGO, TANK, JON |
| RBHWH0093 | RRTNK83425 | RRJON83191 | Manually edited ALGO, TANK, JON, LG. Transcripts of b |
| RBHWH0176 | RRTNK65170 | RRJON64925 | |
| RBHWH0031 | not predicted | not predicted | |
| RBHWH0147 | RRTNK30820 | RRJON30702 | |
| RBHWH0172 | RRTNK75512 | RRJON75291 | |
| RBHWH0221 | RRTNK59567 | RRJON59314 | |
| RBHWH0164 | RRTNK07992 | RRJON07905 | |
| RBHWH0110 | RRTNK65326 | RRJON65069 | |
| RBHWH0055 | RRTNK77193 | RRJON76988 | |
| RBHWH0031 | RRTNK27042 | RRJON05843 | |
| RBHWH0162 | RRTNK45598 | RRJON45427 | |

| | | | | |
|---|---|---|---|---|
| | RBHWH0211 | RRTNK48641 | RRJON48407 | |
| | RBHWH0198 | RRTNK32934 | RRJON32806 | |
| | RBHWH0092 | RRTNK60811 | RRJON60589 | |
| [illegible] | | RRTNK76626 | RRJON76408 | |
| | RBHWH0063 | RRTNK08710 | RRJON08616 | |
| [illegible] | | RRTNK04278 | RRJON04242 | |
| [illegible] | | RRTNK77218 | RRJON77011 | Manually edited TANK, JON. They are missing the first [illegible] |
| | RBHWH0057 | RRTNK14875 | RRJON14817 | |
| | RBHWH0146 | RRTNK42629 | RRJON42431 | |
| [illegible] | | RRTNK78962 | RRJON78733 | |
| | RBHWH0082 | RRTNK11925 | RRJON11803 | |
| | RBHWH0104 | RRTNK60682 | RRJON60455 | |
| | RBHWH0141 | RRTNK33029 | RRJON32903 | |
| [illegible] | | RRTNK10058 | RRJON09931 | |
| | RBHWH0034 | RRTNK71483 | RRJON71240 | |
| [illegible] | | RRTNK76125 | RRJON75901 | |
| | RBHWH0167 | RRTNK12023 | RRJON11904 | |
| [illegible] | | RRTNK75723 | RRJON75502 | Not predicted in LG |
| | RBHWH0049 | RRTNK43381 | RRJON43154 | |
| | RBHWH0092 | RRTNK44013 | RRJON43804 | |
| | RBHWH0058 | RRTNK75093 | RRJON74859 | |
| [illegible] | | RRTNK48896 | RRJON48662 | |
| | RBHWH0190 | RRTNK46513 | RRJON46278 | |
| | RBHWH0074 | RRTNK64155 | RRJON63910 | |
| | RBHWH0118 | RRTNK48751 | RRJON48517 | |
| | RBHWH0170 | RRTNK56057 | RRJON55797 | |
| | RBHWH0066 | RRTNK11827 | RRJON11710 | Manually edited ALGO, TANK, JON, LG. |
| | RBHWH0136 | RRTNK10744 | RRJON10632 | |
| | RBHWH0182 | RRTNK04791 | RRJON04767 | |
| [illegible] | | RRTNK81426 | RRJON81209 | |
| | RBHWH0181 | RRTNK35943 | RRJON35820 | |
| | RBHWH0063 | RRTNK22139 | RRJON22044 | |
| [illegible] | | RRTNK00682 | RRJON35801 | Manually edited ALGO, TANK |
| [illegible] | | RRTNK35924 | RRJON35798 | |
| | RBHWH0119 | RRTNK32272 | RRJON32147 | |
| | RBHWH0055 | RRTNK77190 | RRJON76985 | |
| [illegible] | | RRTNK30767 | RRJON30652 | |
| [illegible] | | not predicted | not predicted | Not predicted in torts |
| | RBHWH0009 | not predicted | not predicted | Not predicted in torts |
| [illegible] | | not predicted | not predicted | Not predicted in torts |
| [illegible] | | not predicted | not predicted | Not predicted in torts |
| [illegible] | | not predicted | not predicted | Not predicted in torts |
| [illegible] | | not predicted | not predicted | Not predicted in torts |
| [illegible] | | not predicted | not predicted | Not predicted in torts |
| | RBHWH0104 | not predicted | not predicted | Not predicted in torts |

| | | | |
|---|---|---|---|
| | not predictec | not predicted | |
| | not predictec | not predicted | |
| RBHWH0181 | RRTNK35932 | RRJON35809 | |
| RBHWH0155 | not predictec | not predicted | |
| RBHWH0022 | RRTNK05671 | RRJON05635 | |
| RBHWH0155 | not predictec | not predicted | |
| RBHWH0039 | not predictec | not predicted | |
| RBHWH0168 | RRTNK12400 | RRJON12272 | |
| | not predictec | not predicted | |
| RBHWH0011 | not predictec | not predicted | |
| | not predictec | not predicted | |
| | not predictec | not predicted | |
| | not predictec | not predicted | |
| RBHWH0011 | RRTNK67649 | RRJON67394 | |
| RBHWH0019 | RRTNK02042 | RRJON02022 | |
| RBHWH0031 | RRTNK05715 | RRJON05667 | |
| RBHWH0153 | RRTNK82769 | RRJON82538 | |
| RBHWH0128 | RRTNK81715 | RRJON81478 | |
| RBHWH0190 | RRTNK08013 | RRJON07923 | |
| RBHWH0059 | RRTNK62839 | RRJON62587 | |
| RBHWH0076 | RRTNK16935 | RRJON16837 | |
| RBHWH0041 | RRTNK61541 | RRJON61293 | |
| | RRTNK64912 | RRJON64674 | Manually edited ALGO, TANK, JON, LG. TANK and JON r |
| | not predictec | not predicted | |
| RBHWH0019 | RRTNK03876 | RRJON03837 | |
| RBHWH0139 | RRTNK31904 | RRJON31790 | |
| RBHWH0020 | RRTNK21706 | RRJON21610 | |
| RBHWH0048 | RRTNK72574 | RRJON72350 | Manually edited ALGO, TANK, JON |
| RBHWH0097 | RRTNK54065 | RRJON53810 | |
| RBHWH0111 | RRTNK18061 | RRJON17968 | Manually edited LG |
| RBHWH0040 | RRTNK53958 | RRJON53694 | |
| RBHWH0184 | RRTNK34150 | RRJON34027 | |
| RBHWH0086 | RRTNK17060 | RRJON16927 | Manually edited ALGO, LG |
| RBHWH0036 | RRTNK85894 | RRJON85619 | |
| | not predictec | not predictec | There is a short hit RRJON17757 |
| RBHWH0022 | RRTNK58208 | RRJON57943 | |
| RBHWH0005 | RRTNK07703 | RRJON07625 | |
| RBHWH0158 | RRTNK20449 | RRJON20368 | |
| RBHWH0077 | RRTNK17664 | RRJON17581 | |
| | RRTNK45943 | RRJON45738 | |
| RBHWH0073 | RRTNK80410 | RRJON80180 | |
| RBHWH0009 | RRTNK66376 | RRJON66131 | |
| RBHWH0063 | RRTNK18634 | RRJON18550 | |
| RBHWH0198 | RRTNK57011 | RRJON56768 | |
| RBHWH0031 | RRTNK04977 | RRJON04920 | |

| | | | |
|---|---|---|---|
| RBHWH0046 | RRTNK24488 | RRJON24369 | |
| RBHWH0194 | RRTNK18508 | RRJON18427 | |
| RBHWH0072 | RRTNK55262 | RRJON54995 | |
| RBHWH0215 | RRTNK52169 | RRJON51929 | |
| RBHWH0008 | RRTNK29147 | RRJON29013 | |
| RBHWH0049 | RRTNK61243 | RRJON61007 | |
| RBHWH0133 | RRTNK05597 | RRJON05558 | |
| RBHWH0202 | RRTNK57121 | RRJON56887 | |
| RBHWH011148 | | RRJON84714 | |
| RBHWH0105 | RRTNK79786 | RRJON79561 | Manually edited LG |
| RBHWH0195 | RRTNK01103 | RRJON01093 | Manually edited ALGO, TANK, JON, LG. All four are miss |
| RBHWH0208 | RRTNK83635 | RRJON83388 | |
| RBHWH0012 | RRTNK82042 | RRJON81811 | |
| RBHWH0014 | RRTNK19839 | RRJON19759 | |
| RBHWH0198 | RRTNK79460 | RRJON79228 | |
| RBHWH0077 | RRTNK30873 | RRJON30761 | |
| RBHWH0106 | RRTNK58385 | RRJON58137 | |
| RBHWH0059 | RRTNK62715 | RRJON62455 | |
| RBHWH0077 | RRTNK30957 | RRJON30843 | Manually edited ALGO |
| RBHWH0069 | RRTNK15053 | RRJON14996 | |
| RBHWH0141 | not predicted | not predicted | |
| RBHWH0018 | RRTNK41209 | RRJON41030 | Manually edited ALGO, TANK, JON, LG |
| [illegible] | RRTNK55931 | RRJON55663 | Manually edited LG |
| RBHWH0126 | RRTNK31967 | RRJON31853 | |
| RBHWH0108 | RRTNK67200 | RRJON66947 | |
| RBHWH0050 | RRTNK49116 | RRJON48884 | |
| RBHWH0031 | RRTNK05747 | RRJON05679 | |
| RBHWH0143 | RRTNK30794 | RRJON30679 | |
| RBHWH0061 | RRTNK64878 | RRJON64640 | |
| RBHWH0077 | RRTNK81955 | RRJON67053 | |
| RBHWH0101 | RRTNK71255 | RRJON71011 | |
| RBHWH0063 | RRTNK09806 | RRJON09681 | CABP's rep is possibly truncated |
| RBHWH0162 | RRTNK38083 | RRJON37904 | |
| RBHWH0030 | RRTNK81340 | RRJON81119 | Manually edited ALGO, TANK, JON, LG. ALGO is missing |
| [illegible] | RRTNK75322 | RRJON75096 | Manually edited ALGO, TANK, JON. |
| [illegible] | RRTNK46547 | RRJON46308 | |
| RBHWH0074 | RRTNK27851 | RRJON27741 | |
| RBHWH0005 | RRTNK08893 | RRJON08802 | |
| [illegible] | RRTNK33947 | RRJON33816 | |
| RBHWH0083 | RRTNK07325 | RRJON07245 | |
| RBHWH0063 | RRTNK08827 | RRJON08718 | |
| RBHWH0089 | RRTNK72119 | RRJON71894 | |
| RBHWH0210 | RRTNK76038 | RRJON75809 | |
| RBHWH0172 | RRTNK22278 | RRJON22170 | |
| RBHWH013879 | | | |

| | | | |
|---|---|---|---|
| RBHWH0155 | RRTNK01067 | RRJON01060 | |
| RBHWH0044 | RRTNK67225 | RRJON66973 | BCL5 is now BCL6 |
| RBHWH0102 | RRTNK17327 | RRJON17237 | Manually edited ALGO, LG |
| RBHWH0137 | RRTNK81161 | RRJON80941 | |
| RBHWH0069 | RRTNK82618 | RRJON82393 | |
| RBHWH0152 | RRTNK32248 | RRJON32130 | |
| RBHWH0130 | RRTNK74919 | RRJON74676 | |
| RBHWH0133 | RRTNK56445 | RRJON56184 | |
| RBHWH0155 | RRTNK28774 | RRJON28653 | |
| RBHWH0219 | RRTNK74410 | RRJON74176 | |
| RBHWH0089 | RRTNK72122 | RRJON71897 | Not predicted in ALGO |
| RBHWH0077 | not predictec | not predicted | |
| RBHWH007892 | | | Aldabra versions of the genes were Diamond BBHs witl |
| RBHWH0078 | RRTNK85113 | RRJON84840 | Not predicted in LG |
| RBHWH0131 | RRTNK81992 | RRJON81759 | |
| RBHWH0172 | RRTNK22098 | RRJON22006 | |
| RBHWH0103 | RRTNK06162 | RRJON06130 | |
| RBHWH0118 | RRTNK38918 | RRJON38756 | |
| RBHWH0008 | RRTNK49369 | RRJON49132 | Manually edited TANK, JON |
| RBHWH0168 | RRTNK51303 | RRJON51086 | |
| RBHWH0219 | RRTNK22641 | RRJON22546 | |
| RBHWH0110 | RRTNK18385 | RRJON22546 | |
| RBHWH0141 | RRTNK49197 | RRJON48958 | |
| RBHWH0157 | RRTNK06453 | RRJON06429 | |
| [illegible] | RRTNK69882 | RRJON69640 | Manually edited TANK, JON, LG |
| RBHWH0211 | RRTNK29144 | RRJON29010 | |
| RBHWH0151 | RRTNK84683 | RRJON84427 | Manually edited LG |
| RBHWH0199 | RRTNK29501 | RRJON29370 | |
| RBHWH0191 | RRTNK28292 | RRJON28165 | |
| [illegible] | RRTNK78872 | RRJON78633 | |
| RBHWH0035 | RRTNK31354 | RRJON31243 | |
| RBHWH0192 | RRTNK65616 | RRJON65354 | |
| RBHWH0190 | RRTNK39507 | RRJON39318 | |
| RBHWH0197 | RRTNK10195 | RRJON10082 | |
| RBHWH0178 | RRTNK78475 | RRJON78228 | |
| RBHWH0202 | not predictec | not predicted | |
| RBHWH0100 | RRTNK43250 | RRJON43034 | |
| [illegible] | RRTNK80895 | RRJON80666 | |
| RBHWH0030 | RRTNK20045 | RRJON19360 | |
| RBHWH0006 | RRTNK65727 | RRJON65467 | |
| RBHWH0144 | not predictec | not predicted | |
| RBHWH0221 | RRTNK30690 | RRJON30572 | |
| RBHWH0216 | RRTNK34316 | RRJON34191 | |
| RBHWH0045 | RRTNK47800 | RRJON47554 | Not predicted in LG. |
| RBHWH0045 | RRTNK47785 | RRJON47533 | |

| | | | |
|---|---|---|---|
| RBHWH0085 | RRTNK20068 | RRJON19984 | |
| | RRTNK47794 | RRJON47539 | |
| | RRTNK32835 | RRJON32715 | |
| RBHWH0114 | RRTNK79020 | RRJON78784 | |
| | RRTNK05326 | RRJON05266 | |
| RBHWH0076 | RRTNK19318 | RRJON19217 | |
| RBHWH0217 | RRTNK05674 | RRJON05641 | |
| RBHWH0055 | RRTNK77093 | RRJON76888 | |
| RBHWH0224 | RRTNK26013 | RRJON25916 | Manually updated LG. Very similar to CR2 |
| | not predicted | not predicted | Not predicted in torts |
| RBHWH0045 | RRTNK32289 | RRJON32167 | |
| RBHWH0065 | RRTNK02546 | RRJON02500 | |
| | not predicted | not predicted | Not predicted in torts |
| | RRTNK69451 | RRJON69196 | |
| RBHWH0060 | RRTNK69469 | RRJON69214 | |
| RBHWH0089 | RRTNK51215 | RRJON51000 | |
| RBHWH0015 | RRTNK26112 | RRJON26012 | |
| RBHWH0015 | RRTNK26184 | RRJON26074 | Not predicted in ALGO |
| | RRTNK09983 | RRJON09877 | Not predicted in ALGO |
| RBHWH0060 | RRTNK41813 | RRJON41629 | |
| RBHWH0060 | RRTNK69679 | RRJON69434 | Not predicted in ALGO |
| | RRTNK72012 | RRJON71786 | |
| | RRTNK68023 | RRJON67767 | Manually edited ALGO, TANK, JON, LG |
| RBHWH0057 | RRTNK18871 | RRJON18770 | |
| | RRTNK36999 | RRJON36837 | |
| | RRTNK35308 | RRJON35176 | |
| RBHWH0097 | RRTNK23864 | RRJON23760 | |
| RBHWH0048 | RRTNK09122 | RRJON09025 | |
| RBHWH0049 | RRTNK12130 | RRJON12020 | |
| RBHWH0055 | RRTNK72967 | RRJON72750 | |
| RBHWH0087 | RRTNK36466 | RRJON36317 | |
| RBHWH0170 | RRTNK57839 | RRJON57584 | |
| RBHWH0102 | RRTNK52032 | RRJON51800 | |
| RBHWH0022 | RRTNK57634 | not predicted | |
| | not predicted | not predicted | Not predicted in torts |
| | not predicted | not predicted | Not predicted in torts |
| RBHWH0060 | not predicted | not predicted | Not predicted in torts |
| | not predicted | not predicted | Not predicted in torts |
| RBHWH0147 | RRTNK31200 | RRJON31089 | |
| RBHWH0205 | RRTNK49186 | RRJON48947 | |
| RBHWH0091 | RRTNK65348 | RRJON65091 | |

| | | | |
|---|---|---|---|
| RBHWH0175 | RRTNK53463 | RRJON53209 | |
| RBHWH0183 | RRTNK59166 | RRJON58898 | |
| RBHWH0024 | RRTNK50109 | RRJON49894 | |
| | RRTNK21143 | RRJON21049 | Manually edited TANK, JON |
| RBHWH0032 | RRTNK32925 | RRJON32803 | |
| | RRTNK80671 | RRJON80446 | |
| RBHWH015531 | | | Not predicted in torts |
| RBHWH0145 | RRTNK71867 | RRJON71645 | |
| RBHWH0040 | RRTNK49933 | RRJON49712 | |
| RBHWH0171 | RRTNK81311 | RRJON81089 | |
| RBHWH0144 | RRTNK73822 | RRJON73578 | |
| | not predicted | not predicted | Not predicted in torts |
| | not predicted | not predicted | Not predicted in torts |
| | not predicted | not predicted | Not predicted in torts. They have lots of CCL/4/5 types |
| | not predicted | not predicted | Not predicted in torts |
| RBHWH0121 | RRTNK18906 | RRJON18808 | |
| | not predicted | not predicted | Not predicted in torts. They have lots of CCL/4/5 types |
| RBHWH0037 | RRTNK57551 | RRJON57323 | |
| | not predicted | RRJON12281 | Manually edited ALGO, JON, LG. TANK not predicted |
| | RRTNK12314 | RRJON12215 | |
| RBHWH0028 | RRTNK12350 | RRJON12245 | |
| | not predicted | not predicted | Not predicted in torts |
| | not predicted | not predicted | Not predicted in torts |
| RBHWH0136 | RRTNK42245 | RRJON36162 | |
| | RRTNK58370 | RRJON58114 | |
| RBHWH0124 | RRTNK76738 | RRJON76534 | |
| RBHWH0000 | RRTNK29962 | RRJON29823 | |
| RBHWH0030 | RRTNK32278 | RRJON32153 | Manually edited LG |
| RBHWH0173 | RRTNK60916 | RRJON60689 | |
| | not predicted | not predicted | Not predicted in torts |
| RBHWH0056 | not predicted | not predicted | Not predicted in torts |
| RBHWH0226 | RRTNK45267 | RRJON45080 | |
| RBHWH0193 | RRTNK74590 | RRJON74358 | |
| RBHWH0135 | RRTNK27137 | RRJON27011 | |
| RBHWH0078 | not predicted | not predicted | Not predicted in torts |
| RBHWH0055 | RRTNK61952 | RRJON61705 | Not predicted in LG. Has CD164L2 versions |
| RBHWH0125 | RRTNK64681 | not predicted | Not predicted in JON |
| RBHWH0008 | RRTNK00574 | RRJON00564 | |
| RBHWH0154 | RRTNK06033 | RRJON03929 | |
| RBHWH0093 | RRTNK80873 | RRJON80646 | |
| | RRTNK09962 | RRJON08345 | |
| RBHWH0221 | not predicted | not predicted | Not predicted in torts |
| RBHWH0122 | RRTNK11674 | RRJON11568 | |
| | RRTNK57297 | RRJON57072 | |
| RBHWH0142 | RRTNK81065 | RRJON80840 | |

| | | | |
|---|---|---|---|
| RBHWH0114 | not predictec | not predictec | Not predicted in torts |
| RBHWH0097 | RRTNK47526 | RRJON47279 | |
| RBHWH0029 | RRTNK54245 | RRJON53986 | |
| RBHWH0194 | not predictec | not predictec | Not predicted in torts |
| d | not predictec | not predictec | Manually edited LG. CD300LF/LB most similar to the o |
| 2 | RRTNK53105 | RRJON52873 | |
| RBHWH0194 | RRTNK53096 | RRJON52830 | |
| RBHWH0055 | RRTNK39724 | RRJON39533 | |
| RBHWH0136 | RRTNK25909 | RRJON25801 | |
| RBHWH0210 | RRTNK34752 | RRJON34617 | |
| RBHWH0063 | RRTNK09866 | RRJON09727 | |
| RBHWH0190 | RRTNK39532 | RRJON39337 | |
| RBHWH0032 | RRTNK36509 | RRJON36355 | |
| RBHWH0032 | not predictec | not predictec | Not predicted in ALGO, TANK, JON |
| RBHWH0114 | RRTNK79095 | RRJON78858 | |
| RBHWH0091 | RRTNK14884 | RRJON14826 | |
| RBHWH0186 | RRTNK59282 | RRJON59028 | |
| RBHWH0159 | not predictec | not predictec | Not predicted in torts |
| RBHWH0159 | not predictec | not predictec | Not predicted in torts |
| RBHWH0159 | not predictec | not predictec | Not predicted in TANK, JON |
| RBHWH0105 | RRTNK79767 | RRJON79542 | |
| RBHWH0211 | RRTNK58958 | RRJON58698 | |
| RBHWH0154 | RRTNK18367 | RRJON18283 | Not predicted in ALGO |
| RBHWH0165 | RRTNK41156 | RRJON40969 | |
| RBHWH0039 | RRTNK19547 | RRJON19447 | Not predicted in LG |
| RBHWH0055 | RRTNK12274 | RRJON12154 | Manually edited ALGO |
| RBHWH0143 | RRTNK68576 | RRJON68333 | |
| RBHWH0173 | RRTNK02201 | RRJON02168 | |
| RBHWH0175 | RRTNK09830 | RRJON09716 | Not predicted in ALGO |
| RBHWH0117 | RRTNK81146 | RRJON80926 | |
| RBHWH0050 | RRTNK48790 | RRJON48549 | |
| RBHWH0006 | RRTNK51777 | RRJON51555 | |
| RBHWH0163 | not predictec | not predictec | Not predicted in torts |
| RBHWH0199 | RRTNK27503 | RRJON27381 | |
| , | RRTNK80810 | RRJON80584 | |
| RBHWH0048 | RRTNK06500 | RRJON06479 | |
| RBHWH0137 | RRTNK13618 | RRJON13538 | |
| RBHWH0059 | RRTNK48321 | RRJON48088 | |
| RBHWH0090 | RRTNK48959 | RRJON48724 | |
| RBHWH0053 | RRTNK44747 | RRJON44554 | |
| RBHWH0173 | RRTNK36701 | RRJON36540 | |
| RBHWH0059 | RRTNK62991 | RRJON62735 | |
| RBHWH005308 | | | Not Predicted |
| 2 | RRTNK20650 | RRJON20577 | |
| RBHWH0013 | RRTNK49601 | RRJON49366 | |

| | | | |
|---|---|---|---|
| RBHWH0126 | RRTNK26107 | RRJON26004 | Manually edited ALGO, TANK, JON, LG |
| | not predicted | not predicted | |
| RBHWH0038 | RRTNK22032 | RRJON21940 | |
| | RRTNK60628 | RRJON60404 | |
| RBHWH0093 | RRTNK64251 | RRJON64010 | |
| RBHWH0040 | RRTNK24632 | RRJON24513 | |
| RBHWH0058 | RRTNK33338 | RRJON33213 | |
| RBHWH0019 | RRTNK76204 | RRJON75986 | |
| RBHWH0130 | RRTNK34567 | RRJON34439 | |
|  | RRTNK34567 | RRJON34439 | Manually edited ALGO, TANK, JON, LG. All torts are ver |
| RBHWH0158 | RRTNK22058 | RRJON21972 | |
| RBHWH0137 | RRTNK13096 | RRJON12998 | Not predicted in ALGO |
| RBHWH0158 | RRTNK32436 | RRJON32312 | |
| RBHWH0145 | RRTNK34700 | RRJON34570 | |
| RBHWH0029 | RRTNK28041 | RRJON27924 | |
| RBHWH0145 | RRTNK34735 | RRJON34600 | |
| RBHWH0053 | RRTNK28309 | RRJON28194 | |
| RBHWH0216 | RRTNK34246 | RRJON34120 | |
| | RRTNK48635 | RRJON48401 | |
| | RRTNK19679 | RRJON19597 | |
| RBHWH0066 | RRTNK14681 | RRJON14623 | |
|  | RRTNK48032 | RRJON47777 | Manually edited TANK, JON. They are significantly sma |
| RBHWH0179 | not predicted | not predicted | Not predicted in torts |
| RBHWH0156 | RRTNK07046 | RRJON06973 | |
| RBHWH0076 | RRTNK16165 | RRJON16068 | |
| RBHWH0157 | RRTNK19682 | RRJON19600 | |
| RBHWH0181 | RRTNK67317 | RRJON67075 | CHCH7 -> CHCHD7 |
| RBHWH0084 | RRTNK23875 | RRJON23766 | |
| RBHWH0016 | RRTNK01287 | RRJON01271 | |
| RBHWH0086 | RRTNK74127 | RRJON73884 | |
| RBHWH0209 | RRTNK29873 | RRJON29700 | |
| RBHWH0178 | RRTNK32390 | RRJON32268 | Manually edited TANK, JON. |
| RBHWH0049 | RRTNK47220 | RRJON46967 | |
| RBHWH0199 | RRTNK29806 | RRJON29677 | |
| RBHWH016820 | | | Not predcited in TANK, JON |
| RBHWH0047 | RRTNK68777 | RRJON68527 | |
| RBHWH0140 | RRTNK56205 | RRJON55935 | |
| RBHWH0173 | RRTNK03447 | RRJON03395 | |
| RBHWH0064 | RRTNK46550 | RRJON46311 | |
| RBHWH0026 | RRTNK70642 | RRJON70391 | |
| RBHWH0170 | RRTNK47306 | RRJON47055 | |
| RBHWH0077 | RRTNK38115 | RRJON37930 | |
| | not predicted | not predicted | Not predicted in torts |
|  | not predicted | not predicted | Not predicted in torts. Lots of CLEC2 but more similar |
| RBHWH0055 | RRTNK10922 | RRJON10793 | |

| | | | |
|---|---|---|---|
| RBHWH0043 | RRTNK78332 | RRJON78090 | |
| RBHWH0014 | RRTNK16772 | RRJON16678 | Not predicted in ALGO |
| RBHWH0003 | RRTNK21608 | RRJON21510 | Manually edited ALGO, TANK, JON, LG |
| RBHWH0058 | RRTNK12323 | RRJON12218 | |
| RBHWH0158 | RRTNK78953 | RRJON78724 | |
| RBHWH0033 | RRTNK01851 | RRJON01836 | |
| | RRTNK37949 | RRJON37769 | Manually edited ALGO |
| RBHWH0097 | RRTNK51635 | RRJON51405 | |
| RBHWH0175 | RRTNK20237 | RRJON20152 | |
| RBHWH0195 | RRTNK32625 | RRJON32498 | |
| RBHWH0138 | RRTNK40261 | RRJON40062 | |
| | RRTNK55141 | RRJON54874 | |
| RBHWH0035 | RRTNK31789 | RRJON31671 | Manually edited ALGO |
| | RRTNK30036 | RRJON29909 | not sure about brandt's bat, or naked mole rat or mouse |
| RBHWH0212 | RRTNK12616 | RRJON12509 | |
| RBHWH0120 | RRTNK83868 | RRJON83626 | |
| RBHWH0169 | not predicted | not predicted | Many proteins with similarity to the GZM/CTH/CMA1 g |
| RBHWH0064 | RRTNK74063 | RRJON73826 | |
| RBHWH0202 | RRTNK12999 | RRJON12909 | |
| RBHWH0052 | RRTNK36006 | RRJON35882 | |
| RBHWH0099 | RRTNK64463 | RRJON64213 | Manually edited TANK, JON, LG. Not predicted in ALGO |
| RBHWH0219 | RRTNK12737 | RRJON12606 | |
| RBHWH0045 | RRTNK19310 | RRJON19220 | Not predicted in LG |
| RBHWH0045 | RRTNK32344 | RRJON32219 | |
| RBHWH0177 | RRTNK69171 | RRJON68915 | |
| RBHWH0066 | RRTNK22732 | RRJON22640 | |
| RBHWH0197 | RRTNK23964 | RRJON23849 | |
| RBHWH0080 | RRTNK50356 | RRJON50139 | |
| RBHWH0128 | RRTNK63150 | RRJON62910 | |
| | RRTNK30143 | RRJON30022 | |
| RBHWH0124 | RRTNK59137 | RRJON58880 | |
| RBHWH0039 | RRTNK61712 | RRJON61460 | |
| RBHWH0062 | RRTNK49422 | RRJON49185 | |
| | RRTNK41660 | RRJON41479 | |
| RBHWH0224 | RRTNK75720 | RRJON75499 | |
| RBHWH0068 | RRTNK31812 | RRJON31694 | |
| RBHWH0055 | RRTNK74910 | RRJON74670 | Not predicted in ALGO |
| | RRTNK19170 | RRJON19080 | |
| RBHWH0172 | RRTNK59610 | RRJON59358 | |
| RBHWH0219 | RRTNK84321 | not predicted | |
| | not predicted | not predicted | Manually edited LG, not predicted in Aldabra. CR like, |
| RBHWH0159 | RRTNK26509 | RRJON26396 | Manually edited ALGO, TANK, JON, LG. Best hit is CR2 |
| RBHWH0016 | RRTNK55147 | RRJON54877 | |
| RBHWH0040 | RRTNK61345 | RRJON61097 | |
| RBHWH0070 | RRTNK70176 | RRJON69934 | |

| | | | |
|---|---|---|---|
| RBHWH0027 | RRTNK40379 | RRJON40189 | |
| RBHWH0019 | RRTNK76148 | RRJON75930 | |
| | RRTNK83215 | RRJON82981 | |
| RBHWH0147 | RRTNK81857 | RRJON81626 | |
| RBHWH0108 | RRTNK30809 | RRJON30694 | Not predicted in ALGO |
| RBHWH0026 | RRTNK06693 | RRJON06654 | Manually edited ALGO, TANK, JON, LG. Torts have a DU |
| RBHWH010308 | | | Not predicted in torts, they have CRTC1 and 2 |
| | RRTNK82471 | RRJON82248 | |
| | RRTNK50891 | RRJON50669 | |
| RBHWH0118 | RRTNK34174 | RRJON34051 | |
| | RRTNK68497 | RRJON68259 | |
| RBHWH0135 | RRTNK26642 | RRJON26533 | |
| RBHWH0107 | RRTNK08417 | RRJON08324 | |
| | RRTNK76635 | RRJON76423 | |
| RBHWH0022 | RRTNK66849 | RRJON66603 | |
| RBHWH0017 | RRTNK79523 | RRJON79296 | |
| | RRTNK07840 | RRJON07744 | |
| RBHWH0029 | RRTNK54277 | RRJON54015 | |
| | RRTNK23475 | RRJON23378 | |
| | RRTNK48060 | RRJON47831 | |
| RBHWH0188 | RRTNK46498 | RRJON46257 | |
| | RRTNK15146 | RRJON15101 | |
| | RRTNK44438 | RRJON44378 | Manually edited ALGO, TANK |
| RBHWH0153 | RRTNK58760 | RRJON58503 | |
| RBHWH0120 | RRTNK18706 | RRJON18619 | Manually edited ALGO, TANK, JON, LG |
| RBHWH0016 | RRTNK45334 | RRJON45138 | |
| | RRTNK75806 | RRJON75582 | |
| RBHWH0185 | RRTNK75356 | not predicted | Not predicted in JON. More similar to other CXCL gene |
| RBHWH0185 | RRTNK75371 | RRJON75146 | |
| RBHWH0141 | RRTNK57199 | RRJON56963 | |
| RBHWH0038 | not predicted | not predicted | Not predicted in TANK, JON |
| RBHWH0086 | not predicted | not predicted | Not predicted in torts |
| | not predicted | not predicted | Not predicted in torts |
| RBHWH0195 | not predicted | not predicted | Not predicted in torts |
| RBHWH0195 | RRTNK75788 | RRJON75567 | |
| RBHWH0185 | not predicted | not predicted | Not predicted in torts |
| | RRTNK09854 | RRJON09743 | |
| RBHWH0083 | RRTNK00241 | RRJON00229 | |
| RBHWH0105 | RRTNK30594 | RRJON30479 | |
| RBHWH0032 | RRTNK36772 | RRJON36608 | |
| RBHWH0075 | RRTNK80111 | RRJON79883 | |
| RBHWH0137 | RRTNK51891 | RRJON51666 | |
| RBHWH0206 | RRTNK33300 | RRJON33179 | |
| RBHWH0177 | RRTNK77951 | RRJON77721 | |
| RBHWH0038 | RRTNK22076 | RRJON21984 | |

| | | | |
|---|---|---|---|
| | not predicted | not predicted | |
| RBHWH0160 | RRTNK84374 | RRJON84115 | |
| RBHWH0045 | RRTNK32741 | RRJON32614 | |
| RBHWH0223 | RRTNK48793 | RRJON48557 | |
| RBHWH0157 | RRTNK20380 | RRJON20299 | |
| RBHWH0142 | RRTNK00586 | RRJON00580 | Manually edited ALGO, TANK, JON, LG |
| RBHWH0030 | RRTNK24048 | RRJON23937 | |
| RBHWH0172 | RRTNK22238 | RRJON22131 | |
| RBHWH0163 | RRTNK30308 | RRJON30190 | |
| RBHWH0167 | RRTNK24072 | RRJON23958 | |
| RBHWH0123 | RRTNK05855 | RRJON05825 | |
| RBHWH0174 | RRTNK41479 | RRJON41298 | |
| RBHWH0041 | RRTNK61686 | RRJON61434 | |
| RBHWH0038 | RRTNK21585 | RRJON21483 | |
| RBHWH0159 | RRTNK36232 | RRJON36091 | |
| | RRTNK84975 | RRJON84698 | |
| RBHWH0043 | RRTNK47379 | RRJON47152 | |
| RBHWH0050 | RRTNK82589 | RRJON82367 | |
| RBHWH0083 | RRTNK70225 | RRJON69982 | |
| RBHWH0065 | RRTNK55196 | RRJON54926 | |
| RBHWH0196 | RRTNK29689 | RRJON29558 | |
| RBHWH0032 | RRTNK36755 | RRJON36594 | |
| RBHWH0195 | RRTNK59364 | RRJON59112 | |
| | not predicted | not predicted | Not predicted in torts |
| | not predicted | not predicted | Not predicted in torts |
| | not predicted | not predicted | Not predicted in torts |
| | not predicted | not predicted | Not predicted in torts |
| | not predicted | not predicted | Not predicted in torts |
| | not predicted | not predicted | Not predicted in torts |
| | RRTNK22701 | RRJON22609 | |
| RBHWH0204 | RRTNK22887 | not predicted | Not predicted in ALGO, JON, LG. TANK is an ok hit |
| | not predicted | not predicted | Not predicted in torts |
| | not predicted | not predicted | Not predicted in torts |
| | not predicted | not predicted | Not predicted in torts |
| | not predicted | not predicted | Not predicted in torts |
| | not predicted | not predicted | Not predicted in torts |
| | not predicted | not predicted | Not predicted in torts |
| | not predicted | not predicted | Not predicted in torts |
| | not predicted | not predicted | Not predicted in torts |
| | not predicted | not predicted | Not predicted in torts |
| | not predicted | not predicted | Not predicted in torts |
| | not predicted | not predicted | Not predicted in torts |
| | not predicted | not predicted | Not predicted in torts |

| | | | | |
|---|---|---|---|---|
| l | | not predicted | not predicted | Not predicted in torts |
| l | | not predicted | not predicted | Not predicted in torts |
| l | | not predicted | not predicted | Not predicted in torts |
| l | | not predicted | not predicted | Not predicted in torts |
| l | | not predicted | not predicted | Not predicted in torts |
| l | | not predicted | not predicted | Not predicted in torts |
| l | | not predicted | not predicted | Not predicted in torts |
| l | | not predicted | not predicted | Not predicted in torts |
| l | | not predicted | not predicted | Not predicted in torts |
| l | | not predicted | not predicted | Not predicted in torts |
| l | | not predicted | not predicted | Not predicted in torts |
| l | | not predicted | not predicted | Not predicted in torts |
| l | | not predicted | not predicted | Not predicted in torts |
| l | | not predicted | not predicted | Not predicted in torts |
| l | | not predicted | not predicted | Not predicted in torts |
| l | | not predicted | not predicted | Not predicted in torts |
| l | | not predicted | not predicted | Not predicted in torts |
| l | | not predicted | not predicted | Not predicted in torts |
| l | | not predicted | not predicted | Not predicted in torts |
| l | | not predicted | not predicted | Not predicted in torts |
| l | | not predicted | not predicted | Not predicted in torts |
| l | | not predicted | not predicted | Not predicted in torts |
| l | | not predicted | not predicted | Not predicted in torts |
| l | | not predicted | RRJON22766 | Manually edited JON, not predicted in other torts |
| l | | not predicted | not predicted | Not predicted in torts |
| | RBHWH0083 | RRTNK52092 | RRJON51857 | |
| | RBHWH0075 | RRTNK80042 | RRJON79812 | |
| | RBHWH0020 | RRTNK42730 | RRJON42529 | |
| | RBHWH0107 | RRTNK67897 | RRJON67638 | |
| | RBHWH0085 | RRTNK20074 | RRJON19993 | |
| | RBHWH0189 | RRTNK60458 | RRJON60227 | |
| | RBHWH0166 | RRTNK28096 | RRJON27984 | Not predicted in ALGO |
| | RBHWH0104 | RRTNK51693 | RRJON51465 | |
| | RBHWH0141 | RRTNK74624 | RRJON74392 | |
| | RBHWH0178 | RRTNK48178 | RRJON47950 | |
| 3 | | RRTNK06927 | RRJON06875 | |
| | RBHWH0080 | RRTNK77828 | RRJON77592 | |
| | RBHWH0158 | RRTNK75294 | RRJON75071 | |
| | RBHWH0011 | RRTNK66743 | RRJON66497 | |
| | RBHWH0027 | RRTNK39821 | RRJON39636 | |
| | RBHWH0136 | RRTNK12578 | RRJON12473 | Manually edited ALGO, TANK, JON |
| | RBHWH0010 | RRTNK06522 | RRJON06496 | Manually edited ALGO, TANK, JON, LG |
| . | | RRTNK34153 | RRJON34030 | Manually edited ALGO. Truncated compared to others |
| 3 | | RRTNK60622 | RRJON60389 | Manually edited ALGO, TANK, JON |
| | RBHWH0037 | RRTNK45849 | RRJON45650 | |

| | | | | |
|---|---|---|---|---|
| | RBHWH0066 | RRTNK22290 | RRJON22192 | |
| | RBHWH0201 | RRTNK34512 | RRJON34387 | |
| 6 | | RRTNK45230 | RRJON45029 | |
| | RBHWH0190 | RRTNK79656 | RRJON79433 | |
| | RBHWH0151 | RRTNK29534 | RRJON29401 | |
| | RBHWH0194 | RRTNK31945 | RRJON31836 | |
| | RBHWH0005 | RRTNK36083 | RRJON35959 | |
| | RBHWH0206 | not predicted | not predicted | |
| 1 | | RRTNK35829 | RRJON35707 | Manually edited ALGO, TANK, JON |
| 6 | | RRTNK35829 | RRJON35707 | |
| | RBHWH0157 | RRTNK19993 | RRJON19905 | |
| | RBHWH0206 | RRTNK56462 | RRJON56198 | |
| 0 | | RRTNK35526 | RRJON35420 | |
| | RBHWH0140 | RRTNK50659 | RRJON50442 | |
| | RBHWH0160 | RRTNK53236 | RRJON52981 | |
| | RBHWH0144 | RRTNK15954 | RRJON15856 | |
| 7 | | RRTNK20360 | RRJON20276 | |
| | RBHWH022064 | | | |
| | RBHWH0071 | RRTNK71676 | RRJON71449 | |
| | RBHWH0080 | RRTNK37599 | RRJON37436 | |
| | RBHWH0112 | not predicted | not predicted | Not predicted in torts |
| | RBHWH0121 | RRTNK70524 | RRJON70271 | |
| | RBHWH0209 | RRTNK50374 | RRJON50159 | |
| | RBHWH0134 | RRTNK77925 | RRJON77692 | |
| | RBHWH0188 | RRTNK26894 | RRJON26783 | |
| | RBHWH0006 | RRTNK65848 | RRJON65593 | |
| | RBHWH0078 | RRTNK62413 | RRJON62145 | |
| | RBHWH0005 | RRTNK36282 | RRJON36139 | |
| | RBHWH0061 | RRTNK82760 | RRJON82529 | |
| | RBHWH0179 | RRTNK76237 | RRJON76019 | |
| | RBHWH0110 | RRTNK65268 | RRJON65016 | |
| | RBHWH0160 | RRTNK69851 | RRJON69616 | |
| | RBHWH0061 | RRTNK72516 | RRJON72295 | |
| 3 | | RRTNK41020 | RRJON40838 | |
| 6 | | RRTNK63017 | RRJON62776 | |
| | RBHWH0184 | RRTNK84215 | RRJON83958 | |
| | RBHWH0162 | RRTNK45676 | RRJON45491 | |
| | RBHWH0023 | RRTNK27894 | RRJON27778 | |
| | RBHWH0026 | RRTNK06039 | RRJON06007 | |
| | RBHWH0092 | RRTNK59439 | RRJON59187 | |
| 3 | | RRTNK40741 | RRJON40552 | Manually edited TANK, JON |
| | RBHWH0062 | RRTNK00926 | RRJON00915 | |
| | RBHWH000379 | | | |
| | RBHWH0008 | RRTNK50382 | RRJON50165 | |
| | RBHWH0186 | RRTNK36127 | RRJON35997 | |

| | | | | |
|---|---|---|---|---|
| | RBHWH0153 | RRTNK76269 | RRJON76051 | |
| | RBHWH0066 | RRTNK53206 | RRJON52947 | |
| | RBHWH001921 | | | |
| | RBHWH0003 | RRTNK80607 | RRJON80379 | |
| | RBHWH0064 | RRTNK46377 | RRJON46139 | |
| | RBHWH0187 | RRTNK26622 | RRJON26502 | |
| | RBHWH0022 | RRTNK57573 | RRJON57338 | |
| | RBHWH0087 | RRTNK42390 | RRJON42194 | |
| i | | RRTNK84263 | RRJON84006 | |
| | RBHWH0128 | RRTNK67988 | RRJON67729 | |
| | RBHWH0051 | RRTNK77390 | RRJON77172 | |
| | RBHWH0087 | RRTNK50814 | RRJON50600 | |
| | RBHWH0009 | RRTNK65700 | RRJON65451 | |
| | RBHWH0185 | RRTNK26018 | RRJON25924 | |
| | RBHWH0205 | not predicted | not predicted | Not predicted in torts |
| | RBHWH0169 | RRTNK80266 | RRJON80036 | |
| i | | RRTNK22357 | RRJON22275 | |
| | RBHWH0094 | RRTNK72589 | RRJON72368 | |
| | RBHWH0101 | not predicted | not predicted | |
| | RBHWH0127 | RRTNK26414 | RRJON26298 | Manually edited ALGO, TANK, JON |
| | RBHWH0162 | RRTNK64865 | RRJON64627 | |
| | RBHWH0039 | RRTNK20060 | RRJON19970 | |
| . | | RRTNK56241 | RRJON55977 | Manually edited ALGO, LG |
| : | | | RRJON71514 | |
| | RBHWH0226 | RRTNK17842 | RRJON17762 | |
| | RBHWH0155 | RRTNK16847 | RRJON16749 | |
| | RBHWH0104 | RRTNK14383 | RRJON14327 | |
| | RBHWH0053 | RRTNK47615 | RRJON47359 | |
| | RBHWH0013 | RRTNK83798 | RRJON83553 | |
| | RBHWH0042 | RRTNK57404 | RRJON57170 | |
| | RBHWH0113 | RRTNK66254 | RRJON66004 | |
| | RBHWH0159 | RRTNK75835 | RRJON75611 | |
| | RBHWH0130 | RRTNK82317 | RRJON82092 | Manually edited ALGO, TANK, JON, LG |
| | RBHWH0045 | RRTNK63520 | RRJON63276 | |
| | RBHWH0065 | RRTNK55100 | RRJON54826 | |
| | RBHWH021361 | | | |
| | RBHWH0129 | RRTNK71907 | RRJON71687 | |
| . | | | | |
| | RBHWH0064 | RRTNK73947 | RRJON73712 | |
| | RBHWH0073 | RRTNK25713 | RRJON25603 | |
| | RBHWH0055 | RRTNK58179 | RRJON57919 | Manually edited ALGO, TANK, JON, LG |
| | RBHWH0152 | RRTNK29372 | RRJON29234 | Manually edited ALGO, TANK, JON, LG |
| | RBHWH0110 | RRTNK51205 | RRJON50997 | |
| | RBHWH0142 | RRTNK75994 | RRJON75761 | |
| | RBHWH0016 | RRTNK38480 | RRJON38306 | |

| | | | |
|---|---|---|---|
| RBHWH0111 | RRTNK76866 | RRJON76658 | |
| RBHWH0028 | RRTNK52915 | RRJON52675 | |
| RBHWH0148 | RRTNK77212 | RRJON77005 | |
| RBHWH0104 | RRTNK60743 | RRJON60519 | Manually edited ALGO, TANK, JON, LG |
| RBHWH0210 | RRTNK35710 | RRJON35586 | |
| RBHWH0137 | RRTNK72834 | RRJON72608 | |
| RBHWH0158 | RRTNK83677 | RRJON83431 | |
| RBHWH000293 | | | |
| RBHWH0132 | RRTNK63385 | RRJON63138 | |
| RBHWH0215 | RRTNK63400 | RRJON63156 | |
| | RRTNK83680 | RRJON83434 | |
| RBHWH0143 | RRTNK58560 | RRJON58308 | Manually edited ALGO. ALGO is a weird fusion |
| RBHWH0174 | RRTNK42050 | RRJON41851 | |
| RBHWH0154 | RRTNK78972 | RRJON78736 | |
| RBHWH0214 | RRTNK73307 | RRJON73078 | |
| | RRTNK32928 | RRJON32800 | |
| RBHWH0164 | RRTNK79454 | RRJON79222 | |
| RBHWH0115 | RRTNK21775 | RRJON21677 | |
| RBHWH0142 | RRTNK44546 | RRJON44350 | |
| RBHWH0012 | RRTNK82979 | RRJON82741 | |
| RBHWH0113 | RRTNK49001 | RRJON48766 | |
| RBHWH0004 | RRTNK59860 | RRJON59612 | |
| RBHWH0109 | RRTNK32400 | RRJON32275 | |
| RBHWH0110 | RRTNK57237 | RRJON57003 | |
| RBHWH0196 | RRTNK75926 | | |
| RBHWH0156 | RRTNK27036 | RRJON26913 | |
| RBHWH0056 | RRTNK81325 | RRJON81107 | |
| RBHWH0204 | RRTNK78553 | RRJON78301 | |
| RBHWH0084 | RRTNK41456 | RRJON41281 | |
| RBHWH0115 | RRTNK55695 | RRJON55428 | |
| RBHWH0101 | RRTNK66516 | RRJON66257 | |
| RBHWH0194 | RRTNK81233 | RRJON81018 | |
| RBHWH0067 | RRTNK42794 | RRJON42590 | |
| RBHWH0030 | RRTNK47979 | RRJON47744 | |
| RBHWH0071 | RRTNK62103 | RRJON61841 | Manually edited TANK, JON |
| | not predicted | not predicted | Not predicted in torts |
| | RRTNK21916 | RRJON21815 | |
| RBHWH0117 | not predicted | not predicted | Not predicted in torts |
| RBHWH0196 | not predicted | not predicted | Not predicted in torts |
| | not predicted | not predicted | Not predicted in torts |
| | not predicted | not predicted | Not predicted in torts |
| | | | Not predicted in torts |
| | not predicted | not predicted | Not predicted in torts |
| | not predicted | not predicted | Not predicted in torts |
| | RRTNK17475 | RRJON17383 | Manually edited ALGO, TANK, JON. FCN1/2/3 are very |

| | | | |
|---|---|---|---|
| | RRTNK17537 | RRJON17434 | Manually edited ALGO, TANK, JON, LG. Hard to distingu |
| RBHWH0019 | RRTNK22384 | RRJON22290 | Not predicted in ALGO, LG. FCN1/2/3 hard to disambig |
| RBHWH015480 | | | Not predicted |
| RBHWH0154 | not predicted | not predicted | Not predicted in torts. FCRL3 most common |
| RBHWH0174 | RRTNK41900 | RRJON41712 | |
| | RRTNK56813 | RRJON56578 | |
| | RRTNK07093 | RRJON07011 | |
| RBHWH0224 | RRTNK61658 | RRJON61408 | Not predicted in ALGO |
| RBHWH0224 | RRTNK61666 | RRJON61417 | |
| RBHWH0143 | RRTNK58395 | RRJON58145 | |
| | RRTNK10555 | RRJON10433 | |
| RBHWH0168 | RRTNK41300 | RRJON41124 | |
| RBHWH0178 | RRTNK74537 | RRJON74305 | Manually edited ALGO |
| RBHWH0070 | RRTNK30299 | RRJON30178 | Manually edited ALGO, TANK, JON |
| RBHWH0054 | RRTNK31245 | RRJON31131 | |
| RBHWH0083 | RRTNK70155 | RRJON69911 | Manually edited TANK, JON, LG |
| RBHWH0111 | RRTNK23797 | RRJON23688 | |
| | RRTNK74368 | RRJON74133 | |
| RBHWH0143 | RRTNK51212 | RRJON50990 | |
| RBHWH0200 | RRTNK53306 | RRJON53057 | |
| RBHWH0073 | RRTNK25878 | RRJON25782 | |
| RBHWH0162 | RRTNK08267 | RRJON08135 | |
| RBHWH0095 | RRTNK84764 | RRJON84507 | |
| RBHWH0203 | RRTNK68440 | RRJON68205 | |
| | RRTNK71235 | RRJON70992 | |
| | RRTNK16245 | RRJON16141 | |
| RBHWH0102 | RRTNK67538 | RRJON67283 | |
| | RRTNK78901 | RRJON78665 | |
| RBHWH0038 | RRTNK70656 | RRJON70405 | |
| | RRTNK82161 | RRJON81929 | |
| RBHWH0137 | RRTNK52137 | RRJON51894 | |
| RBHWH0102 | RRTNK54634 | RRJON54371 | |
| RBHWH0212 | RRTNK84503 | RRJON84241 | |
| RBHWH0055 | RRTNK61758 | RRJON61509 | |
| RBHWH0084 | RRTNK03192 | RRJON03153 | |
| RBHWH011990 | | | |
| RBHWH0176 | RRTNK72562 | RRJON72345 | Manually edited ALGO, TANK, JON, LG |
| RBHWH0166 | not predicted | not predicted | Not predicted in torts |
| RBHWH0185 | RRTNK26264 | RRJON26158 | |
| RBHWH0154 | RRTNK81213 | RRJON80993 | |
| RBHWH0066 | RRTNK15253 | RRJON15189 | |
| RBHWH0131 | RRTNK35692 | RRJON35571 | Manually edited ALGO, TANK, JON, LG |
| RBHWH0007 | RRTNK13146 | RRJON13039 | |

| | | | |
|---|---|---|---|
| RBHWH0170 | RRTNK69687 | RRJON69442 | Not predicted in ALGO |
| RBHWH0103 | RRTNK62609 | RRJON62354 | |
| RBHWH0016 | RRTNK55190 | RRJON54929 | |
| RBHWH0129 | RRTNK24751 | RRJON24621 | |
| RBHWH0131 | RRTNK81040 | RRJON80814 | |
| RBHWH0124 | RRTNK31440 | RRJON31339 | |
| | RRTNK11041 | RRJON10919 | |
| RBHWH0079 | RRTNK50736 | RRJON50519 | |
| RBHWH0114 | RRTNK79183 | RRJON78952 | |
| RBHWH0197 | not predictec | not predicted | |
| | RRTNK83629 | RRJON83382 | |
| RBHWH0016 | RHSPN22410 | RRJON47622 | Manually edited JON, TANK. Missing SH3 domain at sta |
| RBHWH0165 | RRTNK03714 | RRJON03662 | |
| RBHWH0099 | RRTNK69320 | RRJON69057 | Manually edited ALGO. Gene missing the GATA domair |
| RBHWH0006 | RRTNK25804 | RRJON25698 | Not predicted in ALGO |
| RBHWH0172 | RRTNK22293 | RRJON22202 | |
| RBHWH0124 | RRTNK81314 | RRJON81092 | |
| RBHWH0153 | RRTNK58630 | RRJON58372 | |
| RBHWH0128 | RRTNK74667 | RRJON74437 | |
| RBHWH0080 | RRTNK77856 | RRJON77617 | |
| RBHWH0205 | RRTNK48122 | RRJON47890 | |
| RBHWH0053 | RRTNK41288 | RRJON41115 | |
| RBHWH0192 | RRTNK22608 | RRJON22510 | |
| RBHWH0035 | RRTNK31982 | RRJON31868 | |
| RBHWH0201 | RRTNK73686 | RRJON73449 | Manually edited ALGO. Is a fusion protein that is also C |
| RBHWH0038 | RRTNK34912 | RRJON34786 | |
| RBHWH0203 | RRTNK40652 | RRJON40468 | |
| RBHWH0175 | RRTNK09895 | RRJON09794 | |
| | RRTNK31919 | RRJON31811 | |
| | RRTNK71888 | RRJON71666 | |
| RBHWH0004 | RRTNK70744 | RRJON70492 | |
| RBHWH0164 | RRTNK63261 | RRJON63017 | |
| RBHWH0090 | RRTNK68044 | RRJON67785 | |
| RBHWH0151 | RRTNK24137 | RRJON24026 | |
| RBHWH0108 | RRTNK51369 | RRJON51150 | |
| RBHWH0172 | RRTNK60607 | RRJON60374 | Manually edited ALGO, TANK, JON, LG |
| | RRTNK16683 | RRJON16593 | |
| RBHWH0108 | RRTNK42303 | RRJON42103 | Manually edited LG |
| RBHWH0105 | RRTNK22884 | RRJON22795 | Manually edited ALGO, TANK, JON, LG |
| | not predictec | not predictec | Not predicted in torts |
| RBHWH0086 | RRTNK74090 | RRJON73853 | |

| | | | |
|---|---|---|---|
| RBHWH0181 | RRTNK63761 | RRJON63512 | |
| | RRTNK12526 | RRJON12429 | Manually edited ALGO, TANK, JON, LG |
| | RRTNK18073 | RRJON17977 | |
| RBHWH0078 | RRTNK62288 | RRJON62026 | Manually edited ALGO, TANK, JON. Tank and Jon are mi |
| RBHWH0102 | RRTNK53814 | RRJON53572 | |
| RBHWH0120 | RRTNK83883 | RRJON83641 | |
| RBHWH0118 | RRTNK72944 | RRJON72709 | |
| RBHWH0062 | RRTNK49611 | RRJON49378 | |
| RBHWH0059 | RRTNK59985 | RRJON59731 | |
| RBHWH0174 | RRTNK19210 | RRJON19109 | |
| RBHWH0066 | RRTNK13639 | RRJON13559 | |
| RBHWH011200 | | | |
| RBHWH011199 | | | |
| RBHWH0099 | RRTNK23134 | RRJON23035 | |
| RBHWH0196 | RRTNK67934 | RRJON67677 | |
| RBHWH0108 | RRTNK30380 | RRJON30259 | |
| | RRTNK69010 | RRJON68759 | |
| RBHWH0028 | RRTNK46127 | RRJON45928 | Manually edited TANK, JON. |
| RBHWH0019 | RRTNK64615 | RRJON64385 | |
| RBHWH0113 | RRTNK10606 | RRJON10520 | |
| | RRTNK41925 | RRJON41739 | |
| | RRTNK03330 | RRJON03284 | |
| RBHWH0082 | RRTNK77079 | RRJON76874 | |
| RBHWH003084 | | | |
| | RRTNK47880 | RRJON47639 | |
| | RRTNK47900 | RRJON47663 | |
| RBHWH0030 | not predictec | not predictec | probably not predicted though difficult to dis-entangle |
| RBHWH0078 | RRTNK18564 | RRJON18471 | |
| | RRTNK31319 | RRJON31217 | |
| | RRTNK71633 | RRJON71400 | |
| RBHWH0036 | RRTNK49689 | RRJON49453 | |
| RBHWH0093 | RRTNK62349 | RRJON62084 | |
| RBHWH0014 | RRTNK19781 | RRJON19699 | |
| RBHWH0162 | RRTNK13265 | RRJON13146 | |
| RBHWH0200 | RRTNK72751 | RRJON72529 | |
| RBHWH0218 | RRTNK61128 | RRJON60889 | |
| RBHWH0196 | RRTNK67958 | RRJON67699 | |
| RBHWH0168 | RRTNK44420 | RRJON44212 | Manually edited ALGO, TANK, JON, LG |
| | RRTNK44646 | RRJON44236 | Manually edited LG |
| RBHWH0196 | not predictec | not predictec | may not be predicted but it's difficult to tell because of |
| RBHWH0066 | RRTNK11804 | RRJON11684 | |
| RBHWH0032 | RRTNK36947 | RRJON36788 | |
| RBHWH0137 | RRTNK06624 | RRJON06573 | |
| RBHWH0137 | RRTNK06624 | RRJON06573 | |

| | | | |
|---|---|---|---|
| RBHWH0060 | RRTNK42519 | RRJON42321 | |
| | not predicted | not predicted | Not predicted in ALGO, TANK, JON |
| RBHWH0071 | RRTNK77550 | RRJON77331 | |
| RBHWH0147 | RRTNK69528 | RRJON69273 | Manually edited torts |
| RBHWH0078 | RRTNK59032 | RRJON58784 | |
| | RRTNK08070 | RRJON07991 | |
| RBHWH0117 | RRTNK17012 | RRJON16922 | |
| RBHWH0145 | RRTNK45480 | RRJON45295 | |
| RBHWH0184 | RRTNK64884 | RRJON64646 | |
| RBHWH0182 | RRTNK63101 | RRJON62858 | |
| RBHWH0123 | RRTNK58373 | RRJON58123 | |
| | RRTNK60350 | RRJON60120 | |
| RBHWH0151 | RRTNK22464 | RRJON22368 | |
| RBHWH0165 | RRTNK03740 | RRJON03683 | |
| RBHWH0187 | RRTNK26916 | RRJON26792 | |
| | RRTNK47326 | RRJON47090 | |
| | RRTNK56820 | RRJON56585 | |
| RBHWH0221 | RRTNK37992 | RRJON37817 | |
| RBHWH0044 | RRTNK52009 | RRJON51774 | |
| RBHWH0081 | RRTNK59734 | RRJON59483 | |
| RBHWH0207 | RRTNK74268 | RRJON74029 | |
| RBHWH0121 | RRTNK19718 | RRJON19638 | |
| RBHWH0199 | RRTNK71664 | RRJON71440 | Manually edited ALGO, TANK, JON, LG |
| RBHWH0121 | not predicted | not predicted | |
| RBHWH0171 | RRTNK77206 | RRJON76999 | |
| RBHWH0194 | RRTNK11861 | RRJON11753 | |
| RBHWH0111 | RRTNK02383 | RRJON02337 | |
| RBHWH0019 | RRTNK02791 | RDJON03006 | |
| | | | Not predicted in torts |
| RBHWH0030 | not predicted | not predicted | |
| RBHWH0030 | not predicted | not predicted | |
| RBHWH0030 | RRTNK02892 | RRJON02095 | |
| RBHWH0003 | RRTNK54404 | RRJON54142 | Manually edited ALGO, TANK |
| RBHWH0172 | RRTNK59641 | RRJON59392 | |
| RBHWH0011 | RRTNK67712 | RRJON67459 | |
| RBHWH0057 | RRTNK33156 | RRJON33028 | |
| RBHWH0152 | RRTNK55039 | RRJON54772 | Manually edited ALGO, LG |
| RBHWH0161 | RRTNK74448 | RRJON74211 | |
| RBHWH0210 | RRTNK18016 | RRJON17929 | |
| RBHWH0197 | RRTNK54751 | RRJON54482 | |
| RBHWH0197 | RRTNK54713 | RRJON54441 | |
| RBHWH0197 | RRTNK54710 | RRJON54438 | |
| RBHWH0197 | RRTNK54692 | RRJON54432 | |
| | RRTNK54748 | RRJON54479 | |
| RBHWH0197 | RRTNK54745 | RRJON54473 | |

| | | | |
|---|---|---|---|
| RBHWH0197 | RRTNK54736 | RRJON54464 | |
| RBHWH0197 | RRTNK54730 | RRJON54458 | |
| | RRTNK54727 | RRJON54455 | |
| | RRTNK54719 | RRJON54452 | |
| RBHWH0197 | RRTNK54716 | RRJON54449 | |
| RBHWH0097 | RRTNK24129 | RRJON24015 | |
| RBHWH0097 | RRTNK23970 | RRJON23852 | |
| | RRTNK24117 | RRJON24000 | |
| RBHWH0097 | RRTNK24105 | RRJON23997 | |
| RBHWH0097 | RRTNK24085 | RRJON23968 | |
| RBHWH0097 | RRTNK24063 | RRJON23946 | |
| RBHWH0097 | RRTNK24051 | RRJON23940 | |
| | RRTNK24032 | RRJON23915 | |
| | RRTNK24026 | RRJON23904 | |
| RBHWH0097 | RRTNK24006 | RRJON23896 | |
| RBHWH0046 | RRTNK25172 | RRJON25047 | |
| RBHWH0046 | RRTNK25155 | RRJON25033 | |
| RBHWH0046 | RRTNK25127 | RRJON25007 | |
| RBHWH0046 | RRTNK25108 | RRJON24990 | |
| RBHWH0046 | RRTNK25252 | RRJON25129 | |
| RBHWH0046 | RRTNK25238 | RRJON25115 | |
| RBHWH0046 | RRTNK25232 | RRJON25112 | |
| RBHWH0046 | RRTNK25202 | RRJON25076 | |
| RBHWH0046 | RRTNK25187 | RRJON25061 | |
| RBHWH0073 | RRTNK46310 | RRJON46087 | |
| RBHWH0073 | RRTNK46266 | RRJON46044 | |
| | RRTNK46248 | RRJON46028 | |
| RBHWH0073 | RRTNK46245 | RRJON46025 | |
| RBHWH0073 | RRTNK46242 | RRJON46022 | |
| RBHWH0073 | RRTNK46304 | RRJON46070 | |
| RBHWH0073 | RRTNK46280 | RRJON46061 | |
| RBHWH0073 | RRTNK46274 | RRJON46052 | |
| | RRTNK46269 | RRJON46047 | |
| RBHWH0218 | RRTNK46400 | RRJON46162 | |
| RBHWH0151 | RRTNK38624 | RRJON38451 | |
| RBHWH0049 | RRTNK44741 | RRJON44545 | |
| RBHWH0004 | RRTNK63752 | RRJON63503 | |
| | not predictec | not predictec | Not predicted in torts |
| RBHWH0100 | RRTNK50341 | RRJON50121 | |
| RBHWH0142 | RRTNK41374 | RRJON41196 | |
| RBHWH0199 | RRTNK05031 | RRJON05000 | |
| RBHWH0075 | RRTNK80051 | RRJON79821 | |
| RBHWH0156 | RRTNK65030 | RRJON64790 | |
| RBHWH0052 | RRTNK21363 | RRJON21264 | |
| RBHWH0035 | RRTNK08969 | RRJON08884 | |

| | | | | |
|---|---|---|---|---|
| | RBHWH0182 | RRTNK75991 | RRJON75755 | |
| | RBHWH0000 | RRTNK78364 | RRJON78120 | |
| | RBHWH0047 | RRTNK68907 | RRJON68658 | |
| | RBHWH0081 | RRTNK26645 | RRJON26539 | |
| | RBHWH0076 | RRTNK10881 | RRJON10755 | |
| | RBHWH0204 | RRTNK81411 | RRJON81192 | |
| | RBHWH0163 | RRTNK30273 | RRJON30148 | |
| | RBHWH003125 | | | |
| | RBHWH003128 | | | |
| | RBHWH0123 | RRTNK76802 | RRJON76590 | |
| | RBHWH0176 | RRTNK65149 | RRJON64901 | |
| | RBHWH0046 | RRTNK34629 | RRJON34496 | |
| | RBHWH0220 | RRTNK30693 | RRJON30575 | |
| | RBHWH0121 | RRTNK59970 | RRJON59719 | |
| | RBHWH0174 | RRTNK17690 | RRJON17604 | |
| | RBHWH0187 | RRTNK24417 | RRJON24295 | |
| | RBHWH0116 | RRTNK33579 | RRJON33457 | |
| | RBHWH0188 | RRTNK68032 | RRJON67782 | |
| | RBHWH012259 | | | |
| | RBHWH0181 | RRTNK34217 | RRJON34091 | |
| | RBHWH0156 | RRTNK79709 | RRJON79486 | |
| | RBHWH0044 | RRTNK52397 | RRJON52153 | |
| 2 | | RRTNK84703 | RRJON84444 | |
| | RBHWH0123 | RRTNK01585 | RRJON01578 | |
| | RBHWH0054 | RRTNK33452 | RRJON33334 | |
| 3 | | RRTNK33561 | RRJON33439 | |
| | RBHWH0032 | RRTNK36913 | RRJON36748 | |
| 3 | | RRTNK14032 | RRJON13968 | |
| | RBHWH0029 | RRTNK54294 | RRJON54035 | |
| | RBHWH0186 | RRTNK55401 | RRJON55137 | |
| | RBHWH0198 | RRTNK04568 | RRJON04536 | Manually edited LG |
| | RBHWH0060 | RRTNK41755 | RRJON41571 | |
| d | | not predicted | not predicted | Not predicted in torts |
| d | | not predicted | not predicted | Not predicted in torts |
| | RBHWH0026 | RRTNK04776 | RRJON04758 | |
| | RBHWH0211 | RRTNK28937 | RRJON28812 | |
| | RBHWH0019 | RRTNK22735 | RRJON22643 | Not predicted in ALGO |
| | RBHWH0062 | RRTNK40732 | RRJON40543 | Manually edited ALGO. Is a fusion with KATA2A |
| d | | not predicted | not predicted | Not predicted in torts. All similar proteins are closer to |
| d | | not predicted | not predicted | Not predicted in torts. All similar proteins are closer to |
| d | | not predicted | not predicted | Not predicted in torts. All similar proteins are closer to |
| | RBHWH0211 | RRTNK41338 | RRJON41159 | |
| - | | RRTNK43136 | RRJON42921 | |
| d | | not predicted | not predicted | Not predicted in torts. Proteins more similar to IFITM1 |

| | | | |
|---|---|---|---|
| I | not predictec | not predictec | Not predicted in torts. All proteins more similar to IFITI |
| I | not predictec | not predictec | There are a few interferon alpha/betas doesn't appear a |
| I | not predictec | not predictec | Not predicted in torts. Proteins more similar to IFNE/K |
| I | not predictec | not predictec | Not predicted in torts. Proteins more similar to IFNB/E |
| I | not predictec | not predictec | Not predicted in torts. Proteins more similar to IFNE/K |
| I | not predictec | not predictec | Not predicted in torts. Proteins more similar to IFNE/K |
| I | not predictec | not predictec | Not predicted in torts. Proteins more similar to IFNE/K |
| I | not predictec | not predictec | Not predicted in torts. Proteins more similar to IFNE/K |
| I | not predictec | not predictec | Not predicted in torts. Proteins more similar to IFNE/K |
| I | not predictec | not predictec | Not predicted in torts. Proteins more similar to IFNB/E |
| I | not predictec | not predictec | Not predicted in torts. Proteins more similar to IFNE/K |
| I | not predictec | not predictec | Not predicted in torts. Proteins more similar to IFNE/K |
| I | not predictec | not predictec | Not predicted in torts |
| I | not predictec | not predictec | Not predicted in torts. Proteins more similar to IFNE/K |
| RBHWH0012 | RRTNK81961 | RRJON81728 | |
| RBHWH0012 | RRTNK81952 | RRJON81717 | |
| ) | RRTNK40553 | RRJON4036C | Manually edited ALGO, LG |
| RBHWH013C | RRTNK40365 | RRJON40177 | |
| RBHWH0143 | RRTNK57053 | RRJON5681E | Not predicted in ALGO |
| RBHWH0144 | RRTNK69141 | RRJON68888 | |
| RBHWH0012 | RRTNK81974 | RRJON81744 | |
| RBHWH0101 | RRTNK75285 | RRJON75062 | |
| RBHWH0029 | not predictec | not predictec | Not predicted in torts |
| I | not predictec | not predictec | Not predicted in torts |
| ; | RRTNK57291 | RRJON57063 | |
| RBHWH0045 | RRTNK19532 | RRJON19441 | |
| RBHWH013C | not predictec | not predictec | Not predicted in torts |
| RBHWH0082 | not predictec | not predicted | |
| RBHWH006C | RRTNK21306 | RRJON21210 | |
| RBHWH0195 | RRTNK73059 | RRJON72830 | |
| RBHWH0212 | RRTNK34604 | RRJON34481 | |
| RBHWH0135 | RRTNK27025 | RRJON26899 | |
| ) | RRTNK58890 | RRJON58630 | |
| RBHWH0223 | RRTNK08115 | RRJON08022 | |
| RBHWH0096 | RRTNK42364 | RRJON42165 | |
| RBHWH0023 | RRTNK27117 | RRJON26985 | |
| RBHWH0217 | RRTNK00226 | RRJON00214 | |
| RBHWH012C | RRTNK32329 | RRJON32201 | Manually edited ALGO, TANK, JON, LG |
| RBHWH0023 | RRTNK26764 | RRJON26645 | |
| RBHWH0075 | RRTNK61215 | RRJON60976 | |
| RBHWH0064 | RRTNK62982 | RRJON6272E | Not predicted in ALGO |
| RBHWH0026 | RRTNK04238 | RRJON04199 | |
| RBHWH0124 | RRTNK31316 | RRJON31208 | |
| RBHWH0176 | not predictec | not predictec | Not predicted in torts |
| RBHWH0107 | RRTNK10278 | RRJON10183 | |

| | | | |
|---|---|---|---|
| RBHWH0207 | RRTNK63849 | RRJON63597 | |
| RBHWH0074 | RRTNK28498 | RRJON28380 | |
| RBHWH0030 | not predicted | not predicted | Not predicted in torts |
| RBHWH0112 | RRTNK63170 | RRJON62930 | |
| RBHWH0209 | RRTNK43817 | RRJON43607 | |
| RBHWH0030 | RRTNK23932 | RRJON23821 | |
| RBHWH0056 | RRTNK68791 | RRJON68538 | |
| RBHWH0004 | RRTNK75019 | RRJON74782 | |
| RBHWH0116 | RRTNK33739 | RRJON33617 | |
| RBHWH0094 | RRTNK83506 | RRJON83263 | |
| RBHWH0094 | RRTNK83509 | RRJON83266 | Not predicted in ALGO |
| RBHWH0023 | not predicted | not predicted | Not predicted in torts |
| RBHWH0089 | not predicted | not predicted | Not predicted in torts |
| RBHWH0089 | RRTNK31076 | RRJON30968 | |
| RBHWH0176 | RRTNK83490 | RRJON83250 | |
| RBHWH0176 | RRTNK83487 | RRJON83247 | |
| RBHWH0072 | RRTNK67741 | RRJON67498 | |
| RBHWH0094 | RRTNK83498 | RRJON83258 | Not predicted in ALGO. Best hit is closer to IL18 |
| RBHWH0094 | RRTNK83493 | RRJON83255 | Manually edited ALGO |
| RBHWH002484 | | | Not predicted |
| RBHWH0023 | RRTNK26713 | RRJON26602 | |
| RBHWH0081 | RRTNK69103 | RRJON68850 | |
| RBHWH0218 | RRTNK41992 | RRJON41802 | |
| RBHWH0089 | not predicted | not predicted | Not predicted in torts |
| RBHWH0008 | RRTNK56372 | RRJON56114 | Not predicted in ALGO |
| RBHWH0143 | RRTNK57085 | RRJON56851 | |
| RBHWH0039 | RRTNK20527 | RRJON20440 | Not predicted in LG |
| RBHWH0124 | RRTNK31261 | RRJON31156 | |
| RBHWH0089 | not predicted | not predicted | Not predicted in torts |
| RBHWH0138 | RRTNK43649 | RRJON43433 | Not predicted in ALGO, LG |
| RBHWH0143 | RRTNK57076 | RRJON56845 | |
| RBHWH0007 | RRTNK02311 | not predicted | Not predicted in JON, LG |
| RBHWH0048 | RRTNK11618 | RRJON11507 | |
| RBHWH0011 | RRTNK67322 | RRJON67080 | |
| RBHWH0084 | RRTNK02821 | RRJON02778 | |
| RBHWH0089 | not predicted | not predicted | Not predicted in torts |
| [illegible] | RRTNK67744 | RRJON67489 | |
| RBHWH0089 | not predicted | not predicted | Not predicted in torts |
| RBHWH0089 | not predicted | not predicted | Not predicted in torts |
| RBHWH0145 | RRTNK12610 | RRJON12503 | |
| RBHWH0089 | not predicted | not predicted | Not predicted in torts |
| RBHWH0089 | not predicted | not predicted | Not predicted in torts |
| RBHWH0089 | not predicted | not predicted | Not predicted in torts |
| RBHWH0090 | RRTNK31313 | RRJON31211 | |
| RBHWH0089 | not predicted | not predicted | Not predicted in torts |

| | | | |
|---|---|---|---|
| RBHWH0089 | not predictec | not predictec | Not predicted in torts |
| RBHWH0008 | RRTNK56357 | RRJON56099 | Not predicted in ALGO |
| RBHWH0089 | not predictec | not predictec | Not predicted in torts |
| RBHWH0161 | RRTNK71792 | RRJON71567 | |
| RBHWH0208 | RRTNK56013 | RRJON55755 | |
| RBHWH0107 | RRTNK67718 | RRJON67465 | |
| | RRTNK70565 | RRJON70312 | |
| RBHWH0089 | not predictec | not predictec | Not predicted in torts |
| RBHWH0134 | RRTNK06141 | RRJON06107 | |
| RBHWH0136 | RRTNK12873 | RRJON12773 | |
| RBHWH0215 | RRTNK52126 | RRJON51891 | |
| RBHWH0177 | RRTNK02945 | RRJON02919 | |
| RBHWH0044 | RRTNK52248 | RRJON51999 | |
| RBHWH0186 | RRTNK01776 | RRJON01764 | |
| RBHWH0173 | RRTNK54589 | RRJON54323 | |
| RBHWH0004 | RRTNK70758 | RRJON70507 | |
| RBHWH0089 | not predictec | not predictec | Not predicted in torts |
| RBHWH0185 | RRTNK53539 | RRJON53287 | |
| RBHWH0083 | RRTNK65379 | RRJON65125 | |
| RBHWH0003 | RRTNK55561 | RRJON55290 | Not predicted in ALGO |
| RBHWH0065 | RRTNK15105 | RRJON15054 | |
| RBHWH0063 | RRTNK08845 | RRJON08741 | |
| RBHWH0042 | RRTNK47892 | RRJON47657 | |
| RBHWH0080 | RRTNK16380 | RRJON16278 | |
| | RRTNK24660 | RRJON24532 | |
| RBHWH0049 | RRTNK45097 | RRJON44903 | |
| RBHWH0067 | RRTNK42426 | RRJON42221 | |
| | RRTNK43914 | RRJON43709 | Not predicted in ALGO |
| RBHWH0040 | RRTNK84063 | RRJON83809 | |
| RBHWH0089 | not predictec | not predictec | Not predicted in torts |
| RBHWH0037 | RRTNK57278 | RRJON57051 | |
| | RRTNK83746 | RRJON83503 | |
| RBHWH0090 | RRTNK26686 | RRJON26577 | |
| RBHWH0005 | RRTNK36116 | RRJON35983 | Manually edited JON |
| RBHWH0141 | RRTNK68257 | RRJON68015 | |
| RBHWH0042 | RRTNK48282 | RRJON48053 | |
| RBHWH0007 | RRTNK16818 | RRJON16711 | |
| RBHWH0041 | RRTNK66015 | RRJON65765 | |
| RBHWH0184 | RRTNK58292 | RRJON58026 | |
| RBHWH0102 | RRTNK19697 | RRJON19615 | |
| RBHWH0046 | RRTNK23510 | RRJON23413 | |
| RBHWH0147 | RRTNK69543 | RRJON69288 | |
| RBHWH0095 | RRTNK84360 | RRJON84101 | |
| RBHWH0085 | RRTNK81730 | RRJON81493 | |
| RBHWH0087 | RRTNK30185 | RRJON30067 | |

| | | | |
|---|---|---|---|
| RBHWH0097 | RRTNK47403 | RRJON47158 | |
| RBHWH0026 | RRTNK10689 | RRJON10577 | |
| RBHWH0125 | RRTNK29620 | RRJON29483 | |
| RBHWH0181 | RRTNK54477 | RRJON54210 | |
| | RRTNK27087 | RRJON26964 | |
| RBHWH0048 | RRTNK09753 | RRJON09636 | |
| | RRTNK05317 | RRJON05287 | |
| RBHWH0106 | RRTNK55690 | RRJON55416 | |
| RBHWH0106 | RRTNK55690 | RRJON55416 | Manually edited ALGO. LG has KAT6a |
| RBHWH0073 | RRTNK26040 | RRJON25943 | |
| RBHWH0081 | RRTNK27028 | RRJON26902 | |
| | RRTNK54797 | RRJON54525 | |
| RBHWH0192 | RRTNK15044 | RRJON14987 | |
| RBHWH0100 | RRTNK48138 | RRJON47906 | |
| | RRTNK62741 | RRJON62487 | |
| RBHWH0067 | RRTNK71546 | RRJON71311 | |
| RBHWH0202 | RRTNK05291 | RRJON05260 | Manually edited ALGO, TANK, JON. Not predicted in LG |
| RBHWH0070 | RRTNK82520 | RRJON82294 | |
| RBHWH0097 | RRTNK51553 | RRJON51328 | |
| RBHWH0171 | RRTNK46581 | RRJON46340 | |
| RBHWH0043 | RRTNK69616 | RRJON69364 | |
| RBHWH0058 | RRTNK19599 | RRJON19505 | |
| RBHWH0175 | RRTNK65650 | RRJON65394 | |
| | not predicted | not predicted | Not predicted in torts |
| | not predicted | not predicted | Not predicted in torts |
| | not predicted | not predicted | Not predicted in torts |
| | not predicted | not predicted | Not predicted in torts |
| RBHWH0182 | RRTNK51502 | RRJON51277 | |
| RBHWH0151 | RRTNK84664 | RRJON84408 | |
| RBHWH0072 | RRTNK11133 | RRJON11019 | Manually edited ALGO, JON |
| RBHWH0023 | RRTNK67641 | RRJON67386 | |
| RBHWH0125 | RRTNK53382 | RRJON53128 | |
| RBHWH0055 | not predicted | not predicted | Not predicted in torts. Lots of KLRF genes though |
| RBHWH0185 | not predicted | not predicted | Not predicted in torts |
| RBHWH0054 | RRTNK78153 | RRJON77907 | Manually edited TANK, JON |
| RBHWH0055 | RRTNK11304 | RRJON11177 | |
| | RRTNK11146 | RRJON11011 | Manually edited ALGO, TANK, JON, LG. Hard to disting |
| RBHWH0055 | RRTNK77291 | RRJON77084 | |
| RBHWH0043 | not predicted | not predicted | Not predicted in torts |
| | not predicted | not predicted | Not predicted in torts |
| RBHWH0032 | RRTNK36573 | RRJON36420 | |
| RBHWH0117 | RRTNK70290 | RRJON70046 | Manually edited ALGO, TANK, JON, LG |

| | | | |
|---|---|---|---|
| RBHWH0057 | RRTNK19026 | RRJON18932 | Manually edited ALGO, TANK, JON, LG |
| | RRTNK74223 | RRJON73977 | Manually edited ALGO |
| RBHWH0080 | RRTNK77811 | RRJON77572 | |
| RBHWH0114 | RRTNK79101 | RRJON78864 | |
| RBHWH0221 | not predicted | not predicted | Not predicted in torts |
| RBHWH0158 | RRTNK83662 | RRJON83413 | |
| RBHWH0036 | RRTNK85831 | RRJON85560 | |
| RBHWH0033 | RRTNK12849 | RRJON12744 | |
| RBHWH0160 | RRTNK18011 | RRJON17898 | |
| | RRTNK25722 | RRJON25618 | |
| RBHWH0008 | not predicted | RRJON01150 | Not predicted in LG, TANK |
| RBHWH0003 | RRTNK21435 | RRJON21332 | |
| RBHWH0025 | RRTNK71543 | RRJON71306 | |
| RBHWH0091 | RRTNK84880 | RRJON84621 | |
| RBHWH015701 | | | All genes more like BPI than LBP, but there are several |
| RBHWH0117 | RRTNK17089 | RRJON16997 | |
| RBHWH0210 | RRTNK84324 | RRJON84067 | |
| RBHWH0061 | RRTNK60548 | RRJON60318 | |
| RBHWH0140 | RRTNK07698 | RRJON07619 | |
| RBHWH0087 | RRTNK30512 | RRJON30399 | |
| RBHWH0170 | RRTNK51885 | RRJON51660 | |
| | RRTNK77836 | RRJON77600 | Not predicted in ALGO |
| RBHWH0067 | RRTNK56506 | RRJON56244 | |
| RBHWH0150 | RRTNK84509 | RRJON84247 | Manually edited TANK, JON. They are shorter than the |
| RBHWH0070 | RRTNK69825 | RRJON69580 | |
| RBHWH0064 | RRTNK18610 | RRJON18529 | |
| RBHWH0029 | RRTNK15929 | RRJON15831 | |
| RBHWH0179 | RRTNK83763 | RRJON83521 | |
| | not predicted | not predicted | Not predicted in torts |
| | not predicted | not predicted | Not predicted in torts |
| | not predicted | not predicted | Not predicted in torts |
| RBHWH0171 | RRTNK32200 | RRJON32070 | |
| RBHWH0033 | RRTNK13143 | RRJON13036 | |
| | RRTNK52503 | RRJON52257 | |
| RBHWH0066 | RRTNK27268 | RRJON27144 | |
| RBHWH0190 | RRTNK74256 | RRJON74019 | |
| RBHWH0180 | RRTNK51632 | RRJON51399 | |
| RBHWH0211 | RRTNK58999 | RRJON58742 | |
| RBHWH0153 | RRTNK38896 | RRJON38734 | |
| RBHWH0040 | RRTNK35423 | RRJON35300 | |
| | RRTNK67403 | RRJON67158 | Manually edited ALGO, TANK, JON, LG |
| | RRTNK19808 | RRJON19714 | |
| | | | not found |
| RBHWH0117 | RRTNK54637 | RRJON54377 | |

| | | | |
|---|---|---|---|
| [illegible] | RRTNK32466 | RRJON32339 | |
| RBHWH0181 | RRTNK35353 | RRJON35225 | |
| RBHWH0050 | RRTNK48417 | RRJON48190 | |
| RBHWH0169 | RRTNK44244 | RRJON44038 | |
| [illegible] | not predicted | not predicted | Not predicted in torts |
| RBHWH0031 | RRTNK07502 | RRJON07401 | |
| RBHWH0055 | RRTNK39756 | RRJON39565 | |
| RBHWH0008 | RRTNK49007 | RRJON48772 | |
| RBHWH0063 | RRTNK09724 | RRJON09605 | Not predicted in ALGO |
| RBHWH0021 | RRTNK70675 | RRJON70424 | |
| RBHWH0048 | RRTNK08321 | RRJON08228 | |
| [illegible] | RRTNK67173 | RRJON66938 | |
| RBHWH0159 | RRTNK05156 | RRJON05138 | |
| RBHWH0104 | RRTNK76475 | RRJON76251 | |
| RBHWH0079 | RRTNK57309 | RRJON57081 | |
| RBHWH0206 | RRTNK43671 | RRJON43458 | |
| RBHWH0066 | RRTNK15770 | RRJON15669 | |
| [illegible] | RRTNK39141 | RRJON38972 | Manually edited ALGO, TANK, JON, LG |
| RBHWH0072 | RRTNK32657 | RRJON32539 | |
| RBHWH0144 | RRTNK55856 | RRJON55597 | |
| RBHWH0124 | RRTNK85172 | RRJON84898 | Manually edited TANK, JON. These two do not have MA |
| [illegible] | RRTNK15721 | RRJON15628 | |
| RBHWH0157 | RRTNK20145 | RRJON20064 | Manually edited LG |
| RBHWH0199 | RRTNK53076 | RRJON52841 | |
| RBHWH0152 | RRTNK48248 | RRJON48023 | |
| [illegible] | RRTNK56279 | RRJON56018 | |
| RBHWH0117 | RRTNK07805 | RRJON07727 | |
| RBHWH0154 | RRTNK67535 | RRJON67280 | |
| [illegible] | RRTNK51363 | RRJON51147 | |
| RBHWH0113 | RRTNK12356 | RRJON12251 | |
| [illegible] | RRTNK68989 | RRJON68738 | |
| RBHWH0005 | RRTNK57809 | RRJON57558 | |
| RBHWH0204 | RRTNK65053 | RRJON64813 | |
| RBHWH0101 | RRTNK66069 | RRJON65820 | Not predicted in ALGO |
| [illegible] | RRTNK29346 | RRJON29215 | |
| [illegible] | RRTNK73838 | RRJON73594 | |
| [illegible] | RRTNK45684 | RRJON45496 | |
| RBHWH0058 | RRTNK32728 | RRJON32603 | Manually edited ALGO. Only one good copy of MAPK14 |
| RBHWH0007 | RRTNK02717 | RRJON02668 | |
| RBHWH0042 | RRTNK57748 | RRJON57496 | Manually edited ALGO, LG |
| RBHWH0033 | RRTNK68082 | RRJON67829 | |
| RBHWH0023 | RRTNK26807 | RRJON26696 | |
| RBHWH0158 | RRTNK61706 | RRJON61448 | |
| RBHWH0172 | RRTNK60287 | RRJON60051 | |
| RBHWH0042 | RRTNK58194 | RRJON57931 | |

| | | | |
|---|---|---|---|
| RBHWH0085 | RRTNK26118 | RRJON26018 | |
| RBHWH0200 | RRTNK43217 | RRJON42999 | |
| RBHWH0076 | RRTNK21309 | RRJON21207 | |
| RBHWH0174 | RRTNK19315 | RRJON19212 | Manually edited LG |
| RBHWH0011 | RRTNK67664 | RRJON67411 | |
| RBHWH0214 | RRTNK53203 | RRJON52961 | |
| RBHWH0004 | RRTNK70708 | RRJON70455 | |
| | not predictec | not predictec | Not predicted in torts |
| RBHWH0162 | RRTNK08149 | RRJON08041 | |
| RBHWH0143 | RRTNK57208 | RRJON56979 | |
| | | | not found |
| RBHWH0023 | RRTNK28962 | RRJON28837 | Manually edited ALGO |
| | | | no ORF found |
| RBHWH0143 | RRTNK62642 | RRJON62393 | |
| RBHWH0126 | RRTNK26075 | RRJON25978 | |
| RBHWH0084 | RRTNK02561 | RRJON02517 | |
| RBHWH0175 | RRTNK20148 | RRJON20058 | |
| RBHWH0087 | RRTNK60676 | RRJON60445 | |
| RBHWH0027 | not predictec | not predicted | |
| RBHWH0101 | RRTNK66066 | RRJON65812 | |
| RBHWH0000 | RRTNK27776 | RRJON27658 | |
| RBHWH0024 | RRTNK49962 | RRJON49744 | |
| RBHWH0038 | RRTNK22085 | RRJON21993 | |
| RBHWH0038 | RRTNK22125 | RRJON22027 | |
| RBHWH0011 | RRTNK67162 | RRJON66918 | |
| | RRTNK29708 | RRJON29580 | |
| RBHWH0013 | RRTNK35745 | RRJON35621 | |
| RBHWH0019 | RRTNK03641 | RRJON03597 | |
| RBHWH0127 | RRTNK46313 | RRJON46093 | |
| RBHWH0013 | RRTNK31258 | RRJON31153 | |
| RBHWH0150 | RRTNK41983 | RRJON41796 | |
| RBHWH0009 | RRTNK65930 | RRJON65680 | |
| RBHWH0154 | RRTNK43568 | RRJON43361 | |
| RBHWH0101 | RRTNK71198 | RRJON70949 | |
| RBHWH0068 | RRTNK21129 | RRJON21035 | Manually edited ALGO, TANK, JON, LG |
| RBHWH0119 | RRTNK63131 | RRJON62894 | |
| | RRTNK18555 | RRJON18468 | |
| RBHWH0087 | RRTNK40026 | RRJON39849 | Manually edited ALGO, TANK, JON |
| RBHWH0202 | RRTNK74760 | RRJON74535 | Manually edited ALGO |
| RBHWH0139 | RRTNK25471 | RRJON25363 | |
| RBHWH0079 | not predictec | not predicted | |
| RBHWH0113 | RRTNK05755 | RRJON05722 | |
| RBHWH0079 | RRTNK03909 | RRJON03866 | |
| RBHWH0041 | RRTNK77278 | RRJON77068 | |
| RBHWH0018 | RRTNK41646 | RRJON41465 | |

| | | | |
|---|---|---|---|
| RBHWH0165 | RRTNK01446 | RRJON01420 | |
| RBHWH0094 | RRTNK39261 | RRJON39074 | |
| | RRTNK03144 | RRJON29478 | |
| RBHWH0102 | RRTNK19611 | RRJON19517 | |
| RBHWH0150 | not predictec | RRJON20116 | |
| RBHWH0154 | RRTNK85219 | RRJON84954 | |
| RBHWH0164 | RRTNK38660 | RRJON38487 | Not predicted in ALGO |
| RBHWH0175 | RRTNK80939 | RRJON80709 | |
| RBHWH0177 | RRTNK62144 | RRJON61888 | |
| RBHWH0031 | RRTNK67817 | RRJON67563 | |
| RBHWH0194 | RRTNK81238 | RRJON81021 | |
| | RRTNK28733 | RRJON28610 | Manually edited TANK, JON |
| RBHWH0159 | RRTNK33694 | RRJON33569 | Manually edited ALGO, TANK, JON, LG |
| | RRTNK23430 | RRJON23335 | |
| | RRTNK54742 | RRJON54470 | |
| RBHWH0166 | RRTNK50603 | RRJON50389 | |
| | not predictec | not predicted | |
| RBHWH0178 | RRTNK48054 | RRJON47822 | |
| RBHWH0126 | RRTNK18469 | RRJON18389 | |
| RBHWH0200 | RRTNK43390 | RRJON43171 | |
| RBHWH0104 | RRTNK76544 | RRJON76328 | |
| RBHWH0177 | RRTNK70030 | RRJON69788 | |
| RBHWH0191 | RRTNK44951 | RRJON44758 | |
| RBHWH0101 | RRTNK67512 | RRJON67257 | |
| | RRTNK29045 | RRJON28917 | Manually edited ALGO, TANK, JON, LG |
| RBHWH0072 | RRTNK11339 | RRJON11230 | |
| RBHWH0082 | RRTNK11307 | RRJON11192 | |
| RBHWH0123 | RRTNK68706 | RRJON68458 | |
| | RRTNK78002 | RRJON77766 | |
| RBHWH0020 | RRTNK18954 | RRJON18862 | Manually edited ALGO, TANK, JON, LG |
| | RRTNK35983 | RRJON35860 | |
| | RRTNK71387 | RRJON71149 | |
| RBHWH0054 | RRTNK48626 | RRJON48392 | Manually edited TANK, JON, LG. Not predicted in ALGC |
| RBHWH0011 | RRTNK67526 | RRJON67274 | |
| RBHWH0042 | RRTNK46933 | RRJON46692 | |
| RBHWH0203 | RRTNK34030 | RRJON33908 | |
| RBHWH0095 | RRTNK72569 | RRJON72356 | |
| RBHWH0025 | RRTNK02633 | RRJON02590 | |
| RBHWH0179 | RRTNK42614 | RRJON42426 | |
| | RRTNK50712 | RRJON50492 | Manually edited ALGO, TANK, JON, LG |
| RBHWH0038 | RRTNK20971 | RRJON20890 | |
| | RRTNK20343 | RRJON20250 | |

| | | | |
|---|---|---|---|
| RBHWH0205 | RRTNK20741 | RRJON20663 | |
| RBHWH0093 | not predicted | not predicted | Not predicted in torts |
| RBHWH0096 | RRTNK57791 | RRJON57531 | |
| RBHWH0030 | RRTNK84632 | RRJON84376 | |
| RBHWH0080 | RRTNK72853 | RRJON72626 | |
| RBHWH0118 | RRTNK34347 | RRJON34222 | |
| RBHWH0153 | RRTNK81619 | RRJON81396 | |
| RBHWH0003 | RRTNK22035 | RRJON21946 | |
| RBHWH0104 | RRTNK51519 | RRJON51294 | |
| RBHWH0011 | RRTNK67389 | RRJON67144 | |
| RBHWH0046 | RRTNK26086 | RRJON25984 | |
| RBHWH0059 | RRTNK69087 | RRJON68834 | Manually edited ALGO, TANK, JON, LG |
| RBHWH0054 | RRTNK16708 | RRJON16609 | |
| RBHWH0068 | RRTNK69802 | RRJON69562 | |
| RBHWH0042 | RRTNK51496 | RRJON51271 | Manually edited TANK, JON, LG. |
| RBHWH0106 | RRTNK33497 | RRJON33377 | |
| RBHWH0014 | RRTNK20438 | RRJON20358 | |
| RBHWH0093 | not predicted | not predicted | Not predicted in torts |
| RBHWH0031 | RRTNK05949 | RRJON05906 | |
| RBHWH0106 | RRTNK65107 | RRJON64857 | |
| RBHWH0098 | RRTNK78437 | RRJON78193 | |
| RBHWH0136 | RRTNK39677 | RRJON39483 | Manually edited ALGO, TANK, JON, LG. JON has a longe |
| | RRTNK38506 | RRJON38336 | Manually edited ALGO, TANK, JON |
| RBHWH0056 | RRTNK16745 | RRJON16654 | |
| RBHWH0069 | RRTNK28026 | RRJON27906 | |
| RBHWH0219 | RRTNK51711 | RRJON51483 | |
| RBHWH0016 | RRTNK54809 | RRJON54534 | |
| RBHWH0132 | RRTNK13917 | RRJON13851 | |
| RBHWH0015 | RRTNK34533 | RRJON34408 | |
| RBHWH0172 | RRTNK22400 | RRJON22306 | |
| RBHWH0039 | RRTNK57252 | RRJON57016 | |
| RBHWH0170 | RRTNK56611 | RRJON56362 | |
| RBHWH0155 | RRTNK28267 | RRJON28143 | |
| RBHWH0016 | RRTNK38806 | RRJON38638 | Manually edited ALGO |
| | | | not found |
| RBHWH0010 | RRTNK65024 | RRJON64784 | |
| RBHWH0052 | RRTNK44980 | RRJON44790 | |
| RBHWH0165 | RRTNK54522 | RRJON54258 | |
| RBHWH0204 | RRTNK27033 | RRJON26908 | |
| RBHWH0053 | RRTNK23040 | RRJON22933 | |
| RBHWH0169 | RRTNK44289 | RRJON44083 | |
| RBHWH0097 | RRTNK24405 | RRJON24283 | |
| RBHWH0106 | RRTNK46724 | RRJON46489 | |
| RBHWH0135 | RRTNK37547 | RRJON37385 | Manually edited ALGO, TANK, JON |
| RBHWH0150 | RRTNK51126 | RRJON50910 | |

| | | | |
|---|---|---|---|
| RBHWH0026 | RRTNK70578 | RRJON70324 | |
| RBHWH0153 | RRTNK58726 | RRJON58464 | |
| RBHWH0146 | RRTNK70064 | RRJON69820 | |
| RBHWH0162 | RRTNK09552 | RRJON09438 | |
| RBHWH0058 | RRTNK18948 | RRJON18850 | |
| RBHWH0000 | RRTNK78370 | RRJON78126 | |
| RBHWH0198 | RRTNK80644 | RRJON80414 | |
| RBHWH0034 | RRTNK71608 | RRJON71371 | |
| RBHWH0133 | RRTNK06048 | RRJON06001 | |
| RBHWH0061 | RRTNK78124 | | Manually edited ALGO, LG. Not predicted in JON |
| | RRTNK51679 | RRJON51449 | |
| RBHWH0166 | RRTNK27943 | RRJON27827 | |
| RBHWH0038 | RRTNK70399 | RRJON70141 | Manually edited ALGO, TANK, JON, LG |
| RBHWH0162 | RRTNK64941 | RRJON64701 | |
| RBHWH0027 | RRTNK39750 | RRJON39559 | |
| RBHWH0052 | not predicted | not predicted | Not predicted in torts |
| RBHWH0121 | RRTNK19253 | RRJON19450 | |
| | not predicted | not predicted | |
| | RRTNK42945 | RRJON42736 | |
| RBHWH022167 | | | |
| RBHWH012285 | | | |
| | sim to other N | sim to other NLRPs | |
| RBHWH0206 | RRTNK43013 | RRJON42789 | |
| | sim to other N | sim to other NLRPs | |
| RBHWH0221 | sim to other N | sim to other NLRPs | |
| RBHWH0049 | sim to other N | sim to other NLRPs | |
| RBHWH0167 | sim to other N | sim to other NLRPs | |
| RBHWH021412 | | | |
| RBHWH0032 | RRTNK32995 | RRJON32869 | Manually edited ALGO |
| RBHWH0034 | RRTNK36775 | RRJON36611 | |
| RBHWH0203 | RRTNK35075 | RRJON34946 | |
| RBHWH0220 | RRTNK48869 | RRJON48636 | |
| RBHWH0042 | not predicted | not predicted | Not predicted in torts |
| RBHWH0226 | RRTNK54586 | RRJON54320 | |
| RBHWH0084 | RRTNK01641 | RRJON01628 | |
| RBHWH0014 | RRTNK34996 | RRJON34873 | |
| RBHWH0155 | RRTNK28038 | RRJON27921 | |
| RBHWH0147 | RRTNK31065 | RRJON30954 | |
| RBHWH0150 | RRTNK56152 | RRJON55888 | |
| RBHWH0111 | RRTNK84994 | RRJON84717 | |
| RBHWH0044 | RRTNK60290 | RRJON60060 | |

| | | | |
|---|---|---|---|
| RBHWH0099 | RRTNK76257 | RRJON76042 | |
| RBHWH0216 | RRTNK55564 | RRJON55293 | |
| RBHWH0201 | RRTNK64859 | RRJON64621 | |
| | RRTNK21680 | RRJON21570 | Not predicted in LG |
| RBHWH0193 | RRTNK73503 | RRJON73254 | |
| RBHWH0141 | RRTNK17250 | RRJON17164 | Manually edited ALGO |
| RBHWH0023 | RRTNK49433 | RRJON49196 | |
| | RRTNK71879 | RRJON71657 | |
| RBHWH0083 | RRTNK70168 | RRJON69924 | Manually edited LG |
| RBHWH0190 | RRTNK13779 | RRJON13717 | Manually edited ALGO, TANK, JON, LG |
| RBHWH0068 | not predicted | not predicted | |
| RBHWH0182 | RRTNK76647 | RRJON76447 | |
| RBHWH0027 | RRTNK54919 | RRJON54649 | |
| RBHWH0034 | RRTNK10853 | RRJON10730 | Manually edited TANK, JON, LG |
| | RRTNK26209 | RRJON26095 | |
| | RRTNK51723 | RRJON51495 | |
| RBHWH0036 | RRTNK85845 | RRJON85570 | |
| RBHWH0053 | RRTNK27062 | RRJON26942 | |
| RBHWH0180 | RRTNK44738 | RRJON44548 | Manually edited ALGO, TANK, JON, LG |
| | | | Not predicted in torts |
| | | | Not predicted in torts |
| RBHWH0057 | RRTNK60772 | RRJON60548 | |
| RBHWH0004 | RRTNK75316 | RRJON75093 | |
| RBHWH0166 | RRTNK30888 | RRJON30778 | |
| RBHWH0004 | RRTNK75051 | RRJON74817 | |
| | RRTNK81931 | RRJON81698 | |
| RBHWH0180 | RRTNK62865 | RRJON62607 | |
| RBHWH0079 | RRTNK23072 | RRJON22970 | |
| RBHWH0202 | RRTNK66243 | RRJON65993 | |
| RBHWH0184 | RRTNK72236 | RRJON72021 | |
| RBHWH0130 | not predicted | not predicted | |
| RBHWH0134 | RRTNK37065 | RRJON36895 | |
| RBHWH0222 | RRTNK22011 | RRJON21919 | |
| RBHWH0189 | RRTNK61553 | RRJON61316 | |
| | not predicted | not predicted | |
| RBHWH0196 | RRTNK15799 | RRJON15687 | |
| RBHWH0056 | RRTNK21613 | RRJON21519 | |
| RBHWH0200 | RRTNK77733 | RRJON77506 | |
| RBHWH0014 | RRTNK18404 | RRJON18320 | |
| RBHWH0179 | RRTNK83189 | RRJON82955 | |

| | | | |
|---|---|---|---|
| RBHWH0032 | RRTNK36040 | RRJON35916 | |
| RBHWH0213 | RRTNK55706 | RRJON55439 | |
| RBHWH0042 | RRTNK28360 | RRJON28236 | |
| RBHWH0140 | RRTNK73261 | RRJON73022 | |
| RBHWH0104 | RRTNK79325 | RRJON79084 | |
| RBHWH0136 | RRTNK10870 | RRJON10752 | |
| RBHWH0115 | RRTNK61267 | RRJON61028 | |
| [illegible] | RRTNK75177 | RRJON74936 | |
| RBHWH0015 | RRTNK33026 | RRJON32900 | |
| RBHWH0074 | RRTNK59863 | RRJON59615 | |
| RBHWH0165 | RRTNK10073 | RRJON09968 | |
| [illegible] | RRTNK22342 | RRJON22248 | Manually edited TANK, JON. Not predicted in ALGO |
| RBHWH0132 | RRTNK45840 | RRJON45643 | Manually edited TANK, JON, LG. Not predicted in ALGO |
| RBHWH0090 | RRTNK31372 | RRJON31266 | Manually edited TANK, JON, LG. Not predicted in ALGO |
| RBHWH0221 | RRTNK69666 | RRJON69416 | |
| RBHWH0115 | RRTNK62206 | RRJON61949 | Manually edited ALGO, TANK, JON, LG |
| [illegible] | RRTNK54689 | RRJON54426 | |
| RBHWH0002 | RRTNK24057 | RRJON23943 | |
| RBHWH0169 | RRTNK43853 | RRJON43646 | |
| RBHWH0002 | RRTNK53839 | RRJON53591 | |
| RBHWH0025 | RRTNK71560 | RRJON71328 | |
| RBHWH0218 | RRTNK35589 | RRJON35474 | |
| [illegible] | RRTNK76061 | RRJON75832 | |
| RBHWH0134 | not predicted | not predicted | |
| RBHWH0192 | RRTNK59367 | RRJON59117 | |
| RBHWH0141 | RRTNK67019 | RRJON66776 | |
| RBHWH003024 | | | |
| RBHWH0216 | RRTNK30806 | RRJON30691 | |
| RBHWH0054 | RRTNK66860 | RRJON66617 | |
| RBHWH0181 | RRTNK56050 | RRJON55792 | |
| RBHWH0089 | RRTNK51029 | RRJON50816 | |
| RBHWH0009 | RRTNK66544 | RRJON66292 | |
| RBHWH0178 | RRTNK51431 | RRJON51203 | Manually edited ALGO, TANK, JON, LG |
| RBHWH0143 | RRTNK68519 | RRJON68276 | |
| RBHWH0027 | RRTNK56403 | RRJON56155 | Not predicted in ALGO |
| RBHWH0015 | RRTNK81565 | RRJON81342 | |
| RBHWH0017 | RRTNK12335 | RRJON12230 | |
| RBHWH0179 | RRTNK54497 | RRJON54233 | |
| RBHWH0114 | RRTNK79011 | RRJON78775 | |
| RBHWH0191 | RRTNK27815 | RRJON27696 | Manually edited ALGO. Fusion protein with RASAL3 |
| RBHWH0026 | RRTNK05508 | RRJON05469 | |
| RBHWH0067 | RRTNK23531 | RRJON23445 | |
| RBHWH0037 | RRTNK13023 | RRJON12941 | |
| RBHWH0095 | RRTNK49004 | RRJON48769 | |

| | | | |
|---|---|---|---|
| RBHWH0147 | RRTNK09396 | RRJON09290 | |
| RBHWH0009 | RRTNK38441 | RRJON38269 | |
| RBHWH0185 | RRTNK85396 | RRJON85132 | |
| RBHWH0115 | RRTNK55613 | RRJON55345 | |
| RBHWH0023 | RRTNK26654 | RRJON26545 | |
| RBHWH0117 | RRTNK58437 | RRJON58187 | |
| | RRTNK72856 | RRJON72634 | |
| RBHWH0072 | RRTNK64451 | RRJON64218 | |
| RBHWH0143 | RRTNK64540 | RRJON64304 | Manually edited TANK, JON |
| RBHWH0016 | RRTNK49351 | RRJON49099 | |
| RBHWH0058 | RRTNK33018 | RRJON32886 | Manually edited ALGO, TANK, JON, LG |
| RBHWH0151 | RRTNK44182 | RRJON43973 | |
| | RRTNK26034 | RRJON25940 | |
| RBHWH0048 | RRTNK06025 | RRJON05988 | |
| RBHWH0185 | RRTNK39288 | RRJON39101 | |
| RBHWH0085 | not predictec | not predictec | Not predicted in torts. Lots proteins more similar to Pk |
| | RRTNK21512 | RRJON21412 | |
| RBHWH0057 | RRTNK33725 | RRJON33606 | |
| RBHWH0181 | RRTNK67283 | RRJON67038 | |
| RBHWH0096 | RRTNK42350 | RRJON42150 | |
| RBHWH0086 | RRTNK56160 | RRJON55901 | |
| RBHWH0115 | RRTNK53458 | RRJON53204 | |
| RBHWH0070 | RRTNK30944 | RRJON30830 | |
| RBHWH0150 | RRTNK73146 | RRJON72914 | |
| | not predictec | not predicted | |
| RBHWH0119 | RRTNK36545 | RRJON36397 | |
| RBHWH0166 | RRTNK50567 | RRJON50347 | |
| RBHWH0209 | RRTNK50391 | RRJON50174 | |
| RBHWH006336 | | | |
| RBHWH0092 | RRTNK82738 | RRJON82507 | |
| RBHWH0101 | RRTNK66600 | RRJON66343 | |
| RBHWH0096 | RRTNK42432 | RRJON42243 | |
| | RRTNK08379 | RRJON08280 | |
| RBHWH0217 | RRTNK32896 | RRJON32769 | |
| RBHWH0013 | RRTNK85748 | RRJON85475 | |
| RBHWH0077 | RRTNK38026 | RRJON37846 | |
| RBHWH0147 | RRTNK37510 | RRJON37348 | |
| RBHWH0061 | RRTNK78284 | RRJON78040 | |
| RBHWH0051 | RRTNK10414 | RRJON10306 | |
| RBHWH0200 | RRTNK25427 | RRJON25316 | |
| RBHWH0156 | RRTNK27065 | RRJON26948 | |
| RBHWH0065 | RRTNK55193 | RRJON54917 | |
| RBHWH0000 | RRTNK78227 | RRJON77983 | |
| RBHWH0168 | RRTNK72077 | RRJON71859 | |
| RBHWH0128 | RRTNK63181 | RRJON62941 | |

| | | | |
|---|---|---|---|
| RBHWH0126 | RRTNK09476 | RRJON09359 | |
| | RRTNK31652 | RRJON31539 | |
| RBHWH0054 | RRTNK32206 | RRJON32078 | |
| RBHWH0100 | RRTNK79537 | RRJON79310 | |
| RBHWH0049 | RRTNK46667 | RRJON46419 | |
| RBHWH0133 | RRTNK68516 | RRJON68273 | |
| | RRTNK38432 | RRJON38257 | |
| RBHWH0034 | RRTNK71611 | RRJON71381 | |
| | not predictec | not predicted | |
| RBHWH0021 | RRTNK46985 | RRJON46740 | |
| RBHWH0094 | RRTNK81202 | RRJON80982 | |
| RBHWH0116 | RRTNK33116 | RRJON32987 | |
| RBHWH0173 | RRTNK03026 | RRJON02985 | Manually edited LG |
| RBHWH0162 | RRTNK07868 | RRJON07764 | Manually edited ALGO, TANK, JON, LG. Also possibly PC |
| | not predictec | not predicted | |
| RBHWH0020 | RRTNK42507 | RRJON42304 | |
| RBHWH0080 | RRTNK74751 | RRJON74514 | Manually edited ALGO, TANK, JON, LG |
| RBHWH0107 | RRTNK30660 | RRJON30545 | |
| RBHWH0047 | RRTNK26379 | RRJON26266 | |
| RBHWH0222 | RRTNK30699 | RRJON30590 | |
| | RRTNK09502 | RRJON09403 | |
| RBHWH011630 | | | |
| | RRTNK64221 | RRJON63980 | |
| | RRTNK36430 | RRJON36276 | |
| RBHWH0146 | RRTNK69989 | RRJON69741 | |
| RBHWH0086 | RRTNK59463 | RRJON59208 | |
| RBHWH0046 | RRTNK34601 | RRJON34470 | Manually edited ALGO, TANK, JON, LG |
| RBHWH0034 | RRTNK11121 | RRJON10996 | Manually edited ALGO, TANK, JON, LG |
| RBHWH0049 | RRTNK61209 | RRJON60970 | Manually edited ALGO, TANK, JON, LG |
| RBHWH0168 | RRTNK38517 | RRJON38341 | Manually edited ALGO, LG |
| | | | not found |
| RBHWH0029 | RRTNK29307 | RRJON29170 | |
| RBHWH0166 | RRTNK08625 | RRJON08524 | |
| | | | not found |
| | | | not found |
| | | | not found |
| RBHWH0043 | not predictec | RRJON50632 | Not predicted in TANK |
| | RRTNK55619 | RRJON55339 | |
| | RRTNK55435 | RRJON55166 | |
| RBHWH0091 | RRTNK62629 | RRJON62372 | |
| | RRTNK80807 | RRJON80578 | Manually edited TANK, JON |
| RBHWH0213 | RRTNK12353 | RRJON12239 | Manually edited ALGO |
| | RRTNK77224 | RRJON77017 | |
| | RRTNK24666 | RRJON24544 | |
| | RRTNK39617 | RRJON39421 | Manually edited LG |

| | | | |
|---|---|---|---|
| RBHWH0065 | RRTNK13669 | RRJON13601 | |
| RBHWH010564 | | RRJON64452 | |
| RBHWH0143 | RRTNK47157 | RRJON46899 | |
| RBHWH0159 | RRTNK35778 | RRJON35657 | Not predicted in ALGO |
| RBHWH0104 | RRTNK52265 | RRJON52019 | |
| RBHWH0221 | RRTNK39583 | RRJON39390 | |
| RBHWH0136 | not predicted | not predicted | |
| RBHWH0095 | RRTNK80945 | RRJON80715 | |
| RBHWH0164 | RRTNK41110 | RRJON40931 | |
| RBHWH0127 | RRTNK43500 | RRJON43282 | |
| | not predicted | not predicted | |
| RBHWH0120 | RRTNK59587 | RRJON59334 | |
| RBHWH0200 | RRTNK42719 | RRJON42500 | |
| RBHWH0035 | RRTNK63705 | RRJON63462 | |
| RBHWH0217 | RRTNK38123 | RRJON37939 | Manually edited TANK. Truncated protein in TANK |
| RBHWH0193 | RRTNK73471 | RRJON73240 | |
| RBHWH0170 | RRTNK36741 | RRJON36580 | |
| RBHWH0041 | RRTNK61834 | RRJON61587 | |
| RBHWH0112 | RRTNK19122 | RRJON19032 | |
| RBHWH0175 | RRTNK16397 | RRJON16302 | |
| RBHWH0098 | RRTNK77978 | RRJON77745 | |
| RBHWH0127 | RRTNK28876 | RRJON28754 | |
| RBHWH0088 | RRTNK11428 | RRJON11324 | |
| RBHWH0065 | RRTNK55502 | RRJON55219 | |
| RBHWH0049 | RRTNK42395 | RRJON42191 | |
| RBHWH0103 | RRTNK65733 | RRJON65476 | |
| RBHWH0110 | RRTNK19978 | RRJON19887 | Not predicted in LG |
| RBHWH0045 | RRTNK32129 | RRJON32001 | |
| RBHWH0079 | RRTNK31907 | RRJON31793 | |
| RBHWH0133 | RRTNK62556 | RRJON62301 | |
| RBHWH0188 | RRTNK49088 | RRJON48856 | |
| RBHWH0084 | RRTNK41876 | RRJON41689 | |
| RBHWH0060 | RRTNK69549 | RRJON69297 | |
| RBHWH0045 | RRTNK33189 | RRJON33059 | |
| RBHWH0192 | RRTNK50758 | RRJON50539 | |
| RBHWH0039 | RRTNK54006 | RRJON53744 | |
| | RRTNK79031 | RRJON78807 | |
| RBHWH0050 | RRTNK15172 | RRJON15120 | |
| RBHWH0029 | RRTNK27784 | RRJON27669 | |
| RBHWH0099 | RRTNK14104 | RRJON14049 | Manually edited ALGO, TANK, JON, LG |
| RBHWH0024 | RRTNK59346 | RRJON59098 | |
| RBHWH0114 | RRTNK79049 | RRJON78810 | |
| RBHWH0079 | RRTNK35384 | RRJON35264 | Manually edited ALGO, TANK, JON, LG. TANK/JON are [illegible] |
| RBHWH0133 | RRTNK46942 | RRJON46704 | |
| RBHWH0201 | RRTNK49354 | RRJON49117 | |

| | | | |
|---|---|---|---|
| RBHWH0051 | RRTNK66753 | RRJON66506 | Manually edited TANK, JON, LG |
| RBHWH0045 | RRTNK60847 | RRJON60620 | |
| RBHWH0114 | RRTNK26165 | RRJON26054 | |
| | RRTNK33172 | RRJON33049 | |
| RBHWH0160 | RRTNK32524 | RRJON32396 | |
| RBHWH0064 | RRTNK62853 | RRJON62601 | |
| RBHWH0007 | RRTNK13304 | RRJON13199 | |
| | RRTNK57616 | RRJON57377 | |
| RBHWH0132 | RRTNK13020 | RRJON12938 | |
| RBHWH0085 | RRTNK19928 | RRJON19838 | Manually edited ALGO, TANK, JON, LG |
| RBHWH0006 | RRTNK25293 | RRJON25170 | |
| RBHWH0191 | RRTNK50308 | RRJON50091 | Manually edited ALGO, TANK, JON, LG |
| | RRTNK67289 | RRJON67044 | |
| RBHWH0175 | RRTNK70752 | RRJON70504 | |
| RBHWH0093 | RRTNK62310 | RRJON62048 | |
| RBHWH0218 | RRTNK77067 | RRJON76860 | |
| RBHWH0048 | RRTNK09091 | RRJON08993 | |
| RBHWH0072 | RRTNK11254 | RRJON11146 | |
| RBHWH0083 | RRTNK65334 | RRJON65080 | |
| RBHWH0091 | RRTNK65337 | RRJON65072 | |
| | RRTNK62571 | RRJON62316 | |
| RBHWH0035 | RRTNK32237 | RRJON32112 | |
| RBHWH0029 | RRTNK16103 | RRJON16002 | |
| RBHWH0067 | RRTNK71721 | RRJON71493 | |
| RBHWH0205 | RRTNK73695 | RRJON73458 | |
| RBHWH0071 | RRTNK15026 | RRJON14972 | |
| RBHWH0078 | RRTNK38126 | RRJON37945 | |
| RBHWH0002 | RRTNK24250 | RRJON24125 | |
| | RRTNK64313 | RRJON64067 | |
| RBHWH0120 | RRTNK59631 | RRJON59379 | |
| RBHWH0053 | RRTNK28324 | RRJON28200 | |
| | | | not found, closest is RALGDS (above) |
| | RRTNK60311 | RRJON60081 | |
| RBHWH0187 | RRTNK26676 | RRJON26564 | |
| | RRTNK26888 | RRJON26774 | |
| RBHWH0159 | RRTNK67121 | RRJON66871 | |
| RBHWH0095 | RRTNK84365 | RRJON84104 | |
| RBHWH0051 | RRTNK62078 | RRJON61818 | |
| RBHWH0118 | RRTNK33885 | RRJON33754 | |
| RBHWH0020 | RRTNK43206 | RRJON42988 | |
| | RRTNK77201 | RRJON76994 | |
| RBHWH0200 | RRTNK54217 | RRJON53960 | |
| RBHWH0190 | RRTNK33282 | RRJON33161 | |
| RBHWH0108 | RRTNK09919 | RRJON09812 | |
| RBHWH0217 | RRTNK73465 | | |

| | | | |
|---|---|---|---|
| RBHWH0075 | RRTNK79871 | RRJON79644 | |
| RBHWH0153 | RRTNK52486 | RRJON52237 | |
| RBHWH0030 | RRTNK81351 | RRJON81130 | |
| RBHWH0101 | RRTNK67473 | RRJON67218 | |
| RBHWH0155 | RRTNK18044 | RRJON17957 | |
| RBHWH0036 | RRTNK85666 | RRJON85395 | Manually edited ALGO. ALGO is a weird fusion |
| RBHWH0079 | RRTNK53323 | RRJON53066 | |
| RBHWH0093 | RRTNK83413 | RRJON83178 | |
| RBHWH0103 | RRTNK62527 | RRJON62272 | |
| RBHWH0073 | RRTNK48285 | RRJON48066 | |
| RBHWH0048 | RRTNK11516 | RRJON11405 | |
| RBHWH0173 | RRTNK84227 | RRJON83970 | |
| RBHWH0090 | RRTNK49181 | RRJON48942 | |
| RBHWH0059 | RRTNK59997 | RRJON59743 | |
| RBHWH0119 | RRTNK48609 | RRJON48375 | |
| RBHWH0170 | RRTNK69706 | RRJON69466 | |
| RBHWH0034 | RRTNK36809 | RRJON36648 | |
| RBHWH0179 | RRTNK23637 | RRJON23530 | |
| RBHWH0216 | RRTNK48026 | RRJON47800 | |
| RBHWH0080 | RRTNK72811 | RRJON72585 | |
| RBHWH0064 | RRTNK46817 | RRJON46573 | |
| RBHWH0136 | RRTNK08922 | RRJON08828 | |
| RBHWH0088 | RRTNK54971 | RRJON54698 | Not predicted in ALGO |
| RBHWH0064 | RRTNK69743 | RRJON69498 | |
| RBHWH0023 | RRTNK49395 | RRJON49158 | |
| RBHWH0022 | RRTNK57589 | RRJON57348 | Manually edited ALGO, TANK, JON, LG |
| RBHWH0054 | RRTNK32749 | RRJON32619 | |
| RBHWH0035 | RRTNK09265 | RRJON09162 | Manually edited ALGO, TANK, JON, LG |
| RBHWH0058 | RRTNK78394 | RRJON78150 | |
| RBHWH0045 | RRTNK23236 | RRJON23129 | |
| RBHWH0181 | RRTNK63696 | RRJON63451 | |
| RBHWH0001 | RRTNK35390 | RRJON35267 | |
| RBHWH0180 | RRTNK32161 | RRJON32039 | |
| RBHWH0149 | RRTNK59654 | RRJON59403 | |
| RBHWH009697 | | | |
| RBHWH0217 | RRTNK00811 | RRJON00803 | |
| RBHWH0045 | RRTNK19161 | RRJON19068 | |
| RBHWH0095 | RRTNK37319 | RRJON37141 | |
| RBHWH0053 | RRTNK40761 | RRJON40572 | |
| RBHWH0099 | RRTNK69271 | RRJON69018 | |
| RBHWH0032 | RRTNK36863 | RRJON36698 | |
| RBHWH0006 | RRTNK82820 | RRJON82585 | Manually edited LG |
| RBHWH0197 | RRTNK09959 | RRJON09856 | |
| RBHWH0022 | RRTNK57442 | RRJON57205 | |
| | RRTNK75260 | RRJON75022 | |

| | | | | |
|---|---|---|---|---|
| | RBHWH0055 | RRTNK40679 | RRJON40489 | |
| [illegible] | | RRTNK76210 | RRJON75989 | Manually edited ALGO, TANK, JON, LG |
| | RBHWH0051 | RRTNK66493 | RRJON66237 | |
| | RBHWH0144 | RRTNK11818 | RRJON11704 | |
| | RBHWH0176 | not predicted | RRJON66178 | |
| | RBHWH0209 | RRTNK16850 | RRJON16752 | |
| | RBHWH0092 | RRTNK38047 | RRJON37874 | |
| | RBHWH0139 | RRTNK81417 | RRJON81198 | |
| | RBHWH0001 | RRTNK35053 | RRJON34921 | |
| | RBHWH0161 | RRTNK45816 | RRJON45621 | |
| | RBHWH0158 | RRTNK24789 | RRJON24661 | |
| | RBHWH0066 | not predicted | not predicted | Not predicted in TANK, JON. SBNO1 is |
| | RBHWH0087 | RRTNK31686 | RRJON31579 | |
| | RBHWH0174 | RRTNK41586 | RRJON41396 | |
| | RBHWH0081 | RRTNK46595 | RRJON46351 | |
| | RBHWH0063 | RRTNK22406 | RRJON22312 | |
| | RBHWH0116 | RRTNK33719 | RRJON33591 | |
| [illegible] | | RRTNK31849 | RRJON31737 | |
| [illegible] | | RRTNK34668 | RRJON34524 | |
| | RBHWH0033 | RRTNK13295 | RRJON13193 | |
| | RBHWH0079 | RRTNK50664 | RRJON50447 | |
| | RBHWH0072 | RRTNK55047 | RRJON54775 | |
| | RBHWH0119 | RRTNK30169 | RRJON30048 | |
| | RBHWH0087 | RRTNK30931 | RRJON30817 | |
| | RBHWH0177 | RRTNK64201 | RRJON63955 | |
| | RBHWH0120 | RRTNK17730 | RRJON17632 | |
| | RBHWH0174 | RRTNK20335 | RRJON20242 | |
| [illegible] | | RRTNK85632 | RRJON85361 | |
| | RBHWH0085 | RRTNK25341 | RRJON25215 | Manually edited ALGO, TANK, JON, LG |
| | RBHWH0136 | RRTNK73523 | RRJON73286 | |
| | RBHWH0055 | RRTNK72878 | RRJON72654 | Manually edited ALGO, TANK, JON, LG |
| | RBHWH0070 | RRTNK71595 | RRJON71360 | |
| | RBHWH0044 | RRTNK52352 | RRJON52107 | |
| | RBHWH0176 | RRTNK53044 | RRJON52806 | |
| | RBHWH0059 | RRTNK68513 | RRJON68270 | |
| | RBHWH0184 | RRTNK35220 | RRJON35090 | |
| | RBHWH0212 | RRTNK56442 | RRJON56181 | |
| [illegible] | | RRTNK57631 | RRJON57386 | |
| | RBHWH0223 | RRTNK18407 | RRJON18323 | |
| | RBHWH0069 | RRTNK15250 | RRJON15186 | |
| | RBHWH0037 | RRTNK26506 | RRJON26399 | |
| | RBHWH0095 | RRTNK77076 | RRJON76869 | |
| | RBHWH0157 | RRTNK19741 | RRJON19656 | |
| [illegible] | | RRTNK59956 | RRJON59705 | |
| | RBHWH0177 | RRTNK70815 | RRJON70570 | |

| | | | |
|---|---|---|---|
| RBHWH0049 | RRTNK43341 | RRJON43122 | Not predicted in ALGO |
| RBHWH0161 | RRTNK21623 | RRJON21513 | |
| RBHWH0015 | RRTNK29027 | RRJON28907 | |
| RBHWH0043 | RRTNK47877 | RRJON47642 | |
| RBHWH0162 | RRTNK09494 | RRJON09398 | |
| RBHWH0049 | RRTNK42387 | RRJON42182 | |
| RBHWH0057 | RRTNK33748 | RRJON33626 | |
| RBHWH0006 | RRTNK51918 | RRJON51689 | |
| RBHWH0157 | RRTNK20054 | RRJON19973 | |
| RBHWH0022 | RRTNK58121 | RRJON57858 | |
| RBHWH0207 | RRTNK76621 | RRJON76403 | |
| RBHWH0097 | RRTNK24197 | RRJON24080 | |
| RBHWH0062 | RRTNK09294 | RRJON09188 | |
| RBHWH0031 | RRTNK08916 | RRJON08816 | Not predicted in LG |
| RBHWH0077 | RRTNK22271 | RRJON22181 | |
| RBHWH0174 | not predicted | not predicted | |
| RBHWH0001 | RRTNK35021 | RRJON34892 | |
| RBHWH0062 | RRTNK30479 | RRJON30367 | |
| | RRTNK70681 | RRJON70430 | |
| RBHWH0002 | RRTNK25725 | RRJON25621 | |
| RBHWH0038 | RRTNK70468 | RRJON70215 | |
| RBHWH0059 | RRTNK69147 | RRJON68894 | Not predicted in ALGO. Proteins more similar to other |
| RBHWH0020 | RRTNK43036 | RRJON42821 | Manually edited ALGO, TANK, JON, LG |
| RBHWH0044 | RRTNK51859 | RRJON51628 | |
| RBHWH0023 | RRTNK27839 | RRJON27717 | |
| RBHWH0026 | RRTNK54876 | RRJON54604 | |
| RBHWH0027 | RRTNK40613 | RRJON40427 | |
| | RRTNK13859 | RRJON13793 | Manually edited TANK, JON, LG |
| RBHWH0197 | RRTNK72460 | RRJON72240 | |
| | RRTNK46334 | RRJON46102 | |
| RBHWH0167 | RRTNK12250 | RRJON12137 | |
| RBHWH0057 | RRTNK16956 | RRJON16856 | |
| RBHWH0135 | RRTNK27166 | RRJON27046 | |
| RBHWH0080 | RRTNK17781 | RRJON17689 | Manually edited ALGO, TANK, JON, LG |
| RBHWH0046 | RRTNK25424 | RRJON25313 | |
| RBHWH0158 | RRTNK64149 | RRJON63896 | |
| | RRTNK73231 | RRJON72996 | |
| RBHWH0109 | RRTNK31861 | RRJON31746 | |
| RBHWH0195 | not predicted | not predicted | Not predicted in TANK, JON |
| RBHWH0064 | RRTNK46793 | RRJON46549 | |
| RBHWH0035 | RRTNK62507 | RRJON62249 | |
| RBHWH0131 | RRTNK57924 | RRJON57658 | |
| RBHWH0003 | RRTNK80911 | RRJON80682 | |
| RBHWH0193 | RRTNK57188 | RRJON56952 | |

| | | | |
|---|---|---|---|
| | RRTNK19605 | RRJON19502 | |
| RBHWH0174 | RRTNK35541 | RRJON35428 | |
| RBHWH0106 | RRTNK33648 | RRJON33529 | Not predited in ALGO. SPECC1L exists though |
| RBHWH0098 | RRTNK47029 | RRJON46787 | |
| RBHWH0126 | RRTNK18727 | RRJON18640 | |
| RBHWH0002 | RRTNK21179 | RRJON21165 | Manually edited ALGO |
| RBHWH0122 | RRTNK28913 | RRJON28786 | |
| RBHWH0067 | RRTNK23266 | RRJON23161 | |
| RBHWH0130 | RRTNK30823 | RRJON30705 | |
| RBHWH0177 | RRTNK75758 | RRJON75537 | |
| RBHWH0033 | RRTNK68396 | RRJON68150 | |
| RBHWH0092 | RRTNK31818 | RRJON31703 | |
| | RRTNK70810 | RRJON70564 | Manually edited ALGO, TANK, JON, LG. |
| RBHWH0218 | RRTNK51754 | RRJON51524 | |
| RBHWH0058 | RRTNK33355 | RRJON33230 | |
| RBHWH0193 | RRTNK62654 | RRJON62403 | |
| RBHWH0202 | RRTNK20329 | RRJON20253 | Manually edited ALGO, TANK, JON, LG |
| RBHWH0217 | RRTNK74416 | RRJON74179 | |
| RBHWH021743 | | | No predicted genes in any |
| | | | either no sim or other gene hits |
| | | | either no sim or other gene hits |
| RBHWH0207 | RRTNK06380 | RRJON06361 | |
| RBHWH0215 | RRTNK54164 | RRJON53902 | |
| RBHWH0037 | RRTNK51740 | RRJON51515 | Manully edited TANk, JON |
| RBHWH0182 | RRTNK76878 | RRJON76672 | |
| RBHWH0125 | RRTNK50779 | RRJON50572 | |
| RBHWH0156 | RRTNK28301 | RRJON28182 | |
| RBHWH0156 | RRTNK28243 | RRJON28122 | |
| RBHWH0038 | RRTNK20977 | RRJON20907 | |
| RBHWH0066 | RRTNK13204 | RRJON13108 | |
| RBHWH0207 | RRTNK06380 | RRJON06361 | |
| | | RRJON74263 | |
| RBHWH0087 | RRTNK31382 | RRJON31276 | Manually edited ALGO, TANK, JON, LG |
| RBHWH0196 | RRTNK75947 | RRJON75719 | |
| RBHWH0134 | RRTNK77905 | RRJON77672 | Manually edited ALGO. Much shorter transcript than o |
| RBHWH0018 | RRTNK42044 | RRJON41845 | |
| RBHWH0128 | RRTNK68103 | RRJON67853 | |
| RBHWH0040 | RRTNK20037 | RRJON19931 | |
| RBHWH0117 | RRTNK09848 | RRJON09737 | |
| RBHWH0155 | RRTNK46113 | RRJON45911 | |
| RBHWH0095 | RRTNK84349 | RRJON84092 | |
| RBHWH0153 | RRTNK12176 | RRJON12050 | |
| RBHWH0007 | RRTNK76987 | RRJON76779 | |
| | RRTNK71571 | RRJON71339 | |
| RBHWH0215 | RRTNK79563 | RRJON79333 | |

| | | | |
|---|---|---|---|
| RBHWH0165 | RRTNK05884 | RRJON05849 | |
| RBHWH0088 | RRTNK54922 | RRJON54644 | |
| RBHWH0131 | RRTNK28748 | RRJON28627 | |
| RBHWH0044 | RRTNK52368 | RRJON52121 | |
| | RRTNK50990 | RRJON50782 | |
| RBHWH0129 | RRTNK56637 | RRJON56388 | Manually edited LG |
| RBHWH0129 | RRTNK71959 | RRJON71731 | |
| RBHWH0110 | RRTNK61012 | RRJON60780 | |
| RBHWH0025 | RRTNK71475 | RRJON71243 | |
| RBHWH0203 | not predicted | not predicted | Not predicted in TANK, JON |
| RBHWH0028 | RRTNK16727 | RRJON16640 | |
| RBHWH0126 | RRTNK17358 | RRJON17266 | |
| RBHWH0152 | RRTNK40256 | RRJON40065 | |
| RBHWH0142 | RRTNK00763 | RRJON00742 | |
| RBHWH0217 | RRTNK00753 | RRJON00748 | |
| RBHWH0208 | RRTNK56165 | RRJON55908 | |
| | RRTNK63920 | RRJON63671 | |
| | RRTNK69436 | RRJON69181 | |
| RBHWH0102 | RRTNK66368 | RRJON66123 | |
| RBHWH0076 | RRTNK15256 | RRJON15192 | |
| RBHWH0157 | RRTNK23691 | RRJON23588 | |
| RBHWH0119 | not predicted | not predicted | Manually edited LG. Not predicted in ALGO, TANK, JON |
| RBHWH0030 | RRTNK23575 | RRJON23290 | |
| RBHWH0047 | RRTNK68889 | RRJON68643 | |
| RBHWH0125 | RRTNK61373 | RRJON61128 | |
| RBHWH0171 | RRTNK73374 | RRJON73146 | |
| RBHWH0102 | RRTNK53913 | RRJON53655 | |
| | RRTNK50656 | RRJON50439 | |
| RBHWH0101 | RRTNK32596 | RRJON32469 | |
| RBHWH0112 | not predicted | not predicted | |
| RBHWH0136 | RRTNK10819 | RRJON10701 | |
| | RRTNK70474 | RRJON70221 | |
| RBHWH0052 | RRTNK44943 | RRJON44750 | |
| RBHWH0050 | RRTNK05810 | RRJON05775 | |
| RBHWH0207 | RRTNK26476 | RRJON26360 | |
| RBHWH0043 | RRTNK47543 | RRJON47293 | Manually edited ALGO, TANK, JON, LG. Might also be T |
| RBHWH0175 | RRTNK70923 | RRJON70675 | |
| RBHWH0165 | RRTNK05514 | RRJON05466 | |
| RBHWH0000 | RRTNK30224 | RRJON30106 | |
| RBHWH0065 | RRTNK80593 | RRJON80368 | Manually edited TANK, JON |
| | RRTNK12785 | RRJON12678 | Manually edited ALGO, TANK, JON, LG |
| RBHWH0210 | RRTNK34970 | RRJON34844 | |
| RBHWH0112 | RRTNK06144 | RRJON06113 | |
| RBHWH0081 | RRTNK80223 | RRJON79996 | |
| RBHWH0035 | RRTNK71327 | RRJON71083 | |

| | | | |
|---|---|---|---|
| RBHWH005( | RRTNK48711 | RRJON48470 | |
| RBHWH021( | RRTNK66072 | RRJON65823 | |
| RBHWH0051 | RRTNK66675 | RRJON66429 | |
| RBHWH003S | RRTNK61733 | RRJON61481 | |
| RBHWH0107 | RRTNK08913 | RRJON08819 | |
| | RRTNK31011 | RRJON30900 | |
| RBHWH016( | RRTNK32736 | RRJON32611 | |
| RBHWH006S | RRTNK18957 | RRJON18865 | |
| RBHWH0072 | RRTNK55330 | RRJON55061 | |
| RBHWH014S | RRTNK15277 | RRJON15210 | |
| RBHWH0084 | RRTNK24066 | RRJON23952 | |
| RBHWH0174 | RRTNK41500 | RRJON41316 | |
| RBHWH015( | RRTNK20009 | RRJON19928 | |
| RBHWH0073 | RRTNK48133 | RRJON47901 | |
| | not predictec | not predictec | Not predicted in torts. Lots of TLR1 predicted though |
| RBHWH0207 | RRTNK61843 | RRJON61599 | |
| RBHWH0002 | RRTNK54968 | RRJON54701 | |
| RBHWH010E | RRTNK22576 | RRJON22470 | |
| RBHWH021E | RRTNK79582 | RRJON79349 | |
| RBHWH0073 | not predictec | not predictec | Not predicted in torts |
| RBHWH000E | RRTNK83025 | RRJON82787 | |
| RBHWH000E | RRTNK83019 | RRJON82781 | |
| | not predictec | not predictec | Not predicted in torts |
| RBHWH017E | RRTNK10768 | RRJON1065( | Manually edited ALGO, TANK, JON, LG |
| RBHWH0172 | RRTNK60305 | RRJON60075 | |
| RBHWH000( | RRTNK78376 | RRJON78132 | |
| RBHWH007E | RRTNK59485 | RRJON59230 | |
| RBHWH0183 | RRTNK44348 | RRJON44146 | |
| RBHWH021E | RRTNK01609 | RRJON01596 | |
| | RRTNK67336 | RRJON67091 | |
| RBHWH0031 | RRTNK06090 | RRJON06031 | |
| RBHWH0001 | RRTNK27824 | RRJON27708 | |
| RBHWH011( | RRTNK20166 | RRJON20082 | |
| RBHWH009E | RRTNK46165 | RRJON45960 | |
| RBHWH0051 | RRTNK77331 | RRJON77121 | |
| | RRTNK03900 | RRJON03854 | Not predicted in LG |
| RBHWH020( | RRTNK53233 | RRJON52992 | |
| | RRTNK39026 | RRJON38864 | |
| RBHWH0064 | RRTNK48399 | RRJON48178 | |
| RBHWH009S | not predictec | not predictec | Not predicted in torts |
| RBHWH0114 | RRTNK79526 | RRJON79299 | |

| | | | |
|---|---|---|---|
| RBHWH0127 | RRTNK43943 | RRJON43736 | |
| RBHWH0000 | RRTNK27997 | RRJON27877 | |
| RBHWH0099 | RRTNK41224 | RRJON41048 | Not predicted in ALGO |
| RBHWH0144 | RRTNK12047 | RRJON11934 | |
| RBHWH0127 | RRTNK43902 | RRJON43697 | |
| RBHWH0087 | RRTNK36419 | RRJON36265 | |
| RBHWH0164 | RRTNK63267 | RRJON63023 | |
| RBHWH0153 | RRTNK84246 | RRJON83989 | |
| RBHWH0179 | RRTNK83757 | RRJON83515 | |
| RBHWH0217 | RRTNK06738 | RRJON06699 | |
| RBHWH0182 | RRTNK20631 | RRJON20553 | |
| RBHWH0182 | RRTNK20725 | RRJON20655 | |
| | not predictec | not predictec | Not predicted in torts |
| RBHWH0104 | RRTNK68670 | RRJON68422 | |
| RBHWH0054 | RRTNK32989 | RRJON32858 | |
| RBHWH0171 | RRTNK68553 | RRJON68307 | |
| RBHWH0049 | RRTNK47832 | RRJON47577 | |
| RBHWH0155 | RRTNK00712 | RRJON00705 | |
| | RRTNK32405 | RRJON32278 | |
| RBHWH0135 | RRTNK26944 | RRJON26830 | |
| RBHWH0146 | RRTNK29444 | RRJON29314 | |
| RBHWH0047 | RRTNK47665 | RRJON47409 | |
| RBHWH0118 | RRTNK67891 | RRJON67632 | |
| RBHWH0017 | RRTNK14021 | RRJON13965 | |
| RBHWH0108 | RRTNK01157 | RRJON01147 | |
| RBHWH0108 | RRTNK01157 | RRJON01147 | |
| RBHWH0072 | RRTNK67635 | RRJON67380 | |
| RBHWH0214 | RRTNK38260 | RRJON38092 | |
| | RRTNK04899 | RRJON04878 | |
| | RRTNK00194 | RRJON00185 | |
| RBHWH0013 | RRTNK83817 | RRJON83572 | |
| RBHWH0221 | RRTNK84295 | RRJON84038 | |
| RBHWH0052 | RRTNK21318 | RRJON21225 | |
| RBHWH0045 | RRTNK32215 | RRJON32087 | Manually edited ALGO |
| RBHWH0147 | RRTNK09797 | RRJON09684 | |
| RBHWH0058 | RRTNK76139 | RRJON75918 | |
| RBHWH0103 | RRTNK62597 | RRJON62342 | |
| | RRTNK34761 | RRJON34626 | Manually edited ALGO, TANK, JON, LG |
| RBHWH0073 | RRTNK80488 | RRJON80255 | Not predicted in ALGO |
| RBHWH0120 | RRTNK59619 | RRJON59361 | |
| RBHWH0027 | RRTNK55120 | RRJON54858 | |
| RBHWH0057 | RRTNK14521 | RRJON14468 | |
| RBHWH0060 | RRTNK60434 | RRJON60206 | Not predicted in ALGO |
| RBHWH0027 | RRTNK40539 | RRJON40346 | |
| RBHWH0206 | RRTNK80758 | RRJON80523 | |

| | | | |
|---|---|---|---|
| RBHWH0112 | not predictec | not predictec | Not predicted in torts |
| RBHWH0112 | RRTNK30644 | RRJON30526 | |
| RBHWH0112 | RRTNK30681 | RRJON30560 | |
| RBHWH0112 | RRTNK30634 | RRJON30512 | |
| RBHWH005953 | | | |
| RBHWH0195 | RRTNK05405 | RRJON05367 | |
| RBHWH0060 | RRTNK54230 | RRJON53971 | |
| RBHWH0051 | RRTNK84420 | RRJON84163 | |
| RBHWH0018 | RRTNK23444 | RRJON23344 | |
| RBHWH0070 | RRTNK69652 | RRJON69402 | Manually edited ALGO, TANK, JON, LG |
| RBHWH0119 | RRTNK54643 | RRJON54380 | |
| RBHWH0213 | RRTNK16184 | RRJON16051 | |
| RBHWH0088 | RRTNK21979 | RRJON21890 | |
| RBHWH0161 | RRTNK21862 | RRJON21765 | |
| RBHWH0153 | RRTNK12517 | RRJON12410 | |
| RBHWH0086 | RRTNK73962 | RRJON73724 | |
| RBHWH0199 | RRTNK50625 | RRJON50406 | |
| RBHWH0053 | RRTNK28235 | RRJON28117 | |
| RBHWH0027 | RRTNK54936 | RRJON54657 | |
| RBHWH0038 | RRTNK22133 | RRJON22036 | |
| RBHWH0141 | RRTNK72219 | RRJON71999 | |
| RBHWH0100 | RRTNK42974 | RRJON42765 | |
| RBHWH0038 | RRTNK70665 | RRJON70413 | |
| | RRTNK82312 | RRJON82086 | |
| RBHWH0185 | RRTNK63465 | RRJON63221 | |
| RBHWH0162 | RRTNK08222 | RRJON08108 | |
| RBHWH0161 | RRTNK62673 | RRJON62419 | |
| RBHWH0156 | RRTNK28546 | RRJON28428 | |
| | RRTNK53035 | RRJON52797 | |
| | RRTNK50636 | RRJON50422 | |
| RBHWH0042 | RRTNK28636 | RRJON28520 | |
| RBHWH0091 | RRTNK64754 | RRJON64512 | |
| RBHWH0140 | RRTNK18463 | RRJON82245 | Manually edited ALGO. Not predicted in TANK |
| RBHWH0140 | RRTNK82433 | RRJON82205 | |
| | RRTNK24807 | RRJON24682 | |
| RBHWH0037 | RRTNK66535 | RRJON66276 | |
| RBHWH0204 | RRTNK57480 | RRJON57249 | |
| RBHWH0119 | RRTNK48408 | RRJON48187 | |
| RBHWH0105 | RRTNK64762 | RRJON64520 | |
| RBHWH0004 | RRTNK75269 | RRJON75027 | Manually edited ALGO, TANK, JON, LG |
| RBHWH0134 | RRTNK84074 | RRJON83820 | |
| | not predictec | not predicted | |
| RBHWH0013 | RRTNK85700 | RRJON85427 | |
| RBHWH0204 | RRTNK25150 | RRJON25038 | |

| | | | |
|---|---|---|---|
| RBHWH0137 | RRTNK52333 | RRJON52083 | |
| RBHWH0151 | RRTNK19353 | RRJON19258 | |
| RBHWH0082 | RRTNK11908 | RRJON11780 | |
| RBHWH0119 | RRTNK74054 | RRJON73817 | |
| | RRTNK84794 | RRJON84537 | |
| RBHWH0122 | RRTNK29670 | RRJON29533 | |
| RBHWH0213 | RRTNK48517 | RRJON48290 | |
| RBHWH0218 | RRTNK31539 | RRJON31428 | |
| RBHWH0034 | RRTNK71472 | RRJON71232 | |
| RBHWH0177 | RRTNK04725 | RRJON04696 | |
| | RRTNK45800 | RRJON45604 | |
| RBHWH0220 | RRTNK43378 | RRJON43165 | |
| RBHWH0196 | RRTNK75911 | RRJON75684 | |
| RBHWH0000 | RRTNK78278 | RRJON78034 | |
| RBHWH0039 | RRTNK57371 | RRJON57132 | |
| RBHWH0004 | RRTNK70775 | RRJON70527 | |
| RBHWH0170 | RRTNK19189 | RRJON19095 | |
| RBHWH0129 | RRTNK72035 | RRJON71811 | |
| RBHWH0032 | RRTNK72100 | RRJON71871 | |
| RBHWH0127 | RRTNK43977 | not predicted | |
| RBHWH0216 | RRTNK30829 | RRJON30717 | |
| RBHWH0187 | RRTNK46587 | RRJON46343 | Not predicted in ALGO |
| | RRTNK23048 | RRJON22953 | Manually edited ALGO, TANK, JON, LG |
| | RRTNK39744 | RRJON39527 | |
| RBHWH0165 | RRTNK05842 | RRJON05801 | Manually edited TANK, JON, LG. Not predicted in ALGO |
| RBHWH0201 | RRTNK47553 | RRJON47308 | |
| RBHWH0223 | RRTNK75194 | RRJON74956 | |
| RBHWH0164 | RRTNK34521 | RRJON34396 | |
| RBHWH0168 | RRTNK41194 | RRJON41015 | Manually edited TANK, JON, LG |
| RBHWH0067 | RRTNK56329 | RRJON56065 | |
| RBHWH0013 | RRTNK85603 | RRJON85332 | |
| RBHWH0100 | RRTNK48270 | RRJON48047 | |
| RBHWH0017 | RRTNK13680 | RRJON13604 | |
| RBHWH0082 | RRTNK77139 | RRJON76931 | |
| RBHWH0153 | RRTNK55095 | RRJON54831 | Manually edited ALGO |
| RBHWH0114 | RRTNK38657 | RRJON38479 | |
| RBHWH0168 | RRTNK59743 | RRJON59495 | |
| | RRTNK28633 | RRJON28512 | |
| RBHWH0016 | RRTNK00976 | RRJON00968 | |
| RBHWH0052 | RRTNK79236 | RRJON79009 | |
| RBHWH0037 | RRTNK26250 | RRJON26146 | |
| RBHWH0091 | RRTNK21188 | RRJON21091 | |
| RBHWH0207 | RRTNK06690 | RRJON06648 | |
| RBHWH0195 | RRTNK70087 | RRJON69840 | |
| RBHWH0029 | RRTNK81372 | RRJON81153 | |

| | | | | |
|---|---|---|---|---|
| | RBHWH0159 | RRTNK04924 | RRJON04900 | |
| | RBHWH0226 | RRTNK70206 | RRJON69965 | |
| | RBHWH0058 | RRTNK76389 | RRJON76168 | |
| 3 | | RRTNK61652 | RRJON61405 | |
| | RBHWH0164 | RRTNK13741 | RRJON13673 | |
| | RBHWH0010 | RRTNK65437 | RRJON65183 | |
| 3 | | RRTNK85121 | RRJON84846 | Manually edited ALGO, TANK, JON, LG |
| | RBHWH0017 | RRTNK14403 | RRJON14336 | |
| | RBHWH0079 | RRTNK23549 | RRJON23431 | |
| | RBHWH0001 | RRTNK35700 | RRJON35578 | |
| | RBHWH0174 | RRTNK17760 | RRJON17668 | |
| | RBHWH0068 | RRTNK21488 | RRJON21388 | |
| | RBHWH0197 | RRTNK34967 | RRJON34838 | |
| | RBHWH0034 | RRTNK80254 | RRJON80027 | |
| 2 | | RRTNK51306 | RRJON51089 | |
| | RBHWH0133 | RRTNK05793 | RRJON05758 | |
| | RBHWH0035 | RRTNK63676 | RRJON63424 | |
| | RBHWH0054 | RRTNK32454 | RRJON32330 | |
| | RBHWH0020 | RRTNK31236 | RRJON31128 | |
| | RBHWH0109 | RRTNK84924 | RRJON84661 | |
| | | | | not found, replaced with PATZ1 above |
| | | | | Not predicted in torts |
| | RBHWH0114 | RRTNK79125 | RRJON78888 | Manually edited TANK, JON, LG. Not predicted in ALGO |
| | RBHWH0196 | RRTNK80658 | RRJON80428 | |

est hits are shorter for aldabra

PH-like superfamily domain, but have complete SPS1 superfamily missing SH3_GRAF domain

sing the small Cation_ATPase_N domain

; second zinc finger domain set

n BIRC3, nothing was moved.

rthologs predicted in aldabra

y close to CDKN2A also

ller proteins

to human D than B

e

ene but no great hits

)

low identity
but vague

JFF domain in them too

es

close but FCN1 has a collagen domain the other genes do not.  These now all have the collagen d

uish FCN1/2/3
guate

art of genes

ns though

CDH17

issing first group of domains

all the GST paralogs

broad similarity between GZM*

ɔ IFIT5
ɔ IFIT5
ɔ IFIT5

M1 or 5
any are IFNA1
:/B
E/K
:/B
:/B
:/B
:/B
:/B
E/K
:/B
:/B

:/B

;

uish between KLRF1 and 2

other proteins

ML1 domain

), has MYH10

er protein and additional domains.

)
)

[illegible]N1/2

)U5F3

close to non human PTPRV

SLC2

others

1. Lots of proteins with similarity toTF7L2

ET3, hard to say

)

)

omain

**Supplementary Table S4**

| | |
|---|---|
| **A** | Unique Jonathan nsSNVs with amino acid sequence |
| **B** | Protein language models evaluaton of normal human genes |
| **C** | Protein language modesl evaluation with Jonathan's amino a |
| **D** | Protein language models evaluation of normal Jonathan gene |
| **E** | Protein language models evaluation with the human amino a |

### A Unique Jonathan nsSNVs with amino acid sequence

| Ref | wt_sequence |
|---|---|
| KIAA1549 | MYYCNKTFVFYFSDGVSLEQQNLSLLSVGSKLHQSTVGSIAHMALTQSTVEPQENNFLHV |
| ARID1B | MIMAAQVAAAPAGAKNNKGKTPSLPGNLSQDLSSGGAAAAATGGPGPGSMLGVGDGTN |
| GOPC | MSGMGAGSCGSGSGACGGGVAAVASSTGGGVSMFRWLEVLEKEFDKAFVDVDLLLGEI |
| ROS1 | MEGELGINACNVSDISAACTQGCQFWNATVQKSCPLQCNQTYIRACERVSCKFGCSCA |
| RB1 | MPPKPPRKGGAAAKGQRASPDRCTGGTSPPAAARLEFGETDFVALCDTLRVSDVVRANA |
| CBLB | MANSLNGRNPGGRGGNPRKGRILGLIDAIQDAVGPPKQAAADRRTVEKTWKLMDKVVRI |
| BCL9 | MHSSNPKVRNSPSGNTQSSPKSKQEVMVRPPTVMSPSGNPQLDSKFSNPGKQGGSTSQ |
| FLT3 | MTVSVFLFGFAVLAITAMAFTSATSQHISEITCMLINQQSKDSRTLESAVFLEIPSISEDLGCD |
| APC2 | MIDYSSARSYDPIATGYPTLEELKMSGSIASYDQLVRQVEALKKENTHLRRELEDNSSHLSK |
| CASC5 | MDRIYSESNEENDHTERIRRQRRSSILKAPRSPLRDLGSGNEFTQDYDAEKRRKNSRRVS |
| NRG1 | MSEVSAETFPHSSTQLNPAASLEGLPAEDNMSGMQTEDGRVHAIAGLAATCCVCLEAERL |
| CEP89 | MALSFRRGRKGPFKHIAHGLVPAATIAPKPAVPRTPPPRSPNPSPERPRSALAAAILVTTLTGR |
| NDRG1 | MLLKFIFISKEDNPFCNAAKGDAHRGGGSRARYQLLTMSTELQDAHLAEVKPLVEKDEAITI |
| ALK | MISHGSSCYVWTCVPCKIASFLTTLVLCCFLLLCDETRWVLLQKRVGHLKKPFRISLEYVAS |
| NTRK3 | MLAASERKARATGVLSLVSCLGTGLGWLGASQGMQRLVFERRMCYDESRNLSDNRLTTL |
| CIC | MKPARKAASSSSSSSSSSNSRSPPSTRAKTQKRRTAEGGPKSAGEEEGIRGGGEESPAQPL |
| DAXX | MAATVESNIIVLDDEEEGLPPPMPRESPGPAPASPQTNGTPTKKPDPLAQHGGSFKAENER |
| LATS2 | MRPKTFPATTYSGNSRQRLQEIREGLKQPSKSSGQGLPIGPGSETSLDPKILIGKDAARQQ |
| NOTCH1 | MQQLLVQSLLLSLLPVLSRGLRCTQPAESCLNGGKCETYSNGTGVCLCNSAHVGDRCQL |
| BRD4 | MSVESGPGTRLRNLPVMGDGLETTQMSTTQAQTQPQQGNTVGTNPPPPETSNPNKPKR |
| RNF213 | MKCPKCDYISQDASPNFCSSCGFRLVPHISAHSKEKVVGSGNVKPAPVPEWEVESGREL |
| THRAP3 | MSKTNKSKSGSRSSRSRSGSRSRSRSFSKSRSRSRSVSRSRKRRLSSRSRSRSYSPSHNR |
| SETD2 | MGDFYDPEHPTPEEEENEGKTENVQKAGFVKGPVFKGVASSRFLPKGTKTKVNLEEQGR |
| CLIP4 | MTIEDLPEPSQEGDSLIERYQFLFPGSETSVTFSVAAAPMPSDCEFSFFDPNDAACQEILF |
| PTPRB | MELGGAGDHWVQLYADESQSIIRNTSIQRGAAHVDLDGLVPGTRYRVEIVSQAGPHRTS |
| KMT2D | MPLVRYRKVVILGYRSVGKTSLAHQFVDGEFLECYDPTVESTYNKMLMVGKDEFHLQLVD |
| NSD1 | MEPACEVSRRGGLAPFAGPRALAAPESAGSPGPEPPNGCTAPLPAKAAYGQDPASCYIPL |
| IDH1 | MSSGSLRGYPNLHQALAQPPEVPRHGGDNSPPLNPCAKICTEMILAQREQVDRRSQQRI |
| BCL11B | MNFPLGDILVFIEHKKKQCNGTGGVCYEKSLDKSSPPPLSRSELRKVSEPVEIGIQVTPDEE |
| KMT2A | MAHSGRWRFPARPGSGGWGRRGLGGSRLRVPAVRNLRAPEPPPPHAGAGDGELGGP |
| ATRX | MRRGRKAAGGRSSPSSSLRQDGGGAGDGGTSGFVSVQRLSSVPAWPGIPSSGDMTAE |
| NCOA4 | MSPPQDRSGSSCSKESLVKCLQAKKDLETAIAGVVKAEQQMEVNWREVKAQIHSCISRH |
| ERBB2 | MASIVSAPLQSSDHHPHPASLAFPFHPPAVCTGTDMKLLRPSSPENHYATLSRLYQDCQV |
| BRCA1 | MESSVAAIGDVQSVLLAMQKNLECPICLELIKEPVSTKCDHIFCRFCMFKLLSNKKKGPAQ |
| CNOT3 | MADKRKLQGEIDRCLKKVSEGVEQFEDIWQKLHNAANANQKEKYEADLKKEIKKLQRLRD |

| | |
|---|---|
| STRN | MDEQAGPGVFFSNNNNNNALLLPPGGAGPGLEPGEAAAGGSLAAAAPAPVSAARAQYSV |
| TET1 | MAHHARPSRLVKKEELGKRKTSEGKKKSSQVKKKAKKNSKKAVSSGKCKKSMQEKNDVKK |
| TRIM33 | MAENKGGGGEEAAEPGPDGGGQEPAATSPSGSAPPAPAVAPPEERDSPCAAAEKGPAEA |
| SETBP1 | MSSPVGPTHKNKAKQVYDRPPKHSALHYDPGLQQDFTSDTLKSKHQQKSSNQHHLDWS |
| TRIM24 | MEAAVSEAAGAWPADRAGAAALSGENEAESRQGPAERGGEAAQLNLLDTCAVCGQHIQS |
| CDH11 | MKEDSCLHAALICLGMLYYSHAMSTEKLNPLRPSLHGHHEKGKEGQVLHRSKRGWVWN |
| NAB2 | MALPRTLGELQLYRVLQRANLLSYYETFIQQGGDDVQQLCEAGEEEFLEIMALVGMATKPL |
| RALGDS | MVNRARAERRDAGAGSPENMFESSRRMRSLWDGVRLEVAGEGSSPVVLHSFTQLDPDL |
| ACSL6 | MRILESPHGPTLFLGQHSSMFIKEFALSIFEKMQAQEILRSLRLPEFDNFSQFFHSLPATTLV |
| PPFIBP1 | MMSDASDMLAAALEQMDGIIAGSKALEYSNGIFDCQSPTSPFMGSLRALHLVEDLRGLLE |
| TP53BP1 | MDPSQSQLDLGFSQQDTPCLIVEDSQPESAVVEEDTAYGCLSLLTLPLPNGQSPVLEIMTS |
| MET | MKTVAAFTSGIIVFLFSLVQRTSGQCKEAAIKSEMNVSINYHLPHFITETPIQNIVLYKHHIYIG |
| LATS1 | MKRSEKPEGYRQMRPKTFPASNYTGSSRQMLQEIRESLRNLPKPSDAAKAEHNMGKMSA |
| NUTM1 | MTSQPSQGPLLPPLLLPAFPGAALLLTSPNVPGGKLIVKLKPEGAQELPRAQTIILMRAPVGCI |
| MAP3K13 | MHMHGTMASPQEHLSWPSSTNLINKTFSEDKDFSGIHGDHAPTGNHPSPEIIEDVREKG |
| AFF3 | MGTVLLKKGNTVVPQPSSRTSGLSSKRSSSSSSSESESSSESDSESESSSSESDGSKPSHYS |
| KCNT2 | MQFLGPAPQHVTSYDPGIQFGKLKLYFLSFYQTGLRIRLFNFSLKLLSCLLYIIRVLLDDPTQ |
| FOXO3 | MIPEEEDDEEEGGSSPMAIGSAVPGGGEGEALAPEEAGRLLAPLAGVGTEGLGQAPGAAA |
| SSPN | MRKEEKKPQNNTSLKNQNQAGKAPEGEKSTDPANQQKPAMKKKAKGDPKTGQEEESHT |
| POT1 | MGLLYYLHVSLHYRTLINPTSKSRASSNFMDLTFARVNCLPVQKKFALDWARDIRSGAVAL |
| PTCD2 | MVRSEDRMAAAVAGGSSRILRETVKRSMLRSATAAFGCWGCPHGAKRYLLTDNVIRLQE |
| GRN | MQLPGPGARWGAGTCLHAHGQGPGSRPRMWSLLPLWLALSGAVASLRCPDGQLCHG |
| POLQ | MTIPDDQADKLLLASWGLPEVVLEKYHSLGVVHMFAWQAECLMLGQVLEGKNLVYSAPTS |
| LRP1B | MQFNLMRRKLFFTDYGNVAKIERCDMDGMNRTRIVESKIEQPTAVAVDLVNKYVYWIDIY |
| DKC1 | MADGEGCIAKKQKKKKDKKSLPDADVADIQHTEEFFIKPESRVAQLDTSQWPLLLKNFDKL |
| TERF2 | MAGGRGRVSRSRGGPMAAGSGALGLRAPEEAVNRWVLQFYFHQAVGAYRARRNRDFR |
| ACD | MRSAAHPLPAASGGGGDPASRTPRPASAATKPRHVSGLSLCYLPRSTIFQAKNLDRIPESK |
| CYC1 | MAALSAARCLVLRTRGGSLLPPPPGPSLRPPQASMSSLSGMSRGRKVALSALGVLAAGGA |
| LMX1B | MEKIAPTEFVMRALECVYHLSCFSCCVCERQLRKGDEFVLKEGQLLCKSDYEKEKDLLSSV |
| AIF1 | MSSQQNLQGGKAFGALKAQQEEVLDSLNKEFLDDPKYSTDEDLPEKLEAFKKKYMEFDLN |
| CMAS | MDSAAAAAGAVPAAEAEQRPAHMAALVLARGGSKGIPLKNIKPLAGLPLIGWVLRAAIDSGV |
| AGPAT2 | MEPSWLLGGAALLLLLLLPALLALSPAAHFQLKIGFYCVLCVVASALAVPLCLLRHGGRTVRN |
| ATM | MLIDKSIFDASKSLHLQVYLVLLKELPTEENLLPEDDFLLLLKPLSDVCSLYRRDQEVCKAILN |
| KLRF1 | MEDEEGYTVLNLRPKQRQPGRPPGAGPQDSRQCPHWIRIALGVGWAGNIVLLAAVIVLG |
| SYNE1 | VTVNKWEQFERDKETVVKYLKQASSALERILSFSSLESLSSELDQTKELSKQTIAIAAQAENLV |
| MRC2 | MENGAGGGGQEQWDRGWKWHSPSCYWLGEDRVTYSDARKMCIDYGATLVTITNRFE |
| MALSU1 | MLPAARGLRLRPAGPGSGAAVRCAPCWAGLAGRALGGGPSTEVRAGGGAEQERRAPDH |
| IFNGR2 | MPRPGAALLLCLLLLSLPEPPAAESPLQLPTPQSVTIYSYNYQNLLTWAPVKVDNGSVLYTVQ |
| MINPP1 | MASCFRGPLPGRHSLPLFLLLLRVSLAQERAVATSLLSGYFGTKSRYEEVNQHLLQDPLSLG |
| APOA5 | MSHKAALLVLLLATFSELPYATSLLQRHALPPRLFQGHSVKCSCSPHAAAQAERARNGFWD |
| HOXD12 | MCDRGLYRSGYVGSLLNLQSPDSFYFPNLRTNGSQLAALPTISYPRSSIPWTCSSSCTTQP |
| PRUNE2 | MHILLLQVSPPDILCLPVLNIPRREFSYYPETRFILEQLNIPESFHIFQDEINLHQLNDEGKLS |
| SIRT1 | MADEETLLLQPHSGRRAAVAEPLPKRQRLDSDDSGGRGSGKGPGPERGVGTAPTTSSGT |
| IGF2R | MRRAARSSGRLPAPRLPAWPLLLLLLLAGLRGGGEAQSLDLDPLCSYTWEAIDVDKHVRYK |
| PROM1 | MSENSLCFSIDMQQCIGVLCAYAANQHLTSQVRGAKNLVHSNFKDLRIFLNDTPGLPGVS |

| | |
|---|---|
| ATG4C | MEATGTDEVEKLKSKFMSAWHNMKYSWVLKTKTYFSRNSAVVLLGKCYRFKSDEPSELSP |
| TLR4 | MNLEGLVCSKSQCRFAFVVLKKLWKVLMLLLSMVLSVQVLQYCGRMMPDVPFEKDINIC |
| ADAMTSL1 | MDCPPEAGDFRAQQCSAHNDVKYQGQFYEWLPVSNDPDNPCSLKCQAKGTTLVVELA |
| FGF19 | MWRSLCKPYTSLALLGLCFAMVVRSLPFSDAGPHVNYGWGEPIRLRHLYTASKHGLFSYF |
| CGNL1 | MEPYCAGFKHVQRDCGVHLKLANDETDKARNTQHSKPGSYGVSIRVQGIDGHPYIVLN |
| ITGB3 | MPGSPCSGSAHPRLSCPQPHDSSPRRLKSSGSLSPGLQPLLPPHPCKPASEKMQGLRLA |
| NCOR2 | MSGSTQPVTQNWRATEPRYPPHAMSYPVQISRTHADVGLLEYQHHPRDYTSHLPPGSIM |
| MSRA | MFSRWQEGGKSLAPLRCEISSSCQPPPCLVFCRLLSSCPTPSLCPLPGVQKLPEVVGTRR |
| TRIM25 | MAELVRAPSVPSLAGLEEELTCSICLCLFDSPVTTPCGHNFCGPCLDITWSGLLQGFSCP |
| F7 | MVSRHYLVLLLCFLLLLPLFPAAVFIKQDQANSLLKRQRRANSLFEELKIGSLRRECFEETCS |
| TSFM | MQRAALTGVCGPLRGGARPGPHPVGLSCMPLQLLSSPCQQVVSWQPVCLFHTGLRAFA |
| SORCS1 | MGSSKEPDLVHLETKTVDGHAQYIICRMQNSTEATRSKPFPGYIDPDSLIVQDDYVFVQLT |
| AOPEP | MTENLQSLCLANWNMEMKLDPAKDDLPLMANTRHMLVKHYVLDLDVDFKSQVIEGTVV |
| NFKB1 | MAEEDAYIMGVSDHQMFAMDQFMGVNALFNNTGFITSDLPLRTDGPYLQIIEQPKQRG |
| ZBTB20 | MSALGPRLGVIYLKSDTIWKPLKKPKTAENQKASEENEITQTGASSAKPGLPCLNLEAVPPLS |
| FBXO31 | MAVCARLCGVGPARGCRRRGAQRAQQREGPADSEPDTDAEEERIEGARLGQGEALAA |
| BSCL2 | MMSSTGPFLQWAQEVAALLLLRVRRTVLQTAILLCVLLLLLWIAVFLYGSFYYSYMPTVSYVS |
| PPP1R1A | MLLQIISNWSCLSVGQKIDEDRLPNPLLKPGLSMSPRQRKKVPRTTPTMKELQMMVEHHL |
| VWA5A | MMSCGLLNSSNEPVPLRSVSVTVLIRGFVADVGCELLYRNEEQGPVEAVFVFPVDAEAAVY |
| METTL1 | MAGAPPQKRFYRQRAHSNPLADHTLCYPTKPEEMNWAEHYPEFFAPLTKDDRHDDPKD |
| BNIPL | MVPSIKKGRWWGGRRWGLSAALAAWEDSFPVSSSHLTPCIQEVLTHQEFILAIEATLRME |
| PIK3C2G | MKQISISTVDKVIEEAKAICSVLCSVETRAITDAVKKLRSVPLGKTQNSLQNVETPFRVLLENA |
| SVEP1 | MASHPKEEHCYLLHSFAEFEALARRALHEDLPSGSYIQEDISHCSYLCEANENCCDIMAS |
| RAPGEF3 | MHLFRSYSYQIQLEGEAGSLPRIQGIRWTPLPDSESAPDCEQSPTQAHAEKVWRAGRLLF |
| XRCC6 | MSEWMSYYKNEEDEEEEYPEEVETGGEYKYAGRDSLIFLVDASKAMFEYYDDDDEWTP |
| IFNAR1 | MAPPELLLLLLLPLLLAPLAAGQMNLTSPQNVQVNVVNTNFTLRWSWGSENDFNVTFSAQ |
| CYP19A1 | MILETLNLMHCNITSVVPEVMPTATVPILLLVCFLFLIWNYEETSSIPGPGFCMGIGPLISHG |
| METTL21B | MAASRQPLLCHVEFGGAVAGREPPEAAPEGLEAVFPREVGLFADTFPEETRYGFCGHVLTI |
| HIF1A | MELTGEKKRISSDRRKEKSRDAARCRRSKESEVFYELAHQLPLPHTVSAHLDKASIMRLTIS |
| MTFMT | MRGRLLRAWREGAVGVGAAARRGLAELGASRGKPPWRVLFFGTDDFAAESLRALEAARK |
| GZMB | MLILILLPVALLLPPRARAGEIIGGQEAQPHSRPYMAYLEIQRKGKTYICGGFLVSKNFVLTAA |
| GHSR | MRNQSGSPNSSARVPDYENDTWPEYPVHLFPAPVLTGITVTCILLFVVGILGNLMTMLLVS |
| BAG6 | MAEAQNLEVMVKTLDSQTRTFTLEAEVTVKDFKEHIAGSVNIPAEKQRLIYQGRVLQDDKKL |
| ZPLD1 | MERIWLLLLLIIQVLSVTAQFNGYNCEANLHSRFPAERDISVYCGVQAITMKINFCTVLFSG |
| SYT13 | MVLSVPVIALGATLGTATSILALCGLTCFCKCRRPRKGLSGKDQDMDTENAKPSMLQPVQ |
| POLM | MALVPLKRRRRAPSPAAAPLGAGRFPAVVLCLVEKRMGASRRAFLTQLARAKGFRVDGAY |
| AKT1S1 | MADNHRESWAGLIAEAERYRRRTGNEVVLITAFRAPQPGGFAYAQHGCGALGDAARRYL |
| TEP1 | MPPRAQPEPEPGLRDRSQQGGLGPGLLGSPRSGLKSEPGSGAGPAGASHQRFRAAAGR |
| CCL5 | MKATVAALAVLLVAAFCSLASPAPLGSDTTVCCFSYTTRKLPQSHLKEYFYTSGKCSQPAVVF |
| ENOX1 | MLWMEQFLQSHRWQNYSICVPGFDPSLSMMTGITPINPMIPGMGLVPPPPPTDVPVVKE |
| RBPMS | MNSLPADKESGAAEPGLPEEEVRTLFVSGLPLDIKPRELYLLFRPFKGYEGSLIKLTSKQPVG |
| MRE11A | MSCINSGDEGDTFKILVATDIHLGYLEKDPVRGNDTFVTFDEILRHAQEKEVDFILLGGDL |
| CPA2 | MRLILFFSALAGASWCLETFVGDQVLRVKANSEEEINKLQLLETLEHLQLDFWLHPSNPAL |
| NAV2 | MNKVEIYTDWANHYLAKSGHKRLIKDLQQDVTDGVLLAEIIQVVANEKIEDINGCPKNRS |
| LINGO2 | MLHTATSFWQPFLGLAVLLIFMGSTIGCPARCECSAQNKSVSCHRRRLISIPEGIPIETKILD |

| | |
|---|---|
| ZFYVE28 | MRPIHPFSFQDNVLNIVNQIMDECIPHERASRDFCVKFPEEIRHDNLAGQLWFGAECLA |
| ECHS1 | MAVALSSLRRPAWALAPARLLLRAPARLSSAAGTQFQYLMVEKKGAKRNIGVIQLNRPKALN |
| GZMH-aging- | MIQGNWSLASHDPGKGLLGTAQGERSPAMPVTPPEWTQAGDSCFPPSLPADKSISDSRG |
| PPM1D | MEGLYSLGVSVFSDQGGRKYMEDVTQIVVEPELPAEGEEGPPYKGGSVPGSQREAESRGI |
| POLE2 | MAAEKLRSKVGSAFKMHGLLLRAEANKYLTEALQSVNELELEDVIENIIDAVEKQPLSSNMIE |
| TOP2A | MELDESLSPLRPVHENPKVGQGRPAAAAGKRLSVERIYQKKTQLEHILLRPDTYIGSVEFVT |
| PPP1R26 | MFLMNASPLIALKTKWESFGQSRSCRYPVCFSESEEDVTRASVSAKVQMIINNLQSQESSL |
| ERCC1 | MEPMEAAQVPEGRKRFTVQSQDPEHTMVRPIFKSSCSAAEPRAPALPGATSYAEYVVAQVS |
| CSMD3 | MCMGEINLEQCFQTGVAVCVERAPGGPSQFVYLLCPQVHQIVALIGHGSLLQANGNCW |
| CRLS1 | MHRRGAEEPPAATRSWYVLVKKALQYENPWTIPNILSLARIGLAPVLGYLIVEENFNIALGVF |
| APOA4 | MAELRGPCGELSPAGRRGALPGGTGGGAESQTDLVCPTLPCDSEGLNTWWGRCRNSR |
| APOB | MGPPQLWQLLLLLLLSSDVLTQDGPESRNSACSKDATRFKHLRKYIYNYEAEIASGVKGTA |
| HDAC6 | MKRGGRRGPQPRRNPRLPETKRRGRAQRMNKDMEQLESGLQRLDLSLEETVPVGTGLA |
| WDR27 | MESPVDLCFTEEGCSHDIVLEKQVFRSKVPESHVQLACNLHYCAFPLNGNELCIWNTGD |
| HIP1 | MEVGKMLMLTYPFSVFQTVSINKAINTQEVAVKEKHARTCILGTHHEKGAQTFWSVVNRLP |
| CEL | MEPRILKWDVVFSHLVGTLKAKEYKNRCMQVTLTQTGTRGSEDCLYLNIWIPQDKKGVSTI |
| TERT | MAGSGPFGAVLAMLEGCYSEVLGLEQFVQRLREAAACGPGEAELLQRGDSELYQTFVSQ |
| NCAM2 | MYAKCIHAEFTLYFSALLQVTISLSKVELSVGESKFFTCTAIGEPDSIDWYNPQGEKIASSQR |
| LEP | MRCPSLPLCGLLCIWLPLFYSRPVKIDKVKADTKNLTRTIIARIQEHQLFPLNLKINGLDFIPG |
| PARVG | MEDAVVTEELEINPFKVLGVLICKRAKSNTMEPDFFTAFAQPGVLHHSPLKNDAIQGEKRK |
| XDH | MASERTTDQLVFFVNGKKVVEKNVDPETTLLTYLRRKLGLPGTKLGCGEGGCGACTVMIS |
| APOM | MLQRSWSYLLYLYGALISTLSPCAPPAPLPASGLDRDQFPRQYLGTWHFIAAAAATPAALEIF |
| CHIT1 | MGRVFVWIGLAVLLFLQCSSACKLVCYFTNWSQNRPAQGRFLPEDIDPNLCTHLIYAFAG |
| LMNB1 | MAASTPVGQRSRGSAASTPLSPTRISRLQEKEELQHLNDRLAVYIDKVRSLETENSVLQLQV |
| TPP2 | MAAAAEEPFPFHGLLPKKETGAASFLGRYPEYDGRGVLLAVLDTGVDPGAPGMQITTDGK |
| POLE | MVLRNSGRRRAEPAGGDPDTPRDDGSGLSALKRLERSQRTDRLDAQFGFERAKEPGEK |
| GSR | MAAAFDYLVLGGGSGGLASARRAAELGARVAVVEHQRLGGTCVNVGCVPKKVMWNTAV |
| HLA-DRB1 | MGAGWILGAGSCWAGALLVTLTVLRTHLAHCTEPPRGFVYQTNSECYFTNGTERVRFLER |
| HRH1 | MCETKNSTEAYPIPLGMALGSISLITIIMNILVLYAVKTERKLHTVGNLYIVSLSVADLIVGAAV |
| SH2D4A | MLQQILADMYIDPELLAELSEEQKQILFLKMRQEQVRRWKETEAATDKNSAKKVTQRKAHG |
| NEDD4 | MARRLRLHFAAQRSNTDPLSEISGDGLDGNVLVYFQRPSRNSAPDIIQIKVTPRQAIFPTV |
| MARCO | MENPKNYNEDGISNFTKMLTMSDKIRFASSGTTTFELHVFNMQEEMRKLQREDNSHLAE |
| MAPK1 | MAAVSGAGAAGGSANPGGPEMVRGQVFDVGPRYINLSYIGEGAYGMVCSAYDNVNKV |
| CFH | MLPTDCETPPPRRTKEILTGTWDKASYPHGTSAIYNCRPGYIKLGRVTFECRNGLWHHVH |
| GCNT3 | MFRERKYLCQIQRLLFMGFLMLFATVVLKYSSRECNFSDLGKETVHLKSEGCKDHLYQSL |
| DHX33 | MPQGGEVPPAKRLKLAPPPRRAMLLAPPGAKRQPPAPSAGEAQRRSLPIFQARGPLLSQLR |
| NFATC1 | MPSPGFPTSKHFLAGPPDPARRRETLVALQRSSCPFSSSSSSSMKSAEEEHYNYASSNINA |
| PLEKHA1 | MPYVDRQNRICGFLDIEENENSGKFLRRYFILDTKEDNLVWYMDNPQNLPSGSPPVGAI |
| PUM2 | MSIPCVLLGLNEVAWQETRGMMHANIGQEASGVGIGVAPAGTVGGATAGPGPGTSGPS |
| TREML1 | MGSFPAWLLLLALTEPCLAKEELELVTAVQGSPFSTKCYYNDTYSPKEKFWCKVLSDQECN |
| TNFSF8 | MCFSPEKTLFQANDSHEAAMTMTEASVSRRLNVGNKTYFYFTTASLTVCLVFALATIMVLIV |
| PKN2 | MASNTAEREILLTDLQGDARSLLVSENVSIGQKLDFSDTMVQQKLDEIKDQIKREIRKELKI |
| CD302 | MEPPFPALLCLLGLLATASGAAECPSSAWIPFRSSCYIFLQGTLEELDNIDEARNLCKDNAS |
| IRG1 | MLWAGEVLLAKSRPLFFPQTITENFANVIHGLNASHLTDLVIQRSKRMILDTLGVGLLGTSTE |
| MID2 | METLESELTCPICLELFEDPLLLPCAHSLCFSCAHRILLSSCSSSESIEPITAFQCPTCRYVISL |

| | |
|---|---|
| DHX9 | MGDVKNFLYAWCGKRKVTPTYEIRAVGNKNRQKFMCEVRVEGFNYTGMGNSTNKKDA |
| C5AR1 | MSVNEVSEDYDLDSYVNFTAPSFDMPTPTPQTFQLTPILWVSLVLYCLVFLLGVPGNAAVI |
| IL1RL2 | MGEECMVHDVLHSQNLVSVLVGEPLTIDCLLDKTLALQHMNYSLTWYKSGRKMPVTQVK |
| ATF6 | MAAEEVEQGTAEPGPLPPGVGSDWESVLFDELSDLADTDETQLDVDNEIYENSLYNLDFD |
| ST6GAL1 | MVHINLLKKLLCGILLLLVVVTICLWRETRKGYYIPFKTESKSFQIPRALGKGDVLTSHLLSSK |
| LY75 | MRRAPGRSLATCWVALLAACCFACGASAGNGAFTIRHEKTSKCLQVKNSQIMAADCKETI |
| RABGEF1 | EEMSLKSERRGIHLDQSELLCKKGCGYYGNPAWQGFCSKCWREEYHKARQKQIQDD |
| CD1D | MGNTKVRPKVKISPMKSGSQSHFHLLVCTVTGFYPRGIKIKWLKNRQEQTAGVVSTELLQN |
| ADAR | MNTGFGRGKGSYQTPPKHIYCSPNPAFFNNSLVPQENNRDRFRNQQVQFLLGQITEAP |
| ATP7A | MAQGLVTVGGGANVGQGHRGRSRPAAPRGAGCQRCCCGEGAQLADRCWECEDSS |
| PSMD5 | MAEEALELLARVSRLEEPLEELRALGVAVQALPLSALRERAPDIRLGSLFALLNVNDREQVAL |
| TRIM8 | MAENWKNCFEEELICPICLHVFVEPVQLPCKHNFCRGCIGEAWAKETGLVRCPECNQA |
| PIK3AP1 | MASAGAPGGCDVLIVYARDASEWCQYLQNLFFSCRQVRKQRVLSYEVKYGGAIPWEEFD |
| APOBEC2 | MAENQEKPRDTQTGDEQPENSEESEEKKKELEKLPPFEIVGGERFPAFFFKFQFRNVEYSS |
| DAPK1 | MTVFRQENLDDYYETGEELGSGQFAVVKKCREKSTGLQYAAKFIKKRRTKSSRRGVSREDI |
| STK11 | MDVQEPQELSMFAESELMSVGMDTFIHRIDSTEVIYQPRRKRAKLIGKYLMGDLLGEGSY |
| NFKBIZ | MIVESNREAPADGASGSMSSPLNLAYFYSASPYSSDGSCSPAHSAPGSPGSDSELSVCSG |
| TRAT1 | MTIVLLSQILFEIKGEVLLQNSNICTEMEFECHYSIWGLLAFMTMALIVSLVINISYCIANKQK |
| IL17C | MSLLERLLGYMHLSSPISVHQGLLCALLLFSLPAGQAQKRAKDHQRHHHHHHCFSQEQL |
| CD38 | MPLQNRSSKTRQRTILLVIIMVLIVTIVAVLAAVLLANGMKETSMQLLQWKGRGTTKNLQEVI |
| FGA | MEHGIQSDCKQEKSWPFCADDDWGHKCPSGCRMQGLIDETDKDFHHRIQRIRELLLD |
| IFNL3 | MKKCHLSKYKSLPPWELKTFKNVKDRFEDIMLLSDRKCNTKIFHRNWEVKELSVHDRVIL |
| DHX58 | MLTPALPPSYWLCASCRCPVPARMELRDYQWEVILPALEGENIIIWLPTGAGKTRAAAYVCI |
| MADCAM1 | MEPTLLLLLLSLGWTCSGRLTVLPQEPLVQIGGSIQLNCSLDCPDGKPQWKGLDTNLGNII |
| MARCH | MPLHQISVIPARDVTSSRVSRSKTKEKEREEQNEKALGHSVSRSSNISKAGSPTSISAPHSFS |
| CD207 | MSRENICDNMELAEADPWPKGNPPKTSVPWRHRERSIVTAVLLLLCLALATSLLAVTLLYLR |
| MAP4K2 | MSLLRDVSLLDPRLRYELIQRIGAGTYGDVYKARDTQTAELAAIKIVKLDPGDDISSIQQEITT |
| JAG1 | MLLPRRPALAPLLLALLCLRGKVTRASGQFELEILSMQNVNGELQNGNCCDGPRNPGDR |
| ATG9A | MMHFDTQYQRLEASYSDSPPGEEDLLVHVPEGSKSPWHHIENLDLFFSRVYNLHQKNGI |
| VAV3 | MEPWKQCAQWLIHCKVLPPNHRVTWPTAQVFDLAQTLRDGVLLCQLLNNLRSQSINLKE |
| TNIP3 | MIHCGLLKKKNEDAENALELCTSESPEAERCESIELDDEIRNLIEKNSIAQYHDNSEFIAQEI |
| TMEM173 | MSHETERPSVQTSPIIPKPRGRRALYATWGFIVLCIFALYHMEQSWHHPAEYITFYFTALQIG |
| LAMP1 | MARVGGGRGLFLAAGLLVFLQMSSSAFEVKDQSGEVCIIANLSASFSVEYHTKSGQQLTA |
| ITGAL | MQESGCACERQSKRIWQQVRALGSQASPLLGKKGLLGSKGRVRRLSSSSPTLDGDVAA |
| PROS1 | MRLLSRRCVAFHTCLVVVIALSEARTFLSQQYASEFLVRKRRANSFMEESKKGNLERECIEE |
| ATG16L1 | MTSGLRAATFPTWKRHIAEELRRRDRLQRQAFEEIILQYNKLLEKTDLHAVLADKLQAEKYD |
| GCH1 | MEKGKLGALLEQPPRTVSCNGYALEEKQQQPPQPAPSRALEKPRYSQGTSSQCLDSWKEI |
| CTSG | MLILILLPAAFLLPSGAVAGEIIGGREAKPHSRPYMAYLEIQHGDNSSFCGGFLVLENFVLTA |
| COLEC12 | MRMGVPSSFAGIQEGTQCTKCKNNWALKFSIILLYILCALLTITVAILGYKVVEKMDNVTGGI |
| CD97 | MARLCIALCLWGTVAQNNSTQGTTEDCNIHIQCPANAKCVNNTHCTCLDGYQSTGNRF |
| MASP2 | MYGRITSPDFPNVYPNSKERTWNISVPKGYTIRIYFTHFNMELSYLCEYDYVKISSGGKLVA |
| SEMA4A | MPSRALCLLCSLVMPAAFLCADLLPRVTFPKGDARRILTSFSKDGIFNYDTFLLSKDEGTLY |
| LPXN | MDDLDALLAELEQSSKEPKGDYCLLTHSSSEQRPAAAVPQTGTVQNEKLVTSGTHKVQPAL |
| THEMIS | MFGNECCLPTGELIKVTGFKINKIIASICESNEDSQCSARIELPLNFPGLCKIVADKTPYLSIE |
| ANG | MHRAVTPATHKAWSPFWAEARGQTWDHLTGAMALKMTPMLLLLLASALVLLRVSASQEQ |

| | |
|---|---|
| APEX2 | MRLLSWNVAGLRAGPGPAGLGALDGDVICLQETRVTRDLLTEPLAMVDGYNSYFSFCRT |
| OASL | MELYQVPAGSLDAWIAEHLQPSEEFQQQVKDTVRRICDFLKETCFDDIKVFKTVKGGSAG |
| EXOSC3 | MRLRTDVTHSLPVICAFCARGPLSRTRIRLREASRPEEGPDPAMAAAETGALPAEERVGQV |
| CFP | MEGALTSSLLLALCWALAPPADGQEGLCYEQFGAGRCEELLGEDVPQADCCLNPSYGY |
| POU2F2 | MSVQGQRKVQASVQGQGKRQIKAEDPSTLPPAPAPAPPAQQPHLPQAQLMLAGSQLAGL |
| ART1 | MSTRPQSEEQPLLHRSWGKQRRPNYSPPASGARRTQMWMAKMERLRPVLVLLLAGIFM |
| IL18RAP | MFTFCWILLLFIKGAEVREFNLTGCLHKEPERIYHAITDEEFILECVLPSQHRTYMYNNSFL |
| ATF6B | MAAVAPELLVLHDSGRFLADNLLSSEDWDAALYSCLDNGEEPAALFPCLEPSALFDDSFD |
| CD5L | MEALSGKVFSQGNESHPILMTNVQCQGTEAVLGSCPYEDPGDLCQHDEDTGARCQEV |
| EGF | MFLILTAFLPVFFKIDFVSLSALPHWNCPRGYRLGNENSTCIDIDECLEGGLGTCGQTCV |
| ITCH | MLRGGIIQGPAHIMAGSGPRSGTVGGFAMKSQLQITIISAKLKENKNKWFGPSPYVEVSVD |
| IL1RL1 | MLIHALLVQLELGQGYICRSPQRTVMGLLHLILVSTFLSVSMTSESFHAMEDEALVVKCPQ |
| TAPBP | MISPAPVGALALLLPGFTLCLAAAGPPRLSCWFVEEAGGGMGVMPSSVTQRRALLLLREVR |
| C5 | MNTLSFICILAFCGRSLSQEQTYVITAPKIFRVLASEKVVIQAFGYETAFPVSISIKSFPDKRTTY |
| CYP27B1 | MPQTLKLASRASLLTRGFPEICAELGCKAPARVIKTHKTLEDMPGPNALGFFTDLFCKKGLA |
| GZMH-immu | MIQGNWSLASHDPGKGLLGTAQGERSPAMPVTPPEWTQAGDSCFPPSLPADKSISDSR |
| ERBB2IP | MTTKRNLFVRLVPCRCLRGEEETVTALDYSHCSLEQVPKEIFTFEKTLEELYLDANQIEELPK |
| MSH6 | MSRQSTLLRFFPKAQAPGVGPTQAPSPAAAARRRGWASSEAAGDEAVLAPGGGAEEMGR |
| ABL1 | MSRNEINEKLIMGAKNRWLLSARLSLFFSFFSSPEALQRPVVSDFEPQGLSEAARWNSKE |
| SH2B2 | MNGDALCQESSSQLPDWREFCELHAQAAAVDFAQKFCQFLKENPHYDTPGAETSFSHH |
| BMP6 | MTVTILRFTVFQDRVCHNVLSFNVLVEYDKEFTPHQQHHKEFKFNLSQIPEGEAVTAAEFR |
| PLCL2 | MAEFGSRGAATGSTRGGGLPSSPSPVQGKVAFKAGDGEGGGGGGRSCVDSAGGGVS |
| SPPL2A | MSRMRRAWLLWAAFWALLLPRVNAQEGILHASGNGIPPLTKDYCIRYNPSWVSLPNTLD |
| HAVCR2 | MAPYFILQWILMVLLTGLSVSGSPVRGVVGQNVTLPCKYRVNHQNDITTMCWGRGSCPS |
| ATF7 | MQNDLRNHKAEDARFTNEDHLAVHKHKHEMTLKFGPSRTDSVIIADQTPTPTRFLKNCE |
| MR1 | MGLALRLLLLLLGAVALRGGRSRLHSRRIFHTVVSEPGPGLPWYSRIGYVDDQVIFNYTSE |
| APBB1IP | MEQTCDDIDEMFSNLLGEMDLLTQSLGVETVQPPSPKESNKEYNFTVGFKDLNESLNALE |
| PRKCG | MALVASPTELECLTKPLFCRKGALRQKLIHEVKSHKFTARFFKQPTFCSHCTDFIWGIGKQ |
| AQP9 | MNGQSKRTFKEKFALKNSFIKEALSEFLGTFVMIAASRSYKLSSKISYTSWDQHAHVLPFPS |
| TINAGL1 | MRLLWTLPTLWLLAAEDLSAHRARTRRELAPGLHERGIRDAGGSYCERNDACCAGRDD |
| CD200 | MRSYEILQVTWQKKNTGVNENLATYSLSKGVNIVETYRDRLNFTRLSLNDTAITLWRARIQD |
| LY9 | MELSRIVGDSVTFPLGIPAQQFITVSWTVNTTRIIVTVGAGTPPNVIVSDPSYEGRLRVPESN |
| SEMA4D | MLLSNDDKDVLYVGAREVIFALNSVNIAEKHHELYWKVTEDKKTKCADKGKSKQTECLNY |
| ITGB7 | MTGAAWPLGLGPGQSPGHWDPPVPGASFKPAMKGTGLVVGLGLLLAVGQGEKVLEAGS |
| IL31RA | MRNAYISPESPEIERGSTLKLLCILENYDMPYRNASHVIWKLNQNLIAQENYNIINETVAGV |
| CD22 | MVLIQIASVILALLSLVLLVVALVSDNWVKVNEAEESHLGLWKICVSQICILRLTRVPGFFHTS |
| Ref | wt_sequence |
| KIAA1549 | MYYCNKTFVFYFSDGVSLEQQNLSLLSVGSKLHQSTVGSIAHMALTQSTVEPQENNFLHV |
| ARID1B | MIMAAQVAAAPAGAKNNKGKTPSLPGNLSQDLSSGGAAAAATGGPGPGSMLGVGDGTN |
| GOPC | MSGMGAGSCGSGSGACGGGVAAVASSTGGGVSMFRWLEVLEKEFDKAFVDVDLLLGE |
| ROS1 | MEGELGINACNVSDISAACTQGCQFWNATVQKSCPLQCNQTYIRACERVSCKFGCSCA |
| RB1 | MPPKPPRKGGAAAKGQRASPDRCTGGTSPPAAARLEFGETDFVALCDTLRVSDVVRANA |
| CBLB | MANSLNGRNPGGRGGNPRKGRILGLIDAIQDAVGPPKQAAADRRTVEKTWKLMDKVVR |
| BCL9 | MHSSNPKVRNSPSGNTQSSPKSKQEVMVRPPTVMSPSGNPQLDSKFSNPGKQGGSTSQ |
| FLT3 | MTVSVFLFGFAVLAITAMAFTSATSQHISEITCMLINQQSKDSRTLESAVFLEIPSISEDLGCD |

| | |
|---|---|
| APC2 | MIDYSSARSYDPIATGYPTLEELKMSGSIASYDQLVRQVEALKKENTHLRRELEDNSSHLSK |
| CASC5 | MDRIYSESNEENDHTERIRRQRRSSILKAPRSPLRDLGSGNEFTQDYDAEKRRKNSRRVS |
| NRG1 | MSEVSAETFPHSSTQLNPAASLEGLPAEDNMSGMQTEDGRVHAIAGLAATCCVCLEAERL |
| CEP89 | MALSFRRGRKGPFKHIAHGLVPAATIAPKPAVPRTPPPRSPNPSPERPRSALAAAILVTTLTG |
| NDRG1 | MLLKFIFISKEDNPFCNAAKGDAHRGGGSRARYQLLTMSTELQDAHLAEVKPLVEKDEAIT |
| ALK | MISHGSSCYVWTCVPCKIASFLTTLVLCCFLLLCDETRWVLLQKRVGHLKKPFRISLEYVAS |
| NTRK3 | MLAASERKARATGVLSLVSCLGTGLGWLGASQGMQRLVFERRMCYDESRNLSDNRLTTL |
| CIC | MKPARKAASSSSSSSSSSNSRSPPSTRAKTQKRRTAEGGPKSAGEEEGIRGGGEESPAQPL |
| DAXX | MAATVESNIIVLDDEEEGLPPPMPRESPGPAPASPQTNGTPTKKPDPLAQHGGSFKAENER |
| LATS2 | MRPKTFPATTYSGNSRQRLQEIREGLKQPSKSSGQGLPIGPGSETSLDPKILIGKDAARQQ |
| NOTCH1 | MQQLLVQSLLLSLLPVLSRGLRCTQPAESCLNGGKCETYSNGTGVCLCNSAHVGDRCQL |
| BRD4 | MSVESGPGTRLRNLPVMGDGLETTQMSTTQAQTQPQQGNTVGTNPPPPETSNPNKPKR |
| RNF213 | MKCPKCDYISQDASPNFCSSCGFRLVPHISAHSKEKVVGSGNVKPAPVPEWEVESGREL |
| THRAP3 | MSKTNKSKSGSRSSRSRSGSRSRSRSFSKSRSRSRSVSRSRKRRLSSRSRSRSYSPSHNR |
| SETD2 | MGDFYDPEHPTPEEEENEGKTENVQKAGFVKGPVFKGVASSRFLPKGTKTKVNLEEQGR |
| CLIP4 | MTIEDLPEPSQEGDSLIERYQFLFPGSETSVTFSVAAAPMPSDCEFSFFDPNDAACQEILF |
| PTPRB | MELGGAGDHWVQLYADESQSIIRNTSIQRGAAHVDLDGLVPGTRYRVEIVSQAGPHRTS |
| KMT2D | MPLVRYRKVVILGYRSVGKTSLAHQFVDGEFLECYDPTVESTYNKMLMVGKDEFHLQLVD |
| NSD1 | MEPACEVSRRGGLAPFAGPRALAAPESAGSPGPEPPNGCTAPLPAKAAYGQDPASCYIPL |
| IDH1 | MSSGSLRGYPNLHQALAQPPEVPRHGGDNSPPLNPCAKICTEMILAQREQVDRRSQQR |
| BCL11B | MNFPLGDILVFIEHKKKQCNGTGGVCYEKSLDKSSPPPLSRSELRKVSEPVEIGIQVTPDEE |
| KMT2A | MAHSGRWRFPARPGSGGWGRRGLGGSRLRVPAVRNLRAPEPPPPHAGAGDGELGGP |
| ATRX | MRRGRKAAGGRSSPSSSLRQDGGGAGDGGTSGFVSVQRLSSVPAWPGIPSSGDMTAE |
| NCOA4 | MSPPQDRSGSSCSKESLVKCLQAKKDLETAIAGVVKAEQQMEVNWREVKAQIHSCISRH |
| ERBB2 | MASIVSAPLQSSDHHPHPASLAFPFHPPAVCTGTDMKLLRPSSPENHYATLSRLYQDCQV |
| BRCA1 | MESSVAAIGDVQSVLLAMQKNLECPICLELIKEPVSTKCDHIFCRFCMFKLLSNKKKGPAQ |
| CNOT3 | MADKRKLQGEIDRCLKKVSEGVEQFEDIWQKLHNAANANQKEKYEADLKKEIKKLQRLRD |
| STRN | MDEQAGPGVFFSNNNNNNALLLPPGGAGPGLEPGEAAAGGSLAAAAPAPVSAARAQYSV |
| TET1 | MAHHARPSRLVKKEELGKRKTSEGKKKSSQVKKKAKKNSKKAVSSGKCKKSMQEKNDVKK |
| TRIM33 | MAENKGGGGEEAAEPGPDGGGQEPAATSPSGSAPPAPAVAPPEERDSPCAAAEKGPAEA |
| SETBP1 | MSSPVGPTHKNKAKQVYDRPPKHSALHYDPGLQQDFTSDTLKSKHQQKSSNQHHLDW |
| TRIM24 | MEAAVSEAAGAWPADRAGAAALSGENEAESRQGPAERGGEAAQLNLLDTCAVCGQHIQ |
| CDH11 | MKEDSCLHAALICLGMLYYSHAMSTEKLNPLRPSLHGHHEKGKEGQVLHRSKRGWVWN |
| NAB2 | MALPRTLGELQLYRVLQRANLLSYYETFIQQGGDDVQQLCEAGEEEFLEIMALVGMATKPL |
| RALGDS | MVNRARAERRDAGAGSPENMFESSRRMRSLWDGVRLEVAGEGSSPVVLHSFTQLDPDL |
| ACSL6 | MRILESPHGPTLFLGQHSSMFIKEFALSIFEKMQAQEILRSLRLPEFDNFSQFFHSLPATTLV |
| PPFIBP1 | MMSDASDMLAAALEQMDGIIAGSKALEYSNGIFDCQSPTSPFMGSLRALHLVEDLRGLLE |
| TP53BP1 | MDPSQSQLDLGFSQQDTPCLIVEDSQPESAVVEEDTAYGCLSLLTLPLPNGQSPVLEIMTS |
| MET | MKTVAAFTSGIIVFLFSLVQRTSGQCKEAAIKSEMNVSINYHLPHFITETPIQNIVLYKHHIYIG |
| LATS1 | MKRSEKPEGYRQMRPKTFPASNYTGSSRQMLQEIRESLRNLPKPSDAAKAEHNMGKMS |
| NUTM1 | MTSQPSQGPLLPPLLLPAFPGAALLLTSPNVPGGKLIVKLKPEGAQELPRAQTIILMRAPVGC |
| MAP3K13 | MHMHGTMASPQEHLSWPSSTNLINKTFSEDKDFSGIHGDHAPTGNHPSPEIIEDVREKG |
| AFF3 | MGTVLLKKGNTVVPQPSSRTSGLSSKRSSSSSSSESESSSESDSESESSSSESDGSKPSHYS |
| KCNT2 | MQFLGPAPQHVTSYDPGIQFGKLKLYFLSFYQTGLRIRLFNFSLKLLSCLLYIIRVLLDDPTQ |
| FOXO3 | MIPEEEDDEEEGGSSPMAIGSAVPGGGEGEALAPEEAGRLLAPLAGVGTEGLGQAPGAA |

| | |
|---|---|
| SSPN | MRKEEKKPQNNTSLKNQNQAGKAPEGEKSTDPANQQKPAMKKKAKGDPKTGQEEESHT |
| POT1 | MGLLYYLHVSLHYRTLINPTSKSRASSNFMDLTFARVNCLPVQKKFALDWARDIRSGAVAL |
| PTCD2 | MVRSEDRMAAAVAGGSSRILRETVKRSMLRSATAAFGCWGCPHGAKRYLLTDNVIRLQE |
| GRN | MQLPGPGARWGAGTCLHAHGQGPGSRPRMWSLLPLWLALSGAVASLRCPDGQLCHG |
| POLQ | MTIPDDQADKLLLASWGLPEVVLEKYHSLGVVHMFAWQAECLMLGQVLEGKNLVYSAPT |
| LRP1B | MQFNLMRRKLFFTDYGNVAKIERCDMDGMNRTRIVESKIEQPTAVAVDLVNKYVYWIDIY |
| DKC1 | MADGEGCIAKKQKKKKDKKSLPDADVADIQHTEEFFIKPESRVAQLDTSQWPLLLKNFDKL |
| TERF2 | MAGGRGRVSRSRGGPMAAGSGALGLRAPEEAVNRWVLQFYFHQAVGAYRARRNRDF |
| ACD | MRSAAHPLPAASGGGGDPASRTPRPASAATKPRHVSGLSLCYLPRSTIFQAKNLDRIPESK |
| CYC1 | MAALSAARCLVLRTRGGSLLPPPPGPSLRPPQASMSSLSGMSRGRKVALSALGVLAAGGA |
| LMX1B | MEKIAPTEFVMRALECVYHLSCFSCCVCERQLRKGDEFVLKEGQLLCKSDYEKEKDLLSS |
| AIF1 | MSSQQNLQGGKAFGALKAQQEEVLDSLNKEFLDDPKYSTDEDLPEKLEAFKKKYMEFDL |
| CMAS | MDSAAAAAGAVPAAEAEQRPAHMAALVLARGGSKGIPLKNIKPLAGLPLIGWVLRAAIDSG |
| AGPAT2 | MEPSWLLGGAALLLLLLLPALLALSPAAHFQLKIGFYCVLCVVASALAVPLCLLRHGGRTVR |
| ATM | MLIDKSIFDASKSLHLQVYLVLLKELPTEENLLPEDDFLLLLKPLSDVCSLYRRDQEVCKAILN |
| KLRF1 | MEDEEGYTVLNLRPKQRQPGRPPGAGPQDSRQCPHWIRIALGVGWAGNIVLLAAVIVLG |
| SYNE1 | VTVNKWEQFERDKETVVKYLKQASSALERILSFSSLESLSSELDQTKELSKQTIAIAAQAENLV |
| MRC2 | MENGAGGGGQEQWDRGWKWHSPSCYWLGEDRVTYSDARKMCIDYGATLVTITNRFE |
| MALSU1 | MLPAARGLRLRPAGPGSGAAVRCAPCWAGLAGRALGGGPSTEVRAGGGAEQERRAPD |
| IFNGR2 | MPRPGAALLLCLLLLSLPEPPAAESPLQLPTPQSVTIYSYNYQNLLTWAPVKVDNGSVLYTVQ |
| MINPP1 | MASCFRGPLPGRHSLPLFLLLLRVSLAQERAVATSLLSGYFGTKSRYEEVNQHLLQDPLSLG |
| APOA5 | MSHKAALLVLLLATFSELPYATSLLQRHALPPRLFQGHSVKCSCSPHAAAQAERARNGFWD |
| HOXD12 | MCDRGLYRSGYVGSLLNLQSPDSFYFPNLRTNGSQLAALPTISYPRSSIPWTCSSSCTTQP |
| PRUNE2 | MHILLLQVSPPDILCLPVLNIPRREFSYYPETRFILEQLNIPESFHIFQDEINLHQLNDEGKLS |
| SIRT1 | MADEETLLLQPHSGRRAAVAEPLPKRQRLDSDDSGGRGSGKGPGPERGVGTAPTTSSGT |
| IGF2R | MRRAARSSGRLPAPRLPAWPLLLLLLLAGLRGGGEAQSLDLDPLCSYTWEAIDVDKHVRY |
| PROM1 | MSENSLCFSIDMQQCIGVLCAYAANQHLTSQVRGAKNLVHSNFKDLRIFLNDTPGLPGV |
| ATG4C | MEATGTDEVEKLKSKFMSAWHNMKYSWVLKTKTYFSRNSAVVLLGKCYRFKSDEPSELSP |
| TLR4 | MNLEGLVCSKSQCRFAFVVLKKLWKVLMLLLSMVLSVQVLQYCGRMMPDVPFEKDINIC |
| ADAMTSL1 | MDCPPEAGDFRAQQCSAHNDVKYQGQFYEWLPVSNDPDNPCSLKCQAKGTTLVVELA |
| FGF19 | MWRSLCKPYTSLALLGLCFAMVVRSLPFSDAGPHVNYGWGEPIRLRHLYTASKHGLFSYF |
| CGNL1 | MEPYCAGFKHVQRDCGVHLKLANDETDKARNTQHSKPGSYGVSIRVQGIDGHPYIVLN |
| ITGB3 | MPGSPCSGSAHPRLSCPQPHDSSPRRLKSSGSLSPGLQPLLPPHPCKPASEKMQGLRLA |
| NCOR2 | MSGSTQPVTQNWRATEPRYPPHAMSYPVQISRTHADVGLLEYQHHPRDYTSHLPPGSIM |
| MSRA | MFSRWQEGGKSLAPLRCEISSSCQPPPCLVFCRLLSSCPTPSLCPLPGVQKLPEVVGTRR |
| TRIM25 | MAELVRAPSVPSLAGLEEELTCSICLCLFDSPVTTPCGHNFCGPCLDITWSGLLQGFSCP |
| F7 | MVSRHYLVLLLCFLLLLPLFPAAVFIKQDQANSLLKRQRRANSLFEELKIGSLRRECFEETCS |
| TSFM | MQRAALTGVCGPLRGGARPGPHPVGLSCMPLQLLSSPCQQVVSWQPVCLFHTGLRAFA |
| SORCS1 | MGSSKEPDLVHLETKTVDGHAQYIICRMQNSTEATRSKPFPGYIDPDSLIVQDDYVFVQLT |
| AOPEP | MTENLQSLCLANWNMEMKLDPAKDDLPLMANTRHMLVKHYVLDLDVDFKSQVIEGTVV |
| NFKB1 | MAEEDAYIMGVSDHQMFAMDQFMGVNALFNNTGFITSDLPLRTDGPYLQIIEQPKQRG |
| ZBTB20 | MSALGPRLGVIYLKSDTIWKPLKKPKTAENQKASEENEITQTGASSAKPGLPCLNLEAVPPLS |
| FBXO31 | MAVCARLCGVGPARGCRRRGAQRAQQREGPADSEPDTDAEEERIEGARLGQGEALAA |
| BSCL2 | MMSSTGPFLQWAQEVAALLLLRVRRTVLQTAILLCVLLLLLWIAVFLYGSFYYSYMPTVSYVS |
| PPP1R1A | MLLQIISNWSCLSVGQKIDEDRLPNPLLKPGLSMSPRQRKKVPRTTPTMKELQMMVEHHL |

| | |
|---|---|
| VWA5A | MMSCGLLNSSNEPVPLRSVSVTVLIRGFVADVGCELLYRNEEQGPVEAVFVFPVDAEAAVY |
| METTL1 | MAGAPPQKRFYRQRAHSNPLADHTLCYPTKPEEMNWAEHYPEFFAPLTKDDRHDDPKD |
| BNIPL | MVPSIKKGRWWGGRRWGLSAALAAWEDSFPVSSSHLTPCIQEVLTHQEFILAIEATLRME |
| PIK3C2G | MKQISISTVDKVIEEAKAICSVLCSVETRAITDAVKKLRSVPLGKTQNSLQNVETPFRVLLENA |
| SVEP1 | MASHPKEEHCYLLHSFAEFEALARRALHEDLPSGSYIQEDISHCSYLCEANENCCDIMAS |
| RAPGEF3 | MHLFRSYSYQIQLEGEAGSLPRIQGIRWTPLPDSESAPDCEQSPTQAHAEKVWRAGRLLF |
| XRCC6 | MSEWMSYYKNEEDEEEEEYPEEVETGGEYKYAGRDSLIFLVDASKAMFEYYDDDDEWTP |
| IFNAR1 | MAPPELLLLLLLPLLLAPLAAGQMNLTSPQNVQVNVVNTNFTLRWSWGSENDFNVTFSAQ |
| CYP19A1 | MILETLNLMHCNITSVVPEVMPTATVPILLLVCFLFLIWNYEETSSIPGPGFCMGIGPLISHG |
| METTL21B | MAASRQPLLCHVEFGGAVAGREPPEAAPEGLEAVFPREVGLFADTFPEETRYGFCGHVLTI |
| HIF1A | MELTGEKKRISSDRRKEKSRDAARCRRSKESEVFYELAHQLPLPHTVSAHLDKASIMRLTIS |
| MTFMT | MRGRLLRAWREGAVGVGAAARRGLAELGASRGKPPWRVLFFGTDDFAAESLRALEAARK |
| GZMB | MLILILLPVALLLPPRARAGEIIGGQEAQPHSRPYMAYLEIQRKGKTYICGGFLVSKNFVLTAA |
| GHSR | MRNQSGSPNSSARVPDYENDTWPEYPVHLFPAPVLTGITVTCILLFVVGILGNLMTMLLVS |
| BAG6 | MAEAQNLEVMVKTLDSQTRTFTLEAEVTVKDFKEHIAGSVNIPAEKQRLIYQGRVLQDDKKL |
| ZPLD1 | MERIWLLLLLIIQVLSVTAQFNGYNCEANLHSRFPAERDISVYCGVQAITMKINFCTVLFSG |
| SYT13 | MVLSVPVIALGATLGTATSILALCGLTCFCKCRRPRKGLSGKDQDMDTENAKPSMLQPVQ |
| POLM | MALVPLKRRRRAPSPAAAPLGAGRFPAVVLCLVEKRMGASRRAFLTQLARAKGFRVDGAY |
| AKT1S1 | MADNHRESWAGLIAEAERYRRRTGNEVVLITAFRAPQPGGFAYAQHGCGALGDAARRYL |
| TEP1 | MPPRAQPEPEPGLRDRSQQGGLGPGLLGSPRSGLKSEPGSGAGPAGASHQRFRAAAGR |
| CCL5 | MKATVAALAVLLVAAFCSLASPAPLGSDTTVCCFSYTTRKLPQSHLKEYFYTSGKCSQPAVVF |
| ENOX1 | MLWMEQFLQSHRWQNYSICVPGFDPSLSMMTGITPINPMIPGMGLVPPPPPTDVPVVKE |
| RBPMS | MNSLPADKESGAAEPGLPEEEVRTLFVSGLPLDIKPRELYLLFRPFKGYEGSLIKLTSKQPVGI |
| MRE11A | MSCINSGDEGDTFKILVATDIHLGYLEKDPVRGNDTFVTFDEILRHAQEKEVDFILLGGDL |
| CPA2 | MRLILFFSALAGASWCLETFVGDQVLRVKANSEEEINKLQLLETLEHLQLDFWLHPSNPAL |
| NAV2 | MNKVEIYTDWANHYLAKSGHKRLIKDLQQDVTDGVLLAEIIQVVANEKIEDINGCPKNRS |
| LINGO2 | MLHTATSFWQPFLGLAVLLIFMGSTIGCPARCECSAQNKSVSCHRRRLISIPEGIPIETKILD |
| ZFYVE28 | MRPIHPFSFQDNVLNIVNQIMDECIPHERASRDFCVKFPEEIRHDNLAGQLWFGAECL |
| ECHS1 | MAVALSSLRRPAWALAPARLLLRAPARLSSAAGTQFQYLMVEKKGAKRNIGVIQLNRPKALN |
| GZMH-aging- | MIQGNWSLASHDPGKGLLGTAQGERSPAMPVTPPEWTQAGDSCFPPSLPADKSISDSRG |
| PPM1D | MEGLYSLGVSVFSDQGGRKYMEDVTQIVVEPELPAEGEEGPPYKGGSVPGSQREAESRGI |
| POLE2 | MAAEKLRSKVGSAFKMHGLLLRAEANKYLTEALQSVNELELEDVIENIIDAVEKQPLSSNMIE |
| TOP2A | MELDESLSPLRPVHENPKVGQGRPAAAAGKRLSVERIYQKKTQLEHILLRPDTYIGSVEFVT |
| PPP1R26 | MFLMNASPLIALKTKWESFGQSRSCRYPVCFSESEEDVTRASVSAKVQMIINNLQSQESSL |
| ERCC1 | MEPMEAAQVPEGRKRFTVQSQDPEHTMVRPIFKSSCSAAEPRAPALPGATSYAEYVVAQV |
| CSMD3 | MCMGEINLEQCFQTGVAVCVERAPGGPSQFVYLLCPQVHQIVALIGHGSLLQANGNCW |
| CRLS1 | MHRRGAEEPPAATRSWYVLVKKALQYENPWTIPNILSLARIGLAPVLGYLIVEENFNIALGVF |
| APOA4 | MAELRGPCGELSPAGRRGALPGGTGGGAESQTDLVCPTLPCDSEGLNTWWGRCRNSR |
| APOB | MGPPQLWQLLLLLLLSSDVLTQDGPESRNSACSKDATRFKHLRKYIYNYEAEIASGVKGTA |
| HDAC6 | MKRGGRRGPQPRRNPRLPETKRRGRAQRMNKDMEQLESGLQRLDLSLEETVPVGTGLA |
| WDR27 | MESPVDLCFTEEGCSHDIVLEKQVFRSKVPESHVQLACNLHYCAFPLNGNELCIWNTGD |
| HIP1 | MEVGKMLMLTYPFSVFQTVSINKAINTQEVAVKEKHARTCILGTHHEKGAQTFWSVVNRLP |
| CEL | MEPRILKWDVVFSHLVGTLKAKEYKNRCMQVTLTQTGTRGSEDCLYLNIWIPQDKKGVSTI |
| TERT | MAGSGPFGAVLAMLEGCYSEVLGLEQFVQRLREAAACGPGEAELLQRGDSELYQTFVSQ |
| NCAM2 | MYAKCIHAEFTLYFSALLQVTISLSKVELSVGESKFFTCTAIGEPDSIDWYNPQGEKIASSQR |

| | |
|---|---|
| LEP | MRCPSLPLCGLLCIWLPLFYSRPVKIDKVKADTKNLTRTIIARIQEHQLFPLNLKINGLDFIPG |
| PARVG | MEDAVVTEELEINPFKVLGVLICKRAKSNTMEPDFFTAFAQPGVLHHSPLKNDAIQGEKRK |
| XDH | MASERTTDQLVFFVNGKKVVEKNVDPETTLLTYLRRKLGLPGTKLGCGEGGCGACTVMIS |
| APOM | MLQRSWSYLLYLYGALISTLSPCAPPAPLPASGLDRDQFPRQYLGTWHFIAAAAATPAALEI |
| CHIT1 | MGRVFVWIGLAVLLFLQCSSACKLVCYFTNWSQNRPAQGRFLPEDIDPNLCTHLIYAFAG |
| LMNB1 | MAASTPVGQRSRGSAASTPLSPTRISRLQEKEELQHLNDRLAVYIDKVRSLETENSVLQLQV |
| TPP2 | MAAAAEEPFPFHGLLPKKETGAASFLGRYPEYDGRGVLLAVLDTGVDPGAPGMQITTDGK |
| POLE | MVLRNSGRRRAEPAGGDPDTPRDDGSGLSALKRLERSQRTDRLDAQFGFERAKEPGEK |
| GSR | MAAAFDYLVLGGGSGGLASARRAAELGARVAVVEHQRLGGTCVNVGCVPKKVMWNTAV |
| HLA-DRB1 | MGAGWILGAGSCWAGALLVTLTVLRTHLAHCTEPPRGFVYQTNSECYFTNGTERVRFLER |
| HRH1 | MCETKNSTEAYPIPLGMALGSISLITIIMNILVLYAVKTERKLHTVGNLYIVSLSVADLIVGAAV |
| SH2D4A | MLQQILADMYIDPELLAELSEEQKQILFLKMRQEQVRRWKETEAATDKNSAKKVTQRKAHG |
| NEDD4 | MARRLRLHFAAQRSNTDPLSEISGDGLDGNVLVYFQRPSRNSAPDIIQIKVTPRQAIFPTV |
| MARCO | MENPKNYNEDGISNFTKMLTMSDKIRFASSGTTTFELHVFNMQEEMRKLQREDNSHLAE |
| MAPK1 | MAAVSGAGAAGGSANPGGPEMVRGQVFDVGPRYINLSYIGEGAYGMVCSAYDNVNKV |
| CFH | MLPTDCETPPPRRTKEILTGTWDKASYPHGTSAIYNCRPGYIKLGRVTFECRNGLWHHVH |
| GCNT3 | MFRERKYLCQIQRLLFMGFLMLFATVVLKYSSRECNFSDLGKETVHLKSEGCKDHLYQSL |
| DHX33 | MPQGGEVPPAKRLKLAPPPRRAMLLAPPGAKRQPPAPSAGEAQRRSLPIFQARGPLLSQLR |
| NFATC1 | MPSPGFPTSKHFLAGPPDPARRRETLVALQRSSCPFSSSSSSSMKSAEEEHYNYASSNINA |
| PLEKHA1 | MPYVDRQNRICGFLDIEENENSGKFLRRYFILDTKEDNLVWYMDNPQNLPSGSPPVGAI |
| PUM2 | MSIPCVLLGLNEVAWQETRGMMHANIGQEASGVGIGVAPAGTVGGATAGPGPGTSGPS |
| TREML1 | MGSFPAWLLLLALTEPCLAKEELELVTAVQGSPFSTKCYYNDTYSPKEKFWCKVLSDQECN |
| TNFSF8 | MCFSPEKTLFQANDSHEAAMTMTEASVSRRLNVGNKTYFYFTTASLTVCLVFALATIMVLIV |
| PKN2 | MASNTAEREILLTDLQGDARSLLVSENVSIGQKLDFSDTMVQQKLDEIKDQIKREIRKELKI |
| CD302 | MEPPFPALLCLLGLLATASGAAECPSSAWIPFRSSCYIFLQGTLEELDNIDEARNLCKDNAS |
| IRG1 | MLWAGEVLLAKSRPLFFPQTITENFANVIHGLNASHLTDLVIQRSKRMILDTLGVGLLGTST |
| MID2 | METLESELTCPICLELFEDPLLLPCAHSLCFSCAHRILLSSCSSSESIEPITAFQCPTCRYVISL |
| DHX9 | MGDVKNFLYAWCGKRKVTPTYEIRAVGNKNRQKFMCEVRVEGFNYTGMGNSTNKKDA |
| C5AR1 | MSVNEVSEDYDLDSYVNFTAPSFDMPTPTPQTFQLTPILWVSLVLYCLVFLLGVPGNAAVI |
| IL1RL2 | MGEECMVHDVLHSQNLVSVLVGEPLTIDCLLDKTLALQHMNYSLTWYKSGRKMPVTQVK |
| ATF6 | MAAEEVEQGTAEPGPLPPGVGSDWESVLFDELSDLADTDETQLDVDNEIYENSLYNLDFD |
| ST6GAL1 | MVHINLLKKLLCGILLLLVVVTICLWRETRKGYYIPFKTESKSFQIPRALGKGDVLTSHLLSK |
| LY75 | MRRAPGRSLATCWVALLAACCFACGASAGNGAFTIRHEKTSKCLQVKNSQIMAADCKET |
| RABGEF1 | EEMSLKSERRGIHLDQSELLCKKGCGYYGNPAWQGFCSKCWREEYHKARQKQIQDD |
| CD1D | MGNTKVRPKVKISPMKSGSQSHFHLLVCTVTGFYPRGIKIKWLKNRQEQTAGVVSTELLQN |
| ADAR | MNTGFGRGKGSYQTPPKHIYCSPNPAFFNNSLVPQENNRDRFRNQQVQFLLGQITEAP |
| ATP7A | MAQGLVTVGGGANVGQGHRGRSRPAAPRGAGCQRCCCGEGAQLADRCWECEDSS |
| PSMD5 | MAEEALELLARVSRLEEPLEELRALGVAVQALPLSALRERAPDIRLGSLFALLNVNDREQVAL |
| TRIM8 | MAENWKNCFEEELICPICLHVFVEPVQLPCKHNFCRGCIGEAWAKETGLVRCPECNQA |
| PIK3AP1 | MASAGAPGGCDVLIVYARDASEWCQYLQNLFFSCRQVRKQRVLSYEVKYGGAIPWEEFD |
| APOBEC2 | MAENQEKPRDTQTGDEQPENSEESEEKKKELEKLPPFEIVGGERFPAFFFKFQFRNVEYSS |
| DAPK1 | MTVFRQENLDDYYETGEELGSGQFAVVKKCREKSTGLQYAAKFIKKRRTKSSRRGVSREDI |
| STK11 | MDVQEPQELSMFAESELMSVGMDTFIHRIDSTEVIYQPRRKRAKLIGKYLMGDLLGEGSY |
| NFKBIZ | MIVESNREAPADGASGSMSSPLNLAYFYSASPYSSDGSCSPAHSAPGSPGSDSELSVCSG |
| TRAT1 | MTIVLLSQILFEIKGEVLLQNSNICTEMEFECHYSIWGLLAFMTMALIVSLVINISYCIANKQK |

| | |
|---|---|
| IL17C | MSLLERLLGYMHLSSPISVHQGLLCALLLFSLPAGQAQKRAKDHQRHHHHHHCFSQEQL |
| CD38 | MPLQNRSSKTRQRTILLVIIMVLIVTIVAVLAAVLLANGMKETSMQLLQWKGRGTTKNLQEVII |
| FGA | MEHGIQSDCKQEKSWPFCADDDWGHKCPSGCRMQGLIDETDKDFHHRIQRIRELLLD |
| IFNL3 | MKKCHLSKYKSLPPWELKTFKNVKDRFEDIMLLSDRKCNTKIFHRNWEVKELSVHDRVILV |
| DHX58 | MLTPALPPSYWLCASCRCPVPARMELRDYQWEVILPALEGENIIIWLPTGAGKTRAAAYVCI |
| MADCAM1 | MEPTLLLLLLSLGWTCSGRLTVLPQEPLVQIGGSIQLNCSLDCPDGKPQWKGLDTNLGNII |
| MARCH | MPLHQISVIPARDVTSSRVSRSKTKEKEREEQNEKALGHSVSRSSNISKAGSPTSISAPHSFS |
| CD207 | MSRENICDNMELAEADPWPKGNPPKTSVPWRHRERSIVTAVLLLLCLALATSLLAVTLLYLR |
| MAP4K2 | MSLLRDVSLLDPRLRYELIQRIGAGTYGDVYKARDTQTAELAAIKIVKLDPGDDISSIQQEITT |
| JAG1 | MLLPRRPALAPLLLALLCLRGKVTRASGQFELEILSMQNVNGELQNGNCCDGPRNPGDR |
| ATG9A | MMHFDTQYQRLEASYSDSPPGEEDLLVHVPEGSKSPWHHIENLDLFFSRVYNLHQKNGI |
| VAV3 | MEPWKQCAQWLIHCKVLPPNHRVTWPTAQVFDLAQTLRDGVLLCQLLNNLRSQSINLKE |
| TNIP3 | MIHCGLLKKKNEDAENALELCTSESPEAERCESIELDDEIRNLIEKNSIAQYHDNSEFIAQEI |
| TMEM173 | MSHETERPSVQTSPIIPKPRGRRALYATWGFIVLCIFALYHMEQSWHHPAEYITFYFTALQIG |
| LAMP1 | MARVGGGRGLFLAAGLLVFLQMSSSAFEVKDQSGEVCIIANLSASFSVEYHTKSGQQLTAN |
| ITGAL | MQESGCACERQSKRIWQQVRALGSQASPLLGKKGLLGSKGRVRRLSSSSPTLDGDVAAQ |
| PROS1 | MRLLSRRCVAFHTCLVVVIALSEARTFLSQQYASEFLVRKRRANSFMEESKKGNLERECIEE |
| ATG16L1 | MTSGLRAATFPTWKRHIAEELRRRDRLQRQAFEEIILQYNKLLEKTDLHAVLADKLQAEKYD |
| GCH1 | MEKGKLGALLEQPPRTVSCNGYALEEKQQQPPQPAPSRALEKPRYSQGTSSQCLDSWKEI |
| CTSG | MLILILLPAAFLLPSGAVAGEIIGGREAKPHSRPYMAYLEIQHGDNSSFCGGFLVLENFVLTA |
| COLEC12 | MRMGVPSSFAGIQEGTQCTKCKNNWALKFSIILLYILCALLTITVAILGYKVVEKMDNVTGGI |
| CD97 | MARLCIALCLWGTVAQNNSTQGTTEDCNIHIQCPANAKCVNNTHCTCLDGYQSTGNRF |
| MASP2 | MYGRITSPDFPNVYPNSKERTWNISVPKGYTIRIYFTHFNMELSYLCEYDYVKISSGGKLVAT |
| SEMA4A | MPSRALCLLCSLVMPAAFLCADLLPRVTFPKGDARRILTSFSKDGIFNYDTFLLSKDEGTLYV |
| LPXN | MDDLDALLAELEQSSKEPKGDYCLLTHSSSEQRPAAAVPQTGTVQNEKLVTSGTHKVQPAL |
| THEMIS | MFGNECCLPTGELIKVTGFKINKIIASICESNEDSQCSARIELPLNFPGLCKIVADKTPYLSIE |
| ANG | MHRAVTPATHKAWSPFWAEARGQTWDHLTGAMALKMTPMLLLLLASALVLLRVSASQEQY |
| APEX2 | MRLLSWNVAGLRAGPGPAGLGALDGDVICLQETRVTRDLLTEPLAMVDGYNSYFSFCRTF |
| OASL | MELYQVPAGSLDAWIAEHLQPSEEFQQQVKDTVRRICDFLKETCFDDIKVFKTVKGGSAG |
| EXOSC3 | MRLRTDVTHSLPVICAFCARGPLSRTRIRLREASRPEEGPDPAMAAAETGALPAEERVGQV |
| CFP | MEGALTSSLLLALCWALAPPADGQEGLCYEQFGAGRCEELLGEDVPQADCCLNPSYGYR |
| POU2F2 | MSVQGQRKVQASVQGQGKRQIKAEDPSTLPPAPAPAPPAQQPHLPQAQLMLAGSQLAGL |
| ART1 | MSTRPQSEEQPLLHRSWGKQRRPNYSPPASGARRTQMWMAKMERLRPVLVLLLAGIFME |
| IL18RAP | MFTFCWILLLFIKGAEVREFNLTGCLHKEPERIYHAITDEEFILECVLPSQHRTYMYNNSFLY |
| ATF6B | MAAVAPELLVLHDSGRFLADNLLSSEDWDAALYSCLDNGEEPAALFPCLEPSALFDDSFD |
| CD5L | MEALSGKVFSQGNESHPILMTNVQCQGTEAVLGSCPYEDPGDLCQHDEDTGARCQEVF |
| EGF | MFLILTAFLPVFFKIDFVSLSALPHWNCPRGYRLGNENSTCIDIDECLEGGLGTCGQTCVN |
| ITCH | MLRGGIIQGPAHIMAGSGPRSGTVGGFAMKSQLQITIISAKLKENKNKWFGPSPYVEVSVD |
| IL1RL1 | MLIHALLVQLELGQGYICRSPQRTVMGLLHLILVSTFLSVSMTSESFHAMEDEALVVKCPQN |
| TAPBP | MISPAPVGALALLLPGFTLCLAAAGPPRLSCWFVEEAGGGMGVMPSSVTQRRALLLLREVR |
| C5 | MNTLSFICILAFCGRSLSQEQTYVITAPKIFRVLASEKVVIQAFGYETAFPVSISIKSFPDKRTTY |
| CYP27B1 | MPQTLKLASRASLLTRGFPEICAELGCKAPARVIKTHKTLEDMPGPNALGFFTDLFCKKGLA |
| GZMH-immu | MIQGNWSLASHDPGKGLLGTAQGERSPAMPVTPPEWTQAGDSCFPPSLPADKSISDSRG |
| ERBB2IP | MTTKRNLFVRLVPCRCLRGEEETVTALDYSHCSLEQVPKEIFTFEKTLEELYLDANQIEELPK |
| MSH6 | MSRQSTLLRFFPKAQAPGVGPTQAPSPAAAARRRGWASSEAAGDEAVLAPGGGAEEMGR |

| | |
|---|---|
| ABL1 | MSRNEINEKLIMGAKNRWLLSARLSLFFSFFSSPEALQRPVVSDFEPQGLSEAARWNSKEN |
| SH2B2 | MNGDALCQESSSQLPDWREFCELHAQAAAVDFAQKFCQFLKENPHYDTPGAETSFSHH |
| BMP6 | MTVTILRFTVFQDRVCHNVLSFNVLVEYDKEFTPHQQHHKEFKFNLSQIPEGEAVTAAEFR |
| PLCL2 | MAEFGSRGAATGSTRGGGLPSSPSPVQGKVAFKAGDGEGGGGGGRSCVDSAGGGVSN |
| SPPL2A | MSRMRRAWLLWAAFWALLLPRVNAQEGILHASGNGIPPLTKDYCIRYNPSWVSLPNTLDN |
| HAVCR2 | MAPYFILQWILMVLLTGLSVSGSPVRGVVGQNVTLPCKYRVNHQNDITTMCWGRGSCPS |
| ATF7 | MQNDLRNHKAEDARFTNEDHLAVHKHKHEMTLKFGPSRTDSVIIADQTPTPTRFLKNCEE |
| MR1 | MGLALRLLLLLLGAVALRGGRSRLHSRRIFHTVVSEPGPGLPWYSRIGYVDDQVIFNYTSES |
| APBB1IP | MEQTCDDIDEMFSNLLGEMDLLTQSLGVETVQPPSPKESNKEYNFTVGFKDLNESLNALE |
| PRKCG | MALVASPTELECLTKPLFCRKGALRQKLIHEVKSHKFTARFFKQPTFCSHCTDFIWGIGKQC |
| AQP9 | MNGQSKRTFKEKFALKNSFIKEALSEFLGTFVMIAASRSYKLSSKISYTSWDQHAHVLPFPSE |
| TINAGL1 | MRLLWTLPTLWLLAAEDLSAHRARTRRELAPGLHERGIRDAGGSYCERNDACCAGRDDA |
| CD200 | MRSYEILQVTWQKKNTGVNENLATYSLSKGVNIVETYRDRLNFTRLSLNDTAITLWRARIQD |
| LY9 | MELSRIVGDSVTFPLGIPAQQFITVSWTVNTTRIIVTVGAGTPPNVIVSDPSYEGRLRVPESNS |
| SEMA4D | MLLSNDDKDVLYVGAREVIFALNSVNIAEKHHELYWKVTEDKKTKCADKGKSKQTECLNYV |
| ITGB7 | MTGAAWPLGLGPGQSPGHWDPPVPGASFKPAMKGTGLVVGLGLLLAVGQGEKVLEAGSC |
| IL31RA | MRNAYISPESPEIERGSTLKLLCILENYDMPYRNASHVIWKLNQNLIAQENYNIINETVAGV |
| CD22 | MVLIQIASVILALLSLVLLVVALVSDNWVKVNEAEESHLGLWKICVSQICILRLTRVPGFFHTS |

**B** Protein language models evaluaton of normal human genes

| Ref | wt_sequence | mutated_seq | mutant | esm1v_t33_6 | esm1v_t33_6 | esm1v_t33_6 |
|---|---|---|---|---|---|---|
| KIAA1549 | MPGARRRRR | MPGARRRRR | D155N:P196 | 0.56815827 | 0.13537061 | 0.45736825 |
| ARID1B | MAHNAGAAA | MAHNAGAAA | T826A | 0.91868591 | 0.90725231 | 1.32707882 |
| GOPC | MSAGGPCPA | MSAGGPCPA | Q109K | -1.3377624 | -3.856348 | 1.01349902 |
| ROS1 | MKNIYCLIPKI | MKNIYCLIPKI | Y1312C:Q13 | 0.92634439 | 2.63243496 | 1.7125138 |
| RB1 | MPPKTPRKTA | MPPKTPRKTA | H69V:P253A | -2.1347091 | -2.3629174 | -3.9692256 |
| CBLB | MANSMNGRI | MANSMNGRI | L590P:G748S | 2.45987236 | 1.12489915 | 2.52513516 |
| BCL9 | MHSSNPKVR | MHSSNPKVR | M1074V | 0.04656982 | 0.11401248 | 0.13528442 |
| FLT3 | MPALARDGG | MPALARDGG | I21A:V56I:D7 | -4.5167286 | -4.3312438 | -4.4426572 |
| APC2 | MGLLGLLSLL | MGLLGLLSLL | V196M:A154 | -2.1861641 | -1.6606717 | -1.989645 |
| CASC5 | MDGVSSEAN | MDGVSSEAN | H415D:N232 | -3.4055014 | -3.6352692 | -2.1358449 |
| NRG1 | MEIYSPDMSE | MEIYSPDMSE | P49H | -2.118293 | -3.2811284 | -0.9427342 |
| CEP89 | MLLGFRRGR | MLLGFRRGR | Q688H | -3.0332339 | -3.6654406 | -3.1444159 |
| NDRG1 | MSREMQDVI | MSREMQDVI | E38D | -1.3459244 | -1.2072797 | -3.3777874 |
| ALK | MGAIGLLWLI | MISHGLLWLI | G2I:A3S:I4H: | -2.9571358 | -3.1087232 | -3.3170867 |
| NTRK3 | MDVSLCPAK | MDVSLCPAK | F720Y | -0.5141683 | -0.8238664 | -0.3952651 |
| CIC | MKPMKKACT | MKPMKKACT | S1258A:L214 | 2.31638074 | 2.51786566 | 1.9683677 |
| DAXX | MRGSENCGI | MRGSENCGI | S188L | -0.8856511 | -0.7668102 | -1.0536077 |
| LATS2 | MRPKTFPATT | MRPKTFPATT | A325S | -0.4117746 | -0.5848274 | -0.5510826 |
| NOTCH1 | MPPLLAPLLC | MPPLLAPLLC | N2396T | -1.2891409 | -0.9603539 | -1.0540528 |
| BRD4 | MSAESGPGTI | MSAESGPGTI | V997I | -0.8238075 | -1.0777402 | -0.8022606 |
| RNF213 | MECPSCQHV | MECPSCQHV | V1824M:V35 | -1.7812243 | -1.9721115 | -1.4947519 |
| THRAP3 | MSKTNKSKSC | MSKTNKSKSC | H693L | 0.9227612 | 0.53028393 | 1.52836013 |
| SETD2 | MILLMRNLTT | MILLMRNLTT | A1408I:D144 | 0.19061595 | -0.2387407 | 0.04580158 |
| CLIP4 | MTIEDLPDFP | MTIEDLPDFP | P511A | -2.4076977 | -3.9751675 | -1.9585981 |
| PTPRB | MEAEFYMVIL | MEAEFYMVIL | M489V:V613 | -6.0504638 | -8.1408852 | -6.0328024 |

| | | | | | | |
|---|---|---|---|---|---|---|
| KMT2D | MDSQKLAGE | MDSQKLAGE | P1165L | -0.6091868 | -0.7558767 | -0.9885426 |
| NSD1 | MDQTCELPR | MDQTCELPR | A480T:L2513I | -1.6715519 | -1.9027128 | -1.5395658 |
| IDH1 | MSKKISGGSV | MSKKISGGSV | G204S | -1.3525302 | -1.103029 | -1.674109 |
| BCL11B | MSRRKQGNI | MSRRKQGNI | L127R | -0.8546109 | -1.4207979 | -0.579602 |
| KMT2A | MAHSCRWRI | MAHSCRWRI | S570R | -0.4927467 | -0.5282075 | -0.5551405 |
| ATRX | MTAEPMSESI | MTAEPMSESI | R1131Q | -0.7331851 | -0.9560962 | -1.1045108 |
| NCOA4 | MGAQLTTFAL | MGAQLTTFAL | S511R | -1.4340262 | -2.0318582 | -1.7542645 |
| ERBB2 | MELAALCRW | MELAALCRW | T166M | -5.4280338 | -5.4428792 | -5.2057171 |
| BRCA1 | MDLSALRVEE | MDLSALRVEE | M1083R | 1.3450551 | 1.3088398 | 1.31137609 |
| CNOT3 | MADKRKLQG | MADKRKLQG | A442I | -1.81148 | -1.956079 | -1.5161443 |
| STRN | MDEQAGPG\ | MDEQAGPG\ | D226H | -1.0912147 | -2.7924948 | -1.1226907 |
| TET1 | MSRSRHARP | MSRSRHARP | L1329H | -1.1254244 | -1.1552045 | -1.1282465 |
| TRIM33 | MAENKGGGI | MAENKGGGI | G877E | -0.3555288 | 0.89571595 | -0.2091408 |
| SETBP1 | MESRETLSSS | MESRETLSSS | L556I | -0.9343429 | -0.9558482 | -0.4941218 |
| TRIM24 | MEVAVEKAVA | MEVAVEKAVA | Y540S | 0.47321177 | 1.0257287 | 1.40651894 |
| CDH11 | MKENYCLQA | MKENYCLQA | M382V | 2.9765172 | 2.60068655 | 1.94352829 |
| NAB2 | MHRAPSPTAI | MHRAPSPTAI | G494R | -0.1374671 | 0.23674631 | -0.200263 |
| RALGDS | MVQRMWAE, | MVQRMWAE, | D41G | 1.47853601 | 1.62599659 | 0.39746428 |
| ACSL6 | MLTFFLVSGG | MLTFFLVSGG | V519L | -5.5464697 | -8.3445654 | -6.662118 |
| PPFIBP1 | MMSDASDMI | MMSDASDMI | Q249H:V288I | -5.4741936 | -9.6666956 | -3.5793451 |
| TP53BP1 | MPGEQMDPT | MPGEQMDPT | M721V | 1.21791148 | 1.16629887 | 0.40806961 |
| MET | MKSKSKSLAE | MKSKSKSLAE | S122Q | -0.8758252 | -0.6908281 | -0.7083094 |
| LATS1 | MKRSEKPEGY | MKRSEKPEGY | S1023A | 0.384624 | -0.0712914 | 0.69107628 |
| NUTM1 | MVVTLGPGPI | MVVTLGPGPI | S165T | -0.3256001 | 0.13989592 | -0.0617058 |
| MAP3K13 | MANFQEHLS | MANFQEHLS | I745L | 1.60552382 | 1.62463307 | 1.18247128 |
| AFF3 | MSKKLITCLSI | MSKKLITCLSI | R978K | 0.99247289 | 0.94544125 | 0.96801543 |
| KCNT2 | MVDLESEVPI | MVDLESEVPI | I974M | -0.3699148 | -0.4106171 | -0.1280229 |
| FOXO3 | MAEAPASPAP | MAEAPASPAP | A98G | 0.99456882 | 0.84844613 | 0.57848334 |
| SSPN | MGKNKQPR( | MGKNKQPR( | E28K | -0.6977532 | -0.3070512 | -0.4422973 |
| POT1 | MSLVPATNYI | MSLVPATNYI | K440E | 0.98642802 | 0.22716093 | 0.44103599 |
| PTCD2 | MVRDSMAAA | MVRDSMAAA | A316C | -1.5063818 | -4.2727447 | -1.9670663 |
| GRN | MWTLVSWVA | MWTLVSWVA | P430L | -1.069279 | -2.0257127 | -1.8341221 |
| POLQ | MNLLRRSGKI | MNLLRRSGKI | S244I:D2414 | -1.7569904 | -2.2179395 | -1.93274 |
| LRP1B | MSEFLLALLTL | MSEFLLALLTL | T2167M:P294 | -2.295351 | -6.2321978 | -3.8507035 |
| DKC1 | MADAEVIILPK | MADAEVIILPK | E210G | -7.4285874 | -5.5052896 | -6.6383772 |
| TERF2 | MAAGAGTAG | MAAGAGTAG | R322F | -0.4982073 | -1.971648 | -0.7503774 |
| ACD | MPGRCQSD/ | MPGRCQSD/ | S429P | 0.44449639 | 0.70872116 | 0.52379656 |
| CYC1 | MAAAAASLRG | MAAAAASLRG | P38S | -0.6234378 | -0.5053231 | -0.4384776 |
| LMX1B | MDIATGPESL | MDIATGPESL | M196I | -0.1064973 | -0.5103407 | -0.4330382 |
| AIF1 | MSQTRDLQG | MSQTRDLQG | S168G | -1.7150087 | -1.8943981 | -2.2829092 |
| CMAS | MDSVEKGAA | MDSVEKGAA | P43A | -1.073509 | -0.7533555 | -2.2486157 |
| AGPAT2 | MELWPCLAA | MELWPCLAA | P232D | -1.4660363 | -2.1823616 | -1.7073426 |
| ATM | MSLVLNDLLI | MSLVLNDLLI | E1245Q:K145 | -2.2193439 | -2.505863 | -2.4648019 |
| KLRF1 | MQDEERYMT | MQDEERYMT | P202L | -0.828141 | -1.5238621 | -1.5477962 |
| SYNE1 | MATSRGASR( | MATSRGASR( | A2020G:P276 | -7.9793267 | -6.3859808 | -8.3332381 |

| | | | | | | |
|---|---|---|---|---|---|---|
| MRC2 | MGPGRPAPA | MGPGRPAPA | K241R | 0.20470119 | 0.76865196 | 0.45475197 |
| MALSU1 | MGPGGRVAF | MGPGGRVAF | V28S | 0.21666217 | 0.08516645 | 0.42361069 |
| IFNGR2 | MIQQLLMRW | MIQQLLMRW | L242K:S246G | -2.6236652 | -5.9609799 | -1.4266139 |
| MINPP1 | MLRAPGCLLF | MLRAPGCLLF | N424H | -1.592164 | -3.5060763 | -0.9204998 |
| APOA5 | MASMAAVLTV | MASMAAVLTV | A160V | -1.646282 | -1.6387383 | -1.7614985 |
| HOXD12 | MCERSLYRA( | MCERSLYRA( | A215C | -4.9350829 | -4.6727209 | -3.7227006 |
| PRUNE2 | MEEFLQRAKS | MEEFLQRAKS | E78Q:P203L:I | -1.6372051 | -4.4058673 | -3.050884 |
| SIRT1 | MADEAALALC | MADEAALALC | G19V | -1.4521277 | -0.8729548 | -1.8423941 |
| IGF2R | MGAAAGRSP | MGAAAGRSP | K1862S | -0.2378113 | -0.4351761 | -0.6505461 |
| PROM1 | MALVLGSLLLI | MALVLGSLLLI | I169M:V371L | -7.2680544 | -8.1987708 | -2.0681813 |
| ATG4C | MEATGTDEVI | MEATGTDEVI | V245I | -3.8308415 | -2.072629 | -2.7856345 |
| TLR4 | MMSASRLAG | MMSASRLAG | G420C | -0.2384636 | -4.8447151 | -2.8705082 |
| ADAMTSL1 | MGTMARLEA | MGTMARLEA | A1003V | -0.8164368 | 0.14317155 | -0.0329633 |
| FGF19 | MRSGCVVVH | MRSGCVVVH | R63S | -1.778103 | -1.8341045 | -1.6946393 |
| CGNL1 | MELYFGEYQI | MELYFGEYQI | Q27D:C234D | 1.30486608 | 1.24418211 | 1.19570017 |
| ITGB3 | MRARPRPRPI | MRARPRPRPI | A50T:T221A:N | -3.2724488 | -2.9709361 | -2.6426051 |
| NCOR2 | MSGSTQPVA( | MSGSTQPVA( | T946S:H1853 | -0.9427097 | -0.468261 | 0.27008653 |
| MSRA | MLSATRRAC( | MLSATRRAC( | A96S:E177G | -7.4273543 | -6.2269263 | -5.8118072 |
| TRIM25 | MAELCPLAEE | MAELCPLAEE | K332T | 0.06659174 | 2.00424266 | 0.33739185 |
| F7 | MVSQALRLLC | MVSQALRLLC | V24I:K70Q:A2 | -0.5164518 | 2.60595489 | 3.12011528 |
| TSFM | MSLLRSLRVF | MSLLRSLRVF | L143A:I219T | 0.23107445 | -1.2556351 | 1.70519662 |
| SORCS1 | MGKVGAGG( | MGKVGAGG( | G807R | -6.0643835 | -5.5000553 | -5.3993177 |
| AOPEP | MDIQLDPARI | MDIQLDPARI | E61K | -0.3234982 | -0.4100828 | -0.299962 |
| NFKB1 | MKRTTSFRM/ | MKRTTSFRM/ | T212A | 1.96778595 | -0.3398738 | 2.05668354 |
| ZBTB20 | MLERKKPKTA | MLERKKPKTA | N437D | -0.001193 | 0.51978636 | 0.30325484 |
| FBXO31 | MAVCARLCG | MAVCARLCG | G396D:V450I | -1.4762726 | -0.6004418 | -2.7346489 |
| BSCL2 | MSTEKVDQKI | MSTEKVDQKI | R350Q | -0.1640313 | -0.0263252 | -0.3821456 |
| PPP1R1A | MEQDNSPRK | MEQDNSPRK | E49K | -1.7364466 | -6.5457563 | -1.4673872 |
| VWA5A | MSSVCDMSF | MSSVCDMSF | K98R | -2.1557336 | -2.4135861 | -2.7301874 |
| METTL1 | MAAETRNVA( | MAAETRNVA( | Q117H | -4.2752261 | -3.8108242 | -2.316555 |
| BNIPL | MGTIQEAGKI | MGTIQEAGKI | L35F:E43D:E | -75.272571 | -131.38147 | -82.844251 |
| PIK3C2G | MAYSWQTDF | MAYSWQTDF | N426M | -1.2945476 | -1.4244909 | -1.4735508 |
| SVEP1 | MWPRLAFCC | MWPRLAFCC | V494L:T625I: | -6.9680846 | -6.704103 | -6.1820905 |
| RAPGEF3 | MAQREEIKRF | MAQREEIKRF | V906T | 1.00102615 | 0.89562678 | 2.1656189 |
| XRCC6 | MSGWESYYK | MSGWESYYK | Q68W | -6.0374746 | -4.8205347 | -6.8336129 |
| IFNAR1 | MMVVLLGATI | MMVVLLGATI | T68E | -0.0716581 | 1.8702805 | 0.27679634 |
| CYP19A1 | MVLEMLNPIH | MVLEMLNPIH | V158M:K461I | -3.0945404 | -6.9016306 | -3.5510616 |
| METTL21B | MADPGPDPE | MADPGPDPE | H194V:Q198I | -1.1627948 | -4.8653903 | -1.6955018 |
| HIF1A | MSSQCRSLEI | MSSQCRSLEI | S616R | -1.6917436 | -2.1122954 | -1.1611841 |
| MTFMT | MRVLVRRCW | MRVLVRRCW | G16V:S317L | -3.5299137 | -5.4277344 | -4.0192115 |
| GZMB | MQPILLLLAFL | MQPILLLLAFL | K92Y:Q149S: | -6.9028232 | -6.5698714 | -7.2681425 |
| GHSR | MWNATPSEE | MWNATPSEE | R336S | -1.6707393 | -2.4547577 | -2.2798915 |
| BAG6 | MEPNDSTST/ | MEPNDSTST/ | V1055L | 0.65104842 | 0.49916744 | 0.75017381 |
| ZPLD1 | MLLPPPFCLF | MLLPPPFCLF | T38I | -0.6423109 | -0.591311 | -0.4088593 |
| SYT13 | MVLSVPVIALC | MVLSVPVIALC | A159I | -1.5309982 | -0.9629097 | -1.0146022 |

| | | | | | | |
|---|---|---|---|---|---|---|
| POLM | MLPKRRRAR\ | MLPKRRRAR\ | T21A:P146Q:I | -6.4936892 | -2.4697218 | -4.303654 |
| AKT1S1 | MSFEGGDG/ | MSFEGGDG/ | P102T | -2.397162 | -2.4938312 | -2.9879508 |
| TEP1 | MEKLHGHVS | MEKLHGHVS | A1976V:S200 | -2.8633456 | -1.9917806 | -3.2552675 |
| CCL5 | MKVSAAALAV | MKVSAAALAV | A23P | -0.7659194 | 0.38224959 | -1.2055107 |
| ENOX1 | MWRRESSPC | MWRRESSPC | C443S | 1.99824238 | -4.7420707 | -1.8617549 |
| RBPMS | MNNGGKAEI | MNNGGKAEI | N17G | 1.01037574 | -5.1445374 | -3.353101 |
| MRE11A | MSTADALDD | MSTADALDD | N617S:D622 | -5.5219295 | -6.4521022 | 0.34331214 |
| CPA2 | MAMRLILFFG | MAMRLILFFG | N194E | -3.9060349 | -3.6183014 | -3.199707 |
| NAV2 | MPAILVASKM | MPAILVASKM | S1179C | -5.3199463 | -5.5289068 | -4.7572479 |
| LINGO2 | MLHTAISCW( | MLHTAISCW( | I470V:V591I | -2.4627249 | -0.4798442 | -3.1942196 |
| ZFYVE28 | MMNRFRKW | MMNRFRKW | G592K | -1.4396503 | -0.7083044 | -0.5972972 |
| ECHS1 | MAALRVLLSC | MAALRVLLSC | V47I | -1.5738397 | -1.5959144 | -2.5074162 |
| GZMH-aging- | MQPFLLLLAF | MQPFLLFIAFI | L7F:L8I:L11P: | -8.9270529 | -9.3612299 | -8.7381996 |
| PPM1D | MAGLYSLGVS | MAGLYSLGVS | V440I | -0.5973182 | -0.4023066 | -0.393862 |
| POLE2 | MAPERLRSR/ | MAPERLRSR/ | L107V | 0.10704422 | -1.5747212 | 2.53170824 |
| TOP2A | MEVSPLQPVN | MEVSPLQPVN | P681S | 0.23244834 | 0.12210536 | 0.07239842 |
| PPP1R26 | MFLMNASPV | MFLMNASPV | D503G | 1.12145567 | 0.8928721 | 0.85224319 |
| ERCC1 | MDPGKDKEG | MDPGKDKEG | L69S | 0.34839201 | 0.60318923 | 0.48103571 |
| CSMD3 | MKGIRKGESI | AKGIRKGESR | M1A:V1966M | -10.007432 | -7.430424 | -7.1582236 |
| CRLS1 | MLALRVARGS | MLALRVARGS | V298L | -2.4206614 | -6.0840712 | -3.289578 |
| APOA4 | MFLKAVVLTL/ | MFLKAVVLTL/ | A268N | -1.9796941 | -3.6165521 | -3.9444141 |
| APOB | MDPPRPALL/ | MDPPRPALL/ | R2012Q:A42 | 0.83003974 | 0.64892197 | 0.93965626 |
| HDAC6 | MSGANRGR( | MSGANRGR( | N60R | 0.40176558 | 0.70151019 | 0.34319663 |
| WDR27 | MENPQDIFS | MENPQDIFS | E651K | -11.777493 | -3.523026 | -6.5447245 |
| HIP1 | MDRMASSMI | MDRMASSMI | D344G | 1.67059159 | -3.2441642 | -0.3268435 |
| CEL | MLTMGRLQL | MLTMGRLQL | L542M | -0.6231794 | -4.0543909 | -2.1555367 |
| TERT | MPRAPRCRA | MPRAPRCRA | R19S:L593Y | -2.854956 | -2.6637751 | -2.4286342 |
| NCAM2 | MVRSDSGG( | MVRSDSGG( | V163I:R307E | -6.4782581 | -9.8087263 | 0.22629142 |
| LEP | MHWGTLCGI | MHWGTLCGI | L147V | -1.5165672 | -1.1567426 | -1.395516 |
| PARVG | MVPAKEQVP( | MVPAKEQVP( | L339P:N341( | -21.145659 | -26.379897 | -19.605459 |
| XDH | MTADKLVFFV | MTADKLVFFV | T510M | -1.3671184 | -1.9289563 | -0.5390677 |
| APOM | MFHQIWAAL | MFHQIWAAL | V70I | 0.10336637 | -0.0482159 | 0.01784325 |
| CHIT1 | MEKQFERGQ | MEKQFERGQ | D213K | -0.6168451 | -0.1796613 | -1.3722651 |
| LMNB1 | MATATPVPPR | MATATPVPPR | T96M | -5.8964338 | -2.7088282 | -3.1894093 |
| TPP2 | MATAATEEPFI | MATAATEEPFI | A636L | -0.0942411 | -1.0136371 | -0.7455659 |
| POLE | MSLRSGGRR | MSLRSGGRR | V146M:R123 | -4.7300646 | -4.0407677 | -3.863018 |
| GSR | MALLPRALSA( | MALLPRALSA( | S120L | -1.8699265 | -0.3806951 | -1.2767892 |
| HLA-DRB1 | MVCLKLPGG | MVCLKGPGS | L6G:G9S:S10 | -107.1222 | -136.61958 | -130.15496 |
| HRH1 | MSLPNSSCLI | MSLPNSSCLI | V42I | -1.2072635 | -1.4880805 | -1.4380703 |
| SH2D4A | MLKQILSEMY | MLKQILSEMY | R334Q | -5.6816034 | -5.6754303 | -4.1078682 |
| NEDD4 | MAQSLRLHF/ | MAQSLRLHF/ | K261Q:K347T | 2.75702691 | 3.0493474 | -0.1560438 |
| MARCO | MRNKKILKED | MRNKKILKED | L74M | -1.4671597 | -1.5462902 | -1.4709229 |
| MAPK1 | MAAAAAAGA( | MAAAAAAGA( | Y358D | -3.1399646 | -4.9010792 | -0.6835518 |
| CFH | MRLLAKIICLN | MRLLAKIICLN | N718D:S720I | 2.72024632 | 2.68368983 | -1.303975 |
| GCNT3 | MVQWKRLC( | MVQWKYLC( | R6Y | -2.9744875 | -3.599262 | -3.3436537 |

| | | | | | | |
|---|---|---|---|---|---|---|
| DHX33 | MPEEAGFPP/ | MPEGAGFPP/ | E4G | 0.2073667 | 0.21163774 | 0.51712203 |
| NFATC1 | MPSTSFPVPS | MPSTSFPVPS | V928A | -6.5961766 | -8.8952789 | -7.4824295 |
| PLEKHA1 | MPYVDRQNF | MPYVDRQNF | P294S | -5.9852047 | -8.3001242 | -7.8417645 |
| PUM2 | MNHDFQALA | MNHDFQALA | N242S | 0.9522059 | 1.49893773 | 0.54001236 |
| TREML1 | MGLTLLLLLL | MGLTPLLLLL | L5P:G15E:F5 | -1.7985723 | -8.2039343 | -4.0252973 |
| TNFSF8 | MDPGLQQAL | MDPGLQQAL | P69H | -1.0422964 | -1.0687315 | -0.6992197 |
| PKN2 | MASNPERGE | MASNPERGE | E613G | -0.1400659 | -1.2070037 | -0.2791779 |
| CD302 | MLRAALPALLI | MLRAALPALLI | M110V | 0.96353245 | 1.690871 | 1.42631173 |
| IRG1 | MLANSSITESI | MLANSSITESI | K150S | -1.9091974 | -3.8855152 | -1.6772251 |
| MID2 | MGESPASVVL | MGESPASVVL | V59L | -6.0102482 | -6.6641865 | -5.211082 |
| DHX9 | MGDVKNFLY | MGDVKNFLY | T108A | 0.29965091 | 0.06507778 | 0.13633251 |
| C5AR1 | MDSFNYTTPI | MDSFNYTTPI | F182N | -0.3291407 | -4.2072248 | 0.19669271 |
| IL1RL2 | MTGLVSLSYF | MTGLVSLSYF | G127S | 1.27292776 | 0.65097761 | -0.0980263 |
| ATF6 | MGEPAGVAG | MGEPAGVAG | R403S:T606K | -1.5114973 | -5.4593556 | -0.5057467 |
| ST6GAL1 | MIHTNLKKKF | MIHTNLKKKF | D146G | -0.6930439 | -5.1033688 | -0.5436673 |
| LY75 | MRTGWATPR | MRTGWATPR | P153S:K488N | -2.0571718 | -2.816535 | -3.1475077 |
| RABGEF1 | MMASSYHEV | MMASSYHEV | V26L | -4.5316148 | -4.5648232 | -4.3922997 |
| CD1D | MGCLLFLLLW | MGCLLFLLLW | G46T:K232Q | -7.5216231 | -9.3783259 | -8.3380973 |
| ADAR | MMSPICDQT | MMSPICDQT | G488D | -0.320399 | 0.06982803 | 0.58718419 |
| ATP7A | MDPSMGVNS | MDPSMGVNS | R512G:D828 | -1.1934569 | -1.8514243 | -0.0802574 |
| PSMD5 | MAAQALALLR | MAAQALALLR | S266A | -0.6282554 | -2.6188314 | -0.1860745 |
| TRIM8 | MAENWKNCI | MAENWKNCI | V225L | -1.0676324 | -0.6893535 | -0.7782207 |
| PIK3AP1 | MAASGVPRG | MAASGVPRG | R706I:L709M | -3.4599154 | -4.3858733 | -2.1563182 |
| APOBEC2 | MAQKEEAAV/ | MAQKEEAAV/ | L24S | -0.8044679 | -0.9947674 | -1.3300381 |
| DAPK1 | MTVFRQENVI | MTVFRQENVI | P1355S | 0.4518199 | 0.27946782 | 0.02463508 |
| STK11 | MEVVDPQQL | MEVVDPQQL | K108R | -3.4386597 | -1.0013142 | -1.4123034 |
| NFKBIZ | MIVDKLLDDS | MIVDKLLDDS | C177H:I353V | -2.8927364 | -2.2117584 | 0.22303247 |
| TRAT1 | MSGISGCPFI | MSGISGCPFI | K92D | -0.3571913 | -0.308497 | -0.1638064 |
| IL17C | MTLLPGLLFLT | MTLLPGLLFLT | L70V | -1.1182263 | -1.2527426 | -1.1196648 |
| CD38 | MANCEFSPVS | MANCEFSPVS | R47M | -1.3054571 | -1.3721099 | -1.5838795 |
| FGA | MFSMRIVCLV | MFSMRIVCLV | P352E | -3.4284222 | -3.2600408 | -3.8107822 |
| IFNL3 | MKLDMTGDC | MKLDMTGDC | Q52W:Q55K: | -4.2647533 | -3.9792677 | -4.2150993 |
| DHX58 | MELRSYQWE | MELRSYQWE | T395S | 1.27015448 | 1.3752265 | 1.24207902 |
| MADCAM1 | MDFGLALLLA | MDFGLALLLA | E263S:R292S | -6.737103 | -7.4499511 | -6.696951 |
| 8-Mar | MSMPLHQIS/ | MSMPLHQIS/ | T538I | 0.41079807 | 0.61921215 | -0.5089903 |
| CD207 | MTVEKEAPDA | STVEKEDPDA | M1S:A7D:K3 | -3.4222426 | -3.2922162 | -4.5911602 |
| MAP4K2 | MALLRDVSLC | MALLRDVSLC | N349S:V723I | 1.25104046 | 1.77143914 | 0.20379519 |
| JAG1 | MRSPRTRGR | MRSPRTRGR | G1100T | 0.71121335 | 0.51668596 | 0.75824738 |
| ATG9A | MAQFDTEYQ | MMQFDTEYC | A2M:P360L:L | -11.502863 | -12.575901 | -8.9471688 |
| VAV3 | MTQSLSYLAT | MTQSLSYLAT | Q634Y | -4.5801992 | -2.9060926 | -5.3301511 |
| TNIP3 | MIPCGWLMN | MIPCGWLMN | G114T | 0.4603467 | 0.15422058 | 0.19252396 |
| TMEM173 | MPHSSLHPSI | MPHSSLHPSI | E249D | -0.5283582 | -0.6971719 | -0.7362833 |
| LAMP1 | MAAPGSARRI | MAAPGSARRI | T39E | -2.7143948 | -5.3593998 | -3.1691761 |
| ITGAL | MKDSCITVM/ | MKDSCITVM/ | M1125L | 0.92243671 | 1.16111708 | 0.69281459 |
| PROS1 | MRVLGGRCC | MRVLSGRCG | G5S:S428N | -3.9549119 | -4.831996 | -2.9746346 |

| | | | | | | |
|---|---|---|---|---|---|---|
| ATG16L1 | MSSGLRAADI | MSSGLRAADI | G583E | -5.2568803 | -8.0265789 | -9.6190968 |
| GCH1 | MEKGPVRAP/ | MEKGPVRAP/ | P28Q | -1.0546873 | -1.2183113 | -0.9840059 |
| CTSG | MQPLLLLLAFI | MQPLLLLLAFI | R133L:N139S | -3.9907835 | -3.9594716 | -2.4795024 |
| COLEC12 | MKDDFAEEEI | MKDDFAEEEI | P493L:E672A | -3.5455391 | -6.5035965 | -6.0416787 |
| CD97 | MGGRVFLAF( | MGGRVFLAF( | F51T:T59M:S | -8.8717647 | -13.79908 | -12.114711 |
| MASP2 | MRLLTLLGLL( | MRLLTLLGLL( | S112A:S185E | -5.4746442 | -10.066957 | -8.9075098 |
| SEMA4A | MALPALGLDF | MALPALGLDF | A419T:G757F | -4.185663 | -4.4115899 | -2.4513083 |
| LPXN | MSTLLISSSD/ | MSTLLISSSD/ | P383T | -1.6798489 | -2.5665674 | -0.741451 |
| THEMIS | MALSLEEFVH | MALSLEEFVH | D630N | -0.2055144 | 0.01229954 | -0.4239469 |
| ANG | MVMGLGVLL | MVMGLGVMI | L8M | -1.9275757 | -2.7163181 | -2.0650978 |
| APEX2 | MLRVVSWNII | MLRVVSWNII | R132C:A133T | -15.675841 | -15.368213 | -11.98094 |
| OASL | MALMQELYST | MALMQELYST | L119V | -0.566462 | -1.8363489 | -0.4237421 |
| EXOSC3 | MAEPASVAAE | MAEPASVAAE | Q39Y | -3.0307245 | -4.0279717 | -1.8831017 |
| CFP | MITEGAQAPR | MITEGAQAPR | K44E | -0.5150993 | -1.5677679 | -0.324297 |
| POU2F2 | MVHSSMGAF | MVHSSMGAF | G528A | 1.09215724 | -1.1240153 | 2.75892305 |
| ART1 | MQMPAMMS | MQMPAMMS | P126L:S300N | -1.6085038 | -1.8302102 | -2.3164668 |
| IL18RAP | MLCLGWIFL\ | MLCLGWIFL\ | H55Y:Q66R:( | 1.10638237 | 0.72782111 | 1.75821614 |
| ATF6B | MAELMLLSEI/ | MAELMLLSEI/ | L381A | -3.4459467 | -2.6387062 | -3.8620467 |
| CD5L | MALLFSLILAI( | MALLFSLILAI( | E84K | -0.343637 | -0.1886218 | -0.6674082 |
| EGF | MLLTLIILLPVV | MLLTLIILLPVV | R1082S | 0.5641439 | 0.01582265 | 0.63554215 |
| ITCH | MSDSGSQLG | MSDSGSQLG | V24I:N159R | -1.1199489 | -1.4717885 | -1.4966762 |
| IL1RL1 | MGFWILAILTI | MGFWILAILTI | I332T:T391K: | -5.2969126 | -8.7138876 | -2.0626607 |
| TAPBP | MKSLSLLLAV/ | MKSLSLLLAV/ | R57K | -1.4197469 | -1.2699976 | -1.5259435 |
| C5 | MLQTTPDLKS | MLQTTPDLKS | D665G | -0.060061 | -0.0924113 | -0.322087 |
| CYP27B1 | MTQTLKYASR | MTQTLKYASR | V397L | -0.9160514 | -1.8637328 | 0.25821972 |
| GZMH-immu | MQPFLLLLAF | MQLFLLFIAFF | P3L:L7F:L8I:L | -8.7543736 | -8.5209978 | -7.9485179 |
| ERBB2IP | MTTKRSLFVR | MTTKRSLFVR | D724Q | -0.6179061 | -0.5058796 | -0.6431627 |
| MSH6 | MSRQSTLYSF | MSRQSTLYSF | S602A:V1056 | -4.2137849 | -3.2748837 | -1.3194584 |
| ABL1 | MGQQPGKVI | MGQQPGKVI | V753M | -1.1798303 | -1.6373038 | -1.7730572 |
| SH2B2 | MDPSYCPAH | MDPSYCPAH | S639N | -3.7685077 | -3.4326389 | -3.178983 |
| BMP6 | MPGLGRRAQ | MPGLGRRAQ | D402G:K482I | -23.381662 | -31.089759 | -23.743524 |
| PLCL2 | MAECGRGG/ | MAECGRGG/ | S86G | 0.38187671 | 0.37435079 | 0.39838767 |
| SPPL2A | MWLLGCPEG | MWLLGCPEG | T78S | 0.31829906 | 0.67202902 | -0.0637558 |
| HAVCR2 | MFSHLPFDC | MFSHLPFDC | P138Q | -1.6263227 | -2.0758657 | -1.9758909 |
| ATF7 | MGDDRPFV( | AGDDRPFVC | M1A | -8.3136444 | -7.5963783 | -6.0038567 |
| MR1 | MGELMAFLLF | MGELLALRLL | M5L:F7L:L8R: | -113.76583 | -105.12057 | -96.023691 |
| APBB1IP | MGESSEDID( | MGESSEDID( | T349I | -5.3414698 | -5.4044604 | -4.7243958 |
| PRKCG | MAGLGPGVG | MAGLGPGVG | T438N:P602L | 2.64529514 | 3.77467489 | 3.81621718 |
| AQP9 | MQPEGAEKG | MQPEGAEKG | V62A | -2.6605339 | -3.4079123 | -2.5111699 |
| TINAGL1 | MWRCPLGLL | MWRCPLGLL | H467W | 0.05398655 | -1.784519 | -2.4420731 |
| CD200 | MERLTLTRTIG | MERLTLTRTIG | S157R | -4.1126575 | -1.8355906 | -2.2307675 |
| LY9 | MTGLLGLSPV | MTGLLGLSPV | T120Q:I122F: | 0.42388666 | -4.6641626 | -0.0369157 |
| SEMA4D | MRMCTPIRG | MRMCTPIRG | G602H | 1.51670027 | 3.69840884 | 2.65191007 |
| ITGB7 | MVALPMVLVL | MVALPMVLVL | Q102R:R129/ | -3.6432273 | -3.7131958 | -2.2108617 |
| IL31RA | MCIRQLKFFT | MCIRQLKFFT | N536Y | -4.0431886 | -8.6571054 | -5.8645606 |

| | | | | | | |
|---|---|---|---|---|---|---|
| CD22 | MHLLGPWLL | MHLLGPWLL | S123I:H133G | -9.4032631 | -11.568225 | -8.5385233 |

**C** Protein language modesl evaluation with Jonathan's amino a

| Ref | wt_sequence | mutated_seq | mutant | Tranception_ | Tranception_ | Tranception_ |
|---|---|---|---|---|---|---|
| KIAA1549 | MPGARRRRF | MPGARRRRF | D155N | -2.6821413 | -2.6823878 | -0.0002465 |
| KIAA1549 | MPGARRRRF | MPGARRRRF | P196S | -2.6821413 | -2.682081 | 6.03E-05 |
| ARID1B | MAHNAGAAA | MAHNAGAAA | T826A | -2.0061045 | -2.0061162 | -1.18E-05 |
| GOPC | MSAGGPCPA | MSAGGPCPA | Q109K | -0.6684296 | -0.6880972 | -0.0196676 |
| ROS1 | MKNIYCLIPKI | MKNIYCLIPKI | Y1312C | -2.5327781 | -2.5329612 | -0.0001831 |
| ROS1 | MKNIYCLIPKI | MKNIYCLIPKI | Q1368N | -2.5327781 | -2.5319738 | 0.00080427 |
| ROS1 | MKNIYCLIPKI | MKNIYCLIPKI | Y1488A | -2.5327781 | -2.5328671 | -8.90E-05 |
| ROS1 | MKNIYCLIPKI | MKNIYCLIPKI | L1612T | -2.5327781 | -2.5326834 | 9.47E-05 |
| ROS1 | MKNIYCLIPKI | MKNIYCLIPKI | K2100R | -2.5327781 | -2.5335836 | -0.0008055 |
| RB1 | MPPKTPRKTA | MPPKTPRKTA | H69V | -2.5219359 | -2.5198209 | 0.00211501 |
| RB1 | MPPKTPRKTA | MPPKTPRKTA | P253A | -2.5219359 | -2.5232985 | -0.0013626 |
| CBLB | MANSMNGRI | MANSMNGRI | L590P | -1.0984935 | -1.1000594 | -0.0015659 |
| CBLB | MANSMNGRI | MANSMNGRI | G748S | -1.0984935 | -1.0979596 | 0.00053382 |
| BCL9 | MHSSNPKVR | MHSSNPKVR | M1074V | -2.3558865 | -2.3558006 | 8.58E-05 |
| FLT3 | MPALARDGG | MPALARDGG | I21A | -2.1283772 | -2.1282778 | 9.94E-05 |
| FLT3 | MPALARDGG | MPALARDGG | V56I | -2.1283772 | -2.1287766 | -0.0003994 |
| FLT3 | MPALARDGG | MPALARDGG | D779N | -2.1283772 | -2.1308784 | -0.0025012 |
| APC2 | MGLLGLLSLL | MGLLGLLSLL | V196M | -2.4974861 | -2.4977662 | -0.0002801 |
| APC2 | MGLLGLLSLL | MGLLGLLSLL | A1545Q | -2.4974861 | -2.4981686 | -0.0006825 |
| CASC5 | MDGVSSEAN | MDGVSSEAN | H415D | -2.829471 | -2.8297863 | -0.0003153 |
| CASC5 | MDGVSSEAN | MDGVSSEAN | N2321C | -2.829471 | -2.8296118 | -0.0001408 |
| NRG1 | MEIYSPDMSI | MEIYSPDMSI | P49H | -0.7084033 | -0.7174104 | -0.0090071 |
| CEP89 | MLLGFRRGR | MLLGFRRGR | Q688H | -2.5071228 | -2.5108099 | -0.0036871 |
| NDRG1 | MSREMQDVI | MSREMQDVI | E38D | -0.8100705 | -0.8212062 | -0.0111357 |
| ALK | MGAIGLLWLI | MIAIGLLWLLI | G2I | -2.1010849 | -2.1033578 | -0.0022728 |
| ALK | MGAIGLLWLI | MGSIGLLWLI | A3S | -2.1010849 | -2.1006598 | 0.0004251 |
| ALK | MGAIGLLWLI | MGAHGLLWLI | I4H | -2.1010849 | -2.1001275 | 0.00095749 |
| ALK | MGAIGLLWLI | MGAIGLLWLI | L80T | -2.1010849 | -2.1013856 | -0.0003006 |
| ALK | MGAIGLLWLI | MGAIGLLWLI | K81R | -2.1010849 | -2.1000736 | 0.00101137 |
| ALK | MGAIGLLWLI | MGAIGLLWLI | L422M | -2.1010849 | -2.1020017 | -0.0009167 |
| NTRK3 | MDVSLCPAK | MDVSLCPAK | F720Y | -0.3185824 | -0.3194603 | -0.0008779 |
| CIC | MKPMKKACT | MKPMKKACT | S1258A | -2.4847016 | -2.4845525 | 0.00014909 |
| CIC | MKPMKKACT | MKPMKKACT | L2144P | -2.4847016 | -2.4836 | 0.00110165 |
| DAXX | MRGSENCGI | MRGSENCGI | S188L | -2.654047 | -2.6548281 | -0.0007811 |
| LATS2 | MRPKTFPATT | MRPKTFPATT | A325S | -2.3326099 | -2.3327382 | -0.0001283 |
| NOTCH1 | MPPLLAPLLC | MPPLLAPLLC | N2396T | -1.8759292 | -1.8759384 | -9.22E-06 |
| BRD4 | MSAESGPGTI | MSAESGPGTI | V997I | -1.8885357 | -1.8888257 | -0.0002899 |
| RNF213 | MECPSCQHV | MECPSCQHV | V1824M | -2.8245703 | -2.824695 | -0.0001246 |
| RNF213 | MECPSCQHV | MECPSCQHV | V3576N | -2.8245703 | -2.8247732 | -0.0002028 |
| THRAP3 | MSKTNKSKS( | MSKTNKSKS( | H693L | -1.800611 | -1.80826 | -0.0076489 |
| SETD2 | MILLMRNLTT | MILLMRNLTT | A1408I | -2.5838192 | -2.5839264 | -0.0001071 |
| SETD2 | MILLMRNLTT | MILLMRNLTT | D1444E | -2.5838192 | -2.5837083 | 0.00011094 |

| | | | | | | |
|---|---|---|---|---|---|---|
| CLIP4 | MTIEDLPDFP | MTIEDLPDFP | P511A | -1.2078091 | -1.2190548 | -0.0112457 |
| PTPRB | MEAEFYMVIL | MEAEFYMVIL | M489V | -2.1910526 | -2.1905605 | 0.0004921 |
| PTPRB | MEAEFYMVIL | MEAEFYMVIL | V613H | -2.1910526 | -2.1913762 | -0.0003236 |
| PTPRB | MEAEFYMVIL | MEAEFYMVIL | E646A | -2.1910526 | -2.190913 | 0.00013955 |
| PTPRB | MEAEFYMVIL | MEAEFYMVIL | E663T | -2.1910526 | -2.190966 | 8.66E-05 |
| PTPRB | MEAEFYMVIL | MEAEFYMVIL | M750N | -2.1910526 | -2.1909135 | 0.00013908 |
| PTPRB | MEAEFYMVIL | MEAEFYMVIL | A885P | -2.1910526 | -2.1910969 | -4.43E-05 |
| PTPRB | MEAEFYMVIL | MEAEFYMVIL | V889R | -2.1910526 | -2.1928341 | -0.0017815 |
| PTPRB | MEAEFYMVIL | MEAEFYMVIL | H1182V | -2.1910526 | -2.1917079 | -0.0006553 |
| PTPRB | MEAEFYMVIL | MEAEFYMVIL | L1201T | -2.1910526 | -2.1925133 | -0.0014607 |
| PTPRB | MEAEFYMVIL | MEAEFYMVIL | L1442A | -2.1910526 | -2.1917799 | -0.0007273 |
| PTPRB | MEAEFYMVIL | MEAEFYMVIL | R1454Q | -2.1910526 | -2.191346 | -0.0002934 |
| PTPRB | MEAEFYMVIL | MEAEFYMVIL | Y1538W | -2.1910526 | -2.1938697 | -0.0028172 |
| PTPRB | MEAEFYMVIL | MEAEFYMVIL | Q2100G | -2.1910526 | -2.1920234 | -0.0009708 |
| KMT2D | MDSQKLAGE | MDSQKLAGE | P1165L | -2.384999 | -2.3853022 | -0.0003033 |
| NSD1 | MDQTCELPR | MDQTCELPR | A480T | -2.3253816 | -2.3253797 | 1.91E-06 |
| NSD1 | MDQTCELPR | MDQTCELPR | L2513F | -2.3253816 | -2.3259096 | -0.000528 |
| IDH1 | MSKKISGGS\ | MSKKISGGS\ | G204S | -0.395776 | -0.4038902 | -0.0081142 |
| BCL11B | MSRRKQGNI | MSRRKQGNI | L127R | -0.8023987 | -0.8194511 | -0.0170524 |
| KMT2A | MAHSCRWRI | MAHSCRWRI | S570R | -2.5667629 | -2.5668302 | -6.72E-05 |
| ATRX | MTAEPMSESI | MTAEPMSESI | R1131Q | -2.2919094 | -2.2925844 | -0.000675 |
| NCOA4 | MGAQLTTFAL | MGAQLTTFAL | S511R | -2.7558541 | -2.7590039 | -0.0031497 |
| ERBB2 | MELAALCRW | MELAALCRW | T166M | -2.0284977 | -2.0309455 | -0.0024478 |
| BRCA1 | MDLSALRVEE | MDLSALRVEE | M1083R | -0.2244553 | -0.2331983 | -0.008743 |
| CNOT3 | MADKRKLQG | MADKRKLQG | A442I | -1.1813762 | -1.1832558 | -0.0018796 |
| STRN | MDEQAGPG\ | MDEQAGPG\ | D226H | -0.7631508 | -0.7791584 | -0.0160076 |
| TET1 | MSRSRHARP | MSRSRHARP | L1329H | -2.7814051 | -2.7816683 | -0.0002632 |
| TRIM33 | MAENKGGGI | MAENKGGGI | G877E | -1.9123719 | -1.9135797 | -0.0012078 |
| SETBP1 | MESRETLSSS | MESRETLSSS | L556I | -2.6932898 | -2.6937375 | -0.0004478 |
| TRIM24 | MEVAVEKAVA | MEVAVEKAVA | Y540S | -2.1100459 | -2.1126962 | -0.0026503 |
| CDH11 | MKENYCLQA | MKENYCLQA | M382V | -0.5898793 | -0.5854887 | 0.00439066 |
| NAB2 | MHRAPSPTAI | MHRAPSPTAI | G494R | -1.7902986 | -1.7904934 | -0.0001948 |
| RALGDS | MVQRMWAE, | MVQRMWAE, | D41G | -0.6803576 | -0.6801455 | 0.00021207 |
| ACSL6 | MLTFFLVSGG | MLTFFLVSGG | V519L | -0.404587 | -0.4174858 | -0.0128988 |
| PPFIBP1 | MMSDASDMI | MMSDASDMI | Q249H | -1.8046286 | -1.8097155 | -0.0050869 |
| PPFIBP1 | MMSDASDMI | MMSDASDMI | V288M | -1.8046286 | -1.8079104 | -0.0032818 |
| TP53BP1 | MPGEQMDP | MPGEQMDP | M721V | -2.7098217 | -2.7093768 | 0.00044489 |
| MET | MKSKSKSLAE | MKSKSKSLAE | S122Q | -2.0976923 | -2.0980802 | -0.0003879 |
| LATS1 | MKRSEKPEG | MKRSEKPEG | S1023A | -2.2926853 | -2.2956789 | -0.0029936 |
| NUTM1 | MVVTLGPGPI | MVVTLGPGPI | S165T | -2.7141697 | -2.7144375 | -0.0002677 |
| MAP3K13 | MANFQEHLS | MANFQEHLS | I745L | -1.8678291 | -1.8667567 | 0.00107241 |
| AFF3 | MSKKLITCLSI | MSKKLITCLSI | R978K | -2.0903466 | -2.0899489 | 0.00039768 |
| KCNT2 | MVDLESEVPI | MVDLESEVPI | I974M | -1.5852001 | -1.5846328 | 0.00056732 |
| FOXO3 | MAEAPASPAP | MAEAPASPAP | A98G | -1.5849191 | -1.5839924 | 0.00092673 |
| SSPN | MGKNKQPR( | MGKNKQPR( | E28K | -1.0515405 | -1.0524323 | -0.0008918 |

| | | | | | | |
|---|---|---|---|---|---|---|
| POT1 | MSLVPATNYI | MSLVPATNYI | K440E | -2.8792279 | -2.8790174 | 0.00021052 |
| PTCD2 | MVRDSMAAA | MVRDSMAAA | A316C | -2.4047716 | -2.4116988 | -0.0069273 |
| GRN | MWTLVSWVA | MWTLVSWVA | P430L | -1.901294 | -1.9082716 | -0.0069776 |
| POLQ | MNLLRRSGK | MNLLRRSGK | S244I | -2.6026208 | -2.6031257 | -0.000505 |
| POLQ | MNLLRRSGK | MNLLRRSGK | D2414K | -2.6026208 | -2.6038124 | -0.0011916 |
| LRP1B | MSEFLLALLTL | MSEFLLALLTL | T2167M | -2.0059595 | -2.0066921 | -0.0007326 |
| LRP1B | MSEFLLALLTL | MSEFLLALLTL | P2946S | -2.0059595 | -2.0061846 | -0.0002251 |
| LRP1B | MSEFLLALLTL | MSEFLLALLTL | E3112Q | -2.0059595 | -2.0063608 | -0.0004013 |
| DKC1 | MADAEVIILPK | MADAEVIILPK | E210G | -0.7643179 | -0.7776198 | -0.013302 |
| TERF2 | MAAGAGTAG | MAAGAGTAG | R322F | -2.2971981 | -2.2999175 | -0.0027194 |
| ACD | MPGRCQSD/ | MPGRCQSD/ | S429P | -2.7087479 | -2.7089474 | -0.0001996 |
| CYC1 | MAAAAASLRG | MAAAAASLRG | P38S | -1.1457454 | -1.1461378 | -0.0003924 |
| LMX1B | MDIATGPESL | MDIATGPESL | M196I | -0.6630339 | -0.6634661 | -0.0004322 |
| AIF1 | MSQTRDLQG | MSQTRDLQG | S168G | -2.4025819 | -2.4905078 | -0.0879259 |
| CMAS | MDSVEKGAA | MDSVEKGAA | P43A | -1.2593038 | -1.2600718 | -0.0007679 |
| AGPAT2 | MELWPCLAA | MELWPCLAA | P232D | -1.3030779 | -1.3278699 | -0.024792 |
| ATM | MSLVLNDLLI | MSLVLNDLLI | E1245Q | -2.8268925 | -2.8270849 | -0.0001923 |
| ATM | MSLVLNDLLI | MSLVLNDLLI | K1451V | -2.8268925 | -2.827548 | -0.0006555 |
| ATM | MSLVLNDLLI | MSLVLNDLLI | D1452K | -2.8268925 | -2.8268846 | 7.95E-06 |
| ATM | MSLVLNDLLI | MSLVLNDLLI | V1532S | -2.8268925 | -2.8267908 | 0.00010173 |
| ATM | MSLVLNDLLI | MSLVLNDLLI | K2456R | -2.8268925 | -2.8269482 | -5.56E-05 |
| KLRF1 | MQDEERYMT | MQDEERYMT | P202L | -2.2656732 | -2.2722168 | -0.0065436 |
| SYNE1 | MATSRGASR | MATSRGASR | A2020G | -2.5702849 | -2.570471 | -0.0001861 |
| SYNE1 | MATSRGASR | MATSRGASR | P2769I | -2.5702849 | -2.5702523 | 3.26E-05 |
| SYNE1 | MATSRGASR | MATSRGASR | S3321G | -2.5702849 | -2.5704613 | -0.0001763 |
| SYNE1 | MATSRGASR | MATSRGASR | S3915K | -2.5702849 | -2.5703451 | -6.02E-05 |
| SYNE1 | MATSRGASR | MATSRGASR | E4957K | -2.5702849 | -2.5705278 | -0.0002429 |
| SYNE1 | MATSRGASR | MATSRGASR | A5702V | -2.5702849 | -2.5704121 | -0.0001272 |
| SYNE1 | MATSRGASR | MATSRGASR | L5842F | -2.5702849 | -2.570471 | -0.0001861 |
| SYNE1 | MATSRGASR | MATSRGASR | A6283H | -2.5702849 | -2.5704596 | -0.0001746 |
| MRC2 | MGPGRPAPA | MGPGRPAPA | K241R | -2.1623139 | -2.1628852 | -0.0005713 |
| MALSU1 | MGPGGRVAF | MGPGGRVAF | V28S | -1.555312 | -1.553465 | 0.00184703 |
| IFNGR2 | MIQQLLMRW | MIQQLLMRW | L242K | -2.6664093 | -2.665786 | 0.00062323 |
| IFNGR2 | MIQQLLMRW | MIQQLLMRW | S246G | -2.6664093 | -2.669771 | -0.0033617 |
| IFNGR2 | MIQQLLMRW | MIQQLLMRW | L314Q | -2.6664093 | -2.660542 | 0.00586724 |
| IFNGR2 | MIQQLLMRW | MIQQLLMRW | D343G | -2.6664093 | -2.6672668 | -0.0008576 |
| MINPP1 | MLRAPGCLLF | MLRAPGCLLF | N424H | -2.4759398 | -2.4762437 | -0.000304 |
| APOA5 | MASMAAVLTV | MASMAAVLTV | A160V | -2.5726719 | -2.5825138 | -0.0098419 |
| HOXD12 | MCERSLYRA | MCERSLYRA | A215C | -1.731895 | -1.7688733 | -0.0369784 |
| PRUNE2 | MEEFLQRAKS | MEEFLQRAKS | E78Q | -2.7318687 | -2.7319155 | -4.67E-05 |
| PRUNE2 | MEEFLQRAKS | MEEFLQRAKS | P203L | -2.7318687 | -2.7314906 | 0.00037813 |
| PRUNE2 | MEEFLQRAKS | MEEFLQRAKS | P1564A | -2.7318687 | -2.7319241 | -5.53E-05 |
| SIRT1 | MADEAALALQ | MADEAALALQ | G19V | -1.4991884 | -1.5014813 | -0.0022929 |
| IGF2R | MGAAAGRSP | MGAAAGRSP | K1862S | -2.7364883 | -2.7366975 | -0.0002092 |
| PROM1 | MALVLGSLLLI | MALVLGSLLLI | I169M | -1.6574389 | -1.6624537 | -0.0050148 |

| | | | | | | |
|---|---|---|---|---|---|---|
| PROM1 | MALVLGSLLLI | MALVLGSLLLI | V371L | -1.6574389 | -1.6617658 | -0.0043269 |
| PROM1 | MALVLGSLLLI | MALVLGSLLLI | V485S | -1.6574389 | -1.6666808 | -0.0092419 |
| ATG4C | MEATGTDEVI | MEATGTDEVI | V245I | -0.7928526 | -0.8019601 | -0.0091074 |
| TLR4 | MMSASRLAG | MMSASRLAG | G420C | -2.2240021 | -2.2265272 | -0.0025251 |
| ADAMTSL1 | MGTMARLEA | MGTMARLEA | A1003V | -1.810487 | -1.8110745 | -0.0005875 |
| FGF19 | MRSGCVVVH | MRSGCVVVH | R63S | -2.3034539 | -2.317271 | -0.0138171 |
| CGNL1 | MELYFGEYQI | MELYFGEYQI | Q27D | -2.2714884 | -2.2714162 | 7.22E-05 |
| CGNL1 | MELYFGEYQI | MELYFGEYQI | C234D | -2.2714884 | -2.270874 | 0.0006144 |
| ITGB3 | MRARPRPRP | MRARPRPRP | A50T | -0.7087655 | -0.7149769 | -0.0062114 |
| ITGB3 | MRARPRPRP | MRARPRPRP | T221A | -0.7087655 | -0.7125701 | -0.0038046 |
| ITGB3 | MRARPRPRP | MRARPRPRP | M594V | -0.7087655 | -0.7136646 | -0.0048991 |
| NCOR2 | MSGSTQPVA( | MSGSTQPVA( | T946S | -1.8733365 | -1.8726759 | 0.00066058 |
| NCOR2 | MSGSTQPVA( | MSGSTQPVA( | H1853L | -1.8733365 | -1.8753668 | -0.0020304 |
| NCOR2 | MSGSTQPVA( | MSGSTQPVA( | A2237I | -1.8733365 | -1.8740244 | -0.0006879 |
| MSRA | MLSATRRAC( | MLSATRRAC( | A96S | -0.7292565 | -0.7389247 | -0.0096682 |
| MSRA | MLSATRRAC( | MLSATRRAC( | E177G | -0.7292565 | -0.7747379 | -0.0454814 |
| TRIM25 | MAELCPLAEE | MAELCPLAEE | K332T | -2.090394 | -2.0910819 | -0.0006878 |
| F7 | MVSQALRLLC | MVSQALRLLC | V24I | -1.6192336 | -1.6228073 | -0.0035737 |
| F7 | MVSQALRLLC | MVSQALRLLC | K70Q | -1.6192336 | -1.6172248 | 0.0020088 |
| F7 | MVSQALRLLC | MVSQALRLLC | A220T | -1.6192336 | -1.619488 | -0.0002544 |
| F7 | MVSQALRLLC | MVSQALRLLC | P310S | -1.6192336 | -1.6203415 | -0.0011079 |
| TSFM | MSLLRSLRVF | MSLLRSLRVF | L143A | -1.9563199 | -1.983677 | -0.0273571 |
| TSFM | MSLLRSLRVF | MSLLRSLRVF | I219T | -1.9563199 | -1.943333 | 0.0129869 |
| SORCS1 | MGKVGAGG( | MGKVGAGG( | G807R | -1.6300399 | -1.6392945 | -0.0092546 |
| AOPEP | MDIQLDPARI | MDIQLDPARI | E61K | -1.7362792 | -1.7363468 | -6.76E-05 |
| NFKB1 | MKRTTSFRM/ | MKRTTSFRM/ | T212A | -1.5117934 | -1.5204347 | -0.0086414 |
| ZBTB20 | MLERKKPKTA | MLERKKPKTA | N437D | -1.612076 | -1.6126258 | -0.0005498 |
| FBXO31 | MAVCARLCG | MAVCARLCG | G396D | -0.9363843 | -0.9374743 | -0.00109 |
| FBXO31 | MAVCARLCG | MAVCARLCG | V450M | -0.9363843 | -0.9406334 | -0.0042491 |
| BSCL2 | MSTEKVDQKI | MSTEKVDQKI | R350Q | -1.5975564 | -1.5993738 | -0.0018175 |
| PPP1R1A | MEQDNSPRk | MEQDNSPRk | E49K | -1.7316196 | -1.8388416 | -0.107222 |
| VWA5A | MSSVCDMSF | MSSVCDMSF | K98R | -2.2294941 | -2.2320147 | -0.0025206 |
| METTL1 | MAAETRNVA( | MAAETRNVA( | Q117H | -0.9112919 | -0.9390146 | -0.0277227 |
| BNIPL | MGTIQEAGKI | MGTIQEAGKI | L35F | -1.2715396 | -1.2873253 | -0.0157857 |
| BNIPL | MGTIQEAGKI | MGTIQEAGKI | E43D | -1.2715396 | -1.2743458 | -0.0028062 |
| BNIPL | MGTIQEAGKI | MGTIQEAGKI | E51D | -1.2715396 | -1.2729458 | -0.0014062 |
| BNIPL | MGTIQEAGKI | MGTIQEAGKI | A69N | -1.2715396 | -1.2780348 | -0.0064952 |
| BNIPL | MGTIQEAGKI | MGTIQEAGKI | T71E | -1.2715396 | -1.2813275 | -0.0097879 |
| BNIPL | MGTIQEAGKI | MGTIQEAGKI | P72A | -1.2715396 | -1.2847301 | -0.0131905 |
| BNIPL | MGTIQEAGKI | MGTIQEAGKI | S73D | -1.2715396 | -1.2723522 | -0.0008126 |
| BNIPL | MGTIQEAGKI | MGTIQEAGKI | T74I | -1.2715396 | -1.2902558 | -0.0187162 |
| BNIPL | MGTIQEAGKI | MGTIQEAGKI | L77V | -1.2715396 | -1.2805175 | -0.0089779 |
| BNIPL | MGTIQEAGKI | MGTIQEAGKI | C78T | -1.2715396 | -1.2686872 | 0.00285232 |
| BNIPL | MGTIQEAGKI | MGTIQEAGKI | R81Q | -1.2715396 | -1.2749974 | -0.0034578 |
| BNIPL | MGTIQEAGKI | MGTIQEAGKI | P82K | -1.2715396 | -1.2952795 | -0.0237399 |

| | | | | | | |
|---|---|---|---|---|---|---|
| BNIPL | MGTIQEAGK | MGTIQEAGK | M83V | -1.2715396 | -1.2710255 | 0.00051403 |
| BNIPL | MGTIQEAGK | MGTIQEAGK | R86K | -1.2715396 | -1.2785364 | -0.0069969 |
| BNIPL | MGTIQEAGK | MGTIQEAGK | S88M | -1.2715396 | -1.3038703 | -0.0323308 |
| BNIPL | MGTIQEAGK | MGTIQEAGK | L92I | -1.2715396 | -1.282499 | -0.0109594 |
| BNIPL | MGTIQEAGK | MGTIQEAGK | S95T | -1.2715396 | -1.2829859 | -0.0114464 |
| BNIPL | MGTIQEAGK | MGTIQEAGK | T97N | -1.2715396 | -1.2770517 | -0.0055121 |
| BNIPL | MGTIQEAGK | MGTIQEAGK | S119G | -1.2715396 | -1.2862694 | -0.0147299 |
| BNIPL | MGTIQEAGK | MGTIQEAGK | D120E | -1.2715396 | -1.2808133 | -0.0092738 |
| BNIPL | MGTIQEAGK | MGTIQEAGK | L121V | -1.2715396 | -1.2940385 | -0.022499 |
| BNIPL | MGTIQEAGK | MGTIQEAGK | E122D | -1.2715396 | -1.2818282 | -0.0102886 |
| BNIPL | MGTIQEAGK | MGTIQEAGK | I123L | -1.2715396 | -1.2656411 | 0.00589848 |
| BNIPL | MGTIQEAGK | MGTIQEAGK | E125D | -1.2715396 | -1.2847573 | -0.0132177 |
| BNIPL | MGTIQEAGK | MGTIQEAGK | E127D | -1.2715396 | -1.2822901 | -0.0107505 |
| BNIPL | MGTIQEAGK | MGTIQEAGK | S132N | -1.2715396 | -1.2897052 | -0.0181656 |
| BNIPL | MGTIQEAGK | MGTIQEAGK | G138S | -1.2715396 | -1.2947716 | -0.023232 |
| BNIPL | MGTIQEAGK | MGTIQEAGK | H139N | -1.2715396 | -1.286145 | -0.0146054 |
| BNIPL | MGTIQEAGK | MGTIQEAGK | E146D | -1.2715396 | -1.2692858 | 0.00225377 |
| BNIPL | MGTIQEAGK | MGTIQEAGK | R149K | -1.2715396 | -1.2771254 | -0.0055858 |
| BNIPL | MGTIQEAGK | MGTIQEAGK | A150P | -1.2715396 | -1.2954686 | -0.023929 |
| BNIPL | MGTIQEAGK | MGTIQEAGK | E151K | -1.2715396 | -1.2993236 | -0.027784 |
| BNIPL | MGTIQEAGK | MGTIQEAGK | G152T | -1.2715396 | -1.2936563 | -0.0221168 |
| BNIPL | MGTIQEAGK | MGTIQEAGK | L153T | -1.2715396 | -1.2926942 | -0.0211546 |
| BNIPL | MGTIQEAGK | MGTIQEAGK | G154E | -1.2715396 | -1.2921349 | -0.0205953 |
| BNIPL | MGTIQEAGK | MGTIQEAGK | T155G | -1.2715396 | -1.2720996 | -0.00056 |
| BNIPL | MGTIQEAGK | MGTIQEAGK | T158P | -1.2715396 | -1.2928219 | -0.0212823 |
| BNIPL | MGTIQEAGK | MGTIQEAGK | E160Y | -1.2715396 | -1.3301004 | -0.0585608 |
| BNIPL | MGTIQEAGK | MGTIQEAGK | R161T | -1.2715396 | -1.2946243 | -0.0230848 |
| BNIPL | MGTIQEAGK | MGTIQEAGK | L162A | -1.2715396 | -1.291031 | -0.0194914 |
| BNIPL | MGTIQEAGK | MGTIQEAGK | G163A | -1.2715396 | -1.3016313 | -0.0300918 |
| BNIPL | MGTIQEAGK | MGTIQEAGK | G172R | -1.2715396 | -1.3006847 | -0.0291451 |
| BNIPL | MGTIQEAGK | MGTIQEAGK | E173D | -1.2715396 | -1.2772291 | -0.0056895 |
| BNIPL | MGTIQEAGK | MGTIQEAGK | H177R | -1.2715396 | -1.258877 | 0.01266253 |
| BNIPL | MGTIQEAGK | MGTIQEAGK | V180M | -1.2715396 | -1.2766547 | -0.0051152 |
| BNIPL | MGTIQEAGK | MGTIQEAGK | M183I | -1.2715396 | -1.2755971 | -0.0040575 |
| BNIPL | MGTIQEAGK | MGTIQEAGK | R186Q | -1.2715396 | -1.2695918 | 0.00194776 |
| BNIPL | MGTIQEAGK | MGTIQEAGK | E187D | -1.2715396 | -1.2796321 | -0.0080925 |
| BNIPL | MGTIQEAGK | MGTIQEAGK | Q188H | -1.2715396 | -1.2824587 | -0.0109191 |
| BNIPL | MGTIQEAGK | MGTIQEAGK | T193K | -1.2715396 | -1.2847105 | -0.013171 |
| BNIPL | MGTIQEAGK | MGTIQEAGK | V194A | -1.2715396 | -1.2808684 | -0.0093288 |
| BNIPL | MGTIQEAGK | MGTIQEAGK | I216V | -1.2715396 | -1.2777605 | -0.0062209 |
| BNIPL | MGTIQEAGK | MGTIQEAGK | L217V | -1.2715396 | -1.2796769 | -0.0081373 |
| BNIPL | MGTIQEAGK | MGTIQEAGK | S220V | -1.2715396 | -1.3291047 | -0.0575651 |
| BNIPL | MGTIQEAGK | MGTIQEAGK | Y222F | -1.2715396 | -1.2826585 | -0.0111189 |
| BNIPL | MGTIQEAGK | MGTIQEAGK | L223M | -1.2715396 | -1.289445 | -0.0179055 |
| BNIPL | MGTIQEAGK | MGTIQEAGK | R225E | -1.2715396 | -1.2801964 | -0.0086569 |

| | | | | | | |
|---|---|---|---|---|---|---|
| BNIPL | MGTIQEAGKI | MGTIQEAGKI | S227G | -1.2715396 | -1.3010913 | -0.0295517 |
| BNIPL | MGTIQEAGKI | MGTIQEAGKI | I228Q | -1.2715396 | -1.2898401 | -0.0183005 |
| BNIPL | MGTIQEAGKI | MGTIQEAGKI | T232R | -1.2715396 | -1.2844806 | -0.012941 |
| BNIPL | MGTIQEAGKI | MGTIQEAGKI | V234L | -1.2715396 | -1.2841535 | -0.0126139 |
| BNIPL | MGTIQEAGKI | MGTIQEAGKI | E236D | -1.2715396 | -1.2853407 | -0.0138011 |
| BNIPL | MGTIQEAGKI | MGTIQEAGKI | H237N | -1.2715396 | -1.2848504 | -0.0133108 |
| BNIPL | MGTIQEAGKI | MGTIQEAGKI | R240K | -1.2715396 | -1.286651 | -0.0151114 |
| BNIPL | MGTIQEAGKI | MGTIQEAGKI | M242V | -1.2715396 | -1.2820622 | -0.0105226 |
| BNIPL | MGTIQEAGKI | MGTIQEAGKI | L255M | -1.2715396 | -1.2810144 | -0.0094749 |
| BNIPL | MGTIQEAGKI | MGTIQEAGKI | L256I | -1.2715396 | -1.2882284 | -0.0166888 |
| BNIPL | MGTIQEAGKI | MGTIQEAGKI | H258Y | -1.2715396 | -1.3030365 | -0.0314969 |
| BNIPL | MGTIQEAGKI | MGTIQEAGKI | S264T | -1.2715396 | -1.2767594 | -0.0052198 |
| BNIPL | MGTIQEAGKI | MGTIQEAGKI | A266R | -1.2715396 | -1.3010026 | -0.0294631 |
| BNIPL | MGTIQEAGKI | MGTIQEAGKI | Q267K | -1.2715396 | -1.2820622 | -0.0105226 |
| BNIPL | MGTIQEAGKI | MGTIQEAGKI | V268M | -1.2715396 | -1.2911179 | -0.0195783 |
| BNIPL | MGTIQEAGKI | MGTIQEAGKI | I274L | -1.2715396 | -1.2713026 | 0.00023699 |
| BNIPL | MGTIQEAGKI | MGTIQEAGKI | Q276K | -1.2715396 | -1.290259 | -0.0187194 |
| BNIPL | MGTIQEAGKI | MGTIQEAGKI | R279Q | -1.2715396 | -1.2791761 | -0.0076365 |
| BNIPL | MGTIQEAGKI | MGTIQEAGKI | T280Q | -1.2715396 | -1.2793903 | -0.0078508 |
| BNIPL | MGTIQEAGKI | MGTIQEAGKI | A291S | -1.2715396 | -1.2848514 | -0.0133119 |
| BNIPL | MGTIQEAGKI | MGTIQEAGKI | T298S | -1.2715396 | -1.2831726 | -0.011633 |
| BNIPL | MGTIQEAGKI | MGTIQEAGKI | Y300F | -1.2715396 | -1.2959518 | -0.0244123 |
| BNIPL | MGTIQEAGKI | MGTIQEAGKI | K302R | -1.2715396 | -1.2840183 | -0.0124787 |
| BNIPL | MGTIQEAGKI | MGTIQEAGKI | A303T | -1.2715396 | -1.283094 | -0.0115545 |
| BNIPL | MGTIQEAGKI | MGTIQEAGKI | L307I | -1.2715396 | -1.2850885 | -0.013549 |
| BNIPL | MGTIQEAGKI | MGTIQEAGKI | L308T | -1.2715396 | -1.3177693 | -0.0462297 |
| BNIPL | MGTIQEAGKI | MGTIQEAGKI | R309K | -1.2715396 | -1.2804554 | -0.0089158 |
| BNIPL | MGTIQEAGKI | MGTIQEAGKI | T317S | -1.2715396 | -1.2746942 | -0.0031546 |
| BNIPL | MGTIQEAGKI | MGTIQEAGKI | R318Q | -1.2715396 | -1.2932447 | -0.0217052 |
| BNIPL | MGTIQEAGKI | MGTIQEAGKI | F322Y | -1.2715396 | -1.2853823 | -0.0138427 |
| BNIPL | MGTIQEAGKI | MGTIQEAGKI | D324F | -1.2715396 | -1.3281509 | -0.0566113 |
| BNIPL | MGTIQEAGKI | MGTIQEAGKI | S325T | -1.2715396 | -1.2773876 | -0.0058481 |
| BNIPL | MGTIQEAGKI | MGTIQEAGKI | G327A | -1.2715396 | -1.2799079 | -0.0083684 |
| BNIPL | MGTIQEAGKI | MGTIQEAGKI | Q331E | -1.2715396 | -1.2797046 | -0.008165 |
| BNIPL | MGTIQEAGKI | MGTIQEAGKI | L335M | -1.2715396 | -1.2776091 | -0.0060695 |
| BNIPL | MGTIQEAGKI | MGTIQEAGKI | Q337Y | -1.2715396 | -1.2971742 | -0.0256346 |
| BNIPL | MGTIQEAGKI | MGTIQEAGKI | H339G | -1.2715396 | -1.2965473 | -0.0250077 |
| BNIPL | MGTIQEAGKI | MGTIQEAGKI | V344I | -1.2715396 | -1.2756416 | -0.004102 |
| BNIPL | MGTIQEAGKI | MGTIQEAGKI | R345K | -1.2715396 | -1.2761873 | -0.0046477 |
| BNIPL | MGTIQEAGKI | MGTIQEAGKI | L347I | -1.2715396 | -1.2813333 | -0.0097938 |
| BNIPL | MGTIQEAGKI | MGTIQEAGKI | R349Q | -1.2715396 | -1.2708638 | 0.0006758 |
| BNIPL | MGTIQEAGKI | MGTIQEAGKI | D350E | -1.2715396 | -1.2686136 | 0.00292599 |
| BNIPL | MGTIQEAGKI | MGTIQEAGKI | H352D | -1.2715396 | -1.278993 | -0.0074534 |
| BNIPL | MGTIQEAGKI | MGTIQEAGKI | S354K | -1.2715396 | -1.2708527 | 0.00068688 |
| BNIPL | MGTIQEAGKI | MGTIQEAGKI | G355Q | -1.2715396 | -1.2732209 | -0.0016813 |

| | | | | | | |
|---|---|---|---|---|---|---|
| BNIPL | MGTIQEAGK | MGTIQEAGK | G356E | -1.2715396 | -1.2731701 | -0.0016305 |
| BNIPL | MGTIQEAGK | MGTIQEAGK | T357E | -1.2715396 | -1.2680049 | 0.00353467 |
| PIK3C2G | MAYSWQTDF | MAYSWQTDF | N426M | -2.5377605 | -2.5385914 | -0.0008309 |
| SVEP1 | MWPRLAFCC | MWPRLAFCC | V494L | -2.1932523 | -2.1934931 | -0.0002408 |
| SVEP1 | MWPRLAFCC | MWPRLAFCC | T625I | -2.1932523 | -2.1944025 | -0.0011501 |
| SVEP1 | MWPRLAFCC | MWPRLAFCC | G631N | -2.1932523 | -2.1944089 | -0.0011566 |
| SVEP1 | MWPRLAFCC | MWPRLAFCC | S832L | -2.1932523 | -2.1933961 | -0.0001438 |
| SVEP1 | MWPRLAFCC | MWPRLAFCC | S3235V | -2.1932523 | -2.1936188 | -0.0003664 |
| RAPGEF3 | MAQREEIKRF | MAQREEIKRF | V906T | -1.3789129 | -1.3764441 | 0.00246882 |
| XRCC6 | MSGWESYYK | MSGWESYYK | Q68W | -2.5056469 | -2.5360823 | -0.0304353 |
| IFNAR1 | MMVVLLGATI | MMVVLLGATI | T68E | -2.5690446 | -2.5674832 | 0.0015614 |
| CYP19A1 | MVLEMLNPIH | MVLEMLNPIH | V158M | -1.3306938 | -1.334564 | -0.0038701 |
| CYP19A1 | MVLEMLNPIH | MVLEMLNPIH | K461H | -1.3306938 | -1.3353028 | -0.004609 |
| METTL21B | MADPGPDPE | MADPGPDPE | H194V | -1.4093742 | -1.4321102 | -0.022736 |
| METTL21B | MADPGPDPE | MADPGPDPE | Q198P | -1.4093742 | -1.4387376 | -0.0293634 |
| HIF1A | MSSQCRSLEI | MSSQCRSLEI | S616R | -1.4424231 | -1.4455113 | -0.0030882 |
| MTFMT | MRVLVRRCW | MRVLVRRCW | G16V | -1.7106652 | -1.7141645 | -0.0034993 |
| MTFMT | MRVLVRRCW | MRVLVRRCW | S317L | -1.7106652 | -1.7263839 | -0.0157187 |
| GZMB | MQPILLLLAFL | MQPILLLLAFL | K92Y | -1.0996158 | -1.1256998 | -0.0260839 |
| GZMB | MQPILLLLAFL | MQPILLLLAFL | Q149S | -1.0996158 | -1.1128888 | -0.013273 |
| GZMB | MQPILLLLAFL | MQPILLLLAFL | Q214Y | -1.0996158 | -1.136348 | -0.0367322 |
| GHSR | MWNATPSEE | MWNATPSEE | R336S | -1.016259 | -1.0256501 | -0.0093912 |
| BAG6 | MEPNDSTSTA | MEPNDSTSTA | V1055L | -1.6637988 | -1.6633899 | 0.00040889 |
| ZPLD1 | MLLPPPFCLF | MLLPPPFCLF | T38I | -1.0737329 | -1.0712173 | 0.00251555 |
| SYT13 | MVLSVPVIALC | MVLSVPVIALC | A159I | -1.868283 | -1.8720051 | -0.0037221 |
| POLM | MLPKRRRARV | MLPKRRRARV | T21A | -2.0939085 | -2.0919652 | 0.00194335 |
| POLM | MLPKRRRARV | MLPKRRRARV | P146Q | -2.0939085 | -2.0972812 | -0.0033727 |
| POLM | MLPKRRRARV | MLPKRRRARV | L331T | -2.0939085 | -2.1117988 | -0.0178902 |
| POLM | MLPKRRRARV | MLPKRRRARV | L359M | -2.0939085 | -2.104593 | -0.0106845 |
| AKT1S1 | MSFEGGDGA | MSFEGGDGA | P102T | -2.6030049 | -2.6132755 | -0.0102706 |
| TEP1 | MEKLHGHVS | MEKLHGHVS | A1976V | -2.5834891 | -2.5838693 | -0.0003802 |
| TEP1 | MEKLHGHVS | MEKLHGHVS | S2001L | -2.5834891 | -2.5840851 | -0.000596 |
| CCL5 | MKVSAAALAV | MKVSAAALAV | A23P | -2.6190763 | -2.635056 | -0.0159798 |
| ENOX1 | MWRRESSPC | MWRRESSPC | C443S | -0.625618 | -0.629649 | -0.004031 |
| RBPMS | MNNGGKAEK | MNNGGKAEK | N17G | -1.0231057 | -1.04497 | -0.0218643 |
| MRE11A | MSTADALDD | MSTADALDD | N617S | -1.1593493 | -1.1585159 | 0.00083339 |
| MRE11A | MSTADALDD | MSTADALDD | D622G | -1.1593493 | -1.1633286 | -0.0039793 |
| CPA2 | MAMRLILFFG | MAMRLILFFG | N194E | -1.1299925 | -1.1418147 | -0.0118222 |
| NAV2 | MPAILVASKM | MPAILVASKM | S1179C | -1.5068064 | -1.5141732 | -0.0073668 |
| LINGO2 | MLHTAISCWC | MLHTAISCWC | I470V | -0.5846918 | -0.5888481 | -0.0041562 |
| LINGO2 | MLHTAISCWC | MLHTAISCWC | V591I | -0.5846918 | -0.5906238 | -0.005932 |
| ZFYVE28 | MMNRFRKW | MMNRFRKW | G592K | -2.3054521 | -2.306545 | -0.0010929 |
| ECHS1 | MAALRVLLSC | MAALRVLLSC | V47I | -0.7994099 | -0.8121139 | -0.012704 |
| GZMH-aging- | MQPFLLLLAF | MQPFLLFLAF | L7F | -1.2358335 | -1.2462629 | -0.0104294 |
| GZMH-aging- | MQPFLLLLAF | MQPFLLLIAFL | L8I | -1.2358335 | -1.2531158 | -0.0172822 |

| | | | | | | |
|---|---|---|---|---|---|---|
| GZMH-aging- | MQPFLLLLAF | MQPFLLLLAF | L11P | -1.2358335 | -1.2736838 | -0.0378503 |
| GZMH-aging- | MQPFLLLLAF | MQPFLLLLAF | T13L | -1.2358335 | -1.220873 | 0.01496053 |
| GZMH-aging- | MQPFLLLLAF | MQPFLLLLAF | G17Q | -1.2358335 | -1.2421942 | -0.0063607 |
| PPM1D | MAGLYSLGVS | MAGLYSLGVS | V440I | -1.5628524 | -1.5631301 | -0.0002778 |
| POLE2 | MAPERLRSR/ | MAPERLRSR/ | L107V | -1.0827104 | -1.0959533 | -0.013243 |
| TOP2A | MEVSPLQPVN | MEVSPLQPVN | P681S | -1.6991129 | -1.7043962 | -0.0052834 |
| PPP1R26 | MFLMNASPV | MFLMNASPV | D503G | -2.7047558 | -2.704422 | 0.00033379 |
| ERCC1 | MDPGKDKEG | MDPGKDKEG | L69S | -1.4120241 | -1.4103539 | 0.00167024 |
| CSMD3 | MKGIRKGESF | AKGIRKGESR | M1A | -1.615037 | -1.615788 | -0.000751 |
| CSMD3 | MKGIRKGESF | MKGIRKGESF | V1966M | -1.615037 | -1.6152024 | -0.0001655 |
| CRLS1 | MLALRVARGS | MLALRVARGS | V298L | -1.037958 | -1.0570759 | -0.0191178 |
| APOA4 | MFLKAVVLTL/ | MFLKAVVLTL/ | A268N | -1.9898657 | -1.9952734 | -0.0054077 |
| APOB | MDPPRPALL/ | MDPPRPALL/ | R2012Q | -2.7897675 | -2.7897385 | 2.90E-05 |
| APOB | MDPPRPALL/ | MDPPRPALL/ | A4286S | -2.7897675 | -2.7894711 | 0.0002964 |
| HDAC6 | MSGANRGR( | MSGANRGR( | N60R | -2.2413723 | -2.2408814 | 0.0004909 |
| WDR27 | MENPQDIFS! | MENPQDIFS! | E651K | -2.5799966 | -2.5830004 | -0.0030038 |
| HIP1 | MDRMASSMI | MDRMASSMI | D344G | -2.1217833 | -2.1270878 | -0.0053046 |
| CEL | MLTMGRLQL' | MLTMGRLQL' | L542M | -1.3706673 | -1.3713292 | -0.0006618 |
| TERT | MPRAPRCRA | MPRAPRCRA | R19S | -2.7952516 | -2.7955842 | -0.0003326 |
| TERT | MPRAPRCRA | MPRAPRCRA | L593Y | -2.7952516 | -2.7959092 | -0.0006576 |
| NCAM2 | MVRSDSGG( | MVRSDSGG( | V163I | -0.7299464 | -0.7356575 | -0.0057111 |
| NCAM2 | MVRSDSGG( | MVRSDSGG( | R307E | -0.7299464 | -0.7390199 | -0.0090735 |
| LEP | MHWGTLCG | MHWGTLCG | L147V | -2.8554606 | -2.8667202 | -0.0112596 |
| PARVG | MVPAKEQVP( | MVPAKEQVP( | L339P | -1.6979629 | -1.7756474 | -0.0776845 |
| PARVG | MVPAKEQVP( | MVPAKEQVP( | N341Q | -1.6979629 | -1.7219769 | -0.024014 |
| XDH | MTADKLVFFV | MTADKLVFFV | T510M | -1.5678272 | -1.5722091 | -0.0043819 |
| APOM | MFHQIWAAL | MFHQIWAAL | V70I | -2.8068364 | -2.8076649 | -0.0008285 |
| CHIT1 | MEKQFERGQ | MEKQFERGQ | D213K | -1.3390691 | -1.3438258 | -0.0047567 |
| LMNB1 | MATATPVPPR | MATATPVPPR | T96M | -0.6878729 | -0.711997 | -0.0241241 |
| TPP2 | MATAATEEPF | MATAATEEPF | A636L | -2.0641401 | -2.0713553 | -0.0072153 |
| POLE | MSLRSGGRR | MSLRSGGRR | V146M | -2.8298216 | -2.8301277 | -0.0003061 |
| POLE | MSLRSGGRR | MSLRSGGRR | R1231H | -2.8298216 | -2.8308627 | -0.0010411 |
| GSR | MALLPRALSA( | MALLPRALSA( | S120L | -0.9108763 | -0.9261461 | -0.0152698 |
| HLA-DRB1 | MVCLKLPGG! | MVCLKGPGG | L6G | -0.3375464 | -0.4298016 | -0.0922552 |
| HLA-DRB1 | MVCLKLPGG! | MVCLKLPGSS | G9S | -0.3375464 | -0.3989507 | -0.0614043 |
| HLA-DRB1 | MVCLKLPGG! | MVCLKLPGG( | S10C | -0.3375464 | -0.377492 | -0.0399456 |
| HLA-DRB1 | MVCLKLPGG! | MVCLKLPGG! | C11W | -0.3375464 | -0.3861094 | -0.0485629 |
| HLA-DRB1 | MVCLKLPGG! | MVCLKLPGG! | M12A | -0.3375464 | -0.3849722 | -0.0474258 |
| HLA-DRB1 | MVCLKLPGG! | MVCLKLPGG! | M20T | -0.3375464 | -0.3992897 | -0.0617433 |
| HLA-DRB1 | MVCLKLPGG! | MVCLKLPGG! | R33P | -0.3375464 | -0.3616713 | -0.0241249 |
| HLA-DRB1 | MVCLKLPGG! | MVCLKLPGG! | P34R | -0.3375464 | -0.3785293 | -0.0409829 |
| HLA-DRB1 | MVCLKLPGG! | MVCLKLPGG! | R35G | -0.3375464 | -0.3799454 | -0.0423989 |
| HLA-DRB1 | MVCLKLPGG! | MVCLKLPGG! | L37V | -0.3375464 | -0.3680289 | -0.0304824 |
| HLA-DRB1 | MVCLKLPGG! | MVCLKLPGG! | W38Y | -0.3375464 | -0.3367476 | 0.00079888 |
| HLA-DRB1 | MVCLKLPGG! | MVCLKLPGG! | P40T | -0.3375464 | -0.3409475 | -0.0034011 |

| | | | | | | |
|---|---|---|---|---|---|---|
| HLA-DRB1 | MVCLKLPGG | MVCLKLPGG | K41N | -0.3375464 | -0.3649175 | -0.027371 |
| HLA-DRB1 | MVCLKLPGG | MVCLKLPGG | H45Y | -0.3375464 | -0.3672246 | -0.0296781 |
| HLA-DRB1 | MVCLKLPGG | MVCLKLPGG | S66H | -0.3375464 | -0.3498005 | -0.012254 |
| HLA-DRB1 | MVCLKLPGG | MVCLKLPGG | V67L | -0.3375464 | -0.3446681 | -0.0071217 |
| HLA-DRB1 | MVCLKLPGG | MVCLKLPGG | F76W | -0.3375464 | -0.3774477 | -0.0399013 |
| HLA-DRB1 | MVCLKLPGG | MVCLKLPGG | P85A | -0.3375464 | -0.3640129 | -0.0264664 |
| HLA-DRB1 | MVCLKLPGG | MVCLKLPGG | D86T | -0.3375464 | -0.3582272 | -0.0206807 |
| HLA-DRB1 | MVCLKLPGG | MVCLKLPGG | Y89L | -0.3375464 | -0.3616464 | -0.0241 |
| HLA-DRB1 | MVCLKLPGG | MVCLKLPGG | I96S | -0.3375464 | -0.3642221 | -0.0266756 |
| HLA-DRB1 | MVCLKLPGG | MVCLKLPGG | E98Q | -0.3375464 | -0.3665152 | -0.0289688 |
| HLA-DRB1 | MVCLKLPGG | MVCLKLPGG | Q99Y | -0.3375464 | -0.3668081 | -0.0292616 |
| HLA-DRB1 | MVCLKLPGG | MVCLKLPGG | A100K | -0.3375464 | -0.3390537 | -0.0015073 |
| HLA-DRB1 | MVCLKLPGG | MVCLKLPGG | G113Q | -0.3375464 | -0.3635682 | -0.0260218 |
| HLA-DRB1 | MVCLKLPGG | MVCLKLPGG | V114E | -0.3375464 | -0.3703613 | -0.0328149 |
| HLA-DRB1 | MVCLKLPGG | MVCLKLPGG | V115T | -0.3375464 | -0.360355 | -0.0228086 |
| HLA-DRB1 | MVCLKLPGG | MVCLKLPGG | Q121S | -0.3375464 | -0.3704114 | -0.032865 |
| HLA-DRB1 | MVCLKLPGG | MVCLKLPGG | T129K | -0.3375464 | -0.3672974 | -0.029751 |
| HLA-DRB1 | MVCLKLPGG | MVCLKLPGG | Y131S | -0.3375464 | -0.3655245 | -0.027978 |
| HLA-DRB1 | MVCLKLPGG | MVCLKLPGG | L138E | -0.3375464 | -0.3922085 | -0.0546621 |
| HLA-DRB1 | MVCLKLPGG | MVCLKLPGG | H141P | -0.3375464 | -0.3719755 | -0.034429 |
| HLA-DRB1 | MVCLKLPGG | MVCLKLPGG | S149T | -0.3375464 | -0.3528998 | -0.0153533 |
| HLA-DRB1 | MVCLKLPGG | MVCLKLPGG | Q165L | -0.3375464 | -0.3704219 | -0.0328754 |
| HLA-DRB1 | MVCLKLPGG | MVCLKLPGG | T210Q | -0.3375464 | -0.3688959 | -0.0313495 |
| HLA-DRB1 | MVCLKLPGG | MVCLKLPGG | S211G | -0.3375464 | -0.3752684 | -0.037722 |
| HRH1 | MSLPNSSCL | MSLPNSSCL | V42I | -1.7941946 | -1.7949868 | -0.0007923 |
| SH2D4A | MLKQILSEMY | MLKQILSEMY | R334Q | -1.7135504 | -1.7275637 | -0.0140133 |
| NEDD4 | MAQSLRLHF | MAQSLRLHF | K261Q | -1.3014973 | -1.3019328 | -0.0004355 |
| NEDD4 | MAQSLRLHF | MAQSLRLHF | K347T | -1.3014973 | -1.3016798 | -0.0001825 |
| NEDD4 | MAQSLRLHF | MAQSLRLHF | F753A | -1.3014973 | -1.3029398 | -0.0014424 |
| NEDD4 | MAQSLRLHF | MAQSLRLHF | S811A | -1.3014973 | -1.3020991 | -0.0006018 |
| MARCO | MRNKKILKED | MRNKKILKED | L74M | -1.9242253 | -1.9301236 | -0.0058982 |
| MAPK1 | MAAAAAAGA | MAAAAAAGA | Y358D | -0.1609344 | -0.2094409 | -0.0485065 |
| CFH | MRLLAKIICL | MRLLAKIICL | N718D | -2.1752539 | -2.1759202 | -0.0006664 |
| CFH | MRLLAKIICL | MRLLAKIICL | S720D | -2.1752539 | -2.1751533 | 0.00010061 |
| CFH | MRLLAKIICL | MRLLAKIICL | S722R | -2.1752539 | -2.17556 | -0.0003061 |
| CFH | MRLLAKIICL | MRLLAKIICL | T724K | -2.1752539 | -2.1752629 | -9.06E-06 |
| CFH | MRLLAKIICL | MRLLAKIICL | H728S | -2.1752539 | -2.1740718 | 0.00118208 |
| CFH | MRLLAKIICL | MRLLAKIICL | S730K | -2.1752539 | -2.1752429 | 1.10E-05 |
| CFH | MRLLAKIICL | MRLLAKIICL | T732E | -2.1752539 | -2.1759481 | -0.0006943 |
| CFH | MRLLAKIICL | MRLLAKIICL | H735D | -2.1752539 | -2.1733828 | 0.00187111 |
| CFH | MRLLAKIICL | MRLLAKIICL | V745T | -2.1752539 | -2.1757824 | -0.0005286 |
| GCNT3 | MVQWKRLC | MVQWKYLC | R6Y | -1.7005007 | -1.7045522 | -0.0040514 |
| DHX33 | MPEEAGFPP | MPEGAGFPP | E4G | -0.8549162 | -0.854968 | -5.17E-05 |
| NFATC1 | MPSTSFPVPS | MPSTSFPVPS | V928A | -1.2921261 | -1.2974169 | -0.0052909 |
| PLEKHA1 | MPYVDRQN | MPYVDRQN | P294S | -0.9318267 | -0.9496374 | -0.0178107 |

| | | | | | | |
|---|---|---|---|---|---|---|
| PUM2 | MNHDFQALA | MNHDFQALA | N242S | -1.3875917 | -1.3920323 | -0.0044405 |
| TREML1 | MGLTLLLLLLL | MGLTPLLLLLL | L5P | -2.3393931 | -2.3422179 | -0.0028248 |
| TREML1 | MGLTLLLLLLL | MGLTLLLLLLL | G15E | -2.3393931 | -2.3457441 | -0.006351 |
| TREML1 | MGLTLLLLLLL | MGLTLLLLLLL | F54V | -2.3393931 | -2.3474088 | -0.0080156 |
| TREML1 | MGLTLLLLLLL | MGLTLLLLLLL | R110L | -2.3393931 | -2.3480756 | -0.0086825 |
| TNFSF8 | MDPGLQQAL | MDPGLQQAL | P69H | -2.8226457 | -2.8273356 | -0.0046899 |
| PKN2 | MASNPERGE | MASNPERGE | E613G | -0.7849018 | -0.78578 | -0.0008782 |
| CD302 | MLRAALPALLI | MLRAALPALLI | M110V | -2.0321579 | -2.0286369 | 0.00352097 |
| IRG1 | MLANSSITESI | MLANSSITESI | K150S | -1.6902108 | -1.6933913 | -0.0031805 |
| MID2 | MGESPASVVL | MGESPASVVL | V59L | -0.5846902 | -0.5917346 | -0.0070444 |
| DHX9 | MGDVKNFLY | MGDVKNFLY | T108A | -1.9777673 | -1.9775752 | 0.00019217 |
| C5AR1 | MDSFNYTTPI | MDSFNYTTPI | F182N | -1.9267681 | -1.9290183 | -0.0022502 |
| IL1RL2 | MTGLVSLSYF | MTGLVSLSYF | G127S | -2.1878345 | -2.186754 | 0.00108051 |
| ATF6 | MGEPAGVAG | MGEPAGVAG | R403S | -2.1569519 | -2.1543083 | 0.00264359 |
| ATF6 | MGEPAGVAG | MGEPAGVAG | T606K | -2.1569519 | -2.1565864 | 0.0003655 |
| ST6GAL1 | MIHTNLKKKF | MIHTNLKKKF | D146G | -1.7581744 | -1.7668996 | -0.0087252 |
| LY75 | MRTGWATPR | MRTGWATPR | P153S | -2.4958332 | -2.4960701 | -0.000237 |
| LY75 | MRTGWATPR | MRTGWATPR | K488N | -2.4958332 | -2.4967079 | -0.0008748 |
| LY75 | MRTGWATPR | MRTGWATPR | Q959E | -2.4958332 | -2.496037 | -0.0002038 |
| LY75 | MRTGWATPR | MRTGWATPR | T1354H | -2.4958332 | -2.4966426 | -0.0008094 |
| LY75 | MRTGWATPR | MRTGWATPR | K1519Q | -2.4958332 | -2.4962659 | -0.0004327 |
| RABGEF1 | MMASSYHEV | MMASSYHEV | V26L | -0.3226091 | -0.3348524 | -0.0122433 |
| CD1D | MGCLLFLLLW | MGCLLFLLLW | G46T | -1.8123957 | -1.8246678 | -0.0122721 |
| CD1D | MGCLLFLLLW | MGCLLFLLLW | K232Q | -1.8123957 | -1.8251218 | -0.0127261 |
| ADAR | MMSPICDQT | MMSPICDQT | G488D | -2.4387619 | -2.43891 | -0.0001481 |
| ATP7A | MDPSMGVNS | MDPSMGVNS | R512G | -1.4430548 | -1.4463842 | -0.0033294 |
| ATP7A | MDPSMGVNS | MDPSMGVNS | D828G | -1.4430548 | -1.446996 | -0.0039412 |
| ATP7A | MDPSMGVNS | MDPSMGVNS | Q1149N | -1.4430548 | -1.4426651 | 0.0003897 |
| PSMD5 | MAAQALALLR | MAAQALALLR | S266A | -1.3093754 | -1.3211951 | -0.0118197 |
| TRIM8 | MAENWKNCI | MAENWKNCI | V225L | -1.9119438 | -1.9195867 | -0.0076429 |
| PIK3AP1 | MAASGVPRG | MAASGVPRG | R706I | -2.254102 | -2.2545242 | -0.0004222 |
| PIK3AP1 | MAASGVPRG | MAASGVPRG | L709M | -2.254102 | -2.256211 | -0.0021091 |
| APOBEC2 | MAQKEEAAVA | MAQKEEAAVA | L24S | -0.9308188 | -0.9467039 | -0.0158851 |
| DAPK1 | MTVFRQENVI | MTVFRQENVI | P1355S | -2.1845577 | -2.1837261 | 0.0008316 |
| STK11 | MEVVDPQQL | MEVVDPQQL | K108R | -1.2975103 | -1.3042524 | -0.0067421 |
| NFKBIZ | MIVDKLLDDS | MIVDKLLDDS | C177H | -1.9323171 | -1.9310902 | 0.0012269 |
| NFKBIZ | MIVDKLLDDS | MIVDKLLDDS | I353V | -1.9323171 | -1.9315385 | 0.00077868 |
| TRAT1 | MSGISGCPFI | MSGISGCPFI | K92D | -2.8424814 | -2.8424337 | 4.77E-05 |
| IL17C | MTLLPGLLFLT | MTLLPGLLFLT | L70V | -2.1324255 | -2.1377831 | -0.0053575 |
| CD38 | MANCEFSPVS | MANCEFSPVS | R47M | -2.7365932 | -2.7436402 | -0.0070469 |
| FGA | MFSMRIVCLV | MFSMRIVCLV | P352E | -2.314043 | -2.3178394 | -0.0037963 |
| IFNL3 | MKLDMTGDC | MKLDMTGDC | Q52W | -2.6232674 | -2.6464758 | -0.0232084 |
| IFNL3 | MKLDMTGDC | MKLDMTGDC | Q55K | -2.6232674 | -2.6335917 | -0.0103242 |
| IFNL3 | MKLDMTGDC | MKLDMTGDC | A165T | -2.6232674 | -2.6190548 | 0.00421262 |
| DHX58 | MELRSYQWE | MELRSYQWE | T395S | -2.0201998 | -2.0177305 | 0.0024693 |

| | | | | | | |
|---|---|---|---|---|---|---|
| MADCAM1 | MDFGLALLLA | MDFGLALLLA | E263S | -2.354799 | -2.3846476 | -0.0298486 |
| MADCAM1 | MDFGLALLLA | MDFGLALLLA | R292S | -2.354799 | -2.3518982 | 0.00290084 |
| MADCAM1 | MDFGLALLLA | MDFGLALLLA | P355A | -2.354799 | -2.3573496 | -0.0025506 |
| 8-Mar | MSMPLHQIS/ | MSMPLHQIS/ | T538I | -1.6010056 | -1.6080021 | -0.0069965 |
| CD207 | MTVEKEAPDA | STVEKEAPDA | M1S | -2.321537 | -2.3359013 | -0.0143642 |
| CD207 | MTVEKEAPDA | MTVEKEDPD/ | A7D | -2.321537 | -2.3182483 | 0.00328875 |
| CD207 | MTVEKEAPDA | MTVEKEAPDA | K38E | -2.321537 | -2.3216245 | -8.75E-05 |
| CD207 | MTVEKEAPDA | MTVEKEAPDA | P40S | -2.321537 | -2.3194525 | 0.00208449 |
| CD207 | MTVEKEAPDA | MTVEKEAPDA | T50C | -2.321537 | -2.3160479 | 0.00548911 |
| CD207 | MTVEKEAPDA | MTVEKEAPDA | N77E | -2.321537 | -2.3150475 | 0.00648952 |
| CD207 | MTVEKEAPDA | MTVEKEAPDA | G83E | -2.321537 | -2.3156357 | 0.00590134 |
| CD207 | MTVEKEAPDA | MTVEKEAPDA | R84N | -2.321537 | -2.3189905 | 0.00254655 |
| CD207 | MTVEKEAPDA | MTVEKEAPDA | N87D | -2.321537 | -2.3237453 | -0.0022082 |
| CD207 | MTVEKEAPDA | MTVEKEAPDA | Q197S | -2.321537 | -2.3217106 | -0.0001736 |
| CD207 | MTVEKEAPDA | MTVEKEAPDA | T231A | -2.321537 | -2.3143909 | 0.00714612 |
| CD207 | MTVEKEAPDA | MTVEKEAPDA | A246I | -2.321537 | -2.3240213 | -0.0024843 |
| CD207 | MTVEKEAPDA | MTVEKEAPDA | D263T | -2.321537 | -2.3198438 | 0.00169325 |
| CD207 | MTVEKEAPDA | MTVEKEAPDA | S265C | -2.321537 | -2.3308372 | -0.0093002 |
| CD207 | MTVEKEAPDA | MTVEKEAPDA | R279E | -2.321537 | -2.3269112 | -0.0053742 |
| CD207 | MTVEKEAPDA | MTVEKEAPDA | N291T | -2.321537 | -2.3274844 | -0.0059474 |
| MAP4K2 | MALLRDVSLC | MALLRDVSLC | N349S | -0.8766817 | -0.8749176 | 0.00176412 |
| MAP4K2 | MALLRDVSLC | MALLRDVSLC | V723I | -0.8766817 | -0.8806765 | -0.0039948 |
| JAG1 | MRSPRTRGR | MRSPRTRGR | G1100T | -1.918468 | -1.917048 | 0.00142002 |
| ATG9A | MAQFDTEYQ | MMQFDTEYC | A2M | -1.6728463 | -1.6755363 | -0.00269 |
| ATG9A | MAQFDTEYQ | MAQFDTEYQ | P360L | -1.6728463 | -1.7009031 | -0.0280567 |
| ATG9A | MAQFDTEYQ | MAQFDTEYQ | L635V | -1.6728463 | -1.6736611 | -0.0008148 |
| VAV3 | MTQSLSYLATI | MTQSLSYLATI | Q634Y | -0.7548214 | -0.7593246 | -0.0045032 |
| TNIP3 | MIPCGWLMN | MIPCGWLMN | G114T | -2.2846229 | -2.2829618 | 0.00166106 |
| TMEM173 | MPHSSLHPSI | MPHSSLHPSI | E249D | -2.7484498 | -2.7496769 | -0.0012271 |
| LAMP1 | MAAPGSARRI | MAAPGSARRI | T39E | -2.1508224 | -2.1529903 | -0.0021679 |
| ITGAL | MKDSCITVM/ | MKDSCITVM/ | M1125L | -2.5233634 | -2.5182936 | 0.00506973 |
| PROS1 | MRVLGGRCC | MRVLSGRCG | G5S | -1.8471661 | -1.8476477 | -0.0004816 |
| PROS1 | MRVLGGRCC | MRVLGGRCC | S428N | -1.8471661 | -1.8547766 | -0.0076106 |
| ATG16L1 | MSSGLRAADI | MSSGLRAADI | G583E | -0.4942313 | -0.5173606 | -0.0231293 |
| GCH1 | MEKGPVRAP/ | MEKGPVRAP/ | P28Q | -0.7580977 | -0.7629569 | -0.0048591 |
| CTSG | MQPLLLLLAFI | MQPLLLLLAFI | R133L | -1.508136 | -1.5273757 | -0.0192397 |
| CTSG | MQPLLLLLAFI | MQPLLLLLAFI | N139S | -1.508136 | -1.4995759 | 0.00856018 |
| CTSG | MQPLLLLLAFI | MQPLLLLLAFI | R163Q | -1.508136 | -1.5137956 | -0.0056596 |
| CTSG | MQPLLLLLAFI | MQPLLLLLAFI | G191R | -1.508136 | -1.4924231 | 0.01571298 |
| CTSG | MQPLLLLLAFI | MQPLLLLLAFI | S234A | -1.508136 | -1.5106753 | -0.0025393 |
| COLEC12 | MKDDFAEEEI | MKDDFAEEEI | P493L | -1.6348563 | -1.6405724 | -0.0057161 |
| COLEC12 | MKDDFAEEEI | MKDDFAEEEI | E672A | -1.6348563 | -1.6411356 | -0.0062792 |
| CD97 | MGGRVFLAF | MGGRVFLAF | F51T | -1.8091878 | -1.8109006 | -0.0017128 |
| CD97 | MGGRVFLAF | MGGRVFLAF | T59M | -1.8091878 | -1.8105223 | -0.0013345 |
| CD97 | MGGRVFLAF | MGGRVFLAF | S72R | -1.8091878 | -1.8108056 | -0.0016178 |

| | | | | | | |
|---|---|---|---|---|---|---|
| CD97 | MGGRVFLAF( | MGGRVFLAF( | K107T | -1.8091878 | -1.8086029 | 0.00058484 |
| CD97 | MGGRVFLAF( | MGGRVFLAF( | V296H | -1.8091878 | -1.8095734 | -0.0003856 |
| CD97 | MGGRVFLAF( | MGGRVFLAF( | G348N | -1.8091878 | -1.8114314 | -0.0022436 |
| CD97 | MGGRVFLAF( | MGGRVFLAF( | A391H | -1.8091878 | -1.8109159 | -0.0017282 |
| CD97 | MGGRVFLAF( | MGGRVFLAF( | R440H | -1.8091878 | -1.8099098 | -0.0007221 |
| CD97 | MGGRVFLAF( | MGGRVFLAF( | I710M | -1.8091878 | -1.8146019 | -0.0054141 |
| MASP2 | MRLLTLLGLL( | MRLLTLLGLL( | S112A | -1.5764204 | -1.5802222 | -0.0038018 |
| MASP2 | MRLLTLLGLL( | MRLLTLLGLL( | S185E | -1.5764204 | -1.5829737 | -0.0065533 |
| SEMA4A | MALPALGLDF | MALPALGLDF | A419T | -1.86554 | -1.8687345 | -0.0031945 |
| SEMA4A | MALPALGLDF | MALPALGLDF | G757R | -1.86554 | -1.8657688 | -0.0002288 |
| LPXN | MSTLLISSSD/ | MSTLLISSSD/ | P383T | -1.1153804 | -1.1442008 | -0.0288204 |
| THEMIS | MALSLEEFVH | MALSLEEFVH | D630N | -2.8118901 | -2.81195 | -5.98E-05 |
| ANG | MVMGLGVLL | MVMGLGVMI | L8M | -2.7897038 | -2.8092644 | -0.0195606 |
| APEX2 | MLRVVSWNII | MLRVVSWNII | R132C | -2.2575438 | -2.2731647 | -0.0156209 |
| APEX2 | MLRVVSWNII | MLRVVSWNII | A133T | -2.2575438 | -2.2617314 | -0.0041876 |
| OASL | MALMQELYST | MALMQELYST | L119V | -2.7250328 | -2.7263868 | -0.001354 |
| EXOSC3 | MAEPASVAAE | MAEPASVAAE | Q39Y | -1.0500017 | -1.0574179 | -0.0074161 |
| CFP | MITEGAQAPR | MITEGAQAPR | K44E | -2.2638605 | -2.2641749 | -0.0003145 |
| POU2F2 | MVHSSMGAF | MVHSSMGAF | G528A | -0.5741461 | -0.5770394 | -0.0028933 |
| ART1 | MQMPAMMS | MQMPAMMS | P126L | -2.6613748 | -2.6641171 | -0.0027423 |
| ART1 | MQMPAMMS | MQMPAMMS | S300N | -2.6613748 | -2.6694882 | -0.0081134 |
| IL18RAP | MLCLGWIFL\ | MLCLGWIFL\ | H55Y | -2.3941185 | -2.392544 | 0.00157452 |
| IL18RAP | MLCLGWIFL\ | MLCLGWIFL\ | Q66R | -2.3941185 | -2.3938138 | 0.0003047 |
| IL18RAP | MLCLGWIFL\ | MLCLGWIFL\ | G117E | -2.3941185 | -2.3906515 | 0.00346708 |
| IL18RAP | MLCLGWIFL\ | MLCLGWIFL\ | V300F | -2.3941185 | -2.397665 | -0.0035465 |
| ATF6B | MAELMLLSEI/ | MAELMLLSEI/ | L381A | -2.0960419 | -2.1186736 | -0.0226316 |
| CD5L | MALLFSLILAI( | MALLFSLILAI( | E84K | -1.8869226 | -1.8890473 | -0.0021247 |
| EGF | MLLTLIILLPVV | MLLTLIILLPVV | R1082S | -2.3463917 | -2.3446593 | 0.00173235 |
| ITCH | MSDSGSQLG | MSDSGSQLG | V24I | -0.7762986 | -0.7784679 | -0.0021693 |
| ITCH | MSDSGSQLG | MSDSGSQLG | N159R | -0.7762986 | -0.7771508 | -0.0008522 |
| IL1RL1 | MGFWILAILTI | MGFWILAILTI | I332T | -2.4442749 | -2.4474392 | -0.0031643 |
| IL1RL1 | MGFWILAILTI | MGFWILAILTI | T391K | -2.4442749 | -2.4442248 | 5.01E-05 |
| IL1RL1 | MGFWILAILTI | MGFWILAILTI | T436M | -2.4442749 | -2.4512181 | -0.0069432 |
| IL1RL1 | MGFWILAILTI | MGFWILAILTI | V537R | -2.4442749 | -2.4485826 | -0.0043077 |
| TAPBP | MKSLSLLLAV/ | MKSLSLLLAV/ | R57K | -2.5911572 | -2.5941393 | -0.0029821 |
| C5 | MLQTTPDLKS | MLQTTPDLKS | D665G | -2.7860556 | -2.7861443 | -8.87E-05 |
| CYP27B1 | MTQTLKYASR | MTQTLKYASR | V397L | -1.6377439 | -1.635866 | 0.0018779 |
| GZMH-immu | MQPFLLLLAF | MQLFLLLLAFI | P3L | -1.2358335 | -1.2403904 | -0.0045569 |
| GZMH-immu | MQPFLLLLAF | MQPFLLFLAF | L7F | -1.2358335 | -1.2462629 | -0.0104294 |
| GZMH-immu | MQPFLLLLAF | MQPFLLLIAFL | L8I | -1.2358335 | -1.2531158 | -0.0172822 |
| GZMH-immu | MQPFLLLLAF | MQPFLLLLAFI | L11P | -1.2358335 | -1.2736838 | -0.0378503 |
| GZMH-immu | MQPFLLLLAF | MQPFLLLLAFI | T13L | -1.2358335 | -1.220873 | 0.01496053 |
| GZMH-immu | MQPFLLLLAF | MQPFLLLLAFI | G17Q | -1.2358335 | -1.2421942 | -0.0063607 |
| ERBB2IP | MTTKRSLFVR | MTTKRSLFVR | D724Q | -1.6998267 | -1.7027812 | -0.0029545 |
| MSH6 | MSRQSTLYSF | MSRQSTLYSF | S602A | -2.2634575 | -2.2641931 | -0.0007355 |

| | | | | | | |
|---|---|---|---|---|---|---|
| MSH6 | MSRQSTLYSF | MSRQSTLYSF | V1056G | -2.2634575 | -2.2660413 | -0.0025837 |
| ABL1 | MGQQPGKVI | MGQQPGKVI | V753M | -2.081449 | -2.0826161 | -0.0011671 |
| SH2B2 | MDPSYCPAH | MDPSYCPAH | S639N | -1.7670945 | -1.770296 | -0.0032015 |
| BMP6 | MPGLGRRAQ | MPGLGRRAQ | D402G | -1.0858319 | -1.1115059 | -0.025674 |
| BMP6 | MPGLGRRAQ | MPGLGRRAQ | K482E | -1.0858319 | -1.105484 | -0.0196521 |
| BMP6 | MPGLGRRAQ | MPGLGRRAQ | L483V | -1.0858319 | -1.1048772 | -0.0190454 |
| BMP6 | MPGLGRRAQ | MPGLGRRAQ | N484M | -1.0858319 | -1.1130608 | -0.027229 |
| BMP6 | MPGLGRRAQ | MPGLGRRAQ | A485R | -1.0858319 | -1.1119088 | -0.0260769 |
| BMP6 | MPGLGRRAQ | MPGLGRRAQ | S487A | -1.0858319 | -1.104131 | -0.0182991 |
| PLCL2 | MAECGRGG/ | MAECGRGG/ | S86G | -1.9857186 | -1.985683 | 3.56E-05 |
| SPPL2A | MWLLGCPEG | MWLLGCPEG | T78S | -1.3447802 | -1.3450925 | -0.0003123 |
| HAVCR2 | MFSHLPFDC | MFSHLPFDC | P138Q | -2.3676059 | -2.3722882 | -0.0046823 |
| ATF7 | MGDDRPFVC | AGDDRPFVC | M1A | -0.5747371 | -0.6191748 | -0.0444377 |
| MR1 | MGELMAFLLF | MGELLAFLLP | M5L | -1.5061132 | -1.5016222 | 0.00449097 |
| MR1 | MGELMAFLLF | MGELMALLLF | F7L | -1.5061132 | -1.504132 | 0.00198114 |
| MR1 | MGELMAFLLF | MGELMAFRLI | L8R | -1.5061132 | -1.5241822 | -0.018069 |
| MR1 | MGELMAFLLF | MGELMAFLLI | P10L | -1.5061132 | -1.5065393 | -0.0004262 |
| MR1 | MGELMAFLLF | MGELMAFLLF | V17L | -1.5061132 | -1.5063293 | -0.0002161 |
| MR1 | MGELMAFLLF | MGELMAFLLF | K18R | -1.5061132 | -1.5105737 | -0.0044606 |
| MR1 | MGELMAFLLF | MGELMAFLLF | D21R | -1.5061132 | -1.5170436 | -0.0109304 |
| MR1 | MGELMAFLLF | MGELMAFLLF | T24L | -1.5061132 | -1.5179558 | -0.0118426 |
| MR1 | MGELMAFLLF | MGELMAFLLF | L27R | -1.5061132 | -1.5395925 | -0.0334793 |
| MR1 | MGELMAFLLF | MGELMAFLLF | Y29I | -1.5061132 | -1.5512149 | -0.0451018 |
| MR1 | MGELMAFLLF | MGELMAFLLF | R31H | -1.5061132 | -1.5049658 | 0.00114739 |
| MR1 | MGELMAFLLF | MGELMAFLLF | E43W | -1.5061132 | -1.5615522 | -0.055439 |
| MR1 | MGELMAFLLF | MGELMAFLLF | F44Y | -1.5061132 | -1.5220718 | -0.0159587 |
| MR1 | MGELMAFLLF | MGELMAFLLF | I45S | -1.5061132 | -1.5267287 | -0.0206156 |
| MR1 | MGELMAFLLF | MGELMAFLLF | S46R | -1.5061132 | -1.5449363 | -0.0388231 |
| MR1 | MGELMAFLLF | MGELMAFLLF | V47I | -1.5061132 | -1.517208 | -0.0110948 |
| MR1 | MGELMAFLLF | MGELMAFLLF | P54V | -1.5061132 | -1.5252789 | -0.0191658 |
| MR1 | MGELMAFLLF | MGELMAFLLF | T57N | -1.5061132 | -1.522123 | -0.0160098 |
| MR1 | MGELMAFLLF | MGELMAFLLF | D59T | -1.5061132 | -1.5453408 | -0.0392276 |
| MR1 | MGELMAFLLF | MGELMAFLLF | T62S | -1.5061132 | -1.5066665 | -0.0005534 |
| MR1 | MGELMAFLLF | MGELMAFLLF | R63Q | -1.5061132 | -1.5084678 | -0.0023546 |
| MR1 | MGELMAFLLF | MGELMAFLLF | K65V | -1.5061132 | -1.5129375 | -0.0068244 |
| MR1 | MGELMAFLLF | MGELMAFLLF | A69S | -1.5061132 | -1.5144007 | -0.0082875 |
| MR1 | MGELMAFLLF | MGELMAFLLF | P70V | -1.5061132 | -1.5382971 | -0.0321839 |
| MR1 | MGELMAFLLF | MGELMAFLLF | A77D | -1.5061132 | -1.5075964 | -0.0014832 |
| MR1 | MGELMAFLLF | MGELMAFLLF | H80F | -1.5061132 | -1.5096748 | -0.0035616 |
| MR1 | MGELMAFLLF | MGELMAFLLF | R83F | -1.5061132 | -1.5406594 | -0.0345463 |
| MR1 | MGELMAFLLF | MGELMAFLLF | T85N | -1.5061132 | -1.5302736 | -0.0241604 |
| MR1 | MGELMAFLLF | MGELMAFLLF | Q86F | -1.5061132 | -1.5539945 | -0.0478814 |
| MR1 | MGELMAFLLF | MGELMAFLLF | L87W | -1.5061132 | -1.5386363 | -0.0325232 |
| MR1 | MGELMAFLLF | MGELMAFLLF | G90R | -1.5061132 | -1.5242698 | -0.0181566 |
| MR1 | MGELMAFLLF | MGELMAFLLF | V97A | -1.5061132 | -1.5167232 | -0.01061 |

| | | | | | | |
|---|---|---|---|---|---|---|
| MR1 | MGELMAFLLF | MGELMAFLLF | H105L | -1.5061132 | -1.5162252 | -0.010112 |
| MR1 | MGELMAFLLF | MGELMAFLLF | T113I | -1.5061132 | -1.5256323 | -0.0195191 |
| MR1 | MGELMAFLLF | MGELMAFLLF | Y114I | -1.5061132 | -1.5256175 | -0.0195043 |
| MR1 | MGELMAFLLF | MGELMAFLLF | R116M | -1.5061132 | -1.5377806 | -0.0316675 |
| MR1 | MGELMAFLLF | MGELMAFLLF | M117I | -1.5061132 | -1.5161456 | -0.0100324 |
| MR1 | MGELMAFLLF | MGELMAFLLF | I118T | -1.5061132 | -1.5163774 | -0.0102643 |
| MR1 | MGELMAFLLF | MGELMAFLLF | E121D | -1.5061132 | -1.5135623 | -0.0074492 |
| MR1 | MGELMAFLLF | MGELMAFLLF | L123Q | -1.5061132 | -1.518373 | -0.0122598 |
| MR1 | MGELMAFLLF | MGELMAFLLF | G126N | -1.5061132 | -1.5250967 | -0.0189835 |
| MR1 | MGELMAFLLF | MGELMAFLLF | S127T | -1.5061132 | -1.5179095 | -0.0117964 |
| MR1 | MGELMAFLLF | MGELMAFLLF | T129W | -1.5061132 | -1.5448946 | -0.0387814 |
| MR1 | MGELMAFLLF | MGELMAFLLF | Y134D | -1.5061132 | -1.5373091 | -0.0311959 |
| MR1 | MGELMAFLLF | MGELMAFLLF | A135S | -1.5061132 | -1.5166199 | -0.0105067 |
| MR1 | MGELMAFLLF | MGELMAFLLF | I143T | -1.5061132 | -1.5145677 | -0.0084546 |
| MR1 | MGELMAFLLF | MGELMAFLLF | N145H | -1.5061132 | -1.5205492 | -0.014436 |
| MR1 | MGELMAFLLF | MGELMAFLLF | D147E | -1.5061132 | -1.5100546 | -0.0039414 |
| MR1 | MGELMAFLLF | MGELMAFLLF | L149M | -1.5061132 | -1.5163753 | -0.0102621 |
| MR1 | MGELMAFLLF | MGELMAFLLF | N156I | -1.5061132 | -1.5290543 | -0.0229411 |
| MR1 | MGELMAFLLF | MGELMAFLLF | V157G | -1.5061132 | -1.5283073 | -0.0221941 |
| MR1 | MGELMAFLLF | MGELMAFLLF | H159Q | -1.5061132 | -1.502256 | 0.00385714 |
| MR1 | MGELMAFLLF | MGELMAFLLF | Q163R | -1.5061132 | -1.5014638 | 0.0046494 |
| MR1 | MGELMAFLLF | MGELMAFLLF | A164R | -1.5061132 | -1.518227 | -0.0121138 |
| MR1 | MGELMAFLLF | MGELMAFLLF | A167T | -1.5061132 | -1.5122885 | -0.0061753 |
| MR1 | MGELMAFLLF | MGELMAFLLF | N168E | -1.5061132 | -1.5179616 | -0.0118484 |
| MR1 | MGELMAFLLF | MGELMAFLLF | Q169I | -1.5061132 | -1.5167433 | -0.0106301 |
| MR1 | MGELMAFLLF | MGELMAFLLF | H170G | -1.5061132 | -1.5207504 | -0.0146372 |
| MR1 | MGELMAFLLF | MGELMAFLLF | E171D | -1.5061132 | -1.5163908 | -0.0102776 |
| MR1 | MGELMAFLLF | MGELMAFLLF | L172Y | -1.5061132 | -1.5237557 | -0.0176425 |
| MR1 | MGELMAFLLF | MGELMAFLLF | Y174H | -1.5061132 | -1.5246148 | -0.0185016 |
| MR1 | MGELMAFLLF | MGELMAFLLF | N177R | -1.5061132 | -1.5218517 | -0.0157385 |
| MR1 | MGELMAFLLF | MGELMAFLLF | E182R | -1.5061132 | -1.5225073 | -0.0163941 |
| MR1 | MGELMAFLLF | MGELMAFLLF | A185P | -1.5061132 | -1.5273466 | -0.0212334 |
| MR1 | MGELMAFLLF | MGELMAFLLF | K188R | -1.5061132 | -1.5112783 | -0.0051651 |
| MR1 | MGELMAFLLF | MGELMAFLLF | R189S | -1.5061132 | -1.5203692 | -0.014256 |
| MR1 | MGELMAFLLF | MGELMAFLLF | F190S | -1.5061132 | -1.5253426 | -0.0192294 |
| MR1 | MGELMAFLLF | MGELMAFLLF | T201K | -1.5061132 | -1.5100977 | -0.0039846 |
| MR1 | MGELMAFLLF | MGELMAFLLF | E202V | -1.5061132 | -1.5218451 | -0.0157319 |
| MR1 | MGELMAFLLF | MGELMAFLLF | L205T | -1.5061132 | -1.4994578 | 0.00665534 |
| MR1 | MGELMAFLLF | MGELMAFLLF | V206A | -1.5061132 | -1.5159934 | -0.0098802 |
| MR1 | MGELMAFLLF | MGELMAFLLF | E212S | -1.5061132 | -1.5090925 | -0.0029793 |
| MR1 | MGELMAFLLF | MGELMAFLLF | F214H | -1.5061132 | -1.5087818 | -0.0026686 |
| MR1 | MGELMAFLLF | MGELMAFLLF | G216R | -1.5061132 | -1.5285368 | -0.0224236 |
| MR1 | MGELMAFLLF | MGELMAFLLF | H225S | -1.5061132 | -1.5198121 | -0.0136989 |
| MR1 | MGELMAFLLF | MGELMAFLLF | E242S | -1.5061132 | -1.5203835 | -0.0142703 |
| MR1 | MGELMAFLLF | MGELMAFLLF | V244Q | -1.5061132 | -1.503897 | 0.00221622 |

| | | | | | | |
|---|---|---|---|---|---|---|
| MR1 | MGELMAFLLF | MGELMAFLLF | D248Y | -1.5061132 | -1.5159546 | -0.0098414 |
| MR1 | MGELMAFLLF | MGELMAFLLF | Y249S | -1.5061132 | -1.5107875 | -0.0046743 |
| MR1 | MGELMAFLLF | MGELMAFLLF | G250E | -1.5061132 | -1.5234425 | -0.0173293 |
| MR1 | MGELMAFLLF | MGELMAFLLF | A264V | -1.5061132 | -1.5121757 | -0.0060625 |
| MR1 | MGELMAFLLF | MGELMAFLLF | I266M | -1.5061132 | -1.5146693 | -0.0085561 |
| MR1 | MGELMAFLLF | MGELMAFLLF | Q271K | -1.5061132 | -1.505695 | 0.00041819 |
| MR1 | MGELMAFLLF | MGELMAFLLF | S272I | -1.5061132 | -1.514684 | -0.0085708 |
| MR1 | MGELMAFLLF | MGELMAFLLF | S273K | -1.5061132 | -1.5062286 | -0.0001154 |
| MR1 | MGELMAFLLF | MGELMAFLLF | N274A | -1.5061132 | -1.5118488 | -0.0057356 |
| MR1 | MGELMAFLLF | MGELMAFLLF | L275H | -1.5061132 | -1.5081133 | -0.0020001 |
| MR1 | MGELMAFLLF | MGELMAFLLF | Q293S | -1.5061132 | -1.5061585 | -4.53E-05 |
| MR1 | MGELMAFLLF | MGELMAFLLF | S295N | -1.5061132 | -1.5099 | -0.0037868 |
| MR1 | MGELMAFLLF | MGELMAFLLF | E296S | -1.5061132 | -1.5055423 | 0.00057089 |
| MR1 | MGELMAFLLF | MGELMAFLLF | V314I | -1.5061132 | -1.5076215 | -0.0015084 |
| MR1 | MGELMAFLLF | MGELMAFLLF | A316V | -1.5061132 | -1.5077977 | -0.0016845 |
| MR1 | MGELMAFLLF | MGELMAFLLF | G317I | -1.5061132 | -1.5108355 | -0.0047224 |
| MR1 | MGELMAFLLF | MGELMAFLLF | V318I | -1.5061132 | -1.5073174 | -0.0012043 |
| MR1 | MGELMAFLLF | MGELMAFLLF | R324K | -1.5061132 | -1.5075041 | -0.0013909 |
| APBB1IP | MGESSEDID( | MGESSEDID( | T349I | -1.4885432 | -1.4969618 | -0.0084187 |
| PRKCG | MAGLGPGVG | MAGLGPGVG | T438N | -0.9010628 | -0.8962319 | 0.0048309 |
| PRKCG | MAGLGPGVG | MAGLGPGVG | P602L | -0.9010628 | -0.9083685 | -0.0073057 |
| AQP9 | MQPEGAEKG | MQPEGAEKG | V62A | -1.1301471 | -1.1467583 | -0.0166112 |
| TINAGL1 | MWRCPLGLL | MWRCPLGLL | H467W | -1.199859 | -1.2087296 | -0.0088706 |
| CD200 | MERLTLTRTIG | MERLTLTRTIG | S157R | -1.4747841 | -1.4950552 | -0.0202711 |
| LY9 | MTGLLGLSPV | MTGLLGLSPV | T120Q | -2.5127833 | -2.5102136 | 0.00256968 |
| LY9 | MTGLLGLSPV | MTGLLGLSPV | I122F | -2.5127833 | -2.5146844 | -0.0019011 |
| LY9 | MTGLLGLSPV | MTGLLGLSPV | A133I | -2.5127833 | -2.5109034 | 0.00187993 |
| LY9 | MTGLLGLSPV | MTGLLGLSPV | E141T | -2.5127833 | -2.5146811 | -0.0018978 |
| LY9 | MTGLLGLSPV | MTGLLGLSPV | K158P | -2.5127833 | -2.5162439 | -0.0034606 |
| LY9 | MTGLLGLSPV | MTGLLGLSPV | N171G | -2.5127833 | -2.5129778 | -0.0001945 |
| LY9 | MTGLLGLSPV | MTGLLGLSPV | N181I | -2.5127833 | -2.5154054 | -0.0026221 |
| LY9 | MTGLLGLSPV | MTGLLGLSPV | Q287D | -2.5127833 | -2.512924 | -0.0001407 |
| LY9 | MTGLLGLSPV | MTGLLGLSPV | P355F | -2.5127833 | -2.5146844 | -0.0019011 |
| SEMA4D | MRMCTPIRG | MRMCTPIRG | G602H | -1.4904218 | -1.4896916 | 0.00073016 |
| ITGB7 | MVALPMVLVL | MVALPMVLVL | Q102R | -1.5626541 | -1.5610936 | 0.00156057 |
| ITGB7 | MVALPMVLVL | MVALPMVLVL | R129A | -1.5626541 | -1.5675684 | -0.0049143 |
| ITGB7 | MVALPMVLVL | MVALPMVLVL | V675M | -1.5626541 | -1.5645642 | -0.0019101 |
| IL31RA | MCIRQLKFFT | MCIRQLKFFT | N536Y | -2.589622 | -2.5919991 | -0.002377 |
| CD22 | MHLLGPWLL | MHLLGPWLL | S123I | -2.3709903 | -2.3751345 | -0.0041442 |
| CD22 | MHLLGPWLL | MHLLGPWLL | H133G | -2.3709903 | -2.3777826 | -0.0067923 |
| CD22 | MHLLGPWLL | MHLLGPWLL | H213D | -2.3709903 | -2.3709209 | 6.94E-05 |
| CD22 | MHLLGPWLL | MHLLGPWLL | H240C | -2.3709903 | -2.3739009 | -0.0029106 |
| CD22 | MHLLGPWLL | MHLLGPWLL | L352R | -2.3709903 | -2.3691609 | 0.00182939 |

**D** Protein language models evaluation of normal Jonathan gene

| Ref | wt_sequence | mutated_seq | mutant | esm1v_t33_6 | esm1v_t33_6 | esm1v_t33_6 |
|---|---|---|---|---|---|---|

| | | | | | | |
|---|---|---|---|---|---|---|
| KIAA1549 | MYYCNKTFVI | MYYCNKTFVI | N105D:S147I | -0.3655791 | -0.1437199 | 0.30035424 |
| ARID1B | MIMAAQVAA | MIMAAQVAA | A894T | -0.632349 | -0.663434 | -0.9962459 |
| GOPC | MSGMGAGS | MSGMGAGS | K112Q | 1.02608871 | 3.88777018 | -0.4797983 |
| ROS1 | MEGELGINA | MEGELGINA | C1274Y:N13 | 0.2571826 | -0.121621 | -1.0026078 |
| RB1 | MPPKPPRKG | MPPKPPRKG | V54H:A245P | -0.488302 | -1.4736743 | -0.0667653 |
| CBLB | MANSLNGRN | MANSLNGRN | P590L:S749G | -1.1675911 | -0.6739261 | -1.2579427 |
| BCL9 | MHSSNPKVR | MHSSNPKVR | V1073M | 0.03089523 | 0.18389463 | 0.09657717 |
| FLT3 | MTVSVFLFGF | MTVSVFLFGF | A19I:I54V:N7 | 2.94322538 | 3.83268476 | 2.09920788 |
| APC2 | MIDYSSARSY | MIDYSSARSY | M195V:Q169 | 0.82928133 | 0.49585104 | 0.42208242 |
| CASC5 | MDRIYSESNI | MDRIYSESNI | D408H:C240 | 3.43644142 | 3.42743325 | 2.62057567 |
| NRG1 | MSEVSAETFP | MSEVSAETFP | H42P | 2.44906807 | 2.57481599 | 1.31431341 |
| CEP89 | MALSFRRGRI | MALSFRRGRI | H698Q | 2.69421959 | 2.95001864 | 2.43455172 |
| NDRG1 | MLLKFIFISKE | MLLKFIFISKE | D75E | 1.7196902 | 1.33814383 | 2.68234062 |
| ALK | MISHGSSCY | MGAIGSSCY | I2G:S3A:H4I: | 2.26841044 | 1.64047027 | 1.60550189 |
| NTRK3 | MLAASERKAR | MLAASERKAR | Y639F | 0.45058823 | 0.67884159 | -0.1633313 |
| CIC | MKPARKAASS | MKPARKAASS | A1146S:P199 | -2.9563284 | -3.1580096 | -2.8401847 |
| DAXX | MAATVESNIIV | MAATVESNIIV | L156S | 0.79195857 | 0.5623467 | 1.10728741 |
| LATS2 | MRPKTFPATT | MRPKTFPATT | S350A | -0.7759488 | -0.7055809 | -0.4152317 |
| NOTCH1 | MQQLLVQSLI | MQQLLVQSLI | T2395N | 1.47511435 | 1.55930305 | 2.72215796 |
| BRD4 | MSVESGPGTI | MSVESGPGTI | I956V | 0.66281033 | 0.84997272 | 0.73090363 |
| RNF213 | MKCPKCDYIS | MKCPKCDYIS | M1803V:N35 | 1.95066381 | 1.8637886 | 1.57472587 |
| THRAP3 | MSKTNKSKS | MSKTNKSKS | L706H | -1.6243868 | -0.3840847 | -2.3720798 |
| SETD2 | MGDFYDPEH | MGDFYDPEH | I1400A:E143 | 0.39577699 | 0.38576484 | 0.43724036 |
| CLIP4 | MTIEDLPEPS | MTIEDLPEPS | A457P | 1.42871165 | 2.74760675 | 2.38882065 |
| PTPRB | MELGGAGDH | MELGGAGDH | V50M:H178V | -11.838004 | -8.4993247 | -10.483913 |
| KMT2D | MPLVRYRKVV | MPLVRYRKVV | L827P | 0.41278815 | 0.45629668 | 0.6218127 |
| NSD1 | MEPACEVSRI | MEPACEVSRI | T406A:F2368 | 2.85260296 | 3.71175432 | 2.53677058 |
| IDH1 | MSSGSLRGYI | MSSGSLRGYI | S267G | 1.1553545 | 0.84920979 | 1.8396039 |
| BCL11B | MNFPLGDILV | MNFPLGDILV | R65L | 0.8388257 | -0.0134621 | 0.40379286 |
| KMT2A | MAHSGRWRI | MAHSGRWRI | R513S | 0.12568164 | 0.04076886 | 0.34770107 |
| ATRX | MRRGRKAAG | MRRGRKAAG | Q1136R | 2.84079456 | 1.80914402 | 1.83081079 |
| NCOA4 | MSPPQDRSG | MSPPQDRSG | R499S | 1.01789975 | 1.71304035 | 1.28082275 |
| ERBB2 | MASIVSAPLQ | MASIVSAPLQ | M162T | 3.22728515 | 4.42718363 | 3.00272179 |
| BRCA1 | MESSVAAIGD | MESSVAAIGD | R1105M | -0.7994261 | -0.5875645 | -0.6833963 |
| CNOT3 | MADKRKLQG | MADKRKLQG | I418A | 1.48996735 | 1.68590534 | 1.52663088 |
| STRN | MDEQAGPG\ | MDEQAGPG\ | H236D | 1.18885517 | 4.36031055 | 0.21996117 |
| TET1 | MAHHARPSR | MAHHARPSR | H1331L | 0.85612035 | 0.8899188 | 1.0927074 |
| TRIM33 | MAENKGGG | MAENKGGG | E808G | 0.529423 | 4.05411816 | -1.534724 |
| SETBP1 | MSSPVGPTHI | MSSPVGPTHI | I393L | 0.72725511 | 0.73525739 | 0.36570907 |
| TRIM24 | MEAAVSEAAG | MEAAVSEAAG | S539Y | -1.2338142 | -2.439014 | -2.0427604 |
| CDH11 | MKEDSCLHA | MKEDSCLHA | V382M | -2.6894054 | -2.4985368 | -1.6993548 |
| NAB2 | MALPRTLGEL | MALPRTLGEL | R437G | 0.67278898 | 0.4911623 | 0.78376603 |
| RALGDS | MVNRARAER | MVNRARAER | G42D | -0.627041 | 0.15330386 | 1.03029394 |
| ACSL6 | MRILESPHGF | MRILESPHGF | L526V | 6.4389019 | 7.44024658 | 6.77490425 |
| PPFIBP1 | MMSDASDMI | MMSDASDMI | H283Q:M322 | 4.71259165 | 4.94180059 | 3.04966712 |

| | | | | | | |
|---|---|---|---|---|---|---|
| TP53BP1 | MDPSQSQLD | MDPSQSQLD | V678M | -1.3671832 | -1.2886376 | -0.9656925 |
| MET | MKTVAAFTSG | MKTVAAFTSG | Q102S | 0.59953737 | 0.66047692 | 0.64358902 |
| LATS1 | MKRSEKPEG\ | MKRSEKPEG\ | A1031S | -0.6044521 | -0.0146375 | -0.3442307 |
| NUTM1 | MTSQPSQGP | MTSQPSQGP | T79S | 0.76285791 | 0.87899542 | 0.86501837 |
| MAP3K13 | MHMHGTMA | MHMHGTMA | L748I | -1.0006092 | -1.3522899 | -0.988107 |
| AFF3 | MGTVLLKKGI | MGTVLLKKGI | K490R | -0.8340769 | -0.8109727 | -0.8359275 |
| KCNT2 | MQFLGPAPQ | MQFLGPAPQ | M988I | 0.50110102 | 0.63197136 | 0.42372799 |
| FOXO3 | MIPEEEDDEE | MIPEEEDDEE | G38A | 0.50569296 | 0.11112833 | 0.09293389 |
| SSPN | MRKEEKKPQI | MRKEEKKPQI | K37E | -0.8056681 | -0.9834303 | -0.3329794 |
| POT1 | MGLLYYLHVS | MGLLYYLHVS | E634K | 0.09048414 | 0.68482137 | -0.1805677 |
| PTCD2 | MVRSEDRMA | MVRSEDRMA | C318A | 1.71666074 | 3.57674026 | 1.91959739 |
| GRN | MQLPGPGAR | MQLPGPGAR | L469P | 5.39461374 | 5.36171579 | 4.47234583 |
| POLQ | MTIPDDQADI | MTIPDDQADI | I186S:K2402I | 0.00584626 | 0.94970894 | 0.36428857 |
| LRP1B | MQFNLMRRI | MQFNLMRRI | M1837T:S261 | 1.76638067 | 6.11378181 | 3.66391373 |
| DKC1 | MADGEGCIA | MADGEGCIA | G209E | 7.67148066 | 5.31209993 | 6.42133045 |
| TERF2 | MAGGRGRVS | MAGGRGRVS | F360R | 1.07938719 | 2.05297279 | 1.78304338 |
| ACD | MRSAAHPLP/ | MRSAAHPLP/ | P470S | 0.23719454 | 0.12325406 | -0.0065023 |
| CYC1 | MAALSAARCI | MAALSAARCI | S27P | 0.35409975 | 0.611269 | 0.55596101 |
| LMX1B | MEKIAPTEFVI | MEKIAPTEFVI | I78M | -0.2528377 | 0.22211647 | -0.0573001 |
| AIF1 | MSSQQNLQ( | MSSQQNLQ( | G102S | 2.65261173 | 3.66956615 | 3.18789864 |
| CMAS | MDSAAAAAG, | MDSAAAAAG, | A21P | 0.20217872 | 0.32735062 | 1.86210227 |
| AGPAT2 | MEPSWLLGG | MEPSWLLGG | D236P | -1.3539286 | 0.8715651 | 0.36770844 |
| ATM | MLIDKSIFDAS | MLIDKSIFDAS | Q326E:V495I | 0.81111598 | 1.19226813 | 0.79158282 |
| KLRF1 | MEDEEGYTVI | MEDEEGYTVI | L215P | 1.40186 | 0.78647804 | 1.15778136 |
| SYNE1 | VTVNKWEQFI | VTVNKWEQFI | G677A:I1426 | 6.24769962 | 6.85185456 | 6.84190845 |
| MRC2 | MENGAGGG( | MENGAGGG( | R15K | 0.04764509 | -0.0348828 | -0.0345399 |
| MALSU1 | MLPAARGLRI | MLPAARGLRI | S17V | -0.0181606 | 0.08399296 | -0.3691235 |
| IFNGR2 | MPRPGAALLL | MPRPGAALLL | K200L:G203S | -2.1745133 | -1.9110918 | -1.5307407 |
| MINPP1 | MASCFRGPLI | MASCFRGPLI | H407N | 0.90551949 | 1.34488606 | 0.91903877 |
| APOA5 | MSHKAALLVL | MSHKAALLVL | V191A | 0.68206644 | 0.70154881 | 0.8650167 |
| HOXD12 | MCDRGLYRS | MCDRGLYRS | C211A | 3.8816545 | 3.73370647 | 3.25092316 |
| PRUNE2 | MHILLLQVSPI | MHILLLQVSPI | Q38E:L163P:/ | -0.3162969 | -0.7484227 | -0.7497063 |
| SIRT1 | MADEETLLLQ | MADEETLLLQ | V19G | 1.57954359 | 1.15374446 | 1.26233959 |
| IGF2R | MRRAARSSG | MRRAARSSG | S1849K | 0.57743168 | 1.14473462 | 1.09487212 |
| PROM1 | MSENSLCFSI | ISENSLCFSID | M1I:L92V:S1! | -3.6970868 | -1.8505657 | -4.3506374 |
| ATG4C | MEATGTDEVI | MEATGTDEVI | I245V | 3.10086918 | 2.83512592 | 2.06727314 |
| TLR4 | MNLEGLVCSI | MNLEGLVCSI | C819G | 0.54576564 | 1.75077796 | 1.11393046 |
| ADAMTSL1 | MDCPPEAGD | MDCPPEAGD | V874A | 0.66439652 | -0.0486209 | -0.3066185 |
| FGF19 | MWRSLCKPY | MWRSLCKPY | S64R | 1.59884477 | 0.74986696 | 0.57871699 |
| CGNL1 | MEPYCAGFKI | MEPYCAGFKI | D27Q:D235C | -1.6006486 | -1.3402405 | -1.3532758 |
| ITGB3 | MPGSPCSGS | MPGSPCSGS | T96A:A266T:\ | 2.56283426 | 2.77592325 | 2.66859913 |
| NCOR2 | MSGSTQPVT( | MSGSTQPVT( | S899T:L1821I | 0.87430024 | 0.22389317 | 0.45334673 |
| MSRA | MFSRWQEG( | MFSRWQEG( | S129A:G210E | 5.53171206 | 4.33156794 | 4.48105985 |
| TRIM25 | MAELVRAPSV | MAELVRAPSV | T335K | 0.36340857 | 0.09166408 | 0.33689594 |
| F7 | MVSRHYLVLL | MVSRHYLVLL | I26V:Q72K:T1 | -0.4426708 | -1.0792881 | -2.2290052 |

| | | | | | | |
|---|---|---|---|---|---|---|
| TSFM | MQRAALTGV( | MQRAALTGV( | A157L:T211I | -1.5273874 | -0.3509601 | 1.98013592 |
| SORCS1 | MGSSKEPDL\ | MGSSKEPDL\ | R485G | 5.74395227 | 5.21036339 | 4.94365978 |
| AOPEP | MTENLQSLCI | MTENLQSLCI | K75E | 0.32472539 | 0.69857669 | -0.0189571 |
| NFKB1 | MAEEDAYIM( | MAEEDAYIM( | A208T | -0.8732467 | 0.30350733 | -3.2060719 |
| ZBTB20 | MSALGPRLG\ | MSALGPRLG\ | D455N | 0.03487682 | -0.3871555 | -0.1977091 |
| FBXO31 | MAVCARLCG | MAVCARLCG | D424G:M465 | 0.91238141 | -2.5852218 | 2.01047754 |
| BSCL2 | MMSSTGPFL( | MMSSTGPFL( | Q288R | 0.35108328 | 1.33736229 | 0.78626764 |
| PPP1R1A | MLLQIISNWS | MLLQIISNWS | K17E | 0.06079674 | -0.0084026 | 0.43004417 |
| VWA5A | MMSCGLLNS | MMSCGLLNS | R81K | 0.39499331 | -0.1649547 | 0.60347319 |
| METTL1 | MAGAPPQKR | MAGAPPQKR | H108Q | 3.29008484 | 3.1547184 | 1.7941699 |
| BNIPL | MVPSIKKGRV | MVPSIKKGRV | F64L:D72E:D | -83.923219 | -176.67127 | -99.536974 |
| PIK3C2G | MKQISISTVDI | NKQISISTVDI | M1N | -4.0484757 | -4.238657 | -5.0171471 |
| SVEP1 | MASHPKEEH( | MASHPKEEH( | L261V:I392T:I | 5.341429 | 4.07388818 | 3.9503969 |
| RAPGEF3 | MHLFRSYSY( | MHLFRSYSY( | T835V | -0.7462301 | -0.6627133 | -1.6628277 |
| XRCC6 | MSEWMSYYK | MSEWMSYYK | W68Q | 6.83099699 | 5.74687576 | 6.22348404 |
| IFNAR1 | MAPPELLLLL | MAPPELLLLL | E70T | -4.1375189 | -3.083858 | -3.922647 |
| CYP19A1 | MILETLNLMH | MILETLNLMH | M158V:H461 | 0.83128881 | 3.26096296 | 0.54227424 |
| METTL21B | MAASRQPLL( | MAASRQPLL( | V215H:P219( | 3.69391203 | 2.94236946 | 3.12817359 |
| HIF1A | MELTGEKKRI | MELTGEKKRI | R584S | 2.20757484 | 2.25289726 | 1.67792141 |
| MTFMT | MRGRLLRAW | MRGRLLRAW | V16G:L307S | 3.62504005 | 3.80088544 | 3.76371813 |
| GZMB | MLILILLPVALL | MLILILLPVALL | Y93K:S150Q: | 5.47830439 | 4.72221017 | 4.19026661 |
| GHSR | MRNQSGSPN | MRNQSGSPN | S328R | 1.52705622 | 3.16540623 | 1.19575286 |
| BAG6 | MAEAQNLEVI | MAEAQNLEVI | L1075V | -0.6075642 | -0.6134269 | -0.6607907 |
| ZPLD1 | MERIWLLLLLI | MERIWLLLLL | I11T | 0.77820635 | 0.47493458 | 0.43873262 |
| SYT13 | MVLSVPVIAL( | MVLSVPVIAL( | I160A | -0.1231334 | -0.1422381 | -0.6950421 |
| POLM | MALVPLKRRF | MALVPLKRRF | A18T:Q140P: | 2.4490118 | 1.11860204 | 2.2853117 |
| AKT1S1 | MADNHRESV | MADNHRESV | T83P | 1.71512592 | 1.6513207 | 2.04517651 |
| TEP1 | MPPRAQPEPI | MPPRAQPEPI | V1966A:L199 | 2.20801282 | 1.9068532 | 2.31878841 |
| CCL5 | MKATVAALAV | MKATVAALAV | P22A | 1.30634689 | 1.99905348 | 1.26585758 |
| ENOX1 | MLWMEQFL( | MLWMEQFL( | S354C | -2.0621901 | 0.33685303 | -0.5821528 |
| RBPMS | MNSLPADKE! | MNSLPADKE! | G16N | -1.5967226 | 0.19969296 | -0.8290381 |
| MRE11A | MSCINSGDE | MSCINSGDE | S620N:G625I | 3.55497074 | 3.61527872 | 2.23423696 |
| CPA2 | MRLILFFSAL/ | MRLILFFSAL/ | E194N | 3.61307955 | 3.21519279 | 3.1621511 |
| NAV2 | MNKVEIYTDV | MNKVEIYTDV | C1122S | 5.77606583 | 5.96096611 | 4.7753787 |
| LINGO2 | MLHTATSFW( | MLHTATSFW( | V470I:I591V | 2.49247956 | 0.61630255 | 3.16465282 |
| ZFYVE28 | MRPIHPFSFC | MRPIHPFSFC | K556G | -0.5995667 | -0.8129997 | -0.6686583 |
| ECHS1 | MAVALSSLRR | MAVALSSLRR | I50V | 1.32922554 | 2.0333271 | 2.07256031 |
| GZMH-aging- | MIQGNWSLA | MIQGNWSLA | F78L:I80L:P8: | 5.06105542 | 7.28457415 | 5.33248103 |
| PPM1D | MEGLYSLGVS | MEGLYSLGVS | I437V | 0.4063158 | 0.37084484 | 0.32418251 |
| POLE2 | MAAEKLRSKV | MAAEKLRSKV | V107L | -0.2308545 | 3.64018297 | -2.1270604 |
| TOP2A | MELDESLSPL | MELDESLSPL | S685P | -0.2049539 | -0.1618481 | 0.01315951 |
| PPP1R26 | MFLMNASPLI | MFLMNASPLI | G499D | -0.011971 | 0.0057044 | -0.043119 |
| ERCC1 | MEPMEAAQV | MEPMEAAQV | S93L | -1.2213206 | -1.361328 | -0.9138765 |
| CSMD3 | MCMGEINLE | MCMGEINLE | A96M:M1405 | 0.98679447 | 1.56532335 | 0.45902157 |
| CRLS1 | MHRRGAEEP | MHRRGAEEP | L234V | -0.2832611 | -0.5489273 | -0.4013157 |

| | | | | | | |
|---|---|---|---|---|---|---|
| APOA4 | MAELRGPCG | MAELRGPCG | N355A | 2.14819384 | 4.52700996 | 4.18840694 |
| APOB | MGPPQLWQI | MGPPQLWQI | Q2004R:S42[illegible] | -1.1084192 | -0.8487194 | -0.8858812 |
| HDAC6 | MKRGGRRGI | MKRGGRRGI | R16N | -2.4701958 | -1.8366849 | -1.7577894 |
| WDR27 | MESPVDLCF | MESPVDLCF | K630E | 5.82145643 | 4.06643772 | 2.98211312 |
| HIP1 | MEVGKMLML | MEVGKMLML | G319D | -0.9992856 | 3.14573526 | -0.383432 |
| CEL | MEPRILKWD | MEPRILKWD | M483L | 1.31943154 | 1.9277668 | 1.47370672 |
| TERT | MAGSGPFGA | MAGSGPFGA | S19R:Y197L | 2.2852447 | 1.73315907 | 1.46700239 |
| NCAM2 | MYAKCIHAEI | MYAKCIHAEI | I135V:E279R | 1.15104437 | -0.3378019 | -2.0661223 |
| LEP | MRCPSLPLC | MRCPSLPLC | V148L | 0.92389941 | 1.11767018 | 0.62519193 |
| PARVG | MEDAVVTEEL | MEDAVVTEEL | P303L:Q306N | 1.45026767 | 2.20558643 | 0.67182374 |
| XDH | MASERTTDQI | MASERTTDQI | M533T | -0.2659404 | 0.43989015 | 1.69011045 |
| APOM | MLQRSWSYL | MLQRSWSYL | I70V | 0.66271281 | 0.61156821 | 0.76497793 |
| CHIT1 | MGRVFVWIG | MGRVFVWIG | K198D | -0.1831474 | 1.62413776 | 0.58805561 |
| LMNB1 | MAASTPVGQI | MAASTPVGQI | M94T | 6.00239754 | 3.73457432 | 3.41635561 |
| TPP2 | MAAAAEEPFF | MAAAAEEPFF | L635A | 0.43267751 | 1.39587545 | 0.98319602 |
| POLE | MVLRNSGRR | MVLRNSGRR | M145V:H123 | 4.37971735 | 4.29431748 | 4.26300931 |
| GSR | MAAAFDYLVL | MAAAFDYLVL | L60S | 3.70539188 | -0.7754233 | 2.32327461 |
| HLA-DRB1 | MGAGWILGA | MGAGWILLA | G8L:S11G:C1 | 32.2684625 | 30.8595433 | 37.5283316 |
| HRH1 | MCETKNSTE | MCETKNSTE | I27V | 1.1810236 | 0.48670578 | 0.72192574 |
| SH2D4A | MLQQILADM | MLQQILADM | Q337R | 4.30207539 | 5.39921284 | 4.2788744 |
| NEDD4 | MARRLRLHF | MARRLRLHF | Q259K:T346K | -1.6210885 | -2.2493391 | -0.2247825 |
| MARCO | MENPKNYNE | MENPKNYNE | M46L | 1.02562344 | 1.53824306 | 1.68867278 |
| MAPK1 | MAAVSGAGA | MAAVSGAGA | D366Y | 3.17595744 | 6.38219881 | 0.33981514 |
| CFH | MLPTDCETPF | MLPTDCETPF | D250N:D252 | -3.6920928 | -1.0554645 | -1.9659671 |
| GCNT3 | MFRERKYLC | MFRERKRLC | Y7R | 1.46132469 | 2.01117897 | 2.23668242 |
| DHX33 | MPQGGEVPF | MPQEGEVPP | G4E | -0.1855741 | 0.12079787 | 0.32301092 |
| NFATC1 | MPSPGFPTSK | MPSPGFPTSK | A935V | 0.15565729 | 1.51301563 | 0.30546808 |
| PLEKHA1 | MPYVDRQNF | MPYVDRQNF | S296P | 6.06757593 | 7.02827168 | 7.75427723 |
| PUM2 | MSIPCVLLGL | MSIPCVLLGL | S372N | -0.7212336 | -0.9354973 | -0.7022581 |
| TREML1 | MGSFPAWLL | MGSFLAWLL | P5L:E15G:V5 | 3.68801045 | 2.69242656 | 4.89803672 |
| TNFSF8 | MCFSPEKTLF | MCFSPEKTLF | H69P | 1.00411868 | 0.5290091 | 0.84265757 |
| PKN2 | MASNTAEREI | MASNTAEREI | G566E | 0.50200486 | 0.72196937 | 1.06387711 |
| CD302 | MEPPFPALLC | MEPPFPALLC | V113M | -0.2903788 | -1.9373533 | -0.8781624 |
| IRG1 | MLWAGEVLL | MLWAGEVLL | S164K | 0.79431105 | 3.26107049 | 1.65841866 |
| MID2 | METLESELTC | METLESELTC | L39V | 5.71327591 | 6.80236244 | 5.00126171 |
| DHX9 | MGDVKNFLY | MGDVKNFLY | A108T | -1.2868056 | -1.2064719 | -0.9130659 |
| C5AR1 | MSVNEVSED | MSVNEVSED | N184F | -0.1775277 | 1.1302582 | 0.61318445 |
| IL1RL2 | MGEECMVHI | MGEECMVHI | S89G | -1.7312374 | -2.5031216 | -0.3617582 |
| ATF6 | MAAEEVEQG | MAAEEVEQG | S391R:K577T | 2.73495007 | 4.53951263 | -0.7152141 |
| ST6GAL1 | MVHINLLKKL | MVHINLLKKL | G160D | 1.40235436 | 4.36126232 | 2.73347759 |
| LY75 | MRRAPGRSL | MRRAPGRSL | S150P:N486K | 1.42159939 | 1.99762607 | 2.43648481 |
| RABGEF1 | EEMSLKSERR | EEMSLKSERR | L14V | 4.41937876 | 4.18103409 | 5.09858608 |
| CD1D | MGNTKVRPK | MGNTKVRPK | T122G:Q312I | 1.45676541 | 0.31676769 | 1.03117609 |
| ADAR | MNTGFGRGK | MNTGFGRGK | D397G | 0.18991971 | -0.4823489 | -0.8571573 |
| ATP7A | MAQGLVTVG | MAQGLVTVG | G576R:G891 | 1.32816696 | 1.13956368 | 0.71163845 |

| | | | | | | |
|---|---|---|---|---|---|---|
| PSMD5 | MAEEALELLAI | MAEEALELLAI | A266S | 0.49820471 | 1.81840479 | 0.15744925 |
| TRIM8 | MAENWKNCI | MAENWKNCI | L225V | 1.21690702 | 0.55759239 | 0.38380241 |
| PIK3AP1 | MASAGAPGG | MASAGAPGG | I705R:M708L | 1.53287864 | 2.05521965 | 1.40935755 |
| APOBEC2 | MAENQEKPR | MAENQEKPR | S21L | -0.2635124 | -1.0212715 | -0.5183415 |
| DAPK1 | MTVFRQENLI | MTVFRQENLI | S1355P | -0.3962524 | -0.4061823 | -0.3301878 |
| STK11 | MDVQEPQEL | MDVQEPQEL | R108K | 5.1613369 | 2.12538147 | 1.22533369 |
| NFKBIZ | MIVESNREAP | MIVESNREAP | H160C:V334I | 1.37428689 | 0.84892178 | -0.2756455 |
| TRAT1 | MTIVLLSQILF | MTIVLLSQILF | D115K | 0.3447473 | 0.28196263 | -0.2751777 |
| IL17C | MSLLERLLGY | MSLLERLLGY | V87L | 1.02914476 | 0.73247075 | 1.34619904 |
| CD38 | MPLQNRSSKT | MPLQNRSSKT | M40R | 2.606318 | 1.68525124 | 1.90427971 |
| FGA | MEHGIQSDC | MEHGIQSDC | E251P | -0.4752822 | -0.4333024 | -0.6913538 |
| IFNL3 | MKKCHLSKYI | MKKCHLSKYI | W15Q:K18Q: | 1.07650542 | 2.62408614 | 1.74338961 |
| DHX58 | MLTPALPPSY\ | MLTPALPPSY\ | S419T | -1.3538461 | -1.7994478 | -1.2873821 |
| MADCAM1 | MEPTLLLLLLS | MEPTLLLLLLS | S254E:S353R | -2.6759295 | -4.021515 | -2.4434109 |
| 8-Mar | MPLHQISVIP/ | MPLHQISVIP/ | I534T | 0.81426311 | -1.2908425 | -0.506623 |
| CD207 | MSRENICDN | MMRENICDN | S2M:D16A:E3 | -11.170667 | -1.5653664 | -6.0648718 |
| MAP4K2 | MSLLRDVSLL | MSLLRDVSLL | S352N:I749V | -1.1287212 | -3.0765743 | 0.13911533 |
| JAG1 | MLLPRRPALA | MLLPRRPALA | T1095G | -0.7812741 | -0.8477898 | -0.8791797 |
| ATG9A | MMHFDTQY( | MAHFDTQY( | M2A:L360P:V | 10.1685184 | 13.4192295 | 8.23924756 |
| VAV3 | MEPWKQCA( | MEPWKQCA( | Y617Q | 3.84457469 | 1.88881922 | 3.21429825 |
| TNIP3 | MIHCGLLKKK | MIHCGLLKKK | T88G | -0.4169135 | -0.1749413 | -0.3850374 |
| TMEM173 | MSHETERPS\ | MSHETERPS\ | D256E | 0.51193404 | 0.57930255 | 0.40249872 |
| LAMP1 | MARVGGGR( | MARVGGGR( | E35T | 1.33896351 | 0.94491005 | -0.0848644 |
| ITGAL | MQESGCACE | MQESGCACE | L1193M | -0.4909463 | -0.7855978 | -0.4414308 |
| PROS1 | MRLLSRRCV/ | MRLLGRRCV | S5G:N398S | 3.90223956 | 3.6218152 | 2.78750968 |
| ATG16L1 | MTSGLRAATF | MTSGLRAATF | E582G | 5.57267904 | 8.17109776 | 10.9950008 |
| GCH1 | MEKGKLGALI | MEKGKLGALI | Q33P | 1.97572136 | 1.25694728 | 1.81018019 |
| CTSG | MLILILLPAAFL | MLILILLPAAFL | L120R:S126N | 4.62907827 | 3.52711034 | 2.32414746 |
| COLEC12 | MRMGVPSSF | MRMGVPSSF | L484P:A666E | 4.03876185 | 5.95143795 | 4.73760974 |
| CD97 | MARLCIALCL | MARLCIALCL | T54F:M63T:R | 4.05379772 | 5.43879449 | 3.58161569 |
| MASP2 | MYGRITSPDF | MYGRITSPDF | A87S:E160S | 3.72239029 | 6.28090942 | 5.17090821 |
| SEMA4A | MPSRALCLLC | MPSRALCLLC | T410A:R771C | 3.62308526 | 3.61158085 | 2.88524914 |
| LPXN | MDDLDALLAI | MDDLDALLAI | T386P | 0.25087655 | 0.38323343 | -0.1692641 |
| THEMIS | MFGNECCLF | MFGNECCLF | N555D | 0.36163425 | -0.2255657 | 0.22965646 |
| ANG | MHRAVTPATH | MHRAVTPATH | M39L | 2.65868711 | 2.71883988 | 2.32403111 |
| APEX2 | MRLLSWNVA | MRLLSWNVA | C118R:T119/ | 14.4792295 | 12.8869758 | 12.2183743 |
| OASL | MELYQVPAGS | MELYQVPAGS | V108L | -0.2150879 | 0.69544315 | 0.6138072 |
| EXOSC3 | MRLRTDVTHS | MRLRTDVTHS | Y73Q | 3.19033742 | 2.92748713 | 2.32738423 |
| CFP | MEGALTSSLL | MEGALTSSLL | E38K | 0.2001338 | 0.37790585 | -0.6085558 |
| POU2F2 | MSVQGQRK\ | MSVQGQRK\ | A555G | -2.3508937 | 1.53598356 | -2.0859435 |
| ART1 | MSTRPQSEE( | MSTRPQSEE( | L157P:N330S | -0.9932518 | 0.29973483 | 0.24010813 |
| IL18RAP | MFTFCWILLL | MFTFCWILLL | Y55H:R66Q:E | -0.4275997 | -2.5906199 | -0.5824924 |
| ATF6B | MAAVAPELLV | MAAVAPELLV | A340L | 2.02425623 | 1.04490089 | 0.93241179 |
| CD5L | MEALSGKVFS | MEALSGKVFS | K117E | 2.69640589 | 2.03981209 | 1.66126299 |
| EGF | MFLILTAFLPV | MFLILTAFLPV | S1136R | -1.3668042 | -0.9719696 | -2.0240247 |

| | | | | | | |
|---|---|---|---|---|---|---|
| ITCH | MLRGGIIQGI | MLRGGIIQGI | I37V:R126N | 0.87561345 | 1.11585379 | 0.73407978 |
| IL1RL1 | MLIHALLVQLI | MLIHALLVQLI | T356I:K413T:I | 4.42193627 | 8.44315839 | 4.60334253 |
| TAPBP | MISPAPVGAL | MISPAPVGAL | K70R | 1.65529466 | 1.2654593 | 1.8603065 |
| C5 | MNTLSFICILA | MNTLSFICILA | G662D | -0.0246835 | -0.1530213 | 0.03025103 |
| CYP27B1 | MPQTLKLASR | MPQTLKLASR | L394V | 2.38677406 | 3.22503281 | 1.94951499 |
| GZMH-immu | MIQGNWSLA | MIQGNWSLA | L74P:F78L:I8 | 3.99087322 | 6.16688478 | 3.77897525 |
| ERBB2IP | MTTKRNLFVF | MTTKRNLFVF | Q720D | 0.03728509 | -0.0037243 | 0.01017094 |
| MSH6 | MSRQSTLLRF | MSRQSTLLRF | A611S:G106 | 3.48682833 | 2.81058323 | 1.06503105 |
| ABL1 | MSRNEINEKI | MSRNEINEKI | M740V | 1.00349784 | 1.19354916 | 1.45922852 |
| SH2B2 | MNGDALCQI | MNGDALCQI | N644S | 2.41130543 | 1.76431859 | 2.16292143 |
| BMP6 | MTVTILRFTVF | MTVTILRFTVF | G206D:E208I | 1.54107213 | 1.92908812 | 1.37442994 |
| PLCL2 | MAEFGSRGA | MAEFGSRGA | G77S | -0.3147409 | -0.3529074 | -0.2606333 |
| SPPL2A | MSRMRRAW | MSRMRRAW | S62T | -0.6531155 | -0.7447788 | -0.2785742 |
| HAVCR2 | MAPYFILQWI | MAPYFILQWI | Q179P | 1.06360054 | 0.9748683 | 0.74705791 |
| ATF7 | MQNDLRNH | MQNDLRNH | A13M | -0.1605392 | -1.216588 | -0.5287919 |
| MR1 | MGLALRLLLLI | MGMAFLLPLI | L3M:L5F:R6L | 66.321845 | 56.6424472 | 47.0848903 |
| APBB1IP | MEQTCDDID | MEQTCDDID | I318T | 2.2769928 | 1.72790313 | 0.63914609 |
| PRKCG | MALVASPTELI | MALVASPTELI | N427T:L591P | -2.7610428 | -3.9934196 | -3.2814107 |
| AQP9 | MNGQSKRTF | MNGQSKRTF | A90V | 0.99430609 | 2.84076643 | 2.0250752 |
| TINAGL1 | MRLLWTLPTL | MRLLWTLPTL | W465H | -0.1343489 | 2.17521429 | 2.22404909 |
| CD200 | MRSYEILQVT | MRSYEILQVT | R79S | 1.70089746 | 1.34860849 | 0.96678925 |
| LY9 | MELSRIVGDS | MELSRIVGDS | Q20T:F22I:I3 | -1.6545054 | -0.954507 | -1.3581212 |
| SEMA4D | MLLSNDDKD | MLLSNDDKD | H551G | -1.0102859 | -4.3430557 | -2.196969 |
| ITGB7 | MTGAAWPLG | MTGAAWPLG | R116Q:A143I | 2.0748136 | 2.60858357 | 2.34339392 |
| IL31RA | MRNAYISPES | MRNAYISPES | Y675N | 3.22263241 | 4.64729786 | 1.90500259 |
| CD22 | MVLIQIASVIL | MVLIQIASVIL | I264S:G281H | -0.5879977 | -0.4109385 | 3.24501264 |

**E** Protein language models evaluation with the human amino ac

| Ref | wt_sequence | mutated_seq | mutant | Tranception_ | Tranception_ | Tranception_ |
|---|---|---|---|---|---|---|
| KIAA1549 | MYYCNKTFVI | MYYCNKTFVI | N105D | -2.7299855 | -2.7298975 | 8.80E-05 |
| KIAA1549 | MYYCNKTFVI | MYYCNKTFVI | S147P | -2.7299855 | -2.7299738 | 1.17E-05 |
| ARID1B | MIMAAQVAA | MIMAAQVAA | A894T | -1.992675 | -1.9926567 | 1.83E-05 |
| GOPC | MSGMGAGS | MSGMGAGS | K112Q | -0.6377313 | -0.6169946 | 0.02073675 |
| ROS1 | MEGELGINA | MEGELGINA | C1274Y | -2.5050723 | -2.5043691 | 0.00070318 |
| ROS1 | MEGELGINA | MEGELGINA | N1331Q | -2.5050723 | -2.5054445 | -0.0003723 |
| ROS1 | MEGELGINA | MEGELGINA | A1450Y | -2.5050723 | -2.5049896 | 8.27E-05 |
| ROS1 | MEGELGINA | MEGELGINA | T1580L | -2.5050723 | -2.5052169 | -0.0001446 |
| ROS1 | MEGELGINA | MEGELGINA | R2072K | -2.5050723 | -2.5032352 | 0.00183709 |
| RB1 | MPPKPPRKG | MPPKPPRKG | V54H | -2.5810075 | -2.5824957 | -0.0014882 |
| RB1 | MPPKPPRKG | MPPKPPRKG | A245P | -2.5810075 | -2.5811474 | -0.00014 |
| CBLB | MANSLNGRN | MANSLNGRN | P590L | -1.1808499 | -1.1814407 | -0.0005908 |
| CBLB | MANSLNGRN | MANSLNGRN | S749G | -1.1808499 | -1.181426 | -0.0005761 |
| BCL9 | MHSSNPKVR | MHSSNPKVR | V1073M | -2.3700395 | -2.369606 | 0.00043344 |
| FLT3 | MTVSVFLFGF | MTVSVFLFGF | A19I | -2.1745448 | -2.1744068 | 0.00013804 |
| FLT3 | MTVSVFLFGF | MTVSVFLFGF | I54V | -2.1745448 | -2.1742563 | 0.00028849 |
| FLT3 | MTVSVFLFGF | MTVSVFLFGF | N787D | -2.1745448 | -2.1731155 | 0.00142932 |

| | | | | | | |
|---|---|---|---|---|---|---|
| APC2 | MIDYSSARSY | MIDYSSARSY | M195V | -2.6313783 | -2.631179 | 0.00019932 |
| APC2 | MIDYSSARSY | MIDYSSARSY | Q1698A | -2.6313783 | -2.6313233 | 5.50E-05 |
| CASC5 | MDRIYSESNE | MDRIYSESNE | D408H | -2.7402045 | -2.7395039 | 0.00070063 |
| CASC5 | MDRIYSESNE | MDRIYSESNE | C2403N | -2.7402045 | -2.7401435 | 6.10E-05 |
| NRG1 | MSEVSAETFP | MSEVSAETFP | H42P | -0.9756854 | -0.9685033 | 0.00718206 |
| CEP89 | MALSFRRGR | MALSFRRGR | H698Q | -2.5613756 | -2.5583558 | 0.00301981 |
| NDRG1 | MLLKFIFISKE | MLLKFIFISKE | D75E | -1.5244766 | -1.5180894 | 0.00638723 |
| ALK | MISHGSSCY | MGSHGSSCY | I2G | -2.5548282 | -2.5533412 | 0.00148702 |
| ALK | MISHGSSCY | MIAHGSSCY | S3A | -2.5548282 | -2.5556324 | -0.0008042 |
| ALK | MISHGSSCY | MISIGSSCYV | H4I | -2.5548282 | -2.5534868 | 0.00134134 |
| ALK | MISHGSSCY | MISHGSSCY | T37L | -2.5548282 | -2.5542002 | 0.00062799 |
| ALK | MISHGSSCY | MISHGSSCY | R38K | -2.5548282 | -2.5549009 | -7.27E-05 |
| ALK | MISHGSSCY | MISHGSSCY | M75L | -2.5548282 | -2.5527914 | 0.00203681 |
| NTRK3 | MLAASERKAR | MLAASERKAR | Y639F | -0.903671 | -0.9034594 | 0.0002116 |
| CIC | MKPARKAASS | MKPARKAASS | A1146S | -2.5755011 | -2.5756604 | -0.0001593 |
| CIC | MKPARKAASS | MKPARKAASS | P1995L | -2.5755011 | -2.5763334 | -0.0008322 |
| DAXX | MAATVESNII\ | MAATVESNII\ | L156S | -2.6502688 | -2.6486676 | 0.00160122 |
| LATS2 | MRPKTFPATT | MRPKTFPATT | S350A | -2.3569865 | -2.357574 | -0.0005875 |
| NOTCH1 | MQQLLVQSLI | MQQLLVQSLI | T2395N | -1.8964386 | -1.8955112 | 0.00092745 |
| BRD4 | MSVESGPGTI | MSVESGPGTI | I956V | -1.9955032 | -1.9953871 | 0.00011611 |
| RNF213 | MKCPKCDYIS | MKCPKCDYIS | M1803V | -2.8205913 | -2.8203888 | 0.0002025 |
| RNF213 | MKCPKCDYIS | MKCPKCDYIS | N3526V | -2.8205913 | -2.8204168 | 0.00017452 |
| THRAP3 | MSKTNKSKSC | MSKTNKSKSC | L706H | -1.9162755 | -1.9094197 | 0.00685585 |
| SETD2 | MGDFYDPEH | MGDFYDPEH | I1400A | -2.5675462 | -2.5672766 | 0.00026957 |
| SETD2 | MGDFYDPEH | MGDFYDPEH | E1436D | -2.5675462 | -2.567702 | -0.0001558 |
| CLIP4 | MTIEDLPEPS( | MTIEDLPEPS( | A457P | -1.1764956 | -1.1660848 | 0.01041079 |
| PTPRB | MELGGAGDH | MELGGAGDH | V50M | -2.2373569 | -2.2383902 | -0.0010333 |
| PTPRB | MELGGAGDH | MELGGAGDH | H178V | -2.2373569 | -2.2361059 | 0.00125098 |
| PTPRB | MELGGAGDH | MELGGAGDH | A213E | -2.2373569 | -2.2380185 | -0.0006616 |
| PTPRB | MELGGAGDH | MELGGAGDH | T228E | -2.2373569 | -2.2379007 | -0.0005438 |
| PTPRB | MELGGAGDH | MELGGAGDH | N318M | -2.2373569 | -2.2377286 | -0.0003717 |
| PTPRB | MELGGAGDH | MELGGAGDH | P365A | -2.2373569 | -2.2364545 | 0.00090241 |
| PTPRB | MELGGAGDH | MELGGAGDH | R369V | -2.2373569 | -2.2403765 | -0.0030196 |
| PTPRB | MELGGAGDH | MELGGAGDH | V491H | -2.2373569 | -2.2380953 | -0.0007384 |
| PTPRB | MELGGAGDH | MELGGAGDH | T514L | -2.2373569 | -2.2382212 | -0.0008643 |
| PTPRB | MELGGAGDH | MELGGAGDH | A576L | -2.2373569 | -2.2387788 | -0.0014219 |
| PTPRB | MELGGAGDH | MELGGAGDH | Q588R | -2.2373569 | -2.2366614 | 0.00069547 |
| PTPRB | MELGGAGDH | MELGGAGDH | W668Y | -2.2373569 | -2.2373424 | 1.45E-05 |
| PTPRB | MELGGAGDH | MELGGAGDH | G1192Q | -2.2373569 | -2.2365232 | 0.00083375 |
| KMT2D | MPLVRYRKV\ | MPLVRYRKV\ | L827P | -2.4635997 | -2.4636701 | -7.04E-05 |
| NSD1 | MEPACEVSRI | MEPACEVSRI | T406A | -2.0606111 | -2.0606896 | -7.85E-05 |
| NSD1 | MEPACEVSRI | MEPACEVSRI | F2368L | -2.0606111 | -2.057244 | 0.00336711 |
| IDH1 | MSSGSLRGYI | MSSGSLRGYI | S267G | -0.7480305 | -0.7366845 | 0.01134598 |
| BCL11B | MNFPLGDILV | MNFPLGDILV | R65L | -0.8569175 | -0.8458215 | 0.011096 |
| KMT2A | MAHSGRWRI | MAHSGRWRI | R513S | -2.5808938 | -2.5808878 | 5.96E-06 |

| | | | | | | |
|---|---|---|---|---|---|---|
| ATRX | MRRGRKAAG | MRRGRKAAG | Q1136R | -2.2713734 | -2.2710155 | 0.00035795 |
| NCOA4 | MSPPQDRSG | MSPPQDRSG | R499S | -2.7393444 | -2.7361264 | 0.00321794 |
| ERBB2 | MASIVSAPLQ: | MASIVSAPLQ: | M162T | -2.1802125 | -2.1770101 | 0.00320244 |
| BRCA1 | MESSVAAIGD | MESSVAAIGD | R1105M | -2.6460161 | -2.6468158 | -0.0007997 |
| CNOT3 | MADKRKLQG | MADKRKLQG | I418A | -0.9969992 | -0.995566 | 0.00143325 |
| STRN | MDEQAGPG\ | MDEQAGPG\ | H236D | -0.7292972 | -0.7089126 | 0.02038461 |
| TET1 | MAHHARPSR | MAHHARPSR | H1331L | -2.7176743 | -2.7173061 | 0.00036812 |
| TRIM33 | MAENKGGG( | MAENKGGG( | E808G | -1.9176302 | -1.9162133 | 0.00141692 |
| SETBP1 | MSSPVGPTHI | MSSPVGPTHI | I393L | -2.7061698 | -2.7057633 | 0.0004065 |
| TRIM24 | MEAAVSEAAG | MEAAVSEAAG | S539Y | -2.1673803 | -2.1671143 | 0.00026608 |
| CDH11 | MKEDSCLHA | MKEDSCLHA | V382M | -0.6147143 | -0.6199552 | -0.0052409 |
| NAB2 | MALPRTLGEL | MALPRTLGEL | R437G | -2.0033855 | -2.0024333 | 0.00095224 |
| RALGDS | MVNRARAER | MVNRARAER | G42D | -0.611027 | -0.6054659 | 0.00556105 |
| ACSL6 | MRILESPHGF | MRILESPHGF | L526V | -0.6848584 | -0.674082 | 0.01077646 |
| PPFIBP1 | MMSDASDM | MMSDASDM | H283Q | -1.4728371 | -1.4681673 | 0.00466979 |
| PPFIBP1 | MMSDASDM | MMSDASDM | M322V | -1.4728371 | -1.4703543 | 0.00248277 |
| TP53BP1 | MDPSQSQLD | MDPSQSQLD | V678M | -2.7330303 | -2.7337489 | -0.0007186 |
| MET | MKTVAAFTSG | MKTVAAFTSG | Q102S | -2.0276046 | -2.0271749 | 0.00042963 |
| LATS1 | MKRSEKPEG' | MKRSEKPEG' | A1031S | -2.3254704 | -2.32354 | 0.00193048 |
| NUTM1 | MTSQPSQGP | MTSQPSQGP | T79S | -2.717562 | -2.7166564 | 0.00090551 |
| MAP3K13 | MHMHGTMA | MHMHGTMA | L748I | -1.9063194 | -1.9072564 | -0.000937 |
| AFF3 | MGTVLLKKGI | MGTVLLKKGI | K490R | -2.161474 | -2.162286 | -0.0008121 |
| KCNT2 | MQFLGPAPQ | MQFLGPAPQ | M988I | -1.6773593 | -1.6790117 | -0.0016524 |
| FOXO3 | MIPEEEDDEE | MIPEEEDDEE | G38A | -1.7054051 | -1.7048317 | 0.0005734 |
| SSPN | MRKEEKKPQI | MRKEEKKPQI | K37E | -1.4402336 | -1.447982 | -0.0077484 |
| POT1 | MGLLYYLHVS | MGLLYYLHVS | E634K | -2.882823 | -2.882654 | 0.00016904 |
| PTCD2 | MVRSEDRMA | MVRSEDRMA | C318A | -2.4981999 | -2.4928892 | 0.00531077 |
| GRN | MQLPGPGAR | MQLPGPGAR | L469P | -1.6527517 | -1.6466577 | 0.00609398 |
| POLQ | MTIPDDQADI | MTIPDDQADI | I186S | -2.6163565 | -2.615996 | 0.00036049 |
| POLQ | MTIPDDQADI | MTIPDDQADI | K2402D | -2.6163565 | -2.6155167 | 0.00083987 |
| LRP1B | MQFNLMRRK | MQFNLMRRK | M1837T | -2.0637005 | -2.0633789 | 0.00032158 |
| LRP1B | MQFNLMRRK | MQFNLMRRK | S2616P | -2.0637005 | -2.063537 | 0.00016346 |
| LRP1B | MQFNLMRRK | MQFNLMRRK | Q2782E | -2.0637005 | -2.0631254 | 0.00057507 |
| DKC1 | MADGEGCIA | MADGEGCIA | G209E | -1.0348753 | -1.0205598 | 0.01431549 |
| TERF2 | MAGGRGRVS | MAGGRGRVS | F360R | -2.2467225 | -2.245055 | 0.0016675 |
| ACD | MRSAAHPLPA | MRSAAHPLPA | P470S | -2.8269811 | -2.8266716 | 0.00030947 |
| CYC1 | MAALSAARCL | MAALSAARCL | S27P | -1.1471834 | -1.1446936 | 0.00248981 |
| LMX1B | MEKIAPTEFVI | MEKIAPTEFVI | I78M | -0.4367947 | -0.4401834 | -0.0033887 |
| AIF1 | MSSQQNLQ( | MSSQQNLQ( | G102S | -1.3671319 | -1.3089516 | 0.05818033 |
| CMAS | MDSAAAAAG. | MDSAAAAAG. | A21P | -1.9248818 | -1.921643 | 0.0032388 |
| AGPAT2 | MEPSWLLGG | MEPSWLLGG | D236P | -1.3724651 | -1.3827729 | -0.0103078 |
| ATM | MLIDKSIFDA: | MLIDKSIFDA: | Q326E | -2.8614299 | -2.8612137 | 0.00021617 |
| ATM | MLIDKSIFDA: | MLIDKSIFDA: | V495K | -2.8614299 | -2.8611978 | 0.00023206 |
| ATM | MLIDKSIFDA: | MLIDKSIFDA: | K496D | -2.8614299 | -2.8613758 | 5.40E-05 |
| ATM | MLIDKSIFDA: | MLIDKSIFDA: | S576V | -2.8614299 | -2.8614388 | -8.90E-06 |

| | | | | | | |
|---|---|---|---|---|---|---|
| ATM | MLIDKSIFDAS | MLIDKSIFDAS | R1508K | -2.8614299 | -2.8613555 | 7.44E-05 |
| KLRF1 | MEDEEGYTVI | MEDEEGYTVI | L215P | -2.3421099 | -2.3321896 | 0.00992036 |
| SYNE1 | VTVNKWEQFI | VTVNKWEQFI | G677A | -2.5792303 | -2.5791345 | 9.58E-05 |
| SYNE1 | VTVNKWEQFI | VTVNKWEQFI | I1426P | -2.5792303 | -2.5790539 | 0.00017643 |
| SYNE1 | VTVNKWEQFI | VTVNKWEQFI | G1978S | -2.5792303 | -2.5790267 | 0.00020361 |
| SYNE1 | VTVNKWEQFI | VTVNKWEQFI | K2572S | -2.5792303 | -2.579236 | -5.72E-06 |
| SYNE1 | VTVNKWEQFI | VTVNKWEQFI | K3614E | -2.5792303 | -2.5789824 | 0.00024796 |
| SYNE1 | VTVNKWEQFI | VTVNKWEQFI | V4361A | -2.5792303 | -2.5790248 | 0.00020552 |
| SYNE1 | VTVNKWEQFI | VTVNKWEQFI | F4501L | -2.5792303 | -2.5790186 | 0.00021172 |
| SYNE1 | VTVNKWEQFI | VTVNKWEQFI | H4935A | -2.5792303 | -2.5790963 | 0.00013399 |
| MRC2 | MENGAGGG | MENGAGGG | R15K | -2.2213821 | -2.2219946 | -0.0006125 |
| MALSU1 | MLPAARGLRI | MLPAARGLRI | S17V | -1.4890624 | -1.4878311 | 0.00123131 |
| IFNGR2 | MPRPGAALLL | MPRPGAALLL | K200L | -2.5423267 | -2.5404072 | 0.00191951 |
| IFNGR2 | MPRPGAALLL | MPRPGAALLL | G203S | -2.5423267 | -2.5425849 | -0.0002582 |
| IFNGR2 | MPRPGAALLL | MPRPGAALLL | Q269L | -2.5423267 | -2.542973 | -0.0006464 |
| IFNGR2 | MPRPGAALLL | MPRPGAALLL | G297D | -2.5423267 | -2.5396566 | 0.00267005 |
| MINPP1 | MASCFRGPL | MASCFRGPL | H407N | -2.6936755 | -2.6920896 | 0.00158596 |
| APOA5 | MSHKAALLVL | MSHKAALLVL | V191A | -2.6374648 | -2.6344967 | 0.00296807 |
| HOXD12 | MCDRGLYRS | MCDRGLYRS | C211A | -1.7395846 | -1.6987096 | 0.04087496 |
| PRUNE2 | MHILLLQVSP | MHILLLQVSP | Q38E | -2.6424456 | -2.6424287 | 1.68E-05 |
| PRUNE2 | MHILLLQVSP | MHILLLQVSP | L163P | -2.6424456 | -2.6427838 | -0.0003382 |
| PRUNE2 | MHILLLQVSP | MHILLLQVSP | A1554P | -2.6424456 | -2.6419659 | 0.0004797 |
| SIRT1 | MADEETLLLQ | MADEETLLLQ | V19G | -1.5442294 | -1.5424575 | 0.00177193 |
| IGF2R | MRRAARSSG | MRRAARSSG | S1849K | -2.7298619 | -2.72953 | 0.00033188 |
| PROM1 | MSENSLCFSI | ISENSLCFSID | M1I | -2.701205 | -2.7260847 | -0.0248797 |
| PROM1 | MSENSLCFSI | MSENSLCFSI | L92V | -2.701205 | -2.7002401 | 0.00096488 |
| PROM1 | MSENSLCFSI | MSENSLCFSI | S155V | -2.701205 | -2.6902752 | 0.01092982 |
| ATG4C | MEATGTDEVE | MEATGTDEVE | I245V | -1.4267678 | -1.4202048 | 0.00656307 |
| TLR4 | MNLEGLVCS | MNLEGLVCS | C819G | -2.540802 | -2.5398715 | 0.00093055 |
| ADAMTSL1 | MDCPPEAGD | MDCPPEAGD | V874A | -1.8842547 | -1.8836092 | 0.00064552 |
| FGF19 | MWRSLCKPY | MWRSLCKPY | S64R | -1.6514149 | -1.6690829 | -0.017668 |
| CGNL1 | MEPYCAGFK | MEPYCAGFK | D27Q | -2.706332 | -2.7065318 | -0.0001998 |
| CGNL1 | MEPYCAGFK | MEPYCAGFK | D235C | -2.706332 | -2.7081242 | -0.0017922 |
| ITGB3 | MPGSPCSGS | MPGSPCSGS | T96A | -0.9645742 | -0.9580963 | 0.00647789 |
| ITGB3 | MPGSPCSGS | MPGSPCSGS | A266T | -0.9645742 | -0.9637419 | 0.00083226 |
| ITGB3 | MPGSPCSGS | MPGSPCSGS | V639M | -0.9645742 | -0.9590158 | 0.00555837 |
| NCOR2 | MSGSTQPVT( | MSGSTQPVT( | S899T | -2.0702224 | -2.0703882 | -0.0001658 |
| NCOR2 | MSGSTQPVT( | MSGSTQPVT( | L1821H | -2.0702224 | -2.0684853 | 0.00173712 |
| NCOR2 | MSGSTQPVT( | MSGSTQPVT( | I2205A | -2.0702224 | -2.0703874 | -0.000165 |
| MSRA | MFSRWQEG( | MFSRWQEG( | S129A | -1.6826535 | -1.6763567 | 0.00629687 |
| MSRA | MFSRWQEG( | MFSRWQEG( | G210E | -1.6826535 | -1.662836 | 0.01981759 |
| TRIM25 | MAELVRAPSV | MAELVRAPSV | T335K | -2.0691068 | -2.0685751 | 0.00053167 |
| F7 | MVSRHYLVLL | MVSRHYLVLL | I26V | -1.5033691 | -1.5011326 | 0.00223649 |
| F7 | MVSRHYLVLL | MVSRHYLVLL | Q72K | -1.5033691 | -1.5028692 | 0.00049984 |
| F7 | MVSRHYLVLL | MVSRHYLVLL | T186A | -1.5033691 | -1.5035356 | -0.0001665 |

| | | | | | | |
|---|---|---|---|---|---|---|
| F7 | MVSRHYLVLL | MVSRHYLVLL | S274P | -1.5033691 | -1.5064733 | -0.0031042 |
| TSFM | MQRAALTGV( | MQRAALTGV( | A157L | -2.1226108 | -2.1098666 | 0.01274419 |
| TSFM | MQRAALTGV( | MQRAALTGV( | T211I | -2.1226108 | -2.137279 | -0.0146682 |
| SORCS1 | MGSSKEPDL\ | MGSSKEPDL\ | R485G | -0.8618349 | -0.8430778 | 0.0187571 |
| AOPEP | MTENLQSLCI | MTENLQSLCI | K75E | -1.9890624 | -1.9884275 | 0.00063491 |
| NFKB1 | MAEEDAYIM( | MAEEDAYIM( | A208T | -1.6191937 | -1.6089504 | 0.0102433 |
| ZBTB20 | MSALGPRLG\ | MSALGPRLG\ | D455N | -1.6593702 | -1.656994 | 0.0023762 |
| FBXO31 | MAVCARLCG | MAVCARLCG | D424G | -1.0183431 | -1.0177298 | 0.00061333 |
| FBXO31 | MAVCARLCG | MAVCARLCG | M465V | -1.0183431 | -1.0202863 | -0.0019432 |
| BSCL2 | MMSSTGPFL( | MMSSTGPFL( | Q288R | -2.0788021 | -2.0762351 | 0.00256705 |
| PPP1R1A | MLLQIISNWS | MLLQIISNWS | K17E | -2.5935051 | -2.5824795 | 0.01102567 |
| VWA5A | MMSCGLLNS | MMSCGLLNS | R81K | -2.3192601 | -2.3213551 | -0.002095 |
| METTL1 | MAGAPPQKR | MAGAPPQKR | H108Q | -1.3231685 | -1.3068405 | 0.01632798 |
| BNIPL | MVPSIKKGRV | MVPSIKKGRV | F64L | -1.1279103 | -1.1273326 | 0.00057769 |
| BNIPL | MVPSIKKGRV | MVPSIKKGRV | D72E | -1.1279103 | -1.1319436 | -0.0040333 |
| BNIPL | MVPSIKKGRV | MVPSIKKGRV | D80E | -1.1279103 | -1.1285809 | -0.0006707 |
| BNIPL | MVPSIKKGRV | MVPSIKKGRV | N82A | -1.1279103 | -1.1287335 | -0.0008233 |
| BNIPL | MVPSIKKGRV | MVPSIKKGRV | E84T | -1.1279103 | -1.1343017 | -0.0063914 |
| BNIPL | MVPSIKKGRV | MVPSIKKGRV | A85P | -1.1279103 | -1.1351341 | -0.0072238 |
| BNIPL | MVPSIKKGRV | MVPSIKKGRV | D86S | -1.1279103 | -1.1351671 | -0.0072569 |
| BNIPL | MVPSIKKGRV | MVPSIKKGRV | I87T | -1.1279103 | -1.1269904 | 0.00091982 |
| BNIPL | MVPSIKKGRV | MVPSIKKGRV | V90L | -1.1279103 | -1.129504 | -0.0015937 |
| BNIPL | MVPSIKKGRV | MVPSIKKGRV | T91C | -1.1279103 | -1.143652 | -0.0157417 |
| BNIPL | MVPSIKKGRV | MVPSIKKGRV | Q94R | -1.1279103 | -1.1384238 | -0.0105135 |
| BNIPL | MVPSIKKGRV | MVPSIKKGRV | K103P | -1.1279103 | -1.1640673 | -0.036157 |
| BNIPL | MVPSIKKGRV | MVPSIKKGRV | V104M | -1.1279103 | -1.1382015 | -0.0102912 |
| BNIPL | MVPSIKKGRV | MVPSIKKGRV | K107R | -1.1279103 | -1.1448977 | -0.0169874 |
| BNIPL | MVPSIKKGRV | MVPSIKKGRV | M109S | -1.1279103 | -1.1456863 | -0.017776 |
| BNIPL | MVPSIKKGRV | MVPSIKKGRV | I113L | -1.1279103 | -1.1436913 | -0.015781 |
| BNIPL | MVPSIKKGRV | MVPSIKKGRV | T116S | -1.1279103 | -1.139277 | -0.0113667 |
| BNIPL | MVPSIKKGRV | MVPSIKKGRV | N130T | -1.1279103 | -1.1419008 | -0.0139905 |
| BNIPL | MVPSIKKGRV | MVPSIKKGRV | G133S | -1.1279103 | -1.146147 | -0.0182368 |
| BNIPL | MVPSIKKGRV | MVPSIKKGRV | E134D | -1.1279103 | -1.1351722 | -0.007262 |
| BNIPL | MVPSIKKGRV | MVPSIKKGRV | V135L | -1.1279103 | -1.1295342 | -0.001624 |
| BNIPL | MVPSIKKGRV | MVPSIKKGRV | D136E | -1.1279103 | -1.1394926 | -0.0115824 |
| BNIPL | MVPSIKKGRV | MVPSIKKGRV | L137I | -1.1279103 | -1.1456811 | -0.0177709 |
| BNIPL | MVPSIKKGRV | MVPSIKKGRV | D139E | -1.1279103 | -1.1362324 | -0.0083221 |
| BNIPL | MVPSIKKGRV | MVPSIKKGRV | D141E | -1.1279103 | -1.1454929 | -0.0175827 |
| BNIPL | MVPSIKKGRV | MVPSIKKGRV | N146S | -1.1279103 | -1.1433247 | -0.0154145 |
| BNIPL | MVPSIKKGRV | MVPSIKKGRV | S147G | -1.1279103 | -1.1444094 | -0.0164992 |
| BNIPL | MVPSIKKGRV | MVPSIKKGRV | N148H | -1.1279103 | -1.1471646 | -0.0192543 |
| BNIPL | MVPSIKKGRV | MVPSIKKGRV | D155E | -1.1279103 | -1.1406525 | -0.0127423 |
| BNIPL | MVPSIKKGRV | MVPSIKKGRV | K158R | -1.1279103 | -1.1422453 | -0.014335 |
| BNIPL | MVPSIKKGRV | MVPSIKKGRV | P159A | -1.1279103 | -1.1471869 | -0.0192766 |
| BNIPL | MVPSIKKGRV | MVPSIKKGRV | K160E | -1.1279103 | -1.1612263 | -0.033316 |

| | | | | | | |
|---|---|---|---|---|---|---|
| BNIPL | MVPSIKKGRV | MVPSIKKGRV | T161G | -1.1279103 | -1.1613466 | -0.0334363 |
| BNIPL | MVPSIKKGRV | MVPSIKKGRV | T162L | -1.1279103 | -1.1729683 | -0.045058 |
| BNIPL | MVPSIKKGRV | MVPSIKKGRV | E163G | -1.1279103 | -1.1632493 | -0.035339 |
| BNIPL | MVPSIKKGRV | MVPSIKKGRV | G168T | -1.1279103 | -1.1530983 | -0.0251881 |
| BNIPL | MVPSIKKGRV | MVPSIKKGRV | P171T | -1.1279103 | -1.1133687 | 0.01454151 |
| BNIPL | MVPSIKKGRV | MVPSIKKGRV | Y173E | -1.1279103 | -1.1814998 | -0.0535896 |
| BNIPL | MVPSIKKGRV | MVPSIKKGRV | T174R | -1.1279103 | -1.1589378 | -0.0310276 |
| BNIPL | MVPSIKKGRV | MVPSIKKGRV | A175L | -1.1279103 | -1.1598941 | -0.0319839 |
| BNIPL | MVPSIKKGRV | MVPSIKKGRV | A176G | -1.1279103 | -1.14561 | -0.0176997 |
| BNIPL | MVPSIKKGRV | MVPSIKKGRV | R179G | -1.1279103 | -1.158392 | -0.0304817 |
| BNIPL | MVPSIKKGRV | MVPSIKKGRV | D180E | -1.1279103 | -1.1268204 | 0.00108981 |
| BNIPL | MVPSIKKGRV | MVPSIKKGRV | R184H | -1.1279103 | -1.1557362 | -0.027826 |
| BNIPL | MVPSIKKGRV | MVPSIKKGRV | M187V | -1.1279103 | -1.1415377 | -0.0136274 |
| BNIPL | MVPSIKKGRV | MVPSIKKGRV | I190M | -1.1279103 | -1.144662 | -0.0167518 |
| BNIPL | MVPSIKKGRV | MVPSIKKGRV | Q193R | -1.1279103 | -1.1453319 | -0.0174216 |
| BNIPL | MVPSIKKGRV | MVPSIKKGRV | D194E | -1.1279103 | -1.1357197 | -0.0078094 |
| BNIPL | MVPSIKKGRV | MVPSIKKGRV | H195Q | -1.1279103 | -1.1422013 | -0.014291 |
| BNIPL | MVPSIKKGRV | MVPSIKKGRV | K200T | -1.1279103 | -1.1492788 | -0.0213685 |
| BNIPL | MVPSIKKGRV | MVPSIKKGRV | A201V | -1.1279103 | -1.1390872 | -0.0111769 |
| BNIPL | MVPSIKKGRV | MVPSIKKGRV | V223I | -1.1279103 | -1.1317995 | -0.0038892 |
| BNIPL | MVPSIKKGRV | MVPSIKKGRV | V224L | -1.1279103 | -1.147451 | -0.0195408 |
| BNIPL | MVPSIKKGRV | MVPSIKKGRV | V227S | -1.1279103 | -1.1586089 | -0.0306987 |
| BNIPL | MVPSIKKGRV | MVPSIKKGRV | F229Y | -1.1279103 | -1.1455173 | -0.0176071 |
| BNIPL | MVPSIKKGRV | MVPSIKKGRV | M230L | -1.1279103 | -1.1440569 | -0.0161467 |
| BNIPL | MVPSIKKGRV | MVPSIKKGRV | E232R | -1.1279103 | -1.1492246 | -0.0213144 |
| BNIPL | MVPSIKKGRV | MVPSIKKGRV | G234S | -1.1279103 | -1.1323067 | -0.0043964 |
| BNIPL | MVPSIKKGRV | MVPSIKKGRV | Q235I | -1.1279103 | -1.1595331 | -0.0316229 |
| BNIPL | MVPSIKKGRV | MVPSIKKGRV | R239T | -1.1279103 | -1.1571181 | -0.0292078 |
| BNIPL | MVPSIKKGRV | MVPSIKKGRV | L241V | -1.1279103 | -1.1424441 | -0.0145339 |
| BNIPL | MVPSIKKGRV | MVPSIKKGRV | D243E | -1.1279103 | -1.1381758 | -0.0102656 |
| BNIPL | MVPSIKKGRV | MVPSIKKGRV | N244H | -1.1279103 | -1.1450011 | -0.0170908 |
| BNIPL | MVPSIKKGRV | MVPSIKKGRV | K247R | -1.1279103 | -1.1390691 | -0.0111588 |
| BNIPL | MVPSIKKGRV | MVPSIKKGRV | V249M | -1.1279103 | -1.1495143 | -0.0216041 |
| BNIPL | MVPSIKKGRV | MVPSIKKGRV | M262L | -1.1279103 | -1.1421957 | -0.0142854 |
| BNIPL | MVPSIKKGRV | MVPSIKKGRV | I263L | -1.1279103 | -1.1476464 | -0.0197362 |
| BNIPL | MVPSIKKGRV | MVPSIKKGRV | Y265H | -1.1279103 | -1.156379 | -0.0284687 |
| BNIPL | MVPSIKKGRV | MVPSIKKGRV | T271S | -1.1279103 | -1.1369419 | -0.0090317 |
| BNIPL | MVPSIKKGRV | MVPSIKKGRV | R273A | -1.1279103 | -1.1603141 | -0.0324038 |
| BNIPL | MVPSIKKGRV | MVPSIKKGRV | K274Q | -1.1279103 | -1.1503794 | -0.0224692 |
| BNIPL | MVPSIKKGRV | MVPSIKKGRV | M275V | -1.1279103 | -1.1448357 | -0.0169255 |
| BNIPL | MVPSIKKGRV | MVPSIKKGRV | L281I | -1.1279103 | -1.1465886 | -0.0186783 |
| BNIPL | MVPSIKKGRV | MVPSIKKGRV | K283Q | -1.1279103 | -1.1458241 | -0.0179138 |
| BNIPL | MVPSIKKGRV | MVPSIKKGRV | Q286R | -1.1279103 | -1.1460638 | -0.0181535 |
| BNIPL | MVPSIKKGRV | MVPSIKKGRV | Q287T | -1.1279103 | -1.150053 | -0.0221428 |
| BNIPL | MVPSIKKGRV | MVPSIKKGRV | S298A | -1.1279103 | -1.1369033 | -0.008993 |

| | | | | | | |
|---|---|---|---|---|---|---|
| BNIPL | MVPSIKKGRV | MVPSIKKGRV | S305T | -1.1279103 | -1.1414021 | -0.0134919 |
| BNIPL | MVPSIKKGRV | MVPSIKKGRV | F307Y | -1.1279103 | -1.1406186 | -0.0127083 |
| BNIPL | MVPSIKKGRV | MVPSIKKGRV | R309K | -1.1279103 | -1.1433523 | -0.015442 |
| BNIPL | MVPSIKKGRV | MVPSIKKGRV | T310A | -1.1279103 | -1.1417679 | -0.0138576 |
| BNIPL | MVPSIKKGRV | MVPSIKKGRV | I314L | -1.1279103 | -1.1349007 | -0.0069904 |
| BNIPL | MVPSIKKGRV | MVPSIKKGRV | T315L | -1.1279103 | -1.1593529 | -0.0314426 |
| BNIPL | MVPSIKKGRV | MVPSIKKGRV | K316R | -1.1279103 | -1.1338569 | -0.0059466 |
| BNIPL | MVPSIKKGRV | MVPSIKKGRV | S324T | -1.1279103 | -1.1448094 | -0.0168991 |
| BNIPL | MVPSIKKGRV | MVPSIKKGRV | Q325R | -1.1279103 | -1.1426857 | -0.0147754 |
| BNIPL | MVPSIKKGRV | MVPSIKKGRV | Y329F | -1.1279103 | -1.143132 | -0.0152217 |
| BNIPL | MVPSIKKGRV | MVPSIKKGRV | F331D | -1.1279103 | -1.1875172 | -0.0596069 |
| BNIPL | MVPSIKKGRV | MVPSIKKGRV | T332S | -1.1279103 | -1.128701 | -0.0007907 |
| BNIPL | MVPSIKKGRV | MVPSIKKGRV | A334G | -1.1279103 | -1.1428257 | -0.0149155 |
| BNIPL | MVPSIKKGRV | MVPSIKKGRV | E338Q | -1.1279103 | -1.1471794 | -0.0192691 |
| BNIPL | MVPSIKKGRV | MVPSIKKGRV | M342L | -1.1279103 | -1.1442803 | -0.0163701 |
| BNIPL | MVPSIKKGRV | MVPSIKKGRV | Y344Q | -1.1279103 | -1.1701976 | -0.0422873 |
| BNIPL | MVPSIKKGRV | MVPSIKKGRV | G346H | -1.1279103 | -1.165601 | -0.0376908 |
| BNIPL | MVPSIKKGRV | MVPSIKKGRV | I351V | -1.1279103 | -1.139186 | -0.0112758 |
| BNIPL | MVPSIKKGRV | MVPSIKKGRV | K352R | -1.1279103 | -1.1359676 | -0.0080574 |
| BNIPL | MVPSIKKGRV | MVPSIKKGRV | I366L | -1.1279103 | -1.1329736 | -0.0050633 |
| BNIPL | MVPSIKKGRV | MVPSIKKGRV | Q368R | -1.1279103 | -1.1322147 | -0.0043044 |
| BNIPL | MVPSIKKGRV | MVPSIKKGRV | E369D | -1.1279103 | -1.1325192 | -0.004609 |
| BNIPL | MVPSIKKGRV | MVPSIKKGRV | D371H | -1.1279103 | -1.1335206 | -0.0056103 |
| BNIPL | MVPSIKKGRV | MVPSIKKGRV | K373S | -1.1279103 | -1.1364609 | -0.0085506 |
| BNIPL | MVPSIKKGRV | MVPSIKKGRV | Q374G | -1.1279103 | -1.1351695 | -0.0072592 |
| BNIPL | MVPSIKKGRV | MVPSIKKGRV | E375G | -1.1279103 | -1.1359034 | -0.0079931 |
| BNIPL | MVPSIKKGRV | MVPSIKKGRV | E379T | -1.1279103 | -1.1347078 | -0.0067976 |
| PIK3C2G | MKQISISTVDI | NKQISISTVDI | M1N | -2.8635077 | -2.8734505 | -0.0099428 |
| SVEP1 | MASHPKEEH | MASHPKEEH | L261V | -2.12502 | -2.1248293 | 0.00019073 |
| SVEP1 | MASHPKEEH | MASHPKEEH | I392T | -2.12502 | -2.1243868 | 0.00063324 |
| SVEP1 | MASHPKEEH | MASHPKEEH | N398G | -2.12502 | -2.1239913 | 0.00102878 |
| SVEP1 | MASHPKEEH | MASHPKEEH | L599S | -2.12502 | -2.1249311 | 8.89E-05 |
| SVEP1 | MASHPKEEH | MASHPKEEH | V3008S | -2.12502 | -2.1247454 | 0.00027466 |
| RAPGEF3 | MHLFRSYSY( | MHLFRSYSY( | T835V | -1.6936713 | -1.6962665 | -0.0025952 |
| XRCC6 | MSEWMSYYK | MSEWMSYYK | W68Q | -2.6060545 | -2.5818772 | 0.02417731 |
| IFNAR1 | MAPPELLLLL | MAPPELLLLL | E70T | -2.5017288 | -2.5055192 | -0.0037904 |
| CYP19A1 | MILETLNLMH | MILETLNLMH | M158V | -1.35197 | -1.34951 | 0.00246 |
| CYP19A1 | MILETLNLMH | MILETLNLMH | H461K | -1.35197 | -1.35054 | 0.00142992 |
| METTL21B | MAASRQPLL( | MAASRQPLL( | V215H | -2.0149713 | -1.9961571 | 0.01881421 |
| METTL21B | MAASRQPLL( | MAASRQPLL( | P219Q | -2.0149713 | -1.9961604 | 0.01881087 |
| HIF1A | MELTGEKKRI | MELTGEKKRI | R584S | -1.5047575 | -1.5023383 | 0.00241923 |
| MTFMT | MRGRLLRAW | MRGRLLRAW | V16G | -1.8749663 | -1.8635287 | 0.01143754 |
| MTFMT | MRGRLLRAW | MRGRLLRAW | L307S | -1.8749663 | -1.863543 | 0.01142323 |
| GZMB | MLILILLPVALL | MLILILLPVALL | Y93K | -1.6560966 | -1.6360747 | 0.02002192 |
| GZMB | MLILILLPVALL | MLILILLPVALL | S150Q | -1.6560966 | -1.6521921 | 0.00390446 |

| | | | | | | |
|---|---|---|---|---|---|---|
| GZMB | MLILILLPVALL | MLILILLPVALL | Y219Q | -1.6560966 | -1.6402192 | 0.01587737 |
| GHSR | MRNQSGSPN | MRNQSGSPN | S328R | -0.8481347 | -0.8092991 | 0.03883559 |
| BAG6 | MAEAQNLEVI | MAEAQNLEVI | L1075V | -2.208987 | -2.2087283 | 0.00025868 |
| ZPLD1 | MERIWLLLLLI | MERIWLLLLLT | I11T | -1.0980308 | -1.0946571 | 0.00337374 |
| SYT13 | MVLSVPVIALC | MVLSVPVIALC | I160A | -1.9708692 | -1.961511 | 0.00935817 |
| POLM | MALVPLKRRF | MALVPLKRRF | A18T | -2.0268738 | -2.0331533 | -0.0062795 |
| POLM | MALVPLKRRF | MALVPLKRRF | Q140P | -2.0268738 | -2.0264802 | 0.00039363 |
| POLM | MALVPLKRRF | MALVPLKRRF | T320L | -2.0268738 | -2.0220773 | 0.0047965 |
| POLM | MALVPLKRRF | MALVPLKRRF | M348L | -2.0268738 | -2.0216115 | 0.00526237 |
| AKT1S1 | MADNHRESV | MADNHRESV | T83P | -2.7383749 | -2.7309132 | 0.00746179 |
| TEP1 | MPPRAQPEPI | MPPRAQPEPI | V1966A | -2.566374 | -2.5659464 | 0.00042756 |
| TEP1 | MPPRAQPEPI | MPPRAQPEPI | L1991S | -2.566374 | -2.5663686 | 5.40E-06 |
| CCL5 | MKATVAALAV | MKATVAALAV | P22A | -1.1387188 | -1.1254373 | 0.01328158 |
| ENOX1 | MLWMEQFLC | MLWMEQFLC | S354C | -0.6069209 | -0.6129102 | -0.0059893 |
| RBPMS | MNSLPADKES | MNSLPADKES | G16N | -1.5220951 | -1.5249619 | -0.0028669 |
| MRE11A | MSCINSGDE | MSCINSGDE | S620N | -1.3677784 | -1.3681926 | -0.0004141 |
| MRE11A | MSCINSGDE | MSCINSGDE | G625D | -1.3677784 | -1.3647761 | 0.00300229 |
| CPA2 | MRLILFFSALA | MRLILFFSALA | E194N | -1.1475086 | -1.1394378 | 0.00807083 |
| NAV2 | MNKVEIYTDV | MNKVEIYTDV | C1122S | -1.5593932 | -1.5517168 | 0.00767644 |
| LINGO2 | MLHTATSFW( | MLHTATSFW( | V470I | -0.6406263 | -0.6356325 | 0.0049938 |
| LINGO2 | MLHTATSFW( | MLHTATSFW( | I591V | -0.6406263 | -0.6343116 | 0.00631464 |
| ZFYVE28 | MRPIHPFSFC | MRPIHPFSFC | K556G | -1.7154078 | -1.7166224 | -0.0012145 |
| ECHS1 | MAVALSSLRR | MAVALSSLRR | I50V | -0.8212193 | -0.8103257 | 0.01089352 |
| GZMH-aging- | MIQGNWSLA | MIQGNWSLA | F78L | -1.8947619 | -1.8874048 | 0.00735712 |
| GZMH-aging- | MIQGNWSLA | MIQGNWSLA | I80L | -1.8947619 | -1.8861737 | 0.00858819 |
| GZMH-aging- | MIQGNWSLA | MIQGNWSLA | P83L | -1.8947619 | -1.8809023 | 0.01385963 |
| GZMH-aging- | MIQGNWSLA | MIQGNWSLA | L85T | -1.8947619 | -1.8993036 | -0.0045416 |
| GZMH-aging- | MIQGNWSLA | MIQGNWSLA | Q89G | -1.8947619 | -1.8907368 | 0.0040251 |
| PPM1D | MEGLYSLGVS | MEGLYSLGVS | I437V | -1.6314447 | -1.6311216 | 0.00032306 |
| POLE2 | MAAEKLRSKV | MAAEKLRSKV | V107L | -1.1180189 | -1.1084863 | 0.00953257 |
| TOP2A | MELDESLSPL | MELDESLSPL | S685P | -1.7770929 | -1.7739142 | 0.00317872 |
| PPP1R26 | MFLMNASPLI | MFLMNASPLI | G499D | -2.8222785 | -2.8222136 | 6.48E-05 |
| ERCC1 | MEPMEAAQV | MEPMEAAQV | S93L | -1.5318162 | -1.5338446 | -0.0020283 |
| CSMD3 | MCMGEINLE | MCMGEINLE | A96M | -1.815724 | -1.8161751 | -0.0004511 |
| CSMD3 | MCMGEINLE | MCMGEINLE | M1405V | -1.815724 | -1.814754 | 0.00097001 |
| CRLS1 | MHRRGAEEP | MHRRGAEEP | L234V | -1.1788464 | -1.1680998 | 0.0107466 |
| APOA4 | MAELRGPCG | MAELRGPCG | N355A | -2.1100454 | -2.1022289 | 0.00781655 |
| APOB | MGPPQLWQI | MGPPQLWQI | Q2004R | -2.7829035 | -2.783008 | -0.0001045 |
| APOB | MGPPQLWQI | MGPPQLWQI | S4283A | -2.7829035 | -2.7831463 | -0.0002428 |
| HDAC6 | MKRGGRRGI | MKRGGRRGI | R16N | -2.8460567 | -2.8469629 | -0.0009062 |
| WDR27 | MESPVDLCFT | MESPVDLCFT | K630E | -2.5900869 | -2.5889783 | 0.00110865 |
| HIP1 | MEVGKMLML | MEVGKMLML | G319D | -1.1207668 | -1.1073898 | 0.01337695 |
| CEL | MEPRILKWD\ | MEPRILKWD\ | M483L | -1.6347173 | -1.6335312 | 0.00118613 |
| TERT | MAGSGPFGA | MAGSGPFGA | S19R | -2.7507617 | -2.7496011 | 0.00116062 |
| TERT | MAGSGPFGA | MAGSGPFGA | Y197L | -2.7507617 | -2.7503688 | 0.00039291 |

| | | | | | | |
|---|---|---|---|---|---|---|
| NCAM2 | MYAKCIHAEF | MYAKCIHAEF | I135V | -0.6073373 | -0.6048717 | 0.00246561 |
| NCAM2 | MYAKCIHAEF | MYAKCIHAEF | E279R | -0.6073373 | -0.613521 | -0.0061837 |
| LEP | MRCPSLPLC( | MRCPSLPLC( | V148L | -2.8385062 | -2.8303294 | 0.0081768 |
| PARVG | MEDAVVTEEL | MEDAVVTEEL | P303L | -1.9011797 | -1.8803154 | 0.02086425 |
| PARVG | MEDAVVTEEL | MEDAVVTEEL | Q306N | -1.9011797 | -1.9106145 | -0.0094348 |
| XDH | MASERTTDQI | MASERTTDQI | M533T | -1.5939362 | -1.5912173 | 0.00271893 |
| APOM | MLQRSWSYL | MLQRSWSYL | I70V | -2.8315623 | -2.8286543 | 0.00290799 |
| CHIT1 | MGRVFVWIG | MGRVFVWIG | K198D | -1.4959505 | -1.4971154 | -0.0011649 |
| LMNB1 | MAASTPVGQI | MAASTPVGQI | M94T | -0.845464 | -0.825857 | 0.01960695 |
| TPP2 | MAAAAEEPFF | MAAAAEEPFF | L635A | -2.1499972 | -2.1458316 | 0.00416565 |
| POLE | MVLRNSGRR | MVLRNSGRR | M145V | -2.8419685 | -2.8416897 | 0.00027879 |
| POLE | MVLRNSGRR | MVLRNSGRR | H1230R | -2.8419685 | -2.8408515 | 0.00111707 |
| GSR | MAAAFDYLVL | MAAAFDYLVL | L60S | -0.7465237 | -0.7266786 | 0.01984513 |
| HLA-DRB1 | MGAGWILGA | MGAGWILLA( | G8L | -1.2789593 | -1.2809184 | -0.0019591 |
| HLA-DRB1 | MGAGWILGA | MGAGWILGA | S11G | -1.2789593 | -1.2756618 | 0.00329745 |
| HLA-DRB1 | MGAGWILGA | MGAGWILGA | C12S | -1.2789593 | -1.2767736 | 0.0021857 |
| HLA-DRB1 | MGAGWILGA | MGAGWILGA | W13C | -1.2789593 | -1.2796713 | -0.000712 |
| HLA-DRB1 | MGAGWILGA | MGAGWILGA | A14M | -1.2789593 | -1.2830606 | -0.0041013 |
| HLA-DRB1 | MGAGWILGA | MGAGWILGA | T22M | -1.2789593 | -1.2795422 | -0.0005829 |
| HLA-DRB1 | MGAGWILGA | MGAGWILGA | P35R | -1.2789593 | -1.2820439 | -0.0030847 |
| HLA-DRB1 | MGAGWILGA | MGAGWILGA | R36P | -1.2789593 | -1.2820522 | -0.0030929 |
| HLA-DRB1 | MGAGWILGA | MGAGWILGA | G37R | -1.2789593 | -1.2724187 | 0.00654054 |
| HLA-DRB1 | MGAGWILGA | MGAGWILGA | V39L | -1.2789593 | -1.2753606 | 0.00359869 |
| HLA-DRB1 | MGAGWILGA | MGAGWILGA | Y40W | -1.2789593 | -1.2876281 | -0.0086688 |
| HLA-DRB1 | MGAGWILGA | MGAGWILGA | T42P | -1.2789593 | -1.2897249 | -0.0107657 |
| HLA-DRB1 | MGAGWILGA | MGAGWILGA | N43K | -1.2789593 | -1.2602434 | 0.01871586 |
| HLA-DRB1 | MGAGWILGA | MGAGWILGA | Y47H | -1.2789593 | -1.2802988 | -0.0013396 |
| HLA-DRB1 | MGAGWILGA | MGAGWILGA | H68S | -1.2789593 | -1.2784739 | 0.00048542 |
| HLA-DRB1 | MGAGWILGA | MGAGWILGA | L69V | -1.2789593 | -1.276812 | 0.00214732 |
| HLA-DRB1 | MGAGWILGA | MGAGWILGA | W78F | -1.2789593 | -1.2618773 | 0.01708198 |
| HLA-DRB1 | MGAGWILGA | MGAGWILGA | A87P | -1.2789593 | -1.2685114 | 0.01044786 |
| HLA-DRB1 | MGAGWILGA | MGAGWILGA | T88D | -1.2789593 | -1.2744091 | 0.00455022 |
| HLA-DRB1 | MGAGWILGA | MGAGWILGA | L91Y | -1.2789593 | -1.2724049 | 0.00655437 |
| HLA-DRB1 | MGAGWILGA | MGAGWILGA | S98I | -1.2789593 | -1.2631277 | 0.01583159 |
| HLA-DRB1 | MGAGWILGA | MGAGWILGA | Q100E | -1.2789593 | -1.2629215 | 0.01603782 |
| HLA-DRB1 | MGAGWILGA | MGAGWILGA | Y101Q | -1.2789593 | -1.2672104 | 0.01174891 |
| HLA-DRB1 | MGAGWILGA | MGAGWILGA | K102A | -1.2789593 | -1.2769099 | 0.00204933 |
| HLA-DRB1 | MGAGWILGA | MGAGWILGA | Q115G | -1.2789593 | -1.2741997 | 0.00475955 |
| HLA-DRB1 | MGAGWILGA | MGAGWILGA | E116V | -1.2789593 | -1.2602693 | 0.01868999 |
| HLA-DRB1 | MGAGWILGA | MGAGWILGA | T117V | -1.2789593 | -1.2692513 | 0.00970793 |
| HLA-DRB1 | MGAGWILGA | MGAGWILGA | S123Q | -1.2789593 | -1.27196 | 0.00699925 |
| HLA-DRB1 | MGAGWILGA | MGAGWILGA | K131T | -1.2789593 | -1.2825992 | -0.0036399 |
| HLA-DRB1 | MGAGWILGA | MGAGWILGA | S133Y | -1.2789593 | -1.3001078 | -0.0211486 |
| HLA-DRB1 | MGAGWILGA | MGAGWILGA | E140L | -1.2789593 | -1.2657145 | 0.01324475 |
| HLA-DRB1 | MGAGWILGA | MGAGWILGA | P143H | -1.2789593 | -1.2818635 | -0.0029042 |

| | | | | | | |
|---|---|---|---|---|---|---|
| HLA-DRB1 | MGAGWILGA | MGAGWILGA | T151S | -1.2789593 | -1.285918 | -0.0069587 |
| HLA-DRB1 | MGAGWILGA | MGAGWILGA | L167Q | -1.2789593 | -1.255271 | 0.02368832 |
| HLA-DRB1 | MGAGWILGA | MGAGWILGA | Q212T | -1.2789593 | -1.2825087 | -0.0035495 |
| HLA-DRB1 | MGAGWILGA | MGAGWILGA | G213S | -1.2789593 | -1.2769523 | 0.00200701 |
| HRH1 | MCETKNSTE/ | MCETKNSTE/ | I27V | -1.8812164 | -1.8812178 | -1.43E-06 |
| SH2D4A | MLQQILADM | MLQQILADM | Q337R | -1.9469746 | -1.9342523 | 0.01272237 |
| NEDD4 | MARRLRLHF/ | MARRLRLHF/ | Q259K | -1.3886931 | -1.3884208 | 0.00027227 |
| NEDD4 | MARRLRLHF/ | MARRLRLHF/ | T346K | -1.3886931 | -1.3886913 | 1.79E-06 |
| NEDD4 | MARRLRLHF/ | MARRLRLHF/ | A757F | -1.3886931 | -1.3891096 | -0.0004165 |
| NEDD4 | MARRLRLHF/ | MARRLRLHF/ | A815S | -1.3886931 | -1.3880341 | 0.00065899 |
| MARCO | MENPKNYNE | MENPKNYNE | M46L | -2.2910089 | -2.2861521 | 0.00485682 |
| MAPK1 | MAAVSGAGA | MAAVSGAGA | D366Y | -0.3040255 | -0.2577772 | 0.04624829 |
| CFH | MLPTDCETPF | MLPTDCETPF | D250N | -2.2314088 | -2.2293725 | 0.00203633 |
| CFH | MLPTDCETPF | MLPTDCETPF | D252S | -2.2314088 | -2.2339809 | -0.0025721 |
| CFH | MLPTDCETPF | MLPTDCETPF | R254S | -2.2314088 | -2.2297654 | 0.00164342 |
| CFH | MLPTDCETPF | MLPTDCETPF | K256T | -2.2314088 | -2.2314217 | -1.29E-05 |
| CFH | MLPTDCETPF | MLPTDCETPF | S260H | -2.2314088 | -2.236336 | -0.0049272 |
| CFH | MLPTDCETPF | MLPTDCETPF | K262S | -2.2314088 | -2.2297421 | 0.00166678 |
| CFH | MLPTDCETPF | MLPTDCETPF | E264T | -2.2314088 | -2.228761 | 0.00264788 |
| CFH | MLPTDCETPF | MLPTDCETPF | D267H | -2.2314088 | -2.23563 | -0.0042212 |
| CFH | MLPTDCETPF | MLPTDCETPF | T277V | -2.2314088 | -2.226994 | 0.0044148 |
| GCNT3 | MFRERKYLC( | MFRERKRLC( | Y7R | -1.6097413 | -1.6038917 | 0.0058496 |
| DHX33 | MPQGGEVPP | MPQEGEVPP/ | G4E | -0.8623556 | -0.8625923 | -0.0002367 |
| NFATC1 | MPSPGFPTSK | MPSPGFPTSK | A935V | -1.3871887 | -1.3852656 | 0.00192308 |
| PLEKHA1 | MPYVDRQNF | MPYVDRQNF | S296P | -1.0722729 | -1.0567876 | 0.01548529 |
| PUM2 | MSIPCVLLGL | MSIPCVLLGL | S372N | -1.0999531 | -1.0970321 | 0.00292099 |
| TREML1 | MGSFPAWLL | MGSFLAWLLI | P5L | -2.6789854 | -2.6780348 | 0.00095057 |
| TREML1 | MGSFPAWLL | MGSFPAWLLI | E15G | -2.6789854 | -2.6748087 | 0.00417662 |
| TREML1 | MGSFPAWLL | MGSFPAWLLI | V54F | -2.6789854 | -2.6844618 | -0.0054765 |
| TREML1 | MGSFPAWLL | MGSFPAWLLI | L114R | -2.6789854 | -2.6786482 | 0.00033712 |
| TNFSF8 | MCFSPEKTLF | MCFSPEKTLF | H69P | -2.8659182 | -2.8589046 | 0.00701356 |
| PKN2 | MASNTAEREI | MASNTAEREI | G566E | -0.7706791 | -0.7705839 | 9.51E-05 |
| CD302 | MEPPFPALLC | MEPPFPALLC | V113M | -2.4767833 | -2.4838212 | -0.0070379 |
| IRG1 | MLWAGEVLL/ | MLWAGEVLL/ | S164K | -1.7361449 | -1.7344315 | 0.0017134 |
| MID2 | METLESELTC | METLESELTC | L39V | -0.5082177 | -0.5015726 | 0.00664508 |
| DHX9 | MGDVKNFLY | MGDVKNFLY | A108T | -1.9056183 | -1.9060197 | -0.0004014 |
| C5AR1 | MSVNEVSED | MSVNEVSED | N184F | -1.7747191 | -1.7768527 | -0.0021336 |
| IL1RL2 | MGEECMVHI | MGEECMVHI | S89G | -2.1233728 | -2.1261425 | -0.0027697 |
| ATF6 | MAAEEVEQG | MAAEEVEQG | S391R | -2.2637596 | -2.264982 | -0.0012224 |
| ATF6 | MAAEEVEQG | MAAEEVEQG | K577T | -2.2637596 | -2.2642105 | -0.0004508 |
| ST6GAL1 | MVHINLLKKL | MVHINLLKKL | G160D | -1.8300288 | -1.8124281 | 0.01760066 |
| LY75 | MRRAPGRSL | MRRAPGRSL | S150P | -2.480602 | -2.4794216 | 0.00118041 |
| LY75 | MRRAPGRSL | MRRAPGRSL | N486K | -2.480602 | -2.4803226 | 0.00027943 |
| LY75 | MRRAPGRSL | MRRAPGRSL | E957Q | -2.480602 | -2.481051 | -0.0004489 |
| LY75 | MRRAPGRSL | MRRAPGRSL | H1354T | -2.480602 | -2.4800103 | 0.00059175 |

| | | | | | | |
|---|---|---|---|---|---|---|
| LY75 | MRRAPGRSL | MRRAPGRSL | Q1519K | -2.480602 | -2.4804971 | 0.0001049 |
| RABGEF1 | EEMSLKSERR | EEMSLKSERR | L14V | -0.5138707 | -0.5059975 | 0.00787318 |
| CD1D | MGNTKVRPK | MGNTKVRPK | T122G | -2.2292109 | -2.2268674 | 0.00234342 |
| CD1D | MGNTKVRPK | MGNTKVRPK | Q312K | -2.2292109 | -2.2290485 | 0.00016236 |
| ADAR | MNTGFGRGI | MNTGFGRGI | D397G | -2.5995288 | -2.5997434 | -0.0002146 |
| ATP7A | MAQGLVTVG | MAQGLVTVG | G576R | -1.6527474 | -1.6495612 | 0.00318623 |
| ATP7A | MAQGLVTVG | MAQGLVTVG | G891D | -1.6527474 | -1.6487375 | 0.00400984 |
| ATP7A | MAQGLVTVG | MAQGLVTVG | N1212Q | -1.6527474 | -1.6534061 | -0.0006588 |
| PSMD5 | MAEEALELLAI | MAEEALELLAI | A266S | -1.5431502 | -1.5276787 | 0.01547146 |
| TRIM8 | MAENWKNC | MAENWKNC | L225V | -1.9437178 | -1.9362484 | 0.00746942 |
| PIK3AP1 | MASAGAPGG | MASAGAPGG | I705R | -2.3379669 | -2.3371353 | 0.0008316 |
| PIK3AP1 | MASAGAPGG | MASAGAPGG | M708L | -2.3379669 | -2.3367443 | 0.00122261 |
| APOBEC2 | MAENQEKPR | MAENQEKPR | S21L | -1.3881098 | -1.3908967 | -0.0027869 |
| DAPK1 | MTVFRQENLI | MTVFRQENLI | S1355P | -2.1912417 | -2.1922042 | -0.0009625 |
| STK11 | MDVQEPQEL | MDVQEPQEL | R108K | -0.6910493 | -0.6803699 | 0.01067942 |
| NFKBIZ | MIVESNREAP | MIVESNREAP | H160C | -1.9976228 | -1.9982258 | -0.000603 |
| NFKBIZ | MIVESNREAP | MIVESNREAP | V334I | -1.9976228 | -1.9976863 | -6.34E-05 |
| TRAT1 | MTIVLLSQILF | MTIVLLSQILF | D115K | -2.8808963 | -2.8813453 | -0.0004489 |
| IL17C | MSLLERLLGY | MSLLERLLGY | V87L | -2.1848557 | -2.1814892 | 0.00336647 |
| CD38 | MPLQNRSSK | MPLQNRSSK | M40R | -2.1830781 | -2.158541 | 0.02453709 |
| FGA | MEHGIQSDC | MEHGIQSDC | E251P | -2.3860304 | -2.3866 | -0.0005696 |
| IFNL3 | MKKCHLSKYI | MKKCHLSKYI | W15Q | -2.8906658 | -2.8827507 | 0.00791502 |
| IFNL3 | MKKCHLSKYI | MKKCHLSKYI | K18Q | -2.8906658 | -2.8940234 | -0.0033576 |
| IFNL3 | MKKCHLSKYI | MKKCHLSKYI | T125A | -2.8906658 | -2.8871808 | 0.00348496 |
| DHX58 | MLTPALPPSY | MLTPALPPSY | S419T | -2.1516941 | -2.1546824 | -0.0029883 |
| MADCAM1 | MEPTLLLLLLS | MEPTLLLLLLS | S254E | -2.543628 | -2.5473707 | -0.0037427 |
| MADCAM1 | MEPTLLLLLLS | MEPTLLLLLLS | S353R | -2.543628 | -2.5484812 | -0.0048532 |
| MADCAM1 | MEPTLLLLLLS | MEPTLLLLLLS | A474P | -2.543628 | -2.5405684 | 0.00305963 |
| 8-Mar | MPLHQISVIP | MPLHQISVIP | I534T | -1.4058119 | -1.4027472 | 0.00306475 |
| CD207 | MSRENICDN | MMRENICDN | S2M | -2.3115208 | -2.3195145 | -0.0079937 |
| CD207 | MSRENICDN | MSRENICDN | D16A | -2.3115208 | -2.3097053 | 0.00181556 |
| CD207 | MSRENICDN | MSRENICDN | E34K | -2.3115208 | -2.3060975 | 0.00542331 |
| CD207 | MSRENICDN | MSRENICDN | S36P | -2.3115208 | -2.31234 | -0.0008192 |
| CD207 | MSRENICDN | MSRENICDN | C46T | -2.3115208 | -2.3166816 | -0.0051608 |
| CD207 | MSRENICDN | MSRENICDN | E73N | -2.3115208 | -2.3193386 | -0.0078177 |
| CD207 | MSRENICDN | MSRENICDN | E84G | -2.3115208 | -2.309413 | 0.00210786 |
| CD207 | MSRENICDN | MSRENICDN | N85R | -2.3115208 | -2.3111765 | 0.00034428 |
| CD207 | MSRENICDN | MSRENICDN | D88N | -2.3115208 | -2.3122866 | -0.0007658 |
| CD207 | MSRENICDN | MSRENICDN | S150Q | -2.3115208 | -2.3134453 | -0.0019245 |
| CD207 | MSRENICDN | MSRENICDN | A184T | -2.3115208 | -2.320051 | -0.0085301 |
| CD207 | MSRENICDN | MSRENICDN | I199A | -2.3115208 | -2.3142655 | -0.0027447 |
| CD207 | MSRENICDN | MSRENICDN | T216D | -2.3115208 | -2.3149934 | -0.0034726 |
| CD207 | MSRENICDN | MSRENICDN | C218S | -2.3115208 | -2.2914391 | 0.02008176 |
| CD207 | MSRENICDN | MSRENICDN | E232R | -2.3115208 | -2.3072927 | 0.00422812 |
| CD207 | MSRENICDN | MSRENICDN | T246N | -2.3115208 | -2.3120575 | -0.0005367 |

| | | | | | | |
|---|---|---|---|---|---|---|
| MAP4K2 | MSLLRDVSLL | MSLLRDVSLL | S352N | -1.452167 | -1.454248 | -0.0020809 |
| MAP4K2 | MSLLRDVSLL | MSLLRDVSLL | I749V | -1.452167 | -1.4514146 | 0.00075245 |
| JAG1 | MLLPRRPALA | MLLPRRPALA | T1095G | -1.8976099 | -1.9002572 | -0.0026473 |
| ATG9A | MMHFDTQY( | MAHFDTQYC | M2A | -1.6994551 | -1.6979115 | 0.00154364 |
| ATG9A | MMHFDTQY( | MMHFDTQY( | L360P | -1.6994551 | -1.6760449 | 0.0234102 |
| ATG9A | MMHFDTQY( | MMHFDTQY( | V633L | -1.6994551 | -1.6994443 | 1.08E-05 |
| VAV3 | MEPWKQCA( | MEPWKQCA( | Y617Q | -0.6689858 | -0.6643974 | 0.00458843 |
| TNIP3 | MIHCGLLKKK | MIHCGLLKKK | T88G | -2.5017052 | -2.5028348 | -0.0011296 |
| TMEM173 | MSHETERPS\ | MSHETERPS\ | D256E | -2.8382881 | -2.8351824 | 0.00310564 |
| LAMP1 | MARVGGGR( | MARVGGGR( | E35T | -2.2498868 | -2.2502186 | -0.0003319 |
| ITGAL | MQESGCACE | MQESGCACE | L1193M | -2.638716 | -2.6439478 | -0.0052319 |
| PROS1 | MRLLSRRCV/ | MRLLGRRCV | S5G | -1.8189896 | -1.8196119 | -0.0006223 |
| PROS1 | MRLLSRRCV/ | MRLLSRRCV/ | N398S | -1.8189896 | -1.807742 | 0.01124763 |
| ATG16L1 | MTSGLRAATF | MTSGLRAATF | E582G | -0.5570151 | -0.5327456 | 0.02426952 |
| GCH1 | MEKGKLGALI | MEKGKLGALI | Q33P | -0.8795343 | -0.8720823 | 0.00745201 |
| CTSG | MLILILLPAAFI | MLILILLPAAFI | L120R | -1.5524743 | -1.5466863 | 0.00578797 |
| CTSG | MLILILLPAAFI | MLILILLPAAFI | S126N | -1.5524743 | -1.5582824 | -0.0058081 |
| CTSG | MLILILLPAAFI | MLILILLPAAFI | Q150R | -1.5524743 | -1.5423771 | 0.01009715 |
| CTSG | MLILILLPAAFI | MLILILLPAAFI | R184G | -1.5524743 | -1.5711175 | -0.0186433 |
| CTSG | MLILILLPAAFI | MLILILLPAAFI | A227S | -1.5524743 | -1.550101 | 0.00237322 |
| COLEC12 | MRMGVPSSF | MRMGVPSSF | L484P | -1.7391809 | -1.733503 | 0.00567794 |
| COLEC12 | MRMGVPSSF | MRMGVPSSF | A666E | -1.7391809 | -1.7332978 | 0.0058831 |
| CD97 | MARLCIALCL | MARLCIALCL | T54F | -2.0585988 | -2.060415 | -0.0018163 |
| CD97 | MARLCIALCL | MARLCIALCL | M63T | -2.0585988 | -2.0560162 | 0.00258255 |
| CD97 | MARLCIALCL | MARLCIALCL | R76S | -2.0585988 | -2.0575805 | 0.00101829 |
| CD97 | MARLCIALCL | MARLCIALCL | T111K | -2.0585988 | -2.0602212 | -0.0016224 |
| CD97 | MARLCIALCL | MARLCIALCL | H230V | -2.0585988 | -2.0575271 | 0.00107169 |
| CD97 | MARLCIALCL | MARLCIALCL | N262G | -2.0585988 | -2.0592611 | -0.0006623 |
| CD97 | MARLCIALCL | MARLCIALCL | H308A | -2.0585988 | -2.0572112 | 0.0013876 |
| CD97 | MARLCIALCL | MARLCIALCL | H358R | -2.0585988 | -2.0561876 | 0.00241113 |
| CD97 | MARLCIALCL | MARLCIALCL | M609I | -2.0585988 | -2.05563 | 0.00296879 |
| MASP2 | MYGRITSPDF | MYGRITSPDF | A87S | -1.7155421 | -1.7135469 | 0.00199521 |
| MASP2 | MYGRITSPDF | MYGRITSPDF | E160S | -1.7155421 | -1.7113492 | 0.00419283 |
| SEMA4A | MPSRALCLLC | MPSRALCLLC | T410A | -1.9508035 | -1.9496592 | 0.00114429 |
| SEMA4A | MPSRALCLLC | MPSRALCLLC | R771G | -1.9508035 | -1.9509084 | -0.0001049 |
| LPXN | MDDLDALLAI | MDDLDALLAI | T386P | -1.4777367 | -1.4750277 | 0.00270903 |
| THEMIS | MFGNECCLF | MFGNECCLF | N555D | -2.8589687 | -2.8591452 | -0.0001764 |
| ANG | MHRAVTPATH | MHRAVTPATH | M39L | -2.902158 | -2.8863695 | 0.01578856 |
| APEX2 | MRLLSWNVA | MRLLSWNVA | C118R | -2.3498039 | -2.3383989 | 0.01140499 |
| APEX2 | MRLLSWNVA | MRLLSWNVA | T119A | -2.3498039 | -2.3450534 | 0.00475049 |
| OASL | MELYQVPAGS | MELYQVPAGS | V108L | -2.6023827 | -2.6033196 | -0.000937 |
| EXOSC3 | MRLRTDVTHS | MRLRTDVTHS | Y73Q | -1.4418521 | -1.4310427 | 0.01080942 |
| CFP | MEGALTSSLL | MEGALTSSLL | E38K | -2.3437529 | -2.3444793 | -0.0007265 |
| POU2F2 | MSVQGQRK\ | MSVQGQRK\ | A555G | -1.2329011 | -1.2357982 | -0.0028971 |
| ART1 | MSTRPQSEE( | MSTRPQSEE( | L157P | -2.843627 | -2.8481534 | -0.0045264 |

| | | | | | | |
|---|---|---|---|---|---|---|
| ART1 | MSTRPQSEE( | MSTRPQSEE( | N330S | -2.843627 | -2.8402715 | 0.0033555 |
| IL18RAP | MFTFCWILLL | MFTFCWILLL | Y55H | -2.2932878 | -2.2934568 | -0.000169 |
| IL18RAP | MFTFCWILLL | MFTFCWILLL | R66Q | -2.2932878 | -2.2931778 | 0.00010991 |
| IL18RAP | MFTFCWILLL | MFTFCWILLL | E102G | -2.2932878 | -2.2961483 | -0.0028605 |
| IL18RAP | MFTFCWILLL | MFTFCWILLL | F296V | -2.2932878 | -2.2917061 | 0.00158167 |
| ATF6B | MAAVAPELLV | MAAVAPELLV | A340L | -2.304239 | -2.2955174 | 0.00872159 |
| CD5L | MEALSGKVFS | MEALSGKVFS | K117E | -1.8917259 | -1.8828468 | 0.00887907 |
| EGF | MFLILTAFLPV | MFLILTAFLPV | S1136R | -2.2972178 | -2.3020663 | -0.0048485 |
| ITCH | MLRGGIIQGF | MLRGGIIQGF | I37V | -1.8512801 | -1.8504832 | 0.00079691 |
| ITCH | MLRGGIIQGF | MLRGGIIQGF | R126N | -1.8512801 | -1.8529317 | -0.0016516 |
| IL1RL1 | MLIHALLVQLI | MLIHALLVQLI | T356I | -2.3146644 | -2.3101265 | 0.00453782 |
| IL1RL1 | MLIHALLVQLI | MLIHALLVQLI | K413T | -2.3146644 | -2.3102388 | 0.00442553 |
| IL1RL1 | MLIHALLVQLI | MLIHALLVQLI | M457T | -2.3146644 | -2.3113325 | 0.0033319 |
| IL1RL1 | MLIHALLVQLI | MLIHALLVQLI | R553V | -2.3146644 | -2.3130927 | 0.00157166 |
| TAPBP | MISPAPVGAL | MISPAPVGAL | K70R | -2.5312877 | -2.5266888 | 0.00459886 |
| C5 | MNTLSFICILA | MNTLSFICILA | G662D | -2.7895217 | -2.789607 | -8.54E-05 |
| CYP27B1 | MPQTLKLASR | MPQTLKLASR | L394V | -1.6213018 | -1.6178259 | 0.0034759 |
| GZMH-immu | MIQGNWSLA | MIQGNWSLA | L74P | -1.8947619 | -1.9038994 | -0.0091375 |
| GZMH-immu | MIQGNWSLA | MIQGNWSLA | F78L | -1.8947619 | -1.8874048 | 0.00735712 |
| GZMH-immu | MIQGNWSLA | MIQGNWSLA | I80L | -1.8947619 | -1.8861737 | 0.00858819 |
| GZMH-immu | MIQGNWSLA | MIQGNWSLA | P83L | -1.8947619 | -1.8809023 | 0.01385963 |
| GZMH-immu | MIQGNWSLA | MIQGNWSLA | L85T | -1.8947619 | -1.8993036 | -0.0045416 |
| GZMH-immu | MIQGNWSLA | MIQGNWSLA | Q89G | -1.8947619 | -1.8907368 | 0.0040251 |
| ERBB2IP | MTTKRNLFVR | MTTKRNLFVR | Q720D | -1.9292519 | -1.9298649 | -0.000613 |
| MSH6 | MSRQSTLLRF | MSRQSTLLRF | A611S | -2.2897952 | -2.2892942 | 0.00050092 |
| MSH6 | MSRQSTLLRF | MSRQSTLLRF | G1065V | -2.2897952 | -2.2860131 | 0.00378203 |
| ABL1 | MSRNEINEKL | MSRNEINEKL | M740V | -2.0794036 | -2.078769 | 0.00063467 |
| SH2B2 | MNGDALCQI | MNGDALCQI | N644S | -1.863965 | -1.8613988 | 0.00256622 |
| BMP6 | MTVTILRFTVF | MTVTILRFTVF | G206D | -1.8195163 | -1.8099753 | 0.00954103 |
| BMP6 | MTVTILRFTVF | MTVTILRFTVF | E208K | -1.8195163 | -1.8137201 | 0.00579619 |
| BMP6 | MTVTILRFTVF | MTVTILRFTVF | V209L | -1.8195163 | -1.8357382 | -0.0162219 |
| BMP6 | MTVTILRFTVF | MTVTILRFTVF | M210N | -1.8195163 | -1.8089024 | 0.01061392 |
| BMP6 | MTVTILRFTVF | MTVTILRFTVF | R211A | -1.8195163 | -1.8200663 | -0.00055 |
| BMP6 | MTVTILRFTVF | MTVTILRFTVF | A213S | -1.8195163 | -1.7981863 | 0.02133 |
| PLCL2 | MAEFGSRGA | MAEFGSRGA | G77S | -2.0108309 | -2.0112605 | -0.0004296 |
| SPPL2A | MSRMRRAWI | MSRMRRAWI | S62T | -1.1768463 | -1.1764709 | 0.00037539 |
| HAVCR2 | MAPYFILQWI | MAPYFILQWI | Q179P | -2.1555367 | -2.1534097 | 0.00212693 |
| ATF7 | MQNDLRNHI | MQNDLRNHI | A13M | -0.9351937 | -0.9271893 | 0.00800443 |
| MR1 | MGLALRLLLLI | MGMALRLLLLI | L3M | -1.9107206 | -1.9129386 | -0.002218 |
| MR1 | MGLALRLLLLI | MGLAFRLLLLI | L5F | -1.9107206 | -1.9184717 | -0.0077511 |
| MR1 | MGLALRLLLLI | MGLALLLLLLI | R6L | -1.9107206 | -1.9149038 | -0.0041832 |
| MR1 | MGLALRLLLLI | MGLALRLPLLI | L8P | -1.9107206 | -1.9134116 | -0.002691 |
| MR1 | MGLALRLLLLI | MGLALRLLLLI | L17V | -1.9107206 | -1.9106585 | 6.21E-05 |
| MR1 | MGLALRLLLLI | MGLALRLLLLI | R18K | -1.9107206 | -1.9179717 | -0.0072511 |
| MR1 | MGLALRLLLLI | MGLALRLLLLI | R21D | -1.9107206 | -1.9060069 | 0.00471365 |

| | | | | | | |
|---|---|---|---|---|---|---|
| MR1 | MGLALRLLLLI | MGLALRLLLLI | L24T | -1.9107206 | -1.9108108 | -9.02E-05 |
| MR1 | MGLALRLLLLI | MGLALRLLLLI | R27L | -1.9107206 | -1.8951089 | 0.01561165 |
| MR1 | MGLALRLLLLI | MGLALRLLLLI | I29Y | -1.9107206 | -1.9066591 | 0.00406146 |
| MR1 | MGLALRLLLLI | MGLALRLLLLI | H31R | -1.9107206 | -1.9186256 | -0.007905 |
| MR1 | MGLALRLLLLI | MGLALRLLLLI | W43E | -1.9107206 | -1.9013547 | 0.00936592 |
| MR1 | MGLALRLLLLI | MGLALRLLLLI | Y44F | -1.9107206 | -1.9053557 | 0.00536489 |
| MR1 | MGLALRLLLLI | MGLALRLLLLI | S45I | -1.9107206 | -1.9127413 | -0.0020207 |
| MR1 | MGLALRLLLLI | MGLALRLLLLI | R46S | -1.9107206 | -1.9006273 | 0.01009333 |
| MR1 | MGLALRLLLLI | MGLALRLLLLI | I47V | -1.9107206 | -1.9058014 | 0.00491917 |
| MR1 | MGLALRLLLLI | MGLALRLLLLI | V54P | -1.9107206 | -1.9088489 | 0.00187171 |
| MR1 | MGLALRLLLLI | MGLALRLLLLI | N57T | -1.9107206 | -1.9122516 | -0.001531 |
| MR1 | MGLALRLLLLI | MGLALRLLLLI | T59D | -1.9107206 | -1.8945752 | 0.01614535 |
| MR1 | MGLALRLLLLI | MGLALRLLLLI | S62T | -1.9107206 | -1.9096181 | 0.00110245 |
| MR1 | MGLALRLLLLI | MGLALRLLLLI | Q63R | -1.9107206 | -1.9069656 | 0.00375497 |
| MR1 | MGLALRLLLLI | MGLALRLLLLI | V65K | -1.9107206 | -1.9166764 | -0.0059558 |
| MR1 | MGLALRLLLLI | MGLALRLLLLI | S69A | -1.9107206 | -1.9068599 | 0.00386071 |
| MR1 | MGLALRLLLLI | MGLALRLLLLI | V70P | -1.9107206 | -1.9022691 | 0.00845146 |
| MR1 | MGLALRLLLLI | MGLALRLLLLI | D77A | -1.9107206 | -1.9151838 | -0.0044632 |
| MR1 | MGLALRLLLLI | MGLALRLLLLI | F80H | -1.9107206 | -1.913331 | -0.0026104 |
| MR1 | MGLALRLLLLI | MGLALRLLLLI | F83R | -1.9107206 | -1.8969494 | 0.01377118 |
| MR1 | MGLALRLLLLI | MGLALRLLLLI | N85T | -1.9107206 | -1.9056084 | 0.00511217 |
| MR1 | MGLALRLLLLI | MGLALRLLLLI | F86Q | -1.9107206 | -1.9022627 | 0.0084579 |
| MR1 | MGLALRLLLLI | MGLALRLLLLI | W87L | -1.9107206 | -1.9073465 | 0.0033741 |
| MR1 | MGLALRLLLLI | MGLALRLLLLI | R90G | -1.9107206 | -1.9118445 | -0.0011239 |
| MR1 | MGLALRLLLLI | MGLALRLLLLI | A97V | -1.9107206 | -1.9095902 | 0.00113034 |
| MR1 | MGLALRLLLLI | MGLALRLLLLI | L105H | -1.9107206 | -1.9134952 | -0.0027746 |
| MR1 | MGLALRLLLLI | MGLALRLLLLI | I114T | -1.9107206 | -1.9071634 | 0.00355721 |
| MR1 | MGLALRLLLLI | MGLALRLLLLI | I115Y | -1.9107206 | -1.9135964 | -0.0028758 |
| MR1 | MGLALRLLLLI | MGLALRLLLLI | M117R | -1.9107206 | -1.9075074 | 0.00321317 |
| MR1 | MGLALRLLLLI | MGLALRLLLLI | I118M | -1.9107206 | -1.9053192 | 0.00540137 |
| MR1 | MGLALRLLLLI | MGLALRLLLLI | T119I | -1.9107206 | -1.9094126 | 0.00130796 |
| MR1 | MGLALRLLLLI | MGLALRLLLLI | D122E | -1.9107206 | -1.9034518 | 0.00726879 |
| MR1 | MGLALRLLLLI | MGLALRLLLLI | Q124L | -1.9107206 | -1.9107933 | -7.27E-05 |
| MR1 | MGLALRLLLLI | MGLALRLLLLI | N127G | -1.9107206 | -1.898267 | 0.01245356 |
| MR1 | MGLALRLLLLI | MGLALRLLLLI | T128S | -1.9107206 | -1.9086511 | 0.00206947 |
| MR1 | MGLALRLLLLI | MGLALRLLLLI | W130T | -1.9107206 | -1.9057889 | 0.00493169 |
| MR1 | MGLALRLLLLI | MGLALRLLLLI | D135Y | -1.9107206 | -1.9173024 | -0.0065818 |
| MR1 | MGLALRLLLLI | MGLALRLLLLI | S136A | -1.9107206 | -1.9094837 | 0.00123692 |
| MR1 | MGLALRLLLLI | MGLALRLLLLI | T144I | -1.9107206 | -1.913375 | -0.0026544 |
| MR1 | MGLALRLLLLI | MGLALRLLLLI | H146N | -1.9107206 | -1.8996967 | 0.01102388 |
| MR1 | MGLALRLLLLI | MGLALRLLLLI | E148D | -1.9107206 | -1.9084616 | 0.00225902 |
| MR1 | MGLALRLLLLI | MGLALRLLLLI | M150L | -1.9107206 | -1.908574 | 0.0021466 |
| MR1 | MGLALRLLLLI | MGLALRLLLLI | I157N | -1.9107206 | -1.9126178 | -0.0018972 |
| MR1 | MGLALRLLLLI | MGLALRLLLLI | G158V | -1.9107206 | -1.9192007 | -0.0084801 |
| MR1 | MGLALRLLLLI | MGLALRLLLLI | Q160H | -1.9107206 | -1.9222276 | -0.011507 |

| | | | | | | |
|---|---|---|---|---|---|---|
| MR1 | MGLALRLLLLI | MGLALRLLLLI | R164Q | -1.9107206 | -1.9171767 | -0.0064561 |
| MR1 | MGLALRLLLLI | MGLALRLLLLI | R165A | -1.9107206 | -1.9238222 | -0.0131016 |
| MR1 | MGLALRLLLLI | MGLALRLLLLI | T168A | -1.9107206 | -1.9070439 | 0.00367665 |
| MR1 | MGLALRLLLLI | MGLALRLLLLI | E169N | -1.9107206 | -1.9143466 | -0.003626 |
| MR1 | MGLALRLLLLI | MGLALRLLLLI | I170Q | -1.9107206 | -1.906243 | 0.00447762 |
| MR1 | MGLALRLLLLI | MGLALRLLLLI | G171H | -1.9107206 | -1.9183145 | -0.0075939 |
| MR1 | MGLALRLLLLI | MGLALRLLLLI | D172E | -1.9107206 | -1.9124248 | -0.0017042 |
| MR1 | MGLALRLLLLI | MGLALRLLLLI | Y173L | -1.9107206 | -1.9121809 | -0.0014603 |
| MR1 | MGLALRLLLLI | MGLALRLLLLI | H175Y | -1.9107206 | -1.9082887 | 0.00243187 |
| MR1 | MGLALRLLLLI | MGLALRLLLLI | R178N | -1.9107206 | -1.9059381 | 0.00478244 |
| MR1 | MGLALRLLLLI | MGLALRLLLLI | R183E | -1.9107206 | -1.9029922 | 0.00772834 |
| MR1 | MGLALRLLLLI | MGLALRLLLLI | P186A | -1.9107206 | -1.9097395 | 0.00098109 |
| MR1 | MGLALRLLLLI | MGLALRLLLLI | R189K | -1.9107206 | -1.9132706 | -0.00255 |
| MR1 | MGLALRLLLLI | MGLALRLLLLI | S190R | -1.9107206 | -1.9066615 | 0.00405908 |
| MR1 | MGLALRLLLLI | MGLALRLLLLI | S191F | -1.9107206 | -1.9085695 | 0.00215113 |
| MR1 | MGLALRLLLLI | MGLALRLLLLI | K202T | -1.9107206 | -1.9142087 | -0.0034881 |
| MR1 | MGLALRLLLLI | MGLALRLLLLI | V203E | -1.9107206 | -1.9071165 | 0.00360405 |
| MR1 | MGLALRLLLLI | MGLALRLLLLI | T206L | -1.9107206 | -1.9175913 | -0.0068707 |
| MR1 | MGLALRLLLLI | MGLALRLLLLI | A207V | -1.9107206 | -1.9018478 | 0.00887275 |
| MR1 | MGLALRLLLLI | MGLALRLLLLI | S213E | -1.9107206 | -1.9157931 | -0.0050725 |
| MR1 | MGLALRLLLLI | MGLALRLLLLI | H215F | -1.9107206 | -1.9176499 | -0.0069293 |
| MR1 | MGLALRLLLLI | MGLALRLLLLI | R217G | -1.9107206 | -1.8910993 | 0.01962125 |
| MR1 | MGLALRLLLLI | MGLALRLLLLI | S226H | -1.9107206 | -1.9093642 | 0.00135636 |
| MR1 | MGLALRLLLLI | MGLALRLLLLI | S243E | -1.9107206 | -1.9038467 | 0.00687385 |
| MR1 | MGLALRLLLLI | MGLALRLLLLI | Q245V | -1.9107206 | -1.9247144 | -0.0139939 |
| MR1 | MGLALRLLLLI | MGLALRLLLLI | Y249D | -1.9107206 | -1.9213105 | -0.01059 |
| MR1 | MGLALRLLLLI | MGLALRLLLLI | S250Y | -1.9107206 | -1.924161 | -0.0134404 |
| MR1 | MGLALRLLLLI | MGLALRLLLLI | E251G | -1.9107206 | -1.8984765 | 0.01224411 |
| MR1 | MGLALRLLLLI | MGLALRLLLLI | V265A | -1.9107206 | -1.9168596 | -0.006139 |
| MR1 | MGLALRLLLLI | MGLALRLLLLI | M267I | -1.9107206 | -1.905153 | 0.00556755 |
| MR1 | MGLALRLLLLI | MGLALRLLLLI | K272Q | -1.9107206 | -1.9119172 | -0.0011966 |
| MR1 | MGLALRLLLLI | MGLALRLLLLI | I273S | -1.9107206 | -1.9040493 | 0.00667131 |
| MR1 | MGLALRLLLLI | MGLALRLLLLI | K274S | -1.9107206 | -1.9168882 | -0.0061677 |
| MR1 | MGLALRLLLLI | MGLALRLLLLI | A275N | -1.9107206 | -1.9136035 | -0.002883 |
| MR1 | MGLALRLLLLI | MGLALRLLLLI | H276L | -1.9107206 | -1.9116987 | -0.0009781 |
| MR1 | MGLALRLLLLI | MGLALRLLLLI | S297Q | -1.9107206 | -1.9119728 | -0.0012522 |
| MR1 | MGLALRLLLLI | MGLALRLLLLI | N298S | -1.9107206 | -1.907789 | 0.00293159 |
| MR1 | MGLALRLLLLI | MGLALRLLLLI | S299E | -1.9107206 | -1.9162861 | -0.0055655 |
| MR1 | MGLALRLLLLI | MGLALRLLLLI | I314V | -1.9107206 | -1.9109806 | -0.00026 |
| MR1 | MGLALRLLLLI | MGLALRLLLLI | V316A | -1.9107206 | -1.9117163 | -0.0009958 |
| MR1 | MGLALRLLLLI | MGLALRLLLLI | I317G | -1.9107206 | -1.9167682 | -0.0060476 |
| MR1 | MGLALRLLLLI | MGLALRLLLLI | I318V | -1.9107206 | -1.9100801 | 0.00064051 |
| MR1 | MGLALRLLLLI | MGLALRLLLLI | K324R | -1.9107206 | -1.9077452 | 0.00297534 |
| APBB1IP | MEQTCDDID | MEQTCDDID | I318T | -1.8484164 | -1.8306996 | 0.01771688 |
| PRKCG | MALVASPTELI | MALVASPTELI | N427T | -0.9590168 | -0.9628937 | -0.0038769 |

| | | | | | | |
|---|---|---|---|---|---|---|
| PRKCG | MALVASPTELI | MALVASPTELI | L591P | -0.9590168 | -0.9546277 | 0.00438911 |
| AQP9 | MNGQSKRTF | MNGQSKRTF | A90V | -1.5127215 | -1.4975803 | 0.01514125 |
| TINAGL1 | MRLLWTLPTL | MRLLWTLPTL | W465H | -1.281088 | -1.2664669 | 0.01462114 |
| CD200 | MRSYEILQVT | MRSYEILQVT | R79S | -2.3815861 | -2.3563397 | 0.02524638 |
| LY9 | MELSRIVGDS | MELSRIVGDS | Q20T | -2.3462796 | -2.3442981 | 0.0019815 |
| LY9 | MELSRIVGDS | MELSRIVGDS | F22I | -2.3462796 | -2.3492277 | -0.002948 |
| LY9 | MELSRIVGDS | MELSRIVGDS | I34A | -2.3462796 | -2.3418803 | 0.0043993 |
| LY9 | MELSRIVGDS | MELSRIVGDS | T42E | -2.3462796 | -2.3471379 | -0.0008583 |
| LY9 | MELSRIVGDS | MELSRIVGDS | P60K | -2.3462796 | -2.348758 | -0.0024784 |
| LY9 | MELSRIVGDS | MELSRIVGDS | G73N | -2.3462796 | -2.3461502 | 0.00012946 |
| LY9 | MELSRIVGDS | MELSRIVGDS | I83N | -2.3462796 | -2.3385322 | 0.00774741 |
| LY9 | MELSRIVGDS | MELSRIVGDS | D184Q | -2.3462796 | -2.3477607 | -0.0014811 |
| LY9 | MELSRIVGDS | MELSRIVGDS | F313P | -2.3462796 | -2.3422613 | 0.00401831 |
| SEMA4D | MLLSNDDKD | MLLSNDDKD | H551G | -1.6654401 | -1.6658871 | -0.000447 |
| ITGB7 | MTGAAWPLG | MTGAAWPLG | R116Q | -1.8194293 | -1.8200147 | -0.0005854 |
| ITGB7 | MTGAAWPLG | MTGAAWPLG | A143R | -1.8194293 | -1.8218037 | -0.0023744 |
| ITGB7 | MTGAAWPLG | MTGAAWPLG | M683V | -1.8194293 | -1.8176336 | 0.00179565 |
| IL31RA | MRNAYISPES | MRNAYISPES | Y675N | -2.636323 | -2.6354914 | 0.0008316 |
| CD22 | MVLIQIASVIL | MVLIQIASVIL | I264S | -2.6991963 | -2.6975543 | 0.00164199 |
| CD22 | MVLIQIASVIL | MVLIQIASVIL | G281H | -2.6991963 | -2.6977344 | 0.00146198 |
| CD22 | MVLIQIASVIL | MVLIQIASVIL | D356H | -2.6991963 | -2.6976957 | 0.00150061 |
| CD22 | MVLIQIASVIL | MVLIQIASVIL | C383H | -2.6991963 | -2.6998191 | -0.0006227 |
| CD22 | MVLIQIASVIL | MVLIQIASVIL | R542L | -2.6991963 | -2.6995809 | -0.0003846 |

cid inserted into the human protein
s
cid inserted into Jonathan's protein

VLSSLTTPFDVTSFDQERMSRSSEGRTMPVRDVTVTSNEAFLSDNGLISSFLNEQWTPSLKSFTVEYRSDNS
NSSLNNVDHHLLHHHNELNMANNATSAAATGNNNNNANTSNSSSSSSESALKEGGSSSGLPSSSSSSSN
IDPDQADITYEGRQKMTSLSSCFAQLCHKAQTVSQINHKLEAQLVDLKSELTETKAEKVVLEKEVHDQLLQLH
AEEAYGAEVQDYLNTPTAPFASTIGSHNVTLRWKLANISGVTYIIQWKYNQLPGDWRYAEVVSEHSYTVSNLQ
WKTYEKVSSADGTLVGYSEKKKGMWGICIFIAAVDLDEMTFTFTELLKSIGMRVCTCFQLLREMDVNMDTIS
LCQNPKLQLKNSPPYILDILPDTYQHLRLILGNCDDNQKLAQLSENEYFKIYIDSLMKKSKRAIRLFKEGKERN
SSQPSPCDPKSGGHPPKVLPGPGGSMGLKNGSGNGAKGKGKRERSISADSFEQRDAGTPSDDSEIKDCN
LSSQSTENVYETAKSVEVDVFGDIELKVIMNISETISCLWVFKENQAGCKPSLDSENGYVVSLAFSKIKETQAC
LENETSDMKEVLKHLQGKLEQEARVMVSSGQTEVLDQLKALQMDITSLYNLKFQPPALAPELACPEGSPVHS
SFADTIKVFQTDLQINVVEPENTERAANIRNQILFNQNEGPETVQCEITGMDTLLHAPIQTPVQQTEWHDTDI
LRGCLNSEKICIVPILACLLSLCLCIAGLKWVFVDKIFEYDSPTHLDPGRIGQDPVISVEPTALSAWVPSVVHTS
RTIAIPQPRQRSLSESDTSCLEQECFIEPYATVTELKLGPNWKHDDSERSPVQSLEGSGNLCEDENMDTYLS
RLLPDFDVQEQDVDTLHGSVHVTMYGTPKGNRPAILTYHDIGLNHKTCFNPLFNFEDMQEITQHFAVCHV
GNRSVAAIDSFAMKNCSSGSHSGSKMALQGSFSCWNGTEIKLGQACDFIRDCAEGEDEGDMCKKLPPC
SWQLFQTLRLSELTLERNLFNCSCDIRWIQLWQEKGEANLQKQELYCMNMEAAVIPLQEMNITQCDLPEI
LPLPGPHLMPVKMDPGSRELPGRQSDPYEPPDGEPGCDDPPGEPPLSRKTATFKSRAPRKNPAAGEEWSG
LLFAEFLECCSRLTEEHPEVIPFLASRHQKASAEFLASAEFRNVLGRCLTRVQSKRSKVYVYINELCTVLKAHTQ
QMRPTAKFGPYQKALREIRYSLLPFANESGTPASTEVNRQMLQELVNVGCDQEMAVRALKQTGSRSIEAALE
LPNPCLSAPCKNAGTCHPVIRGNAVDYTCSCRVGFTDRLCLTPVENACLNNPCRNGGTCDLLTLTEYKCR
QTNQLQYLLKVVLKTLWKHQFAWPFHHPVDAVKLNLPDYYKIIKTPMDMGTIRKRLENNYYWNAQECIQDF
KVEGTFSVPAADGQPLAKETQGKPGELLSQSLGTQSGVSKKKKKRKRNKSKKNPSDSGMQDSLTSVSSAPC
ERNHPRVYQSRDFRGHNRGYRRPYYFRGRNRGFYPWGQYNRGGYGNYRSNWQNYRQAYSPRRGRS
QKVSFSFSLTKKTLQNRFLTVLGNEKQNETQNSPATPLPTDFAPKIKMDLGDTSGSAEESSPPKPKVELGKIHF
NPKTSVSELFAILRQWVPQVQQNIDIIGNEIIKRGCSVNDRDGLTDMTLLHYTCKSGAHGIGDVETAAKFASH
SQTAIGYTAPLMLLSPAMINGGSTSALVVHWEAPAGQRDGYLVSVYEEGSLAAGRRVNVGKDSTNVSLAGLV
TAGQDEYTILPPSFSIGIHGYLLVYSVTSLRSFQVVKTLHNRLYESRGKTRMPVVLVGNKADLSMESREVKTDI
RRLQDLASMISAEYLNGAADGPGASRAASPRPPEPAALAGPQGDGGAAAGSPQAPAELGRRSLAGATCCEI
KPEFKMSKKINGGSVVEMQGDEMTRVIWELIKEKLIFPYVDLDLHSYDLGMENRDATNDKVTVEAAEAIRKY
DRRLTPTKGICPKQENIAGPSRPAKLLGAPIAASSHSHTSVITSPLRALASLPPCLSLPCCGARAVSVGGTQTE
GSSGAAGSGGGGAAPGSGGSGGPAAGVGPGFDAALQVSAAIGINLRRFRAACGSEAGSGEDEQFLGFGS
PMSESKLNTLVQKLHDFLAHSSEESEDANSPPRLSMSKSIGKSSGPENSPTPMENNREEGNSSSERSKSLGS
MECLRSREVWLLEQVDLIQQLKEETLQQQTYQLYWLLGQFNCLIHQLEHPHTNDLVNQISVCLERLGSLTLI
VQGNLEITYLGPDADMSFLKDIKEVQGYVLIAQNQVSRVQLESLRIIRGTQLYEEKYALAVLANSDPTGTVGLC
CPLCKTEVTKRSLQENPRFKQLIEGLLETIHAFELDTGIKFLNSQDFSKKPVETSSTELLWKEGSIIQSKGYRNF
DQIKTWVASNEIKDKRQLIDNRKLIETQMERFKVVERETKTKAYSKEGLGLAQKVDPAQKEKEEVGQWLTNTII

PGILHFLQHEWGRFEAERAEWEAERAELQAQIAFLQGERKGQENLKKDLVRRIKMLEYALKQERAKYHKL
KQQEKKPPLSSSVRFLRSSVTRTFSGTSWVNMEKTDILFHNQDSLNVNGFTMSLRSRSFSHRLSQTAVITK
EAAAGPGPGPSPVPPLPAAAPGPLSLLDTCAVCAQSLQSRREAEPKLLPCLHSFCRRCLPEPERQLSVPVPG
ASSDTGSASQSCFLNTESSREAASATKASAQEPVSFSKPQTKKSGAGSTWSQLSTNNKELAVCSAVPSPNN
RRPKLLPCLHSFCLRCLPQPDRYLMLPPVAPSPSPGPGTPKELQPPASPSSPAPSPTSPLHCTPVGVIRCPV
IQFFVIEEYTGPDPVLVGKLHSDIDSGDGNIKYILSGEGAGIIFVIDDKSGNIHATKTLDREERAQYILTAQAV
HVRRLQKALREWAANPGLFSQPMASLPVSSIPLFKISESGGRKSLSNGHASPGEAPGKATPPACSPSELGE
PPLESSTQEIGEELGDGVIYSISLRKVQLHHTANKGQRWLGFENESALNLYETCKVRTIKAGTLEKLVEYLVS
GIGAFAAVLAYWFASRPKALKPPCDLLMQSEEVEGWEGARRSVIGGSTELLTHYYDDARTMYEVFRRGFN
MMEADEKEGLRCQVPDSTAEALIEWLQSQMTNGHISGNGDIYQERLARLENDKESLVLQVSVLTDQVEA
PQGSKLAAGGGGEDGVLEEKLLTEPMDSSTHLDTTGSLSQVIDRLPQPPKNNRVPGMVAAAAEGGGEDA
AVNKIYVLNENLQNISVYKTGPVLENPDCAPCEDCKQRANLSNSFWKDNVNMALLLETYYDDQLISCGS
EDPRQGRNPPKFVTYHKALQEIRNSLLPFANETSSSARGTSEINRQMLQDLQAAGFDEDMVVRALRQTNS
PPPRRDGAPICLQLATGIRTVLLSKAGEEQRPTETPPAQAPPTSDPSASCTSKGVYENYRRWQRYKALARRH
LLQADLIENVGSPVTTAVLTSVSEDSRDQFENSVLQLREQDEAETAVPQGHSNTMDGENNSGTDDIKIQFS
SPEPEQPSSNKWQLDKWLNKVNPHKTSILNQNDNHGLEGNQYYSSIKDEGQECEKIPAICQTELSDKEIK
HGCKELSKGCSSQNYTPAKIDWSPIIWVNRSLPLWGLQVSVALISLFETMLLGYLSYKGNIWEQVLRIPFL
AGGSGLSGGQPAAAPRKCSSRRNAWGNLSYADLITRAIESAPDKRLTLSQIYDWMVRCVPYFKDKGDSN
CCGCRFPLLVALLQLVLGISITVLGFIMAGISSSVLVRDTPYWAGIIVSVVALVGMVMLCVSYQPEEKTCFQF
FWLGYERGSAKGEAAAWPVIITDKAVMPVQVIDGTPENLLPSHLQKVDLKYLQMGCNQANKYFQGTVILSY
FQQKKIATEHQIYGNKDSYFRNMEEKMKKNGIILKDELKTLLHLCKTPDDVELAKNVIYRYHTENRNVAFGE
LDICCRDPGGNGYACCAPSLGSPHSLPMIQASSVAKPAGTVCPDGTQCPAEYSCLRTPAGSFSCCPLSE
AGKTLVAELLILKRVLETHKKALFILPFVSVAKEKKYYLQTLFQEVGVRVEGYMGSTSPAGRFSALDVAVCTIER
LDYIEVVDYQGRNRHTVIQGRQVRHLYSLTVFENYLYATNSDNFSILQINKYNRTDVQSLIRLENAREIHVY
NVRTTHYTPLPSGCNPLKRDIAEYVRTGFINLDKPSNPSSHEVVAWIRRILRVEKTGHSGTLDPKVTGCLIVC
EFRDVMQALLLRPLVKEPAVSRLLRIMQFLSRIEEGENLDCTFDTESELTPLESAIGVLELIEREFPVSEVAME
GMQRLEDEKRWLSVPKVYVLQPWILNLLMKYEQLDSRNTQVPGHILKVVNDLKAADQGSAAVLHISDGSY
LALALHTAVSASDLQLHPPSYAWSHNGLLASLDHSSIRRGYQVYKQVCAACHSMDYLAFRNLIGVSHTEA
SPDDSDSVKSDDEDGDIKPTKGQANQGKGSDDGKDPRRPKRPRTILTTQQRRAFKASFEVSSKPCRKVR
GEGDIDIMALKRMLEKLGAAKTHLELKKMITEVTGGLGETIGYRDFVRMMLGKRSAIFKLILMYEDKAKEKE
FQSVVVSTDHDEIEKVAKKFGVKVHRRSAEVSKDTSSSLETIQEFLRHHNEIDIIGHIQATSPCLHPGDLIK
IMRIIKNIVKTFKYIYGLRFEVKGLENFQIERPCVIVSNHQSILDMMGLMEILPDNCVQIGKKELLYTGPVGLI
NLLPAAVGLGQANPNVEETGEVQGQFLTVVGAFWHLTKEGRCTAPVRVALLNCMKALLEADPHSNWAVL
LWVFQLQVPCEEQTRAAMNTLHSDGASHRPTNGTKSDTGQEDFLSRLKQFLCEPPYSSSPEDSRCRLCP
KNSSEILLGSKNKQSLQQQAKSIQEQVKKVEVTLEEDIKTMEMVKSKWEHFGNNFEALSIWITEKEKELDTLE
QAYVSSLIYGWDREYFWTALQDINETGAFRWLSGDEVMYTHWNRDQPGYNKGGCVALATGSSIGLWEV
ALSKFNIDLIVSLLRQENASNICVIQVPPEMRYSDYFIIVSGSSSRHIHAMAHYMLKMYKQLKQENDPHIQI
YNTLACDWWEEINCTNITKTECNFTDPDIKRFWTITLRVRAELGQVKSDWVETNKFVADKNTTIGPPVIVN
PPEPGYLLPSAACAPLQLRALIRHGTRFPTEKQIRKLGQLHRLLRSQERPCAAAQQLAHWDMWYQPDMD
YLSQLTHDKDSMAHSQSTKLDQEIKKVKESIQEGVNYMGNLLEKLAPLNRGLQPRFYQDSDSLRRLIRKEL
QNHAFSGSSPPYLTGSVPININSSSNKECLEENNKYYAHEASSKQEERCRQRLSFANDPSITHPTNLKPVK
LTLVNSNMLASDDKSLESAVVKVINPDEQCDVNLGLQASSSSLVVKEILQEAPELITQQLAHLLRGSILFQCM
RAAAVVVRLQGEGAADWSGGENGAGLQGLPHEEPPPQQREGAETAPSGDAVETAIGYRRSQCSHRAAE
INPCGGIEDCDGSSAICAYDLNKKVYLSVGDSSLKRSSKSLLEFNTTKPCSQQGAQHVVQSNINFLCGKT
TQLAKVNDVLKIDLSSLVQKGYAAFNDTPDLLLNQTRNILSDIKNLLDLVGSNITSFTQMLPVQKILADLTIYL

DGSCLDIVDKVISGNVEEFRKDFISRIWLTYREDFPQIKDSALTTDCGWGCTLRTGQMLLAQGLVLHFLGR
RKHFAWRKPILGRFEPDWKPFLTSKELFWQDDTPARRQLNDLLHLHRLPFGLNGSVLHAFHTLQLLPQQG
PKVLDGTRCYTESLDMCISGLCQIVGCDHQLGSTVKEDNCGVCNGDGSTCRLVRGQYKSQLSANKLED
LRISNDGKVDGTSIQSPHSLLEIRAVAVRTVAIKGVHSSRYLCMEEDGKLHGLLRYAAEDCSFEEEIRPDGY
NTERCLSENPSLPENGLQVASQTVNRSAPNSNTALKLEEKNSEAVKFPGIQSTEPSMFKNLKQANINEGENG
LRTLVLALCAAGLQGNICTTRGVNSCRQCLAVSPLCTWCSKEDLRQGTPRCDLKTNLLQNGCTQDFIEYP
QPQRRRPSLLSEFQPGNERSQELHLRAEPHSYLSDMSKQSEIEFIETKRPRLDLLQEPLLRHSPLLNQNQQ
ASRRGPMGIGPHGGGRPMLRARHAVSGNATLPPFPEGMQMAVFGMGCFWGAEKMFWSLPGVFSTQV
QCRASFQRRPELQKNTVLCRVVEQFHSTQPAGPAEEKRGWDGSRSEVACDSCLEAAAAAKTCLTCMASF
YEEAREIFQDDKRTMEFWQNYVEANQCDSDPCQNGGTCVDHFQAYVCLCPEEYKGKNCEKGLNEKPT
ADKELLVKLRKKTGYSFVNCKKALEKFSGDSKQAEAWLHEQAQKEGWSKASKLQGRKTKEGLIGLLQEGN
SGGRPHYYVSYRRSAFAQMKLPKYSLPKDMHVISTDENQVFAAVQEWNQNDTYNLYISDTRGVYFTLALE
LFLEPGSRCKKGGSVGKEACLSQSYETCKIRASEPCHSPVTNVSARSSKTEYNDFAVCAKGEKDTSDKNGN
FRFRYVCEGPSHGGLPGASSEKNKKSYPQVKIFNYVGPAKVIVQLVTNGKNVHLHAHSLVGKYCEEGVCTV
PALTHSSQSPTNLPAHTGSSDCNISCKGMTERIHSINLHNFSNSVLETLNEQRNRGHFCDVTVRIHGSML
ARTPLSLLELPPELLVQIFGSLPGTDLPNLARVCSTFRRILCTDIIWRRRCREEYGVCENLRKLEITGVSCRDVY
PVHYTFRTDCGSPGPDLCSFPIANISLVKNNRDRVLMYGQPYRISLELELPESPVNQELGMFMVMMSCYT
LCKQQAEEEGASVGSQEPHSPGCCHHGNTDPSHSPGGTCPIAQAKTRLESQEGNHAGPDGKEESTGAC
YAFQARLGGTCIQAQLREKKRAQEMYGDALAGGQSSFLLQQEGAGGDVFSCSLGNLPPGEEAALTLFYVC
SEERAKPAAQVEFADIGCGYGGLLVELSPLFPNTLMLGLEIRVKVSDYVHDRIQSLRASHPGQYQNIACIRS
GVEFKEEWQDEDFPRPLPEDDNVEADILAVTGQQGQHEVNGNKVRKKLMAPDISLTLDHSEESVLSDDLN
VTELSAAISHLIQVFSSSFQTDFQPAKILRSPSCADVVLDPHFSFTVYAAHNIPAAWLKSYKLFSFSCSLTYGG
CKCGTHTGQFECICEKGYYGKGLQHECTACPSGTYKPEGSPGGISTCISCPDDNHTSPPGSTSIEDCVC
SHLVSSCPGLIRDHKYHLRHHRHCCSGKELVDRLLSAGLSIQTRGQAIGVCQVLVDGGVLTHVKQEWHF
FDMTIQCIWSVYTNKIISSDRDLLSVVFYGTEKHKNSVDFKHVYVLQELDNPGAKRVLELDKYKGDQGKAL
YQRFENYEEEEEEEETDWKEISGCQNVTVLECDFSSAMTAYLDSYNVRIRAEKREEKSLWSTTLSFVPYSVA
RFLWMGVGNACNYYNKMYGEFMRVWISGEETLIISKSSSIFHVMKHGQYICRFGSKLGLQCIGMHENGI
AQHHGARLGVAAPVWEAALTLCKYFEKQKLNFWGKKVIELGAGTGIVGILATLLGGDVTITDLPLALEQIQEN
YLRMRKLLDASDLETEAKMEKEMNCFYLKALDGFVMVLSEDGDMIYMSENVNKCMGLTQFELTGHSVFD
PREEPLIDRLEVVTLPSPLPKGLPVKNCAQQFQLPTHDWPDTGPSGQFDVGVVASFGRLLSEDLILQFPYG
LHCKGDEITVTLGAHNIRQQEQSQQEIPVYHQIPHPHYNKTTMNNDVMLLQLVKTAKLNRWVDTIALPCAG
RFRDMRTTTNLYLSSMAFSDLLIFLCMPLDLFRLWQYRPWNFGDLLCKLFQFVSESCTYSTILNITALSVERY
LKEYNVGGKVIHLVERAPPQAQSPTSGGSSGASSASAPHNGIGGRGAGATVHDRNANSYVMVGTFNLPVS
YSETDLALNGKHGDAHCKGFINNNTFPAVVIFIINLSTLESCGSILVVSSVPGINAYGNVSVVQTGNVSGYID
QFSVKKTTEPIQPRTLLKFPNIYGPKPVVTAPEIVNYVDYALKTTEEPTRGKDDIQDQNRMKIQVNEELFVLPQ
SAAVTHVVSEQNSGDEVARWLEQQREECGSGGDPALLDISWFTESMGAGRPVEIESRHRLKVTALAPLRQ
LEDIAVVHKTTAFTYAARPPPAPRTPPSPYSRSCPPQDPPAGPHWTAGAQGAPAEAAGGQQEEEEEEEKNM
LGAVLPGDAAMKTPGPALSPRPGSCWLENPFLGGHSPLPAWVEETRARLLRPLPPPAPWLSSLTLTPCLASS
FVTRKNRQVCTNPDTKWVKDYVNFLEMK
IIHCKSCTLFPPNPNLPPPSTRERPPGCKTVFVGGLPENATEEIIQEVFEQCGDITAIRKSKKNFCHIRFAEEF
FVSFDSRSEAEAAKNALNGIRFDPEIPQTLRLEFAKANTKMAKSKLVGTPNPSTPLPSTVPQFITREQYEVVLS
FHENKPSRKTLHNSIELLRKYCMGDHPVQFEILSDQSVNFGFSKFPWVNYQDGNLNISIPVFSIHGNHDD
PVDIRVPFPSLQSVKVFLESHHIEYSILLEDLQVIVEEEKQEMVNNQRRERGTSNFNYGAYHPLESIYAALDD
QMKFSCCYGSNIKNLFSTAVNKGIENIDACLSFLAAKGINIQGLSAEEIRNGNLKAILGLFFSLSRYKQQQQQ
LSKNRLKSVNPEEFTSYPLLEEIDLSDNIVANVEPGAFNNLFNLRSLRLKGNRLKLVPLGVFTGLSNLTKLDIS

AGSIIMNREIESMAMRPLAKDLTRSLEEVRNIIRDQALRDLNLYTEKMKESLKHFDVLFAEFELSYVSAMVPV
IALCEGLMNEMNQALDFFEKDPDVGAIVITGSEKAFAAGADIKEMQNRTFQECYGGSFLAHWNRVSLVRK
RHIVSFPGLPLKMLLLLLFHIAFPLLPGAQTGEIIGGQEAQPHSRPYMAYLNIRREDKRFRCGGFLVAENFV
RHDPLPASCRRRGASPKAAEEEEQSPGSSQCLRRSVAFFAVCDGHGGREAAQFARENLWGFIQKQKGF
RATVEAAVQECSQSLDETIEHIFNIIGAFDVPRYVYNSERKKFVPLSMTNHPAPNLFGTARDKAELFRERYTIL
QQMWVYDEDEGISCRDVTFVPGLYKIFDEILVNAADNKQRDKNMSCIKITIDPENNTISVWNNGKGIPVVE
DMNNDYDCITQKKRKGEKRSNRLTSSTRVLQQHTKYSKCGCPADSDGTEVEDSSEFGTLLLDSDSDDSV
ASSASMPVSRPARMTAASSGSREGAASGALPSGTEASTVPPAASGDSDPSPTLKLGAKSNCIIVSARQRGN
KRSRLRDLLVAASSSLHWPAVANRGQWEVRSDGPADGAEIEKESCGDPGTPLYGIREGDGFSNRDILRF
ALAGITDLLDGFIARNWASQKSALGSALDPLADKILISVLYVSLSCASLIPVPLTTLIILRDIVLIAAVFCVRYKTLS
VSAHRETDASCSLCRVAHFAWPVPEAKQHRMSLKAVTLALALLAITGAHADVSAQQVADVVWNYFTQLTG
SRSGSKISCKVELEVPQLCSFILKTSECTLREMSGVDAEGKTLLKQSKNSDDFARAMAQYELKFSTQDGKK
FDEQLMEFHCLWDDSFPERPERLAAVKEKLQQCGLWERCVPVEARLATEEEILLAHSQEYVELMKSTELM
SPACHQPLHLVGHHHSITALAFGSEINPLLVCSASCDYVIVWNLDECTKKALQGLMPRGIVIGTLLGMVLY
LSSNAVLCWKFCHVFHKLLRDGHPNVLKDSVRYKNELNDMSRMWSHLSEGYGQLCSIYLKLLRTKMEF
NLPVMIWIYGGAFLLGGGQGANFLNNYLYDGQEIAVRGKVIVVTFNYRVGPLGFLSTGDASIPGNYGLKD
CVVCVPRGARAIPQPLSFQQLSSESEVVIRVIQRLCEKKKKNVLAFGFSLQELMWKMRVNDCTWLRVIKGD
VIQKEGVRSRLTIYNANVEDAGIYRCQATDAKGQTQEATVVLEIYQKLTFRDIITPQEFRQGEDAEVVCHITS
EQPLESLDSMDETLQIFQRILPSLPMENVPVAQIFNDIENLRSLIQTLGSHLGCTFHKSSNLDALGNLTELLT
INPTSCNNPKLEELQMLLIDWINTTLKQEHIVVKSLEEDLFDGLILHHLLEKLGSLRLDVEKIALTEKKQKHL
FDNSQKKIIHYSVNACLFPICALHHVAVTTVEGIGSTKTRLHPVQERIAKSHGSQCGFCTPGIVMSMYALLR
VGTDSSIFHMGQGSQPQHLQLRATLRLKSGDCVPRSWIYLLNEGSTDLATEGRPGMRTELFSSKCPDTII
MDDNRITTVEGNDEQHYKTFNGLKSKNPGLKTLLAVGGWNFGSEKFSSMASTPGNRRTFILSVVEFLHKY
TEREEVRGRELTGLKGLYEAELADARRALDDMARERAKLQIELGKIRTEHEHLLGSYAKKESDHNAAQIKLRE
KIIDIIDTTGSGDVNTSTIVEPKDGEIVGLSGRTLKIPTNWVNPSGKYHIGIKNGYDFYPKALKERIQKERKEK
TGWLVNMHPTEILDDDKRLVSSVDYYFIQEDGSRFKVALPYKPYFYIATRKGCEREVSSFLSKKYQGKIAKLE
HLEFIHDHADYGFETPNVKFNWRVIKEKRDAYVSRLNGIYQNNLNKAHIETIRGHAVFTSDPEPTVEVNGK
YIYNRQQHLHFDSDLGVWVADTELGRATAELWNSQPESLQYKRAAVDRFCRHNYQETEPFTVSRRVQPK
PLNIMYLLKHRWVLGKPACQFWLSMDYVASTASIFNLFILCLDRYRSVQQPLKYLKYRTKSRALVMISGAW
KSVKWKLGADSEVWVWVMGEHASDKPYDVICEEIQAWRAQQEAGEARGRQTFLELAAPGSLHSQPGAL
QKIINQQQHSTVSSLQKNNKKSTSLQISLQSRRYSDCFQSDTVVASEDTFISVGNEGNACNRNMADYRS
ELVSPFYNETLPERSFYRKNSSGEEPLIESLEEEINTIRVSNHHLWMRMNNITLAAGNPGEKGAEGLSGPGG
VAIKKISPFEHQTYCQRTLREIKILLRFRHENIIGINDIIRAPTIEQMKDVYIVQDLMETDLYKLLKTQHLSNDH
PYVECRNKPCGHPGDIQFGSFELTAGTEFVFGARVQYKCDEGYQMLSQRNFRECQADGWSNELPHCE
KLSPRGQINCSRIIRGDVEAVEEALLSRLTTKTKAEPLAPKDYVNLTKDCTHFKTRRRFVEFPMSKEEEDFPIA
SLDSAILIGETGSGKTTQIPQYLYEAGIGRQGIIAVTQPRRVAAISLATRVSDEKKTELGKLVGYTVRFEDFTSE
GLPLNVAHSSLSTPCHGLQTSNPVISIVPSTNHPSGYGGHVDGGASGYYLSPNVRPNGAPALESPRIEITSY
KLTYISKVSDATKLRPKAEFCFVMNAGMRKYFLQANDQQDLVEWVNVLNKAIKITVPKQSDLLCQPDNPNR
GGNNPHTIPTAAVSGAFSVQVPLSGRSQDDATVGYFFQRQAGEQLSGYSSKHRWPTGDSIHVDHSVRS
SRIFLSIQAKGGEYLNIAPKRRVHLTNLGTGWISVSMTELQIEDSGTYLCGVLDHQTIILLKMIKVVVSYDAPM
RKGLAHPEDGITSTLRRENELKAVITTLQNASFMKAAAYMTVSKPINSPKLKWTNSRILQNITYNSKGDPVIH
EGAENLRKVTTDKKNLAYVDNILKKSNKKLEELHHKLQELNAHIVVTDPEDVTECPRTPDTPNSDPRFSTNN
GADIISIKNKEENTFIQETFKNHWQGPDYISLGMFFDTDDNAFKWFDKSEVKFTNWIEEESNEELLNTCAS
VFHKAIQYSKIYRSDVSSTVWGHSDFRLPPLYAAFVNGVAVHSMDFDDTWHPATHPSGAVLPALLALSEVL
NHRGLEGLKRNVTLQNIIDRFQKASLSGPNSPSESRRERTYRKSPAMSSERIACQFCEQDPPRDAVKTCIT

QSNAARDFLNYLVRLNEIKSEEIPALGAGAADAPDGIVPEARGDAGERVSALGGPLPPHLALKSETGGGLAP
VTGFGMKRTVNTVWFLNLAVADLLCCLALPFLAVPVARGNRWELGVFACKLLPSLTILNMFASVLLLMGISV
LSRIHQHKNLLWFMPAILEDSGSYECVMWSLTSCNKIYFSVTVFKNTAGLCFNGDFLYAQEISASSNEKIVC
LDLMSWDSGPWDVMSHTCIDENLKEEPQSPATSVCSVPSPLSDSSVQHVPEDLDFSTSSQMSPLSLYSK
VNEKGQLSAALLVKLNKVNGIISSTPPGNNAVAKSKKSLLSVNVWNEDNSSKNLIPRLQKVKNNYLAMNKY
NETLWKWVSQNRLFHLGSRQCLGLNIAKPAAPLKMVNCDSQLMLWWRCVDASIFGASQYKLTLKDNVV
ELAERLQREEEEAYASSQSTQGAQSLTFSKFEEKKSNEKTRKVTTVKKFFSASSRTGAKKEIHEAKAPSPSINR
IRDWTFQILVMLEMSPRRGGIYTCQVEHISLRGPLTVHWASVTLQLFQINVFHNASSTDVATMALLGDLETH
YPFWEQRPQVEQHIRGPRAAVPIPRGSGHNSWGTAGSFRPPYISQQTRSLSSYPPAQYNYRKSEREFDH
MDPSVVARSMVAFVEGMTCNSCVQTIEQHVGKLNGIHNIKVSLEEKNVTIIYDPRLQTPDSLQKVIVDMG
VSILERFLQAVDPLYVIQNFREELQKGLFHPDDSVKLLTIAQVGRIVENSDAVREILNNPELLRQIIYCIGGE
NQKPNLEKNLKLTNIVEKFNSLNLEKTPSVLHCVFCRRGPPLPAQKICLRCEAPCCQSHVQTHLQQPSTA
RFSSSRCIIVLLSAELVQNFYLPDVLGSLQEALKPPHKVVKLFCGVEDCDDYSGFFRDWAQWKELTYDDE
GRNKTFLCYVVETQGKESGTFRGYLEDEHAAAHVEDAFFNNILPTCDPNLRYNVTWYVSSSPCVPCAGRI
EREVNILKEIQHPNVITLHDVYENKTDVILILELVAGGELFDFLAEKESLTEEEATEFLKQILNGVYYLHTLQIAH
KVKEMLDSETLCRRAVKILKKKKLRRIPNGEANVKKEIQLLRRLRHRNVIQLVDVLYNEEKQKMYMVMEYC
GGGRRRDSLESGHQALQAEHRMGGGKQHRGPFQGVRVKNSVKELLLHIRSNKKMSSGHVADEFKAQA
GYTYRDYQEYNPSYDDDYTEECPVYGNLNQEILDECCYEQMKAQPRRYANDVKVDVESQMCYASLDHS
QAGELPTHFVSRTMKWDRYAPVQLVPHVEKMQQEGGQRRKRQVDGCPALQLQAIVNSEPNERSLSPW
GRCYNYITTVNPELRDKDCQNIWETMQKAFIYKKPCNITTEDYQPLLDLASHPIPCNKSLFWSKTSDLVHR
NENKYKKSNVLTVETINFLKGNLASAQEIDTKYGQVSEDLRRRLVTLKQKVLIQVNRIKALQNNIRDQVVEM
KEELDLVIDVLENIRDPNLSEPLGRPLEILRHIREDLKNCIRHQPDNHQRSERLTSWLQKFHATKKMETPG
RHLETRERAKVAVLVNRVHLVEQHYTQEFHALQDRFRVTPVSSNSDQKFFFSKEVKLSDVVICTAKILQNA
STPTYTLLLITNAAVAMEGTKFCTGNCQGKSHQGSTNLQVYSLPDTLQLETQPKELVAGQPAHLHCSISKVY
RTSVTPSNQDICSSSAVFSECCHHSPVQSAVVLKNPHCQNSLTQGVTVTVICQDTLQASKRNSHGQDW
AQSKLQAVEEEVQKLQVSLQPENASDASAKGSDSVQLLDEAQAGIQMLKDLLGNVSADPVQIKIRLDISAAR
LRDCGHPNVVAYFGSYLRNDRLWICMEFCGGGSLQEIYHTTGPLSEKQIAYVCRETLQGLSHLHAKGKM
KCTKDECDTYFKVCLKEYQSRVSAGGHCSFGSGSTPVIGGNTFNLKYNRNNERNRIVLPFNFAWPRSYTL
FTCMLIGEIFELMQFIFVVAFTTFLVSCVDYDILFANKMVNHSQHASELVKVTLPDAFLPPSVCSARIQQNGF
INLRPQMSQFLCLKNIRTFLSACCEIFGMKKNELFEAFDLFDVRDFGKVIETLSKLSRTSVALATGIRPFPTEE
VLKQMTSCSLKTCSKTDIQSLFRLTQTGMNGPETSADQLQLAEVKAKLGASQRSISELEKERHQNQQECE
ALLRGVCNFMEEIQHVQSRYRSSWWRVVVACLSPSLHRHMLLLLISFCAYMALFGDTGLQLFLNLILLCL
FTLAQNAEVLANKSSCGKENVSAPILAIGFGAGHLLQMSFTKTSKLYSVDELTFIYNLSDTTVFPNSTGGER
EDALAPAVKAALTQFPSISHPPVGPAPSVGYNIDTIHPEVLAPEASRTFGYQVLQFKEASIVVGAPGDQNSTG
LCNKEEAREIFENNPETEYFYPKYLSCLSSHRAGIFHVTYVISDSSTDLRNCVKEISNQCSPLPCNEDGYME
LQSRHEISPGHDGSRNEAQLQEMAQLRIKHQEELTELHKKRGELAQSVIDLNNQMQQKDREMQINEAKI
ERTRSEEDNEMNLPSLAAAYTTILRSLGEDPERQGLLRTPWRAATAMQFFTKGYQETISDVLNDAIFDEDHD
AHCNGDKINVTLGAHDITKQERSQQVIRVRRAYPHQDYNKKSHNNDIMLLKLEHNALLNEYVSPIPLPMA
METSHRRYAEKLSEVESDLKKLDDQTGQKDMSTNKELSSFRSDILALRQQLQEIAEKTTKNKDTLEKLQDSG
FTDTMETCDDINECLGPRLPDCGRNRNCNNAPGSYYCTCINGYEPSSGKANFTHASENSCQDIDECK
TLCGRESTDTEEAPGNRTYHSIDNALTVTFRSDYSNENPFTGFEAFYAAEDVDECELLIDDEPLCDHHCH
GARDTLLSLRVDSAGSMELTGSIVWQPVPEKKQECVFKKKSNLTECFNFIRVLVPVNQTRLYTCGTYAFSPT
STRLPESDDKNVYSEVPTPQLSLPSPPPPTAAQQLDDLLSHLGQMQSKISVVGSTAEPERAKGPSAVGPSPG
EITRKVSIGPTRLGHPCFYCNEDIKLANFTIKQGEQIMFNSVEEVNGTLTVNCGVVRDNQSHSFTLPLSQEG
ERFLLQHFDAKPKGRDDRYCNNRMRYMMKEDRRRDRNPGPKVCKETNTFIHGNSDDIKAICTSHGG

RSGYSGVATFCKDSATPLAAEEGLSGLLARPGEGGAVGCYGDLGAFTPQELKALDSEGCTVITHHRIRTSAQ
KGTALKNNSDADLVLFLSCFSSYQDQMENRAAVLHTIKQKLNRCRQTVAFSIDVEVSQPKEKGTCPRSLSIT
VLPGDVLLLPSYPEADGERLWLGPGAALPGRLVCGPGLRSCGAGLLVTKCGLLRHRQPGGAGGGSGGAY
RPRAGAPCQSCRQAEWSDWTPWGPCSVSCGEGVQRRSQTCLGQGQCSNGQHKRLEIQACSLKDCCP
TALMPAQQQLLLQQAQAQLLAAAVQQSNAAAAANAASQLPGGEAPQSQPQTPQLALSQPIQLTAQDIQQLL
ESPQVQCFSLRKRDLFSVRESLLDMALSSFDDQYKDCVSMMEAELGELNRTELANNRNYAETWLEAASSW
YKSRVLWFWQQKDGEPLKIISNESTNPTQEGNALWFRPIEINASGAYICTIREEIPCLSNVIKVQAKNNTNCS
LGMDTSPPELPWDQMPTSLFPDLQVKSEPMSPASSHSSDSSILSMTPDPPRQAASPDDVMGVKAEHPPTP
RLVGGPHRCAGRVEVRLDGRWGTVCDDGWDLDDVAVVCRQLGCGPALKSLTSKVYGPGSKSDPVLMVI
NVPGSYLCSCLPGYTLDNDKWSCYADDPAPFLIFSHGNAIFRIDREGINHERLVANAGVPVLIDFHYKEEKN
DGQSKKTEKCNNTNSPKWKQHLTVIVTPVSKLNFRVWSHQTLKSDILLGSATLEVHETLKSNNMKRPTQSDG
NCQNRTIKWYITQTNQLIPAEEGGRIHSSGRFLWFLPTSKEDSGNYTCVTRYSNYTKQSTVSVMVYPHKQGI
GGHPEMESKLPPDLEPGMIFDIVDPAGALHRVPLPAGGRPKEGELPSCEINAYMPQEAHVPWAAGLTPEH
YASGHTQLSPANKFQDSVTLTIQPKDLPTSASSVLYVYLEASSSHFTREKKMPVTYKNGFLFIQTDKPIYTPDQS
RLHELQIEGKAKYGPVWKASFGPILTVHVAEPSLIEQVLRQEGKHPIRSDLSSWKDYRVCRGHAYGLLTAEG
GRHIVSFPGLPLKMLLLLLFHIAFPLLPGAQTGEIIGGQEAQPHSRPYMAYLNIRREDKRFRCGGFLVAENFV
QLFNCQSLHKLSLPDNDLTTLPASIANLINLRELDVSKNGIQEFPENIKNCKVLAIVEASVNPISKLPDGFSQLI
LSGGVEAQGARKERTKAARAPSVSCDYSPGDLVWAKMEGYPWWPCLVYNHPTEGTLVRAKGKSSRVHVQ
NLLSGPSENDPNLFVALYDFVASGDNTLSITKGEKLRVLGYNHNGEWCEAQTKNGQGWVPSNYITPVNSLI
IFAANFLDIFSLEVSRVFVSDSPTKYNIVPFAGLQNCHMPYGRDVLQRKEDTSTESLDSMDNPVNSSRYLSC
IYKDCVMGSFKNQTFLISIYQVLQEHQNRDSDLFLLDTRAVWASEEGWLEFDITATSNMWVMNPQHNMC
NGDCALAGDETGSAQLGSPGGKARGCSTVQQRESKSGIPRRSSIIKDGTKQKRERKKTVSFSSMPTEKKISS
NISSSSLENLTSKVFCSNSEVPSTLVIKDKAVVVMRGNCSFVDKARTAQSHGAKILLIASKSGLSSPSGNKTDV
SSQCSQTILWTDGRRVTGRQSSRYRLEGNLAQGDVSLTIVNAAEADRGMYCCRVEIPGWFNDQLINLEVV
EVGLFNELASSFEHEFKKATEDDEKKNAGPLDMSLPSTPDIKIKEEEPVEVDSSPPDSPVSSPQSPQIKEKEITS
SQRVEPRSVWMAQNEDPEFWDFQNFWAQRWQAWFNASLNTLRELYNQSEGFHIIQMITGCDLQEDNT
DNDLDALMADLVADLTEVEQKTVQAQKTSHYSQSEVTQPSTGVCAAASYDYKSNVPVTTTGYFEDDLPPPP
GLQCLVCSFVVHKRCHEFVTFECPGAGKGPQTDDPRNKHKFKIHSYSSPTFCDHCGSLLYGLVHQGMKC
ELQVLGCGSVAQAVLSRGAMGGTVTVSAGFAMAVTMAVYVSGGVSGGHINPAVSLAMCATGRLKWTKLPF
ACTVPYLDTICYCDLFCNRTVSDCCPDFWEYCLGIPPPVAVHKGCDRGGQKYPTGATYRENCNLCTCSG
DDGCYTCLFNTFPNGAIRQHTCLTVYEQLRGSLHYVISEGNLTATCSANAWPRPVISWIVPKAESEQTEEVVT
SLQITSLRMGDTGTYGAEIITATGTIHRPFLLRVYKQVPEPTILCDSVTCVIETCNYNLRCTVGDGGEHVTYSW
VRVLQPLNDDVLYVCGTNAFQPICDHLNLMSFELRGKNEDGKGRCPFDPAQSYTSVMVDGELYSGTSYN
CRPRASCRDCIRSQPSCAWCKALNFTKAGESDATRCATREELLRRGCALEEVVEPRGRHQVLEDTPLSDS
TIHNFTYESAHVTCLIKYPAKEQFDKILTEVKSGFPPSTPGNISCIYHYNVKFTCSWIPGKETYLTTKYTLIRKLPI
RFFMILAMFAGFLSFFALIASFFCSHFRSVSLTLVSTVGSFSAGVCAVIALAVFTSADCQLVMPGILSYSWSYG

VLSSLTTPFDVTSFDQERMSRSSEGRTMPVRDVTVTSNEAFLSDNGLISSFLNEQWTPSLKSFTVEYRSDNS
NSSLNNVDHHLLHHHNELNMANNATSAAATGNNNNNANTSNSSSSSSESALKEGGSSSGLPSSSSSSSN
IDPDQADITYEGRQKMTSLSSCFAQLCHKAQTVSQINHKLEAQLVDLKSELTETKAEKVVLEKEVHDQLLQLH
AEEAYGAEVQDYLNTPTAPFASTIGSHNVTLRWKLANISGVTYIIQWKYNQLPGDWRYAEVVSEHSYTVSNLQ
WKTYEKVSSADGTLVGYSEKKKGMWGICIFIAAVDLDEMTFTFTELLKSIGMRVCTCFQLLREMDVNMDTIS
LCQNPKLQLKNSPPYILDILPDTYQHLRLILGNCDDNQKLAQLSENEYFKIYIDSLMKKSKRAIRLFKEGKERN
QSQPSPCDPKSGGHPPKVLPGPGGSMGLKNGSGNGAKGKGKRERSISADSFEQRDAGTPSDDSEIKDCN
LSSQSTENVYETAKSVEVDVFGDIELKVIMNISETISCLWVFKENQAGCKPSLDSENGYVVSLAFSKIKETQAC

LENETSDMKEVLKHLQGKLEQEARVMVSSGQTEVLDQLKALQMDITSLYNLKFQPPALAPELACPEGSPVHS
SFADTIKVFQTDLQINVVEPENTERAANIRNQILFNQNEGPETVQCEITGMDTLLHAPIQTPVQQTEWHDTDI
LRGCLNSEKICIVPILACLLSLCLCIAGLKWVFVDKIFEYDSPTHLDPGRIGQDPVISVEPTALSAWVPSVVHTS
RTIAIPQPRQRSLSESDTSCLEQECFIEPYATVTELKLGPNWKHDDSERSPVQSLEGSGNLCEDENMDTYLS
RLLPDFDVQEQDVDTLHGSVHVTMYGTPKGNRPAILTYHDIGLNHKTCFNPLFNFEDMQEITQHFAVCHV
GNRSVAAIDSFAMKNCSSGSHSGSKMALQGSFSCWNGTEIKLGQACDFIRDCAEGEDEGDMCKKLPPC
SWQLFQTLRLSELTLERNLFNCSCDIRWIQLWQEKGEANLQKQELYCMNMEAAVIPLQEMNITQCDLPEI
PLPGPHLMPVKMDPGSRELPGRQSDPYEPPDGEPGCDDPPGEPPLSRKTATFKSRAPRKNPAAGEEWSG
LFAEFLECCSRLTEEHPEVIPFLASRHQKASAEFLASAEFRNVLGRCLTRVQSKRSKVYVYINELCTVLKAHTQ
QMRPTAKFGPYQKALREIRYSLLPFANESGTPASTEVNRQMLQELVNVGCDQEMAVRALKQTGSRSIEAALE
LPNPCLSAPCKNAGTCHPVIRGNAVDYTCSCRVGFTDRLCLTPVENACLNNPCRNGGTCDLLTLTEYKCR
QTNQLQYLLKVVLKTLWKHQFAWPFHHPVDAVKLNLPDYYKIIKTPMDMGTIRKRLENNYYWNAQECIQDF
KVEGTFSVPAADGQPLAKETQGKPGELLSQSLGTQSGVSKKKKKRKRNKSKKNPSDSGMQDSLTSVSSAPC
ERNHPRVYQSRDFRGHNRGYRRPYYFRGRNRGFYPWGQYNRGGYGNYRSNWQNYRQAYSPRRGRS
QKVSFSFSLTKKTLQNRFLTVLGNEKQNETQNSPATPLPTDFAPKIKMDLGDTSGSAEESSPPKPKVELGKIHF
NPKTSVSELFAILRQWVPQVQQNIDIIGNEIIKRGCSVNDRDGLTDMTLLHYTCKSGAHGIGDVETAAKFASH
SQTAIGYTAPLMLLSPAMINGGSTSALVVHWEAPAGQRDGYLVSVYEEGSLAAGRRVNVGKDSTNVSLAGLV
TAGQDEYTILPPSFSIGIHGYLLVYSVTSLRSFQVVKTLHNRLYESRGKTRMPVVLVGNKADLSMESREVKTDI
RRLQDLASMISAEYLNGAADGPGASRAASPRPPEPAALAGPQGDGGAAAGSPQAPAELGRRSLAGATCCEI
KPEFKMSKKINGGSVVEMQGDEMTRVIWELIKEKLIFPYVDLDLHSYDLGMENRDATNDKVTVEAAEAIRKY
DRRLTPTKGICPKQENIAGPSRPAKLLGAPIAASSHSHTSVITSPLRALASLPPCLSLPCCGARAVSVGGTQTE
GSSGAAGSGGGGAAPGSGGSGGPAAGVGPGFDAALQVSAAIGINLRRFRAACGSEAGSGEDEQFLGFGS
PMSESKLNTLVQKLHDFLAHSSEESEDANSPPRLSMSKSIGKSSGPENSPTPMENNREEGNSSSERSKSLGS
MECLRSREVWLLEQVDLIQQLKEETLQQQTYQLYWLLGQFNCLIHQLEHPHTNDLVNQISVCLERLGSLTLI
VQGNLEITYLGPDADMSFLKDIKEVQGYVLIAQNQVSRVQLESLRIIRGTQLYEEKYALAVLANSDPTGTVGLC
CPLCKTEVTKRSLQENPRFKQLIEGLLETIHAFELDTGIKFLNSQDFSKKPVETSSTELLWKEGSIIQSKGYRNF
DQIKTWVASNEIKDKRQLIDNRKLIETQMERFKVVERETKTKAYSKEGLGLAQKVDPAQKEKEEVGQWLTNTII
VPGILHFLQHEWGRFEAERAEWEAERAELQAQIAFLQGERKGQENLKKDLVRRIKMLEYALKQERAKYHKLK
KQQEKKPPLSSSVRFLRSSVTRTFSGTSWVNMEKTDILFHNQDSLNVNGFTMSLRSRSFSHRLSQTAVITKF
EAAAGPGPGPSPVPPLPAAAPGPLSLLDTCAVCAQSLQSRREAEPKLLPCLHSFCRRCLPEPERQLSVPVPG
SASSDTGSASQSCFLNTESSREAASATKASAQEPVSFSKPQTKKSGAGSTWSQLSTNNKELAVCSAVPSPNN
SRRPKLLPCLHSFCLRCLPQPDRYLMLPPVAPSPSPGPGTPKELQPPASPSSPAPSPTSPLHCTPVGVIRCPV
IQFFVIEEYTGPDPVLVGKLHSDIDSGDGNIKYILSGEGAGIIFVIDDKSGNIHATKTLDREERAQYILTAQAVD
LHVRRLQKALREWAANPGLFSQPMASLPVSSIPLFKISESGGRKSLSNGHASPGEAPGKATPPACSPSELGEK
LPPLESSTQEIGEELGDGVIYSISLRKVQLHHTANKGQRWLGFENESALNLYETCKVRTIKAGTLEKLVEYLVSA
YGIGAFAAVLAYWFASRPKALKPPCDLLMQSEEVEGWEGARRSVIGGSTELLTHYYDDARTMYEVFRRGFNI
MMEADEKEGLRCQVPDSTAEALIEWLQSQMTNGHISGNGDIYQERLARLENDKESLVLQVSVLTDQVEAQ
PQGSKLAAGGGGEDGVLEEKLLTEPMDSSTHLDTTGSLSQVIDRLPQPPKNNRVPGMVAAAAEGGGEDAS
AVNKIYVLNENLQNISVYKTGPVLENPDCAPCEDCKQRANLSNSFWKDNVNMALLLETYYDDQLISCGSV
AEDPRQGRNPPKFVTYHKALQEIRNSLLPFANETSSSARGTSEINRQMLQDLQAAGFDEDMVVRALRQTNS
PPPRRDGAPICLQLATGIRTVLLSKAGEEQRPTETPPAQAPPTSDPSASCTSKGVYENYRRWQRYKALARRH
LLQADLIENVGSPVTTAVLTSVSEDSRDQFENSVLQLREQDEAETAVPQGHSNTMDGENNSGTDDIKIQFS
SSPEPEQPSSNKWQLDKWLNKVNPHKTSILNQNDNHGLEGNQYYSSIKDEGQECEKIPAICQTELSDKEIK
GHGCKELSKGCSSQNYTPAKIDWSPIIWVNRSLPLWGLQVSVALISLFETMLLGYLSYKGNIWEQVLRIPFLL
AGGSGLSGGQPAAAPRKCSSRRNAWGNLSYADLITRAIESAPDKRLTLSQIYDWMVRCVPYFKDKGDSNS

CCGCRFPLLVALLQLVLGISITVLGFIMAGISSSVLVRDTPYWAGIIVSVVALVGMVMLCVSYQPEEKTCFQFA
FWLGYERGSAKGEAAAWPVIITDKAVMPVQVIDGTPENLLPSHLQKVDLKYLQMGCNQANKYFQGTVILSY
FQQKKIATEHQIYGNKDSYFRNMEEKMKKNGIILKDELKTLLHLCKTPDDVELAKNVIYRYHTENRNVAFGEI
LDICCRDPGGNGYACCAPSLGSPHSLPMIQASSVAKPAGTVCPDGTQCPAEYSCLRTPAGSFSCCPLSEA
SAGKTLVAELLILKRVLETHKKALFILPFVSVAKEKKYYLQTLFQEVGVRVEGYMGSTSPAGRFSALDVAVCTIER
LDYIEVVDYQGRNRHTVIQGRQVRHLYSLTVFENYLYATNSDNFSILQINKYNRTDVQSLIRLENAREIHVYG
LNVRTTHYTPLPSGCNPLKRDIAEYVRTGFINLDKPSNPSSHEVVAWIRRILRVEKTGHSGTLDPKVTGCLIVC
REFRDVMQALLLRPLVKEPAVSRLLRIMQFLSRIEEGENLDCTFDTESELTPLESAIGVLELIEREFPVSEVAMES
GMQRLEDEKRWLSVPKVYVLQPWILNLLMKYEQLDSRNTQVPGHILKVVNDLKAADQGSAAVLHISDGSYY
GLALALHTAVSASDLQLHPPSYAWSHNGLLASLDHSSIRRGYQVYKQVCAACHSMDYLAFRNLIGVSHTEA
VSPDDSDSVKSDDEDGDIKPTKGQANQGKGSDDGKDPRRPKRPRTILTTQQRRAFKASFEVSSKPCRKVR
NGEGDIDIMALKRMLEKLGAAKTHLELKKMITEVTGGLGETIGYRDFVRMMLGKRSAIFKLILMYEDKAKEKEE
VFQSVWVSTDHDEIEKVAKKFGVKVHRRSAEVSKDTSSSLETIQEFLRHHNEIDIIGHIQATSPCLHPGDLIKV
IMRIIKNIVKTFKYIYGLRFEVKGLENFQIERPCVIVSNHQSILDMMGLMEILPDNCVQIGKKELLYTGPVGLIN
NLLPAAVGLGQANPNVEETGEVQGQFLTVVGAFWHLTKEGRCTAPVRVALLNCMKALLEADPHSNWAVLN
LWVFQLQVPCEEQTRAAMNTLHSDGASHRPTNGTKSDTGQEDFLSRLKQFLCEPPYSSSPEDSRCRLCPT
KNSSEILLGSKNKQSLQQQAKSIQEQVKKVEVTLEEDIKTMEMVKSKWEHFGNNFEALSIWITEKEKELDTLE
QAYVSSLIYGWDREYFWTALQDINETGAFRWLSGDEVMYTHWNRDQPGYNKGGCVALATGSSIGLWEV
HALSKFNIDLIVSLLRQENASNICVIQVPPEMRYSDYFIIVSGSSSRHIHAMAHYMLKMYKQLKQENDPHIQII
YNTLACDWWEEINCTNITKTECNFTDPDIKRFWTITLRVRAELGQVKSDWVETNKFVADKNTTIGPPVIVNV
GPPEPGYLLPSAACAPLQLRALIRHGTRFPTEKQIRKLGQLHRLLRSQERPCAAAQQLAHWDMWYQPDMD
DYLSQLTHDKDSMAHSQSTKLDQEIKKVKESIQEGVNYMGNLLEKLAPLNRGLQPRFYQDSDSLRRLIRKEL
PQNHAFSGSSPPYLTGSVPININSSSNKECLEENNKYYAHEASSKQEERCRQRLSFANDPSITHPTNLKPVKY
SLTLVNSNMLASDDKSLESAVVKVINPDEQCDVNLGLQASSSSLVVKEILQEAPELITQQLAHLLRGSILFQCM
RAAAVVVRLQGEGAADWSGGENGAGLQGLPHEEPPPQQREGAETAPSGDAVETAIGYRRSQCSHRAAEA
KINPCGGIEDCDGSSAICAYDLNKKVYLSVGDSSLKRSSKSLLEFNTTKPCSQQGAQHVVQSNINFLCGKTI
STQLAKVNDVLKIDLSSLVQKGYAAFNDTPDLLLNQTRNILSDIKNLLDLVGSNITSFTQMLPVQKILADLTIYLT
DGSCLDIVDKVISGNVEEFRKDFISRIWLTYREDFPQIKDSALTTDCGWGCTLRTGQMLLAQGLVLHFLGR
RKHFAWRKPILGRFEPDWKPFLTSKELFWQDDTPARRQLNDLLHLHRLPFGLNGSVLHAFHTLQLLPQQG
PKVLDGTRCYTESLDMCISGLCQIVGCDHQLGSTVKEDNCGVCNGDGSTCRLVRGQYKSQLSANKLEDT
LRISNDGKVDGTSIQSPHSLLEIRAVAVRTVAIKGVHSSRYLCMEEDGKLHGLLRYAAEDCSFEEEIRPDGYI
NTERCLSENPSLPENGLQVASQTVNRSAPNSNTALKLEEKNSEAVKFPGIQSTEPSMFKNLKQANINEGENG
LRTLVLALCAAGLQGNICTTRGVNSCRQCLAVSPLCTWCSKEDLRQGTPRCDLKTNLLQNGCTQDFIEYPV
QPQRRRPSLLSEFQPGNERSQELHLRAEPHSYLSDMSKQSEIEFIETKRPRLDLLQEPLLRHSPLLNQNQQG
ASRRGPMGIGPHGGGRPMLRARHAVSGNATLPPFPEGMQMAVFGMGCFWGAEKMFWSLPGVFSTQVG
QCRASFQRRPELQKNTVLCRVVEQFHSTQPAGPAEEKRGWDGSRSEVACDSCLEAAAAAKTCLTCMASFC
YEEAREIFQDDKRTMEFWQNYVEANQCDSDPCQNGGTCVDHFQAYVCLCPEEYKGKNCEKGLNEKPT
ADKELLVKLRKKTGYSFVNCKKALEKFSGDSKQAEAWLHEQAQKEGWSKASKLQGRKTKEGLIGLLQEGNA
SGGRPHYYVSYRRSAFAQMKLPKYSLPKDMHVISTDENQVFAAVQEWNQNDTYNLYISDTRGVYFTLALEN
LFLEPGSRCKKGGSVGKEACLSQSYETCKIRASEPCHSPVTNVSARSSKTEYNDFAVCAKGEKDTSDKNGN
FRFRYVCEGPSHGGLPGASSEKNKKSYPQVKIFNYVGPAKVIVQLVTNGKNVHLHAHSLVGKYCEEGVCTV
PALTHSSQSPTNLPAHTGSSDCNISCKGMTERIHSINLHNFSNSVLETLNEQRNRGHFCDVTVRIHGSMLI
ARTPLSLLELPPELLVQIFGSLPGTDLPNLARVCSTFRRILCTDIIWRRRCREEYGVCENLRKLEITGVSCRDVY
PVHYTFRTDCGSPGPDLCSFPIANISLVKNNRDRVLMYGQPYRISLELELPESPVNQELGMFMVMMSCYTK
LCKQQAEEEGASVGSQEPHSPGCCHHGNTDPSHSPGGTCPIAQAKTRLESQEGNHAGPDGKEESTGACA

YAFQARLGGTCIQAQLREKKRAQEMYGDALAGGQSSFLLQQEGAGGDVFSCSLGNLPPGEEAALTLFYVCI
SEERAKPAAQVEFADIGCGYGGLLVELSPLFPNTLMLGLEIRVKVSDYVHDRIQSLRASHPGQYQNIACIRS
GVEFKEEWQDEDFPRPLPEDDNVEADILAVTGQQGQHEVNGNKVRKKLMAPDISLTLDHSEESVLSDDLN
VTELSAAISHLIQVFSSSFQTDFQPAKILRSPSCADVVLDPHFSFTVYAAHNIPAAWLKSYKLFSFSCSLTYGGK
CKCGTHTGQFECICEKGYYGKGLQHECTACPSGTYKPEGSPGGISTCISCPDDNHTSPPGSTSIEDCVCK
SHLVSSCPGLIRDHKYHLRHHRHCCSGKELVDRLLSAGLSIQTRGQAIGVCQVLVDGGVLTHVKQEWHF
FDMTIQCIWSVYTNKIISSDRDLLSVVFYGTEKHKNSVDFKHVYVLQELDNPGAKRVLELDKYKGDQGKALF
YQRFENYEEEEEEEETDWKEISGCQNVTVLECDFSSAMTAYLDSYNVRIRAEKREEKSLWSTTLSFVPYSVAK
RFLWMGVGNACNYYNKMYGEFMRVWISGEETLIISKSSSIFHVMKHGQYICRFGSKLGLQCIGMHENGII
AQHHGARLGVAAPVWEAALTLCKYFEKQKLNFWGKKVIELGAGTGIVGILATLLGGDVTITDLPLALEQIQEN
YLRMRKLLDASDLETEAKMEKEMNCFYLKALDGFVMVLSEDGDMIYMSENVNKCMGLTQFELTGHSVFD
PREEPLIDRLEVVTLPSPLPKGLPVKNCAQQFQLPTHDWPDTGPSGQFDVGVVASFGRLLSEDLILQFPYGV
HCKGDEITVTLGAHNIRQQEQSQQEIPVYHQIPHPHYNKTTMNNDVMLLQLVKTAKLNRWVDTIALPCAG
RFRDMRTTTNLYLSSMAFSDLLIFLCMPLDLFRLWQYRPWNFGDLLCKLFQFVSESCTYSTILNITALSVERY
LKEYNVGGKVIHLVERAPPQAQSPTSGGSSGASSASAPHNGIGGRGAGATVHDRNANSYVMVGTFNLPVS
YSETDLALNGKHGDAHCKGFINNNTFPAVVIFIINLSTLESCGSILVVSSVPGINAYGNVSVVQTGNVSGYIDT
QFSVKKTTEPIQPRTLLKFPNIYGPKPVVTAPEIVNYVDYALKTTEEPTRGKDDIQDQNRMKIQVNEELFVLPQN
SAAVTHVVSEQNSGDEVARWLEQQREECGSGGDPALLDISWFTESMGAGRPVEIESRHRLKVTALAPLRQG
LEDIAVVHKTTAFTYAARPPPAPRTPPSPYSRSCPPQDPPAGPHWTAGAQGAPAEAAGGQQEEEEEEEKNMI
LGAVLPGDAAMKTPGPALSPRPGSCWLENPFLGGHSPLPAWVEETRARLLRPLPPPAPWLSSLTLTPCLASSP
FVTRKNRQVCTNPDTKWVKDYVNFLEMK
IIHCKSCTLFPPNPNLPPPSTRERPPGCKTVFVGGLPENATEEIIQEVFEQCGDITAIRKSKKNFCHIRFAEEF
FVSFDSRSEAEAAKNALNGIRFDPEIPQTLRLEFAKANTKMAKSKLVGTPNPSTPLPSTVPQFITREQYEVVLSM
FHENKPSRKTLHNSIELLRKYCMGDHPVQFEILSDQSVNFGFSKFPWVNYQDGNLNISIPVFSIHGNHDD
PVDIRVPFPSLQSVKVFLESHHIEYSILLEDLQVIVEEEKQEMVNNQRRERGTSNFNYGAYHPLESIYAALDDL
QMKFSCCYGSNIKNLFSTAVNKGIENIDACLSFLAAKGINIQGLSAEEIRNGNLKAILGLFFSLSRYKQQQQQ
LSKNRLKSVNPEEFTSYPLLEEIDLSDNIVANVEPGAFNNLFNLRSLRLKGNRLKLVPLGVFTGLSNLTKLDISI
AGSIIMNREIESMAMRPLAKDLTRSLEEVRNIIRDQALRDLNLYTEKMKESLKHFDVLFAEFELSYVSAMVPVK
IALCEGLMNEMNQALDFFEKDPDVGAIVITGSEKAFAAGADIKEMQNRTFQECYGGSFLAHWNRVSLVRK
GRHIVSFPGLPLKMLLLLLFHIAFPLLPGAQTGEIIGGQEAQPHSRPYMAYLNIRREDKRFRCGGFLVAENFV
RHDPLPASCRRRGASPKAAEEEEQSPGSSQCLRRSVAFFAVCDGHGGREAAQFARENLWGFIQKQKGFG
ERATVEAAVQECSQSLDETIEHIFNIIGAFDVPRYVYNSERKKFVPLSMTNHPAPNLFGTARDKAELFRERYTIL
QQMWVYDEDEGISCRDVTFVPGLYKIFDEILVNAADNKQRDKNMSCIKITIDPENNTISVWNNGKGIPVVE
LDMNNDYDCITQKKRKGEKRSNRLTSSTRVLQQHTKYSKCGCPADSDGTEVEDSSEFGTLLLDSDSDDSV
SASSASMPVSRPARMTAASSGSREGAASGALPSGTEASTVPPAASGDSDPSPTLKLGAKSNCIIVSARQRGNP
YKRSRLRDLLVAASSSLHWPAVANRGQWEVRSDGPADGAEIEKESCGDPGTPLYGIREGDGFSNRDILRFI
ALAGITDLLDGFIARNWASQKSALGSALDPLADKILISVLYVSLSCASLIPVPLTTLIILRDIVLIAAVFCVRYKTLSI
LVSAHRETDASCSLCRVAHFAWPVPEAKQHRMSLKAVTLALALLAITGAHADVSAQQVADVVWNYFTQLTGN
DSRSGSKISCKVELEVPQLCSFILKTSECTLREMSGVDAEGKTLLKQSKNSDDFARAMAQYELKFSTQDGKK
LFDEQLMEFHCLWDDSFPERPERLAAVKEKLQQCGLWERCVPVEARLATEEEILLAHSQEYVELMKSTELMS
SPACHQPLHLVGHHHSITALAFGSEINPLLVCSASCDYVIVWNLDECTKKALQGLMPRGIVIGTLLGMVLYV
LSSNAVLCWKFCHVFHKLLRDGHPNVLKDSVRYKNELNDMSRMWSHLSEGYGQLCSIYLKLLRTKMEFH
NLPVMIWIYGGAFLLGGGQGANFLNNYLYDGQEIAVRGKVIVVTFNYRVGPLGFLSTGDASIPGNYGLKDQ
CVVCVPRGARAIPQPLSFQQLSSESEVVIRVIQRLCEKKKKNVLAFGFSLQELMWKMRVNDCTWLRVIKGD
VVIQKEGVRSRLTIYNANVEDAGIYRCQATDAKGQTQEATVVLEIYQKLTFRDIITPQEFRQGEDAEVVCHITS

EQPLESLDSMDETLQIFQRILPSLPMENVPVAQIFNDIENLRSLIQTLGSHLGCTFHKSSNLDALGNLTELLT
INPTSCNNPKLEELQMLLIDWINTTLKQEHIVVKSLEEDLFDGLILHHLLEKLGSLRLDVEKIALTEKKQKHKL
FDNSQKKIIHYSVNACLFPICALHHVAVTTVEGIGSTKTRLHPVQERIAKSHGSQCGFCTPGIVMSMYALLR
VGTDSSIFHMGQGSQPQHLQLRATLRLKSGDCVPRSWIYLLNEGSTDLATEGRPGMRTELFSSKCPDTII
MDDNRITTVEGNDEQHYKTFNGLKSKNPGLKTLLAVGGWNFGSEKFSSMASTPGNRRTFILSVVEFLHKY
TEREEVRGRELTGLKGLYEAELADARRALDDMARERAKLQIELGKIRTEHEHLLGSYAKKESDHNAAQIKLRE
KIIDIIDTTGSGDVNTSTIVEPKDGEIVGLSGRTLKIPTNWVNPSGKYHIGIKNGYDFYPKALKERIQKERKEK
GWLVNMHPTEILDDDKRLVSSVDYYFIQEDGSRFKVALPYKPYFYIATRKGCEREVSSFLSKKYQGKIAKLE
HLEFIHDHADYGFETPNVKFNWRVIKEKRDAYVSRLNGIYQNNLNKAHIETIRGHAVFTSDPEPTVEVNGK
YIYNRQQHLHFDSDLGVWVADTELGRATAELWNSQPESLQYKRAAVDRFCRHNYQETEPFTVSRRVQPK
PLNIMYLLKHRWVLGKPACQFWLSMDYVASTASIFNLFILCLDRYRSVQQPLKYLKYRTKSRALVMISGAW
KSVKWKLGADSEVWVWVMGEHASDKPYDVICEEIQAWRAQQEAGEARGRQTFLELAAPGSLHSQPGAL
QKIINQQQHSTVSSLQKNNKKSTSLQISLQSRRYSDCFQSDTVVASEDTFISVGNEGNACNRNMADYRS
ELVSPFYNETLPERSFYRKNSSGEEPLIESLEEEINTIRVSNHHLWMRMNNITLAAGNPGEKGAEGLSGPGG
VAIKKISPFEHQTYCQRTLREIKILLRFRHENIIGINDIIRAPTIEQMKDVYIVQDLMETDLYKLLKTQHLSNDH
PYVECRNKPCGHPGDIQFGSFELTAGTEFVFGARVQYKCDEGYQMLSQRNFRECQADGWSNELPHCE
LSPRGQINCSRIIRGDVEAVEEALLSRLTTKTKAEPLAPKDYVNLTKDCTHFKTRRRFVEFPMSKEEEDFPIA
SLDSAILIGETGSGKTTQIPQYLYEAGIGRQGIIAVTQPRRVAAISLATRVSDEKKTELGKLVGYTVRFEDFTSEE
GLPLNVAHSSLSTPCHGLQTSNPVISIVPSTNHPSGYGGHVDGGASGYYLSPNVRPNGAPALESPRIEITSY
LTYISKVSDATKLRPKAEFCFVMNAGMRKYFLQANDQQDLVEWVNVLNKAIKITVPKQSDLLCQPDNPNR
GGNNPHTIPTAAVSGAFSVQVPLSGRSQDDATVGYFFQRQAGEQLSGYSSKHRWPTGDSIHVDHSVRS
SRIFLSIQAKGGEYLNIAPKRRVHLTNLGTGWISVSMTELQIEDSGTYLCGVLDHQTIILLKMIKVVVSYDAPM
RKGLAHPEDGITSTLRRENELKAVITTLQNASFMKAAAYMTVSKPINSPKLKWTNSRILQNITYNSKGDPVIH
EGAENLRKVTTDKKNLAYVDNILKKSNKKLEELHHKLQELNAHIVVTDPEDVTECPRTPDTPNSDPRFSTNN
GADIISIKNKEENTFIQETFKNHWQGPDYISLGMFFDTDDNAFKWFDKSEVKFTNWIEEESNEELLNTCAS
VFHKAIQYSKIYRSDVSSTVWGHSDFRLPPLYAAFVNGVAVHSMDFDDTWHPATHPSGAVLPALLALSEVL
NHRGLEGLKRNVTLQNIIDRFQKASLSGPNSPSESRRERTYRKSPAMSSERIACQFCEQDPPRDAVKTCIT
QSNAARDFLNYLVRLNEIKSEEIPALGAGAADAPDGIVPEARGDAGERVSALGGPLPPHLALKSETGGGLAP
VTGFGMKRTVNTVWFLNLAVADLLCCLALPFLAVPVARGNRWELGVFACKLLPSLTILNMFASVLLLMGISV
LSRIHQHKNLLWFMPAILEDSGSYECVMWSLTSCNKIYFSVTVFKNTAGLCFNGDFLYAQEISASSNEKIVC
LDLMSWDSGPWDVMSHTCIDENLKEEPQSPATSVCSVPSPLSDSSVQHVPEDLDFSTSSQMSPLSLYSK
VNEKGQLSAALLVKLNKVNGIISSTPPGNNAVAKSKKSLLSVNVWNEDNSSKNLIPRLQKVKNNYLAMNKY
NETLWKWVSQNRLFHLGSRQCLGLNIAKPAAPLKMVNCDSQLMLWWRCVDASIFGASQYKLTLKDNVVT
VELAERLQREEEEAYASSQSTQGAQSLTFSKFEEKKSNEKTRKVTTVKKFFSASSRTGAKKEIHEAKAPSPSINR
IRDWTFQILVMLEMSPRRGGIYTCQVEHISLRGPLTVHWASVTLQLFQINVFHNASSTDVATMALLGDLETH
YPFWEQRPQVEQHIRGPRAAVPIPRGSGHNSWGTAGSFRPPYISQQTRSLSSYPPAQYNYRKSEREFDH
MDPSVVARSMVAFVEGMTCNSCVQTIEQHVGKLNGIHNIKVSLEEKNVTIIYDPRLQTPDSLQKVIVDMG
CVSILERFLQAVDPLYVIQNFREELQKGLFHPDDSVKLLTIAQVGRIVENSDAVREILNNPELLRQIIYCIGGE
NQKPNLEKNLKLTNIVEKFNSLNLEKTPSVLHCVFCRRGPPLPAQKICLRCEAPCCQSHVQTHLQQPSTA
RFSSSRCIIVLLSAELVQNFYLPDVLGSLQEALKPPHKVVKLFCGVEDCDDYSGFFRDWAQWKELTYDDE
GRNKTFLCYVVETQGKESGTFRGYLEDEHAAAHVEDAFFNNILPTCDPNLRYNVTWYVSSSPCVPCAGRI
EREVNILKEIQHPNVITLHDVYENKTDVILILELVAGGELFDFLAEKESLTEEEATEFLKQILNGVYYLHTLQIAH
GKVKEMLDSETLCRRAVKILKKKKLRRIPNGEANVKKEIQLLRRLRHRNVIQLVDVLYNEEKQKMYMVMEYC
GGGRRRDSLESGHQALQAEHRMGGGKQHRGPFQGVRVKNSVKELLLHIRSNKKMSSGHVADEFKAQA
GYTYRDYQEYNPSYDDDYTEECPVYGNLNQEILDECCYEQMKAQPRRYANDVKVDVESQMCYASLDHS

LQAGELPTHFVSRTMKWDRYAPVQLVPHVEKMQQEGGQRRKRQVDGCPALQLQAIVNSEPNERSLSPWR
LGRCYNYITTVNPELRDKDCQNIWETMQKAFIYKKPCNITTEDYQPLLDLASHPIPCNKSLFWSKTSDLVHR
DNENKYKKSNVLTVETINFLKGNLASAQEIDTKYGQVSEDLRRRLVTLKQKVLIQVNRIKALQNNIRDQVVEM
VKEELDLVIDVLENIRDPNLSEPLGRPLEILRHIREDLKNCIRHQPDNHQRSERLTSWLQKFHATKKMETPG
KRHLETRERAKVAVLVNRVHLVEQHYTQEFHALQDRFRVTPVSSNSDQKFFFSKEVKLSDVVICTAKILQNAL
STPTYTLLLITNAAVAMEGTKFCTGNCQGKSHQGSTNLQVYSLPDTLQLETQPKELVAGQPAHLHCSISKVYI
SRTSVTPSNQDICSSSAVFSECCHHSPVQSAVVLKNPHCQNSLTQGVTVTVICQDTLQASKRNSHGQDWV
AQSKLQAVEEEVQKLQVSLQPENASDASAKGSDSVQLLDEAQAGIQMLKDLLGNVSADPVQIKIRLDISAAR
TLRDCGHPNVVAYFGSYLRNDRLWICMEFCGGGSLQEIYHTTGPLSEKQIAYVCRETLQGLSHLHAKGKM
KCTKDECDTYFKVCLKEYQSRVSAGGHCSFGSGSTPVIGGNTFNLKYNRNNERNRIVLPFNFAWPRSYTL
FTCMLIGEIFELMQFIFVVAFTTFLVSCVDYDILFANKMVNHSQHASELVKVTLPDAFLPPSVCSARIQQNGF
EINLRPQMSQFLCLKNIRTFLSACCEIFGMKKNELFEAFDLFDVRDFGKVIETLSKLSRTSVALATGIRPFPTEE
RVLKQMTSCSLKTCSKTDIQSLFRLTQTGMNGPETSADQLQLAEVKAKLGASQRSISELEKERHQNQQECE
GALLRGVCNFMEEIQHVQSRYRSSWWRVVVACLSPSLHRHMLLLLISFCAYMALFGDTGLQLFLNLILLCLC
NFTLAQNAEVLANKSSCGKENVSAPILAIGFGAGHLLQMSFTKTSKLYSVDELTFIYNLSDTTVFPNSTGGERN
QEDALAPAVKAALTQFPSISHPPVGPAPSVGYNIDTIHPEVLAPEASRTFGYQVLQFKEASIVVGAPGDQNSTG
ELCNKEEAREIFENNPETEYFYPKYLSCLSSHRAGIFHVTYVISDSSTDLRNCVKEISNQCSPLPCNEDGYME
LQSRHEISPGHDGSRNEAQLQEMAQLRIKHQEELTELHKKRGELAQSVIDLNNQMQQKDREMQINEAKI
ERTRSEEDNEMNLPSLAAAYTTILRSLGEDPERQGLLRTPWRAATAMQFFTKGYQETISDVLNDAIFDEDHD
AHCNGDKINVTLGAHDITKQERSQQVIRVRRAYPHQDYNKKSHNNDIMLLKLEHNALLNEYVSPIPLPMA
METSHRRYAEKLSEVESDLKKLDDQTGQKDMSTNKELSSFRSDILALRQQLQEIAEKTTKNKDTLEKLQDSG
FTDTMETCDDINECLGPRLPDCGRNRNCNNAPGSYYCTCINGYEPSSGKANFTHASENSCQDIDECK
TLCGRESTDTEEAPGNRTYHSIDNALTVTFRSDYSNENPFTGFEAFYAAEDVDECELLIDDEPLCDHHCHN
VGARDTLLSLRVDSAGSMELTGSIVWQPVPEKKQECVFKKKSNLTECFNFIRVLVPVNQTRLYTCGTYAFSPT
LSTRLPESDDKNVYSEVPTPQLSLPSPPPPTAAQQLDDLLSHLGQMQSKISVVGSTAEPERAKGPSAVGPSPG
EITRKVSIGPTRLGHPCFYCNEDIKLANFTIKQGEQIMFNSVEEVNGTLTVNCGVVRDNQSHSFTLPLSQEG
YERFLLQHFDAKPKGRDDRYCNNRMRYMMKEDRRRDRNPGPKVCKETNTFIHGNSDDIKAICTSHGGR
RSGYSGVATFCKDSATPLAAEEGLSGLLARPGEGGAVGCYGDLGAFTPQELKALDSEGCTVITHHRIRTSAQ
KGTALKNNSDADLVLFLSCFSSYQDQMENRAAVLHTIKQKLNRCRQTVAFSIDVEVSQPKEKGTCPRSLSIT
VLPGDVLLLPSYPEADGERLWLGPGAALPGRLVCGPGLRSCGAGLLVTKCGLLRHRQPGGAGGGSGGAY
RPRAGAPCQSCRQAEWSDWTPWGPCSVSCGEGVQRRSQTCLGQGQCSNGQHKRLEIQACSLKDCCR
TALMPAQQQLLLQQAQAQLLAAAVQQSNAAAAANAASQLPGGEAPQSQPQTPQLALSQPIQLTAQDIQQLL
ESPQVQCFSLRKRDLFSVRESLLDMALSSFDDQYKDCVSMMEAELGELNRTELANNRNYAETWLEAASSW
YKSRVLWFWQQKDGEPLKIISNESTNPTQEGNALWFRPIEINASGAYICTIREEIPCLSNVIKVQAKNNTNCS
LGMDTSPPELPWDQMPTSLFPDLQVKSEPMSPASSHSSDSSILSMTPDPPRQAASPDDVMGVKAEHPPTP
RLVGGPHRCAGRVEVRLDGRWGTVCDDGWDLDDVAVVCRQLGCGPALKSLTSKVYGPGSKSDPVLMVI
NVPGSYLCSCLPGYTLDNDKWSCYADDPAPFLIFSHGNAIFRIDREGINHERLVANAGVPVLIDFHYKEEKN
DGQSKKTEKCNNTNSPKWKQHLTVIVTPVSKLNFRVWSHQTLKSDILLGSATLEVHETLKSNNMKRPTQSDG
NCQNRTIKWYITQTNQLIPAEEGGRIHSSGRFLWFLPTSKEDSGNYTCVTRYSNYTKQSTVSVMVYPHKQGI
GGHPEMESKLPPDLEPGMIFDIVDPAGALHRVPLPAGGRPKEGELPSCEINAYMPQEAHVPWAAGLTPEH
YASGHTQLSPANKFQDSVTLTIQPKDLPTSASSVLYVYLEASSSHFTREKKMPVTYKNGFLFIQTDKPIYTPDQS
RLHELQIEGKAKYGPVWKASFGPILTVHVAEPSLIEQVLRQEGKHPIRSDLSSWKDYRVCRGHAYGLLTAEG
GRHIVSFPGLPLKMLLLLLFHIAFPLLPGAQTGEIIGGQEAQPHSRPYMAYLNIRREDKRFRCGGFLVAENFV
QLFNCQSLHKLSLPDNDLTTLPASIANLINLRELDVSKNGIQEFPENIKNCKVLAIVEASVNPISKLPDGFSQLI
LSGGVEAQGARKERTKAARAPSVSCDYSPGDLVWAKMEGYPWWPCLVYNHPTEGTLVRAKGKSSRVHVQ

NLLSGPSENDPNLFVALYDFVASGDNTLSITKGEKLRVLGYNHNGEWCEAQTKNGQGWVPSNYITPVNSLI
IFAANFLDIFSLEVSRVFVSDSPTKYNIVPFAGLQNCHMPYGRDVLQRKEDTSTESLDSMDNPVNSSRYLSC
IYKDCVMGSFKNQTFLISIYQVLQEHQNRDSDLFLLDTRAVWASEEGWLEFDITATSNMWVMNPQHNMC
NGDCALAGDETGSAQLGSPGGKARGCSTVQQRESKSGIPRRSSIIKDGTKQKRERKKTVSFSSMPTEKKISS
NISSSSLENLTSKVFCSNSEVPSTLVIKDKAVVVMRGNCSFVDKARTAQSHGAKILLIASKSGLSSPSGNKTDV
SSQCSQTILWTDGRRVTGRQSSRYRLEGNLAQGDVSLTIVNAAEADRGMYCCRVEIPGWFNDQLINLEVV
EVGLFNELASSFEHEFKKATEDDEKKNAGPLDMSLPSTPDIKIKEEEPVEVDSSPPDSPVSSPQSPQIKEKEITS
SQRVEPRSVWMAQNEDPEFWDFQNFWAQRWQAWFNASLNTLRELYNQSEGFHIIQMITGCDLQEDNT
DNDLDALMADLVADLTEVEQKTVQAQKTSHYSQSEVTQPSTGVCAAASYDYKSNVPVTTTGYFEDDLPPPP
GLQCLVCSFVVHKRCHEFVTFECPGAGKGPQTDDPRNKHKFKIHSYSSPTFCDHCGSLLYGLVHQGMKC
ELQVLGCGSVAQAVLSRGAMGGTVTVSAGFAMAVTMAVYVSGGVSGGHINPAVSLAMCATGRLKWTKLPF
ACTVPYLDTICYCDLFCNRTVSDCCPDFWEYCLGIPPPVAVHKGCDRGGQKYPTGATYRENCNLCTCSG
DDGCYTCLFNTFPNGAIRQHTCLTVYEQLRGSLHYVISEGNLTATCSANAWPRPVISWIVPKAESEQTEEVVT
SLQITSLRMGDTGTYGAEIITATGTIHRPFLLRVYKQVPEPTILCDSVTCVIETCNYNLRCTVGDGGEHVTYSW
VRVLQPLNDDVLYVCGTNAFQPICDHLNLMSFELRGKNEDGKGRCPFDPAQSYTSVMVDGELYSGTSYN
CRPRASCRDCIRSQPSCAWCKALNFTKAGESDATRCATREELLRRGCALEEVVEPRGRHQVLEDTPLSDS
TIHNFTYESAHVTCLIKYPAKEQFDKILTEVKSGFPPSTPGNISCIYHYNVKFTCSWIPGKETYLTTKYTLIRKLPI
RFFMILAMFAGFLSFFALIASFFCSHFRSVSLTLVSTVGSFSAGVCAVIALAVFTSADCQLVMPGILSYSWSYG

| esm1v_t33_6 | esm1v_t33_6 | ESM1v_score | Tranception_ | Tranception_ | Tranception_ | Progen2_scor |
|---|---|---|---|---|---|---|
| 0.59673452 | 0.20892823 | 0.39331198 | -2.6821413 | -2.6823082 | -0.0001669 | -0.0007968 |
| 0.93670702 | 1.08547473 | 1.03503976 | -2.0061045 | -2.0061162 | -1.18E-05 | -0.000277 |
| 0.53111696 | 2.24252367 | -0.2813941 | -0.6684296 | -0.6880972 | -0.0196676 | -0.0021024 |
| -0.3508558 | 2.66346228 | 1.51677992 | -2.5327781 | -2.5329959 | -0.0002178 | -0.0007384 |
| -3.2215102 | -2.4588549 | -2.8294435 | -2.5219359 | -2.5211892 | 0.00074673 | -0.0015624 |
| 2.91607368 | 1.49647367 | 2.1044908 | -1.0984935 | -1.0994586 | -0.0009651 | -0.0008758 |
| -0.0659285 | 0.18799305 | 0.08358626 | -2.3558865 | -2.3558006 | 8.58E-05 | -0.0011935 |
| -4.7024608 | -2.7222056 | -4.1430592 | -2.1283772 | -2.1312451 | -0.0028679 | -0.0013059 |
| -2.2755139 | -2.3993518 | -2.1022693 | -2.4974861 | -2.4984484 | -0.0009623 | -0.0014783 |
| -3.6640544 | -3.921725 | -3.352479 | -2.829471 | -2.8299271 | -0.0004562 | -0.0022161 |
| -5.4864731 | -3.315594 | -3.0288445 | -0.7084033 | -0.7174104 | -0.0090071 | -0.0006234 |
| -2.186157 | -3.5101595 | -3.1078814 | -2.5071228 | -2.5108099 | -0.0036871 | -0.0024214 |
| -2.3177443 | -2.5406156 | -2.1578702 | -0.8100705 | -0.8212062 | -0.0111357 | -0.0027125 |
| -3.303447 | -3.939636 | -3.3252057 | -2.1010849 | -2.1022041 | -0.0011191 | -0.0006863 |
| -0.6065021 | -1.5895264 | -0.7858656 | -0.3185824 | -0.3194603 | -0.0008779 | -0.0004268 |
| 2.18707538 | 1.98650551 | 2.195239 | -2.4847016 | -2.4834509 | 0.00125074 | -0.0002644 |
| -0.9658525 | -1.0438187 | -0.943148 | -2.654047 | -2.6548281 | -0.0007811 | -0.0023933 |
| -0.2744212 | -0.5391381 | -0.4722488 | -2.3326099 | -2.3327382 | -0.0001283 | -0.0010594 |
| -1.5996066 | -1.7187452 | -1.3243799 | -1.8759292 | -1.8759384 | -9.22E-06 | -0.0005104 |
| -0.3936849 | -0.8001187 | -0.7795224 | -1.8885357 | -1.8888257 | -0.0002899 | -0.0001528 |
| -1.620887 | -1.5424495 | -1.6822848 | -2.8245703 | -2.8248971 | -0.0003268 | -0.0008653 |
| 1.5823369 | 0.62937021 | 1.03862247 | -1.800611 | -1.80826 | -0.0076489 | -0.0010927 |
| -0.3798428 | 0.27782965 | -0.0208673 | -2.5838192 | -2.5838122 | 6.99E-06 | -0.0007542 |
| -4.3247809 | -4.7960291 | -3.4924547 | -1.2078091 | -1.2190548 | -0.0112457 | -0.0013696 |
| -4.4384862 | -7.4530441 | -6.4231363 | -2.1910526 | -2.1990291 | -0.0079765 | -0.0006308 |

| | | | | | | |
|---|---|---|---|---|---|---|
| -0.6562409 | -0.6703563 | -0.7360406 | -2.384999 | -2.3853022 | -0.0003033 | -0.0001401 |
| -1.5688841 | -1.69502 | -1.6755469 | -2.3253816 | -2.3259077 | -0.0005261 | -0.0010149 |
| -1.7386281 | -1.6598442 | -1.5056281 | -0.395776 | -0.4038902 | -0.0081142 | -0.0012731 |
| 0.01655006 | 0.1099143 | -0.5457093 | -0.8023987 | -0.8194511 | -0.0170524 | -0.0004284 |
| -0.7642269 | -0.2917649 | -0.5264173 | -2.5667629 | -2.5668302 | -6.72E-05 | -0.0001223 |
| -0.8873892 | -0.7450366 | -0.8852436 | -2.2919094 | -2.2925844 | -0.000675 | -0.0002391 |
| -2.6396699 | -1.0405874 | -1.7800812 | -2.7558541 | -2.7590039 | -0.0031497 | -0.0038283 |
| -3.8659444 | -5.3304291 | -5.0546007 | -2.0284977 | -2.0309455 | -0.0024478 | -0.0007293 |
| 1.42928171 | 1.38349843 | 1.35561023 | -0.2244553 | -0.2331983 | -0.008743 | -0.0009964 |
| -2.1480474 | -1.7657032 | -1.8394908 | -1.1813762 | -1.1832558 | -0.0018796 | -0.0007694 |
| -2.2226329 | -2.110909 | -1.8679884 | -0.7631508 | -0.7791584 | -0.0160076 | -0.0007264 |
| -1.2495551 | -1.017611 | -1.1352083 | -2.7814051 | -2.7816683 | -0.0002632 | -0.0019296 |
| -0.1300213 | -0.354815 | -0.030758 | -1.9123719 | -1.9135797 | -0.0012078 | -0.000489 |
| -0.7514477 | -0.805732 | -0.7882985 | -2.6932898 | -2.6937375 | -0.0004478 | -0.0008713 |
| 1.29161978 | 1.7785027 | 1.19511638 | -2.1100459 | -2.1126962 | -0.0026503 | -0.0004746 |
| 3.3343811 | 3.57640696 | 2.88630402 | -0.5898793 | -0.5854887 | 0.00439066 | -0.00111 |
| -0.1010175 | -0.1610286 | -0.072606 | -1.7902986 | -1.7904934 | -0.0001948 | -0.0036306 |
| 0.36170626 | 1.07791471 | 0.98832357 | -0.6803576 | -0.6801455 | 0.00021207 | -0.0005967 |
| -5.7639422 | -6.6132431 | -6.5860677 | -0.404587 | -0.4174858 | -0.0128988 | -0.0008232 |
| -11.4651 | -7.2797232 | -7.4930115 | -1.8046286 | -1.8129375 | -0.0083089 | -0.000192 |
| 1.10016966 | 0.95928097 | 0.97034612 | -2.7098217 | -2.7093768 | 0.00044489 | -0.0004331 |
| -0.7046304 | -0.9005609 | -0.7760308 | -2.0976923 | -2.0980802 | -0.0003879 | -0.000732 |
| 0.41664147 | -0.0654802 | 0.27111402 | -2.2926853 | -2.2956789 | -0.0029936 | -0.0008141 |
| -0.3967218 | 0.11407375 | -0.1060116 | -2.7141697 | -2.7144375 | -0.0002677 | -0.0020741 |
| 1.34253931 | 1.35241842 | 1.42151718 | -1.8678291 | -1.8667567 | 0.00107241 | -0.0011286 |
| 1.38107693 | 0.74437213 | 1.00627573 | -2.0903466 | -2.0899489 | 0.00039768 | -0.0004406 |
| -0.4327168 | -0.5208907 | -0.3724325 | -1.5852001 | -1.5846328 | 0.00056732 | -0.0001764 |
| 0.72480071 | 0.91533339 | 0.81232648 | -1.5849191 | -1.5839924 | 0.00092673 | -0.0019806 |
| -0.5511553 | -0.8345459 | -0.5665606 | -1.0515405 | -1.0524323 | -0.0008918 | -0.0078093 |
| 0.11467957 | 0.28035474 | 0.40993185 | -2.8792279 | -2.8790174 | 0.00021052 | -0.0028936 |
| -2.2306151 | -2.230165 | -2.4413946 | -2.4047716 | -2.4116988 | -0.0069273 | -0.0046596 |
| -2.6132379 | -1.1239836 | -1.733267 | -1.901294 | -1.9082716 | -0.0069776 | -0.0024509 |
| -2.6262378 | -1.9658203 | -2.0999456 | -2.6026208 | -2.6043173 | -0.0016966 | -0.001006 |
| -4.198786 | -3.419034 | -3.9992145 | -2.0059595 | -2.0073183 | -0.0013588 | -0.0003509 |
| -8.2430077 | -6.0823288 | -6.7795181 | -0.7643179 | -0.7776198 | -0.013302 | -0.0014993 |
| 0.2153945 | -0.7246742 | -0.7459025 | -2.2971981 | -2.2999175 | -0.0027194 | -0.0036725 |
| 0.52475023 | 0.95301723 | 0.63095632 | -2.7087479 | -2.7089474 | -0.0001996 | -0.0049323 |
| -0.4371519 | -0.8772736 | -0.5763328 | -1.1457454 | -1.1461378 | -0.0003924 | -0.00385 |
| 1.26971841 | 1.09459424 | 0.26288729 | -0.6630339 | -0.6634661 | -0.0004322 | -0.0023618 |
| -1.0168853 | -0.4095063 | -1.4637415 | -2.4025819 | -2.4905078 | -0.0879259 | -0.0143452 |
| -0.941566 | -0.7440608 | -1.1522214 | -1.2593038 | -1.2600718 | -0.0007679 | -0.0034009 |
| -3.9611142 | -2.7997947 | -2.4233299 | -1.3030779 | -1.3278699 | -0.024792 | -0.0047587 |
| -3.1576948 | -1.9027839 | -2.4500975 | -2.8268925 | -2.8274632 | -0.0005706 | -0.000285 |
| -0.8326426 | -1.363367 | -1.2191618 | -2.2656732 | -2.2722168 | -0.0065436 | -0.0098031 |
| -9.7148529 | -8.07964 | -8.0986077 | -2.5702849 | -2.5714065 | -0.0011215 | -0.0002154 |

| | | | | | | |
|---|---|---|---|---|---|---|
| 0.22286415 | 0.49024272 | 0.4282424 | -2.1623139 | -2.1628852 | -0.0005713 | -0.0006546 |
| 0.19310951 | 0.15502954 | 0.21471567 | -1.555312 | -1.553465 | 0.00184703 | -0.0076339 |
| -4.5254266 | -1.8958659 | -3.2865103 | -2.6664093 | -2.6625795 | 0.00382972 | -0.0064486 |
| -4.3685389 | -1.3610159 | -2.349659 | -2.4759398 | -2.4762437 | -0.000304 | -0.0027897 |
| -2.1492066 | -1.758683 | -1.7908817 | -2.5726719 | -2.5825138 | -0.0098419 | -0.0068269 |
| -4.203599 | -6.8711472 | -4.8810501 | -1.731895 | -1.7688733 | -0.0369784 | -0.0080941 |
| -4.1040622 | -2.7953854 | -3.1986808 | -2.7318687 | -2.7315955 | 0.00027323 | -0.001442 |
| -1.9928007 | -1.4233594 | -1.5167274 | -1.4991884 | -1.5014813 | -0.0022929 | -0.0013827 |
| 0.40025067 | -0.1435544 | -0.2133675 | -2.7364883 | -2.7366975 | -0.0002092 | -0.0009115 |
| -4.3155861 | -7.0069952 | -5.7715175 | -1.6574389 | -1.6758369 | -0.018398 | -0.0019137 |
| -2.9049873 | -3.2321749 | -2.9652534 | -0.7928526 | -0.8019601 | -0.0091074 | -0.0022182 |
| -1.7006652 | -4.1080794 | -2.7524863 | -2.2240021 | -2.2265272 | -0.0025251 | -0.0021911 |
| 0.16485667 | 0.14114881 | -0.0800446 | -1.810487 | -1.8110745 | -0.0005875 | -0.000555 |
| -2.3856194 | -0.6831086 | -1.675115 | -2.3034539 | -2.317271 | -0.0138171 | -0.0116704 |
| 1.28282714 | 1.06241131 | 1.21799736 | -2.2714884 | -2.2707949 | 0.00069356 | -0.0009557 |
| -4.6642516 | -3.4159904 | -3.3932464 | -0.7087655 | -0.7236556 | -0.0148901 | -0.0008195 |
| -0.3019433 | 0.25358176 | -0.2378491 | -1.8733365 | -1.8753945 | -0.002058 | -0.0002024 |
| -4.6713233 | -6.7150278 | -6.1704878 | -0.7292565 | -0.7845294 | -0.0552729 | -0.0046153 |
| 0.17449856 | 0.54939318 | 0.6264236 | -2.090394 | -2.0910819 | -0.0006878 | -0.0024117 |
| -1.0892725 | -1.8396884 | 0.45613148 | -1.6192336 | -1.6222254 | -0.0029918 | -0.0028602 |
| 3.30004859 | 2.82429695 | 1.36099629 | -1.9563199 | -1.9709864 | -0.0146664 | -0.004834 |
| -1.1564598 | -6.7654719 | -4.9771377 | -1.6300399 | -1.6392945 | -0.0092546 | -0.0004827 |
| 0.38431287 | -1.2216781 | -0.3741817 | -1.7362792 | -1.7363468 | -6.76E-05 | -0.0011963 |
| -1.2805347 | -2.0513659 | 0.07053902 | -1.5117934 | -1.5204347 | -0.0086414 | -0.0010794 |
| 1.05782509 | 0.41904306 | 0.45974326 | -1.612076 | -1.6126258 | -0.0005498 | -0.0025995 |
| -3.3727834 | 3.72115076 | -0.8925992 | -0.9363843 | -0.9418461 | -0.0054618 | -0.0016558 |
| -0.5792577 | -0.7530828 | -0.3809685 | -1.5975564 | -1.5993738 | -0.0018175 | -0.0030882 |
| -1.4562526 | -3.3681781 | -2.9148042 | -1.7316196 | -1.8388416 | -0.107222 | -0.0115544 |
| -2.2301958 | -2.2484815 | -2.3556369 | -2.2294941 | -2.2320147 | -0.0025206 | -0.0018041 |
| -3.1568241 | -4.2848892 | -3.5688637 | -0.9112919 | -0.9390146 | -0.0277227 | -0.0040133 |
| -107.49349 | -128.1077 | -105.0199 | -1.2715396 | -1.6595203 | -0.3879807 | -0.0033868 |
| -1.307209 | -1.2439823 | -1.3487561 | -2.5377605 | -2.5385914 | -0.0008309 | -0.0009272 |
| -7.3564936 | -6.8489521 | -6.8119447 | -2.1932523 | -2.1962414 | -0.0029891 | -0.0004643 |
| -1.2270478 | 0.31645584 | 0.63033597 | -1.3789129 | -1.3764441 | 0.00246882 | -0.0008012 |
| -5.810576 | -6.9529123 | -6.0910221 | -2.5056469 | -2.5360823 | -0.0304353 | -0.001944 |
| 2.07273293 | 0.07648802 | 0.84492793 | -2.5690446 | -2.5674832 | 0.0015614 | -0.004085 |
| -2.7760634 | -4.0293334 | -4.0705259 | -1.3306938 | -1.3392339 | -0.00854 | -0.002708 |
| -2.3147235 | -4.4774399 | -2.9031701 | -1.4093742 | -1.4757183 | -0.066344 | -0.0066924 |
| -1.3713305 | -2.2189646 | -1.7111036 | -1.4424231 | -1.4455113 | -0.0030882 | -0.0016971 |
| -7.0291567 | -2.6159074 | -4.5243847 | -1.7106652 | -1.7298713 | -0.019206 | -0.0035441 |
| -3.3948965 | -9.2833233 | -6.6838114 | -1.0996158 | -1.1747061 | -0.0750903 | -0.0044843 |
| -2.1674237 | -2.9615726 | -2.306877 | -1.016259 | -1.0256501 | -0.0093912 | -0.0037495 |
| 0.74298525 | 0.62338018 | 0.65335102 | -1.6637988 | -1.6633899 | 0.00040889 | -0.0002357 |
| -0.1521804 | -0.577852 | -0.4745027 | -1.0737329 | -1.0712173 | 0.00251555 | -0.0030626 |
| -0.9590929 | -0.8024549 | -1.0540116 | -1.868283 | -1.8720051 | -0.0037221 | -0.0053258 |

| | | | | | | |
|---|---|---|---|---|---|---|
| -5.0453138 | -5.6006739 | -4.7826105 | -2.0939085 | -2.1209853 | -0.0270767 | -0.0030846 |
| -2.7402031 | -2.6702995 | -2.6578893 | -2.6030049 | -2.6132755 | -0.0102706 | -0.0075179 |
| -2.9047743 | -2.8215916 | -2.7673519 | -2.5834891 | -2.5844515 | -0.0009624 | -0.0010565 |
| -0.8303703 | -0.7531052 | -0.6345312 | -2.6190763 | -2.635056 | -0.0159798 | -0.0188481 |
| 0.26411688 | -0.4594135 | -0.960176 | -0.625618 | -0.629649 | -0.004031 | -0.0008165 |
| -1.1392541 | -5.2290206 | -2.7711075 | -1.0231057 | -1.04497 | -0.0218643 | -0.0044064 |
| -2.7026178 | -5.1769638 | -3.9020602 | -1.1593493 | -1.1625319 | -0.0031825 | -0.0013888 |
| -2.4297855 | -2.4546261 | -3.121691 | -1.1299925 | -1.1418147 | -0.0118222 | -0.0025195 |
| -4.833293 | -5.2657738 | -5.1410336 | -1.5068064 | -1.5141732 | -0.0073668 | -0.0002366 |
| -3.9851012 | -1.842185 | -2.392815 | -0.5846918 | -0.5948538 | -0.0101619 | -0.0024287 |
| -1.0129852 | -1.0108922 | -0.9538259 | -2.3054521 | -2.306545 | -0.0010929 | -0.0014373 |
| -1.9533125 | -0.4431186 | -1.6147203 | -0.7994099 | -0.8121139 | -0.012704 | -0.0035201 |
| -8.0044844 | -8.9856025 | -8.8033139 | -1.2358335 | -1.2861961 | -0.0503626 | -0.0050398 |
| -0.4123557 | -0.2036502 | -0.4018985 | -1.5628524 | -1.5631301 | -0.0002778 | -0.0026381 |
| 0.3826704 | -4.3886728 | -0.5883942 | -1.0827104 | -1.0959533 | -0.013243 | -0.002233 |
| 0.77722788 | 0.18438816 | 0.27771363 | -1.6991129 | -1.7043962 | -0.0052834 | -0.0005903 |
| 0.94872355 | 0.93875456 | 0.95080981 | -2.7047558 | -2.704422 | 0.00033379 | -0.0019164 |
| 0.32814145 | 0.82941914 | 0.51803551 | -1.4120241 | -1.4103539 | 0.00167024 | -0.0045715 |
| -7.3812494 | -7.4806833 | -7.8916025 | -1.615037 | -1.6159534 | -0.0009165 | -0.0001543 |
| -1.2308048 | -4.4849615 | -3.5020154 | -1.037958 | -1.0570759 | -0.0191178 | -0.0044718 |
| -2.4463949 | -2.4524086 | -2.8878928 | -1.9898657 | -1.9952734 | -0.0054077 | -0.0044424 |
| 0.30676794 | 0.58040786 | 0.66115875 | -2.7897675 | -2.7894421 | 0.00032539 | -0.0014838 |
| 0.34718013 | 0.48662376 | 0.45605526 | -2.2413723 | -2.2408814 | 0.0004909 | -0.0009528 |
| -5.9941878 | -8.5816746 | -7.2842211 | -2.5799966 | -2.5830004 | -0.0030038 | -0.0016142 |
| -0.8643084 | -1.2371316 | -0.8003712 | -2.1217833 | -2.1270878 | -0.0053046 | -0.0006628 |
| -2.2709236 | -2.7229078 | -2.3653877 | -1.3706673 | -1.3713292 | -0.0006618 | -0.0012595 |
| -2.4736443 | -2.3142912 | -2.5470602 | -2.7952516 | -2.7962441 | -0.0009925 | -0.0015873 |
| -6.0902429 | -6.6226838 | -5.7547239 | -0.7299464 | -0.7447966 | -0.0148502 | -0.0010242 |
| -1.2131445 | -1.1599206 | -1.2883782 | -2.8554606 | -2.8667202 | -0.0112596 | -0.0172031 |
| -19.103286 | -22.611027 | -21.769066 | -1.6979629 | -1.7786453 | -0.0806824 | -0.0036032 |
| -1.0254707 | -0.9023461 | -1.1525918 | -1.5678272 | -1.5722091 | -0.0043819 | -0.0004622 |
| -0.5052357 | -0.176899 | -0.1218282 | -2.8068364 | -2.8076649 | -0.0008285 | -0.0152691 |
| -2.2688756 | -2.3988218 | -1.3672938 | -1.3390691 | -1.3438258 | -0.0047567 | -0.002773 |
| -6.2931614 | -4.3797112 | -4.4935088 | -0.6878729 | -0.711997 | -0.0241241 | -0.0011151 |
| -0.2440243 | 0.28285193 | -0.3629233 | -2.0641401 | -2.0713553 | -0.0072153 | -0.0004678 |
| -3.2656515 | -4.1169379 | -4.0032879 | -2.8298216 | -2.8311685 | -0.0013469 | -0.0009984 |
| 0.84328973 | -4.2168665 | -1.3801975 | -0.9108763 | -0.9261461 | -0.0152698 | -0.001748 |
| -128.87037 | -128.11277 | -126.17597 | -0.3375464 | -0.9698723 | -0.6323259 | -0.003031 |
| -0.1047206 | 0.06229305 | -0.8351684 | -1.7941946 | -1.7949868 | -0.0007923 | -0.0032998 |
| -6.0016465 | -4.1379766 | -5.120905 | -1.7135504 | -1.7275637 | -0.0140133 | -0.0035416 |
| -0.8082616 | 1.83379102 | 1.33517199 | -1.3014973 | -1.3041871 | -0.0026897 | -0.0003354 |
| -2.8445742 | -1.7854595 | -1.8228813 | -1.9242253 | -1.9301236 | -0.0058982 | -0.003566 |
| 2.19447899 | 0.57143664 | -1.191736 | -0.1609344 | -0.2094409 | -0.0485065 | -0.0009121 |
| 0.43472409 | -0.7940162 | 0.7481338 | -2.1752539 | -2.173965 | 0.00128889 | -0.0010377 |
| -2.9525819 | -2.3149772 | -3.0369925 | -1.7005007 | -1.7045522 | -0.0040514 | -0.0035056 |

| | | | | | | |
|---|---|---|---|---|---|---|
| 0.35748005 | 0.36226749 | 0.3311748 | -0.8549162 | -0.854968 | -5.17E-05 | -0.0013073 |
| -8.4462194 | -6.8625393 | -7.6565288 | -1.2921261 | -1.2974169 | -0.0052909 | -0.0010403 |
| -5.8544345 | -6.3814282 | -6.8725912 | -0.9318267 | -0.9496374 | -0.0178107 | -0.0025491 |
| -1.414782 | -0.2779303 | 0.25968874 | -1.3875917 | -1.3920323 | -0.0044405 | -0.0001732 |
| -2.2616789 | -4.7971661 | -4.2173298 | -2.3393931 | -2.3680139 | -0.0286207 | -0.0070983 |
| -1.0940385 | -0.9154923 | -0.9639557 | -2.8226457 | -2.8273356 | -0.0046899 | -0.0116321 |
| -0.4682846 | -0.8268218 | -0.5842708 | -0.7849018 | -0.78578 | -0.0008782 | -0.0005582 |
| 1.28646076 | 1.17887652 | 1.30921049 | -2.0321579 | -2.0286369 | 0.00352097 | -0.0079792 |
| -1.6020032 | -2.371191 | -2.2890264 | -1.6902108 | -1.6933913 | -0.0031805 | -0.0027725 |
| -5.5919008 | -5.524847 | -5.8004529 | -0.5846902 | -0.5917346 | -0.0070444 | -0.0009218 |
| 0.34393215 | 0.28278828 | 0.22555633 | -1.9777673 | -1.9775752 | 0.00019217 | -0.0004488 |
| 1.23197627 | -2.1683428 | -1.0552079 | -1.9267681 | -1.9290183 | -0.0022502 | -0.0047225 |
| 0.3246057 | -1.0516975 | 0.21975746 | -2.1878345 | -2.186754 | 0.00108051 | -0.0031494 |
| -3.9545243 | -2.0049 | -2.6872048 | -2.1569519 | -2.1537931 | 0.00315881 | -0.0022968 |
| -1.6496406 | 0.02392173 | -1.5931598 | -1.7581744 | -1.7668996 | -0.0087252 | -0.003661 |
| -3.2434169 | -2.7698967 | -2.8069056 | -2.4958332 | -2.4983692 | -0.0025361 | -0.0007669 |
| -6.2264471 | -5.4895682 | -5.0409506 | -0.3226091 | -0.3348524 | -0.0122433 | -0.0009626 |
| -11.762614 | -10.970958 | -9.5943238 | -1.8123957 | -1.8374991 | -0.0251034 | -0.0050599 |
| -0.0617785 | 0.0468421 | 0.06433535 | -2.4387619 | -2.43891 | -0.0001481 | -0.0008606 |
| -0.1961366 | -1.2904345 | -0.9223419 | -1.4430548 | -1.4498997 | -0.0068449 | -0.0004586 |
| -0.138432 | -1.3382342 | -0.9819655 | -1.3093754 | -1.3211951 | -0.0118197 | -0.0037957 |
| -0.4589953 | -0.3417187 | -0.6671841 | -1.9119438 | -1.9195867 | -0.0076429 | -0.0040029 |
| -2.5204463 | -5.3926136 | -3.5830334 | -2.254102 | -2.2566617 | -0.0025597 | -0.0018337 |
| 0.10195374 | -1.33122 | -0.8717079 | -0.9308188 | -0.9467039 | -0.0158851 | -0.0051373 |
| -0.1735971 | -0.3792266 | 0.04061983 | -2.1845577 | -2.1837261 | 0.0008316 | -0.0008378 |
| -0.461978 | -3.9079695 | -2.0444449 | -1.2975103 | -1.3042524 | -0.0067421 | -0.0022045 |
| -2.8060472 | -3.3837008 | -2.2142421 | -1.9323171 | -1.9303155 | 0.00200164 | -0.0024651 |
| 0.13892722 | -0.2724719 | -0.1926079 | -2.8424814 | -2.8424337 | 4.77E-05 | -0.0153109 |
| -2.1806593 | -2.3332083 | -1.6009003 | -2.1324255 | -2.1377831 | -0.0053575 | -0.0113538 |
| -0.2782168 | -1.6116567 | -1.230264 | -2.7365932 | -2.7436402 | -0.0070469 | -0.0091425 |
| -3.6174276 | -2.9355857 | -3.4104517 | -2.314043 | -2.3178394 | -0.0037963 | -0.0020566 |
| -5.1760111 | -5.2643495 | -4.5798962 | -2.6232674 | -2.6483188 | -0.0250514 | -0.0138288 |
| 0.79021001 | 0.94604468 | 1.12474294 | -2.0201998 | -2.0177305 | 0.0024693 | -0.0019866 |
| -6.6237494 | -7.8101298 | -7.0635768 | -2.354799 | -2.3840244 | -0.0292253 | -0.0054091 |
| 0.68667054 | -1.4669094 | -0.0518438 | -1.6010056 | -1.6080021 | -0.0069965 | -0.002918 |
| -7.2165767 | -6.8571609 | -5.0758713 | -2.321537 | -2.3406637 | -0.0191267 | -0.006654 |
| 1.64882392 | 0.64897591 | 1.10481492 | -0.8766817 | -0.8789144 | -0.0022327 | -0.0006666 |
| 0.60895443 | 0.54324794 | 0.62766981 | -1.918468 | -1.917048 | 0.00142002 | -0.0004901 |
| -7.6458561 | -6.5377567 | -9.4419091 | -1.6728463 | -1.704108 | -0.0312617 | -0.0014429 |
| -1.5970416 | -5.3085904 | -3.944415 | -0.7548214 | -0.7593246 | -0.0045032 | -0.0008745 |
| 0.29592967 | 0.41203976 | 0.30301213 | -2.2846229 | -2.2829618 | 0.00166106 | -0.0049209 |
| -0.2166405 | -0.6519442 | -0.5660796 | -2.7484498 | -2.7496769 | -0.0012271 | -0.0064729 |
| -2.3819175 | -4.6350632 | -3.6519903 | -2.1508224 | -2.1529903 | -0.0021679 | -0.0044687 |
| 1.4001441 | 0.35393977 | 0.90609045 | -2.5233634 | -2.5182936 | 0.00506973 | -0.0012714 |
| -0.8064245 | -2.5105548 | -3.0157044 | -1.8471661 | -1.8552625 | -0.0080965 | -0.0018213 |

| | | | | | | |
|---|---|---|---|---|---|---|
| -7.3415766 | -7.6969404 | -7.5882146 | -0.4942313 | -0.5173606 | -0.0231293 | -0.0014707 |
| -0.7065618 | -1.4059675 | -1.0739068 | -0.7580977 | -0.7629569 | -0.0048591 | -0.0040797 |
| -2.9426041 | -2.9656974 | -3.2676118 | -1.508136 | -1.5109671 | -0.0028311 | -0.0053911 |
| -6.7668822 | -7.9584861 | -6.1632365 | -1.6348563 | -1.6469376 | -0.0120813 | -0.0016515 |
| -16.983474 | -13.798294 | -13.113465 | -1.8091878 | -1.8241901 | -0.0150024 | -0.0017179 |
| -9.7825663 | -8.9660666 | -8.6395489 | -1.5764204 | -1.586695 | -0.0102745 | -0.0013859 |
| -3.3262391 | -4.5628614 | -3.7875323 | -1.86554 | -1.8689982 | -0.0034581 | -0.0019536 |
| -2.8388801 | -1.053471 | -1.7760437 | -1.1153804 | -1.1442008 | -0.0288204 | -0.0031746 |
| 0.07714272 | -0.2991745 | -0.1678387 | -2.8118901 | -2.81195 | -5.98E-05 | -0.0036912 |
| -2.6853848 | -2.834975 | -2.4458703 | -2.7897038 | -2.8092644 | -0.0195606 | -0.0195448 |
| -14.418182 | -13.30355 | -14.149345 | -2.2575438 | -2.277674 | -0.0201302 | -0.0031427 |
| -0.9542956 | -2.2753758 | -1.2112449 | -2.7250328 | -2.7263868 | -0.001354 | -0.0037343 |
| -1.7473464 | -3.1823332 | -2.7742955 | -1.0500017 | -1.0574179 | -0.0074161 | -0.0047532 |
| -0.4607263 | -2.6988969 | -1.1133574 | -2.2638605 | -2.2641749 | -0.0003145 | -0.0038228 |
| 0.06515312 | 1.01754069 | 0.76195176 | -0.5741461 | -0.5770394 | -0.0028933 | -0.0007085 |
| -1.767663 | -3.762629 | -2.2570946 | -2.6613748 | -2.6724255 | -0.0110507 | -0.0052794 |
| 2.36032677 | 1.06264758 | 1.40307879 | -2.3941185 | -2.3922098 | 0.00190878 | -0.0034036 |
| -0.7151117 | -2.1185088 | -2.556064 | -2.0960419 | -2.1186736 | -0.0226316 | -0.0028385 |
| 0.00432897 | -0.2155092 | -0.2821694 | -1.8869226 | -1.8890473 | -0.0021247 | -0.0049963 |
| 0.67276812 | 0.36811829 | 0.45127902 | -2.3463917 | -2.3446593 | 0.00173235 | -0.0011091 |
| -2.6430392 | -3.097169 | -1.9657244 | -0.7762986 | -0.7792996 | -0.003001 | -0.0007486 |
| -6.1436429 | -9.3349669 | -6.3104141 | -2.4442749 | -2.458293 | -0.0140181 | -0.0038399 |
| -1.4959817 | -2.0512834 | -1.5525906 | -2.5911572 | -2.5941393 | -0.0029821 | -0.0042622 |
| -0.0832593 | -0.3657403 | -0.1847118 | -2.7860556 | -2.7861443 | -8.87E-05 | -0.0006312 |
| -0.5164886 | -0.0977716 | -0.6271649 | -1.6377439 | -1.635866 | 0.0018779 | -0.0025543 |
| -7.3889475 | -8.5213081 | -8.226829 | -1.2358335 | -1.2813443 | -0.0455108 | -0.0050398 |
| -0.6085937 | -0.8633142 | -0.6477713 | -1.6998267 | -1.7027812 | -0.0029545 | -0.0003983 |
| -3.5571704 | -3.5236659 | -3.1777927 | -2.2634575 | -2.266777 | -0.0033195 | -0.0009154 |
| -1.3425043 | -1.3174872 | -1.4500366 | -2.081449 | -2.0826161 | -0.0011671 | -0.0009021 |
| -4.3012929 | -8.2089043 | -4.5780653 | -1.7670945 | -1.770296 | -0.0032015 | -0.0020114 |
| -25.175454 | -27.155024 | -26.109085 | -1.0858319 | -1.1960535 | -0.1102216 | -0.0023634 |
| 0.50933814 | 0.43047547 | 0.41888576 | -1.9857186 | -1.985683 | 3.56E-05 | -0.0008894 |
| 0.05722022 | 0.03999591 | 0.20475769 | -1.3447802 | -1.3450925 | -0.0003123 | -0.0024244 |
| -1.5117682 | -2.0122793 | -1.8404254 | -2.3676059 | -2.3722882 | -0.0046823 | -0.0078558 |
| -5.9169312 | -10.003619 | -7.5668859 | -0.5747371 | -0.6191748 | -0.0444377 | -0.001822 |
| -106.02804 | -114.16421 | -107.02047 | -1.5061132 | -2.1493754 | -0.6432623 | -0.0043621 |
| -3.6838861 | -6.4979715 | -5.1304367 | -1.4885432 | -1.4969618 | -0.0084187 | -0.0017709 |
| 2.54649282 | 1.58894253 | 2.87432451 | -0.9010628 | -0.9034739 | -0.0024111 | -0.0012167 |
| -2.8419206 | -2.6952474 | -2.8233568 | -1.1301471 | -1.1467583 | -0.0166112 | -0.0040987 |
| -1.668551 | -1.7224839 | -1.5127281 | -1.199859 | -1.2087296 | -0.0088706 | -0.0020206 |
| -3.2600563 | -2.9501629 | -2.877847 | -1.4747841 | -1.4950552 | -0.0202711 | -0.0059174 |
| 2.44047999 | 0.2952683 | -0.3082887 | -2.5127833 | -2.5201645 | -0.0073812 | -0.0034051 |
| 0.77240658 | -0.395999 | 1.64868536 | -1.4904218 | -1.4896916 | 0.00073016 | -0.0013535 |
| -0.5430293 | -2.286631 | -2.479389 | -1.5626541 | -1.5678431 | -0.0051889 | -0.0013557 |
| -7.5986013 | -8.7185955 | -6.9764103 | -2.589622 | -2.5919991 | -0.002377 | -0.0027046 |

| | | | | | | |
|---|---|---|---|---|---|---|
| -14.873096 | -14.66458 | -11.809537 | -2.3709903 | -2.3820713 | -0.011081 | -0.0023475 |

cid inserted into the human protein

| esm1v_t33_6 | esm1v_t33_6 | esm1v_t33_6 | esm1v_t33_6 | esm1v_t33_6 | ESM1v_score | Progen2_sco |
|---|---|---|---|---|---|---|
| -0.2573504 | -0.403203 | -0.3247275 | -0.3934729 | -0.4694467 | -0.3696401 | -0.0007968 |
| 0.82550871 | 0.53857362 | 0.78209579 | 0.99020743 | 0.67837489 | 0.76295209 | -0.0007968 |
| 0.91868591 | 0.90725231 | 1.32707882 | 0.93670702 | 1.08547473 | 1.03503976 | -0.000277 |
| -1.3377624 | -3.856348 | 1.01349902 | 0.53111696 | 2.24252367 | -0.2813941 | -0.0021024 |
| -2.0701008 | 0.0801363 | -0.8836777 | -2.1304393 | -1.0663109 | -1.2140785 | -0.0007384 |
| 0.99665952 | 1.16737616 | 1.16878903 | 1.02376151 | 1.61297214 | 1.19391167 | -0.0007384 |
| 1.1577208 | 1.10524678 | 1.30216694 | 1.11317873 | 1.31516623 | 1.1986959 | -0.0007384 |
| 0.73767233 | 0.61408377 | 0.84226203 | 0.69767094 | 0.44919157 | 0.66817613 | -0.0007384 |
| 0.10439253 | -0.334408 | -0.7170265 | -1.0550277 | 0.35244322 | -0.3299253 | -0.0007384 |
| 0.28086209 | -0.6550756 | -0.2048247 | 0.36655092 | 0.20016932 | -0.0024636 | -0.0015624 |
| -2.4155712 | -1.7078419 | -3.764401 | -3.5880611 | -2.6590242 | -2.8269799 | -0.0015624 |
| 1.60308111 | -0.1050355 | 1.07572317 | 1.6154567 | 0.73676848 | 0.98519878 | -0.0008758 |
| 0.85679126 | 1.22993469 | 1.44941199 | 1.30061698 | 0.75970519 | 1.11929202 | -0.0008758 |
| 0.04656982 | 0.11401248 | 0.13528442 | -0.0659285 | 0.18799305 | 0.08358626 | -0.0011935 |
| -0.0683944 | -0.3821321 | -0.415621 | -0.1194344 | 0.12756991 | -0.1716024 | -0.0013059 |
| -0.3639712 | -0.2654626 | -0.2806673 | -0.42801 | -0.40815 | -0.3492522 | -0.0013059 |
| -4.084363 | -3.6836491 | -3.7463689 | -4.1550164 | -2.4416256 | -3.6222046 | -0.0013059 |
| -0.5921111 | 0.03453112 | -0.1965227 | -0.4741843 | -0.3851135 | -0.3226801 | -0.0014783 |
| -1.594053 | -1.6952028 | -1.7931223 | -1.8013296 | -2.0142384 | -1.7795892 | -0.0014783 |
| -2.3350463 | -2.666384 | -1.6664863 | -2.3757291 | -3.1803713 | -2.4448034 | -0.0022161 |
| -1.0704551 | -0.9688852 | -0.4693587 | -1.2883253 | -0.7413538 | -0.9076756 | -0.0022161 |
| -2.118293 | -3.2811284 | -0.9427342 | -5.4864731 | -3.315594 | -3.0288445 | -0.0006234 |
| -3.0332339 | -3.6654406 | -3.1444159 | -2.186157 | -3.5101595 | -3.1078814 | -0.0024214 |
| -1.3459244 | -1.2072797 | -3.3777874 | -2.3177443 | -2.5406156 | -2.1578702 | -0.0027125 |
| -2.772145 | -2.5405588 | -3.2560544 | -3.1123028 | -2.8039701 | -2.8970062 | -0.0006863 |
| -0.388196 | 0.26396394 | -0.2317982 | -0.2450728 | -0.1341143 | -0.1470435 | -0.0006863 |
| 0.37836361 | 1.03092146 | 1.00250149 | 1.08759308 | 0.58527136 | 0.8169302 | -0.0006863 |
| -0.3627799 | -0.7766726 | -0.8380466 | -0.5370634 | -0.7978978 | -0.662492 | -0.0006863 |
| 2.04314733 | 2.00789142 | 2.06076193 | 1.63619661 | 1.61465716 | 1.87253089 | -0.0006863 |
| -1.8555259 | -3.0942686 | -2.054451 | -2.1327977 | -2.4035823 | -2.3081251 | -0.0006863 |
| -0.5141683 | -0.8238664 | -0.3952651 | -0.6065021 | -1.5895264 | -0.7858656 | -0.0004268 |
| 0.43975163 | 0.71985459 | 0.45349574 | 0.48049629 | 0.51287413 | 0.52129447 | -0.0002644 |
| 1.87662911 | 1.79801106 | 1.51487195 | 1.70657909 | 1.47363138 | 1.67394452 | -0.0002644 |
| -0.8856511 | -0.7668102 | -1.0536077 | -0.9658525 | -1.0438187 | -0.943148 | -0.0023933 |
| -0.4117746 | -0.5848274 | -0.5510826 | -0.2744212 | -0.5391381 | -0.4722488 | -0.0010594 |
| -1.2891409 | -0.9603539 | -1.0540528 | -1.5996066 | -1.7187452 | -1.3243799 | -0.0005104 |
| -0.8238075 | -1.0777402 | -0.8022606 | -0.3936849 | -0.8001187 | -0.7795224 | -0.0001528 |
| -0.7702012 | -0.8233471 | -0.7103572 | -0.6442447 | -0.6242483 | -0.7144797 | -0.0008653 |
| -1.011023 | -1.1487644 | -0.7843947 | -0.9766424 | -0.9182012 | -0.9678051 | -0.0008653 |
| 0.9227612 | 0.53028393 | 1.52836013 | 1.5823369 | 0.62937021 | 1.03862247 | -0.0010927 |
| -0.209163 | -0.0698998 | -0.2894797 | 0.0266521 | 0.03703809 | -0.1009705 | -0.0007542 |
| 0.3997789 | -0.1688409 | 0.33528131 | -0.4064949 | 0.24079156 | 0.08010321 | -0.0007542 |

| | | | | | | |
|---|---|---|---|---|---|---|
| -2.4076977 | -3.9751675 | -1.9585981 | -4.3247809 | -4.7960291 | -3.4924547 | -0.0013696 |
| 1.31018639 | 1.06321263 | 0.81775141 | 0.87575436 | 1.20758915 | 1.05489879 | -0.0006308 |
| -0.4727108 | 0.25639343 | -0.3732591 | 0.44538617 | -0.1661341 | -0.0620649 | -0.0006308 |
| -0.003674 | 0.53141165 | 0.5947268 | 0.9756515 | 0.99110568 | 0.61784432 | -0.0006308 |
| 0.61012423 | 0.38798392 | 1.08474433 | 0.34617877 | 0.38140512 | 0.56208727 | -0.0006308 |
| 2.34340668 | 2.73054147 | 2.29316497 | 1.60862017 | 1.94572806 | 2.18429227 | -0.0006308 |
| 0.28283453 | -0.1998248 | 0.21872288 | -0.1152139 | -1.0211687 | -0.16693 | -0.0006308 |
| -6.0816031 | -6.2158017 | -5.4097004 | -5.479764 | -6.2456145 | -5.8864967 | -0.0006308 |
| -0.5400994 | -0.6426079 | -1.1899271 | 0.26528597 | -0.8645394 | -0.5943776 | -0.0006308 |
| -2.3681903 | -2.741431 | -2.3685443 | -1.9992664 | -2.6350756 | -2.4225015 | -0.0006308 |
| 1.02769637 | 0.7766068 | 0.78228831 | 1.10315907 | 1.30813837 | 0.99957778 | -0.0006308 |
| 0.03165627 | -0.4119477 | -0.6457288 | -0.3745842 | -0.8927104 | -0.458663 | -0.0006308 |
| -1.2761049 | -1.8743973 | -1.3853741 | -1.5625942 | -1.638715 | -1.5474371 | -0.0006308 |
| -0.9139857 | -1.8010247 | -0.4516673 | -0.5270996 | 0.17694736 | -0.703366 | -0.0006308 |
| -0.6091868 | -0.7558767 | -0.9885426 | -0.6562409 | -0.6703563 | -0.7360406 | -0.0001401 |
| 0.20188308 | 0.0846405 | 0.26231456 | 0.19758439 | 0.16035676 | 0.18135586 | -0.0010149 |
| -1.873435 | -1.9873533 | -1.8018804 | -1.7664685 | -1.8553767 | -1.8569028 | -0.0010149 |
| -1.3525302 | -1.103029 | -1.674109 | -1.7386281 | -1.6598442 | -1.5056281 | -0.0012731 |
| -0.8546109 | -1.4207979 | -0.579602 | 0.01655006 | 0.1099143 | -0.5457093 | -0.0004284 |
| -0.4927467 | -0.5282075 | -0.5551405 | -0.7642269 | -0.2917649 | -0.5264173 | -0.0001223 |
| -0.7331851 | -0.9560962 | -1.1045108 | -0.8873892 | -0.7450366 | -0.8852436 | -0.0002391 |
| -1.4340262 | -2.0318582 | -1.7542645 | -2.6396699 | -1.0405874 | -1.7800812 | -0.0038283 |
| -5.4280338 | -5.4428792 | -5.2057171 | -3.8659444 | -5.3304291 | -5.0546007 | -0.0007293 |
| 1.3450551 | 1.3088398 | 1.31137609 | 1.42928171 | 1.38349843 | 1.35561023 | -0.0009964 |
| -1.81148 | -1.956079 | -1.5161443 | -2.1480474 | -1.7657032 | -1.8394908 | -0.0007694 |
| -1.0912147 | -2.7924948 | -1.1226907 | -2.2226329 | -2.110909 | -1.8679884 | -0.0007264 |
| -1.1254244 | -1.1552045 | -1.1282465 | -1.2495551 | -1.017611 | -1.1352083 | -0.0019296 |
| -0.3555288 | 0.89571595 | -0.2091408 | -0.1300213 | -0.354815 | -0.030758 | -0.000489 |
| -0.9343429 | -0.9558482 | -0.4941218 | -0.7514477 | -0.805732 | -0.7882985 | -0.0008713 |
| 0.47321177 | 1.0257287 | 1.40651894 | 1.29161978 | 1.7785027 | 1.19511638 | -0.0004746 |
| 2.9765172 | 2.60068655 | 1.94352829 | 3.3343811 | 3.57640696 | 2.88630402 | -0.00111 |
| -0.1374671 | 0.23674631 | -0.200263 | -0.1010175 | -0.1610286 | -0.072606 | -0.0036306 |
| 1.47853601 | 1.62599659 | 0.39746428 | 0.36170626 | 1.07791471 | 0.98832357 | -0.0005967 |
| -5.5464697 | -8.3445654 | -6.662118 | -5.7639422 | -6.6132431 | -6.5860677 | -0.0008232 |
| -0.8586073 | -3.8129082 | -2.1909041 | -5.7438536 | -4.4063931 | -3.4025332 | -0.000192 |
| -4.6155863 | -5.8537874 | -1.388441 | -5.7212467 | -2.8733301 | -4.0904783 | -0.000192 |
| 1.21791148 | 1.16629887 | 0.40806961 | 1.10016966 | 0.95928097 | 0.97034612 | -0.0004331 |
| -0.8758252 | -0.6908281 | -0.7083094 | -0.7046304 | -0.9005609 | -0.7760308 | -0.000732 |
| 0.384624 | -0.0712914 | 0.69107628 | 0.41664147 | -0.0654802 | 0.27111402 | -0.0008141 |
| -0.3256001 | 0.13989592 | -0.0617058 | -0.3967218 | 0.11407375 | -0.1060116 | -0.0020741 |
| 1.60552382 | 1.62463307 | 1.18247128 | 1.34253931 | 1.35241842 | 1.42151718 | -0.0011286 |
| 0.99247289 | 0.94544125 | 0.96801543 | 1.38107693 | 0.74437213 | 1.00627573 | -0.0004406 |
| -0.3699148 | -0.4106171 | -0.1280229 | -0.4327168 | -0.5208907 | -0.3724325 | -0.0001764 |
| 0.99456882 | 0.84844613 | 0.57848334 | 0.72480071 | 0.91533339 | 0.81232648 | -0.0019806 |
| -0.6977532 | -0.3070512 | -0.4422973 | -0.5511553 | -0.8345459 | -0.5665606 | -0.0078093 |

| | | | | | | |
|---|---|---|---|---|---|---|
| 0.98642802 | 0.22716093 | 0.44103599 | 0.11467957 | 0.28035474 | 0.40993185 | -0.0028936 |
| -1.5063818 | -4.2727447 | -1.9670663 | -2.2306151 | -2.230165 | -2.4413946 | -0.0046596 |
| -1.069279 | -2.0257127 | -1.8341221 | -2.6132379 | -1.1239836 | -1.733267 | -0.0024509 |
| -1.6662159 | -1.9866046 | -1.1459079 | -2.0809505 | -1.593886 | -1.694713 | -0.001006 |
| -0.0907745 | -0.2313349 | -0.7868321 | -0.5452873 | -0.3719343 | -0.4052326 | -0.001006 |
| -1.0885417 | -2.6324737 | -1.8966358 | -1.8404968 | -1.4661651 | -1.7848626 | -0.0003509 |
| -1.8909032 | -2.348175 | -2.2939966 | -2.008863 | -2.5055459 | -2.2094967 | -0.0003509 |
| 0.68409395 | -1.251549 | 0.33992887 | -0.3494263 | 0.55267692 | -0.0048551 | -0.0003509 |
| -7.4285874 | -5.5052896 | -6.6383772 | -8.2430077 | -6.0823288 | -6.7795181 | -0.0014993 |
| -0.4982073 | -1.971648 | -0.7503774 | 0.2153945 | -0.7246742 | -0.7459025 | -0.0036725 |
| 0.44449639 | 0.70872116 | 0.52379656 | 0.52475023 | 0.95301723 | 0.63095632 | -0.0049323 |
| -0.6234378 | -0.5053231 | -0.4384776 | -0.4371519 | -0.8772736 | -0.5763328 | -0.00385 |
| -0.1064973 | -0.5103407 | -0.4330382 | 1.26971841 | 1.09459424 | 0.26288729 | -0.0023618 |
| -1.7150087 | -1.8943981 | -2.2829092 | -1.0168853 | -0.4095063 | -1.4637415 | -0.0143452 |
| -1.073509 | -0.7533555 | -2.2486157 | -0.941566 | -0.7440608 | -1.1522214 | -0.0034009 |
| -1.4660363 | -2.1823616 | -1.7073426 | -3.9611142 | -2.7997947 | -2.4233299 | -0.0047587 |
| -0.523237 | -0.5913653 | -0.4889903 | -0.8359246 | -0.6678839 | -0.6214802 | -0.000285 |
| -1.3116894 | -1.1815622 | -1.6524695 | -1.3361597 | -1.3326108 | -1.3628983 | -0.000285 |
| -0.3898616 | -0.573807 | -0.2580135 | -0.7590725 | 0.1817832 | -0.3597943 | -0.000285 |
| 0.28111243 | 0.30287218 | 0.19574928 | 0.18386316 | 0.32672501 | 0.25806441 | -0.000285 |
| -0.2756684 | -0.4620006 | -0.2610779 | -0.4104011 | -0.4107974 | -0.3639891 | -0.000285 |
| -0.828141 | -1.5238621 | -1.5477962 | -0.8326426 | -1.363367 | -1.2191618 | -0.0098031 |
| -1.3414202 | -1.2037535 | -1.3382568 | -1.4250081 | -1.3649616 | -1.33468 | -0.0002154 |
| -0.2796354 | 1.70106792 | 0.2400806 | -0.1641514 | 0.8546164 | 0.47039561 | -0.0002154 |
| -0.7349186 | -1.0366659 | -1.1105306 | -1.3279625 | -1.4551693 | -1.1330494 | -0.0002154 |
| 0.27833939 | 0.16887581 | -0.1399431 | -0.3906972 | -0.3239031 | -0.0814656 | -0.0002154 |
| -2.2575898 | -1.8146232 | -1.9100218 | -2.3591652 | -2.2748432 | -2.1232486 | -0.0002154 |
| -0.7232544 | -0.7241542 | -1.0640538 | -0.7970233 | -0.4100697 | -0.7437111 | -0.0002154 |
| -1.6594524 | -1.476434 | -1.3978131 | -1.6730106 | -1.5122719 | -1.5437964 | -0.0002154 |
| -1.2613952 | -2.0002937 | -1.6126995 | -1.5778346 | -1.5930376 | -1.6090521 | -0.0002154 |
| 0.20470119 | 0.76865196 | 0.45475197 | 0.22286415 | 0.49024272 | 0.4282424 | -0.0006546 |
| 0.21666217 | 0.08516645 | 0.42361069 | 0.19310951 | 0.15502954 | 0.21471567 | -0.0076339 |
| -1.1168704 | -1.3851764 | -0.7852433 | 0.10236621 | -0.3239784 | -0.7017805 | -0.0064486 |
| -1.0591819 | -0.3314834 | -0.5211048 | -0.7691183 | -0.3461418 | -0.605406 | -0.0064486 |
| 1.76544893 | 0.31532741 | 0.88591433 | 0.1763134 | 1.9613266 | 1.02086613 | -0.0064486 |
| -2.2130618 | -4.5596476 | -1.0061802 | -4.0349879 | -3.1870723 | -3.0001899 | -0.0064486 |
| -1.592164 | -3.5060763 | -0.9204998 | -4.3685389 | -1.3610159 | -2.349659 | -0.0027897 |
| -1.646282 | -1.6387383 | -1.7614985 | -2.1492066 | -1.758683 | -1.7908817 | -0.0068269 |
| -4.9350829 | -4.6727209 | -3.7227006 | -4.203599 | -6.8711472 | -4.8810501 | -0.0080941 |
| 0.17382431 | -0.7545717 | -1.1376626 | -1.2622429 | 0.50571799 | -0.494987 | -0.001442 |
| -1.9373932 | -3.7141252 | -1.8605983 | -2.8491993 | -3.6108129 | -2.7944258 | -0.001442 |
| 0.12636375 | 0.06282949 | -0.052623 | 0.00738001 | 0.30970955 | 0.09073195 | -0.001442 |
| -1.4521277 | -0.8729548 | -1.8423941 | -1.9928007 | -1.4233594 | -1.5167274 | -0.0013827 |
| -0.2378113 | -0.4351761 | -0.6505461 | 0.40025067 | -0.1435544 | -0.2133675 | -0.0009115 |
| -3.4643683 | -3.6770854 | -0.0088155 | -2.0464454 | -2.2155576 | -2.2824544 | -0.0019137 |

| | | | | | | |
|---|---|---|---|---|---|---|
| -2.5941606 | -3.1193924 | -0.6069102 | -2.2413807 | -3.34975 | -2.3823188 | -0.0019137 |
| -1.2095255 | -1.402293 | -1.4524555 | -0.02776 | -1.4416876 | -1.1067443 | -0.0019137 |
| -3.8308415 | -2.072629 | -2.7856345 | -2.9049873 | -3.2321749 | -2.9652534 | -0.0022182 |
| -0.2384636 | -4.8447151 | -2.8705082 | -1.7006652 | -4.1080794 | -2.7524863 | -0.0021911 |
| -0.8164368 | 0.14317155 | -0.0329633 | 0.16485667 | 0.14114881 | -0.0800446 | -0.000555 |
| -1.778103 | -1.8341045 | -1.6946393 | -2.3856194 | -0.6831086 | -1.675115 | -0.0116704 |
| -0.1487114 | -0.0578523 | -0.0605175 | 0.00221419 | -0.1591926 | -0.0848119 | -0.0009557 |
| 1.45357752 | 1.30203438 | 1.25621772 | 1.28061295 | 1.22160387 | 1.30280929 | -0.0009557 |
| -4.106988 | -4.4136496 | -4.3054581 | -5.1738591 | -3.3192956 | -4.2638501 | -0.0008195 |
| -0.3969104 | 0.37459636 | 1.18114424 | 0.11515117 | -0.7502418 | 0.10474792 | -0.0008195 |
| 1.2314496 | 1.06811714 | 0.48170877 | 0.39445639 | 0.65354705 | 0.76585579 | -0.0008195 |
| 0.56413507 | 1.10427761 | 1.1227529 | 0.66626191 | 1.00375962 | 0.89223742 | -0.0002024 |
| 0.00060081 | 0.08373308 | 0.62861967 | 0.44499612 | 0.73775673 | 0.37914128 | -0.0002024 |
| -1.5074456 | -1.6562717 | -1.481286 | -1.4132013 | -1.4879346 | -1.5092278 | -0.0002024 |
| -1.7494121 | -1.4172349 | -1.0491004 | -0.2037463 | -1.611186 | -1.2061359 | -0.0046153 |
| -5.6779423 | -4.8096914 | -4.7627068 | -4.467577 | -5.1038418 | -4.9643518 | -0.0046153 |
| 0.06659174 | 2.00424266 | 0.33739185 | 0.17449856 | 0.54939318 | 0.6264236 | -0.0024117 |
| -1.0402355 | 0.77635956 | 0.92967844 | -1.0228345 | -1.1105108 | -0.2935086 | -0.0028602 |
| 0.73162389 | 1.58808017 | 1.34257269 | 1.28648317 | 1.42746234 | 1.27524445 | -0.0028602 |
| 0.06423402 | 0.25758839 | 1.52238989 | 0.12843633 | -1.0095345 | 0.19262283 | -0.0028602 |
| -0.2720742 | -0.0160732 | -0.6745257 | -1.4813575 | -1.1471055 | -0.7182272 | -0.0028602 |
| -0.1230119 | -1.8239163 | 0.07028127 | 0.5938623 | 0.42836571 | -0.1708838 | -0.004834 |
| 0.3540864 | 0.56828117 | 1.63491535 | 2.70618629 | 2.39593124 | 1.53188009 | -0.004834 |
| -6.0643835 | -5.5000553 | -5.3993177 | -1.1564598 | -6.7654719 | -4.9771377 | -0.0004827 |
| -0.3234982 | -0.4100828 | -0.299962 | 0.38431287 | -1.2216781 | -0.3741817 | -0.0011963 |
| 1.96778595 | -0.3398738 | 2.05668354 | -1.2805347 | -2.0513659 | 0.07053902 | -0.0010794 |
| -0.001193 | 0.51978636 | 0.30325484 | 1.05782509 | 0.41904306 | 0.45974326 | -0.0025995 |
| -1.0322211 | -1.134699 | -0.983325 | -1.3239701 | -0.8894187 | -1.0727268 | -0.0016558 |
| -0.4440515 | 0.53425717 | -1.7513239 | -2.0488133 | 4.61056948 | 0.18012757 | -0.0016558 |
| -0.1640313 | -0.0263252 | -0.3821456 | -0.5792577 | -0.7530828 | -0.3809685 | -0.0030882 |
| -1.7364466 | -6.5457563 | -1.4673872 | -1.4562526 | -3.3681781 | -2.9148042 | -0.0115544 |
| -2.1557336 | -2.4135861 | -2.7301874 | -2.2301958 | -2.2484815 | -2.3556369 | -0.0018041 |
| -4.2752261 | -3.8108242 | -2.316555 | -3.1568241 | -4.2848892 | -3.5688637 | -0.0040133 |
| -2.927218 | -6.3819103 | -2.0228009 | -2.4516475 | -2.4998913 | -3.2566936 | -0.0033868 |
| 0.35143518 | 1.04720688 | -0.0998292 | -0.0414665 | 0.23308778 | 0.29808683 | -0.0033868 |
| -0.2329434 | -0.4314157 | -0.2386904 | -0.3534415 | -0.2829611 | -0.3078904 | -0.0033868 |
| -2.2820354 | -1.3772814 | -1.8341738 | -2.3825636 | -2.2889102 | -2.0329929 | -0.0033868 |
| -0.7565331 | -0.3529062 | -0.6620264 | -0.1526101 | -0.3000662 | -0.4448284 | -0.0033868 |
| -0.2442886 | -0.3261175 | -0.3739964 | -0.7306551 | -0.0994198 | -0.3548955 | -0.0033868 |
| -0.1418679 | -0.2623248 | -0.7832441 | -0.5111279 | -0.2767527 | -0.3950635 | -0.0033868 |
| -2.0058 | -2.442697 | -2.0722299 | -2.4452877 | -1.7547302 | -2.144149 | -0.0033868 |
| -0.5436962 | -0.5780485 | -0.5289106 | -0.4665446 | -0.819685 | -0.587377 | -0.0033868 |
| 1.4710288 | 1.62851787 | 1.12688637 | 1.650244 | -0.5886769 | 1.05760002 | -0.0033868 |
| -1.6178364 | -1.7932274 | -1.4943557 | -1.1133702 | -1.3992902 | -1.483616 | -0.0033868 |
| -0.108194 | -1.7610106 | -0.0307124 | -1.5797476 | -1.1833539 | -0.9326037 | -0.0033868 |

| | | | | | | |
|---|---|---|---|---|---|---|
| 1.41480684 | 1.13779998 | 1.03395844 | 2.20550609 | 1.53015757 | 1.46444578 | -0.0033868 |
| -1.6690149 | -0.0987043 | -1.0871491 | -1.687379 | -1.4710298 | -1.2026554 | -0.0033868 |
| -4.9143023 | -0.79281 | -4.5600328 | -3.5820613 | -2.9584172 | -3.3615247 | -0.0033868 |
| 0.1909647 | -0.1070147 | -3.0984979 | -1.5866194 | -0.8326902 | -1.0867715 | -0.0033868 |
| -0.9233505 | -2.201112 | -0.4639351 | -1.41081 | -0.5484233 | -1.1095262 | -0.0033868 |
| -0.461472 | -0.2645822 | -0.7470293 | -1.0347731 | -0.4711919 | -0.5958097 | -0.0033868 |
| -1.1694291 | 0.00981092 | -1.1460264 | -0.689703 | -1.1956713 | -0.8382038 | -0.0033868 |
| -1.8996257 | -1.525299 | -0.811192 | -1.8528426 | -2.8319931 | -1.7841905 | -0.0033868 |
| -3.2505593 | -4.3239713 | -0.5485197 | -0.5133533 | -5.6375923 | -2.8547992 | -0.0033868 |
| 1.56365895 | 0.90411401 | 0.58311772 | 0.64954436 | -1.0545216 | 0.5291827 | -0.0033868 |
| 1.29984188 | 1.90385628 | 3.06222057 | -0.0165215 | -0.4498501 | 1.15990944 | -0.0033868 |
| -0.2560306 | 2.12715673 | 0.6530447 | 2.35456729 | -2.5431399 | 0.46711965 | -0.0033868 |
| 0.09106731 | -0.5653816 | 1.53702593 | 0.91800213 | -0.0038581 | 0.39537114 | -0.0033868 |
| -3.2081444 | -7.3299837 | -3.2075696 | -3.1495438 | -3.847379 | -4.1485241 | -0.0033868 |
| -0.8982928 | -1.586437 | -1.5261594 | -1.8657498 | 1.75859368 | -0.8236091 | -0.0033868 |
| 3.79354763 | -2.4951122 | 1.01152706 | 1.23526466 | -3.4609995 | 0.01684554 | -0.0033868 |
| -1.8638363 | 1.29348898 | 0.42656708 | 1.5166769 | -1.6865511 | -0.0627309 | -0.0033868 |
| -0.8242168 | -2.5707581 | -0.7648211 | -1.5478978 | -3.4541063 | -1.83236 | -0.0033868 |
| -0.8946137 | -2.4315627 | 0.22764528 | -0.6753521 | -1.2250239 | -0.9997814 | -0.0033868 |
| -2.5704336 | -4.3121033 | -1.904521 | -3.0886188 | 0.53905034 | -2.2673253 | -0.0033868 |
| -0.7664163 | -0.6815586 | -0.1865249 | -3.0052586 | -2.1977148 | -1.3674946 | -0.0033868 |
| 0.22503114 | -0.2108068 | -0.950233 | 0.04071283 | 0.37310386 | -0.1044384 | -0.0033868 |
| -0.4543215 | -2.8970656 | -0.1415128 | 0.89916861 | -1.9432919 | -0.9074046 | -0.0033868 |
| 1.3522923 | 0.51047993 | 0.9960171 | -0.6410904 | 1.10491252 | 0.66452229 | -0.0033868 |
| 0.33493996 | -1.4001684 | 0.34953499 | -1.0555625 | 0.59785056 | -0.2346811 | -0.0033868 |
| -4.2095661 | -8.6302881 | -6.1511927 | -5.3084445 | -8.7255001 | -6.6049983 | -0.0033868 |
| -1.9104847 | -4.087903 | -2.1832948 | -2.8026228 | -1.4392152 | -2.4847041 | -0.0033868 |
| 1.33052778 | -3.5089333 | 0.15319777 | -2.6573582 | -4.9731565 | -1.9311445 | -0.0033868 |
| -0.410809 | -1.3724511 | -0.0643144 | -2.5215704 | -0.3129618 | -0.9364213 | -0.0033868 |
| -2.2960658 | -1.9605802 | -4.8868971 | -5.6328988 | -2.4841714 | -3.4521227 | -0.0033868 |
| -1.1990936 | -1.4700196 | -0.097003 | 0.45921946 | -2.5290103 | -0.9671814 | -0.0033868 |
| 4.65649509 | 3.05967212 | 3.26616955 | 3.6471715 | 5.52156591 | 4.03021483 | -0.0033868 |
| -1.6527772 | -1.9338102 | -0.6568952 | -3.6294162 | -2.2800293 | -2.0305856 | -0.0033868 |
| 2.61621332 | 3.55654764 | 2.99704504 | 2.7645793 | 3.76427794 | 3.13973265 | -0.0033868 |
| -0.2075658 | 2.99667692 | 3.63888288 | -1.4736562 | 2.12458897 | 1.41578536 | -0.0033868 |
| -1.4226604 | -2.7687306 | -3.1352143 | -3.519228 | -1.9407263 | -2.5573119 | -0.0033868 |
| 0.31522655 | 0.37472391 | -0.1155763 | 0.04307556 | -0.5276113 | 0.0179677 | -0.0033868 |
| -1.0730602 | 2.3942914 | 2.89105487 | 1.60231709 | 3.85636711 | 1.93419406 | -0.0033868 |
| -3.2437234 | 0.41725737 | -1.1176605 | -1.3742943 | -1.2469702 | -1.3130782 | -0.0033868 |
| -0.3847837 | -1.8223481 | -0.878253 | -0.9314803 | -0.9263696 | -0.988647 | -0.0033868 |
| 0.33845425 | -3.1459069 | 1.31405926 | 2.36136341 | 0.53087902 | 0.2797698 | -0.0033868 |
| -5.6661615 | 1.57718742 | -4.4127016 | -3.8181062 | -2.7900767 | -3.0219717 | -0.0033868 |
| 0.10431147 | -0.9490871 | -3.2161436 | -2.0073476 | -2.8769736 | -1.7890481 | -0.0033868 |
| -3.4227347 | -2.9603829 | -3.3555765 | -5.0480509 | -5.1485081 | -3.9870506 | -0.0033868 |
| -0.088335 | -5.5566978 | 0.5453496 | 1.13190246 | 4.13923025 | 0.03428988 | -0.0033868 |

| | | | | | | |
|---|---|---|---|---|---|---|
| -4.0826092 | -3.0320492 | -3.7575035 | -3.6502724 | -2.5732079 | -3.4191284 | -0.0033868 |
| -1.9421213 | 0.96164751 | -2.7345564 | 0.00359344 | -3.4131944 | -1.4249262 | -0.0033868 |
| 2.05980945 | -2.6307638 | 0.97848177 | 0.09203291 | 1.79149377 | 0.45821083 | -0.0033868 |
| -3.4700017 | -2.241622 | -1.1179032 | -2.0693569 | -2.2066174 | -2.2211002 | -0.0033868 |
| -0.4026303 | -0.1335316 | -0.5626431 | -0.3927217 | -0.4115949 | -0.3806243 | -0.0033868 |
| 2.2414217 | 0.79538727 | 2.4698348 | 2.05454493 | -2.6868482 | 0.97486811 | -0.0033868 |
| -5.0740862 | -3.4092286 | -3.7686727 | -1.7416748 | -3.0601158 | -3.4107556 | -0.0033868 |
| 1.97353745 | 0.77726269 | 5.03601551 | -0.9772611 | 2.98505688 | 1.95892229 | -0.0033868 |
| -1.0216737 | 1.46964312 | -2.4703484 | -0.7936263 | -0.0698786 | -0.5771768 | -0.0033868 |
| -0.737216 | -3.6918359 | -2.425477 | -1.5849619 | -0.9196315 | -1.8718245 | -0.0033868 |
| 2.01859045 | -0.2124929 | -0.6605358 | 3.1570859 | 0.55280161 | 0.97108984 | -0.0033868 |
| -1.8145334 | 0.14483452 | -1.0067096 | -2.3848872 | -3.0926604 | -1.6307912 | -0.0033868 |
| -0.4486111 | -4.6552353 | 1.5897398 | -0.9346535 | 0.1284759 | -0.8640568 | -0.0033868 |
| 1.51715064 | -2.3527384 | 0.65112352 | -2.6931744 | -1.4267131 | -0.8608703 | -0.0033868 |
| -0.6443977 | -1.1035872 | 2.20106196 | -0.0046272 | 2.09802675 | 0.50929532 | -0.0033868 |
| 2.51761866 | 2.17791176 | 5.09290743 | 2.35997581 | 0.48206276 | 2.52609528 | -0.0033868 |
| -1.9161768 | -3.4009306 | -2.5912836 | -3.7250175 | -2.8978863 | -2.906259 | -0.0033868 |
| 0.72045922 | 1.40288115 | 1.94018507 | 3.2989738 | 1.94420052 | 1.86133995 | -0.0033868 |
| 0.32434022 | -0.991447 | -1.3922989 | -1.6230254 | -1.2140889 | -0.979304 | -0.0033868 |
| -0.3447771 | -1.490221 | -1.4769182 | 0.62501425 | -0.6166716 | -0.6607147 | -0.0033868 |
| 2.22137213 | -0.7236124 | -0.4256291 | -0.6429319 | -1.027647 | -0.1196897 | -0.0033868 |
| -0.8496151 | -2.1834722 | -3.5290205 | -0.7901264 | -3.0291772 | -2.0762823 | -0.0033868 |
| 2.19102621 | -0.4330025 | 1.51573849 | -1.1612997 | -1.0180597 | 0.21888056 | -0.0033868 |
| -0.6751013 | -2.3897467 | -2.2302365 | -2.2765884 | 0.03182697 | -1.5079692 | -0.0033868 |
| -0.4769082 | -1.3385668 | -1.7190796 | -1.6138096 | -2.8882046 | -1.6073138 | -0.0033868 |
| -0.4334702 | -1.4498954 | 1.49100006 | -0.6546855 | -4.3752775 | -1.0844657 | -0.0033868 |
| -0.2733898 | -3.5561032 | -2.6789656 | -1.65905 | -4.8004136 | -2.5935844 | -0.0033868 |
| 0.95879674 | 0.57232332 | 2.59132481 | 2.82020783 | -1.4826007 | 1.0920104 | -0.0033868 |
| -0.8946126 | -3.2273691 | -2.643507 | -3.075655 | -1.0483334 | -2.1778954 | -0.0033868 |
| -2.0394287 | -3.4101219 | -1.0500903 | -1.378859 | -1.9713949 | -1.969979 | -0.0033868 |
| -4.8381872 | -6.5876684 | -6.2323685 | -3.0709074 | -4.6878543 | -5.0833972 | -0.0033868 |
| -1.1566045 | -2.1776972 | -2.5381298 | -3.3455729 | -2.6793542 | -2.3794717 | -0.0033868 |
| 0.34793234 | 0.35655379 | 0.17630041 | 0.71933198 | 0.46574235 | 0.41317217 | -0.0033868 |
| 0.12428522 | -1.3337777 | 1.0465579 | -0.2812794 | -2.4879448 | -0.5864318 | -0.0033868 |
| 0.34860563 | 1.03600883 | 0.88010317 | -1.1520691 | -1.781502 | -0.1337707 | -0.0033868 |
| -3.2765903 | -2.778852 | -6.2065277 | -3.3208022 | -1.4085486 | -3.3982642 | -0.0033868 |
| -3.7803712 | -4.7544227 | -5.7826395 | -4.8650837 | -0.6808934 | -3.9726821 | -0.0033868 |
| -2.1912584 | -1.3088112 | -3.3575044 | -2.3828654 | -1.5662937 | -2.1613466 | -0.0033868 |
| -2.8980465 | -3.6353996 | -2.0900884 | -3.8775704 | -1.7298064 | -2.8461823 | -0.0033868 |
| -5.2535825 | -0.6824794 | -5.4591274 | -4.0155287 | -4.8464551 | -4.0514346 | -0.0033868 |
| 0.0491209 | 0.57964045 | 0.51912379 | -0.2279778 | 1.89622176 | 0.56322583 | -0.0033868 |
| 2.66411352 | 1.67969155 | 0.26443338 | 0.86889458 | -0.196167 | 1.05619321 | -0.0033868 |
| -1.1373439 | -1.1535647 | -1.7794018 | -1.1816273 | -5.0940328 | -2.0691941 | -0.0033868 |
| -1.7591791 | -1.1563957 | -2.5616846 | -3.0438399 | -3.6714211 | -2.4385041 | -0.0033868 |
| -0.8668418 | -1.1523694 | -0.2699575 | 0.94280839 | -0.2584194 | -0.3209559 | -0.0033868 |

| | | | | | | |
|---|---|---|---|---|---|---|
| -0.7982843 | -1.5088271 | -0.6827996 | -0.6389611 | -1.2185432 | -0.969483 | -0.0033868 |
| 0.56934953 | -0.3176966 | -0.3444622 | -0.3268499 | -0.5113549 | -0.1862028 | -0.0033868 |
| -1.2945476 | -1.4244909 | -1.4735508 | -1.307209 | -1.2439823 | -1.3487561 | -0.0009272 |
| -1.9486926 | -0.8945217 | -0.9092088 | -1.4595028 | -1.0144024 | -1.2452657 | -0.0004643 |
| -2.2946501 | -2.9401608 | -3.1518896 | -2.860465 | -2.3686173 | -2.7231565 | -0.0004643 |
| -2.7684729 | -2.406534 | -2.073103 | -2.6907649 | -2.786263 | -2.5450275 | -0.0004643 |
| -0.4738305 | -0.4553154 | -0.3602397 | -0.0838513 | -0.2948494 | -0.3336173 | -0.0004643 |
| 0.51756144 | -0.0075712 | 0.31235051 | -0.2619095 | -0.38482 | 0.03512225 | -0.0004643 |
| 1.00102615 | 0.89562678 | 2.1656189 | -1.2270478 | 0.31645584 | 0.63033597 | -0.0008012 |
| -6.0374746 | -4.8205347 | -6.8336129 | -5.810576 | -6.9529123 | -6.0910221 | -0.001944 |
| -0.0716581 | 1.8702805 | 0.27679634 | 2.07273293 | 0.07648802 | 0.84492793 | -0.004085 |
| -1.0670984 | -2.7494829 | -0.5661173 | 0.09904861 | -1.0610895 | -1.0689479 | -0.002708 |
| -2.027442 | -4.1521478 | -2.9849443 | -2.8751121 | -2.9682438 | -3.001578 | -0.002708 |
| -1.296418 | -2.6797767 | -0.1433933 | -1.6720285 | -0.8648934 | -1.331302 | -0.0066924 |
| 0.13362312 | -2.1856136 | -1.5521085 | -0.642695 | -3.6125464 | -1.5718681 | -0.0066924 |
| -1.6917436 | -2.1122954 | -1.1611841 | -1.3713305 | -2.2189646 | -1.7111036 | -0.0016971 |
| -0.8233645 | -0.1615038 | -0.584655 | -0.3871479 | -0.708688 | -0.5330719 | -0.0035441 |
| -2.7065492 | -5.2662306 | -3.4345565 | -6.6420088 | -1.9072194 | -3.9913129 | -0.0035441 |
| -4.6273098 | -4.8349648 | -6.3608336 | -3.6102471 | -5.977459 | -5.0821629 | -0.0044843 |
| -0.5516944 | -0.3844447 | 0.77714801 | 0.38057995 | 0.94526005 | 0.23336978 | -0.0044843 |
| -1.723819 | -1.350462 | -1.6844568 | -0.1652293 | -4.2511244 | -1.8350183 | -0.0044843 |
| -1.6707393 | -2.4547577 | -2.2798915 | -2.1674237 | -2.9615726 | -2.306877 | -0.0037495 |
| 0.65104842 | 0.49916744 | 0.75017381 | 0.74298525 | 0.62338018 | 0.65335102 | -0.0002357 |
| -0.6423109 | -0.591311 | -0.4088593 | -0.1521804 | -0.577852 | -0.4745027 | -0.0030626 |
| -1.5309982 | -0.9629097 | -1.0146022 | -0.9590929 | -0.8024549 | -1.0540116 | -0.0053258 |
| 1.04819632 | 1.04981661 | 0.91579103 | 0.73548913 | 1.26220202 | 1.00229902 | -0.0030846 |
| -0.4740829 | 1.26776981 | 0.6100471 | -1.3320951 | -0.912045 | -0.1680812 | -0.0030846 |
| -1.883094 | 0.34337282 | -0.5300527 | -0.3737948 | -1.0204182 | -0.6927974 | -0.0030846 |
| -5.1847086 | -5.130681 | -5.2994394 | -4.074913 | -4.9304128 | -4.924031 | -0.0030846 |
| -2.397162 | -2.4938312 | -2.9879508 | -2.7402031 | -2.6702995 | -2.6578893 | -0.0075179 |
| -1.0563784 | -1.1540277 | -1.8029313 | -0.9594036 | -1.1401111 | -1.2225704 | -0.0010565 |
| -1.8069673 | -0.8377529 | -1.4523362 | -1.9453707 | -1.6814805 | -1.5447815 | -0.0010565 |
| -0.7659194 | 0.38224959 | -1.2055107 | -0.8303703 | -0.7531052 | -0.6345312 | -0.0188481 |
| 1.99824238 | -4.7420707 | -1.8617549 | 0.26411688 | -0.4594135 | -0.960176 | -0.0008165 |
| 1.01037574 | -5.1445374 | -3.353101 | -1.1392541 | -5.2290206 | -2.7711075 | -0.0044064 |
| 0.06052899 | 0.37677097 | 0.86558545 | 0.6095475 | 0.10240221 | 0.40296702 | -0.0013888 |
| -5.5824585 | -6.8288732 | -0.5222733 | -3.3121653 | -5.279366 | -4.3050272 | -0.0013888 |
| -3.9060349 | -3.6183014 | -3.199707 | -2.4297855 | -2.4546261 | -3.121691 | -0.0025195 |
| -5.3199463 | -5.5289068 | -4.7572479 | -4.833293 | -5.2657738 | -5.1410336 | -0.0002366 |
| -2.203002 | 0.61644739 | -3.033298 | -2.754703 | -1.0017424 | -1.6752596 | -0.0024287 |
| -0.2597229 | -1.0962915 | -0.1609216 | -1.2303982 | -0.8404427 | -0.7175554 | -0.0024287 |
| -1.4396503 | -0.7083044 | -0.5972972 | -1.0129852 | -1.0108922 | -0.9538259 | -0.0014373 |
| -1.5738397 | -1.5959144 | -2.5074162 | -1.9533125 | -0.4431186 | -1.6147203 | -0.0035201 |
| -2.0151777 | -2.0118716 | -2.2209585 | -2.1378193 | -2.2236371 | -2.1218928 | -0.0050398 |
| -3.3174901 | -3.6875267 | -3.7080355 | -3.4148145 | -3.5167003 | -3.5289134 | -0.0050398 |

| | | | | | | |
|---|---|---|---|---|---|---|
| -3.8710048 | -4.2862844 | -3.9673114 | -4.5588794 | -3.5622969 | -4.0491554 | -0.0050398 |
| 0.99447191 | 1.65436161 | 1.62858248 | 2.20472527 | 1.53480351 | 1.60338895 | -0.0050398 |
| -0.7178521 | -1.0299088 | -0.4704767 | -0.0976965 | -1.2177718 | -0.7067412 | -0.0050398 |
| -0.5973182 | -0.4023066 | -0.393862 | -0.4123557 | -0.2036502 | -0.4018985 | -0.0026381 |
| 0.10704422 | -1.5747212 | 2.53170824 | 0.3826704 | -4.3886728 | -0.5883942 | -0.002233 |
| 0.23244834 | 0.12210536 | 0.07239842 | 0.77722788 | 0.18438816 | 0.27771363 | -0.0005903 |
| 1.12145567 | 0.8928721 | 0.85224319 | 0.94872355 | 0.93875456 | 0.95080981 | -0.0019164 |
| 0.34839201 | 0.60318923 | 0.48103571 | 0.32814145 | 0.82941914 | 0.51803551 | -0.0045715 |
| -7.816638 | -4.9054847 | -5.077589 | -5.1620221 | -5.0791368 | -5.6081741 | -0.0001543 |
| -2.190794 | -2.5249393 | -2.0806346 | -2.2192273 | -2.4015465 | -2.2834283 | -0.0001543 |
| -2.4206614 | -6.0840712 | -3.289578 | -1.2308048 | -4.4849615 | -3.5020154 | -0.0044718 |
| -1.9796941 | -3.6165521 | -3.9444141 | -2.4463949 | -2.4524086 | -2.8878928 | -0.0044424 |
| 0.22408032 | -0.0877249 | 0.35253501 | -0.0050354 | -0.0770237 | 0.08136625 | -0.0014838 |
| 0.60595942 | 0.73664689 | 0.58712125 | 0.31180334 | 0.6574316 | 0.5797925 | -0.0014838 |
| 0.40176558 | 0.70151019 | 0.34319663 | 0.34718013 | 0.48662376 | 0.45605526 | -0.0009528 |
| -11.777493 | -3.523026 | -6.5447245 | -5.9941878 | -8.5816746 | -7.2842211 | -0.0016142 |
| 1.67059159 | -3.2441642 | -0.3268435 | -0.8643084 | -1.2371316 | -0.8003712 | -0.0006628 |
| -0.6231794 | -4.0543909 | -2.1555367 | -2.2709236 | -2.7229078 | -2.3653877 | -0.0012595 |
| -1.1816076 | -0.8850204 | -0.6919706 | -0.7911043 | -0.7446269 | -0.858866 | -0.0015873 |
| -1.6733484 | -1.7787547 | -1.7366636 | -1.6825399 | -1.5696644 | -1.6881942 | -0.0015873 |
| -2.7351599 | -4.5772257 | -0.2099504 | -1.764935 | -2.9290442 | -2.4432631 | -0.0010242 |
| -3.7430983 | -5.2315006 | 0.43624187 | -4.3253078 | -3.6936395 | -3.3114609 | -0.0010242 |
| -1.5165672 | -1.1567426 | -1.395516 | -1.2131445 | -1.1599206 | -1.2883782 | -0.0172031 |
| -8.338397 | -10.757898 | -8.1716175 | -8.7595482 | -9.5509176 | -9.1156757 | -0.0036032 |
| -12.807262 | -15.621999 | -11.433842 | -10.343738 | -13.060109 | -12.65339 | -0.0036032 |
| -1.3671184 | -1.9289563 | -0.5390677 | -1.0254707 | -0.9023461 | -1.1525918 | -0.0004622 |
| 0.10336637 | -0.0482159 | 0.01784325 | -0.5052357 | -0.176899 | -0.1218282 | -0.0152691 |
| -0.6168451 | -0.1796613 | -1.3722651 | -2.2688756 | -2.3988218 | -1.3672938 | -0.002773 |
| -5.8964338 | -2.7088282 | -3.1894093 | -6.2931614 | -4.3797112 | -4.4935088 | -0.0011151 |
| -0.0942411 | -1.0136371 | -0.7455659 | -0.2440243 | 0.28285193 | -0.3629233 | -0.0004678 |
| -1.7746503 | -1.3594842 | -1.1275527 | -0.8266733 | -1.1899459 | -1.2556613 | -0.0009984 |
| -2.9554143 | -2.6812835 | -2.7354653 | -2.4389782 | -2.9269919 | -2.7476266 | -0.0009984 |
| -1.8699265 | -0.3806951 | -1.2767892 | 0.84328973 | -4.2168665 | -1.3801975 | -0.001748 |
| -5.4897194 | -8.9620638 | -3.5875359 | -6.2101507 | -7.8609524 | -6.4220844 | -0.003031 |
| -4.8817792 | -3.0488353 | -2.7879424 | -4.6721859 | -4.2229819 | -3.9227449 | -0.003031 |
| -3.3216095 | -5.5221014 | -2.4301367 | -2.066071 | -1.914022 | -3.0507881 | -0.003031 |
| 1.53648925 | 0.42920345 | -0.3735351 | 0.30216265 | 1.52076292 | 0.68301663 | -0.003031 |
| -0.1912348 | -4.0127096 | -6.3218775 | -0.4364939 | -1.742888 | -2.5410408 | -0.003031 |
| -2.3245368 | -5.7172966 | -4.5579243 | -4.2571435 | -4.0173078 | -4.1748418 | -0.003031 |
| 2.64757109 | -4.6007185 | -1.616394 | 2.21115589 | -2.2371542 | -0.719108 | -0.003031 |
| -4.9445829 | -6.3797083 | -4.724112 | -7.1515369 | -4.4020329 | -5.5203946 | -0.003031 |
| -4.0991244 | -1.3777673 | -4.0591149 | -3.1715791 | -6.1280661 | -3.7671304 | -0.003031 |
| -1.1689711 | -2.8479934 | -1.3502402 | -3.1879282 | -2.7777014 | -2.2665668 | -0.003031 |
| 2.78755188 | 2.40424299 | 2.6887691 | 1.76161575 | 1.1210494 | 2.15264583 | -0.003031 |
| 2.58011627 | 4.01674652 | 2.231812 | 1.68916941 | 2.36970949 | 2.57751074 | -0.003031 |

| | | | | | | |
|---|---|---|---|---|---|---|
| -5.8805637 | -4.4100037 | -4.1072502 | -5.103344 | -4.4185076 | -4.7839338 | -0.003031 |
| -2.7770076 | -2.36975 | -1.6000762 | -2.2289691 | -4.3755522 | -2.670271 | -0.003031 |
| 1.1793685 | -0.0056856 | 0.6714921 | 1.60906196 | 2.84821486 | 1.26049037 | -0.003031 |
| -2.8633311 | -1.8577204 | -1.9430366 | -2.7653503 | -2.6277471 | -2.4114371 | -0.003031 |
| -6.5617285 | -6.4394236 | -6.82584 | -6.1754565 | -6.8791895 | -6.5763276 | -0.003031 |
| -4.8812747 | -5.3396544 | -5.3844233 | -4.024684 | -5.6989007 | -5.0657874 | -0.003031 |
| -1.883705 | -4.3000069 | -3.0210853 | -4.6194868 | -3.4600067 | -3.4568581 | -0.003031 |
| -4.4896002 | -3.7373271 | -6.3050809 | -6.3446107 | -4.4323673 | -5.0617972 | -0.003031 |
| -3.2016208 | -1.8428771 | -1.2154918 | -4.342885 | -2.9946876 | -2.7195125 | -0.003031 |
| -4.6151333 | -6.1610227 | -6.551115 | -4.5247126 | -6.7787218 | -5.7261411 | -0.003031 |
| -4.5800686 | -2.6371539 | -7.0042515 | -6.3683248 | -3.4258485 | -4.8031294 | -0.003031 |
| 1.91373038 | 1.83930016 | 4.44498253 | -1.4318931 | 3.70454597 | 2.09413319 | -0.003031 |
| -1.0517378 | -4.7084703 | -3.0590136 | -4.826035 | -3.245575 | -3.3781663 | -0.003031 |
| -5.7968326 | -4.4202957 | -3.3781147 | -3.9653633 | -4.0651083 | -4.3251429 | -0.003031 |
| -2.1424489 | -2.0374575 | -1.4234562 | -0.9473026 | -2.5250981 | -1.8151526 | -0.003031 |
| -6.5688515 | -7.5978298 | -8.0781298 | -6.3179283 | -5.2481942 | -6.7621867 | -0.003031 |
| -4.1834784 | -5.5491967 | -4.9371591 | -4.8437386 | -5.2777748 | -4.9582695 | -0.003031 |
| 3.73837852 | -5.0418634 | -6.4519091 | -0.8079133 | -4.151412 | -2.5429439 | -0.003031 |
| -11.221227 | -11.432168 | -11.493489 | -10.833626 | -11.058847 | -11.207871 | -0.003031 |
| -7.4974794 | -8.2903519 | -8.9163961 | -9.4381618 | -8.0429688 | -8.4370716 | -0.003031 |
| 0.51206064 | -0.8869205 | 0.11866498 | -0.5381432 | -2.554945 | -0.6698566 | -0.003031 |
| -6.8223152 | -4.2504859 | -6.3662739 | -7.5850477 | -5.4061623 | -6.086057 | -0.003031 |
| -3.8586886 | -3.0195608 | -3.9102471 | -2.4961514 | -1.5501564 | -2.9669609 | -0.003031 |
| -6.7188144 | -6.504652 | -6.5300255 | -4.7613163 | -6.156177 | -6.134197 | -0.003031 |
| -1.2072635 | -1.4880805 | -1.4380703 | -0.1047206 | 0.06229305 | -0.8351684 | -0.0032998 |
| -5.6816034 | -5.6754303 | -4.1078682 | -6.0016465 | -4.1379766 | -5.120905 | -0.0035416 |
| -0.4537079 | -0.603693 | -0.9314594 | -0.3579624 | 0.02080464 | -0.4652036 | -0.0003354 |
| 0.14178753 | 0.5009203 | 0.11214089 | 0.23611116 | 0.13259768 | 0.22471151 | -0.0003354 |
| 3.81197596 | 3.70851994 | 2.1292367 | -0.0624671 | 1.88197064 | 2.29384723 | -0.0003354 |
| -0.7430286 | -0.5563998 | -1.4659619 | -0.6239433 | -0.201582 | -0.7181831 | -0.0003354 |
| -1.4671597 | -1.5462902 | -1.4709229 | -2.8445742 | -1.7854595 | -1.8228813 | -0.003566 |
| -3.1399646 | -4.9010792 | -0.6835518 | 2.19447899 | 0.57143664 | -1.191736 | -0.0009121 |
| 0.16445589 | 0.06367874 | -1.4940438 | -0.9673128 | -0.6737351 | -0.5813914 | -0.0010377 |
| 0.69921255 | 1.75049889 | 1.41391754 | 1.348701 | 0.69836926 | 1.18213985 | -0.0010377 |
| -0.484303 | -0.4818738 | -1.1212721 | -0.8785379 | -0.7347989 | -0.7401571 | -0.0010377 |
| 0.48302662 | -0.007592 | -0.4622215 | 0.0664463 | -0.0708748 | 0.00175693 | -0.0010377 |
| 2.14898181 | 2.12744856 | 2.43448782 | 2.16849637 | 1.42879534 | 2.06164198 | -0.0010377 |
| -0.202512 | -0.1566546 | -0.3036777 | 0.04538822 | -0.0886846 | -0.1412281 | -0.0010377 |
| -1.7854882 | -1.9487057 | -1.4780515 | -1.9430733 | -0.8768883 | -1.6064414 | -0.0010377 |
| 1.81181073 | 2.60885048 | 0.71135092 | 1.6197933 | 1.6181097 | 1.67398303 | -0.0010377 |
| -0.114938 | -1.2719609 | -1.0044646 | -1.0251771 | -2.0943089 | -1.1021699 | -0.0010377 |
| -2.9744875 | -3.599262 | -3.3436537 | -2.9525819 | -2.3149772 | -3.0369925 | -0.0035056 |
| 0.2073667 | 0.21163774 | 0.51712203 | 0.35748005 | 0.36226749 | 0.3311748 | -0.0013073 |
| -6.5961766 | -8.8952789 | -7.4824295 | -8.4462194 | -6.8625393 | -7.6565288 | -0.0010403 |
| -5.9852047 | -8.3001242 | -7.8417645 | -5.8544345 | -6.3814282 | -6.8725912 | -0.0025491 |

| | | | | | | |
|---|---|---|---|---|---|---|
| 0.9522059 | 1.49893773 | 0.54001236 | -1.414782 | -0.2779303 | 0.25968874 | -0.0001732 |
| -1.3769658 | -1.32516 | -1.118732 | -0.5081801 | -0.5508802 | -0.9759836 | -0.0070983 |
| -2.6620212 | -2.1508942 | -1.7698641 | -2.6063008 | -2.2485805 | -2.2875321 | -0.0070983 |
| 1.83332348 | -3.0381877 | -1.0487958 | 1.50795984 | -1.4163537 | -0.4324108 | -0.0070983 |
| 0.40709114 | -1.6896924 | -0.0879054 | -0.6551578 | -0.5813518 | -0.5214032 | -0.0070983 |
| -1.0422964 | -1.0687315 | -0.6992197 | -1.0940385 | -0.9154923 | -0.9639557 | -0.0116321 |
| -0.1400659 | -1.2070037 | -0.2791779 | -0.4682846 | -0.8268218 | -0.5842708 | -0.0005582 |
| 0.96353245 | 1.690871 | 1.42631173 | 1.28646076 | 1.17887652 | 1.30921049 | -0.0079792 |
| -1.9091974 | -3.8855152 | -1.6772251 | -1.6020032 | -2.371191 | -2.2890264 | -0.0027725 |
| -6.0102482 | -6.6641865 | -5.211082 | -5.5919008 | -5.524847 | -5.8004529 | -0.0009218 |
| 0.29965091 | 0.06507778 | 0.13633251 | 0.34393215 | 0.28278828 | 0.22555633 | -0.0004488 |
| -0.3291407 | -4.2072248 | 0.19669271 | 1.23197627 | -2.1683428 | -1.0552079 | -0.0047225 |
| 1.27292776 | 0.65097761 | -0.0980263 | 0.3246057 | -1.0516975 | 0.21975746 | -0.0031494 |
| 0.74108958 | 0.10590672 | -0.2313195 | 1.39913964 | -0.075449 | 0.38787348 | -0.0022968 |
| -2.2525868 | -5.5652623 | -0.2744272 | -5.3536639 | -1.929451 | -3.0750782 | -0.0022968 |
| -0.6930439 | -5.1033688 | -0.5436673 | -1.6496406 | 0.02392173 | -1.5931598 | -0.003661 |
| 0.48320746 | 0.20113897 | 0.03602338 | -0.2330251 | 0.10198402 | 0.11786575 | -0.0007669 |
| -0.7883825 | -1.3617287 | -0.6636233 | -1.2580769 | -0.730036 | -0.9603695 | -0.0007669 |
| 0.12711763 | -0.1349373 | -0.8169034 | -0.0413011 | -0.3825536 | -0.2497155 | -0.0007669 |
| -1.1048567 | -0.6344495 | -1.08641 | -0.9553893 | -0.8954656 | -0.9353142 | -0.0007669 |
| -0.7742577 | -0.8865585 | -0.6165943 | -0.7556245 | -0.8638256 | -0.7793721 | -0.0007669 |
| -4.5316148 | -4.5648232 | -4.3922997 | -6.2264471 | -5.4895682 | -5.0409506 | -0.0009626 |
| -2.280673 | -4.1767015 | -3.7733538 | -4.5566626 | -5.5995703 | -4.0773922 | -0.0050599 |
| -5.2409501 | -5.2016244 | -4.5647435 | -7.2059517 | -5.371388 | -5.5169315 | -0.0050599 |
| -0.320399 | 0.06982803 | 0.58718419 | -0.0617785 | 0.0468421 | 0.06433535 | -0.0008606 |
| -1.2995586 | -1.627827 | -0.8313322 | -1.0271745 | -0.9854603 | -1.1542705 | -0.0004586 |
| -0.603092 | -1.0948088 | -0.3804536 | -0.3631873 | -1.3984563 | -0.7679996 | -0.0004586 |
| 0.70919371 | 0.87121153 | 1.13152838 | 1.19422519 | 1.09348214 | 0.99992819 | -0.0004586 |
| -0.6282554 | -2.6188314 | -0.1860745 | -0.138432 | -1.3382342 | -0.9819655 | -0.0037957 |
| -1.0676324 | -0.6893535 | -0.7782207 | -0.4589953 | -0.3417187 | -0.6671841 | -0.0040029 |
| -2.2022357 | -2.5260062 | -0.7377996 | -1.3460195 | -4.3909898 | -2.2406102 | -0.0018337 |
| -1.2576797 | -1.8598671 | -1.4185185 | -1.1744268 | -1.0016239 | -1.3424232 | -0.0018337 |
| -0.8044679 | -0.9947674 | -1.3300381 | 0.10195374 | -1.33122 | -0.8717079 | -0.0051373 |
| 0.4518199 | 0.27946782 | 0.02463508 | -0.1735971 | -0.3792266 | 0.04061983 | -0.0008378 |
| -3.4386597 | -1.0013142 | -1.4123034 | -0.461978 | -3.9079695 | -2.0444449 | -0.0022045 |
| -3.3348556 | -2.5387831 | -0.1419916 | -3.2545538 | -3.683692 | -2.5907752 | -0.0024651 |
| 0.44211912 | 0.3270247 | 0.36502409 | 0.44850659 | 0.29999113 | 0.37653313 | -0.0024651 |
| -0.3571913 | -0.308497 | -0.1638064 | 0.13892722 | -0.2724719 | -0.1926079 | -0.0153109 |
| -1.1182263 | -1.2527426 | -1.1196648 | -2.1806593 | -2.3332083 | -1.6009003 | -0.0113538 |
| -1.3054571 | -1.3721099 | -1.5838795 | -0.2782168 | -1.6116567 | -1.230264 | -0.0091425 |
| -3.4284222 | -3.2600408 | -3.8107822 | -3.6174276 | -2.9355857 | -3.4104517 | -0.0020566 |
| -2.3121228 | -2.8814406 | -2.503469 | -3.2429199 | -2.5643449 | -2.7008595 | -0.0138288 |
| -0.7730627 | -0.4341216 | -0.82268 | -0.8218982 | -0.877125 | -0.7457775 | -0.0138288 |
| -1.1795678 | -0.6637055 | -0.8889503 | -1.1111929 | -1.8228796 | -1.1332592 | -0.0138288 |
| 1.27015448 | 1.3752265 | 1.24207902 | 0.79021001 | 0.94604468 | 1.12474294 | -0.0019866 |

| | | | | | | |
|---|---|---|---|---|---|---|
| -7.2674932 | -7.9579835 | -7.3543682 | -7.6959748 | -8.1921425 | -7.6935925 | -0.0054091 |
| 1.4387219 | 0.97939157 | 1.13777149 | 1.44431007 | 1.04245186 | 1.20852938 | -0.0054091 |
| -0.9083316 | -0.4713591 | -0.4803543 | -0.3720846 | -0.6604391 | -0.5785138 | -0.0054091 |
| 0.41079807 | 0.61921215 | -0.5089903 | 0.68667054 | -1.4669094 | -0.0518438 | -0.002918 |
| -5.724227 | -6.1736035 | -6.2754831 | -5.6366982 | -5.5707283 | -5.876148 | -0.006654 |
| 0.58043838 | 0.93263578 | 0.59624362 | 1.00621033 | 0.62569714 | 0.74824505 | -0.006654 |
| -1.8522184 | -1.4194112 | -2.0836039 | -1.1959956 | -1.1045361 | -1.531153 | -0.006654 |
| 0.95290661 | 0.6649332 | 0.93037844 | 0.84224582 | 1.35715592 | 0.949524 | -0.006654 |
| 0.60468554 | 0.66297555 | 0.8532958 | 0.03921103 | 1.15999675 | 0.66403294 | -0.006654 |
| 1.67888355 | 2.72039032 | 1.6480689 | 1.63291621 | 2.17233276 | 1.97051835 | -0.006654 |
| 0.92144513 | 0.62315464 | 0.55063105 | 0.14003134 | 0.41671038 | 0.53039451 | -0.006654 |
| 1.68391347 | 1.03801572 | 1.46328533 | 0.8678751 | 0.87567139 | 1.1857522 | -0.006654 |
| -2.4348273 | -1.2860513 | -1.8920752 | -1.429801 | -2.449379 | -1.8984267 | -0.006654 |
| 0.7139864 | 0.30033112 | 0.61591196 | 0.86883736 | 0.58841825 | 0.61749701 | -0.006654 |
| 2.56877518 | 2.68782687 | 1.86655807 | 1.91213417 | 2.81432796 | 2.36992445 | -0.006654 |
| -0.7424588 | -0.1691673 | -0.7820382 | -1.3344078 | -1.6491435 | -0.9354431 | -0.006654 |
| 1.81080794 | 1.43328428 | 1.31573331 | 1.80506766 | 0.33740234 | 1.34045911 | -0.006654 |
| -2.054924 | -2.3930516 | -2.4733486 | -3.2517798 | -2.7544732 | -2.5855155 | -0.006654 |
| -1.2547648 | -2.3481736 | -0.0273545 | -0.2179222 | -2.8404555 | -1.3377341 | -0.006654 |
| -0.8746645 | -0.5663052 | -0.8973632 | -3.2645011 | -0.8361583 | -1.2877985 | -0.006654 |
| 2.43212175 | 2.62706995 | 1.95734477 | 1.85034347 | 1.35109615 | 2.04359522 | -0.0006666 |
| -1.1810813 | -0.8556308 | -1.7535496 | -0.2015195 | -0.7021202 | -0.9387803 | -0.0006666 |
| 0.71121335 | 0.51668596 | 0.75824738 | 0.60895443 | 0.54324794 | 0.62766981 | -0.0004901 |
| -6.4342132 | -5.5174451 | -3.071702 | -3.5728798 | -2.0274067 | -4.1247293 | -0.0014429 |
| -4.6205707 | -6.3457494 | -5.3181114 | -3.3829646 | -3.3676734 | -4.6070139 | -0.0014429 |
| -0.4480791 | -0.7127063 | -0.5573554 | -0.6900117 | -1.1426766 | -0.7101658 | -0.0014429 |
| -4.5801992 | -2.9060926 | -5.3301511 | -1.5970416 | -5.3085904 | -3.944415 | -0.0008745 |
| 0.4603467 | 0.15422058 | 0.19252396 | 0.29592967 | 0.41203976 | 0.30301213 | -0.0049209 |
| -0.5283582 | -0.6971719 | -0.7362833 | -0.2166405 | -0.6519442 | -0.5660796 | -0.0064729 |
| -2.7143948 | -5.3593998 | -3.1691761 | -2.3819175 | -4.6350632 | -3.6519903 | -0.0044687 |
| 0.92243671 | 1.16111708 | 0.69281459 | 1.4001441 | 0.35393977 | 0.90609045 | -0.0012714 |
| -0.5068882 | -0.1538048 | -0.0836635 | -0.1249249 | -0.2058196 | -0.2150202 | -0.0018213 |
| -3.4480238 | -4.6781912 | -2.8909712 | -0.6814996 | -2.3047352 | -2.8006842 | -0.0018213 |
| -5.2568803 | -8.0265789 | -9.6190968 | -7.3415766 | -7.6969404 | -7.5882146 | -0.0014707 |
| -1.0546873 | -1.2183113 | -0.9840059 | -0.7065618 | -1.4059675 | -1.0739068 | -0.0040797 |
| -3.0985851 | -5.0672398 | -3.9894528 | -3.5567775 | -4.0235481 | -3.9471207 | -0.0053911 |
| 0.50754213 | 1.49558902 | 2.74276757 | 1.37189484 | 1.63173819 | 1.54990635 | -0.0053911 |
| -1.0533073 | -1.0005466 | -1.5707158 | -1.6545229 | -0.9933227 | -1.2544831 | -0.0053911 |
| -0.0708191 | 0.83833218 | 0.82110727 | 0.76579952 | 0.63069105 | 0.59702218 | -0.0053911 |
| -0.275614 | -0.2256064 | -0.4832087 | 0.13100195 | -0.2112558 | -0.2129366 | -0.0053911 |
| -3.2312522 | -2.8243055 | -3.6565154 | -3.2967677 | -3.1226907 | -3.2263063 | -0.0016515 |
| -0.3142869 | -3.679291 | -2.3851633 | -3.4701145 | -4.8357954 | -2.9369302 | -0.0016515 |
| -2.0462375 | 2.55060983 | 0.88262558 | 3.69845438 | 0.37358427 | 1.09180732 | -0.0017179 |
| -0.3261092 | -1.6743187 | -1.7087593 | -4.3190956 | 1.31606865 | -1.3424428 | -0.0017179 |
| -1.3696475 | -3.4189401 | -1.592021 | -2.7747626 | -1.0754442 | -2.0461631 | -0.0017179 |

| | | | | | | |
|---|---|---|---|---|---|---|
| 0.65460753 | 1.16232061 | -2.6995296 | -2.653501 | -1.5326785 | -1.0137562 | -0.0017179 |
| -1.2787259 | -2.3120041 | -0.6563871 | 0.17628336 | -1.6077678 | -1.1357203 | -0.0017179 |
| 0.28224611 | -2.7202551 | -0.7705116 | -5.0722785 | -5.0267539 | -2.6615106 | -0.0017179 |
| -2.4978046 | -3.2840965 | -1.6192837 | -3.588012 | -2.6103344 | -2.7199062 | -0.0017179 |
| -0.0009027 | -0.2219844 | -0.6029096 | -0.6683016 | -1.1170738 | -0.5222344 | -0.0017179 |
| -2.289191 | -3.8804111 | -3.3479347 | -1.7822604 | -2.5178947 | -2.7635384 | -0.0017179 |
| -1.7586186 | -3.2937207 | -2.2541704 | -1.1516964 | -2.5172226 | -2.1950858 | -0.0013859 |
| -3.7160256 | -6.7732368 | -6.6533394 | -8.6308699 | -6.448844 | -6.4444631 | -0.0013859 |
| -3.2239888 | -3.6628668 | -2.9283218 | -2.7842941 | -3.4724064 | -3.2143756 | -0.0019536 |
| -0.9616742 | -0.748723 | 0.47701359 | -0.541945 | -1.0904551 | -0.5731567 | -0.0019536 |
| -1.6798489 | -2.5665674 | -0.741451 | -2.8388801 | -1.053471 | -1.7760437 | -0.0031746 |
| -0.2055144 | 0.01229954 | -0.4239469 | 0.07714272 | -0.2991745 | -0.1678387 | -0.0036912 |
| -1.9275757 | -2.7163181 | -2.0650978 | -2.6853848 | -2.834975 | -2.4458703 | -0.0195448 |
| -8.4710541 | -10.40258 | -6.5668149 | -7.6407666 | -7.8504257 | -8.1863283 | -0.0031427 |
| -7.2047873 | -4.9656324 | -5.4141254 | -6.7774158 | -5.453124 | -5.963017 | -0.0031427 |
| -0.566462 | -1.8363489 | -0.4237421 | -0.9542956 | -2.2753758 | -1.2112449 | -0.0037343 |
| -3.0307245 | -4.0279717 | -1.8831017 | -1.7473464 | -3.1823332 | -2.7742955 | -0.0047532 |
| -0.5150993 | -1.5677679 | -0.324297 | -0.4607263 | -2.6988969 | -1.1133574 | -0.0038228 |
| 1.09215724 | -1.1240153 | 2.75892305 | 0.06515312 | 1.01754069 | 0.76195176 | -0.0007085 |
| 0.54220629 | 0.51837444 | 0.03988266 | 0.46711969 | -1.1953473 | 0.07444715 | -0.0052794 |
| -2.1507101 | -2.3485847 | -2.3563495 | -2.2347827 | -2.5672817 | -2.3315417 | -0.0052794 |
| 0.11039019 | 0.33972406 | 0.84774137 | 0.92962623 | 0.37222815 | 0.519942 | -0.0034036 |
| 0.18277383 | 0.11882257 | 0.37099147 | 0.33881879 | -0.0773709 | 0.18680716 | -0.0034036 |
| 1.77110815 | 1.05621362 | 1.14762282 | 1.15758777 | 1.67386079 | 1.36127863 | -0.0034036 |
| -0.9578898 | -0.7869391 | -0.6081395 | -0.065706 | -0.9060705 | -0.664949 | -0.0034036 |
| -3.4459467 | -2.6387062 | -3.8620467 | -0.7151117 | -2.1185088 | -2.556064 | -0.0028385 |
| -0.343637 | -0.1886218 | -0.6674082 | 0.00432897 | -0.2155092 | -0.2821694 | -0.0049963 |
| 0.5641439 | 0.01582265 | 0.63554215 | 0.67276812 | 0.36811829 | 0.45127902 | -0.0011091 |
| -0.7639661 | -0.7006293 | -0.6872492 | -1.3822293 | -1.7761798 | -1.0620507 | -0.0007486 |
| -0.3559828 | -0.7711592 | -0.809427 | -1.2608099 | -1.3209893 | -0.9036736 | -0.0007486 |
| -1.5643584 | -1.0299348 | -0.7189357 | -0.7653501 | -0.391726 | -0.894061 | -0.0038399 |
| -0.0208144 | -0.9433846 | 0.21377802 | -0.5215781 | -0.7730556 | -0.4090109 | -0.0038399 |
| -1.2621195 | -2.1422691 | -1.1870613 | -2.3486018 | -3.3048427 | -2.0489789 | -0.0038399 |
| -2.4496202 | -4.598299 | -0.3704417 | -2.5081129 | -4.8653426 | -2.9583633 | -0.0038399 |
| -1.4197469 | -1.2699976 | -1.5259435 | -1.4959817 | -2.0512834 | -1.5525906 | -0.0042622 |
| -0.060061 | -0.0924113 | -0.322087 | -0.0832593 | -0.3657403 | -0.1847118 | -0.0006312 |
| -0.9160514 | -1.8637328 | 0.25821972 | -0.5164886 | -0.0977716 | -0.6271649 | -0.0025543 |
| 0.17267931 | 0.84023213 | 0.78968167 | 0.61553693 | 0.46429443 | 0.57648489 | -0.0050398 |
| -2.0151777 | -2.0118716 | -2.2209585 | -2.1378193 | -2.2236371 | -2.1218928 | -0.0050398 |
| -3.3174901 | -3.6875267 | -3.7080355 | -3.4148145 | -3.5167003 | -3.5289134 | -0.0050398 |
| -3.8710048 | -4.2862844 | -3.9673114 | -4.5588794 | -3.5622969 | -4.0491554 | -0.0050398 |
| 0.99447191 | 1.65436161 | 1.62858248 | 2.20472527 | 1.53480351 | 1.60338895 | -0.0050398 |
| -0.7178521 | -1.0299088 | -0.4704767 | -0.0976965 | -1.2177718 | -0.7067412 | -0.0050398 |
| -0.6179061 | -0.5058796 | -0.6431627 | -0.6085937 | -0.8633142 | -0.6477713 | -0.0003983 |
| -2.137887 | -1.8703637 | -1.4257349 | -2.4552088 | -2.1076965 | -1.9993782 | -0.0009154 |

| | | | | | | |
|---|---|---|---|---|---|---|
| -2.0758979 | -1.40452 | 0.10627651 | -1.1019616 | -1.4159694 | -1.1784145 | -0.0009154 |
| -1.1798303 | -1.6373038 | -1.7730572 | -1.3425043 | -1.3174872 | -1.4500366 | -0.0009021 |
| -3.7685077 | -3.4326389 | -3.178983 | -4.3012929 | -8.2089043 | -4.5780653 | -0.0020114 |
| 0.53983164 | 0.62038279 | 0.04951072 | 0.83072281 | 0.40492201 | 0.48907399 | -0.0023634 |
| -3.5906446 | -2.8726888 | -2.8504288 | -3.7578599 | -1.7197194 | -2.9582683 | -0.0023634 |
| -6.5594091 | -8.3971691 | -7.3509007 | -7.0409942 | -6.3426647 | -7.1382276 | -0.0023634 |
| -3.1626952 | -6.0791626 | -3.0937102 | -3.3932509 | -6.3083773 | -4.4074392 | -0.0023634 |
| -6.1606069 | -9.5939465 | -5.8606491 | -6.5804758 | -9.4373856 | -7.5266128 | -0.0023634 |
| -4.4481382 | -4.7671747 | -4.6373463 | -5.2335963 | -3.7517991 | -4.5676109 | -0.0023634 |
| 0.38187671 | 0.37435079 | 0.39838767 | 0.50933814 | 0.43047547 | 0.41888576 | -0.0008894 |
| 0.31829906 | 0.67202902 | -0.0637558 | 0.05722022 | 0.03999591 | 0.20475769 | -0.0024244 |
| -1.6263227 | -2.0758657 | -1.9758909 | -1.5117682 | -2.0122793 | -1.8404254 | -0.0078558 |
| -8.3136444 | -7.5963783 | -6.0038567 | -5.9169312 | -10.003619 | -7.5668859 | -0.001822 |
| 0.92909765 | 1.39145041 | 1.04602408 | 1.36948848 | 1.10274386 | 1.1677609 | -0.0043621 |
| 1.338238 | 1.0706085 | 1.37346268 | 1.87550974 | 1.55239666 | 1.44204311 | -0.0043621 |
| -2.1517868 | -2.2252812 | -3.5326309 | -3.5106363 | -3.1107965 | -2.9062263 | -0.0043621 |
| 1.17471695 | 0.94386375 | 0.92036295 | 0.60281372 | 0.54660892 | 0.83767326 | -0.0043621 |
| 0.05206275 | 0.92638898 | 0.24918032 | 0.3174932 | 0.07947922 | 0.32492089 | -0.0043621 |
| 0.28266525 | 0.18706322 | 0.41325378 | 0.21954536 | 0.20512843 | 0.26153121 | -0.0043621 |
| -1.6677926 | -2.6935239 | -1.7877235 | -3.1510525 | -2.1782489 | -2.2956683 | -0.0043621 |
| -1.4255196 | -1.9447548 | -3.3573623 | -1.7485758 | -0.7225724 | -1.839757 | -0.0043621 |
| -7.5253496 | -7.2256646 | -7.1122918 | -7.2089138 | -9.0867968 | -7.6318033 | -0.0043621 |
| -6.6401291 | -5.3922906 | -5.4889116 | -4.7122612 | -6.4548264 | -5.7376838 | -0.0043621 |
| 1.93123484 | 2.09529161 | 1.5697968 | 2.66329646 | 2.19130564 | 2.09018507 | -0.0043621 |
| -5.2847919 | -6.5767479 | -4.5921555 | -7.4077358 | -5.8732486 | -5.9469359 | -0.0043621 |
| -2.5843005 | -1.3036041 | -1.188343 | -1.8507608 | -2.0755301 | -1.8005077 | -0.0043621 |
| -1.7960172 | -1.0184202 | -2.4456306 | -2.7818654 | -1.8489614 | -1.9781789 | -0.0043621 |
| -3.8027992 | -2.31039 | -4.1013083 | -3.912466 | -3.9347177 | -3.6123363 | -0.0043621 |
| -2.6893229 | -2.639327 | -0.7812464 | -2.0649238 | -3.5302203 | -2.3410081 | -0.0043621 |
| -1.4909178 | -2.3128643 | -1.943572 | -2.1634755 | -2.5723925 | -2.0966444 | -0.0043621 |
| 1.54392838 | 0.72578311 | 0.96884036 | -0.4623923 | 0.92010307 | 0.73925252 | -0.0043621 |
| -5.9483776 | -5.398695 | -6.1476917 | -4.5009422 | -5.8213468 | -5.5634107 | -0.0043621 |
| -1.1870532 | -0.9265654 | 0.03630662 | 0.35569143 | -0.8041275 | -0.5051496 | -0.0043621 |
| 0.37896585 | -0.121644 | 0.22487617 | 0.27338743 | -0.5775323 | 0.03561063 | -0.0043621 |
| 0.51298296 | 0.32703638 | 0.25222158 | 1.07841682 | 1.28689802 | 0.69151115 | -0.0043621 |
| -1.3263116 | -2.7285318 | -0.9679713 | -2.4528813 | -2.129415 | -1.9210222 | -0.0043621 |
| -2.1150739 | -3.0677085 | -1.6484456 | -2.7788858 | -2.2700043 | -2.3760236 | -0.0043621 |
| 1.2309804 | 1.30129695 | 1.06136107 | 0.60960317 | 1.43356609 | 1.12736154 | -0.0043621 |
| -0.3888063 | -0.8933198 | -0.1984782 | -0.3806076 | -0.7374873 | -0.5197399 | -0.0043621 |
| -3.8809271 | -2.5218463 | -3.0343199 | -4.6327724 | -2.5676699 | -3.3275071 | -0.0043621 |
| -5.7720175 | -4.5671101 | -5.2126441 | -5.4319949 | -4.384697 | -5.0736927 | -0.0043621 |
| -5.9958267 | -5.6160798 | -3.8352759 | -6.1479173 | -5.8541265 | -5.4898452 | -0.0043621 |
| -1.0881305 | -2.0131598 | -2.4797683 | -3.0797291 | -1.8260734 | -2.0973722 | -0.0043621 |
| -3.6376295 | -1.3874438 | -0.9081967 | -3.5159469 | -2.1198249 | -2.3138083 | -0.0043621 |
| -1.341481 | -0.5905784 | -1.538435 | -0.5227163 | -0.5205793 | -0.902758 | -0.0043621 |

| | | | | | | |
|---|---|---|---|---|---|---|
| -1.8847311 | -0.7273319 | -0.4518242 | -0.1664975 | -1.5262077 | -0.9513185 | -0.0043621 |
| -1.4700089 | -1.8903319 | -0.7966585 | -0.9700924 | -0.4142747 | -1.1082733 | -0.0043621 |
| 1.51740456 | 0.07812715 | -0.5277814 | -0.3397613 | 1.45407629 | 0.43641307 | -0.0043621 |
| -3.1204922 | -1.4146752 | -1.3853235 | -0.8761162 | -2.5955596 | -1.8784333 | -0.0043621 |
| -2.8331492 | -2.6998332 | -1.9071389 | -2.0766065 | -3.1787975 | -2.5391051 | -0.0043621 |
| -0.5983548 | -1.503288 | -1.4034989 | 0.41757202 | -0.8985634 | -0.7972266 | -0.0043621 |
| -1.6530819 | -2.6414804 | -0.6576862 | -0.8220124 | -0.6166119 | -1.2781746 | -0.0043621 |
| 0.24125791 | -0.0217235 | -0.2840919 | 0.79419804 | -0.257936 | 0.0943409 | -0.0043621 |
| -2.4908757 | -1.027844 | -3.7016869 | -1.8246533 | -2.5368505 | -2.3163821 | -0.0043621 |
| -1.1281023 | -0.4863116 | -0.1016865 | -0.2495713 | -0.3459406 | -0.4623225 | -0.0043621 |
| -0.951304 | -2.155282 | -1.3994689 | -1.1664584 | -2.9030442 | -1.7151115 | -0.0043621 |
| -3.6800797 | -1.4258121 | -2.2401085 | -1.4360335 | -1.6680022 | -2.0900072 | -0.0043621 |
| -3.4912133 | -2.5494964 | -2.3166738 | -2.060245 | -1.9253292 | -2.4685915 | -0.0043621 |
| 2.25286365 | 1.619187 | 1.69394875 | 2.55853319 | 1.6672101 | 1.95834854 | -0.0043621 |
| -1.5449286 | -1.714047 | -2.3678818 | -2.0041142 | -2.3325911 | -1.9927125 | -0.0043621 |
| 0.03395605 | 0.35755962 | -0.0712528 | 0.35481542 | 1.4482224 | 0.42466013 | -0.0043621 |
| -1.9975911 | -1.1072519 | -1.5424179 | -2.0853753 | -1.7308559 | -1.6926984 | -0.0043621 |
| 0.02383256 | -1.45191 | -1.4633756 | -0.2792308 | -0.641748 | -0.7624864 | -0.0043621 |
| -1.3207119 | -2.5134816 | 0.69165492 | -0.0439675 | -0.6290064 | -0.7631025 | -0.0043621 |
| 1.92522085 | 1.23813224 | 1.52160001 | 1.85026562 | 1.96828544 | 1.70070083 | -0.0043621 |
| 0.95051718 | 1.2858212 | 0.8421247 | 0.42889726 | -0.6769704 | 0.566078 | -0.0043621 |
| 2.72556949 | 2.75354385 | 3.19914365 | 2.74386239 | 1.82357228 | 2.64913833 | -0.0043621 |
| -0.6818383 | -0.5608582 | -1.6635557 | -0.2478642 | -0.3437681 | -0.6995769 | -0.0043621 |
| -1.2050734 | -2.1972733 | -1.9229081 | -1.6074506 | -0.4518876 | -1.4769186 | -0.0043621 |
| -1.1869786 | -2.5515742 | -2.7335434 | -2.5391407 | -3.7903609 | -2.5603196 | -0.0043621 |
| 0.39036298 | 1.42870235 | 1.42310381 | 2.25030804 | 1.23391819 | 1.34527907 | -0.0043621 |
| -0.6637013 | -0.7791183 | 0.0394423 | -0.4416547 | -0.7243111 | -0.5138686 | -0.0043621 |
| -2.1652987 | -1.8018348 | -2.2486377 | -2.4759309 | -1.3039722 | -1.9991349 | -0.0043621 |
| -2.1190932 | -0.6293221 | -0.1540365 | -0.797622 | -0.4886501 | -0.8377448 | -0.0043621 |
| -1.6011887 | -1.0478965 | 0.30895698 | -1.7973878 | -1.608431 | -1.1491894 | -0.0043621 |
| -1.5833195 | -2.7236061 | -2.3706508 | -2.7022002 | -3.2375393 | -2.5234632 | -0.0043621 |
| -1.515496 | -1.223331 | -3.1869226 | -2.3225238 | -2.1406794 | -2.0777905 | -0.0043621 |
| -0.7884636 | -0.1592212 | -0.2659931 | -0.1380603 | -0.8473824 | -0.4398241 | -0.0043621 |
| -3.1229904 | -2.9816215 | -2.7675035 | -2.5849147 | -2.5761328 | -2.8066326 | -0.0043621 |
| -2.7287221 | -3.736479 | -2.4486475 | -4.4184389 | -4.0548987 | -3.4774373 | -0.0043621 |
| -1.3012173 | 0.21762824 | 1.7362951 | -1.1166291 | -1.2945349 | -0.3516916 | -0.0043621 |
| -0.7594485 | 0.01263452 | -0.6986816 | -1.9145813 | -1.5331905 | -0.9786535 | -0.0043621 |
| 1.85456157 | 1.77729464 | 0.33625507 | 1.4931314 | 1.62270737 | 1.41679001 | -0.0043621 |
| -2.4497015 | -2.7662396 | -1.6348543 | -1.7649775 | -0.904108 | -1.9039762 | -0.0043621 |
| 0.21363783 | -1.4877466 | -1.1431037 | 0.35088396 | -1.3189377 | -0.6770532 | -0.0043621 |
| 1.02109909 | 0.43148375 | 1.63720536 | 1.29284811 | 0.93742085 | 1.06401143 | -0.0043621 |
| -3.2218566 | -2.9920125 | -1.9533625 | -2.6619051 | -4.0500636 | -2.97584 | -0.0043621 |
| -0.0201545 | 0.00460958 | 1.44228888 | -0.391315 | -0.4930413 | 0.10847754 | -0.0043621 |
| -3.0486822 | -2.2245436 | -3.5118604 | -2.5778961 | -2.4579358 | -2.7641836 | -0.0043621 |
| 1.09653759 | 1.32052374 | 1.65758085 | 2.16158438 | 1.12091351 | 1.47142801 | -0.0043621 |

| | | | | | | |
|---|---|---|---|---|---|---|
| -1.6731908 | -2.1860723 | -3.0373232 | -3.0994463 | -2.053669 | -2.4099403 | -0.0043621 |
| -0.5691226 | -0.6436789 | -0.2771475 | 0.4763298 | -0.1431735 | -0.2313585 | -0.0043621 |
| -3.2647932 | -3.3451338 | -2.5562587 | -3.5204945 | -3.4471793 | -3.2267719 | -0.0043621 |
| 0.05890059 | 0.37749839 | 0.80232793 | 0.16131687 | 0.15961719 | 0.31193219 | -0.0043621 |
| -3.1398201 | -2.6973233 | -1.6121626 | -1.4341273 | -2.640444 | -2.3047755 | -0.0043621 |
| 0.0742023 | 0.99618077 | -0.0569158 | -0.0912619 | 0.52346706 | 0.2891345 | -0.0043621 |
| -1.3642302 | -1.4033852 | -0.889401 | -1.4109161 | -1.3090723 | -1.2754009 | -0.0043621 |
| 0.34173751 | 0.64857078 | 1.24478769 | 0.10735631 | 0.47058916 | 0.56260829 | -0.0043621 |
| -1.2185285 | -0.8162799 | -1.9157715 | -1.0939116 | 0.39937353 | -0.9290236 | -0.0043621 |
| -0.6280627 | -0.2134478 | -0.3589301 | -0.8510685 | -0.0413384 | -0.4185695 | -0.0043621 |
| 0.42653322 | 0.74022007 | 0.4056139 | 0.58277464 | 0.55854321 | 0.54273701 | -0.0043621 |
| -0.2106371 | -0.919029 | -1.1156776 | -1.4588101 | -0.989362 | -0.9387032 | -0.0043621 |
| 0.66142237 | 2.94070387 | 1.14843881 | 1.94485116 | 1.55496871 | 1.65007699 | -0.0043621 |
| -0.0529146 | 0.06484365 | -0.5227966 | 0.23855019 | -0.0342746 | -0.0613184 | -0.0043621 |
| 0.4973011 | 0.40752172 | 0.71358728 | 0.64602137 | 0.35120106 | 0.52312651 | -0.0043621 |
| 1.25603867 | 0.49571657 | 1.25169683 | 0.64655495 | 0.27202439 | 0.78440628 | -0.0043621 |
| -0.6046104 | -0.3718147 | -0.5193405 | -0.5520039 | -0.7962437 | -0.5688026 | -0.0043621 |
| 0.09046412 | -0.0213623 | 0.25151539 | -0.0612147 | -0.0944843 | 0.03298364 | -0.0043621 |
| -5.3414698 | -5.4044604 | -4.7243958 | -3.6838861 | -6.4979715 | -5.1304367 | -0.0017709 |
| 6.69975853 | 6.87610006 | 5.77189445 | 5.18248177 | 5.91564226 | 6.08917542 | -0.0012167 |
| -4.0544634 | -3.1014252 | -1.9556773 | -2.635989 | -4.3266997 | -3.2148509 | -0.0012167 |
| -2.6605339 | -3.4079123 | -2.5111699 | -2.8419206 | -2.6952474 | -2.8233568 | -0.0040987 |
| 0.05398655 | -1.784519 | -2.4420731 | -1.668551 | -1.7224839 | -1.5127281 | -0.0020206 |
| -4.1126575 | -1.8355906 | -2.2307675 | -3.2600563 | -2.9501629 | -2.877847 | -0.0059174 |
| 1.68615925 | 1.06059039 | 0.87950242 | 2.98901415 | 2.07257223 | 1.73756769 | -0.0034051 |
| -2.0273533 | -2.0406995 | -1.8083985 | -2.1408861 | -0.6828046 | -1.7400284 | -0.0034051 |
| 2.49043322 | 2.5536809 | 1.95399702 | 3.07277346 | 0.7582736 | 2.16583164 | -0.0034051 |
| -0.4860597 | 0.80790758 | -0.972569 | -0.8923893 | -0.8888748 | -0.486397 | -0.0034051 |
| -0.6595924 | -1.272498 | 0.75297964 | 0.09199572 | 0.40463603 | -0.1364958 | -0.0034051 |
| -0.6673951 | -0.0561285 | -1.1323128 | -0.4133389 | -0.1742554 | -0.4886861 | -0.0034051 |
| 1.44906664 | -2.6887457 | 0.41387963 | 0.85958934 | 2.47119999 | 0.50099797 | -0.0034051 |
| 0.21491098 | -3.1115351 | -0.1220951 | -0.9988012 | -1.1974548 | -1.042995 | -0.0034051 |
| -1.576283 | 0.0832653 | -0.001899 | -0.1274772 | -2.468024 | -0.8180836 | -0.0034051 |
| 1.51670027 | 3.69840884 | 2.65191007 | 0.77240658 | -0.395999 | 1.64868536 | -0.0013535 |
| 0.42918491 | 0.50292373 | 0.28550673 | 0.21219301 | 1.12374294 | 0.51071026 | -0.0013557 |
| -1.6787171 | -1.1298802 | -0.5361562 | 1.01770115 | 0.28834748 | -0.407741 | -0.0013557 |
| -2.3936951 | -3.0862393 | -1.9602122 | -1.7729235 | -3.6987214 | -2.5823583 | -0.0013557 |
| -4.0431886 | -8.6571054 | -5.8645606 | -7.5986013 | -8.7185955 | -6.9764103 | -0.0027046 |
| -0.9120302 | -2.1228127 | -1.6431473 | -2.4884765 | -1.620312 | -1.7573558 | -0.0023475 |
| -2.3426933 | -2.3331177 | -1.739974 | -2.682704 | -3.5506704 | -2.5298319 | -0.0023475 |
| -3.32585 | -2.5075598 | -1.4007039 | -3.9918883 | -2.7468832 | -2.794577 | -0.0023475 |
| -3.1094952 | -6.6855211 | -4.1537356 | -4.7815275 | -5.727931 | -4.8916421 | -0.0023475 |
| 0.28680563 | 2.08078671 | 0.3990376 | -0.9284999 | -1.0187831 | 0.16386938 | -0.0023475 |

es

| esm1v_t33_6 | esm1v_t33_6 | ESM1v_score | Tranception_ | Tranception_ | Tranception_ | Progen2_scor |
|---|---|---|---|---|---|---|

| | | | | | | |
|---|---|---|---|---|---|---|
| -0.2010553 | 0.09675193 | -0.0626496 | -2.7299855 | -2.7298894 | 9.61E-05 | -0.0008884 |
| -0.601337 | -0.9044282 | -0.7595588 | -1.992675 | -1.9926567 | 1.83E-05 | -0.0003379 |
| -0.8239117 | -2.6737413 | 0.18728151 | -0.6377313 | -0.6169946 | 0.02073675 | -0.0018941 |
| 1.14156127 | -0.7939053 | -0.103878 | -2.5050723 | -2.5029411 | 0.00213114 | -0.0013389 |
| -0.4369411 | -0.1761947 | -0.5283755 | -2.5810075 | -2.5826411 | -0.0016336 | -0.0017237 |
| -1.2470453 | -0.9602093 | -1.0613429 | -1.1808499 | -1.1820103 | -0.0011604 | -0.0010476 |
| 0.07545137 | -0.2308688 | 0.03118992 | -2.3700395 | -2.369606 | 0.00043344 | -0.0011935 |
| 2.89273953 | 3.10399389 | 2.97437029 | -2.1745448 | -2.172765 | 0.00177979 | -0.0018399 |
| 0.66319871 | 0.56394839 | 0.59487238 | -2.6313783 | -2.631124 | 0.00025431 | -0.0014411 |
| 3.16257405 | 3.30052948 | 3.18951077 | -2.7402045 | -2.7394431 | 0.00076135 | -0.0021786 |
| 2.45089912 | 2.44946885 | 2.24771309 | -0.9756854 | -0.9685033 | 0.00718206 | -0.0014254 |
| 1.88390064 | 3.24878383 | 2.64229488 | -2.5613756 | -2.5583558 | 0.00301981 | -0.002403 |
| 1.51285696 | 2.73190594 | 1.99698751 | -1.5244766 | -1.5180894 | 0.00638723 | -0.0029394 |
| 2.40181971 | 3.06621528 | 2.19648352 | -2.5548282 | -2.5518658 | 0.00296235 | -0.0019415 |
| 0.1357255 | 0.80535793 | 0.3814364 | -0.903671 | -0.9034594 | 0.0002116 | -0.001 |
| -2.525942 | -2.7424765 | -2.8445882 | -2.5755011 | -2.5764923 | -0.0009912 | -0.0012295 |
| 0.85503197 | 1.00149441 | 0.86362381 | -2.6502688 | -2.6486676 | 0.00160122 | -0.0027362 |
| -0.7219167 | -0.7128742 | -0.6663105 | -2.3569865 | -2.357574 | -0.0005875 | -0.0011759 |
| 1.33908248 | 1.95763493 | 1.81065855 | -1.8964386 | -1.8955112 | 0.00092745 | -0.0005867 |
| 0.51888466 | 0.60468888 | 0.67345204 | -1.9955032 | -1.9953871 | 0.00011611 | -0.0004173 |
| 1.72819734 | 1.84879446 | 1.79323401 | -2.8205913 | -2.8202146 | 0.0003767 | -0.0009289 |
| -2.2183623 | -0.6768293 | -1.4551486 | -1.9162755 | -1.9094197 | 0.00685585 | -0.0013495 |
| 0.55236506 | 0.26547956 | 0.40732536 | -2.5675462 | -2.5674257 | 0.00012048 | -0.0008837 |
| 2.09522724 | 4.99762869 | 2.731599 | -1.1764956 | -1.1660848 | 0.01041079 | -0.0014199 |
| -10.904434 | -9.0361135 | -10.152358 | -2.2373569 | -2.2422857 | -0.0049288 | -0.0010095 |
| 0.43609166 | 0.37809658 | 0.46101716 | -2.4635997 | -2.4636701 | -7.04E-05 | -0.0007406 |
| 3.11883593 | 2.57333446 | 2.95865965 | -2.0606111 | -2.0573228 | 0.00328827 | -0.0013611 |
| 1.29710698 | 1.39988995 | 1.30823302 | -0.7480305 | -0.7366845 | 0.01134598 | -0.0020378 |
| -0.2361195 | -0.2039046 | 0.15782647 | -0.8569175 | -0.8458215 | 0.011096 | -0.0008494 |
| 0.01686311 | -0.0861089 | 0.08898115 | -2.5808938 | -2.5808878 | 5.96E-06 | -0.0003989 |
| 1.64292455 | 2.29325962 | 2.08338671 | -2.2713734 | -2.2710155 | 0.00035795 | -0.000516 |
| 2.14151955 | 1.45160103 | 1.52097669 | -2.7393444 | -2.7361264 | 0.00321794 | -0.0040844 |
| 3.65480375 | 4.54281759 | 3.77096238 | -2.1802125 | -2.1770101 | 0.00320244 | -0.0010707 |
| -0.5253496 | -0.7676063 | -0.6726686 | -2.6460161 | -2.6468158 | -0.0007997 | -0.0012262 |
| 1.33842182 | 1.65676367 | 1.53953781 | -0.9969992 | -0.995566 | 0.00143325 | -0.0011472 |
| 2.20251417 | 3.48465204 | 2.29125862 | -0.7292972 | -0.7089126 | 0.02038461 | -0.000751 |
| 1.15200377 | 0.90305257 | 0.97876058 | -2.7176743 | -2.7173061 | 0.00036812 | -0.0014391 |
| -0.2608752 | -0.7558831 | 0.40641177 | -1.9176302 | -1.9162133 | 0.00141692 | -0.0003965 |
| 0.56808496 | 0.65605712 | 0.61047273 | -2.7061698 | -2.7057633 | 0.0004065 | -0.0009909 |
| -1.2306035 | -0.7765794 | -1.5445543 | -2.1673803 | -2.1671143 | 0.00026608 | -0.0009506 |
| -3.2727942 | -3.280983 | -2.6882149 | -0.6147143 | -0.6199552 | -0.0052409 | -0.0011463 |
| 0.75866365 | 0.69122076 | 0.67952034 | -2.0033855 | -2.0024333 | 0.00095224 | -0.0040337 |
| 0.2426002 | 0.09515667 | 0.17886274 | -0.611027 | -0.6054659 | 0.00556105 | -0.0007266 |
| 5.45729017 | 5.9052248 | 6.40331354 | -0.6848584 | -0.674082 | 0.01077646 | -0.0011184 |
| 7.24696207 | 4.79054999 | 4.94831429 | -1.4728371 | -1.4656814 | 0.00715566 | -0.0006488 |

| | | | | | | |
|---|---|---|---|---|---|---|
| -1.1589415 | -1.1987176 | -1.1958345 | -2.7330303 | -2.7337489 | -0.0007186 | -0.0009678 |
| 0.72212911 | 0.89828873 | 0.70480423 | -2.0276046 | -2.0271749 | 0.00042963 | -0.0008232 |
| -0.5559182 | -0.2512689 | -0.3541015 | -2.3254704 | -2.32354 | 0.00193048 | -0.0009253 |
| 0.73813868 | 1.06216359 | 0.86143479 | -2.717562 | -2.7166564 | 0.00090551 | -0.002469 |
| -1.0339258 | -1.1739757 | -1.1097815 | -1.9063194 | -1.9072564 | -0.000937 | -0.0013135 |
| -0.9157913 | -0.9129241 | -0.8619385 | -2.161474 | -2.162286 | -0.0008121 | -0.0016497 |
| 0.74717832 | 0.80053043 | 0.62090182 | -1.6773593 | -1.6790117 | -0.0016524 | -0.0003766 |
| 0.02736521 | -0.0277648 | 0.14187112 | -1.7054051 | -1.7048317 | 0.0005734 | -0.0026664 |
| -0.4421737 | -0.302784 | -0.5734071 | -1.4402336 | -1.447982 | -0.0077484 | -0.0081358 |
| 0.70222902 | -0.5159631 | 0.15620074 | -2.882823 | -2.882654 | 0.00016904 | -0.002999 |
| 2.30584192 | 2.96069455 | 2.49590697 | -2.4981999 | -2.4928892 | 0.00531077 | -0.0054477 |
| 5.00729799 | 4.80278444 | 5.00775156 | -1.6527517 | -1.6466577 | 0.00609398 | -0.0018404 |
| 0.94154704 | 0.54596949 | 0.56147206 | -2.6163565 | -2.6151562 | 0.00120036 | -0.0011486 |
| 3.44362271 | 3.83831632 | 3.76520305 | -2.0637005 | -2.0626402 | 0.0010603 | -0.0005903 |
| 7.97780561 | 5.98752785 | 6.6740489 | -1.0348753 | -1.0205598 | 0.01431549 | -0.0019529 |
| 1.13383627 | 0.42472291 | 1.29479251 | -2.2467225 | -2.245055 | 0.0016675 | -0.0036162 |
| 0.31098557 | 0.087345 | 0.15045538 | -2.8269811 | -2.8266716 | 0.00030947 | -0.0039595 |
| 0.39414179 | 0.62910533 | 0.50891538 | -1.1471834 | -1.1446936 | 0.00248981 | -0.0038648 |
| -1.9184434 | -1.1932125 | -0.6399354 | -0.4367947 | -0.4401834 | -0.0033887 | -0.0026118 |
| 1.80821383 | 1.95971322 | 2.65560071 | -1.3671319 | -1.3089516 | 0.05818033 | -0.0136611 |
| 0.64052105 | 0.61847329 | 0.73012519 | -1.9248818 | -1.921643 | 0.0032388 | -0.0046456 |
| 2.50024223 | 2.32303429 | 0.9417243 | -1.3724651 | -1.3827729 | -0.0103078 | -0.0054044 |
| 1.84650636 | 1.87041664 | 1.30237799 | -2.8614299 | -2.8608634 | 0.00056648 | -0.0014478 |
| 0.88326955 | 0.93245065 | 1.03236792 | -2.3421099 | -2.3321896 | 0.00992036 | -0.0093348 |
| 6.54888833 | 4.8701973 | 6.27210965 | -2.5792303 | -2.5779595 | 0.00127077 | -0.0004134 |
| -0.2054527 | -0.532182 | -0.1518825 | -2.2213821 | -2.2219946 | -0.0006125 | -0.0014487 |
| -0.0576425 | 0.24543738 | -0.0230992 | -1.4890624 | -1.4878311 | 0.00123131 | -0.0091347 |
| -0.1249347 | -0.387172 | -1.2256905 | -2.5423267 | -2.5389397 | 0.00338697 | -0.0071124 |
| 4.06144333 | 0.14409781 | 1.47499709 | -2.6936755 | -2.6920896 | 0.00158596 | -0.0038035 |
| 1.31445265 | 0.95915151 | 0.90444722 | -2.6374648 | -2.6344967 | 0.00296807 | -0.0061726 |
| 4.07692385 | 4.49962902 | 3.8885674 | -1.7395846 | -1.6987096 | 0.04087496 | -0.0087184 |
| -0.151998 | 0.36878967 | -0.3195269 | -2.6424456 | -2.6422863 | 0.00015926 | -0.0015533 |
| 1.16952801 | 1.25397944 | 1.28382702 | -1.5442294 | -1.5424575 | 0.00177193 | -0.0016978 |
| 0.19272089 | 0.78641129 | 0.75923412 | -2.7298619 | -2.72953 | 0.00033188 | -0.0010222 |
| -3.8603128 | -3.8390613 | -3.5195328 | -2.701205 | -2.7212811 | -0.020076 | -0.0071953 |
| 2.38681507 | 3.21325874 | 2.72066841 | -1.4267678 | -1.4202048 | 0.00656307 | -0.0030304 |
| 0.0902617 | 0.35125065 | 0.77039728 | -2.540802 | -2.5398715 | 0.00093055 | -0.0018089 |
| -0.380625 | -0.2962689 | -0.0735474 | -1.8842547 | -1.8836092 | 0.00064552 | -0.0006721 |
| 1.18847752 | -0.0260427 | 0.81797271 | -1.6514149 | -1.6690829 | -0.017668 | -0.0112793 |
| -1.3041105 | -1.7143295 | -1.462521 | -2.706332 | -2.7083106 | -0.0019786 | -0.0029864 |
| 4.70381904 | 2.66630888 | 3.07549691 | -0.9645742 | -0.9516304 | 0.01294374 | -0.0012028 |
| -0.0221684 | -0.2119899 | 0.26347637 | -2.0702224 | -2.0688162 | 0.00140619 | -0.0005843 |
| 5.46177101 | 6.1194005 | 5.18510227 | -1.6826535 | -1.656567 | 0.02608657 | -0.006331 |
| 0.22286463 | 0.00484419 | 0.20393548 | -2.0691068 | -2.0685751 | 0.00053167 | -0.0025842 |
| -1.3559275 | -1.8957797 | -1.4005343 | -1.5033691 | -1.5038338 | -0.0004647 | -0.0033755 |

| | | | | | | |
|---|---|---|---|---|---|---|
| -2.2184002 | -2.2781935 | -0.8789611 | -2.1226108 | -2.1234004 | -0.0007896 | -0.0052328 |
| 3.13840032 | 5.92938423 | 4.993152 | -0.8618349 | -0.8430778 | 0.0187571 | -0.0006241 |
| 0.12163162 | 0.9729923 | 0.41979377 | -1.9890624 | -1.9884275 | 0.00063491 | -0.0017446 |
| 1.3039422 | 1.84074783 | -0.1262242 | -1.6191937 | -1.6089504 | 0.0102433 | -0.0012575 |
| -0.6846609 | -0.2962155 | -0.3061728 | -1.6593702 | -1.656994 | 0.0023762 | -0.0026043 |
| 2.29923165 | -1.8530626 | 0.15676124 | -1.0183431 | -1.0192786 | -0.0009356 | -0.0017306 |
| 0.70340049 | 1.07676768 | 0.85097628 | -2.0788021 | -2.0762351 | 0.00256705 | -0.0043125 |
| 0.08859324 | 0.40684605 | 0.19557552 | -2.5935051 | -2.5824795 | 0.01102567 | -0.0161528 |
| 0.26924801 | 0.16086912 | 0.25272579 | -2.3192601 | -2.3213551 | -0.002095 | -0.0020644 |
| 2.28021932 | 3.92853522 | 2.88954554 | -1.3231685 | -1.3068405 | 0.01632798 | -0.0049317 |
| -87.74816 | -149.3978 | -119.45548 | -1.1279103 | -1.7560124 | -0.6281022 | -0.0032143 |
| -4.5547285 | -4.1792502 | -4.4076517 | -2.8635077 | -2.8734505 | -0.0099428 | -0.0064392 |
| 5.09512734 | 3.26297021 | 4.34476233 | -2.12502 | -2.1226847 | 0.00233531 | -0.000596 |
| 0.47462797 | -1.7689042 | -0.8732095 | -1.6936713 | -1.6962665 | -0.0025952 | -0.0016112 |
| 5.54295921 | 7.07930326 | 6.28472385 | -2.6060545 | -2.5818772 | 0.02417731 | -0.00238 |
| -4.592227 | -3.9561629 | -3.9384828 | -2.5017288 | -2.5055192 | -0.0037904 | -0.0037171 |
| 1.18682241 | 1.13795424 | 1.39186053 | -1.35197 | -1.3480742 | 0.00389576 | -0.0026509 |
| 4.00660634 | 5.84266663 | 3.92274561 | -2.0149713 | -1.9796748 | 0.03529644 | -0.008202 |
| 1.89772725 | 1.78457689 | 1.96413953 | -1.5047575 | -1.5023383 | 0.00241923 | -0.0019503 |
| 6.2041173 | 2.56598341 | 3.99194887 | -1.8749663 | -1.852156 | 0.02281022 | -0.0041843 |
| 5.0824157 | 5.29291904 | 4.95322318 | -1.6560966 | -1.6167556 | 0.03934097 | -0.0060546 |
| 2.3669095 | 2.8587904 | 2.22278304 | -0.8481347 | -0.8092991 | 0.03883559 | -0.0037908 |
| -0.7932091 | -0.5192943 | -0.638857 | -2.208987 | -2.2087283 | 0.00025868 | -0.0013034 |
| 0.42129946 | 0.84350348 | 0.5913353 | -1.0980308 | -1.0946571 | 0.00337374 | -0.0032149 |
| -0.1120179 | 0.06656337 | -0.2011736 | -1.9708692 | -1.961511 | 0.00935817 | -0.0055917 |
| 3.22299647 | 3.12290621 | 2.43976564 | -2.0268738 | -2.022922 | 0.00395179 | -0.0036205 |
| 2.01492834 | 1.78971434 | 1.84325316 | -2.7383749 | -2.7309132 | 0.00746179 | -0.0097025 |
| 2.16647649 | 2.55459404 | 2.23094499 | -2.566374 | -2.5659291 | 0.00044489 | -0.0017387 |
| 2.31288576 | 2.63855028 | 1.9045388 | -1.1387188 | -1.1254373 | 0.01328158 | -0.0316486 |
| -1.1367397 | -1.1184058 | -0.9125271 | -0.6069209 | -0.6129102 | -0.0059893 | -0.0008513 |
| -2.5418553 | 0.704211 | -0.8127424 | -1.5220951 | -1.5249619 | -0.0028669 | -0.007704 |
| 1.19716907 | 2.39045167 | 2.59842143 | -1.3677784 | -1.3650622 | 0.00271618 | -0.0016724 |
| 1.55044889 | 1.67338324 | 2.64285111 | -1.1475086 | -1.1394378 | 0.00807083 | -0.0024758 |
| 4.83242893 | 5.52302647 | 5.37357321 | -1.5593932 | -1.5517168 | 0.00767644 | -0.0003419 |
| 3.8625257 | 1.39943719 | 2.30707957 | -0.6406263 | -0.6293933 | 0.01123297 | -0.0024019 |
| -0.5026584 | 0.07980442 | -0.5008157 | -1.7154078 | -1.7166224 | -0.0012145 | -0.0015089 |
| 1.39344645 | 0.41092825 | 1.44789753 | -0.8212193 | -0.8103257 | 0.01089352 | -0.0030737 |
| 6.71213365 | 5.68685627 | 6.0154201 | -1.8947619 | -1.8669943 | 0.02776766 | -0.0056595 |
| 0.30779791 | 0.28337383 | 0.33850298 | -1.6314447 | -1.6311216 | 0.00032306 | -0.0026996 |
| -0.0696275 | 3.09756422 | 0.86204095 | -1.1180189 | -1.1084863 | 0.00953257 | -0.0020442 |
| -0.7101436 | -0.0175173 | -0.2162607 | -1.7770929 | -1.7739142 | 0.00317872 | -0.0006109 |
| -0.0721095 | 0.08477068 | -0.0073449 | -2.8222785 | -2.8222136 | 6.48E-05 | -0.0018089 |
| -1.3102062 | -1.0645847 | -1.1742632 | -1.5318162 | -1.5338446 | -0.0020283 | -0.0047174 |
| 1.23831654 | 1.22793031 | 1.09547725 | -1.815724 | -1.8152051 | 0.00051892 | -0.0003119 |
| -0.2973263 | -0.5221183 | -0.4105897 | -1.1788464 | -1.1680998 | 0.0107466 | -0.0059831 |

| | | | | | | |
|---|---|---|---|---|---|---|
| 2.80818748 | 2.79446793 | 3.29325323 | -2.1100454 | -2.1022289 | 0.00781655 | -0.00431 |
| -0.9556532 | -0.7234035 | -0.9044153 | -2.7829035 | -2.7832508 | -0.0003473 | -0.0014861 |
| -1.2714968 | -1.8787237 | -1.8429781 | -2.8460567 | -2.8469629 | -0.0009062 | -0.0014208 |
| 6.52794409 | 2.54060507 | 4.38771129 | -2.5900869 | -2.5889783 | 0.00110865 | -0.0020026 |
| 1.47685492 | 1.1157279 | 0.87112008 | -1.1207668 | -1.1073898 | 0.01337695 | -0.0007688 |
| 0.7002629 | 1.24435091 | 1.33310378 | -1.6347173 | -1.6335312 | 0.00118613 | -0.0022914 |
| 1.45697141 | 1.37246394 | 1.6629683 | -2.7507617 | -2.7492156 | 0.00154614 | -0.0023405 |
| 0.2474227 | -2.5051832 | -0.7021281 | -0.6073373 | -0.6110507 | -0.0037134 | -0.0009128 |
| 0.86195028 | 0.86340189 | 0.87842274 | -2.8385062 | -2.8303294 | 0.0081768 | -0.0169261 |
| 0.95086074 | 0.89376211 | 1.23446014 | -1.9011797 | -1.884358 | 0.01682162 | -0.0062107 |
| 1.08236647 | 0.33961678 | 0.65720868 | -1.5939362 | -1.5912173 | 0.00271893 | -0.0006696 |
| 0.59190297 | 0.9059968 | 0.70743175 | -2.8315623 | -2.8286543 | 0.00290799 | -0.0146976 |
| 0.61743557 | -0.7677164 | 0.37575302 | -1.4959505 | -1.4971154 | -0.0011649 | -0.0030007 |
| 6.96130753 | 5.24526024 | 5.07197905 | -0.845464 | -0.825857 | 0.01960695 | -0.0013639 |
| 0.66265774 | 0.30147243 | 0.75517583 | -2.1499972 | -2.1458316 | 0.00416565 | -0.0005451 |
| 3.78338766 | 4.16156268 | 4.1763989 | -2.8419685 | -2.840573 | 0.00139554 | -0.0008615 |
| 0.36550808 | 6.07956982 | 2.33966422 | -0.7465237 | -0.7266786 | 0.01984513 | -0.0017258 |
| 32.6123536 | 33.5314003 | 33.3600183 | -1.2789593 | -1.0824498 | 0.19650948 | -0.0052023 |
| 0.11566257 | -0.6609621 | 0.36887112 | -1.8812164 | -1.8812178 | -1.43E-06 | -0.0036612 |
| 6.31952572 | 3.41179705 | 4.74229708 | -1.9469746 | -1.9342523 | 0.01272237 | -0.0045135 |
| -0.7676264 | -1.7438524 | -1.3213378 | -1.3886931 | -1.3881404 | 0.00055265 | -0.0009867 |
| 1.30603182 | 1.48690212 | 1.40909464 | -2.2910089 | -2.2861521 | 0.00485682 | -0.0056763 |
| -2.2533505 | -0.7630076 | 1.37632265 | -0.3040255 | -0.2577772 | 0.04624829 | -0.0010739 |
| -3.1294991 | -2.7119857 | -2.5110018 | -2.2314088 | -2.2272389 | 0.00416994 | -0.0037536 |
| 1.93835771 | 1.81326199 | 1.89216115 | -1.6097413 | -1.6038917 | 0.0058496 | -0.0033294 |
| 0.19654918 | -0.0732527 | 0.07630625 | -0.8623556 | -0.8625923 | -0.0002367 | -0.0013769 |
| -1.3318224 | -0.5365582 | 0.02115209 | -1.3871887 | -1.3852656 | 0.00192308 | -0.0012553 |
| 3.88401508 | 5.70128679 | 6.08708534 | -1.0722729 | -1.0567876 | 0.01548529 | -0.0028502 |
| 0.7159425 | -0.5765831 | -0.4439259 | -1.0999531 | -1.0970321 | 0.00292099 | -0.0003143 |
| 2.39498186 | 2.98519289 | 3.3317297 | -2.6789854 | -2.6806555 | -0.0016701 | -0.0064415 |
| 0.89245987 | 0.8552289 | 0.82469482 | -2.8659182 | -2.8589046 | 0.00701356 | -0.0110811 |
| 1.3173492 | 0.86456752 | 0.89395361 | -0.7706791 | -0.7705839 | 9.51E-05 | -0.0005804 |
| -0.6857793 | -1.0293909 | -0.9642129 | -2.4767833 | -2.4838212 | -0.0070379 | -0.0100319 |
| 3.32332325 | 2.78667307 | 2.3647593 | -1.7361449 | -1.7344315 | 0.0017134 | -0.0031094 |
| 6.01676273 | 5.95810509 | 5.89835358 | -0.5082177 | -0.5015726 | 0.00664508 | -0.0007869 |
| -1.0587418 | -1.0233142 | -1.0976799 | -1.9056183 | -1.9060197 | -0.0004014 | -0.0004022 |
| -1.6299791 | -0.3163443 | -0.0760817 | -1.7747191 | -1.7768527 | -0.0021336 | -0.004551 |
| -0.947763 | 0.24053288 | -1.0606695 | -2.1233728 | -2.1261425 | -0.0027697 | -0.0035858 |
| 4.43588591 | -0.5889727 | 2.08123236 | -2.2637596 | -2.2654133 | -0.0016537 | -0.0028694 |
| 3.55957723 | 2.74204373 | 2.95974305 | -1.8300288 | -1.8124281 | 0.01760066 | -0.0041532 |
| 1.47500467 | 2.72586739 | 2.01131647 | -2.480602 | -2.4788978 | 0.00170422 | -0.0009817 |
| 7.09465122 | 4.57793617 | 5.07431726 | -0.5138707 | -0.5059975 | 0.00787318 | -0.0012569 |
| 1.82294869 | -0.0191648 | 0.92169862 | -2.2292109 | -2.2266598 | 0.00255108 | -0.0048851 |
| -0.250524 | -0.4246957 | -0.3649613 | -2.5995288 | -2.5997434 | -0.0002146 | -0.0012716 |
| 0.78617203 | 1.41061759 | 1.07523174 | -1.6527474 | -1.6462085 | 0.00653887 | -0.0007095 |

| | | | | | | |
|---|---|---|---|---|---|---|
| -0.1664 | 0.76548266 | 0.61462829 | -1.5431502 | -1.5276787 | 0.01547146 | -0.003776 |
| 0.44124436 | 0.49170637 | 0.61825051 | -1.9437178 | -1.9362484 | 0.00746942 | -0.0040457 |
| 1.24287343 | 2.28674984 | 1.70541582 | -2.3379669 | -2.3356972 | 0.00226974 | -0.0023194 |
| -0.5589449 | -1.0107865 | -0.6745714 | -1.3881098 | -1.3908967 | -0.0027869 | -0.0072187 |
| -0.198777 | -0.1342282 | -0.2931255 | -2.1912417 | -2.1922042 | -0.0009625 | -0.000857 |
| 0.92587376 | 4.08032227 | 2.70364962 | -0.6910493 | -0.6803699 | 0.01067942 | -0.001801 |
| 2.15974569 | 2.41556549 | 1.30457487 | -1.9976228 | -1.9982861 | -0.0006633 | -0.002792 |
| 0.44314098 | 0.22599888 | 0.20413442 | -2.8808963 | -2.8813453 | -0.0004489 | -0.0143127 |
| 1.15011883 | 1.16434324 | 1.08445532 | -2.1848557 | -2.1814892 | 0.00336647 | -0.0113588 |
| 2.25383806 | 1.97016788 | 2.08397098 | -2.1830781 | -2.158541 | 0.02453709 | -0.008475 |
| -0.6229312 | -0.912559 | -0.6270857 | -2.3860304 | -2.3866 | -0.0005696 | -0.0025678 |
| 1.43953586 | 1.75364804 | 1.72743301 | -2.8906658 | -2.8828471 | 0.0078187 | -0.0181883 |
| -1.1723886 | -1.0806491 | -1.3387427 | -2.1516941 | -2.1546824 | -0.0029883 | -0.0021181 |
| -3.4279985 | -3.0576929 | -3.1253094 | -2.543628 | -2.5490537 | -0.0054257 | -0.0046107 |
| -0.4024267 | 1.39949811 | 0.00277379 | -1.4058119 | -1.4027472 | 0.00306475 | -0.002444 |
| -6.0559704 | -0.386161 | -5.0486073 | -2.3115208 | -2.3174987 | -0.0059779 | -0.008104 |
| -0.8903574 | -0.881062 | -1.1675199 | -1.452167 | -1.4535013 | -0.0013343 | -0.0016411 |
| -0.9942226 | -0.7090945 | -0.8423121 | -1.8976099 | -1.9002572 | -0.0026473 | -0.0005538 |
| 8.05960155 | 5.12932444 | 9.00318429 | -1.6994551 | -1.6742787 | 0.02517641 | -0.0015049 |
| 1.25575113 | 3.64697552 | 2.77008376 | -0.6689858 | -0.6643974 | 0.00458843 | -0.0007958 |
| -0.2655883 | -0.2146332 | -0.2914227 | -2.5017052 | -2.5028348 | -0.0011296 | -0.0063794 |
| 0.3868854 | 0.30447364 | 0.43701887 | -2.8382881 | -2.8351824 | 0.00310564 | -0.007179 |
| 0.09713745 | 1.23146367 | 0.70552206 | -2.2498868 | -2.2502186 | -0.0003319 | -0.0051494 |
| -0.5010149 | -0.9568994 | -0.6351779 | -2.638716 | -2.6439478 | -0.0052319 | -0.0017434 |
| -0.1852818 | 3.6588769 | 2.75703192 | -1.8189896 | -1.8083411 | 0.01064849 | -0.0021945 |
| 7.57535839 | 8.77568626 | 8.21796446 | -0.5570151 | -0.5327456 | 0.02426952 | -0.0014895 |
| 1.76898694 | 2.37855768 | 1.83807869 | -0.8795343 | -0.8720823 | 0.00745201 | -0.0041701 |
| 1.08579671 | 2.04129696 | 2.72148595 | -1.5524743 | -1.5584773 | -0.006003 | -0.0059735 |
| 6.1138134 | 7.2630105 | 5.62092669 | -1.7391809 | -1.7277414 | 0.01143956 | -0.0020566 |
| 2.68558884 | 2.76222456 | 3.70440426 | -2.0585988 | -2.0510325 | 0.00756621 | -0.0026059 |
| 7.76177597 | 5.02824581 | 5.59284594 | -1.7155421 | -1.7093533 | 0.00618875 | -0.0021362 |
| 3.14080811 | 4.00783229 | 3.45371113 | -1.9508035 | -1.9497579 | 0.00104558 | -0.0022668 |
| 1.62308502 | 0.90199852 | 0.59798589 | -1.4777367 | -1.4750277 | 0.00270903 | -0.0038781 |
| -0.024647 | -0.0584712 | 0.05652137 | -2.8589687 | -2.8591452 | -0.0001764 | -0.0041975 |
| 2.69026041 | 2.36792779 | 2.55194926 | -2.902158 | -2.8863695 | 0.01578856 | -0.0154931 |
| 12.7079988 | 9.71664095 | 12.4018438 | -2.3498039 | -2.3347468 | 0.01505709 | -0.0040199 |
| 0.58413267 | 1.89914751 | 0.71548853 | -2.6023827 | -2.6033196 | -0.000937 | -0.0035637 |
| 2.3377707 | 3.74251938 | 2.90509977 | -1.4418521 | -1.4310427 | 0.01080942 | -0.0048313 |
| 0.96995401 | 1.70226324 | 0.52834022 | -2.3437529 | -2.3444793 | -0.0007265 | -0.004376 |
| -0.6305747 | -0.4999161 | -0.8062689 | -1.2329011 | -1.2357982 | -0.0028971 | -0.0018945 |
| -0.2682307 | -0.5333813 | -0.2510042 | -2.843627 | -2.8449614 | -0.0013344 | -0.0059677 |
| -1.6038918 | 0.27255654 | -0.9864094 | -2.2932878 | -2.2949209 | -0.0016332 | -0.0035365 |
| 0.60210848 | 0.09022474 | 0.93878043 | -2.304239 | -2.2955174 | 0.00872159 | -0.0031981 |
| 1.96898007 | 1.48556137 | 1.97040448 | -1.8917259 | -1.8828468 | 0.00887907 | -0.0048131 |
| -2.1315837 | -1.0735972 | -1.5135959 | -2.2972178 | -2.3020663 | -0.0048485 | -0.0012707 |

| | | | | | | |
|---|---|---|---|---|---|---|
| 0.89523196 | 2.43439794 | 1.21103538 | -1.8512801 | -1.852088 | -0.0008079 | -0.0030983 |
| 9.0546602 | 10.2264011 | 7.3498997 | -2.3146644 | -2.3010056 | 0.01365876 | -0.0037831 |
| 1.72236872 | 1.8737731 | 1.67544045 | -2.5312877 | -2.5266888 | 0.00459886 | -0.0053965 |
| -0.0739827 | 0.11147237 | -0.0219928 | -2.7895217 | -2.789607 | -8.54E-05 | -0.0008011 |
| 1.72888517 | 0.63379097 | 1.9847996 | -1.6213018 | -1.6178259 | 0.0034759 | -0.0025543 |
| 5.36434329 | 3.98088825 | 4.65639296 | -1.8947619 | -1.8690294 | 0.02573252 | -0.0056595 |
| -0.0074592 | 0.27004933 | 0.06126437 | -1.9292519 | -1.9298649 | -0.000613 | -0.0006098 |
| 2.98776078 | 3.40404129 | 2.75084894 | -2.2897952 | -2.285512 | 0.00428319 | -0.0009867 |
| 1.08287978 | 0.99382114 | 1.14659529 | -2.0794036 | -2.078769 | 0.00063467 | -0.001021 |
| 3.35891557 | 4.22255373 | 2.78400295 | -1.863965 | -1.8613988 | 0.00256622 | -0.0022749 |
| 1.66519356 | 1.38891196 | 1.57973914 | -1.8195163 | -1.7709546 | 0.04856169 | -0.0072195 |
| -0.4251397 | -0.1854051 | -0.3077653 | -2.0108309 | -2.0112605 | -0.0004296 | -0.0009345 |
| -0.2124145 | 0.10847902 | -0.3560808 | -1.1768463 | -1.1764709 | 0.00037539 | -0.0024908 |
| 1.02991199 | 0.75595403 | 0.91427855 | -2.1555367 | -2.1534097 | 0.00212693 | -0.0078778 |
| -0.4587986 | -0.6888356 | -0.6107107 | -0.9351937 | -0.9271893 | 0.00800443 | -0.0022907 |
| 55.1178843 | 64.2124202 | 57.8758974 | -1.9107206 | -1.6225842 | 0.28813636 | -0.0047351 |
| 0.18251657 | 1.33188462 | 1.23168864 | -1.8484164 | -1.8306996 | 0.01771688 | -0.0043641 |
| -1.4606926 | -2.7697058 | -2.8532543 | -0.9590168 | -0.9586461 | 0.00037074 | -0.0013682 |
| 2.73649383 | 1.70137191 | 2.05960269 | -1.5127215 | -1.4975803 | 0.01514125 | -0.0049086 |
| 1.45635295 | 1.1798296 | 1.38021941 | -1.281088 | -1.2664669 | 0.01462114 | -0.0024692 |
| 1.17515576 | 0.83349478 | 1.20498915 | -2.3815861 | -2.3563397 | 0.02524638 | -0.0130085 |
| -2.1914215 | 6.73094463 | 0.11447792 | -2.3462796 | -2.3366609 | 0.00961876 | -0.0038876 |
| -1.1089277 | -3.6900074 | -2.4698492 | -1.6654401 | -1.6658871 | -0.000447 | -0.0014752 |
| 0.45186079 | 0.62733197 | 1.62119677 | -1.8194293 | -1.8203897 | -0.0009605 | -0.0019693 |
| 3.40063667 | 4.20883465 | 3.47688084 | -2.636323 | -2.6354914 | 0.0008316 | -0.0026099 |
| -2.8112292 | -2.2063372 | -0.554298 | -2.6991963 | -2.6963403 | 0.00285602 | -0.0022051 |

cid inserted into Jonathan's protein

| esm1v_t33_6 | esm1v_t33_6 | esm1v_t33_6 | esm1v_t33_6 | esm1v_t33_6 | ESM1v_score | Progen2_scor |
|---|---|---|---|---|---|---|
| 0.48966169 | 0.66403961 | 0.66729975 | 0.54080701 | 0.62521505 | 0.59740462 | -0.0008884 |
| -0.8552408 | -0.8077595 | -0.3669455 | -0.7418623 | -0.5284631 | -0.6600543 | -0.0008884 |
| -0.632349 | -0.663434 | -0.9962459 | -0.601337 | -0.9044282 | -0.7595588 | -0.0003379 |
| 1.02608871 | 3.88777018 | -0.4797983 | -0.8239117 | -2.6737413 | 0.18728151 | -0.0018941 |
| 2.30216312 | 1.36650276 | 0.50906324 | 1.89431012 | 1.40196848 | 1.49480155 | -0.0013389 |
| -1.1536312 | -0.7937465 | -0.7879331 | -0.7736574 | -1.4711863 | -0.9960309 | -0.0013389 |
| -0.8340931 | -0.3896632 | -0.607506 | -0.6376362 | -0.5054514 | -0.59487 | -0.0013389 |
| -0.4280472 | -0.4630427 | -0.4933448 | -0.1341906 | -0.3702135 | -0.3777678 | -0.0013389 |
| 0.37079096 | 0.15832865 | 0.37711287 | 0.79273534 | 0.15097749 | 0.36998906 | -0.0013389 |
| -0.6422627 | -1.0367296 | -0.6918893 | -0.5562439 | -0.5579448 | -0.697014 | -0.0017237 |
| 0.1539607 | -0.4369447 | 0.62512398 | 0.11930275 | 0.38175011 | 0.16863856 | -0.0017237 |
| -0.6635178 | 0.10415125 | -0.2885898 | -0.4302657 | -0.3145955 | -0.3185635 | -0.0010476 |
| -0.5040733 | -0.7780774 | -0.9693528 | -0.8167796 | -0.6456138 | -0.7427794 | -0.0010476 |
| 0.03089523 | 0.18389463 | 0.09657717 | 0.07545137 | -0.2308688 | 0.03118992 | -0.0011935 |
| -0.0761843 | 0.29932714 | 0.02263021 | -0.0389616 | -0.0645354 | 0.02845521 | -0.0018399 |
| 0.17629743 | -0.0283275 | -0.0103736 | -0.0654769 | -0.0460818 | 0.00520754 | -0.0018399 |
| 2.84311223 | 3.56168509 | 2.08695126 | 2.99717808 | 3.21461105 | 2.94070754 | -0.0018399 |

| | | | | | | |
|---|---|---|---|---|---|---|
| 0.79823565 | 0.5347743 | 0.33924198 | 0.08008933 | 0.46252418 | 0.44297309 | -0.0014411 |
| 0.03104568 | -0.0389233 | 0.08284044 | 0.58310938 | 0.10142422 | 0.15189929 | -0.0014411 |
| 2.51187038 | 1.93327713 | 1.5793705 | 2.26712704 | 2.30200791 | 2.11873059 | -0.0021786 |
| 0.92457104 | 1.49415612 | 1.04120517 | 0.89544702 | 0.99852157 | 1.07078018 | -0.0021786 |
| 2.44906807 | 2.57481599 | 1.31431341 | 2.45089912 | 2.44946885 | 2.24771309 | -0.0014254 |
| 2.69421959 | 2.95001864 | 2.43455172 | 1.88390064 | 3.24878383 | 2.64229488 | -0.002403 |
| 1.7196902 | 1.33814383 | 2.68234062 | 1.51285696 | 2.73190594 | 1.99698751 | -0.0029394 |
| 1.124542 | 0.23345351 | 0.93300939 | 0.85235739 | 0.99515104 | 0.82770267 | -0.0019415 |
| -0.8229222 | -0.6669083 | -0.724194 | -0.7133274 | -0.6928561 | -0.7240416 | -0.0019415 |
| 0.49979925 | 0.4323864 | 0.21375346 | 0.36755776 | 0.23135805 | 0.34897099 | -0.0019415 |
| 0.08292747 | -0.9212728 | -0.2363846 | 0.45420909 | -0.0305686 | -0.1302179 | -0.0019415 |
| -0.1485016 | 0.01425123 | -0.2590578 | -0.1805491 | -0.0608749 | -0.1269464 | -0.0019415 |
| 1.53256559 | 2.54856014 | 1.67837548 | 1.62157202 | 2.62400579 | 2.00101581 | -0.0019415 |
| 0.45058823 | 0.67884159 | -0.1633313 | 0.1357255 | 0.80535793 | 0.3814364 | -0.001 |
| -0.3977785 | -0.8159033 | -0.5097938 | -0.5341108 | -0.3301125 | -0.5175398 | -0.0012295 |
| -2.5585499 | -2.3421063 | -2.3303909 | -1.9918312 | -2.412364 | -2.3270485 | -0.0012295 |
| 0.79195857 | 0.5623467 | 1.10728741 | 0.85503197 | 1.00149441 | 0.86362381 | -0.0027362 |
| -0.7759488 | -0.7055809 | -0.4152317 | -0.7219167 | -0.7128742 | -0.6663105 | -0.0011759 |
| 1.47511435 | 1.55930305 | 2.72215796 | 1.33908248 | 1.95763493 | 1.81065855 | -0.0005867 |
| 0.66281033 | 0.84997272 | 0.73090363 | 0.51888466 | 0.60468888 | 0.67345204 | -0.0004173 |
| 0.97449136 | 0.85608125 | 0.91638565 | 0.91725016 | 0.96303463 | 0.92544861 | -0.0009289 |
| 0.97617245 | 1.00770736 | 0.65834022 | 0.81094718 | 0.88575983 | 0.86778541 | -0.0009289 |
| -1.6243868 | -0.3840847 | -2.3720798 | -2.2183623 | -0.6768293 | -1.4551486 | -0.0013495 |
| 0.76612186 | 0.30434132 | 0.57701063 | 0.44839525 | 0.39976454 | 0.49912672 | -0.0008837 |
| -0.3703449 | 0.08142352 | -0.1397703 | 0.10396981 | -0.134285 | -0.0918014 | -0.0008837 |
| 1.42871165 | 2.74760675 | 2.38882065 | 2.09522724 | 4.99762869 | 2.731599 | -0.0014199 |
| -2.5608478 | -2.8581939 | -1.5034502 | -2.3717799 | -2.4031913 | -2.3394926 | -0.0010095 |
| 0.70475602 | 1.22442341 | 1.04393625 | 0.9102757 | 0.57472658 | 0.89162359 | -0.0010095 |
| -1.8950318 | -0.9448308 | -1.4089131 | -2.0724654 | -1.5060308 | -1.5654544 | -0.0010095 |
| -1.4926791 | -1.1564627 | -1.4634941 | -1.0640273 | -1.1965835 | -1.2746493 | -0.0010095 |
| -1.1208112 | -0.7314916 | -0.859863 | -1.0634253 | 0.80872035 | -0.5933742 | -0.0010095 |
| 0.04660606 | 0.41799235 | -0.2169724 | 1.0513773 | -0.0843205 | 0.24293656 | -0.0010095 |
| -3.9880667 | -3.1648231 | -4.2712336 | -4.5132408 | -3.5524139 | -3.8979556 | -0.0010095 |
| -0.0233588 | -0.5315347 | 0.5579617 | -0.6501818 | 0.08734941 | -0.1119528 | -0.0010095 |
| -1.3344338 | -1.7470635 | -1.7918987 | -2.0599768 | -1.5091376 | -1.6885021 | -0.0010095 |
| -1.7801268 | -1.8498315 | -1.9042687 | -1.2245677 | -1.9536521 | -1.7424894 | -0.0010095 |
| 0.4220252 | 0.739079 | 1.02416372 | 1.08881426 | 0.77350092 | 0.80951662 | -0.0010095 |
| 0.24445868 | 0.80169392 | 0.69626808 | 0.74805927 | 0.38253546 | 0.57460308 | -0.0010095 |
| 0.93950605 | 1.30171824 | -0.3861494 | 0.31670451 | 0.54238367 | 0.54283261 | -0.0010095 |
| 0.41278815 | 0.45629668 | 0.6218127 | 0.43609166 | 0.37809658 | 0.46101716 | -0.0007406 |
| -0.1393113 | 0.16678882 | -0.0897868 | -0.0887518 | -0.0463932 | -0.0394908 | -0.0013611 |
| 2.99191427 | 3.54496551 | 2.62655735 | 3.20758772 | 2.61972761 | 2.99815049 | -0.0013611 |
| 1.1553545 | 0.84920979 | 1.8396039 | 1.29710698 | 1.39988995 | 1.30823302 | -0.0020378 |
| 0.8388257 | -0.0134621 | 0.40379286 | -0.2361195 | -0.2039046 | 0.15782647 | -0.0008494 |
| 0.12568164 | 0.04076886 | 0.34770107 | 0.01686311 | -0.0861089 | 0.08898115 | -0.0003989 |

| | | | | | | |
|---|---|---|---|---|---|---|
| 2.84079456 | 1.80914402 | 1.83081079 | 1.64292455 | 2.29325962 | 2.08338671 | -0.000516 |
| 1.01789975 | 1.71304035 | 1.28082275 | 2.14151955 | 1.45160103 | 1.52097669 | -0.0040844 |
| 3.22728515 | 4.42718363 | 3.00272179 | 3.65480375 | 4.54281759 | 3.77096238 | -0.0010707 |
| -0.7994261 | -0.5875645 | -0.6833963 | -0.5253496 | -0.7676063 | -0.6726686 | -0.0012262 |
| 1.48996735 | 1.68590534 | 1.52663088 | 1.33842182 | 1.65676367 | 1.53953781 | -0.0011472 |
| 1.18885517 | 4.36031055 | 0.21996117 | 2.20251417 | 3.48465204 | 2.29125862 | -0.000751 |
| 0.85612035 | 0.8899188 | 1.0927074 | 1.15200377 | 0.90305257 | 0.97876058 | -0.0014391 |
| 0.529423 | 4.05411816 | -1.534724 | -0.2608752 | -0.7558831 | 0.40641177 | -0.0003965 |
| 0.72725511 | 0.73525739 | 0.36570907 | 0.56808496 | 0.65605712 | 0.61047273 | -0.0009909 |
| -1.2338142 | -2.439014 | -2.0427604 | -1.2306035 | -0.7765794 | -1.5445543 | -0.0009506 |
| -2.6894054 | -2.4985368 | -1.6993548 | -3.2727942 | -3.280983 | -2.6882149 | -0.0011463 |
| 0.67278898 | 0.4911623 | 0.78376603 | 0.75866365 | 0.69122076 | 0.67952034 | -0.0040337 |
| -0.627041 | 0.15330386 | 1.03029394 | 0.2426002 | 0.09515667 | 0.17886274 | -0.0007266 |
| 6.4389019 | 7.44024658 | 6.77490425 | 5.45729017 | 5.9052248 | 6.40331354 | -0.0011184 |
| 1.32660341 | 1.80703259 | 1.98965645 | 4.4807744 | 2.66769981 | 2.45435333 | -0.0006488 |
| 3.38598824 | 3.13476801 | 1.06001067 | 2.76618767 | 2.12285018 | 2.49396095 | -0.0006488 |
| -1.3671832 | -1.2886376 | -0.9656925 | -1.1589415 | -1.1987176 | -1.1958345 | -0.0009678 |
| 0.59953737 | 0.66047692 | 0.64358902 | 0.72212911 | 0.89828873 | 0.70480423 | -0.0008232 |
| -0.6044521 | -0.0146375 | -0.3442307 | -0.5559182 | -0.2512689 | -0.3541015 | -0.0009253 |
| 0.76285791 | 0.87899542 | 0.86501837 | 0.73813868 | 1.06216359 | 0.86143479 | -0.002469 |
| -1.0006092 | -1.3522899 | -0.988107 | -1.0339258 | -1.1739757 | -1.1097815 | -0.0013135 |
| -0.8340769 | -0.8109727 | -0.8359275 | -0.9157913 | -0.9129241 | -0.8619385 | -0.0016497 |
| 0.50110102 | 0.63197136 | 0.42372799 | 0.74717832 | 0.80053043 | 0.62090182 | -0.0003766 |
| 0.50569296 | 0.11112833 | 0.09293389 | 0.02736521 | -0.0277648 | 0.14187112 | -0.0026664 |
| -0.8056681 | -0.9834303 | -0.3329794 | -0.4421737 | -0.302784 | -0.5734071 | -0.0081358 |
| 0.09048414 | 0.68482137 | -0.1805677 | 0.70222902 | -0.5159631 | 0.15620074 | -0.002999 |
| 1.71666074 | 3.57674026 | 1.91959739 | 2.30584192 | 2.96069455 | 2.49590697 | -0.0054477 |
| 5.39461374 | 5.36171579 | 4.47234583 | 5.00729799 | 4.80278444 | 5.00775156 | -0.0018404 |
| 0.22010803 | 0.78480577 | 0.39267397 | 0.39022636 | 0.44599056 | 0.44676094 | -0.0011486 |
| -0.2142618 | 0.16490316 | -0.0283854 | 0.55132067 | 0.09997892 | 0.11471112 | -0.0011486 |
| 1.08943129 | 2.62727809 | 1.80525064 | 1.79502249 | 1.63884079 | 1.79116466 | -0.0005903 |
| 1.20137393 | 2.25117016 | 2.06275845 | 1.38036072 | 2.37746954 | 1.85462656 | -0.0005903 |
| -0.5244246 | 1.23533356 | -0.2040954 | 0.2682395 | -0.177994 | 0.11941183 | -0.0005903 |
| 7.67148066 | 5.31209993 | 6.42133045 | 7.97780561 | 5.98752785 | 6.6740489 | -0.0019529 |
| 1.07938719 | 2.05297279 | 1.78304338 | 1.13383627 | 0.42472291 | 1.29479251 | -0.0036162 |
| 0.23719454 | 0.12325406 | -0.0065023 | 0.31098557 | 0.087345 | 0.15045538 | -0.0039595 |
| 0.35409975 | 0.611269 | 0.55596101 | 0.39414179 | 0.62910533 | 0.50891538 | -0.0038648 |
| -0.2528377 | 0.22211647 | -0.0573001 | -1.9184434 | -1.1932125 | -0.6399354 | -0.0026118 |
| 2.65261173 | 3.66956615 | 3.18789864 | 1.80821383 | 1.95971322 | 2.65560071 | -0.0136611 |
| 0.20217872 | 0.32735062 | 1.86210227 | 0.64052105 | 0.61847329 | 0.73012519 | -0.0046456 |
| -1.3539286 | 0.8715651 | 0.36770844 | 2.50024223 | 2.32303429 | 0.9417243 | -0.0054044 |
| 0.52225447 | 0.59026814 | 0.31486416 | 0.8006568 | 0.76229286 | 0.59806728 | -0.0014478 |
| 0.4196732 | 0.55931544 | 0.26793456 | 0.53832364 | 0.56075144 | 0.46919966 | -0.0014478 |
| -0.065542 | -0.2128682 | -0.0199914 | 0.15077901 | 0.02310109 | -0.0249043 | -0.0014478 |
| -0.3540559 | -0.2550025 | -0.0159903 | 0.03077507 | 0.04993081 | -0.1088686 | -0.0014478 |

| | | | | | | |
|---|---|---|---|---|---|---|
| 0.28878617 | 0.51055527 | 0.24476576 | 0.32597184 | 0.47434044 | 0.3688839 | -0.0014478 |
| 1.40186 | 0.78647804 | 1.15778136 | 0.88326955 | 0.93245065 | 1.03236792 | -0.0093348 |
| 0.77595401 | 1.02892399 | 0.76708364 | 0.86354899 | 0.685395 | 0.82418113 | -0.0004134 |
| -1.3130677 | -1.8502269 | -1.2958851 | -2.3485966 | -2.9826992 | -1.9580951 | -0.0004134 |
| 1.01589525 | 1.14353621 | 1.26682425 | 1.28876996 | 1.3863399 | 1.22027311 | -0.0004134 |
| 0.2520566 | 0.23231578 | 0.52095509 | 0.73479187 | 0.49306154 | 0.44663618 | -0.0004134 |
| 2.16841745 | 1.88664877 | 1.78057826 | 2.1999526 | 2.32885456 | 2.07289033 | -0.0004134 |
| 0.6585567 | 1.26714718 | 1.31874263 | 0.69561815 | 0.38882351 | 0.86577764 | -0.0004134 |
| 1.63560724 | 1.75730646 | 1.32255363 | 1.73977733 | 1.66247058 | 1.62354305 | -0.0004134 |
| 1.05428004 | 1.38620305 | 1.16105604 | 1.37502599 | 0.90795135 | 1.1769033 | -0.0004134 |
| 0.04764509 | -0.0348828 | -0.0345399 | -0.2054527 | -0.532182 | -0.1518825 | -0.0014487 |
| -0.0181606 | 0.08399296 | -0.3691235 | -0.0576425 | 0.24543738 | -0.0230992 | -0.0091347 |
| -0.7842042 | -1.0365505 | -1.2667947 | -0.9400265 | -1.0791023 | -1.0213356 | -0.0071124 |
| 0.27056956 | 0.18709278 | 0.03839445 | -0.0105186 | -0.2346559 | 0.05017648 | -0.0071124 |
| -2.4813242 | -2.457072 | -1.9377544 | -1.0088375 | -1.5836725 | -1.8937321 | -0.0071124 |
| 0.82044554 | 1.39543796 | 1.63541389 | 1.83444786 | 2.51025867 | 1.63920078 | -0.0071124 |
| 0.90551949 | 1.34488606 | 0.91903877 | 4.06144333 | 0.14409781 | 1.47499709 | -0.0038035 |
| 0.68206644 | 0.70154881 | 0.8650167 | 1.31445265 | 0.95915151 | 0.90444722 | -0.0061726 |
| 3.8816545 | 3.73370647 | 3.25092316 | 4.07692385 | 4.49962902 | 3.8885674 | -0.0087184 |
| 0.65856361 | 0.31172776 | 0.48920822 | 0.46283197 | 0.44641352 | 0.47374902 | -0.0015533 |
| -1.0917159 | -1.2519869 | -0.8885765 | -0.949497 | -1.0766149 | -1.0516782 | -0.0015533 |
| 0.11685538 | 0.19183636 | -0.350338 | 0.33466697 | 0.99899101 | 0.25840235 | -0.0015533 |
| 1.57954359 | 1.15374446 | 1.26233959 | 1.16952801 | 1.25397944 | 1.28382702 | -0.0016978 |
| 0.57743168 | 1.14473462 | 1.09487212 | 0.19272089 | 0.78641129 | 0.75923412 | -0.0010222 |
| -4.1560755 | -3.359271 | -4.2062244 | -4.1722894 | -3.7502213 | -3.9288163 | -0.0071953 |
| -0.0680718 | 0.73844022 | -0.1093962 | 0.89282846 | 0.10275626 | 0.31131138 | -0.0071953 |
| 0.52706051 | 0.7702651 | -0.0350168 | -0.5808519 | -0.1915963 | 0.09797213 | -0.0071953 |
| 3.10086918 | 2.83512592 | 2.06727314 | 2.38681507 | 3.21325874 | 2.72066841 | -0.0030304 |
| 0.54576564 | 1.75077796 | 1.11393046 | 0.0902617 | 0.35125065 | 0.77039728 | -0.0018089 |
| 0.66439652 | -0.0486209 | -0.3066185 | -0.380625 | -0.2962689 | -0.0735474 | -0.0006721 |
| 1.59884477 | 0.74986696 | 0.57871699 | 1.18847752 | -0.0260427 | 0.81797271 | -0.0112793 |
| 0.16337943 | 0.09371471 | 0.06724787 | 0.1797657 | -0.0198376 | 0.09685402 | -0.0029864 |
| -1.7640281 | -1.4339552 | -1.4205236 | -1.4838762 | -1.6944919 | -1.559375 | -0.0029864 |
| 3.50636196 | 4.11248016 | 4.42909956 | 4.63798523 | 3.40250373 | 4.01768613 | -0.0012028 |
| 0.17830443 | -0.0919247 | -0.9831192 | -0.4473596 | 0.61399364 | -0.1460211 | -0.0012028 |
| -1.1218321 | -1.2446322 | -0.7773812 | 0.51319337 | -1.3501885 | -0.7961681 | -0.0012028 |
| -0.7470825 | -0.7121241 | -0.8575499 | -0.6811836 | -0.8745146 | -0.7744909 | -0.0005843 |
| 0.74028969 | 0.61574936 | 0.39130497 | 0.27183151 | 0.05170536 | 0.41417618 | -0.0005843 |
| 0.88109303 | 0.32026792 | 0.91959167 | 0.38718367 | 0.61081934 | 0.62379112 | -0.0005843 |
| 1.60094023 | 0.43086761 | 0.58212048 | 0.85669041 | 1.3780365 | 0.96973104 | -0.006331 |
| 3.93077183 | 3.90070033 | 3.89893937 | 4.6050806 | 4.741364 | 4.21537123 | -0.006331 |
| 0.36340857 | 0.09166408 | 0.33689594 | 0.22286463 | 0.00484419 | 0.20393548 | -0.0025842 |
| -0.6907377 | -0.0215735 | -1.904749 | -0.6468232 | -0.840825 | -0.8209417 | -0.0033755 |
| 0.08733892 | -0.4996095 | -0.2683742 | -0.2740841 | -0.6118286 | -0.3133115 | -0.0033755 |
| -0.2368705 | -0.4339588 | -0.3211751 | -0.2980869 | -0.4576187 | -0.349542 | -0.0033755 |

| | | | | | | |
|---|---|---|---|---|---|---|
| 0.39759851 | -0.1241463 | 0.26529312 | -0.1369333 | 0.01449251 | 0.08326089 | -0.0033755 |
| 0.13519382 | 0.49617803 | 2.3310194 | -0.9728434 | -0.4971097 | 0.29848764 | -0.0052328 |
| -1.6625812 | -0.8471382 | -0.3508835 | -1.2455568 | -1.7810838 | -1.1774487 | -0.0052328 |
| 5.74395227 | 5.21036339 | 4.94365978 | 3.13840032 | 5.92938423 | 4.993152 | -0.0006241 |
| 0.32472539 | 0.69857669 | -0.0189571 | 0.12163162 | 0.9729923 | 0.41979377 | -0.0017446 |
| -0.8732467 | 0.30350733 | -3.2060719 | 1.3039422 | 1.84074783 | -0.1262242 | -0.0012575 |
| 0.03487682 | -0.3871555 | -0.1977091 | -0.6846609 | -0.2962155 | -0.3061728 | -0.0026043 |
| 0.42090344 | 0.63789415 | 0.32069397 | 0.92201185 | 0.43384218 | 0.54706912 | -0.0017306 |
| 0.49147797 | -3.2231159 | 1.68978357 | 1.3772198 | -2.2869048 | -0.3903079 | -0.0017306 |
| 0.35108328 | 1.33736229 | 0.78626764 | 0.70340049 | 1.07676768 | 0.85097628 | -0.0043125 |
| 0.06079674 | -0.0084026 | 0.43004417 | 0.08859324 | 0.40684605 | 0.19557552 | -0.0161528 |
| 0.39499331 | -0.1649547 | 0.60347319 | 0.26924801 | 0.16086912 | 0.25272579 | -0.0020644 |
| 3.29008484 | 3.1547184 | 1.7941699 | 2.28021932 | 3.92853522 | 2.88954554 | -0.0049317 |
| 1.19991064 | 3.57732677 | 1.69529533 | 1.29266453 | 4.43656349 | 2.44035215 | -0.0032143 |
| -0.0489352 | -2.0959296 | -0.341215 | -0.3465295 | -0.3519728 | -0.6369164 | -0.0032143 |
| 0.53484988 | -0.2235715 | 0.83414185 | 0.20167661 | 0.40582108 | 0.35058358 | -0.0032143 |
| 0.86654878 | 0.56328368 | 0.71844196 | 1.14051771 | 1.01347804 | 0.86045403 | -0.0032143 |
| -1.1447184 | -0.4585567 | -1.1357307 | -0.3283606 | -0.8535585 | -0.784185 | -0.0032143 |
| -0.1461668 | -0.1533499 | -0.311341 | -0.5600221 | -0.2010694 | -0.2743898 | -0.0032143 |
| -0.5012941 | -0.6296273 | -0.7304494 | -1.438217 | -1.268638 | -0.9136452 | -0.0032143 |
| 0.67089915 | 0.95682311 | 0.74121594 | 0.93882322 | 0.14809632 | 0.69117155 | -0.0032143 |
| 0.31634498 | 0.16231847 | -0.245563 | 0.30243373 | 0.09378982 | 0.12586479 | -0.0032143 |
| -1.369463 | -1.6372542 | -1.3137136 | -1.6249065 | -1.6934154 | -1.5277505 | -0.0032143 |
| -0.1514857 | -0.3801515 | -0.3291721 | -0.7748048 | -0.3896744 | -0.4050577 | -0.0032143 |
| 0.33854055 | -0.6144588 | -1.5117511 | 0.38082886 | 0.12063217 | -0.2572417 | -0.0032143 |
| -0.514941 | 0.70142603 | -0.4851532 | -0.6984811 | -0.7435513 | -0.3481401 | -0.0032143 |
| 0.77211547 | -0.1085086 | -0.3330126 | 0.77499205 | -0.2671552 | 0.16768624 | -0.0032143 |
| 5.08547783 | 1.12597156 | 1.82049716 | 4.96652412 | -0.4965234 | 2.50038946 | -0.0032143 |
| 0.29508829 | -2.514884 | 2.10701132 | -2.8223572 | -3.2788925 | -1.2428068 | -0.0032143 |
| 1.91584063 | -1.0588665 | 0.54470158 | -2.5066895 | 1.89697027 | 0.15839128 | -0.0032143 |
| -1.2759798 | -1.8678408 | 0.69501185 | -1.6672639 | -2.4390099 | -1.3110165 | -0.0032143 |
| -0.170217 | -1.7951016 | 2.16853809 | -0.3620172 | -1.1596873 | -0.263697 | -0.0032143 |
| -0.4367757 | 0.34990859 | -0.1256881 | 1.57649851 | 1.63559771 | 0.59990821 | -0.0032143 |
| 1.07684112 | 2.96887016 | 1.13651216 | 0.82004929 | 4.64687061 | 2.12982867 | -0.0032143 |
| -0.9685504 | -2.1554224 | -1.3077879 | -1.7728724 | -1.5417013 | -1.5492669 | -0.0032143 |
| -0.4741912 | -1.3506355 | -0.2506483 | 1.53430223 | -1.4626924 | -0.400773 | -0.0032143 |
| 0.48103672 | -1.1905229 | -1.0635495 | -2.5882583 | -0.3427501 | -0.9408088 | -0.0032143 |
| -2.629354 | -0.4770937 | -1.9155264 | -0.1303868 | -1.5284572 | -1.3361636 | -0.0032143 |
| 2.54415584 | -3.4568086 | -1.5035818 | 0.65809679 | 1.0602417 | -0.1395792 | -0.0032143 |
| -0.238821 | 0.84874916 | 0.98821831 | -1.2807837 | -2.0986712 | -0.3562617 | -0.0032143 |
| -5.6570544 | -2.3874512 | -5.2179689 | -3.8718843 | -2.8202918 | -3.9909301 | -0.0032143 |
| 0.35506582 | -0.419713 | -1.438271 | -3.3881149 | -3.1026263 | -1.5987319 | -0.0032143 |
| -0.5663929 | -2.4119635 | -2.6924381 | 1.0927999 | 0.54236364 | -0.8071262 | -0.0032143 |
| -2.16816 | -2.614532 | -0.710682 | 0.11337948 | -0.6062242 | -1.1972437 | -0.0032143 |
| -0.0783542 | -4.2098122 | 2.206604 | 1.25167704 | 0.92931747 | 0.01988642 | -0.0032143 |

| | | | | | | |
|---|---|---|---|---|---|---|
| -0.4401648 | -4.0035243 | -0.3649096 | -0.1491337 | -0.1228325 | -1.016113 | -0.0032143 |
| 0.33873773 | -1.568758 | 0.26694632 | 0.63151407 | -0.6579235 | -0.1978967 | -0.0032143 |
| -0.1942332 | -1.264372 | -0.5785748 | -0.1648116 | -0.6501987 | -0.5704381 | -0.0032143 |
| -0.2927742 | -5.2585888 | -0.5445824 | 0.45255184 | -2.2726669 | -1.5832121 | -0.0032143 |
| -0.5439572 | 1.45013726 | 0.08185387 | -0.1002345 | 0.75735235 | 0.32903035 | -0.0032143 |
| 3.48213363 | 1.71288228 | 2.22027659 | 3.38828945 | 1.24641991 | 2.41000037 | -0.0032143 |
| -1.0410061 | -2.1484914 | -0.7199202 | -1.6765513 | -2.6579816 | -1.6487901 | -0.0032143 |
| -1.3254777 | -1.6259733 | 1.11507821 | -0.6046019 | -0.6792498 | -0.6240449 | -0.0032143 |
| -2.5058486 | -1.5769721 | 3.18182802 | -0.035593 | 0.09727073 | -0.167863 | -0.0032143 |
| -0.0109649 | -4.1670203 | -0.1349835 | 3.03371859 | -3.8804293 | -1.0319359 | -0.0032143 |
| 1.27088833 | 1.26668024 | 2.16811752 | 1.09208524 | 2.68869638 | 1.69729354 | -0.0032143 |
| -3.4172306 | -6.0350866 | -4.856122 | -3.3039842 | -4.9519057 | -4.5128658 | -0.0032143 |
| 1.48509789 | 0.650805 | -0.1915472 | 3.8160224 | 2.02208853 | 1.55649333 | -0.0032143 |
| -4.7163839 | -5.6667724 | -5.9073472 | -3.8924112 | -5.3553972 | -5.1076624 | -0.0032143 |
| -3.5567303 | -4.8911901 | -5.5273275 | -3.4027152 | -4.8956609 | -4.4547248 | -0.0032143 |
| 4.05034256 | 1.59584141 | 2.18635845 | 3.55441666 | 1.9561162 | 2.66861506 | -0.0032143 |
| -1.7736897 | -2.0467033 | -2.2226157 | -3.2144601 | -2.3674345 | -2.3249807 | -0.0032143 |
| -0.3996555 | -3.6534805 | -4.0770054 | -3.0916922 | -4.6414566 | -3.172658 | -0.0032143 |
| 1.21198988 | 1.04590225 | 2.22313023 | 1.89740002 | -0.794014 | 1.11688168 | -0.0032143 |
| 3.02455664 | 3.51012421 | 1.66797256 | 3.79843998 | 1.97768497 | 2.79575567 | -0.0032143 |
| -3.7046919 | -6.9219356 | -1.2565584 | -4.0682273 | -3.9444699 | -3.9791766 | -0.0032143 |
| -1.7446083 | -7.0653839 | -0.0289578 | 1.62141275 | -5.7598906 | -2.5954856 | -0.0032143 |
| -1.8256378 | -0.762228 | 1.7010107 | 1.38432789 | -1.3225465 | -0.1650147 | -0.0032143 |
| -0.1509309 | -3.1386223 | 0.1355896 | 2.22496367 | 1.39524174 | 0.09324837 | -0.0032143 |
| -7.4734592 | -6.1075583 | -1.8972311 | -6.5426083 | -7.8286381 | -5.969899 | -0.0032143 |
| 4.09803009 | 4.77969074 | 0.51887512 | 4.20026684 | 3.5945034 | 3.43827324 | -0.0032143 |
| -4.8677073 | -4.0758991 | -1.9812058 | -2.5783153 | -6.8333406 | -4.0672936 | -0.0032143 |
| -4.5812373 | -4.737258 | -0.5851467 | -4.0342708 | -4.5289454 | -3.6933716 | -0.0032143 |
| 2.96320009 | 3.79111004 | 1.42783403 | 1.98503757 | 2.8780241 | 2.60904117 | -0.0032143 |
| 0.51036596 | -0.3663378 | -0.0590553 | 0.28712177 | 0.12234497 | 0.09888792 | -0.0032143 |
| -4.2047248 | -5.2729969 | -4.3213353 | -3.8531871 | -4.6397791 | -4.4584046 | -0.0032143 |
| 2.69342637 | 0.02032518 | 0.65585351 | -1.2155851 | -1.7489452 | 0.08101494 | -0.0032143 |
| -4.1723242 | -4.4181976 | -6.8955078 | -1.9910703 | -4.9806519 | -4.4915504 | -0.0032143 |
| -3.0463967 | -3.5717645 | -1.5468769 | -2.2286711 | -4.3037858 | -2.939499 | -0.0032143 |
| -3.3972235 | -4.4117646 | -2.2848678 | -4.8480721 | -3.2752624 | -3.6434381 | -0.0032143 |
| -4.7198234 | -6.4583836 | -6.9089861 | -6.680398 | -5.9183192 | -6.137182 | -0.0032143 |
| 3.54829621 | -0.2857246 | 0.23714042 | 1.35729361 | 2.50370789 | 1.4721427 | -0.0032143 |
| -1.1144891 | -4.3184967 | -7.6537132 | -2.4699645 | -8.3321657 | -4.7777658 | -0.0032143 |
| -4.765202 | -7.076869 | -5.188375 | -4.2222862 | -5.4148006 | -5.3335066 | -0.0032143 |
| -2.0807338 | -2.319715 | -5.4017029 | -1.594687 | -5.3842368 | -3.3562151 | -0.0032143 |
| -3.1362762 | -3.9690053 | -6.2389841 | -6.1778827 | -6.0969234 | -5.1238143 | -0.0032143 |
| -2.1125889 | -4.656827 | -1.6655492 | -1.0017753 | -4.5467529 | -2.7966986 | -0.0032143 |
| -6.3392601 | -6.1401391 | -4.0705986 | -4.8990192 | -5.4489508 | -5.3795936 | -0.0032143 |
| -3.7572458 | -6.50844 | 2.78789544 | -4.8292637 | -4.2237654 | -3.3061639 | -0.0032143 |
| -1.7484189 | -2.3341141 | -1.9233091 | -4.2654896 | -3.2248473 | -2.6992358 | -0.0032143 |

| | | | | | | |
|---|---|---|---|---|---|---|
| -2.7592359 | -3.6245499 | -3.0141659 | 0.54600096 | -2.7404823 | -2.3184866 | -0.0032143 |
| -1.9939947 | -1.9888431 | -2.3650703 | -1.9998264 | -1.566329 | -1.9828127 | -0.0032143 |
| -2.065908 | -3.8453894 | -4.5158215 | -2.3075037 | -2.8653755 | -3.1199996 | -0.0032143 |
| -3.9425721 | -1.7991381 | -1.9728035 | -3.2377324 | -3.3437271 | -2.8591946 | -0.0032143 |
| -4.0063934 | 0.00170612 | -3.2460051 | -1.6325617 | 0.03355503 | -1.7699398 | -0.0032143 |
| -3.8721948 | -6.19416 | -3.4666057 | -5.6864705 | -3.1011167 | -4.4641095 | -0.0032143 |
| 0.32579994 | 0.83396918 | 2.02885389 | 1.47016668 | 3.29581881 | 1.5909217 | -0.0032143 |
| -1.963541 | 0.08338159 | -1.6714234 | -3.7653079 | -0.8892334 | -1.6412248 | -0.0032143 |
| -1.2872543 | -1.6471449 | -1.2363881 | 0.57388449 | -3.9352846 | -1.5064375 | -0.0032143 |
| 2.00807047 | -0.8842239 | -2.843049 | -2.1991882 | -0.0263157 | -0.7889413 | -0.0032143 |
| 1.08819008 | 3.05274153 | 2.45442486 | 1.36510706 | 1.96820807 | 1.98573432 | -0.0032143 |
| 2.3411572 | 0.37756777 | 1.32199264 | 4.22563744 | 2.72458887 | 2.19818878 | -0.0032143 |
| -1.4949466 | -2.7904267 | -2.344173 | -2.0503788 | -1.3912106 | -2.0142271 | -0.0032143 |
| -4.04492 | -0.2212386 | -4.981873 | -3.0949001 | -1.8103261 | -2.8306516 | -0.0032143 |
| -3.2334676 | -4.1263552 | -3.1550074 | -3.3659058 | -1.9604335 | -3.1682339 | -0.0032143 |
| -1.7893363 | -6.9306107 | -1.2292366 | -4.0150585 | -5.6924295 | -3.9313343 | -0.0032143 |
| 3.72828317 | -4.0463743 | 4.08195877 | 0.23325872 | -3.8436408 | 0.03069711 | -0.0032143 |
| 1.92436647 | -0.2527266 | -0.6690402 | -1.2587919 | -2.4866285 | -0.5485641 | -0.0032143 |
| -0.655304 | -1.0041326 | -1.7057736 | -0.4397254 | -0.5029328 | -0.8615737 | -0.0032143 |
| 1.49420404 | 1.24753594 | 1.35374045 | 1.70859361 | 0.92053056 | 1.34492092 | -0.0032143 |
| 0.19671273 | 0.14006376 | -0.2026069 | -0.2058284 | -0.8524792 | -0.1848276 | -0.0032143 |
| -0.7233374 | -0.4885645 | -0.3862134 | -0.6090652 | -1.0527482 | -0.6519857 | -0.0032143 |
| -0.4693797 | -0.5608063 | -1.0753961 | -0.5599451 | -0.4757271 | -0.6282508 | -0.0032143 |
| 0.28917718 | 0.05188513 | 0.33114934 | 0.66780519 | 0.7844882 | 0.42490101 | -0.0032143 |
| 0.37516046 | 0.22387218 | 0.09231496 | -0.1344476 | 0.07079268 | 0.12553854 | -0.0032143 |
| -0.143646 | -0.1270516 | -0.5211225 | -0.1458294 | -0.0479374 | -0.1971174 | -0.0032143 |
| -0.6767387 | -0.1937788 | -0.8354435 | -0.3013611 | -0.4498894 | -0.4914423 | -0.0032143 |
| -4.0484757 | -4.238657 | -5.0171471 | -4.5547285 | -4.1792502 | -4.4076517 | -0.0064392 |
| 1.40275764 | 0.54679108 | -0.0180583 | 0.86078262 | 0.61875725 | 0.68220606 | -0.000596 |
| 2.20048261 | 2.07041526 | 2.49902678 | 2.04824018 | 0.68429279 | 1.90049152 | -0.000596 |
| 1.95547152 | 1.5470202 | 1.58011687 | 2.37824082 | 1.87101364 | 1.86637261 | -0.000596 |
| 0.19906783 | 0.28600121 | 0.08012295 | -0.0370059 | 0.22027898 | 0.14969301 | -0.000596 |
| -0.4163506 | -0.3763396 | -0.1908114 | -0.1551304 | -0.1313725 | -0.2540009 | -0.000596 |
| -0.7462301 | -0.6627133 | -1.6628277 | 0.47462797 | -1.7689042 | -0.8732095 | -0.0016112 |
| 6.83099699 | 5.74687576 | 6.22348404 | 5.54295921 | 7.07930326 | 6.28472385 | -0.00238 |
| -4.1375189 | -3.083858 | -3.922647 | -4.592227 | -3.9561629 | -3.9384828 | -0.0037171 |
| 0.72399712 | 1.11655045 | -0.1052279 | 0.18993235 | 0.86757565 | 0.55856552 | -0.0026509 |
| 0.1072917 | 2.14441252 | 0.64750218 | 0.99689007 | 0.27037859 | 0.83329501 | -0.0026509 |
| 2.05991602 | 1.33294868 | 0.86487985 | 1.34838939 | 1.4954958 | 1.42032595 | -0.008202 |
| 1.63399601 | 1.60942078 | 2.26329374 | 2.65821695 | 4.34717083 | 2.50241966 | -0.008202 |
| 2.20757484 | 2.25289726 | 1.67792141 | 1.89772725 | 1.78457689 | 1.96413953 | -0.0019503 |
| 1.57520795 | 1.3458035 | 1.05882931 | 1.29764986 | 1.39897275 | 1.33529267 | -0.0041843 |
| 2.04983211 | 2.45508194 | 2.70488882 | 4.90646744 | 1.16701066 | 2.65665619 | -0.0041843 |
| 3.54512978 | 4.47578526 | 4.58527279 | 3.63886929 | 4.6902523 | 4.18706188 | -0.0060546 |
| 0.07898498 | -0.0372112 | -0.8119307 | -0.4428198 | -1.1122274 | -0.4650408 | -0.0060546 |

| | | | | | | |
|---|---|---|---|---|---|---|
| 1.85418963 | 0.28363609 | 0.41692448 | 1.88636625 | 1.71489418 | 1.23120213 | -0.0060546 |
| 1.52705622 | 3.16540623 | 1.19575286 | 2.3669095 | 2.8587904 | 2.22278304 | -0.0037908 |
| -0.6075642 | -0.6134269 | -0.6607907 | -0.7932091 | -0.5192943 | -0.638857 | -0.0013034 |
| 0.77820635 | 0.47493458 | 0.43873262 | 0.42129946 | 0.84350348 | 0.5913353 | -0.0032149 |
| -0.1231334 | -0.1422381 | -0.6950421 | -0.1120179 | 0.06656337 | -0.2011736 | -0.0055917 |
| -2.1060252 | -2.1384175 | -1.5701597 | -1.4872503 | -2.08359 | -1.8770885 | -0.0036205 |
| 0.42782593 | -0.1559319 | 0.43310523 | 1.17633176 | 1.39947224 | 0.65616064 | -0.0036205 |
| 0.87364388 | 0.19167471 | -0.0760739 | 0.64515734 | 0.99446535 | 0.52577348 | -0.0036205 |
| 3.25356722 | 3.22127676 | 3.49844003 | 2.88875771 | 2.81255865 | 3.13492007 | -0.0036205 |
| 1.71512592 | 1.6513207 | 2.04517651 | 2.01492834 | 1.78971434 | 1.84325316 | -0.0097025 |
| 0.99632645 | 1.37361217 | 1.59618199 | 0.99547887 | 1.33344269 | 1.25900843 | -0.0017387 |
| 1.21168637 | 0.53324103 | 0.72260642 | 1.17099762 | 1.22115135 | 0.97193656 | -0.0017387 |
| 1.30634689 | 1.99905348 | 1.26585758 | 2.31288576 | 2.63855028 | 1.9045388 | -0.0316486 |
| -2.0621901 | 0.33685303 | -0.5821528 | -1.1367397 | -1.1184058 | -0.9125271 | -0.0008513 |
| -1.5967226 | 0.19969296 | -0.8290381 | -2.5418553 | 0.704211 | -0.8127424 | -0.007704 |
| -0.4292767 | -0.2321625 | -0.8070633 | -0.9151499 | -0.5413404 | -0.5849986 | -0.0016724 |
| 3.98424745 | 3.8474412 | 3.0413003 | 2.11231899 | 2.93179202 | 3.18341999 | -0.0016724 |
| 3.61307955 | 3.21519279 | 3.1621511 | 1.55044889 | 1.67338324 | 2.64285111 | -0.0024758 |
| 5.77606583 | 5.96096611 | 4.7753787 | 4.83242893 | 5.52302647 | 5.37357321 | -0.0003419 |
| 2.22665024 | -0.5048637 | 2.95782995 | 2.54039526 | 0.36728287 | 1.51745893 | -0.0024019 |
| 0.26582932 | 1.12116623 | 0.20682287 | 1.32213044 | 1.03215432 | 0.78962064 | -0.0024019 |
| -0.5995667 | -0.8129997 | -0.6686583 | -0.5026584 | 0.07980442 | -0.5008157 | -0.0015089 |
| 1.32922554 | 2.0333271 | 2.07256031 | 1.39344645 | 0.41092825 | 1.44789753 | -0.0030737 |
| 2.31833553 | 2.42124867 | 2.31110311 | 2.50049996 | 2.25760436 | 2.36175833 | -0.0056595 |
| 2.40117669 | 2.90627098 | 2.78895712 | 3.33476782 | 2.80927086 | 2.84808869 | -0.0056595 |
| 1.02710485 | 1.48256612 | 0.96671212 | 1.60736966 | 1.35700369 | 1.28815129 | -0.0056595 |
| -0.8125396 | -0.9508367 | -0.5882182 | -1.0480788 | -1.3001454 | -0.9399637 | -0.0056595 |
| 0.12697792 | 1.42532504 | -0.1460731 | 0.31757498 | 0.56312275 | 0.45738552 | -0.0056595 |
| 0.4063158 | 0.37084484 | 0.32418251 | 0.30779791 | 0.28337383 | 0.33850298 | -0.0026996 |
| -0.2308545 | 3.64018297 | -2.1270604 | -0.0696275 | 3.09756422 | 0.86204095 | -0.0020442 |
| -0.2049539 | -0.1618481 | 0.01315951 | -0.7101436 | -0.0175173 | -0.2162607 | -0.0006109 |
| -0.011971 | 0.0057044 | -0.043119 | -0.0721095 | 0.08477068 | -0.0073449 | -0.0018089 |
| -1.2213206 | -1.361328 | -0.9138765 | -1.3102062 | -1.0645847 | -1.1742632 | -0.0047174 |
| -1.260669 | -0.9052062 | -1.6426482 | -0.9212084 | -1.2635643 | -1.1986592 | -0.0003119 |
| 2.24746346 | 2.47052956 | 2.10166979 | 2.15952492 | 2.49149466 | 2.29413648 | -0.0003119 |
| -0.2832611 | -0.5489273 | -0.4013157 | -0.2973263 | -0.5221183 | -0.4105897 | -0.0059831 |
| 2.14819384 | 4.52700996 | 4.18840694 | 2.80818748 | 2.79446793 | 3.29325323 | -0.00431 |
| -0.4584715 | -0.2277615 | -0.3859208 | -0.3203638 | -0.2786789 | -0.3342393 | -0.0014861 |
| -0.6499476 | -0.6209579 | -0.4999604 | -0.6352894 | -0.4447246 | -0.570176 | -0.0014861 |
| -2.4701958 | -1.8366849 | -1.7577894 | -1.2714968 | -1.8787237 | -1.8429781 | -0.0014208 |
| 5.82145643 | 4.06643772 | 2.98211312 | 6.52794409 | 2.54060507 | 4.38771129 | -0.0020026 |
| -0.9992856 | 3.14573526 | -0.383432 | 1.47685492 | 1.1157279 | 0.87112008 | -0.0007688 |
| 1.31943154 | 1.9277668 | 1.47370672 | 0.7002629 | 1.24435091 | 1.33310378 | -0.0022914 |
| 1.03441453 | 0.6769278 | 0.25696826 | 0.53816438 | 0.35347462 | 0.57198992 | -0.0023405 |
| 1.25083017 | 1.05623126 | 1.21003413 | 0.91880703 | 1.01898932 | 1.09097838 | -0.0023405 |

| | | | | | | |
|---|---|---|---|---|---|---|
| 2.48272848 | 3.95077419 | 1.52333689 | 1.75123453 | 2.90912533 | 2.52343988 | -0.0009128 |
| -1.3316841 | -4.2885761 | -3.5894592 | -1.5038118 | -5.4143085 | -3.225568 | -0.0009128 |
| 0.92389941 | 1.11767018 | 0.62519193 | 0.86195028 | 0.86340189 | 0.87842274 | -0.0169261 |
| 1.52580726 | 2.55183411 | 0.89136481 | 0.93508935 | 1.01432967 | 1.38368504 | -0.0062107 |
| -0.0755396 | -0.3462477 | -0.2195411 | 0.01577139 | -0.1205676 | -0.1492249 | -0.0062107 |
| -0.2659404 | 0.43989015 | 1.69011045 | 1.08236647 | 0.33961678 | 0.65720868 | -0.0006696 |
| 0.66271281 | 0.61156821 | 0.76497793 | 0.59190297 | 0.9059968 | 0.70743175 | -0.0146976 |
| -0.1831474 | 1.62413776 | 0.58805561 | 0.61743557 | -0.7677164 | 0.37575302 | -0.0030007 |
| 6.00239754 | 3.73457432 | 3.41635561 | 6.96130753 | 5.24526024 | 5.07197905 | -0.0013639 |
| 0.43267751 | 1.39587545 | 0.98319602 | 0.66265774 | 0.30147243 | 0.75517583 | -0.0005451 |
| 1.58980322 | 1.24116778 | 1.03911519 | 0.62356091 | 1.05954576 | 1.11063857 | -0.0008615 |
| 2.78991413 | 3.0531497 | 3.22389412 | 3.15982676 | 3.10201693 | 3.06576033 | -0.0008615 |
| 3.70539188 | -0.7754233 | 2.32327461 | 0.36550808 | 6.07956982 | 2.33966422 | -0.0017258 |
| -1.9625399 | 0.08470035 | -0.5325437 | -1.043354 | -0.8346364 | -0.8576747 | -0.0052023 |
| 0.87006128 | 0.92782032 | 0.83533192 | 0.43460822 | 4.23425198 | 1.46041474 | -0.0052023 |
| 0.61645222 | 0.60610294 | 0.21540761 | 0.96112084 | -0.5506787 | 0.36968098 | -0.0052023 |
| -2.2846313 | -2.0523491 | -1.5943674 | -2.2116337 | -2.2786868 | -2.0843336 | -0.0052023 |
| -1.9644686 | -2.3740304 | -0.1871042 | -2.1964393 | -2.6687052 | -1.8781495 | -0.0052023 |
| -1.0989629 | 2.52306318 | 0.33705664 | 0.54281759 | 1.78629863 | 0.81805463 | -0.0052023 |
| -2.4757502 | -3.370141 | -2.9174919 | -4.1636143 | 0.57939887 | -2.4695197 | -0.0052023 |
| 1.34527171 | -0.438262 | -1.6262398 | -0.1585388 | -1.3488754 | -0.4453289 | -0.0052023 |
| -1.8069227 | -2.454253 | -0.6853456 | -0.986429 | -1.9232464 | -1.5712393 | -0.0052023 |
| 0.31836075 | 0.7655068 | -0.0368681 | 0.36410952 | -0.756651 | 0.1308916 | -0.0052023 |
| -0.3245776 | -2.6136131 | -1.2047431 | -2.4164827 | -0.3063793 | -1.3731592 | -0.0052023 |
| -3.2177947 | -4.5568624 | -2.890322 | -2.4875376 | -3.6162024 | -3.3537438 | -0.0052023 |
| 4.87722015 | 3.99092436 | 3.61658382 | 3.66205788 | 3.44998884 | 3.91935501 | -0.0052023 |
| 0.48696089 | 1.07194185 | -0.3172274 | 1.03166151 | 0.3396821 | 0.5226038 | -0.0052023 |
| -1.7291315 | -0.4629538 | -0.7426848 | -1.2110994 | -2.3613501 | -1.3014439 | -0.0052023 |
| 1.42144871 | 0.272614 | 0.77874374 | 2.33840275 | 0.19433022 | 1.00110788 | -0.0052023 |
| 5.0876689 | 6.42817879 | 4.79992008 | 3.5153265 | 5.71240425 | 5.1086997 | -0.0052023 |
| 2.75241327 | 2.65104866 | 2.7847662 | 2.6762557 | 3.58345175 | 2.88958712 | -0.0052023 |
| 0.89518094 | 1.5254873 | 0.65617585 | 2.00526333 | 1.56567287 | 1.32955606 | -0.0052023 |
| 3.91431117 | 2.94511509 | 3.61076617 | 2.48844528 | 2.62573242 | 3.11687403 | -0.0052023 |
| 3.21904659 | 2.3045392 | 1.85652971 | 1.91926718 | 3.4875679 | 2.55739012 | -0.0052023 |
| 2.46724653 | 1.7466774 | 2.94625998 | 2.89839625 | 4.72513103 | 2.95674224 | -0.0052023 |
| 1.38287592 | 0.97761714 | 3.67957997 | 1.91075826 | 0.79016972 | 1.7482002 | -0.0052023 |
| -0.5135789 | -0.6276469 | -1.4370422 | 1.03717709 | -2.3166089 | -0.77154 | -0.0052023 |
| 0.11657596 | 1.32205284 | 0.04013634 | 1.42083251 | 0.59350634 | 0.6986208 | -0.0052023 |
| 5.86614895 | 5.21453524 | 3.26302004 | 4.32457066 | 4.51119232 | 4.63589344 | -0.0052023 |
| 1.89138198 | 1.0197686 | 0.79949903 | -0.0857029 | 1.79336691 | 1.08366272 | -0.0052023 |
| 3.66000843 | 3.8220365 | 4.21572971 | 3.90694261 | 1.37506783 | 3.39595702 | -0.0052023 |
| 0.34591699 | 2.19695497 | 0.3551755 | 0.91814423 | 1.23351121 | 1.00994058 | -0.0052023 |
| -4.8899317 | -1.5849528 | 1.76949143 | -3.8039186 | -0.6274042 | -1.8273432 | -0.0052023 |
| 4.92793083 | 5.02778196 | 2.51548576 | 5.46987534 | 6.9570713 | 4.97962904 | -0.0052023 |
| 1.09424686 | 1.88560557 | 4.85102463 | 3.905581 | 0.8621428 | 2.51972017 | -0.0052023 |

| | | | | | | |
|---|---|---|---|---|---|---|
| -1.2667663 | -1.7967939 | -1.1780882 | -0.6443902 | -0.400311 | -1.0572699 | -0.0052023 |
| 3.94909024 | 4.47633362 | 5.3114953 | 5.00388098 | 4.17451954 | 4.58306394 | -0.0052023 |
| 2.30204105 | -2.4837275 | 0.84277534 | -1.1928768 | -3.6697812 | -0.8403138 | -0.0052023 |
| 1.99565828 | 1.88872242 | 2.7974453 | 2.47887564 | 2.61645842 | 2.35543201 | -0.0052023 |
| 1.1810236 | 0.48670578 | 0.72192574 | 0.11566257 | -0.6609621 | 0.36887112 | -0.0036612 |
| 4.30207539 | 5.39921284 | 4.2788744 | 6.31952572 | 3.41179705 | 4.74229708 | -0.0045135 |
| 0.77236843 | 0.69967437 | 0.64585328 | 0.5346086 | 0.06097341 | 0.54269562 | -0.0009867 |
| 0.13208342 | 0.10898447 | -0.014801 | -0.0679414 | 0.18788719 | 0.06924253 | -0.0009867 |
| -3.3588605 | -3.8502908 | -2.2543221 | -2.2225108 | -2.4383259 | -2.824862 | -0.0009867 |
| 0.83332014 | 0.79229283 | 1.39848733 | 0.98821723 | 0.44561291 | 0.89158609 | -0.0009867 |
| 1.02562344 | 1.53824306 | 1.68867278 | 1.30603182 | 1.48690212 | 1.40909464 | -0.0056763 |
| 3.17595744 | 6.38219881 | 0.33981514 | -2.2533505 | -0.7630076 | 1.37632265 | -0.0010739 |
| 0.81531024 | 0.93569565 | 1.85220885 | 1.0939374 | 1.14459419 | 1.16834927 | -0.0037536 |
| -1.0235169 | -0.8162763 | -1.1696143 | -1.7450337 | -0.8144758 | -1.1137834 | -0.0037536 |
| -0.3695064 | 0.73326159 | 0.69063377 | -0.0213437 | -0.3849459 | 0.12961988 | -0.0037536 |
| -0.2060635 | 0.44942605 | -0.0916696 | 0.26166034 | -0.0359223 | 0.07548621 | -0.0037536 |
| -2.5516052 | -2.4658623 | -2.4969268 | -2.6577377 | -2.4163198 | -2.5176904 | -0.0037536 |
| -0.9671502 | -1.0135504 | 0.29478645 | -0.7389985 | -0.1302755 | -0.5110376 | -0.0037536 |
| 1.77850711 | 1.89313459 | 0.92205989 | 1.33450627 | 0.52907681 | 1.29145694 | -0.0037536 |
| -1.9173028 | -1.468636 | -2.6072712 | -2.0248055 | -1.6035999 | -1.9243231 | -0.0037536 |
| 0.74923491 | 0.69734263 | 0.63982582 | 1.36831617 | 0.99988246 | 0.8909204 | -0.0037536 |
| 1.46132469 | 2.01117897 | 2.23668242 | 1.93835771 | 1.81326199 | 1.89216115 | -0.0033294 |
| -0.1855741 | 0.12079787 | 0.32301092 | 0.19654918 | -0.0732527 | 0.07630625 | -0.0013769 |
| 0.15565729 | 1.51301563 | 0.30546808 | -1.3318224 | -0.5365582 | 0.02115209 | -0.0012553 |
| 6.06757593 | 7.02827168 | 7.75427723 | 3.88401508 | 5.70128679 | 6.08708534 | -0.0028502 |
| -0.7212336 | -0.9354973 | -0.7022581 | 0.7159425 | -0.5765831 | -0.4439259 | -0.0003143 |
| 1.23215294 | 1.50123298 | 1.50896525 | 0.50990987 | 1.47131109 | 1.24471443 | -0.0064415 |
| 3.09595394 | 2.26246881 | 2.76214457 | 1.72804499 | 1.81154668 | 2.3320318 | -0.0064415 |
| -1.5959673 | -2.1154127 | 0.1736598 | -1.3652754 | -1.2120314 | -1.2230054 | -0.0064415 |
| 0.95587087 | 1.04413748 | 0.4532671 | 1.52230239 | 0.91436648 | 0.97798886 | -0.0064415 |
| 1.00411868 | 0.5290091 | 0.84265757 | 0.89245987 | 0.8552289 | 0.82469482 | -0.0110811 |
| 0.50200486 | 0.72196937 | 1.06387711 | 1.3173492 | 0.86456752 | 0.89395361 | -0.0005804 |
| -0.2903788 | -1.9373533 | -0.8781624 | -0.6857793 | -1.0293909 | -0.9642129 | -0.0100319 |
| 0.79431105 | 3.26107049 | 1.65841866 | 3.32332325 | 2.78667307 | 2.3647593 | -0.0031094 |
| 5.71327591 | 6.80236244 | 5.00126171 | 6.01676273 | 5.95810509 | 5.89835358 | -0.0007869 |
| -1.2868056 | -1.2064719 | -0.9130659 | -1.0587418 | -1.0233142 | -1.0976799 | -0.0004022 |
| -0.1775277 | 1.1302582 | 0.61318445 | -1.6299791 | -0.3163443 | -0.0760817 | -0.004551 |
| -1.7312374 | -2.5031216 | -0.3617582 | -0.947763 | 0.24053288 | -1.0606695 | -0.0035858 |
| -0.7575843 | -0.527041 | 0.28590834 | -0.673533 | -0.9732062 | -0.5290912 | -0.0028694 |
| 3.4925344 | 5.06655359 | -1.0011225 | 5.10941887 | 0.38423347 | 2.61032357 | -0.0028694 |
| 1.40235436 | 4.36126232 | 2.73347759 | 3.55957723 | 2.74204373 | 2.95974305 | -0.0041532 |
| -0.4876001 | -0.3256376 | 0.29393816 | -0.2582905 | 0.41303277 | -0.0729115 | -0.0009817 |
| 0.82454848 | 1.42359328 | 0.80491328 | 1.12265718 | 0.80211484 | 0.99556541 | -0.0009817 |
| -0.0433631 | -0.1283448 | 0.31024909 | -0.4203533 | -0.4323149 | -0.1428254 | -0.0009817 |
| 0.70905066 | 0.54124212 | 0.94357347 | 0.91506124 | 1.33903551 | 0.8895926 | -0.0009817 |

| | | | | | | |
|---|---|---|---|---|---|---|
| 0.41896343 | 0.48677301 | 0.08381081 | 0.11593008 | 0.60399914 | 0.34189529 | -0.0009817 |
| 4.41937876 | 4.18103409 | 5.09858608 | 7.09465122 | 4.57793617 | 5.07431726 | -0.0012569 |
| 0.35981655 | 0.5574646 | 0.10501957 | 0.50550461 | 0.41505766 | 0.3885726 | -0.0048851 |
| 1.09694886 | -0.2406969 | 0.92615652 | 1.31744409 | -0.4342225 | 0.53312602 | -0.0048851 |
| 0.18991971 | -0.4823489 | -0.8571573 | -0.250524 | -0.4246957 | -0.3649613 | -0.0012716 |
| 0.98231339 | 1.466681 | 0.93407202 | 0.74581146 | 0.81103516 | 0.98798261 | -0.0007095 |
| 0.99074292 | 0.78292382 | 0.67885351 | 1.05133688 | 1.49663377 | 1.00009818 | -0.0007095 |
| -0.6448894 | -1.1100411 | -0.9012871 | -1.0109763 | -0.8970513 | -0.912849 | -0.0007095 |
| 0.49820471 | 1.81840479 | 0.15744925 | -0.1664 | 0.76548266 | 0.61462829 | -0.003776 |
| 1.21690702 | 0.55759239 | 0.38380241 | 0.44124436 | 0.49170637 | 0.61825051 | -0.0040457 |
| 0.50741291 | 0.95428419 | 0.43811512 | 0.26851249 | 1.36359024 | 0.70638299 | -0.0023194 |
| 1.02546573 | 1.10093546 | 0.97124243 | 0.97436094 | 0.9231596 | 0.99903283 | -0.0023194 |
| -0.2635124 | -1.0212715 | -0.5183415 | -0.5589449 | -1.0107865 | -0.6745714 | -0.0072187 |
| -0.3962524 | -0.4061823 | -0.3301878 | -0.198777 | -0.1342282 | -0.2931255 | -0.000857 |
| 5.1613369 | 2.12538147 | 1.22533369 | 0.92587376 | 4.08032227 | 2.70364962 | -0.001801 |
| 1.46746683 | 0.8760643 | -0.161047 | 2.34730148 | 2.46834993 | 1.39962711 | -0.002792 |
| -0.0931799 | -0.0271425 | -0.1145985 | -0.1875558 | -0.0527844 | -0.0950522 | -0.002792 |
| 0.3447473 | 0.28196263 | -0.2751777 | 0.44314098 | 0.22599888 | 0.20413442 | -0.0143127 |
| 1.02914476 | 0.73247075 | 1.34619904 | 1.15011883 | 1.16434324 | 1.08445532 | -0.0113588 |
| 2.606318 | 1.68525124 | 1.90427971 | 2.25383806 | 1.97016788 | 2.08397098 | -0.008475 |
| -0.4752822 | -0.4333024 | -0.6913538 | -0.6229312 | -0.912559 | -0.6270857 | -0.0025678 |
| 1.48531389 | 2.08375454 | 1.70702362 | 1.75058675 | 1.62382746 | 1.73010125 | -0.0181883 |
| -0.6354325 | -0.2017488 | -0.5038054 | -0.4889307 | -0.3694017 | -0.4398638 | -0.0181883 |
| 0.22662401 | 0.74208045 | 0.54017138 | 0.17787981 | 0.49922228 | 0.43719559 | -0.0181883 |
| -1.3538461 | -1.7994478 | -1.2873821 | -1.1723886 | -1.0806491 | -1.3387427 | -0.0021181 |
| -2.3294516 | -2.946887 | -2.1132274 | -2.7130134 | -2.9163818 | -2.6037922 | -0.0046107 |
| -0.1334548 | -0.7204152 | -0.5173159 | -0.2300735 | -0.4409698 | -0.4084458 | -0.0046107 |
| -0.2130232 | -0.3542128 | 0.18713236 | -0.4849117 | 0.29965878 | -0.1130713 | -0.0046107 |
| 0.81426311 | -1.2908425 | -0.506623 | -0.4024267 | 1.39949811 | 0.00277379 | -0.002444 |
| -1.9435253 | -2.2144589 | -3.0803392 | -2.3242691 | -1.914886 | -2.2954957 | -0.008104 |
| -0.1461058 | 0.76857185 | 0.35541821 | 0.68427134 | 0.72331595 | 0.47709432 | -0.008104 |
| 0.40161514 | 1.60729027 | 2.12450409 | 0.29118347 | 0.53970146 | 0.99285889 | -0.008104 |
| -0.5484424 | 0.10615373 | -0.4004576 | -0.2473152 | -0.342077 | -0.2864277 | -0.008104 |
| -4.0869083 | -2.1341617 | -3.2113824 | -2.1081364 | -1.0803695 | -2.5241917 | -0.008104 |
| -2.5281861 | -0.8488284 | -1.7830718 | -1.5992503 | -0.1112721 | -1.3741217 | -0.008104 |
| 0.31882024 | 1.03206897 | 0.74712753 | 0.7719655 | 1.14151967 | 0.80230038 | -0.008104 |
| -0.3592412 | -0.0021632 | -0.2937641 | -0.5298736 | -0.5660927 | -0.350227 | -0.008104 |
| 0.41280389 | 0.73315668 | 0.58586168 | -0.36059 | 0.35397887 | 0.34504223 | -0.008104 |
| -0.3148484 | -0.5404019 | -0.4290576 | -0.9538922 | -0.7379866 | -0.5952374 | -0.008104 |
| -2.3270333 | -1.9038782 | -1.5330606 | -2.2831552 | -2.66677 | -2.1427794 | -0.008104 |
| -1.6185384 | -0.235486 | 0.17822242 | 0.18651366 | 0.38387609 | -0.2210824 | -0.008104 |
| -2.2399335 | -1.803899 | -2.9574921 | -2.6602097 | -1.1353805 | -2.159383 | -0.008104 |
| 2.49651766 | 2.1634531 | 1.54555368 | 2.89493227 | 2.93281841 | 2.40665503 | -0.008104 |
| 0.8176899 | 1.27594185 | 1.38077569 | 0.87597299 | 1.53496408 | 1.1770689 | -0.008104 |
| 0.49464893 | 0.43127465 | 0.70629025 | 1.30588198 | 0.55849886 | 0.69931893 | -0.008104 |

| | | | | | | |
|---|---|---|---|---|---|---|
| -1.6193342 | -2.3383641 | -1.601913 | -1.5984778 | -1.5191598 | -1.7354498 | -0.0016411 |
| 0.49061298 | -0.7382102 | 1.74102831 | 0.70812041 | 0.63809776 | 0.56792985 | -0.0016411 |
| -0.7812741 | -0.8477898 | -0.8791797 | -0.9942226 | -0.7090945 | -0.8423121 | -0.0005538 |
| 5.07467365 | 6.35750008 | 2.87350941 | 3.85703659 | 0.84592009 | 3.80172796 | -0.0015049 |
| 4.11894655 | 6.36771774 | 4.46624327 | 3.75388479 | 3.31800985 | 4.40496044 | -0.0015049 |
| 0.97489822 | 0.69401169 | 0.89949489 | 0.44868016 | 0.9653945 | 0.79649589 | -0.0015049 |
| 3.84457469 | 1.88881922 | 3.21429825 | 1.25575113 | 3.64697552 | 2.77008376 | -0.0007958 |
| -0.4169135 | -0.1749413 | -0.3850374 | -0.2655883 | -0.2146332 | -0.2914227 | -0.0063794 |
| 0.51193404 | 0.57930255 | 0.40249872 | 0.3868854 | 0.30447364 | 0.43701887 | -0.007179 |
| 1.33896351 | 0.94491005 | -0.0848644 | 0.09713745 | 1.23146367 | 0.70552206 | -0.0051494 |
| -0.4909463 | -0.7855978 | -0.4414308 | -0.5010149 | -0.9568994 | -0.6351779 | -0.0017434 |
| -0.3860748 | -0.0804231 | -0.4088902 | -0.3943791 | -0.3530211 | -0.3245577 | -0.0021945 |
| 4.28831434 | 3.70223832 | 3.19639993 | 0.20909739 | 4.01189804 | 3.0815896 | -0.0021945 |
| 5.57267904 | 8.17109776 | 10.9950008 | 7.57535839 | 8.77568626 | 8.21796446 | -0.0014895 |
| 1.97572136 | 1.25694728 | 1.81018019 | 1.76898694 | 2.37855768 | 1.83807869 | -0.0041701 |
| 1.59347725 | 3.30752373 | 1.21899891 | 1.12332869 | 2.0009079 | 1.84884729 | -0.0059735 |
| 0.31008816 | -0.8294446 | -2.1900144 | -1.5995674 | -1.8727059 | -1.2363288 | -0.0059735 |
| 1.52488172 | 0.70087457 | 1.95731425 | 1.62152302 | 1.63909554 | 1.48873782 | -0.0059735 |
| 0.67439675 | -0.1831312 | 0.66944027 | -0.4627481 | -0.3095062 | 0.07769032 | -0.0059735 |
| 0.52623439 | 0.53128791 | 0.66840839 | 0.40326047 | 0.58350563 | 0.54253936 | -0.0059735 |
| 3.34810472 | 2.99669075 | 3.28680348 | 3.34144306 | 3.36189318 | 3.26698704 | -0.0020566 |
| 0.69065714 | 2.9547472 | 1.45080626 | 2.77237034 | 3.90111732 | 2.35393965 | -0.0020566 |
| -1.3271687 | -3.2021327 | -2.6941848 | -3.0047064 | -1.8062034 | -2.4068792 | -0.0026059 |
| 1.16040802 | 1.98371613 | 1.81647539 | 2.48392582 | 0.66819191 | 1.62254345 | -0.0026059 |
| 0.5062573 | 1.42393565 | 1.12939262 | 0.65131879 | 0.39765811 | 0.82171249 | -0.0026059 |
| -0.3765574 | -0.6082343 | -1.3470676 | -1.9067397 | -2.8081682 | -1.4093534 | -0.0026059 |
| 0.24961185 | 0.43975258 | 0.27173495 | 0.03451324 | 0.38417721 | 0.27595797 | -0.0026059 |
| -0.6576757 | 0.27075768 | -0.0188198 | 1.03222668 | 0.87048721 | 0.2993952 | -0.0026059 |
| 2.01662564 | 1.30674291 | 1.47076035 | 1.26649332 | 1.41224432 | 1.49457331 | -0.0026059 |
| 0.98096657 | 1.13534009 | 1.87980676 | 0.99484861 | 1.042961 | 1.20678461 | -0.0026059 |
| 1.50133014 | 2.68891644 | 1.0735178 | 1.13370848 | 2.60087633 | 1.79966984 | -0.0026059 |
| 1.82734776 | 1.78089321 | 0.55268836 | 1.8641572 | 1.76685178 | 1.55838766 | -0.0021362 |
| 1.89504254 | 4.50001621 | 4.61821985 | 5.89761877 | 3.26139402 | 4.03445828 | -0.0021362 |
| 3.10447168 | 3.61833763 | 2.82991648 | 2.41436434 | 3.00274038 | 2.9939661 | -0.0022668 |
| 0.51861358 | -0.0067568 | 0.05533266 | 0.72644377 | 1.00509191 | 0.45974503 | -0.0022668 |
| 0.25087655 | 0.38323343 | -0.1692641 | 1.62308502 | 0.90199852 | 0.59798589 | -0.0038781 |
| 0.36163425 | -0.2255657 | 0.22965646 | -0.024647 | -0.0584712 | 0.05652137 | -0.0041975 |
| 2.65868711 | 2.71883988 | 2.32403111 | 2.69026041 | 2.36792779 | 2.55194926 | -0.0154931 |
| 7.82660341 | 7.66321325 | 6.75176334 | 6.40353012 | 5.96831799 | 6.92268562 | -0.0040199 |
| 6.65262604 | 5.22376251 | 5.46661091 | 6.30446863 | 3.74832296 | 5.47915821 | -0.0040199 |
| -0.2150879 | 0.69544315 | 0.6138072 | 0.58413267 | 1.89914751 | 0.71548853 | -0.0035637 |
| 3.19033742 | 2.92748713 | 2.32738423 | 2.3377707 | 3.74251938 | 2.90509977 | -0.0048313 |
| 0.2001338 | 0.37790585 | -0.6085558 | 0.96995401 | 1.70226324 | 0.52834022 | -0.004376 |
| -2.3508937 | 1.53598356 | -2.0859435 | -0.6305747 | -0.4999161 | -0.8062689 | -0.0018945 |
| -1.6991462 | -1.2884401 | -1.2257693 | -1.7192366 | -1.0678265 | -1.4000837 | -0.0059677 |

| | | | | | | |
|---|---|---|---|---|---|---|
| 0.70589435 | 1.58817494 | 1.46587741 | 1.45100594 | 0.53444517 | 1.14907956 | -0.0059677 |
| -0.3184173 | -1.5686411 | -0.6943936 | -0.820204 | -0.8860917 | -0.8575495 | -0.0035365 |
| -0.6572032 | -0.1884265 | 0.31970549 | -0.0782015 | 0.51674199 | -0.0174767 | -0.0035365 |
| -0.5144858 | -0.5352225 | -1.2424943 | -0.9041457 | -0.1908712 | -0.6774439 | -0.0035365 |
| 1.06250668 | -0.2983298 | 1.03469014 | 0.19865942 | 0.8327775 | 0.56606078 | -0.0035365 |
| 2.02425623 | 1.04490089 | 0.93241179 | 0.60210848 | 0.09022474 | 0.93878043 | -0.0031981 |
| 2.69640589 | 2.03981209 | 1.66126299 | 1.96898007 | 1.48556137 | 1.97040448 | -0.0048131 |
| -1.3668042 | -0.9719696 | -2.0240247 | -2.1315837 | -1.0735972 | -1.5135959 | -0.0012707 |
| 1.13241577 | 1.14182711 | 0.92792982 | 1.47026157 | 2.05237627 | 1.34496211 | -0.0030983 |
| -0.2568023 | -0.0259733 | -0.19385 | -0.5750296 | 0.38202167 | -0.1339267 | -0.0030983 |
| 2.56921744 | 1.83404136 | 2.38511038 | 1.82428443 | 3.12968779 | 2.34846828 | -0.0037831 |
| -0.7374175 | -0.0080712 | -1.4879541 | 0.38602018 | 0.49814725 | -0.2698551 | -0.0037831 |
| 1.19192147 | 1.82930255 | 0.84409285 | 2.02830648 | 2.47773647 | 1.67427197 | -0.0037831 |
| 1.39821482 | 4.78788567 | 2.86209345 | 4.8160491 | 4.12082958 | 3.59701452 | -0.0037831 |
| 1.65529466 | 1.2654593 | 1.8603065 | 1.72236872 | 1.8737731 | 1.67544045 | -0.0053965 |
| -0.0246835 | -0.1530213 | 0.03025103 | -0.0739827 | 0.11147237 | -0.0219928 | -0.0008011 |
| 2.38677406 | 3.22503281 | 1.94951499 | 1.72888517 | 0.63379097 | 1.9847996 | -0.0025543 |
| -1.0701822 | -1.1176894 | -1.5535058 | -1.3477904 | -1.705968 | -1.3590271 | -0.0056595 |
| 2.31833553 | 2.42124867 | 2.31110311 | 2.50049996 | 2.25760436 | 2.36175833 | -0.0056595 |
| 2.40117669 | 2.90627098 | 2.78895712 | 3.33476782 | 2.80927086 | 2.84808869 | -0.0056595 |
| 1.02710485 | 1.48256612 | 0.96671212 | 1.60736966 | 1.35700369 | 1.28815129 | -0.0056595 |
| -0.8125396 | -0.9508367 | -0.5882182 | -1.0480788 | -1.3001454 | -0.9399637 | -0.0056595 |
| 0.12697792 | 1.42532504 | -0.1460731 | 0.31757498 | 0.56312275 | 0.45738552 | -0.0056595 |
| 0.03728509 | -0.0037243 | 0.01017094 | -0.0074592 | 0.27004933 | 0.06126437 | -0.0006098 |
| 1.45825243 | 1.34039414 | 0.92476654 | 1.8627975 | 1.63587141 | 1.4444164 | -0.0009867 |
| 2.0285759 | 1.47018909 | 0.14026451 | 1.12496328 | 1.76816988 | 1.30643253 | -0.0009867 |
| 1.00349784 | 1.19354916 | 1.45922852 | 1.08287978 | 0.99382114 | 1.14659529 | -0.001021 |
| 2.41130543 | 1.76431859 | 2.16292143 | 3.35891557 | 4.22255373 | 2.78400295 | -0.0022749 |
| 0.08285022 | 0.1192708 | 0.13528514 | 0.057652 | -0.2160201 | 0.03580761 | -0.0072195 |
| -0.4208131 | -0.3196177 | -0.5756779 | -0.4300532 | -0.0951676 | -0.3682659 | -0.0072195 |
| 0.08553815 | 0.07276011 | -0.0545976 | 0.05258512 | 0.16080213 | 0.06341758 | -0.0072195 |
| 1.18597364 | 1.10675788 | 1.11989546 | 1.07193923 | 0.90032625 | 1.07697849 | -0.0072195 |
| -0.3982942 | 0.12871146 | -0.1611378 | -0.086376 | -0.1282256 | -0.1290644 | -0.0072195 |
| 1.00581741 | 0.82120562 | 0.91066265 | 0.99944639 | 0.76719689 | 0.90086579 | -0.0072195 |
| -0.3147409 | -0.3529074 | -0.2606333 | -0.4251397 | -0.1854051 | -0.3077653 | -0.0009345 |
| -0.6531155 | -0.7447788 | -0.2785742 | -0.2124145 | 0.10847902 | -0.3560808 | -0.0024908 |
| 1.06360054 | 0.9748683 | 0.74705791 | 1.02991199 | 0.75595403 | 0.91427855 | -0.0078778 |
| -0.1605392 | -1.216588 | -0.5287919 | -0.4587986 | -0.6888356 | -0.6107107 | -0.0022907 |
| -2.1755166 | -1.3736949 | -1.4701853 | -2.1635323 | -1.8214455 | -1.8008749 | -0.0047351 |
| -3.2149692 | -2.6159062 | -2.8042967 | -2.9879551 | -3.0170035 | -2.9280262 | -0.0047351 |
| 1.2472465 | 1.03676021 | 0.62927198 | 1.07135355 | 0.70765066 | 0.93845658 | -0.0047351 |
| -1.3987179 | -1.9943838 | -1.7472334 | -1.358005 | -2.5402255 | -1.8077131 | -0.0047351 |
| -0.0088665 | -0.3122077 | 0.20943367 | 0.20763183 | 0.2094028 | 0.06107881 | -0.0047351 |
| -3.1550584 | -2.6186934 | -2.2562802 | -2.2622957 | -1.8008757 | -2.4186407 | -0.0047351 |
| 1.33538151 | 1.55610836 | 1.31093526 | 2.00747299 | 1.04753447 | 1.45148652 | -0.0047351 |

| | | | | | | |
|---|---|---|---|---|---|---|
| 0.5287168 | 0.92795229 | 1.12123775 | 2.18316746 | 1.39761758 | 1.23173838 | -0.0047351 |
| 6.65266991 | 6.08591652 | 5.38004541 | 6.22679377 | 7.75513506 | 6.42011213 | -0.0047351 |
| 6.60546112 | 3.99845076 | 3.81535912 | 3.86445045 | 5.51581955 | 4.7599082 | -0.0047351 |
| -0.9721155 | -2.0879595 | -1.5509372 | -1.0828569 | -0.7693357 | -1.292641 | -0.0047351 |
| 3.11732507 | 3.46761012 | 2.77093697 | 4.80812073 | 1.18763089 | 3.07032475 | -0.0047351 |
| 2.36397362 | 0.90960699 | 0.60472065 | 1.24887037 | 1.02878237 | 1.2311908 | -0.0047351 |
| 2.38013864 | -0.0975423 | 1.13436389 | 0.22007656 | 0.53273964 | 0.83395529 | -0.0047351 |
| 2.48218989 | 2.16761088 | 3.16764212 | 2.69921494 | 2.25117779 | 2.55356712 | -0.0047351 |
| 2.21521854 | 2.27236652 | 1.02303123 | 2.85634089 | 2.83339477 | 2.24007039 | -0.0047351 |
| 1.32768679 | 1.90160024 | 1.22832632 | 0.94650054 | 1.18285632 | 1.31739404 | -0.0047351 |
| -1.4367185 | -1.0078344 | 0.17860603 | 0.98754406 | -0.660429 | -0.3877664 | -0.0047351 |
| 4.34296417 | 5.33979607 | 5.61062574 | 5.80147314 | 7.20671558 | 5.66031494 | -0.0047351 |
| 0.87100959 | 0.66354555 | -0.3970199 | 0.16797209 | 0.47947693 | 0.35699686 | -0.0047351 |
| -0.3780293 | 0.35229975 | 0.58735228 | 1.2444551 | 0.86058182 | 0.53333192 | -0.0047351 |
| -1.3188571 | -0.8469398 | -0.5354441 | -1.5673994 | -4.1145043 | -1.6766289 | -0.0047351 |
| 1.94733799 | 1.59420812 | 0.39602399 | 2.16912055 | 2.65547895 | 1.75243392 | -0.0047351 |
| 2.13250661 | 2.45462751 | 1.02801847 | 2.79362822 | 3.10384989 | 2.30252614 | -0.0047351 |
| -1.9677041 | -0.9202912 | -2.0361793 | -1.1720735 | -2.5923254 | -1.7377147 | -0.0047351 |
| -1.820087 | -0.5545678 | -1.915442 | -1.5098445 | -0.9605315 | -1.3520946 | -0.0047351 |
| 3.52769375 | 2.27895594 | 3.30843854 | 3.95772552 | 2.66591454 | 3.14774566 | -0.0047351 |
| 3.90835333 | 5.54818249 | 2.39212942 | 2.47984505 | 2.78177094 | 3.42205625 | -0.0047351 |
| 1.80074477 | 2.37480783 | 2.19770217 | 3.08872843 | 3.50798059 | 2.59399276 | -0.0047351 |
| 0.978091 | 0.92913747 | 1.48175263 | 1.16691089 | 1.36893082 | 1.18496456 | -0.0047351 |
| 1.67447925 | 0.33165812 | 0.62116075 | 0.18429685 | -0.2101181 | 0.52029538 | -0.0047351 |
| 1.34786773 | 0.37853932 | 1.36763406 | 1.05768335 | 1.24824834 | 1.07999456 | -0.0047351 |
| 1.38060522 | 0.76151848 | 0.31763482 | 0.47123575 | 0.93512774 | 0.7732244 | -0.0047351 |
| 0.56435323 | 0.44183683 | 1.43806267 | 1.12785578 | 0.97004938 | 0.90843158 | -0.0047351 |
| 1.25972331 | 0.45966434 | 0.96243632 | 1.00719166 | -0.57056 | 0.62369113 | -0.0047351 |
| 2.21952224 | 2.07867718 | 1.26841903 | 1.5381093 | 2.36691451 | 1.89432845 | -0.0047351 |
| 1.4442178 | 0.93487096 | 1.18198049 | 1.55369127 | 1.64229178 | 1.35141046 | -0.0047351 |
| -0.3617353 | -0.3744588 | -0.278774 | -0.1236069 | 0.04946947 | -0.2178211 | -0.0047351 |
| 3.41350412 | 2.60995436 | 0.95663071 | 2.48171473 | 2.11268616 | 2.31489801 | -0.0047351 |
| 0.83505023 | 0.35014009 | -0.338163 | -0.5355151 | 1.09080076 | 0.2804626 | -0.0047351 |
| 2.45743084 | 2.1104331 | 3.96629357 | 0.72075987 | 3.14227009 | 2.47943749 | -0.0047351 |
| 0.45066381 | 0.30428267 | -0.7609301 | 0.02388906 | 0.01032591 | 0.00564626 | -0.0047351 |
| 0.51279521 | 2.26356053 | 0.8238256 | -0.0918674 | 1.56270909 | 1.0142046 | -0.0047351 |
| 4.21037769 | -0.003722 | 3.0781951 | 1.50333679 | 1.78148985 | 2.11393549 | -0.0047351 |
| 2.53115559 | 2.56383371 | 1.62249732 | 1.29440355 | 2.04150605 | 2.01067924 | -0.0047351 |
| -2.058125 | -1.864733 | -1.1974204 | -2.2964466 | -1.5331409 | -1.7899732 | -0.0047351 |
| 2.71697044 | 1.83064747 | 2.10165358 | 2.69148159 | 3.56242108 | 2.58063483 | -0.0047351 |
| 0.16188622 | -0.3250601 | 0.45112658 | -0.1319656 | -0.6964655 | -0.1080957 | -0.0047351 |
| 1.84985864 | 1.54396105 | 1.03452373 | 1.2333293 | 0.75620747 | 1.28357604 | -0.0047351 |
| -1.3945286 | 0.00095606 | 0.38680339 | -1.0728574 | -0.4619927 | -0.5083239 | -0.0047351 |
| 0.46456957 | 0.29024935 | -1.5240636 | -0.3818281 | -1.1296761 | -0.4561498 | -0.0047351 |
| -3.4167361 | -2.618252 | -3.1613126 | -3.9600894 | -3.1518281 | -3.2616436 | -0.0047351 |

| | | | | | | |
|---|---|---|---|---|---|---|
| -1.2326268 | -0.4215405 | -1.2961433 | 0.38463616 | -0.4822977 | -0.6095944 | -0.0047351 |
| -2.5512433 | -3.5788307 | -2.4175358 | -2.1859932 | -2.0908883 | -2.5648983 | -0.0047351 |
| -0.1930008 | -0.1022019 | 1.11800587 | 1.04232216 | 1.18914461 | 0.61085398 | -0.0047351 |
| -0.4955242 | 0.27657461 | 0.20221353 | -0.8587273 | -1.0613813 | -0.3873689 | -0.0047351 |
| 0.97147727 | 1.08064485 | 1.66892982 | 0.79004478 | 1.50573301 | 1.20336595 | -0.0047351 |
| -1.0305603 | -1.8730545 | -1.844036 | -1.300005 | -0.9980595 | -1.409143 | -0.0047351 |
| 0.73110199 | 0.21465206 | -0.1283333 | 0.22996116 | 0.22458816 | 0.25439401 | -0.0047351 |
| 1.62943041 | 0.80465865 | 1.37332726 | 2.19548631 | 1.54665351 | 1.50991123 | -0.0047351 |
| 1.76730895 | 0.97010422 | 0.77213478 | 0.79675245 | 0.51746273 | 0.96475263 | -0.0047351 |
| 1.60751522 | 1.65645993 | 1.71782517 | 0.34687424 | 2.99883413 | 1.66550174 | -0.0047351 |
| 2.08930421 | 1.38802052 | 2.2624929 | 1.17372239 | 1.73427212 | 1.72956243 | -0.0047351 |
| 1.40091991 | 0.83684587 | 2.69534373 | 0.43863726 | 2.01865244 | 1.47807984 | -0.0047351 |
| 0.11663628 | -0.3867419 | 0.33028865 | -0.0197301 | 0.12030703 | 0.032152 | -0.0047351 |
| 2.34133577 | 3.10172534 | 2.52841544 | 2.57353687 | 2.78663135 | 2.66632895 | -0.0047351 |
| 3.05086327 | 2.32190084 | 2.24153638 | 4.18233871 | 2.81761551 | 2.92285094 | -0.0047351 |
| 0.40188503 | 0.03937054 | -2.3724012 | -0.0897667 | 0.03416204 | -0.3973501 | -0.0047351 |
| -0.532623 | -0.0704684 | 0.21634197 | 0.17020607 | 0.0890398 | -0.0255007 | -0.0047351 |
| -2.2822223 | -2.147028 | -1.159807 | -2.5246956 | -1.4738963 | -1.9175298 | -0.0047351 |
| 3.90442705 | 2.76989269 | 1.53699863 | 1.62813854 | 0.77745622 | 2.12338263 | -0.0047351 |
| -1.212839 | 0.38524842 | -0.4923131 | -1.0914352 | 0.59481883 | -0.363304 | -0.0047351 |
| -0.5114551 | -0.4320168 | -1.6769471 | -0.4253564 | -0.6786022 | -0.7448755 | -0.0047351 |
| 1.93903375 | 3.08961797 | 1.76377904 | 1.77019322 | 3.92400265 | 2.49732533 | -0.0047351 |
| 1.00886536 | 1.54731703 | -1.1657922 | 1.54266596 | 0.99923921 | 0.78645906 | -0.0047351 |
| 2.52755046 | 2.82994366 | 2.63020062 | 1.01339841 | 1.15249944 | 2.03071852 | -0.0047351 |
| -0.9897652 | -0.2993407 | 0.3081212 | -0.5062315 | -1.6919777 | -0.6358388 | -0.0047351 |
| -0.3430738 | 0.53529263 | 0.80207849 | 0.60460329 | -0.6844363 | 0.18289285 | -0.0047351 |
| -2.7769842 | -2.2676411 | -2.4485855 | -3.4964395 | -2.4883542 | -2.6956009 | -0.0047351 |
| 2.01311684 | 2.87650394 | 1.65146601 | 1.82114804 | 3.45392036 | 2.36323104 | -0.0047351 |
| -1.1467386 | -1.5233836 | -1.6908185 | -2.3213916 | -0.401216 | -1.4167097 | -0.0047351 |
| 3.26945567 | 2.149827 | 1.59289122 | 1.27321434 | 2.30268478 | 2.1176146 | -0.0047351 |
| -0.0938387 | -1.0473137 | 0.14470935 | 0.28105736 | -0.3280206 | -0.2086812 | -0.0047351 |
| 2.07499743 | 1.82914543 | 1.04481888 | 2.45893621 | 1.99161315 | 1.87990222 | -0.0047351 |
| -0.6823123 | -1.3969712 | -1.4005828 | -0.7841573 | -0.7711415 | -1.007033 | -0.0047351 |
| 1.31277943 | 1.03426886 | 3.03720117 | 1.91820621 | 1.33130336 | 1.7267518 | -0.0047351 |
| -0.5012326 | -0.5307422 | 0.0720861 | 0.52945232 | -1.271302 | -0.3403477 | -0.0047351 |
| -1.2023867 | -0.7373276 | -1.8917969 | -0.38712 | -1.5419011 | -1.1521065 | -0.0047351 |
| 1.64391303 | 1.59663796 | 2.17144656 | 2.12988758 | 2.23575401 | 1.95552783 | -0.0047351 |
| -3.331527 | -3.6242752 | -3.8224704 | -2.4877439 | -3.1404138 | -3.281286 | -0.0047351 |
| -0.0116386 | 0.32646132 | 0.27290773 | -0.5917149 | 0.53046918 | 0.10529695 | -0.0047351 |
| -0.1582494 | -0.4458704 | -0.3217421 | -0.3756557 | 0.02928972 | -0.2544456 | -0.0047351 |
| -3.713639 | -2.3414245 | -3.2799652 | -3.4076147 | -1.6414813 | -2.8768249 | -0.0047351 |
| 0.66046625 | 0.89052159 | 0.93618399 | 0.66097343 | 0.67240763 | 0.76411058 | -0.0047351 |
| 0.67096668 | -0.1810961 | 0.22126675 | -0.3729677 | -0.1432672 | 0.0389805 | -0.0047351 |
| 2.2769928 | 1.72790313 | 0.63914609 | 0.18251657 | 1.33188462 | 1.23168864 | -0.0043641 |
| -6.5226216 | -6.3863649 | -5.6681232 | -5.1480551 | -5.9967337 | -5.9443797 | -0.0013682 |

| | | | | | | |
|---|---|---|---|---|---|---|
| 3.7615788 | 2.39294529 | 2.38671255 | 3.68736243 | 3.22702789 | 3.09112539 | -0.0013682 |
| 0.99430609 | 2.84076643 | 2.0250752 | 2.73649383 | 1.70137191 | 2.05960269 | -0.0049086 |
| -0.1343489 | 2.17521429 | 2.22404909 | 1.45635295 | 1.1798296 | 1.38021941 | -0.0024692 |
| 1.70089746 | 1.34860849 | 0.96678925 | 1.17515576 | 0.83349478 | 1.20498915 | -0.0130085 |
| 0.74325132 | -0.138823 | 0.77773952 | 0.18926787 | 1.43528175 | 0.60134349 | -0.0038876 |
| 1.37734437 | 1.10733354 | 1.29653192 | -0.6616828 | 2.25883198 | 1.07567179 | -0.0038876 |
| -1.3185825 | -0.670893 | -0.6638451 | -1.3527174 | -0.2669115 | -0.8545899 | -0.0038876 |
| -0.7402389 | 0.09481931 | -0.1180644 | 0.21178102 | -0.0581598 | -0.1219726 | -0.0038876 |
| -2.261713 | -3.098474 | -2.0903397 | -3.8283734 | -1.4314756 | -2.5420752 | -0.0038876 |
| 0.20330143 | 1.20861316 | 0.25958157 | 0.06122589 | 1.81195331 | 0.70893507 | -0.0038876 |
| 1.1227541 | 0.8197037 | 1.31184316 | 1.07872653 | 1.71913028 | 1.21043155 | -0.0038876 |
| -0.8104972 | -1.571364 | -2.5099242 | -0.8432224 | -2.3502936 | -1.6170603 | -0.0038876 |
| 0.02987504 | 1.29457736 | 0.37835598 | 2.95357323 | 3.61258793 | 1.65379391 | -0.0038876 |
| -1.0102859 | -4.3430557 | -2.196969 | -1.1089277 | -3.6900074 | -2.4698492 | -0.0014752 |
| -0.0889902 | -0.2865036 | -0.0622406 | -0.6203597 | -0.9997399 | -0.4115668 | -0.0019693 |
| 0.99355149 | 0.90060937 | 0.98196828 | -0.2457975 | -0.0608485 | 0.51389663 | -0.0019693 |
| 1.17025232 | 1.99447775 | 1.42366624 | 1.31801796 | 1.68792033 | 1.51886692 | -0.0019693 |
| 3.22263241 | 4.64729786 | 1.90500259 | 3.40063667 | 4.20883465 | 3.47688084 | -0.0026099 |
| -0.9544528 | -0.0691683 | 0.59139943 | -0.7829387 | -0.7785201 | -0.3987361 | -0.0022051 |
| 1.85862684 | 2.14883423 | 2.06456637 | 2.06659889 | 1.48812819 | 1.9253509 | -0.0022051 |
| 3.40268564 | 1.75322127 | 3.69824409 | 3.74415255 | 2.26921606 | 2.97350392 | -0.0022051 |
| -2.8693492 | -1.3206117 | -1.1825358 | -6.0031309 | -3.9037805 | -3.0558816 | -0.0022051 |
| -2.0255082 | -2.923214 | -1.9266615 | -1.835911 | -1.2813809 | -1.9985351 | -0.0022051 |

VEPPRETLEAVLLTSSLSVLLLPYDEMKTAQQMTTSRAASGHSRAFTQLKPFAPDGIILTTSRDLLLYPTNTDIYL
IPGSAMETGLLPNHKLKNVGGDHPAAPPHHHAHHPHHHAHHHHAHHQHHHHHALQQQLNQFQQQ
AIQLQLHAKTGQSVDSGAIKAKLSVLSVDELERELEANKKEKVKEAQLEAEVKLLRKENEALRRHIAVLQAEV
AFTEYVFRVVWIITSQMQLYSPSSPSYRTHAFGVPVTAPIIKDIQSSGPDTVEISWSPPLFPSGLIVGYNLLLTS
TKVDSSVSRLNKKYDVLSALYHKFERTCALIYMIQASSQTSAELSLPLVLKTSWITFLLAKGKVLQVEDDLVISF
YEEQSQDRRNLTKLSLIFSHMLAEIKGIFPNGQFQGDNFRITKADAAEFWRKFFGDKTIVPWKVFRQCL
SADHMKTQDSQHTSHSVTPSTASAPTSSTPSHGLTATSEPASGQKTPSKVVYVFSTDMANKAAEAVLKGQA
KYTLAVQSENRNYTIMFTVLVKSKPRKPYFVKGENLGIINCTSEGYPTPDIEWLFCRTADDSCINTNRKDSK
AAPQKKDSMGELSRATIRLLEELDRERCFLLSEIEKEEKEKLWYYTQLQSLSTRLDELPHMETFSMQMDLIR
NQMNRNDRTLIFLDENEMDMTASHTAVITRNLKSNEEIDKPRKIDIKSFLAGLTSKNEGPKMNKEFRFFC
PFPVPSPETKVEVTVQADSSLVPSKPFLLPSLHNRVLDFSLLSSAAPSFPPPTLEPEVREATTLSQAQTTETNLQI
ETEKEPEHNYQGTEKREESFSNDAIYAVPYKNKKVKLPVSSSVDLDENEISSHEAAFPILVDQSNIQVEEDEP
DAPGQQDGAASFPAGYMYPSMDQLAEMLPGILKQFGLKSVIGMGTGAGAYILTRFALNNPELVEGLVLIN
FYCSFEEGDCGWTQSSPTSLTFPWQIGNLEYSHFPLLEGSALLLSTSKVPVAETIIVTSAVFPAPIRNSPCEL
SVSHINLTVREGDNAVITCNGSGAPLPDVDWTVADLHSITTHQTNLNWTNVHAINLTLVNVTSEDNGFLLT
SEGSGAPGEEGTGHCPSSSTDTASEHSADEDAKGAGGELWGTAPGPSDYDLRLLRSQRVLARRQGIFQ
RRKLVLSSAPATPHQPEPAALEEEGEGTSGQRGTGSKRQIRYLENLLRVYMGEIRRLQERELDLAELDSED
YISKMGYLDPRNEQIVRVIKQTSPGKGIVPNNVTRRPSFEGSNESFPSYHQISNTAYEGTNFGTEGANMLS
CPPGWSGKTCLQADPCASNPCANGGHCVPFDAHYVCKCTAGFHGANCKQDVNECNTSPPICKNGG
NTMFTNCYIYNKPGDDIVLMAEALEKLFLQKISEMTEEETEIVIVQAKGRGRGRKEAATPKPGASTVQNVTQ
SLDSPLSSELADAKNPAQESQIQENEVAKAEAESSPYNNESQIKSDHSLCPLGSEGQSPETIVEDNPSSSDT
SRSRSPKRRSPSPRSRSHSRNSDKSSSDRSRRSSSSRSSSNHSRVESSKRRSIKEKKSSSKDTRASQAVGD
KKHLLSVTAKPLSAAVAPPPSLASVAPAEIPAAAELPLTSPPLLSQTASPPPPVLTAVAFPQPPIAVSVTAAPLLLPP
LIDLGADTSLRSRWTNMNALHYAAYFDVPELIRVILKNAKPKDVDATCSDFDFGTALHIAAFNLCTAAVKC
PGSCYHVGISALAGPYSSDPQNVTGCTVPAAPANVSLTNSGSSSVLHAAWDEPPGGRDHYRVILYSLVP
GKKLADSWGAVFLESSAKENQMTRGIFTKIIEEIDRVDNSYCTERSCLLILLAVSIPPGQLSGFGAVFQNQL
PAAELGRAGPAGGSAGRGPHEPISAVLCSPSPPAERQGQAPRDAAVGKRTERVELRPVPGPLGQTRGKA
NVGIKCATITPDEKRVEEFKLKQMWKSPNGTIRNILGGTVFREAIICKNIPRLVSGWVKPIIIGRHAYGDQYR
TQTETSGTFGCHCQLSGKDEPSSYICTTCKQPFNSAWFLLQHAQNTHGFRIYLETSPSNSSLTPRITIPPPL
SDEEVKVRSPTKSPTVKSSPRKPRRRPRSSPDRSSAAHSDSSSVCSPLSKSETMSMEKVKKKEIKSGEKRRG
SSRSKRKPTVVTKYVGSDDEQTLPETVNEDVSNENSENDVDMKSLPKGTVVVQPEPVLNEDKDDFKGPEF
KPEESSNLNFEADASFLRQAIMSFGSIKTMQTSETENASPWMSVQNCTLAQHEQKKLLSGTLGSPLSDWLL
ELGMRHLTEILKGGVHIEGNPQLCFQEMILWGDIFHKHNDLALHQQINSSRERTCADCRRFCSQGHC
QKRIQEQENPTLEANAGAQVADSMVRRSSLRNKRQKCESGKAVYIEFGSDSSDDLFKKAGGGGMGDTE
TLNMQVDQFESEVESLSVQTRKKKGDKDKQDRIEGLKRHIEKHRYHIRMLETILRMLDNDSIQVDSIRKIK

KYGTELNQGDMKPPNYDSDEGNETEVQPQQNSQLIWKQGRQLLRQYLQEVGYTDTILDVKSKRVRALLGIS
PKKVTAQKRLEKRDKYEQETLTILEEKKVEAAERVSKQDSAQNELTALILCDTDIEPSIKCDSQDKKIGRPESSVS
GANGDVQQVGVIRCPICRQECRQIDLVDNYFVKDTSEAPSSSDEKSEQVCTSCEDNASAVGFCVECGEV
ITAASQPSNPSECNGLSPLVDKEGGGQMEIPDQLTGNTKKKSSKKDLINHTIQNTDVEWVKNAQKAFESKT
CSQECAERHIIDNFFVKDTTEVPSSTVEKSNQVCTSCEDNAEANGFCVECVEWLCKTCIRAHQRVKFTKD
DRNTNRPLEPPSEFIVKVQDINDNPPEFLHENYHANVPERSNVGTSVIQVTASDADDPTYGNSAKLVYSILEG
KLSPLPTPQWPGRSTPESEGGGVEEDGGPPPFSPGAGGDPAPEQLDPELARTVADSVERLLQSCPRGGEPG
AFKGNDSTYITIFLCTYRAFATTKQVLDLLLNRYGKLNVQVNGDHARHAVDERLELKNTISSILGAWLDQYSE
ISENGPCLGFRKPKQPYQWLSYKEVSERAEALGSGLLQQGCKPSASQFIGVFAQNRPEWIISELACYTYSMV
QGEKIRDLEFCLEEHREKLNATEEMLQQELLSRTSLETQKLNLMAEISNLKLKLTAVEKDQLDYEDRYGDTESI
SHCATPLLGAEDSGTSQLGFGALELSQSQDVEGDSTLSEEGRGHQLQPAGTVPTCCNPKRNDHNEDLHIV
VTRGVCQRHIIDPDNPADIESEVHCMYSKQRDGESDSCPDCVVSTLGTKVLVTEKNRFVNFFVGNTVKSSF
SRSIEAAIEFISKMSYQDPRREQMAAAAARPVNAGMKPPAGGVQQSINRKQSWKGSKESLVPQRHGPPLGD
FPGTPDAEALACFFIPVLRSLTRLRPEMALEEGVPRAVQEWERSSNFERMIFYEMAEKFMEFEAEEELQIQKN
SRSGSGSSGFLEGLFGCLRPVWNIIGKAYSTDYKLQQQDTWEVPFEEISELQWLGSGAQGAVFLGKFRAEE
LNPSKEELRPRTANKAPGNKGGKQKSPPTAAAVSGDSGPQRRTTCKKQPRRVERTSGGDALNCHTSEDLAL
LEIINAVPFIITIFWPALRNLFIPVFLNCWLAKHALENMINDLHRAIQRTQSAMFNQVLILISTLLCLIFTCICGIQ
SSAGWKNSIRHNLSLHSRFIRVQNEGTGKSSWWMINPDGGKGGKPPRRRAVSMDNSNKYTKSRGRAAK
AIKLVYFLLSTLSLITCVLAVAFVAHHYSKIMQFTCDVVLDSCQCKLETSDPLSRTFVYQDVSDCTSVTGTFSLY
YSLTKLVNGTEFFKIVLQESENSCSKTSTINVLFFGRLAGDCGKVVHQGDTMVVAGFNVAKSPSANEDNRHG
FKFGPLFMRLCYELGLEVPAIELIKDQTLHGFFSDSTSFNILMDMLFKKGHYESALEVLGEMKKQHIKIIKETY
AVACADRHHCCPRGSHCSADGRSCFMLPGLPAAGAVQCPDGESECPNESTCCMMPTGSWGCCPMPI
ANGLINRLIEENKMDSLGMVVVDELHMLGDSHRGYLLELLLTKVRFVTEKIAKRQTKENGPVFDSIQIVGMS
QKRMQPSVRSHACEVDPYGMPGGCSHICLLSSNYKTRTCRCRTGFVLGSDGRSCKRPKNELFLFYGKGR
CIERATRLVKSQQSAGKEYVGIIRLHNAIESEIQLSRAIETLTGALFQRPPLIAAVKRQLRVRTIYESKLVEYDPGR
SVRKMVKQAAVIICIKKREFDKASKILKKNMGKDPSTQKLRTELLSIIREKNLSHTMIRNFSYKTFQQSAFQFLE
YIRVVVSPEAVQMAECVQLQSGFSNIIGRIIILQKYMVCFQEETKAEECEFYLSVHQFIVLPLQRQRMEALNC
EAKALAEEAEVVDGPDENGEMFTRPGKLSDYFPKPYPNPEAARAANNGALPPDLSYIISARHGGEDYVFSLL
LETLAAETGLSVRVVQVWFQNQRAKMKKLARRHQQQQEQQNSQRLGQEVMSNRMEGMMTSYTPLAPPQ
EKPAGPPAKRAIADLP
VSKMIREDGYDSVFSVVRRHLFRWSEIKKGVNKVTTPQNLDPAKRPRRQDWDGELYENGSFYFAKKNLIEK
MYLGGVIFINRKSTSSAKTVMSGVVQTMLDENVKVWVYPEGTRNGNGDLLPFKKGAFHIAVQAQVPIIPVVYS
NVRDKHLPVSDAFPHFLADSHHQVCMLAAKSIASLFQDMKQRDSSGLPKALPLKFQQKAFENVYLRVQEG
TEWLMHEGKCYWASKESRIWHKSHDDCSAKSSRLLVIQDQKEMDFISSVTQDTNPVWLGLTMTSPEWKV
TSSSPLDMQTSQMKVISKEIDGKINGIPRLEEEAQSFCQFVTSGESAHVRAKLAQLRRYWEEVRDHARCLEG
VKNCSMFKAKYICRQNLGTPVNPELPAPYPTPSLTGSCPQGWSSDPKLRHCYKVFNYDKLQDKKTWTLAQL
EGKDTDDWMCIDFGSIVIHFMLPETREIYELEKLWTLRSYDDQLVQMVPESLPEDFIVGLTHDQQ
VDSTPDSLFLNFLHTVKEDTLQYLVRYWEQSTNKQHETEWTKNTRIGLLNLKQLTVYCFQIRAKVKVFGDNIG
GKLAPKGRLDMEQLAQRLAARFPGLFSPQRRFAFTSSSKHRCMESSAAFRKGLQLALHRQPPARDIENEET
LEGLRVKLSPYVDEVQQKISKNLEVLRFQMIPFTQELLDQVMLKARELQQHLTPTRDMKVQFLEGIDEVQRFV
YDHSSLQRGLHGSSALFEVNSSPSSLKEDIKNSVNLNLTVQSAAVQSCLRPSVQDGLPWCPAQGRLRKKRI
ISVEPERITEEQEEILSILEEKFPELLPREDIISVFQQTPFKAQGFNIEEAMLKDLKELSDGEIKIAVSTVYVTLEDC
AAAARPDDILFRDEIIANGFHSCDSDEDDRASRASSSDWTPRPRIGPYTFVQQHLMIGTDPRTILKDLLPETI
LGTPELVTATECVHYFEWRTFIACKKGSFKAKKEVPCYVFDDLKKHDLNPLIKTSGGYLVDDSDEDSLYINVC
TQSEAYVQDYFPLVEQYDFYSGVGFSFLFSWVLMITVVITFITGGNVEKLVCEPFEDKTLFKYTEDVSLKIEHIN

AWTWPGALNIENSESESWTSHTVKKLTASFDASLTGEREPKVLSNDHKEALKRDCDDSEMRDEVYHRKIIS
MPKSIERLQCRRCVVVGNSNSIRGRGLGAVIDSHNIIIRLNNAPVKEHKEDVGEKTSIRLFFPESALPNPLD
TVVAVPYGSRQVRLVLKGPDHLYLETKTLQGVKSENSLSSTGSFLVDNSSIDFQKFPDKEILRISGPLTADFTV
NVYKSKKYGVSVSLSSAKQRQQFKGKDFLPLSHFLPMINTVPVESMDFVEYGDYSHTFESDVFSSPLETDS
VSDSKYGILKPSNLLNFQRHPELLQPYDPEKNNLNLDSYQSSVCNKSKSLADEGKIETKPWTSSSKVVTLQ
VSSTEILEDKPLSNTGSGQTVATQMSPQKIKLHLRPDDSQIFSVQVRQVEDYPVDIYYLMDLSNSMKDDLH
GGMEDLSKERSLTGKLEPVSPVSPAHPDSELDLLPSRLSKEELIQNMDRVDREITMVEQQISKLKKKQQLE
GYSGGFTPNPTYKEVCTGLTGHAEVVRVVYDPQKTSYEALLKAFWENHDPTQGMQQEENVGTQYRSVIHT
CSEHLRPHQESPAFRDHQLCPPLRDLQQRKCQEHNKLLEFYCRDHASCICCVCLVSHKACATGPVQEAK
CINENGYCEHYCIDSPTMIRQCFCAEGYILASDEVSCTPQVDYPCGKIHVLAKQSTSPQGRIVGGYVCPP
AAVMVEVNCETDFVARNVKFQQLVQQVAAGTMFYHQGAKDHLTTYIKRFMKAAELSQLRTGPAGCLLSDQ
NVKSSRGLEGNVMIDLYEVAGIKGMFLANKKIDNQVKTFITYNKGRDWRLLQAPDTDLRGNPVHCLLPYC
HSNGEQASGISSSKNCYDIENHGSEDFLLVLDCCDLSVLKVEEVDVAAVPGIEKFTRSAKLTDAEELRNLR
NAGPKDMVVGFPNLGILHVTKKKVFETLEARMTDACKKGYNPGLLVHPELNYIQAEGGGERQLTEREREII
RAHRCVLAAGSPFFQDKLLLGYSDIEIPSVVSVQSVQKLIDFMYSGVLRVSQSEALQILTAASILQIKTVIDECT
AKCINPRVKSGRFMKILPDYEHMEYRDVYTCLLHRYRHILGLWQPDIGPYGGLLNVVVDGLFIIGWMYLP
KGGRIVSSTTRSTMLHYRSGLLQTLDTLVFAGLFFLGFAEQKQMVEVELYSDYREDSYVPTVGAVIEIQTKRIQ
AKSEDNGRPGLSSEKQSVHILPKDSSGKQAPASV
ELPLEPDGAARYMLPAVLRPRYTPHGWEGEDVTQGVPRVPQGKLPYTLSLSATLQSTHGIDRVLSNCSLSPL
NAMKHLPNFFHKGQLAKMFFLFPDPHFKKTKHKWRIISSTLLAEYAYVLRVGGLVYTITDVEEVHEWMVGH
IESGEVDLDDLDTPSENSNEFEWEDDLPKPKTTEVMRKGSLPEYTAAEERDDGRRWRMFRIGEQDHRVD
KICQVKNCKNIPVKRSFFFLANWNEKINFPVQIKSLPNETILTIKLFGENSVSKNTEILAWTCSPLYQKQKFIH
KDGYRTAGQACELVHCPELQPPENGYFIQNICNNYFNAACGIRCKAGFDLVGSSIRLCQPNGLWSGSET
QDKDAQFYRFAEGELTLEPGAWDAEELLEALAFLAQLGPDALLTMALRKPPAQRTEDELELIFEELLHIKAVA
FRESFGHNADYSLGEALWACCNLFSDVQLKMSHKRIMLFTNEDNPHGHDSTKAKLTRTKAADLRDTGIFL
KIGPPGVHLESIDGVVKINISPPEENQNRKMWTNDISFTYNAVFWENSSNAEPKSKTINRRDTIYDLAPETTY
IFNNNPALWKEIRPFFTKALSGPGLMRMIAICVESTKDHLDRLENVTTGLGNINALNFMRQITLDTSNTLFLG
VHRNVPAAHLARARVCALSWGLDHEEFLRDYDFILGADIVYLKDTYPLLIRTLRHLCGPHSTIYLSSKMRQE
FTHPCDHEELREMLTHRNGPTKKGKELNTERSFFLRMKCTLTSRGRTVNIKSAAWKVLHCTGHVRVYDTC
VLNVHPSYLPRWRGPAPIVHAVLHGDTVTGVTIMQIMPKRFDVGPIIKQERYAVPSQCTAKELEAILSKLGAN
KRVKPGAVCSVAGWGSTSTDRKSSSTRLQEVDVVVMPDAACPRKPNGPYRFYDASTMMCVGDPKMGKD
FAVCFPLRAKVVITKGKVKLVILVLWAVSFASAGPIFVLVGVEHENGTNPLDTNECRATEYAIKSGLLTIMVWT
IMDPQPPSQIDGSSVDVHINMEQAPIQSEPRVRLVMAQHMLRDVQGILGRLEGRTNGQPQPTPENAPSA
TPDPSTIISYLPGLLYKFSCSYPLEYLVNNTQLASSSAAISVKENNGTFISTLNLLLYNDSTYSQLLVIPSIGLPLKT
NVPGAVKDVCVTEHLNPEKAANCNQIPELCYSLGYDRQKAELCVAFLQAMHGSMIGDQDAGCDCYILGT
QMAPYACQRQTPLLHSNQLLTEALETLAEEAGFSGSEGRSLAFTRAASVLKALPDRLSTLEELGPLPGIGEHS
PVPREVEAPGKQQPPSDTTGLFVMDEDSPSQDYEPFFDSDPESTDDGSLSEEAPAQPPPAPPNHQYAKSL
PISTPSRVPLAPGGEDGTPAHSEELELPWWADAAPDPSGSLPQYTLAELPAEPSPEPLRQRDPSTQQALRKG

MVDKAIYLSGYRMRLGSSTDKKDSGRLHVDFAQARDDFYEWECKQRMLAREERHRRKMEEDRLRPPSP
MTKSFVCRPGSGGDLLPLAGCLGSRFRERHSMIQWLEVGARQIQTGNTIIF
PTGADALCALDILSSAGLLNHFGRSSSVEKIDISPILLHKGSTKIALYGLGSIPDERLYRMFVNKRVTMLRPKE
LAAEYPSLVSKQEIGESYEKRPMYVLKFSTGGTNRPAIWLDAGIHSREWITQATALWTAEKIASEYGNNPSVTS
QQPQKQHPQHHPSQQPPTVFQQPGPPSQCPIGVQQQQMPASPQTQCQQPQQTAHHQLKAQPEMQS
ENKIVILLDYMFQDLHNLKSLEVGDNDLVYISHRAFSGLLSLEQLTLEKCNLTAVPTEALSHLHNLISLHLRHL

KSPKEYYVQQEVIVLFCETVERALKLGYLTQDMIDDYEPALMFTIPRLAIVCGLVVYSEGPLNLDRKPEDMSEL
[illegible]PVIAAVNGYALGGGCELAMMCDIIYAGEKAQFGQPEILLGTIPGAGGTQRLTRAVGKSLAMEMVLTGDRIS[illegible]
LTAAHCNGDKITVYLGAHNIKQQEQSQQKISVRRQIPHCKYNRKIFNNDIMLLQLKHKAELNEQVGLISLP[illegible]
[illegible]SAEPTEVCAALRKGFIACHRAMWKKLPEWPKTMTGLPSTSGTTASVVIIRGSKMYVAHVGDSGVVLGVQD
[illegible]QQRTHRHELFTPSAIGAHPDDSRSKFQLKTIETLLGNTAKVGEVIVLGMITQLKEGKFFLEDPTGVVQLDTL
[illegible]HKVEKVYVPALIFGQLLTSSNYDDDEKKVTGGRNGYGAKLCNIFSTKFTVETACREYKKLFKQTWTDNMGK
[illegible]DRDIEEAIQEYLKKKSKSIQPLPSKAECSDRTDVDKRVKRELSRNKVTSNLLPMKFKAEMVTEGFISDHVGIC
[illegible]ILKFVRNVPWEFGEIVPDYVLGQSTCALFLSLRYHNLNPNYIHARLQLLGKTYAVQVLLVQVDVKDPHQAL[illegible]
[illegible]CQFGFELIGEKSIVCQENNQWSANIPICIFPCLSNFTAPMGTVLSPDYPEGYGNNLNCIWTIISDPGSRIH
[illegible]PRTVGKYFNPCYATAQLKPTFISKMNTAVQLILVAASLAAPVFDYVDSIYLQTLWCITAFTTAASAYSYYHYGR
[illegible]AKETMEQIQQSEISKQLNTLFQDNLQNVNSYAEDLQKKLIPFTTELHAKLSQDSEKLKEQIRQELEDLRLKL[illegible]
[illegible]VQLYSEKDEPMNILNIKRGIISALMVPMETEDNLKTISMDTIYGKCDSEVEVKDRRGNTASVISINRNLKTCG
[illegible]EQELRTLSETFDSVFLHPGSYRGARLAGGCVLRLLEKVQDGELRNGLALVRPPGHHAHRDKMDGYCMF
[illegible]RFSPDDQAVAVCAGNRIYMLNAKNEAILTELEGHLAPVTAAEFCTWERNMLISVSEDRSFKVWDYCTGLL[illegible]
[illegible]MKNPRFPGNLQMSDRQLDEAGENDVNNFFQLTVEMFDYLECELNLFQTVFSSLDMSRSVSVTAAGQC[illegible]
[illegible]HMAIAWVKRNIKAFGGDPENITIFGESAGAVSVSLQILSPYNKGLIKRAISQSGVGLCTWAIQENPLFWATK[illegible]
[illegible]HFVPALEHHYREEILAKFLYWLMDTYIVQLLRSFFYITETMFQKNMLFFYRKLIWSKLQNIGMRNHYAKVHL
[illegible]SPAPVVSWLYRNEEVTTIADNRFAMLSNNNLQILNINKSDEGIYRCEGRVEARGEIDFRDITVIVNVPPAITF
[illegible]TSPYTTAVVAVDRLQKCLHSIIKHLDHIQSC
[illegible]VILEAVAQSLQLEESQLKWSVDSIFQKDLLSTLYLLIAIAKHFQPDLPIPSNISVEVIIIERTLTGLRTENAVEYITE
[illegible]NRPQPNMEEIEENFQGNLCRCTGYRPILQGYRTFAKDWSSCGKGINGPGCCMAGKDGGCCGRKVKD
[illegible]QETDRDYQRILLYSRTPHLADDCIEDFQSQAYCLDMEEFLLIPRSQDQPGSSSG
[illegible]EFDGLDLNWEYPGARGSPPEDKQHFTALVQEMAEEFKEEAKRTGKERLLLTAAVAAGKANIDGAYEIKNISK
[illegible]FEAALNSKEAALATALGDKKSLEGELEDLRDQIAQLEASLAAAKKQLVDETLMKVDLENRCQSLTEDLEFRK[illegible]
[illegible]QWDPVHRLLLAEACRKLDEFDAVHNSPSQTTKLMKEEFQSQVELLNSLEKKYNDPGPVYDCLVWHDGET[illegible]
[illegible]TVPKEDLDLPNHLMGLKRNYIKLSFNTVDDLVKVRREISPAVKKNRERDQASDTYTSMLSSALIGGSLTMDD
[illegible]KYTAPHILIATGGRPSITRDSEIPGASLGITSDGFFELEELPRRSVVVGAGYIAVEIAGILSTLGSKTSLLIRYDKVL
[illegible]VKVSPTKSGSEPHPNLLVCSVTGFYPGGIEIKWLKNGLEQTAGVVSTELLQNGDWTFQILVMLEMSPRRGD
[illegible]LISFLWIIPILGWRTFTNGREQQVEEGTCETEFSTVTWFKVLTAIVNFYVPSFLMLWFYAKIYKAVRKHCQHR[illegible]
[illegible]RGADAKREEEETGDREKEDVITAAKKERSWDCAQAAPSKGRDVQQMLADSKAQTRKYNFQQGKDLARR[illegible]
[illegible]NMLNSSSPLSNTTMCTGNLDSVAANNSGSLSSTGSDYESCTGITSDGGSYSNSIISDSSSCTLSTGDDICS[illegible]
[illegible]AQKGEKGSSGKDGTPGIPGGVGAPGAKGDKGEAGLRGPLGEDGSKGDRGLAGPKGEPGMKGDTGPT[illegible]
[illegible]ICYFLYQILRGLKYIHSANVLHRDLKPSNLLLNTTCDLKICDFGLARVADPDHDHTGFLTEYVATRWYRAPE
[illegible]VITCEPPRINNGNLMTPMKTYRENERLQFTCNAGYTFGERSDAECTEYGWRPKPFCKVKNVNRNETEKC[illegible]
[illegible]SMVVHEKIEMFERLLRSIYTPQNIYCVHVDLKSPDEFKEAVRAIASCFENVFVATKLERVVYASWSRVQADL[illegible]
[illegible]TRIKFLTDGMLLREAIGDPLLRKYSVVILDEAHERTIHTDVLFGVVKAALKKRKDLGKFPLKVVIMSATMDVD[illegible]
[illegible]PGLHHNNNQFFHDAEVEEAIPNAKRSPSTATLNLPNIEAYRDPSCLSPASSLSSRSCNSEASSYESNYSYPY
[illegible]QVENPGGKKQVSYRTEIVGGVPIITPTQREELNECGEGGDRNYLKRSQSHLPYFSAKPPPDNAVIKAGYCV
[illegible]DEMNHEFQALALESRGMGELLPAKKFWEPDDSAKDGQKGIFLGDEWRETAWGTPHHSMSQPIMVQR[illegible]
[illegible]RLSAEEGSFISLNCSYFVMDDSRRPNNFTWCKMVTATRCQPVVSVEIDQIVNTQGRTRIKIDRWNRVIIVT
[illegible]TPGLYLVYCNLHFHLTNFSSNTIFLKMELLVNNITNRQTLFTPCSSTNRQTKFKQCASVTDNDRIFQELSILH
[illegible]RLMVLKKQLDIELKVKQGAENMIQMYSNGSSKDRKLLATAQQMLQDSKTKIEVIRMQILQAVQTNELAFDN
[illegible]MYTKSGEWKKISCEDLPRTGTLCKTAFAYEKKYLPDKTALTTTLVIIFTIVVTVSAALFWFLYKRNMSSGALCIT[illegible]
[illegible]PQKQKFSGLDLLLAFNVGIEVQGRLMHFSSEASYVPKRFHPPSVVGTIGSAAATAKLLALDQFKCKAALAVA[illegible]
[illegible]CEVSYCDRCLRATHPNKKPFTSHRLVEPVPDTHFRGLTCLEHENEKVNMYCVVDDQLICALCKLVGRHR[illegible]

GVGSSGYGGAQWDRGANLKDYYSRKEEQEAQATLESEEVDLNAGLHGNWTLENAKARLNQFFQKEKVQ
VDRCALVTRPIWCQNHRTTRLAWGLSGAAWLLALLMTLPTLIFRTTHTEKNSNKTMCVLEYARVANYQTTVEV
CPHLDHFRDEKNTLPIHWYKECNLIDNERFLSWDDDLIINSANVNDSGNYSCKMTYNYMEKEYNISRNIY
RSGSQKDKKLTSSAAHQSANSSVGTPKRSSQTTSKPSIQHKPLLLPAVPEAQASLGIQAKAIIIQALPALLPLTKQ
NVTYKGQRNTAKLSPEKLLCQLRGRVNVTMIHGSDGPFNTSEWQMYLPRKNLSQEVGHFRQCAVVSSAG
TASINSSDAWRRSNSSDDICEYSYREIYTRDGNSYGRPCEFPFLVNKTWYHDCIRDETHTDGDWCATTSS
QVSIETDRASKEFIEFLRTCHKSGQEIYKQCKIFLEAMHHKRELSIEEQSECAQDFYQNVAEKLQTRSKVSPE
ISLACGTCKIRFLQPWAQQGLTPKQWQDLELVIHHYLFNFNRIVNRIVQQHGMGYPFVTQVSLGCEKHPN
FSLNFQRLSVAEQNREQEILTILGQLRQGESCTGRELARKLKTQKKEVNRYLYKLLREGKLHKEGETPPFWR
FGATLADSDPQPVLLDTTFLTIPTQSHVTCEQICNVLLKNKGILDVKISSDQKTAVVTFISSIINGNQIIQMVPGT
KIAVAKEAIKSLSRIAQTQDGLEALFASSFLVDMKNVMATSDVVRYRVYELVVEISSVSADSLNYCANSGLISEL
RGHLLVEADDVRAWSCPQHNAYRLYHCEAEQVAVCQFCCYYSGAHQGHSVCDVEIRRNEIRKMLMKQC
PDAYIATVKKTISEDSGCDSVTDTEAEDEKNPVYSRKLAMSQHHETSESTVGDLVVQPDRIHCGVQTTLYIIM
AQILQKNKNLRLSILVSRLFMWEEPEMQAALKKLKTSGCKLKIMKPQDFEYIWQNFVEQEEGEAKAFEPWEI
FDLKPENIMLLDRNVPKPRIKIIDFGLAHKIDFGNEFKNIFGTPEFVAPEIVNYEPLGLEADMWSIGVITYILLSG
VCGMQEMLDSVPEKRFPVFQAHGYFCQLIDGLEYLHSQGIVHKDIKPGNLLLTTNGTLKISDLGVAEALHP
AGLMNYEQFTELKNILSHSGKRKAHESLCDGPSHKRPATFHSQLLTPPQTPTSMDNMEEAHKNDPKHDNC
SIKEKHRKPRKKKAPSLEVNEEMPSKNSTLASQTSIYLNSDQLAAENKPTEEAIHDDPIRLFGLIHAT
RYRIDEDENRYPQKLAFAECLCAGCIDVKTGRETSSLNSVPMHQTMMVLHRKPCPHNASPGTFAFEVDYIK
AYTKTNHNFLTLEDTLLGYMADGVSWCGNPSSPGINYESCPKWSECESNPSSVYWKMASKMFAEAACGV
KRLEVDIDIKIRSCKGSCAKSFSYQVDTESYENIQKQLTQASSINLQSELETSSLKVLKIRPLKDSTVPEHYKTLP
CLEASVVLNLFRLLNEDLKCTAYMESCT
LTSQDEDMHVELTDFSLLVIDECHHTHKDGVYNQIMEDYLERKLRGQRKLPQILGLTASPGTGGATSVEGA
PPDSLTLSWYWGDQRLESPDPEGAADDEELFSYVSKLEVPGEKVTEGMEFRCEVELLLPSDRSFHRATAVTV
VHALESAKNAKATRALKFSKSLNDMDQKAQSITEGSNYGERTCSESKLTPTQEQCLRINRFNQKEKKSLNLRI
PAVKDCCSDVLTKLSSGWRFYGGNLYYFSQEKKSFDEAEQFCVSQDSHLASISSEEEQEFLSSEIQGQDH
HRDIKGANILLTLFGDVKLADFGVSAELTASVAKRKSFIGTPYWMAPEVAAVEKKGGYNQLCDIWAIGITAIEL
LLVEAWDYSNDTANPDSLIEKASHSGMINPSRQWQTLKQNAGIAHFEYQIRVTCDEHYYGFGCNKFCRPR
LISILVIAGVFWIHRLIKFIYNICCYWEIHSFYINALKIPMSNLPYYTWQEVQARIVQIQKEHQICIHKKELTELD
SVDDEDIYKGLPDLLDESGGDEDEELYDCVYGEDEGGEVYEDLMKDEAAHQPKCTENDIRSCCLAEIKQT
RLQALVRDGQLQEVKEKEILNEALNELKEENKVLKEQKASVTKKKEYYKCEISRLNKALLDALQKQHPSAFVPH
CQLLSFAFGLQSPSAVEMSEICEKNNRNVAHGLAWSYYVGYLKLVLPRLKDLISEFNRANNNLLKCKETWKL
IVSQKTNIQADLDTRYRCFSTNRINMSIVTITLSNVTLQAYLMNHTLSGNETVCAADKTPTTPVATTPTSQTPTT
QLYQCAMKKGSCQAIPLPASAQMAHLGMSLASSDAEAQMIACGSGLSRTCDANQYLSGVCYLFKGNLEQ
ECIDGQARYTCVCKPGWQGERCEEDINECEDPANKNGGCSHKCTNFPGSYRCSCDSGYYMLPNKQDC
AEYMQKISELETECQELRSKLQELERANQTLKDEYDALQITFTALEEKLRKTTEDNQELVTRWMAEKAQEANR
DEMVIVKDIDMFSLCEHHLVPFVGKVHIGYLPNKQVLGLSKLARIVEIYSRRLQVQERLTKQIAVAITEALQPTG
QQRMRPGTVCSVAGWGQTSAHNNLSSDTLREVEVVVMQDAECLREPDGIYRNYNASTMMCVGDPEMG
NVLDDRQSQMKGVLDSNSFMIISVNKTLQAYTGYINNLQQDTSNIQTNLQSQMHSHNVVIMNLNNLNLT
KIPDVCGPKATCINTYGSYLCECQAGYVSSNRNKTLCQELTCPPLLGDDNSAGAKNLLGSFQAQVGHLCQ
NYLGGFYCSCRVGYVLHENQRTCAAQCENNAFTKRSGVITSPNYPKPYPKLSTCSYTIRVEDGFMIILEFVET
TCSYINLENFSLVSVGRGQPLLQDGKGQCPFDPRHKYTAILVDGELYTGTMNNFQGNEPIISRTLGSRTLLKT
GSTLESMLGDLTQELQDLGITAVPKGDCASCRKPIAGKMITALGKTWHPEHFTCTQCRQEIGSSTFYERGGL
GEFYECEDDHIYTLKEIVEWKIPKSRNRIVKFTNMLHTWDSSNPLPENFDGCLILAPIYEVQGVMKFRRDIVH
RNYVTRTGQAMRQSLRSFQVTTCTWHGGKPLDNCQYRATRDSRMIVIACDEHEHPVHFAESQI

QETTLTVINVYCPRADPQRPERAAYQASFYRLLQGRAEALLRDGGHVVILGDINTSHRPIDHCEPGDLEAF
IQSKRRSESIEVDVLPAYDALGQVTPGNKPSPQVYEDLIQAGARPGEFCTSFTELQRDFVKRRPAKLKNLLRL
WVDSQQKRYVPVKGDHVIGIVTLKAGDIFKVDVGGSEQASLSYLAFEGATKRNRPNVQVGDLVYGQFVVA
VMGGWSAWGPWSSCSVTCAEGVRKRIRSCTEPVPICGGSCPSPSLSTDTLPCDTNKICPTHGNWGSW
QLQQLVLVPGHHLQPPAQFLLPQAQQGQQGLLPTPNLFQLPQQSPGGLLPAPSRAGLPAQPVTRPGLPES
KEMKDKSYMPRDFKPEYAIAILAYTSQGLLNKQPLHQEFNAAVREAGRTRDFYLNNFNFKTLHFLLTRALH
DNGRNELSLLIEKGHSISCPGIHCYNHLQRSPVTWYKNGIRIMFKEIRPSLKLKDDKINLSSTYEQDAGIYV
PCMFGDVLAPPVGTVQISLLPAPETQTVPGKPVTPIKAESILSRKPPIQPKPVVVATVPVQPAATPSPTILLQTVP
DVQCQGTEAALGNCLYTDPEYLCQHDEDAGARCQAVELPTPPVRLVGNNKRCSGQVEIYYDGRWGAV
YWVDSEKGLLQRVYLNGTKQERLRYVEKGVAGFAINWINQDILWANHQKATIEATDLNGNNSRVLLKDV
SRARNDTRANANGLESYEDAGSSESRTVNGSDSPLLTNGVFKPARPPRPSRPPPPTPRRPSSAAGCVNG
CFPSRIRYPNDTGTLTSGRIVCPTIDNYAKATIVKWYKDCKPLQGPREKYSMGERYLFIKRPQKTDEGHYTC
RSPLGLGGPWFINSIQAPARGYGVSAVLKSEEAAMSLQPAVTMATAVLSVFSHTPHLDSRLGRDVLLDCGF
VKVRVYSLNEELRPAQRETVLTFVDPEGVKVDIIEERDFTGILSFPDFKIPSNPKYGVWKIEAKYKKSFTTSAVA
EEWQKIRSILGKHMLKPKEVESYTGTLNSVVSDLIRRLQHQRSCHQQHVVKDVAGEFYKFGLEGISSVLFD
LTAAHCNGDKITVYLGAHNIKQQEQSQQKISVRRQIPHCKYNRKIFNNDIMLLQLKHKAELNEQVGLISLP
LNLTQLYLNDAFLEFLPANFGRLTKLQILELRENQLKMLPKTMNRLTQLERLDLGSNEFTEVPEVLEQLSGLK
FFDDPPTRGWVSIKYLKPYKGSAVRETQRGGMFYSSKPEIKRAMEMADDAMRKDKTKRLELAVCNEPSDT
EKHSWYHGPVSRNAAEYLLSSGINGSFLVRESESSPGQRSISLRYEGRVYHYRINTASDGKLYVSSESRFNT
QSQQTQAHKVSSYGHSRSSEDVSVHATSKPKFKKGFSLRNMSLCVVDGMKEMWHRKSSPEPGMEATQG
GLQLSVVTRDGLSVNPRAAGLIGRNGPYDKQPFMVAFFKVSEVHVRTTRSATSRRRQQNRNRSTQSQDVS
ASDCINAMVEGSELRKFRSNSRVYHRYFLLDADMQSLHWEPSKKDSEKAKIDIKSVKEVRTGKNTDIFRSN
QNVTIPVALIRYYDIMDMQQALGARINVTLYSPSLPEFDYSMVVIFLIAVLTVGLGGYWSGVSELEDLKAVTN
IKKARTSTAGPHPYTPEHTSAAANASDTSSAITPQWPLVLSSEAPEDVSVQTMLTSIHQQMEPESTRAGVHVG
KPVIISTPTPTIMRPGSLPLHLGYDPLHPTLPSPTSVITQAPPSNRQLGSPTGSLPLVMHLANGQTMPVLPGP
TWGFYQDSYDGRDFLTFHKETMTWVAADIGAQITKRRWDTEIGDYQHWKRYLEERCIPWLRSSLEYGKE
PDPAFNLPLPPPPPEPLTQLVVKVHMNDNSTKSLMVDERQVARDVLDNLFEKTHCDCNVDWCLYEIYPEL
SCCEMNVHKRCVQSVPSLCGVDHTERRGRLQLRIETLGGEEIRVFVGEARNLIPMDPNGLSDPYVKLKLI
YILAQFLGAFVGAAAVFGIYYDAFMNYSGGELIVTGSNATAHIFATYPAPHLSLISGFADQVMSTALLVLVVFA
NGKWDCEQHACLIDGEMIDAINRGNYGWRAANYSHFWGMTLDEGIRYRLGTIKPPPVVMHMNELQM
THPNGTVSVISKLYVNSSTSLAGQLLTCRVRHMERDRDYSVKVKEGQ
THTAGGAVEPNGSILHISLSPRDAHLNVTCTAQNPASNSATTASAKDLCAETLGSPSSATEGSQARPASQRS
FLGSEPIISRNSHQSPLRTEYAIPWLNEPSFVFADVIRADRNSTEEDDDKVYFFFTEMSVEYEFVGKLMIPRI
AEQETVTQLAPQKIALWLRPGEQHSFSVRFKRAEGYPVDVYYLMDLSYSMKDDLENVKKLGNDLLAALRN
FMVTDCDNICKTNQSICYSFYPVLAQDYVNVREEDALVEDKLDFHSMQDIKSVSADKIKNITCIYFYDGCC
LGWLSFFLYLGTGFLCQCDPPVEVPETLIAWTGACLSIWCRYKSCVVNPIKTNKLIISSLTWYLNPVYDSEK

VEPPRETLEAVLLTSSLSVLLLPYDEMKTAQQMTTSRAASGHSRAFTQLKPFAPDGIILTTSRDLLLYPTNTDIYL
IPGSAMETGLLPNHKLKNVGGDHPAAPPHHHAHHPHHHAHHHHAHHQHHHHHALQQQLNQFQQQ
AIQLQLHAKTGQSVDSGAIKAKLSVLSVDELERELEANKKEKVKEAQLEAEVKLLRKENEALRRHIAVLQAEV
AFTEYVFRVVWIITSQMQLYSPSSPSYRTHAFGVPVTAPIIKDIQSSGPDTVEISWSPPLFPSGLIVGYNLLLTS
TKVDSSVSRLNKKYDVLSALYHKFERTCALIYMIQASSQTSAELSLPLVLKTSWITFLLAKGKVLQVEDDLVISF
YEEQSQDRRNLTKLSLIFSHMLAEIKGIFPNGQFQGDNFRITKADAAEFWRKFFGDKTIVPWKVFRQCLH
SADHMKTQDSQHTSHSVTPSTASAPTSSTPSHGLTATSEPASGQKTPSKVVYVFSTDMANKAAEAVLKGQA
KYTLAVQSENRNYTIMFTVLVKSKPRKPYFVKGENLGIINCTSEGYPTPDIEWLFCRTADDSCINTNRKDSK

SAAPQKKDSMGELSRATIRLLEELDRERCFLLSEIEKEEKEKLWYYTQLQSLSTRLDELPHMETFSMQMDLIR
NNQMNRNDRTLIFLDENEMDMTASHTAVITRNLKSNEEIDKPRKIDIKSFLAGLTSKNEGPKMNKEFRFFC
PFPVPSPETKVEVTVQADSSLVPSKPFLLPSLHNRVLDFSLLSSAAPSFPPPTLEPEVREATTLSQAQTTETNLQI
ETEKEPEHNYQGTEKREESFSNDAIYAVPYKNKKVKLPVSSSVDLDENEISSHEAAFPILVDQSNIQVEEDEP
VDAPGQQDGAASFPAGYMYPSMDQLAEMLPGILKQFGLKSVIGMGTGAGAYILTRFALNNPELVEGLVLINI
GFYCSFEEGDCGWTQSSPTSLTFPWQIGNLEYSHFPLLEGSALLLSTSKVPVAETIIVTSAVFPAPIRNSPCELF
SVSHINLTVREGDNAVITCNGSGAPLPDVDWTVADLHSITTHQTNLNWTNVHAINLTLVNVTSEDNGFLLTG
SEGSGAPGEEGTGHCPSSSTDTASEHSADEDAKGAGGELWGTAPGPSDYDLRLLRSQRVLARRQGIFQF
QRRKLVLSSAPATPHQPEPAALEEEGEGTSGQRGTGSKRQIRYLENLLRVYMGEIRRLQERELDLAELDSEDS
EYISKMGYLDPRNEQIVRVIKQTSPGKGIVPNNVTRRPSFEGSNESFPSYHQISNTAYEGTNFGTEGANMLS
CPPGWSGKTCLQADPCASNPCANGGHCVPFDAHYVCKCTAGFHGANCKQDVNECNTSPPICKNGGS
FNTMFTNCYIYNKPGDDIVLMAEALEKLFLQKISEMTEEETEIVIVQAKGRGRGRKEAATPKPGASTVQNVTQA
SLDSPLSSELADAKNPAQESQIQENEVAKAEAESSPYNNESQIKSDHSLCPLGSEGQSPETIVEDNPSSSDT
SRSRSPKRRSPSPRSRSHSRNSDKSSSDRSRRSSSSRSSSNHSRVESSKRRSIKEKKSSSKDTRASQAVGD
KKHLLSVTAKPLSAAVAPPPSLASVAPAEIPAAAELPLTSPPLLSQTASPPPPVLTAVAFPQPPIAVSVTAAPLLLPPV
HLIDLGADTSLRSRWTNMNALHYAAYFDVPELIRVILKNAKPKDVDATCSDFDFGTALHIAAFNLCTAAVKCL
VPGSCYHVGISALAGPYSSDPQNVTGCTVPAAPANVSLTNSGSSSVLHAAWDEPPGGRDHYRVILYSLVPW
EGKKLADSWGAVFLESSAKENQMTRGIFTKIIEEIDRVDNSYCTERSCLLILLAVSIPPGQLSGFGAVFQNQLI
EPAAELGRAGPAGGSAGRGPHEPISAVLCSPSPPAERQGQAPRDAAVGKRTERVELRPVPGPLGQTRGKAE
TNVGIKCATITPDEKRVEEFKLKQMWKSPNGTIRNILGGTVFREAIICKNIPRLVSGWVKPIIIGRHAYGDQYR
TQTETSGTFGCHCQLSGKDEPSSYICTTCKQPFNSAWFLLQHAQNTHGFRIYLETSPSNSSLTPRITIPPPLG
SDEEVKVRSPTKSPTVKSSPRKPRRRPRSSPDRSSAAHSDSSSVCSPLSKSETMSMEKVKKKEIKSGEKRRGI
SSRSKRKPTVVTKYVGSDDEQTLPETVNEDVSNENSENDVDMKSLPKGTVVVQPEPVLNEDKDDFKGPEFF
KPEESSNLNFEADASFLRQAIMSFGSIKTMQTSETENASPWMSVQNCTLAQHEQKKLLSGTLGSPLSDWLL
QELGMRHLTEILKGGVHIEGNPQLCFQEMILWGDIFHKHNDLALHQQINSSRERTCADCRRFCSQGHCV
RQKRIQEQENPTLEANAGAQVADSMVRRSSLRNKRQKCESGKAVYIEFGSDSSDDLFKKAGGGGMGDTE
DTLNMQVDQFESEVESLSVQTRKKKGDKDKQDRIEGLKRHIEKHRYHIRMLETILRMLDNDSIQVDSIRKIK
KYGTELNQGDMKPPNYDSDEGNETEVQPQQNSQLIWKQGRQLLRQYLQEVGYTDTILDVKSKRVRALLGIS
PKKVTAQKRLEKRDKYEQETLTILEEKKVEAAERVSKQDSAQNELTALILCDTDIEPSIKCDSQDKKIGRPESSVS
GANGDVQQVGVIRCPICRQECRQIDLVDNYFVKDTSEAPSSSDEKSEQVCTSCEDNASAVGFCVECGEW
ITAASQPSNPSECNGLSPLVDKEGGGQMEIPDQLTGNTKKKSSKKDLINHTIQNTDVEWVKNAQKAFESKT
TCSQECAERHIIDNFFVKDTTEVPSSTVEKSNQVCTSCEDNAEANGFCVECVEWLCKTCIRAHQRVKFTKD
DRNTNRPLEPPSEFIVKVQDINDNPPEFLHENYHANVPERSNVGTSVIQVTASDADDPTYGNSAKLVYSILEG
KLSPLPTPQWPGRSTPESEGGGVEEDGGPPPFSPGAGGDPAPEQLDPELARTVADSVERLLQSCPRGGEPG
AFKGNDSTYITIFLCTYRAFATTKQVLDLLLNRYGKLNVQVNGDHARHAVDERLELKNTISSILGAWLDQYSE
ISENGPCLGFRKPKQPYQWLSYKEVSERAEALGSGLLQQGCKPSASQFIGVFAQNRPEWIISELACYTYSMV
QGEKIRDLEFCLEEHREKLNATEEMLQQELLSRTSLETQKLNLMAEISNLKLKLTAVEKDQLDYEDRYGDTESI
SHCATPLLGAEDSGTSQLGFGALELSQSQDVEGDSTLSEEGRGHQLQPAGTVPTCCNPKRNDHNEDLHIV
VTRGVCQRHIIDPDNPADIESEVHCMYSKQRDGESDSCPDCVVSTLGTKVLVTEKNRFVNFFVGNTVKSSF
SRSIEAAIEFISKMSYQDPRREQMAAAAARPVNAGMKPPAGGVQQSINRKQSWKGSKESLVPQRHGPPLGD
FPGTPDAEALACFFIPVLRSLTRLRPEMALEEGVPRAVQEWERSSNFERMIFYEMAEKFMEFEAEEELQIQKN
SRSGSGSSGFLEGLFGCLRPVWNIIGKAYSTDYKLQQQDTWEVPFEEISELQWLGSGAQGAVFLGKFRAEE
NPSKEELRPRTANKAPGNKGGKQKSPPTAAAVSGDSGPQRRTTCKKQPRRVERTSGGDALNCHTSEDLAL
LEIINAVPFIITIFWPALRNLFIPVFLNCWLAKHALENMINDLHRAIQRTQSAMFNQVLILISTLLCLIFTCICGIQ
SSAGWKNSIRHNLSLHSRFIRVQNEGTGKSSWWMINPDGGKGGKPPRRRAVSMDNSNKYTKSRGRAAK

IKLVYFLLSTLSLITCVLAVAFVAHHYSKIMQFTCDVVLDSCQCKLETSDPLSRTFVYQDVSDCTSVTGTFSLY
SLTKLVNGTEFFKIVLQESENSCSKTSTINVLFFGRLAGDCGKVVHQGDTMVVAGFNVAKSPSANEDNRH
FKFGPLFMRLCYELGLEVPAIELIKDQTLHGFFSDSTSFNILMDMLFKKGHYESALEVLGEMKKQHIKIIKETY
VACADRHHCCPRGSHCSADGRSCFMLPGLPAAGAVQCPDGESECPNESTCCMMPTGSWGCCPMP
ANGLINRLIEENKMDSLGMVVVDELHMLGDSHRGYLLELLLTKVRFVTEKIAKRQTKENGPVFDSIQIVGMS
KRMQPSVRSHACEVDPYGMPGGCSHICLLSSNYKTRTCRCRTGFVLGSDGRSCKRPKNELFLFYGKG
IERATRLVKSQQSAGKEYVGIIRLHNAIESEIQLSRAIETLTGALFQRPPLIAAVKRQLRVRTIYESKLVEYDPGR
VRKMVKQAAVIICIKKREFDKASKILKKNMGKDPSTQKLRTELLSIIREKNLSHTMIRNFSYKTFQQSAFQFLE
IRVVVSPEAVQMAECVQLQSGFSNIIGRIIILQKYMVCFQEETKAEECEFYLSVHQFIVLPLQRQRMEALNC
EAKALAEEAEVVDGPDENGEMFTRPGKLSDYFPKPYPNPEAARAANNGALPPDLSYIISARHGGEDYVFSLL
ETLAAETGLSVRVVQVWFQNQRAKMKKLARRHQQQQEQQNSQRLGQEVMSNRMEGMMTSYTPLAPP
KPAGPPAKRAIADLP
SKMIREDGYDSVFSVVRRHLFRWSEIKKGVNKVTTPQNLDPAKRPRRQDWDGELYENGSFYFAKKNLIEK
YLGGVIFINRKSTSSAKTVMSGVVQTMLDENVKVWVYPEGTRNGNGDLLPFKKGAFHIAVQAQVPIIPVVY
VRDKHLPVSDAFPHFLADSHHQVCMLAAKSIASLFQDMKQRDSSGLPKALPLKFQQKAFENVYLRVQEG
TEWLMHEGKCYWASKESRIWHKSHDDCSAKSSRLLVIQDQKEMDFISSVTQDTNPVWLGLTMTSPEWKV
TSSSPLDMQTSQMKVISKEIDGKINGIPRLEEEAQSFCQFVTSGESAHVRAKLAQLRRYWEEVRDHARCLE
KNCSMFKAKYICRQNLGTPVNPELPAPYPTPSLTGSCPQGWSSDPKLRHCYKVFNYDKLQDKKTWTLAQ
GKDTDDWMCIDFGSIVIHFMLPETREIYELEKLWTLRSYDDQLVQMVPESLPEDFIVGLTHDQQ
DSTPDSLFLNFLHTVKEDTLQYLVRYWEQSTNKQHETEWTKNTRIGLLNLKQLTVYCFQIRAKVKVFGDNI
GKLAPKGRLDMEQLAQRLAARFPGLFSPQRRFAFTSSSKHRCMESSAAFRKGLQLALHRQPPARDIENEE
EGLRVKLSPYVDEVQQKISKNLEVLRFQMIPFTQELLDQVMLKARELQQHLTPTRDMKVQFLEGIDEVQRF
DHSSLQRGLHGSSALFEVNSSPSSLKEDIKNSVNLNLTVQSAAVQSCLRPSVQDGLPWCPAQGRLRKKR
ISVEPERITEEQEEILSILEEKFPELLPREDIISVFQQTPFKAQGFNIEEAMLKDLKELSDGEIKIAVSTVYVTLEDC
AAARPDDILFRDEIIANGFHSCDSDEDDRASRASSSDWTPRPRIGPYTFVQQHLMIGTDPRTILKDLLPETI
LGTPELVTATECVHYFEWRTFIACKKGSFKAKKEVPCYVFDDLKKHDLNPLIKTSGGYLVDDSDEDSLYINV
TQSEAYVQDYFPLVEQYDFYSGVGFSFLFSWVLMITVVITFITGGNVEKLVCEPFEDKTLFKYTEDVSLKIEHI
AWTWPGALNIENSESESWTSHTVKKLTASFDASLTGEREPKVLSNDHKEALKRDCDDSEMRDEVYHRKIIS
MPKSIERLQCRRCVVVGNSNSIRGRGLGAVIDSHNIIIRLNNAPVKEHKEDVGEKTSIRLFFPESALPNPLD
TVVAVPYGSRQVRLVLKGPDHLYLETKTLQGVKSENSLSSTGSFLVDNSSIDFQKFPDKEILRISGPLTADFTV
NVYKSKKYGVSVSLSSAKQRQQFKGKDFLPLSHFLPMINTVPVESMDFVEYGDYSHTFESDVFSSPLETDS
VSDSKYGILKPSNLLNFQRHPELLQPYDPEKNNLNLDSYQSSVCNKSKSLADEGKIETKPWTSSSKVVTLQ
VSSTEILEDKPLSNTGSGQTVATQMSPQKIKLHLRPDDSQIFSVQVRQVEDYPVDIYYLMDLSNSMKDDLH
GGMEDLSKERSLTGKLEPVSPVSPAHPDSELDLLPSRLSKEELIQNMDRVDREITMVEQQISKLKKKQQLE
YSGGFTPNPTYKEVCTGLTGHAEVVRVVYDPQKTSYEALLKAFWENHDPTQGMQQEENVGTQYRSVIHT
SEHLRPHQESPAFRDHQLCPPLRDLQQRKCQEHNKLLEFYCRDHASCICCVCLVSHKACATGPVQEAK
CINENGYCEHYCIDSPTMIRQCFCAEGYILASDEVSCTPQVDYPCGKIHVLAKQSTSPQGRIVGGYVCPP
AVMVEVNCETDFVARNVKFQQLVQQVAAGTMFYHQGAKDHLTTYIKRFMKAAELSQLRTGPAGCLLSDQ
NVKSSRGLEGNVMIDLYEVAGIKGMFLANKKIDNQVKTFITYNKGRDWRLLQAPDTDLRGNPVHCLLPYC
HSNGEQASGISSSKNCYDIENHGSEDFLLVLDCCDLSVLKVEEVDVAAVPGIEKFTRSAKLTDAEELRNLR
NAGPKDMVVGFPNLGILHVTKKKVFETLEARMTDACKKGYNPGLLVHPELNYIQAEGGGERQLTEREREII
RAHRCVLAAGSPFFQDKLLLGYSDIEIPSVVSVQSVQKLIDFMYSGVLRVSQSEALQILTAASILQIKTVIDECT
AKCINPRVKSGRFMKILPDYEHMEYRDVYTCLLHRYRHILGLWQPDIGPYGGLLNVVVDGLFIIGWMYLP
GGRIVSSTTRSTMLHYRSGLLQTLDTLVFAGLFFLGFAEQKQMVEVELYSDYREDSYVPTVGAVIEIQTKRIQ
KSEDNGRPGLSSEKQSVHILPKDSSGKQAPASV

ELPLEPDGAARYMLPAVLRPRYTPHGWEGEDVTQGVPRVPQGKLPYTLSLSATLQSTHGIDRVLSNCSLSPL
NAMKHLPNFFHKGQLAKMFFLFPDPHFKKTKHKWRIISSTLLAEYAYVLRVGGLVYTITDVEEVHEWMVGH
IESGEVDLDDLDTPSENSNEFEWEDDLPKPKTTEVMRKGSLPEYTAAEERDDGRRWRMFRIGEQDHRVD
KKICQVKNCKNIPVKRSFFFLANWNEKINFPVQIKSLPNETILTIKLFGENSVSKNTEILAWTCSPLYQKQKFIH
KDGYRTAGQACELVHCPELQPPENGYFIQNICNNYFNAACGIRCKAGFDLVGSSIRLCQPNGLWSGSETT
QDKDAQFYRFAEGELTLEPGAWDAEELLEALAFLAQLGPDALLTMALRKPPAQRTEDELELIFEELLHIKAVAH
FRESFGHNADYSLGEALWACCNLFSDVQLKMSHKRIMLFTNEDNPHGHDSTKAKLTRTKAADLRDTGIFL
KIGPPGVHLESIDGVVKINISPPEENQNRKMWTNDISFTYNAVFWENSSNAEPKSKTINRRDTIYDLAPETTY
IFNNNPALWKEIRPFFTKALSGPGLMRMIAICVESTKDHLDRLENVTTGLGNINALNFMRQITLDTSNTLFLG
VHRNVPAAHLARARVCALSWGLDHEEFLRDYDFILGADIVYLKDTYPLLIRTLRHLCGPHSTIYLSSKMRQE
FTHPCDHEELREMLTHRNGPTKKGKELNTERSFFLRMKCTLTSRGRTVNIKSAAWKVLHCTGHVRVYDTC
VLNVHPSYLPRWRGPAPIVHAVLHGDTVTGVTIMQIMPKRFDVGPIIKQERYAVPSQCTAKELEAILSKLGANN
KRVKPGAVCSVAGWGSTSTDRKSSSTRLQEVDVVVMPDAACPRKPNGPYRFYDASTMMCVGDPKMGKD
TFAVCFPLRAKVVITKGKVKLVILVLWAVSFASAGPIFVLVGVEHENGTNPLDTNECRATEYAIKSGLLTIMVWTS
IMDPQPPSQIDGSSVDVHINMEQAPIQSEPRVRLVMAQHMLRDVQGILGRLEGRTNGQPQPTPENAPSAI
TPDPSTIISYLPGLLYKFSCSYPLEYLVNNTQLASSSAAISVKENNGTFISTLNLLLYNDSTYSQLLVIPSIGLPLKT
NVPGAVKDVCVTEHLNPEKAANCNQIPELCYSLGYDRQKAELCVAFLQAMHGSMIGDQDAGCDCYILGT
QMAPYACQRQTPLLHSNQLLTEALETLAEEAGFSGSEGRSLAFTRAASVLKALPDRLSTLEELGPLPGIGEHSP
PVPREVEAPGKQQPPSDTTGLFVMDEDSPSQDYEPFFDSDPESTDDGSLSEEAPAQPPPAPPNHQYAKSLP
PISTPSRVPLAPGGEDGTPAHSEELELPWWADAAPDPSGSLPQYTLAELPAEPSPEPLRQRDPSTQQALRKG

MVDKAIYLSGYRMRLGSSTDKKDSGRLHVDFAQARDDFYEWECKQRMLAREERHRRKMEEDRLRPPSP
MTKSFVCRPGSGGDLLPLAGCLGSRFRERHSMIQWLEVGARQIQTGNTIIF
PTGADALCALDILSSAGLLNHFGRSSSVEKIDISPILLHKGSTKIALYGLGSIPDERLYRMFVNKRVTMLRPKEI
LAAEYPSLVSKQEIGESYEKRPMYVLKFSTGGTNRPAIWLDAGIHSREWITQATALWTAEKIASEYGNNPSVTS
QQPQKQHPQHHPSQQPPTVFQQPGPPSQCPIGVQQQQMPASPQTQCQQPQQTAHHQLKAQPEMQSS
ENKIVILLDYMFQDLHNLKSLEVGDNDLVYISHRAFSGLLSLEQLTLEKCNLTAVPTEALSHLHNLISLHLRHL
KSPKEYYVQQEVIVLFCETVERALKLGYLTQDMIDDYEPALMFTIPRLAIVCGLVVYSEGPLNLDRKPEDMSEL
KPVIAAVNGYALGGGCELAMMCDIIYAGEKAQFGQPEILLGTIPGAGGTQRLTRAVGKSLAMEMVLTGDRISA
LTAAHCNGDKITVYLGAHNIKQQEQSQQKISVRRQIPHCKYNRKIFNNDIMLLQLKHKAELNEQVGLISLPP
CSAEPTEVCAALRKGFIACHRAMWKKLPEWPKTMTGLPSTSGTTASVVIIRGSKMYVAHVGDSGVVLGVQD
LQQRTHRHELFTPSAIGAHPDDSRSKFQLKTIETLLGNTAKVGEVIVLGMITQLKEGKFFLEDPTGVVQLDLTL
EHKVEKVYVPALIFGQLLTSSNYDDDEKKVTGGRNGYGAKLCNIFSTKFTVETACREYKKLFKQTWTDNMGK
YDRDIEEAIQEYLKKKSKSIQPLPSKAECSDRTDVDKRVKRELSRNKVTSNLLPMKFKAEMVTEGFISDHVGIC
PILKFVRNVPWEFGEIVPDYVLGQSTCALFLSLRYHNLNPNYIHARLQLLGKTYAVQVLLVQVDVKDPHQALK
ECQFGFELIGEKSIVCQENNQWSANIPICIFPCLSNFTAPMGTVLSPDYPEGYGNNLNCIWTIISDPGSRIH
PPRTVGKYFNPCYATAQLKPTFISKMNTAVQLILVAASLAAPVFDYVDSIYLQTLWCITAFTTAASAYSYYHYGR
NAKETMEQIQQSEISKQLNTLFQDNLQNVNSYAEDLQKKLIPFTTELHAKLSQDSEKLKEQIRQELEDLRLKLS
VQLYSEKDEPMNILNIKRGIISALMVPMETEDNLKTISMDTIYGKCDSEVEVKDRRGNTASVISINRNLKTCG
SEQELRTLSETFDSVFLHPGSYRGARLAGGCVLRLLEKVQDGELRNGLALVRPPGHHAHRDKMDGYCMF
VRFSPDDQAVAVCAGNRIYMLNAKNEAILTELEGHLAPVTAAEFCTWERNMLISVSEDRSFKVWDYCTGLLI
HMKNPRFPGNLQMSDRQLDEAGENDVNNFFQLTVEMFDYLECELNLFQTVFSSLDMSRSVSVTAAGQCP
QHMAIAWVKRNIKAFGGDPENITIFGESAGAVSVSLQILSPYNKGLIKRAISQSGVGLCTWAIQENPLFWATK
HFVPALEHHYREEILAKFLYWLMDTYIVQLLRSFFYITETMFQKNMLFFYRKLIWSKLQNIGMRNHYAKVHL
SSPAPVVSWLYRNEEVTTIADNRFAMLSNNNLQILNINKSDEGIYRCEGRVEARGEIDFRDITVIVNVPPAITF

TSPYTTAVVAVDRLQKCLHSIIKHLDHIQSC

AVILEAVAQSLQLEESQLKWSVDSIFQKDLLSTLYLLIAIAKHFQPDLPIPSNISVEVIIIERTLTGLRTENAVEYITE

NRPQPNMEEIEENFQGNLCRCTGYRPILQGYRTFAKDWSSCGKGINGPGCCMAGKDGGCCGRKVKD

VQETDRDYQRILLYSRTPHLADDCIEDFQSQAYCLDMEEFLLIPRSQDQPGSSSG

EFDGLDLNWEYPGARGSPPEDKQHFTALVQEMAEEFKEEAKRTGKERLLLTAAVAAGKANIDGAYEIKNISK

EFEAALNSKEAALATALGDKKSLEGELEDLRDQIAQLEASLAAAKKQLVDETLMKVDLENRCQSLTEDLEFRKN

QWDPVHRLLLAEACRKLDEFDAVHNSPSQTTKLMKEEFQSQVELLNSLEKKYNDPGPVYDCLVWHDGET

TVPKEDLDLPNHLMGLKRNYIKLSFNTVDDLVKVRREISPAVKKNRERDQASDTYTSMLSSALIGGSLTMDD

KYTAPHILIATGGRPSITRDSEIPGASLGITSDGFFELEELPRRSVVVGAGYIAVEIAGILSTLGSKTSLLIRYDKVL

VKVSPTKSGSEPHPNLLVCSVTGFYPGGIEIKWLKNGLEQTAGVVSTELLQNGDWTFQILVMLEMSPRRGD

LISFLWIIPILGWRTFTNGREQQVEEGTCETEFSTVTWFKVLTAIVNFYVPSFLMLWFYAKIYKAVRKHCQHRI

RGADAKREEEETGDREKEDVITAAKKERSWDCAQAAPSKGRDVQQMLADSKAQTRKYNFQQGKDLARR

NMLNSSSPLSNTTMCTGNLDSVAANNSGSLSSTGSDYESCTGITSDGGSYSNSIISDSSSCTLSTGDDICS

AQKGEKGSSGKDGTPGIPGGVGAPGAKGDKGEAGLRGPLGEDGSKGDRGLAGPKGEPGMKGDTGPTG

ICYFLYQILRGLKYIHSANVLHRDLKPSNLLLNTTCDLKICDFGLARVADPDHDHTGFLTEYVATRWYRAPE

VITCEPPRINNGNLMTPMKTYRENERLQFTCNAGYTFGERSDAECTEYGWRPKPFCKVKNVNRNETEKCS

YSMVVHEKIEMFERLLRSIYTPQNIYCVHVDLKSPDEFKEAVRAIASCFENVFVATKLERVVYASWSRVQADLI

ETRIKFLTDGMLLREAIGDPLLRKYSVVILDEAHERTIHTDVLFGVVKAALKKRKDLGKFPLKVVIMSATMDVDQ

PGLHHNNNQFFHDAEVEEAIPNAKRSPSTATLNLPNIEAYRDPSCLSPASSLSSRSCNSEASSYESNYSYPY

QVENPGGKKQVSYRTEIVGGVPIITPTQREELNECGEGGDRNYLKRSQSHLPYFSAKPPPDNAVIKAGYCV

VDEMNHEFQALALESRGMGELLPAKKFWEPDDSAKDGQKGIFLGDEWRETAWGTPHHSMSQPIMVQRK

RLSAEEGSFISLNCSYFVMDDSRRPNNFTWCKMVTATRCQPVVSVEIDQIVNTQGRTRIKIDRWNRVIIVT

TPGLYLVYCNLHFHLTNFSSNTIFLKMELLVNNITNRQTLFTPCSSTNRQTKFKQCASVTDNDRIFQELSILH

RLMVLKKQLDIELKVKQGAENMIQMYSNGSSKDRKLLATAQQMLQDSKTKIEVIRMQILQAVQTNELAFDN

MYTKSGEWKKISCEDLPRTGTLCKTAFAYEKKYLPDKTALTTTLVIIFTIVVTVSAALFWFLYKRNMSSGALCITH

PQKQKFSGLDLLLAFNVGIEVQGRLMHFSSEASYVPKRFHPPSVVGTIGSAAATAKLLALDQFKCKAALAVA

CEVSYCDRCLRATHPNKKPFTSHRLVEPVPDTHFRGLTCLEHENEKVNMYCVVDDQLICALCKLVGRHRI

GVGSSGYGGAQWDRGANLKDYYSRKEEQEAQATLESEEVDLNAGLHGNWTLENAKARLNQFFQKEKVQ

VDRCALVTRPIWCQNHRTTRLAWGLSGAAWLLALLMTLPTLIFRTTHTEKNSNKTMCVLEYARVANYQTTVE

PHLDHFRDEKNTLPIHWYKECNLIDNERFLSWDDDLIINSANVNDSGNYSCKMTYNYMEKEYNISRNIY

RSGSQKDKKLTSSAAHQSANSSVGTPKRSSQTTSKPSIQHKPLLLPAVPEAQASLGIQAKAIIIQALPALLPLTKQ

NVTYKGQRNTAKLSPEKLLCQLRGRVNVTMIHGSDGPFNTSEWQMYLPRKNLSQEVGHFRQCAVVSSAG

ASINSSDAWRRSNSSDDICEYSYREIYTRDGNSYGRPCEFPFLVNKTWYHDCIRDETHTDGDWCATTSS

QVSIETDRASKEFIEFLRTCHKSGQEIYKQCKIFLEAMHHKRELSIEEQSECAQDFYQNVAEKLQTRSKVSPE

ISLACGTCKIRFLQPWAQQGLTPKQWQDLELVIHHYLFNFNRIVNRIVQQHGMGYPFVTQVSLGCEKHPN

FSLNFQRLSVAEQNREQEILTILGQLRQGESCTGRELARKLKTQKKEVNRYLYKLLREGKLHKEGETPPFWR

FGATLADSDPQPVLLDTTFLTIPTQSHVTCEQICNVLLKNKGILDVKISSDQKTAVVTFISSIINGNQIIQMVPGT

KIAVAKEAIKSLSRIAQTQDGLEALFASSFLVDMKNVMATSDVVRYRVYELVVEISSVSADSLNYCANSGLISEL

RGHLLVEADDVRAWSCPQHNAYRLYHCEAEQVAVCQFCCYYSGAHQGHSVCDVEIRRNEIRKMLMKQ

PDAYIATVKKTISEDSGCDSVTDTEAEDEKNPVYSRKLAMSQHHETSESTVGDLVVQPDRIHCGVQTTLYIIM

AQILQKNKNLRLSILVSRLFMWEEPEMQAALKKLKTSGCKLKIMKPQDFEYIWQNFVEQEEGEAKAFEPWE

FDLKPENIMLLDRNVPKPRIKIIDFGLAHKIDFGNEFKNIFGTPEFVAPEIVNYEPLGLEADMWSIGVITYILLS

VCGMQEMLDSVPEKRFPVFQAHGYFCQLIDGLEYLHSQGIVHKDIKPGNLLLTTNGTLKISDLGVAEALHP

GLMNYEQFTELKNILSHSGKRKAHESLCDGPSHKRPATFHSQLLTPPQTPTSMDNMEEAHKNDPKHDN

IKEKHRKPRKKKAPSLEVNEEMPSKNSTLASQTSIYLNSDQLAAENKPTEEAIHDDPIRLFGLIHAT

RYRIDEDENRYPQKLAFAECLCAGCIDVKTGRETSSLNSVPMHQTMMVLHRKPCPHNASPGTFAFEVDYI
YTKTNHNFLTLEDTLLGYMADGVSWCGNPSSPGINYESCPKWSECESNPSSVYWKMASKMFAEAACG
KRLEVDIDIKIRSCKGSCAKSFSYQVDTESYENIQKQLTQASSINLQSELETSSLKVLKIRPLKDSTVPEHYKTL
LEASVVLNLFRLLNEDLKCTAYMESCT
TSQDEDMHVELTDFSLLVIDECHHTHKDGVYNQIMEDYLERKLRGQRKLPQILGLTASPGTGGATSVEGA
PPDSLTLSWYWGDQRLESPDPEGAADDEELFSYVSKLEVPGEKVTEGMEFRCEVELLLPSDRSFHRATAVT
HALESAKNAKATRALKFSKSLNDMDQKAQSITEGSNYGERTCSESKLTPTQEQCLRINRFNQKEKKSLNLR
PAVKDCCSDVLTKLSSGWRFYGGNLYYFSQEKKSFDEAEQFCVSQDSHLASISSEEEQEFLSSEIQGQDH
HRDIKGANILLTLFGDVKLADFGVSAELTASVAKRKSFIGTPYWMAPEVAAVEKKGGYNQLCDIWAIGITAIEL
LVEAWDYSNDTANPDSLIEKASHSGMINPSRQWQTLKQNAGIAHFEYQIRVTCDEHYYGFGCNKFCRP
LISILVIAGVFWIHRLIKFIYNICCYWEIHSFYINALKIPMSNLPYYTWQEVQARIVQIQKEHQICIHKKELTELD
SVDDEDIYKGLPDLLDESGGDEDEELYDCVYGEDEGGEVYEDLMKDEAAHQPKCTENDIRSCCLAEIKQ
RLQALVRDGQLQEVKEKEILNEALNELKEENKVLKEQKASVTKKKEYYKCEISRLNKALLDALQKQHPSAFVP
QLLSFAFGLQSPSAVEMSEICEKNNRNVAHGLAWSYYVGYLKLVLPRLKDLISEFNRANNNLLKCKETWK
IVSQKTNIQADLDTRYRCFSTNRINMSIVTITLSNVTLQAYLMNHTLSGNETVCAADKTPTTPVATTPTSQTPTT
QLYQCAMKKGSCQAIPLPASAQMAHLGMSLASSDAEAQMIACGSGLSRTCDANQYLSGVCYLFKGNLE
CIDGQARYTCVCKPGWQGERCEEDINECEDPANKNGGCSHKCTNFPGSYRCSCDSGYYMLPNKQD
EYMQKISELETECQELRSKLQELERANQTLKDEYDALQITFTALEEKLRKTTEDNQELVTRWMAEKAQEAN
EMVIVKDIDMFSLCEHHLVPFVGKVHIGYLPNKQVLGLSKLARIVEIYSRRLQVQERLTKQIAVAITEALQPTG
QRMRPGTVCSVAGWGQTSAHNNLSSDTLREVEVVVMQDAECLREPDGIYRNYNASTMMCVGDPEMG
NVLDDRQSQMKGVLDSNSFMIISVNKTLQAYTGYINNLQQDTSNIQTNLQSQMHSHNVVIMNLNNLNLT
IPDVCGPKATCINTYGSYLCECQAGYVSSNRNKTLCQELTCPPLLGDDNSAGAKNLLGSFQAQVGHLCQ
YLGGFYCSCRVGYVLHENQRTCAAQCENNAFTKRSGVITSPNYPKPYPKLSTCSYTIRVEDGFMIILEFVET
CSYINLENFSLVSVGRGQPLLQDGKGQCPFDPRHKYTAILVDGELYTGTMNNFQGNEPIISRTLGSRTLLK
STLESMLGDLTQELQDLGITAVPKGDCASCRKPIAGKMITALGKTWHPEHFTCTQCRQEIGSSTFYERGGL
EFYECEDDHIYTLKEIVEWKIPKSRNRIVKFTNMLHTWDSSNPLPENFDGCLILAPIYEVQGVMKFRRDIV
NYVTRTGQAMRQSLRSFQVTTCTWHGGKPLDNCQYRATRDSRMIVIACDEHEHPVHFAESQI
QETTLTVINVYCPRADPQRPERAAYQASFYRLLQGRAEALLRDGGHVVILGDINTSHRPIDHCEPGDLEAF
IQSKRRSESIEVDVLPAYDALGQVTPGNKPSPQVYEDLIQAGARPGEFCTSFTELQRDFVKRRPAKLKNLLR
WVDSQQKRYVPVKGDHVIGIVTLKAGDIFKVDVGGSEQASLSYLAFEGATKRNRPNVQVGDLVYGQFVVA
VMGGWSAWGPWSSCSVTCAEGVRKRIRSCTEPVPICGGSCPSPSLSTDTLPCDTNKICPTHGNWGSW
QLQQLVLVPGHHLQPPAQFLLPQAQQGQQGLLPTPNLFQLPQQSPGGLLPAPSRAGLPAQPVTRPGLPES
KEMKDKSYMPRDFKPEYAIAILAYTSQGLLNKQPLHQEFNAAVREAGRTRDFYLNNFNFKTLHFLLTRALH
DNGRNELSLLIEKGHSISCPGIHCYNHLQRSPVTWYKNGIRIMFKEIRPSLKLKDDKINLSSTYEQDAGIYV
PCMFGDVLAPPVGTVQISLLPAPETQTVPGKPVTPIKAESILSRKPPIQPKPVVVATVPVQPAATPSPTILLQTVP
DVQCQGTEAALGNCLYTDPEYLCQHDEDAGARCQAVELPTPPVRLVGNNKRCSGQVEIYYDGRWGAVC
YWVDSEKGLLQRVYLNGTKQERLRYVEKGVAGFAINWINQDILWANHQKATIEATDLNGNNSRVLLKDV
SRARNDTRANANGLESYEDAGSSESRTVNGSDSPLLTNGVFKPARPPRPSRPPPPTPRRPSSAAGCVNG
CFPSRIRYPNDTGTLTSGRIVCPTIDNYAKATIVKWYKDCKPLQGPREKYSMGERYLFIKRPQKTDEGHYTC
RSPLGLGGPWFINSIQAPARGYGVSAVLKSEEAAMSLQPAVTMATAVLSVFSHTPHLDSRLGRDVLLDCGF
VKVRVYSLNEELRPAQRETVLTFVDPEGVKVDIIEERDFTGILSFPDFKIPSNPKYGVWKIEAKYKKSFTTSAVA
EEWQKIRSILGKHMLKPKEVESYTGTLNSVVSDLIRRLQHQRSCHQQHVVKDVAGEFYKFGLEGISSVLFD
LTAAHCNGDKITVYLGAHNIKQQEQSQQKISVRRQIPHCKYNRKIFNNDIMLLQLKHKAELNEQVGLISLP
LNLTQLYLNDAFLEFLPANFGRLTKLQILELRENQLKMLPKTMNRLTQLERLDLGSNEFTEVPEVLEQLSGLK
FFDDPPTRGWVSIKYLKPYKGSAVRETQRGGMFYSSKPEIKRAMEMADDAMRKDKTKRLELAVCNEPSDT

EKHSWYHGPVSRNAAEYLLSSGINGSFLVRESESSPGQRSISLRYEGRVYHYRINTASDGKLYVSSESRFNT
QSQQTQAHKVSSYGHSRSSEDVSVHATSKPKFKKGFSLRNMSLCVVDGMKEMWHRKSSPEPGMEATQG
GLQLSVVTRDGLSVNPRAAGLIGRNGPYDKQPFMVAFFKVSEVHVRTTRSATSRRRQQNRNRSTQSQDVS
ASDCINAMVEGSELRKFRSNSRVYHRYFLLDADMQSLHWEPSKKDSEKAKIDIKSVKEVRTGKNTDIFRSN
VQNVTIPVALIRYYDIMDMQQALGARINVTLYSPSLPEFDYSMVVIFLIAVLTVGLGGYWSGVSELEDLKAVTNI
TIKKARTSTAGPHPYTPEHTSAAANASDTSSAITPQWPLVLSSEAPEDVSVQTMLTSIHQQMEPESTRAGVHVG
SKPVIISTPTPTIMRPGSLPLHLGYDPLHPTLPSPTSVITQAPPSNRQLGSPTGSLPLVMHLANGQTMPVLPGPP
TWGFYQDSYDGRDFLTFHKETMTWVAADIGAQITKRRWDTEIGDYQHWKRYLEERCIPWLRSSLEYGKE
PDPAFNLPLPPPPPEPLTQLVVKVHMNDNSTKSLMVDERQVARDVLDNLFEKTHCDCNVDWCLYEIYPEL
CSCCEMNVHKRCVQSVPSLCGVDHTERRGRLQLRIETLGGEEIRVFVGEARNLIPMDPNGLSDPYVKLKLI
YILAQFLGAFVGAAAVFGIYYDAFMNYSGGELIVTGSNATAHIFATYPAPHLSLISGFADQVMSTALLVLVVFAI
GNGKWDCEQHACLIDGEMIDAINRGNYGWRAANYSHFWGMTLDEGIRYRLGTIKPPPVVMHMNELQMS
THPNGTVSVISKLYVNSSTSLAGQLLTCRVRHMERDRDYSVKVKEGQ
VTHTAGGAVEPNGSILHISLSPRDAHLNVTCTAQNPASNSATTASAKDLCAETLGSPSSATEGSQARPASQRS
FLGSEPIISRNSHQSPLRTEYAIPWLNEPSFVFADVIRADRNSTEEDDDKVYFFFTEMSVEYEFVGKLMIPRIS
AEQETVTQLAPQKIALWLRPGEQHSFSVRFKRAEGYPVDVYYLMDLSYSMKDDLENVKKLGNDLLAALRNI
FMVTDCDNICKTNQSICYSFYPVLAQDYVNVREEDALVEDKLDFHSMQDIKSVSADKIKNITCIYFYDGCC
GLGWLSFFLYLGTGFLCQCDPPVEVPETLIAWTGACLSIWCRYKSCVVNPIKTNKLIISSLTWYLNPVYDSEKK

| Progen2_scoı | Progen2_fina | RITA_score_v | RITA_score_r | RITA_final_score |
|---|---|---|---|---|
| -0.0007967 | 7.46E-08 | -5.7761756 | -5.7763287 | -0.0001531 |
| -0.0002777 | -6.77E-07 | -1.1480411 | -1.1555094 | -0.0074683 |
| -0.0021234 | -2.10E-05 | -0.909482 | -0.9326511 | -0.0231691 |
| -0.0007554 | -1.70E-05 | -6.1483403 | -6.2438149 | -0.0954746 |
| -0.0015775 | -1.51E-05 | -0.768777 | -0.8207616 | -0.0519845 |
| -0.0008883 | -1.25E-05 | -0.2800521 | -0.3450576 | -0.0650056 |
| -0.0011941 | -5.42E-07 | -0.795298 | -0.8468308 | -0.0515328 |
| -0.0013108 | -4.86E-06 | -1.5605069 | -1.58072 | -0.0202131 |
| -0.0014805 | -2.24E-06 | -7.8435887 | -7.8665599 | -0.0229711 |
| -0.0022127 | 3.34E-06 | -16.503493 | -16.511009 | -0.0075154 |
| -0.0006566 | -3.32E-05 | -0.6270494 | -0.6503557 | -0.0233063 |
| -0.0024263 | -4.87E-06 | -2.3552071 | -2.3733878 | -0.0181807 |
| -0.0027324 | -1.99E-05 | -1.4829732 | -1.5087059 | -0.0257326 |
| -0.0006879 | -1.65E-06 | -3.502847 | -3.5039282 | -0.0010811 |
| -0.0004274 | -6.55E-07 | -0.4019826 | -0.4032089 | -0.0012263 |
| -0.0002644 | 1.24E-09 | -1.3239557 | -1.3269317 | -0.002976 |
| -0.0023968 | -3.46E-06 | -2.900107 | -2.912194 | -0.012087 |
| -0.0010597 | -3.86E-07 | -5.0344251 | -5.0373703 | -0.0029452 |
| -0.0005118 | -1.38E-06 | -2.4483284 | -2.4662896 | -0.0179612 |
| -0.0001571 | -4.31E-06 | -1.1290543 | -1.1392764 | -0.0102221 |
| -0.0008658 | -4.32E-07 | -20.552388 | -20.556313 | -0.0039246 |
| -0.0011099 | -1.72E-05 | -0.4449093 | -0.4670334 | -0.0221241 |
| -0.0007551 | -8.64E-07 | -3.8675048 | -3.8920173 | -0.0245126 |
| -0.0013997 | -3.01E-05 | -0.956029 | -1.0023501 | -0.0463211 |
| -0.0006946 | -6.38E-05 | -3.6611753 | -4.0220606 | -0.3608853 |

| | | | | |
|---|---|---|---|---|
| -0.0001426 | -2.49E-06 | -2.9223296 | -2.9811999 | -0.0588704 |
| -0.0010144 | 5.20E-07 | -4.3244708 | -4.3458529 | -0.0213821 |
| -0.0012888 | -1.57E-05 | -0.7905396 | -0.8133472 | -0.0228076 |
| -0.0004551 | -2.67E-05 | -0.3147297 | -0.3489361 | -0.0342064 |
| -0.0001267 | -4.36E-06 | -1.2305499 | -1.2688979 | -0.038348 |
| -0.0002442 | -5.05E-06 | -1.428465 | -1.467354 | -0.038889 |
| -0.003837 | -8.74E-06 | -2.6240703 | -2.6732416 | -0.0491713 |
| -0.0007353 | -6.01E-06 | -2.5569598 | -2.5760515 | -0.0190917 |
| -0.0009988 | -2.46E-06 | -3.179351 | -3.2928262 | -0.1134751 |
| -0.0008055 | -3.61E-05 | -1.7497606 | -1.7655132 | -0.0157526 |
| -0.0007479 | -2.15E-05 | -0.4553627 | -0.5013803 | -0.0460176 |
| -0.0019294 | 1.71E-07 | -11.307464 | -11.309479 | -0.0020143 |
| -0.0004983 | -9.31E-06 | -2.0907101 | -2.1296644 | -0.0389543 |
| -0.0008713 | 1.65E-08 | -1.8203478 | -1.8167188 | 0.00362903 |
| -0.0005028 | -2.83E-05 | -4.7844269 | -4.8337323 | -0.0493053 |
| -0.0011064 | 3.63E-06 | -0.4144071 | -0.4183699 | -0.0039628 |
| -0.0036351 | -4.56E-06 | -1.0966194 | -1.1407531 | -0.0441337 |
| -0.0006065 | -9.81E-06 | -0.6497772 | -0.6568594 | -0.0070822 |
| -0.0008389 | -1.56E-05 | -0.3730513 | -0.398612 | -0.0255607 |
| -0.0002027 | -1.08E-05 | -5.987494 | -6.0195446 | -0.0320507 |
| -0.0004339 | -7.90E-07 | -1.5965817 | -1.6069037 | -0.010322 |
| -0.0007327 | -6.90E-07 | -1.8065445 | -1.8096137 | -0.0030692 |
| -0.0008151 | -1.04E-06 | -2.81196 | -2.819927 | -0.007967 |
| -0.0020742 | -1.34E-07 | -10.067 | -10.067887 | -0.0008869 |
| -0.001127 | 1.57E-06 | -1.174509 | -1.175121 | -0.000612 |
| -0.000448 | -7.46E-06 | -1.5591125 | -1.5822682 | -0.0231557 |
| -0.0001794 | -3.01E-06 | -1.802992 | -1.8061398 | -0.0031478 |
| -0.0019814 | -8.27E-07 | -1.0988454 | -1.0974442 | 0.00140119 |
| -0.0077922 | 1.71E-05 | -1.9902009 | -2.0058901 | -0.0156892 |
| -0.0028976 | -3.98E-06 | -1.9840342 | -2.032913 | -0.0488787 |
| -0.0046921 | -3.25E-05 | -3.4195845 | -3.432229 | -0.0126445 |
| -0.0024611 | -1.02E-05 | -2.3679322 | -2.3961196 | -0.0281874 |
| -0.0010116 | -5.52E-06 | -7.2152555 | -7.2844883 | -0.0692328 |
| -0.000354 | -3.10E-06 | -2.6109387 | -2.6653247 | -0.054386 |
| -0.0015318 | -3.25E-05 | -1.2527143 | -1.287239 | -0.0345247 |
| -0.003682 | -9.48E-06 | -2.1158047 | -2.1593046 | -0.0434999 |
| -0.0049296 | 2.71E-06 | -4.957653 | -4.9563878 | 0.00126529 |
| -0.0038584 | -8.46E-06 | -1.8451612 | -1.8515058 | -0.0063446 |
| -0.0023812 | -1.94E-05 | -1.1163919 | -1.1308352 | -0.0144433 |
| -0.0147289 | -0.0003837 | -4.940866 | -5.0625293 | -0.1216633 |
| -0.0034077 | -6.73E-06 | -1.4850341 | -1.5050884 | -0.0200543 |
| -0.0049089 | -0.0001501 | -2.6534158 | -2.7368958 | -0.08348 |
| -0.0002944 | -9.38E-06 | -1.8749188 | -1.9484986 | -0.0735798 |
| -0.0098503 | -4.72E-05 | -4.569036 | -4.5865607 | -0.0175247 |
| -0.0002193 | -3.91E-06 | -3.5323832 | -3.6810507 | -0.1486675 |

| | | | | |
|---|---|---|---|---|
| -0.0006559 | -1.29E-06 | -1.6267801 | -1.6425428 | -0.0157627 |
| -0.0075896 | 4.43E-05 | -3.4411291 | -3.435216 | 0.00591314 |
| -0.0065312 | -8.25E-05 | -4.888731 | -4.9196475 | -0.0309165 |
| -0.0028 | -1.03E-05 | -1.5363495 | -1.5546269 | -0.0182774 |
| -0.0068558 | -2.89E-05 | -4.7834325 | -4.7909698 | -0.0075374 |
| -0.0082141 | -0.00012 | -2.9864361 | -3.0705392 | -0.0841031 |
| -0.0014512 | -9.15E-06 | -7.4127843 | -7.4816777 | -0.0688934 |
| -0.0013888 | -6.06E-06 | -1.6657202 | -1.67192 | -0.0061998 |
| -0.0009118 | -3.16E-07 | -5.0401531 | -5.0531421 | -0.012989 |
| -0.0019305 | -1.69E-05 | -1.2581683 | -1.3300681 | -0.0718997 |
| -0.0022363 | -1.81E-05 | -0.6970943 | -0.7269341 | -0.0298398 |
| -0.0022017 | -1.06E-05 | -2.8286794 | -2.86096 | -0.0322806 |
| -0.0005577 | -2.75E-06 | -1.3578547 | -1.3718305 | -0.0139758 |
| -0.0116803 | -9.84E-06 | -4.2449923 | -4.2612259 | -0.0162337 |
| -0.0009608 | -5.10E-06 | -2.3794524 | -2.43561 | -0.0561575 |
| -0.0008443 | -2.48E-05 | -0.6194317 | -0.6594021 | -0.0399704 |
| -0.0002235 | -2.10E-05 | -1.0097306 | -1.1113297 | -0.1015991 |
| -0.0047523 | -0.000137 | -1.6299783 | -1.71623 | -0.0862517 |
| -0.0024114 | 3.11E-07 | -2.8693186 | -2.8672384 | 0.0020802 |
| -0.0028816 | -2.14E-05 | -2.5239444 | -2.5309724 | -0.007028 |
| -0.0048469 | -1.28E-05 | -2.5258427 | -2.5129021 | 0.01294053 |
| -0.0004916 | -8.93E-06 | -2.6256427 | -2.6488921 | -0.0232494 |
| -0.0011984 | -2.02E-06 | -1.1088262 | -1.1266348 | -0.0178086 |
| -0.0010857 | -6.32E-06 | -1.0281827 | -1.0359121 | -0.0077294 |
| -0.0025983 | 1.24E-06 | -0.5123502 | -0.5569021 | -0.0445519 |
| -0.0016649 | -9.08E-06 | -1.3613333 | -1.3882045 | -0.0268713 |
| -0.0030931 | -4.90E-06 | -1.3240166 | -1.3479517 | -0.0239351 |
| -0.0123485 | -0.0007941 | -3.5523084 | -3.7631464 | -0.210838 |
| -0.001807 | -2.85E-06 | -2.6761651 | -2.6853393 | -0.0091742 |
| -0.0040904 | -7.70E-05 | -1.5730819 | -1.6276278 | -0.0545459 |
| -0.0052083 | -0.0018215 | -2.1696168 | -3.4161844 | -1.2465676 |
| -0.0009297 | -2.48E-06 | -3.9869752 | -4.0126865 | -0.0257113 |
| -0.0004704 | -6.03E-06 | -5.0274558 | -5.1018876 | -0.0744318 |
| -0.0008014 | -1.64E-07 | -0.9612685 | -0.9684643 | -0.0071958 |
| -0.0020312 | -8.71E-05 | -1.0714771 | -1.1977351 | -0.126258 |
| -0.0040835 | 1.50E-06 | -4.5495367 | -4.5457981 | 0.00373864 |
| -0.0027266 | -1.85E-05 | -1.8184353 | -1.836888 | -0.0184527 |
| -0.0069629 | -0.0002704 | -3.4321432 | -3.5167758 | -0.0846326 |
| -0.0016994 | -2.30E-06 | -0.7578108 | -0.7753595 | -0.0175487 |
| -0.0036472 | -0.0001031 | -2.3028153 | -2.3805072 | -0.0776919 |
| -0.0048138 | -0.0003295 | -2.3888377 | -2.5631369 | -0.1742992 |
| -0.0037901 | -4.06E-05 | -1.513676 | -1.5543166 | -0.0406405 |
| -0.0002408 | -5.10E-06 | -1.325442 | -1.4873026 | -0.1618606 |
| -0.0030614 | 1.26E-06 | -1.2722265 | -1.2850339 | -0.0128074 |
| -0.005327 | -1.23E-06 | -2.3013797 | -2.3602325 | -0.0588528 |

| | | | | |
|---|---|---|---|---|
| -0.003138 | -5.34E-05 | -2.6345102 | -2.7396474 | -0.1051372 |
| -0.0075788 | -6.09E-05 | -4.8850434 | -4.9058392 | -0.0207958 |
| -0.0010591 | -2.58E-06 | -7.0708419 | -7.0984657 | -0.0276238 |
| -0.0188874 | -3.93E-05 | -5.5169268 | -5.5280938 | -0.011167 |
| -0.0008313 | -1.49E-05 | -0.684289 | -0.7134912 | -0.0292022 |
| -0.0045098 | -0.0001033 | -2.0171154 | -2.0976983 | -0.080583 |
| -0.0014154 | -2.67E-05 | -0.9565216 | -1.0345602 | -0.0780386 |
| -0.0025427 | -2.32E-05 | -1.5747667 | -1.6051734 | -0.0304067 |
| -0.0002403 | -3.68E-06 | -1.0535637 | -1.0785337 | -0.02497 |
| -0.0024388 | -1.01E-05 | -0.5462767 | -0.5872137 | -0.040937 |
| -0.001439 | -1.69E-06 | -7.8185436 | -7.8257231 | -0.0071795 |
| -0.0035463 | -2.62E-05 | -1.7015079 | -1.716794 | -0.0152861 |
| -0.0052181 | -0.0001783 | -2.638581 | -2.7471552 | -0.1085742 |
| -0.0026394 | -1.33E-06 | -1.7291405 | -1.7297158 | -0.0005753 |
| -0.0022366 | -3.56E-06 | -0.9272602 | -0.9495686 | -0.0223083 |
| -0.0005944 | -4.09E-06 | -3.3412147 | -3.3653787 | -0.024164 |
| -0.0019162 | 1.51E-07 | -9.8785892 | -9.879168 | -0.0005789 |
| -0.0045525 | 1.90E-05 | -2.5116304 | -2.5072314 | 0.00439906 |
| -0.0001553 | -1.04E-06 | -1.4789754 | -1.5100015 | -0.0310261 |
| -0.0045159 | -4.41E-05 | -2.1787543 | -2.2161884 | -0.037434 |
| -0.0044663 | -2.39E-05 | -3.5233126 | -3.5393151 | -0.0160025 |
| -0.0014838 | -2.14E-08 | -24.903048 | -24.898335 | 0.00471306 |
| -0.0009638 | -1.11E-05 | -5.0646605 | -5.075349 | -0.0106885 |
| -0.0016339 | -1.97E-05 | -2.9709574 | -2.9972404 | -0.026283 |
| -0.0006841 | -2.13E-05 | -6.4183677 | -6.449926 | -0.0315584 |
| -0.0012655 | -5.99E-06 | -1.6380005 | -1.6414738 | -0.0034732 |
| -0.0015926 | -5.35E-06 | -8.0951571 | -8.1185447 | -0.0233876 |
| -0.0010464 | -2.22E-05 | -0.9477808 | -0.9980796 | -0.0502988 |
| -0.0172389 | -3.58E-05 | -5.7369568 | -5.7508795 | -0.0139227 |
| -0.0037806 | -0.0001774 | -2.6077371 | -2.7325709 | -0.1248338 |
| -0.0004756 | -1.34E-05 | -1.8096185 | -1.8263183 | -0.0166998 |
| -0.0152934 | -2.43E-05 | -5.773191 | -5.7745426 | -0.0013516 |
| -0.0027822 | -9.20E-06 | -2.3869611 | -2.3992813 | -0.0123202 |
| -0.0011429 | -2.78E-05 | -0.5937798 | -0.62664 | -0.0328602 |
| -0.000475 | -7.21E-06 | -2.401434 | -2.4237048 | -0.0222708 |
| -0.0010022 | -3.83E-06 | -3.9409951 | -3.9979588 | -0.0569637 |
| -0.0017932 | -4.52E-05 | -1.4710156 | -1.5269802 | -0.0559646 |
| -0.0045835 | -0.0015525 | -0.7639301 | -2.0337979 | -1.2698679 |
| -0.0033068 | -7.00E-06 | -2.910821 | -2.919492 | -0.008671 |
| -0.0035822 | -4.06E-05 | -2.2975897 | -2.3276248 | -0.0300351 |
| -0.0003691 | -3.36E-05 | -2.2960039 | -2.3902441 | -0.0942402 |
| -0.0035753 | -9.29E-06 | -3.6944085 | -3.7034751 | -0.0090666 |
| -0.0009603 | -4.83E-05 | -0.4380997 | -0.5259381 | -0.0878385 |
| -0.0010528 | -1.52E-05 | -5.1459557 | -5.1787921 | -0.0328364 |
| -0.0035113 | -5.72E-06 | -2.6226554 | -2.6352743 | -0.0126189 |

| | | | | |
|---|---|---|---|---|
| -0.0013068 | 4.60E-07 | -0.9524018 | -0.953017 | -0.0006151 |
| -0.0010573 | -1.70E-05 | -1.5069317 | -1.5350773 | -0.0281456 |
| -0.0026132 | -6.41E-05 | -1.9005635 | -1.9444394 | -0.0438758 |
| -0.0001801 | -6.91E-06 | -2.1198822 | -2.1343678 | -0.0144856 |
| -0.0072052 | -0.0001069 | -4.4908752 | -4.6059351 | -0.1150599 |
| -0.011651 | -1.89E-05 | -5.6973722 | -5.7056158 | -0.0082436 |
| -0.0005693 | -1.11E-05 | -0.6869611 | -0.7187499 | -0.0317888 |
| -0.0080406 | -6.15E-05 | -2.3378622 | -2.3911169 | -0.0532547 |
| -0.0027794 | -6.92E-06 | -2.0395138 | -2.0599695 | -0.0204557 |
| -0.0009311 | -9.35E-06 | -0.7234929 | -0.7424633 | -0.0189704 |
| -0.0004482 | 6.55E-07 | -2.7418649 | -2.7448485 | -0.0029836 |
| -0.0047232 | -7.43E-07 | -3.4703749 | -3.4807332 | -0.0103582 |
| -0.0031517 | -2.33E-06 | -3.2397258 | -3.2371608 | 0.00256503 |
| -0.0023168 | -2.00E-05 | -1.7621403 | -1.8027049 | -0.0405645 |
| -0.0037114 | -5.04E-05 | -2.4688362 | -2.507272 | -0.0384358 |
| -0.0007749 | -8.06E-06 | -3.7010784 | -3.8026901 | -0.1016118 |
| -0.000992 | -2.94E-05 | -0.6326177 | -0.6613495 | -0.0287317 |
| -0.0051321 | -7.22E-05 | -3.3852556 | -3.4423031 | -0.0570475 |
| -0.0008696 | -9.04E-06 | -2.7332928 | -2.7779737 | -0.0446808 |
| -0.0004673 | -8.74E-06 | -1.7999096 | -1.8827115 | -0.0828019 |
| -0.003805 | -9.35E-06 | -0.9404358 | -0.9659359 | -0.0255001 |
| -0.004016 | -1.31E-05 | -0.9075314 | -0.935944 | -0.0284126 |
| -0.0018533 | -1.95E-05 | -0.9276683 | -1.002188 | -0.0745197 |
| -0.005236 | -9.87E-05 | -2.3630136 | -2.3621871 | 0.00082648 |
| -0.0008402 | -2.39E-06 | -0.862428 | -0.8942331 | -0.0318051 |
| -0.0022079 | -3.33E-06 | -2.3732115 | -2.381009 | -0.0077975 |
| -0.0024657 | -5.10E-07 | -2.685766 | -2.7069935 | -0.0212275 |
| -0.0152771 | 3.38E-05 | -5.7556026 | -5.7474601 | 0.00814247 |
| -0.0113591 | -5.25E-06 | -4.2692192 | -4.2846057 | -0.0153866 |
| -0.0091229 | 1.96E-05 | -5.1913033 | -5.2127733 | -0.0214701 |
| -0.0020593 | -2.62E-06 | -3.8007272 | -3.8063498 | -0.0056225 |
| -0.0139662 | -0.0001374 | -5.5617664 | -5.6252408 | -0.0634744 |
| -0.001986 | 5.36E-07 | -1.9768283 | -1.9757153 | 0.001113 |
| -0.0055009 | -9.18E-05 | -4.2536428 | -4.3081722 | -0.0545294 |
| -0.0029293 | -1.14E-05 | -1.9877014 | -2.0032766 | -0.0155752 |
| -0.006784 | -0.0001301 | -4.6323006 | -4.7077641 | -0.0754635 |
| -0.0006765 | -9.96E-06 | -0.3761401 | -0.403517 | -0.0273769 |
| -0.000492 | -1.91E-06 | -1.8281673 | -1.8649904 | -0.0368231 |
| -0.0014638 | -2.09E-05 | -0.471567 | -0.5491727 | -0.0776057 |
| -0.0008862 | -1.17E-05 | -0.7929353 | -0.7972625 | -0.0043272 |
| -0.0049086 | 1.23E-05 | -4.2811604 | -4.2791626 | 0.00199771 |
| -0.0064717 | 1.20E-06 | -5.1953313 | -5.2007217 | -0.0053904 |
| -0.0044834 | -1.47E-05 | -2.5967596 | -2.6510245 | -0.0542649 |
| -0.0012751 | -3.72E-06 | -5.0329672 | -5.0641057 | -0.0311385 |
| -0.0018269 | -5.53E-06 | -1.968936 | -1.9887379 | -0.019802 |

| | | | | |
|---|---|---|---|---|
| -0.0014966 | -2.59E-05 | -0.5590315 | -0.5912526 | -0.0322212 |
| -0.0041008 | -2.12E-05 | -1.820273 | -1.8292862 | -0.0090131 |
| -0.0054475 | -5.64E-05 | -2.9584706 | -2.9758506 | -0.01738 |
| -0.0016715 | -2.00E-05 | -0.4332443 | -0.4808843 | -0.0476401 |
| -0.0017416 | -2.36E-05 | -1.5206687 | -1.6342247 | -0.113556 |
| -0.0014086 | -2.27E-05 | -2.1424934 | -2.1932073 | -0.0507139 |
| -0.0019656 | -1.19E-05 | -2.1803623 | -2.1939689 | -0.0136065 |
| -0.0031403 | 3.44E-05 | -1.6977017 | -1.7585671 | -0.0608654 |
| -0.0036909 | 2.87E-07 | -2.1532099 | -2.1511445 | 0.00206542 |
| -0.0196617 | -0.0001169 | -5.7784798 | -5.8021338 | -0.023654 |
| -0.0032257 | -8.30E-05 | -2.8066473 | -2.9174521 | -0.1108048 |
| -0.0037419 | -7.59E-06 | -4.4451437 | -4.450218 | -0.0050743 |
| -0.0047997 | -4.64E-05 | -2.0469307 | -2.117374 | -0.0704433 |
| -0.0038208 | 1.96E-06 | -4.1691189 | -4.1686571 | 0.00046182 |
| -0.0007037 | 4.87E-06 | -0.3868574 | -0.4015357 | -0.0146783 |
| -0.0053269 | -4.76E-05 | -3.9419196 | -4.0530649 | -0.1111454 |
| -0.0034094 | -5.85E-06 | -3.5949917 | -3.6068532 | -0.0118616 |
| -0.0028615 | -2.30E-05 | -1.0910285 | -1.1269784 | -0.0359498 |
| -0.005002 | -5.76E-06 | -3.536881 | -3.5433372 | -0.0064563 |
| -0.0011088 | 3.73E-07 | -6.5867815 | -6.5787601 | 0.00802135 |
| -0.0007514 | -2.76E-06 | -0.792472 | -0.7998446 | -0.0073726 |
| -0.0038748 | -3.48E-05 | -3.9030195 | -3.9526346 | -0.049615 |
| -0.0042699 | -7.77E-06 | -3.765528 | -3.7735192 | -0.0079912 |
| -0.0006336 | -2.42E-06 | -4.6312806 | -4.6479481 | -0.0166675 |
| -0.0025602 | -5.87E-06 | -1.989177 | -2.0015798 | -0.0124028 |
| -0.0050851 | -4.53E-05 | -2.638581 | -2.6976659 | -0.0590849 |
| -0.0004011 | -2.88E-06 | -1.1399609 | -1.1645483 | -0.0245874 |
| -0.0009291 | -1.37E-05 | -2.2683671 | -2.4331954 | -0.1648282 |
| -0.0009065 | -4.40E-06 | -2.0653144 | -2.0807258 | -0.0154114 |
| -0.0020391 | -2.77E-05 | -1.8010639 | -1.8287292 | -0.0276653 |
| -0.0025275 | -0.0001642 | -1.8864185 | -2.0787426 | -0.1923242 |
| -0.0008881 | 1.28E-06 | -2.7815961 | -2.7788619 | 0.00273415 |
| -0.0024281 | -3.76E-06 | -1.6561757 | -1.6666728 | -0.0104972 |
| -0.0078859 | -3.01E-05 | -4.740658 | -4.7495806 | -0.0089226 |
| -0.0018516 | -2.96E-05 | -0.5895518 | -0.6311555 | -0.0416037 |
| -0.0064363 | -0.0020743 | -2.5686424 | -4.2601602 | -1.6915178 |
| -0.0017937 | -2.28E-05 | -2.03149 | -2.0608676 | -0.0293776 |
| -0.0012308 | -1.41E-05 | -0.8377688 | -0.864923 | -0.0271543 |
| -0.0041518 | -5.31E-05 | -2.1969886 | -2.2311136 | -0.034125 |
| -0.00204 | -1.93E-05 | -1.1381428 | -1.1873072 | -0.0491644 |
| -0.0059698 | -5.24E-05 | -2.5214831 | -2.5825021 | -0.0610191 |
| -0.0034184 | -1.33E-05 | -4.5323098 | -4.5974689 | -0.0651591 |
| -0.0013578 | -4.25E-06 | -1.4492629 | -1.455106 | -0.005843 |
| -0.0013677 | -1.20E-05 | -1.0606045 | -1.0933716 | -0.0327671 |
| -0.0027318 | -2.72E-05 | -3.9737203 | -4.0207973 | -0.0470769 |

| | | | | |
|---|---|---|---|---|
| -0.0023847 | -3.72E-05 | -3.3864609 | -3.4435612 | -0.0571003 |

| Progen2_scoı | Progen2_fina | RITA_score_v | RITA_score_r | RITA_final_score |
|---|---|---|---|---|
| -0.000797 | -1.81E-07 | -5.7761756 | -5.7773766 | -0.0012009 |
| -0.0007965 | 2.55E-07 | -5.7761756 | -5.7751628 | 0.0010128 |
| -0.0002777 | -6.77E-07 | -1.1480411 | -1.1555094 | -0.0074683 |
| -0.0021234 | -2.10E-05 | -0.909482 | -0.9326511 | -0.0231691 |
| -0.0007435 | -5.09E-06 | -6.1483403 | -6.1791222 | -0.0307819 |
| -0.0007397 | -1.36E-06 | -6.1483403 | -6.1578207 | -0.0094804 |
| -0.0007404 | -2.05E-06 | -6.1483403 | -6.1652117 | -0.0168713 |
| -0.0007393 | -9.71E-07 | -6.1483403 | -6.1652745 | -0.0169342 |
| -0.0007459 | -7.48E-06 | -6.1483403 | -6.1682673 | -0.0199269 |
| -0.001567 | -4.55E-06 | -0.768777 | -0.8002546 | -0.0314775 |
| -0.0015732 | -1.07E-05 | -0.768777 | -0.7892605 | -0.0204835 |
| -0.0008844 | -8.65E-06 | -0.2800521 | -0.3281193 | -0.0480673 |
| -0.0008798 | -4.00E-06 | -0.2800521 | -0.2969154 | -0.0168633 |
| -0.0011941 | -5.42E-07 | -0.795298 | -0.8468308 | -0.0515328 |
| -0.0013059 | 1.45E-08 | -1.5605069 | -1.5607784 | -0.0002715 |
| -0.0013061 | -1.41E-07 | -1.5605069 | -1.5620294 | -0.0015225 |
| -0.0013106 | -4.67E-06 | -1.5605069 | -1.5785624 | -0.0180554 |
| -0.0014798 | -1.57E-06 | -7.8435887 | -7.8586862 | -0.0150975 |
| -0.0014789 | -6.68E-07 | -7.8435887 | -7.8514624 | -0.0078737 |
| -0.0022164 | -3.38E-07 | -16.503493 | -16.507315 | -0.0038216 |
| -0.0022124 | 3.68E-06 | -16.503493 | -16.507187 | -0.0036938 |
| -0.0006566 | -3.32E-05 | -0.6270494 | -0.6503557 | -0.0233063 |
| -0.0024263 | -4.87E-06 | -2.3552071 | -2.3733878 | -0.0181807 |
| -0.0027324 | -1.99E-05 | -1.4829732 | -1.5087059 | -0.0257326 |
| -0.0006874 | -1.07E-06 | -3.502847 | -3.5201389 | -0.0172919 |
| -0.0006856 | 6.69E-07 | -3.502847 | -3.5004875 | 0.00235951 |
| -0.0006855 | 7.39E-07 | -3.502847 | -3.4929378 | 0.00990927 |
| -0.0006865 | -2.44E-07 | -3.502847 | -3.501112 | 0.00173509 |
| -0.0006856 | 7.29E-07 | -3.502847 | -3.4994427 | 0.00340438 |
| -0.0006882 | -1.89E-06 | -3.502847 | -3.5112376 | -0.0083905 |
| -0.0004274 | -6.55E-07 | -0.4019826 | -0.4032089 | -0.0012263 |
| -0.0002658 | -1.32E-06 | -1.3239557 | -1.3367253 | -0.0127695 |
| -0.0002631 | 1.32E-06 | -1.3239557 | -1.3141622 | 0.00979356 |
| -0.0023968 | -3.46E-06 | -2.900107 | -2.912194 | -0.012087 |
| -0.0010597 | -3.86E-07 | -5.0344251 | -5.0373703 | -0.0029452 |
| -0.0005118 | -1.38E-06 | -2.4483284 | -2.4662896 | -0.0179612 |
| -0.0001571 | -4.31E-06 | -1.1290543 | -1.1392764 | -0.0102221 |
| -0.0008654 | -9.23E-08 | -20.552388 | -20.554197 | -0.0018091 |
| -0.0008657 | -3.40E-07 | -20.552388 | -20.554504 | -0.0021155 |
| -0.0011099 | -1.72E-05 | -0.4449093 | -0.4670334 | -0.0221241 |
| -0.0007546 | -3.53E-07 | -3.8675048 | -3.8819757 | -0.0144709 |
| -0.0007547 | -5.06E-07 | -3.8675048 | -3.8777579 | -0.0102532 |

| | | | | |
|---|---|---|---|---|
| -0.0013997 | -3.01E-05 | -0.956029 | -1.0023501 | -0.0463211 |
| -0.0006298 | 9.81E-07 | -3.6611753 | -3.6514991 | 0.00967622 |
| -0.000634 | -3.24E-06 | -3.6611753 | -3.6863374 | -0.0251621 |
| -0.000632 | -1.26E-06 | -3.6611753 | -3.6755767 | -0.0144014 |
| -0.0006334 | -2.65E-06 | -3.6611753 | -3.6855317 | -0.0243564 |
| -0.0006325 | -1.73E-06 | -3.6611753 | -3.6765908 | -0.0154155 |
| -0.0006331 | -2.30E-06 | -3.6611753 | -3.6767055 | -0.0155302 |
| -0.0006344 | -3.66E-06 | -3.6611753 | -3.6907937 | -0.0296184 |
| -0.0006338 | -3.07E-06 | -3.6611753 | -3.6868189 | -0.0256436 |
| -0.0006337 | -2.98E-06 | -3.6611753 | -3.6821772 | -0.0210019 |
| -0.0006333 | -2.50E-06 | -3.6611753 | -3.6813717 | -0.0201964 |
| -0.0006326 | -1.88E-06 | -3.6611753 | -3.6816359 | -0.0204606 |
| -0.0006358 | -5.03E-06 | -3.6611753 | -3.6907989 | -0.0296236 |
| -0.0006656 | -3.48E-05 | -3.6611753 | -3.8020649 | -0.1408896 |
| -0.0001426 | -2.49E-06 | -2.9223296 | -2.9811999 | -0.0588704 |
| -0.001016 | -1.11E-06 | -4.3244708 | -4.3343662 | -0.0098954 |
| -0.0010133 | 1.63E-06 | -4.3244708 | -4.3359575 | -0.0114867 |
| -0.0012888 | -1.57E-05 | -0.7905396 | -0.8133472 | -0.0228076 |
| -0.0004551 | -2.67E-05 | -0.3147297 | -0.3489361 | -0.0342064 |
| -0.0001267 | -4.36E-06 | -1.2305499 | -1.2688979 | -0.038348 |
| -0.0002442 | -5.05E-06 | -1.428465 | -1.467354 | -0.038889 |
| -0.003837 | -8.74E-06 | -2.6240703 | -2.6732416 | -0.0491713 |
| -0.0007353 | -6.01E-06 | -2.5569598 | -2.5760515 | -0.0190917 |
| -0.0009988 | -2.46E-06 | -3.179351 | -3.2928262 | -0.1134751 |
| -0.0008055 | -3.61E-05 | -1.7497606 | -1.7655132 | -0.0157526 |
| -0.0007479 | -2.15E-05 | -0.4553627 | -0.5013803 | -0.0460176 |
| -0.0019294 | 1.71E-07 | -11.307464 | -11.309479 | -0.0020143 |
| -0.0004983 | -9.31E-06 | -2.0907101 | -2.1296644 | -0.0389543 |
| -0.0008713 | 1.65E-08 | -1.8203478 | -1.8167188 | 0.00362903 |
| -0.0005028 | -2.83E-05 | -4.7844269 | -4.8337323 | -0.0493053 |
| -0.0011064 | 3.63E-06 | -0.4144071 | -0.4183699 | -0.0039628 |
| -0.0036351 | -4.56E-06 | -1.0966194 | -1.1407531 | -0.0441337 |
| -0.0006065 | -9.81E-06 | -0.6497772 | -0.6568594 | -0.0070822 |
| -0.0008389 | -1.56E-05 | -0.3730513 | -0.398612 | -0.0255607 |
| -0.0001987 | -6.72E-06 | -5.987494 | -6.0055252 | -0.0180313 |
| -0.000196 | -4.03E-06 | -5.987494 | -6.0026053 | -0.0151113 |
| -0.0004339 | -7.90E-07 | -1.5965817 | -1.6069037 | -0.010322 |
| -0.0007327 | -6.90E-07 | -1.8065445 | -1.8096137 | -0.0030692 |
| -0.0008151 | -1.04E-06 | -2.81196 | -2.819927 | -0.007967 |
| -0.0020742 | -1.34E-07 | -10.067 | -10.067887 | -0.0008869 |
| -0.001127 | 1.57E-06 | -1.174509 | -1.175121 | -0.000612 |
| -0.000448 | -7.46E-06 | -1.5591125 | -1.5822682 | -0.0231557 |
| -0.0001794 | -3.01E-06 | -1.802992 | -1.8061398 | -0.0031478 |
| -0.0019814 | -8.27E-07 | -1.0988454 | -1.0974442 | 0.00140119 |
| -0.0077922 | 1.71E-05 | -1.9902009 | -2.0058901 | -0.0156892 |

| | | | | |
|---|---|---|---|---|
| -0.0028976 | -3.98E-06 | -1.9840342 | -2.032913 | -0.0488787 |
| -0.0046921 | -3.25E-05 | -3.4195845 | -3.432229 | -0.0126445 |
| -0.0024611 | -1.02E-05 | -2.3679322 | -2.3961196 | -0.0281874 |
| -0.0010066 | -5.73E-07 | -7.2152555 | -7.2445961 | -0.0293406 |
| -0.001011 | -4.95E-06 | -7.2152555 | -7.2551476 | -0.0398921 |
| -0.0003517 | -7.60E-07 | -2.6109387 | -2.6259028 | -0.014964 |
| -0.0003522 | -1.22E-06 | -2.6109387 | -2.6303264 | -0.0193877 |
| -0.0003521 | -1.13E-06 | -2.6109387 | -2.6310167 | -0.020078 |
| -0.0015318 | -3.25E-05 | -1.2527143 | -1.287239 | -0.0345247 |
| -0.003682 | -9.48E-06 | -2.1158047 | -2.1593046 | -0.0434999 |
| -0.0049296 | 2.71E-06 | -4.957653 | -4.9563878 | 0.00126529 |
| -0.0038584 | -8.46E-06 | -1.8451612 | -1.8515058 | -0.0063446 |
| -0.0023812 | -1.94E-05 | -1.1163919 | -1.1308352 | -0.0144433 |
| -0.0147289 | -0.0003837 | -4.940866 | -5.0625293 | -0.1216633 |
| -0.0034077 | -6.73E-06 | -1.4850341 | -1.5050884 | -0.0200543 |
| -0.0049089 | -0.0001501 | -2.6534158 | -2.7368958 | -0.08348 |
| -0.0002867 | -1.68E-06 | -1.8749188 | -1.8898151 | -0.0148963 |
| -0.0002892 | -4.17E-06 | -1.8749188 | -1.9004414 | -0.0255226 |
| -0.0002885 | -3.42E-06 | -1.8749188 | -1.9030474 | -0.0281286 |
| -0.0002853 | -2.35E-07 | -1.8749188 | -1.8824318 | -0.007513 |
| -0.0002859 | -8.52E-07 | -1.8749188 | -1.8840797 | -0.009161 |
| -0.0098503 | -4.72E-05 | -4.569036 | -4.5865607 | -0.0175247 |
| -0.0002158 | -3.80E-07 | -3.5323832 | -3.5486661 | -0.0162828 |
| -0.000216 | -5.93E-07 | -3.5323832 | -3.5555987 | -0.0232154 |
| -0.0002159 | -4.46E-07 | -3.5323832 | -3.5473501 | -0.0149669 |
| -0.0002158 | -3.17E-07 | -3.5323832 | -3.5551348 | -0.0227515 |
| -0.0002161 | -6.51E-07 | -3.5323832 | -3.5505887 | -0.0182054 |
| -0.0002159 | -4.79E-07 | -3.5323832 | -3.5467772 | -0.0143939 |
| -0.0002162 | -7.22E-07 | -3.5323832 | -3.5469266 | -0.0145433 |
| -0.0002158 | -3.26E-07 | -3.5323832 | -3.5569801 | -0.0245969 |
| -0.0006559 | -1.29E-06 | -1.6267801 | -1.6425428 | -0.0157627 |
| -0.0075896 | 4.43E-05 | -3.4411291 | -3.435216 | 0.00591314 |
| -0.00648 | -3.13E-05 | -4.888731 | -4.8980412 | -0.0093102 |
| -0.0064618 | -1.32E-05 | -4.888731 | -4.9013546 | -0.0126235 |
| -0.0064663 | -1.76E-05 | -4.888731 | -4.8869495 | 0.00178146 |
| -0.0064669 | -1.83E-05 | -4.888731 | -4.9001989 | -0.0114679 |
| -0.0028 | -1.03E-05 | -1.5363495 | -1.5546269 | -0.0182774 |
| -0.0068558 | -2.89E-05 | -4.7834325 | -4.7909698 | -0.0075374 |
| -0.0082141 | -0.00012 | -2.9864361 | -3.0705392 | -0.0841031 |
| -0.0014435 | -1.51E-06 | -7.4127843 | -7.4311578 | -0.0183735 |
| -0.0014498 | -7.80E-06 | -7.4127843 | -7.4475252 | -0.0347409 |
| -0.001442 | 3.23E-09 | -7.4127843 | -7.4292495 | -0.0164652 |
| -0.0013888 | -6.06E-06 | -1.6657202 | -1.67192 | -0.0061998 |
| -0.0009118 | -3.16E-07 | -5.0401531 | -5.0531421 | -0.012989 |
| -0.0019155 | -1.84E-06 | -1.2581683 | -1.27201 | -0.0138416 |

| | | | | |
|---|---|---|---|---|
| -0.0019177 | -4.02E-06 | -1.2581683 | -1.2816513 | -0.023483 |
| -0.0019246 | -1.10E-05 | -1.2581683 | -1.2931107 | -0.0349424 |
| -0.0022363 | -1.81E-05 | -0.6970943 | -0.7269341 | -0.0298398 |
| -0.0022017 | -1.06E-05 | -2.8286794 | -2.86096 | -0.0322806 |
| -0.0005577 | -2.75E-06 | -1.3578547 | -1.3718305 | -0.0139758 |
| -0.0116803 | -9.84E-06 | -4.2449923 | -4.2612259 | -0.0162337 |
| -0.0009615 | -5.72E-06 | -2.3794524 | -2.4018237 | -0.0223712 |
| -0.0009552 | 5.86E-07 | -2.3794524 | -2.4134411 | -0.0339887 |
| -0.0008293 | -9.79E-06 | -0.6194317 | -0.6369941 | -0.0175624 |
| -0.0008255 | -5.95E-06 | -0.6194317 | -0.6271943 | -0.0077626 |
| -0.0008286 | -9.12E-06 | -0.6194317 | -0.6343808 | -0.0149491 |
| -0.0002046 | -2.11E-06 | -1.0097306 | -1.0282524 | -0.0185218 |
| -0.000207 | -4.57E-06 | -1.0097306 | -1.0475715 | -0.0378409 |
| -0.0002168 | -1.44E-05 | -1.0097306 | -1.0549669 | -0.0452364 |
| -0.0046557 | -4.04E-05 | -1.6299783 | -1.6733587 | -0.0433804 |
| -0.0047129 | -9.76E-05 | -1.6299783 | -1.6734896 | -0.0435113 |
| -0.0024114 | 3.11E-07 | -2.8693186 | -2.8672384 | 0.0020802 |
| -0.0028693 | -9.09E-06 | -2.5239444 | -2.5302151 | -0.0062708 |
| -0.002853 | 7.18E-06 | -2.5239444 | -2.5146999 | 0.00924444 |
| -0.0028639 | -3.71E-06 | -2.5239444 | -2.5267588 | -0.0028144 |
| -0.0028754 | -1.52E-05 | -2.5239444 | -2.5308969 | -0.0069525 |
| -0.004845 | -1.10E-05 | -2.5258427 | -2.5716056 | -0.0457629 |
| -0.0048361 | -2.05E-06 | -2.5258427 | -2.4654841 | 0.06035852 |
| -0.0004916 | -8.93E-06 | -2.6256427 | -2.6488921 | -0.0232494 |
| -0.0011984 | -2.02E-06 | -1.1088262 | -1.1266348 | -0.0178086 |
| -0.0010857 | -6.32E-06 | -1.0281827 | -1.0359121 | -0.0077294 |
| -0.0025983 | 1.24E-06 | -0.5123502 | -0.5569021 | -0.0445519 |
| -0.0016561 | -2.86E-07 | -1.3613333 | -1.3639048 | -0.0025715 |
| -0.0016654 | -9.60E-06 | -1.3613333 | -1.3858476 | -0.0245144 |
| -0.0030931 | -4.90E-06 | -1.3240166 | -1.3479517 | -0.0239351 |
| -0.0123485 | -0.0007941 | -3.5523084 | -3.7631464 | -0.210838 |
| -0.001807 | -2.85E-06 | -2.6761651 | -2.6853393 | -0.0091742 |
| -0.0040904 | -7.70E-05 | -1.5730819 | -1.6276278 | -0.0545459 |
| -0.0034245 | -3.77E-05 | -2.1696168 | -2.2095162 | -0.0398993 |
| -0.0034078 | -2.09E-05 | -2.1696168 | -2.1854495 | -0.0158327 |
| -0.0033872 | -3.30E-07 | -2.1696168 | -2.1740463 | -0.0044295 |
| -0.0034025 | -1.57E-05 | -2.1696168 | -2.1783057 | -0.0086889 |
| -0.0033947 | -7.83E-06 | -2.1696168 | -2.168847 | 0.00076985 |
| -0.0033926 | -5.80E-06 | -2.1696168 | -2.1857692 | -0.0161524 |
| -0.0033753 | 1.15E-05 | -2.1696168 | -2.1761689 | -0.0065521 |
| -0.0033935 | -6.67E-06 | -2.1696168 | -2.1922287 | -0.0226119 |
| -0.0033954 | -8.57E-06 | -2.1696168 | -2.1750288 | -0.005412 |
| -0.0033712 | 1.56E-05 | -2.1696168 | -2.1792364 | -0.0096196 |
| -0.0034096 | -2.28E-05 | -2.1696168 | -2.2123562 | -0.0427394 |
| -0.0034759 | -8.91E-05 | -2.1696168 | -2.2191682 | -0.0495514 |

| | | | | |
|---|---|---|---|---|
| -0.0033715 | 1.54E-05 | -2.1696168 | -2.1735767 | -0.0039599 |
| -0.0034069 | -2.01E-05 | -2.1696168 | -2.2102113 | -0.0405945 |
| -0.0034671 | -8.03E-05 | -2.1696168 | -2.2382364 | -0.0686196 |
| -0.0034029 | -1.61E-05 | -2.1696168 | -2.206912 | -0.0372952 |
| -0.0033988 | -1.19E-05 | -2.1696168 | -2.1940483 | -0.0244315 |
| -0.0033837 | 3.13E-06 | -2.1696168 | -2.1921256 | -0.0225087 |
| -0.0034752 | -8.83E-05 | -2.1696168 | -2.2214639 | -0.0518471 |
| -0.0034426 | -5.57E-05 | -2.1696168 | -2.201762 | -0.0321451 |
| -0.0034566 | -6.98E-05 | -2.1696168 | -2.2064521 | -0.0368353 |
| -0.003414 | -2.72E-05 | -2.1696168 | -2.1849544 | -0.0153376 |
| -0.0034267 | -3.98E-05 | -2.1696168 | -2.1847157 | -0.0150989 |
| -0.003413 | -2.62E-05 | -2.1696168 | -2.1845523 | -0.0149355 |
| -0.0034294 | -4.25E-05 | -2.1696168 | -2.207063 | -0.0374461 |
| -0.0035076 | -0.0001208 | -2.1696168 | -2.2127236 | -0.0431068 |
| -0.0034878 | -0.0001009 | -2.1696168 | -2.2210238 | -0.051407 |
| -0.0034605 | -7.36E-05 | -2.1696168 | -2.2652534 | -0.0956366 |
| -0.0033979 | -1.11E-05 | -2.1696168 | -2.1914846 | -0.0218678 |
| -0.0034121 | -2.53E-05 | -2.1696168 | -2.1979626 | -0.0283458 |
| -0.003412 | -2.52E-05 | -2.1696168 | -2.1836228 | -0.014006 |
| -0.003481 | -9.42E-05 | -2.1696168 | -2.2024183 | -0.0328015 |
| -0.0034828 | -9.60E-05 | -2.1696168 | -2.1852727 | -0.0156559 |
| -0.0034665 | -7.97E-05 | -2.1696168 | -2.2125859 | -0.0429691 |
| -0.0034221 | -3.52E-05 | -2.1696168 | -2.1882492 | -0.0186324 |
| -0.0034229 | -3.60E-05 | -2.1696168 | -2.1704061 | -0.0007893 |
| -0.0034417 | -5.49E-05 | -2.1696168 | -2.1925493 | -0.0229325 |
| -0.0035813 | -0.0001944 | -2.1696168 | -2.2467699 | -0.0771531 |
| -0.0034654 | -7.86E-05 | -2.1696168 | -2.2123429 | -0.042726 |
| -0.0034982 | -0.0001113 | -2.1696168 | -2.220969 | -0.0513521 |
| -0.0034521 | -6.52E-05 | -2.1696168 | -2.2157309 | -0.0461141 |
| -0.0035123 | -0.0001255 | -2.1696168 | -2.2243687 | -0.0547519 |
| -0.0034657 | -7.89E-05 | -2.1696168 | -2.1995893 | -0.0299724 |
| -0.0034058 | -1.90E-05 | -2.1696168 | -2.1626941 | 0.00692272 |
| -0.003442 | -5.52E-05 | -2.1696168 | -2.2051218 | -0.0355049 |
| -0.0034156 | -2.88E-05 | -2.1696168 | -2.1670493 | 0.00256753 |
| -0.0033668 | 2.01E-05 | -2.1696168 | -2.1459842 | 0.02363265 |
| -0.0034211 | -3.42E-05 | -2.1696168 | -2.1914142 | -0.0217974 |
| -0.0034301 | -4.32E-05 | -2.1696168 | -2.1872514 | -0.0176346 |
| -0.0034682 | -8.13E-05 | -2.1696168 | -2.2067884 | -0.0371716 |
| -0.00341 | -2.32E-05 | -2.1696168 | -2.2135431 | -0.0439262 |
| -0.0034042 | -1.73E-05 | -2.1696168 | -2.1921079 | -0.0224911 |
| -0.0034263 | -3.95E-05 | -2.1696168 | -2.1880784 | -0.0184616 |
| -0.0036263 | -0.0002395 | -2.1696168 | -2.2752678 | -0.105651 |
| -0.0035062 | -0.0001193 | -2.1696168 | -2.2000729 | -0.0304561 |
| -0.0034656 | -7.88E-05 | -2.1696168 | -2.2176682 | -0.0480514 |
| -0.0034846 | -9.77E-05 | -2.1696168 | -2.2221086 | -0.0524918 |

| | | | | |
|---|---|---|---|---|
| -0.0034767 | -8.99E-05 | -2.1696168 | -2.2019202 | -0.0323033 |
| -0.0035457 | -0.0001589 | -2.1696168 | -2.2435429 | -0.0739261 |
| -0.0034522 | -6.54E-05 | -2.1696168 | -2.2291442 | -0.0595274 |
| -0.003423 | -3.61E-05 | -2.1696168 | -2.2043384 | -0.0347216 |
| -0.0034287 | -4.18E-05 | -2.1696168 | -2.1932619 | -0.023645 |
| -0.0034581 | -7.13E-05 | -2.1696168 | -2.203683 | -0.0340662 |
| -0.0034565 | -6.97E-05 | -2.1696168 | -2.2020327 | -0.0324159 |
| -0.0034557 | -6.88E-05 | -2.1696168 | -2.1674221 | 0.00219476 |
| -0.0034086 | -2.18E-05 | -2.1696168 | -2.177188 | -0.0075712 |
| -0.0034371 | -5.03E-05 | -2.1696168 | -2.2070343 | -0.0374175 |
| -0.0035564 | -0.0001695 | -2.1696168 | -2.268514 | -0.0988972 |
| -0.0034232 | -3.63E-05 | -2.1696168 | -2.1814405 | -0.0118237 |
| -0.0034493 | -6.25E-05 | -2.1696168 | -2.2183102 | -0.0486934 |
| -0.0034296 | -4.28E-05 | -2.1696168 | -2.2031722 | -0.0335554 |
| -0.0034006 | -1.38E-05 | -2.1696168 | -2.1956292 | -0.0260124 |
| -0.0034226 | -3.58E-05 | -2.1696168 | -2.1799999 | -0.0103831 |
| -0.0034308 | -4.40E-05 | -2.1696168 | -2.212929 | -0.0433122 |
| -0.0034183 | -3.14E-05 | -2.1696168 | -2.1954594 | -0.0258425 |
| -0.0034711 | -8.43E-05 | -2.1696168 | -2.2082876 | -0.0386708 |
| -0.003429 | -4.22E-05 | -2.1696168 | -2.1963044 | -0.0266876 |
| -0.0034268 | -3.99E-05 | -2.1696168 | -2.2244923 | -0.0548755 |
| -0.0034736 | -8.68E-05 | -2.1696168 | -2.2416849 | -0.0720681 |
| -0.0034158 | -2.90E-05 | -2.1696168 | -2.1954342 | -0.0258174 |
| -0.0034248 | -3.79E-05 | -2.1696168 | -2.2178528 | -0.048236 |
| -0.0034026 | -1.58E-05 | -2.1696168 | -2.1890205 | -0.0194037 |
| -0.0035199 | -0.0001331 | -2.1696168 | -2.2464439 | -0.076827 |
| -0.0034556 | -6.87E-05 | -2.1696168 | -2.2005291 | -0.0309123 |
| -0.0034012 | -1.44E-05 | -2.1696168 | -2.1796175 | -0.0100007 |
| -0.0034305 | -4.37E-05 | -2.1696168 | -2.2238276 | -0.0542108 |
| -0.0034255 | -3.86E-05 | -2.1696168 | -2.1953903 | -0.0257735 |
| -0.00357 | -0.0001832 | -2.1696168 | -2.2758242 | -0.1062074 |
| -0.0034136 | -2.68E-05 | -2.1696168 | -2.1911998 | -0.021583 |
| -0.0033909 | -4.11E-06 | -2.1696168 | -2.2056755 | -0.0360587 |
| -0.0034108 | -2.40E-05 | -2.1696168 | -2.1937082 | -0.0240914 |
| -0.0034315 | -4.47E-05 | -2.1696168 | -2.1961217 | -0.0265049 |
| -0.0034988 | -0.0001119 | -2.1696168 | -2.2445111 | -0.0748943 |
| -0.0034816 | -9.48E-05 | -2.1696168 | -2.2085258 | -0.038909 |
| -0.0034199 | -3.30E-05 | -2.1696168 | -2.1855484 | -0.0159316 |
| -0.0034483 | -6.15E-05 | -2.1696168 | -2.1982595 | -0.0286427 |
| -0.0034261 | -3.93E-05 | -2.1696168 | -2.1976458 | -0.028029 |
| -0.0034024 | -1.55E-05 | -2.1696168 | -2.1948413 | -0.0252244 |
| -0.0033834 | 3.43E-06 | -2.1696168 | -2.1742854 | -0.0046686 |
| -0.0034029 | -1.61E-05 | -2.1696168 | -2.1951237 | -0.0255069 |
| -0.003426 | -3.92E-05 | -2.1696168 | -2.1989293 | -0.0293125 |
| -0.0034087 | -2.18E-05 | -2.1696168 | -2.1769933 | -0.0073764 |

| | | | | |
|---|---|---|---|---|
| -0.0033872 | -3.50E-07 | -2.1696168 | -2.1874229 | -0.0178061 |
| -0.0033692 | 1.77E-05 | -2.1696168 | -2.1899061 | -0.0202893 |
| -0.0009297 | -2.48E-06 | -3.9869752 | -4.0126865 | -0.0257113 |
| -0.0004648 | -4.51E-07 | -5.0274558 | -5.0359053 | -0.0084495 |
| -0.0004664 | -2.04E-06 | -5.0274558 | -5.0369311 | -0.0094753 |
| -0.0004649 | -6.15E-07 | -5.0274558 | -5.0329111 | -0.0054553 |
| -0.0004663 | -1.98E-06 | -5.0274558 | -5.0467866 | -0.0193308 |
| -0.0004648 | -4.53E-07 | -5.0274558 | -5.0582082 | -0.0307524 |
| -0.0008014 | -1.64E-07 | -0.9612685 | -0.9684643 | -0.0071958 |
| -0.0020312 | -8.71E-05 | -1.0714771 | -1.1977351 | -0.126258 |
| -0.0040835 | 1.50E-06 | -4.5495367 | -4.5457981 | 0.00373864 |
| -0.0027223 | -1.43E-05 | -1.8184353 | -1.8290734 | -0.0106381 |
| -0.0027123 | -4.26E-06 | -1.8184353 | -1.8260676 | -0.0076323 |
| -0.0069301 | -0.0002377 | -3.4321432 | -3.5022473 | -0.0701041 |
| -0.0067147 | -2.22E-05 | -3.4321432 | -3.4408425 | -0.0086993 |
| -0.0016994 | -2.30E-06 | -0.7578108 | -0.7753595 | -0.0175487 |
| -0.0035621 | -1.79E-05 | -2.3028153 | -2.3272315 | -0.0244162 |
| -0.003629 | 8.49E-05 | -2.3028153 | -2.3566469 | -0.0538316 |
| -0.0046431 | -0.0001587 | -2.3888377 | -2.4460739 | -0.0572362 |
| -0.0045168 | -3.25E-05 | -2.3888377 | -2.3960043 | -0.0071666 |
| -0.004642 | -0.0001577 | -2.3888377 | -2.5016582 | -0.1128205 |
| -0.0037901 | -4.06E-05 | -1.513676 | -1.5543166 | -0.0406405 |
| -0.0002408 | -5.10E-06 | -1.325442 | -1.4873026 | -0.1618606 |
| -0.0030614 | 1.26E-06 | -1.2722265 | -1.2850339 | -0.0128074 |
| -0.005327 | -1.23E-06 | -2.3013797 | -2.3602325 | -0.0588528 |
| -0.0030778 | 6.79E-06 | -2.6345102 | -2.6255916 | 0.00891852 |
| -0.0030918 | -7.24E-06 | -2.6345102 | -2.6578749 | -0.0233648 |
| -0.0031184 | -3.38E-05 | -2.6345102 | -2.6977197 | -0.0632095 |
| -0.0031053 | -2.08E-05 | -2.6345102 | -2.6629881 | -0.0284779 |
| -0.0075788 | -6.09E-05 | -4.8850434 | -4.9058392 | -0.0207958 |
| -0.0010565 | -3.40E-08 | -7.0708419 | -7.0804412 | -0.0095993 |
| -0.0010591 | -2.56E-06 | -7.0708419 | -7.090158 | -0.0193161 |
| -0.0188874 | -3.93E-05 | -5.5169268 | -5.5280938 | -0.011167 |
| -0.0008313 | -1.49E-05 | -0.684289 | -0.7134912 | -0.0292022 |
| -0.0045098 | -0.0001033 | -2.0171154 | -2.0976983 | -0.080583 |
| -0.0013937 | -4.93E-06 | -0.9565216 | -0.9682726 | -0.0117509 |
| -0.0014114 | -2.27E-05 | -0.9565216 | -1.0247434 | -0.0682217 |
| -0.0025427 | -2.32E-05 | -1.5747667 | -1.6051734 | -0.0304067 |
| -0.0002403 | -3.68E-06 | -1.0535637 | -1.0785337 | -0.02497 |
| -0.0024331 | -4.40E-06 | -0.5462767 | -0.5599455 | -0.0136689 |
| -0.0024345 | -5.79E-06 | -0.5462767 | -0.5734825 | -0.0272058 |
| -0.001439 | -1.69E-06 | -7.8185436 | -7.8257231 | -0.0071795 |
| -0.0035463 | -2.62E-05 | -1.7015079 | -1.716794 | -0.0152861 |
| -0.005151 | -0.0001112 | -2.638581 | -2.6820251 | -0.043444 |
| -0.0051859 | -0.0001461 | -2.638581 | -2.7002553 | -0.0616742 |

| | | | | |
|---|---|---|---|---|
| -0.0052583 | -0.0002185 | -2.638581 | -2.7061999 | -0.0676188 |
| -0.0049684 | 7.14E-05 | -2.638581 | -2.6181026 | 0.02047849 |
| -0.0049918 | 4.80E-05 | -2.638581 | -2.6502907 | -0.0117097 |
| -0.0026394 | -1.33E-06 | -1.7291405 | -1.7297158 | -0.0005753 |
| -0.0022366 | -3.56E-06 | -0.9272602 | -0.9495686 | -0.0223083 |
| -0.0005944 | -4.09E-06 | -3.3412147 | -3.3653787 | -0.024164 |
| -0.0019162 | 1.51E-07 | -9.8785892 | -9.879168 | -0.0005789 |
| -0.0045525 | 1.90E-05 | -2.5116304 | -2.5072314 | 0.00439906 |
| -0.0001546 | -2.97E-07 | -1.4789754 | -1.4936187 | -0.0146433 |
| -0.000155 | -7.49E-07 | -1.4789754 | -1.4953582 | -0.0163828 |
| -0.0045159 | -4.41E-05 | -2.1787543 | -2.2161884 | -0.037434 |
| -0.0044663 | -2.39E-05 | -3.5233126 | -3.5393151 | -0.0160025 |
| -0.0014842 | -3.58E-07 | -24.903048 | -24.902229 | 0.00081873 |
| -0.0014835 | 3.37E-07 | -24.903048 | -24.899154 | 0.00389433 |
| -0.0009638 | -1.11E-05 | -5.0646605 | -5.075349 | -0.0106885 |
| -0.0016339 | -1.97E-05 | -2.9709574 | -2.9972404 | -0.026283 |
| -0.0006841 | -2.13E-05 | -6.4183677 | -6.449926 | -0.0315584 |
| -0.0012655 | -5.99E-06 | -1.6380005 | -1.6414738 | -0.0034732 |
| -0.0015885 | -1.19E-06 | -8.0951571 | -8.1132967 | -0.0181396 |
| -0.0015915 | -4.18E-06 | -8.0951571 | -8.100342 | -0.0051849 |
| -0.0010297 | -5.48E-06 | -0.9477808 | -0.9624424 | -0.0146616 |
| -0.0010412 | -1.69E-05 | -0.9477808 | -0.9833105 | -0.0355296 |
| -0.0172389 | -3.58E-05 | -5.7369568 | -5.7508795 | -0.0139227 |
| -0.0037858 | -0.0001826 | -2.6077371 | -2.6802868 | -0.0725497 |
| -0.0037154 | -0.0001122 | -2.6077371 | -2.6959804 | -0.0882434 |
| -0.0004756 | -1.34E-05 | -1.8096185 | -1.8263183 | -0.0166998 |
| -0.0152934 | -2.43E-05 | -5.773191 | -5.7745426 | -0.0013516 |
| -0.0027822 | -9.20E-06 | -2.3869611 | -2.3992813 | -0.0123202 |
| -0.0011429 | -2.78E-05 | -0.5937798 | -0.62664 | -0.0328602 |
| -0.000475 | -7.21E-06 | -2.401434 | -2.4237048 | -0.0222708 |
| -0.0009997 | -1.28E-06 | -3.9409951 | -3.9580669 | -0.0170718 |
| -0.0010009 | -2.55E-06 | -3.9409951 | -3.980887 | -0.0398919 |
| -0.0017932 | -4.52E-05 | -1.4710156 | -1.5269802 | -0.0559646 |
| -0.0030445 | -1.34E-05 | -0.7639301 | -0.9358685 | -0.1719384 |
| -0.003037 | -5.99E-06 | -0.7639301 | -0.8427097 | -0.0787796 |
| -0.0030643 | -3.33E-05 | -0.7639301 | -0.8571676 | -0.0932375 |
| -0.0030176 | 1.34E-05 | -0.7639301 | -0.8666672 | -0.1027372 |
| -0.0030487 | -1.76E-05 | -0.7639301 | -0.8464371 | -0.082507 |
| -0.0030882 | -5.71E-05 | -0.7639301 | -0.8847151 | -0.120785 |
| -0.0030382 | -7.11E-06 | -0.7639301 | -0.7888854 | -0.0249553 |
| -0.0031314 | -0.0001004 | -0.7639301 | -0.8275325 | -0.0636024 |
| -0.0030691 | -3.80E-05 | -0.7639301 | -0.8231223 | -0.0591923 |
| -0.0030313 | -2.80E-07 | -0.7639301 | -0.8041485 | -0.0402184 |
| -0.0030142 | 1.68E-05 | -0.7639301 | -0.7943667 | -0.0304367 |
| -0.002997 | 3.40E-05 | -0.7639301 | -0.7683506 | -0.0044206 |

| | | | | |
|---|---|---|---|---|
| -0.0030834 | -5.24E-05 | -0.7639301 | -0.8208835 | -0.0569535 |
| -0.0030794 | -4.83E-05 | -0.7639301 | -0.8079137 | -0.0439837 |
| -0.0029888 | 4.22E-05 | -0.7639301 | -0.7815076 | -0.0175776 |
| -0.0030822 | -5.12E-05 | -0.7639301 | -0.7943377 | -0.0304076 |
| -0.0031923 | -0.0001612 | -0.7639301 | -0.8471354 | -0.0832054 |
| -0.0031084 | -7.73E-05 | -0.7639301 | -0.8150746 | -0.0511445 |
| -0.003085 | -5.39E-05 | -0.7639301 | -0.8114003 | -0.0474703 |
| -0.0031219 | -9.08E-05 | -0.7639301 | -0.8225337 | -0.0586036 |
| -0.0030948 | -6.37E-05 | -0.7639301 | -0.8122744 | -0.0483443 |
| -0.0030903 | -5.93E-05 | -0.7639301 | -0.8224536 | -0.0585236 |
| -0.0031402 | -0.0001092 | -0.7639301 | -0.8204493 | -0.0565193 |
| -0.003014 | 1.70E-05 | -0.7639301 | -0.7678967 | -0.0039667 |
| -0.0030952 | -6.41E-05 | -0.7639301 | -0.8277803 | -0.0638502 |
| -0.0031327 | -0.0001016 | -0.7639301 | -0.8142647 | -0.0503346 |
| -0.0030664 | -3.54E-05 | -0.7639301 | -0.798694 | -0.0347639 |
| -0.0031237 | -9.26E-05 | -0.7639301 | -0.8258938 | -0.0619637 |
| -0.0031381 | -0.0001071 | -0.7639301 | -0.8307079 | -0.0667779 |
| -0.003131 | -1.00E-04 | -0.7639301 | -0.8418497 | -0.0779197 |
| -0.0032564 | -0.0002253 | -0.7639301 | -0.8722758 | -0.1083458 |
| -0.0032549 | -0.0002238 | -0.7639301 | -0.8488629 | -0.0849328 |
| -0.0030478 | -1.68E-05 | -0.7639301 | -0.8054032 | -0.0414732 |
| -0.0031438 | -0.0001127 | -0.7639301 | -0.8124847 | -0.0485546 |
| -0.0031416 | -0.0001105 | -0.7639301 | -0.8355337 | -0.0716036 |
| -0.0030959 | -6.48E-05 | -0.7639301 | -0.8251744 | -0.0612444 |
| -0.0033068 | -7.00E-06 | -2.910821 | -2.919492 | -0.008671 |
| -0.0035822 | -4.06E-05 | -2.2975897 | -2.3276248 | -0.0300351 |
| -0.0003427 | -7.23E-06 | -2.2960039 | -2.3312213 | -0.0352174 |
| -0.0003479 | -1.25E-05 | -2.2960039 | -2.3188408 | -0.0228369 |
| -0.0003459 | -1.05E-05 | -2.2960039 | -2.3279658 | -0.0319619 |
| -0.0003387 | -3.28E-06 | -2.2960039 | -2.3028603 | -0.0068564 |
| -0.0035753 | -9.29E-06 | -3.6944085 | -3.7034751 | -0.0090666 |
| -0.0009603 | -4.83E-05 | -0.4380997 | -0.5259381 | -0.0878385 |
| -0.001041 | -3.34E-06 | -5.1459557 | -5.1506643 | -0.0047086 |
| -0.0010386 | -9.73E-07 | -5.1459557 | -5.1510103 | -0.0050546 |
| -0.0010418 | -4.14E-06 | -5.1459557 | -5.1508633 | -0.0049076 |
| -0.0010412 | -3.55E-06 | -5.1459557 | -5.1503867 | -0.004431 |
| -0.0010386 | -9.75E-07 | -5.1459557 | -5.148562 | -0.0026063 |
| -0.0010416 | -3.96E-06 | -5.1459557 | -5.1615335 | -0.0155778 |
| -0.0010417 | -4.04E-06 | -5.1459557 | -5.1542888 | -0.0083331 |
| -0.0010383 | -6.02E-07 | -5.1459557 | -5.150575 | -0.0046194 |
| -0.0010405 | -2.83E-06 | -5.1459557 | -5.1528465 | -0.0068908 |
| -0.0035113 | -5.72E-06 | -2.6226554 | -2.6352743 | -0.0126189 |
| -0.0013068 | 4.60E-07 | -0.9524018 | -0.953017 | -0.0006151 |
| -0.0010573 | -1.70E-05 | -1.5069317 | -1.5350773 | -0.0281456 |
| -0.0026132 | -6.41E-05 | -1.9005635 | -1.9444394 | -0.0438758 |

| | | | | |
|---|---|---|---|---|
| -0.0001801 | -6.91E-06 | -2.1198822 | -2.1343678 | -0.0144856 |
| -0.0071042 | -5.94E-06 | -4.4908752 | -4.4995205 | -0.0086453 |
| -0.0071186 | -2.04E-05 | -4.4908752 | -4.5073252 | -0.0164499 |
| -0.0071293 | -3.10E-05 | -4.4908752 | -4.5566487 | -0.0657735 |
| -0.0071372 | -3.90E-05 | -4.4908752 | -4.5151577 | -0.0242825 |
| -0.011651 | -1.89E-05 | -5.6973722 | -5.7056158 | -0.0082436 |
| -0.0005693 | -1.11E-05 | -0.6869611 | -0.7187499 | -0.0317888 |
| -0.0080406 | -6.15E-05 | -2.3378622 | -2.3911169 | -0.0532547 |
| -0.0027794 | -6.92E-06 | -2.0395138 | -2.0599695 | -0.0204557 |
| -0.0009311 | -9.35E-06 | -0.7234929 | -0.7424633 | -0.0189704 |
| -0.0004482 | 6.55E-07 | -2.7418649 | -2.7448485 | -0.0029836 |
| -0.0047232 | -7.43E-07 | -3.4703749 | -3.4807332 | -0.0103582 |
| -0.0031517 | -2.33E-06 | -3.2397258 | -3.2371608 | 0.00256503 |
| -0.0022981 | -1.28E-06 | -1.7621403 | -1.7845692 | -0.0224289 |
| -0.0023158 | -1.90E-05 | -1.7621403 | -1.780957 | -0.0188166 |
| -0.0037114 | -5.04E-05 | -2.4688362 | -2.507272 | -0.0384358 |
| -0.0007696 | -2.75E-06 | -3.7010784 | -3.7215242 | -0.0204459 |
| -0.0007683 | -1.36E-06 | -3.7010784 | -3.7163547 | -0.0152764 |
| -0.0007685 | -1.65E-06 | -3.7010784 | -3.7129869 | -0.0119085 |
| -0.0007692 | -2.33E-06 | -3.7010784 | -3.7525892 | -0.0515108 |
| -0.0007669 | 3.34E-08 | -3.7010784 | -3.703671 | -0.0025927 |
| -0.000992 | -2.94E-05 | -0.6326177 | -0.6613495 | -0.0287317 |
| -0.0050714 | -1.14E-05 | -3.3852556 | -3.4100015 | -0.0247459 |
| -0.0051213 | -6.13E-05 | -3.3852556 | -3.41664 | -0.0313845 |
| -0.0008696 | -9.04E-06 | -2.7332928 | -2.7779737 | -0.0446808 |
| -0.0004618 | -3.18E-06 | -1.7999096 | -1.8305773 | -0.0306677 |
| -0.0004612 | -2.64E-06 | -1.7999096 | -1.8154328 | -0.0155233 |
| -0.0004616 | -2.96E-06 | -1.7999096 | -1.8366666 | -0.036757 |
| -0.003805 | -9.35E-06 | -0.9404358 | -0.9659359 | -0.0255001 |
| -0.004016 | -1.31E-05 | -0.9075314 | -0.935944 | -0.0284126 |
| -0.0018463 | -1.26E-05 | -0.9276683 | -0.9919566 | -0.0642882 |
| -0.0018424 | -8.71E-06 | -0.9276683 | -0.9466348 | -0.0189664 |
| -0.005236 | -9.87E-05 | -2.3630136 | -2.3621871 | 0.00082648 |
| -0.0008402 | -2.39E-06 | -0.862428 | -0.8942331 | -0.0318051 |
| -0.0022079 | -3.33E-06 | -2.3732115 | -2.381009 | -0.0077975 |
| -0.002467 | -1.90E-06 | -2.685766 | -2.707618 | -0.021852 |
| -0.0024638 | 1.38E-06 | -2.685766 | -2.6852146 | 0.00055134 |
| -0.0152771 | 3.38E-05 | -5.7556026 | -5.7474601 | 0.00814247 |
| -0.0113591 | -5.25E-06 | -4.2692192 | -4.2846057 | -0.0153866 |
| -0.0091229 | 1.96E-05 | -5.1913033 | -5.2127733 | -0.0214701 |
| -0.0020593 | -2.62E-06 | -3.8007272 | -3.8063498 | -0.0056225 |
| -0.0139378 | -0.000109 | -5.5617664 | -5.6066618 | -0.0448954 |
| -0.0138515 | -2.26E-05 | -5.5617664 | -5.5748374 | -0.0130711 |
| -0.0138388 | -9.98E-06 | -5.5617664 | -5.5703173 | -0.0085509 |
| -0.001986 | 5.36E-07 | -1.9768283 | -1.9757153 | 0.001113 |

| | | | | |
|---|---|---|---|---|
| -0.0054804 | -7.13E-05 | -4.2536428 | -4.3094625 | -0.0558197 |
| -0.0054075 | 1.61E-06 | -4.2536428 | -4.2492917 | 0.00435114 |
| -0.0054289 | -1.98E-05 | -4.2536428 | -4.2583358 | -0.004693 |
| -0.0029293 | -1.14E-05 | -1.9877014 | -2.0032766 | -0.0155752 |
| -0.0066591 | -5.17E-06 | -4.6323006 | -4.655935 | -0.0236344 |
| -0.0066453 | 8.66E-06 | -4.6323006 | -4.6301091 | 0.00219154 |
| -0.0066874 | -3.34E-05 | -4.6323006 | -4.6467159 | -0.0144153 |
| -0.0066451 | 8.84E-06 | -4.6323006 | -4.6226223 | 0.00967836 |
| -0.0066611 | -7.11E-06 | -4.6323006 | -4.6336942 | -0.0013936 |
| -0.0066396 | 1.44E-05 | -4.6323006 | -4.6225984 | 0.00970221 |
| -0.0066536 | 3.43E-07 | -4.6323006 | -4.6318514 | 0.00044918 |
| -0.0066492 | 4.81E-06 | -4.6323006 | -4.6212513 | 0.01104927 |
| -0.006664 | -1.00E-05 | -4.6323006 | -4.6345196 | -0.002219 |
| -0.0066609 | -6.90E-06 | -4.6323006 | -4.6349552 | -0.0026546 |
| -0.0066497 | 4.25E-06 | -4.6323006 | -4.621314 | 0.01098657 |
| -0.0066793 | -2.53E-05 | -4.6323006 | -4.6389363 | -0.0066357 |
| -0.0066687 | -1.47E-05 | -4.6323006 | -4.6298306 | 0.00247002 |
| -0.0066736 | -1.97E-05 | -4.6323006 | -4.6564329 | -0.0241323 |
| -0.0066771 | -2.31E-05 | -4.6323006 | -4.6486359 | -0.0163352 |
| -0.0066802 | -2.62E-05 | -4.6323006 | -4.6411097 | -0.0088091 |
| -0.0006678 | -1.26E-06 | -0.3761401 | -0.3882884 | -0.0121483 |
| -0.0006753 | -8.78E-06 | -0.3761401 | -0.3914287 | -0.0152887 |
| -0.000492 | -1.91E-06 | -1.8281673 | -1.8649904 | -0.0368231 |
| -0.001451 | -8.02E-06 | -0.471567 | -0.4981895 | -0.0266225 |
| -0.0014549 | -1.20E-05 | -0.471567 | -0.5059641 | -0.034397 |
| -0.0014436 | -6.80E-07 | -0.471567 | -0.4884243 | -0.0168573 |
| -0.0008862 | -1.17E-05 | -0.7929353 | -0.7972625 | -0.0043272 |
| -0.0049086 | 1.23E-05 | -4.2811604 | -4.2791626 | 0.00199771 |
| -0.0064717 | 1.20E-06 | -5.1953313 | -5.2007217 | -0.0053904 |
| -0.0044834 | -1.47E-05 | -2.5967596 | -2.6510245 | -0.0542649 |
| -0.0012751 | -3.72E-06 | -5.0329672 | -5.0641057 | -0.0311385 |
| -0.0018174 | 3.93E-06 | -1.968936 | -1.9674031 | 0.00153285 |
| -0.0018307 | -9.41E-06 | -1.968936 | -1.990282 | -0.021346 |
| -0.0014966 | -2.59E-05 | -0.5590315 | -0.5912526 | -0.0322212 |
| -0.0041008 | -2.12E-05 | -1.820273 | -1.8292862 | -0.0090131 |
| -0.0054672 | -7.61E-05 | -2.9584706 | -2.9966843 | -0.0382137 |
| -0.0053645 | 2.66E-05 | -2.9584706 | -2.9441473 | 0.01432323 |
| -0.0054013 | -1.02E-05 | -2.9584706 | -2.9714475 | -0.0129769 |
| -0.0053986 | -7.49E-06 | -2.9584706 | -2.936597 | 0.02187359 |
| -0.0053873 | 3.81E-06 | -2.9584706 | -2.9590614 | -0.0005908 |
| -0.0016613 | -9.85E-06 | -0.4332443 | -0.4607666 | -0.0275223 |
| -0.0016616 | -1.01E-05 | -0.4332443 | -0.4533439 | -0.0200997 |
| -0.0017159 | 2.03E-06 | -1.5206687 | -1.5154649 | 0.00520378 |
| -0.0017223 | -4.39E-06 | -1.5206687 | -1.5342629 | -0.0135942 |
| -0.0017202 | -2.21E-06 | -1.5206687 | -1.5368347 | -0.016166 |

| | | | | |
|---|---|---|---|---|
| -0.0017227 | -4.72E-06 | -1.5206687 | -1.529933 | -0.0092643 |
| -0.0017186 | -6.94E-07 | -1.5206687 | -1.5344458 | -0.0137771 |
| -0.0017244 | -6.46E-06 | -1.5206687 | -1.5479137 | -0.027245 |
| -0.0017232 | -5.28E-06 | -1.5206687 | -1.5320539 | -0.0113852 |
| -0.0017185 | -5.52E-07 | -1.5206687 | -1.5277492 | -0.0070805 |
| -0.0017235 | -5.59E-06 | -1.5206687 | -1.5425641 | -0.0218954 |
| -0.0013924 | -6.50E-06 | -2.1424934 | -2.1596909 | -0.0171975 |
| -0.0014022 | -1.63E-05 | -2.1424934 | -2.1767325 | -0.0342392 |
| -0.0019642 | -1.06E-05 | -2.1803623 | -2.1946723 | -0.01431 |
| -0.001955 | -1.36E-06 | -2.1803623 | -2.1796793 | 0.00068307 |
| -0.0031403 | 3.44E-05 | -1.6977017 | -1.7585671 | -0.0608654 |
| -0.0036909 | 2.87E-07 | -2.1532099 | -2.1511445 | 0.00206542 |
| -0.0196617 | -0.0001169 | -5.7784798 | -5.8021338 | -0.023654 |
| -0.0032021 | -5.94E-05 | -2.8066473 | -2.8981769 | -0.0915296 |
| -0.0031746 | -3.19E-05 | -2.8066473 | -2.8363506 | -0.0297033 |
| -0.0037419 | -7.59E-06 | -4.4451437 | -4.450218 | -0.0050743 |
| -0.0047997 | -4.64E-05 | -2.0469307 | -2.117374 | -0.0704433 |
| -0.0038208 | 1.96E-06 | -4.1691189 | -4.1686571 | 0.00046182 |
| -0.0007037 | 4.87E-06 | -0.3868574 | -0.4015357 | -0.0146783 |
| -0.0053644 | -8.50E-05 | -3.9419196 | -4.034318 | -0.0923984 |
| -0.005242 | 3.73E-05 | -3.9419196 | -3.9578635 | -0.0159439 |
| -0.0034024 | 1.16E-06 | -3.5949917 | -3.5950795 | -8.79E-05 |
| -0.0034039 | -3.33E-07 | -3.5949917 | -3.5967965 | -0.0018048 |
| -0.0034017 | 1.87E-06 | -3.5949917 | -3.5961306 | -0.0011389 |
| -0.0034117 | -8.17E-06 | -3.5949917 | -3.6042851 | -0.0092934 |
| -0.0028615 | -2.30E-05 | -1.0910285 | -1.1269784 | -0.0359498 |
| -0.005002 | -5.76E-06 | -3.536881 | -3.5433372 | -0.0064563 |
| -0.0011088 | 3.73E-07 | -6.5867815 | -6.5787601 | 0.00802135 |
| -0.0007514 | -2.79E-06 | -0.792472 | -0.7993774 | -0.0069053 |
| -0.0007485 | 9.29E-08 | -0.792472 | -0.7930204 | -0.0005484 |
| -0.0038461 | -6.20E-06 | -3.9030195 | -3.9104183 | -0.0073987 |
| -0.003839 | 9.49E-07 | -3.9030195 | -3.9052767 | -0.0022571 |
| -0.0038477 | -7.75E-06 | -3.9030195 | -3.9180812 | -0.0150616 |
| -0.0038603 | -2.04E-05 | -3.9030195 | -3.9287547 | -0.0257351 |
| -0.0042699 | -7.77E-06 | -3.765528 | -3.7735192 | -0.0079912 |
| -0.0006336 | -2.42E-06 | -4.6312806 | -4.6479481 | -0.0166675 |
| -0.0025602 | -5.87E-06 | -1.989177 | -2.0015798 | -0.0124028 |
| -0.0050162 | 2.36E-05 | -2.638581 | -2.6375679 | 0.00101316 |
| -0.005151 | -0.0001112 | -2.638581 | -2.6820251 | -0.043444 |
| -0.0051859 | -0.0001461 | -2.638581 | -2.7002553 | -0.0616742 |
| -0.0052583 | -0.0002185 | -2.638581 | -2.7061999 | -0.0676188 |
| -0.0049684 | 7.14E-05 | -2.638581 | -2.6181026 | 0.02047849 |
| -0.0049918 | 4.80E-05 | -2.638581 | -2.6502907 | -0.0117097 |
| -0.0004011 | -2.88E-06 | -1.1399609 | -1.1645483 | -0.0245874 |
| -0.0009191 | -3.69E-06 | -2.2683671 | -2.279692 | -0.0113248 |

| | | | | |
|---|---|---|---|---|
| -0.0009255 | -1.01E-05 | -2.2683671 | -2.4218705 | -0.1535034 |
| -0.0009065 | -4.40E-06 | -2.0653144 | -2.0807258 | -0.0154114 |
| -0.0020391 | -2.77E-05 | -1.8010639 | -1.8287292 | -0.0276653 |
| -0.0024053 | -4.20E-05 | -1.8864185 | -1.9277239 | -0.0413055 |
| -0.0023923 | -2.89E-05 | -1.8864185 | -1.9374799 | -0.0510614 |
| -0.0023924 | -2.90E-05 | -1.8864185 | -1.9273024 | -0.0408839 |
| -0.0024072 | -4.39E-05 | -1.8864185 | -1.9521259 | -0.0657074 |
| -0.0023954 | -3.20E-05 | -1.8864185 | -1.9368936 | -0.0504752 |
| -0.0023857 | -2.23E-05 | -1.8864185 | -1.9265212 | -0.0401027 |
| -0.0008881 | 1.28E-06 | -2.7815961 | -2.7788619 | 0.00273415 |
| -0.0024281 | -3.76E-06 | -1.6561757 | -1.6666728 | -0.0104972 |
| -0.0078859 | -3.01E-05 | -4.740658 | -4.7495806 | -0.0089226 |
| -0.0018516 | -2.96E-05 | -0.5895518 | -0.6311555 | -0.0416037 |
| -0.0043381 | 2.39E-05 | -2.5686424 | -2.5547811 | 0.0138613 |
| -0.0043258 | 3.63E-05 | -2.5686424 | -2.5687637 | -0.0001214 |
| -0.0044251 | -6.31E-05 | -2.5686424 | -2.5939099 | -0.0252675 |
| -0.0043356 | 2.65E-05 | -2.5686424 | -2.5719402 | -0.0032978 |
| -0.0043615 | 5.90E-07 | -2.5686424 | -2.5697343 | -0.001092 |
| -0.0043656 | -3.52E-06 | -2.5686424 | -2.578428 | -0.0097857 |
| -0.0043567 | 5.34E-06 | -2.5686424 | -2.6155565 | -0.0469141 |
| -0.0043782 | -1.61E-05 | -2.5686424 | -2.5958188 | -0.0271764 |
| -0.0044288 | -6.67E-05 | -2.5686424 | -2.6553372 | -0.0866948 |
| -0.0044307 | -6.87E-05 | -2.5686424 | -2.6658294 | -0.097187 |
| -0.0043442 | 1.79E-05 | -2.5686424 | -2.5725857 | -0.0039433 |
| -0.0046556 | -0.0002935 | -2.5686424 | -2.7055393 | -0.136897 |
| -0.0044105 | -4.85E-05 | -2.5686424 | -2.6221038 | -0.0534614 |
| -0.0044814 | -0.0001193 | -2.5686424 | -2.6531942 | -0.0845518 |
| -0.004458 | -9.60E-05 | -2.5686424 | -2.6666151 | -0.0979728 |
| -0.0043921 | -3.01E-05 | -2.5686424 | -2.5936058 | -0.0249634 |
| -0.0044646 | -0.0001025 | -2.5686424 | -2.6396292 | -0.0709869 |
| -0.0044071 | -4.50E-05 | -2.5686424 | -2.5968746 | -0.0282322 |
| -0.0044456 | -8.35E-05 | -2.5686424 | -2.6210585 | -0.0524161 |
| -0.0043756 | -1.35E-05 | -2.5686424 | -2.5784849 | -0.0098425 |
| -0.0043747 | -1.26E-05 | -2.5686424 | -2.5930367 | -0.0243943 |
| -0.0044156 | -5.35E-05 | -2.5686424 | -2.6036634 | -0.0350211 |
| -0.0043915 | -2.94E-05 | -2.5686424 | -2.6097333 | -0.041091 |
| -0.0044443 | -8.22E-05 | -2.5686424 | -2.6331971 | -0.0645547 |
| -0.0043452 | 1.69E-05 | -2.5686424 | -2.5605656 | 0.00807679 |
| -0.0044232 | -6.12E-05 | -2.5686424 | -2.6425612 | -0.0739188 |
| -0.0044593 | -9.73E-05 | -2.5686424 | -2.6498657 | -0.0812234 |
| -0.0044311 | -6.90E-05 | -2.5686424 | -2.6200171 | -0.0513747 |
| -0.0044727 | -0.0001106 | -2.5686424 | -2.6430818 | -0.0744394 |
| -0.0044506 | -8.85E-05 | -2.5686424 | -2.6065024 | -0.03786 |
| -0.0044187 | -5.66E-05 | -2.5686424 | -2.6057353 | -0.0370929 |
| -0.0043681 | -6.07E-06 | -2.5686424 | -2.592132 | -0.0234896 |

| | | | | |
|---|---|---|---|---|
| -0.0043919 | -2.98E-05 | -2.5686424 | -2.5979346 | -0.0292922 |
| -0.0044249 | -6.28E-05 | -2.5686424 | -2.6107354 | -0.042093 |
| -0.0044129 | -5.08E-05 | -2.5686424 | -2.6316959 | -0.0630535 |
| -0.004454 | -9.19E-05 | -2.5686424 | -2.6461061 | -0.0774637 |
| -0.0044022 | -4.01E-05 | -2.5686424 | -2.5947043 | -0.0260619 |
| -0.004408 | -4.59E-05 | -2.5686424 | -2.603423 | -0.0347806 |
| -0.0044001 | -3.81E-05 | -2.5686424 | -2.58788 | -0.0192376 |
| -0.0044151 | -5.30E-05 | -2.5686424 | -2.6100168 | -0.0413744 |
| -0.0044265 | -6.44E-05 | -2.5686424 | -2.6057217 | -0.0370793 |
| -0.0043874 | -2.54E-05 | -2.5686424 | -2.5881726 | -0.0195302 |
| -0.0044558 | -9.38E-05 | -2.5686424 | -2.6668559 | -0.0982136 |
| -0.0045499 | -0.0001879 | -2.5686424 | -2.712006 | -0.1433636 |
| -0.0043964 | -3.43E-05 | -2.5686424 | -2.5858998 | -0.0172575 |
| -0.0043907 | -2.86E-05 | -2.5686424 | -2.6074286 | -0.0387862 |
| -0.0044132 | -5.11E-05 | -2.5686424 | -2.5911547 | -0.0225123 |
| -0.0043725 | -1.04E-05 | -2.5686424 | -2.5897589 | -0.0211165 |
| -0.0043711 | -9.06E-06 | -2.5686424 | -2.5786318 | -0.0099894 |
| -0.0044101 | -4.80E-05 | -2.5686424 | -2.6331323 | -0.06449 |
| -0.0044022 | -4.01E-05 | -2.5686424 | -2.6236557 | -0.0550133 |
| -0.0043574 | 4.72E-06 | -2.5686424 | -2.5614135 | 0.00722885 |
| -0.0043474 | 1.47E-05 | -2.5686424 | -2.5617522 | 0.00689018 |
| -0.0043774 | -1.53E-05 | -2.5686424 | -2.5871584 | -0.0185161 |
| -0.004392 | -2.99E-05 | -2.5686424 | -2.5847696 | -0.0161272 |
| -0.0043679 | -5.87E-06 | -2.5686424 | -2.6031332 | -0.0344908 |
| -0.004384 | -2.19E-05 | -2.5686424 | -2.5929112 | -0.0242689 |
| -0.0043851 | -2.30E-05 | -2.5686424 | -2.5861436 | -0.0175012 |
| -0.0043729 | -1.09E-05 | -2.5686424 | -2.5861926 | -0.0175502 |
| -0.0044215 | -5.94E-05 | -2.5686424 | -2.6043451 | -0.0357027 |
| -0.0043963 | -3.42E-05 | -2.5686424 | -2.5985923 | -0.0299499 |
| -0.004394 | -3.19E-05 | -2.5686424 | -2.6058683 | -0.037226 |
| -0.0044186 | -5.65E-05 | -2.5686424 | -2.6146476 | -0.0460052 |
| -0.0044299 | -6.78E-05 | -2.5686424 | -2.6433229 | -0.0746806 |
| -0.0043748 | -1.28E-05 | -2.5686424 | -2.5714266 | -0.0027843 |
| -0.0044146 | -5.25E-05 | -2.5686424 | -2.6106061 | -0.0419637 |
| -0.0045047 | -0.0001426 | -2.5686424 | -2.6503041 | -0.0816617 |
| -0.0043918 | -2.97E-05 | -2.5686424 | -2.5960916 | -0.0274493 |
| -0.0044134 | -5.13E-05 | -2.5686424 | -2.6162877 | -0.0476453 |
| -0.0043663 | -4.20E-06 | -2.5686424 | -2.5695554 | -0.000913 |
| -0.0044122 | -5.01E-05 | -2.5686424 | -2.5983531 | -0.0297108 |
| -0.0043738 | -1.17E-05 | -2.5686424 | -2.5899186 | -0.0212762 |
| -0.00435 | 1.20E-05 | -2.5686424 | -2.576969 | -0.0083266 |
| -0.0044352 | -7.32E-05 | -2.5686424 | -2.6454519 | -0.0768095 |
| -0.0044013 | -3.92E-05 | -2.5686424 | -2.6153582 | -0.0467159 |
| -0.004412 | -4.99E-05 | -2.5686424 | -2.6074868 | -0.0388445 |
| -0.0043693 | -7.27E-06 | -2.5686424 | -2.5841199 | -0.0154775 |

| | | | | |
|---|---|---|---|---|
| -0.0043999 | -3.78E-05 | -2.5686424 | -2.6023344 | -0.033692 |
| -0.0043922 | -3.01E-05 | -2.5686424 | -2.5959238 | -0.0272814 |
| -0.0043983 | -3.62E-05 | -2.5686424 | -2.6148814 | -0.046239 |
| -0.0043561 | 5.94E-06 | -2.5686424 | -2.5516971 | 0.01694524 |
| -0.004381 | -1.89E-05 | -2.5686424 | -2.5864824 | -0.01784 |
| -0.0043906 | -2.86E-05 | -2.5686424 | -2.5715021 | -0.0028597 |
| -0.0043981 | -3.61E-05 | -2.5686424 | -2.5952302 | -0.0265878 |
| -0.0043636 | -1.55E-06 | -2.5686424 | -2.5748897 | -0.0062473 |
| -0.0043753 | -1.33E-05 | -2.5686424 | -2.588446 | -0.0198036 |
| -0.0043756 | -1.36E-05 | -2.5686424 | -2.5923859 | -0.0237435 |
| -0.0043579 | 4.17E-06 | -2.5686424 | -2.5678672 | 0.00077522 |
| -0.0043691 | -7.07E-06 | -2.5686424 | -2.5705994 | -0.0019571 |
| -0.0043497 | 1.23E-05 | -2.5686424 | -2.566364 | 0.00227833 |
| -0.0043659 | -3.81E-06 | -2.5686424 | -2.5712719 | -0.0026295 |
| -0.0043592 | 2.84E-06 | -2.5686424 | -2.5689592 | -0.0003169 |
| -0.0043598 | 2.24E-06 | -2.5686424 | -2.5725474 | -0.0039051 |
| -0.0043644 | -2.30E-06 | -2.5686424 | -2.5685767 | 6.57E-05 |
| -0.0043639 | -1.80E-06 | -2.5686424 | -2.5710866 | -0.0024443 |
| -0.0017937 | -2.28E-05 | -2.03149 | -2.0608676 | -0.0293776 |
| -0.0012195 | -2.75E-06 | -0.8377688 | -0.8487431 | -0.0109743 |
| -0.0012281 | -1.14E-05 | -0.8377688 | -0.8537588 | -0.0159901 |
| -0.0041518 | -5.31E-05 | -2.1969886 | -2.2311136 | -0.034125 |
| -0.00204 | -1.93E-05 | -1.1381428 | -1.1873072 | -0.0491644 |
| -0.0059698 | -5.24E-05 | -2.5214831 | -2.5825021 | -0.0610191 |
| -0.0034071 | -1.99E-06 | -4.5323098 | -4.529896 | 0.00241375 |
| -0.0034084 | -3.32E-06 | -4.5323098 | -4.5517404 | -0.0194306 |
| -0.0034006 | 4.55E-06 | -4.5323098 | -4.5274477 | 0.00486207 |
| -0.0034072 | -2.10E-06 | -4.5323098 | -4.5403891 | -0.0080793 |
| -0.0034023 | 2.79E-06 | -4.5323098 | -4.5419652 | -0.0096555 |
| -0.0034083 | -3.24E-06 | -4.5323098 | -4.5379543 | -0.0056446 |
| -0.0034052 | -1.27E-07 | -4.5323098 | -4.5422137 | -0.0099039 |
| -0.0034067 | -1.63E-06 | -4.5323098 | -4.5410137 | -0.0087039 |
| -0.003414 | -8.90E-06 | -4.5323098 | -4.5484788 | -0.0161691 |
| -0.0013578 | -4.25E-06 | -1.4492629 | -1.455106 | -0.005843 |
| -0.0013536 | 2.11E-06 | -1.0606045 | -1.0617855 | -0.0011811 |
| -0.0013642 | -8.48E-06 | -1.0606045 | -1.0806673 | -0.0200628 |
| -0.0013614 | -5.72E-06 | -1.0606045 | -1.072717 | -0.0121125 |
| -0.0027318 | -2.72E-05 | -3.9737203 | -4.0207973 | -0.0470769 |
| -0.002354 | -6.52E-06 | -3.3864609 | -3.3973904 | -0.0109295 |
| -0.0023555 | -8.01E-06 | -3.3864609 | -3.3887185 | -0.0022576 |
| -0.0023537 | -6.17E-06 | -3.3864609 | -3.4010315 | -0.0145706 |
| -0.0023609 | -1.34E-05 | -3.3864609 | -3.4087669 | -0.022306 |
| -0.0023484 | -8.57E-07 | -3.3864609 | -3.3903 | -0.0038391 |
| Progen2_sco | Progen2_fina | RITA_score_v | RITA_score_r | RITA_final_score |

| | | | | |
|---|---|---|---|---|
| -0.0008882 | 2.50E-07 | -6.4390417 | -6.439102 | -6.03E-05 |
| -0.000337 | 8.59E-07 | -2.3434079 | -2.3361149 | 0.00729305 |
| -0.0018649 | 2.92E-05 | -0.9718568 | -0.9504507 | 0.02140611 |
| -0.001331 | 7.89E-06 | -9.7287054 | -9.6813532 | 0.04735219 |
| -0.0017215 | 2.16E-06 | -1.8331247 | -1.8070011 | 0.02612364 |
| -0.0010408 | 6.72E-06 | -1.0097367 | -0.9870164 | 0.02272028 |
| -0.0011926 | 8.72E-07 | -1.65412 | -1.6271994 | 0.02692068 |
| -0.0018374 | 2.50E-06 | -3.3894198 | -3.3836411 | 0.00577867 |
| -0.00144 | 1.10E-06 | -10.09513 | -10.087838 | 0.00729233 |
| -0.0021789 | -3.01E-07 | -16.271221 | -16.260752 | 0.01046944 |
| -0.0014101 | 1.53E-05 | -1.3045034 | -1.2847674 | 0.01973599 |
| -0.002395 | 7.92E-06 | -3.2212473 | -3.2066203 | 0.01462698 |
| -0.0029276 | 1.18E-05 | -2.8073002 | -2.7902782 | 0.01702201 |
| -0.0019391 | 2.40E-06 | -3.2303723 | -3.2221088 | 0.00826347 |
| -0.0009995 | 4.86E-07 | -1.3263996 | -1.324002 | 0.00239754 |
| -0.00123 | -5.65E-07 | -9.4428061 | -9.4480748 | -0.0052687 |
| -0.0027329 | 3.28E-06 | -4.2898488 | -4.2859974 | 0.00385141 |
| -0.0011763 | -3.57E-07 | -3.6040708 | -3.6126527 | -0.0085818 |
| -0.000586 | 6.57E-07 | -2.9197316 | -2.9008079 | 0.01892376 |
| -0.0004165 | 8.24E-07 | -2.6504609 | -2.6455849 | 0.00487608 |
| -0.0009286 | 2.57E-07 | -23.284598 | -23.279563 | 0.00503469 |
| -0.0013381 | 1.14E-05 | -1.2852365 | -1.2675939 | 0.01764262 |
| -0.0008829 | 7.65E-07 | -6.6892018 | -6.6769469 | 0.01225495 |
| -0.0013908 | 2.91E-05 | -1.3852651 | -1.3514793 | 0.03378576 |
| -0.0010184 | -8.83E-06 | -6.5544376 | -6.5716864 | -0.0172487 |
| -0.0007389 | 1.63E-06 | -16.240251 | -16.221931 | 0.01831996 |
| -0.0013576 | 3.57E-06 | -9.8017685 | -9.7773341 | 0.02443445 |
| -0.0020177 | 2.01E-05 | -1.5629956 | -1.5410077 | 0.02198786 |
| -0.0008304 | 1.90E-05 | -0.8019912 | -0.7571612 | 0.04482993 |
| -0.0003956 | 3.31E-06 | -5.2172873 | -5.1853416 | 0.03194577 |
| -0.0005152 | 8.00E-07 | -3.963347 | -3.9536018 | 0.00974524 |
| -0.0040819 | 2.53E-06 | -3.3413576 | -3.3216872 | 0.01967037 |
| -0.0010658 | 4.83E-06 | -7.474027 | -7.4615392 | 0.01248783 |
| -0.0012261 | 3.57E-08 | -9.3787434 | -9.3799174 | -0.001174 |
| -0.001142 | 5.19E-06 | -1.6517534 | -1.6461651 | 0.00558829 |
| -0.0007179 | 3.31E-05 | -0.7203402 | -0.6776901 | 0.04265013 |
| -0.001439 | 1.52E-07 | -9.6503721 | -9.6497658 | 0.0006063 |
| -0.0003859 | 1.06E-05 | -2.9767831 | -2.9389171 | 0.037866 |
| -0.0009918 | -8.73E-07 | -2.8514222 | -2.8560442 | -0.004622 |
| -0.0009397 | 1.09E-05 | -6.3694062 | -6.3472135 | 0.02219266 |
| -0.0011494 | -3.17E-06 | -0.8055149 | -0.8083717 | -0.0028568 |
| -0.0040317 | 1.94E-06 | -2.2792813 | -2.2467211 | 0.03256011 |
| -0.0007252 | 1.36E-06 | -0.769554 | -0.7652974 | 0.00425655 |
| -0.0011063 | 1.21E-05 | -1.1268061 | -1.1061676 | 0.02063853 |
| -0.0006394 | 9.36E-06 | -3.7080747 | -3.6888341 | 0.01924065 |

| | | | | |
|---|---|---|---|---|
| -0.0009682 | -3.16E-07 | -6.6496345 | -6.6524702 | -0.0028358 |
| -0.0008228 | 3.95E-07 | -2.5503785 | -2.5473519 | 0.00302655 |
| -0.0009247 | 5.40E-07 | -3.6778395 | -3.6546465 | 0.023193 |
| -0.0024683 | 7.56E-07 | -4.9960942 | -4.994617 | 0.00147724 |
| -0.001314 | -4.37E-07 | -1.8609445 | -1.8630245 | -0.00208 |
| -0.0016345 | 1.53E-05 | -2.2494159 | -2.2319114 | 0.01750445 |
| -0.0003752 | 1.46E-06 | -2.5279352 | -2.5257889 | 0.0021463 |
| -0.0026647 | 1.64E-06 | -1.6452838 | -1.6405819 | 0.00470185 |
| -0.0082157 | -7.99E-05 | -2.8870831 | -2.8907764 | -0.0036933 |
| -0.0029989 | 1.17E-07 | -4.2506518 | -4.2139065 | 0.03674531 |
| -0.0054316 | 1.61E-05 | -3.8585483 | -3.8477401 | 0.01080823 |
| -0.0018336 | 6.82E-06 | -2.8712335 | -2.8588655 | 0.01236796 |
| -0.0011457 | 2.92E-06 | -9.3495153 | -9.3311049 | 0.01841038 |
| -0.0005875 | 2.82E-06 | -5.0816348 | -5.0322017 | 0.04943314 |
| -0.0019258 | 2.71E-05 | -1.8361189 | -1.8008912 | 0.03522766 |
| -0.0036127 | 3.46E-06 | -3.6871473 | -3.6805098 | 0.00663745 |
| -0.0039595 | 7.05E-08 | -5.6582158 | -5.6584427 | -0.000227 |
| -0.0038566 | 8.22E-06 | -2.0634927 | -2.0601102 | 0.00338244 |
| -0.0026061 | 5.70E-06 | -0.8595581 | -0.8508215 | 0.00873661 |
| -0.0126512 | 0.00100983 | -3.2667494 | -3.0314156 | 0.2353338 |
| -0.0046465 | -8.88E-07 | -3.1956809 | -3.1860987 | 0.00958216 |
| -0.0053878 | 1.67E-05 | -2.8072529 | -2.8014092 | 0.00584364 |
| -0.0014441 | 3.69E-06 | -5.9076757 | -5.871762 | 0.03591377 |
| -0.0093082 | 2.66E-05 | -4.8694561 | -4.8656783 | 0.00377774 |
| -0.0004093 | 4.13E-06 | -9.9702165 | -9.8971102 | 0.07310635 |
| -0.0014495 | -7.65E-07 | -2.1135607 | -2.1110379 | 0.00252283 |
| -0.0091518 | -1.71E-05 | -3.2048043 | -3.2080948 | -0.0032905 |
| -0.0070991 | 1.33E-05 | -4.7806289 | -4.7924304 | -0.0118015 |
| -0.0037969 | 6.58E-06 | -3.0865149 | -3.0761365 | 0.01037848 |
| -0.006159 | 1.36E-05 | -5.1807091 | -5.1686177 | 0.0120914 |
| -0.0086002 | 0.00011816 | -3.0490208 | -2.9582869 | 0.09073389 |
| -0.0015503 | 3.07E-06 | -12.807871 | -12.748096 | 0.05977476 |
| -0.0016917 | 6.05E-06 | -2.2513477 | -2.2470793 | 0.00426841 |
| -0.0010225 | -2.99E-07 | -7.1350336 | -7.1364101 | -0.0013765 |
| -0.0071841 | 1.12E-05 | -4.1961834 | -4.2230911 | -0.0269077 |
| -0.0030169 | 1.35E-05 | -2.2903301 | -2.274725 | 0.01560509 |
| -0.001807 | 1.87E-06 | -7.5051646 | -7.493505 | 0.01165962 |
| -0.0006704 | 1.74E-06 | -2.5945129 | -2.5757558 | 0.0187571 |
| -0.0112627 | 1.66E-05 | -3.5969728 | -3.6363949 | -0.039422 |
| -0.0029754 | 1.10E-05 | -2.8992147 | -2.8859156 | 0.01329911 |
| -0.001186 | 1.68E-05 | -1.4873038 | -1.4646681 | 0.0226357 |
| -0.0005624 | 2.19E-05 | -3.0392226 | -2.9776095 | 0.06161311 |
| -0.006235 | 9.60E-05 | -3.5429775 | -3.5006919 | 0.04228556 |
| -0.0025932 | -8.98E-06 | -3.1500336 | -3.1447115 | 0.0053221 |
| -0.0033753 | 1.98E-07 | -2.6264769 | -2.6238841 | 0.0025928 |

| | | | | |
|---|---|---|---|---|
| -0.0052725 | -3.97E-05 | -3.4495574 | -3.440449 | 0.00910842 |
| -0.0006105 | 1.36E-05 | -0.9723271 | -0.9451578 | 0.02716926 |
| -0.001745 | -4.03E-07 | -2.2105173 | -2.2070788 | 0.00343847 |
| -0.0012538 | 3.63E-06 | -1.6136152 | -1.6059715 | 0.00764364 |
| -0.0026055 | -1.21E-06 | -1.0073244 | -0.9753737 | 0.03195068 |
| -0.0017233 | 7.35E-06 | -1.5788817 | -1.5761423 | 0.00273949 |
| -0.0043078 | 4.71E-06 | -3.2570525 | -3.2499361 | 0.00711644 |
| -0.0161014 | 5.14E-05 | -5.131901 | -5.100198 | 0.031703 |
| -0.0020665 | -2.16E-06 | -3.5347781 | -3.5335379 | 0.00124025 |
| -0.0048732 | 5.85E-05 | -2.382865 | -2.347967 | 0.03489792 |
| -0.0043394 | -0.001125 | -2.0530571 | -3.2782241 | -1.225167 |
| -0.0064087 | 3.04E-05 | -4.1594572 | -4.1533877 | 0.00606954 |
| -0.0005938 | 2.26E-06 | -6.6293839 | -6.5902165 | 0.0391674 |
| -0.0016118 | -6.12E-07 | -2.4901638 | -2.4926544 | -0.0024906 |
| -0.002326 | 5.40E-05 | -2.1878717 | -2.0772835 | 0.11058819 |
| -0.0037246 | -7.51E-06 | -4.4150956 | -4.4268639 | -0.0117683 |
| -0.0026397 | 1.12E-05 | -1.965931 | -1.9585288 | 0.00740224 |
| -0.0081014 | 0.00010057 | -4.345454 | -4.3163211 | 0.02913284 |
| -0.0019464 | 3.91E-06 | -1.6313696 | -1.620644 | 0.01072556 |
| -0.0040793 | 0.00010499 | -2.9058316 | -2.8623275 | 0.04350412 |
| -0.0059825 | 7.21E-05 | -3.2628428 | -3.1688948 | 0.09394801 |
| -0.0037134 | 7.74E-05 | -1.5197471 | -1.4639351 | 0.05581194 |
| -0.0013023 | 1.13E-06 | -6.3520499 | -6.3054413 | 0.04660869 |
| -0.0032082 | 6.69E-06 | -1.3846945 | -1.3669464 | 0.01774806 |
| -0.0055946 | -2.94E-06 | -3.0372242 | -2.9936999 | 0.04352427 |
| -0.003584 | 3.65E-05 | -3.3368195 | -3.3195722 | 0.01724732 |
| -0.0096787 | 2.39E-05 | -5.3990593 | -5.3798192 | 0.01924014 |
| -0.0017382 | 5.17E-07 | -14.026059 | -14.02132 | 0.00473952 |
| -0.0312312 | 0.00041736 | -3.1645474 | -3.0926611 | 0.0718863 |
| -0.0008511 | 1.99E-07 | -0.6783376 | -0.6824348 | -0.0040973 |
| -0.007801 | -9.70E-05 | -2.8531959 | -2.8595849 | -0.006389 |
| -0.0016566 | 1.58E-05 | -1.9560472 | -1.9411591 | 0.01488805 |
| -0.0024639 | 1.19E-05 | -1.953825 | -1.9261752 | 0.02764976 |
| -0.0003384 | 3.46E-06 | -2.142764 | -2.1249915 | 0.0177725 |
| -0.0023906 | 1.13E-05 | -0.8694702 | -0.8318598 | 0.03761041 |
| -0.0015074 | 1.43E-06 | -1.534805 | -1.5174433 | 0.0173617 |
| -0.0030626 | 1.11E-05 | -1.6528968 | -1.6338795 | 0.01901728 |
| -0.005612 | 4.75E-05 | -3.8005583 | -3.7573385 | 0.0432198 |
| -0.0026983 | 1.31E-06 | -2.2123461 | -2.2138695 | -0.0015234 |
| -0.0020344 | 9.81E-06 | -1.1977739 | -1.1697027 | 0.02807117 |
| -0.0006072 | 3.67E-06 | -4.6277581 | -4.6168378 | 0.01092032 |
| -0.0018096 | -7.29E-07 | -9.7776465 | -9.7803624 | -0.0027158 |
| -0.0047377 | -2.03E-05 | -3.0257996 | -3.0365376 | -0.010738 |
| -0.0003106 | 1.27E-06 | -5.8612205 | -5.8453589 | 0.01586159 |
| -0.0059662 | 1.69E-05 | -2.5947028 | -2.6005174 | -0.0058146 |

| | | | | |
|---|---|---|---|---|
| -0.004295 | 1.50E-05 | -3.912486 | -3.9016838 | 0.01080215 |
| -0.001486 | 6.95E-08 | -25.425731 | -25.429346 | -0.0036156 |
| -0.0014216 | -7.71E-07 | -9.4074762 | -9.4092108 | -0.0017346 |
| -0.0019769 | 2.57E-05 | -3.2441757 | -3.207677 | 0.03649867 |
| -0.0007485 | 2.04E-05 | -1.1672033 | -1.1447541 | 0.02244925 |
| -0.0022842 | 7.17E-06 | -2.1805161 | -2.1765538 | 0.00396228 |
| -0.0023374 | 3.11E-06 | -3.0268186 | -3.006098 | 0.0207206 |
| -0.0009109 | 1.82E-06 | -0.7328092 | -0.7250372 | 0.007772 |
| -0.016904 | 2.21E-05 | -5.7053301 | -5.6910274 | 0.01430273 |
| -0.0062356 | -2.49E-05 | -3.2744529 | -3.2820177 | -0.0075648 |
| -0.0006645 | 5.18E-06 | -3.1415467 | -3.1221898 | 0.01935691 |
| -0.0146754 | 2.22E-05 | -5.7171688 | -5.7086112 | 0.00855756 |
| -0.0029953 | 5.32E-06 | -2.7965685 | -2.7903352 | 0.00623333 |
| -0.0013255 | 3.84E-05 | -1.0882928 | -1.0631289 | 0.02516383 |
| -0.0005386 | 6.46E-06 | -1.8162707 | -1.8062388 | 0.01003194 |
| -0.0008573 | 4.16E-06 | -4.0866706 | -4.0510757 | 0.03559485 |
| -0.0016835 | 4.24E-05 | -1.1892807 | -1.1557599 | 0.03352082 |
| -0.0044011 | 0.00080118 | -2.5771707 | -2.201313 | 0.37585771 |
| -0.0036636 | -2.41E-06 | -3.4686086 | -3.47179 | -0.0031813 |
| -0.004488 | 2.55E-05 | -3.602026 | -3.5857718 | 0.01625419 |
| -0.0009811 | 5.66E-06 | -3.8487695 | -3.8202761 | 0.0284934 |
| -0.0056679 | 8.41E-06 | -4.5140228 | -4.5059834 | 0.00803947 |
| -0.0010263 | 4.77E-05 | -0.6854979 | -0.5971113 | 0.08838657 |
| -0.0037604 | -6.79E-06 | -3.6459848 | -3.6621234 | -0.0161387 |
| -0.0033227 | 6.70E-06 | -2.7836981 | -2.7757792 | 0.00791883 |
| -0.0013753 | 1.70E-06 | -1.0337586 | -1.0337312 | 2.74E-05 |
| -0.0012548 | 5.47E-07 | -1.2869626 | -1.2915722 | -0.0046096 |
| -0.002806 | 4.42E-05 | -1.9903347 | -1.9482592 | 0.04207546 |
| -0.0003102 | 4.09E-06 | -1.3434725 | -1.3321735 | 0.01129907 |
| -0.0064313 | 1.03E-05 | -5.366035 | -5.3550737 | 0.01096129 |
| -0.0110614 | 1.97E-05 | -5.7706823 | -5.7652533 | 0.00542903 |
| -0.0005684 | 1.20E-05 | -0.8994807 | -0.8639139 | 0.03556684 |
| -0.0100359 | -3.96E-06 | -4.6658745 | -4.6582534 | 0.00762105 |
| -0.0031018 | 7.65E-06 | -2.4009733 | -2.3908886 | 0.01008475 |
| -0.0007743 | 1.26E-05 | -0.5624813 | -0.5410397 | 0.02144158 |
| -0.0004015 | 6.96E-07 | -2.6902702 | -2.6905009 | -0.0002307 |
| -0.0045319 | 1.91E-05 | -3.475304 | -3.4745961 | 0.00070786 |
| -0.0035862 | -3.15E-07 | -3.4950849 | -3.4979621 | -0.0028772 |
| -0.0028666 | 2.85E-06 | -3.0763273 | -3.0739696 | 0.00235772 |
| -0.0041506 | 2.57E-06 | -2.8032361 | -2.6716806 | 0.13155556 |
| -0.0009775 | 4.16E-06 | -6.1610487 | -6.1085591 | 0.05248952 |
| -0.0012284 | 2.86E-05 | -0.8556025 | -0.8333705 | 0.022232 |
| -0.0048665 | 1.87E-05 | -4.2531805 | -4.2354035 | 0.01777697 |
| -0.0012733 | -1.76E-06 | -6.6402045 | -6.6382499 | 0.00195467 |
| -0.0007025 | 6.96E-06 | -4.2541022 | -4.2201305 | 0.03397173 |

| | | | | |
|---|---|---|---|---|
| -0.0037741 | 1.86E-06 | -1.5930979 | -1.5754369 | 0.01766104 |
| -0.0040304 | 1.53E-05 | -0.9076615 | -0.875371 | 0.03229043 |
| -0.0023102 | 9.25E-06 | -2.1830587 | -2.1401812 | 0.04287755 |
| -0.0072481 | -2.94E-05 | -2.8418188 | -2.8484268 | -0.006608 |
| -0.0008554 | 1.56E-06 | -1.2829938 | -1.3029347 | -0.0199409 |
| -0.0017883 | 1.27E-05 | -0.967878 | -0.9542368 | 0.01364118 |
| -0.0027937 | -1.71E-06 | -3.1934713 | -3.1888224 | 0.00464892 |
| -0.0143317 | -1.90E-05 | -5.8164129 | -5.8185887 | -0.0021758 |
| -0.0113471 | 1.16E-05 | -4.7024236 | -4.7032564 | -0.0008328 |
| -0.008443 | 3.19E-05 | -2.9469135 | -2.8849655 | 0.06194794 |
| -0.0025676 | 1.50E-07 | -3.8751553 | -3.8752548 | -9.94E-05 |
| -0.0181091 | 7.92E-05 | -5.8143361 | -5.7946672 | 0.01966882 |
| -0.0021216 | -3.52E-06 | -2.8728584 | -2.8767267 | -0.0038683 |
| -0.0046268 | -1.61E-05 | -4.8763578 | -4.8974426 | -0.0210848 |
| -0.0024353 | 8.74E-06 | -1.8055692 | -1.7926703 | 0.01289892 |
| -0.0080586 | 4.54E-05 | -4.7185683 | -4.7040205 | 0.01454782 |
| -0.0016424 | -1.29E-06 | -2.6097411 | -2.6087126 | 0.00102854 |
| -0.0005582 | -4.32E-06 | -2.2488417 | -2.2745679 | -0.0257263 |
| -0.0014807 | 2.41E-05 | -1.0979036 | -1.0741683 | 0.02373534 |
| -0.0007906 | 5.21E-06 | -0.8395574 | -0.8348114 | 0.00474605 |
| -0.006382 | -2.65E-06 | -4.7073927 | -4.7061734 | 0.00121927 |
| -0.0071756 | 3.43E-06 | -5.4879959 | -5.4874039 | 0.00059199 |
| -0.0051348 | 1.46E-05 | -3.5835761 | -3.5717151 | 0.01186097 |
| -0.0017435 | -8.65E-08 | -9.5841079 | -9.5800757 | 0.00403214 |
| -0.0021889 | 5.59E-06 | -2.2215521 | -2.2049211 | 0.01663101 |
| -0.0014536 | 3.59E-05 | -0.7571855 | -0.7198916 | 0.03729388 |
| -0.004153 | 1.71E-05 | -1.958903 | -1.9434823 | 0.01542062 |
| -0.0059159 | 5.76E-05 | -3.1241008 | -3.105423 | 0.01867783 |
| -0.0020456 | 1.10E-05 | -1.3761867 | -1.3459034 | 0.03028333 |
| -0.0026071 | -1.15E-06 | -3.7747123 | -3.7553397 | 0.01937258 |
| -0.0021145 | 2.17E-05 | -2.9664243 | -2.9418839 | 0.02454042 |
| -0.0022633 | 3.48E-06 | -3.5932918 | -3.5897164 | 0.00357533 |
| -0.0039008 | -2.27E-05 | -2.9092921 | -2.9348532 | -0.0255611 |
| -0.0041985 | -1.08E-06 | -3.2395291 | -3.2389674 | 0.00056171 |
| -0.0154281 | 6.50E-05 | -5.9271495 | -5.898735 | 0.02841449 |
| -0.0039587 | 6.11E-05 | -4.0706265 | -3.9902526 | 0.08037388 |
| -0.003559 | 4.74E-06 | -4.0985315 | -4.0959101 | 0.00262141 |
| -0.0048114 | 1.99E-05 | -2.9161085 | -2.8882259 | 0.02788258 |
| -0.0043735 | 2.43E-06 | -4.5226986 | -4.5259774 | -0.0032787 |
| -0.0019057 | -1.12E-05 | -2.267019 | -2.2669404 | 7.87E-05 |
| -0.0059617 | 5.97E-06 | -5.0133686 | -4.9961643 | 0.01720428 |
| -0.0035351 | 1.39E-06 | -3.7914685 | -3.7929997 | -0.0015312 |
| -0.0031894 | 8.70E-06 | -3.7984121 | -3.7928652 | 0.00554693 |
| -0.0048019 | 1.12E-05 | -3.6552789 | -3.6429633 | 0.01231563 |
| -0.0012721 | -1.39E-06 | -6.8993119 | -6.9198266 | -0.0205147 |

| | | | | |
|---|---|---|---|---|
| -0.003092 | 6.23E-06 | -3.0378985 | -3.0293757 | 0.00852287 |
| -0.0037632 | 1.99E-05 | -4.1817975 | -4.150064 | 0.03173351 |
| -0.0053876 | 8.85E-06 | -4.8311579 | -4.821692 | 0.00946593 |
| -0.0007993 | 1.78E-06 | -5.468096 | -5.4543515 | 0.01374447 |
| -0.0025398 | 1.45E-05 | -2.4813159 | -2.4723686 | 0.00894725 |
| -0.0056162 | 4.33E-05 | -3.8005583 | -3.7633398 | 0.03721857 |
| -0.0006089 | 8.66E-07 | -2.8973174 | -2.8941188 | 0.00319862 |
| -0.0009738 | 1.29E-05 | -3.9833233 | -3.8569163 | 0.12640697 |
| -0.0010189 | 2.06E-06 | -3.2420771 | -3.2304442 | 0.01163286 |
| -0.0022584 | 1.65E-05 | -2.0416963 | -2.0228676 | 0.01882863 |
| -0.0071263 | 9.33E-05 | -2.2434704 | -2.2466384 | -0.003168 |
| -0.0009345 | 3.25E-08 | -3.3111343 | -3.313069 | -0.0019348 |
| -0.0024879 | 2.87E-06 | -1.6658168 | -1.6578374 | 0.00797945 |
| -0.007863 | 1.48E-05 | -4.2863357 | -4.28301 | 0.0033257 |
| -0.002302 | -1.12E-05 | -1.2070313 | -1.2188146 | -0.0117834 |
| -0.0041689 | 0.00056621 | -3.7984385 | -3.1822172 | 0.61622131 |
| -0.004358 | 6.16E-06 | -2.4044405 | -2.3666633 | 0.03777719 |
| -0.0013668 | 1.39E-06 | -1.5141555 | -1.5043216 | 0.00983393 |
| -0.0048738 | 3.49E-05 | -3.031247 | -2.9986825 | 0.03256452 |
| -0.0024467 | 2.25E-05 | -2.0498269 | -2.0059381 | 0.04388881 |
| -0.0129071 | 0.00010138 | -4.566308 | -4.5308132 | 0.0354948 |
| -0.0038625 | 2.51E-05 | -4.6171238 | -4.5885952 | 0.02852869 |
| -0.0014811 | -5.89E-06 | -2.3043573 | -2.3184948 | -0.0141375 |
| -0.0019612 | 8.07E-06 | -3.102077 | -3.101596 | 0.00048101 |
| -0.002609 | 9.31E-07 | -4.7533844 | -4.7388844 | 0.0144999 |
| -0.0021964 | 8.67E-06 | -10.494191 | -10.491521 | 0.00267005 |

| Progen2_scoi | Progen2_fina | RITA_score_v | RITA_score_r | RITA_final_score |
|---|---|---|---|---|
| -0.0008882 | 2.05E-07 | -6.4390417 | -6.438613 | 0.00042868 |
| -0.0008884 | 4.59E-08 | -6.4390417 | -6.4395931 | -0.0005515 |
| -0.000337 | 8.59E-07 | -2.3434079 | -2.3361149 | 0.00729305 |
| -0.0018649 | 2.92E-05 | -0.9718568 | -0.9504507 | 0.02140611 |
| -0.0013371 | 1.85E-06 | -9.7287054 | -9.6921614 | 0.03654397 |
| -0.0013397 | -7.93E-07 | -9.7287054 | -9.733328 | -0.0046226 |
| -0.0013389 | 5.41E-08 | -9.7287054 | -9.7285259 | 0.00017953 |
| -0.0013381 | 8.62E-07 | -9.7287054 | -9.7304767 | -0.0017713 |
| -0.001333 | 5.93E-06 | -9.7287054 | -9.7124337 | 0.01627171 |
| -0.0017244 | -7.65E-07 | -1.8331247 | -1.833796 | -0.0006713 |
| -0.0017208 | 2.87E-06 | -1.8331247 | -1.8063238 | 0.02680087 |
| -0.0010404 | 7.18E-06 | -1.0097367 | -0.983563 | 0.02617362 |
| -0.001048 | -4.54E-07 | -1.0097367 | -1.0130838 | -0.0033471 |
| -0.0011926 | 8.72E-07 | -1.65412 | -1.6271994 | 0.02692068 |
| -0.0018401 | -2.14E-07 | -3.3894198 | -3.3902044 | -0.0007846 |
| -0.0018396 | 2.60E-07 | -3.3894198 | -3.3901711 | -0.0007513 |
| -0.0018375 | 2.43E-06 | -3.3894198 | -3.3822407 | 0.00717914 |

| | | | | |
|---|---|---|---|---|
| -0.0014402 | 8.98E-07 | -10.09513 | -10.085732 | 0.00939769 |
| -0.0014409 | 1.95E-07 | -10.09513 | -10.097235 | -0.0021054 |
| -0.0021781 | 5.25E-07 | -16.271221 | -16.264086 | 0.00713539 |
| -0.0021794 | -8.26E-07 | -16.271221 | -16.267887 | 0.00333405 |
| -0.0014101 | 1.53E-05 | -1.3045034 | -1.2847674 | 0.01973599 |
| -0.002395 | 7.92E-06 | -3.2212473 | -3.2066203 | 0.01462698 |
| -0.0029276 | 1.18E-05 | -2.8073002 | -2.7902782 | 0.01702201 |
| -0.0019412 | 3.09E-07 | -3.2303723 | -3.2270824 | 0.00328994 |
| -0.0019417 | -2.47E-07 | -3.2303723 | -3.231499 | -0.0011266 |
| -0.0019392 | 2.24E-06 | -3.2303723 | -3.2296696 | 0.00070274 |
| -0.0019417 | -2.46E-07 | -3.2303723 | -3.2305832 | -0.0002109 |
| -0.0019421 | -6.72E-07 | -3.2303723 | -3.2291559 | 0.00121641 |
| -0.0019386 | 2.88E-06 | -3.2303723 | -3.2254275 | 0.0049448 |
| -0.0009995 | 4.86E-07 | -1.3263996 | -1.324002 | 0.00239754 |
| -0.0012295 | -1.75E-08 | -9.4428061 | -9.4442018 | -0.0013957 |
| -0.00123 | -5.48E-07 | -9.4428061 | -9.4466734 | -0.0038673 |
| -0.0027329 | 3.28E-06 | -4.2898488 | -4.2859974 | 0.00385141 |
| -0.0011763 | -3.57E-07 | -3.6040708 | -3.6126527 | -0.0085818 |
| -0.000586 | 6.57E-07 | -2.9197316 | -2.9008079 | 0.01892376 |
| -0.0004165 | 8.24E-07 | -2.6504609 | -2.6455849 | 0.00487608 |
| -0.0009288 | 1.32E-07 | -23.284598 | -23.282992 | 0.00160563 |
| -0.0009288 | 1.25E-07 | -23.284598 | -23.281168 | 0.00342906 |
| -0.0013381 | 1.14E-05 | -1.2852365 | -1.2675939 | 0.01764262 |
| -0.0008835 | 2.01E-07 | -6.6892018 | -6.6864296 | 0.00277227 |
| -0.0008831 | 5.63E-07 | -6.6892018 | -6.6798293 | 0.00937253 |
| -0.0013908 | 2.91E-05 | -1.3852651 | -1.3514793 | 0.03378576 |
| -0.0010112 | -1.61E-06 | -6.5544376 | -6.5607284 | -0.0062908 |
| -0.0010113 | -1.77E-06 | -6.5544376 | -6.559616 | -0.0051783 |
| -0.0010105 | -9.39E-07 | -6.5544376 | -6.5578505 | -0.0034128 |
| -0.0010107 | -1.15E-06 | -6.5544376 | -6.5586932 | -0.0042555 |
| -0.0010097 | -1.81E-07 | -6.5544376 | -6.5522355 | 0.00220215 |
| -0.0010083 | 1.27E-06 | -6.5544376 | -6.5492871 | 0.00515056 |
| -0.0010126 | -3.06E-06 | -6.5544376 | -6.5645833 | -0.0101457 |
| -0.0010098 | -2.56E-07 | -6.5544376 | -6.5529969 | 0.00144076 |
| -0.0010104 | -8.86E-07 | -6.5544376 | -6.5624254 | -0.0079877 |
| -0.0010105 | -9.69E-07 | -6.5544376 | -6.5586542 | -0.0042166 |
| -0.0010096 | -7.90E-08 | -6.5544376 | -6.5504171 | 0.00402057 |
| -0.0010073 | 2.22E-06 | -6.5544376 | -6.5446014 | 0.0098362 |
| -0.0010104 | -8.31E-07 | -6.5544376 | -6.5534744 | 0.00096321 |
| -0.0007389 | 1.63E-06 | -16.240251 | -16.221931 | 0.01831996 |
| -0.0013608 | 3.83E-07 | -9.8017685 | -9.7963257 | 0.00544286 |
| -0.001358 | 3.18E-06 | -9.8017685 | -9.782777 | 0.01899159 |
| -0.0020177 | 2.01E-05 | -1.5629956 | -1.5410077 | 0.02198786 |
| -0.0008304 | 1.90E-05 | -0.8019912 | -0.7571612 | 0.04482993 |
| -0.0003956 | 3.31E-06 | -5.2172873 | -5.1853416 | 0.03194577 |

| | | | | |
|---|---|---|---|---|
| -0.0005152 | 8.00E-07 | -3.963347 | -3.9536018 | 0.00974524 |
| -0.0040819 | 2.53E-06 | -3.3413576 | -3.3216872 | 0.01967037 |
| -0.0010658 | 4.83E-06 | -7.474027 | -7.4615392 | 0.01248783 |
| -0.0012261 | 3.57E-08 | -9.3787434 | -9.3799174 | -0.001174 |
| -0.001142 | 5.19E-06 | -1.6517534 | -1.6461651 | 0.00558829 |
| -0.0007179 | 3.31E-05 | -0.7203402 | -0.6776901 | 0.04265013 |
| -0.001439 | 1.52E-07 | -9.6503721 | -9.6497658 | 0.0006063 |
| -0.0003859 | 1.06E-05 | -2.9767831 | -2.9389171 | 0.037866 |
| -0.0009918 | -8.73E-07 | -2.8514222 | -2.8560442 | -0.004622 |
| -0.0009397 | 1.09E-05 | -6.3694062 | -6.3472135 | 0.02219266 |
| -0.0011494 | -3.17E-06 | -0.8055149 | -0.8083717 | -0.0028568 |
| -0.0040317 | 1.94E-06 | -2.2792813 | -2.2467211 | 0.03256011 |
| -0.0007252 | 1.36E-06 | -0.769554 | -0.7652974 | 0.00425655 |
| -0.0011063 | 1.21E-05 | -1.1268061 | -1.1061676 | 0.02063853 |
| -0.0006425 | 6.32E-06 | -3.7080747 | -3.6967713 | 0.01130348 |
| -0.0006457 | 3.09E-06 | -3.7080747 | -3.7004471 | 0.0076277 |
| -0.0009682 | -3.16E-07 | -6.6496345 | -6.6524702 | -0.0028358 |
| -0.0008228 | 3.95E-07 | -2.5503785 | -2.5473519 | 0.00302655 |
| -0.0009247 | 5.40E-07 | -3.6778395 | -3.6546465 | 0.023193 |
| -0.0024683 | 7.56E-07 | -4.9960942 | -4.994617 | 0.00147724 |
| -0.001314 | -4.37E-07 | -1.8609445 | -1.8630245 | -0.00208 |
| -0.0016345 | 1.53E-05 | -2.2494159 | -2.2319114 | 0.01750445 |
| -0.0003752 | 1.46E-06 | -2.5279352 | -2.5257889 | 0.0021463 |
| -0.0026647 | 1.64E-06 | -1.6452838 | -1.6405819 | 0.00470185 |
| -0.0082157 | -7.99E-05 | -2.8870831 | -2.8907764 | -0.0036933 |
| -0.0029989 | 1.17E-07 | -4.2506518 | -4.2139065 | 0.03674531 |
| -0.0054316 | 1.61E-05 | -3.8585483 | -3.8477401 | 0.01080823 |
| -0.0018336 | 6.82E-06 | -2.8712335 | -2.8588655 | 0.01236796 |
| -0.0011488 | -1.66E-07 | -9.3495153 | -9.3480005 | 0.00151479 |
| -0.0011455 | 3.09E-06 | -9.3495153 | -9.3326197 | 0.01689559 |
| -0.0005895 | 8.02E-07 | -5.0816348 | -5.0665026 | 0.01513225 |
| -0.0005893 | 1.04E-06 | -5.0816348 | -5.0682243 | 0.01341057 |
| -0.0005893 | 9.71E-07 | -5.0816348 | -5.0607714 | 0.02086341 |
| -0.0019258 | 2.71E-05 | -1.8361189 | -1.8008912 | 0.03522766 |
| -0.0036127 | 3.46E-06 | -3.6871473 | -3.6805098 | 0.00663745 |
| -0.0039595 | 7.05E-08 | -5.6582158 | -5.6584427 | -0.000227 |
| -0.0038566 | 8.22E-06 | -2.0634927 | -2.0601102 | 0.00338244 |
| -0.0026061 | 5.70E-06 | -0.8595581 | -0.8508215 | 0.00873661 |
| -0.0126512 | 0.00100983 | -3.2667494 | -3.0314156 | 0.2353338 |
| -0.0046465 | -8.88E-07 | -3.1956809 | -3.1860987 | 0.00958216 |
| -0.0053878 | 1.67E-05 | -2.8072529 | -2.8014092 | 0.00584364 |
| -0.0014456 | 2.20E-06 | -5.9076757 | -5.8877977 | 0.01987803 |
| -0.0014471 | 6.99E-07 | -5.9076757 | -5.9035584 | 0.00411737 |
| -0.0014475 | 2.69E-07 | -5.9076757 | -5.9055674 | 0.00210834 |
| -0.0014485 | -7.04E-07 | -5.9076757 | -5.9082973 | -0.0006216 |

| | | | | |
|---|---|---|---|---|
| -0.001447 | 8.06E-07 | -5.9076757 | -5.9021168 | 0.00555891 |
| -0.0093082 | 2.66E-05 | -4.8694561 | -4.8656783 | 0.00377774 |
| -0.0004129 | 4.82E-07 | -9.9702165 | -9.964286 | 0.00593054 |
| -0.0004128 | 6.05E-07 | -9.9702165 | -9.9584643 | 0.01175219 |
| -0.0004129 | 4.73E-07 | -9.9702165 | -9.960373 | 0.00984347 |
| -0.0004132 | 2.33E-07 | -9.9702165 | -9.9637269 | 0.00648957 |
| -0.0004124 | 1.02E-06 | -9.9702165 | -9.9601842 | 0.01003236 |
| -0.0004126 | 8.18E-07 | -9.9702165 | -9.9549192 | 0.01529729 |
| -0.000413 | 3.94E-07 | -9.9702165 | -9.960013 | 0.01020354 |
| -0.0004133 | 1.09E-07 | -9.9702165 | -9.9665331 | 0.00368345 |
| -0.0014495 | -7.65E-07 | -2.1135607 | -2.1110379 | 0.00252283 |
| -0.0091518 | -1.71E-05 | -3.2048043 | -3.2080948 | -0.0032905 |
| -0.0071091 | 3.27E-06 | -4.7806289 | -4.793612 | -0.0129831 |
| -0.0070989 | 1.34E-05 | -4.7806289 | -4.7711756 | 0.0094533 |
| -0.0071231 | -1.08E-05 | -4.7806289 | -4.7951233 | -0.0144944 |
| -0.0070984 | 1.40E-05 | -4.7806289 | -4.7748668 | 0.0057621 |
| -0.0037969 | 6.58E-06 | -3.0865149 | -3.0761365 | 0.01037848 |
| -0.006159 | 1.36E-05 | -5.1807091 | -5.1686177 | 0.0120914 |
| -0.0086002 | 0.00011816 | -3.0490208 | -2.9582869 | 0.09073389 |
| -0.0015527 | 5.82E-07 | -12.807871 | -12.794132 | 0.01373851 |
| -0.0015509 | 2.43E-06 | -12.807871 | -12.763673 | 0.04419768 |
| -0.0015532 | 9.00E-08 | -12.807871 | -12.805972 | 0.00189829 |
| -0.0016917 | 6.05E-06 | -2.2513477 | -2.2470793 | 0.00426841 |
| -0.0010225 | -2.99E-07 | -7.1350336 | -7.1364101 | -0.0013765 |
| -0.0071942 | 1.15E-06 | -4.1961834 | -4.2065864 | -0.0104029 |
| -0.0071977 | -2.44E-06 | -4.1961834 | -4.175936 | 0.02024746 |
| -0.0071845 | 1.08E-05 | -4.1961834 | -4.2321107 | -0.0359273 |
| -0.0030169 | 1.35E-05 | -2.2903301 | -2.274725 | 0.01560509 |
| -0.001807 | 1.87E-06 | -7.5051646 | -7.493505 | 0.01165962 |
| -0.0006704 | 1.74E-06 | -2.5945129 | -2.5757558 | 0.0187571 |
| -0.0112627 | 1.66E-05 | -3.5969728 | -3.6363949 | -0.039422 |
| -0.0029761 | 1.02E-05 | -2.8992147 | -2.8869196 | 0.01229513 |
| -0.0029861 | 2.71E-07 | -2.8992147 | -2.897992 | 0.00122273 |
| -0.0011969 | 5.94E-06 | -1.4873038 | -1.4758958 | 0.01140797 |
| -0.0011994 | 3.40E-06 | -1.4873038 | -1.4841077 | 0.00319606 |
| -0.0011954 | 7.48E-06 | -1.4873038 | -1.4793108 | 0.00799304 |
| -0.0005844 | -9.37E-08 | -3.0392226 | -3.0388248 | 0.00039774 |
| -0.0005803 | 4.01E-06 | -3.0392226 | -3.0225313 | 0.0166913 |
| -0.0005663 | 1.80E-05 | -3.0392226 | -2.9946985 | 0.04452407 |
| -0.0063036 | 2.73E-05 | -3.5429775 | -3.5307984 | 0.01217902 |
| -0.0062622 | 6.87E-05 | -3.5429775 | -3.5138327 | 0.02914476 |
| -0.0025932 | -8.98E-06 | -3.1500336 | -3.1447115 | 0.0053221 |
| -0.0033643 | 1.12E-05 | -2.6264769 | -2.6161723 | 0.01030457 |
| -0.0033786 | -3.05E-06 | -2.6264769 | -2.6318644 | -0.0053875 |
| -0.0033754 | 1.40E-07 | -2.6264769 | -2.6241637 | 0.00231314 |

| | | | | |
|---|---|---|---|---|
| -0.0033834 | -7.83E-06 | -2.6264769 | -2.6313182 | -0.0048413 |
| -0.0052088 | 2.40E-05 | -3.4495574 | -3.3992748 | 0.0502826 |
| -0.0052944 | -6.17E-05 | -3.4495574 | -3.4903989 | -0.0408415 |
| -0.0006105 | 1.36E-05 | -0.9723271 | -0.9451578 | 0.02716926 |
| -0.001745 | -4.03E-07 | -2.2105173 | -2.2070788 | 0.00343847 |
| -0.0012538 | 3.63E-06 | -1.6136152 | -1.6059715 | 0.00764364 |
| -0.0026055 | -1.21E-06 | -1.0073244 | -0.9753737 | 0.03195068 |
| -0.0017301 | 5.06E-07 | -1.5788817 | -1.5735858 | 0.00529593 |
| -0.0017241 | 6.53E-06 | -1.5788817 | -1.5814283 | -0.0025466 |
| -0.0043078 | 4.71E-06 | -3.2570525 | -3.2499361 | 0.00711644 |
| -0.0161014 | 5.14E-05 | -5.131901 | -5.100198 | 0.031703 |
| -0.0020665 | -2.16E-06 | -3.5347781 | -3.5335379 | 0.00124025 |
| -0.0048732 | 5.85E-05 | -2.382865 | -2.347967 | 0.03489792 |
| -0.0031935 | 2.09E-05 | -2.0530571 | -2.0379421 | 0.01511502 |
| -0.0032218 | -7.50E-06 | -2.0530571 | -2.0677935 | -0.0147364 |
| -0.003203 | 1.13E-05 | -2.0530571 | -2.0749607 | -0.0219036 |
| -0.0032142 | 1.42E-07 | -2.0530571 | -2.0565292 | -0.0034721 |
| -0.0032366 | -2.23E-05 | -2.0530571 | -2.0978428 | -0.0447857 |
| -0.0032298 | -1.55E-05 | -2.0530571 | -2.0865904 | -0.0335333 |
| -0.0032284 | -1.40E-05 | -2.0530571 | -2.0939415 | -0.0408844 |
| -0.0032181 | -3.77E-06 | -2.0530571 | -2.0703949 | -0.0173378 |
| -0.0032078 | 6.50E-06 | -2.0530571 | -2.058018 | -0.0049609 |
| -0.0032221 | -7.78E-06 | -2.0530571 | -2.0773605 | -0.0243034 |
| -0.0032313 | -1.70E-05 | -2.0530571 | -2.0630829 | -0.0100259 |
| -0.0032368 | -2.25E-05 | -2.0530571 | -2.1066911 | -0.053634 |
| -0.0032475 | -3.32E-05 | -2.0530571 | -2.0760317 | -0.0229746 |
| -0.0032356 | -2.13E-05 | -2.0530571 | -2.0645864 | -0.0115293 |
| -0.0032039 | 1.04E-05 | -2.0530571 | -2.1102479 | -0.0571908 |
| -0.0032357 | -2.14E-05 | -2.0530571 | -2.0867651 | -0.033708 |
| -0.0032129 | 1.43E-06 | -2.0530571 | -2.0765839 | -0.0235268 |
| -0.0032267 | -1.24E-05 | -2.0530571 | -2.0870017 | -0.0339446 |
| -0.0032391 | -2.48E-05 | -2.0530571 | -2.0845745 | -0.0315174 |
| -0.003239 | -2.47E-05 | -2.0530571 | -2.0688406 | -0.0157835 |
| -0.003185 | 2.93E-05 | -2.0530571 | -2.0557629 | -0.0027058 |
| -0.0032682 | -5.39E-05 | -2.0530571 | -2.0911038 | -0.0380467 |
| -0.003236 | -2.17E-05 | -2.0530571 | -2.07726 | -0.0242029 |
| -0.0032395 | -2.52E-05 | -2.0530571 | -2.0725669 | -0.0195098 |
| -0.0032289 | -1.46E-05 | -2.0530571 | -2.0856214 | -0.0325643 |
| -0.0032642 | -4.99E-05 | -2.0530571 | -2.0886486 | -0.0355915 |
| -0.0032927 | -7.84E-05 | -2.0530571 | -2.1113648 | -0.0583078 |
| -0.0032742 | -5.98E-05 | -2.0530571 | -2.0955461 | -0.0424891 |
| -0.0032545 | -4.02E-05 | -2.0530571 | -2.0843627 | -0.0313057 |
| -0.0032507 | -3.64E-05 | -2.0530571 | -2.0874521 | -0.034395 |
| -0.0032408 | -2.65E-05 | -2.0530571 | -2.0866342 | -0.0335771 |
| -0.0032641 | -4.98E-05 | -2.0530571 | -2.1245718 | -0.0715147 |

| | | | | |
|---|---|---|---|---|
| -0.0032614 | -4.71E-05 | -2.0530571 | -2.1070323 | -0.0539752 |
| -0.003284 | -6.97E-05 | -2.0530571 | -2.0940809 | -0.0410239 |
| -0.0032894 | -7.50E-05 | -2.0530571 | -2.0937184 | -0.0406613 |
| -0.0032905 | -7.61E-05 | -2.0530571 | -2.0994517 | -0.0463946 |
| -0.003177 | 3.73E-05 | -2.0530571 | -2.0382261 | 0.01483095 |
| -0.0033452 | -0.0001309 | -2.0530571 | -2.1493775 | -0.0963204 |
| -0.0032591 | -4.48E-05 | -2.0530571 | -2.092557 | -0.0394999 |
| -0.0032741 | -5.97E-05 | -2.0530571 | -2.0796297 | -0.0265726 |
| -0.0032415 | -2.71E-05 | -2.0530571 | -2.0855939 | -0.0325369 |
| -0.0032588 | -4.45E-05 | -2.0530571 | -2.0870122 | -0.0339551 |
| -0.0031966 | 1.77E-05 | -2.0530571 | -2.0525216 | 0.00053549 |
| -0.0032779 | -6.36E-05 | -2.0530571 | -2.0901053 | -0.0370482 |
| -0.0032432 | -2.89E-05 | -2.0530571 | -2.0680457 | -0.0149887 |
| -0.0032593 | -4.50E-05 | -2.0530571 | -2.0956162 | -0.0425591 |
| -0.0032641 | -4.98E-05 | -2.0530571 | -2.089523 | -0.0364659 |
| -0.0032373 | -2.30E-05 | -2.0530571 | -2.0701993 | -0.0171422 |
| -0.003253 | -3.87E-05 | -2.0530571 | -2.0864357 | -0.0333786 |
| -0.0032534 | -3.90E-05 | -2.0530571 | -2.0834647 | -0.0304077 |
| -0.0032827 | -6.84E-05 | -2.0530571 | -2.0977551 | -0.044698 |
| -0.003223 | -8.68E-06 | -2.0530571 | -2.0654473 | -0.0123903 |
| -0.0032734 | -5.91E-05 | -2.0530571 | -2.0929039 | -0.0398468 |
| -0.0033843 | -0.0001699 | -2.0530571 | -2.1560459 | -0.1029888 |
| -0.0032602 | -4.58E-05 | -2.0530571 | -2.0919464 | -0.0388893 |
| -0.0032526 | -3.83E-05 | -2.0530571 | -2.0866344 | -0.0335773 |
| -0.0033149 | -0.0001005 | -2.0530571 | -2.1272327 | -0.0741756 |
| -0.0032117 | 2.64E-06 | -2.0530571 | -2.0493478 | 0.00370932 |
| -0.0033018 | -8.75E-05 | -2.0530571 | -2.1072168 | -0.0541598 |
| -0.0033197 | -0.0001054 | -2.0530571 | -2.1095188 | -0.0564617 |
| -0.0032449 | -3.06E-05 | -2.0530571 | -2.0809902 | -0.0279331 |
| -0.0032576 | -4.33E-05 | -2.0530571 | -2.0800161 | -0.0269591 |
| -0.0032528 | -3.85E-05 | -2.0530571 | -2.0918677 | -0.0388106 |
| -0.0032409 | -2.66E-05 | -2.0530571 | -2.082463 | -0.029406 |
| -0.0032545 | -4.02E-05 | -2.0530571 | -2.0956932 | -0.0426362 |
| -0.003248 | -3.37E-05 | -2.0530571 | -2.0865326 | -0.0334755 |
| -0.0032574 | -4.31E-05 | -2.0530571 | -2.0826114 | -0.0295544 |
| -0.0033182 | -0.0001039 | -2.0530571 | -2.1243607 | -0.0713036 |
| -0.0032229 | -8.58E-06 | -2.0530571 | -2.0692006 | -0.0161436 |
| -0.0032935 | -7.92E-05 | -2.0530571 | -2.1057602 | -0.0527031 |
| -0.0032913 | -7.70E-05 | -2.0530571 | -2.0882252 | -0.0351682 |
| -0.003264 | -4.97E-05 | -2.0530571 | -2.0905156 | -0.0374585 |
| -0.003259 | -4.46E-05 | -2.0530571 | -2.0946518 | -0.0415947 |
| -0.0032554 | -4.11E-05 | -2.0530571 | -2.0902125 | -0.0371554 |
| -0.0032593 | -4.49E-05 | -2.0530571 | -2.0949677 | -0.0419106 |
| -0.0032568 | -4.25E-05 | -2.0530571 | -2.0997275 | -0.0466704 |
| -0.0032474 | -3.31E-05 | -2.0530571 | -2.0771697 | -0.0241126 |

| | | | | |
|---|---|---|---|---|
| -0.0032669 | -5.25E-05 | -2.0530571 | -2.0884877 | -0.0354307 |
| -0.0032557 | -4.14E-05 | -2.0530571 | -2.0982437 | -0.0451866 |
| -0.0032491 | -3.47E-05 | -2.0530571 | -2.0975622 | -0.0445051 |
| -0.0032689 | -5.46E-05 | -2.0530571 | -2.0995432 | -0.0464861 |
| -0.0032172 | -2.89E-06 | -2.0530571 | -2.0633433 | -0.0102862 |
| -0.003298 | -8.37E-05 | -2.0530571 | -2.1140362 | -0.0609791 |
| -0.003216 | -1.66E-06 | -2.0530571 | -2.0615027 | -0.0084456 |
| -0.0032337 | -1.94E-05 | -2.0530571 | -2.0863268 | -0.0332698 |
| -0.0032621 | -4.78E-05 | -2.0530571 | -2.0875396 | -0.0344825 |
| -0.0032543 | -4.00E-05 | -2.0530571 | -2.0822399 | -0.0291828 |
| -0.0033513 | -0.000137 | -2.0530571 | -2.1868739 | -0.1338168 |
| -0.0032127 | 1.58E-06 | -2.0530571 | -2.0581257 | -0.0050687 |
| -0.0032598 | -4.55E-05 | -2.0530571 | -2.0829809 | -0.0299238 |
| -0.0032728 | -5.85E-05 | -2.0530571 | -2.0876161 | -0.034559 |
| -0.0032487 | -3.44E-05 | -2.0530571 | -2.0884566 | -0.0353996 |
| -0.0033274 | -0.0001131 | -2.0530571 | -2.154045 | -0.1009879 |
| -0.0032593 | -4.49E-05 | -2.0530571 | -2.1042647 | -0.0512077 |
| -0.0032183 | -3.96E-06 | -2.0530571 | -2.0695442 | -0.0164871 |
| -0.0032359 | -2.16E-05 | -2.0530571 | -2.0593001 | -0.006243 |
| -0.0032116 | 2.71E-06 | -2.0530571 | -2.0480403 | 0.0050168 |
| -0.003229 | -1.47E-05 | -2.0530571 | -2.0715942 | -0.0185372 |
| -0.0032146 | -3.15E-07 | -2.0530571 | -2.0581886 | -0.0051315 |
| -0.0032211 | -6.74E-06 | -2.0530571 | -2.056672 | -0.0036149 |
| -0.0032139 | 4.30E-07 | -2.0530571 | -2.0659726 | -0.0129155 |
| -0.0032101 | 4.26E-06 | -2.0530571 | -2.0634226 | -0.0103655 |
| -0.0032174 | -3.05E-06 | -2.0530571 | -2.0797976 | -0.0267406 |
| -0.0032197 | -5.36E-06 | -2.0530571 | -2.0825409 | -0.0294838 |
| -0.0064087 | 3.04E-05 | -4.1594572 | -4.1533877 | 0.00606954 |
| -0.0005961 | -4.32E-08 | -6.6293839 | -6.6208618 | 0.00852209 |
| -0.0005954 | 6.10E-07 | -6.6293839 | -6.6210706 | 0.00831336 |
| -0.0005959 | 1.12E-07 | -6.6293839 | -6.6234031 | 0.00598085 |
| -0.0005942 | 1.83E-06 | -6.6293839 | -6.609367 | 0.02001691 |
| -0.0005963 | -2.64E-07 | -6.6293839 | -6.6338645 | -0.0044806 |
| -0.0016118 | -6.12E-07 | -2.4901638 | -2.4926544 | -0.0024906 |
| -0.002326 | 5.40E-05 | -2.1878717 | -2.0772835 | 0.11058819 |
| -0.0037246 | -7.51E-06 | -4.4150956 | -4.4268639 | -0.0117683 |
| -0.0026475 | 3.42E-06 | -1.965931 | -1.957583 | 0.00834805 |
| -0.0026432 | 7.71E-06 | -1.965931 | -1.9668045 | -0.0008735 |
| -0.0080738 | 0.00012822 | -4.345454 | -4.318593 | 0.02686095 |
| -0.0081709 | 3.11E-05 | -4.345454 | -4.3233767 | 0.02207732 |
| -0.0019464 | 3.91E-06 | -1.6313696 | -1.620644 | 0.01072556 |
| -0.0041631 | 2.12E-05 | -2.9058316 | -2.9025425 | 0.0032891 |
| -0.0041018 | 8.25E-05 | -2.9058316 | -2.8649875 | 0.04084408 |
| -0.0059838 | 7.08E-05 | -3.2628428 | -3.22755 | 0.03529274 |
| -0.0060408 | 1.38E-05 | -3.2628428 | -3.2673464 | -0.0045036 |

| | | | | |
|---|---|---|---|---|
| -0.0060699 | -1.53E-05 | -3.2628428 | -3.1996136 | 0.0632292 |
| -0.0037134 | 7.74E-05 | -1.5197471 | -1.4639351 | 0.05581194 |
| -0.0013023 | 1.13E-06 | -6.3520499 | -6.3054413 | 0.04660869 |
| -0.0032082 | 6.69E-06 | -1.3846945 | -1.3669464 | 0.01774806 |
| -0.0055946 | -2.94E-06 | -3.0372242 | -2.9936999 | 0.04352427 |
| -0.0036249 | -4.43E-06 | -3.3368195 | -3.3492835 | -0.0124639 |
| -0.0036077 | 1.28E-05 | -3.3368195 | -3.3345814 | 0.00223815 |
| -0.0036088 | 1.17E-05 | -3.3368195 | -3.3255658 | 0.01125371 |
| -0.0036057 | 1.48E-05 | -3.3368195 | -3.3206924 | 0.01612711 |
| -0.0096787 | 2.39E-05 | -5.3990593 | -5.3798192 | 0.01924014 |
| -0.0017385 | 1.86E-07 | -14.026059 | -14.023352 | 0.002707 |
| -0.0017383 | 3.38E-07 | -14.026059 | -14.024024 | 0.00203514 |
| -0.0312312 | 0.00041736 | -3.1645474 | -3.0926611 | 0.0718863 |
| -0.0008511 | 1.99E-07 | -0.6783376 | -0.6824348 | -0.0040973 |
| -0.007801 | -9.70E-05 | -2.8531959 | -2.8595849 | -0.006389 |
| -0.0016741 | -1.68E-06 | -1.9560472 | -1.9574842 | -0.0014371 |
| -0.0016551 | 1.73E-05 | -1.9560472 | -1.9429731 | 0.01307404 |
| -0.0024639 | 1.19E-05 | -1.953825 | -1.9261752 | 0.02764976 |
| -0.0003384 | 3.46E-06 | -2.142764 | -2.1249915 | 0.0177725 |
| -0.0023964 | 5.54E-06 | -0.8694702 | -0.8549423 | 0.01452792 |
| -0.0023962 | 5.72E-06 | -0.8694702 | -0.8462447 | 0.02322552 |
| -0.0015074 | 1.43E-06 | -1.534805 | -1.5174433 | 0.0173617 |
| -0.0030626 | 1.11E-05 | -1.6528968 | -1.6338795 | 0.01901728 |
| -0.0056332 | 2.62E-05 | -3.8005583 | -3.787413 | 0.01314533 |
| -0.0056334 | 2.60E-05 | -3.8005583 | -3.7785919 | 0.02196646 |
| -0.0056232 | 3.62E-05 | -3.8005583 | -3.7721257 | 0.02843261 |
| -0.0056651 | -5.60E-06 | -3.8005583 | -3.8096066 | -0.0090482 |
| -0.0056759 | -1.64E-05 | -3.8005583 | -3.8159539 | -0.0153955 |
| -0.0026983 | 1.31E-06 | -2.2123461 | -2.2138695 | -0.0015234 |
| -0.0020344 | 9.81E-06 | -1.1977739 | -1.1697027 | 0.02807117 |
| -0.0006072 | 3.67E-06 | -4.6277581 | -4.6168378 | 0.01092032 |
| -0.0018096 | -7.29E-07 | -9.7776465 | -9.7803624 | -0.0027158 |
| -0.0047377 | -2.03E-05 | -3.0257996 | -3.0365376 | -0.010738 |
| -0.0003117 | 1.22E-07 | -5.8612205 | -5.8605905 | 0.00063002 |
| -0.0003107 | 1.15E-06 | -5.8612205 | -5.8459889 | 0.01523156 |
| -0.0059662 | 1.69E-05 | -2.5947028 | -2.6005174 | -0.0058146 |
| -0.004295 | 1.50E-05 | -3.912486 | -3.9016838 | 0.01080215 |
| -0.0014859 | 1.66E-07 | -25.425731 | -25.426913 | -0.0011828 |
| -0.0014862 | -9.65E-08 | -25.425731 | -25.428164 | -0.0024328 |
| -0.0014216 | -7.71E-07 | -9.4074762 | -9.4092108 | -0.0017346 |
| -0.0019769 | 2.57E-05 | -3.2441757 | -3.207677 | 0.03649867 |
| -0.0007485 | 2.04E-05 | -1.1672033 | -1.1447541 | 0.02244925 |
| -0.0022842 | 7.17E-06 | -2.1805161 | -2.1765538 | 0.00396228 |
| -0.0023389 | 1.55E-06 | -3.0268186 | -3.0238552 | 0.00296342 |
| -0.0023389 | 1.54E-06 | -3.0268186 | -3.0089117 | 0.0179069 |

| | | | | |
|---|---|---|---|---|
| -0.0009086 | 4.15E-06 | -0.7328092 | -0.7166088 | 0.01620039 |
| -0.0009152 | -2.46E-06 | -0.7328092 | -0.7411764 | -0.0083672 |
| -0.016904 | 2.21E-05 | -5.7053301 | -5.6910274 | 0.01430273 |
| -0.0061916 | 1.91E-05 | -3.2744529 | -3.2545171 | 0.01993585 |
| -0.0062524 | -4.17E-05 | -3.2744529 | -3.2942775 | -0.0198246 |
| -0.0006645 | 5.18E-06 | -3.1415467 | -3.1221898 | 0.01935691 |
| -0.0146754 | 2.22E-05 | -5.7171688 | -5.7086112 | 0.00855756 |
| -0.0029953 | 5.32E-06 | -2.7965685 | -2.7903352 | 0.00623333 |
| -0.0013255 | 3.84E-05 | -1.0882928 | -1.0631289 | 0.02516383 |
| -0.0005386 | 6.46E-06 | -1.8162707 | -1.8062388 | 0.01003194 |
| -0.0008597 | 1.77E-06 | -4.0866706 | -4.0722564 | 0.01441422 |
| -0.0008591 | 2.38E-06 | -4.0866706 | -4.0654899 | 0.02118063 |
| -0.0016835 | 4.24E-05 | -1.1892807 | -1.1557599 | 0.03352082 |
| -0.0052029 | -6.08E-07 | -2.5771707 | -2.5784831 | -0.0013124 |
| -0.0052024 | -5.00E-08 | -2.5771707 | -2.5719409 | 0.00522983 |
| -0.0051902 | 1.21E-05 | -2.5771707 | -2.5759881 | 0.00118268 |
| -0.0052114 | -9.07E-06 | -2.5771707 | -2.5727206 | 0.00445008 |
| -0.0052209 | -1.86E-05 | -2.5771707 | -2.5874687 | -0.010298 |
| -0.0051965 | 5.75E-06 | -2.5771707 | -2.5768811 | 0.00028968 |
| -0.0052103 | -8.03E-06 | -2.5771707 | -2.5881541 | -0.0109833 |
| -0.0051685 | 3.38E-05 | -2.5771707 | -2.5717657 | 0.00540507 |
| -0.0051733 | 2.90E-05 | -2.5771707 | -2.5910124 | -0.0138416 |
| -0.0052166 | -1.43E-05 | -2.5771707 | -2.5701813 | 0.00698948 |
| -0.0052297 | -2.74E-05 | -2.5771707 | -2.5962211 | -0.0190504 |
| -0.0052368 | -3.45E-05 | -2.5771707 | -2.5907801 | -0.0136094 |
| -0.0051448 | 5.75E-05 | -2.5771707 | -2.53862 | 0.03855073 |
| -0.005173 | 2.93E-05 | -2.5771707 | -2.5852309 | -0.0080602 |
| -0.005232 | -2.97E-05 | -2.5771707 | -2.5970749 | -0.0199041 |
| -0.0051691 | 3.32E-05 | -2.5771707 | -2.5733846 | 0.00378609 |
| -0.0050347 | 0.00016756 | -2.5771707 | -2.5112628 | 0.06590796 |
| -0.0051479 | 5.44E-05 | -2.5771707 | -2.5549741 | 0.02219665 |
| -0.00517 | 3.23E-05 | -2.5771707 | -2.5568578 | 0.02031291 |
| -0.0051507 | 5.16E-05 | -2.5771707 | -2.5599154 | 0.01725531 |
| -0.0051328 | 6.95E-05 | -2.5771707 | -2.5649577 | 0.01221299 |
| -0.0051826 | 1.97E-05 | -2.5771707 | -2.5430362 | 0.03413451 |
| -0.0051424 | 5.99E-05 | -2.5771707 | -2.5589614 | 0.01820934 |
| -0.0052425 | -4.02E-05 | -2.5771707 | -2.5859828 | -0.0088121 |
| -0.0051765 | 2.58E-05 | -2.5771707 | -2.5676891 | 0.00948167 |
| -0.0051406 | 6.17E-05 | -2.5771707 | -2.5326191 | 0.04455161 |
| -0.0052002 | 2.11E-06 | -2.5771707 | -2.5659027 | 0.01126802 |
| -0.005163 | 3.94E-05 | -2.5771707 | -2.55885 | 0.01832068 |
| -0.0052275 | -2.52E-05 | -2.5771707 | -2.5741178 | 0.00305295 |
| -0.0052592 | -5.69E-05 | -2.5771707 | -2.5963011 | -0.0191303 |
| -0.0051368 | 6.55E-05 | -2.5771707 | -2.5647866 | 0.01238418 |
| -0.0051898 | 1.25E-05 | -2.5771707 | -2.6049656 | -0.0277948 |

| | | | | |
|---|---|---|---|---|
| -0.00522 | -1.77E-05 | -2.5771707 | -2.5965873 | -0.0194166 |
| -0.0050946 | 0.00010766 | -2.5771707 | -2.5290428 | 0.04812789 |
| -0.0052198 | -1.75E-05 | -2.5771707 | -2.6157042 | -0.0385334 |
| -0.0052046 | -2.33E-06 | -2.5771707 | -2.582317 | -0.0051463 |
| -0.0036636 | -2.41E-06 | -3.4686086 | -3.47179 | -0.0031813 |
| -0.004488 | 2.55E-05 | -3.602026 | -3.5857718 | 0.01625419 |
| -0.0009824 | 4.39E-06 | -3.8487695 | -3.8263461 | 0.02242339 |
| -0.0009853 | 1.45E-06 | -3.8487695 | -3.8458268 | 0.00294268 |
| -0.0009864 | 3.02E-07 | -3.8487695 | -3.8466425 | 0.00212705 |
| -0.0009872 | -4.85E-07 | -3.8487695 | -3.8487564 | 1.31E-05 |
| -0.0056679 | 8.41E-06 | -4.5140228 | -4.5059834 | 0.00803947 |
| -0.0010263 | 4.77E-05 | -0.6854979 | -0.5971113 | 0.08838657 |
| -0.003752 | 1.63E-06 | -3.6459848 | -3.641849 | 0.00413573 |
| -0.0037587 | -5.11E-06 | -3.6459848 | -3.6484737 | -0.002489 |
| -0.0037499 | 3.67E-06 | -3.6459848 | -3.6448493 | 0.00113547 |
| -0.0037549 | -1.36E-06 | -3.6459848 | -3.649492 | -0.0035073 |
| -0.0037617 | -8.10E-06 | -3.6459848 | -3.6554528 | -0.0094681 |
| -0.0037637 | -1.02E-05 | -3.6459848 | -3.6509113 | -0.0049266 |
| -0.0037474 | 6.23E-06 | -3.6459848 | -3.644389 | 0.00159574 |
| -0.0037703 | -1.67E-05 | -3.6459848 | -3.6663754 | -0.0203906 |
| -0.0037473 | 6.26E-06 | -3.6459848 | -3.6376439 | 0.00834084 |
| -0.0033227 | 6.70E-06 | -2.7836981 | -2.7757792 | 0.00791883 |
| -0.0013753 | 1.70E-06 | -1.0337586 | -1.0337312 | 2.74E-05 |
| -0.0012548 | 5.47E-07 | -1.2869626 | -1.2915722 | -0.0046096 |
| -0.002806 | 4.42E-05 | -1.9903347 | -1.9482592 | 0.04207546 |
| -0.0003102 | 4.09E-06 | -1.3434725 | -1.3321735 | 0.01129907 |
| -0.0064334 | 8.13E-06 | -5.366035 | -5.3612995 | 0.00473547 |
| -0.006423 | 1.85E-05 | -5.366035 | -5.3530991 | 0.01293588 |
| -0.0064565 | -1.50E-05 | -5.366035 | -5.3683403 | -0.0023053 |
| -0.0064357 | 5.78E-06 | -5.366035 | -5.3638527 | 0.00218225 |
| -0.0110614 | 1.97E-05 | -5.7706823 | -5.7652533 | 0.00542903 |
| -0.0005684 | 1.20E-05 | -0.8994807 | -0.8639139 | 0.03556684 |
| -0.0100359 | -3.96E-06 | -4.6658745 | -4.6582534 | 0.00762105 |
| -0.0031018 | 7.65E-06 | -2.4009733 | -2.3908886 | 0.01008475 |
| -0.0007743 | 1.26E-05 | -0.5624813 | -0.5410397 | 0.02144158 |
| -0.0004015 | 6.96E-07 | -2.6902702 | -2.6905009 | -0.0002307 |
| -0.0045319 | 1.91E-05 | -3.475304 | -3.4745961 | 0.00070786 |
| -0.0035862 | -3.15E-07 | -3.4950849 | -3.4979621 | -0.0028772 |
| -0.0028715 | -2.13E-06 | -3.0763273 | -3.0786458 | -0.0023185 |
| -0.0028645 | 4.92E-06 | -3.0763273 | -3.072091 | 0.00423634 |
| -0.0041506 | 2.57E-06 | -2.8032361 | -2.6716806 | 0.13155556 |
| -0.0009801 | 1.59E-06 | -6.1610487 | -6.144327 | 0.01672161 |
| -0.0009802 | 1.45E-06 | -6.1610487 | -6.1566315 | 0.00441718 |
| -0.0009813 | 4.33E-07 | -6.1610487 | -6.1595745 | 0.00147414 |
| -0.0009813 | 4.33E-07 | -6.1610487 | -6.1366789 | 0.02436972 |

| | | | | |
|---|---|---|---|---|
| -0.0009815 | 2.34E-07 | -6.1610487 | -6.1557547 | 0.00529397 |
| -0.0012284 | 2.86E-05 | -0.8556025 | -0.8333705 | 0.022232 |
| -0.0048697 | 1.54E-05 | -4.2531805 | -4.2483348 | 0.00484574 |
| -0.0048815 | 3.60E-06 | -4.2531805 | -4.2400231 | 0.01315737 |
| -0.0012733 | -1.76E-06 | -6.6402045 | -6.6382499 | 0.00195467 |
| -0.000706 | 3.50E-06 | -4.2541022 | -4.2334298 | 0.02067244 |
| -0.000707 | 2.49E-06 | -4.2541022 | -4.2471603 | 0.00694191 |
| -0.0007085 | 9.52E-07 | -4.2541022 | -4.2477415 | 0.00636071 |
| -0.0037741 | 1.86E-06 | -1.5930979 | -1.5754369 | 0.01766104 |
| -0.0040304 | 1.53E-05 | -0.9076615 | -0.875371 | 0.03229043 |
| -0.0023168 | 2.65E-06 | -2.1830587 | -2.1499453 | 0.03311348 |
| -0.0023147 | 4.75E-06 | -2.1830587 | -2.1722332 | 0.01082551 |
| -0.0072481 | -2.94E-05 | -2.8418188 | -2.8484268 | -0.006608 |
| -0.0008554 | 1.56E-06 | -1.2829938 | -1.3029347 | -0.0199409 |
| -0.0017883 | 1.27E-05 | -0.967878 | -0.9542368 | 0.01364118 |
| -0.0027933 | -1.36E-06 | -3.1934713 | -3.187925 | 0.00554633 |
| -0.0027923 | -3.42E-07 | -3.1934713 | -3.1944981 | -0.0010267 |
| -0.0143317 | -1.90E-05 | -5.8164129 | -5.8185887 | -0.0021758 |
| -0.0113471 | 1.16E-05 | -4.7024236 | -4.7032564 | -0.0008328 |
| -0.008443 | 3.19E-05 | -2.9469135 | -2.8849655 | 0.06194794 |
| -0.0025676 | 1.50E-07 | -3.8751553 | -3.8752548 | -9.94E-05 |
| -0.0181085 | 7.98E-05 | -5.8143361 | -5.7956977 | 0.01863837 |
| -0.0182168 | -2.85E-05 | -5.8143361 | -5.8205967 | -0.0062606 |
| -0.0181772 | 1.11E-05 | -5.8143361 | -5.8096662 | 0.0046699 |
| -0.0021216 | -3.52E-06 | -2.8728584 | -2.8767267 | -0.0038683 |
| -0.0046196 | -8.88E-06 | -4.8763578 | -4.8852057 | -0.008848 |
| -0.0046186 | -7.88E-06 | -4.8763578 | -4.8862867 | -0.0099289 |
| -0.0046098 | 9.25E-07 | -4.8763578 | -4.8790977 | -0.0027399 |
| -0.0024353 | 8.74E-06 | -1.8055692 | -1.7926703 | 0.01289892 |
| -0.0081258 | -2.17E-05 | -4.7185683 | -4.7335916 | -0.0150232 |
| -0.0080899 | 1.42E-05 | -4.7185683 | -4.7108626 | 0.00770569 |
| -0.0080855 | 1.85E-05 | -4.7185683 | -4.7029309 | 0.0156374 |
| -0.0080906 | 1.34E-05 | -4.7185683 | -4.7276344 | -0.0090661 |
| -0.0081244 | -2.04E-05 | -4.7185683 | -4.7353809 | -0.0168126 |
| -0.0081322 | -2.82E-05 | -4.7185683 | -4.7263741 | -0.0078058 |
| -0.0080953 | 8.71E-06 | -4.7185683 | -4.7134373 | 0.00513101 |
| -0.0081 | 4.02E-06 | -4.7185683 | -4.7159982 | 0.00257015 |
| -0.0081041 | -9.83E-08 | -4.7185683 | -4.7194541 | -0.0008857 |
| -0.0081098 | -5.72E-06 | -4.7185683 | -4.7205741 | -0.0020058 |
| -0.0081146 | -1.05E-05 | -4.7185683 | -4.7267072 | -0.0081389 |
| -0.0080921 | 1.19E-05 | -4.7185683 | -4.7194262 | -0.0008578 |
| -0.0081152 | -1.12E-05 | -4.7185683 | -4.7274501 | -0.0088818 |
| -0.008043 | 6.10E-05 | -4.7185683 | -4.6725283 | 0.04604006 |
| -0.0080703 | 3.37E-05 | -4.7185683 | -4.7061753 | 0.012393 |
| -0.0081018 | 2.23E-06 | -4.7185683 | -4.7135851 | 0.00498319 |

| | | | | |
|---|---|---|---|---|
| -0.0016458 | -4.68E-06 | -2.6097411 | -2.6112989 | -0.0015578 |
| -0.0016377 | 3.40E-06 | -2.6097411 | -2.6071563 | 0.00258482 |
| -0.0005582 | -4.32E-06 | -2.2488417 | -2.2745679 | -0.0257263 |
| -0.0014923 | 1.25E-05 | -1.0979036 | -1.0856737 | 0.01222992 |
| -0.001491 | 1.39E-05 | -1.0979036 | -1.0821217 | 0.01578194 |
| -0.0015072 | -2.29E-06 | -1.0979036 | -1.1023435 | -0.0044399 |
| -0.0007906 | 5.21E-06 | -0.8395574 | -0.8348114 | 0.00474605 |
| -0.006382 | -2.65E-06 | -4.7073927 | -4.7061734 | 0.00121927 |
| -0.0071756 | 3.43E-06 | -5.4879959 | -5.4874039 | 0.00059199 |
| -0.0051348 | 1.46E-05 | -3.5835761 | -3.5717151 | 0.01186097 |
| -0.0017435 | -8.65E-08 | -9.5841079 | -9.5800757 | 0.00403214 |
| -0.0022028 | -8.32E-06 | -2.2215521 | -2.2232262 | -0.0016741 |
| -0.0021806 | 1.39E-05 | -2.2215521 | -2.203234 | 0.01831818 |
| -0.0014536 | 3.59E-05 | -0.7571855 | -0.7198916 | 0.03729388 |
| -0.004153 | 1.71E-05 | -1.958903 | -1.9434823 | 0.01542062 |
| -0.0059341 | 3.93E-05 | -3.1241008 | -3.1083297 | 0.01577115 |
| -0.0059839 | -1.05E-05 | -3.1241008 | -3.1322274 | -0.0081266 |
| -0.0059611 | 1.24E-05 | -3.1241008 | -3.107224 | 0.01687682 |
| -0.0059644 | 9.05E-06 | -3.1241008 | -3.1320754 | -0.0079746 |
| -0.0059616 | 1.18E-05 | -3.1241008 | -3.1233369 | 0.00076389 |
| -0.0020505 | 6.14E-06 | -1.3761867 | -1.3508954 | 0.02529132 |
| -0.0020516 | 4.99E-06 | -1.3761867 | -1.3712891 | 0.00489759 |
| -0.0026117 | -5.80E-06 | -3.7747123 | -3.7842764 | -0.009564 |
| -0.0026025 | 3.39E-06 | -3.7747123 | -3.7679234 | 0.00678897 |
| -0.0026042 | 1.76E-06 | -3.7747123 | -3.7686986 | 0.00601375 |
| -0.0026123 | -6.37E-06 | -3.7747123 | -3.7798046 | -0.0050923 |
| -0.0026067 | -8.22E-07 | -3.7747123 | -3.7761657 | -0.0014534 |
| -0.0026065 | -5.68E-07 | -3.7747123 | -3.772537 | 0.00217533 |
| -0.0026052 | 7.70E-07 | -3.7747123 | -3.769853 | 0.00485933 |
| -0.0026041 | 1.86E-06 | -3.7747123 | -3.770617 | 0.00409532 |
| -0.0026018 | 4.11E-06 | -3.7747123 | -3.7628191 | 0.01189327 |
| -0.0021289 | 7.32E-06 | -2.9664243 | -2.9574689 | 0.00895548 |
| -0.0021219 | 1.43E-05 | -2.9664243 | -2.9510415 | 0.01538289 |
| -0.0022635 | 3.29E-06 | -3.5932918 | -3.5908508 | 0.00244093 |
| -0.0022666 | 1.96E-07 | -3.5932918 | -3.592151 | 0.00114071 |
| -0.0039008 | -2.27E-05 | -2.9092921 | -2.9348532 | -0.0255611 |
| -0.0041985 | -1.08E-06 | -3.2395291 | -3.2389674 | 0.00056171 |
| -0.0154281 | 6.50E-05 | -5.9271495 | -5.898735 | 0.02841449 |
| -0.0039833 | 3.66E-05 | -4.0706265 | -4.0124202 | 0.05820632 |
| -0.0040004 | 1.95E-05 | -4.0706265 | -4.0502037 | 0.02042282 |
| -0.003559 | 4.74E-06 | -4.0985315 | -4.0959101 | 0.00262141 |
| -0.0048114 | 1.99E-05 | -2.9161085 | -2.8882259 | 0.02788258 |
| -0.0043735 | 2.43E-06 | -4.5226986 | -4.5259774 | -0.0032787 |
| -0.0019057 | -1.12E-05 | -2.267019 | -2.2669404 | 7.87E-05 |
| -0.005983 | -1.53E-05 | -5.0133686 | -5.0107055 | 0.00266314 |

| | | | | |
|---|---|---|---|---|
| -0.0059464 | 2.12E-05 | -5.0133686 | -4.9986105 | 0.01475811 |
| -0.0035358 | 7.09E-07 | -3.7914685 | -3.7895157 | 0.00195277 |
| -0.0035368 | -3.18E-07 | -3.7914685 | -3.7943767 | -0.0029082 |
| -0.0035335 | 2.94E-06 | -3.7914685 | -3.7920564 | -0.0005879 |
| -0.0035374 | -8.75E-07 | -3.7914685 | -3.7917945 | -0.000326 |
| -0.0031894 | 8.70E-06 | -3.7984121 | -3.7928652 | 0.00554693 |
| -0.0048019 | 1.12E-05 | -3.6552789 | -3.6429633 | 0.01231563 |
| -0.0012721 | -1.39E-06 | -6.8993119 | -6.9198266 | -0.0205147 |
| -0.0030946 | 3.69E-06 | -3.0378985 | -3.0309485 | 0.00695002 |
| -0.0030958 | 2.44E-06 | -3.0378985 | -3.0365032 | 0.00139534 |
| -0.0037804 | 2.62E-06 | -4.1817975 | -4.1774023 | 0.00439525 |
| -0.0037769 | 6.20E-06 | -4.1817975 | -4.1750553 | 0.00674224 |
| -0.0037804 | 2.65E-06 | -4.1817975 | -4.172771 | 0.00902653 |
| -0.0037742 | 8.89E-06 | -4.1817975 | -4.1704233 | 0.01137424 |
| -0.0053876 | 8.85E-06 | -4.8311579 | -4.821692 | 0.00946593 |
| -0.0007993 | 1.78E-06 | -5.468096 | -5.4543515 | 0.01374447 |
| -0.0025398 | 1.45E-05 | -2.4813159 | -2.4723686 | 0.00894725 |
| -0.0056763 | -1.69E-05 | -3.8005583 | -3.8260635 | -0.0255052 |
| -0.0056332 | 2.62E-05 | -3.8005583 | -3.787413 | 0.01314533 |
| -0.0056334 | 2.60E-05 | -3.8005583 | -3.7785919 | 0.02196646 |
| -0.0056232 | 3.62E-05 | -3.8005583 | -3.7721257 | 0.02843261 |
| -0.0056651 | -5.60E-06 | -3.8005583 | -3.8096066 | -0.0090482 |
| -0.0056759 | -1.64E-05 | -3.8005583 | -3.8159539 | -0.0153955 |
| -0.0006089 | 8.66E-07 | -2.8973174 | -2.8941188 | 0.00319862 |
| -0.0009837 | 2.93E-06 | -3.9833233 | -3.9728724 | 0.01045084 |
| -0.0009767 | 9.99E-06 | -3.9833233 | -3.8673671 | 0.11595613 |
| -0.0010189 | 2.06E-06 | -3.2420771 | -3.2304442 | 0.01163286 |
| -0.0022584 | 1.65E-05 | -2.0416963 | -2.0228676 | 0.01882863 |
| -0.0070251 | 0.00019439 | -2.2434704 | -2.2320018 | 0.01146865 |
| -0.0072774 | -5.78E-05 | -2.2434704 | -2.2703313 | -0.0268608 |
| -0.0073152 | -9.56E-05 | -2.2434704 | -2.2362226 | 0.00724781 |
| -0.0071433 | 7.62E-05 | -2.2434704 | -2.2338345 | 0.00963593 |
| -0.0071398 | 7.98E-05 | -2.2434704 | -2.2610313 | -0.0175608 |
| -0.0071344 | 8.52E-05 | -2.2434704 | -2.2388846 | 0.00458586 |
| -0.0009345 | 3.25E-08 | -3.3111343 | -3.313069 | -0.0019348 |
| -0.0024879 | 2.87E-06 | -1.6658168 | -1.6578374 | 0.00797945 |
| -0.007863 | 1.48E-05 | -4.2863357 | -4.28301 | 0.0033257 |
| -0.002302 | -1.12E-05 | -1.2070313 | -1.2188146 | -0.0117834 |
| -0.0047457 | -1.06E-05 | -3.7984385 | -3.8091816 | -0.010743 |
| -0.0047616 | -2.65E-05 | -3.7984385 | -3.8177929 | -0.0193543 |
| -0.0047329 | 2.22E-06 | -3.7984385 | -3.7963896 | 0.00204897 |
| -0.004763 | -2.79E-05 | -3.7984385 | -3.8079984 | -0.0095599 |
| -0.004729 | 6.08E-06 | -3.7984385 | -3.8019626 | -0.0035241 |
| -0.0047524 | -1.73E-05 | -3.7984385 | -3.811404 | -0.0129654 |
| -0.0047179 | 1.71E-05 | -3.7984385 | -3.7999471 | -0.0015086 |

| | | | | |
|---|---|---|---|---|
| -0.0047275 | 7.56E-06 | -3.7984385 | -3.8034371 | -0.0049986 |
| -0.0047157 | 1.94E-05 | -3.7984385 | -3.7486613 | 0.04977727 |
| -0.0047014 | 3.36E-05 | -3.7984385 | -3.7782897 | 0.02014887 |
| -0.0047376 | -2.52E-06 | -3.7984385 | -3.7984064 | 3.22E-05 |
| -0.0047014 | 3.37E-05 | -3.7984385 | -3.7681643 | 0.03027427 |
| -0.0047265 | 8.57E-06 | -3.7984385 | -3.7963636 | 0.00207496 |
| -0.0047635 | -2.85E-05 | -3.7984385 | -3.7814612 | 0.01697731 |
| -0.0047063 | 2.88E-05 | -3.7984385 | -3.7787271 | 0.01971149 |
| -0.0047286 | 6.50E-06 | -3.7984385 | -3.7907507 | 0.00768781 |
| -0.0047546 | -1.95E-05 | -3.7984385 | -3.8049833 | -0.0065447 |
| -0.0047403 | -5.26E-06 | -3.7984385 | -3.7983755 | 6.31E-05 |
| -0.0046999 | 3.52E-05 | -3.7984385 | -3.7678897 | 0.03054881 |
| -0.00473 | 5.04E-06 | -3.7984385 | -3.7942047 | 0.00423384 |
| -0.0047393 | -4.20E-06 | -3.7984385 | -3.7933897 | 0.00504887 |
| -0.0047561 | -2.10E-05 | -3.7984385 | -3.7873881 | 0.01105046 |
| -0.0047133 | 2.18E-05 | -3.7984385 | -3.7875247 | 0.01091385 |
| -0.0047329 | 2.23E-06 | -3.7984385 | -3.7703758 | 0.0280627 |
| -0.0047511 | -1.61E-05 | -3.7984385 | -3.8083936 | -0.009955 |
| -0.0047463 | -1.12E-05 | -3.7984385 | -3.818871 | -0.0204325 |
| -0.0047084 | 2.67E-05 | -3.7984385 | -3.7788763 | 0.01956224 |
| -0.0047049 | 3.02E-05 | -3.7984385 | -3.7769637 | 0.02147484 |
| -0.0047127 | 2.23E-05 | -3.7984385 | -3.7790868 | 0.01935172 |
| -0.0047133 | 2.18E-05 | -3.7984385 | -3.7898703 | 0.00856829 |
| -0.0047276 | 7.47E-06 | -3.7984385 | -3.7877158 | 0.01072276 |
| -0.0047362 | -1.13E-06 | -3.7984385 | -3.7890077 | 0.00943089 |
| -0.0047216 | 1.34E-05 | -3.7984385 | -3.7969368 | 0.0015018 |
| -0.0047107 | 2.44E-05 | -3.7984385 | -3.7892876 | 0.00915098 |
| -0.0047416 | -6.48E-06 | -3.7984385 | -3.7920994 | 0.00633919 |
| -0.0047315 | 3.56E-06 | -3.7984385 | -3.7762642 | 0.02217436 |
| -0.0047239 | 1.12E-05 | -3.7984385 | -3.7940131 | 0.00442541 |
| -0.0047389 | -3.84E-06 | -3.7984385 | -3.7944067 | 0.0040319 |
| -0.0047263 | 8.82E-06 | -3.7984385 | -3.7884738 | 0.0099647 |
| -0.0047329 | 2.21E-06 | -3.7984385 | -3.7953776 | 0.00306094 |
| -0.0047181 | 1.70E-05 | -3.7984385 | -3.7704661 | 0.02797246 |
| -0.0047341 | 1.03E-06 | -3.7984385 | -3.795938 | 0.00250053 |
| -0.00469 | 4.51E-05 | -3.7984385 | -3.77707 | 0.0213685 |
| -0.0047351 | -2.96E-08 | -3.7984385 | -3.7719855 | 0.02645302 |
| -0.0047351 | -1.79E-08 | -3.7984385 | -3.7990048 | -0.0005662 |
| -0.0047554 | -2.03E-05 | -3.7984385 | -3.815219 | -0.0167805 |
| -0.004711 | 2.41E-05 | -3.7984385 | -3.7704411 | 0.02799749 |
| -0.0047365 | -1.37E-06 | -3.7984385 | -3.7956845 | 0.00275409 |
| -0.0047307 | 4.37E-06 | -3.7984385 | -3.796674 | 0.00176454 |
| -0.0047352 | -1.11E-07 | -3.7984385 | -3.8075441 | -0.0091056 |
| -0.004753 | -1.79E-05 | -3.7984385 | -3.8094755 | -0.011037 |
| -0.0047609 | -2.58E-05 | -3.7984385 | -3.8225814 | -0.0241429 |

| | | | | |
|---|---|---|---|---|
| -0.0047382 | -3.10E-06 | -3.7984385 | -3.8079478 | -0.0095092 |
| -0.0047633 | -2.82E-05 | -3.7984385 | -3.827105 | -0.0286665 |
| -0.0047285 | 6.63E-06 | -3.7984385 | -3.7942548 | 0.00418377 |
| -0.0047352 | -1.08E-07 | -3.7984385 | -3.8054849 | -0.0070463 |
| -0.0047397 | -4.62E-06 | -3.7984385 | -3.7890676 | 0.00937092 |
| -0.0047527 | -1.76E-05 | -3.7984385 | -3.8066595 | -0.0082209 |
| -0.0047337 | 1.40E-06 | -3.7984385 | -3.7983004 | 0.00013816 |
| -0.0047239 | 1.12E-05 | -3.7984385 | -3.8043685 | -0.0059299 |
| -0.0047375 | -2.46E-06 | -3.7984385 | -3.7996441 | -0.0012056 |
| -0.0047224 | 1.27E-05 | -3.7984385 | -3.7854104 | 0.01302814 |
| -0.0047135 | 2.16E-05 | -3.7984385 | -3.7881255 | 0.01031303 |
| -0.0047189 | 1.62E-05 | -3.7984385 | -3.7850577 | 0.01338089 |
| -0.0047358 | -7.54E-07 | -3.7984385 | -3.7987671 | -0.0003285 |
| -0.0047201 | 1.50E-05 | -3.7984385 | -3.7826767 | 0.01576185 |
| -0.0047143 | 2.08E-05 | -3.7984385 | -3.7852345 | 0.0132041 |
| -0.0047667 | -3.16E-05 | -3.7984385 | -3.8122362 | -0.0137976 |
| -0.0047522 | -1.71E-05 | -3.7984385 | -3.7913138 | 0.00712478 |
| -0.0047638 | -2.87E-05 | -3.7984385 | -3.821444 | -0.0230055 |
| -0.0047351 | -7.18E-09 | -3.7984385 | -3.7848504 | 0.01358819 |
| -0.0047416 | -6.48E-06 | -3.7984385 | -3.8151976 | -0.016759 |
| -0.0047573 | -2.22E-05 | -3.7984385 | -3.80999 | -0.0115515 |
| -0.0046912 | 4.39E-05 | -3.7984385 | -3.780807 | 0.01763153 |
| -0.0047323 | 2.78E-06 | -3.7984385 | -3.7836982 | 0.01474035 |
| -0.0047213 | 1.38E-05 | -3.7984385 | -3.791679 | 0.00675952 |
| -0.0047452 | -1.01E-05 | -3.7984385 | -3.819315 | -0.0208764 |
| -0.0047648 | -2.97E-05 | -3.7984385 | -3.8389144 | -0.0404758 |
| -0.004764 | -2.89E-05 | -3.7984385 | -3.8468037 | -0.0483651 |
| -0.0047228 | 1.23E-05 | -3.7984385 | -3.7922322 | 0.00620639 |
| -0.0047608 | -2.57E-05 | -3.7984385 | -3.8262557 | -0.0278171 |
| -0.0047273 | 7.73E-06 | -3.7984385 | -3.7901108 | 0.00832772 |
| -0.0047461 | -1.10E-05 | -3.7984385 | -3.798686 | -0.0002475 |
| -0.004733 | 2.10E-06 | -3.7984385 | -3.7929773 | 0.00546122 |
| -0.0047591 | -2.40E-05 | -3.7984385 | -3.8081566 | -0.0097181 |
| -0.0047488 | -1.38E-05 | -3.7984385 | -3.8031006 | -0.004662 |
| -0.0047702 | -3.52E-05 | -3.7984385 | -3.8113668 | -0.0129282 |
| -0.0047389 | -3.81E-06 | -3.7984385 | -3.7949373 | 0.0035013 |
| -0.0047184 | 1.66E-05 | -3.7984385 | -3.7838523 | 0.01458621 |
| -0.0047517 | -1.67E-05 | -3.7984385 | -3.8063258 | -0.0078872 |
| -0.004739 | -3.95E-06 | -3.7984385 | -3.7975717 | 0.00086689 |
| -0.0047403 | -5.17E-06 | -3.7984385 | -3.8068423 | -0.0084038 |
| -0.004766 | -3.09E-05 | -3.7984385 | -3.8146148 | -0.0161762 |
| -0.004737 | -1.94E-06 | -3.7984385 | -3.7967514 | 0.00168717 |
| -0.0047377 | -2.63E-06 | -3.7984385 | -3.8051527 | -0.0067141 |
| -0.004358 | 6.16E-06 | -2.4044405 | -2.3666633 | 0.03777719 |
| -0.0013799 | -1.17E-05 | -1.5141555 | -1.51558 | -0.0014245 |

| | | | | |
|---|---|---|---|---|
| -0.0013549 | 1.33E-05 | -1.5141555 | -1.5030694 | 0.01108611 |
| -0.0048738 | 3.49E-05 | -3.031247 | -2.9986825 | 0.03256452 |
| -0.0024467 | 2.25E-05 | -2.0498269 | -2.0059381 | 0.04388881 |
| -0.0129071 | 0.00010138 | -4.566308 | -4.5308132 | 0.0354948 |
| -0.0038859 | 1.70E-06 | -4.6171238 | -4.6086044 | 0.00851941 |
| -0.0038927 | -5.04E-06 | -4.6171238 | -4.6064417 | 0.01068211 |
| -0.0038769 | 1.07E-05 | -4.6171238 | -4.6083679 | 0.00875592 |
| -0.0038875 | 8.28E-08 | -4.6171238 | -4.6232965 | -0.0061727 |
| -0.0038877 | -8.53E-08 | -4.6171238 | -4.6197181 | -0.0025942 |
| -0.0038827 | 4.91E-06 | -4.6171238 | -4.6067495 | 0.01037431 |
| -0.0038762 | 1.14E-05 | -4.6171238 | -4.6154428 | 0.00168109 |
| -0.0038898 | -2.19E-06 | -4.6171238 | -4.6203368 | -0.0032129 |
| -0.003878 | 9.59E-06 | -4.6171238 | -4.6124597 | 0.00466418 |
| -0.0014811 | -5.89E-06 | -2.3043573 | -2.3184948 | -0.0141375 |
| -0.0019695 | -1.95E-07 | -3.102077 | -3.1022577 | -0.0001807 |
| -0.0019636 | 5.68E-06 | -3.102077 | -3.1046031 | -0.002526 |
| -0.0019667 | 2.55E-06 | -3.102077 | -3.0986627 | 0.00341427 |
| -0.002609 | 9.31E-07 | -4.7533844 | -4.7388844 | 0.0144999 |
| -0.0022043 | 7.85E-07 | -10.494191 | -10.492832 | 0.00135827 |
| -0.0022006 | 4.50E-06 | -10.494191 | -10.491055 | 0.0031352 |
| -0.0022021 | 2.96E-06 | -10.494191 | -10.487273 | 0.00691724 |
| -0.002205 | 1.17E-07 | -10.494191 | -10.499077 | -0.0048866 |
| -0.0022068 | -1.73E-06 | -10.494191 | -10.498139 | -0.0039485 |

SSTTSEVVVGLVGNVDGSYASSSDSSLFLPFDTTSEESSVLTRQLNEDSQVMPDALAKSTDQYTEAYPDVNS
QPPPQHHPMQNNNSLGGGGGSNNNNGGGGAAQPSADMEQQQHGGKDGVLGNQAEPPPQQMLNK
GARLAAKYLDKELAGRVQQIQLLGRDMKGPAHDKLWNQLEAEIHLHRHKTVIRACRGRNDLKRPMQAP
EKQELRTASRGQSFQFYSTFPNTTYRFTIVAVNEVGDGPPAEANITTPEPTVKEKVKWLFLARNQSLRKRSME
QLLLCVLDYFIKLSPPSMLKEPYKSHVSAMSINGSTRAARRGQNRSTRTSKQTDTDTKVIEVLCKEYGCNLDE
IEIHQISSGLEAMALKSTIDLTCNDYISVFEFDIFTRLFQPWGSILRNWNFLAVTHPGYMAFLTYDEVKARLQ
ETIVSFHIQNISNSKAERNTVPLNTQVTALRNETKPLPQPQPQPPASQEHNPPQNAKMQPTPPVSAPAPKST
RNGKVYEVQEVPRGYSANKLFQTDIWCCATNELGRECTKLYTIDLNEKLGGSLPELLLKVGEPLLIRCRAFY
QQLEFEAQHIRSLMEERFGTTDEMVQRAQIRASRLEQIDKELMEAQDKVQMSEPQLCGKLSGVEGDGSL
DPTEANYACPSLKQKPDFDPVKKIDFSEFLTSLKSSEMFASHTLEGPDKENLFFVSSQLPECNAFPPVECVY
VPKLSTSTSTTGTSHLTKCDVKQKAFCVNGGECYMVKDLPNPPRYLCRCPNEFTGDRCQNYVMASFYKH
PVLNLKEQKLAVENPLNEKPPRSPDVSVRTRQGQLETTKEKFQELKQENWSLINANQTMALQLEELKQQIKE
NPCAEGWMDWAATKISGWAHALPDMVISHLFGKEEIHNNHDLIHTYRQHIVNDMNQTNLHLFVNSYN
RMSWLIHGFLLGNISLVLVENKMGKEQSRTFWHFTNSEGLGVWQSMVLSLPAVSDRFWLQIITSWEEESD
CIAENVVGMTNASVLLTVYYPPRIQSLEEPELRLEHCIEFVVHGNPAPNLHWLHNGQVLRETEIIRMEFYQQ
PAVVKHIRRGQDLGVQFTGDRAITYYEGALGSGALDVVLDTTPLAGTVAVGAHVCVCVAPEETAYYEGIVLEV
STYLQESRLKRKMMHIFKRLCQLKQCSSLTGRVIEQRIPYRGTRYPEVNRRIERFINRPDVFPDYTDILKVIQ
EVPRTYMDYLLSASQSAAMNAAVQRPSGVGSHSTPGSHQQKTYSANMDSPVINYPVTNHNSQALQLQAT
SCTNEVGTYQCSCKPAYTGQNCEHLYVPCNPSPCQNGGTCRQTSDTTYDCTCLPGFAGQNCEDNIDD
ASSPAQTQASQQNLQSVQTTHSFPSVTPDLIVQPPVMTVVPPQPPAPPPPPPPPSVPPTPAPQPIQAHPPVIPT
RHQSVQNGINDELAQPAGSRAAEEKMEDESSPSDGQMAREYKGSDKKESESSPTEVDQPSENRSDTQTS
NQGDDSKDQPFSGGATQDVKVSEGSKPWQDMTTYSTSSVSRVSAMPELSPRERSPALKSPLQSVVVRRR
VESEVQTSKELSNAVAKETLELDVKQDSASNSSEEHVTQNVSEQMEITSQQEDSHIGKEEEISNSSRSVSLCS
LLEHGANPAFRNDKGQIPADVVPDPVDMPLEMADAAASAKEIKQMLLDAVPLSCDISKAMLPNYDHVTGK
VGIETAQSLGPNAQNITWTHLAAGSRFTAQVTAVKGPYESSSVNVTQWTYPLAPANLTLASPSASALQVSWTG
HFDQRVRVSGRLCKAVSLGGTAGDGMDDRKLPSQAKDCEVPADGGASSEEWGGAEGGPLKPPEGAST
EQPFSAAAGSDQEDPDPSAGTLRSAIDIPEPAAGTQLNRAKKKPAPVKYEVGDLVWAKFNRRPWWPCTICH
ATDFVVPGPGKVEMTYTPKDGGKPVTYLVHNFEGCGGVAMGMYNLDQSIRDFAHSSFQMALSKSWPLY
GPETVAQSPLMNFLGDNNPFSLLRMTGPILRDHPGFGEGRLPNTPPLFSPPPRHHLDPHRLSAEEMGLVA
RPPALSSVRFKLSQVKDTSDIHKGSKEAKENLKKIKRAPSTAFQHAAKIKKLRTGKLSPLKSKFKTGGKLQIGRK
RSRNIVKMKPEAPKKRGEEGLHGIVSCTACGQQVNHFQKDSIYRHPTLKVLICKTCYKYYMSDDISRDSDG
GSKSASICQAPYAPSINPEDWLMKKLVPETNQDLNSSKVCYLEEAWGNLKDLENWLLQNQQREETSEKTD
WGEGPQYCQTLTQSICDGCPRCKGTKPTDCCHEQCAAGCTGPKHSDCLACLHVNRSGICELHCPPLFI
LFQTSPRKGECEEEQKYAEKESESPYNMWPLKSDAGEILSSDILGECGVSEEDSKGIWNGIEDLEAAQVNV
DDVEYYVDSSQDPDFEENEFLYDDLDLEDIPQALVATSPPSHSHLEDEIFNQSSSTPTSTTSSSPIPPSPAN

SSDASDRENDKNQESMVNGTETEIKENAMIGKPELTHSASLLETFKFLENAAADFSDEDEDEDIEGREKTM
STSKSLPSIYNPEVSEEEYECEEPPYGWISSKINTGSSDEGFQNCELVPAPFHSDSVITSEINTKDHLEDSGSI
WLCKTCIEAHQRVKFTKDHMIRKKEDVSSEAVGASGQRPVFCPVHKQEQLKLFCETCDRLTCRDCQLLEH
THDSMERKKESYSTDKMLEASPSRQNPNSANGSESDTSHVRITIPIKTPTADMSGHKRKKRQSAKSSADKTIC
HTVRQKEEVSPEAVGVNSQRPVFCPYHKKEQLKLYCETCDKLTCRDCQLLEHKEHRYQFIEEAFQNQKVII
GQPYFSVEAQTGIIRTALPNMDREAKEEYHVVIQAKDMGGHMGGLSGTTKVTITLTDVNDNPPKFPQSVYQN
GLRTLMKLNKKLAKAVGHIFQLDDGDRRKEEEIRRHSAIYGRGEARRRDGKPLTLHELTINEAAAQFCLRDH
DFRKPPDFTCLKQLISYVRHNIPGSDLERRARILLAQFQQQEHSEAKPDAVDQTSCTFRIVEENGLGEEKPD
VVVPLYDTLGHGAIRYIINTADISTVICDKPEKARILLDHVEKRETPGLKSIILMDPFEKDLKERGKRCGVHVQT
LHNIIADLRQQMEKVEVELALFQEKRLQYGDQEGLMQEISELRLRVGEMDNERLQYEKKLKSTKSLMAKLSS
VRSEALSTGESYTGAQCQQEKSGRSEVMSSSPRILQQHKEISRIQMSVCDVLESELPAEKRPDPCLQESEILS
FQPQHAQHSVSVRRLKETQDGFEFLTDHSYMDILPQFRDSYPIKYIHAFENDHFVYFLTVQRESLDSQAFHT
DSIAYRSESPSSQSDVGRPLSGSGLAAFVQAHPGNGQRVNPPPPPQVRSVTPPPPPPRGQTPPPRGTTPPP
MKLLSSSQFQPPPADPPRLPSPPAPDAGQQQVYIPKKAASKGRQPRRRQRRLPGPLDPGEPREIPPEAVRQY
EVAIKKVREQNETDIKHLRKLKHPNIIAFKGVCTQAPCYCIIMEYCAHGQLYEVLRAGRKVTPQLLVDWSTGIA
LSQVFLENNICEQPKARPCNNRTGHRKESRSVAACEKRRTRGPSKTAPKSKEFIETESSSSSSSSDSASDSEC
QHLERAGNKLTLFDSLYFCIVTFSTVGFGDVTPKIWPSKLLVVIMICVALVVLPIQFEQLAYLWMERQKSGGNY
KKKAALQAAQEATEDSPSQLSKWPGSPTSRSSDELDAWTDFRSRTNSNASTISGRLSPILASTELDDVQDDD
LLLQMVLNLITAIVCLLACFVMWKHRYQVFFVGVRLHSLTGTEIQQQKV
CCHLEVSEDAGSAVYIYAKPSIVTASETASMPVAPKYTYTPLNHLKGGTIVNLYGAVKFFKPPYISKGTDYCSVV
LLAFAICYKLNSPESCKVCTKLLEEAQLTGDNMPRRAYCFAVAFALKQNDVVKARSFYSQIMNTENRLCSNL
EAACCADKVHCCPHATTCDLEHARCISATGEQPLGSKVPAWKRAPWQGVPVSHSVITCPDHQSVCRDGT
SATLPNLGLLASWLNAELYYTDFRPVPLMEWVKIGRNIYDSSMKLVREFQPMLQVKGDEDHVVSLCYETLRD
RPGIIRGMDLNSKVADEYMIPIENLVNPRALDFHAETNYIYFADTTSFLIGRQKIDGTERETILKDDLDNVEGIA
RLGIFWVSCEAGTYIRTLCVHLGLLLGVGGQMQELRRVRSGIMGEKDHMVTMHDVLDAQWQYDNNKD
ETYLEDSEPFLLTMAKKTLNSEYIDETKKLALVPEPVEVATPKPVEVAKQPSAAAPDSVEVAKQPSAGTPDSVEVA
NQEPSILQKIKELWQRGLTLKTVSSSGMSVSQLMIDIGQRRLKVLKQNVEDCLDLRDPSEVPAIGERLPVTSV
LTGYCDPPPGVVIREELHYNPYFPGQAITMAPPIYNEILEFEDGTPATMSQIAKDVCTFLRWAAEPEHDHRKF
QQQIVAMDQSSYGTDPFQQGLTPPQMPGDHMNPYGNDSIFHDIDSDTSLTSLSDCFLASSEVNSMQARV

KNYLQGGKVIYYEMCAECSVDIDIDIDWPIAEQRVLRYGYLGKEMKLLVCSIDGWLTNGHSCLSEDQKQISS
SSFSSFYNPAKKLFISGKIQVEVLQPIDTKGLKDEDVTDLTERCYCTMRETFFRLSGMTSEANGSMQGPQQ
MRELACGGRSPEDLLDEQCNRRAVLLMLITMVLCCSPVCEKQALFAICQSMKENGLDPHVIKKAVLLIVQV
VTWLDNSLLDQTLFPVVGLGERNSCGVIKGKKIHSETCNAVVKWICEKEALLV
GIIAGNASQQQKYEENLKQVQQAVSRFEAKLAEAATSCFSSADTYKALQDHMDLCQAVEKLNNTVTFLLASA
LFCRDLGAQLLSLGSYEEEHFVANTLNKIFGESEPEIHEQHWFWIGLNRRDPGAERSWRWSDGLGFFYH

GPLSKTKCHKTAASGAMRAGFVVVIFVAATVTVVILAIVCFLFIQKFHQVVKYWFQPPFKIPQHIEEYLKEPEMPY
TEINDKLMRFFDHCEKFVTYVEENATAMYEVDAFKQGPEMKRVLEKIAATLCVPVRDLNADLVQVAFFTCSF
VSQYANKIAFHTDQVKEIFHPYAARLVTEIHRNMEELHRNVTPHTKVSPEKLNQYIQELSEKLTQNARDLHQK
KPYTKQQICELENEFLLNEFINRQKRKELSNRLNLSDQQVKIWFQNRRMKKKRVVMREQALSMY
CIFHRNITGDLKGFIDKYGFDVLVILANYLSEDEQTKRQIVVYSENLELCNQICCELEECQNPCLELDPLECG
PPPELDDMTLWQIVINILSEPPKRKKRKDINTIEDAVKFLQECKKIIVLTGAGVSVSCGIPDFRSRDGIYARLAV
CRDIGSSSSETRSCPVGSAACLLSEGHAFDVGSPQEQLKLRDKDRLVLSYEKTYSEEEKPNFCQGHNPAVTI
NVNLSNIVLLDEIGKRNLLDFSSSGIDGINFAAYLTEINKSVTKVDLLSFANDLEARADQLPKGALENALKGHA

WFGDSPPSSFGLHQLIEYGKKSGKTAGDWYGPAVVAHILRKAIEEARDPELQGITIYVAQDCTVYSSDVVD
NNDNETLMVLIAFKSLDFTWVKEMLLQTRNKTVEGFWRRPPSEWKWKASHLRILHPYVTYEATYKLLQLEKT
KVHYAGAADSTVQFIFYQPIIHRWRETDFFPCSASCGGGYQLTSAECYDLRSSRVVADQYCHYYPENIKPK
DPFGITSKISPVKSPSFQK
TPQLVESAKSNVKTIHLNRSIAVEDQNKHVSKDPRVTSTPNEVHVFESLSSSGGSPHISPTGQPEARKSRPD
NIQSLGTKLATEMRKLTSNLRIGFGAFVDKPISPYMYISPQEAIDNPCYELHQTCLPMFGYKHVLALTDEVTR
EEAAKPPEPEKPISPPPIESKHRSLVQIIYDENRKKAEAAHRILEGLGPQVELPLYNQPSDTRQYHENIKINQA
FGAQQRDAALKSKGMYQQELAKRQLGPITTEIQAAGEFYYAEDYHQQYLHKTPKGYCSQKGTGVACPIRP
AEKESQLKNRLTELYNQSEKASLALDEVRVLQRQANETASRKMDLLRSEFLEIKALIDEEEYQAMKKVAEEEK
ECPWQALIIENKKAKCGGSLLGPSWIVTAAHCLENTHPEQLQVKLGEYRIDHEDEGEQERKVSEIIMHEK
LALAIGKLGENMTIKRAAWVAVPATYYIGSYVHGALPADSPSLSNVALGKYGALVICQASEQSQKSNLADLG
LHLHLKVSENPYTSGNIASRDTAPSIIVASGNIGTKLSDNDISMFVSSDAGNTWRQIFEEEHSVLYLDQGG
NQIVHELVTLPADHWREQLYYYTHCSQAPGCGELLFSTGPWSLMIRKAEVQAPRDFPHAIRIWYKTKPEGR
QAAVQQAKEMDLSVVRLMFTAFLPDSTGSFTRRLDPVVSDAIYDSKAPNASNLKIVRMDRTAGCVTGGEE
RIVSQNVGEVYPVIQDSGQETPRGTPESGTSGQSSDTESGYLQSHSQHSVDRIYSALYACSMQNGSGERS
PHDPHVDDPMRFKPLFRIHLMERKSATVECMYGHKGPHNGHIQIVKKDEFSTKCNQTDHHRMSGGRQ
IYGAQLRIHAHFTGLRYLLYNFPVTSAILGVASNFTFLSVIVFLSYLQWIWGSMWPREPLLVQVNTRDGASL

NYTAGNRTSAQVSLAEAPPWDRDVELLVYYAEPHKPSVVLEVGLPGAKPGSLMGDPAVMMTLLPSLPEAVP
FSGHPLFVQVPLEDLGGDPIVGRLGTSTEEGKKVQRNSGKTFPAVFQRIEAEALQGALGTEPCQGAPDT
MKAIEPYKKVISHGGYYGDGLNAIVVFAVCFMPESGQPNYRYLMDNLFKYVIGTLELLVAENYMIVYLNGAT
GTVLLSMTLYCEPPTTMIAPGICDTSLPALVTLQVDFPDTDLEFIKPDPEEKRDDLEEPAKECLKHIARLSQKH
TCRVRTCPKLHAPENGRINCSSGDTSYKTVCFVTCDEGYELEGNSQLTCQGNSQWDGKEATCLEVHCPI
HLSHSVKRELASVLLFESHPKAGTVLFSQGDKGTSWYIIWKGSVNVVTHGKGLVTTLHEGDDFGQLALVND
DLMHLKKPGGFDISLFYRDIIHTAEDEDLGVYTDASSKLEDLMKKVRAKETKKRALVRLNFHLGKDLSLTVG
CLKVQARITSERKVGSFSPVHCIKTTDKAWNVLPYPMNVEVHALNMKFLLSWDNQYDQNVSFTVQYLSGY
IPLDENAIVLKIQNYFDAWQALLLKPDIFFKISWLYKKYEKSAKDLKEAIEILIEQKRQRLSTVEKLEEHMDFAS
HSTAVFYEVLLPPHFASELAYRDENENINIYRVTSKQNS
NNQTLCGYKKPPMTCLVLICVPIPHPSNIEVPLDSKTFLSRHSLDMKFSYCDERITELMGYEPEELLGHSIYE
MLISVLKNLPESLKNKREQPNEGITFAPKVTIAMSCIKWEEQTPEQILRLQRAIGATLPLQTLWMGNTVKLLDF
SWKGDSGGPLVCGKKAYGIVSWGPPTPPGVYTRVSTFIPWIRATMRRLLP
SSVFFFLPVFCLTVLYSLIGRKIWRRKRKNMGPNTSVRDKNNKQTVKMLAVVVFAFILCWLPFHVGRYLFSK
PDEPPPSTSPGQREPMEAADSEPPAAPGEENPQPGPEPPPAAEGAPNHPAPAEYVEVLLELRRVEEQLQPFL
KIYAAVRATNLDGRWNVLMDYCYTTPSGNPNDELRYDLFFSCDKDPQTTVIENGKSQMGRFSFEVFRFV
LVSMSGTTEAQTVLKKKVHHIIWEEALLFPLTEEEVLEGMLTLTLRNCDKFSRHSIVGEIKLSLANVGEPFKMA
RVIQDVLEDGVSVEVERVKLSERYRTMKLFTKIFGVGVRTASRWYQEGLRTLVDLQERNTKLTRQQQAGLR
PVSVPVWAFRQAGPEPRSSDEENAKHSSPDLEKIAASMRALALRGSDGTEMFGDLPRPRLNTGDLQKLQR
PQQHLRDHKLALVNLVCCSLVEGPDLGAPNDATRGRLVELCEELAQLEPEFILKVALYTRQELNIRGTANFL

PAIMHYSEHEATLLAEKLKDDSKFSEAITVLLTWIERGEVNRRSANQFYSMVQSANSHVRRLMNERATHEQ

DEDIWFNLFVIHQNRSKHGATNYIPEQFLDDFIDLVIWGHEHECKIAPTRNEQQLFYVSQPGSSVVTSLSP
LLNRMDIFLLVVTNPDGYAFTHTNNRMWRKTRSKNSGSLCVGVDPNRNWDAGFGGPGASTNPCSESY
SASPRDSSQSKIIRFTLGQKKTSRLPGPTTRVSAAGSETKTRGSISANNRRSQSFNNYDKSKPVNSQPPPSN
NINLLPVNAFKRLFHLKDLEISYWPLLDMMPANSLYGLNLTSLSVTNTNLSAIPYSALKHLVYLTHLNLSFNPIS

LFRPFHTLLRKIRDLLQTLTDDELHTLECNLCISQDVDFPVRTDPEVPTVSTTALAATLPAEELSAKAENTEAELA
AQEAKEAGLVSKIYPTEKLLEEAISCGEKIAGNSKLVTAMAKEAVNAAFEMTLTEGNKMEKRLFYATFATDDRK
PAHQRVQPGTMCSVAGWGRTNALSEVLPDTLQEVDLKVLDDEACLKYPGGIYRNYNTCTMMCVGDPKEH
DPKDDFVRAVEVTQDHKPELPKEKERIEGLGGSVINKSGVNRVVWKRPRLTHNGPVRRSTVIDQIPFLAVAF
HQFHSGLYTESCFVLAEGWYEDQIFHVNAFGFPPTEPSATTRAYYGNINFFGGPSSTSVKASAKLKQLEDEN
KAGEMKLKYFDGEDYTCITFQPDLSKFKMTSLDKDIVALMSRRAYDIAGCTKDVKVFLNGKRLPVKGFRSYVI
EKLRSASPLSISSDDSFEQSIHAEIVQFLNEKKQQETNKCVIVEDKNLEQKETQVKSMFKYNKESANKANRS
KELAKMCILADCTLILAWSPEEAGRYLETYKAYEQKPADLLKEKVDQDFLSIVTDCLTSVKSVNKTDALSLLSTF
ILSFNDFDLESQFDFLAVKDGDSPDSPILGTFTGAEVPSHLTSNSHILRLEFQADHSMSGRGFNITYNTFGH
KTVQDSCFNSNHVTTIGLI
SPYADEVHQQISKNIQELRLKLSPYAEELRGQVNQNADLLRQQLAPYAQELRDKLQENVDGLQAALAPYAE
SFNPIRDYVSPIALIKGLNSPLATLLSSSQSCQYTIDLRRKHVAEVDCRENHLFLPSSYKNQYGMMAQVTQT
NHLAIGARYAQQKLGVERVLIVDWDVHHGQGTQFLFEEDPSVLYFSVHRYEQGRFWPHLPASDSPAVGQ
YQSAVLTAFPLLSLFIDEENKQIITGCADGQLWIFSLLSGHQYRCVVHINLKKEREKFYSKTGKSRQQLDQIQ
RLAPLIQVILDCSHLYDYTVKLLFKLHSCLPADTLQGHRDRFLEQFRKLKDLFYRSSNLQYFKRLIQIPQLPEN
VAEKVGCPTDDTTALGNCLRVTDPQAVTLAYHLEMVNLPYPLVHYLAFTPVVDGDFLPDRPENLFANAADID
RALSAEEIEKVQQKKCVPMASKLRFIPKPNGLRPIVKVNTVFGAKTLNRKSRDKKVHYFNTQLKNLFSVLNYE
LQKSFNATADRGEAITLFCRATGSPEPEISWHRNGKLIDESEKYILRGSNTELTIRDIKNFDAGPYVCEAKNK

NRENLEGQSKADAFDELFTHAPDKLDAVKKVLLQFVNKHVGKLGLNVKDAESQFADGVILLLLIGQLGGYF
SGCCMNEKKDNAAMVSSILFNPAEFQPLDPTQEPIFPPELLTQKDKPQEKLCFKGERVMWIRPTTLKELLAL

ALDFINLMTYDFHGSWEKITGHISPLYKGKTDTGSAVYSNIDYAVTYWRNKGAPAEKIIMGIPTYGRSFTLSS
NMYEEEINETRRKHETRLVEVDSGRQIEYEHKLAQALHEMREQHDAQVKLYKEELEQTYHAKLENARLSSEN
WRACIDSSECGDFTKCTVLGAYRERQEYGSFGISEMLNYSVNIYDEGNLLSIVTSGGAHGTHVASIAAGYFI
EGPSKKTVNQMDNIVDMREYDVPYHIRLSIDLKIHVAHWYNVRYRGSIYPPEITRRDDLVERPDPVVLAFD
RTFDSMISANCTQELENIGIEVWKHSLVKSVTKSSLGLLEVTVISTAPGHKPTVSTIPDVDCLLWATGRDPNTE
VYTCQVEHISLQGPLAVHWEAQSDSARSKMLTGVGGFVLGLIFLAPGLLIYLKNKKGRPIPQPAGLLS
ELINGSFRSFSESKSIHHDKVQKKQTICHQKPNKDINTGLSKGSSVDPCCNVNLYFPQSKSMGAKPELSSF
RPGGMRASAERALHTNTDRKEAQESEQKSDKDDPEWQESLRKSKAADERRRSMAKQAREDYNRLSLQGI
SIVYSTSSYRSTANKCNSYRNNIYEEDLCRSKNTTFDQDGLEQVKLPMRRKTKIPLQRFSSLVIFPRSPSTTPF
GRVGAPGLPGQKGEAGIPGPKGEQGDSGKALAVIDDYTIGYFTRNVSLTFEGFRGQYGESRRYFGIPGPKG
IMLNSKGYTKSIDIWSVGCILAEMLSNRPIFPGKHYLDQLNHILGILGSPSQEDLNCIINVKARNYLLSLPHKN
SPGYLENGRFRPSYRNTYKEEETKACGEPPSITNGNIISELHENYQHGDSVEYDCDERFKIIGSRKIECIDGI
NCMEELLKSNVPWRYLLNTCGTDFPIKTNAEIVRTLKFLNGRNNMESEKPSSIKAARWKYHHHVESSVIRTT
QFSQYFNGAPVLYLEGRQHPIQIFYTKQPQSDYLQAALVSVFQIHQEAPSSQDILVFLTGQEEIEAMTKTCRD
TSPQTSPWQSPCVSPKTTDTEEGYQRSMVACTLLNSPRQSPSTSPRTSITEENWLSACNSRPSSPCNKRKY
YKQGAVMKNWKRRYFQLDENTIGYFKSEQEKEPLRIIPLKEVHKVQECKQSDIMMRDNLFEIVTTSRTFYVQA
KPGQGFHGNSEVNAVLSPRSESGGLGVSMVEYVLSSSPADKLDSRFRKGAFGTRDAETDGPEKGDQKGKA
LVALQLRDSGEYHCETHVQGSTVLLKMITLNVLGKSLCYDTVNEATSPPTNEGETAHPTVASNKEQRILYVVL
LVNLDADDWLSVKTEHYEYLDLDFLPKDNVLAVLRYSDKSEECTDCFL
IAKPVISPLELRMEELRHHFRIEYAVAEGAKNVMKLLGSGKVTDRKALSEAQARFNESSQKLDLLKYSLEQRLN
HSTAEMPYSDETVLIDEENEYTA
ASYAGAPMANAATQTKPLHVGNAARHGLEATCFASLGLQGNKQILDMESGLGAFYTDYTPQFLPTLQSYSW
DHQVASLSDRFEKLKQTLETNLTNLVKRNSELENQMAKLIQICQQVEVNTAMHEAKLMEECDELMEIIRQRI

GEYKYTQVGPDHNRSFIAEMTIYVKQLGRRIFAREHGSNKKLAAQSCALSLVRQLYHLGVIEPYTGQTKKKE
VSVATFRFAAGFLVPFVVIATCYGLVLARVYRSRFTRSHKTIKVILVVIIGFFVCWLPYHLVGLIIAAHKTTTRIYKG
VTVIETPLRKKTEILYPRNNTIEVELGSNVIVDCNVSSSKDNSIGISWRVNNTLVNFLFMGRVQEGNQEDSF
HPVVSIQPAPPKGQPVVLSQSTVVQLQTPGVQPASQPVIAVTGGTTQLCSHTVNVLAPATGSSSVSGKIALAK
SLKSSHLGPEIDSHDAVLRFNGAPTIGFQNDVGEKTTVRLVNSQLVTVEEQKFLTDTLYNTGILIVWDPAPYH
YNQDAKWGICLKPEDGCRDTWEHDPELESCYQLNTQSALSWKEAYISCQTQGGDLLSITNTSELSYIQAK
KVEKTMDQIEKYIMTRLYKYVFCPETTDDEKKDLVVQKRIRALHWVTPQMLCVPVNEEIPEVSDMVVKAITD
NGTSRGFYDAAGNGEDIVSFEADTGKWVARQGDKLALYIQDLLNQDKGTAITLQFLLRTTCVNQLNSFVLH
IVDKSGSIGVVCEGSATAHRSHWRETASESQETESSSVDSEDSAESPIMAEVKEKICNYLFNVTDSTAYNLAK
TDLNTAAPEKTSETGEDSSWSQACDVVLRMKVEGMTCHSCTSTIEGKIGKLQGVQRIKVSLDNQEAVVVYQ
IEELTGDDVLVRATSIEMVTSLAYTKHGRQYLAQQGIIDKISNIIIGADSDPFAGFYLPGFVKFFGNLAIVDSPQ
QDRLEEREQDIEEQLYKLESDKRLVEEKLSQLKDEVRLQYEKMHQLLDEDLRKTMEILEKAQAKFCNENAAQ
IKCKLDCQVRTEVEFFPENAQSVRVPAKMENEYTVSVQAPDLTSGAASLRIYSGDLMVGEANISYYSDMEEI
DIQENFLYYEEKLSEVLH
GASPFLGETKQETLANVSAVNYDFEEEFFSNTSALAKDFIRRLLVKDPKKRMTILDSLQHPWIKPKDTQQALS
FAEDDTCRTSQGSPAFQPPEIANGLDTFSGFKVDIWSAGVTLYNITTGLYPFEGDNIYKLFENIGKGDYTIPE
CSDMLQNIINIKNESNPVSLNTVQVSWLHSISNHSSPGERYQDIHGAQAFSPSENYQAFQAHSPQQMLDP

KVPVGCTCVLPRSSG
VQVMLNGSIEAGAFRSNSIFGSVEIFNLDPDKVSTVQIWLMQNIGGPASESCTGRSITKLKSILEERNLRVSC
PLPGTQELNILNDIKQVVMTLEQAEAEAKGSSVETSHVSVGSGVGGELQDGKIVTSGYGKESVGLGERPSSS

RQHILQICANLDTAKIMSSQKHRVHLEAQVPQPRKQYDVSHERPQDPLGRKLKETMAHIHQYLDVRDLPQ
VSTKAVAEQPTTESVTSQENPRTESLATTENPPATDCSPRTGLGATTGSSTAELTSAESPTTEFIVTSHKSSAKATS
PSFSNSKHSYMPSLSTTYSSALGAAKNHHAVHMPLLDDKVDNETSARSKKLLRYLFSFSHSSSASSLHKFHE
WIGLTDLGTEGTWCWVDGTEYRADASREFWEKNQPDNWDQGTGGREDCVHIYPVNRNFWNDANCT
AELQPPLFDLHPMRALMLMSKSSFQPPKLKDKAHWSPEFHHFLKLALTKNPKKRPTAEKLLQHLFTSQPLAR
RDDFFGHHTCDQNGNKTCLEGWMGPECSKAICRQGCSPKHGSCKTPGECRCQYGWQGQYCDKCIP
IYHRILRFKNYMVAMVNKSLLPIRFHLPGLGDTIFYTRGLKYNFELIFFWGPGSLFENEWSLKAEYKRGGNR
TEEKYTETLESIEKFFMVPLKRFLSASEFDIVFINIPELVKTHRSLMQDIFDSIVNKNDQNLYQVFINYKERLVIY
HLDKLNRNCNHEEMRTQLEVLRQQVQIYEEDFKKERSDRERLNEEKEALQKLNERSQSQLNKLNSQIKAC
LHILLPVSCEIYDDLQKADSHIQYWKDLPALQLDRAGTKWRSYKQSIYTILDEDKKLNYCIVEYAPPLQSLYAN
SQAPTSPPQNPMTGKYNVTGTNGTCVLAHMGLQLNVTYLRKDGKIGLDVLNFVPVNTSFSGSCDHSSAFL
QPKEMTPGFEDCLKGNVDLVFLFDGSGSMNAYQFNAIKEFMIEVMEKLGNSTIHFAAVQFSLNVETEFDFN
CKDRDECTIHPNICGTAQCKNSPGKYKCECAEGYTYNSTSKNCEDVDECAENTCAQECVNSPGSYACYG
RLNAENEKDSRRRQAKLQKELAEAAKEPLPVEQDDDIEVLVDETSDPAEETSPVRAVSRTSSKRLSQPAGGLL
VGVVIEATHMCMVMRGVQKMNSKTVTSTMLGVFREDAKTREEFLTLIRS
GKSSFKGDSGGPLVCQQTAQGIVSWGSAKGTPPAVYTRVSTFIPWIQKTMRMLQP
TQVQQRNLISVLQRSVDDTSQAIQRIKNDFQNLQQVVLQARKDTDWLKEKVQNLQMLAVNNSALAKANND
VTLEELKQMNSQNAEGKLQGFLDILEKQINLVRQHGESVEWRHRIATELMAIVEKLLRTLALTLRDNMISIAS
TFNVESHVETVCPYDALKIKTPKKEYGPFCGQNVPPKIETGSNAVDITFTTDISGDHTGWKISYTTTALPCPSPK
TDAFLRWLNVDAGFVASANIPGDEKVYFFFEETAEEFDFFEKLTVSRVARVCKNDVGGEKVLQKKWTTFLKA
AYCQEDYHQLFSPRCAYCAAPILEKVLTAMDKTWHPEHFFCAHCGKVFSNDGFHEKNGKPYCQKDFLAI
IILSDLDIEVIDVTDSYDISCFLHPLSMEDVFERTSKEFPMVAEVIEGPTGRQQPCNLLCPGKEIVIHKKYQAK

QEHPGRRWLDGFLWVPDGQGLFVDTFRLLHPSQPYAFTCWRIDTGARLTNHGTRLDYVLADRALALAEL
LVKHWYKELSKTTSGLPPKYALELLTIYTWESGTERAEKFSTAEGFRTVMELLCRYQELRIYWTEFYDLQSSMI
NKDMEPEMVCIDSSGRANGMGVIGHEGLLFKVSLGLIRKLLAPNCEIIQELGQLYPFEVVFGMNGRIWVK
GQWGPCSATCSPEGAGQKPQQRRWRVCNNPPPSSIPPGLPCSGSTSEDQSCLGLPFCPVDGAWSNW
HLAHPQPPKCLEPPAHPEEPSDLEELEQFARTFKQRRIKLGFTQGDVGLAMGKLYGNDFSQTTISRFEALN
VLKASEPTKCHQVYRGVRGIRFKAESRKPVRFGHFTSTSLKNESALHFGQDTFFSIKTCHGVNIKNFSFFPG
CDYILSDNSIQWTMRRVVRVKIIARNTLHSPNILSPSGMKTLEVEVGKPFKIECKVFFGFESDFLPMITWYRD
GAQPQPLKPAITVPAPSVVISQSDLLHLPLSGLLKVQPPGKEGPPARLPPSAPKPEAKTIVPAPSQLGPCNPEID
CGLGWDLTDAKVLCRELGCGSPRYVTHHCSKLSRSSAPVLLGQLECTGQEASLTQCQAQAWEGRHCPH
GHPTSIAVDPGKRFIFWSSDGALSSVHRAVLDGTDMRIVLRTTEKIKTISLDFIDTRLFWIQYDSKEDVSRIGS
PSSMSIENDESSAGSSPSTTPNAPSEQSPEGATAGTTVPGASALTTPPVSRQVQPVNPPSQPHSTVSQGPLPP
CHFTYIHSGNVYNVSATRPFIVKEEVSSPISPQIQYPTDNTVIQVEFGSPVNISCRALLGIGKPNIAVVTWEVNE
TAPPGPFSVEWRYQYRGAGRVVLAYDGVSGRAHVAEEGAQLFLEEGRADADTNVSLRLSQVGVRHEGTYI
KFEVKEYAMPSFFISIEPEKNIISSEKFETFRITIKARYFYNKKVTRADVYMRFGIMEGNEKKMMKRAMYVTEI
SRIGCLEPEVPKETETFIRSINRMFVMTLLTMAMPKFLHHIFPKPWKIFCESWDYMFAFAKGHIDKRMSEVA
PAHQRVQPGTMCSVAGWGRTNALSEVLPDTLQEVDLKVLDDEACLKYPGGIYRNYNTCTMMCVGDPKEH
EFWMDGNRLTLIPGLIGSLKQLTYLDVSKNNIEIVEEGVSGCESLQDLLLSSNSLQQLPESIGSLKKVTTLKIDE
EDEEEMEYIRSRKLAEQSGHWLIKVSESASGNSEDYDSEEEEKKKKLLASRWRSSGKAKKRRVVDSDSDSE
TLAELVHHHSTVADGLITTLHYPAPKRNKPTIYGVSPNYDKWEIERTDITMKHKLGGGQYGEVYEGVWKKYN
GRRSEREQCPSANKEHNDTRERWTHKLRLSKVQSSKVDLVDIQREGTLRYMVADDTNCVGSSQWQKC
RVSSITGTEVMRIAR
GIYDQISEDCAFSIIYGDNYESLDLVANSADVANIWVTGLRYLISYGKHTLDMIESSQDNMRTSWVSQMF
PEDRETRRKKEENVTFTPLTVIIFVVICCAMLLLLYFFYKWLVYVIISVFCLASAMSLYNCLSALIRKISFGQCRI
GISIFVVLLVILVLALIISKRYFPNRLKLKTLASSVSFSNPEHGGLQSTLEAGDHAEENIYTMG
PVQMPSVISLARPVSMVPNIPGIPGPPINSSGSISPSGLPVHSEAKMRLKASLTHQVSSSLNGSSMVVGSAST
ETLQRKVRPTARVSDRSSHDRLTTLSCKVSGFYPRDITVTWLKNGESRQQETYSEGILPSGDGTYQTWVTME
QIERFFEDHENVIEILSDWTRDSENKVLFLEKNEKYAVFKNPQNFYLANKGKNEGKEMNEKNKEFLLEESF
PDPKNVSKQKTRTVKSTLNPVWNETFLFTLRPGDVERRLSIEVWDWDRTSRNDFMGAMSFGVSELLKGH
IFDTKNLGVPKGLEPIAVGLLILLLTCSLGMNSGCAMNPARDLSPRIFTAIAGWGFEVFTAGNNWWWVPTIA
SMDTNEQLPSHFNAADKWPGLVLEPLDQGNCAGSWAFSTAAVASDRISIHSMGHMTPALSPQNLISCDT

SPSEPLQALAPFGEDLRHLLPHAVRMEGAPLRPSFLLLLLLLQPGEYRAAADETELTGILGGSVTFPLRIPAAQK
SRVCKRDQGGLRTLQKKWTSFLKARLICTTPDKNLIFNIINDVFILKSPSLKEPVIYGVFTPQLNNMGLSAVCA
TKSVKIGFGSFVDKTVLPYVSTVPAKLRNPCPDRYEQCDSPVSFRHVLPLTDNASEFESRVSQQRISGNLDT
VRRWTARKVTYLKKTKYAQVNIQDQNLMDAFQEDTCQSNNGSCSFFPPNIPFAEDYCCQVKAENVLGV
KDFNGTILYKHNVAISPDFAGRVRFLGDLEKDCSLQLSDLRASENGSYGLRLIILNPRNQKEEKKWMTKIGV

LSSTTSEVVVGLVGNVDGSYASSSDSSLFLPFDTTSEESSVLTRQLNEDSQVMPDALAKSTDQYTEAYPDVNS
QPPPQHHPMQNNNSLGGGGGSNNNNGGGGAAQPSADMEQQQHGGKDGVLGNQAEPPPQQMLNK
YGARLAAKYLDKELAGRVQQIQLLGRDMKGPAHDKLWNQLEAEIHLHRHKTVIRACRGRNDLKRPMQAP
EKQELRTASRGQSFQFYSTFPNTTYRFTIVAVNEVGDGPPAEANITTPEPTVKEKVKWLFLARNQSLRKRSME
QLLLCVLDYFIKLSPPSMLKEPYKSHVSAMSINGSTRAARRGQNRSTRTSKQTDTDTKVIEVLCKEYGCNLDE
IEIHQISSGLEAMALKSTIDLTCNDYISVFEFDIFTRLFQPWGSILRNWNFLAVTHPGYMAFLTYDEVKARLQ
ETIVSFHIQNISNSKAERNTVPLNTQVTALRNETKPLPQPQPQPPASQEHNPPQNAKMQPTPPVSAPAPKST
RNGKVYEVQEVPRGYSANKLFQTDIWCCATNELGRECTKLYTIDLNEKLGGSLPELLLKVGEPLLIRCRAFY

QQLEFEAQHIRSLMEERFGTTDEMVQRAQIRASRLEQIDKELMEAQDKVQMSEPQLCGKLSGVEGDGSLD
DPTEANYACPSLKQKPDFDPVKKIDFSEFLTSLKSSEMFASHTLEGPDKENLFFVSSQLPECNAFPPVECVYS
VPKLSTSTSTTGTSHLTKCDVKQKAFCVNGGECYMVKDLPNPPRYLCRCPNEFTGDRCQNYVMASFYKH
PVLNLKEQKLAVENPLNEKPPRSPDVSVRTRQGQLETTKEKFQELKQENWSLINANQTMALQLEELKQQIKE
NPCAEGWMDWAATKISGWAHALPDMVISHLFGKEEIHNNHDLIHTYRQHIVNDMNQTNLHLFVNSYNS
RMSWLIHGFLLGNISLVLVENKMGKEQSRTFWHFTNSEGLGVWQSMVLSLPAVSDRFWLQIITSWEEESDA
CIAENVVGMTNASVLLTVYYPPRIQSLEEPELRLEHCIEFVVHGNPAPNLHWLHNGQVLRETEIIRMEFYQQ
PAVVKHIRRGQDLGVQFTGDRAITYYEGALGSGALDVVLDTTPLAGTVAVGAHVCVCVAPEETAYYEGIVLEVS
STYLQESRLKRKMMHIFKRLCQLKQCSSLTGRVIEQRIPYRGTRYPEVNRRIERFINRPDVFPDYTDILKVIQK
EVPRTYMDYLLSASQSAAMNAAVQRPSGVGSHSTPGSHQQKTYSANMDSPVINYPVTNHNSQALQLQATH
SCTNEVGTYQCSCKPAYTGQNCEHLYVPCNPSPCQNGGTCRQTSDTTYDCTCLPGFAGQNCEDNIDDC
ASSPAQTQASQQNLQSVQTTHSFPSVTPDLIVQPPVMTVVPPQPPAPPPPPPPPSVPPTPAPQPIQAHPPVIPT
RHQSVQNGINDELAQPAGSRAAEEKMEDESSPSDGQMAREYKGSDKKESESSPTEVDQPSENRSDTQTS
NQGDDSKDQPFSGGATQDVKVSEGSKPWQDMTTYSTSSVSRVSAMPELSPRERSPALKSPLQSVVVRRRS
VESEVQTSKELSNAVAKETLELDVKQDSASNSSEEHVTQNVSEQMEITSQQEDSHIGKEEEISNSSRSVSLCS
LLEHGANPAFRNDKGQIPADVVPDPVDMPLEMADAAASAKEIKQMLLDAVPLSCDISKAMLPNYDHVTGKA
VGIETAQSLGPNAQNITWTHLAAGSRFTAQVTAVKGPYESSSVNVTQWTYPLAPANLTLASPSASALQVSWTG
HFDQRVRVSGRLCKAVSLGGTAGDGMDDRKLPSQAKDCEVPADGGASSEEWGGAEGGPLKPPEGASTA
EQPFSAAAGSDQEDPDPSAGTLRSAIDIPEPAAGTQLNRAKKKPAPVKYEVGDLVWAKFNRRPWWPCTICH
ATDFVVPGPGKVEMTYTPKDGGKPVTYLVHNFEGCGGVAMGMYNLDQSIRDFAHSSFQMALSKSWPLYN
GPETVAQSPLMNFLGDNNPFSLLRMTGPILRDHPGFGEGRLPNTPPLFSPPPRHHLDPHRLSAEEMGLVAC
RPPALSSVRFKLSQVKDTSDIHKGSKEAKENLKKIKRAPSTAFQHAAKIKKLRTGKLSPLKSKFKTGGKLQIGRK
RSRNIVKMKPEAPKKRGEEGLHGIVSCTACGQQVNHFQKDSIYRHPTLKVLICKTCYKYYMSDDISRDSDG
GSKSASICQAPYAPSINPEDWLMKKLVPETNQDLNSSKVCYLEEAWGNLKDLENWLLQNQQREETSEKTD
WGEGPQYCQTLTQSICDGCPRCKGTKPTDCCHEQCAAGCTGPKHSDCLACLHVNRSGICELHCPPLFI
LFQTSPRKGECEEEQKYAEKESESPYNMWPLKSDAGEILSSDILGECGVSEEDSKGIWNGIEDLEAAQVNV
KDDVEYYVDSSQDPDFEENEFLYDDLDLEDIPQALVATSPPSHSHLEDEIFNQSSSTPTSTTSSSPIPPSPANC
SSDASDRENDKNQESMVNGTETEIKENAMIGKPELTHSASLLETFKFLENAAADFSDEDEDEDIEGREKTM
STSKSLPSIYNPEVSEEEYECEEPPYGWISSKINTGSSDEGFQNCELVPAPFHSDSVITSEINTKDHLEDSGSI
WLCKTCIEAHQRVKFTKDHMIRKKEDVSSEAVGASGQRPVFCPVHKQEQLKLFCETCDRLTCRDCQLLEH
THDSMERKKESYSTDKMLEASPSRQNPNSANGSESDTSHVRITIPIKTPTADMSGHKRKKRQSAKSSADKTIC
HTVRQKEEVSPEAVGVNSQRPVFCPYHKKEQLKLYCETCDKLTCRDCQLLEHKEHRYQFIEEAFQNQKVII
GQPYFSVEAQTGIIRTALPNMDREAKEEYHVVIQAKDMGGHMGGLSGTTKVTITLTDVNDNPPKFPQSVYQN
GLRTLMKLNKKLAKAVGHIFQLDDGDRRKEEEIRRHSAIYGRGEARRRDGKPLTLHELTINEAAAQFCLRDH
DFRKPPDFTCLKQLISYVRHNIPGSDLERRARILLAQFQQQEHSEAKPDAVDQTSCTFRIVEENGLGEEKPD
VVVPLYDTLGHGAIRYIINTADISTVICDKPEKARILLDHVEKRETPGLKSIILMDPFEKDLKERGKRCGVHVQT
LHNIIADLRQQMEKVEVELALFQEKRLQYGDQEGLMQEISELRLRVGEMDNERLQYEKKLKSTKSLMAKLSS
VRSEALSTGESYTGAQCQQEKSGRSEVMSSSPRILQQHKEISRIQMSVCDVLESELPAEKRPDPCLQESEILS
FQPQHAQHSVSVRRLKETQDGFEFLTDHSYMDILPQFRDSYPIKYIHAFENDHFVYFLTVQRESLDSQAFHT
DSIAYRSESPSSQSDVGRPLSGSGLAAFVQAHPGNGQRVNPPPPPQVRSVTPPPPPPRGQTPPPRGTTPPP
MKLLSSSQFQPPPADPPRLPSPPAPDAGQQQVYIPKKAASKGRQPRRRQRRLPGPLDPGEPREIPPEAVRQ
EVAIKKVREQNETDIKHLRKLKHPNIIAFKGVCTQAPCYCIIMEYCAHGQLYEVLRAGRKVTPQLLVDWSTGIA
LSQVFLENNICEQPKARPCNNRTGHRKESRSVAACEKRRTRGPSKTAPKSKEFIETESSSSSSSSDSASDSEC
QHLERAGNKLTLFDSLYFCIVTFSTVGFGDVTPKIWPSKLLVVIMICVALVVLPIQFEQLAYLWMERQKSGGNY
KKKAALQAAQEATEDSPSQLSKWPGSPTSRSSDELDAWTDFRSRTNSNASTISGRLSPILASTELDDVQDDD

LLLQMVLNLITAIVCLLACFVMWKHRYQVFFVGVRLHSLTGTEIQQQKV
CCHLEVSEDAGSAVYIYAKPSIVTASETASMPVAPKYTYTPLNHLKGGTIVNLYGAVKFFKPPYISKGTDYCSVV
LLAFAICYKLNSPESCKVCTKLLEEAQLTGDNMPRRAYCFAVAFALKQNDVVKARSFYSQIMNTENRLCSNL
EAACCADKVHCCPHATTCDLEHARCISATGEQPLGSKVPAWKRAPWQGVPVSHSVITCPDHQSVCRDG
SATLPNLGLLASWLNAELYYTDFRPVPLMEWVKIGRNIYDSSMKLVREFQPMLQVKGDEDHVVSLCYETLRD
RPGIIRGMDLNSKVADEYMIPIENLVNPRALDFHAETNYIYFADTTSFLIGRQKIDGTERETILKDDLDNVEGI
RLGIFWVSCEAGTYIRTLCVHLGLLLGVGGQMQELRRVRSGIMGEKDHMVTMHDVLDAQWQYDNNKD
ETYLEDSEPFLLTMAKKTLNSEYIDETKKLALVPEPVEVATPKPVEVAKQPSAAAPDSVEVAKQPSAGTPDSVEVA
NQEPSILQKIKELWQRGLTLKTVSSSGMSVSQLMIDIGQRRLKVLKQNVEDCLDLRDPSEVPAIGERLPVTS
LTGYCDPPPGVVIREELHYNPYFPGQAITMAPPIYNEILEFEDGTPATMSQIAKDVCTFLRWAAEPEHDHRK
QQQIVAMDQSSYGTDPFQQGLTPPQMPGDHMNPYGNDSIFHDIDSDTSLTSLSDCFLASSEVNSMQAR

KNYLQGGKVIYYEMCAECSVDIDIDIDWPIAEQRVLRYGYLGKEMKLLVCSIDGWLTNGHSCLSEDQKQIS
SSFSSFYNPAKKLFISGKIQVEVLQPIDTKGLKDEDVTDLTERCYCTMRETFFRLSGMTSEANGSMQGPQQ
MRELACGGRSPEDLLDEQCNRRAVLLMLITMVLCCSPVCEKQALFAICQSMKENGLDPHVIKKAVLLIVQV
VTWLDNSLLDQTLFPVVGLGERNSCGVIKGKKIHSETCNAVVKWICEKEALLV
GIIAGNASQQQKYEENLKQVQQAVSRFEAKLAEAATSCFSSADTYKALQDHMDLCQAVEKLNNTVTFLLAS
LFCRDLGAQLLSLGSYEEEHFVANTLNKIFGESEPEIHEQHWFWIGLNRRDPGAERSWRWSDGLGFFYH

GPLSKTKCHKTAASGAMRAGFVVVIFVAATVTVVILAIVCFLFIQKFHQVVKYWFQPPFKIPQHIEEYLKEPEMP
TEINDKLMRFFDHCEKFVTYVEENATAMYEVDAFKQGPEMKRVLEKIAATLCVPVRDLNADLVQVAFFTCSF
VSQYANKIAFHTDQVKEIFHPYAARLVTEIHRNMEELHRNVTPHTKVSPEKLNQYIQELSEKLTQNARDLHQ
KPYTKQQICELENEFLLNEFINRQKRKELSNRLNLSDQQVKIWFQNRRMKKKRVVMREQALSMY
CIFHRNITGDLKGFIDKYGFDVLVILANYLSEDEQTKRQIVVYSENLELCNQICCELEECQNPCLELDPLECG
PPPELDDMTLWQIVINILSEPPKRKKRKDINTIEDAVKFLQECKKIIVLTGAGVSVSCGIPDFRSRDGIYARLA
CRDIGSSSSETRSCPVGSAACLLSEGHAFDVGSPQEQLKLRDKDRLVLSYEKTYSEEEKPNFCQGHNPAVT
NVNLSNIVLLDEIGKRNLLDFSSSGIDGINFAAYLTEINKSVTKVDLLSFANDLEARADQLPKGALENALKGHA
SWFGDSPPSSFGLHQLIEYGKKSGKTAGDWYGPAVVAHILRKAIEEARDPELQGITIYVAQDCTVYSSDVVD
NNDNETLMVLIAFKSLDFTWVKEMLLQTRNKTVEGFWRRPPSEWKWKASHLRILHPYVTYEATYKLLQLEKT
KVHYAGAADSTVQFIFYQPIIHRWRETDFFPCSASCGGGYQLTSAECYDLRSSRVVADQYCHYYPENIKPK
MDPFGITSKISPVKSPSFQK
KTPQLVESAKSNVKTIHLNRSIAVEDQNKHVSKDPRVTSTPNEVHVFESLSSSGGSPHISPTGQPEARKSRPD
NIQSLGTKLATEMRKLTSNLRIGFGAFVDKPISPYMYISPQEAIDNPCYELHQTCLPMFGYKHVLALTDEVTR
EEAAKPPEPEKPISPPPIESKHRSLVQIIYDENRKKAEAAHRILEGLGPQVELPLYNQPSDTRQYHENIKINQA
FGAQQRDAALKSKGMYQQELAKRQLGPITTEIQAAGEFYYAEDYHQQYLHKTPKGYCSQKGTGVACPIRP
AEKESQLKNRLTELYNQSEKASLALDEVRVLQRQANETASRKMDLLRSEFLEIKALIDEEEYQAMKKVAEEEK
GECPWQALIIENKKAKCGGSLLGPSWIVTAAHCLENTHPEQLQVKLGEYRIDHEDEGEQERKVSEIIMHEK
LALAIGKLGENMTIKRAAWVAVPATYYIGSYVHGALPADSPSLSNVALGKYGALVICQASEQSQKSNLADLG
SLHLHLKVSENPYTSGNIASRDTAPSIIVASGNIGTKLSDNDISMFVSSDAGNTWRQIFEEEHSVLYLDQGG
NQIVHELVTLPADHWREQLYYYTHCSQAPGCGELLFSTGPWSLMIRKAEVQAPRDFPHAIRIWYKTKPEGR
RQAAVQQAKEMDLSVVRLMFTAFLPDSTGSFTRRLDPVVSDAIYDSKAPNASNLKIVRMDRTAGCVTGGEE
RIVSQNVGEVYPVIQDSGQETPRGTPESGTSGQSSDTESGYLQSHSQHSVDRIYSALYACSMQNGSGERS
PHDPHVDDPMRFKPLFRIHLMERKSATVECMYGHKGPHNGHIQIVKKDEFSTKCNQTDHHRMSGGRQ
IYGAQLRIHAHFTGLRYLLYNFPVTSAILGVASNFTFLSVIVFLSYLQWIWGSMWPREPLLVQNTRDGASL

NYTAGNRTSAQVSLAEAPPWDRDVELLVYYAEPHKPSVVLEVGLPGAKPGSLMGDPAVMMTLLPSLPEAVP
FSGHPLFVQVPLEDLGGDPIVGRLGTSTEEGKKVQRNSGKTFPAVFQRIEAEALQGALGTEPCQGAPDT
MKAIEPYKKVISHGGYYGDGLNAIVVFAVCFMPESGQPNYRYLMDNLFKYVIGTLELLVAENYMIVYLNGAT
GTVLLSMTLYCEPPTTMIAPGICDTSLPALVTLQVDFPDTDLEFIKPDPEEKRDDLEEPAKECLKHIARLSQKH
TCRVRTCPKLHAPENGRINCSSGDTSYKTVCFVTCDEGYELEGNSQLTCQGNSQWDGKEATCLEVHCPI
LSHSVKRELASVLLFESHPKAGTVLFSQGDKGTSWYIIWKGSVNVVTHGKGLVTTLHEGDDFGQLALVNDA
DLMHLKKPGGFDISLFYRDIIHTAEDEDLGVYTDASSKLEDLMKKVRAKETKKRALVRLNFHLGKDLSLTVG
CLKVQARITSERKVGSFSPVHCIKTTDKAWNVLPYPMNVEVHALNMKFLLSWDNQYDQNVSFTVQYLSGY
IPLDENAIVLKIQNYFDAWQALLLKPDIFFKISWLYKKYEKSAKDLKEAIEILIEQKRQRLSTVEKLEEHMDFAS
HSTAVFYEVLLPPHFASELAYRDENENINIYRVTSKQNS
NNQTLCGYKKPPMTCLVLICVPIPHPSNIEVPLDSKTFLSRHSLDMKFSYCDERITELMGYEPEELLGHSIYE
MLISVLKNLPESLKNKREQPNEGITFAPKVTIAMSCIKWEEQTPEQILRLQRAIGATLPLQTLWMGNTVKLLDF
SWKGDSGGPLVCGKKAYGIVSWGPPTPPGVYTRVSTFIPWIRATMRRLLP
SVFFFLPVFCLTVLYSLIGRKIWRRKRKNMGPNTSVRDKNNKQTVKMLAVVVFAFILCWLPFHVGRYLFSK
PDEPPPSTSPGQREPMEAADSEPPAAPGEENPQPGPEPPPAAEGAPNHPAPAEYVEVLLELRRVEEQLQPFL
KIYAAVRATNLDGRWNVLMDYCYTTPSGNPNDELRYDLFFSCDKDPQTTVIENGKSQMGRFSFEVFRFV
LVSMSGTTEAQTVLKKKVHHIIWEEALLFPLTEEEVLEGMLTLTLRNCDKFSRHSIVGEIKLSLANVGEPFKMA
RVIQDVLEDGVSVEVERVKLSERYRTMKLFTKIFGVGVRTASRWYQEGLRTLVDLQERNTKLTRQQQAGLR
VSVPVWAFRQAGPEPRSSDEENAKHSSPDLEKIAASMRALALRGSDGTEMFGDLPRPRLNTGDLQKLQR
PQQHLRDHKLALVNLVCCSLVEGPDLGAPNDATRGRLVELCEELAQLEPEFILKVALYTRQELNIRGTANFL

PAIMHYSEHEATLLAEKLKDDSKFSEAITVLLTWIERGEVNRRSANQFYSMVQSANSHVRRLMNERATHEQ

DEDIWFNLFVIHQNRSKHGATNYIPEQFLDDFIDLVIWGHEHECKIAPTRNEQQLFYVSQPGSSVVTSLSP
LLNRMDIFLLVVTNPDGYAFTHTNNRMWRKTRSKNSGSLCVGVDPNRNWDAGFGGPGASTNPCSESY
ASPRDSSQSKIIRFTLGQKKTSRLPGPTTRVSAAGSETKTRGSISANNRRSQSFNNYDKSKPVNSQPPPSN
NINLLPVNAFKRLFHLKDLEISYWPLLDMMPANSLYGLNLTSLSVTNTNLSAIPYSALKHLVYLTHLNLSFNPI
FRPFHTLLRKIRDLLQTLTDDELHTLECNLCISQDVDFPVRTDPEVPTVSTTALAATLPAEELSAKAENTEAELA
QEAKEAGLVSKIYPTEKLLEEAISCGEKIAGNSKLVTAMAKEAVNAAFEMTLTEGNKMEKRLFYATFATDDRK
AHQRVQPGTMCSVAGWGRTNALSEVLPDTLQEVDLKVLDDEACLKYPGGIYRNYNTCTMMCVGDPKE
DPKDDFVRAVEVTQDHKPELPKEKERIEGLGGSVINKSGVNRVVWKRPRLTHNGPVRRSTVIDQIPFLAVA
HQFHSGLYTESCFVLAEGWYEDQIFHVNAFGFPPTEPSATTRAYYGNINFFGGPSSTSVKASAKLKQLEDE
AGEMKLKYFDGEDYTCITFQPDLSKFKMTSLDKDIVALMSRRAYDIAGCTKDVKVFLNGKRLPVKGFRSYV
EKLRSASPLSISSDDSFEQSIHAEIVQFLNEKKQQETNKCVIVEDKNLEQKETQVKSMFKYNKESANKANRS
ELAKMCILADCTLILAWSPEEAGRYLETYKAYEQKPADLLKEKVDQDFLSIVTDCLTSVKSVNKTDALSLLSTF
LSFNDFDLESQFDFLAVKDGDSPDSPILGTFTGAEVPSHLTSNSHILRLEFQADHSMSGRGFNITYNTFG
KTVQDSCFNSNHVTTIGLI
PYADEVHQQISKNIQELRLKLSPYAEELRGQVNQNADLLRQQLAPYAQELRDKLQENVDGLQAALAPYAE
SFNPIRDYVSPIALIKGLNSPLATLLSSSQSCQYTIDLRRKHVAEVDCRENHLFLPSSYKNQYGMMAQVTQT
NHLAIGARYAQQKLGVERVLIVDWDVHHGQGTQFLFEEDPSVLYFSVHRYEQGRFWPHLPASDSPAVGQ
YQSAVLTAFPLLSLFIDEENKQIITGCADGQLWIFSLLSGHQYRCVVHINLKKEREKFYSKTGKSRQQLDQIQ
LAPLIQVILDCSHLYDYTVKLLFKLHSCLPADTLQGHRDRFLEQFRKLKDLFYRSSNLQYFKRLIQIPQLPEN
VAEKVGCPTDDTTALGNCLRVTDPQAVTLAYHLEMVNLPYPLVHYLAFTPVVDGDFLPDRPENLFANAADID
RALSAEEIEKVQQKKCVPMASKLRFIPKPNGLRPIVKVNTVFGAKTLNRKSRDKKVHYFNTQLKNLFSVLNYE
LQKSFNATADRGEAITLFCRATGSPEPEISWHRNGKLIDESEKYILRGSNTELTIRDIKNFDAGPYVCEAKN

NRENLEGQSKADAFDELFTHAPDKLDAVKKVLLQFVNKHVGKLGLNVKDAESQFADGVILLLLIGQLGGYF
SGCCMNEKKDNAAMVSSILFNPAEFQPLDPTQEPIFPPELLTQKDKPQEKLCFKGERVMWIRPTTLKELLAL

ALDFINLMTYDFHGSWEKITGHISPLYKGKTDTGSAVYSNIDYAVTYWRNKGAPAEKIIMGIPTYGRSFTLSS
NMYEEEINETRRKHETRLVEVDSGRQIEYEHKLAQALHEMREQHDAQVKLYKEELEQTYHAKLENARLSSEN
WRACIDSSECGDFTKCTVLGAYRERQEYGSFGISEMLNYSVNIYDEGNLLSIVTSGGAHGTHVASIAAGYFI
EGPSKKTVNQMDNIVDMREYDVPYHIRLSIDLKIHVAHWYNVRYRGSIYPPEITRRDDLVERPDPVVLAFD
RTFDSMISANCTQELENIGIEVWKHSLVKSVTKSSLGLLEVTVISTAPGHKPTVSTIPDVDCLLWATGRDPNTE
VYTCQVEHISLQGPLAVHWEAQSDSARSKMLTGVGGFVLGLIFLAPGLLIYLKNKKGRPIPQPAGLLS
ELINGSFRSFSESKSIHHDKVQKKQTICHQKPNKDINTGLSKGSSVDPCCNVNLYFPQSKSMGAKPELSSF
RPGGMRASAERALHTNTDRKEAQESEQKSDKDDPEWQESLRKSKAADERRRSMAKQAREDYNRLSLQGI
SIVYSTSSYRSTANKCNSYRNNIYEEDLCRSKNTTFDQDGLEQVKLPMRRKTKIPLQRFSSLVIFPRSPSTTPF
GRVGAPGLPGQKGEAGIPGPKGEQGDSGKALAVIDDYTIGYFTRNVSLTFEGFRGQYGESRRYFGIPGPKG
IMLNSKGYTKSIDIWSVGCILAEMLSNRPIFPGKHYLDQLNHILGILGSPSQEDLNCIINVKARNYLLSLPHKN
SPGYLENGRFRPSYRNTYKEEETKACGEPPSITNGNIISELHENYQHGDSVEYDCDERFKIIGSRKIECIDGI
NCMEELLKSNVPWRYLLNTCGTDFPIKTNAEIVRTLKFLNGRNNMESEKPSSIKAARWKYHHHVESSVIRTT
QFSQYFNGAPVLYLEGRQHPIQIFYTKQPQSDYLQAALVSVFQIHQEAPSSQDILVFLTGQEEIEAMTKTCRD
TSPQTSPWQSPCVSPKTTDTEEGYQRSMVACTLLNSPRQSPSTSPRTSITEENWLSACNSRPSSPCNKRKY
KQGAVMKNWKRRYFQLDENTIGYFKSEQEKEPLRIIPLKEVHKVQECKQSDIMMRDNLFEIVTTSRTFYVQA
KPGQGFHGNSEVNAVLSPRSESGGLGVSMVEYVLSSSPADKLDSRFRKGAFGTRDAETDGPEKGDQKGKA
LVALQLRDSGEYHCETHVQGSTVLLKMITLNVLGKSLCYDTVNEATSPPTNEGETAHPTVASNKEQRILYVVL
LVNLDADDWLSVKTEHYEYLDLDFLPKDNVLAVLRYSDKSEECTDCFL
AKPVISPLELRMEELRHHFRIEYAVAEGAKNVMKLLGSGKVTDRKALSEAQARFNESSQKLDLLKYSLEQRLN
HSTAEMPYSDETVLIDEENEYTA
SYAGAPMANAATQTKPLHVGNAARHGLEATCFASLGLQGNKQILDMESGLGAFYTDYTPQFLPTLQSYSW
DHQVASLSDRFEKLKQTLETNLTNLVKRNSELENQMAKLIQICQQVEVNTAMHEAKLMEECDELMEIIRQRI
GEYKYTQVGPDHNRSFIAEMTIYVKQLGRRIFAREHGSNKKLAAQSCALSLVRQLYHLGVIEPYTGQTKKKE
VSVATFRFAAGFLVPFVVIATCYGLVLARVYRSRFTRSHKTIKVILVVIIGFFVCWLPYHLVGLIIAAHKTTTRIYKG
VTVIETPLRKKTEILYPRNNTIEVELGSNVIVDCNVSSSKDNSIGISWRVNNTLVNFLFMGRVQEGNQEDSFF
HPVVSIQPAPPKGQPVVLSQSTVVQLQTPGVQPASQPVIAVTGGTTQLCSHTVNVLAPATGSSSVSGKIALAK
SLKSSHLGPEIDSHDAVLRFNGAPTIGFQNDVGEKTTVRLVNSQLVTVEEQKFLTDTLYNTGILIVWDPAPYH
YNQDAKWGICLKPEDGCRDTWEHDPELESCYQLNTQSALSWKEAYISCQTQGGDLLSITNTSELSYIQAK
KVEKTMDQIEKYIMTRLYKYVFCPETTDDEKKDLVVQKRIRALHWVTPQMLCVPVNEEIPEVSDMVVKAITD
NGTSRGFYDAAGNGEDIVSFEADTGKWVARQGDKLALYIQDLLNQDKGTAITLQFLLRTTCVNQLNSFVLH
IVDKSGSIGVVCEGSATAHRSHWRETASESQETESSSVDSEDSAESPIMAEVKEKICNYLFNVTDSTAYNLAK
TDLNTAAPEKTSETGEDSSWSQACDVVLRMKVEGMTCHSCTSTIEGKIGKLQGVQRIKVSLDNQEAVVVYQ
IEELTGDDVLVRATSIEMVTSLAYTKHGRQYLAQQGIIDKISNIIIGADSDPFAGFYLPGFVKFFGNLAIVDSPQ
QDRLEEREQDIEEQLYKLESDKRLVEEKLSQLKDEVRLQYEKMHQLLDEDLRKTMEILEKAQAKFCNENAAQ
IKCKLDCQVRTEVEFFPENAQSVRVPAKMENEYTVSVQAPDLTSGAASLRIYSGDLMVGEANISYYSDMEEI
DIQENFLYYEEKLSEVLH
GASPFLGETKQETLANVSAVNYDFEEEFFSNTSALAKDFIRRLLVKDPKKRMTILDSLQHPWIKPKDTQQALS
FAEDDTCRTSQGSPAFQPPEIANGLDTFSGFKVDIWSAGVTLYNITTGLYPFEGDNIYKLFENIGKGDYTIPE
CSDMLQNIINIKNESNPVSLNTVQVSWLHSISNHSSPGERYQDIHGAQAFSPSENYQAFQAHSPQQMLDP

KVPVGCTCVLPRSSG
VQVMLNGSIEAGAFRSNSIFGSVEIFNLDPDKVSTVQIWLMQNIGGPASESCTGRSITKLKSILEERNLRVSC
PLPGTQELNILNDIKQVVMTLEQAEAEAKGSSVETSHVSVGSGVGGELQDGKIVTSGYGKESVGLGERPSSS

RQHILQICANLDTAKIMSSQKHRVHLEAQVPQPRKQYDVSHERPQDPLGRKLKETMAHIHQYLDVRDLPQ
VSTKAVAEQPTTESVTSQENPRTESLATTENPPATDCSPRTGLGATTGSSTAELTSAESPTTEFIVTSHKSSAKATS
PSFSNSKHSYMPSLSTTYSSALGAAKNHHAVHMPLLDDKVDNETSARSKKLLRYLFSFSHSSSASSLHKFHE
WIGLTDLGTEGTWCWVDGTEYRADASREFWEKNQPDNWDQGTGGREDCVHIYPVNRNFWNDANCT
LAELQPPLFDLHPMRALMLMSKSSFQPPKLKDKAHWSPEFHHFLKLALTKNPKKRPTAEKLLQHLFTSQPLAR
RDDFFGHHTCDQNGNKTCLEGWMGPECSKAICRQGCSPKHGSCKTPGECRCQYGWQGQYCDKCIP
IYHRILRFKNYMVAMVNKSLLPIRFHLPGLGDTIFYTRGLKYNFELIFFWGPGSLFENEWSLKAEYKRGGNR
TEEKYTETLESIEKFFMVPLKRFLSASEFDIVFINIPELVKTHRSLMQDIFDSIVNKNDQNLYQVFINYKERLVIY
HLDKLNRNCNHEEMRTQLEVLRQQVQIYEEDFKKERSDRERLNEEKEALQKLNERSQSQLNKLNSQIKAC
LHILLPVSCEIYDDLQKADSHIQYWKDLPALQLDRAGTKWRSYKQSIYTILDEDKKLNYCIVEYAPPLQSLYAN
TSQAPTSPPQNPMTGKYNVTGTNGTCVLAHMGLQLNVTYLRKDGKIGLDVLNFVPVNTSFSGSCDHSSAFL
QPKEMTPGFEDCLKGNVDLVFLFDGSGSMNAYQFNAIKEFMIEVMEKLGNSTIHFAAVQFSLNVETEFDFN
CKDRDECTIHPNICGTAQCKNSPGKYKCECAEGYTYNSTSKNCEDVDECAENTCAQECVNSPGSYACYC
RLNAENEKDSRRRQAKLQKELAEAAKEPLPVEQDDDIEVLVDETSDPAEETSPVRAVSRTSSKRLSQPAGGLL
VGVVIEATHMCMVMRGVQKMNSKTVTSTMLGVFREDAKTREEFLTLIRS
GKSSFKGDSGGPLVCQQTAQGIVSWGSAKGTPPAVYTRVSTFIPWIQKTMRMLQP
TQVQQRNLISVLQRSVDDTSQAIQRIKNDFQNLQQVVLQARKDTDWLKEKVQNLQMLAVNNSALAKANND
VTLEELKQMNSQNAEGKLQGFLDILEKQINLVRQHGESVEWRHRIATELMAIVEKLLRTLALTLRDNMISIAS
TFNVESHVETVCPYDALKIKTPKKEYGPFCGQNVPPKIETGSNAVDITFTTDISGDHTGWKISYTTTALPCPSPK
TDAFLRWLNVDAGFVASANIPGDEKVYFFFEETAEEFDFFEKLTVSRVARVCKNDVGGEKVLQKKWTTFLKA
AYCQEDYHQLFSPRCAYCAAPILEKVLTAMDKTWHPEHFFCAHCGKVFSNDGFHEKNGKPYCQKDFLAI
IILSDLDIEVIDVTDSYDISCFLHPLSMEDVFERTSKEFPMVAEVIEGPTGRQQPCNLLCPGKEIVIHKKYQAK

QEHPGRRWLDGFLWVPDGQGLFVDTFRLLHPSQPYAFTCWRIDTGARLTNHGTRLDYVLADRALALAELP
LVKHWYKELSKTTSGLPPKYALELLTIYTWESGTERAEKFSTAEGFRTVMELLCRYQELRIYWTEFYDLQSSMIC
NKDMEPEMVCIDSSGRANGMGVIGHEGLLFKVSLGLIRKLLAPNCEIIQELGQLYPFEVVFGMNGRIWVKA
GQWGPCSATCSPEGAGQKPQQRRWRVCNNPPPSSIPPGLPCSGSTSEDQSCLGLPFCPVDGAWSNWI
HLAHPQPPKCLEPPAHPEEPSDLEELEQFARTFKQRRIKLGFTQGDVGLAMGKLYGNDFSQTTISRFEALNI
VLKASEPTKCHQVYRGVRGIRFKAESRKPVRFGHFTSTSLKNESALHFGQDTFFSIKTCHGVNIKNFSFFPG
CDYILSDNSIQWTMRRVVRVKIIARNTLHSPNILSPSGMKTLEVEVGKPFKIECKVFFGFESDFLPMITWYRD
GAQPQPLKPAITVPAPSVVISQSDLLHLPLSGLLKVQPPGKEGPPARLPPSAPKPEAKTIVPAPSQLGPCNPEID
CGLGWDLTDAKVLCRELGCGSPRYVTHHCSKLSRSSAPVLLGQLECTGQEASLTQCQAQAWEGRHCPHH
GHPTSIAVDPGKRFIFWSSDGALSSVHRAVLDGTDMRIVLRTTEKIKTISLDFIDTRLFWIQYDSKEDVSRIGS
PSSMSIENDESSAGSSPSTTPNAPSEQSPEGATAGTTVPGASALTTPPVSRQVQPVNPPSQPHSTVSQGPLPP
CHFTYIHSGNVYNVSATRPFIVKEEVSSPISPQIQYPTDNTVIQVEFGSPVNISCRALLGIGKPNIAVVTWEVNE
TAPPGPFSVEWRYQYRGAGRVVLAYDGVSGRAHVAEEGAQLFLEEGRADADTNVSLRLSQVGVRHEGTYI
AKFEVKEYAMPSFFISIEPEKNIISSEKFETFRITIKARYFYNKKVTRADVYMRFGIMEGNEKKMMKRAMYVTEIV
SRIGCLEPEVPKETETFIRSINRMFVMTLLTMAMPKFLHHIFPKPWKIFCESWDYMFAFAKGHIDKRMSEVA
PAHQRVQPGTMCSVAGWGRTNALSEVLPDTLQEVDLKVLDDEACLKYPGGIYRNYNTCTMMCVGDPKEH
EFWMDGNRLTLIPGLIGSLKQLTYLDVSKNNIEIVEEGVSGCESLQDLLLSSNSLQQLPESIGSLKKVTTLKIDE
EDEEEMEYIRSRKLAEQSGHWLIKVSESASGNSEDYDSEEEEKKKKLLASRWRSSGKAKKRRVVDSDSDSE

LAELVHHHSTVADGLITTLHYPAPKRNKPTIYGVSPNYDKWEIERTDITMKHKLGGGQYGEVYEGVWKKYN
GRRSEREQCPSANKEHNDTRERWTHKLRLSKVQSSKVDLVDIQREGTLRYMVADDTNCVGSSQWQKC
RVSSITGTEVMRIAR
GIYDQISEDCAFSIIYGDNYESLDLVANSADVANIWVTGLRYLISYGKHTLDMIESSQDNMRTSWVSQMF
PEDRETRRKKEENVTFTPLTVIIFVVICCAMLLLLYFFYKWLVYVIISVFCLASAMSLYNCLSALIRKISFGQCRI
ISIFVVLLVILVLALIISKRYFPNRLKLKTLASSVSFSNPEHGGLQSTLEAGDHAEENIYTMG
VQMPSVISLARPVSMVPNIPGIPGPPINSSGSISPSGLPVHSEAKMRLKASLTHQVSSSLNGSSMVVGSAST
TLQRKVRPTARVSDRSSHDRLTTLSCKVSGFYPRDITVTWLKNGESRQQETYSEGILPSGDGTYQTWVTME
QIERFFEDHENVIEILSDWTRDSENKVLFLEKNEKYAVFKNPQNFYLANKGKNEGKEMNEKNKEFLLEESF
PDPKNVSKQKTRTVKSTLNPVWNETFLFTLRPGDVERRLSIEVWDWDRTSRNDFMGAMSFGVSELLKGH
IFDTKNLGVPKGLEPIAVGLLILLLTCSLGMNSGCAMNPARDLSPRIFTAIAGWGFEVFTAGNNWWWVPTI
SMDTNEQLPSHFNAADKWPGLVLEPLDQGNCAGSWAFSTAAVASDRISIHSMGHMTPALSPQNLISCDT

PSEPLQALAPFGEDLRHLLPHAVRMEGAPLRPSFLLLLLLLQPGEYRAAADETELTGILGGSVTFPLRIPAAQK
RVCKRDQGGLRTLQKKWTSFLKARLICTTPDKNLIFNIINDVFILKSPSLKEPVIYGVFTPQLNNMGLSAVC
TKSVKIGFGSFVDKTVLPYVSTVPAKLRNPCPDRYEQCDSPVSFRHVLPLTDNASEFESRVSQQRISGNLDT
VRRWTARKVTYLKKTKYAQVNIQDQNLMDAFQEDTCQSNNGSCSFFPPNIPFAEDYCCQVKAENVLGV
DFNGTILYKHNVAISPDFAGRVRFLGDLEKDCSLQLSDLRASENGSYGLRLIILNPRNQKEEKKWMTKIGV

DATETLDLRTYSVASISRAAPVLVTSMEPALFSISLPYVSFPVHSAAGSTDSDTFVTDDLFTQVFSHKSSSTTPN
GGEEEELPSKMGEQIGGRYDHPSLGALGGQQQQQQPQSAGSGSGNSSLGSGGGSGGGGGPSAVGVS
GHEPDSLKKSQGVGPIRKVLLVKEDHEGLGISITGGKEHGVPILISEIHPGQPADRCGGLHVGDAILAVNG
SFLEVAQCLQNDVIHHSITGISVNIYQQVVYFSEGNSIWAKGAASMSDVSDLKLFYAGPGKITSISVDWLY
VKNVYFTSFIPFLNSVGIVTSNGLPEVEDLSKQYEELYLKNKDIDARLFLDNDETLQRNHIECLQLERTPRKN
YSTKPGSYIFRLSCTRLGQWAIGYVTGDGNILQTIPHNKPLFQALIDGSREGFYLYPDGRSYNPDLTGLCE
PPCPMDQDSPGVESKVMSVGSPANSTPLQTEGFGQSSTPNNRAVSPVSQGSNSSTTDPKGPTQQVSGG
QNYKFRLKWHFENRELQQERWFEETDYQQDSSMVRILYAFASSMGRSDNGHYTCSSTEHPNKTTLVTVL
IPTHPEEGGSHLSNNKVEVVFWLLSMLATRDKDDMSRTLLAMSSSQESCLAMRKSGCLPLLVQLLHDTD
HAQENTGNVTRLFRGQDDGMDITKCHVSDNTMFSNTSEALSKQLECGDVTVACGDMDLTASQTVKMS
LGIEFMEAEELYQKRVLTITGICIALLVVGIVCVVAYCKTKKQRKKLHDHLRQSLRSEQNNMVNMANGPHH
LQLKLKTLEKENKKCRKAAESSHKEEEVAELLSLRQQAQELVDENDGLKMTVHRLNVELSRYQTKFRPLSQD
RRDLEIERPMPGANVVTLQCPSLLVVGDSSPAVDAVVECNSKLDPTKTTLLKMADSGGLPQVSQPAKLAEA
AIIAFDNVSLSLNCYLTINREKTSRDAAPDSSNLFIKGDSQKRIRLEQKILESLQLSTVPVSGPADQWLFTTCG
GEVSEGCLLFNKPTHYNNGNYTLVATNQLGTANQTIKGHFLEKPFPESTDNFVSIGDYEVSPTPPITVTHKPE
SLKPASYKVHFAGPSPVPCKDTVWVPRSGLRLLRPPWSPAVAAPPKPEEEEPPCPEPKQPEDAEVSKISAGV
ASARHSLGLAKRQMQSMALDAFREVGNRLQERRHLDLVYNFGSHLTDQYRPDTDPARRDSALARRLRE
HGSNSQHYSRQHMMVQGEPIGYGVQRSPSFQNKMQQEGGYANLPNKGTVQNNSGHAFQQAPASLYI
PGNNCKNGGTCVDGVNTYNCQCLPEWTGQYCTEDVDECQLMPNACQNGGTCHNNHGGYNCVC
AAQPVKTKKGVKRKADTTTPTTIDAIHESSSLPTEPKSAKMGPRRESSRPVKPPKKDVPDSQQHVVEKSNSN
EGDSNALASKEGESVSLGLCSNMEALASKDSNSDADHQSQKKKVSFPEAETTQSRKVEQESKQQASAPVS
SPRPSPLQKPSPPLSNPSQLGSVLQSSASFQTGSHQGPFDHGSSGLSPTRKSPVCKSPTAISSIYGASQKEE
KKQGSKRKFSQSDATLLGSESDEDSIRTSSSQRSHELKISTSTEKEGDPKKSSTSLKSEDSVKSSSRSKSDRD
MLLSLGLKLGDRVVIAGQKVGTLRFCGTTEFASGQWAGVELDEPEGKNDGSVGKVQYFKCAPRCGIFAP
AEGRGEGYTVDLYNASSGRHVGHTLLPWGARNHTFQSLTPGTRYNVGVSTTAGPYHSSTPNLTHWTHPL
QEPEAKEPPQTCSALPRRCVLCNCGERSLHGQGMLQRFELAPDWAEHLSGLPDGRGPGETWAPGDDL
IDPVLDCHSKMKVSNRRPYREYYVDALGDPSEKAWVAGKAIVLFEGRYQFEELPVLRRRGKQKEKGYKHK
MSTKNTILKRYDGRFKDIFQEIYEKEYRPQFEAQKIWYEHRLIDDMVAQALKSEGGFIWACKNYDGDVQSD
HPSAFDRVMRLNPMAIESPAMDFSRRLRELAGNNSTPPPVSPNRTNPMHRLLNPFQPSPKSPFLSTPPLP
AVQIVRRRGRPPSSERLKTASPLVINSQLEKHQRVCKEKDGTPPLTKEEKTAVRQSPRRIKPVRIIPSTKRTDA
MDEQCRWCAEGGNLICCDFCHNAFCKKCILRNLGRKELSAIMDENNQWHCYICHPEPLLDLVTACDS
THKHKSSYSTSSSFISTEKMDQLEFLEQEEMDISDWLLTPSDPENGSNASEEKWKCVFKPFREEYNINDW
YNSDIFQSVPNRDGRYTFGASCVRKCPYNYLATEVGSCTLVCPQDNQEVSTDSVQKCEKCSKPCPEGCY
TEGHLKESQGISASNSQAEQCDRTANASSLLNEDTSLFYSAEEMDAGETPFSSKNKESGLAHSSEKQLDKS
TTENSEDDKKRGRSTDSEVSQSPAKNGSKSVHNNHPPPSPAVTPSYQLGSSSSTSSSLGNGPGSSSNGA

IDTSTMVRRRVVSDSSEDRDTEEALKEFDFLVGSDDGDGESRSSGDGTDWEKEDQCLMPEAWNVDQG
FAESHVKMNLNLKPANKEPHSNSDSLVTTESNSIIYLEDLGSFVGSHTKNNLDLMPVNKEPDSNSDLCCKR
IKEHRYQFLEEAFQNQKGAIENLLAKLLEKKNYVNFAANQVQNRIKEVNETNKRVEQEIKVAIFTLINEINKKG
QEKSLPSGLPMSSEVANRTSSQLEASKKDLRVTKQGKVPENESPLTAIESSGVIDKAPPNSASRLRKSAAASSM
DTLITKLMEKTKYIKYTGKQIQNRILEANQNQKQVEQDIKVAIFTLMVEINKKGKALLHQLESLSKDHRIKLLQ
MSISEASIPGEEVGRLKAKDPDIGENGLVTYSIIDGDGVEMFEITTDYETQEGVVKLKKLLDYETKKSYSLKVEA
HTLLLRRVELFSLSRQVARESTYLSSLKGARSHPDESGAVQPKKLKQEVGEQSRPEPLQQPGGSESYAPPYR
DFLAFSQEMVAEQFTLMDAELFKKVVPYHCLGCIWSQRDKKGKEHLAPTIRATVSQFNSVTNCVIASCLGD
TMQEVEASGRESWHVPVPPQPEDLSIVCFTSGTTGNPKGAMLTHRNVVADFSGFLKVTEKVIFPRQDDVLIS
SMKIKVGQMHYEKQRMEQKCQILKDELASWKEKLEQKEAEVRKLQEKLMCKMKGGGVEITDRDENCKKK
STQEDMFSQSHTTASESDSTITRVEVGGSVACTPAGTLHLLHLSGQGSLGQESLSTYSSGLITPSPGAFGSH
TRIIRFCSFDSELRSYMEMPLECIFTEKRRKRSIRNEIFNVLQAAYVSKPGAVLAHEMGLSVNDDILYGVFAQ
PSWEPNAQTKRYSGSMEYMISRISPVPPGAWQDGYPPPPMNPSPISSSTQGQRGINSVPLGRQPIIMQSS
YAEIMEGLRPSWEEEAGEAGGRGEPQSQAQGALPDPTLLQYVEQLCEDESFICKVEAVIHPQFLAQLLSPE
ASGMNYLHLHKIIHRDLKSPNVLVTHTDAVKISDFGTSKELSDKSTKMSFAGTVAWMAPEVIRNEPVSEKVDI
QEEYLLPKSQTVASASGGSDQRLKDSVSSNTNKTNSSCSAFGSINAKTSNDIARELEEQFYTLVPFGRNELLS
YSRHRAQTEKHVVLCVSSLKIDLLMDFLNEFYAHPRLQDYYVVILCPTEMDAQVRRVLQIPMWSQRVIYLQ
APLSPMLYNSPSSMSPSVNKPCTVELPRLTDMAGTMNLNDGLTDNLIMDDLLDNITLPSPQQSPSGGLM

VTIVDQSNVKLTCTLFNGNLHALPKIYKIGDIVRFHRIKIREYNKQMQGITGIGFASLTFDGTVGAPIVPRTSSK
LNILVLAKSGALEDLMQTLEAALDMGASQLVKRTEFSEQVLCTVREELEENSVFRARLEDICAKLQASGQVTK
TTCCQLPTSQYGCCPLQNAVCCSDRLHCCPQGTTCDLRHSKCASTLGQMQPLTRLLDASEGAREVRCD
DGHSVLLFCPSKNWCEKLADIIAREFYNLQAERFTKTSALSPVALDKEGIEEVMDQLRRSLSGLDTVLQRTLP
AVDWIGNNLYWTNDGHRKTITVARLEKASQSRKILLEGDMSHPRGIVVDPVNGWMYWTDWEEDEIDDS
ESYLRRVIFPLEKLLTSHKRLVMKDSAVNAICYGAKIMLPGVLRYEDGIEINQEIVVITTKGEAICMAIALMTTA
KQPPVAAPDSVEVAKQPRAAAPDSVEVAKQPRAAAPDSVEVAKQPPAAAPDSVEVAKQPRAVAPDSVEMSS
WEVERKRELSKDVFTVPVKFLVIGPEEKAAIRDACLAEAPHTVPGRSHLDRDLADCSVVSCPSDPGAMSRSL
RMGMKMLMMIGLLTPLIYYLKRHKWSVLKSRKIAYRPPK
VGNPIDRLYSMQSSYFAS

SGVRGADGISLLKKRGVEVWLISEEECSKTISALKLDCEVKVNVTHKRQVIEDWRNKMSLSWKEVAYLGYE

VLERVSNIFGYKSIEDFMTSQLDYLVLEWLRMKDNGYSLSAFPYIVLNYSSLQEFYRSCYKVLVPHLVIRSQFE

AHKVSNKEKAIQEVTVLQERYEKLLGIAKEKQTLLETLLAQWQRQEKERSIFLTWLERCETAAGPTEQYVSADR
NFDRSNHDDDDIRNCAVLDLASLQWMPMQCESQLDWICKLPKGSKEWVKYQDAEYKFFEHHSTWVQ

VLEELQNNCAEEDPWGSLSVVSNAEESQSLVSDTLNTNVHSNSRSIESEKT
FELAVKNVNSPWCLLFNEEDAKVLEYLNDLKQYWKRGYGYDINSRSSCILFQDIFKHLDKAIAESKSSMPISS
KIQRNLNHLKEQLCLYPRSLRERIAGTTVDPNWAVDPYMEGLAQEVQERIEEFRRDTFLQIDDFTRTIDRET

GCDQFLVYHQDNTLVTGDQIFLIIKEVINKRQPEMVPNSRTSSTEAVAGSAPLSQGSSGIMELYGSDIEPQPS
VDFPDLPDPQAMFDIEYFRKDPRPFFKFAKEIYPGQFQPSLCHKFIALMDKEGKLLRNYTQNIDTLEQVAGI
ITFVCPSKRRGERTGPKLTAKLKCRYEIEWVTEYACHRDYLESRNCSLTSEQHDVSIDLTPLTQQPSYVTPYV
NSIRMIHNQQVVPLEQAMVKVRNVISAVDAAQFLISNDASLVIIQETKKYMDMIIGYFEQYVEWVKESPISHV

RQCSSMGSVKADMKAVIILVPVRLGGERTNMDYLEFVKGILSLEYCVGVIGGKPKQSYYFAGFQDDSLIYM
GRKYSTTGIIALNLALHICHEVNIIGFGYPDNGDNTMPVHYYDTKPLSQGVLVCSTLCVAIYAPFRFGFDSA
PKLQECNLDPCPASDGYKQIMPYDLYHPLPRWESTPWTACSSSCGGGIQSRTVSCVEEDIQGHISPVEEV

VLSFGRQDSAGSVLDGSRRSSSSSTTPTSANSLYKFLLDDQEYAIYADNVNRHENRRYIPFLPGTGRDIDT
FNEEVRKQSVSRNRDAPEGGFDAIIQATVCDEKIGWRNDASHLLVFTTDAKTHIALDGRLAGIVQPNDGQ
MRKKLILYFKRRNHARKQWEQKFCQRYDQLMEAWEKKVERIENNPRRRAKESKVREYYEKQFPEIRKQRE
AAQQES
RVCDKFDYVHNVLGKKKNEIQVLRDQIEMALTEDDDIAFLKNAAKLRHTSTKDVFIPRIDLDQTLIHTIYQAA
YISQTTDHDIALLRLEVPVNFSDYVVPICLPEKQFATSELSSIRFSTVSGWGRLIEGGATAAVLMRVDLPRVKT
RRLGQHVVGMAPLSVGSLEDVPGGDTETKMLAQPFLLDPSITLGQYVQPQGVSVLDFVRFECGEDTEIPEV
VLVAMKHTSLPIRHLWLSFDEGRSWSKYSFTSIPLFVDGVLGEPGEETLIMTVFGHFSHRSEWQLVKVDYKS
SVTWTTDQSSRPCVYTMGSPINNRALLPCQEPPVAMSTWQTTVRTASCFVVLMSGENSAEPVQLQEGLLS
YLLCDKVQKDDIQIRFYEEDENGGIWEGFGDFSPTDVHRQFAIVFKTPKYRDVNITKPASVFVQLRRKSDL
SFYSGAVVSHHETALGLPRDHHMEDPSWITRIHERSQQMERYLSTTPETTHCRKQPRPVRIQTLVGNIHIKQ
EEFRTWLREEWGRTLEDIFHEHMQELILMKFIYTSQYDNCLTYRRIYLPPSNPDDLLNPGLFKGTYGSHGL
QRREEARQRMATPRPEDQEELTTLQREALGLGKDDQERVPEGSASPPQLPGEVLETIGPTGDTAQAKELD

GQSSAGEFIFLLDRSGSMECPMDGRNHSPQRIDSAKETLVLLLKSLPLGCYFNIYSFGSQFESFYPQSVAYT

TRRKMPSLGWLRKCYQQIDRRLRKNLKSLIIVHPSWFIRTLLAITKPFISSKFSQKIRYVFTLAELAELIPMEYV
SLLLLLEQERRFLWFYRSYCNNENCSLSLVLGSAPSWDRKSISEMYKVLREWSFSNPLEALGLLTFSFASVS
FQKPESVTVHPSSCGQGPSKSGTVCQLSCPRGFTLSGVREELRCTPSGRWSENIQKALCKDFEAPQISCP
APRAATIILREDNCHFLRVDKRDFNRILKDVEANTMRLKEHGKVVLVLEKNLQGSSSSNQSPSAGSSRYSVM
YNLVQKAFKPSPVKLYRETNEPVKTKTRTFNRETGSLLLPSDTKKAQTYGNRQIVLEKEEVEEVKRFDSPGLV
LKSLSEDYSDKWITISGCQNITTTQCDFSSDIAISGIYFLRVQAMNGYNKSRWSNEIKVDPCVVNEIGPPSV
QLIFAQSRGDLTGENVNQCVLEMMIAAPDTLSVTLFFMLVLIAEHPKVEEDMMKEIQAVIGDRDIQSNDMS

YYHALDSDHLTKTHHDMFTKGQVTTGQYRMLAKHGGYVWVETQATVIYSTKNSQPQCIVCVNYVLSEIVQ
VEVHNICSFADQLLNEHGIVPGSVLYHKSLQTLVVRCKEGWVGVKTVILKKKLTAADFYNGYLHPWFQQNS

SFEAGSLEIAVISQYCNLVSFVLFYLSAAINPILYNIMSKKYRVAACSLFGLKALPKNRFSVTKQEHSCAWTES
LQRYEEILATASTTDYNNNTEGREEDQRIINLVGESMRLLGNTFVALSDLRCNLSCSAPRHLHVVRPMSHYA
KHKNQKMSTVFLHCVTKLCRTDDCPFLAPICSNRERRDAVRRTTWSAQSTSGNAVISAGPIITRSDETPTNN
QWEKLKTPEKELDAGYGEILLSISYLPAANRLLVVLIKAKNLHSKQLKDLLGKDVSVKVTLKHQSLKLKKKQTK
HYEDLNTPVERGEAESIGQMVQEAVQRFLPGASVTLAGGFRRGKTRGHDVDLLLMHPEAGREVGLLSRV
KY
LALAARLPPCRSHLRRYFCATLQLPSDWIQVVALFQSLAGAGRPLAPLPRCLRAGLADKFQQFDAYQLAKY

EMEQAKESFKNALTGILTQFEQIVAVFNASTRQKAWDHFSKAQRKNIDIWRKHSEELRNAHSEQLMGIRR

GEAIKKHIGLLHIKGKKMNMEKIPLETVRSFYIEDIVLADHPDIFNPDNPKVTQAVQAFCIEKVETLLDNAERE
HGPSANSEVEVKNVVNFIKSHGKIKAFLTLHSYSQLLLFPFGYKCTNPADYNELDTVGKAAASALASLYGTTY
CNEKEPADSTASQSTGMNENVPSSVQSSSSTNAVSCNNSSAIPQPSSAIKPWRSKSLSAKHTATSSMLSVK
STIEAGMFADLVRLQELHMVGAQLRTIEPHAFQGLRFLRVLNVSQNLLETLEENVFHSPQTLEILCINNNPLA

CSMQYDAQELEQLNRMVHRVGDEMSSLLSPPSVSQSPAHRPGIRNGSSTEASPNRQHLDNLIDEEERVF
EGMSAFVEKRKANFTDH
IKASFKGDSGGPLVCGETAQGIVSWGPVNGSPPRVYVRLSTFIPWIRKMMKKLQP
RALGDLWSYDFYSGEFVVSPEPDTSVYTLDPQKHKYIILGSDGLWNMIPPQDAISMCQDHEEKQYFMGEH
NEDAMFVFVSDVWLDQVEVLEKLHTMFSGYSSVPPTCFFFCGNFSSAPYGKNQIQSLKDSLKALADIICEYF
DLYVKDKVDETGNALKVIHEEANIRWEVCLTLSEKGFQQVSFVNSIATTKGGRHVDYVADQIVNKLIEVVKKK
TLKQGCKALLLRHQPKLRKTNVQSKCLKSKISEEPTDFSKANQTHFKMTATSQPWVVELNKEGEGKRHFCK
GSLANIAQASKEDLSLCPGIGPQKAKRLFDTLHEPFLKIPK
INECPDPGVPINARRFGDNFQLGSSISVICEEGFIKTQGTETISCILMDEKVMWSGPIPRCGAPCGGHFSS

QLQQQIDKNVADMKEKLAPLADELQVKIDQNMEELRKQLAPYAQDVQDKLNHQLEGLSFQMKKGAEHLR
LKLVETPKINSRNFDEDKLEKKKLSLESADTKSLRHGDAVLKVLQELQKLSASQQNQQRASLFYKFVTGLRSL
GRGEGFNINVPWNAVGMRDGDYLAAFLHVLLPVAFEFQPQLVLVAAGFDSVLGDPKGEMAASAACFAHL
KFSKLYSTNDMRSEEIVETTLPVLRIEHCDQSINLGKEEKFFPAINARCLWIGSSTGLFIINLANFELEAVLHYK
PPNFLRASALSEHISPVVVIPAEASSPDSEPITDLVETDTASQQSLFGNKFDDIFGSSFSNDPFNFSSQNGM
YIAGTNNMDGHIFAGIDMPAINRPLVKITADEVYNLVRGLTLQKGVRAANATFSLYTQLWNDNPDQEVMK
RTENTSLLGSSVFGKDDIYKIWRKFVSEVLESNDKIPRFYYVKADVTGAYDAIPHDKLVEVVSQILDPEKRTVY
AGNDKKQTFLQVFVQPQIIHLKNETTFENGQTTLICEAEGEPVPEITWKRAIDGITFSEGDKSPDGRIEVKGQ

LNLREFFLTPNCATEMPHFQ
KAQYPDAKLVVGNTEVGIEMRLKNMLYPVIIAPAWIPEMTFVQHTKEGITFGAACTLSCVEVVLKKAVAELPSY

SETGVGAPAFGPGSLGPFTREAGFWAYYEICTFQKGATTKRIKEQKVPYSFKENQWVGYDDQQSIKTKVQY
MSSSAANTVREELTESRMRIESLSSQLSSLQKESRAWLDKMQELEDTLANERDNYRRMLSDKEREMAEIRD
PEEPERNGVAPGAQILAIKIGDTRLSTMETGTGLIRAMIEAMKYKCDLVNYSYGEATHWPNSGRICEVINEA
IETTKLPLKFPDAETDQIMMISYMIDGQGYLITNREIVSEDIEDFEFTPKPEYEGPFCVFNEPDEAHLIQRWFE
EGLNLDKLGLKLDAKGHIIVDEFQNTSREGIYAVGDVCGRALLTPVAIAAGRKLAHRLFEGKQDSKLDYNNI

DKPPKALITEGDSKVVKLRCFPLAIAKTQLGPDKVGMKYVPVKKGSCSQAPELSDTSDEHTFIEGAPCKNDS
ICKGKVAEVSKNFGSKERSLPPRPSLPPKPKTLNSGVANGRPHRNPGVQRTISTSTQESIIRWFKEEQFPLQA
TSPTYLGPLPSRGSYQTSHQFVISSTELAHNEDQRTSSKGFLSTAVNGLRLSKTICSPGETRDIRPFYLTNAL
EPGMKGSKGDMGSRAESYSDSKIRLIGTAKGGRVEILYKGKWGTICDDKWDKQDGTVVCKMLGYSHAV
NKVPWNRLFPNADPKALDLLDKMLTFNPHKRIEVEQALAHPYLEQYYDPSDEPVAEAPFKFDMELDDLPKE
EWTSLPSCTDATCSAPPKIPNANIINEKEGRYLPGDKVQYKCQEGFENMESNYVVCKDGDWSQPPICKEA
TEKSLPPHGSPMFTGSAYIAVTRDFVQALFENPRAQQLLEWSKDTYSPDEHVWATLHRLPGMPGSVPYND
IAKHLPDGCPQMVVMPLYASLPYTQQLRVFQAAPKGYRKVILSTNIAETSITIAGIKYIVDTGMVKAKKFSPESG
SLNGRQTSYSPHHSPTPSPQSSPRVSVTDDTWLGNTNQYTSSAIVAAINALTTDSTIDMNDGIPIKSRKTAID
ANSPEDMHSWIKAISGAIVAQRGSGRSAASEHSSSSSESSYAFYFANIAATSHSTASHNNALVSNLNAKRLA
ASPFEEDKNRDLKQGDDEDVTKINGRGLPNGMDADCKDFNRTPGSRQASPTEVAERLGPNPNSTEGFG
ALGSFLVDSALIAIIAIIFIIGRKRVGNGLNFGEYPNCRLAAEQDEGHSRDATPDGELKNDIIYATLRLQPKPKL

NELPKNHPKSSIIIEELSLVSSPTLSPRQSIISTQNQYSTLSKPAALTGTLEVRLMGCQDILENVPGRSKASSITLP

LLDQQDVAIKRFPAHLGTHWVADAVSSLRKYLVDSDDSVPISRIKRIVVKVPDVRYVNRPFPNSEHEARHSF
KQVIAVKIKETKVMKLRKLAQQVANCRQCLERSTVLINQAEHILKENDHARFLQTARNVAERVAMATASSQVL

GETVEPYEVVISPDLENQLNSIIQELSLEVMPQEDPSYPVSLNIGKLAHFEPSVRQNQMGVVPWSPPQSN
AAESDPLIVGLAYVNSCLNPVIYVIMGQDFKDRFQRSLKTILRNVLNEDSLSMGDNRKKTRSTQETKSTME
DRYLFSTVTLNITEVKQEDYGHEFVCHAGEVAAYIALRQPVRNFQGYLIGGLIAPVFVIIVAILIYKYFKVDIVL
PLLQATTQTTGVDINILRRQQRMIKNRESACQSRKKKKEYMLGLESRLKAALLENEQLKRENGSLKRQLEEL
AEIHEWYRKPDYNFFVNYKAYRTRHPEQPFYILNPKMQWQLWDILQENSLEEIQPNPPSSGMLGIVIMM
DGIAEVFWIGLNQLDISGGWQWSDHTPLNFLNWNPDMQNVSPLDGSSCVAMNAASGQWQSYHCGT
IIEMDSKRVPRDKLACITQCSKHIFSAIKITKNEPASADDFLPTLIYIVLKGNPPRLQSNIQYITRFCNPSRLMT
GKESLERQERPVAVVFARAPPPAGTPTPVLLVCRVTGFYPQPVRVAWLQDGEEVTPGWWLNSSGILPNAD
NIGFSKAKDVNTTLRALEKLGEVHKENTNPPKWSLTEKKRERMQIKIKANEVTEMPPPAPEPEFPAACVEPD
PHLITAEEIKNQIEAAGFTAIFKKQPKPIKLGVIDVQRLKNTQTKSSEGTLQKSPKHINDLKTIVFRVDGMHCN
QICERYPIFMEKVFGMAEGHDPTMIGVAVDTLAILGSNVEGKQVLQKAGSRFQQLLNRIGHQAKNPPAEL
VLHLNERMQEAKKLLSSVQVMFDKTEDINFMKNTKSVKILMDRTQNCTGGSLPPPKIGHLNSKLFLNEIAK
GNLLANAANPVEFMCQAFKIVPYNIEALDKLLTESLKNNIPASGLHLFGINQLEEEDMTTNQRDEELPTLLH

RKASAVNMEKFKKFAARRKWKQSVRLISLCQRLSRSFLSRSNMSVARSDDTLDEEDSFVMKAIIHAINDDN
DCGPPLSDLLRGMLEYDPAKRFSIQQIRQHNWFRKKHTQAEALVPIPPSPETKDRWRSMTVVPYLEDLHG
PQNYQFSSSQTQDLPQKYTQDSLLEYRPFSGNDQSPTYQQNVFESHELPYCPGQSFASLLNESEHSENV

EDNYRPVQLLQCASDPDHADCRLCTNSVQ
VIHRTCTKTITKKIITGPGGPREEIVETMTSPDGSDCSHFQGLRKEEIAGQGGTLHVRVTSQGERGDLSEFD

DFGTQSYEQRVVELGKEGAQSFCRKKRTCAMHLRKYNDALLINDTVRMIDAFNALDEFYQLEKATKLLQD
SVPWLVTTETLATESRAASQHPTAGTATTDWSPSSSLSSATENPTTGDVASTENPLTKKATSVQSPRTESVCSL
LESYPSHFQTAKSSSVLVESTGFCSDDMGDDDVFEDSASVRLKSKELRAPLCSVEKDSDLDCPSPLSEKFP
LPSRWICKQAHGPAGL
SLAIELLDKVNSPDLAAPSLDDGDFEVYDIFPDKIHSCGAHSQAERTLSEIQFNQVKFGPPLRKETEPYCGL
HPGCVHGTCIEPWQCLCETNWGGQLCDKDLNYCGTHPPCLNGGTCSNTGPDKYQCSCPEGYSGQ
LELADKLSARILWIGIANFILCPLILIWQILYAFFSYTEILKREPGSLGARCWSLYGRCYLRHFNELDHELHSR
GQYCSQVEAAISCLDNISKTKEDVKLKLEECSKRANNGKFTLRDLLVVPMQRVLKYHLLLQELVKHTIDPME
WQEKEFLEKQLKQQTKDLPVSSEKCCFLPQMFLPPCVNYENCGLVVHYQDPRVQMSSRGMYDQQQH
SQDESAAFSREDRLEQAQLFCRTLGEILQHSNECAGCYRLIVYEESGDSDSHFLSKEILRHIRQQHLEEY
SLTFGRTTLVFQFLLNASTDKFFLQGVNVTTTLPSEAKETKIDVFNHSMSELRAKLGNSYKCNAEEKLWVTD
IDYKRNRNPRELLKEVKHMASLTHTFNAINYVANQVFTPERGAREDVARVMIIITDGDPSDPGGNVNAAKE
DGKKGFKLSKDRKHCEAVPVCVPLNLERNYELLYLAEQFIAIPVLYLRFRLPEVTRFSAEFDFRTYDTEGVIL
DSITNIFGLSESSLLGHHSSDAARRRSLSSFPAPQENVDVHPGSSKEVRVPTTAVYVFDAHDGEVNAVQFS

TLEDMNSQLSSFSGQMENITTVAQANEQNLKDLQEHHKDYANRTSAKFNQLEERFQLFETDIVNIISNISY
PTGTELGLAVRQAGNQSQETVTLQQSKTQMELKWAGAPGHKNEGFTLAGLLTYQGMSPILDGAERVKAP
APPHGRINPVQAKYILKDYYTLSCDTGYVLLENELVLKSFKAECQKSGSWNKPMAKCIIVNCGQPEDIDN
QLTCFQPGHFPFNVIHHSFVLPQPGVGAVFYGVFTSQWQMGDAGSTAVCAFNLEDIERVFSGKYKELNK
MFSPKCRACERPVMENYLSALQGVWHPECFVCGDCFSSFTTGSFFELDGRPYCELHFHQRQGSLCHG
RILASEIRSDSHKRHFLIPASYKGKFKRRPREFLTAYDLEIARSDKEQLHVVATKAFDSPHAELFSVSVGDQFL

RDARILPEVPGSDHCPVLAELAATFVPAPCCPSLCTRHMPEFAGTQQKLSRFLVRVERGVGAVGGHGMCR
GPHVKRLLREPCPVILDPADPTGTLGQEKRWDLLAREAAKCRDQLCCRNGLAPIRCWDVQPARPMQVMV
AKTIQQTLIVANILEACEHMTPEQRTQAFAKLSER
ETSSPCSVTCGLGRITQKRLCNNPAPRHGGKACLGLNTRSHFCNTKIPCPVDGQWSEWGEWSTCSRPG
LSFKNMCKLKPLLEKWLNDAETMSVDSTLPSPNQLSSPSLGFDGLPGRRRKKRTSIETNVRFALEKSFLANQ
EDEVLIPPFEKFKVTNFTRAQDRTFIQLLSLDSYSNYNCEYVKEKRCKARRCVFSSGMNSFSSSIQCLYLLGC
NKESKSEPLLQNTTRFHVEELEGKSFVHVATLREVTERDLNSNFICFAQNSVGNSTGVLKLKRKERALFIFILY
VKVLKRQQRMIKNRESACQSRRKKKEYLQGLECRLREALAENERLRRENALLRRRLDCALGESRELKFSPG
HRDTRVMCQEPFALQLVNGPSKCSGRLEVRYDGVWGTVCDDDWSDTNAQVVCQELGCGPAEPLAPKLG
SCDYNGGSIHLPKHSARHHLLGISLFAEHFYYPDPKTDRIRRANKFTGKVIVTISLKPPFLPPAEIKVVHPFKQP
GWERRVDNMGRIYYVDHFTRTTTWQRPTLESVRNYEQWQLQRSQLQGAMQQFNQRFIYGNQDVTAAC
IKAEYLNTSRFREDHQFFMRRDREYYGESILTIEEVTEEDLLSNFTCVALNEIENVVLTVTLQLKVPCKGDNHS
CTIYLPHLHAQQALELRITEPPVVTLRPDPLSVAPGTPAELVCEISGYYPLDVTVSWLRRGPGETGGDLLEYVT
VDGVAEVDFNSKNAAKEIGYQSLEELDGSYLYIAVTVLESTGGLSEESEFAGVRYALSPYKLNLVATPLFVKPGL
AEKVSRGETVEGKYLTYYLAQEKLSMKSIYGNVTELLLAGVDTISSTLSWSLYELSRHPQVQAALHEEITGVMKG
IKASFKGDSGGPLVCGETAQGIVSWGPVNGSPPRVYVRLSTFIPWIRKMMKKLQP
ENQLMYLPDSIGGLISVEELDCSFNEIEALPSSIGQLTNIRTFASDHNFLTQLPSEIGNWKNVTVLFLHSNKLE
EGSDEEFKPDAKDEASSDEASSGMGEGESASSETDTKMDVDSPVKVPSKRKRETDRKPAKRNSLENDLSET
LTVAVKTLKEDTMEVEEFLKEAAVMKEIKHPNLVQLLGVCTREPPFYIITEFMTYGNLLDYLRECNRQEVNAVV
RLMLRKAVKMEGERFLLEFYVPPKASKPKVSIPLSAIIEVRTTMPLEMPDKDNTFVLKVENGAEYILETIDSLQKH

SETDVDNFGHIALYKAVQFIKKLNPGLKTSKIELKFKELHRSKDKTGTEVTKEEFIEVFHELCTRPEIYFLLVQFSS
SCCNKSIEVRLIFLAGFCIATAVVWAVFRNEDRWAWILQDILGVAFCLNFIKTLKMPNFKSCVILLGLLLLYDV

MVTSRPEQGQILLQHPDAPSPAQPQVSPAQPTPSTGGRRRRTMDEDPDERRQRFLERNRAAASRCRQKR
EIDPKIKAHYSCHVEHESLLEPLSVPWEPSNSLIPIVAGVITAVLLIGVIIGVVVWKKKHPGRKGDGYTVTQGND
CGTSVIVPELEGALYLKEDGKKSWKRRYFLLRASGIYYVPKGKTKI
ADGWYKLLNQEEGEYYNVPVADAENCGLLQKFEACNYPLELYQKVRHGPGPSPTPSPTDSKHGSFFGTSR
APMMGGVIGAFTYIIFIEIHHSDTHLEEDTDMYEKYELTNMEKAENLRSAPCLDLKQNDYIGHSANTVCLS
RNQQGCSGGRIDRAWWYLRRRGVVTDSCYPFTSREKDSASAGRPCMMQSRSTGRGKRQATQRCPNP

FENGAWMVNTSSLVTIKAGRPPTILVFHSFYKGRLRFSDESYSLQITDLRLEDTGSYAVEFNTDKEQFAYREF
AYNLSTVEDVFSKGKYMQSATVEQSHTKWVRYNGEIPKPRPGACINKEARAANYMSSLNLPDKTLQFVKDH
PEGGFDAIMQAAVCEEQIGWRNVTRLLVFTSDDVFHTAGDGKLGGIYLPNDNQCHLDTDGLYSKSHIYDY
GISDCVPINMNKIVKTKPPKIVSVENIASVKQLLNVTWDKPDPTLLDLNCRLQYRNTSSNTSVTVNFTVNQNE
NVTDSPPAPQIKAPSELRESVPAQAVCSIAYYCPDYPITLTWVGLGHRTPEPTTRMDSGRTENTLTFTPTWQD

DATETLDLRTYSVASISRAAPVLVTSMEPALFSISLPYVSFPVHSAAGSTDSDTFVTDDLFTQVFSHKSSSTTPN
GGEEEELPSKMGEQIGGRYDHPSLGALGGQQQQQPQSAGSGSGNSSLGSGGGSGGGGGPSAVGVS
PGHEPDSLKKSQGVGPIRKVLLVKEDHEGLGISITGGKEHGVPILISEIHPGQPADRCGGLHVGDAILAVNGV
ESFLEVAQCLQNDVIHHSITGISVNIYQQVVYFSEGNSIWAKGAASMSDVSDLKLFYAGPGKITSISVDWLYQ
EVKNVYFTSFIPFLNSVGIVTSNGLPEVEDLSKQYEELYLKNKDIDARLFLDNDETLQRNHIECLQLERTPRKN
KYSTKPGSYIFRLSCTRLGQWAIGYVTGDGNILQTIPHNKPLFQALIDGSREGFYLYPDGRSYNPDLTGLCEI
GPPCPMDQDSPGVESKVMSVGSPANSTPLQTEGFGQSSTPNNRAVSPVSQGSNSSTTDPKGPTQQVSGG
QNYKFRLKWHFENRELQQERWFEETDYQQDSSMVRILYAFASSMGRSDNGHYTCSSTEHPNKTTLVTVLI

DIPTHPEEGGSHLSNNKVEVVFWLLSMLATRDKDDMSRTLLAMSSSQESCLAMRKSGCLPLLVQLLHDTDI
SHAQENTGNVTRLFRGQDDGMDITKCHVSDNTMFSNTSEALSKQLECGDVTVACGDMDLTASQTVKMS
LGIEFMEAEELYQKRVLTITGICIALLVVGIVCVVAYCKTKKQRKKLHDHLRQSLRSEQNNMVNMANGPHHI
LQLKLKTLEKENKKCRKAAESSHKEEEVAELLSLRQQAQELVDENDGLKMTVHRLNVELSRYQTKFRPLSQD
SRRDLEIERPMPGANVVTLQCPSLLVVGDSSPAVDAVVECNSKLDPTKTTLLKMADSGGLPQVSQPAKLAEAI
AIIAFDNVSLSLNCYLTINREKTSRDAAPDSSNLFIKGDSQKRIRLEQKILESLQLSTVPVSGPADQWLFTTCG
GEVSEGCLLFNKPTHYNNGNYTLVATNQLGTANQTIKGHFLEKPFPESTDNFVSIGDYEVSPTPPITVTHKPE
SLKPASYKVHFAGPSPVPCKDTVWVPRSGLRLLRPPWSPAVAAPPKPEEEEPPCPEPKQPEDAEVSKISAGVI
KASARHSLGLAKRQMQSMALDAFREVGNRLQERRHLDLVYNFGSHLTDQYRPDTDPARRDSALARRLREI
HGSNSQHYSRQHMMVQGEPIGYGVQRSPSFQNKMQQEGGYANLPNKGTVQNNSGHAFQQAPASLYIH
CPGNNCKNGGTCVDGVNTYNCQCLPEWTGQYCTEDVDECQLMPNACQNGGTCHNNHGGYNCVCV
TAAQPVKTKKGVKRKADTTTPTTIDAIHESSSLPTEPKSAKMGPRRESSRPVKPPKKDVPDSQQHVVEKSNSNK
SEGDSNALASKEGESVSLGLCSNMEALASKDSNSDADHQSQKKKVSFPEAETTQSRKVEQESKQQASAPVS
SPRPSPLQKPSPPLSNPSQLGSVLQSSASFQTGSHQGPFDHGSSGLSPTRKSPVCKSPTAISSIYGASQKEEA
KKQGSKRKFSQSDATLLGSESDEDSIRTSSSQRSHELKISTSTEKEGDPKKSSTSLKSEDSVKSSSRSKSDRD
AMLLSLGLKLGDRVVIAGQKVGTLRFCGTTEFASGQWAGVELDEPEGKNDGSVGKVQYFKCAPRCGIFAPL
GAEGRGEGYTVDLYNASSGRHVGHTLLPWGARNHTFQSLTPGTRYNVGVSTTAGPYHSSTPNLTHWTHPLA
AQEPEAKEPPQTCSALPRRCVLCNCGERSLHGQGMLQRFELAPDWAEHLSGLPDGRGPGETWAPGDDL
IDPVLDCHSKMKVSNRRPYREYYVDALGDPSEKAWVAGKAIVLFEGRYQFEELPVLRRRGKQKEKGYKHKV
MSTKNTILKRYDGRFKDIFQEIYEKEYRPQFEAQKIWYEHRLIDDMVAQALKSEGGFIWACKNYDGDVQSD
QHPSAFDRVMRLNPMAIESPAMDFSRRLRELAGNNSTPPPVSPNRTNPMHRLLNPFQPSPKSPFLSTPPLP
KAVQIVRRRGRPPSSERLKTASPLVINSQLEKHQRVCKEKDGTPPLTKEEKTAVRQSPRRIKPVRIIPSTKRTDAT
GMDEQCRWCAEGGNLICCDFCHNAFCKKCILRNLGRKELSAIMDENNQWHCYICHPEPLLDLVTACDS
THKHKSSYSTSSSFISTEKMDQLEFLEQEEMDISDWLLTPSDPENGSNASEEKWKCVFKPFREEYNINDWI
YNSDIFQSVPNRDGRYTFGASCVRKCPYNYLATEVGSCTLVCPQDNQEVSTDSVQKCEKCSKPCPEGCY
VTEGHLKESQGISASNSQAEQCDRTANASSLLNEDTSLFYSAEEMDAGETPFSSKNKESGLAHSSEKQLDKS
CTTENSEDDKKRGRSTDSEVSQSPAKNGSKSVHNNHPPPSPAVTPSYQLGSSSSTSSSLGNGPGSSSNGAV
IDTSTMVRRRVVSDSSEDRDTEEALKEFDFLVGSDDGDGESRSSGDGTDWEKEDQCLMPEAWNVDQG
FAESHVKMNLNLKPANKEPHSNSDSLVTTESNSIIYLEDLGSFVGSHTKNNLDLMPVNKEPDSNSDLCCKR
IKEHRYQFLEEAFQNQKGAIENLLAKLLEKKNYVNFAANQVQNRIKEVNETNKRVEQEIKVAIFTLINEINKKG
QEKSLPSGLPMSSEVANRTSSQLEASKKDLRVTKQGKVPENESPLTAIESSGVIDKAPPNSASRLRKSAAASSM
DTLITKLMEKTKYIKYTGKQIQNRILEANQNQKQVEQDIKVAIFTLMVEINKKGKALLHQLESLSKDHRIKLLQC
MSISEASIPGEEVGRLKAKDPDIGENGLVTYSIIDGDGVEMFEITTDYETQEGVVKLKKLLDYETKKSYSLKVEA
HTLLLRRVELFSLSRQVARESTYLSSLKGARSHPDESGAVQPKKLKQEVGEQSRPEPLQQPGGSESYAPPYRA
DFLAFSQEMVAEQFTLMDAELFKKVVPYHCLGCIWSQRDKKGKEHLAPTIRATVSQFNSVTNCVIASCLGDF
TMQEVEASGRESWHVPVPPQPEDLSIVCFTSGTTGNPKGAMLTHRNVVADFSGFLKVTEKVIFPRQDDVLIS
SMKIKVGQMHYEKQRMEQKCQILKDELASWKEKLEQKEAEVRKLQEKLMCKMKGGGVEITDRDENCKKKI
STQEDMFSQSHTTASESDSTITRVEVGGSVACTPAGTLHLLHLSGQGSLGQESLSTYSSGLITPSPGAFGSHF
TRIIRFCSFDSELRSYMEMPLECIFTEKRRKRSIRNEIFNVLQAAYVSKPGAVLAHEMGLSVNDDILYGVFAQ
PSWEPNAQTKRYSGSMEYMISRISPVPPGAWQDGYPPPPMNPSPISSSTQGQRGINSVPLGRQPIIMQSSA
YAEIMEGLRPSWEEEAGEAGGRGEPQSQAQGALPDPTLLQYVEQLCEDESFICKVEAVIHPQFLAQLLSPEK
ASGMNYLHLHKIIHRDLKSPNVLVTHTDAVKISDFGTSKELSDKSTKMSFAGTVAWMAPEVIRNEPVSEKVDIY
QEEYLLPKSQTVASASGGSDQRLKDSVSSNTNKTNSSCSAFGSINAKTSNDIARELEEQFYTLVPFGRNELLS
YSRHRAQTEKHVVLCVSSLKIDLLMDFLNEFYAHPRLQDYYVVILCPTEMDAQVRRVLQIPMWSQRVIYLQG
APLSPMLYNSPSSMSPSVNKPCTVELPRLTDMAGTMNLNDGLTDNLIMDDLLDNITLPSPQQSPSGGLMQ

TIVDQSNVKLTCTLFNGNLHALPKIYKIGDIVRFHRIKIREYNKQMQGITGIGFASLTFDGTVGAPIVPRTSSKS
NILVLAKSGALEDLMQTLEAALDMGASQLVKRTEFSEQVLCTVREELEENSVFRARLEDICAKLQASGQVTKI
TTCCQLPTSQYGCCPLQNAVCCSDRLHCCPQGTTCDLRHSKCASTLGQMQPLTRLLDASEGAREVRCD
GHSVLLFCPSKNWCEKLADIIAREFYNLQAERFTKTSALSPVALDKEGIEEVMDQLRRSLSGLDTVLQRTLP
AVDWIGNNLYWTNDGHRKTITVARLEKASQSRKILLEGDMSHPRGIVVDPVNGWMYWTDWEEDEIDDS
ESYLRRVIFPLEKLLTSHKRLVMKDSAVNAICYGAKIMLPGVLRYEDGIEINQEIVVITTKGEAICMAIALMTTA
KQPPVAAPDSVEVAKQPRAAAPDSVEVAKQPRAAAPDSVEVAKQPPAAAPDSVEVAKQPRAVAPDSVEMSS
WEVERKRELSKDVFTVPVKFLVIGPEEKAAIRDACLAEAPHTVPGRSHLDRDLADCSVVSCPSDPGAMSRSL
RMGMKMLMMIGLLTPLIYYLKRHKWSVLKSRKIAYRPPK
GNPIDRLYSMQSSYFAS

SGVRGADGISLLKKRGVEVWLISEEECSKTISALKLDCEVKVNVTHKRQVIEDWRNKMSLSWKEVAYLGYEE

LERVSNIFGYKSIEDFMTSQLDYLVLEWLRMKDNGYSLSAFPYIVLNYSSLQEFYRSCYKVLVPHLVIRSQFE

AHKVSNKEKAIQEVTVLQERYEKLLGIAKEKQTLLETLLAQWQRQEKERSIFLTWLERCETAAGPTEQYVSADR
NFDRSNHDDDDIRNCAVLDLASLQWMPMQCESQLDWICKLPKGSKEWVKYQDAEYKFFEHHSTWVQ

VLEELQNNCAEEDPWGSLSVVSNAEESQSLVSDTLNTNVHSNSRSIESEKT
ELAVKNVNSPWCLLFNEEDAKVLEYLNDLKQYWKRGYGYDINSRSSCILFQDIFKHLDKAIAESKSSMPISS
KIQRNLNHLKEQLCLYPRSLRERIAGTTVDPNWAVDPYMEGLAQEVQERIEEFRRDTFLQIDDFTRTIDRET

CDQFLVYHQDNTLVTGDQIFLIIKEVINKRQPEMVPNSRTSSTEAVAGSAPLSQGSSGIMELYGSDIEPQPS
DFPDLPDPQAMFDIEYFRKDPRPFFKFAKEIYPGQFQPSLCHKFIALMDKEGKLLRNYTQNIDTLEQVAGI
TFVCPSKRRGERTGPKLTAKLKCRYEIEWVTEYACHRDYLESRNCSLTSEQHDVSIDLTPLTQQPSYVTPYV
NSIRMIHNQQVVPLEQAMVKVRNVISAVDAAQFLISNDASLVIIQETKKYMDMIIGYFEQYVEWVKESPISHV
RQCSSMGSVKADMKAVIILVPVRLGGERTNMDYLEFVKGILSLEYCVGVIGGKPKQSYYFAGFQDDSLIYM
GRKYSTTGIIALNLALHICHEVNIIGFGYPDNGDNTMPVHYYDTKPLSQGVLVCSTLCVAIYAPFRFGFDSA
PKLQECNLDPCPASDGYKQIMPYDLYHPLPRWESTPWTACSSSCGGGIQSRTVSCVEEDIQGHISPVEEV

VLSFGRQDSAGSVLDGSRRSSSSSTTPTSANSLYKFLLDDQEYAIYADNVNRHENRRYIPFLPGTGRDIDT
FNEEVRKQSVSRNRDAPEGGFDAIIQATVCDEKIGWRNDASHLLVFTTDAKTHIALDGRLAGIVQPNDGQ
MRKKLILYFKRRNHARKQWEQKFCQRYDQLMEAWEKKVERIENNPRRRAKESKVREYYEKQFPEIRKQRE
AAQQES
RVCDKFDYVHNVLGKKKNEIQVLRDQIEMALTEDDDIAFLKNAAKLRHTSTKDVFIPRIDLDQTLIHTIYQAA
YISQTTDHDIALLRLEVPVNFSDYVVPICLPEKQFATSELSSIRFSTVSGWGRLIEGGATAAVLMRVDLPRVKT
RRLGQHVVGMAPLSVGSLEDVPGGDTETKMLAQPFLLDPSITLGQYVQPQGVSVLDFVRFECGEDTEIPEV
VLVAMKHTSLPIRHLWLSFDEGRSWSKYSFTSIPLFVDGVLGEPGEETLIMTVFGHFSHRSEWQLVKVDYKS
SVTWTTDQSSRPCVYTMGSPINNRALLPCQEPPVAMSTWQTTVRTASCFVVLMSGENSAEPVQLQEGLLS
YLLCDKVQKDDIQIRFYEEDENGGIWEGFGDFSPTDVHRQFAIVFKTPKYRDVNITKPASVFVQLRRKSDL
FYSGAVVSHHETALGLPRDHHMEDPSWITRIHERSQQMERYLSTTPETTHCRKQPRPVRIQTLVGNIHIKQ
EEFRTWLREEWGRTLEDIFHEHMQELILMKFIYTSQYDNCLTYRRIYLPPSNPDDLLNPGLFKGTYGSHGL
QRREEARQRMATPRPEDQEELTTLQREALGLGKDDQERVPEGSASPPQLPGEVLETIGPTGDTAQAKELDL

GQSSAGEFIFLLDRSGSMECPMDGRNHSPQRIDSAKETLVLLLKSLPLGCYFNIYSFGSQFESFYPQSVAYT

TRRKMPSLGWLRKCYQQIDRRLRKNLKSLIIVHPSWFIRTLLAITKPFISSKFSQKIRYVFTLAELAELIPMEYV
SLLLLLEQERRFLWFYRSYCNNENCSLSLVLGSAPSWDRKSISEMYKVLREWSFSNPLEALGLLTFSFASVS
FQKPESVTVHPSSCGQGPSKSGTVCQLSCPRGFTLSGVREELRCTPSGRWSENIQKALCKDFEAPQISCP
APRAATIILREDNCHFLRVDKRDFNRILKDVEANTMRLKEHGKVVLVLEKNLQGSSSSNQSPSAGSSRYSV
YNLVQKAFKPSPVKLYRETNEPVKTKTRTFNRETGSLLLPSDTKKAQTYGNRQIVLEKEEVEEVKRFDSPGLV
LKSLSEDYSDKWITISGCQNITTTQCDFSSDIAISGIYFLRVQAMNGYNKSRWSNEIKVDPCVVNEIGPPSV
QLIFAQSRGDLTGENVNQCVLEMMIAAPDTLSVTLFFMLVLIAEHPKVEEDMMKEIQAVIGDRDIQSNDMS

YYHALDSDHLTKTHHDMFTKGQVTTGQYRMLAKHGGYVWVETQATVIYSTKNSQPQCIVCVNYVLSEIVQ
VEVHNICSFADQLLNEHGIVPGSVLYHKSLQTLVVRCKEGWVGVKTVILKKKLTAADFYNGYLHPWFQQNS

SFEAGSLEIAVISQYCNLVSFVLFYLSAAINPILYNIMSKKYRVAACSLFGLKALPKNRFSVTKQEHSCAWTES
LQRYEEILATASTTDYNNNTEGREEDQRIINLVGESMRLLGNTFVALSDLRCNLSCSAPRHLHVVRPMSHYA
KHKNQKMSTVFLHCVTKLCRTDDCPFLAPICSNRERRDAVRRTTWSAQSTSGNAVISAGPIITRSDETPTN
QWEKLKTPEKELDAGYGEILLSISYLPAANRLLVVLIKAKNLHSKQLKDLLGKDVSVKVTLKHQSLKLKKKQTK
HYEDLNTPVERGEAESIGQMVQEAVQRFLPGASVTLAGGFRRGKTRGHDVDLLLMHPEAGREVGLLSRV
KY
LALAARLPPCRSHLRRYFCATLQLPSDWIQVVALFQSLAGAGRPLAPLPRCLRAGLADKFQQFDAYQLAKY

EMEQAKESFKNALTGILTQFEQIVAVFNASTRQKAWDHFSKAQRKNIDIWRKHSEELRNAHSEQLMGIRR

GEAIKKHIGLLHIKGKKMNMEKIPLETVRSFYIEDIVLADHPDIFNPDNPKVTQAVQAFCIEKVETLLDNAERE
HGPSANSEVEVKNVVNFIKSHGKIKAFLTLHSYSQLLLFPFGYKCTNPADYNELDTVGKAAASALASLYGTTY
CNEKEPADSTASQSTGMNENVPSSVQSSSSTNAVSCNNSSAIPQPSSAIKPWRSKSLSAKHTATSSMLSVK
STIEAGMFADLVRLQELHMVGAQLRTIEPHAFQGLRFLRVLNVSQNLLETLEENVFHSPQTLEILCINNNPLA
CSMQYDAQELEQLNRMVHRVGDEMSSLLSPPSVSQSPAHRPGIRNGSSTEASPNRQHLDNLIDEEERV
EGMSAFVEKRKANFTDH
IKASFKGDSGGPLVCGETAQGIVSWGPVNGSPPRVYVRLSTFIPWIRKMMKKLQP
RALGDLWSYDFYSGEFVVSPEPDTSVYTLDPQKHKYIILGSDGLWNMIPPQDAISMCQDHEEKQYFMGE
NEDAMFVFVSDVWLDQVEVLEKLHTMFSGYSSVPPTCFFFCGNFSSAPYGKNQIQSLKDSLKALADIICEY
DLYVKDKVDETGNALKVIHEEANIRWEVCLTLSEKGFQQVSFVNSIATTKGGRHVDYVADQIVNKLIEVVKK
TLKQGCKALLLRHQPKLRKTNVQSKCLKSKISEEPTDFSKANQTHFKMTATSQPWVVELNKEGEGKRHFCK
GSLANIAQASKEDLSLCPGIGPQKAKRLFDTLHEPFLKIPK
INECPDPGVPINARRFGDNFQLGSSISVICEEGFIKTQGTETISCILMDEKVMWSGPIPRCGAPCGGHFSS

QLQQQIDKNVADMKEKLAPLADELQVKIDQNMEELRKQLAPYAQDVQDKLNHQLEGLSFQMKKGAEHLR
TLKLVETPKINSRNFDEDKLEKKKLSLESADTKSLRHGDAVLKVLQELQKLSASQQNQQRASLFYKFVTGLRSL
GRGEGFNINVPWNAVGMRDGDYLAAFLHVLLPVAFEFQPQLVLVAAGFDSVLGDPKGEMAASAACFAHL
KFSKLYSTNDMRSEEIVETTLPVLRIEHCDQSINLGKEEKFFPAINARCLWIGSSTGLFIINLANFELEAVLHY
PPNFLRASALSEHISPVVVIPAEASSPDSEPITDLVETDTASQQSLFGNKFDDIFGSSFSNDPFNFSSQNGM
YIAGTNNMDGHIFAGIDMPAINRPLVKITADEVYNLVRGLTLQKGVRAANATFSLYTQLWNDNPDQEVMK
RTENTSLLGSSVFGKDDIYKIWRKFVSEVLESNDKIPRFYYVKADVTGAYDAIPHDKLVEVVSQILDPEKRTV
AGNDKKQTFLQVFVQPQIIHLKNETTFENGQTTLICEAEGEPVPEITWKRAIDGITFSEGDKSPDGRIEVKGQ

LNLREFFLTPNCATEMPHFQ
KAQYPDAKLVVGNTEVGIEMRLKNMLYPVIIAPAWIPEMTFVQHTKEGITFGAACTLSCVEVVLKKAVAELPSY

SETGVGAPAFGPGSLGPFTREAGFWAYYEICTFQKGATTKRIKEQKVPYSFKENQWVGYDDQQSIKTKVQY
MSSSAANTVREELTESRMRIESLSSQLSSLQKESRAWLDKMQELEDTLANERDNYRRMLSDKEREMAEIRD
PEEPERNGVAPGAQILAIKIGDTRLSTMETGTGLIRAMIEAMKYKCDLVNYSYGEATHWPNSGRICEVINEAV
IETTKLPLKFPDAETDQIMMISYMIDGQGYLITNREIVSEDIEDFEFTPKPEYEGPFCVFNEPDEAHLIQRWFE
EGLNLDKLGLKLDAKGHIIVDEFQNTSREGIYAVGDVCGRALLTPVAIAAGRKLAHRLFEGKQDSKLDYNNI

DKPPKALITEGDSKVVKLRCFPLAIAKTQLGPDKVGMKYVPVKKGSCSQAPELSDTSDEHTFIEGAPCKNDS
ICKGKVAEVSKNFGSKERSLPPRPSLPPKPKTLNSGVANGRPHRNPGVQRTISTSTQESIIRWFKEEQFPLQA
PTSPTYLGPLPSRGSYQTSHQFVISSTELAHNEDQRTSSKGFLSTAVNGLRLSKTICSPGETRDIRPFYLTNAL
EPGMKGSKGDMGSRAESYSDSKIRLIGTAKGGRVEILYKGKWGTICDDKWDKQDGTVVCKMLGYSHAV
NKVPWNRLFPNADPKALDLLDKMLTFNPHKRIEVEQALAHPYLEQYYDPSDEPVAEAPFKFDMELDDLPKE
EWTSLPSCTDATCSAPPKIPNANIINEKEGRYLPGDKVQYKCQEGFENMESNYVVCKDGDWSQPPICKEA
TEKSLPPHGSPMFTGSAYIAVTRDFVQALFENPRAQQLLEWSKDTYSPDEHVWATLHRLPGMPGSVPYND
IAKHLPDGCPQMVVMPLYASLPYTQQLRVFQAAPKGYRKVILSTNIAETSITIAGIKYIVDTGMVKAKKFSPES
SLNGRQTSYSPHHSPTPSPQSSPRVSVTDDTWLGNTNQYTSSAIVAAINALTTDSTIDMNDGIPIKSRKTAID
ANSPEDMHSWIKAISGAIVAQRGSGRSAASEHSSSSSESSYAFYFANIAATSHSTASHNNALVSNLNAKRLA
ASPFEEDKNRDLKQGDDEDVTKINGRGLPNGMDADCKDFNRTPGSRQASPTEVAERLGPNPNSTEGFG
ALGSFLVDSALIAIIAIIFIIGRKRVGNGLNFGEYPNCRLAAEQDEGHSRDATPDGELKNDIIYATLRLQPKPKL

NELPKNHPKSSIIIEELSLVSSPTLSPRQSIISTQNQYSTLSKPAALTGTLEVRLMGCQDILENVPGRSKASSITLP

LLDQQDVAIKRFPAHLGTHWVADAVSSLRKYLVDSDDSVPISRIKRIVVKVPDVRYVNRPFPNSEHEARHSF
KQVIAVKIKETKVMKLRKLAQQVANCRQCLERSTVLINQAEHILKENDHARFLQTARNVAERVAMATASSQVL
GETVEPYEVVISPDLENQLNSIIQELSLEVMPQEDPSYPVSLNIGKLAHFEPSVRQNQMGVVPWSPPQSNV
AAESDPLIVGLAYVNSCLNPVIYVIMGQDFKDRFQRSLKTILRNVLNEDSLSMGDNRKKTRSTQETKSTMED
PDRYLFSTVTLNITEVKQEDYGHEFVCHAGEVAAYIALRQPVRNFQGYLIGGLIAPVFVIIVAILIYKYFKVDIVLV
PLLQATTQTTGVDINILRRQQRMIKNRESACQSRKKKKEYMLGLESRLKAALLENEQLKRENGSLKRQLEELV
HAEIHEWYRKPDYNFFVNYKAYRTRHPEQPFYILNPKMQWQLWDILQENSLEEIQPNPPSSGMLGIVIMMT
DGIAEVFWIGLNQLDISGGWQWSDHTPLNFLNWNPDMQNVSPLDGSSCVAMNAASGQWQSYHCGT
IIEMDSKRVPRDKLACITQCSKHIFSAIKITKNEPASADDFLPTLIYIVLKGNPPRLQSNIQYITRFCNPSRLMT
GKESLERQERPVAVVFARAPPPAGTPTPVLLVCRVTGFYPQPVRVAWLQDGEEVTPGWWLNSSGILPNADL
KNIGFSKAKDVNTTLRALEKLGEVHKENTNPPKWSLTEKKRERMQIKIKANEVTEMPPPAPEPEFPAACVEPD
PHLITAEEIKNQIEAAGFTAIFKKQPKPIKLGVIDVQRLKNTQTKSSEGTLQKSPKHINDLKTIVFRVDGMHCN
QQICERYPIFMEKVFGMAEGHDPTMIGVAVDTLAILGSNVEGKQVLQKAGSRFQQLLNRIGHQAKNPPAEL
QVLHLNERMQEAKKLLSSVQVMFDKTEDINFMKNTKSVKILMDRTQNCTGGSLPPPKIGHLNSKLFLNEIAK
GNLLANAANPVEFMCQAFKIVPYNIEALDKLLTESLKNNIPASGLHLFGINQLEEEDMTTNQRDEELPTLLHF

RKASAVNMEKFKKFAARRKWKQSVRLISLCQRLSRSFLSRSNMSVARSDDTLDEEDSFVMKAIIHAINDDN
DCGPPLSDLLRGMLEYDPAKRFSIQQIRQHNWFRKKHTQAEALVPIPPSPETKDRWRSMTVVPYLEDLHG
PQNYQFSSSQTQDLPQKYTQDSLLEYRPFSGNDQSPTYQQNVFESHELPYCPGQSFASLLNESEHSENV

EDNYRPVQLLQCASDPDHADCRLCTNSVQ
VIHRTCTKTITKKIITGPGGPREEIVETMTSPDGSDCSHFQGLRKEEIAGQGGTLHVRVTSQGERGDLSEFD

DFGTQSYEQRVVELGKEGAQSFCRKKRTCAMHLRKYNDALLINDTVRMIDAFNALDEFYQLEKATKLLQD
SVPWLVTTETLATESRAASQHPTAGTATTDWSPSSSLSSATENPTTGDVASTENPLTKKATSVQSPRTESVCSL
LESYPSHFQTAKSSSVLVESTGFCSDDMGDDDVFEDSASVRLKSKELRAPLCSVEKDSDLDCPSPLSEKFP
TLPSRWICKQAHGPAGL
SLAIELLDKVNSPDLAAPSLDDGDFEVYDIFPDKIHSCGAHSQAERTLSEIQFNQVKFGPPLRKETEPYCGL
HPGCVHGTCIEPWQCLCETNWGGQLCDKDLNYCGTHPPCLNGGTCSNTGPDKYQCSCPEGYSGQN
LELADKLSARILWIGIANFILCPLILIWQILYAFFSYTEILKREPGSLGARCWSLYGRCYLRHFNELDHELHSR
GQYCSQVEAAISCLDNISKTKEDVKLKLEECSKRANNGKFTLRDLLVVPMQRVLKYHLLLQELVKHTIDPME
WQEKEFLEKQLKQQTKDLPVSSEKCCFLPQMFLPPCVNYENCGLVVHYQDPRVQMSSRGMYDQQQH
SQDESAAFSREDRLEQAQLFCRTLGEILQHSNECAGCYRLIVYEESGDSDSHFLSKEILRHIRQQHLEEY
SLTFGRTTLVFQFLLNASTDKFFLQGVNVTTTLPSEAKETKIDVFNHSMSELRAKLGNSYKCNAEEKLWVTD
IDYKRNRNPRELLKEVKHMASLTHTFNAINYVANQVFTPERGAREDVARVMIIITDGDPSDPGGNVNAAKE
DGKKGFKLSKDRKHCEAVPVCVPLNLERNYELLYLAEQFIAIPVLYLRFRLPEVTRFSAEFDFRTYDTEGVIL
DSITNIFGLSESSLLGHHSSDAARRRSLSSFPAPQENVDVHPGSSKEVRVPTTAVYVFDAHDGEVNAVQFS

TLEDMNSQLSSFSGQMENITTVAQANEQNLKDLQEHHKDYANRTSAKFNQLEERFQLFETDIVNIISNISY
PTGTELGLAVRQAGNQSQETVTLQQSKTQMELKWAGAPGHKNEGFTLAGLLTYQGMSPILDGAERVKAPV
APPHGRINPVQAKYILKDYYTLSCDTGYVLLENELVLKSFKAECQKSGSWNKPMAKCIIVNCGQPEDIDN
QLTCFQPGHFPFNVIHHSFVLPQPGVGAVFYGVFTSQWQMGDAGSTAVCAFNLEDIERVFSGKYKELNK
MFSPKCRACERPVMENYLSALQGVWHPECFVCGDCFSSFTTGSFFELDGRPYCELHFHQRQGSLCHG
RILASEIRSDSHKRHFLIPASYKGKFKRRPREFLTAYDLEIARSDKEQLHVVATKAFDSPHAELFSVSVGDQFL

RDARILPEVPGSDHCPVLAELAATFVPAPCCPSLCTRHMPEFAGTQQKLSRFLVRVERGVGAVGGHGMCR
GPHVKRLLREPCPVILDPADPTGTLGQEKRWDLLAREAAKCRDQLCCRNGLAPIRCWDVQPARPMQVM
AKTIQQTLIVANILEACEHMTPEQRTQAFAKLSER
ETSSPCSVTCGLGRITQKRLCNNPAPRHGGKACLGLNTRSHFCNTKIPCPVDGQWSEWGEWSTCSRPG
LSFKNMCKLKPLLEKWLNDAETMSVDSTLPSPNQLSSPSLGFDGLPGRRRKKRTSIETNVRFALEKSFLANQ
EDEVLIPPFEKFKVTNFTRAQDRTFIQLLSLDSYSNYNCEYVKEKRCKARRCVFSSGMNSFSSSIQCLYLLG
NKESKSEPLLQNTTRFHVEELEGKSFVHVATLREVTERDLNSNFICFAQNSVGNSTGVLKLKRKERALFIFILY
VKVLKRQQRMIKNRESACQSRRKKKEYLQGLECRLREALAENERLRRENALLRRRLDCALGESRELKFSPG
HRDTRVMCQEPFALQLVNGPSKCSGRLEVRYDGVWGTVCDDDWSDTNAQVVCQELGCGPAEPLAPKL
CDYNGGSIHLPKHSARHHLLGISLFAEHFYYPDPKTDRIRRANKFTGKVIVTISLKPPFLPPAEIKVVHPFKQ
GWERRVDNMGRIYYVDHFTRTTTWQRPTLESVRNYEQWQLQRSQLQGAMQQFNQRFIYGNQDVTAA
IKAEYLNTSRFREDHQFFMRRDREYYGESILTIEEVTEEDLLSNFTCVALNEIENVVLTVTLQLKVPCKGDNH
CTIYLPHLHAQQALELRITEPPVVTLRPDPLSVAPGTPAELVCEISGYYPLDVTVSWLRRGPGETGGDLLEYVT
DGVAEVDFNSKNAAKEIGYQSLEELDGSYLYIAVTVLESTGGLSEESEFAGVRYALSPYKLNLVATPLFVKPGL
EKVSRGETVEGKYLTYYLAQEKLSMKSIYGNVTELLLAGVDTISSTLSWSLYELSRHPQVQAALHEEITGVMK
IKASFKGDSGGPLVCGETAQGIVSWGPVNGSPPRVYVRLSTFIPWIRKMMKKLQP
NQLMYLPDSIGGLISVEELDCSFNEIEALPSSIGQLTNIRTFASDHNFLTQLPSEIGNWKNVTVLFLHSNKLE
GSDEEFKPDAKDEASSDEASSGMGEGESASSETDTKMDVDSPVKVPSKRKRETDRKPAKRNSLENDLSET

LTVAVKTLKEDTMEVEEFLKEAAVMKEIKHPNLVQLLGVCTREPPFYIITEFMTYGNLLDYLRECNRQEVNAVV
RLMLRKAVKMEGERFLLEFYVPPKASKPKVSIPLSAIIEVRTTMPLEMPDKDNTFVLKVENGAEYILETIDSLQKH

SETDVDNFGHIALYKAVQFIKKLNPGLKTSKIELKFKELHRSKDKTGTEVTKEEFIEVFHELCTRPEIYFLLVQFSS
SCCNKSIEVRLIFLAGFCIATAVVWAVFRNEDRWAWILQDILGVAFCLNFIKTLKMPNFKSCVILLGLLLLYDV

MVTSRPEQGQILLQHPDAPSPAQPQVSPAQPTPSTGGRRRRTMDEDPDERRQRFLERNRAAASRCRQKR
EIDPKIKAHYSCHVEHESLLEPLSVPWEPSNSLIPIVAGVITAVLLIGVIIGVVVWKKKHPGRKGDGYTVTQGND
CGTSVIVPELEGALYLKEDGKKSWKRRYFLLRASGIYYVPKGKTKI
ADGWYKLLNQEEGEYYNVPVADAENCGLLQKFEACNYPLELYQKVRHGPGPSPTPSPTDSKHGSFFGTSR
APMMGGVIGAFTYIIFIEIHHSDTHLEEDTDMYEKYELTNMEKAENLRSAPCLDLKQNDYIGHSANTVCLS
RNQQGCSGGRIDRAWWYLRRRGVVTDSCYPFTSREKDSASAGRPCMMQSRSTGRGKRQATQRCPNP

FENGAWMVNTSSLVTIKAGRPPTILVFHSFYKGRLRFSDESYSLQITDLRLEDTGSYAVEFNTDKEQFAYREF
AYNLSTVEDVFSKGKYMQSATVEQSHTKWVRYNGEIPKPRPGACINKEARAANYMSSLNLPDKTLQFVKDH
PEGGFDAIMQAAVCEEQIGWRNVTRLLVFTSDDVFHTAGDGKLGGIYLPNDNQCHLDTDGLYSKSHIYDT
GISDCVPINMNKIVKTKPPKIVSVENIASVKQLLNVTWDKPDPTLLDLNCRLQYRNTSSNTSVTVNFTVNQNE
NVTDSPPAPQIKAPSELRESVPAQAVCSIAYYCPDYPITLTWVGLGHRTPEPTTRMDSGRTENTLTFTPTWQD

LPGSLEIPSQAELFSEFISPVPFTRPYASCSYCDFASVPPEQVFSAGHTEHDVGSGDYIETLSFTASGEKGITP
SEFNSYYGNSVPASSATASRAGPCFDQHGGQQSPGMGLMHPTAAPNSMDPLQNSHEGYPNSQYNHYP
VNLRDAKHKEAVTILSQQRGEIEFEVVYVAPEVDSDDENVEYEDESGHRYRLYLDELEEGGNSGSNRKHAS
QRMYFVMNEKVWVCNLENCTVAEEIASPSITSPKKIVADPYNGYVFYLLEDGIYRASLPVYPGQVPEALPIVRL
INPDEELNLILPQTPVRAAMNTIQQLMMILNSASDKPSDNLLSYFNNCTVNPKQGILARVEDLGNIFKEKFAE
PTPHDHIKVTQEQYELYCEMGSTFQLCKICAENDKDVKIEPCGHLMCTSCLTAWQESDGQGCPFCRCEI
DPSSLGENPDGLSQEQLEHRERSLQTLRDIQRMLFPDEKEFTGGQSGGPQQNSGVLDGPQKKSEGPIQA
EKGFINVTDSREDFEIDLLEKFCFEIKLKAYPPIRCIWVFSQKSFPCEQRYSADGYSISSRFCDHKYQSGEYV
REPGGPQSPGSGKDSRMRANAALHNIIFSQPDEGQAKKEMRVLHVLEQIRSYSETCWDWLQMQLQSKD
LSKNNTFNAEKYSQTIRMDLSSSFVSDHSRLKRTSNNQMIVPQDPQRHVEKKTVSVEDEEGITQSDTILIDS
PNPPPENVQLVNYVSKNVISSEHVIERETEMSFSTSHYTSTTHHSTTVTQTPSHSWSNGQTESIISESHSVLVS
DENIKIGGLPAKGPPPPWLLDMKYLSPLLLSYEDRMREKEDLHLAHEEDMKKFRIRVEELVKENEQLHQQLN
FKYFVQGMGYMPSASMTRLMRSRTASGSSVTSLEGQRSRSHTGEGTRSRSHTGDLMVRDGYCWSPEGA
ASGPQGPTQTQCNDAYRNFNLSVIVGAEGILQGVQIWRVPATNTYSISGYGAAGGKGGKNTMVRSHGVS
EEDTFGVSIAVGLAAFACVLLVVLFIMINKYGRRSKFGMKGPVAVISGEEDSASPLHHINHGITTPSSLDAGPD
RGAGEEKPPAAAPQLLSPPKSPAFARLPEQASPLLSPEAGGSRSSSVVSVEKCSTPGSARSRTPLTAAQQKYKK
NRKVALTRLDSVVSRYAQLQDESEEEERVRKRAAQARGSAAPRQEGPLPPPCPMAPDPGEGQPHSQVSTS
HAHHKQTSPSSHQMHVIARGPAFANDFSDSPPQNLIAPSRNSLNMDLYDMNNPQVQQWQTAAPSHRDS
VNGWTGEDCSENIDDCAMAACFQGATCHDRVASFYCECPHGRTGLLCHLNDACISNPCNEGSNCDT
KVSEQLKYCSGIIKEMFAKKHAAYAWPFYKPVDVEALGLHDYCDIIKHPMDLSTIKSKLENREYRDAQEFAAD
SKNMEAVTKGTPRKGTPKEMEKKTNQTKAIEAPKTDYAAAVSGNNKKDKKQQKNQEANANRIKPSQVNLGE
AAAPGGAAFSKRYLEEQKTENGKDKEQKLTNVEKEKMKEKGGFSELGLADGKTKSDLYTSKTDPEKPYRSSQ
DEKHSSHSKPERDSRYMSSRSRSDRDRRRSRSHSRSRSDRGSRTSSSYSRSERSHYYESDRRYHRSSPYR
LSKISKASDHKKNFVRSPSTRSTLVKSKKIDVTHITSKVNTGLAALKKDTSSESTFLATEEGMKIMPGLNNASPS
APLTVSVANQGHPDRLSASWGAPAGGREGYVLTLYHAGSGAVAARAIAREDTHNFTFAELAPGYRHSLEVV
LSLIGFAEGVSPGELFGPAGHCWAHHWCAAWSAGVQPQAGAGLSDVDKAVFSGISQKCEHCQRMGATI
VPQRFMAKWEVSVGHAEEFLLKETEDQKCSQNSSKLNSEDVLLESCTNGPVMEPDRQLNGCFKSLTFDSK
DSVAQGYGSLGMMTSVLICPDGKTVEAEAAHGTVTRHYRLHQKGQETSTNPIASIFAWTRGLAHRAKLDNN
PMPPSSTTPPQPQAKSKSCEFCGKTFKFQSNLIVHRRSHTGEKPYKCQLCDHACSQASKLKRHMKTHMH
TIAKQLLQRAKKGAQKKIEKEAAKLQGRKGRTQLKNIRQFIMPVVSAISSRIIKTPKRFIEDEDYDPPIKIARLEST
SVFENLEQLLQQNKKKIRVESEKSKIYDHTLKFSPKRNISNCNGEEKKLDDSYSGSLTYSYKALMVPKDMLKK
LLKADSCTNCHGSQAKSVEIENLGNLKCLNEHHDVKKSPTVSTNVWLVQHPKTPFKVEDVCKANEPCTSFS
YGLGVDFLKGVRAVNSSNIQHFAHCTKIFGSLAFLPETFTGDPATNTAPLKPEQLKSFGSLEELTGFLYIESWP
SKKTGNGVQHTEAVETCQSESNSLHEREPALESLWQPKTFHSNPLSQVSGKRLKRSIQKVNEWFSKSNEML
VSNSSGGAGAKAVPATGHTPSTPTPYAQAVAPPTPGANASQPRPPSAQQNASKQNGATSYSSVVADSTSDSE

VITKLKEQYKKERKGKKGVKRKTTEDMFQPQSYIKQSKQGCGKMMAQSTGPNRSKLQDMLANLRDVDDL
DDSESDTQNVVTVCEPVNLAVQQPVNYIGKETLVLDSGYVPISSSNIEKSIVALVSTRTAVFKENTGHSSEGS
KSLLQQLENVTKERQMKLIQQQSDITGLSRQVKHVMNFTNWAIASGSSTALLYSKRLITFQLRHILKARCDP
ISAPNNLSASSKLPEVQHPKLAAKRRWTCSKPKNMLQETSMATSEKLMIEPPSAYPITPSSPLYTNTDSLTVITP
QQQEVASLSKQLEHVMNFSKWAVSSGSSTALLYSKRLITYRLRYLLRARCDASPVTNTTVQFHCDPSFWAQ
ANVHIDPKFISNGPFKDTVTVKVAVEDADEPPVFLAPSYILEVQENAAAGAVVGRVHAKDPDAANSAIRYSID
SLEEDAASLSGESLDGHLQAVGSCPRLTPPPSATPDLPLSLPPHGLWSRHILQQTLMDEGLRLARLVSHER
RSLKPQQRAKVVERWIEVARECRILKNFSSLRAILSALQCNAVHRLKKTWDEVSRESFRTFHELSEIFSDENN
FLPLAHMFERVIQSVVYCHGGRIGFFQGDIRLLSDDMKALRPTIFPVVPRLLNRMYDKIFSQADTPLKRWL
LKDKNIEVQKMKKAVESLMAANEEKDRKIEELRQSLNRYKKVQDMVILAQGKKGKEGECEEFLNSGSVSAV
LIIASSPTEQEEAKDEQMNISAPAEGGDLLQEKQTGDEPVEVGNPPVPPGPHVLPQVSTPVSQNAPAFVPG
SKPDSSEPTNRSAVCAVSVRTINEFFNKIVDKQNMKCLQHFYGKDSKYCLKRTFSRNASYCEAQDDEYRL
ANSKFSFPSGRAGMQNGNCQTDFMVHQNVISGNSVNRQPPPPYPISPTNRQSPTALQMQAGGSVPPSA
KRCDPLDLSEELEQELGLTPSQLVEKRLLALSEEELSLPACPAPHLDSTPSQSEEEDEGSSQKGPASHRQAM
WSFGVVLWELLTGEIPYKDVDSSAIIWGVGSNSLHLPVPSTCPDGFKILMKQTWQSKPRNRPSFRQTLMH
PLKDSDEVKSLWVKIDLTLLSRIPECLPEEPLAMSTGAKEEASLQQNNYTYPPAEKTVPKSRRKRKCENEED
SVLKDQDLLRAKMDDAEACFILSSRCEVDRTAADHQTILRAWAVKDFAPKCPLYVQILKPENKFHIKFADH
RSSSFPYGSKGSGLGSPSSNFNSAVFGPSSLNSLRQSPMQTIQENKQATFSSISHYNNQTLQDLLASESLS

SYSFTEEEQKTIEELRKAEVDGSSFLLKVWDGTKCPYPAWKVLVEESDLEADKVLIRQLRNLIVDILVYDNHV
LTLDDMLCQIPYGKKRHMQLLKQKQVSHRTFRPLKSALLAE
EQTSCPDGSTCCRLSTGAWACCPLVEAVCCPDHIHCCPKGYTCDPVEGSCQQALLSVPWLEKTPARVR
WGVAFHHAGLTFDERDIIEGAFRQGLIRVLAATSTLSSGVNLPAHRVIIRTPVFGGRLLDILTYKQMAGRAGR
VGRIEKAWMDGYSRQIFVTSKMLWPNGLTLDYHTNVLYWCDAYYDHIERIFLNGTDRKVVYNGKELNHP
ISTCDHGIVAKIKRVIMERDTYPRKWGLGPKASQKMMMIKQGLLSKHGKPNESTPDSWKKEYVDYSVAVK
KDSERQPFGTVTTYGISALREAFKTLSNSQDSDAAFTKLDETDFSFPKQLSPSVSHRLKRQREEENPHLETSE
LDVSLDNPWDKLQSVSVTLSSSLEEKSIIQPSLPSTQEAQLPNSTEEAAITQPDSNTPDFLEPSCQQSPCSLA

ESDVDCLREAGMSGVPADACAAAQKAAGYICKSNGGHGAIQEFAGYVCLLMKTVNL

EDVKSIANMIEKDWRQLLADCFPKILVNILPYFACESHRDSEITQQRDTAYKVYDMLKDESCLGKQQIDNLS

VKLEGELQSAQDLQADITSHASLYTSLLQLNESLFTAASEESVKAMKQEFKELDGRWKALPQTVSKRISFLQS
AQRICTWFQAELVSVHNQAELDFLGQSLKKGCGALHWLVHEATDWHQRKHLIGPAKLLHEEKGGGGGW

SPVILQFGHAETLQPLLALMGFFKDEEPLTANNYKKQMHRKFRSGRIVPYASNLIFVLYHCDHAKTPEEEYQ
EEMKFKLSPLSSYSEEFQDSLAPLEDLHVRLDSLWKDISQSLNEKSSDLQ

SVNFIENPQDLNGSIQAHVDVNIDLVSPDSGLATIRSSRSSKESSVFLSDDSPVAEGAHHSLLPGFDSYSPI
QRIIQCHGSFATASCLICKYKVDCEAVRGDIFNQVVPRCPRCPPDEPLAIMKPEIVFFGENLPEQFHRAMK
AKDEKEEYYYYLNICGKTSAGNCRDDTESVSSCQVKTRGQQEKVAGKFKNQTLRYSDGDLTLTYPDGDEC
VGLLGFLILIYKIFDSRVNSTGTSENFNVKDF

DPHYCQSFVDVSIKDFPLESFHCPSPKKMSFKNMDPSCTIGFYCRNMKDFERISEEITKKFQKVLTCPQILL
PSSPGMSLHLTLSSVNFLPFGPTSVQAKPLCTDVYPVYIKLKCVTLMFGREAVSFPTCFVQSLATLFLSQLMAS
VKCMYTPKMPIVQPCNIFDCPKWLAQEWSPCTVTCGQGLRYRVVLCIDHRGMHTGGCSPKTKPHIKEEC

GSIPTVDQLIEKFDRKIGPQRRGRSGKRNRIHPDDRKRSRSVDSVLSLGLEVNSDYLDEFSKNLGRSTEHLL
CHIDKDNLYSASTTLDYPSLGLMTEKLSQKNINLIFAVTQAVVGLYKNYSELIPGTTVGTLSSDSSNVLQLILDS
LQERMQRVGQRGSGLSMSAARSEHEVSEIIDGLSEQENLEKQMRQLAVIPPMLYDADQQRIKFINMNGL

FSLKEIVKQTVTQPQEKKMEAGGQSRPSVEEKATPQWKPKAFQINPSNKLSASGRKAVGSHAPKIGKAPQKP
QECIQKASLNITENMFCAGDLSGTKDSCKGDSGGPHVTKYKNTSFLTGVVSWGKGCATEGTYGVYTRVSK
EETQQRLAQEENSS
IFDRRCAEEDYRPWQLHSQGEACIMGAKRIYRKRKSEKKCMQGKYAGAMASEPCVCTEADFDCDYGYE
WYYYVTMPMPASTFTIAIGCWTEVKQESHIADIRTTNEFSSVFSRADLRLIEGSCGHGAYPCRFQNPTASSQ
ETSEPKPFLYYPEIKDKEEVQRKRQKLMPNFSDGYGGGSGAGGGGGGMYGGGGSGAGSGYGYPSYGYS
QEMEDDYDYYGQQRVQILERNESEECTEDTDQAEGTESEPKGESFDSGVSSSIGTEPDSMEQHFMANLG
EIVMLSFHGKKAKGTKITGDPNIPAGQQTVEIDLAHPIQLPDIENLRNFNELSRIVLEVQEQVRREQQEEEEG
LSEETEFESVDKSSLFTDASFPPPAGTLLTLPQPGEGAEAIAGAEIRQRHICSSS

TQQTMAKSLQHVQQLQADLGGTEILAPLQAIYRSPCRDGHPRQLFVFTDGEVGNTQEVITEVQRHRGAHR

GIPECIKQYEEEKFRKKHKRIDQELDGKQEQKSEQREVVQLLRTVRAGEARSRYRPGRDLSPAGLQ
E
DNIEARTLEHHNSANISWQVPIAKDNSGDEVSIHVTPAFVPPSLFPIGEVVITYIAVDKSNNRASCTFSVKVID
MAGTPDKILEHLLEAMRLDATFSDPVDTLVGDFLLTYNVFMPTTQLCRALLHHFHAEPSEGSEQEKAVYSLSK
LIGFKPLVILKQHHYIKPSQFIYPEESFISGSTTLFNALLTKCLDKEVMALCRYTPCRNTPPRFVALVPQEEELDE
TVSSSKDSLRIQLTPPGESENKSMSEFYDLSYRVLYWKNSSDTKEKIQEEQQLLFTISDLTPLVLYCLKVQAFS
NLKVVENFINESMRYQPVVDLVMRKALQDDVIDGYPVKRGTNIILNIGRMHKLEFFPKPNEFTLENFEKNVF

KELFLSLAQTECVLKPVESPDMKIFSKADLEDTNSLFEKLKKEPDALTLLAPAPGDTIISLDFSSNESDEQQYDE
SKMLHGDCRFHTLKLTTVKKTERKGNANAQCKTKSA

NVIT
APMVLQQAAIPIQINVGTTVTMTGNGARPSQPGSEGPAPTQPSPPVLAGAPSPGEPGRGPESQLGPGAAPT
NSQLAHLNGSPFQLNSVTSALISGIVILGIMSIFFFVCSLTILRRKWPSSSVLGGVRNPVFN
CAKHKINPVWNEMIMFEVPHELLRASSVELEMLSQDGLGQSQSLGKCSLGLHATGTERNHWEEMLRNPR
VSWMDSQGLILYQHSQENSFTLHEGPEAPGSSDVLDRFERCFSIFRLEVRGSPGAARGWKAVRVDLVVAP

NSRKSRGKARGRPRPRQPRRELRSPMLGPQRCQPILPPGAAPQRPLPWTRRFRLRQAFGKLQTKFESGQF

EEEMEMSDDDSSDNPSKKLRVDESALTAQAYALKEENDSLRWQLDAYRNEVELLKQEKGQLFRTEESLTKD

RLGNPRQPEKPLIRLRVDYSGGFEPFSIHRFSQKFVERTANPKDIIHFFRHREQKEKNEDEISFGKLFRRPAS
KVGTICSTIYQASGGSIDWSYDYGIKYSFAFELRDTGRYGFLLPASQIIPTAEETWLGLLKIMEHVRDNQS
QPIPESPKPPSEMVKASPNGQKSMLEKLKLFNSKGGSKAGGTSLECSGSRDTSCERLEALPSFEESEEIDAAN
CDCRLLWLLQRQTTLHFGSQQPMCASPDSVREKLFKDFHSTALSFYFTCKKPKIRDKKLQYLVVEEGQTVQ

FMDDLDGAGEALAGLDSANDTFAWVNNPCHNSKQSVQHREPLVYDNGHQTEETLLLREIESDLSNNNN

IRQSCAKMLVNRALGRWRQRMLRADNTSAIVICISPVLDSRSNWENEEELYLNLAEGPCYSSPDMCMMT
PSIHKSSRFVFVPGPEDPGPGSVLPRPPLAENITSEFRQQVPFSVFTTNPCRVQYCTQEIIIFREDLVNKMCRI
KNKGGVGVKPFQVKNHMWVFVNSLIENPTFDSQTKENMTLQAKNFGSTCKLSEKFIKGAVSCGIVESILNW
LTEGEQIIESTHLSDSSSDDGIEEAIQLYQLEKVRKEVNPTTDFQKGVADISASLTVSSTKCASAEDHKKSLSTKI

PSGVILSPGWPGYYKDSLNCEWVIEAEPGHSIKITFERFQTELNYDVLEVHDGPNLLSPLLGSYNGTQVPQI

AKLSESAEELRLKLNPYAKELKEKLRTDAEGLRQSLGPYVDGLSGQMEQKIEEFRQTVGPYGEAFNKQLVQK
LHNDTLRSLVPKMMETSSSITVQGLIQCGTPDCMGSILQIIRTGNVDPLVADAVTYSLGLLPFPCTKRIREILNI
LTHLLMALAGGKVLLSLEGGYNLSSLAEGVCAVLRTLLGDPCPQLEPPVAPCRSALSSVSRTLAIHEKYWRSLI
KDYSGLTIQFAGSCALMRKAVNGKVFCLLTSMFENTIALLEINLAALMRSQQSDFLLCGKEKRLSVIASCSLW
NKDDKDRLIEQLYQKINALKEELENFKAESHRLGVQLRGRISELEAELAEQQHLKQQALDESDFLRTELEELK
RTVVDLETDYIFLIPTQQTLALHNANARNAKTYSYVFSQPSRMPVYPSWVGADHADDLQYVFGKPFSTPLGY
YCIRRYAVIKITMSGQIRKYYRRHVSTFKDFMSHMRSFVSHLQESTSLQNAIVVEQSLSLNETSSSLFAFFLQM
HGKSSLHIKDVKLSDSGRYDCEAASRIGGHQKSMYLDIEYAPTFVSNQTLYYSWEGNPINISCEVLSNPIAS

YQTELFQAVLEQLRWFAGPQIKNVAALGGNIMTASPISDLNPVFMACGSKLTLVSNEGRRTIRMDEKFFTGY

TLKKKSLGGAMVWAIDLDDFSDSFCGQGEYPLLRTLKQELRCGPVQTTPVPETTTTSSTTSPPTVSSSTLSFTTL
QMQQQLNDYEQLLDVKLALDMEINAYRKLLEGEEERLKLSPSPSSRVTVSRASSSRSMRTTRGKRKRIDVEE
VWKHNVIYVSSAGNNGPCLSTVGCPGGTTSSVIGVGAYVSPDMMVAEYSLREKLPANQYTWSSRGPSTDG
EHVQETKPTIMVTYNGDFFDWPFVEARAAVHGMNMYQEIGFQKDSQGEYKASQCIHMDCLRWVKRDSY
PTVVFSHPPIGTVGLTEEEAISTYGRENVKIYTTSFTPMYHAVTQRKTKCVMKLVCAGKEEKVVGLHMQGLGC

NLTLIPAASSAEAKANKDSAGLSYLKSTWWRLRTQSMPGVHVNRERKAAKQLGFIMAAFMVCWIPYFVLFM
AGYEKSTDRIAPWFHGIVTLRQAEELLSKSVPGSFLVRVSEKIKGYALSLLSPEGCKHFLIDASSNSYSFLGVDC
QDKNQAASSAEQTTCKKISNSFPCISVHFTQQRESTSRDEMVNGCYSPELSEKELNLYSDAKYPNLKLILSTS
RIMTTNTALTLQKDSVPIWLDNVECSGTEDSIFDCEKNDWGEHNCSHSEDAGVECA
KLKELIFEETARFQPGDRS
AGKCGPPPVTDNGDTIEFVKQVYESGSAVKYKCKSLHVMKGPQFVRCEYGHWTDPPVCLEPCTASPEEM
KYEVSDVNAVARLVKWSYLEGDVSKGAPYPPCTGAHRRAVCVYGAGDLHWMLQNHHLLANKFDPTVDE
GLEVLAVQRVSKAQAWQRTGRAGREDSGICYRLYTEDEFEKFDKMTIPEIQRCNLASVMLQLLALRIPNVLTF
HTPSMTLKTEPSGEDHGTISPTADLSPEEFSGFQHIRKGNFCDQYLSVPQHPYQWAKPKSLCPTSYMSPSL
LEKRGFHESFTKAKPGNFKVQAVPPREPASKVTVQGLLEPQSKNGTQEQDPGPVDLDDASLPVSDV
PLPNPATHKPLVEDFSNPETQNLDAMEQVGLDSLQFDYPGSQVQMDSSGATVGLFDYSSQQQLFQRTNA
LDDVMHVNVELPPKPKGAKHSAEPSAVLFSSGTMEYATIIFGDSPPRSGTKEKQCP

GWSPSEARSSFMSRTSKNEICAVLKLDNTVVGQTSWKPISNQSWDQKFTLELDRSRELEISVYWRDWRSL

FQFAACTALLDGSISIQSFSDQNISRPELKELLSKTKLEHPADNKPSFDNLYCEVNVTLQDGKVFSECCDTFY
LIPDINFNDAFENFALDFSREKKLLEGLDYLTAPNPPSVREELCTASHDTITVHWISEDEFSVSSYELQYTIFTG

VNPWTSCNIDEGPLAYATPEQISMDLKNDLMYRLEQDQELQRILQERETLPVKKFETEILDAIHHSPVVVIRG
DQSIGV
WYRKSCRPFLSKEVSDGKIYDAYVLYPKINKVDCIYTPDNFVLKLLPQVLERQCGYNLFILGRDDLPGKALVN
VLENQNLKIPSPKRRAVCVMAILAFVMLNYGPLSVFERDSSGAEPSVSLAHHSRNLLEFSASQKLSTAQQMA
TLCDEVNVYEFLPSKRQTDICHYYQRYHDRACTMGAYHPLLFEKNLVKHINQGQNELIYTHGKVTLPGFRN
ALPYVCKKPLHNTKSELPDVWRYLETQCDSGWLPYSGFCYMLMNNQASWNMASQSCRENKSDLISIHS
GEDGYYFTNLCCAVAFIEKLDAQSLNLSQEDFDRYMSGQTSPKKQESENWSSDVCLGVKQMYKNLDLLSG
LTYQLRSSLAVGPGTGHSYACRVQHSSLGDQSLLIPWEQSRPWGPGLAVGITLGVLAVAAVAVVLWRSRRR
PQESVLPFTEVKMEEEENVKNGQQASEQTEQTDTSAPDPGLRARKSRYRFMDYDNSENGKWATDDIPDD
SCVLNIQSSISALPSVASIVVSLEKKTATVKYDPDFISIDALRRAIEAVSPGTFKVSLPDEYENVELFSAQISPVKPS
RLRCLDAISSLLYLPPDQQTEDLLGMTESWFAALSSQPLELFRSISTQPFPDLHCGALRVFTAIANQPWAQKI
KEKQLRKLLEGPFSTPVPFLQSIPMYPCPSVSSSGAEKRKHSAAFPEGSFLEPSSGTVASQYMSQGASAGEG
FAAKYGLKNLTALLLTCPGALQAYSVANKYGYYPNNIAEIHGFKDLRQFIDEYVETADMLKSHIKEELMQGEED

IVPGLQHLLGSLTNYDVNQPNKHGTPPLLIAAGCGNIQMLQLLIKRGSRIDIQDKAGSNAVYWASRHGHVG
YNEEEDDDLYDIEDDIIYTQDFTVPGQVPDEEAAPNGQNRGKGLPKPVCMNGTEPGQLSTKSKVERRASA
LQPLTTAHQQSDVNAPAQNFGLVQNNNCSTLDGHNSSLAALNVSLQHQGIVRNTTQLGKSFFQWQVEQI

GLIPDFGKIFTPEFGSTGGKFHTTHTSTSGTDRGTGTGSFSSGGTASHTVTFEGSESFTDLGEGEEDIFDTSR

PTERFLVSTFEEIKAALLALASDPRYENPKLSKLEEILQEQFQTLGHSRGIIFTKMRQSAHALHKWIQENAKLAS
QIRPVPSKGTTGEALKIICEAECGENITIRWLKTPVELSQYQEEESKSKSILTVDRVDLNHQGIYECVMLSRRLG
PLSPVSTSGDACRICHCEGDEESPLITPCHCTGSLHFVHQACLQQWIKSSDTRCCELCKYEFIMETKLKPL

SEEEDDWTLLGAEESLLQSVEEALEERSLTIRPASGRVPTPEDQPVDPHCIPENETGTIKRVSLWDLTKWELG
ICEIAEHACLSDPCHNGGSCLETSTGFECVCTPGWAGPTCTLNIDDCSPNPCGHGGTCQDLVDGFKCIC
LSKGYKLASKYMNCFISPLLTIVAKNVAFFAGSILAVLIALTIYDEDVLAVEHVLTTVTLLGVGVTICRSFIPDQHL
KENLKLALDAMKDLAQYVNEVKRDNESLREIKQFQLSIENLNHSLLIYGRPQGDGEIRITTLDKRARQDRHII
PPDYQWYVPDQFPPDVQHKANDTPSEKEVHH
TMCEGNHESTTFLSTEPNILISDSELPRPLRSDGF
HVFVNLFDVQVQVFKIDENKFGAVEECQLDENNMVIPIIVGAALAGLVLIVLIAYLIGRKRSHAGYQTI
KKIVRYIIGIGNNFNWNRSQAFLTQMASEPTSQFVKVLDSFEKLKGLFSELQAKIYAIEGTSRRSSFHLELSSSG
YAESSDSTAWFLLALRDGKIEIQFKNELGTKVTSGGKAINDGLWHIVSVEELEHSINVKIAKEAVMNINSPRSL
PASRLLATGGMDRRVKLWEVFGDRCELKSSLSGSNAGITSIEFDSAGSYLLAASNDFASRIWTVDDYRLRHT

TAHHLRTLTSNLNEVRTTCTDTLSKHTDDLTFMNNTLANIRLDSTSLRMQQDVMRSRLDIEVANLSLIMEEM
WDEIGRTGKWAWEPGRPSYHVLSPVVSAFISDPNPQALNLSVSIRFSHPVPEKKTDLRLLCAYWEPGSRRV
GTVTYITGPEVTTYKAEIQYECESTFYTMKASNNGRYQCGADGYWKNSKGEKVLPICEPVCGIQTRTTLERIIG
LESSRWTTYSNGVPEPRPGSCSVPPSADKVLTFMKNHFLMDGKVVPRNKQPLLVKQNVKYTRITVDQTRSVS
CGKPITGRCISAMGRRFHPEHFICAFCLGQLQKGTFRDHGDKAYCHTCYSKLFV
LVQQHQTGEVIYQGRKKVIDVLACEQILSEAYKEVLLPMYMEGGFVEVIHDKRQYQLSELCKEFRLPFNVKVS

GAGVGPRCSTRRGRRLPRGQGDLLRYFRAAQGQLGTGMGQNGQESGQNASGHGTQEPAQDAMEQ
KQLSGISLALLLSPSTTIWEIKEELEREWGISPYTQRLALQEPGLGDQQLLDDQTLASHSIFYDTTVLMLTTEP

YTGSISCQEIVGQQRRTRECKGRSPDGKRCEGSYIHIRSCYNMYRCKLEGQWSDWSSWGLCIPPCGK
KPTSEEILLIAEQLNMEKEVIRVWFCNRRQKEKRINPCSSTPMLPAQGKTPGYGPHMVRSDTPGTVPPRG
VLLVGSALGSPDFL
GAIAVLAGLLVGSAFVYQHWIEIVLMYRNYLAKDETIGDRKEFDAFVSYAKQDSFERDSTFLKEEQFALEVLP
NRKVVCVMVVLLFIAFNFGPVSISEQKAVETVLPSQGAGAGQSRRHLLGVVEQPPKEEPGELREEVAPASVS
DRPRFGQGTGKIWLDDIRCKGTEKTLQNCAHRFWSYHDCTHREDISVVCQDN
GARTDARDVEHRSCHLNKENCKRRICRDDSKTHRCKCSNGFVLSRNRQYCEDVNECAFWNHGCTLG
NAEFDPLGTLPHGWEKRTDSNGRVYFVNHNTRITQWEDPRNQGDNGPQIAYVRDFKAKVQYFRFWC
STYTTIGVLVLLVIIALLLILYQFFRIDIVLLCQGIFKPYKTQDDGKIYDAYVIYPKNRTKEANFVEYFVHQILPDVL
ETWESGHRRGPDGTYSLSSFARLAPVQPHDHGATYSCHVTHAALATPARRSIRLCVAGATGPSLEDAIGM
PFFIKVQVKDTLDQPVGNLPVILGANVYDEQMDSTTLVQEGSDAGRRQTSMNDGTALFVVNIPSASKALEF
GAIPSAADVAQMPLLKAVMKEALRLYPVIPGNARVISDRDIQLGEYIIPRKTLITLCHYATSRDARFFPEPDSF

FLPEEMGDMQKLKVINLSDNRLKNLPYSFTKLQQLTAMWLSDNQSKPLIPLQKEVDPETQKTVLTNYMFPQ
PKRAVPVSLETKSRLTSFAAPENFESQPNVSSSSIWEHDKLEWLQEGKKRDGCRRRQSDPHYDPCTLHV
LLYMATQISSAMEYLEKKNFIHRDLAARNCLVGENHLVKVADFGLSRLMTGDTYTAHAGAKFPIKWTAPES
SWVADIQDCIDPGDSGDDIELSSCVQGACLPGRVPSCSCEFVADGANCIGDNTPSVHRTPDRCSVLAT

SNKEFLDTKDLMMFLEAEQGMAHVTEEISLEIIHKYEPSKEGQEKGWLSIDGFTNYLTSPDCHIFDPEHKKV
FFVFITPFITKNGESIMVEVAAGPFGNSEKNDGNLVEGPAERSAPHEKLPVVIRVPRLEYSALTICGMPFSLL

KLWVSSLEKKAEELTTQNIQLSNEVTLLRNEVAQLKQLLLAHKDCPVTALQKKTQGYLESPKESSEPTGSPAP
QGSSDSDISANAFLWHLEHPVRDFWPHLSAHWTTFYFSDVLQLQCLWV

FHISDFNFLMVLGKGSFGKVMLAERRGTDELYAIKILKKDVIIQDDDVECTMVEKRVLALGDRPHFLTQLH

QTHSNEIYQSTPAYRLSSSEKEIMKELMENGPVQAIMEVHEDFFVYKSGVYRHTPVMAGKPEHHRRHGTH

TLRVSKGQLPPTIACDSVACKDDACIYTLRCAIRNGRENVTYSWSNTASSVLSIDSIVRIPQRHQAARERVT
PLMDDSVTPIGDRPRLVKQDVKYTQIVVDRVTALNGSVYDVMFVSTDKGALHKAISYENGMHIIEEIQLFPN
YPSVGHLAQVLSASNIQPIFAVTGSTLPVYQELSRLIPKSVVGELKEDSSNVVQLISDAYNSLSSTVNLEHFQL
RRSSFNLTDLWDCTEYAIAIQCVGVESEIWSEWSEEKIGRTEQQAPSRKVDLWRVIESSPSAENRYVHLM
DGTNVTCRLSTPAGTPSSESSMVLDVKCESGGRGFELGLGLTAPAPPAALALTHKTLYGLLFLGGTWEVD

LPGSLEIPSQAELFSEFISPVPFTRPYASCSYCDFASVPPEQVFSAGHTEHDVGSGDYIETLSFTASGEKGITP
EFNSYYGNSVPASSATASRAGPCFDQHGGQQSPGMGLMHPTAAPNSMDPLQNSHEGYPNSQYNHYP
NLRDAKHKEAVTILSQQRGEIEFEVVYVAPEVDSDDENVEYEDESGHRYRLYLDELEEGGNSGSNRKHAS
RMYFVMNEKVWVCNLENCTVAEEIASPSITSPKKIVADPYNGYVFYLLEDGIYRASLPVYPGQVPEALPIVRL
INPDEELNLILPQTPVRAAMNTIQQLMMILNSASDKPSDNLLSYFNNCTVNPKQGILARVEDLGNIFKEKFA
PTPHDHIKVTQEQYELYCEMGSTFQLCKICAENDKDVKIEPCGHLMCTSCLTAWQESDGQGCPFCRCE
DPSSLGENPDGLSQEQLEHRERSLQTLRDIQRMLFPDEKEFTGGQSGGPQQNSGVLDGPQKKSEGPIQ
KGFINVTDSREDFEIDLLEKFCFEIKLKAYPPIRCIWVFSQKSFPCEQRYSADGYSISSRFCDHKYQSGEY

REPGGPQSPGSGKDSRMRANAALHNIIFSQPDEGQAKKEMRVLHVLEQIRSYSETCWDWLQMQLQSKD
LSKNNTFNAEKYSQTIRMDLSSSFVSDHSRLKRTSNNQMIVPQDPQRHVEKKTVSVEDEEGITQSDTILIDS
PNPPPENVQLVNYVSKNVISSEHVIERETEMSFSTSHYTSTTHHSTTVTQTPSHSWSNGQTESIISESHSVLVS
ENIKIGGLPAKGPPPPWLLDMKYLSPLLLSYEDRMREKEDLHLAHEEDMKKFRIRVEELVKENEQLHQQLN
FKYFVQGMGYMPSASMTRLMRSRTASGSSVTSLEGQRSRSHTGEGTRSRSHTGDLMVRDGYCWSPEGA
ASGPQGPTQTQCNDAYRNFNLSVIVGAEGILQGVQIWRVPATNTYSISGYGAAGGKGGKNTMVRSHGVS
EDTFGVSIAVGLAAFACVLLVVLFIMINKYGRRSKFGMKGPVAVISGEEDSASPLHHINHGITTPSSLDAGPD
RGAGEEKPPAAAPQLLSPPKSPAFARLPEQASPLLSPEAGGSRSSSVVSVEKCSTPGSARSRTPLTAAQQKYK
NRKVALTRLDSVVSRYAQLQDESEEEERVRKRAAQARGSAAPRQEGPLPPPCPMAPDPGEGQPHSQVSTS
HAHHKQTSPSSHQMHVIARGPAFANDFSDSPPQNLIAPSRNSLNMDLYDMNNPQVQQWQTAAPSHRD
VNGWTGEDCSENIDDCAMAACFQGATCHDRVASFYCECPHGRTGLLCHLNDACISNPCNEGSNCDT
KVSEQLKYCSGIIKEMFAKKHAAYAWPFYKPVDVEALGLHDYCDIIKHPMDLSTIKSKLENREYRDAQEFAAD
KNMEAVTKGTPRKGTPKEMEKKTNQTKAIEAPKTDYAAAVSGNNKKDKKQQKNQEANANRIKPSQVNLGE
AAAPGGAAFSKRYLEEQKTENGKDKEQKLTNVEKEKMKEKGGFSELGLADGKTKSDLYTSKTDPEKPYRSS
EKHSSHSKPERDSRYMSSRSRSDRDRRRSRSHSRSRSDRGSRTSSSYSRSERSHYYESDRRYHRSSPY
LSKISKASDHKKNFVRSPSTRSTLVKSKKIDVTHITSKVNTGLAALKKDTSSESTFLATEEGMKIMPGLNNASPS
APLTVSVANQGHPDRLSASWGAPAGGREGYVLTLYHAGSGAVAARAIAREDTHNFTFAELAPGYRHSLEV
LSLIGFAEGVSPGELFGPAGHCWAHHWCAAWSAGVQPQAGAGLSDVDKAVFSGISQKCEHCQRMGATI
VPQRFMAKWEVSVGHAEEFLLKETEDQKCSQNSSKLNSEDVLLESCTNGPVMEPDRQLNGCFKSLTFDSK
DSVAQGYGSLGMMTSVLICPDGKTVEAEAAHGTVTRHYRLHQKGQETSTNPIASIFAWTRGLAHRAKLDNN
PMPPSSTTPPQPQAKSKSCEFCGKTFKFQSNLIVHRRSHTGEKPYKCQLCDHACSQASKLKRHMKTHMH
TIAKQLLQRAKKGAQKKIEKEAAKLQGRKGRTQLKNIRQFIMPVVSAISSRIIKTPKRFIEDEDYDPPIKIARLES
SVFENLEQLLQQNKKKIRVESEKSKIYDHTLKFSPKRNISNCNGEEKKLDDSYSGSLTYSYKALMVPKDMLKK
LLKADSCTNCHGSQAKSVEIENLGNLKCLNEHHDVKKSPTVSTNVWLVQHPKTPFKVEDVCKANEPCTSF
YGLGVDFLKGVRAVNSSNIQHFAHCTKIFGSLAFLPETFTGDPATNTAPLKPEQLKSFGSLEELTGFLYIESWP
SKKTGNGVQHTEAVETCQSESNSLHEREPALESLWQPKTFHSNPLSQVSGKRLKRSIQKVNEWFSKSNEML
VSNSSGGAGAKAVPATGHTPSTPTPYAQAVAPPTPGANASQPRPPSAQQNASKQNGATSYSSVVADSTSDS
VITKLKEQYKKERKGKKGVKRKTTEDMFQPQSYIKQSKQGCGKMMAQSTGPNRSKLQDMLANLRDVDDL
DDSESDTQNVVTVCEPVNLAVQQPVNYIGKETLVLDSGYVPISSSNIEKSIVALVSTRTAVFKENTGHSSEGS
KSLLQQLENVTKERQMKLIQQQSDITGLSRQVKHVMNFTNWAIASGSSTALLYSKRLITFQLRHILKARCDP
ISAPNNLSASSKLPEVQHPKLAAKRRWTCSKPKNMLQETSMATSEKLMIEPPSAYPITPSSPLYTNTDSLTVITP
QQQEVASLSKQLEHVMNFSKWAVSSGSSTALLYSKRLITYRLRYLLRARCDASPVTNTTVQFHCDPSFWAQ
ANVHIDPKFISNGPFKDTVTVKVAVEDADEPPVFLAPSYILEVQENAAAGAVVGRVHAKDPDAANSAIRYSID
ASLEEDAASLSGESLDGHLQAVGSCPRLTPPPSATPDLPLSLPPHGLWSRHILQQTLMDEGLRLARLVSHER
RSLKPQQRAKVVERWIEVARECRILKNFSSLRAILSALQCNAVHRLKKTWDEVSRESFRTFHELSEIFSDENN
SFLPLAHMFERVIQSVVYCHGGRIGFFQGDIRLLSDDMKALRPTIFPVVPRLLNRMYDKIFSQADTPLKRWL
LKDKNIEVQKMKKAVESLMAANEEKDRKIEELRQSLNRYKKVQDMVILAQGKKGKEGECEEFLNSGSVSAV
PLIIASSPTEQEEAKDEQMNISAPAEGGDLLQEKQTGDEPVEVGNPPVPPGPHVLPQVSTPVSQNAPAFVPG
SKPDSSEPTNRSAVCAVSVRTINEFFNKIVDKQNMKCLQHFYGKDSKYCLKRTFSRNASYCEAQDDEYRL
ANSKFSFPSGRAGMQNGNCQTDFMVHQNVISGNSVNRQPPPPYPISPTNRQSPTALQMQAGGSVPPSA
KRCDPLDLSEELEQELGLTPSQLVEKRLLALSEEELSLPACPAPHLDSTPSQSEEEDEGSSQKGPASHRQAM
WSFGVVLWELLTGEIPYKDVDSSAIIWGVGSNSLHLPVPSTCPDGFKILMKQTWQSKPRNRPSFRQTLMH
SPLKDSDEVKSLWVKIDLTLLSRIPECLPEEPLAMSTGAKEEASLQQNNYTYPPAEKTVPKSRRKRKCENEED
GSVLKDQDLLRAKMDDAEACFILSSRCEVDRTAADHQTILRAWAVKDFAPKCPLYVQILKPENKFHIKFADH
QRSSSFPYGSKGSGLGSPSSNFNSAVFGPSSLNSLRQSPMQTIQENKQATFSSISHYNNQTLQDLLASESLS

SYSFTEEEQKTIEELRKAEVDGSSFLLKVWDGTKCPYPAWKVLVEESDLEADKVLIRQLRNLIVDILVYDNHV
LTLDDMLCQIPYGKKRHMQLLKQKQVSHRTFRPLKSALLAE
EQTSCPDGSTCCRLSTGAWACCPLVEAVCCPDHIHCCPKGYTCDPVEGSCQQALLSVPWLEKTPARVR
WGVAFHHAGLTFDERDIIEGAFRQGLIRVLAATSTLSSGVNLPAHRVIIRTPVFGGRLLDILTYKQMAGRAGR
VGRIEKAWMDGYSRQIFVTSKMLWPNGLTLDYHTNVLYWCDAYYDHIERIFLNGTDRKVVYNGKELNHP
ISTCDHGIVAKIKRVIMERDTYPRKWGLGPKASQKMMMIKQGLLSKHGKPNESTPDSWKKEYVDYSVAVK
KDSERQPFGTVTTYGISALREAFKTLSNSQDSDAAFTKLDETDFSFPKQLSPSVSHRLKRQREEENPHLETSE
LDVSLDNPWDKLQSVSVTLSSSLEEKSIIQPSLPSTQEAQLPNSTEEAAITQPDSNTPDFLEPSCQQSPCSLA

ESDVDCLREAGMSGVPADACAAAQKAAGYICKSNGGHGAIQEFAGYVCLLMKTVNL

EDVKSIANMIEKDWRQLLADCFPKILVNILPYFACESHRDSEITQQRDTAYKVYDMLKDESCLGKQQIDNLS

VKLEGELQSAQDLQADITSHASLYTSLLQLNESLFTAASEESVKAMKQEFKELDGRWKALPQTVSKRISFLQS
AQRICTWFQAELVSVHNQAELDFLGQSLKKGCGALHWLVHEATDWHQRKHLIGPAKLLHEEKGGGGGW

SPVILQFGHAETLQPLLALMGFFKDEEPLTANNYKKQMHRKFRSGRIVPYASNLIFVLYHCDHAKTPEEEYQV
EEMKFKLSPLSSYSEEFQDSLAPLEDLHVRLDSLWKDISQSLNEKSSDLQ

SVNFIENPQDLNGSIQAHVDVNIDLVSPDSGLATIRSSRSSKESSVFLSDDSPVAEGAHHSLLPGFDSYSPI
QRIIQCHGSFATASCLICKYKVDCEAVRGDIFNQVVPRCPRCPPDEPLAIMKPEIVFFGENLPEQFHRAMK
AKDEKEEYYYYLNICGKTSAGNCRDDTESVSSCQVKTRGQQEKVAGKFKNQTLRYSDGDLTLTYPDGDEC
VGLLGFLILIYKIFDSRVNSTGTSENFNVKDF
DPHYCQSFVDVSIKDFPLESFHCPSPKKMSFKNMDPSCTIGFYCRNMKDFERISEEITKKFQKVLTCPQILL
PSSPGMSLHLTLSSVNFLPFGPTSVQAKPLCTDVYPVYIKLKCVTLMFGREAVSFPTCFVQSLATLFLSQLMAS
VKCMYTPKMPIVQPCNIFDCPKWLAQEWSPCTVTCGQGLRYRVVLCIDHRGMHTGGCSPKTKPHIKEEC

GSIPTVDQLIEKFDRKIGPQRRGRSGKRNRIHPDDRKRSRSVDSVLSLGLEVNSDYLDEFSKNLGRSTEHLL
CHIDKDNLYSASTTLDYPSLGLMTEKLSQKNINLIFAVTQAVVGLYKNYSELIPGTTVGTLSSDSSNVLQLILDS
LQERMQRVGQRGSGLSMSAARSEHEVSEIIDGLSEQENLEKQMRQLAVIPPMLYDADQQRIKFINMNGL

FSLKEIVKQTVTQPQEKKMEAGGQSRPSVEEKATPQWKPKAFQINPSNKLSASGRKAVGSHAPKIGKAPQKP
QECIQKASLNITENMFCAGDLSGTKDSCKGDSGGPHVTKYKNTSFLTGVVSWGKGCATEGTYGVYTRVSK
EETQQRLAQEENSS
IFDRRCAEEDYRPWQLHSQGEACIMGAKRIYRKRKSEKKCMQGKYAGAMASEPCVCTEADFDCDYGYE
WYYYVTMPMPASTFTIAIGCWTEVKQESHIADIRTTNEFSSVFSRADLRLIEGSCGHGAYPCRFQNPTASSQ
ETSEPKPFLYYPEIKDKEEVQRKRQKLMPNFSDGYGGGSGAGGGGGGMYGGGGSGAGSGYGYPSYGY
QEMEDDYDYYGQQRVQILERNESEECTEDTDQAEGTESEPKGESFDSGVSSSIGTEPDSMEQHFMANLG
EIVMLSFHGKKAKGTKITGDPNIPAGQQTVEIDLAHPIQLPDIENLRNFNELSRIVLEVQEQVRREQQEEEEG
LSEETEFESVDKSSLFTDASFPPPAGTLLTLPQPGEGAEAIAGAEIRQRHICSSS

TQQTMAKSLQHVQQLQADLGGTEILAPLQAIYRSPCRDGHPRQLFVFTDGEVGNTQEVITEVQRHRGAHR

GIPECIKQYEEEKFRKKHKRIDQELDGKQEQKSEQREVVQLLRTVRAGEARSRYRPGRDLSPAGLQ
E
DNIEARTLEHHNSANISWQVPIAKDNSGDEVSIHVTPAFVPPSLFPIGEVVITYIAVDKSNNRASCTFSVKVIDT
MAGTPDKILEHLLEAMRLDATFSDPVDTLVGDFLLTYNVFMPTTQLCRALLHHFHAEPSEGSEQEKAVYSLSK
LIGFKPLVILKQHHYIKPSQFIYPEESFISGSTTLFNALLTKCLDKEVMALCRYTPCRNTPPRFVALVPQEEELDE
TVSSSKDSLRIQLTPPGESENKSMSEFYDLSYRVLYWKNSSDTKEKIQEEQQLLFTISDLTPLVLYCLKVQAFS
NLKVVENFINESMRYQPVVDLVMRKALQDDVIDGYPVKRGTNIILNIGRMHKLEFFPKPNEFTLENFEKNVP

KELFLSLAQTECVLKPVESPDMKIFSKADLEDTNSLFEKLKKEPDALTLLAPAPGDTIISLDFSSNESDEQQYDE
SKMLHGDCRFHTLKLTTVKKTERKGNANAQCKTKSA

NVIT
APMVLQQAAIPIQINVGTTVTMTGNGARPSQPGSEGPAPTQPSPPVLAGAPSPGEPGRGPESQLGPGAAPT
NSQLAHLNGSPFQLNSVTSALISGIVILGIMSIFFFVCSLTILRRKWPSSSVLGGVRNPVFN
CAKHKINPVWNEMIMFEVPHELLRASSVELEMLSQDGLGQSQSLGKCSLGLHATGTERNHWEEMLRNPRI
VSWMDSQGLILYQHSQENSFTLHEGPEAPGSSDVLDRFERCFSIFRLEVRGSPGAARGWKAVRVDLVVAP

NSRKSRGKARGRPRPRQPRRELRSPMLGPQRCQPILPPGAAPQRPLPWTRRFRLRQAFGKLQTKFESGQP

EEEMEMSDDDSSDNPSKKLRVDESALTAQAYALKEENDSLRWQLDAYRNEVELLKQEKGQLFRTEESLTKD

RLGNPRQPEKPLIRLRVDYSGGFEPFSIHRFSQKFVERTANPKDIIHFFRHREQKEKNEDEISFGKLFRRPAS
KVGTICSTIYQASGGSIDWSYDYGIKYSFAFELRDTGRYGFLLPASQIIPTAEETWLGLLKIMEHVRDNQS
QPIPESPKPPSEMVKASPNGQKSMLEKLKLFNSKGGSKAGGTSLECSGSRDTSCERLEALPSFEESEEIDAAN
CDCRLLWLLQRQTTLHFGSQQPMCASPDSVREKLFKDFHSTALSFYFTCKKPKIRDKKLQYLVVEEGQTVQ
FMDDLDGAGEALAGLDSANDTFAWVNNPCHNSKQSVQHREPLVYDNGHQTEETLLLREIESDLSNNNN

IRQSCAKMLVNRALGRWRQRMLRADNTSAIVICISPVLDSRSNWENEEELYLNLAEGPCYSSPDMCMMT
PSIHKSSRFVFVPGPEDPGPGSVLPRPPLAENITSEFRQQVPFSVFTTNPCRVQYCTQEIIIFREDLVNKMCRI
KNKGGVGVKPFQVKNHMWVFVNSLIENPTFDSQTKENMTLQAKNFGSTCKLSEKFIKGAVSCGIVESILNW
LTEGEQIIESTHLSDSSSDDGIEEAIQLYQLEKVRKEVNPTTDFQKGVADISASLTVSSTKCASAEDHKKSLSTKP

PSGVILSPGWPGYYKDSLNCEWVIEAEPGHSIKITFERFQTELNYDVLEVHDGPNLLSPLLGSYNGTQVPQP

AKLSESAEELRLKLNPYAKELKEKLRTDAEGLRQSLGPYVDGLSGQMEQKIEEFRQTVGPYGEAFNKQLVQK
LHNDTLRSLVPKMMETSSSITVQGLIQCGTPDCMGSILQIIRTGNVDPLVADAVTYSLGLLPFPCTKRIREILNI
LTHLLMALAGGKVLLSLEGGYNLSSLAEGVCAVLRTLLGDPCPQLEPPVAPCRSALSSVSRTLAIHEKYWRSLP
KDYSGLTIQFAGSCALMRKAVNGKVFCLLTSMFENTIALLEINLAALMRSQQSDFLLCGKEKRLSVIASCSLW
NKDDKDRLIEQLYQKINALKEELENFKAESHRLGVQLRGRISELEAELAEQQHLKQQALDESDFLRTELEELK
LRTVVDLETDYIFLIPTQQTLALHNANARNAKTYSYVFSQPSRMPVYPSWVGADHADDLQYVFGKPFSTPLGY
YCIRRYAVIKITMSGQIRKYYRRHVSTFKDFMSHMRSFVSHLQESTSLQNAIVVEQSLSLNETSSSLFAFFLQM
QHGKSSLHIKDVKLSDSGRYDCEAASRIGGHQKSMYLDIEYAPTFVSNQTLYYSWEGNPINISCEVLSNPIAS

QTELFQAVLEQLRWFAGPQIKNVAALGGNIMTASPISDLNPVFMACGSKLTLVSNEGRRTIRMDEKFFTGY

LKKKSLGGAMVWAIDLDDFSDSFCGQGEYPLLRTLKQELRCGPVQTTPVPETTTTSSTTSPPTVSSSTLSFTTL
QMQQQLNDYEQLLDVKLALDMEINAYRKLLEGEEERLKLSPSPSSRVTVSRASSSRSMRTTRGKRKRIDVEE
WKHNVIYVSSAGNNGPCLSTVGCPGGTTSSVIGVGAYVSPDMMVAEYSLREKLPANQYTWSSRGPSTDG
HVQETKPTIMVTYNGDFFDWPFVEARAAVHGMNMYQEIGFQKDSQGEYKASQCIHMDCLRWVKRDSY
TVVFSHPPIGTVGLTEEEAISTYGRENVKIYTTSFTPMYHAVTQRKTKCVMKLVCAGKEEKVVGLHMQGLGC

NLTLIPAASSAEAKANKDSAGLSYLKSTWWRLRTQSMPGVHVNRERKAAKQLGFIMAAFMVCWIPYFVLFM
GYEKSTDRIAPWFHGIVTLRQAEELLSKSVPGSFLVRVSEKIKGYALSLLSPEGCKHFLIDASSNSYSFLGVD
DKNQAASSAEQTTCKKISNSFPCISVHFTQQRESTSRDEMVNGCYSPELSEKELNLYSDAKYPNLKLILSTS
RIMTTNTALTLQKDSVPIWLDNVECSGTEDSIFDCEKNDWGEHNCSHSEDAGVECA
KLKELIFEETARFQPGDRS
AGKCGPPPVTDNGDTIEFVKQVYESGSAVKYKCKSLHVMKGPQFVRCEYGHWTDPPVCLEPCTASPEEM
KYEVSDVNAVARLVKWSYLEGDVSKGAPYPPCTGAHRRAVCVYGAGDLHWMLQNHHLLANKFDPTVDE
LEVLAVQRVSKAQAWQRTGRAGREDSGICYRLYTEDEFEKFDKMTIPEIQRCNLASVMLQLLALRIPNVLT
HTPSMTLKTEPSGEDHGTISPTADLSPEEFSGFQHIRKGNFCDQYLSVPQHPYQWAKPKSLCPTSYMSPSL
LEKRGFHESFTKAKPGNFKVQAVPPREPASKVTVQGLLEPQSKNGTQEQDPGPVDLDDASLPVSDV
PLPNPATHKPLVEDFSNPETQNLDAMEQVGLDSLQFDYPGSQVQMDSSGATVGLFDYSSQQQLFQRTN
DDVMHVNVELPPKPKGAKHSAEPSAVLFSSGTMEYATIIFGDSPPRSGTKEKQCP

GWSPSEARSSFMSRTSKNEICAVLKLDNTVVGQTSWKPISNQSWDQKFTLELDRSRELEISVYWRDWRSL

QFAACTALLDGSISIQSFSDQNISRPELKELLSKTKLEHPADNKPSFDNLYCEVNVTLQDGKVFSECCDTFY
IPDINFNDAFENFALDFSREKKLLEGLDYLTAPNPPSVREELCTASHDTITVHWISEDEFSVSSYELQYTIFTG
VNPWTSCNIDEGPLAYATPEQISMDLKNDLMYRLEQDQELQRILQERETLPVKKFETEILDAIHHSPVVIRG
QSIGV
WYRKSCRPFLSKEVSDGKIYDAYVLYPKINKVDCIYTPDNFVLKLLPQVLERQCGYNLFILGRDDLPGKALV
LENQNLKIPSPKRRAVCVMAILAFVMLNYGPLSVFERDSSGAEPSVSLAHHSRNLLEFSASQKLSTAQQMA
LCDEVNVYEFLPSKRQTDICHYYQRYHDRACTMGAYHPLLFEKNLVKHINQGQNELIYTHGKVTLPGFR
ALPYVCKKPLHNTKSELPDVWRYLETQCDSGWLPYSGFCYMLMNNQASWNMASQSCRENKSDLISIHS
EDGYYFTNLCCAVAFIEKLDAQSLNLSQEDFDRYMSGQTSPKKQESENWSSDVCLGVKQMYKNLDLLS
TYQLRSSLAVGPGTGHSYACRVQHSSLGDQSLLIPWEQSRPWGPGLAVGITLGVLAVAAVAVVLWRSRRR
PQESVLPFTEVKMEEEENVKNGQQASEQTEQTDTSAPDPGLRARKSRYRFMDYDNSENGKWATDDIPDD
SCVLNIQSSISALPSVASIVVSLEKKTATVKYDPDFISIDALRRAIEAVSPGTFKVSLPDEYENVELFSAQISPVKP
RLRCLDAISSLLYLPPDQQTEDLLGMTESWFAALSSQPLELFRSISTQPFPDLHCGALRVFTAIANQPWAQK
KEKQLRKLLEGPFSTPVPFLQSIPMYPCPSVSSSGAEKRKHSAAFPEGSFLEPSSGTVASQYMSQGASAGEG
AAKYGLKNLTALLLTCPGALQAYSVANKYGYYPNNIAEIHGFKDLRQFIDEYVETADMLKSHIKEELMQGEE

VPGLQHLLGSLTNYDVNQPNKHGTPPLLIAAGCGNIQMLQLLIKRGSRIDIQDKAGSNAVYWASRHGHV
YNEEEDDDLYDIEDDIIYTQDFTVPGQVPDEEAAPNGQNRGKGLPKPVCMNGTEPGQLSTKSKVERRASA
LQPLTTAHQQSDVNAPAQNFGLVQNNNCSTLDGHNSSLAALNVSLQHQGIVRNTTQLGKSFFQWQVEQ

GLIPDFGKIFTPEFGSTGGKFHTTHTSTSGTDRGTGTGSFSSGGTASHTVTFEGSESFTDLGEGEEDIFDTSR

PTERFLVSTFEEIKAALLALASDPRYENPKLSKLEEILQEQFQTLGHSRGIIFTKMRQSAHALHKWIQENAKLAS
QIRPVPSKGTTGEALKIICEAECGENITIRWLKTPVELSQYQEEESKSKSILTVDRVDLNHQGIYECVMLSRRL
PLSPVSTSGDACRICHCEGDEESPLITPCHCTGSLHFVHQACLQQWIKSSDTRCCELCKYEFIMETKLKPL

SEEEDDWTLLGAEESLLQSVEEALEERSLTIRPASGRVPTPEDQPVDPHCIPENETGTIKRVSLWDLTKWELG
ICEIAEHACLSDPCHNGGSCLETSTGFECVCTPGWAGPTCTLNIDDCSPNPCGHGGTCQDLVDGFKCI
LSKGYKLASKYMNCFISPLLTIVAKNVAFFAGSILAVLIALTIYDEDVLAVEHVLTTVTLLGVGVTICRSFIPDQHL
KENLKLALDAMKDLAQYVNEVKRDNESLREIKQFQLSIENLNHSLLIYGRPQGDGEIRITTLDKRARQDRHI
PPDYQWYVPDQFPPDVQHKANDTPSEKEVHH
TMCEGNHESTTFLSTEPNILISDSELPRPLRSDGF
HVFVNLFDVQVQVFKIDENKFGAVEECQLDENNMVIPIIVGAALAGLVLIVLIAYLIGRKRSHAGYQTI
KKIVRYIIGIGNNFNWNRSQAFLTQMASEPTSQFVKVLDSFEKLKGLFSELQAKIYAIEGTSRRSSFHLELSSS
YAESSDSTAWFLLALRDGKIEIQFKNELGTKVTSGGKAINDGLWHIVSVEELEHSINVKIAKEAVMNINSPRSL
PASRLLATGGMDRRVKLWEVFGDRCELKSSLSGSNAGITSIEFDSAGSYLLAASNDFASRIWTVDDYRLRH

TAHHLRTLTSNLNEVRTTCTDTLSKHTDDLTFMNNTLANIRLDSTSLRMQQDVMRSRLDIEVANLSLIMEE
WDEIGRTGKWAWEPGRPSYHVLSPVVSAFISDPNPQALNLSVSIRFSHPVPEKKTDLRLLCAYWEPGSRRV
GTVTYITGPEVTTYKAEIQYECESTFYTMKASNNGRYQCGADGYWKNSKGEKVLPICEPVCGIQTRTTLERIIG
LESSRWTTYSNGVPEPRPGSCSVPPSADKVLTFMKNHFLMDGKVVPRNKQPLLVKQNVKYTRITVDQTRSV
CGKPITGRCISAMGRRFHPEHFICAFCLGQLQKGTFRDHGDKAYCHTCYSKLFV
LVQQHQTGEVIYQGRKKVIDVLACEQILSEAYKEVLLPMYMEGGFVEVIHDKRQYQLSELCKEFRLPFNVKV

RGAGVGPRCSTRRGRRLPRGQGDLLRYFRAAQGQLGTGMGQNGQESGQNASGHGTQEPAQDAMEQ
VKQLSGISLALLLSPSTTIWEIKEELEREWGISPYTQRLALQEPGLGDQQLLDDQTLASHSIFYDTTVLMLTTEP

YTGSISCQEIVGQQRRTRECKGRSPDGKRCEGSYIHIRSCYNMYRCKLEGQWSDWSSWGLCIPPCGK
KPTSEEILLIAEQLNMEKEVIRVWFCNRRQKEKRINPCSSTPMLPAQGKTPGYGPHMVRSDTPGTVPPRG
GVLLVGSALGSPDFL
YGAIAVLAGLLVGSAFVYQHWIEIVLMYRNYLAKDETIGDRKEFDAFVSYAKQDSFERDSTFLKEEQFALEVLP
GNRKVVCVMVVLLFIAFNFGPVSISEQKAVETVLPSQGAGAGQSRRHLLGVVEQPPKEEPGELREEVAPASVS
QDRPRFGQGTGKIWLDDIRCKGTEKTLQNCAHRFWSYHDCTHREDISVVCQDN
PGARTDARDVEHRSCHLNKENCKRRICRDDSKTHRCKCSNGFVLSRNRQYCEDVNECAFWNHGCTLG
QNAEFDPLGTLPHGWEKRTDSNGRVYFVNHNTRITQWEDPRNQGDNGPQIAYVRDFKAKVQYFRFWC
STYTTIGVLVLLVIIALLLILYQFFRIDIVLLCQGIFKPYKTQDDGKIYDAYVIYPKNRTKEANFVEYFVHQILPDVL
TETWESGHRRGPDGTYSLSSFARLAPVQPHDHGATYSCHVTHAALATPARRSIRLCVAGATGPSLEDAIGM
LPFFIKVQVKDTLDQPVGNLPVILGANVYDEQMDSTTLVQEGSDAGRRQTSMNDGTALFVVNIPSASKALEF
GGAIPSAADVAQMPLLKAVMKEALRLYPVIPGNARVISDRDIQLGEYIIPRKTLITLCHYATSRDARFFPEPDSF

FLPEEMGDMQKLKVINLSDNRLKNLPYSFTKLQQLTAMWLSDNQSKPLIPLQKEVDPETQKTVLTNYMFPQ
TPKRAVPVSLETKSRLTSFAAPENFESQPNVSSSSIWEHDKLEWLQEGKKRDGCRRRQSDPHYDPCTLHVP

VLLYMATQISSAMEYLEKKNFIHRDLAARNCLVGENHLVKVADFGLSRLMTGDTYTAHAGAKFPIKWTAPESL
HSWVADIQDCIDPGDSGDDIELSSCVQGACLPGRVPSCSCEFVADGANCIGDNTPSVHRTPDRCSVLAT

SNKEFLDTKDLMMFLEAEQGMAHVTEEISLEIIHKYEPSKEGQEKGWLSIDGFTNYLTSPDCHIFDPEHKKV
TFFVFITPFITKNGESIMVEVAAGPFGNSEKNDGNLVEGPAERSAPHEKLPVVIRVPRLEYSALTICGMPFSLLG

KLWVSSLEKKAEELTTQNIQLSNEVTLLRNEVAQLKQLLLAHKDCPVTALQKKTQGYLESPKESSEPTGSPAPV
DQGSSDSDISANAFLWHLEHPVRDFWPHLSAHWTTFYFSDVLQLQCLWV

RFHISDFNFLMVLGKGSFGKVMLAERRGTDELYAIKILKKDVIIQDDDVECTMVEKRVLALGDRPHFLTQLHS

QTHSNEIYQSTPAYRLSSSEKEIMKELMENGPVQAIMEVHEDFFVYKSGVYRHTPVMAGKPEHHRRHGTH

FTLRVSKGQLPPTIACDSVACKDDACIYTLRCAIRNGRENVTYSWSNTASSVLSIDSIVRIPQRHQAARERVTC
PLMDDSVTPIGDRPRLVKQDVKYTQIVVDRVTALNGSVYDVMFVSTDKGALHKAISYENGMHIIEEIQLFPN
YPSVGHLAQVLSASNIQPIFAVTGSTLPVYQELSRLIPKSVVGELKEDSSNVVQLISDAYNSLSSTVNLEHFQLP
RRSSFNLTDLWDCTEYAIAIQCVGVESEIWSEWSEEKIGRTEQQAPSRKVDLWRVIESSPSAENRYVHLMW
DDGTNVTCRLSTPAGTPSSESSMVLDVKCESGGRGFELGLGLTAPAPPAALALTHKTLYGLLFLGGTWEVDW

LTVVTDLYELEESTPEIFDTTFPSRPVVSFSSRLIEVSESNLFFSNSVDVGLNIMSTTIFPSYRQPSEGISLDITSV
GYGRPSGAAAGGAGAAAVAAAAAAGAGGGGYGGSSSGYGVLSSPRQQGSSMMMGPGGGASLGKAAAA
AAVKPSEVFVKKPSVDGHENGDLGNSVETPLEETAPKLADSAESLP
NALQDFVINFLSKRLIYFNKTDQSFVSGFLDGSEFNILRPQVSFRDIESFVYENNTFVVTDGRAVFREELSPT
AVGQGCAEIGSQRYKLGVRLYYRVMESMLKSEEERLSVHNFSKLLNNNIFHTSLLACAVEVVMATYGRNA
KGTEPIIVDPFDPRDENSRCCSIIDSFSMPMLDLDDDDDREESLMMNRLASVRKCNERQNSPVTSPGSS
MMAQSQSLGKGPGPRTDGGAPFGPQGHRDMPFSPDEMVPPSLNSQSGPVGPDHLDHMTPEQVAWL
FYAENDDTYVNKTMMLYVKRKPEVTIQSTSKGISCYSDSYPAPSWTWRKCPGPDSSNCTEEITEGIWSDT
GGKGPEGSAVPVPIEPQICQATCAIMKLSFDEEYRRAMNELGGLQAVAELLQVDYEMHKMTSDPLNLALR
HKASQTSNQGCRTRTVVPGCVSFEPVFHGDKTVFFSSCNDMEVTGNCTVVNCNETSKATGKSCYEAFQ
SSVENSRHMSPAGARGRLSGIGGPRDCNSFLRHARETPDSYRDSPHSERYVSAMTTPARMSPVDFHTPS
KKSSITLKEWQQLQTQAKLVLEENGVLMKQLEIQQAKAKESYNQHLQEVSKLTKQLMVLEAKKQSQEEEISE
QYILGTISGTTTCCLGYQAHSAILCPGTTHFSASPFEPDSQATMHLDTVCFSIVHFSTTHPGAAYSCTEICGT
ILGIFDLQKDDALYILVGQQGEDACPSMENGEPVPLIIAAGGGGRAYRAKTDTFHPERLENDSSVPGLNG
TVVIGMTRIPVIENPQYFRQGHNCHKPDTYVQHIKRRDIVLKRELGEGAFGKVFLAECYNLSPTKDKMLVA
GDVVCTPNGIRKKFNGKQWRRLCSREGCMKESQRRGYCSRHLSMRTKELESLSEGRASGLREGSAEFD
AASQGAPAVSEEEDDDESEEEEPSSDTDIEEELQQSQEGGEEEEEDAAMQEAAGEETAGDQEDQCMELG
SLQNPGIETSPRQHVSFRPDASVPSRTNSFNNHQQQQVPVSIRQVPAGKPDPSITSPNTITAVTSAHILQP
NPVNGKAICTCPSGYTGPACNQDIDECSLGANPCEHAGKCINTQGSFQCQCLQGYSGPRCEIDVNECL
VRLMFSNCYKYNPADHEVVAMARKLQDVFEMRFAKMPDEPEEPMIPASSPVVVPPPTKVAPPSSSDSSSD
GVTVYFHAILSKDFKLDPNQHKVFIRAQGISGYLNWKDNICELTCTKDLGEHGLLIEGWVTISKDNMNKQF
SPKRYKFRDDFEKLKMAEFHKEGHYGKEEMDEQDKKDKAKGRKDSEFDDEPKFMSKVVATSSKNQDED
ERTRYSRSYADSRTRESSDSEDEYRRTHSKSSDSRRTSSHSSSYRDSRTSSYSKSDRDSKTESSYAETERR
TSKNPFDEGEIQLGDKVLVVGQRTGTVRFYGTTKFAAGFWCGIELNKPHGKNDGSVGGVQYFSCLPRYGI
SIAGPYRAAADNVSDWTYPLPPPNISLTNQGHSDRLTTSWGAAAGGRDGYTLTLYHAGSGAVAARVSVGN
QCHSEGCPRHYHFPCAAASGSFQSMKTLRLLCPEHVAEAVQMEDSRCMVCDGLGELRDLLFCTSCGL
HLAREKGRSCTKSQVKKSSDSRKRTRVKKSSTRLEASRREIGGKAPENIVRNRVAGDLPDKHASHELCRIA
ADLKNFATALEEVCVETIEAGFMTKDLAACIKGLPNVKRSDYLNTFEFMEKLAENLSLKLNSQPRL
KAGSMTGRSDDGLSTTSSPEPGTSELTGEGLKSSETDFRNESDPSLGHDNEEEEEEEEEEEELENESRPES
TPNSRFSATSCGSSEKSSAASQHSSQMSSDSSRSSSPSVDTSTDSQASEEMQALSEECSNTPEVHTPLPISQ
TKKLVETTTNMNASFVSFLKTAAENAEVNSIIQLRQLKAFKSVLSDIKKAHLALEEGLNQEIQALDVKIKEKNTK
ECVCDDSCEKEALCKWLLKKEGKDKNGMPVSQIPVVNPESEKPKSAVKIWLHPSRQLPEESATTTVKLDSS
ETFPDLSIFQNLRIIRGRVLHNGAYSLTLQNLSIVSLGLRSLQEISSGMVLVHQNPNLCFLQKVPWKAIFRNR
SSGSSQDAPVGATDTEDGDSYLSDRESCISEKTDLMVNCTEVVMGCENDKSLSKSIVHNIEDKIFGKTYK
ILGSSGPALPSQPVAHNPPSSAAKEPSGAPAPPSASSGLLVPMTVNTPSSPTPSFSEAKPSQPLLNGPPQFSS

PSLQPSVTSPSRPSSSRLIEHEINRTDEVEALTFPPTSGKSFIMGADEALESELGLGELAGLTVANEADSLTYDI
ISGLSSNTGSSTLTSDREINVHSPTDSKLWYKDSSSQSGSVGINLNSVQDKSNRIGSTEASEPSSFTNPVYSY
PAANGAIRFHCDPTFWAKNVVNLGNLVIENKPTPGYTPNVVVGQVPPGTNHINKAPGQINLAQLRLQHM
IKKKRGRPKKQPLLTVETIHEGTSTSPVSPINREFPGTKKRKRRRNLAKLAQMVPGEDKSTSELKFHKKVGKL
NIFNLGSLVIEDKETPPHMPKSPVMEPNLQPQSGLPSNQLSKFPTQISLAQLRLQHMQQQVMAHRQQAQ
RHTDLERYFSISPEDGYIKTIKALDREEIAWHNISVFASEVYNRHQEAKVPVAIKVLDVNDNAPKFASTYEAFI
VGRLSPCLPGKSPGPEFEDRLAEGAPPAQAESHRSGVKVEQETSRQ
IHSLSRELLIKEGTSKFATLEMNPKRAQKRQQQQREMGVMQGTIPYLGTFLTDLVMLDTAMKDFLDGGLIN
LEFAAKRKKAEVQSGIIRNDSLWDKLFFNKIQASLGGCVRMIVTGAAPASPTVLGFLRAALGCQIYEGYGQT
LLDMQSLTDQEKSPSPIPVISSPSHEEFNTTVHEENSLQIHTSILQVSIPAFPPAPESSEATTEKVEAQPKPEVSS
SFPIPSQPEFSHDIFISTQSLEEKPSGEEKGGVLLGSCLPADVSETTDALPKIPTGDPELSCKLVLSASECSQPT
EVTTPLQRVDLFMGQFNSVLLTSISVFTKGNLTVANLGTSEGRFMQIVISRSEATSPHVNFPLDSHPVSPQVI
YTNGNIPQSMMVSNRNSHNMELYNINLPGIPVSWSQSSSVPPQSPPGSGHEIPTWQPNIPVRSNSFNN
SGASPPRGSRAKGGALGTETPETPTPTPVQPSQARQCQTWDAAAPRCLSPHKCGGEGRLLASLRSRVERV
LDIASADVLATPQETYFKSQAEWREEVKKHFEKIKSEGTCIHRLDEELIRRRREELRHALDIREHYERKLERA
EIKKSHLERECSSRLIASAPGISTANHCNRNEHSLAIPINKNEKILRSPISPLSDASKHKYSTDDLTSSNKPNG
VVCEEEFKYAMLALNCICPATSTLITLLVHTSRGQTAPWEAFRHLTGTKQTLTREGQQSPEQWQKMYGRCS
HSDVMMTQSDPLMSQASTAVSAQNSRRSIMLRSDPMMSFAAQSNQGSLVNQNLLHHQHQSQNSSLG

LAKTLEVGNFLRIYSLHTKQARVENQPNVSYVEFHLHGGTCYGRGISILPENSTDVKELKAVMDSVNLTEF

PSRGRDVQCDEQTSCPDSSTCCRLSTGAWGCCPLVEAVCCQDHVHCCPKGYTCDPAEGSCQPPAV
KGVDIVGESILVCKSSERSKGIALLQGSLKPVHSCLLRGEGEGVTSSMIRAILEIIVGGVASTPDDVRTYASCT
GLSHHGNYIFWTDYMNGSIFQLDLMMRNVTLLRSERPPLFGLQIYDRYKQQGDNACRVNNGGCSTLC
KEAEAPAAIAERVPKRKRETESENEETAVAPSSPATPLAEEQSRKEKKKKKKEKKAKQAAESEGELMEVVGSW
TSDVPKSPPKSKRLLTISRLIMEQDSQSTELSETPDSSQEPVVSSAFRTPVQKPHVQCVSSRSPKPLKARWNT
DEPVETVSPSLLSFSSTCPPEAASRGTAAAMFQRVPDVDKHPDASHDLPCGQPSGAFQAHLCPFSPALPIS

HSNLPEVVVELLMTLHEPVNTTAGEGADLSKFTGELDPAPNPPHFPSYVIMATLDYISSCHKTKLKSIVAVLSK

LVAEHGEFDELLLSFSEWINQFLGELQDTSEINTADQHPAVCRNQDHSMEVESKKQELQSLREHVEKLCS
FSRGQEQHWWIGLHTYENDGRFKWSDGSLLNFISWAPGKPRPISKEKKCVYVTASREDWGDQKCLTAL

QILLNEKLLPFPHSEETVSLYTDLKNYYKDILQNCHFSEECELSKTNNTSTDEL

EGAIPEEQQPQPRNNSDNFDLFSFDLAPMVTIRSESSSHSVDYSPADDFFLNSDSSEGQPPIVQKELDET
YDKDEVDLLIVIGSSLKVRPVALIPSSIPHEVPQILINREPLPHLHFDVELLGDCDVIINELCQRLGGEYTKLCY
SSGFQRMTVINFECNETAGNEGKGAPVFTGEVDCTYFFTWDTKYACVQEKEDLLCRVADKKKRYDLSPLI

AIQEMTEDMKTVLQLVTHECKLIEFCIYVVVQFASVKHVARMGPETPSVAIMLKSSSKEKYPLFTFVKGHSRD
SFNPCVEEITNITYRCMELNLSGVPAEIPPSTQNLDLSFNPLESLTSNYFSAVPALRFLDLTRCHIQTIEDYAF
IVPTPCYKPKEKLPVEAKLPWYKQAQELEDGAMVSEEPSFIPEAWSACSVTCGVGSQVRPVKCQVLLSFSQ

RPSKICLQKQLSLEQTLPSPDTQVAAGTVSKLQPTAQDIQNNHYNSLQHHKENKVNGASLMFRSSTFPVQ
YGKIRSKVELEVRDLPEELSLSFNATCLNDEVIQGLKSCVGLKIGDTVSFSIEAKVRGCPQQQEKSFTIKPVG
MDDPMKVYKDRQVMNMWSEQEKETFREKFMQHPKNFGVIASFLDRKTVADCVLYYYLTKKSENYKNLVR

QPELSEDADTRDKRKPAKSAPNTVNPCATSLESFLTKSREELLEYAAKITLDFNTAHNKVLLSDRNTKMSVSE
LEWLNNHMDS

RHTNSQCLPAFWYNPSSLSKDCSFGQSYLNSTGYRKVVSNNCTDGVREQYTAKPQQCPGKAPHGLRIIT
LVIPHRVFAPWCLKDRCAEILLELIPRCLSAAHGTLGTHPFSRLDVLIVPSNFSSLGMASPHIIFLSQSVLTGG
AYGGMPFHPGTTKSSSGMKHGISNDMGEQDGESCNKHDDKPDMKHNTTVEITERRESRVCSGDNKE
EGQQEPSQAEQNEVPAEGTQTQPQHLDANTSSPDRSDDIEMDSKVLTVSNNSEKGTLQPSVNTTVAQPL
QDEEEGRSRQDDLQQVEQPAQRDDAGAEGTEAAGAAAAQESTLTSQPFVLPMGVISRNEDYPRTCQMC

CFSFGIGEGASTALVKGIARAAGGSAEFITGQDRMQPKALQSLKRALQPAVTGISLSWDLPPGMEAALLGR

TESPVIDRCRSPPPVQVLENEYRATWEEPQFSDNSGAALTITKSHSPGDLFLKGETTVQYTATDPSGNNRTC
RQKILRLVSQWVLLYGPLLQADHSAINLLQHLSDFVSRDPRLCNLLRDPTQDRRRNRTLENGGGSTSPQL
QKVQTAPPGFHLIFLPYADDKRKIDFTEKVPANDEQVDKMKEIIQKLRFKYRNDSFENPVLQQHFRNLEAL
PAYKKDGYFSDVECIKTLNDKTSPLTILKTFAIAMVAFFFFASFLIFGVYYISRRIRYAFFPSCKPPSNIECLGG
SRYFQPFGFGPRGCVGKFIAMVMMKAILVTLLRRYRVHTPKGRGLKNIQKSNDLSMHPNERQPSLEMF

VPLYNDVMLPSSSEKLKNINLAMSPLPVSEITKPLRSNADPALNREVVSKLEPNTEPLELAFSSRRRQEQPAS

QTQGSQTSHPRVIRISHQTVEPVVMMHMNIQDSGSQAPGGGSGGTVSMGTAGPAGHGPGIGGAAHVP

RQIAMWHQLHM
VSQFPFALLGWTGSRHFERELRRFAGHERKMVLNSHTLYDTRRNLFLPAASEEEIFQHLGLEYVPPAQRN

SKEPRETKPFSLKALIRWLHISKPAQHVMSLLGCRYPSDLNSFSRSHLPGPWDPSQAGTRMKLPQPQTWD

QQLKFLQQTMQGMQQQLLSIQEELNNKKSELEQAKEEQTHTQARLKVLQEQLKSAKELAEINGHDESQA

EGTTLRVEDLVKQYFQTAEKKVQLSLLTERGMGEAVQEFVDKEEKDAIEELVKFQLEKTQRFLKERHTDAEE

RNTNNPGPTSSSPKIALKGIAQRTFSRALTNKKSSPKGSEKEKQKEKEKDKSKDIAKRISVTEKQEIKEEPKEE
LVCNADGDPQPTISWVTPRRRLITTKSNGRATVLGDGMLEVRFAQDQDTGIYVCIASNAAGNDTYSASLT

NIEDGKLMTCISIQNSCSCLEGPDSQLYLSGWDTYADAETAEMIAHRTGGMKISATVIFNPKSPATSESPVTT

PSRCCTPPIKLLEEDLWPRLNSSEPVPSLIHSNAILDKYSELSSEIAKSSIPLAVASSRDLDPQEENCSKTLALR
NCVRFPSTNLDIPTHFVKTILSQGHLTPLPLYVSPVFWAYDYTLRVYPVPDLIVTADKYDPFTVTNTDCLCINP
VKFKAQNQLNKKCSAVKHTRIKGIPKLDDANDAGSKNSTDCTLILTEGDSAKTLAVSGLGVVGRDRYGVFP
RREMSTKSTQFKGTGNEFNKLFKPLKKARSCASPVNKIAKCKLTLQASCTADTSTELMCAEAILDISKTIMPPQ

FLFSSSNFIYLLFTTDNSRSNNGFKIHYESVTVNTYSCLDPGIPLHGRRYGHDFSIGSTVSFSCDPGYRLSH

VEEMRQKLGPYAGEVEDHLSFLEKDVRDMVASFFSTIKQIEN
MAQYQQSRATFYALSHAVTNFYEDKKTVTQEIMDVANFMESMIGNECSGNDELTYLTLRAIGNMGKAMEV
RRAEADIPPEDVEEEPAPTTPPDSPEVPRPLPAARTGLVYDERMTEHYNMWDSQHPELPQRVSRIFQRHGE
TPNSPLCTESFKKESCKPTNKKNLGTKSTMKDQSLVFHSKIKSSGYTTAPQMTMFSPKTNLKKNNKISEWKSS
KKQWEDTEKAQRSLTEIERKAQANEQRYTKLKEKYSELVQNHADLLRKNAEVTKQVTVARQAQGDVEREKKE
YWPKHRTVSKALIAYWTNFARTGDPNKGESTVPIKWIPYTWEESYYLEINSDINYDSVKQSLRASYMQFWN
MIHNNILEIEGRYYLQCCGIPQGSILSTLLCSLCYGDMENKLFCGIQRDGVLIRLIDDFLLLTPHLTLAKTFLRT
SVHWRRGKLVLPAKNTTHLKTYSAGRKLILEIAPRSDNDFGRYNCTATNRIGTRHQEYTLGQADVPSSPYGV

TRKTVLKPKEILLSIEMPYSRKDEYFSAFKQASRREDDIAIVTCGMRVLFQEGSNRVQEIKLSYGGMAPTTVLA

LPSIPATTTNSTTSRFCLHKANGIYPVPKDHAMFYICANRRTFRMSCPEGLIFDYLCKCCNWP
ESEASSSVSISHSASATGNICIEEIDVDGKFIRLKNTSEQDQPMGGWEMIRKIGDISVSYKYTSRYVLKAGQIV
GALGVSISAPGGAIASVPNWTLRGTQLMNGTSMSSPNACGGIALVLSGLKANDVHYTVHSVRRALENTAVKA
YLPVGSHNLKAAAKAKLGYDPVELDPEEMCRMATEEPQTLATYSVSDAVATYYMYMKYVHPFIFALCTIIPME
CDEMLQGFAVAVKMGATKASFDETVAIHPTSAEELVTLR

VIACCQSCCNYSFHMFTIWLGYVNSTLNPFIYPLCNENFKKTFKKIFHIHS
QLQHATLADLVNYHKEEPITSLGKELLLYPCGQQGPEPDYLDLFE
SACLPSKKHSEYQILINELERPNTQMRTTHHILQRSVSLEGSCQKVSCCLSSLKYKCPKAETSQLHIQFTPGN

EKNNIELKWRETTKLYSESGDFIEFNCKEGYERDPTSSALRVQCMEGKLAYPKCKKKGK
NAIQCLEEYLRYKSVYGKDL
FDFMSKPSPDAIQAAIEQLDLLGAVDRKEDQVTLTPLGRKMAAFPLEPKFSKTILMSHKFHCTEEILTIVSLLSV
LPALDWQLPSHSGPYELRIEEQPKSHHRAHYETEGSRGAVKASAGGHPVVQLHGYLESEPLTLQLFIGTADD

ALTVQQLTAAQQQQYALAAAQQPHIAGVFSAGLAPAAFVPNPYIISAAPPGTDPYTAAGLAAAATLAGPAVVPP

LCAVKFLRLEDFLDNQRHGMCLYLEPQGTLFAEVTFFNPVIERRPKLQRQKKIFSKQQGKTFLRAPQMNIN

YGHWRKPLSTEDLKKKFRSNASSVLSVEATEGIIETVDNLEKVEDCFVLSAFLKRDTVNQSTFKNVLTLKNN
GQANFISLYNSMDSWMIVPNIKQNHYTVHGLQSGTRYIFLVKAINQAGSRNSEPARLKTNSQPFKLDPKMA

ATGCGKTTQVPQYILDEYIKNGKAAECNIVVTQPRRISAVSVAERVSYERGEEPGQSCGYSVRFESVLPRPH

IVVDETIKQSRRLIIVLVPESSSCSLLEDIFEQQLAVYNALIRDGIKVILIELDKIKDYANMPESIKYIKQKHGAIR
NIISQNKERNERSVSDNKALMVVTEDPLLYISPPPCQPLINRTESLRLNHELRGWVHRHEVERTRSKRMSE
IMHC
LADVELVVTKLHNETKEEIWMGFMNENTPALFKWSDGSKVVFTYWDQNEPKIPFNNTPNCVSYSGKLGR
QLNERQERIVSEAKKLEKDLIDWTDGITKEVQDIVEKYPLEIKPKNQPLAAIDSENVENDKLPPPLQPQVYAG
GYQDVRPGESSA
LNAINKQADELRCIMESPSSPSYAAQFDTAFQCTPLEKLIACQEKNPVSGLIEYSQYTYQRCEFALLEQSGP
SQTALNSTNQSVTQVTVVNIEGMTCNSCVQSIEGVISQKPGVKSICVSLVNHNGTIKYDPLLTCPEDLKSAIE
LMLDSPGFVEYIVDRSVEPDKASKDAKYELVKALVNSKTIAEIFGNQYYLRLRAYLHEGPYYVKAVSTTAVEGA
QSAQSMVPCTSTQHIVGLPSGPQPVHSGSVFNPSHYPNSTSSQQSVLSQYGGRKILVCSVDNCYCSSVS
DSVYESMAHLSTDLLMKCSLNPGCDDELYESMASFAPDATDDLYVEMLQSKPKAPVARDQLSLNIKDSM

QTLKFLCESKCPLDVKDKSGETALHVAARYGHVDVVQYLCSIGSNPNFQDKEEETPLHCAAWHGYYSVAK
SSNPSRKACSTSSKIRKLSTCKQQ
EENKLANISQEQILAKDSDGDTFLHIAVAQGRRALSYVLARKMAALHMLDIKEHNGQSAFQVAVAANQHLI

FPSSKVQSGSSAHSKTIKTSSSSSSTSFNKGGSTFETKSLKSRAVTEELGGLQHDESGEDTPDFRARSLRTG

SLGIKAAVLTGAGYSNQTKHMTQNEQQDVIKLFRKGDLNLLFSTSVAEEGLDIPECNIVVRYGLMTNEIAMV
QASLHIAVSAATFSTDSAIVIGTASSLLGLIVTAFVSHRLWRRLHALGMTSPKGNSV
RKWEKLQMTASERRKIMCSVTFHIIAITCVVWSLYVLIDRTAEEIKQGQTTGILEWPFWTKLVVVAIGFTGGLL

GMEPGGGSETMRRQAPAGSPESGDVQPPLHTGSPEPGASLLSTEWATMKRREETERSSCHGLPPTPKVH
CPPQWTGKTCQLDANECEGKPCVNANSCRNLIGSYYCDCIAGWSGQNCDININDCLGQCQNGGSC
VFCPEQLLRVILAHIHYMPDHWQANAHRYETRDEFSQLFQYKAAFILEELLSPIVTPLILIFCLRPKSLEIIDFF
FLFDLAVIVCKRRGDNYEMKEIIDLQKYKITNNPTTDKENKKWSYGFYLIHIQGQNGLEFYCKTKDLKKKWL

FSVDLSKERVVLGAVGADDWAGGLIELRGNQALETFIRSPSASEQIKDAYLGYAVKSMQHQNRTFYAAGA
FKPSNGFLETKVYIAGLPRKVGDTLIKPLNPRLDGCIRAWNLMNQGSSGIKEVIQEKQSKHCLVTVGKGSY
TLTGHSGKVLSAKFLLDNARIVSGSHDRTLKLWDLRSKVCIKTVFAGSSCNDIVCTEQCVMSGHFDKKIRF

KMVDSKHGQLIKNFTILQGPPGPRGPKGERGPQGPFGLTGLKGQKGEKGEPGPPGPAGEKGPVGPIGP
VSTDGCTLQMLNTTITHCQCNHLTSFAVLMAFYELEDWTLDVITKVGLVISLLCLLLAIITFLFCRALKGPRTTI
GKTAKPGQFPWQVMLITEDERAGAGALLYDNWVLTAAHVVISQKNPSTLTIKMGFLNKSSVHYQQAWAEE
GISYEVMFLGTDQGFLHKAVALASGAHIIEEIQLFKDPEPVQNLLFSPGKGILYVGYSKGVLQVPLANCSVY

VRDLSIEEDVLAAMPGLQLEEEITDSYLLISSAGDPADSWEIPVYRLNMSVQLLSKSVDETVSSPTRTIVEEISE

RGQEAAHNAPGQSRQLEPTLGQAAGTQLSSTAHWRMVLHGPTLPPCCPAHKEPCALRTVRKPGPNRGRR
QEMEIFVKDHNSRTTLYGVSASDTVLNLKKKIEDRTGVSASQQRLTFNSKELQDDYTLAHYRIRSKSTVFLLLF

NPMKSRKRVCQPTYPKFSMVTQGVAATGPVNVSFWGKPWPQCEPINGKTLEVEEKEPCLNVPPCPEEEEEY
IYRHPRYRPPRGTYRHPGYGPHMVTSPGPGVSLPLSQASSSLSTTVTTLSSAVGSLTPSRTGVGGGPATINSA

PEVLENKYGYKLCLLERDILPGGAYTDDIVTAIQQSRRAIIILSPGYVNGPSIFELQAAVNCALEDNMLKLILIKYH
SKTRFRNLTAALSGMKDLMLRDLDKLFLASDCRHFNRTESLRLADELSGWVRRHQINRRKPLQARKPRANC

GCVNIPGSYYCTCPRGFVLLPDRKTCHELISCANNHTECSHGCLETSRGPVCFCPEGSVLRVDEKTCSGC
QIMSFSPQDLRRRLWVIFPGEEGLDYGGVARLVAEWRLSRGVEEQTQAFFEGFNEILPQQYLQYFDAKELE
LENKHGYKLCIYGRDMLPGEDAANALEMRIQKSRRLIIILTPQLIQCEEFGYEHQIALYNALIQNNIKVILLEME
FLAAFLLYGLFRLLGNKVCPSGAEEKSKKSE
QLKTADASLSEENQATGNYEAKAYSSLSKSYLYIDWASNYKLLHVGDYMSIKVYPHSHYIHKIHHYSYLILSKC
KPERWLQKDASHHPYASIPFGVGKRSCIGRRIAELEVHLALARILMQFEVKPEPDGGRVRPMTRTLLVPEKD

QPRSEDVLFISDNESFNPSLWEEQRKQRAQVAFECDEDKDEREAPPREGNLKRYPTPYPDELKNMVKTVQ
PEDYLNKCTPGMRKWWQLKSQNFDAVIFYKVGKFYELYHMDAVTGVNELGLVFMKGIWAHSGFPEIAFGF
LAYNKFSIKSDVWAFGVLLWEIATYGMSPYPGIDLSQVYELLEKDYRMERPEGCPEKVYELMRACWQWSPS
HTTTPIAATVISAPHGRARDSSGEHLAHVPLENFLQTLESSATSSHLAGEESETEAEINLSDFPWFHGTLSRIKA

CQDMKQPLSHYFINSSHNTYLIEDQFRGPSDITGYIRALKMGCRSVELDVWDGPDNEPVIYTGHTMTTQIV
GFGDIIVPGLLVAYCRRFDIQTRSSSVYYVSCTIAYAVGMVLTFVVLALMKMGQPALLYLVPCTLITSSLVAWRR

VIQHSSVTASPNELNVRSAAEAVATSVLTQMASQRTELSMPVQWHVIMAPQSQSAGR

STFQTTDRLYFVMEYVNGGDLMHHIQQVGKFKEPHAMFYAAEIAIGLFFLHNKGIIYRDLKLDNVMLDAEGH

SVKITGWGEVRTPQGQTQKYWVAANSWGKHWGEGGYFRIARGANECEIETFVVGVWGRVSMEDMRRW

CTARLPDGNSSTTVSPKEFCAEQVPATTIVCDSVTCANGTCHYSMRCVAGERGGRVAYSWTQEDSGVVVS
FEPVQTLLLSSKKGKRYLYAGSSSGVVQSPVAFCEKYSTCVDCVLARDPYCAWKSHEASCINILKEDEIERDV
PPGVSVAYESHCEDATRSVGGHSGVCSGVRINQLVSFTVTVQADACLEGPHSFALRVLGFTEEVRVELQTLC
VKVLKEYPSSECLLGYKIQYCPENNASFKMTHNTTDKVAKLLLTGEAYVISVVAYNSVGESPEATLRIPSLGEKT
VGPDAPKGVQVKATPGETIQEGDRLVLMCITESSNPPIFTYKWYKDSQWLHQRQEQKLEFVAKGDEHSGSY

LTVVTDLYELEESTPEIFDTTFPSRPVVSFSSRLIEVSESNLFFSNSVDVGLNIMSTTIFPSYRQPSEGISLDITSV
GYGRPSGAAAGGAGAAAVAAAAAAGAGGGGYGGSSSGYGVLSSPRQQGSSMMMGPGGGASLGKAAAA
SAAVKPSEVFVKKPSVDGHENGDLGNSVETPLEETAPKLADSAESLP
NALQDFVINFLSKRLIYFNKTDQSFVSGFLDGSEFNILRPQVSFRDIESFVYENNTFVVTDGRAVFREELSPT
EAVGQGCAEIGSQRYKLGVRLYYRVMESMLKSEEERLSVHNFSKLLNNNIFHTSLLACAVEVVMATYGRNAS
KGTEPIIVDPFDPRDENSRCCSIIDSFSMPMLDLDDDDDREESLMMNRLASVRKCNERQNSPVTSPGSSI
AMMAQSQSLGKGPGPRTDGGAPFGPQGHRDMPFSPDEMVPPSLNSQSGPVGPDHLDHMTPEQVAWL
VFYAENDDTYVNKTMMLYVKRKPEVTIQSTSKGISCYSDSYPAPSWTWRKCPGPDSSNCTEEITEGIWSDTH

GGKGPEGSAVPVPIEPQICQATCAIMKLSFDEEYRRAMNELGGLQAVAELLQVDYEMHKMTSDPLNLALR
HKASQTSNQGCRTRTVVPGCVSFEPVFHGDKTVFFSSCNDMEVTGNCTVVNCNETSKATGKSCYEAFQ
SSVENSRHMSPAGARGRLSGIGGPRDCNSFLRHARETPDSYRDSPHSERYVSAMTTPARMSPVDFHTPS
KKSSITLKEWQQLQTQAKLVLEENGVLMKQLEIQQAKAKESYNQHLQEVSKLTKQLMVLEAKKQSQEEISE
QYILGTISGTTTCCLGYQAHSAILCPGTTHFSASPFEPDSQATMHLDTVCFSIVHFSTTHPGAAYSCTEICGT
ILGIFDLQKDDALYILVGQQGEDACPSMENGEPVPLIIAAGGGGRAYRAKTDTFHPERLENDSSVPGLNG
TVVIGMTRIPVIENPQYFRQGHNCHKPDTYVQHIKRRDIVLKRELGEGAFGKVFLAECYNLSPTKDKMLVA
GDVVCTPNGIRKKFNGKQWRRLCSREGCMKESQRRGYCSRHLSMRTKELESLSEGRASGLREGSAEFD
AASQGAPAVSEEEDDDESEEEEPSSDTDIEEELQQSQEGGEEEEEDAAMQEAAGEETAGDQEDQCMEL
SLQNPGIETSPRQHVSFRPDASVPSRTNSFNNHQQQQVPVSIRQVPAGKPDPSITSPNTITAVTSAHILQP
NPVNGKAICTCPSGYTGPACNQDIDECSLGANPCEHAGKCINTQGSFQCQCLQGYSGPRCEIDVNEC
VRLMFSNCYKYNPADHEVVAMARKLQDVFEMRFAKMPDEPEEPMIPASSPVVVPPPTKVAPPSSSDSSSD
GVTVYFHAILSKDFKLDPNQHKVFIRAQGISGYLNWKDNICELTCTKDLGEHGLLIEGWVTISKDNMNKQF
SPKRYKFRDDFEKLKMAEFHKEGHYGKEEMDEQDKKDKAKGRKDSEFDDEPKFMSKVVATSSKNQDED
ERTRYSRSYADSRTRESSDSEDEYRRTHSKSSDSRRTSSHSSSYRDSRTSSYSKSDRDSKTESSYAETERR
TSKNPFDEGEIQLGDKVLVVGQRTGTVRFYGTTKFAAGFWCGIELNKPHGKNDGSVGGVQYFSCLPRYG
SIAGPYRAAADNVSDWTYPLPPPNISLTNQGHSDRLTTSWGAAAGGRDGYTLTLYHAGSGAVAARVSVGN
QCHSEGCPRHYHFPCAAASGSFQSMKTLRLLCPEHVAEAVQMEDSRCMVCDGLGELRDLLFCTSCGL
HLAREKGRSCTKSQVKKSSDSRKRTRVKKSSTRLEASRREIGGKAPENIVRNRVAGDLPDKHASHELCRIA
ADLKNFATALEEVCVETIEAGFMTKDLAACIKGLPNVKRSDYLNTFEFMEKLAENLSLKLNSQPRL
KAGSMTGRSDDGLSTTSSPEPGTSELTGEGLKSSETDFRNESDPSLGHDNEEEEEEEEEEEEELENESRPES
TPNSRFSATSCGSSEKSSAASQHSSQMSSDSSRSSSPSVDTSTDSQASEEMQALSEECSNTPEVHTPLPISQ
TKKLVETTTNMNASFVSFLKTAAENAEVNSIIQLRQLKAFKSVLSDIKKAHLALEEGLNQEIQALDVKIKEKNTK
SECVCDDSCEKEALCKWLLKKEGKDKNGMPVSQIPVVNPESEKPKSAVKIWLHPSRQLPEESATTTVKLDSS
ETFPDLSIFQNLRIIRGRVLHNGAYSLTLQNLSIVSLGLRSLQEISSGMVLVHQNPNLCFLQKVPWKAIFRNR
SSGSSQDAPVGATDTEDGDSYLSDRESCISEKTDLMVNCTEVVMGCENDKSLSKSIVHNIEDKIFGKTYK
ILGSSGPALPSQPVAHNPPSSAAKEPSGAPAPPSASSGLLVPMTVNTPSSPTPSFSEAKPSQPLLNGPPQFSST
PSLQPSVTSPSRPSSSRLIEHEINRTDEVEALTFPPTSGKSFIMGADEALESELGLGELAGLTVANEADSLTYDI
ISGLSSNTGSSTLTSDREINVHSPTDSKLWYKDSSSQSGSVGINLNSVQDKSNRIGSTEASEPSSFTNPVYSY
PAANGAIRFHCDPTFWAKNVVNLGNLVIENKPTPGYTPNVVVGQVPPGTNHINKAPGQINLAQLRLQHM
IKKKRGRPKKQPLLTVETIHEGTSTSPVSPINREFPGTKKRKRRRNLAKLAQMVPGEDKSTSELKFHKKVGKL
NIFNLGSLVIEDKETPPHMPKSPVMEPNLQPQSGLPSNQLSKFPTQISLAQLRLQHMQQQVMAHRQQAQ
RHTDLERYFSISPEDGYIKTIKALDREEIAWHNISVFASEVYNRHQEAKVPVAIKVLDVNDNAPKFASTYEAFI
VGRLSPCLPGKSPGPEFEDRLAEGAPPAQAESHRSGVKVEQETSRQ
IHSLSRELLIKEGTSKFATLEMNPKRAQKRQQQQREMGVMQGTIPYLGTFLTDLVMLDTAMKDFLDGGLIN
LEFAAKRKKAEVQSGIIRNDSLWDKLFFNKIQASLGGCVRMIVTGAAPASPTVLGFLRAALGCQIYEGYGQT
LLDMQSLTDQEKSPSPIPVISSPSHEEFNTTVHEENSLQIHTSILQVSIPAFPPAPESSEATTEKVEAQPKPEVSS
SFPIPSQPEFSHDIFISTQSLEEKPSGEEKGGVLLGSCLPADVSETTDALPKIPTGDPELSCKLVLSASECSQPT
EVTTPLQRVDLFMGQFNSVLLTSISVFTKGNLTVANLGTSEGRFMQIVISRSEATSPHVNFPLDSHPVSPQVI
YTNGNIPQSMMVSNRNSHNMELYNINLPGIPVSWSQSSSVPPQSPPGSGHEIPTWQPNIPVRSNSFNN
SGASPPRGSRAKGGALGTETPETPTPTPVQPSQARQCQTWDAAAPRCLSPHKCGGEGRLLASLRSRVERV
LDIASADVLATPQETYFKSQAEWREEVKKHFEKIKSEGTCIHRLDEELIRRRREELRHALDIREHYERKLERA
EIKKSHLERECSSRLIASAPGISTANHCNRNEHSLAIPINKNEKILRSPISPLSDASKHKYSTDDLTSSNKPNG
VVCEEEFKYAMLALNCICPATSTLITLLVHTSRGQTAPWEAFRHLTGTKQTLTREGQQSPEQWQKMYGRCS
HSDVMMTQSDPLMSQASTAVSAQNSRRSIMLRSDPMMSFAAQSNQGSLVNQNLLHHQHQSQNSSLG

KLAKTLEVGNFLRIYSLHTKQARVENQPNVSYVEFHLHGGTCYGRGISILPENSTDVKELKAVMDSVNLTEFQ

APSRGRDVQCDEQTSCPDSSTCCRLSTGAWGCCPLVEAVCCQDHVHCCPKGYTCDPAEGSCQPPAVS
KGVDIVGESILVCKSSERSKGIALLQGSLKPVHSCLLRGEGEGVTSSMIRAILEIIVGGVASTPDDVRTYASCT
FGLSHHGNYIFWTDYMNGSIFQLDLMMRNVTLLRSERPPLFGLQIYDRYKQQGDNACRVNNGGCSTLC
KEAEAPAAIAERVPKRKRETESENEETAVAPSSPATPLAEEQSRKEKKKKKKEKKAKQAAESEGELMEVVGSWM
TSDVPKSPPKSKRLLTISRLIMEQDSQSTELSETPDSSQEPVVSSAFRTPVQKPHVQCVSSRSPKPLKARWNT
QDEPVETVSPSLLSFSSTCPPEAASRGTAAAMFQRVPDVDKHPDASHDLPCGQPSGAFQAHLCPFSPALPIS

HSNLPEVVVELLMTLHEPVNTTAGEGADLSKFTGELDPAPNPPHFPSYVIMATLDYISSCHKTKLKSIVAVLSK

LVAEHGEFDELLLSFSEWINQFLGELQDTSEINTADQHPAVCRNQDHSMEVESKKQELQSLREHVEKLCS
VFSRGQEQHWWIGLHTYENDGRFKWSDGSLLNFISWAPGKPRPISKEKKCVYVTASREDWGDQKCLTAL

VQILLNEKLLPFPHSEETVSLYTDLKNYYKDILQNCHFSEECELSKTNNTSTDEL

PEGAIPEEQQPQPRNNSDNFDLFSFDLAPMVTIRSESSSHSVDYSPADDFFLNSDSSEGQPPIVQKELDETI
YDKDEVDLLIVIGSSLKVRPVALIPSSIPHEVPQILINREPLPHLHFDVELLGDCDVIINELCQRLGGEYTKLCY
SSGFQRMTVINFECNETAGNEGKGAPVFTGEVDCTYFFTWDTKYACVQEKEDLLCRVADKKKRYDLSPLI

AIQEMTEDMKTVLQLVTHECKLIEFCIYVVVQFASVKHVARMGPETPSVAIMLKSSSKEKYPLFTFVKGHSRD
SFNPCVEEITNITYRCMELNLSGVPAEIPPSTQNLDLSFNPLESLTSNYFSAVPALRFLDLTRCHIQTIEDYAF
CIVPTPCYKPKEKLPVEAKLPWYKQAQELEDGAMVSEEPSFIPEAWSACSVTCGVGSQVRPVKCQVLLSFSQ

LRPSKICLQKQLSLEQTLPSPDTQVAAGTVSKLQPTAQDIQNNHYNSLQHHKENKVNGASLMFRSSTFPVQ
YGKIRSKVELEVRDLPEELSLSFNATCLNDEVIQGLKSCVGLKIGDTVSFSIEAKVRGCPQQQEKSFTIKPVGF
MDDPMKVYKDRQVMNMWSEQEKETFREKFMQHPKNFGVIASFLDRKTVADCVLYYYLTKKSENYKNLVR

QPELSEDADTRDKRKPAKSAPNTVNPCATSLESFLTKSREELLEYAAKITLDFNTAHNKVLLSDRNTKMSVSE
YLEWLNNHMDS

ERHTNSQCLPAFWYNPSSLSKDCSFGQSYLNSTGYRKVVSNNCTDGVREQYTAKPQQCPGKAPHGLRIIT
LVIPHRVFAPWCLKDRCAEILLELIPRCLSAAHGTLGTHPFSRLDVLIVPSNFSSLGMASPHIIFLSQSVLTGG
SAYGGMPFHPGTTKSSSGMKHGISNDMGEQDGESCNKHDDKPDMKHNTTVEITERRESRVCSGDNKE
REGQQEPSQAEQNEVPAEGTQTQPQHLDANTSSPDRSDDIEMDSKVLTVSNNSEKGTLQPSVNTTVAQPL
QDEEEGRSRQDDLQQVEQPAQRDDAGAEGTEAAGAAAAQESTLTSQPFVLPMGVISRNEDYPRTCQMC

CFSFGIGEGASTALVKGIARAAGGSAEFITGQDRMQPKALQSLKRALQPAVTGISLSWDLPPGMEAALLGR

TESPVIDRCRSPPPVQVLENEYRATWEEPQFSDNSGAALTITKSHSPGDLFLKGETTVQYTATDPSGNNRTC
RQKILRLVSQWVLLYGPLLQADHSAINLLQHLSDFVSRDPRLCNLLRDPTQDRRRNRTLENGGGSTSPQL
EQKVQTAPPGFHLIFLPYADDKRKIDFTEKVPANDEQVDKMKEIIQKLRFKYRNDSFENPVLQQHFRNLEAL
PAYKKDGYFSDVECIKTLNDKTSPLTILKTFAIAMVAFFFFASFLIFGVYYISRRIRYAFFPSCKPPSNIECLGG
SRYFQPFGFGPRGCVGKFIAMVMMKAILVTLLRRYRVHTPKGRGLKNIQKSNDLSMHPNERQPSLEMFF

EVPLYNDVMLPSSSEKLKNINLAMSPLPVSEITKPLRSNADPALNREVVSKLEPNTEPLELAFSSRRRQEQPAS

QTQGSQTSHPRVIRISHQTVEPVVMMHMNIQDSGSQAPGGGSGGTVSMGTAGPAGHGPGIGGAAHVP

RQIAMWHQLHM
VSQFPFALLGWTGSRHFERELRRFAGHERKMVLNSHTLYDTRRNLFLPAASEEEIFQHLGLEYVPPAQRN

SKEPRETKPFSLKALIRWLHISKPAQHVMSLLGCRYPSDLNSFSRSHLPGPWDPSQAGTRMKLPQPQTWD

QQLKFLQQTMQGMQQQLLSIQEELNNKKSELEQAKEEQTHTQARLKVLQEQLKSAKELAEINGHDESQA

EGTTLRVEDLVKQYFQTAEKKVQLSLLTERGMGEAVQEFVDKEEKDAIEELVKFQLEKTQRFLKERHTDAEE

IRNTNNPGPTSSSPKIALKGIAQRTFSRALTNKKSSPKGSEKEKQKEKEKDKSKDIAKRISVTEKQEIKEEPKEE
LVCNADGDPQPTISWVTPRRRLITTKSNGRATVLGDGMLEVRFAQDQDTGIYVCIASNAAGNDTYSASLT
NIEDGKLMTCISIQNSCSCLEGPDSQLYLSGWDTYADAETAEMIAHRTGGMKISATVIFNPKSPATSESPVTT

PSRCCTPPIKLLEEDLWPRLNSSEPVPSLIHSNAILDKYSELSSEIAKSSIPLAVASSRDLDPQEENCSKTLALR
NCVRFPSTNLDIPTHFVKTILSQGHLTPLPLYVSPVFWAYDYTLRVYPVPDLIVTADKYDPFTVTNTDCLCINP
VKFKAQNQLNKKCSAVKHTRIKGIPKLDDANDAGSKNSTDCTLILTEGDSAKTLAVSGLGVVGRDRYGVFP
RREMSTKSTQFKGTGNEFNKLFKPLKKARSCASPVNKIAKCKLTLQASCTADTSTELMCAEAILDISKTIMPPQ

FLFSSSNFIYLLFTTDNSRSNNGFKIHYESVTVNTYSCLDPGIPLHGRRYGHDFSIGSTVSFSCDPGYRLSH

VEEMRQKLGPYAGEVEDHLSFLEKDVRDMVASFFSTIKQIEN
MAQYQQSRATFYALSHAVTNFYEDKKTVTQEIMDVANFMESMIGNECSGNDELTYLTLRAIGNMGKAMEV
RRAEADIPPEDVEEEPAPTTPPDSPEVPRPLPAARTGLVYDERMTEHYNMWDSQHPELPQRVSRIFQRHGE
TPNSPLCTESFKKESCKPTNKKNLGTKSTMKDQSLVFHSKIKSSGYTTAPQMTMFSPKTNLKKNNKISEWKSS
KKQWEDTEKAQRSLTEIERKAQANEQRYTKLKEKYSELVQNHADLLRKNAEVTKQVTARQAQGDVEREKK
YWPKHRTVSKALIAYWTNFARTGDPNKGESTVPIKWIPYTWEESYYLEINSDINYDSVKQSLRASYMQFWN
MIHNNILEIEGRYYLQCCGIPQGSILSTLLCSLCYGDMENKLFCGIQRDGVLIRLIDDFLLLTPHLTLAKTFLRT
SVHWRRGKLVLPAKNTTHLKTYSAGRKLILEIAPRSDNDFGRYNCTATNRIGTRHQEYTLGQADVPSSPYGV

RKTVLKPKEILLSIEMPYSRKDEYFSAFKQASRREDDIAIVTCGMRVLFQEGSNRVQEIKLSYGGMAPTTVLA

LPSIPATTTNSTTSRFCLHKANGIYPVPKDHAMFYICANRRTFRMSCPEGLIFDYLCKCCNWP
ESEASSSVSISHSASATGNICIEEIDVDGKFIRLKNTSEQDQPMGGWEMIRKIGDISVSYKYTSRYVLKAGQIV
ALGVSISAPGGAIASVPNWTLRGTQLMNGTSMSSPNACGGIALVLSGLKANDVHYTVHSVRRALENTAVKA
LPVGSHNLKAAAKAKLGYDPVELDPEEMCRMATEEPQTLATYSVSDAVATYYMYMKYVHPFIFALCTIIPME
DEMLQGFAVAVKMGATKASFDETVAIHPTSAEELVTLR

VIACCQSCCNYSFHMFTIWLGYVNSTLNPFIYPLCNENFKKTFKKIFHIHS
QLQHATLADLVNYHKEEPITSLGKELLLYPCGQQGPEPDYLDLFE
ACLPSKKHSEYQILINELERPNTQMRTTHHILQRSVSLEGSCQKVSCCLSSLKYKCPKAETSQLHIQFTPGN

EKNNIELKWRETTKLYSESGDFIEFNCKEGYERDPTSSALRVQCMEGKLAYPKCKKKGK
NAIQCLEEYLRYKSVYGKDL
FDFMSKPSPDAIQAAIEQLDLLGAVDRKEDQVTLTPLGRKMAAFPLEPKFSKTILMSHKFHCTEEILTIVSLLSV
LPALDWQLPSHSGPYELRIEEQPKSHHRAHYETEGSRGAVKASAGGHPVVQLHGYLESEPLTLQLFIGTAD

ALTVQQLTAAQQQQYALAAAQQPHIAGVFSAGLAPAAFVPNPYIISAAPPGTDPYTAAGLAAAATLAGPAVVPP

CAVKFLRLEDFLDNQRHGMCLYLEPQGTLFAEVTFFNPVIERRPKLQRQKKIFSKQQGKTFLRAPQMNINI

GHWRKPLSTEDLKKKFRSNASSVLSVEATEGIIETVDNLEKVEDCFVLSAFLKRDTVNQSTFKNVLTLKNN
QANFISLYNSMDSWMIVPNIKQNHYTVHGLQSGTRYIFLVKAINQAGSRNSEPARLKTNSQPFKLDPKMA
ATGCGKTTQVPQYILDEYIKNGKAAECNIVVTQPRRISAVSVAERVSYERGEEPGQSCGYSVRFESVLPRPH

IVVDETIKQSRRLIIVLVPESSSCSLLEDIFEQQLAVYNALIRDGIKVILIELDKIKDYANMPESIKYIKQKHGAIRV
NIISQNKERNERSVSDNKALMVVTEDPLLYISPPPCQPLINRTESLRLNHELRGWVHRHEVERTRSKRMSE
IMHC
LADVELVVTKLHNETKEEIWMGFMNENTPALFKWSDGSKVVFTYWDQNEPKIPFNNTPNCVSYSGKLGRV
QLNERQERIVSEAKKLEKDLIDWTDGITKEVQDIVEKYPLEIKPKNQPLAAIDSENVENDKLPPPLQPQVYAG
GYQDVRPGESSA
LNAINKQADELRCIMESPSSPSYAAQFDTAFQCTPLEKLIACQEKNPVSGLIEYSQYTYQRCEFALLEQSGP
SQTALNSTNQSVTQVTVVNIEGMTCNSCVQSIEGVISQKPGVKSICVSLVNHNGTIKYDPLLTCPEDLKSAIE
LMLDSPGFVEYIVDRSVEPDKASKDAKYELVKALVNSKTIAEIFGNQYYLRLRAYLHEGPYYVKAVSTTAVEGA
QSAQSMVPCTSTQHIVGLPSGPQPVHSGSVFNPSHYPNSTSSQQSVLSQYGGRKILVCSVDNCYCSSVS
DDSVYESMAHLSTDLLMKCSLNPGCDDELYESMASFAPDATDDLYVEMLQSKPKAPVARDQLSLNIKDSM

QTLKFLCESKCPLDVKDKSGETALHVAARYGHVDVVQYLCSIGSNPNFQDKEEETPLHCAAWHGYYSVAKA
SSNPSRKACSTSSKIRKLSTCKQQ
EENKLANISQEQILAKDSDGDTFLHIAVAQGRRALSYVLARKMAALHMLDIKEHNGQSAFQVAVAANQHLI

FPSSKVQSGSSAHSKTIKTSSSSSSTSFNKGGSTFETKSLKSRAVTEELGGLQHDESGEDTPDFRARSLRTG

LGIKAAVLTGAGYSNQTKHMTQNEQQDVIKLFRKGDLNLLFSTSVAEEGLDIPECNIVVRYGLMTNEIAMV
ASLHIAVSAATFSTDSAIVIGTASSLLGLIVTAFVSHRLWRRLHALGMTSPKGNSV
RKWEKLQMTASERRKIMCSVTFHIIAITCVVWSLYVLIDRTAEEIKQGQTTGILEWPFWTKLVVVAIGFTGGLL

GMEPGGGSETMRRQAPAGSPESGDVQPPLHTGSPEPGASLLSTEWATMKRREETERSSCHGLPPTPKVH
CPPQWTGKTCQLDANECEGKPCVNANSCRNLIGSYYCDCIAGWSGQNCDININDCLGQCQNGGSC
VFCPEQLLRVILAHIHYMPDHWQANAHRYETRDEFSQLFQYKAAFILEELLSPIVTPLILIFCLRPKSLEIIDFF
FLFDLAVIVCKRRGDNYEMKEIIDLQKYKITNNPTTDKENKKWSYGFYLIHIQGQNGLEFYCKTKDLKKKWL

FSVDLSKERVVLGAVGADDWAGGLIELRGNQALETFIRSPSASEQIKDAYLGYAVKSMQHQNRTFYAAGA
FKPSNGFLETKVYIAGLPRKVGDTLIKPLNPRLDGCIRAWNLMNQGSSGIKEVIQEKQSKHCLVTVGKGSY
TLTGHSGKVLSAKFLLDNARIVSGSHDRTLKLWDLRSKVCIKTVFAGSSCNDIVCTEQCVMSGHFDKKIRF

KMVDSKHGQLIKNFTILQGPPGPRGPKGERGPQGPFGLTGLKGQKGEKGEPGPPGPAGEKGPVGPIGP
STDGCTLQMLNTTITHCQCNHLTSFAVLMAFYELEDWTLDVITKVGLVISLLCLLLAIITFLFCRALKGPRTTI
GKTAKPGQFPWQVMLITEDERAGAGALLYDNWVLTAAHVVISQKNPSTLTIKMGFLNKSSVHYQQAWAEE
GISYEVMFLGTDQGFLHKAVALASGAHIIEEIQLFKDPEPVQNLLFSPGKGILYVGYSKGVLQVPLANCSVY

VRDLSIEEDVLAAMPGLQLEEEITDSYLLISSAGDPADSWEIPVYRLNMSVQLLSKSVDETVSSPTRTIVEEISE

GQEAAHNAPGQSRQLEPTLGQAAGTQLSSTAHWRMVLHGPTLPPCCPAHKEPCALRTVRKPGPNRGRR
EMEIFVKDHNSRTTLYGVSASDTVLNLKKKIEDRTGVSASQQRLTFNSKELQDDYTLAHYRIRSKSTVFLLL

PMKSRKRVCQPTYPKFSMVTQGVAATGPVNVSFWGKPWPQCEPINGKTLEVEEKEPCLNVPPCPEEEEE
YRHPRYRPPRGTYRHPGYGPHMVTSPGPGVSLPLSQASSSLSTTVTTLSSAVGSLTPSRTGVGGGPATINS

EVLENKYGYKLCLLERDILPGGAYTDDIVTAIQQSRRAIIILSPGYVNGPSIFELQAAVNCALEDNMLKLILIKY
KTRFRNLTAALSGMKDLMLRDLDKLFLASDCRHFNRTESLRLADELSGWVRRHQINRRKPLQARKPRAN

CVNIPGSYYCTCPRGFVLLPDRKTCHELISCANNHTECSHGCLETSRGPVCFCPEGSVLRVDEKTCSGC
IMSFSPQDLRRRLWVIFPGEEGLDYGGVARLVAEWRLSRGVEEQTQAFFEGFNEILPQQYLQYFDAKELE
ENKHGYKLCIYGRDMLPGEDAANALEMRIQKSRRLIIILTPQLIQCEEFGYEHQIALYNALIQNNIKVILLEME
LAAFLLYGLFRLLGNKVCPSGAEEKSKKSE
QLKTADASLSEENQATGNYEAKAYSSLSKSYLYIDWASNYKLLHVGDYMSIKVYPHSHYIHKIHHYSYLILSK
KPERWLQKDASHHPYASIPFGVGKRSCIGRRIAELEVHLALARILMQFEVKPEPDGGRVRPMTRTLLVPEKD

QPRSEDVLFISDNESFNPSLWEEQRKQRAQVAFECDEDKDEREAPPREGNLKRYPTPYPDELKNMVKTV
EDYLNKCTPGMRKWWQLKSQNFDAVIFYKVGKFYELYHMDAVTGVNELGLVFMKGIWAHSGFPEIAFG

LAYNKFSIKSDVWAFGVLLWEIATYGMSPYPGIDLSQVYELLEKDYRMERPEGCPEKVYELMRACWQWSPS
HTTTPIAATVISAPHGRARDSSGEHLAHVPLENFLQTLESSATSSHLAGEESETEAEINLSDFPWFHGTLSRIK

CQDMKQPLSHYFINSSHNTYLIEDQFRGPSDITGYIRALKMGCRSVELDVWDGPDNEPVIYTGHTMTTQI
GFGDIIVPGLLVAYCRRFDIQTRSSSVYYVSCTIAYAVGMVLTFVVLALMKMGQPALLYLVPCTLITSSLVAWRR

VIQHSSVTASPNELNVRSAAEAVATSVLTQMASQRTELSMPVQWHVIMAPQSQSAGR

STFQTTDRLYFVMEYVNGGDLMHHIQQVGKFKEPHAMFYAAEIAIGLFFLHNKGIIYRDLKLDNVMLDAEG

SVKITGWGEVRTPQGQTQKYWVAANSWGKHWGEGGYFRIARGANECEIETFVVGVWGRVSMEDMRR

CTARLPDGNSSTTVSPKEFCAEQVPATTIVCDSVTCANGTCHYSMRCVAGERGGRVAYSWTQEDSGVVVS
FEPVQTLLLSSKKGKRYLYAGSSSGVVQSPVAFCEKYSTCVDCVLARDPYCAWKSHEASCINILKEDEIERD
PPGVSVAYESHCEDATRSVGGHSGVCSGVRINQLVSFTVTVQADACLEGPHSFALRVLGFTEEVRVELQTL
VKVLKEYPSSECLLGYKIQYCPENNASFKMTHNTTDKVAKLLLTGEAYVISVVAYNSVGESPEATLRIPSLGEKT
VGPDAPKGVQVKATPGETIQEGDRLVLMCITESSNPPIFTYKWYKDSQWLHQRQEQKLEFVAKGDEHSGS

QFPVPITETVVLTSGIRAKSSDITSILLESVFTTSENVSSVTIKHTGLLESPEFMPSESVLLLDKTPASVSTDKSSIN
SGAGGFQRFSGQNQHPSGATPTLNQLLTSPSPMMRSYGSSYPDYSNPSAPPQQPQPQAATAAPGSQQS

GIKAFNEFLVDCDFTQADYFGYNNLLSYGAATQPYPLPTHPQLVTVLFGSDQAVVCWRPPQPTIGTSPSAW
SQSDGTSTETDLSFPWILNVLDLKAFDFYKVIESFIKAEPSLTREMIKHLERCEHRIMESLAWQSDSSLFDLIK
PLTQRRKPLPDPLQIPHLSLPPVPPRLDLIQKGVARSPCASPTGSPKSSPCMVRKQDKPLPAPPPPLRDPPPP
KLQQEFYEEKRRKQEQVVVQQCSLQDMMVHQHGPRGVVRGPPPPYQMSPGEGWGPGGPESFADGM
HERKTFGSWHSSSILDIQDTVIGFSVECCANNSVGSSCAKRFITQQEAVFSLQDNVSFYATAGFCLPFIAVLL
RYAGMALTNLTFGDVVNKATLSSRRGCMQAIVAQLASENEELHQVVSSILRNLSWRADINSKKILREVGSVT
KPGNAISLLMEKTVSRDDNMDITKCQITSDNQVPREYKMNAHTISSVPDNGSEKGIQNCRAASETLGLDSN
SPKSPPSEMSPPVSSLTVSVPSVAVSPFMEEERPLLLVTPPRLREKYDHHIQQFNSFHHNPAHESNSPPPSP
NQKQLKSLCSTCEELKANLNGRVTAEEHVALVNALRSQLQQEQEKQSAEMEDLTERLASLQAQKKSLLLEK
TITELRTFVVPGDLHVTADMAFLLISMVLNIPPMVVTDTATGACATTWFHWYQLGLML
NSGAAGGNAAADNDPEMDGEDGVSFISPLGTLYTPALKADPVAHLPLSLVLSVVTSALVAALILAFSGIMIVYR
VKALKDPTLAARKDFQREAELLTNLQHEHIVKFYGVCGDGDPLIMVFEYMKHGDLNKFLRAHGPDAMILVD
DWDETSRDSEASSARGDSRPRLVAPADLSRFEFDECEAAVMLVSLGSSRSGTPSFSPVSTQSPFSPAPSPSP
GLEPLPCSSGGEDLEDEDEDDEDEEDEEEDEDPLPLAEGQGPPSPKEGEGPPFSPMQELFVLEIEALPLELS
VKSMRVMRPEPQTAVGPSHPGWLPAQAPPVDSLEIIEQHPLPVGGANAYQLDVDYSNQEPRCPPPPYPKH
LSNPCQNDATCLDQIGEFQCICMPGYEGVYCEINTDECASSPCLHNGNCIDKINEFHCECPTGFNGHL
SSSDSDSSSDDSEEERAQRLAELQEQLKAVHEQLAALSQPQQSKPKKKEKDKKEKKKEKHKKKEEVEENKK
PYKYFISRGKNGDYEFIYKPCPEGVFVNRCLFIPSKYLSSGDWHQYDDIVCIRPPDGLVRRLKSLVGWDD
RPGKWEGIVFLPSAKEKQRKAEEMEEDPYSERSKKEDKLASKRAEAGHRGFVPEKNFRVTTYKASQEKSTS
GKSSKSDRESKRTSESEVSKRCSPLNELGYRRGSSHSKPDSNVNSSHYKYTPTKTSAPKSDKSKSSFCCTES
FAPPSRVQRLTGSLDSLTEVSTGKMNHSFPGFRRSLSTTSASSQKEINRRNSFVKSRTSMLRRSWSNTTTTN
DTHNFTFVNLAPGSQYLLELASTAGPYGMSTGNVSNWTYPLAPPNVYLRNQGHSDRLSASWGAGAGERD
HYHGTCLDVPVTPRKRSGWQCPECKVCQTCRQPGADSMMLVCEACDKGYHTFCLKPAIESLPTDSWKC
NSLTAANGAVENYLLSSFGERRFEKTALKEEYQSPHGDSLRDTLRVDLLSSSSAERRKSPVGLLCSSKLQVH

SFSMDSELSRNRENGSKSLPDEKSLVLGKVIENVSLGAIQQYNDMLAEKQKRSSFMKRSSDQRDLCPQDL
SPENDNGDRRSRRFSMSERRFGQRAAKKLSTLQSVSQQQSSSSPPPPLLTPPPPLQPATGISDHTPWLMP
EKKIDIKLEKNDVKRDEGKEHAELKEDTVKSVKKAVSEQPENESMDQNVPATQQTESKNTSSEEKKSETNEE
SVKVVEPPKALLESPLRTWLAKSVHKAASTEEKANQEKAEVSFKSPFLDLLAPFHLPLNVNSWTLSSKNIENVN
RQTLFQTNNKPVEQCENEGLVCFHLCAKGQCWGPGPTQCVACEQFLRGQECVESCNLLDGAVREHV
RERKSNPLVHTREMPRSPKMEDVTTDVKPSSNSRASKLKRKRKTASDLQPVDFIKKRDEKDLKKHSEGVNQ
TPEIKAPEPLSSLKSMAERAAIGASIEDPVPSLHLSERDILISSTTSQPASSQPPIQLSEVNIPLSLGVCPLGPVPLT

ANNKDALRKTWNPKFTLRSHFDGIRGLAFHPVEPVLITASEDHTLKMWNLQKTAPAKKSASLDVEPIYTFRA
PLSYSSLLPMLEKKKRRRCGVCEPCLRKTNCEECSCCRKRKTSHRICKKRKCEELKKPPPPIMLPFEVLTE
QQQVYAQKHQQLQQMRMGQPAGSLPRQAGPQMLQQQPPRLISMQTMQRGNLNCGAFQAHQMRMA
VLDKKTIKTINKMKTLKRKNILNQILSCSSNIALKTKAQPPSSTGPATIDARLGKQINVSKRGTIYIGKKRGRKP
RVGPVGLPNPRILGPVQHTPTSHQQPPPRLINFQNHNPNSNGSILPAQQMRSPQNQNVPRLTIKPNPLQ
CENGQSNKAFITITADDKDDAANGPRFIFSLPPEIIHNPNFTLTDNRDNTAGIYVRRSGFSRQKQDLYLLPI

FEKRRKEFEVIAQIKLLQSACNNYSFMQEDSFVEWFHGVERLSEAESYSLSCEIEPLSESASNTLKAKKNTGI
CTAGCTFTTPGDWTSGHVGAPLPCNLIKLMDLEEMNYFASKGEGEICVKGPNVFKGYLKDGEKTAEALD
EVSEGTTPCSSVETQPCDSPLMSALQKSSSLGNLKKETSEVDKEPVQKPAETKAPMEGNKSGTLPPKSPSHS
RTEGNASSLKTNQGSENIQVEMDSTLQASELKPCSTESYSMQLRLSEDSQALLSENCEKPKREAVCATGEP
DHSLGADGYTLAVTGMKITKIPLNGPGCDHFKSCSQCLSAPPFMQCGWCSDRCVRSWECVKRMWTQ
HGNRQNHSSSSQPSATTVTAITPAPIQQPVKSMRVLKPELQTALAPTHPSWMPQPMQAIQPVPFSESPSTN
SQQPQGQGQDGGRREGSAVTLGTTEPVSQAWSIRSQGDNLVGSANLASTPSGYRKQDGGQEGKVKSQ
INLYMELSAIMLQLEVREKELIKREQAVEKKYPGTYKRHPVRPIIHPIAVEKLMKKKGVSHKPGSQTKRPDLLK
SSLLPSMSSSKKHKSEIQAQPHPTDCNKAVHINSESVLCNKPHSQTEPWPSLSRAHKDCRRTKLIFDDIPR
GNEVYHINLEDSTFFAEYEGKSFTYASFHAHKKFGVCLIGVRREDNKNILLNPGPRYIMSATDICFYINITKE
GSRALSNSISNMGLNDTNNLGSAKHQQQSPINQSMQTLSDSLSGSLYSASVNLPVMGHDKFPSDLDLDI

NVESTSSMELDGIFSNLSHQSYSQRCQQLSVTVLTDHQHLDDTALNTILNSSVPQQYRIRAKLRTFKPEKLY

IPWLEKTLARVRAPSRGQDVQCDEQTSCPDGNTCCRLSTGAWGCCPLVEAVCCPDHIHCCPKGYTCD
LLAASLKDKDQESERDEDRAQNGAVEACVRWLLENEFIQILDSGNGMKAEVYRPTHLGTATLSSSLSPAEA
LAIPGGRVCACADHQLLEENSTVCMINPGEALPQLCKPGEFQCNNKRCIQDRWRCDGDDDCLDASD
VPQSRPRSVAAWHDLYIPPCDTYKTRHLFSLSMYVAAFPHQYRVTALQASSI
MNEREEKDTWSEEDELFLDKKSPGKNSHSSSGLGSKKQRWTVQESEWIREGVKKFGEGNWKIICQKFPF
GSINSGAAPPANGGQADLGGAASLPRVENSGLASSEWDTELGGGAGGEDLSRSRQHRVMKRKLLVDD

NPVKIKDGLGGAWAFVLRDIIYTLIHYINSRPSLGRDMSMRSFSLCCDLLNLVCHAAVTYCKDALENHLHV

FSRPEEHHMLQGKAEDCILLFQEASQITSRRQKALSQLGVFLELHAAASSVLDHVRHTVETTGNMDKAQSE
PYICKRSNTTAVKPSLPPLPAPVPGGCARGWVPFINKCFSIKGYNKAEQVKWQEAKQACESQGAVLATISN

LSENDMANYSADLLMTTNEEDNLVEFDDGFRQENPGDLSEKTSSLTDLVEDDSSSPGVLKNVETRIPPTP
NSVKLSEITEKPSRMHKELEMHSAELPPTPLKISEDSSSPNRMTPPDPLVLLSEHPTEGKAENADPPSESKGN
RSSESAQNWEAVNSSPVEADKRFFINVCHKVLKKGAAVVCSDDAAVCSVDKNNKPKNLGTVMSSLKVGE

YDFASSPLHEENDLFSEEEKKRLKRFSTEEFVLL
EDLHNLLSLILTANPLRHLGPKAFYSLGSLQKLVAVEDNISSLADLPIGHLHTLKELNVAKNNVDSLKLPKYFSN
SVADLPIDECEGPKPASQRACYSGPCSGEAAESNPEEIELIYGGLQDFDELYDWEYEGFTECSESCGGGV

NKKEEVKIVTSTFLLPNRITIPPDPGTKKILVKTFSSAPNTQATPDLLKGQQDLTQQTNEETAKQILYNYLKEGST
FKDSLTVLVNFECNCTCQEHAEPDSPACHQGNGTFECGVCRCSPGRLGSHCECSEEEYSPTQQDNCSP
RNYRRRGKNQQQQQMPRSSQEEKDEKEKEKEKEAEKEEDKQEAENDKEELTKEKNDDTSGEDNDEKEA

PQKYSPHPQRFTYCSQVLGFQCFKRGIHYWEVELQRNNFCGIGICYGSMDRQGADSRLGRNSSSWCII

SDRKLTAEQGHNITFIVQLEEGDVQRMIIQVDFGDGIAVSYANLSSTEDGIKHVYQNVGIFRVTVQVENSLG
SQLCGTRLCHEIAHAWFGLAIGARDWTEEWISEGFATYLEDVFWAVAQQLPHDEAREQQELKALLRWRRI
EDVTLSCTAGVTEADCRLQDDLFLEKAMQLAKRHANALFDYAVTGDVKMLLAVQRHLTAVQDENGDNVLH
PSTQLYLHQTETLTSNLRMPLTLTSNTQVIGTAGNTYLPALFTTQSAGSGPKPFLFSLPQPLAGQQTQFVTVTQ
FYGTGLIAGHGFTSPERTPGIFILFDDDRFGFIWLELKSFSLYSRIKVSFQNAEAPSQEAFDEMLKNIQSMTT

GPEVIFPGQRCLIYAQLCGQPQPPDTAMGGVTLQYRIQDQTYKETLQFPLQPQDGDRLPVHRLAAKSLLLEL

EIHIVIKGSPCEISFTPMNGNFICIEDEAGVNCTLRCMDGYNFTAASSENYYCAYEDGIWKPPYSTEWPDC
KVRSTGNWIASLEDASLNNSYAIRAQDKVPYDIYRADHSFLTVVLPVNASVRDVLQSLACQEGAHGECMLV
ALDLIEPEQAEDLTQPKVEAMNHRLGDLVEQFKQLVYPPDYNPEGKAMKRKQGEGADQTEKKPRMEVSKD
QPFNSPYLSTSEEPTENCCIIESIVIEETDQTDFKDYKHSKQSTRDSGNYSNDDDTSGTKVLEETLEQETT
TPRRNVDKCQDN

PSDASTNQSSPEPCSPNDYCFDVDNDMANEFKLELVEKLFAIDTEAKNPFTTQETDLDLEMLAPYIPMDDD

LPSLPPELMQAVAHQITQQAMAAAASAATGQQVPGFQSAPTRVVIARPSTPSARPTHPGAPQAPGPTTPGIG

[illegible]

RQLSQHGNKAAVWEELIDSGKLPFMAMLRNLRNIIRAGVSERHHQRVLNRLQNKESVIRSRQFPFRFLAA

NEINLLTVALVNQEREKNSEKRSQGLKSEKEALLIGIISTFLHVHPFGANIEYLWSYMQQLDSRISANEIEMLLN

AKIDEEVRKFREARKENIEEEEEEVREAMTRARAHRSHEAALASGSSDDDLMDMGMSEQKTNDDSDAGTS

QTAPATTEMPKKSSKIASFIPKGGKLNSAKKEASAPSHSGIPKPGMKNTTGKSSSAPTSAKEGERSRSGKLGSG
VKGFTSDRFLYANRTPMYMTDSNDTSSNGTNVNTFSLDLKTILVSTAMGCFTFLGVVLFCFLLLFVWSRGKG

PEVAISCVPGSSDSLADEEESESHKLSIAATNCLINSCVCCGSCEDSREDNIVGLKTRHSPGKVIKASYSLVK

IHESYSNGLSVTLSTSNSPNSGMDQKSLKMSPSGQTKSQEAERTPTGNFKRTLEESNSGPALKKPRRSLSRN

GSFPRSGFSFKVFYPSNKTVEDR

LRGKMLNVREASHKQIMENAEINNIIKIVGLQYKKNYDDQESLKTLRYGKIMIMTDQDQDGSHIKGLLINFII

MGCEGKSFAANPFFYPQSMPSSHCESDSSPVDSDDSIEQEIRAFLALKAQTENLVTKPDNLSHSVQRPLSS

EEPLLCEKNHWWSHPLPNCDALCGGDVRGPSGTILSPGYPELYPNSLNCTWTVDVTHGKGVQFTFHTFH

ANPNLKSALKTCIKSEVASISVQKAAIQALRRMTITDEDRTVLLKAFQNANAPVDKRLAAYLMLMKDPSASDLN

LRLTERCHRIPARSAEEEELRMCHSPAYVQRVKATSSMKPRELHRQGDQYNSIYICASSYECARLAAGSAFI

CKRKNKEYLLENSPPTKSERQISVANKPTSICCIQYSGDGELLACGLADKSLLVFNSNLTGTPSVFSGHDGA

ELEDSFQRVSEQAQRKTQEQAEVLETLKRELTASKQELQALQSTLQSSLQSEAERSTRTAGLEQERDTLVVAVT

MVYQSLPDVPATFLD

TLATGIPEYGFLINPKKTMMNFPVDEDIPGCSKFKQLPYCSVIPWCGLLLDIQTLEVYCDYSSYACTSIRSSLA

VRIIELSQTIAKVSFNKPDSHGGVPIHHYQVDVKEVASETWKVVRSHGVQTMVVLSNLEPNTTYEIKVAAVNGI

EKTCRDLISREWNENLLQEACRLLADEMNLSPSAPGGMVEFRRMLTLSFFFKFYLIVLQKLNKPPNGNNNL

TVWAANAGVTASPPSDLIWKNQNSWGTGKDVKVILKNSQGEEVAQRTTVFKTTLREGEEEEEAEEEAVEAIE

AENIEVFAQGHGIIQVDKAYDYLIQNSSFTSNIGFTVTVGNNRGIYLRDPVQIAAPSDRGVGIEPVFPENTENT

PDEVLRKGSGTLCEALLMVQAYHANIIFPNKQEQEFNKLTEDGHVLDSETYVGGHVEALESGVFRSDIPCRF

EGKTRGFPKNASICKRESNSTLWRPHKTRDDFLGQVDVPLHQLPTENPNMERPYTFKDFVLHPRSHKSRV

DSILYNPPSRRDEVQSVRKKFISSEGDHLTLLNVYRAFKNVKGNKEWCRENFVNNRNMMLVSDVRAQLRI

DRLLRPHAFYQVHRITGKTVSTTSHETILSNTKVLEIPLLPENKMKAIIDCAGILKLRNSDIELRKGETDIGRKNTI

QYYGVPWGVYPANLFQQQATAANTTANQQAASQAQGQQQVLRAGASQRPLTPNQGQQGQQAESLAAAA

ATWGRLVRRAVPTVNHSGTFSPQASVPATVPVVDTCIPELARSASDSPVAKLDFEPEPPPAPPRASSLGGICD

HKKLKLSNDGLQMEKDESSLKKSHTPERFSSTGCYGAAGNIFIDSGCHYWEVMVGSSTWYAIGIAYKSAPK

IASVMFCTVGVLLRKLEAGIRGISHVIVDEIHERDINTDFLMVVLRDVVEAYPEIRVILMSATIDTSMFCEYFFN

VKGDFTEKSHSANTKFWKNVRYRMPPRCPTFSELHLLPTASNTFQTTER
NQQQKTQVVQSSTGENADSQLIAMPNTDPSTSRNSGNELQVYYASLRNYQDFFEAIHRRGDTFYVVSFRR

VRVRSCEEKLKYVCMKNGEVLNETSDKHCFPKEEWKRHGDFCYKIHNNKVSFGAQCNLTIMNRFEQEFI

SHEPRFKFQAMINGRRFPPAEAGSKKLAKQEAAANAMKILMREAENQGEDGMEGEESFVPDSSEAELPARI
DMGFDASLSAKTEPLVLITQPSLEMLLDSQDNMLYITAPHSKVSPVHDDKQEAKTPSKCYIQVTGMTCASCV
E
NHSGHQPYPRSGHFPWTVSSQEYSHPLPPAPAVPQSLPGLAVREWIDASQQHGRQDFYRVYGQPSTKHY
IRKFLEGSSKDVPDSEDDPYQQCGEDELYYTVEQDDFPQEMVNRPPVPVPRPDSSSLHQDKELYISKVFA

ALCEAGCNVNIKNKEGETPLLTASARGYHDIVECLAEHGADLDATDKDGHIALHLAVRRCQMEVVKTLIGQ

VQDLVSLGAQVNTTDCWGRTPLHVCAEKGHSQVLQAIQKGADGSNQYVALEATNYDGFTALHCAVLAHN

REKLGTSYTGTDCDDVRQKHTSGAKSGIFRIKPAGSNKVFSVYCDQDTTLGGWLLIQQRLDGSVNFNRTW

QARGRARAENSLYSVLAKANSKEVSRERLNETLEELTQRVIEEVQAMPEKEYRDTITDLQRSAIISRKVKKAEGI

FMYVQCKVYVQLWKRLKAYNRVIYVQNCPETSKKSLFEKSALIEPNIENKEMLGVHHSDTNSSHYTEPEDC

IMGACFSKVFNGCPLKINSAVTWIHSETRDQYLLVGAEEGIYTLNLHELHEDTMEKLLPHRCSWLYCMNNV
RDLVNGYRCICPPGYAGDHCEKDINECASNPCLNGGHCQDEINGFQCLCPAGFSGNLCQLDIDYCEP
RNFTVEVVGVGDTCSFAQMDVRQHGHPAWMSAGKTEASIYQQAEDGKTELSLMHFAITNPKWQPPREST
EQFEMALSNIRPDNTNTNFHEFKMHTFTRVTSCKVCQMLLRGTFYQGYLCSKCGAGAHKECLGRLDNC

APRYQHIGMVIVFEIDPGSSNWTDTQHLMGKQIGSYFGSVLCSVDVDRDGDTDVLLVGAPLYYEERSGGR
YPGTGMAQFHINYNNFTNDKEWLINITLTIRPSTGTGVMFALVSGKTVPLALSIVDSNSPNSQEIIVTIENIIVAF
WDIRTESIVQELELFGRITALDLNPERTELLTCSRDDLLKIIDLRVNAVKQTFSAQGFKCGSDWTRVVFSPDG

LGEKGGKGSRGSPGSKGQRGSTGKTGLPGPNGDPGPPGPPGKDGPPGPPGPPGFQGLQGTVGEPGVP
HLHLCLALFIAYSIFLTGTSSTGNRVVCGVVAGLLHYFFLAAFCWMCLEGAELYLLVVQVFTPHGLRRRYMFL
IFVHEGYIYGTNYNNDIALIKLKHKVPINANITPICLPRKEERFHVKTDDMGTVAGWGTTEKRRPSRYLLYVEI
RSCADCVLARDPYCAWDRHSQLCQETRGTDKNTSDWLQDVERGRPGATCQQISGKGRATPRSRDDLG

EEQYYMVRRFENQVLHPPPRPPKKPTTLESKPLLTLPERSVSDAPHAVKNLYVNTTKKLRSAQATAVSELQVSC

FYVCARPPGRSSDPRACCNFFRWAGKGHS
RLRGG

YDQPLLSLARLTSPDLRPTTGFEFMEVPDDV
ATPPPPNSANLSPQSSSHSAIGLSGLNPSTGSTVLGVNAGLNPALMATNPLATIQALASGGTLPITSLDAGGNL

KSFQEPESLPPIVKKALKILPVVTWKSSNSNSPNKQFWKYMRYHMPLKNVGGLGKNSLKFFFHRLFSMVINS
QSQESPQREAPSPSRYVPASPPERPERDSLSELQVYQRPDRTEHHFMDAIDRREDTFYVVSFRRDHLLLPAI

TSLDNGECSQICTSLSPSSWECGCFPGYKLQADKRHCTAIGPKPFLLFANGQDIRRISFDGTDYTSLLDW
EFVKEIDNEKRMRLLQFVTGTCRLPVGGFADLMGSNGPQKFCIEKVGKENWLPRSHTCFNRLDLPPYKSYI
ETIGNYEGLQESLRYIIKQQGTIKWKEKHKECPHASNSKFWKLVRYHMPRRHKPPQLNYAVV

GKIVSFGTQERIMSSDYENLSFQLTQEMVPSARLLVYYIVTGEETAELVADSVWLNIEQKCGNSLQVQLLSKQ
DINLQFMSR

QTIVHRLKDEETTEDGGRGSKKEGQQVSVKDVGVKTAEMASAKSKTEDGENYMMENCVQKPTEPEAEHST
RFSDVLVQKGYKVVRVEQTETPEMMEARCRSLSHPTKFDRVVRREICRIITKGTQTYSVLDGDPSENLNKYLI
SDRPSFAEIHQAFETMFQESSISDEVEKELGKKGMRSVVSNFLQAPELPTKTRSSRRAAESKDASENTEMAH
AAQLVLFGGARSHGLFVIRQSETRPGEYVLTFNFQGKAKHLRLSLNENGQCHVQHLWFQTIFDMLRHFHT

VFRSVIDIINKYAFFASEYPLILCLENHCSIKQQKVMVQHMKKILGDKLHTQSPNIEESYLPSPDSLKGKILIKAI
KEMKKFWQGSSYQMMEHLDYTANEESTTTANEQSGEQ

HIKITDFGMCKENIFPGVTTRTFCGTPDYIAPEIIAYQPYGKSVDWWSFGVLLYEMLAGQPPFDGEDEDELF

VK

TGPILHVSVRPFETLPAVICTARHPASNSSRLISPKDLCPDSPLNRSLALSLSYHLIILLLALGALSAGVIA
WIQNIGGDASSCPDKVRENSLQHTFKHGSTAELKCSQKSNLAQVVWKFKDDVLKVESPKYHLLEKALLIFN
CECPCDQPEPHATHCSGGHGTLTCGVCSCNPGRVGKLCECELAEAMDLEAGCRGGNGTGPMCSGKG
PKGIDTVRTLNLNEQMVVEWEASVSEINRYVIEWYEELETEPLGRSWQYILNTTKWTAQKDAFKPFKCYNIS
YVCEAVNEISSIRSPTLQIDVQYAPKDVRVELVTGSQIQEETTVVLRCSCKAHPPVSNYTWYRNGQHIPEQTQ

QFPVPITETVVLTSGIRAKSSDITSILLESVFTTSENVSSVTIKHTGLLESPEFMPSESVLLLDKTPASVSTDKSSINI
ASGAGGFQRFSGQNQHPSGATPTLNQLLTSPSPMMRSYGSSYPDYSNPSAPPQQPQPQAATAAPGSQQSA

GIKAFNEFLVDCDFTQADYFGYNNLLSYGAATQPYPLPTHPQLVTVLFGSDQAVVCWRPPQPTIGTSPSAW
SQSDGTSTETDLSFPWILNVLDLKAFDFYKVIESFIKAEPSLTREMIKHLERCEHRIMESLAWQSDSSLFDLIK
PLTQRRKPLPDPLQIPHLSLPPVPPRLDLIQKGVARSPCASPTGSPKSSPCMVRKQDKPLPAPPPPLRDPPPP
KLQQEFYEEKRRKQEQVVVQQCSLQDMMVHQHGPRGVVRGPPPPYQMSPGEGWGPGGPESFADGMI
HERKTFGSWHSSSILDIQDTVIGFSVECCANNSVGSSCAKRFITQQEAVFSLQDNVSFYATAGFCLPFIAVLL

RYAGMALTNLTFGDVVNKATLSSRRGCMQAIVAQLASENEELHQVVSSILRNLSWRADINSKKILREVGSVT
KPGNAISLLMEKTVSRDDNMDITKCQITSDNQVPREYKMNAHTISSVPDNGSEKGIQNCRAASETLGLDSN
SPKSPPSEMSPPVSSLTVSVPSVAVSPFMEEERPLLLVTPPRLREKYDHHIQQFNSFHHNPAHESNSPPPSP
NQKQLKSLCSTCEELKANLNGRVTAEEHVALVNALRSQLQQEQEKQSAEMEDLTERLASLQAQKKSLLLEKN
ITELRTFVVPGDLHVTADMAFLLISMVLNIPPMVVTDTATGACATTWFHWYQLGLML
NSGAAGGNAAADNDPEMDGEDGVSFISPLGTLYTPALKADPVAHLPLSLVLSVVTSALVAALILAFSGIMIVYR
VKALKDPTLAARKDFQREAELLTNLQHEHIVKFYGVCGDGDPLIMVFEYMKHGDLNKFLRAHGPDAMILVD
DWDETSRDSEASSARGDSRPRLVAPADLSRFEFDECEAAVMLVSLGSSRSGTPSFSPVSTQSPFSPAPSPSP
GLEPLPCSSGGEDLEDEDEDDEDEEDEEEDEDPLPLAEGQGPPSPKEGEGPPFSPMQELFVLEIEALPLELS
VKSMRVMRPEPQTAVGPSHPGWLPAQAPPVDSLEIIEQHPLPVGGANAYQLDVDYSNQEPRCPPPPYPKH
LSNPCQNDATCLDQIGEFQCICMPGYEGVYCEINTDECASSPCLHNGNCIDKINEFHCECPTGFNGHL
SSSDSDSSSDDSEEERAQRLAELQEQLKAVHEQLAALSQPQQSKPKKKEKDKKEKKKEKHKKKEEVEENKKS
PYKYFISRGKNGDYEFIYKPCPEGVFVNRCLFIPSKYLSSGDWHQYDDIVCIRPPDGLVRRLKSLVGWDD
RPGKWEGIVFLPSAKEKQRKAEEMEEDPYSERSKKEDKLASKRAEAGHRGFVPEKNFRVTTYKASQEKSTS
GKSSKSDRESKRTSESEVSKRCSPLNELGYRRGSSHSKPDSNVNSSHYKYTPTKTSAPKSDKSKSSFCCTES
FAPPSRVQRLTGSLDSLTEVSTGKMNHSFPGFRRSLSTTSASSQKEINRRNSFVKSRTSMLRRSWSNTTTTN
DTHNFTFVNLAPGSQYLLELASTAGPYGMSTGNVSNWTYPLAPPNVYLRNQGHSDRLSASWGAGAGERD
HYHGTCLDVPVTPRKRSGWQCPECKVCQTCRQPGADSMMLVCEACDKGYHTFCLKPAIESLPTDSWKC
NSLTAANGAVENYLLSSFGERRFEKTALKEEYQSPHGDSLRDTLRVDLLSSSSAERRKSPVGLLCSSKLQVHY

SFSMDSELSRNRENGSKSLPDEKSLVLGKVIENVSLGAIQQYNDMLAEKQKRSSFMKRSSDQRDLCPQDL
QSPENDNGDRRSRRFSMSERRFGQRAAKKLSTLQSVSQQQSSSSPPPPLLTPPPPLQPATGISDHTPWLMP
EKKIDIKLEKNDVKRDEGKEHAELKEDTVKSVKKAVSEQPENESMDQNVPATQQTESKNTSSEEKKSETNEE
SVKVVEPPKALLESPLRTWLAKSVHKAASTEEKANQEKAEVSFKSPFLDLLAPFHLPLNVNSWTLSSKNIENVN
RRQTLFQTNNKPVEQCENEGLVCFHLCAKGQCWGPGPTQCVACEQFLRGQECVESCNLLDGAVREHVN
RERKSNPLVHTREMPRSPKMEDVTTDVKPSSNSRASKLKRKRKTASDLQPVDFIKKRDEKDLKKHSEGVNQ
TPEIKAPEPLSSLKSMAERAAIGASIEDPVPSLHLSERDILISSTTSQPASSQPPIQLSEVNIPLSLGVCPLGPVPLT
ANNKDALRKTWNPKFTLRSHFDGIRGLAFHPVEPVLITASEDHTLKMWNLQKTAPAKKSASLDVEPIYTFRA
YPLSYSSLLPMLEKKKRRRCGVCEPCLRKTNCEECSCCRKRKTSHRICKKRKCEELKKPPPPIMLPFEVLTEN
QQQVYAQKHQQLQQMRMGQPAGSLPRQAGPQMLQQQPPRLISMQTMQRGNLNCGAFQAHQMRMA
GVLDKKTIKTINKMKTLKRKNILNQILSCSSNIALKTKAQPPSSTGPATIDARLGKQINVSKRGTIYIGKKRGRKP
RRVGPVGLPNPRILGPVQHTPTSHQQPPPRLINFQNHNPNSNGSILPAQQMRSPQNQNVPRLTIKPNPLQ
CENGQSNKAFITITADDKDDAANGPRFIFSLPPEIIHNPNFTLTDNRDNTAGIYVRRSGFSRQKQDLYLLPIV

FEKRRKEFEVIAQIKLLQSACNNYSFMQEDSFVEWFHGVERLSEAESYSLSCEIEPLSESASNTLKAKKNTGI
ECTAGCTFTTPGDWTSGHVGAPLPCNLIKLMDLEEMNYFASKGEGEICVKGPNVFKGYLKDGEKTAEALDK
EVSEGTTPCSSVETQPCDSPLMSALQKSSSLGNLKKETSEVDKEPVQKPAETKAPMEGNKSGTLPPKSPSHS
TRTEGNASSLKTNQGSENIQVEMDSTLQASELKPCSTESYSMQLRLSEDSQALLSENCEKPKREAVCATGEP
DHSLGADGYTLAVTGMKITKIPLNGPGCDHFKSCSQCLSAPPFMQCGWCSDRCVRSWECVKRMWTQE
PHGNRQNHSSSSQPSATTVTAITPAPIQQPVKSMRVLKPELQTALAPTHPSWMPQPMQAIQPVPFSESPSTN
SQQPQGQGQDGGRREGSAVTLGTTEPVSQAWSIRSQGDNLVGSANLASTPSGYRKQDGGQEGKVKSQS
INLYMELSAIMLQLEVREKELIKREQAVEKKYPGTYKRHPVRPIIHPIAVEKLMKKKGVSHKPGSQTKRPDLLKS
GSSLLPSMSSSKKHKSEIQAQPHPTDCNKAVHINSESVLCNKPHSQTEPWPSLSRAHKDCRRTKLIFDDIPR
SGNEVYHINLEDSTFFAEYEGKSFTYASFHAHKKFGVCLIGVRREDNKNILLNPGPRYIMSATDICFYINITKE
GSRALSNSISNMGLNDTNNLGSAKHQQQSPINQSMQTLSDSLSGSLYSASVNLPVMGHDKFPSDLDLDIF

NVESTSSMELDGIFSNLSHQSYSQRCQQLSVTVLTDHQHLDDTALNTILNSSVPQQYRIRAKLRTFKPEKLY

SIPWLEKTLARVRAPSRGQDVQCDEQTSCPDGNTCCRLSTGAWGCCPLVEAVCCPDHIHCCPKGYTCD
LLAASLKDKDQESERDEDRAQNGAVEACVRWLLENEFIQILDSGNGMKAEVYRPTHLGTATLSSSLSPAEAL
LAIPGGRVCACADHQLLEENSTVCMINPGEALPQLCKPGEFQCNNKRCIQDRWRCDGDDDCLDASDE
VPQSRPRSVAAWHDLYIPPCDTYKTRHLFSLSMYVAAFPHQYRVTALQASSI
MNEREEKDTWSEEDELFLDKKSPGKNSHSSSGLGSKKQRWTVQESEWIREGVKKFGEGNWKIICQKFPF
SGSINSGAAPPANGGQADLGGAASLPRVENSGLASSEWDTELGGGAGGEDLSRSRQHRVMKRKLLVDDG

NPVKIKDGLGGAWAFVLRDIIYTLIHYINSRPSLGRDMSMRSFSLCCDLLNLVCHAAVTYCKDALENHLHV

FSRPEEHHMLQGKAEDCILLFQEASQITSRRQKALSQLGVFLELHAAASSVLDHVRHTVETTGNMDKAQSE
LPYICKRSNTTAVKPSLPPLPAPVPGGCARGWVPFINKCFSIKGYNKAEQVKWQEAKQACESQGAVLATISNH

NLSENDMANYSADLLMTTNEEDNLVEFDDGFRQENPGDLSEKTSSLTDLVEDDSSSPGVLKNVETRIPPTP
NSVKLSEITEKPSRMHKELEMHSAELPPTPLKISEDSSSPNRMTPPDPLVLLSEHPTEGKAENADPPSESKGN
RSSESAQNWEAVNSSPVEADKRFFINVCHKVLKKGAAVVCSDDAAVCSVDKNNKPKNLGTVMSSLKVGEF

YDFASSPLHEENDLFSEEEKKRLKRFSTEEFVLL
EDLHNLLSLILTANPLRHLGPKAFYSLGSLQKLVAVEDNISSLADLPIGHLHTLKELNVAKNNVDSLKLPKYFSN
SVADLPIDECEGPKPASQRACYSGPCSGEAAESNPEEIELIYGGLQDFDELYDWEYEGFTECSESCGGGVG

NKKEEVKIVTSTFLLPNRITIPPDPGTKKILVKTFSSAPNTQATPDLLKGQQDLTQQTNEETAKQILYNYLKEGST
FKDSLTVLVNFECNCTCQEHAEPDSPACHQGNGTFECGVCRCSPGRLGSHCECSEEEYSPTQQDNCSP
RNYRRRGKNQQQQQMPRSSQEEKDEKEKEKEKEAEKEEDKQEAENDKEELTKEKNDDTSGEDNDEKEA

PQKYSPHPQRFTYCSQVLGFQCFKRGIHYWEVELQRNNFCGIGICYGSMDRQGADSRLGRNSSSWCII

SDRKLTAEQGHNITFIVQLEEGDVQRMIIQVDFGDGIAVSYANLSSTEDGIKHVYQNVGIFRVTVQVENSLG
SQLCGTRLCHEIAHAWFGLAIGARDWTEEWISEGFATYLEDVFWAVAQQLPHDEAREQQELKALLRWRRI
EDVTLSCTAGVTEADCRLQDDLFLEKAMQLAKRHANALFDYAVTGDVKMLLAVQRHLTAVQDENGDNVLH
PSTQLYLHQTETLTSNLRMPLTLTSNTQVIGTAGNTYLPALFTTQSAGSGPKPFLFSLPQPLAGQQTQFVTVTQ
FYGTGLIAGHGFTSPERTPGIFILFDDDRFGFIWLELKSFSLYSRIKVSFQNAEAPSQEAFDEMLKNIQSMTT

GPEVIFPGQRCLIYAQLCGQPQPPDTAMGGVTLQYRIQDQTYKETLQFPLQPQDGDRLPVHRLAAKSLLLEL

EIHIVIKGSPCEISFTPMNGNFICIEDEAGVNCTLRCMDGYNFTAASSENYYCAYEDGIWKPPYSTEWPDC
KVRSTGNWIASLEDASLNNSYAIRAQDKVPYDIYRADHSFLTVVLPVNASVRDVLQSLACQEGAHGECMLV
ALDLIEPEQAEDLTQPKVEAMNHRLGDLVEQFKQLVYPPDYNPEGKAMKRKQGEGADQTEKKPRMEVSKD
QPFNSPYLSTSEEPTENCCIIESIVIEETDQTDFKDYKHSKQSTRDSGNYSNDDDTSGTKVLEETLEQETT
TPRRNVDKCQDN

PSDASTNQSSPEPCSPNDYCFDVDNDMANEFKLELVEKLFAIDTEAKNPFTTQETDLDLEMLAPYIPMDDD

LPSLPPELMQAVAHQITQQAMAAAASAATGQQVPGFQSAPTRVVIARPSTPSARPTHPGAPQAPGPTTPGIGC

A

DRQLSQHGNKAAVWEELIDSGKLPFMAMLRNLRNIIRAGVSERHHQRVLNRLQNKESVIRSRQFPFRFLAA

NEINLLTVALVNQEREKNSEKRSQGLKSEKEALLIGIISTFLHVHPFGANIEYLWSYMQQLDSRISANEIEMLLN

AKIDEEVRKFREARKENIEEEEEEVREAMTRARAHRSHEAALASGSSDDDLMDMGMSEQKTNDDSDAGTS

QTAPATTEMPKKSSKIASFIPKGGKLNSAKKEASAPSHSGIPKPGMKNTTGKSSSAPTSAKEGERSRSGKLGSG
VKGFTSDRFLYANRTPMYMTDSNDTSSNGTNVNTFSLDLKTILVSTAMGCFTFLGVVLFCFLLLFVWSRGKG
PEVAISCVPGSSDSLADEEESESHKLSIAATNCLINSCVCCGSCEDSREDNIVGLKTRHSPGKVIKASYSLVK

IHESYSNGLSVTLSTSNSPNSGMDQKSLKMSPSGQTKSQEAERTPTGNFKRTLEESNSGPALKKPRRSLSRN
GSFPRSGFSFKVFYPSNKTVEDR
LRGKMLNVREASHKQIMENAEINNIIKIVGLQYKKNYDDQESLKTLRYGKIMIMTDQDQDGSHIKGLLINFIH
MGCEGKSFAANPFFYPQSMPSSHCESDSSPVDSDDSIEQEIRAFLALKAQTENLVTKPDNLSHSVQRPLSS

EEPLLCEKNHWWSHPLPNCDALCGGDVRGPSGTILSPGYPELYPNSLNCTWTVDVTHGKGVQFTFHTFH

ANPNLKSALKTCIKSEVASISVQKAAIQALRRMTITDEDRTVLLKAFQNANAPVDKRLAAYLMLMKDPSASDLN
LRLTERCHRIPARSAEEEELRMCHSPAYVQRVKATSSMKPRELHRQGDQYNSIYICASSYECARLAAGSAFI
CKRKNKEYLLENSPPTKSERQISVANKPTSICCIQYSGDGELLACGLADKSLLVFNSNLTGTPSVFSGHDGA
ELEDSFQRVSEQAQRKTQEQAEVLETLKRELTASKQELQALQSTLQSSLQSEAERSTRTAGLEQERDTLVVAVT
MVYQSLPDVPATFLD
TLATGIPEYGFLINPKKTMMNFPVDEDIPGCSKFKQLPYCSVIPWCGLLLDIQTLEVYCDYSSYACTSIRSSLA
VRIIELSQTIAKVSFNKPDSHGGVPIHHYQVDVKEVASETWKVVRSHGVQTMVVLSNLEPNTTYEIKVAAVNGI

EKTCRDLISREWNENLLQEACRLLADEMNLSPSAPGGMVEFRRMLTLSFFFKFYLIVLQKLNKPPNGNNNL

TVWAANAGVTASPPSDLIWKNQNSWGTGKDVKVILKNSQGEEVAQRTTVFKTTLREGEEEEEAEEEAVEAIE
AENIEVFAQGHGIIQVDKAYDYLIQNSSFTSNIGFTVTVGNNRGIYLRDPVQIAAPSDRGVGIEPVFPENTENT
PDEVLRKGSGTLCEALLMVQAYHANIIFPNKQEQEFNKLTEDGHVLDSETYVGGHVEALESGVFRSDIPCRF

EGKTRGFPKNASICKRESNSTLWRPHKTRDDFLGQVDVPLHQLPTENPNMERPYTFKDFVLHPRSHKSRV

DSILYNPPSRRDEVQSVRKKFISSEGDHLTLLNVYRAFKNVKGNKEWCRENFVNNRNMMLVSDVRAQLRI
DRLLRPHAFYQVHRITGKTVSTTSHETILSNTKVLEIPLLPENKMKAIIDCAGILKLRNSDIELRKGETDIGRKNTI

QYYGVPWGVYPANLFQQQATAANTTANQQAASQAQGQQQVLRAGASQRPLTPNQGQQGQQAESLAAAA

ATWGRLVRRAVPTVNHSGTFSPQASVPATVPVVDTCIPELARSASDSPVAKLDFEPEPPPAPPRASSLGGICD

HKKLKLSNDGLQMEKDESSLKKSHTPERFSSTGCYGAAGNIFIDSGCHYWEVMVGSSTWYAIGIAYKSAPK
IASVMFCTVGVLLRKLEAGIRGISHVIVDEIHERDINTDFLMVVLRDVVEAYPEIRVILMSATIDTSMFCEYFFN

VKGDFTEKSHSANTKFWKNVRYRMPPRCPTFSELHLLPTASNTFQTTER
NQQQKTQVVQSSTGENADSQLIAMPNTDPSTSRNSGNELQVYYASLRNYQDFFEAIHRRGDTFYVVSFRR

VRVRSCEEKLKYVCMKNGEVLNETSDKHCFPKEEWKRHGDFCYKIHNNKVSFGAQCNLTIMNRFEQEFI

SHEPRFKFQAMINGRRFPPAEAGSKKLAKQEAAANAMKILMREAENQGEDGMEGEESFVPDSSEAELPARI
DMGFDASLSAKTEPLVLITQPSLEMLLDSQDNMLYITAPHSKVSPVHDDKQEAKTPSKCYIQVTGMTCASCV
E
NHSGHQPYPRSGHFPWTVSSQEYSHPLPPAPAVPQSLPGLAVREWIDASQQHGRQDFYRVYGQPSTKHY
IRKFLEGSSKDVPDSEDDPYQQCGEDELYYTVEQDDFPQEMVNRPPVPVPRPDSSSLHQDKELYISKVFAG

ALCEAGCNVNIKNKEGETPLLTASARGYHDIVECLAEHGADLDATDKDGHIALHLAVRRCQMEVVKTLIGQG

VQDLVSLGAQVNTTDCWGRTPLHVCAEKGHSQVLQAIQKGADGSNQYVALEATNYDGFTALHCAVLAHN

REKLGTSYTGTDCDDVRQKHTSGAKSGIFRIKPAGSNKVFSVYCDQDTTLGGWLLIQQRLDGSVNFNRTW

QARGRARAENSLYSVLAKANSKEVSRERLNETLEELTQRVIEEVQAMPEKEYRDTITDLQRSAIISRKVKKAEG

FMYVQCKVYVQLWKRLKAYNRVIYVQNCPETSKKSLFEKSALIEPNIENKEMLGVHHSDTNSSHYTEPEDC

IMGACFSKVFNGCPLKINSAVTWIHSETRDQYLLVGAEEGIYTLNLHELHEDTMEKLLPHRCSWLYCMNN
RDLVNGYRCICPPGYAGDHCEKDINECASNPCLNGGHCQDEINGFQCLCPAGFSGNLCQLDIDYCEP
RNFTVEVVGVGDTCSFAQMDVRQHGHPAWMSAGKTEASIYQQAEDGKTELSLMHFAITNPKWQPPREST
EQFEMALSNIRPDNTNTNFHEFKMHTFTRVTSCKVCQMLLRGTFYQGYLCSKCGAGAHKECLGRLDNC

APRYQHIGMVIVFEIDPGSSNWTDTQHLMGKQIGSYFGSVLCSVDVDRDGDTDVLLVGAPLYYEERSGGR
YPGTGMAQFHINYNNFTNDKEWLINITLTIRPSTGTGVMFALVSGKTVPLALSIVDSNSPNSQEIIVTIENIIVA
WDIRTESIVQELELFGRITALDLNPERTELLTCSRDDLLKIIDLRVNAVKQTFSAQGFKCGSDWTRVVFSPDG

LGEKGGKGSRGSPGSKGQRGSTGKTGLPGPNGDPGPPGPPGKDGPPGPPGPPGFQGLQGTVGEPGVP
HLHLCLALFIAYSIFLTGTSSTGNRVVCGVVAGLLHYFFLAAFCWMCLEGAELYLLVVQVFTPHGLRRRYMFL
EIFVHEGYIYGTNYNNDIALIKLKHKVPINANITPICLPRKEERFHVKTDDMGTVAGWGTTEKRRPSRYLLYVE
RSCADCVLARDPYCAWDRHSQLCQETRGTDKNTSDWLQDVERGRPGATCQQISGKGRATPRSRDDLG

EEQYYMVRRFENQVLHPPPRPPKKPTTLESKPLLTLPERSVSDAPHAVKNLYVNTTKKLRSAQATAVSELQVSC

FYVCARPPGRSSDPRACCNFFRWAGKGHS
RLRGG

YDQPLLSLARLTSPDLRPTTGFEFMEVPDDV
ATPPPPNSANLSPQSSSHSAIGLSGLNPSTGSTVLGVNAGLNPALMATNPLATIQALASGGTLPITSLDAGGNL

KSFQEPESLPPIVKKALKILPVVTWKSSNSNSPNKQFWKYMRYHMPLKNVGGLGKNSLKFFFHRLFSMVINS
QSQESPQREAPSPSRYVPASPPERPERDSLSELQVYQRPDRTEHHFMDAIDRREDTFYVVSFRRDHLLLPA

TSLDNGECSQICTSLSPSSWECGCFPGYKLQADKRHCTAIGPKPFLLFANGQDIRRISFDGTDYTSLLDW
EFVKEIDNEKRMRLLQFVTGTCRLPVGGFADLMGSNGPQKFCIEKVGKENWLPRSHTCFNRLDLPPYKSY
ETIGNYEGLQESLRYIIKQQGTIKWKEKHKECPHASNSKFWKLVRYHMPRRHKPPQLNYAVV

GKIVSFGTQERIMSSDYENLSFQLTQEMVPSARLLVYYIVTGEETAELVADSVWLNIEQKCGNSLQVQLLSKQ
INLQFMSR

QTIVHRLKDEETTEDGGRGSKKEGQQVSVKDVGVKTAEMASAKSKTEDGENYMMENCVQKPTEPEAEHST
RFSDVLVQKGYKVVRVEQTETPEMMEARCRSLSHPTKFDRVVRREICRIITKGTQTYSVLDGDPSENLNKYL

SDRPSFAEIHQAFETMFQESSISDEVEKELGKKGMRSVVSNFLQAPELPTKTRSSRRAAESKDASENTEMAH
AAQLVLFGGARSHGLFVIRQSETRPGEYVLTFNFQGKAKHLRLSLNENGQCHVQHLWFQTIFDMLRHFHT

VFRSVIDIINKYAFFASEYPLILCLENHCSIKQQKVMVQHMKKILGDKLHTQSPNIEESYLPSPDSLKGKILIKAI
KEMKKFWQGSSYQMMEHLDYTANEESTTTANEQSGEQ

HIKITDFGMCKENIFPGVTTRTFCGTPDYIAPEIIAYQPYGKSVDWWSFGVLLYEMLAGQPPFDGEDEDELF

VK

STGPILHVSVRPFETLPAVICTARHPASNSSRLISPKDLCPDSPLNRSLALSLSYHLIILLLALGALSAGVIA
WIQNIGGDASSCPDKVRENSLQHTFKHGSTAELKCSQKSNLAQVVWKFKDDVLKVESPKYHLLEKALLIFN
CECPCDQPEPHATHCSGGHGTLTCGVCSCNPGRVGKLCECELAEAMDLEAGCRGGNGTGPMCSGKG
PKGIDTVRTLNLNEQMVVEWEASVSEINRYVIEWYEELETEPLGRSWQYILNTTKWTAQKDAFKPFKCYNISY
YVCEAVNEISSIRSPTLQIDVQYAPKDVRVELVTGSQIQEETTVVLRCSCKAHPPVSNYTWYRNGQHIPEQTQ

ISDFSSNLLSTPFLSEPSSWSSLSTFYSRMPSDLSTLLPQVFPTLLGPSVVFDDSSSWDSTEFPAADSTSIEVSH
AAGMGMGMGMGKDMGSQYGSANPAWAAAQQRAHPAMSPGSTGQTINRTQGSPMDPMVMKRPQLY

VQNWTYIVKVSAQHGADEDRIVSNISDTRFTVKGLTYSTAYQVSVRALSPAGEGPWTESFRGATLKEVEEEPY
QSKEREGQVDQPEPTCTLNFPLQHNHTAADLYLSPVRSPKKKGSTASLNPTPIIDAQPVLAPQTQKPQKSTSL
PPERPPPIPPDNRISRHLHHPESVPSRDQPVPPEAWCPRDVFGTSPLIGCRILNDSSPKPGLTASSNINGRH
NVSHSLPPRGMAPHPNMPGSQMRLPGFAGMMNPDMDSPSVPNPTSRPGLSGVSWPEDVPKIPDGRN
LIFIFHKYKKQFRYESQMQMIQIVGPSDNEYIYIDFSEFEYDLKWEFPRENLEFGQALGSGAFGKVVNATAY
GLMQCALRATKESTLKSVLSALWNLSAHSTENKAAICLVQGALGFLVSTLTYKCQSNSLAIIESGGGILRNVSS
TNPGSWTSKSRFQHSTVTSLLVSGPSDKTIVFSGEDMDLTRSYVEKGIASQLPPVSAGMSTSVTFKPQSFAN
LRIVEDEEYETTQEYEPAQEPPKKLVNSRRAKRTKPNGHISNRLEVDTDTSSESSSSESETEDERIGEDTPFLS
NDLVAENKILEAELKMAQKTNRKSQKKICLLKQQLDEAMEKEVAAHQYLANLISLAENITQERDHLIHLAKCLE

RKHQELQAMQLELQSPEYKLSRLRTSTIMTDYNPNYCFAGKTTSISDLKEVPRKNISLISKFNHQNIVRCIGVS
DGQPRQAKGELGLSQMLHIASQIASGMVYLASQHFVHRDLATRNCLVGANLLVKIGDFGMSRDVYSTDYY
SPLFGFRPANFSPINASPVIQRAAARSRHVSASTPKGAGGGVLSPEMLHHAPHRERHSSGIVPTFQTNLTFT
ETPPPSPQIPGGSEATPKAAPSSPEPSEATPLGSASPETVSSRTEWQVSLKEPACFAVKQERGSMGDSGLRT
ILLLPGSSEQFDINCLCMGVEQTLRGVPNSTCNKAEENSERNDKSSKNAKVEKTSKDKKQIQTSPVPVRKN
CQYDIDECASTPCKNGAKCIDGPNTYTCECTEGFAGAHCEIDKNECDPDPCHYGTCRDGVAAFTCLCQ
SKTKEPPLKKAKKNNSNSSNSSSKKEPVMVKNSKPPPVYESEEEEKCKPMSYEEKRQLSLDINKLPGEKLGRV
WNKGVERGKQIAAEIMLETIFSILSTWTDINMKNFLNQLHQFYVVTKTPMVYENEPKEWALLEFGEKQVKEL
PPLRKASESREKLGTRGDGLATGKSSFSITREAQVNVRMDSFDEDLARPSGLLAQERKLCRDLVHSNKKEQ
SVEVKQQSNFLDLETPCLKSNELRMTIIKKLEREKTVSPFSQLNDSPTFKKLDESKDGSPDLKSELTGNDCHD
NTEGPVKLHEGSQVLLTSSNEMGTIRYIGPTDFAPGIWLGLELRSAKGKNDGSVGDKRYFTCKTNHGVLVRP
DGYTLTLYHAQAGTVAANASVRKDAHKFTFSGLAPGQKYLLEVASVAGPYRTFAGNISDWTMNFSAMPGESP
KNCRVCSDCGSRPAALDPGCHWYENYSVCEGCQRQRSAAGGTDATRCCQSCQKPLPLDSDATGGST
YSTADTSIEEKKQTESDSLTSFSDDGTSDLDTGDQSSEAACSALEISDSADKEEKEFCTGMRGNSKCSMYLS

LCQRDTGDEDSVAGELDRTEEGTVNGRNFGPGEPFPGLFPRKPTPITSPSLNNSSKRIKIEKDLDLPPATIIPS
PTIPLASPFLPTSAAPMQEKRKSILREPTFRWTSLKHSRSEPQYFSSAKYAKEGLIRKPVFDNFRPPPLTPEDV
PQYEPNSTEALDMDIVSVPSSVPEDIFETLESAMETQSAADEQGNGSTATELETVNMNIKCTTASKDIKGGTK
QPSVEDKWLLRKKAQDYGLPTVCDLFACMKLNGDREKWLYQTPLQK
NGTRCLSCHPECLPQNGTETCHGSEADQCMACAHYRDAQYCVERCPSGVKADSSFVPIWKYPDEHGV
CLVGRNTGRKKSGEGSTVVNGSPMGVKNGDSIPAELPLQEGKSILKNDTEKVTCNNTDGEPAQNTCDRKS
TKEQLYQQAMEEAAWHHMPHPSDSERIRQYLPRNPCPTPPYHHQMPPLHSDTVEFYQRLSTETLFFIFYYL

HSGPVLCVVMSSNGEQCYSGGTDGLIQAWNTTNPNIDPYDSYDPSVLRGAFVGHTDAVWGLVYSGAHQ
IKRPQRRKQRVLKADLENKPVNGLRPELMECSICGHGEKYRVKTNQTVSFENVQSNEKESMTGLEAEKWT
QNAARIPGIPRHNGPQYSMMQPHLQRQHSNPGHAGPFPVVSVHNNTINPTSPTTATMANANRGPTSPS
RAELQPPPEEPKTVIKHSRPVSSQPENPAVPSNLQSLVALSPAAIHPLSTQLVGSNGNLSPASTETNFSELKTM
MAFLAQQAIKQWQISSGQASAASSTASSTTSTPSSPTVTSAAGYDGKTFGPPMIDLSSPVGSSYILPSLPDIEC
ISDGGIPPMSSTNTLTIRVCGCDSNGSMLSCNAEAYILNAGLSTGALIAILACIVILLVIVVLFVTLKRQKKEPLI

IKRWSDRQVPSTEPCTSGSSHSKSFDQLKCGQYLCSGDTADSVSVTSAGSSSSDVEEINISFIPESPDCQE
EGWLHTGDIGKWLPTGTLKIIDRKKHIFKLAQGEYIAPEKIENIYIRSEPIAQIYVHGDSLQAFLVGIVVPDSEV
GTVDEDSFGTRKARSSFGRGFFKIKSNKRTASAPNLDRSRSASAPTLAETEKGSADHLDLAGLPPRPKETDS
TICPVAEGVAAEGSVAQMACIDLSCDSGSQTAATLSVQSETSSHTLGQEALTEIKEKLSGEASEFEAVPETLCE
ETCLPSVYEIVPRSAPLEGGTRLTVCGWDFGFSKNNRFDLKNTAVLIGGKPCTLELKVSNKNKLECTVAAAK
MAVMTPVAEAPNYQGPPPPYPKHLLHQNPSVSLYETGTKPSKEDHPNLPKEEESEKNNECVDGTDKEKKQ
SEDNADGEGLRDAPMGTKCCCQGDNLVEAGALESGLNGHGQQDGCQGDTVKGQPQAGWEPLGDIP
SEGISNAEVAPNGSPVSGSPKMSTLSNKGRYRCKPRHRRGNSKGSYNEFAGILKNQPAHDNSPQPPPSH
NADHFMQEAKQKKHKADAMVEKFGKALNYAEAALSFIECGNAMEQGPMESKSPYTMYSETVELIRYAVRL
ENSAFKKQEQHRKNNESKSYYHGPSRLPLHSVIASMGTVAIDLQDTGCRSSSGPTLTLPSEGSKNGRRSSI
FNGSLECDMESIIRSELMDADGVGCQADGFLRKGEMGKAGP

QSVKLYCSKCNSLQEVPNDDDFDLILRDCAATAPNPEVHNSLWYDSAVWDTENQGQRKIAVHFVEHDE

PVEGSCQQALLSVPWLEKTPARVRAPSRSRDVQCDEQTSCPDGSTCCRLSTGTWGCCPLVEAVCCQDH
LGIFADLQRAMKGFVLENDLHILYLVTPVYEEWTTIDWYQFFCLWEKLPSSMKRVAELVGIEEGFLARSVKG
EHSEICFNHSCPDDQFKCQNNRCIPKRWLCDGANDCGNHEDESNETCAARTCQVDQFSCGNGRCIP

ENRTTVMIKDRWRTMKKLGLH
GKAHGEVLPNPGKTHQRAQRKPPHVDVPMGDERCMDANVVGQNPGVELVSAQRAKKSRLEQVGPAQEF

VVGTLIPLVRDQSEIQKQVLGLLKYLVIDNKDNENLYQTIKHLDPFPEHPAFEDLRATQQKIKYSKGAFSLLEE

LLEKELREVVQDINKLASAAISLDGALTKAQYHLRHGSSEQRSSCRAMADNLCLELESVQNLLGTKQNEAE
HLEQAFITSILPNVTFDLWIGLHDTKQEFQWVQSEPLRYVNWAPGEPSGYGTSTANNKPTNCVLVWHGSP

MNSLVESSPLDNGPPLFFPQDVIKKINEIDSASYSQSRARYGSWWDGFELDSKNADAWSSSEQESVFQSP
CMENKLQEVQACSENSESITGQLMNSEHMKDNGANKEENKEKNEITSSVETLRKCLANRFAKEQISKRLD
RIQLTYSNGDSCGNNNMIKTIITLMCKPGDLESAPVFNLENDECLYVFEWYTAAACVLSKTEGNNCRVSDA

NLTFFQFLSLKSNKISSISIGDLDVLQGEKGLNLTLVLSMNDIKYIQLGSFEEIHLHELSLRACFENTSVMKACI
QEAVVTCLNKQTREIADETLCVTSRRPPQLLKACNLEPCPPRWEIGKWSSCSLTCGVGLQTRDVFCSHLLS

DNDDATKRKVNLVFEKIQTLKSRAAGSAQGSNSSTEVKALLEQKTELETKVAELQKKLDLEIRNQQNVKEERD
PGAGQPLCSQRGECLCGQCVCHASDFGKVTGKYCECDDFSCVRFKGEMCSGHGQCSCGDCLCDSD
VASKGRKTANSQGRRKGRITRSMANEANHEEAAAPQQNTELASLEMNESSRWTEEEMETAKKGLLEHGR

EWFNSKISAWHNDLEKCLPNTKATKIGVLLNYEGGFVIFLGVGEKHNLIYKFKAQFAEALYPAFWVFSSGTM

LDSAVLYLHVTCPLEHVHLSLPFVTTKNKEVNATAVLWPSQVGTLTYIWWFGNNTEPLVTLEGSITFTFSVEG
LRDEVQNSEEELQVLRPNKENTGEVSESGTCIIKHGLNAEKIFMQVHYLKGYFLLQSLAKRIGEATYFEFLRK
HLAIIHLHSELVRNLLEVVLDFNLDDIINMRNDLYQAPLHLAVITKQAEVVEDLLRAGADVSLLDRHGNSILHL
PGLSTFTAQLPAPQPLAPSAGHSTAGGQGEKKPYECTLCNKTFTAKQNYVKHMFVHTGEKPHQCSICWRS

LEGAVDTGSEGDRRRALETSLSSGVVCSLTAYVGVDTERGQPVQGPLVRRDIPLAGFGAASHVWKMQGFS

SLNRFANHGFKSFEMLYRTTRCDDSNLLKNFVEAFQSALGKMVPLFCNDVDDIDCRLEDLPQKHCLEY
KVNSAGDKVGLQMDTIGVYTSLGLNERLFVVNIQELHTLTPHPEQLGPTVGSSETLDFISSKDLANQLTDYD
ELRSHVQKGTLGKFTVPVLKEACRVYGLKGGGKKQELLDALTEYFSKF

FQLRSFDQPPSLDSGCPSSQRVATSTIFQQTHTLSTAAEEITPVKTECVDDIKTLIVSQSPVLIINETNSAPPSP

GNASLAQMISGLLGQLLMQPVLVAQGSGSSSASSSSSSSSSTTTTTSSASFPAPASPPTASASAATTNTATTAA

NKIIQDMEEQLRKKDSPCPSNVELLRVLLRHEKKQPGALPGVRRQPPWTRRELRAALSIPFLFRAVKSEKQ

RLPRMFKQEFTGVGATLEKRWKFCAFEGIKTT

ASFSRGRGRGRGQARGGRGQISTSRGASRRGRGNTTQEQSNTSRNYKPVSTPTKSMNIIGAFKAAQQ

LSHQKTQLDSRNSSSTSSLASSEGKGIGGVSALNSSSSTQAVNGPVSTTQTTGSNTVSVQLPHPQQQYNH
KHKTSIDLEYVPRKNNGAVIEGEVAGPRRFNMKMI

SKEKDHIDKLDSAVSAQEALKTETSPALLAERDSGREEQKLSNSSKCLAHSSGSQLGAENGSQVEVEISNQC

NISVQPESLSTTPQRKNSNKLSMRRSLRGQKKLGNSLFHQTRKAVCVC

HHNWPSLLRHRFLEEFITPIIKVSKNKEEFSFYSIPEFEEWKNNTPNHKSWKTKYYKGLGTSTSKEAKEYFADI
SERNSVTGGLEPSLPKTLNLSLSRKRRLKGESKVAKQSIDNKTEGLEKGCTHAGYGNSAKMLVLQDGSGLSC

ILEDHHDYLLITENGSFTQPLARLTGSELPLAINAGLYGNFRAQLRFISDFSISYEGFNITFSEYNLEPCEDPGI

NKVTRALLRDKSEQVKSFVATHIANILDSEEVGIEDLKNKVQEALKGSQIPAANDFRKFSQNYQISKRISLPGN
NAVQAVLEGEVQNAVAIVRPPGHHAESDTACGFCFFNSVALAARFAQRLAGRPLRVMILDWDVHHGNGT
AVNSVGWSHDKRWLVSASEDRTLRIWSVRNTEPALFLDKEMFHKSIRFAQFYYIDTFILLCCGAKFHLLSYYL
TQRSQEVTALQAELQQLKDALDSEKESSSKSLETLQTRLGEKESHEQALQQRLLDEQFALLQCTVQEAEQMV

FNSSNTAGKNMKYKLLTVLKLKCHSLFLNLQVNSLRTVFINIYKIFLLQAYRFHACVLQLPFNQQVRKNPGFF
KGQGEYSKIEIFQTLPVREPSPPSIHGQPGSGKSFKLSITKQDDGGAPILEYIVKYRSKDKEDQWLEKKVQGS

LDELVPSKDVSATEVFHKDPVANVQLFQEVPAGQSVEDMVGRPMVHLSAAKQACGEAVYCDDIPCYENER

EEDLFHQQGDPRAANRSCAIM
TKRISLQLHLALTSNAAWVQCPSHLELMNQCRHINVRVDPRGLREGVHYTEVCGYDIAAPNAGPLFRVPITL
FKMNPVAFDFLLQRVEKTLRHAIEEEEGLPLDQVTNFQEVCEEIKVKLNSLKDIPNRIECPLIYHLDVGAMYPN

VKGHLRLKMTYLPKNNGSEEENTEQVEDLEPGWIILDQPDAACQQQQESPPLPPGWEERQDILGRTYYVNI

DICLKLSLPIESSRSDTGNIRRCLAHSLFMNAAELQPDGTYATVDSHQLVAIHPSSVLFHCKPACVMYNELLI
RVRLVFRVHIPQSNGRTLSLQVASNPIECSQRSAQELPLVEKQSADSFPVIGGKKIILSGHNFLQDSKVIFVEK

AAANPALAFGQGLATGMPGYQVLAPTAYYDQTGALVVGPGARTGLGAPVRLVASTPVIISSAAAQAAAAASAG

DSSSEKKGPDTPIQVAEATFDFENERDSSVSKLQAEIICDPEIPHTDVKSTNIREPEERRTQRFQFSLQDFKCC

KNEWIGKNSSSWVFSRCSSNFVVRHNSKEMLVDVHPQMKRLGILLDYDNNILSFYDPASSLHLHTFNVSFI

ICPIIEVYGRTFPVEEYFLEDCIQMTQFVPPLKDKKKKDRDEEGGEDDDANCNLICSDEYGPETKRCMAQL

RDHLLLPATTHNKTKRPKMSIVLPAINISENVINGQDYEVMMQIDCEVMDTRILHIKSSSIPRYLREQQGNQT

NSLIGKYSKGKEKYFWTGLQDISRSGKYTWGDADGRKNEAVMYANWNSLEPAFPGGCVAMSSGKSLGK

PEPEPLSAVAQLHLSSGKNPISILMEYGQKSGSACEFQLLSQEGPPHDPKFTYCVKVGEQTFPAVVANSKKG
VANIERNLRGEDGICSVLVALMAGKAEVRYDPAIIQPSVIADLIRELGFGATVMENYDGDGILELVVRGMTCA

YVTS
GKAHERPGNLYVSAGKVKKEPAVRLVRDQSQSSIYDPFAGMKTPGQRQLITLQEQVKMGILNVDEAVLHFK

GCFVDFQDRHGNTPMHVACKDGNVPIVIALCEASCNLDVTNKYGRTPLHLAANNGILDVVRYLCLTGANV

AVVHELRNSQPPHSPEVQELLLKNKSLVDTIKALIQMGASVEAKDGKSGRSALHLAAEEANLELIRFFLDQPN

VQDYKKGFGSVDDKGKGEFWLGNENIHLLTQKDTVLRVELEDWAGNDAYAEYNIHVGSESEGYVLSVSAY

NQKRQLHNPDAVRLHCINCNMAVCHGSDLRTVEKMHHVNINPDFKIYYNATSNNVEIPRNFKDWKPGG

GAEILHV

VLLSLAGKSSQISAHNLLGLFEQRRQLQKRQVPLSIATNRLTERIIPRKFALSSKIADTKGCLKCRVVRNPYSGI
NPCQNGAQCFNLNTDYFCNCSEDYEGKNCSHLKDHCRTTPCEVIDSCTVAVASNSTPEGVRYISSNVCC
AFIGFLKERVLRDSSVALAQQVVLPENALFTSIQSLQSESEPHSLIANVIAGSSALGYVSHELQASRQLSEVASA
GRANSGEHGTLKHEKRTNGIRRNSRQMDSGLPKMQVIRNYNGIPQPAPHDGPPLYIQTGDTIELIKGDAH

VHIYRWDQAGLTSVAVLQGALGNPLGRFGAAITALADLNGDGWADVAVGAPLESEEHGAVYIYNGHQREI
RLESERLCTSKRMLLGLRVNKQLLELTADSQTQNINSEQEQQLSILDQLMKGPIVTYLGGIPDVPVGATLLTAF
SYVAGGSADGVLYIWNVLTEKVERTLAKHHSSSINAVAWSPAGAHVVSVDKGNKAVLWSEF

GPRGLPGLPGIPGLPGPPGPPGPPGPGMPLALKSEPTSIPEANGCSAHWKNFTEKCYYFSSEREVLEDAKVI
LLGYGVPALIVGISAATYHQGYGTARHCWLSLENQFIWSFLAPVCIIIAVNAVMFVVTVWKLSLKFADINPNMS
LVVIDNRKCKEAYAKLPAGKSLLITENMLCAGDEEGGKDSCQGDSGGPLTFFDNLTKKWFVGGIVSWGVE
SSLVKTLSPALNSVIWLPCPQLSALANYNWSYPGDKIPAGLTLLDNKTLVIIMQRQAAGEYTCWASENGHRC

QNDLVHWENEDNVTRTATLLDTSVIKGPINNMLLEEKHSYEYADENVIENIRKKFQESTIHNKMYPLKKKRE

LVLGTTASTPGSQSIVTSPLFLNHAGLPLLSATPGGVGLVSAAVAASLSSKSPSLTSASSSSSSSTCSSCSSSSEA

SRKVTGRS
ISHNKTTRPKMSLVMPAMALNESLYNASRGFEVMMQIDCEVMDTRVIHIKSSTVPPSLRRPPAPDNRTQAPS

QMGMVLALDYDPVESKIYFAHTALKWIERADLDGSNREKIICEAIDIPEGLAVDWINRKLYWTDRGKSYIGR
EQLKEKLLFAIEETEGFGQE

QDSHKPGMVISINMKTQFSSFVALSSIDKAIYGITGTRKRSMERVILDLEKSDLGCGAGGGQNNVDVFRSAG

GNSQMTSHVKTLENTKIIVNHEDTLEESEELSSDEEMKMAEMRPPLIETSINQPKVVALSNNKKDKDADSLSD
LSVKEKVEDSSGHHRAYGVCFVDTSVGKFYVGQFPDDRHCSRFRTLVAHYTPVQILFEKGNLAADTHKILKC
PKGQGECDPLEHEPAVSPLLPRKERVALESSLNEDERLLPKDKKTNLFSALIKKKKKTAPAPPKRSSSFREMDC
THPIPLESGGSADITLRSYVVAQGPFSDASPAPAVASQQPSCRSDLLSQHYFSSFVPGPFQPTSPVEGAVASS

KKLSSNCSGLEGDVTDEDEGAEMSQRVGKEGAEQQNTVIVKRFQLCKELSELVSICKSVQFKEFQVSFQLQ

FQAIMEHTVSYPKSLSREAVSICKGFLTKHPLKRLGSGLAGEQDIREHGFFRWMDWESLERLEIPPPFTPRPC

LSEGDSGVYQCLSEEKVKNKKFSQVVAKHVLELKKIQPSTVDPIQPAAQTEGNTDVPKMSTVPTEGSTAQTST
QCVCGQCQCNSNVRGQHCECDDTSCERHNGQLCAGQGRCQCGSCFCNEGYTGSACDCSLDTRA
VYPLHENKIGTPHSIQAYVQEGRPSVGPVAKTNNVGKNEAMIKWNKIPKDKRNGFIINYTIFYKPKGGKELNE
QQELHFNKIHPNQSGSYHCEPQNRVGMSESPAITVDVQYPPKEVRITLENLQLIREGDAVTLNCSVGSSNPP

ISDFSSNLLSTPFLSEPSSWSSLSTFYSRMPSDLSTLLPQVFPTLLGPSVVFDDSSSWDSTEFPAADSTSIEVSH
AAGMGMGMGMGKDMGSQYGSANPAWAAAQQRAHPAMSPGSTGQTINRTQGSPMDPMVMKRPQLYG

VQNWTYIVKVSAQHGADEDRIVSNISDTRFTVKGLTYSTAYQVSVRALSPAGEGPWTESFRGATLKEVEEEPY
QSKEREGQVDQPEPTCTLNFPLQHNHTAADLYLSPVRSPKKKGSTASLNPTPIIDAQPVLAPQTQKPQKSTSL
PPERPPPIPPDNRISRHLHHPESVPSRDQPVPPEAWCPRDVFGTSPLIGCRILNDSSPKPGLTASSNINGRH
NVSHSLPPRGMAPHPNMPGSQMRLPGFAGMMNPDMDSPSVPNPTSRPGLSGVSWPEDVPKIPDGRNF
LIFIFHKYKKQFRYESQMQMIQIVGPSDNEYIYIDFSEFEYDLKWEFPRENLEFGQALGSGAFGKVVNATAYG

GLMQCALRATKESTLKSVLSALWNLSAHSTENKAAICLVQGALGFLVSTLTYKCQSNSLAIIESGGGILRNVSS
TNPGSWTSKSRFQHSTVTSLLVSGPSDKTIVFSGEDMDLTRSYVEKGIASQLPPVSAGMSTSVTFKPQSFAN
LRIVEDEEYETTQEYEPAQEPPKKLVNSRRAKRTKPNGHISNRLEVDTDTSSESSSSESETEDERIGEDTPFLS
NDLVAENKILEAELKMAQKTNRKSQKKICLLKQQLDEAMEKEVAAHQYLANLISLAENITQERDHLIHLAKCLE

RKHQELQAMQLELQSPEYKLSRLRTSTIMTDYNPNYCFAGKTTSISDLKEVPRKNISLISKFNHQNIVRCIGVS
DGQPRQAKGELGLSQMLHIASQIASGMVYLASQHFVHRDLATRNCLVGANLLVKIGDFGMSRDVYSTDYY
PSPLFGFRPANFSPINASPVIQRAAARSRHVSASTPKGAGGGVLSPEMLHHAPHRERHSSGIVPTFQTNLTFT
ETPPPSPQIPGGSEATPKAAPSSPEPSEATPLGSASPETVSSRTEWQVSLKEPACFAVKQERGSMGDSGLRTG
ILLLPGSSEQFDINCLCMGVEQTLRGVPNSTCNKAEENSERNDKSSKNAKVEKTSKDKKQIQTSPVPVRKNG
CQYDIDECASTPCKNGAKCIDGPNTYTCECTEGFAGAHCEIDKNECDPDPCHYGTCRDGVAAFTCLCQ
SKTKEPPLKKAKKNNSNSSNSSSKKEPVMVKNSKPPPVYESEEEEKCKPMSYEEKRQLSLDINKLPGEKLGRV
WNKGVERGKQIAAEIMLETIFSILSTWTDINMKNFLNQLHQFYVVTKTPMVYENEPKEWALLEFGEKQVKELI
PPLRKASESREKLGTRGDGLATGKSSFSITREAQVNVRMDSFDEDLARPSGLLAQERKLCRDLVHSNKKEQ
SVEVKQQSNFLDLETPCLKSNELRMTIIKKLEREKTVSPFSQLNDSPTFKKLDESKDGSPDLKSELTGNDCHD
NTEGPVKLHEGSQVLLTSSNEMGTIRYIGPTDFAPGIWLGLELRSAKGKNDGSVGDKRYFTCKTNHGVLVRP
DGYTLTLYHAQAGTVAANASVRKDAHKFTFSGLAPGQKYLLEVASVAGPYRTFAGNISDWTMNFSAMPGESP
KNCRVCSDCGSRPAALDPGCHWYENYSVCEGCQRQRSAAGGTDATRCCQSCQKPLPLDSDATGGSTG
YSTADTSIEEKKQTESDSLTSFSDDGTSDLDTGDQSSEAACSALEISDSADKEEKEFCTGMRGNSKCSMYLS

LCQRDTGDEDSVAGELDRTEEGTVNGRNFGPGEPFPGLFPRKPTPITSPSLNNSSKRIKIEKDLDLPPATIIPS
PTIPLASPFLPTSAAPMQEKRKSILREPTFRWTSLKHSRSEPQYFSSAKYAKEGLIRKPVFDNFRPPPLTPEDVG
PQYEPNSTEALDMDIVSVPSSVPEDIFETLESAMETQSAADEQGNGSTATELETVNMNIKCTTASKDIKGGTK
QPSVEDKWLLRKKAQDYGLPTVCDLFACMKLNGDREKWLYQTPLQK
NGTRCLSCHPECLPQNGTETCHGSEADQCMACAHYRDAQYCVERCPSGVKADSSFVPIWKYPDEHGVG
CLVGRNTGRKKSGEGSTVVNGSPMGVKNGDSIPAELPLQEGKSILKNDTEKVTCNNTDGEPAQNTCDRKS
TKEQLYQQAMEEAAWHHMPHPSDSERIRQYLPRNPCPTPPYHHQMPPLHSDTVEFYQRLSTETLFFIFYYL
HSGPVLCVVMSSNGEQCYSGGTDGLIQAWNTTNPNIDPYDSYDPSVLRGAFVGHTDAVWGLVYSGAHQ
NKRPQRRKQRVLKADLENKPVNGLRPELMECSICGHGEKYRVKTNQTVSFENVQSNEKESMTGLEAEKWT
AQNAARIPGIPRHNGPQYSMMQPHLQRQHSNPGHAGPFPVVSVHNNTINPTSPTTATMANANRGPTSPSV
RAELQPPPEEPKTVIKHSRPVSSQPENPAVPSNLQSLVALSPAAIHPLSTQLVGSNGNLSPASTETNFSELKTM
MAFLAQQAIKQWQISSGQASAASSTASSTTSTPSSPTVTSAAGYDGKTFGPPMIDLSSPVGSSYILPSLPDIEC
VISDGGIPPMSSTNTLTIRVCGCDSNGSMLSCNAEAYILNAGLSTGALIAILACIVILLVIVVLFVTLKRQKKEPLIV

IKRWSDRQVPSTEPCTSGSSHSKSFDQLKCGQYLCSGDTADSVSVTSAGSSSSDVEEINISFIPESPDCQEI
KEGWLHTGDIGKWLPTGTLKIIDRKKHIFKLAQGEYIAPEKIENIYIRSEPIAQIYVHGDSLQAFLVGIVVPDSEV
SGTVDEDSFGTRKARSSFGRGFFKIKSNKRTASAPNLDRSRSASAPTLAETEKGSADHLDLAGLPPRPKETDS
TICPVAEGVAAEGSVAQMACIDLSCDSGSQTAATLSVQSETSSHTLGQEALTEIKEKLSGEASEFEAVPETLCE
ETCLPSVYEIVPRSAPLEGGTRLTVCGWDFGFSKNNRFDLKNTAVLIGGKPCTLELKVSNKNKLECTVAAAKN
MAVMTPVAEAPNYQGPPPPYPKHLLHQNPSVSLYETGTKPSKEDHPNLPKEEESEKNNECVDGTDKEKKQI
SEDNADGEGLRDAPMGTKCCCQGDNLVEAGALESGLNGHGQQDGCQGDTVKGQPQAGWEPLGDIPS
SEGISNAEVAPNGSPVSGSPKMSTLSNKGRYRCKPRHRRGNSKGSYNEFAGILKNQPAHDNSPQPPPSHH
NADHFMQEAKQKKHKADAMVEKFGKALNYAEAALSFIECGNAMEQGPMESKSPYTMYSETVELIRYAVRLI
ENSAFKKQEQHRKNNESKSYYHGPSRLPLHSVIASMGTVAIDLQDTGCRSSSGPTLTLPSEGSKNGRRSSIA
FNGSLECDMESIIRSELMDADGVGCQADGFLRKGEMGKAGP

YQSVKLYCSKCNSLQEVPNDDDFDLILRDCAATAPNPEVHNSLWYDSAVWDTENQGQRKIAVHFVEHDE

PVEGSCQQALLSVPWLEKTPARVRAPSRSRDVQCDEQTSCPDGSTCCRLSTGTWGCCPLVEAVCCQDH
LGIFADLQRAMKGFVLENDLHILYLVTPVYEEWTTIDWYQFFCLWEKLPSSMKRVAELVGIEEGFLARSVKGK
EHSEICFNHSCPDDQFKCQNNRCIPKRWLCDGANDCGNHEDESNETCAARTCQVDQFSCGNGRCIP

ENRTTVMIKDRWRTMKKLGLH
GKAHGEVLPNPGKTHQRAQRKPPHVDVPMGDERCMDANVVGQNPGVELVSAQRAKKSRLEQVGPAQEP

VVGTLIPLVRDQSEIQKQVLGLLKYLVIDNKDNENLYQTIKHLDPFPEHPAFEDLRATQQKIKYSKGAFSLLEE

LLEKELREVVQDINKLASAAISLDGALTKAQYHLRHGSSEQRSSCRAMADNLCLELESVQNLLGTKQNEAEA
HLEQAFITSILPNVTFDLWIGLHDTKQEFQWVQSEPLRYVNWAPGEPSGYGTSTANNKPTNCVLVWHGSPE

MNSLVESSPLDNGPPLFFPQDVIKKINEIDSASYSQSRARYGSWWDGFELDSKNADAWSSSEQESVFQSP
CMENKLQEVQACSENSESITGQLMNSEHMKDNGANKEENKEKNEITSSVETLRKCLANRFAKEQISKRLD
RIQLTYSNGDSCGNNNMIKTIITLMCKPGDLESAPVFNLENDECLYVFEWYTAAACVLSKTEGNNCRVSDA

NLTFFQFLSLKSNKISSISIGDLDVLQGEKGLNLTLVLSMNDIKYIQLGSFEEIHLHELSLRACFENTSVMKACIC
QEAVVTCLNKQTREIADETLCVTSRRPPQLLKACNLEPCPPRWEIGKWSSCSLTCGVGLQTRDVFCSHLLS

DNDDATKRKVNLVFEKIQTLKSRAAGSAQGSNSSTEVKALLEQKTELETKVAELQKKLDLEIRNQQNVKEERD
PGAGQPLCSQRGECLCGQCVCHASDFGKVTGKYCECDDFSCVRFKGEMCSGHGQCSCGDCLCDSD
AVASKGRKTANSQGRRKGRITRSMANEANHEEAAAPQQNTELASLEMNESSRWTEEEMETAKKGLLEHGRN

EWFNSKISAWHNDLEKCLPNTKATKIGVLLNYEGGFVIFLGVGEKHNLIYKFKAQFAEALYPAFWVFSSGTM

GLDSAVLYLHVTCPLEHVHLSLPFVTTKNKEVNATAVLWPSQVGTLTYIWWFGNNTEPLVTLEGSITFTFSVEGE
LRDEVQNSEEELQVLRPNKENTGEVSESGTCIIKHGLNAEKIFMQVHYLKGYFLLQSLAKRIGEATYFEFLRKE
HLAIIHLHSELVRNLLEVVLDFNLDDIINMRNDLYQAPLHLAVITKQAEVVEDLLRAGADVSLLDRHGNSILHL
DPGLSTFTAQLPAPQPLAPSAGHSTAGGQGEKKPYECTLCNKTFTAKQNYVKHMFVHTGEKPHQCSICWRS

LEGAVDTGSEGDRRRALETSLSSGVVCSLTAYVGVDTERGQPVQGPLVRRDIPLAGFGAASHVWKMQGFS

CSLNRFANHGFKSFEMLYRTTRCDDSNLLKNFVEAFQSALGKMVPLFCNDVDDIDCRLEDLPQKHCLEYN
VKVNSAGDKVGLQMDTIGVYTSLGLNERLFVVNIQELHTLTPHPEQLGPTVGSSETLDFISSKDLANQLTDYD
DELRSHVQKGTLGKFTVPVLKEACRVYGLKGGGKKQELLDALTEYFSKF

DFQLRSFDQPPSLDSGCPSSQRVATSTIFQQTHTLSTAAEEITPVKTECVDDIKTLIVSQSPVLIINETNSAPPSP

GNASLAQMISGLLGQLLMQPVLVAQGSGSSSASSSSSSSSTTTTTSSASFPAPASPPTASASAATTNTATTAAG

ANKIIQDMEEQLRKKDSPCPSNVELLRVLLRHEKKQPGALPGVRRQPPWTRRELRAALSIPFLFRAVKSEKQP

MRLPRMFKQEFTGVGATLEKRWKFCAFEGIKTT

SASFSRGRGRGRGQARGGRGQISTSRGASRRGRGNTTQEQSNTSRNYKPVSTPTKSMNIIGAFKAAQQQ

GLSHQKTQLDSRNSSSTSSLASSEGKGIGGVSALNSSSSTQAVNGPVSTTQTTGSNTVSVQLPHPQQQYNH
GKHKTSIDLEYVPRKNNGAVIEGEVAGPRRFNMKMI
SKEKDHIDKLDSAVSAQEALKTETSPALLAERDSGREEQKLSNSSKCLAHSSGSQLGAENGSQVEVEISNQC

NISVQPESLSTTPQRKNSNKLSMRRSLRGQKKLGNSLFHQTRKAVCVC

HHNWPSLLRHRFLEEFITPIIKVSKNKEEFSFYSIPEFEEWKNNTPNHKSWKTKYYKGLGTSTSKEAKEYFADI
SERNSVTGGLEPSLPKTLNLSLSRKRRLKGESKVAKQSIDNKTEGLEKGCTHAGYGNSAKMLVLQDGSGLSC

ILEDHHDYLLITENGSFTQPLARLTGSELPLAINAGLYGNFRAQLRFISDFSISYEGFNITFSEYNLEPCEDPG

NKVTRALLRDKSEQVKSFVATHIANILDSEEVGIEDLKNKVQEALKGSQIPAANDFRKFSQNYQISKRISLPGN
NAVQAVLEGEVQNAVAIVRPPGHHAESDTACGFCFFNSVALAARFAQRLAGRPLRVMILDWDVHHGNGT
AVNSVGWSHDKRWLVSASEDRTLRIWSVRNTEPALFLDKEMFHKSIRFAQFYYIDTFILLCCGAKFHLLSYYL
TQRSQEVTALQAELQQLKDALDSEKESSSKSLETLQTRLGEKESHEQALQQRLLDEQFALLQCTVQEAEQMV

FNSSNTAGKNMKYKLLTVLKLKCHSLFLNLQVNSLRTVFINIYKIFLLQAYRFHACVLQLPFNQQVRKNPGFF
KGQGEYSKIEIFQTLPVREPSPPSIHGQPGSGKSFKLSITKQDDGGAPILEYIVKYRSKDKEDQWLEKKVQGS

DELVPSKDVSATEVFHKDPVANVQLFQEVPAGQSVEDMVGRPMVHLSAAKQACGEAVYCDDIPCYENER

EEDLFHQQGDPRAANRSCAIM
KRISLQLHLALTSNAAWVQCPSHLELMNQCRHINVRVDPRGLREGVHYTEVCGYDIAAPNAGPLFRVPITL
KMNPVAFDFLLQRVEKTLRHAIEEEEGLPLDQVTNFQEVCEEIKVKLNSLKDIPNRIECPLIYHLDVGAMYP

KGHLRLKMTYLPKNNGSEEENTEQVEDLEPGWIILDQPDAACQQQQESPPLPPGWEERQDILGRTYYVN

DICLKLSLPIESSRSDTGNIRRCLAHSLFMNAAELQPDGTYATVDSHQLVAIHPSSVLFHCKPACVMYNELL
VRLVFRVHIPQSNGRTLSLQVASNPIECSQRSAQELPLVEKQSADSFPVIGGKKIILSGHNFLQDSKVIFVE

AANPALAFGQGLATGMPGYQVLAPTAYYDQTGALVVGPGARTGLGAPVRLVASTPVIISSAAAQAAAAASAG

SSSEKKGPDTPIQVAEATFDFENERDSSVSKLQAEIICDPEIPHTDVKSTNIREPEERRTQRFQFSLQDFKC

NEWIGKNSSSWVFSRCSSNFVVRHNSKEMLVDVHPQMKRLGILLDYDNNILSFYDPASSLHLHTFNVSF
CPIIEVYGRTFPVEEYFLEDCIQMTQFVPPLKDKKKKDRDEEGGEDDDANCNLICSDEYGPETKRCMAQL

DHLLLPATTHNKTKRPKMSIVLPAINISENVINGQDYEVMMQIDCEVMDTRILHIKSSSIPRYLREQQGNQT

NSLIGKYSKGKEKYFWTGLQDISRSGKYTWGDADGRKNEAVMYANWNSLEPAFPGGCVAMSSGKSLGK

PEPEPLSAVAQLHLSSGKNPISILMEYGQKSGSACEFQLLSQEGPPHDPKFTYCVKVGEQTFPAVVANSKKG
ANIERNLRGEDGICSVLVALMAGKAEVRYDPAIIQPSVIADLIRELGFGATVMENYDGDGILELVVRGMTCA

VTS
GKAHERPGNLYVSAGKVKKEPAVRLVRDQSQSSIYDPFAGMKTPGQRQLITLQEQVKMGILNVDEAVLHF

CFVDFQDRHGNTPMHVACKDGNVPIVIALCEASCNLDVTNKYGRTPLHLAANNGILDVVRYLCLTGAN

AVVHELRNSQPPHSPEVQELLLKNKSLVDTIKALIQMGASVEAKDGKSGRSALHLAAEEANLELIRFFLDQPN

QDYKKGFGSVDDKGKGEFWLGNENIHLLTQKDTVLRVELEDWAGNDAYAEYNIHVGSESEGYVLSVSAY

NQKRQLHNPDAVRLHCINCNMAVCHGSDLRTVEKMHHVNINPDFKIYYNATSNNVEIPRNFKDWKPGG

GAEILHV

LLSLAGKSSQISAHNLLGLFEQRRQLQKRQVPLSIATNRLTERIIPRKFALSSKIADTKGCLKCRVVRNPYSG
NPCQNGAQCFNLNTDYFCNCSEDYEGKNCSHLKDHCRTTPCEVIDSCTVAVASNSTPEGVRYISSNVC
AFIGFLKERVLRDSSVALAQQVVLPENALFTSIQSLQSESEPHSLIANVIAGSSALGYVSHELQASRQLSEVAS
GRANSGEHGTLKHEKRTNGIRRNSRQMDSGLPKMQVIRNYNGIPQPAPHDGPPLYIQTGDTIELIKGDAH

VHIYRWDQAGLTSVAVLQGALGNPLGRFGAAITALADLNGDGWADVAVGAPLESEEHGAVYIYNGHQRE
LESERLCTSKRMLLGLRVNKQLLELTADSQTQNINSEQEQQLSILDQLMKGPIVTYLGGIPDVPVGATLLTA
SYVAGGSADGVLYIWNVLTEKVERTLAKHHSSSINAVAWSPAGAHVVSVDKGNKAVLWSEF

PRGLPGLPGIPGLPGPPGPPGPPGPGMPLALKSEPTSIPEANGCSAHWKNFTEKCYYFSSEREVLEDAKV
LGYGVPALIVGISAATYHQGYGTARHCWLSLENQFIWSFLAPVCIIIAVNAVMFVVTVWKLSLKFADINPNM
VVIDNRKCKEAYAKLPAGKSLLITENMLCAGDEEGGKDSCQGDSGGPLTFFDNLTKKWFVGGIVSWGVE
SSLVKTLSPALNSVIWLPCPQLSALANYNWSYPGDKIPAGLTLLDNKTLVIIMQRQAAGEYTCWASENGHR

QNDLVHWENEDNVTRTATLLDTSVIKGPINNMLLEEKHSYEYADENVIENIRKKFQESTIHNKMYPLKKKRE

VLGTTASTPGSQSIVTSPLFLNHAGLPLLSATPGGVGLVSAAVAASLSSKSPSLTSASSSSSSSTCSSCSSSSE

RKVTGRS
SHNKTTRPKMSLVMPAMALNESLYNASRGFEVMMQIDCEVMDTRVIHIKSSTVPPSLRRPPAPDNRTQAP

MGMVLALDYDPVESKIYFAHTALKWIERADLDGSNREKIICEAIDIPEGLAVDWINRKLYWTDRGKSYIGR
QLKEKLLFAIEETEGFGQE

DSHKPGMVISINMKTQFSSFVALSSIDKAIYGITGTRKRSMERVILDLEKSDLGCGAGGGQNNVDVFRSA

NSQMTSHVKTLENTKIIVNHEDTLEESEELSSDEEMKMAEMRPPLIETSINQPKVVALSNNKKDKDADSLS
SVKEKVEDSSGHHRAYGVCFVDTSVGKFYVGQFPDDRHCSRFRTLVAHYTPVQILFEKGNLAADTHKILK

PKGQGECDPLEHEPAVSPLLPRKERVALESSLNEDERLLPKDKKTNLFSALIKKKKKTAPAPPKRSSSFREMD
THPIPLESGGSADITLRSYVVAQGPFSDASPAPAVASQQPSCRSDLLSQHYFSSFVPGPFQPTSPVEGAVAS

KKLSSNCSGLEGDVTDEDEGAEMSQRVGKEGAEQQNTVIVKRFQLCKELSELVSICKSVQFKEFQVSFQL

QAIMEHTVSYPKSLSREAVSICKGFLTKHPLKRLGSGLAGEQDIREHGFFRWMDWESLERLEIPPPFTPRP

LSEGDSGVYQCLSEEKVKNKKFSQVVAKHVLELKKIQPSTVDPIQPAAQTEGNTDVPKMSTVPTEGSTAQTS
QCVCGQCQCNSNVRGQHCECDDTSCERHNGQLCAGQGRCQCGSCFCNEGYTGSACDCSLDTRA
VYPLHENKIGTPHSIQAYVQEGRPSVGPVAKTNNVGKNEAMIKWNKIPKDKRNGFIINYTIFYKPKGGKELN
QELHFNKIHPNQSGSYHCEPQNRVGMSESPAITVDVQYPPKEVRITLENLQLIREGDAVTLNCSVGSSNP

HLSPWQTSYLYSNTFSVPNTHNPEVISSSFYSYSSILLEGTSVLLMDSEVSFTSIFTEATSQLESSLFSLESAVPTL
GMGNNPHSQTQQSSPYPGQSYGPPGPQRYPMGMQGRTSGALGGMQYPQQQMPPQYGQQGVSGYC

ILAVGDNGLWKQPLDSFGPGLFFSPDIRNVSDMDWYNDTLYWSNSTGEVHMWSLTRRAGAIEKFHISDI
LSLFYKKVYRLAYLRLNTLCLRLLFDHPELEHLIWTLFQHTLQNEYELMKDRHLDQIMMCSMYGICKVKNVD
HNRMGSDPGFMRKHRRHDLPLEGAKVFSNGHLGNEEYDVPPRLSPPLTAASILPGIKCVVSVTNSLGEKLR
FPPGQGVFSGPGRGERFPNPQGLPEEIFQQQLAEKQLGLPPGLSIEGIRPGIEMNRMIPSQRHMEPGNNP
GISKTGVSVQVAVKMLKEKSDPSEEEALMSELKMMTHIGSHENIVNLLGACTTSGPVYLIFEYCCYGDLLNYL
SLIATREDYRQVLRDHNCLQTLLQHLKSHSLTIVSNACGTLWNLSARSPKDQELLWDLGAVSMLRNLIHSKH
KSSSSNLNEQEEMEITKCHAVIIDNQNTGITIIQQQMQPKTIPGENWNKEVNEAVSLLNLNKESPLSGMSDI
IQNPMATSLEAATACRLADSRTNPTSRFSTQEELQARLSSVIANQDPIAV
ESEKHGVLNKVIEGNVRLGRLEEKVRVYKKKAAGKLGDINLKMTEQEKEFAGKASQYQHEMKHLQRLLQDK

SLQALPRFILLELMAGGDLKSFLRETRPRPSQPSSLCMLDLLHVARDIACGCQYLEENHFIHRDIAARNCLL
REGPYPKGQYSAAWQRQRLARPAAKTGVCTLFTTHHATSAVKEERMLWIPPSFMTHHAISAVNEVGGHTM
VPMSPGKRKAEAPHGGPGGGADFQKSDSLDSGVESVSHTPTPSTPAGFRAVSPPGPFSSRSQQASPLLLLS
QPPAKRRLSVPRSPLARPLENGGSIISSTSFNGSAGPPACKRGCQELPCGNSCLEVHSIGSEDEDLELPLDG
GKDEEKRESRIKSYSPFAFKFYMEQHVENVLKTYQQKINRRLQLEQEMAKAGLCEAEQEQMRKILYQKESN
PGYTGHLCEFNINECQSQPCKNGGTCQDRDNAYNCLCLKGTTGPNCEINLDDCASNPCDYGKCLDKI
VVHIIQSREPSLKNSNPDEIEIDFETLKPSTLRELERYVTSCLRKKRKPQAEKVDVIAGSSKMKGFSSSESESTSE
LVNYLRKVARPFLDQNGANAPPKDVTVKSRLGLGLISVILGECYALPTCKEDLSRVCSLLCLEQKPQEVLLLEN
EFRSIFHHIQSAQSQRSPSELFAQHIVTIVHHVKEHHFGSSGMTLNERFTKYLKRGMEQDQAKNKKSPEIH
DSIKEPENLIKVKRDQLRTFCPIELNINGSPEGRSDDLATFCVSKTEDIAMSFDDSLGRSEESPLIKSNKFYVLP
PSRVTYRGINGAKLVDDIC
RTVLAVSWGQAPSQQDYCQLWLRSPGNNTLPQRHTLAGGQVQHFFWGLVPGRNYSVSLSCVAGPYWSS
CQACQRQQKHPQPDGAGPQEEMLLECDAKPLDPPVATSPTRPEEPPLDAKEDVARTQPAGPPAITAASLEP
SQNSNRRARKRSSRDHRTPASPVGGRSDAESVDGELQEAASDTEMPVIALECTSAMRNDHLSLAHKHPMI

SENVYSQWLVGYAASRHFMKDPFLGFTDSRQSPFATSSEHSSENGSLRFSTPPGDMLDGGLSGRSGTASG
GFASGFSTSGATAPARLFSALHSGTRFDMHKRSPLLRAPRFTPSEAHSRIFESVTLPPSVSRTTAGTSVTGISSP
LKASAKVRKELFVKLTPVSLSDSPVKAEDQDSNPVKGDQNADIIPKGENCDSGKENHYIGNEHSPENETVPS

CRLCPTNCSHSCTIRDEDGCPMDQKPSQVTSIIAGVVGALLALVLILIIVVCIKRRRQQERKHAMRRLLQETE
SIKKMRSSQTKRSRHSGRPVCSLQLVVDRNSSCPDQTEMQIDSYPSSEEPRKGDSEKKRVRRSRRLQLLTE
EGTKAQYLAAKALKKQSWRFHTKYMMWFQRHEEPKTITDEFEQGTYIYFDYEKWGQRKKEGFTFEYRYLEI

RLLSCSADGTIRLWKASEIAPALNIFNDNQEMGIPSSVDLVSSDPSHMVASFNSGHTCIFNIETRQRILTLES
DNEKPSFSICVNGDIHGTVTDNEHLQNFEEERAVSPSELVEPKKLFVQTSRNGIKNIHYPPSEADVSQTKVII
TAIELIPSVTNPENLPSLPDIPPIQLEDAGSSNLDNLLSRYISGSHLPPQPTSTMNPSPGPSALSPGSSGLSNS
PNLQPISALPTKTQKGMHSGTWKLSPPRLMANSPSHLCEIGSLKEVTLSPVSESHSEETIPSDSGIGTDNNST
TGNITLDSIVRKDGIVEQSQPKPPSNRTVQSPNSSVPSPGLLGPVTMTSVHPPIRSPSASSVGSRGSSGSSSF
VFEEEDVRENIITYDDEGGGEEDTEAFDIATLQNPDGINGFIPRKDIKPEYQYMPRPGLRPAPNSVDVDDFI

KKFWECTSLSSLDTSGIGSGSSSASSSSVSSTPVTASRTHKRSVSGISSYSSLSLPLYNQQIDDCCIIRVSLAVD
MPGWSRKRGFEGTYAELCKNTELTQAIMEDMVRLGKEGGLHSFEQVKAIYIHSDTFSIQNGLLTPTLKARR
GQMSPCSPDSKKKARGIKKLFGRLKRSQSTTFNPDDMSEAEFKRGGTRATAGPRLGWSRDLGQSNSDLD
EQGGERQAEEKQLNEEGSSDPLTVSEGLILLRETQEHKGQEVVAVESSEAFCKNSEKLCDLRHGLVQMDL
IPSFNITSSVSSGHGISSFNTFSYVNPVITSISPTYGPKFGGTLLTLSGKYLKSGKSREIYIGGKPCTLKSESDNT
TTSPVPVRKNKRDEERRESRIQSYSPQAFKFFMEQHVENILKSHQQRLHRKKQLENEMMRVGLSQDAQD
SATGQANIVRSHRDNLVEKRALQSTVNGHGKQDSGLEGKVRGQYKKEADWSEATLRERDLQLGSMKLSG
HHHHHPHSHLSSLQQHSHPHYHGHHPRLNIQGQDIANCANNLRYFGPAAALRSPLSNHAQRHMSGSS
THAGPNATPEDKQLAALCYRCLALLYWRMFRLKRDHAVKYSKALIDYFKNSSKAAQAPSPWGTSGKSTGTI
APVLEVADSSSLQACDLLSDQSEDEATPSDDDISTGLEYAKGYPPYSPYIGSSPTFCHLLHEKVPFCCLRLDK

LLQEPEDTLIMVQGATLKEIRKLSGRFKCIIPVRSTKEDLELLDLSAPFLLQGKIWHYGCKQCSNQKTIQNLSS

IVHCCPKGYTCDPAKGSCQQALLSVPWAPKSPAHIIGTAQSRGVKCNQTVSCEDGQTCCKSRMGSWAC
KIIAKTEKQHRQIAIHKRFFTSLALLDLINEIPLKDITKKYNCSRGQLQSLQQSAATYAGMVTVFSNRLGWHNN
KSWLCDREDDCGDETDEMASCEFPTCEPLTQFICKNGRCVSSKWHCDSDDDCGDGSDELGCVHSC

ALPRDPEGLPGRGNERAAVRSMAASGSGQHRAQEQKYVKTTPLQYTYKAPTPDLCAQVNSTRISKAMLRW

INHFLSVSFCDSLPFARLEGLNDLRKQLEQHKDQMKDLMKNFQEKPQDYVIVKLVVSLLQLAKTAVNHTGE

LGALQKSFMERKEQMLKSIEDLEERADKEGLKEPTLQGLQQRLKIFNQLEDELNSHQHELQWLMDKAKQ
PHFTGRWDDRNCLEEKHGYICQRSIDPALSSSRTSSLPSPSSPLSYLNSTYRILQKPLKWQEALLLCETRNT

DSWKDRKASPLLREHFDRRASDSVFLQKQPRQMEYSRTGLWENQFNSVRDKHSLELQNKTTEHSHGQ
GTQYLFLPPNRYIFHGAEVYSDSEDDVLSSSSCGSSSDTGSCHSPSLDVEDESEIEEFYNGIEEEEEDAPEF
QAGFSFDLSPLTKKSGSYVINTTDYDFYINICSSVPEEHCLLKAAAACQVAKNKEHTWSLGEPSSKLSYYDGI

QGLAGLQVKKLVLGEFRNIERVEDFSNGLLDGLCQVQLQEFVFISFQRFDNCTDTLFDCLVNTSTIRLVDIY
REMNETVILADELCHKPKPTLVQACNRFDCPPSWYPREWQQCSQTCGAGTQKRDVLCKQRMADGSFLI

TTRATLKDLQQQLDKSVQEKDALKKHLGESEKELRQNLEE
WTGEYCNCTTRTDTCVSSNGLVCSGHGHCQCGKCECTQAGSYGDTCEKCPTCPDACTIKKDCVECK
NWSAIARMVGSKTVSQCKNFYFNYKKRQNLDEILQQHKLKMEKERNARRKKKKAPAVQNEEAAFPPTAED

LSLCQLK

MNTITVQVSAGNTILQDTKTIAVYERFQSLRLSFSPNLDEYNPDIPEWRRDVSRVIKRALVEATSISSKHILVAV
FVQKFHGQLILSQDFLYMLLEDIPEQKGYGLSVENIIQAWLDTSGIPKPLLEECQTWDRCRLVQQVNDEVT
AAKEGDDKILNILLTHEKTSPMINYSNGEGLNVIHMAVMANSMPCLRQMIAAGADVNAQEQKSGRTALHL
FSLKDYLIKHMVTHTGVRAYQCSICNKRFTQKSSLNVHMRLHRGEKSYECYICKKKFSHKTLLERHVALHS

NTNFSPPIAMCCSASIPTDCDEDLCFESADCAEDCHFDDYSLDMPERAPEESPLLRLVSLQNADGSWDL

YDYENGFAIGPGGWGAANRLDYSYDDFLDIVQEEQLQMPLSGSVKTAPSRVKRHKKLNIPMTDHKIKLIF
WNLFSSVHQVELIYYIFGQQTFPSATMANLERVMRRFNELQYWIATELCLCVEPGKRAQLLRKFIKLAAHLK

YGGNRSRTASPIRAGKGTLDQTEKTYPGTPNLLTVTLSKRSAVMDEELNPKMLALHNSQRKRKMEYDGSLF

GSSAGAPGGPMPSASSSAQPPPPTDDLHFSQLLGNLLGAAGPGVAGVGSPTITVAMPGMPGMPTFLQG

LQKARELRPDPTLLLRYRWALDAAIHISARHNLPPLPGRTLILCCTSNAMETPYHRARELRGPRPEPMTALE

PPRNSSTKNYSEDIIDDDSDVEEVSFTAASKVTQRLSSTSTSSKRSSQSQITRGVDFESDEDEFDPFKSTATS
PNTATVAPFMYRSQTDNEGNVTAESGTGGVSMDTALYTKTGQPGLEDLSGEDPETRRLRTVKNIADLRQN

QKKWEKRRQSSQKQSENNEESSERMAKDDIKSRASSSSQDGLRDSPLSSISSSDYDSVSVATYSLSSIYALSS

MARHRIGFKYSGPEDDAAITLAFSKKKIDERKEWLTNFMEGRRQRKLHGLSEDFVYGKSTSYLTYNDFIHKE
CHRKACEAGRSGNKETSQQLISTEFTGLVDKHVALDFANSLMQVQSNTRELIKNTVQAQERDGTDDKSSSL

IPQFGNRIGFNFGVGDTLTFSCSSGYRLEGALEIICLGGGRRVWSAPLPRCVAECGASATNNEGILLSPNYI

IDPVSAKIEGNLIFDPNTYIPKETMLKTTLQLYGLKPMDIFEIGMEGKSFEPTVEALFGQKGFFPDSASKALYWY
TQHMFEEDPSVLYVSLHRYDHGAFFPTSEDADADRVGEGPGRGFTLNVPWNSPRMGDAEYLAAFHRLVLI
LDTSKDDLKRYKQKSICKSVQKFPMASTVEITSLSAINDFYSYIILAAGSNRTLEVFDLNAGCSAAVIPDAHSRS
VQDSLNRLDDPAHISCTSSADYLLCRTLAASDCIERLQGAHGKYLSNRSDVSSLLPCVALFAHLISDTILQGSA

FLRIISDTASCCYSIMKTKNTGIALGTKGASGLFPSEAAEWLCYRAFIIKLTNHKVVYKCLLRFLKICKYRVTLKVI
SRDHIILEHLQWTMGYEVQITAANRLGYSEPTLYEFSMPPKPNIITDTLFNGLGLGAVIGLGVAALLLILVVTDVS

LYLTLVTSTKAHAKIISIDTTEAQSVPGFVCFVSATDIPGSNVTGVAHDEPVFADNTVTCIGHIIGGVVADTHEHS

LIILARIDESSHYDLSYTDVHFKPGQIRRHFIEVPQGATWAEVTVCSCSTDVTAKFVLHAVQLVKQKAYRSHEF
NIILTNRLQPSAIVDEATCAACDFNKPGANCQRRMTWQWRGEFMPASRSEYHRIQQQLESEKLPPLYPDG

HESRRTQWKRPTAQDNLADVDNGNIHLEAQHAFTHRRQITEEAENLDNRDSPESWEVITEDEATMYSNQ

HTNKCYMRDLCVVDADWLYDAAPDYFRRKLRAAKN
KAPDGHHVWEMEAKTDKEMCKPNSLVVEIPPFRNQRITSPVQVNFYVCNGKRKRSQYQLFTYLPANVPIIKT

GTANNLAGATNGLFRPLGAQPQQPQQQTNSSLQSNSFYGSNSLTNNSQSSSLFSHGPGQPGSTSLGFGS

CAVLGRGHFGKVLLAEYKNTNEMFAIKALKKGDIVARDEVDSLMCEKRIFETVNSVRHPFLVNLFACFQTKD

LPVCPTFTIWNKSLMILSGLPAPDFIDYPEQQECNCRPQESPYVSGMKACH

NEKETPFELIEALLKYIETLNVPGAVLVFLPGWNLIYTMQKHLEMNPHFGSHRYRILPLHSQIPRDEQRRVFD

NPFYSAAPTVPETAHPMRAIVESLQ

WEAKDCKTFTALSICKNYTGPPRQPEVLPKPTDPCPAGWQNGSGLACYKFFHSERLLRTRTWEEAERFCE

AKQMAAEVAVKILRGEAGGQFFPEQPSGEVLSEPSLDPPVAQPLNPDESKAAKAKGVGELIKYLNANPISGLI
SCVHKIESTLMKTKGVLYSSVALATNKAHIKYDPEIVGPRDIIQTVEDLGFETSLVKKDRSASHLDHKQEIRQW

EWQLNQKKRSESFRFQQENLKRLRDSITRRQIEKQKTGKRSDLEITVPIRPSHNAPGKLECGIYEFGPRKN

EALTSDGKTAEDLARTEQHEHVASLLARLKKDTHRGLFIQQLRPTQNPQPRIKLKLFGHSGSGKTTLVESLK

ICLSFVNAKAYNGNTALHVAASLQYRLTQLDAVRLLMRKGADPSARNLENEQPVHLVPDGPVGEQIRRILK

EGTAGDALIIGSVEEGSEYTAHDNMKFSTFDRDSDQWEENCSEVYGGGWWYNNCQAANLNGIYYRGG

SISCASCGQAWGMEMIYRSVTLPILSIKNFVVATPDGNKQPKQWSRVTFTIEEFDYVEYCNRNLGM

NTFLCAALPSSLVLLQWYEPLQKFMLLKHFPAALPMPLSLFELLVVETEEYPQVCVRVVDRDRPGQELEFNVL
GPHGKCRSQAGGKFTCECNKGFTGTYCHENINDCESNPCKNGGTCIDGINSYKCICSDGWEGIYCETN
LRSFSPLQSAQQTHSGFQTSGSHVEGAPPRAHSTMTASGTDARTASSGSSAWEGQLQSLILSEYASTEMSI
ISRFWQGRNLVTGELGYFPSDAVKPCPCVPKPVDYSCQPWYAGAMERLQAESELINRVNRTYLVRHRTKE

NTHYSQRIEGATISSGVKYFGQSIHGQIDLSGDGLPDLSVGALGEVIMLRSQPILTVVPKMSFSPEEIPVKQV
YNGCMEVKVNDIQLDLDEAFSKHNDIRSHSCPLIANEMNEFLN

FEDAKVFCDEKSSHLVIINNKEEQQWIKRQIAGKGNFWIGLTDSAQENEWRWLDGTLPEYTNWKSGQPD
SQLKKLRVLTITAIAQLCILGITWIFGMFQFNQRSLVASYIFTILNSLQGLFIFLLHCLLKKQVRDEYRRWLSCG
CGVAGQYGVYTKVINYISWIENTILKNS
QAVARYVVQDPTGVNILSESGGDSMVGEGLEPEGSPKDGHHSYWAQFVTVTVLLSVTLAIAMAMAGFSYHD

ETSKME

AAGPPPGHTGPAASEPDGKTA

SVPPRPGRGPLRSSRHPRTFYLGEQGEG

SELNGMHREMIIKEDIFQPRGIAVHPLAKRLFWTDLGVNPRIESASLQGIDRLVIASIDLVWPSGITVDCLAD

GLTFLTNANAGDSKVADESCKAIVRPRRSSFEEEIRNKASKYLHPEIRKCCMSGVKESPLGQTCDQRVEHIKS

DEATHNSNHNNSNCSSPSRMSDSVSLNTNSSNDTSVCSPDKEIHSAAISKIRQEDENFNSLLQNGGEFNI
GSLSSSMQEGLISGSQFWNASKTLKTLIEEGYFEDRQNVDNRFILPPVIKSMTSESDSLGLTPGENCELALSA
GHLDRKGAGEDESRDASNGAFIMPPADAMDASKCQGPSNGAGVTNGAVTGNSGFLSPHLWKKNSAMT
SSSTGHPLYHRMEGTLSARSRNNSTERLLESPSVSVDDYHESDGSRNRTRAVENQYSFY

QKYWEVCSFNEVLASKYANENPGDFVNYNKRFLARVFPSPMRIDSSNMNPQDFWKCGCQIVAMNFQTP

CGRSGENFDKFFTRAVPALTPPDQLVMAGLDQSEFQGFTYINPDFVHLSLRPPTSPISMV

TTQIIPITTGRIIRKPIGPILASATSNPEIFNSVPDGVPEKAMFLKSNDNFLLIFLFLFFFVLFLSLLSYNCYKGYLPI
ACLQDGVECSGHGSCICNKCQCQPGYFDRLCSRCPSCRTPCEEHRDCADCQAFRMGPLSQNCSIACI
ETVNSAVLQYRLKSLQANTQYTAHVMAATKAGGTYGNRITFSTLKFSKEDFIFVTISFGFCLLLLLILGITCTLKKI
PVTKYTWYKDDTQCQETQESVLTFPATKERSGSYRCEAQNAIGYRQSLPVSVDVQYAPQHVAVVRQPSGSI

HLSPWQTSYLYSNTFSVPNTHNPEVISSSFYSYSSILLEGTSVLLMDSEVSFTSIFTEATSQLESSLFSLESAVPTL
GMGNNPHSQTQQSSPYPGQSYGPPGPQRYPMGMQGRTSGALGGMQYPQQMPPQYGQQGVSGYC

ILAVGDNGLWKQPLDSFGPGLFFSPDIRNVSDMDWYNDTLYWSNSTGEVHMWSLTRRAGAIEKFHISDI
LSLFYKKVYRLAYLRLNTLCLRLLFDHPELEHLIWTLFQHTLQNEYELMKDRHLDQIMMCSMYGICKVKNVD
HNRMGSDPGFMRKHRRHDLPLEGAKVFSNGHLGNEEYDVPPRLSPPLTAASILPGIKCVVSVTNSLGEKLR
FPPGQGVFSGPGRGERFPNPQGLPEEIFQQQLAEKQLGLPPGLSIEGIRPGIEMNRMIPSQRHMEPGNNF
GISKTGVSVQVAVKMLKEKSDPSEEEALMSELKMMTHIGSHENIVNLLGACTTSGPVYLIFEYCCYGDLLNYL

SLIATREDYRQVLRDHNCLQTLLQHLKSHSLTIVSNACGTLWNLSARSPKDQELLWDLGAVSMLRNLIHSKH
KSSSSNLNEQEEMEITKCHAVIIDNQNTGITIIQQQMQPKTIPGENWNKEVNEAVSLLNLNKESPLSGMSDI
IQNPMATSLEAATACRLADSRTNPTSRFSTQEELQARLSSVIANQDPIAV
ESEKHGVLNKVIEGNVRLGRLEEKVRVYKKKAAGKLGDINLKMTEQEKEFAGKASQYQHEMKHLQRLLQDK

SLQALPRFILLELMAGGDLKSFLRETRPRPSQPSSLCMLDLLHVARDIACGCQYLEENHFIHRDIAARNCLL
REGPYPKGQYSAAWQRQRLARPAAKTGVCTLFTTHHATSAVKEERMLWIPPSFMTHHAISAVNEVGGHTM
VPMSPGKRKAEAPHGGPGGGADFQKSDSLDSGVESVSHTPTPSTPAGFRAVSPPGPFSSRSQQASPLLLLS
QPPAKRRLSVPRSPLARPLENGGSIISSTSFNGSAGPPACKRGCQELPCGNSCLEVHSIGSEDEDLELPLDG
GKDEEKRESRIKSYSPFAFKFYMEQHVENVLKTYQQKINRRLQLEQEMAKAGLCEAEQEQMRKILYQKESN
PGYTGHLCEFNINECQSQPCKNGGTCQDRDNAYNCLCLKGTTGPNCEINLDDCASNPCDYGKCLDKI
VHIIQSREPSLKNSNPDEIEIDFETLKPSTLRELERYVTSCLRKKRKPQAEKVDVIAGSSKMKGFSSSESESTSE
LVNYLRKVARPFLDQNGANAPPKDVTVKSRLGLGLISVILGECYALPTCKEDLSRVCSLLCLEQKPQEVLLLEN
EFRSIFHHIQSAQSQRSPSELFAQHIVTIVHHVKEHHFGSSGMTLNERFTKYLKRGMEQDQAKNKKSPEIH
DSIKEPENLIKVKRDQLRTFCPIELNINGSPEGRSDDLATFCVSKTEDIAMSFDDSLGRSEESPLIKSNKFYVLP
PSRVTYRGINGAKLVDDIC
RTVLAVSWGQAPSQQDYCQLWLRSPGNNTLPQRHTLAGGQVQHFFWGLVPGRNYSVSLSCVAGPYWSS
CQACQRQQKHPQPDGAGPQEEMLLECDAKPLDPPVATSPTRPEEPPLDAKEDVARTQPAGPPAITAASLEP
SQNSNRRARKRSSRDHRTPASPVGGRSDAESVDGELQEAASDTEMPVIALECTSAMRNDHLSLAHKHPMI

SENVYSQWLVGYAASRHFMKDPFLGFTDSRQSPFATSSEHSSENGSLRFSTPPGDMLDGGLSGRSGTASG
GFASGFSTSGATAPARLFSALHSGTRFDMHKRSPLLRAPRFTPSEAHSRIFESVTLPPSVSRTTAGTSVTGISSP
LKASAKVRKELFVKLTPVSLSDSPVKAEDQDSNPVKGDQNADIIPKGENCDSGKENHYIGNEHSPENETVPS

CRLCPTNCSHSCTIRDEDGCPMDQKPSQVTSIIAGVVGALLALVLILIIVVCIKRRRQQERKHAMRRLLQETE
SIKKMRSSQTKRSRHSGRPVCSLQLVVDRNSSCPDQTEMQIDSYPSSEEPRKGDSEKKRVRRSRRLQLLTE
EGTKAQYLAAKALKKQSWRFHTKYMMWFQRHEEPKTITDEFEQGTYIYFDYEKWGQRKKEGFTFEYRYLEI
DRLLSCSADGTIRLWKASEIAPALNIFNDNQEMGIPSSVDLVSSDPSHMVASFNSGHTCIFNIETRQRILTLES
DNEKPSFSICVNGDIHGTVTDNEHLQNFEEERAVSPSELVEPKKLFVQTSRNGIKNIHYPPSEADVSQTKVII
VTAIELIPSVTNPENLPSLPDIPPIQLEDAGSSNLDNLLSRYISGSHLPPQPTSTMNPSPGPSALSPGSSGLSNS
PNLQPISALPTKTQKGMHSGTWKLSPPRLMANSPSHLCEIGSLKEVTLSPVSESHSEETIPSDSGIGTDNNST
TGNITLDSIVRKDGIVEQSQPKPPSNRTVQSPNSSVPSPGLLGPVTMTSVHPPIRSPSASSVGSRGSSGSSSP
VFEEEDVRENIITYDDEGGGEEDTEAFDIATLQNPDGINGFIPRKDIKPEYQYMPRPGLRPAPNSVDVDDFI

KKFWECTSLSSLDTSGIGSGSSSASSSSVSSTPVTASRTHKRSVSGISSYSSLSLPLYNQQIDDCCIIRVSLAVD
VMPGWSRKRGFEGTYAELCKNTELTQAIMEDMVRLGKEGGLHSFEQVKAIYIHSDTFSIQNGLLTPTLKARR
SGQMSPCSPDSKKKARGIKKLFGRLKRSQSTTFNPDDMSEAEFKRGGTRATAGPRLGWSRDLGQSNSDLD
EEQGGERQAEEKQLNEEGSSDPLTVSEGLILLRETQEHKGQEVVAVESSEAFCKNSEKLCDLRHGLVQMDL
NPSFNITSSVSSGHGISSFNTFSYVNPVITSISPTYGPKFGGTLLTLSGKYLKSGKSREIYIGGKPCTLKSESDNT
TTSPVPVRKNKRDEERRESRIQSYSPQAFKFFMEQHVENILKSHQQRLHRKKQLENEMMRVGLSQDAQD
SATGQANIVRSHRDNLVEKRALQSTVNGHGKQDSGLEGKVRGQYKKEADWSEATLRERDLQLGSMKLSG
HHHHHPHSHLSSLQQHSHPHYHGHHPRLNIQGQDIANCANNLRYFGPAAALRSPLSNHAQRHMSGSS
KTHAGPNATPEDKQLAALCYRCLALLYWRMFRLKRDHAVKYSKALIDYFKNSSKAAQAPSPWGTSGKSTGTI
APVLEVADSSSLQACDLLSDQSEDEATPSDDDISTGLEYAKGYPPYSPYIGSSPTFCHLLHEKVPFCCLRLDK

LLQEPEDTLIMVQGATLKEIRKLSGRFKCIIPVRSTKEDLELLDLSAPFLLQGKIWHYGCKQCSNQKTIQNLSS

IVHCCPKGYTCDPAKGSCQQALLSVPWAPKSPAHIIGTAQSRGVKCNQTVSCEDGQTCCKSRMGSWAC
KIIAKTEKQHRQIAIHKRFFTSLALLDLINEIPLKDITKKYNCSRGQLQSLQQSAATYAGMVTVFSNRLGWHNM
PKSWLCDREDDCGDETDEMASCEFPTCEPLTQFICKNGRCVSSKWHCDSDDDCGDGSDELGCVHSC

PALPRDPEGLPGRGNERAAVRSMAASGSGQHRAQEQKYVKTTPLQYTYKAPTPDLCAQVNSTRISKAMLRW

INHFLSVSFCDSLPFARLEGLNDLRKQLEQHKDQMKDLMKNFQEKPQDYVIVKLVVSLLQLAKTAVNHTGE

ALGALQKSFMERKEQMLKSIEDLEERADKEGLKEPTLQGLQQRLKIFNQLEDELNSHQHELQWLMDKAKQ
PHFTGRWDDRNCLEEKHGYICQRSIDPALSSSRTSSLPSPSSPLSYLNSTYRILQKPLKWQEALLLCETRNT

PDSWKDRKASPLLREHFDRRASDSVFLQKQPRQMEYSRTGLWENQFNSVRDKHSLELQNKTTEHSHGQS
GTQYLFLPPNRYIFHGAEVYSDSEDDVLSSSSCGSSSDTGSCHSPSLDVEDESEIEEFYNGIEEEEEDAPEF
QAGFSFDLSPLTKKSGSYVINTTDYDFYINICSSVPEEHCLLKAAAACQVAKNKEHTWSLGEPSSKLSYYDGI

QGLAGLQVKKLVLGEFRNIERVEDFSNGLLDGLCQVQLQEFVFISFQRFDNCTDTLFDCLVNTSTIRLVDIY
REMNETVILADELCHKPKPTLVQACNRFDCPPSWYPREWQQCSQTCGAGTQKRDVLCKQRMADGSFLI

DTTRATLKDLQQQLDKSVQEKDALKKHLGESEKELRQNLEE
DWTGEYCNCTTRTDTCVSSNGLVCSGHGHCQCGKCECTQAGSYGDTCEKCPTCPDACTIKKDCVECKK
NWSAIARMVGSKTVSQCKNFYFNYKKRQNLDEILQQHKLKMEKERNARRKKKKAPAVQNEEAAFPPTAEDI

LSLCQLK

MNTITVQVSAGNTILQDTKTIAVYERFQSLRLSFSPNLDEYNPDIPEWRRDVSRVIKRALVEATSISSKHILVAV
FVQKFHGQLILSQDFLYMLLEDIPEQKGYGLSVENIIQAWLDTSGIPKPLLEECQTWDRCRLVQQVNDEVTI
LAAKEGDDKILNILLTHEKTSPMINYSNGEGLNVIHMAVMANSMPCLRQMIAAGADVNAQEQKSGRTALHL
SFSLKDYLIKHMVTHTGVRAYQCSICNKRFTQKSSLNVHMRLHRGEKSYECYICKKKFSHKTLLERHVALHS

NTNFSPPIAMCCSASIPTDCDEDLCFESADCAEDCHFDDYSLDMPERAPEESPLLRLVSLQNADGSWDL

NYDYENGFAIGPGGWGAANRLDYSYDDFLDIVQEEQLQMPLSGSVKTAPSRVKRHKKLNIPMTDHKIKLIF
WNLFSSVHQVELIYYIFGQQTFPSATMANLERVMRRFNELQYWIATELCLCVEPGKRAQLLRKFIKLAAHLK

YGGNRSRTASPIRAGKGTLDQTEKTYPGTPNLLTVTLSKRSAVMDEELNPKMLALHNSQRKRKMEYDGSLF

GSSAGAPGGPMPSASSSAQPPPPTDDLHFSQLLGNLLGAAGPGVAGVGSPTITVAMPGMPGMPTFLQGM

RLQKARELRPDPTLLLRYRWALDAAIHISARHNLPPLPGRTLILCCTSNAMETPYHRARELRGPRPEPMTALE

PPRNSSTKNYSEDIIDDDSDVEEVSFTAASKVTQRLSSTSTSSKRSSQSQITRGVDFESDEDEFDPFKSTATS

PNTATVAPFMYRSQTDNEGNVTAESGTGGVSMDTALYTKTGQPGLEDLSGEDPETRRLRTVKNIADLRQNL

QKKWEKRRQSSQKQSENNEESSERMAKDDIKSRASSSSQDGLRDSPLSSISSSDYDSVSVATYSLSSIYALS

MARHRIGFKYSGPEDDAAITLAFSKKKIDERKEWLTNFMEGRRQRKLHGLSEDFVYGKSTSYLTYNDFIHKE
CHRKACEAGRSGNKETSQQLISTEFTGLVDKHVALDFANSLMQVQSNTRELIKNTVQAQERDGTDDKSSSL

IPQFGNRIGFNFGVGDTLTFSCSSGYRLEGALEIICLGGGRRVWSAPLPRCVAECGASATNNEGILLSPNYI

IDPVSAKIEGNLIFDPNTYIPKETMLKTTLQLYGLKPMDIFEIGMEGKSFEPTVEALFGQKGFFPDSASKALYW
TQHMFEEDPSVLYVSLHRYDHGAFFPTSEDADADRVGEGPGRGFTLNVPWNSPRMGDAEYLAAFHRLVL
LDTSKDDLKRYKQKSICKSVQKFPMASTVEITSLSAINDFYSYIILAAGSNRTLEVFDLNAGCSAAVIPDAHSRS
VQDSLNRLDDPAHISCTSSADYLLCRTLAASDCIERLQGAHGKYLSNRSDVSSLLPCVALFAHLISDTILQGS

FLRIISDTASCCYSIMKTKNTGIALGTKGASGLFPSEAAEWLCYRAFIIKLTNHKVVYKCLLRFLKICKYRVTLKV
SRDHIILEHLQWTMGYEVQITAANRLGYSEPTLYEFSMPPKPNIITDTLFNGLGLGAVIGLGVAALLLILVVTDVS

YLTLVTSTKAHAKIISIDTTEAQSVPGFVCFVSATDIPGSNVTGVAHDEPVFADNTVTCIGHIIGGVVADTHEH

LIILARIDESSHYDLSYTDVHFKPGQIRRHFIEVPQGATWAEVTVCSCSTDVTAKFVLHAVQLVKQKAYRSHEF
NIILTNRLQPSAIVDEATCAACDFNKPGANCQRRMTWQWRGEFMPASRSEYHRIQQQLESEKLPPLYPDG

HESRRTQWKRPTAQDNLADVDNGNIHLEAQHAFTHRRQITEEAENLDNRDSPESWEVITEDEATMYSNQ

HTNKCYMRDLCVVDADWLYDAAPDYFRRKLRAAKN
APDGHHVWEMEAKTDKEMCKPNSLVVEIPPFRNQRITSPVQVNFYVCNGKRKRSQYQLFTYLPANVPIIKT

GTANNLAGATNGLFRPLGAQPQQPQQQTNSSLQSNSFYGSNSLTNNSQSSSLFSHGPGQPGSTSLGFG

CAVLGRGHFGKVLLAEYKNTNEMFAIKALKKGDIVARDEVDSLMCEKRIFETVNSVRHPFLVNLFACFQTKD

LPVCPTFTIWNKSLMILSGLPAPDFIDYPEQQECNCRPQESPYVSGMKACH
NEKETPFELIEALLKYIETLNVPGAVLVFLPGWNLIYTMQKHLEMNPHFGSHRYRILPLHSQIPRDEQRRVFD

NPFYSAAPTVPETAHPMRAIVESLQ

WEAKDCKTFTALSICKNYTGPPRQPEVLPKPTDPCPAGWQNGSGLACYKFFHSERLLRTRTWEEAERFCE

AKQMAAEVAVKILRGEAGGQFFPEQPSGEVLSEPSLDPPVAQPLNPDESKAAKAKGVGELIKYLNANPISGLI
SCVHKIESTLMKTKGVLYSSVALATNKAHIKYDPEIVGPRDIIQTVEDLGFETSLVKKDRSASHLDHKQEIRQW

EWQLNQKKRSESFRFQQENLKRLRDSITRRQIEKQKTGKRSDLEITVPIRPSHNAPGKLECGIYEFGPRKN

VEALTSDGKTAEDLARTEQHEHVASLLARLKKDTHRGLFIQQLRPTQNPQPRIKLKLFGHSGSGKTTLVESLK

ICLSFVNAKAYNGNTALHVAASLQYRLTQLDAVRLLMRKGADPSARNLENEQPVHLVPDGPVGEQIRRILK

EGTAGDALIIGSVEEGSEYTAHDNMKFSTFDRDSDQWEENCSEVYGGGWWYNNCQAANLNGIYYRG

SISCASCGQAWGMEMIYRSVTLPILSIKNFVVATPDGNKQPKQWSRVTFTIEEFDYVEYCNRNLGM

NTFLCAALPSSLVLLQWYEPLQKFMLLKHFPAALPMPLSLFELLVVETEEYPQVCVRVVDRDRPGQELEFNVL
GPHGKCRSQAGGKFTCECNKGFTGTYCHENINDCESNPCKNGGTCIDGINSYKCICSDGWEGIYCET
ALRSFSPLQSAQQTHSGFQTSGSHVEGAPPRAHSTMTASGTDARTASSGSSAWEGQLQSLILSEYASTEMS
ISRFWQGRNLVTGELGYFPSDAVKPCPCVPKPVDYSCQPWYAGAMERLQAESELINRVNRTYLVRHRTKE

LNTHYSQRIEGATISSGVKYFGQSIHGQIDLSGDGLPDLSVGALGEVIMLRSQPILTVVPKMSFSPEEIPVKQ
FYNGCMEVKVNDIQLDLDEAFSKHNDIRSHSCPLIANEMNEFLN

FEDAKVFCDEKSSHLVIINNKEEQQWIKRQIAGKGNFWIGLTDSAQENEWRWLDGTLPEYTNWKSGQPD
SQLKKLRVLTITAIAQLCILGITWIFGMFQFNQRSLVASYIFTILNSLQGLFIFLLHCLLKKQVRDEYRRWLSC
CGVAGQYGVYTKVINYISWIENTILKNS
QAVARYVVQDPTGVNILSESGGDSMVGEGLEPEGSPKDGHHSYWAQFVTVTVLLSVTLAIAMAMAGFSYH

ETSKME

AAGPPPGHTGPAASEPDGKTA

SVPPRPGRGPLRSSRHPRTFYLGEQGEG

SELNGMHREMIIKEDIFQPRGIAVHPLAKRLFWTDLGVNPRIESASLQGIDRLVIASIDLVWPSGITVDCLAD

LTFLTNANAGDSKVADESCKAIVRPRRSSFEEEIRNKASKYLHPEIRKCCMSGVKESPLGQTCDQRVEHIK

EATHNSNHNNSNCSSPSRMSDSVSLNTNSSNDTSVCSPDKEIHSAAISKIRQEDENFNSLLQNGGEFN
SLSSSMQEGLISGSQFWNASKTLKTLIEEGYFEDRQNVDNRFILPPVIKSMTSESDSLGLTPGENCELALS

GHLDRKGAGEDESRDASNGAFIMPPADAMDASKCQGPSNGAGVTNGAVTGNSGFLSPHLWKKNSAMT
SSTGHPLYHRMEGTLSARSRNNSTERLLESPSVSVDDYHESDGSRNRTRAVENQYSFY

QKYWEVCSFNEVLASKYANENPGDFVNYNKRFLARVFPSPMRIDSSNMNPQDFWKCGCQIVAMNFQTP

CGRSGENFDKFFTRAVPALTPPDQLVMAGLDQSEFQGFTYINPDFVHLSLRPPTSPISMV

TQIIPITTGRIIRKPIGPILASATSNPEIFNSVPDGVPEKAMFLKSNDNFLLIFLFLFFFVLFLSLLSYNCYKGYLPI
ACLQDGVECSGHGSCICNKCQCQPGYFDRLCSRCPSCRTPCEEHRDCADCQAFRMGPLSQNCSIACI
ETVNSAVLQYRLKSLQANTQYTAHVMAATKAGGTYGNRITFSTLKFSKEDFIFVTISFGFCLLLLLILGITCTLKK
PVTKYTWYKDDTQCQETQESVLTFPATKERSGSYRCEAQNAIGYRQSLPVSVDVQYAPQHVAVVRQPSGSI

LVLSISDSESILTSNILGDETHLTSWFTTFQTTPVLSVSSSLLSTVTASDEVTGATANHTLLLTSPSKATASTAHFTT
QQGQQPYYNQQPQPQHLPPQAQYLSQAQQRYQTQQDMSQEGYGTRSQPPIASGKPNHEDLNLIQQEF

KKARIVAFDWLGQYLYWAGKANTIYRKSLLVGHTDVVVHVAFLVKDLVVDAINGYLYWATTYSVESTRLNGE
DLRFKIIVTAYKELTNTNQETFKRVLIREGQYDSIIVFYNLVFMQKLKTNILQYASNRPPTLSPIPHIPRSPYKFSN
REHVEDDDEYKIPSSHPVSLSSQPPHCHNIKPHIRGCENGQCAGIMNGTHSSASEMKKSKTPELGDAFDA
PIFSRMPVEGPLSPSRGDFPKGMPPQMGPGRELEFGMGPGSMKGDISLNVSMGSNPSMIPQKMREAGV
RNKREIFHRTWTDIFKQHNFSFYHNFHLDRNSRMDDFGGMQLRQDSDLTSGSNGITLYSEEEEIKYINR
HKMIAMGSAAALRNLLTNRPLKYKDTAVISPGSCMPSLYVRKQKALEAELDAKHLAEAFDTKKPLRHMDSLAK
HMDIKRSHPNLKTIIFSSQEQEREVLQSNTIATNNCVLKPLQRQNCEIPQQLRKSLVNPSLAYTNDKTVVFPD

QETLDEVLQQKREVEGELEIVWESTSKENRRMKEFLHKSLKNNNNMWSTVRVYESYLDEISQKDFISGYDF

TCQGPGRVAKIGDFGMARDIYRYRIMTQCWQHQPEDRPNFAIILERIEYCTQDPDVITTALPVEYGPSTEEI
LPIRWMPPESIMYRKFTTESDVWSFGVILWEIFTYGKQPWFQLSNTEVIECITQGRVLERPRVCPKEVYDIM
SPPAGLTSDPSPTVRRVPAVQRDSPVIVRNPDVPLPSKFTEKPREGQAGSPRLGKVGTKEQLQVPVPINMGR
GLLPHSPLPDSTRADSPFQSLVSSSQGSPQLCQRLLPKACKTSVATQCDPEEIIVLSDSD
YNRLKRAKMDKSMFVKIKTLGIGAFGEVCLACKVDTHALYAMKTLRKKDVLNRNQVAHVKAERDILAEADN
NGYECTCKPGYTGRMCNINIDECAGNPCHNGGTCKDGINGFTCICPEGYHDPMCVLEVNECNSNPCI
ESSSSDSEDSETEMAPKLKKKGHSGREQKKHHHHHHQQVQQQQMQPLPPQPVPQQAPPPMKSSPPPFV
MNHNKEVFSVVSCMKLYLSHLCQRCIDEQVDRWVCILPVLHLFTASFHLDEGMRVESPLESEEIWAGLEG
RRIDISPSTFRKHGFTLEEMKSTREPANKCLEIHSEGKYKDDPVDLRLDIERRKKHKERDLKREKSRESVDSF
PSNGFQSVYKSKEQDLNDSHIQSNEFNSSFKEREAPVPSEQDEVISSPVTNVDHSEMLVKKLDNLGTPCNF

STGILVVPIAPDPVEDLQCQPDSKGFSLSWTFPTGDVEACELAAERLSGGSPQAEARVIAPRSKTNLWGLEP
PAGSPQAPMEKEPAAASPPRTETPPAAPQLCPLQPREAMGCGEEEGMDTAGGEAGSLASPAGDGTQPTPAV
PQNVCRESSHPGITNQLNLHLKNLKVPRLISIRSKHKESAAGAEPPLLEEECGVNCCSSDTKGFTGPPAKSG

GGSTPHISGPGPGRPSSKEGRRSDTCEYCGKVFKNCSNLTVHRRSHTGERPYKCELCNYACAQSSKLTRH
RKRKRRVFSPIRSEPRSPSHSMRTRSGRLSTSDLTTLTPQSSVSSSLTSMSVSSLATSALNSTFTFSSHPLTQSG
SVEESDLRRSPRVKTTPLRRQTDMNPSTSNSEEDSNDAYCEKHKQKSSTQSRRKKDKRNSSDSTTDHPKP

ELVEPLTPSGALPNQAQMRILKETELKKVKVLGSGAFGTVYKGIWIPDGESVKIPVAIKVLRENTSPKANKEILDE
EKMTKETRRRGEPNERARKHASESKDSSSGLQRNVLVHVPECKDPDEQQDALNHGIPLSDTDLKGNDLQA
DRDLQ

NTDTTANSSCQINRVVSHPTLPISITAHEDRHIKFYDNNTGKLIHAMVAHLDAVTSLAVDPNGLYLMSGSH
EERADLTSNCRENNSPSHQTNTANLNNVVSTLTTGVCCAHPEKQGNVFLRQKSNCTCETFQGSDNSVL
HTPVRPPSTSSTGSRGSCGSSGRTAEKIGINFKSDHVKVKQEPGTEEEICNFSGAVKQEKTEDGRRSACML
TSDQAEKSSESRRRYSFDFCTLDNPEAIPSDTSTKNRHGHRQKHLIVDNFLAHESIKKPKHKRKRKSLQNR
PPGADSTHRVPVVMLEPIRIKQESGTSHENYDFPIVIVKQETDEESRPRNTNFSGSILTSLLLDSRHNTTSD
NTRIQEADNDPTAPPYDSIQIYGYEGRGSVAGSLSSLESATTDSDLDYDYLQNWGPRFKKLADLYGSKDT

NGNMYKSILVTSQDKTPVVIRKAMAKHNLDGDRPEDYELVQIISEERELKIPDNANVFYAMNSAANYDFVL
RELRDYFKKQIEELYSSVSV
MPFAKWTKEQVCNWLQGQGLGSYMNHGKHWILSGQTLLQASQQDLEKELGIKHPLHRKKLQLALQAL
ETRQKETSKYSNVVAEVKSMKDLSLKAAPEDVPKLSEVPPQPSMGESLQQQDDSWPQVKMDVAPGLASC
VVCYTPAQGTSSEYPVKMEIDLAIREAKGHFTYKEDPIVFKIHPAKSFLSGGSTITVYGIHLNSVYLPQMVITVP
QMRKMLCQKESNYIRLKRAKMDKSMFVKIKTLGIGAFGEVCLARKVDTNALYATKTLRKKDVLLRNQVAHV
NNLKEGSPASAPTRPGEQDGSQAGQPTKLESRVGDGHPTVLLLSRANGQPLGLFARPLGADRRGEGELP
PDLISTAMEADCRRNLENEEHKVDHWQCCKADPHNPCPQCRQANGGQKQLPSIEPGMNHPESTTCG
PSPISPSPSPVSSVGSQGSTGAPSPSSTISIPQRIHQMAANHVSITNSILHSYDYWEMADNLAKENREFFNEL
GCQHNYYEDAKAYGFKNKLIIVAAETAGNGLYNFIVPLRAYYRPKKELNPIVLLLDNPPDMHFLDAICWFP

LAEEEPSWEPTEIAQVLGVEPLQYVFVMKFTLDDGTGALDAYLFDYEKFFQIPASEVLTSSLLQEKMEMIMN

CHLPNAVCCEDHQHCCPWGYTCNLATQSCEKHQARRVLPVGLSLSASLLPASPEPSNDVTCDAQHYCC
EQLLSQFQSRLTFGVQRELCDLVRVSLLNAQHARALYNAGFITVADLAKGNTAEVETALKSAVPFKSIRRAV
SSNQFRCSNGRCIPSHWACDGDNDCGDFSDETKTNCTKEETPSPGGCNEREFQCNPDGNCVPVLW

ACWVITKAEQP

KAVLEAVGSCLGELGPIDFSTITLQHSKNAIYSKAVDLFEDRELQWVFIMLTLINNTLIDDCIGVRSAAAACL

MAQKDITLAPEIDKEINRLEATWEDTKKLIHEKQEQCCVLVDLMKEYQCLKSAVTKVIENGDNVTMTKFILKD
TLASVLDPYTQAYLTQAISSLRAPLWIGLANEEGGRSYSWLTEESFSYVNWQDGEPRQVTGCSYVDVDGT

SASLEETRQQMEAFTDLWKSSQLTPVTSAPWYNIVGESGQLDAESYDVVTEFDQEDGTKPSENVWSMPK
EEETGFGEDGVVQDAVGEPAYINEAAGIDRPSNKL
IQLTYKNGTTYNNEEKTERSTLITFLCDYEAGIGQPEYQVEDNYTYNFKWYTKYACPEMLLECVVTDPNTM

LNQVSAVPTASRIHQLECRTCKFSEVPALKLSSFKELRVLRITKSKYLTGFRQKFMGLPKLEVLDLSENRLSFIK
ELPETFCSTPRPVAQQACKNKDCPSEWLLSEWTQCSVSCGEGTQTRNAICRKMLKTGGSVIINSSMCPPL

KFERGTLMEQQTCSYTCRDEIELVQELGDTGKDAVNCTYKDENDCVVRFQYYEDSSGKSILYVIEEPDCPK
EEMEVSGTSGNEEEMAEEAEAAVNHSSDTESIPSPRPEAKESGENGPKPTPGPGRDSSPEPALKAEDADAAT

LPGLPTSAELFILPCQEATGENKRPAEDLDQISEILIHKLNQNLVQFELKPGVRVLVHAAHLSAVPLVDLTPSH
KWIQVNQRARKRSKRKQRQDEVAFQKLLPDQLVLLLESLLEEKTLCLRTLQRLEKTYHLDEQDAEVRHRWY
AVEQENISLAGCLLLEGDANVDSTTYDGTTPLHIAAGRGSTKLTALLKAAGADPHIENFEPLFDLEDVRDKD
ATNGTTGAAGTGTRPVPAGVVACTEGTTYVCSVCPAKFDQIEHFNDHMRMHVSDG

DPRLAAALGVGETDARGMMPSKDVPPSVWATVLAVVWLHSRAAGQRDEWELLEAKAVGWVRGRAGSRL

NITASVPLPNERNDSVESENQQRLLKTLETITNRLKRTLNKEPMYSFQFASELVVADSNSLETEKAFLFCRPG
EQKNLNSFFAVMFGLSNTAVSRLTKTWERLPHKTRKLYSALERMLDPSWNHRVYRLAIAKLTPPIIPFMPLLL

QAVGIGSLFQQSEDCGANASLCWKRVKGCKSNGQNGVGQKTIILLSTDIASKLLGQSMDETGLPQLTSYD

TDFLQATQMGAAPPPVPEQAPPPAPPPPPQPEAPPAPALPLMSERVSGGERGRGRPPGGPEALTPEFFISV

VALLLGLMGRAVAERARLVLYGRGHWEEVQALAGSLLENVQSLHQQVQATTEPSSSSSTVSDVLWDLVARG

RRKHR

LEETMSSLRGTQVSHSTLETTFDTNVTTEISGRSILSLTGRPTPLSWRLGQSSPRLQAGDAPSMGNGYPPRAN

SLMTSSCSSDDIDQEEIQLALQAAKIATREKIRSRFHGSNDLIHRLFVCISGVADQLQTNYASDLRSILKTLFE

ELVLFSNSDNERSIPSLVDGLKPGQRKVLFTCFKRNDKREVKVAQLAGSVAEMSSYHHGEASLMMTIINLAQ
LDSDEDLDSAIKDLLRSKRKLKKKSKDQKIQCKKKVRFGDTETQLLDELSSLQRKDWKCKSPILLKSCLSKSR

PLNYENNHECIYSIQVQAGKGINVSARTFHLAQGDVLKIYDGKDNTAHLLGAFTGASLRGLTLSSTSNHLWL

VDGRVPEHVSKVLFDYFGYSKDDRQEQDVMKGIMLNFEKLIKEMGNKEVPEARAYLRILGEELGYMKLND
PIAYQFAPELVLVSAGFDAARGDPLGGCLVSPEGYAHMTHLLLGLAGGRVVLVLEGGYNLVSISESMTMCTP
SVHQICQNKGSAFSTQPLEAYNLFMTTATGDGIKLWDLRTLRCERRFEGHINRCHPCGIAVTPCGQFIACG
ATSHMVPMEPADLLSEMCKQCGSEALHYLGALKDEASLGGADCTAVRNCLSRISALGEELRPKGLDVEQE

KLFKYR
SCFFVRQCGLLMCITRRICGKKSGSSGKSKELEEGKAAYLKDGSKEPIVEMRTEDERITNHEDGSPVNEPNI

SRRAAKAVKIEYEALTPIVTIQEAIEKQSFYQSPKKIEKGNVQKGFEKSDCVVEGEMYIGGQEHFYLETNCAIA

FYKFSSLPEKGSVTEAFPVLGGKTIEFCIARWWASLSDVSIDYTISFHGIVCATPQLNIHASEGIVRFDVQSTLK
GPPRAFHELSREEQAKFEKKRLADYCRKAYKKLHVTKVEERITTICQRENSFYVDTVRAFRDRRYEFKGLHKV

QNLQTLLPQSSSDMQSHLAEELNIRLTTSGSPATGQSTSSNNHSSRRHTELQTYRAEEQPALPVLLSTSSGLPP

TEPSDDYEPAQTCGPMTQGLNPLSKPYYSQQILMPPDPGSCLVAGFTSCQQRNTVMSSSPNSSPKLHDLS

SSSSLGAAIGSALGGFGSSVGSSASSSATRRESLSTSSDLYKRSSSSLAPIGQPFYNSLGFSSSPSPIGMPLPSG

HVCFVMEYAAGGDLMMHIHTDVFSEPRAVFYAACVVLGLQYLHEHKIVYRDLKLDNLLLDTEGFVKIADFG

DPVPSGVTKVILSTNIAETSITINDVVYVIDSCKQKVKLFTAHNNMTNYATVWASKTNLEQRKGRAGRVRAGFC

ALGGHLPSFSDIAEMKELHYILRNIISNDRWVWVGLNKRNPDSLGTWQWSDNNPETTVVMSHDFQRDV

LEYARSNGFAAEFKMVDQTGPPHDPKFVYQAKVGGRWFPAVTAHNKKQGKQEAADAALRVLIGETEKAERT
VKRSFLVSLSFCIPVMALMIYMMVMDSHLADVHQHRNMSKEEMEAYHSSMFLERQILPGLSVMNLLSFLLC

VFPPIKEMKRGEWKTESTSSTASSASNRSSTRSTLSVSSGMEGDSEDNEILEATRSRSPIPQQVERFPLTLPER

CGLIRSFFRRRRPRLSSTNSTRFPPSPLSTKPSVSVSITNLYPGCENVSVRSRSMMFEPGLTKGVLEVFVSPSH

GKAVQQRASPY

GQYDPRNNVPYEIENGVVWIPFRASDYSLKVVRMKIRPIETL

LSLSTSSSLSTEPSADVSSSVACHMTQMDRDTVLVCFEHYIRTVNLHGAPKRALAPELTFNFPIETIVCLQDSV
NINDCSKNPCHNGGTCRDLVNDFFCECKNGWKGKTCHSRDSQCDEATCNNGGTCYDEGDTFKCMC
LHALYMHELHKQHTQTEPERHVWHRRESDESGESAPEEPDAQKGAPAVIPRSASYPFSSPRQAVEESATLQ
SGEYAISIKYNNEVKHIKILTRDGFFHIAENKKFKSLMELVEYYKHHSLKEGFRSLDTTLQVPYKESENSDGQ

VECSGSVSSRQGVQSNLTICFSVSRATMRYQEPLSANLTYWLEIDANRMRSRGVFGNGQRNMSGTMEISE

NWSTGQGPGEDCAGLISAGLWNDFHCEDVNSFICEKDMDKEDEKEKGKRAKSYID
GLKGQAKSSEFSSSITITRGLQRSQESG

DRLKAKSKVQGCNTPEATKVSNQEKVPLNGGRSPQLQGPIQEGAKDPRDCCVQMEGLHQEIDMDNNRL

KLYWCDAKQSVIETANLDGSKRRILTQNDVGHPFDVAVFEDHVWFSDWAKPSLTRVDKKTGQNRVRLRG

SIPRCIKAFKDCCEFATKLRENETHKVLILARMQLKALLGLDKPEIRSYFPESWLWEVHQVPPGSKALSVTLPD

ITAEEEISAQQNVTNLPEYELRIEERLALIEKGVDLNPTTEESHKLDQINMNINKLTSEDTIQPSVMERLKTQEIV
ALGGCIFYLKKCLIDQELLSLANFEEYVPVDADVAKAKGSSSFFAKTNQRMVLDGVTLANLEILQNGTTGTTEG
SSRPTTGEEECSSNASKRFLRSCSASCVPHGAKDTEWRSMTLPRDLQSTVRQFDSSTFGGHKSEKPALPRI

GLMMDLNIGWFRQNGNCGYVLRPAIMREEVSFFSANTKDTVPGVSPQLLHVKIISGQNFPKPKGSGAKGD

DQCLKFRSAMLLGKKRPKSDFSDCEQSVKETLVEQGSIQNQTGEQPKPAHDTGYETEPDCGNGQVQHD
NHTMATLVPEATVNEQWCKEKTDDGRILIFLIEGTEGGKVPLKVKDKAAVADRTSMFVLGVVLGIVFIGLLIIG
HTLKKVCWPDIPNPAESIAVEWPTDMLKNNSPLKKVQSEDGTVDPDGICVLESCFPAEDHTALLVESHENY
KEGETVTLECSVGSANPSALHFAWYKDEARQGENSKRLQLVAVVSADSGQYRCEAKNDVGTTHAEPIPLDV

LVLSISDSESILTSNILGDETHLTSWFTTFQTTPVLSVSSSLLSTVTASDEVTGATANHTLLLTSPSKATASTAHFTT
QQGQQPYYNQQPQPQHLPPQAQYLSQAQQRYQTQQDMSQEGYGTRSQPPIASGKPNHEDLNLIQQEF

KKARIVAFDWLGQYLYWAGKANTIYRKSLLVGHTDVVVHVAFLVKDLVVDAINGYLYWATTYSVESTRLNGE
DLRFKIIVTAYKELTNTNQETFKRVLIREGQYDSIIVFYNLVFMQKLKTNILQYASNRPPTLSPIPHIPRSPYKFSN
REHVEDDDEYKIPSSHPVSLSSQPPHCHNIKPHIRGCENGQCAGIMNGTHSSASEMKKSKTPELGDAFDA
PIFSRMPVEGPLSPSRGDFPKGMPPQMGPGRELEFGMGPGSMKGDISLNVSMGSNPSMIPQKMREAGV
RNKREIFHRTWTDIFKQHNFSFYHNFHLDRNSRMDDFGGMQLRQDSDLTSGSNGITLYSEEEEIKYINR

HKMIAMGSAAALRNLLTNRPLKYKDTAVISPGSCMPSLYVRKQKALEAELDAKHLAEAFDTKKPLRHMDSLAK
HMDIKRSHPNLKTIIFSSQEQEREVLQSNTIATNNCVLKPLQRQNCEIPQQLRKSLVNPSLAYTNDKTVVFPD

QETLDEVLQQKREVEGELEIVWESTSKENRRMKEFLHKSLKNNNNMWSTVRVYESYLDEISQKDFISGYDF

TCQGPGRVAKIGDFGMARDIYRYRIMTQCWQHQPEDRPNFAIILERIEYCTQDPDVITTALPVEYGPSTEEI
LPIRWMPPESIMYRKFTTESDVWSFGVILWEIFTYGKQPWFQLSNTEVIECITQGRVLERPRVCPKEVYDIM
SPPAGLTSDPSPTVRRVPAVQRDSPVIVRNPDVPLPSKFTEKPREGQAGSPRLGKVGTKEQLQVPVPINMGR
GLLPHSPLPDSTRADSPFQSLVSSSQGSPQLCQRLLPKACKTSVATQCDPEEIIVLSDSD
YNRLKRAKMDKSMFVKIKTLGIGAFGEVCLACKVDTHALYAMKTLRKKDVLNRNQVAHVKAERDILAEADN
NGYECTCKPGYTGRMCNINIDECAGNPCHNGGTCKDGINGFTCICPEGYHDPMCVLEVNECNSNPCI
ESSSSDSEDSETEMAPKLKKKGHSGREQKKHHHHHHQQVQQQQMQPLPPQPVPQQAPPPMKSSPPPFV
MNHNKEVFSVVSCMKLYLSHLCQRCIDEQVDRWVCILPVLHLFTASFHLDEGMRVESPLESEEIWAGLEG
RRIDISPSTFRKHGFTLEEMKSTREPANKCLEIHSEGKYKDDPVDLRLDIERRKKHKERDLKREKSRESVDSF
PSNGFQSVYKSKEQDLNDSHIQSNEFNSSFKEREAPVPSEQDEVISSPVTNVDHSEMLVKKLDNLGTPCNF

STGILVVPIAPDPVEDLQCQPDSKGFSLSWTFPTGDVEACELAAERLSGGSPQAEARVIAPRSKTNLWGLEP
PAGSPQAPMEKEPAAASPPRTETPPAAPQLCPLQPREAMGCGEEEGMDTAGGEAGSLASPAGDGTQPTPAV
PQNVCRESSHPGITNQLNLHLKNLKVPRLISIRSKHKESAAGAEPPLLEEECGVNCCSSDTKGFTGPPAKSG

GGSTPHISGPGPGRPSSKEGRRSDTCEYCGKVFKNCSNLTVHRRSHTGERPYKCELCNYACAQSSKLTRH
RKRKRRVFSPIRSEPRSPSHSMRTRSGRLSTSDLTTLTPQSSVSSSLTSMSVSSLATSALNSTFTFSSHPLTQSG
SVEESDLRRSPRVKTTPLRRQTDMNPSTSNSEEDSNDAYCEKHKQKSSTQSRRKKDKRNSSDSTTDHPKP

ELVEPLTPSGALPNQAQMRILKETELKKVKVLGSGAFGTVYKGIWIPDGESVKIPVAIKVLRENTSPKANKEILDE
EKMTKETRRRGEPNERARKHASESKDSSSGLQRNVLVHVPECKDPDEQQDALNHGIPLSDTDLKGNDLQA
DRDLQ
SNTDTTANSSCQINRVVSHPTLPISITAHEDRHIKFYDNNTGKLIHAMVAHLDAVTSLAVDPNGLYLMSGSHD
EERADLTSNCRENNSPSHQTNTANLNNVVSTLTTGVCCAHPEKQGNVFLRQKSNCTCETFQGSDNSVLQ
SHTPVRPPSTSSTGSRGSCGSSGRTAEKIGINFKSDHVKVKQEPGTEEEICNFSGAVKQEKTEDGRRSACML
TSDQAEKSSESRRRYSFDFCTLDNPEAIPSDTSTKNRHGHRQKHLIVDNFLAHESIKKPKHKRKRKSLQNRI
RPPGADSTHRVPVVMLEPIRIKQESGTSHENYDFPIVIVKQETDEESRPRNTNFSGSILTSLLLDSRHNTTSDK
NTRIQEADNDPTAPPYDSIQIYGYEGRGSVAGSLSSLESATTDSDLDYDYLQNWGPRFKKLADLYGSKDTF

DNGNMYKSILVTSQDKTPVVIRKAMAKHNLDGDRPEDYELVQIISEERELKIPDNANVFYAMNSAANYDFVL
RELRDYFKKQIEELYSSVSV
DMPFAKWTKEQVCNWLQGQGLGSYMNHGKHWILSGQTLLQASQQDLEKELGIKHPLHRKKLQLALQALG
LETRQKETSKYSNVVAEVKSMKDLSLKAAPEDVPKLSEVPPQPSMGESLQQQDDSWPQVKMDVAPGLASCA
VVCYTPAQGTSSEYPVKMEIDLAIREAKGHFTYKEDPIVFKIHPAKSFLSGGSTITVYGIHLNSVYLPQMVITVP
QMRKMLCQKESNYIRLKRAKMDKSMFVKIKTLGIGAFGEVCLARKVDTNALYATKTLRKKDVLLRNQVAHVI
GNNLKEGSPASAPTRPGEQDGSQAGQPTKLESRVGDGHPTVLLLSRANGQPLGLFARPLGADRRGEGELPI
SPDLISTAMEADCRRNLENEEHKVDHWQCCKADPHNPCPQCRQANGGQKQLPSIEPGMNHPESTTCGI
PSPISPSPSPVSSVGSQGSTGAPSPSSTISIPQRIHQMAANHVSITNSILHSYDYWEMADNLAKENREFFNEL
KGCQHNYYEDAKAYGFKNKLIIVAAETAGNGLYNFIVPLRAYYRPKKELNPIVLLLDNPPDMHFLDAICWFPV

SLAEEEPSWEPTEIAQVLGVEPLQYVFVMKFTLDDGTGALDAYLFDYEKFFQIPASEVLTSSLLQEKMEMIMNI

CHLPNAVCCEDHQHCCPWGYTCNLATQSCEKHQARRVLPVGLSLSASLLPASPEPSNDVTCDAQHYCQ
MEQLLSQFQSRLTFGVQRELCDLVRVSLLNAQHARALYNAGFITVADLAKGNTAEVETALKSAVPFKSIRRAVD
SSNQFRCSNGRCIPSHWACDGDNDCGDFSDETKTNCTKEETPSPGGCNEREFQCNPDGNCVPVLW

VACWVITKAEQP

EKAVLEAVGSCLGELGPIDFSTITLQHSKNAIYSKAVDLFEDRELQWVFIMLTLINNTLIDDCIGVRSAAAACLK

MAQKDITLAPEIDKEINRLEATWEDTKKLIHEKQEQCCVLVDLMKEYQCLKSAVTKVIENGDNVTMTKFILKD
TLASVLDPYTQAYLTQAISSLRAPLWIGLANEEGGRSYSWLTEESFSYVNWQDGEPRQVTGCSYVDVDGTW

SASLEETRQQMEAFTDLWKSSQLTPVTSAPWYNIVGESGQLDAESYDVWTEFDQEDGTKPSENVWSMPKI
REEETGFGEDGVVQDAVGEPAYINEAAGIDRPSNKL
MIQLTYKNGTTYNNEEKTERSTLITFLCDYEAGIGQPEYQVEDNYTYNFKWYTKYACPEMLLECVVTDPNTM

YLNQVSAVPTASRIHQLECRTCKFSEVPALKLSSFKELRVLRITKSKYLTGFRQKFMGLPKLEVLDLSENRLSFIK
ELPETFCSTPRPVAQQACKNKDCPSEWLLSEWTQCSVSCGEGTQTRNAICRKMLKTGGSVIINSSMCPPLI

KFERGTLMEQQTCSYTCRDEIELVQELGDTGKDAVNCTYKDENDCVVRFQYYEDSSGKSILYVIEEPDCPK
EEMEVSGTSGNEEEMAEEAEAAVNHSSDTESIPSPRPEAKESGENGPKPTPGPGRDSSPEPALKAEDADAAT

TLPGLPTSAELFILPCQEATGENKRPAEDLDQISEILIHKLNQNLVQFELKPGVRVLVHAAHLSAVPLVDLTPSHS
KWIQVNQRARKRSKRKQRQDEVAFQKLLPDQLVLLLESLLEEKTLCLRTLQRLEKTYHLDEQDAEVRHRWY
AVEQENISLAGCLLLEGDANVDSTTYDGTTPLHIAAGRGSTKLTALLKAAGADPHIENFEPLFDLEDVRDKDS
ATNGTTGAAGTGTRPVPAGVVACTEGTTYVCSVCPAKFDQIEHFNDHMRMHVSDG

DPRLAAALGVGETDARGMMPSKDVPPSVWATVLAVVWLHSRAAGQRDEWELLEAKAVGWVRGRAGSRL

NITASVPLPNERNDSVESENQQRLLKTLETITNRLKRTLNKEPMYSFQFASELVVADSNSLETEKAFLFCRPG
EQKNLNSFFAVMFGLSNTAVSRLTKTWERLPHKTRKLYSALERMLDPSWNHRVYRLAIAKLTPPIIPFMPLLL

QAVGIGSLFQQSEDCGANASLCWKRVKGCKSNGQNGVGQKTIILLSTDIASKLLGQSMDETGLPQLTSYD

TDFLQATQMGAAPPPVPEQAPPPAPPPPPQPEAPPAPALPLMSERVSGGERGRGRPPGGPEALTPEFFISV

ALLLGLMGRAVAERARLVLYGRGHWEEVQALAGSLLENVQSLHQQVQATTEPSSSSSTVSDVLWDLVAR

RKHR

EETMSSLRGTQVSHSTLETTFDTNVTTEISGRSILSLTGRPTPLSWRLGQSSPRLQAGDAPSMGNGYPPRA

SLMTSSCSSDDIDQEEIQLALQAAKIATREKIRSRFHGSNDLIHRLFVCISGVADQLQTNYASDLRSILKTLFE

LVLFSNSDNERSIPSLVDGLKPGQRKVLFTCFKRNDKREVKVAQLAGSVAEMSSYHHGEASLMMTIINLAQ
DSDEDLDSAIKDLLRSKRKLKKKSKDQKIQCKKKVRFGDTETQLLDELSSLQRKDWKCKSPILLKSCLSKSR

LNYENNHECIYSIQVQAGKGINVSARTFHLAQGDVLKIYDGKDNTAHLLGAFTGASLRGLTLSSTSNHLW

VDGRVPEHVSKVLFDYFGYSKDDRQEQDVMKGIMLNFEKLIKEMGNKEVPEARAYLRILGEELGYMKLND
PIAYQFAPELVLVSAGFDAARGDPLGGCLVSPEGYAHMTHLLLGLAGGRVVLVLEGGYNLVSISESMTMCT
VHQICQNKGSAFSTQPLEAYNLFMTTATGDGIKLWDLRTLRCERRFEGHINRCHPCGIAVTPCGQFIAC
ATSHMVPMEPADLLSEMCKQCGSEALHYLGALKDEASLGGADCTAVRNCLSRISALGEELRPKGLDVEQE

KLFKYR
CFFVRQCGLLMCITRRICGKKSGSSGKSKELEEGKAAYLKDGSKEPIVEMRTEDERITNHEDGSPVNEPN

SRRAAKAVKIEYEALTPIVTIQEAIEKQSFYQSPKKIEKGNVQKGFEKSDCVVEGEMYIGGQEHFYLETNCAIA

FYKFSSLPEKGSVTEAFPVLGGKTIEFCIARWWASLSDVSIDYTISFHGIVCATPQLNIHASEGIVRFDVQSTLK
PPRAFHELSREEQAKFEKKRLADYCRKAYKKLHVTKVEERITTICQRENSFYVDTVRAFRDRRYEFKGLHKV

NLQTLLPQSSSDMQSHLAEELNIRLTTSGSPATGQSTSSNNHSSRRHTELQTYRAEEQPALPVLLSTSSGLP

EPSDDYEPAQTCGPMTQGLNPLSKPYYSQQILMPPDPGSCLVAGFTSCQQRNTVMSSSPNSSPKLHDLS

SSSSLGAAIGSALGGFGSSVGSSASSSATRRESLSTSSDLYKRSSSSLAPIGQPFYNSLGFSSSPSPIGMPLPS

HVCFVMEYAAGGDLMMHIHTDVFSEPRAVFYAACVVLGLQYLHEHKIVYRDLKLDNLLLDTEGFVKIADF

PVPSGVTKVILSTNIAETSITINDVVYVIDSCKQKVKLFTAHNNMTNYATVWASKTNLEQRKGRAGRVRAGF

ALGGHLPSFSDIAEMKELHYILRNIISNDRWVWVGLNKRNPDSLGTWQWSDNNPETTVVMSHDFQRDV

LEYARSNGFAAEFKMVDQTGPPHDPKFVYQAKVGGRWFPAVTAHNKKQGKQEAADAALRVLIGETEKAER
VKRSFLVSLSFCIPVMALMIYMMVMDSHLADVHQHRNMSKEEMEAYHSSMFLERQILPGLSVMNLLSFLL

VFPPIKEMKRGEWKTESTSSTASSASNRSSTRSTLSVSSGMEGDSEDNEILEATRSRSPIPQQVERFPLTLPE

GLIRSFFRRRRPRLSSTNSTRFPPSPLSTKPSVSVSITNLYPGCENVSVRSRSMMFEPGLTKGVLEVFVSPS

GKAVQQRASPY

GQYDPRNNVPYEIENGVVWIPFRASDYSLKVVRMKIRPIETL

LSLSTSSSLSTEPSADVSSSVACHMTQMDRDTVLVCFEHYIRTVNLHGAPKRALAPELTFNFPIETIVCLQDSV
NINDCSKNPCHNGGTCRDLVNDFFCECKNGWKGKTCHSRDSQCDEATCNNGGTCYDEGDTFKCMC
LHALYMHELHKQHTQTEPERHVWHRRESDESGESAPEEPDAQKGAPAVIPRSASYPFSSPRQAVEESATLQ
SGEYAISIKYNNEVKHIKILTRDGFFHIAENKKFKSLMELVEYYKHHSLKEGFRSLDTTLQVPYKESENSDGQI

VECSGSVSSRQGVQSNLTICFSVSRATMRYQEPLSANLTYWLEIDANRMRSRGVFGNGQRNMSGTMEISE

NWSTGQGPGEDCAGLISAGLWNDFHCEDVNSFICEKDMDKEDEKEKGKRAKSYID
GLKGQAKSSEFSSSITITRGLQRSQESG

DRLKAKSKVQGCNTPEATKVSNQEKVPLNGGRSPQLQGPIQEGAKDPRDCCVQMEGLHQEIDMDNNRL

KLYWCDAKQSVIETANLDGSKRRILTQNDVGHPFDVAVFEDHVWFSDWAKPSLTRVDKKTGQNRVRLRG

SIPRCIKAFKDCCEFATKLRENETHKVLILARMQLKALLGLDKPEIRSYFPESWLWEVHQVPPGSKALSVTLPD

ITAEEEISAQQNVTNLPEYELRIEERLALIEKGVDLNPTTEESHKLDQINMNINKLTSEDTIQPSVMERLKTQEIV
ALGGCIFYLKKCLIDQELLSLANFEEYVPVDADVAKAKGSSSFFAKTNQRMVLDGVTLANLEILQNGTTGTTEG

SSRPTTGEEECSSNASKRFLRSCSASCVPHGAKDTEWRSMTLPRDLQSTVRQFDSSTFGGHKSEKPALPR

GLMMDLNIGWFRQNGNCGYVLRPAIMREEVSFFSANTKDTVPGVSPQLLHVKIISGQNFPKPKGSGAKG

QCLKFRSAMLLGKKRPKSDFSDCEQSVKETLVEQGSIQNQTGEQPKPAHDTGYETEPDCGNGQVQHD
NHTMATLVPEATVNEQWCKEKTDDGRILIFLIEGTEGGKVPLKVKDKAAVADRTSMFVLGVVLGIVFIGLLIIIG
HTLKKVCWPDIPNPAESIAVEWPTDMLKNNSPLKKVQSEDGTVDPDGICVLESCFPAEDHTALLVESHENY
KEGETVTLECSVGSANPSALHFAWYKDEARQGENSKRLQLVAVVSADSGQYRCEAKNDVGTTHAEPIPLD

NPHTLLLHTDVNSITASPTESVSVVTSFVPTQLPPSINTSTEFGTQTKTNVTEVTNRTSATTAITTTNSSIVGHRTT
RPSSLPVEVLASEDAAFGLKDLSGSIDDLPTGTEATLSSAVSASGSTSSQGEQSNPAQSPFSPHASPHLSGIPG

HYLMLQEQPRFSGKQVVALTLDLSDGFLFWLVQDSLCLNLYRVSLCGESCGDAVVTEFAEWSSSDVSHSA
SPLRVPAGNNIYISPLKSPYKFSEGMLSPTKMTPRSRILVSIGESFGTSEKFQKINQMVGSSDHQLKRSAEVS
STAPVSLPPARPSARDNPKHSPLLNRTPSDYDLLVPPLGEDSFDSSPPSLPPPPPPARHSLTEHSKPLGSGSR
GPEEMMKLRPGVSEMLSSQQKMVSLPFGEHPQQEYGMGPRPFLPMSQGPGGGLRNLREQIGPDQRTN
RVYEEENFNMLTFEDLLCFSYQVAKGMEFLESKLCIHRDLAARNVLVTCGKVVKICDFGLARDIMNDSNYV
KDYASDSGCFDDDEVPNVSAGAETGNASILSMFLNSSFLQGQALPRAIAQRRGPEPEKDESSKLAEPKKLPL
DDQSMDITRNHTAALDFAPILVAQEYESTHIQNSVISKPKQLQNKTLLFSGDSDIDITRSQTAAIECGSLRMNS

FSYCDVKPSSPTKNQSKQEPPC

EEKVPMRPEDPKAIPPLMVSPHQKKEGDKQALPSPPPLPSAVTAGKPLAKVAAPEPPVPVHVRSALEGGHINN
LGCWQREPQQRLNIKEIYKILHALGKATPIYLDILG
ALPCSQPSAVPAPEQAPPVFSVSSPFQPVAFHPSPAALLPVIVPNDYSGPPAPKKEIIMGRPGTVWTNVEPRS

EWVVKLYYSFQDKENLYFVMDYIPGGDMMSLLIRMEVFPEHLARFYIAELTLAIESVHKMGFIHRDIKPDNII
IHGKCNDGLNGYRCDCDPGWSGTNCDINNNECESNPCMNGGTCKDMTSGYICTCREGFSGPNCQT
VPAQVPIMEHPLPGNMFDAITHFTQPIMHLPQPEMPPHLPQQPEHSTPPHLNQHAVVPPPALHNALPQQPS
FPFIEFRKKQEKSCKNRALLQLMKGKKHLMEVDTTLVRSWISLLPLGDLAEFLTDLSVELLDCLLGIYHRLQN
RDSSHSRERSTEKPERSHKGSKKKKHRRVRERSRSTSSSSPSSRSYKGEDYPEESEEREECTTGFDKSRLGT
RTKDSDFCYISDDTSPSLYHSEVEIEVEPTDIKVTQETLMDIHVEPQTVTCDYKNTEDQNSTKSACEDELYDIS

DSSYQVSVSTVGRNGLRSRAVTLVCNTSAEVLPPPVRADVPRVEPGSRVLISPDTFSEENGRIEYYAVIVTSNE
VALGPFPGLTGEEEEEAAAGLEQAGAAGRCSPQLSEDGLEESRGGGDPESSQTPVSFNSEPKGSPMLLEPE
GKLDSQLNNVRERARDSAEIETAVVKHVLSELKELSFRSLGDEASGYGTPKTTAPLLFSSASGQSCLPIEPDYKI

IMKTHGQIGKEVYRCDICQMPFSVYSTLEKHMKKWHGEHLLTNDVKIEQAERS
GESAERSQRPRKQISTPAEPFSSTNPTPLFPWFTPSPQTERGRNKDRATEELSKDKDIDKSMEKDKSRERDRI
NKLSKSKKSDILDQSSDSDEMPAVLKEVAMMSHSSSDIDSNSETSNNIQKNALNDLTKQPVKYENGKRKRI

EAYVMAGVGSPYVSRLLGICLTSTVQLVTQLMPYGCLLDYVRENKDRIGSQDLLNWCVQIAKGMSYLEDVR
ADKTHRNPENCSDGMQTGKGLFYVKISMEDLNACSVVPDTDSQVCENQGSQSLLQPHSTNVDQAACPV

DCSIRLWNLESKTCIQEFTAHRKKFDESIHDVAFHPSKCYIASAGADALAKVFV
QTNSIVNTRERSCYTPVPEKEVHVEKDAVKIPNKETQLENSSLQPTFLSLIKNRNLTVEQVVAIEALTKLSEAPLEA
LSSPESSLTPPLSTNLHLESELEALGSLENHVKTEPTDLSESCKQSEHSLVNGKSPVRSLMHRSARVGGEGNN
DDLQFLADLEELINKFQVFRISHRSYTFYHENPYPSIFRINFDHYYPVPYIQYDPLLYLRRTSDMKSKKKRGR
KTVLRTDAPDSTGDRPEILQENTSSGKTAWMGTPHIGDIRKEDDPNEDWCAVCQNGGELLCCEKCPKVFH
FDDDS

LKKRGFSKGVKIKHGSSSTLPRMKQKGLKIAKGIF

GSEEENNHGKLDYNWVTRWLDDIGLPQYKTQFDEGRVDGRMLHYMSVDDLLSLKVVSVLHHLSIKRAIQV
AKMCPGNQGNRSQAEGDLKLKQVQKDQQPPPASLERGDEVPATPQESQQPHLMSLEKEADMPEKPLKTS
KLGKNFTVTCNHRSNTEIICCTTPSLKSFNLQLPFATKVFFIFDGVSKQYFDFDYVNNPVFKPFEKPVVISRGI
KAERDILAEADNEWVVRLYYSFQDKDNLYFVMDYIPGGDMMSLLIRMGVFPENLARFYIAELTCAVESVHK
PVSNIHSQRPDPPPLGLSGPTGPTEAQMASEAKSSCRAEPEWRFGIEMGSRQPHLQDGGNSEGTGVKTTA
LLYEKAQLPEKMEKEDFNSCKSVSLLDTPQHQAAPVLPCKPRPIQKSGDDSSEQEEGEVDSEVEFPRRQRP
LDALMGPVTLHSSMEHLVQYTQQGLHWVRNSAHAKDCVPFKCTA
VVYYMVGSIDNLDDLLRCGVTFAANMVVVDKESTMSAEEDYMADAKTIVNVQTLFRLFSSLSIITELTHPANM

ILCPPGTKLDDLPWLECFIKSYNVKDGMEQQVCYQIYDTMVAEDVI

QAHQTCCKAATGTWACCPYHWGFCCSDKRHCCPWGYRCSKSGLDCKRRWALRWDVGTHQPHRLL
DEDEEAAEERRNTHCIWMVGMKGLTESEAASLIVEEARILLQQDLAAMGVQWSPDSFLESDTSSMTSSESE
HCDGEKDCEDGSDEKGCNGTIRLCDERTKFSCKNTGRCISKAWLCDGDIDCDDQSDEDNCDSYMCG

KNILATKAGNEFWEVYKNKADPMLIYLHPFRMSRKKFLEVPSNGRESPSENLDDASVWIPQSVSHDTWIKNI

QEDIRRALSKHEAVKNELNDRQKELDAFTNKGKHLLAELKRIHNCDPMLVKTDMNSTVDKWLDVSERTED
VRTAGCDTKLQGAICKVNTGSPPSHKWSYSGSCPKSLEDSSWIPFRNHCYTFHMEVTLGQKDAMKRCQK

LDTDQTSVTSPEGWAMSKTSLSYSSEITAENESENPNNEVSLEAWNKGRAYPGEEYSTFENTKSSIDNVQNN

EQYDLSSLSKSEDRGENWYAMDNSEPNERRKYYINVCRPLVPVAGCDRYASSCQMKFEKDDDSFSETTS

KCCSFYLDGTPNLKHLNLSFNSVISMSGGLCLNELVSLDFQHSKLNSLGIVPVFLSLGKLIYLDISYTNTHVES
PLSSLVRSCALVACTRHNRLAQKQSPHIMALKNVYIQTKKQKKLHFIVGGYAYLLPKTSVVLRCPVRRFRKSM

GPDILMVLLLVAGAILLIGLVALFIWKLLITVHDRREFAKFEEERSRAKWDTANNPLYKGATSTFTNITYRGNS
TAANPTANTTSQPQEAAAEPANSEEKPQEELVVDVVKTEEPEEKPSEEPCKSEIKKEESEDCPERDKANEKKTE

SGSAMLMLLSVVFVGLAVFVIYKFKRKIPGINVYAQMQNEKDQEMVSPVSQRESMPNIPQSELMSPEQLVD
ELVVKHKYTRRYGDIEKFLKEDQAMGVYLYGELMVNEDAKQQELAYKCFAATQKQMDVSSAKVVAEMLF
SEDEGIVPGTTPLDMASSWEVFDILNGKPYESKLASDDLLLQGDMKKLNEDSKQQLYKLLEVPDPNKNWST

SEYLEAANTLLGCSVGPAVFGL

SVLRGRMCVNCPLGTYYSLEHRTCESCWTGSYQDEEGQLECKSCPSGSYTEYLHSRSISECKAQCKQGT
KDMTFIHEGNRTLAENLINFEKMRMMAKAVRIIHHCQSHTYAPISPLRSRATQLLEDPQTLRISTCSEQSLSG

CEVNAPIQGSRNLLQGEELLRALDQVN

VQGVLSSMMGSLNAQQGSTESIASFIQRLSGTSNLFEPGTEGAIGFFGDLLSLICQNFSMVDVVMLLHGQ

EPVDTLLVLSPRSEASLAGPALRLVRDRVAPHCLFVNLCLETRGCQGGPVDEVVLAGFNEQVLRFLAVRGT

NASRFINTESGRYMYSAPLRRQLASRGSNVCHMDISEKGGDEIDLETISMDATGYMSDGDVLGKNIRTDD

VMATKPETEDKEKQKKVNQGLRNVALEDCALCQETISSSELAAKARDGDFEDPPDWVPDEVCSYCTACKA

QNFVGSNNLNLLQPIGQFGTRLHGGKDSASPRYIFTMLSSLARLIFPATDDSILRFLYDDNQRVEPEWYMPII
KDTKENAVRSPQNNVNGRLSKERSESINDLPFGLQFKKECKPKPVCNKNNLEAAKNKRCSLTATSATDDSS

LEFNSDTEGTNEGFQLVYTSFELSHCEDPGVPQFGYKISDQGHFAGSTIIYGCNPGYTLHGSSLLKCMTGE

FKLLGNVILKSIKTLQAIPEKIVQAISKGTEGDLFVHYIFMDNEFELPTGAGLQLQVAFSGIATPGAKAGMKIQP
RTLLGDPLPELGRLRAPHPSALQSLARASAKHGKYWACLRLDVPIREDSRPRSGQAGPVREDSPPEDPTGG
GSEDKSVRNNIL
ELGDLVDKEMAATSAAVETAVARIEDMLSKSRAGDTGVKLEVNERILGSCTDLMQAIQVLVLASKDLQREIVE

ETTPLTEPEKLPLKEENGKEALNPETIEIKVSNDIIQSKEDDSKA

VPKGEDGEMELFVSTQNPMKTQECVASALGVPANRILVRVKRMGGGFGGKESRSTLLSTAVAVAAFKTGHP

KYEDTAPCINLKSWVQTLRPVSAKIKPLGSRDVLPNNRQLYEMILTYNFHQPKSGEVTPSCPLLCELLYESEFD
WKKKLSAATEMGDATEVKRCKNMEILYDSLQLAHKCILNSFYGYVMRKGARWYSMEMAGIVCFTGANIITC

PGWEERQDEKGRSYYIDHNSRTTTWVKPVVQTTIETSQLSAVQSTPTGRQPQTSSDSSERSSNQQPEVEQG

SSAYKCITNPGHSHLGLQQPAGGVPTIQEVPRSIVVHPSSPEQSSLIMLQPQVSQHLSSSCSLGYQQTLYPN

QTPGHSLTPPPSLSSHGSSSSLHLGGLTNGSGRYISAAPGAEAKYRSATSTSSLFSSSSQLFPPSRLRYSRSD

GLCKEGMGFGDRTSTFCGTPEFLAPEVLTETSYTRAVDWWGLGVLIYEMLVGESPFPGDDEEEVFDSIVND

CFHLCSRARFERLETHMTPEMFRTPLHEIALSIKLLRLGGIGQFLAKAIEPPPLDAVIEAEHTLRELDALDSNDE

YDIRDCAALKTTRMSRRYFWRLHFHDDRQLDFYLKPFLCDARLEWVCQITKGSTPKTPEWYTPDVDGIHC

TEGPSITELPVSGSTLHDQIAMLSHQRFNALTARIQHSLLGRKILAAIIMRRGEEGLGVVVSIGTGNRCVKGEI
CVPVQLFGGWYFYIQAYKALKHKTANMDVLIVLATTIAFVYSFVILLVAMAEKAKVNPVTFFDTPPMLFVFIALG

RPPRIPPRGASRPVSNENFFPPPVPPRGR

HHSHCSADDQSTKAIDIQNAYLNGIGDFSVWEFSGNPVYFCCYDYFAANDPTAIHIVLFSLEEPYEIQLNQV

TLAFWKHGMQGRSLEGNEVTQEITDKSRVFRVLGANRDIILESTPTENPLAHSNLYILTGHESSY
CPAGWEGATCNIARNSSCLPNPCHNGGTCVVSGDSFTCVCKEGWEGPTCTQNTNDCSPHPCYNSGTC
QMGFQRRYGGITDPGTVHRAPSHFSRLPLGGWAEDGQSARHPEPVPEESSEDELPPQIHKV
RSNKASSNLLSPKAIGIAIARYDFCARDMRELSLLKGDVVKIYTKMSANGWWRGEVNGRVGWFPSTYVEED

EGMLCIAEMIHISNCIEDFVTAIKVSLHFALHEDNGSSRTPHLVLNPLYNSSMTEIPFEKNCGEDGVCNADL

LNASLRAGEGS

SMLRPSSVIIVHPLAKPGADPCLHENGGCDQICEDRFGVAHCSCHKGFMKTQNGKTCQTLNIPETAAG

SLTTWEIQGVGISDKGICVADALQVQVLKDVFLSMHIPYSVVRGEQIELKGSVYNYRDSRMMFCVKATVG

SVKGCSNYNMKEETEHLENRNQYSNLCSINKVNGHSERAPQSPRKAELKKSITVDTDSSVDICISKSTEDL
TLLERIDVCCTPFGKRLLKQWICAPLCNPHSINDRLDAVEDLLAFPDKMFEVSELLKKLPDLERLLSKIHSIG
KRANENKTDLAVRGTVTPPPRLLKKNEETTDDVFKDTESSPGSSPPSLTPKLGRRQPIIAPSSSLSHKDDGIKS

VVDPYVYVEIHGIPADCAEKRTKTVHQNGDNPIFDESFEFQINLPELAVMRFVVLDDDYIGDEFIGQYTIP

DDVSSQAHKEVKDKPFDVKCELKYADSDAEASSSIPFPPTGTSFLSCVGSDSPPSPTNTMPDVWSSTELVM
YRISVEILDRREYTRFVKEQERAKGNEVNNPLFQSATTTVINPRYNED
DSKHTNVCKKDMGNGHKKILYGEEYEVIKPLSPSIPYIVTEQDFRSQMLLSALMPTRRIQHQHLEISKECLC
WYGPRDVQLSMAPQGKIVEGMEVMLRCWANARPPVLGYDWYRDGQLQQRQERQSQAVLSILAVKVE

NPHTLLLHTDVNSITASPTESVSVVTSFVPTQLPPSINTSTEFGTQTKTNVTEVTNRTSATTAITTTNSSIVGHRTT
PSSLPVEVLASEDAAFGLKDLSGSIDDLPTGTEATLSSAVSASGSTSSQGEQSNPAQSPFSPHASPHLSGIP

HYLMLQEQPRFSGKQVVALTLDLSDGFLFWLVQDSLCLNLYRVSLCGESCGDAVVTEFAEWSSSDVSHS
SPLRVPAGNNIYISPLKSPYKFSEGMLSPTKMTPRSRILVSIGESFGTSEKFQKINQMVGSSDHQLKRSAEVS
STAPVSLPPARPSARDNPKHSPLLNRTPSDYDLLVPPLGEDSFDSSPPSLPPPPPPARHSLTEHSKPLGSGS
GPEEMMKLRPGVSEMLSSQQKMVSLPFGEHPQQEYGMGPRPFLPMSQGPGGGLRNLREQIGPDQRTN
RVYEEENFNMLTFEDLLCFSYQVAKGMEFLESKLCIHRDLAARNVLVTCGKVVKICDFGLARDIMNDSNY

KDYASDSGCFDDDEVPNVSAGAETGNASILSMFLNSSFLQGQALPRAIAQRRGPEPEKDESSKLAEPKKLPL
DDQSMDITRNHTAALDFAPILVAQEYESTHIQNSVISKPKQLQNKTLLFSGDSDIDITRSQTAAIECGSLRMNS

FSYCDVKPSSPTKNQSKQEPPC

EEKVPMRPEDPKAIPPLMVSPHQKKEGDKQALPSPPPLPSAVTAGKPLAKVAAPEPPVPVHVRSALEGGHINN
LGCWQREPQQRLNIKEIYKILHALGKATPIYLDILG
ALPCSQPSAVPAPEQAPPVFSVSSPFQPVAFHPSPAALLPVIVPNDYSGPPAPKKEIIMGRPGTVWTNVEPRS

EWVVKLYYSFQDKENLYFVMDYIPGGDMMSLLIRMEVFPEHLARFYIAELTLAIESVHKMGFIHRDIKPDNII
IHGKCNDGLNGYRCDCDPGWSGTNCDINNNECESNPCMNGGTCKDMTSGYICTCREGFSGPNCQT
VPAQVPIMEHPLPGNMFDAITHFTQPIMHLPQPEMPPHLPQQPEHSTPPHLNQHAVVPPPALHNALPQQPS
FPFIEFRKKQEKSCKNRALLQLMKGKKHLMEVDTTLVRSWISLLPLGDLAEFLTDLSVELLDCLLGIYHRLQN
RDSSHSRERSTEKPERSHKGSKKKKHRRVRERSRSTSSSSPSSRSYKGEDYPEESEEREECTTGFDKSRLGT
RTKDSDFCYISDDTSPSLYHSEVEIEVEPTDIKVTQETLMDIHVEPQTVTCDYKNTEDQNSTKSACEDELYDIS

DSSYQVSVSTVGRNGLRSRAVTLVCNTSAEVLPPPVRADVPRVEPGSRVLISPDTFSEENGRIEYYAVIVTSNE
VALGPFPGLTGEEEEEAAAGLEQAGAAGRCSPQLSEDGLEESRGGGDPESSQTPVSFNSEPKGSPMLLEPE
GKLDSQLNNVRERARDSAEIETAVVKHVLSELKELSFRSLGDEASGYGTPKTTAPLLFSSASGQSCLPIEPDYKI

IMKTHGQIGKEVYRCDICQMPFSVYSTLEKHMKKWHGEHLLTNDVKIEQAERS
GESAERSQRPRKQISTPAEPFSSTNPTPLFPWFTPSPQTERGRNKDRATEELSKDKDIDKSMEKDKSRERDRI
NKLSKSKKSDILDQSSDSDEMPAVLKEVAMMSHSSSDIDSNSETSNNIQKNALNDLTKQPVKYENGKRKRI

EAYVMAGVGSPYVSRLLGICLTSTVQLVTQLMPYGCLLDYVRENKDRIGSQDLLNWCVQIAKGMSYLEDVR
ADKTHRNPENCSDGMQTGKGLFYVKISMEDLNACSVVPDTDSQVCENQGSQSLLQPHSTNVDQAACPV

DCSIRLWNLESKTCIQEFTAHRKKFDESIHDVAFHPSKCYIASAGADALAKVFV
QTNSIVNTRERSCYTPVPEKEVHVEKDAVKIPNKETQLENSSLQPTFLSLIKNRNLTVEQVVAIEALTKLSEAPLEA
LSSPESSLTPPLSTNLHLESELEALGSLENHVKTEPTDLSESCKQSEHSLVNGKSPVRSLMHRSARVGGEGNN
DDLQFLADLEELINKFQVFRISHRSYTFYHENPYPSIFRINFDHYYPVPYIQYDPLLYLRRTSDMKSKKKRGR
KTVLRTDAPDSTGDRPEILQENTSSGKTAWMGTPHIGDIRKEDDPNEDWCAVCQNGGELLCCEKCPKVFH
DDDS

LKKRGFSKGVKIKHGSSSTLPRMKQKGLKIAKGIF

GSEEENNHGKLDYNWVTRWLDDIGLPQYKTQFDEGRVDGRMLHYMSVDDLLSLKVVSVLHHLSIKRAIQV
AKMCPGNQGNRSQAEGDLKLKQVQKDQQPPPASLERGDEVPATPQESQQPHLMSLEKEADMPEKPLKTS
KLGKNFTVTCNHRSNTEIICCTTPSLKSFNLQLPFATKVFFIFDGVSKQYFDFDYVNNPVFKPFEKPVVISRGI
KAERDILAEADNEWVVRLYYSFQDKDNLYFVMDYIPGGDMMSLLIRMGVFPENLARFYIAELTCAVESVHK
PVSNIHSQRPDPPPLGLSGPTGPTEAQMASEAKSSCRAEPEWRFGIEMGSRQPHLQDGGNSEGTGVKTTA
LLYEKAQLPEKMEKEDFNSCKSVSLLDTPQHQAAPVLPCKPRPIQKSGDDSSEQEEGEVDSEVEFPRRQRP
LDALMGPVTLHSSMEHLVQYTQQGLHWVRNSAHAKDCVPFKCTA
VVYYMVGSIDNLDDLLRCGVTFAANMVVVDKESTMSAEEDYMADAKTIVNVQTLFRLFSSLSIITELTHPANM

ILCPPGTKLDDLPWLECFIKSYNVKDGMEQQVCYQIYDTMVAEDVI

QAHQTCCKAATGTWACCPYHWGFCCSDKRHCCPWGYRCSKSGLDCKRRWALRWDVGTHQPHRLL
DEDEEAAEERRNTHCIWMVGMKGLTESEAASLIVEEARILLQQDLAAMGVQWSPDSFLESDTSSMTSSESE
HCDGEKDCEDGSDEKGCNGTIRLCDERTKFSCKNTGRCISKAWLCDGDIDCDDQSDEDNCDSYMC

KNILATKAGNEFWEVYKNKADPMLIYLHPFRMSRKKFLEVPSNGRESPSENLDDASVWIPQSVSHDTWIKN

QEDIRRALSKHEAVKNELNDRQKELDAFTNKGKHLLAELKRIHNCDPMLVKTDMNSTVDKWLDVSERTED
VRTAGCDTKLQGAICKVNTGSPPSHKWSYSGSCPKSLEDSSWIPFRNHCYTFHMEVTLGQKDAMKRCQK

LDTDQTSVTSPEGWAMSKTSLSYSSEITAENESENPNNEVSLEAWNKGRAYPGEEYSTFENTKSSIDNVQN

EQYDLSSLSKSEDRGENWYAMDNSEPNERRKYYINVCRPLVPVAGCDRYASSCQMKFEKDDDSFSETTS

KCCSFYLDGTPNLKHLNLSFNSVISMSGGLCLNELVSLDFQHSKLNSLGIVPVFLSLGKLIYLDISYTNTHVES
PLSSLVRSCALVACTRHNRLAQKQSPHIMALKNVYIQTKKQKKLHFIVGGYAYLLPKTSVVLRCPVRRFRKSM

GPDILMVLLLVAGAILLIGLVALFIWKLLITVHDRREFAKFEEERSRAKWDTANNPLYKGATSTFTNITYRGNS
TAANPTANTTSQPQEAAAEPANSEEKPQEELVVDVVKTEEPEEKPSEEPCKSEIKKEESEDCPERDKANEKKTE

SGSAMLMLLSVVFVGLAVFVIYKFKRKIPGINVYAQMQNEKDQEMVSPVSQRESMPNIPQSELMSPEQLVD
ELVVKHKYTRRYGDIEKFLKEDQAMGVYLYGELMVNEDAKQQELAYKCFAATQKQMDVSSAKVVAEMLF
SEDEGIVPGTTPLDMASSWEVFDILNGKPYESKLASDDLLLQGDMKKLNEDSKQQLYKLLEVPDPNKNWST

SEYLEAANTLLGCSVGPAVFGL

SVLRGRMCVNCPLGTYYSLEHRTCESCWTGSYQDEEGQLECKSCPSGSYTEYLHSRSISECKAQCKQGT
KDMTFIHEGNRTLAENLINFEKMRMMAKAVRIIHHCQSHTYAPISPLRSRATQLLEDPQTLRISTCSEQSLSG

CEVNAPIQGSRNLLQGEELLRALDQVN

VQGVLSSMMGSLNAQQGSTESIASFIQRLSGTSNLFEPGTEGAIGFFGDLLSLICQNFSMVDVVMLLHGQ

EPVDTLLVLSPRSEASLAGPALRLVRDRVAPHCLFVNLCLETRGCQGGPVDEVVLAGFNEQVLRFLAVRGT

NASRFINTESGRYMYSAPLRRQLASRGSNVCHMDISEKGGDEIDLETISMDATGYMSDGDVLGKNIRTDD

VMATKPETEDKEKQKKVNQGLRNVALEDCALCQETISSSELAAKARDGDFEDPPDWVPDEVCSYCTACK

NFVGSNNLNLLQPIGQFGTRLHGGKDSASPRYIFTMLSSLARLIFPATDDSILRFLYDDNQRVEPEWYMPI
KDTKENAVRSPQNNVNGRLSKERSESINDLPFGLQFKKECKPKPVCNKNNLEAAKNKRCSLTATSATDDS

LEFNSDTEGTNEGFQLVYTSFELSHCEDPGVPQFGYKISDQGHFAGSTIIYGCNPGYTLHGSSLLKCMTGE

FKLLGNVILKSIKTLQAIPEKIVQAISKGTEGDLFVHYIFMDNEFELPTGAGLQLQVAFSGIATPGAKAGMKIQ
RTLLGDPLPELGRLRAPHPSALQSLARASAKHGKYWACLRLDVPIREDSRPRSGQAGPVREDSPPEDPTGG
GSEDKSVRNNIL
ELGDLVDKEMAATSAAVETAVARIEDMLSKSRAGDTGVKLEVNERILGSCTDLMQAIQVLVLASKDLQREIVE

ETTPLTEPEKLPLKEENGKEALNPETIEIKVSNDIIQSKEDDSKA

VPKGEDGEMELFVSTQNPMKTQECVASALGVPANRILVRVKRMGGGFGGKESRSTLLSTAVAVAAFKTGH

YEDTAPCINLKSWVQTLRPVSAKIKPLGSRDVLPNNRQLYEMILTYNFHQPKSGEVTPSCPLLCELLYESEF
WKKKLSAATEMGDATEVKRCKNMEILYDSLQLAHKCILNSFYGYVMRKGARWYSMEMAGIVCFTGANIIT

GWEERQDEKGRSYYIDHNSRTTTWVKPVVQTTIETSQLSAVQSTPTGRQPQTSSDSSERSSNQQPEVEQ

SSAYKCITNPGHSHLGLQQPAGGVPTIQEVPRSIVVHPSSPEQSSLIMLQPQVSQHLSSSCSLGYQQTLYP

QTPGHSLTPPPSLSSHGSSSSLHLGGLTNGSGRYISAAPGAEAKYRSATSTSSLFSSSSQLFPPSRLRYSRSD

LCKEGMGFGDRTSTFCGTPEFLAPEVLTETSYTRAVDWWGLGVLIYEMLVGESPFPGDDEEEVFDSIVND

FHLCSRARFERLETHMTPEMFRTPLHEIALSIKLLRLGGIGQFLAKAIEPPPLDAVIEAEHTLRELDALDSND

YDIRDCAALKTTRMSRRYFWRLHFHDDRQLDFYLKPFLCDARLEWVCQITKGSTPKTPEWYTPDVDGIH

TEGPSITELPVSGSTLHDQIAMLSHQRFNALTARIQHSLLGRKILAAIIMRRGEEGLGVVVSIGTGNRCVKGE
VPVQLFGGWYFYIQAYKALKHKTANMDVLIVLATTIAFVYSFVILLVAMAEKAKVNPVTFFDTPPMLFVFIALG

PPRIPPRGASRPVSNENFFPPPVPPRGR

HSHCSADDQSTKAIDIQNAYLNGIGDFSVWEFSGNPVYFCCYDYFAANDPTAIHIVLFSLEEPYEIQLNQ

LAFWKHGMQGRSLEGNEVTQEITDKSRVFRVLGANRDIILESTPTENPLAHSNLYILTGHESSY
PAGWEGATCNIARNSSCLPNPCHNGGTCVVSGDSFTCVCKEGWEGPTCTQNTNDCSPHPCYNSGT
MGFQRRYGGITDPGTVHRAPSHFSRLPLGGWAEDGQSARHPEPVPEESSEDELPPQIHKV
RSNKASSNLLSPKAIGIAIARYDFCARDMRELSLLKGDVVKIYTKMSANGWWRGEVNGRVGWFPSTYVEE

GMLCIAEMIHISNCIEDFVTAIKVSLHFALHEDNGSSRTPHLVLNPLYNSSMTEIPFEKNCGEDGVCNADL

NASLRAGEGS

SMLRPSSVIIVHPLAKPGADPCLHENGGCDQICEDRFGVAHCSCHKGFMKTQNGKTCQTLNIPETAAG

SLTTWEIQGVGISDKGICVADALQVQVLKDVFLSMHIPYSVVRGEQIELKGSVYNYRDSRMMFCVKATVG

SVKGCSNYNMKEETEHLENRNQYSNLCSINKVNGHSERAPQSPRKAELKKSITVDTDSSVDICISKSTEDL
TLLERIDVCCTPFGKRLLKQWICAPLCNPHSINDRLDAVEDLLAFPDKMFEVSELLKKLPDLERLLSKIHSIG

KRANENKTDLAVRGTVTPPPRLLKKNEETTDDVFKDTESSPGSSPPSLTPKLGRRQPIIAPSSSLSHKDDGIKS

DVVDPYVYVEIHGIPADCAEKRTKTVHQNGDNPIFDESFEFQINLPELAVMRFVVLDDDYIGDEFIGQYTIPF

DDVSSQAHKEVKDKPFDVKCELKYADSDAEASSSIPFPPTGTSFLSCVGSDSPPSPTNTMPDVWSSTELVM

YRISVEILDRREYTRFVKEQERAKGNEVNNPLFQSATTTVINPRYNED

DSKHTNVCKKDMGNGHKKILYGEEYEVIKPLSPSIPYIVTEQDFRSQMLLSALMPTRRIQHQHLEISKECLC

WYGPRDVQLSMAPQGKIVEGMEVMLRCWANARPPVLGYDWYRDGQLQQRQERQSQAVLSILAVKVEA

TESSSSTSSFPSTAVVATTPSEAIIVTSAMTTTRQPYVCDITVPDTYLIMAVLARRAVLENISDSIKEVLRVQFRRS
GPSPSPVGSPVGSNQSRSGPISPASIPGSQMPPQPPGSQSESSSHPALSQSPMPQDRGFMAGIQRNPQ

LLYYSGRLFWINGFRFITTQEVNQTVSIPFSQPAEFAAFTLAQRSLKPLPGNFSFPPKVIPDSILESSFRIEGN
TAPKPLKRLRFDIEGQDEADGSKHLPGESKFQQKLAEMTSTRTRMQKQKLNDGTDTSATEEK
PSSGHELFLPPSDLFFDPATSVIPLPPARRLTVENVKTNRTSQDYDPLPSISDGSQAPARPPKPLPRRTAPEI
NRLSHMPPLPLNPTSNPNSLNTAPPVQRSLGRKPLDISVAASQVHSPGINPLKSPTMRQVQSPMLGSPSG
IRGNARLPVKWMAPESLFDRIYTIKSDVWSYGILLWEIFSLGVNPYPGIQVDSNFYKLIQNGFKMDQPFYA
PEDEVSLVAEKLANKISTTVAKIDKLVEDISTLHTSSDDSFSLSSEDHCLDWQYGSEEVHEARAQSCSPCRL
SNQKNDTPGNSQIPALAAPRFSFTSGDKAIPSDIKGNVHFRKTAGIHIDSKNSKLEQQSRTMVTLNKVLLNS

AYAQSTPASDLQKGRGSRNKPTNLWNPTYGSWFAEKPVVKNSSLLEKEVSERENLGHEGNCTVGPSIVT

VAVFPWHSLVPFLAPSQPDSSVQPSEGQQPVSHPLVSNQNKEPPESAAIAHDLPGAVAGAELGASRPNSP

LIDLDGHIKLTDFGLCTGFRWTHNSKYYQKGSHSRQDSMEPSDLWDDVSNCRCGDRLKTLEQRAKKQ
TNINECASNPCLNQGTCIDDVAGYKCNCLLPYTGATCEDVLAPCAAGPCKNGGECRESEDYESFSCVCP
RPSNRAAALPPKPSRPPATSPAISQQPMLPQPPLPQPPQVLLEDEEPSTPHHSLPLSTSQMQLYLQQLQKV
TRIRWCSMEMIEKLLETPLHVLDRNQHRMLQVNSWKSCLHICLKLHRMICLQMKELKYYKLPATNAEIISR
TKEFAGPNERGRARGTFFRARGRGWNRGTFSGNNNNNNSSSSNDFQKRNRDEEWDPEYTPKSKKYYL
SSHEPSLTVDSLSRDSSQEECSLEIKSDNEIITPQPEENLASRLDRAMQNESEECLQHSETEMMIELDAQQ

SLSRPTQETVSSTWYDHYYGQEDSYVAVLLPNPFHPDKRSTPQTWSVPVGTDECGQSREICNGKLKANA
ELGPATLMEVYRTGGKLLERGEEVGPEPPVLRPRPTLIKSDIVNEISNLSQSDTRSSFPGSEPLLGSPDPEGG
STLLMMLKDMHDSKTKEQQLLSSQNVVPYRNAGMSSSSGSNSASGLKALGTKGPTYKSEKNGDCVLEAT

EREKENKRESRKEKRKKGLEIQSSSALFPVGRVSKEKVVNEDVAAAASSAKKTAGRKKPTAIDPVADASSVAL
KSSTSGSDLVAKKGKSTKGSTCAKKKRQNPSESSNYDSELEREIKSLSKIESAKKAKKKYPRKEENYDSSSEE

LVHRDLAARNVLVKSPNHVKITDFGLARLLDIDETEYHADGGKVPIKWMALESILRRRFTHQSDVWSYGVT
QNLSENNQCAATEPKKKELCAGSPKTNRHCTENDLERFRNRKSHEAKFISELNTETEDSEIDTQYLRNMF

ASAADKTKNIEHAEQTTSHLLHNYKRDISCSLTSTLKEIKETYLQNEQHLFSRYARSQKQPLNKAVLYNGQSTV
NKEDDPNEDWCAVCQNGGDLLCCEKCPKVFHLTCHVPTLLSFPSGEWICTFCRDLSKPEVEYDCDNLQ
PAKTNDTMTKVPFLQGFGYPIPSGSYYAPYGMPYTSMPMMNLGYYGQYPAPLYLSHTLGAASPFMRPAVPP
HLSCHIPSLMTFPRGEWICTFCRDLSKPEVEYDCDNPNQSSEKRKLEGYMGLAPIDRRKCERLLLYLYCHE

VLRINNFEPNCLRRRPSDENNITPSEVTQWTNHRVMEWLRSVDLAEYAPNLRGSGVHGGLMVLEPRFNV
SNLEQEEERAAEQSPCPSTGTLFHFALPKEGDVIQPLTSVTPSLIGQLKMGPRRHSTPIVDGSCPDRTIVTSDL
NQNILEIKGNYIDSEAIKGEVLKVGNRSCESIRLQSESIFCMVPRDLLKSHSELNIEWKQEVLSTLIGKVLVLQI
MGFIHRDIKPDNILIDRDGHIKLTDFGLCTGFRWTHDSKYYQSGDHSRQDSMDFSNEWGDPANCRCG
AGPGDPEQCEAEATRGCGNGTSSRDDRKEVGEASSAMGDGPSKMATIASSEFLPGGMEDVGKHDDDD
PHRCISSCQSYSTFSSENFSVSDGEEGNTSDHSNSPDEMANRLEDQLTEKLEEMLSQTPEIPIEISTQSDGLS

IRFMQFRAKDCYSLALSKLEKKEREKGSNLAFMFRLPFAAGRVFSISMLDTLLYQSFVKDYMISITRLLLGLDT

LEDRKCKPNGECSLGILKVFSNERYRILPDQGTNFQPGNSSNIKKGNKRLLSASMSNPRPQNESQEDHTC
GPPKYPCANDTSICLQPDKLCNGRRDCPDGSDEGDLCDECSLNNGGCSHQCSVVPGRRIVCFCPPGF

LTCSLLDSGGVRNEVLQLLKPMCEVKTDFCQTLLPYLIHDILLHDSNESWRNLLSMYIQRFFTTCCRYSSSS

DNVEKLSVSVSLWDDVLTIGDEIDVWCSNVISQLNEGISNLNSSQKMEAFLKEFQSEVINKGQKLEQLSSKVS
KAGGTVLSIQDEMENVFVWEHLQAYESQSKGAWLGMTFNPKGGTLVWHDDTAMNYSNWGQHDTGPSN

NSNLAIENQHILGDTKCRPKQFENMDIWDLYEKDIEKDVLETLVPWEDSVLSYRCSDFSSSNAGEDLVVSP

SISNLGIASKGLVIEGRGRVLLTYTNGSVCISADGKRTSYTTIIHFTCSKGTLNSSPRFIDNRECVVTFLWETEA

SQYTFCGLDSLQVLKMAGNSFKDNTLGDSFMKLTQLLTLDVSNCKLIQVSLSTFSSLGKLQNLNISHNKLLTF
ITWEKDNKQLLSSVHITIAPYGYMKIHRLKPSDAGTYTCTAGPVREHFVIKLIGGNNKLIINQPAGIREEEGTR

ESVSSTAGVEAAPKAEKKESHKPGKAAGLNPDSDSSATCSADEMDEQDAVDKNRLLSPRPSLLSAATDARLN

EKLDTWPIGKARTATLGIDLNAKELKSPRLQ

LAQKLGLGILSNAFRLSPTPAKTLLDNYEVSGGTVKELVDALRLMDYTEAIEIIQKVLSIPLNQSSNEDNVLKAP

YSPNGLETCESCPLGTYQPVFGSKNCIACPENMSTVKRGAVDISACGVPCPAGEFSRSGLMPCYPCPRDY
GRSPASTRAYLQHLKVIDNQKELSRLSRELES

FQPLQRVQPQLRRFFHQEYLGGREPSDHNVRVATHSLITGLQEYIRESFASVRVQDGVDITRTNLEFLQEHI

SCLLEHVGRMDKIHGLPPRLGAPSRHPELSSALPTLTPAPWSRWRSIRLFISSPVRDMQGERDVLLRAVLPA

ITSGYMTDGGLGLYARRLNRLPDGMAAVRETLQRNTSLGLGDADSWDDSSSVSSGISDTIDNLSTDDINT

APFTVIRRKHHCRSCGKIFCSRCSSHSAPLPRYGQMKPVRVCTHCYMFHVTPFYSDKAGI

IPLVLINGAEGIGTGWSSRIPNFDIREVVNNIRLLMDGEEPLPMLPSYKNFKGTIDELGPNQYLINGEVSILDS
SSVDSDDSIEQEIRKFLAEKAKESANSMEIRRDNLTADSLRVTKPRANKGKTKHQLIKNETNILSGQSEKTKKV

RRAWDYPLPSCIAECGGRFKGESSGRILSPGYPFPYDNNLRCMWMIEVDPGNIVSLQFLAFDTEASHDIL

PKNMQADLIIKPSMAIEFITHVGANIPAFARSGVQMISKIYHESGIEARVGLRAGQLKLSIPAPKTPTKLFSFSNT
SQSPPVSEGSVDDILRLGALCLSDKAGADSGSPQLSPDSALVGRASTGGAEPPAAEESEDPRSLLDEGAALI

SGRGAASPKEFYARNSRWTEGLISASKAVGWGATVMVDAADLVVQGKGKFEELMVCSHEIAASTAQLVAAS

PVRCMLDRDEDMLITGGRHSFLGRYKVGFMKNGKINSLEVSYYSNGGNSEDLSVAVMDRALFHLDNSYN

DSQLWIIFDQNKRQMGSGDAYPHQYSLKLEKGDYTIRLQIRHEQNSELDRIKDLPFVVSHRLSNTLSLDIYE
QARELIEQIGRPLELDTDGIWCVLPNSFPENFVIKSTNAKKPKVTISYPGAMLNILVKEGFTNDQYQELVDPAS

GFLPKGWEVRHAPNGRPFFIDHNTKTTTWEDPRLKIPAHLRKKTSLDPVDLGPLPPGWEERTHTDGRIFYIN

NSPSSPVPSVTQEPTYLQSCSPTHSSIMGQQQHQLPKVQRNESPTNQPLSLPELHEDSNHNLAPIPVMIKRE

IMPSGRSRLLEDFRNNRFPNLQLRDLIGHIVEFSQDQHGSRFIQQKLERATPAERQMVFAEILQAAYQLMT

EVRYPRFLSTEAISIMRRLLRRNPERRLGAGEKDAEDVKKHHFFRVIDWNALLAKKVKPPFVPTIRGREDVSN

LTPLGRILAKLPIEPRLGKMMIMGCIFYVGDAVCTISAATCFPEPFISEGKRLGYVHRNFAGNRFSDHVALL

PPLVIDGAEFWFVPDKNLSYQEAAVYCANNDSDLASVDSYTKLRALLTKIEKLSGAEQKWWLKYMDRGTS

LSLKGETVNDCHAEIISRRGFVRFLYSELMKYNPTDPSSAEQSIFESAGENRLKIKDDVTFHLYVSTAPCGD
RWLEHMAKGKTSEALSRLISLQATEATLVNLGPGNLLLSEDQVDVELVQRGDIVKVVPGGKFPVDGCVIEG

TFWLNFLKSLVPVEEPIAFGGRLKNPLHVVLVATHADIVNLPRPVGGEFGYDKDMSLLKEIRNRFGNDLQI

VDGDNWYRCECAPGFAGPDCRININECQSSPCAFGATCVDEINGYRCICPPGRSGPRCQEVTGRPCI

E

QIHFHQNTSQQLLVSPSTRLEVVLELTNRGEDAYHTAVHLPQLPGLSFRKASVLESSVQARVSCNGLETPDT

GISVHKKAAPQNKTATLETLYQSPLNSGTYGDTNNGEELKTQQLLMAEIMVSDEDDCAVLECNVNAQCVPL

GICLFKGSASAQQGLQSCKFQNLAGSSVASVTFRILPLELGLHTINFTLLTEQNSEVLVKTLRVVPEGIKREL

SPQRSGPVVKSHSITNIDNGALKIFDIVNDNGSQESSSVVKSASAGPDGKNIVRSKSATLLYDQSLQVFPGS
SPLKSETHPDNRAIFYEETRYSKKKITDFLSALEGFKVMLETIAIMEEIAADFKSKILKQLVTLKSKNPEGCFPDL
SALGTAGMADQGPTVKPNSAGVVVASKASTEEPRMRRLKQTSESSGKDKVKLSKLKPAPPPPLSSASIGKPG

ECLQAGYRHVPLQSLTGEILAHAFLFVHVAITNRRGGGKPHKRGLSVRKGKKSREYASMRTLWIKTIDEVF

VMLLPPFLSDQAQHVSVHGTFHLFCQATGPSDAHFVWKKNGQKMEMCVSEQSHALADGRVHILSWVK

ESLHSSIENCKENEKEEVLEHTDFEEETKINPYLKNSLIQRNF
SGHYHCRARNQISYGDSSPIRLTVYFSSMTIAKNSVLGVGVVVAFITLLLALGFTLKRWRKRRLPELVVAPLG

TESSSSTSSFPSTAVVATTPSEAIIVTSAMTTTRQPYVCDITVPDTYLIMAVLARRAVLENISDSIKEVLRVQFRRS
GPSPSPVGSPVGSNQSRSGPISPASIPGSQMPPQPPGSQSESSSHPALSQSPMPQDRGFMAGIQRNPQ

LLYYSGRLFWINGFRFITTQEVNQTVSIPFSQPAEFAAFTLAQRSLKPLPGNFSFPPKVIPDSILESSFRIEGN
TAPKPLKRLRFDIEGQDEADGSKHLPGESKFQQKLAEMTSTRTRMQKQKLNDGTDTSATEEK
PSSGHELFLPPSDLFFDPATSVIPLPPARRLTVENVKTNRTSQDYDPLPSISDGSQAPARPPKPLPRRTAPEI
NRLSHMPPLPLNPTSNPNSLNTAPPVQRSLGRKPLDISVAASQVHSPGINPLKSPTMRQVQSPMLGSPSG
IRGNARLPVKWMAPESLFDRIYTIKSDVWSYGILLWEIFSLGVNPYPGIQVDSNFYKLIQNGFKMDQPFYA

LPEDEVSLVAEKLANKISTTVAKIDKLVEDISTLHTSSDDSFSLSSEDHCLDWQYGSEEVHEARAQSCSPCRLS
SNQKNDTPGNSQIPALAAPRFSFTSGDKAIPSDIKGNVHFRKTAGIHIDSKNSKLEQQSRTMVTLNKVLLNSA

MAYAQSTPASDLQKGRGSRNKPTNLWNPTYGSWFAEKPVVKNSSLLEKEVSERENLGHEGNCTVGPSIVTN

SVAVFPWHSLVPFLAPSQPDSSVQPSEGQQPVSHPLVSNQNKEPPESAAIAHDLPGAVAGAELGASRPNSPA

LIDLDGHIKLTDFGLCTGFRWTHNSKYYQKGSHSRQDSMEPSDLWDDVSNCRCGDRLKTLEQRAKKQH
TNINECASNPCLNQGTCIDDVAGYKCNCLLPYTGATCEDVLAPCAAGPCKNGGECRESEDYESFSCVCP
SRPSNRAAALPPKPSRPPATSPAISQQPMLPQPPLPQPPQVLLEDEEPSTPHHSLPLSTSQMQLYLQQLQKVQ
TRIRWCSMEMIEKLLETPLHVLDRNQHRMLQVNSWKSCLHICLKLHRMICLQMKELKYYKLPATNAEIISRI
TKEFAGPNERGRARGTFFRARGRGWNRGTFSGNNNNNNSSSSNDFQKRNRDEEWDPEYTPKSKKYYLI
SSHEPSLTVDSLSRDSSQEECSLEIKSDNEIITPQPEENLASRLDRAMQNESEECLQHSETEMMIELDAQQ

ESLSRPTQETVSSTWYDHYYGQEDSYVAVLLPNPFHPDKRSTPQTWSVPVGTDECGQSREICNGKLKANAG
ELGPATLMEVYRTGGKLLERGEEVGPEPPVLRPRPTLIKSDIVNEISNLSQSDTRSSFPGSEPLLGSPDPEGG
FSTLLMMLKDMHDSKTKEQQLLSSQNVVPYRNAGMSSSSGSNSASGLKALGTKGPTYKSEKNGDCVLEATG

EREKENKRESRKEKRKKGLEIQSSSALFPVGRVSKEKVVNEDVAAAASSAKKTAGRKKPTAIDPVADASSVALV
KSSTSGSDLVAKKGKSTKGSTCAKKKRQNPSESSNYDSELEREIKSLSKIESAKKAKKKYPRKEENYDSSSEEQ

LVHRDLAARNVLVKSPNHVKITDFGLARLLDIDETEYHADGGKVPIKWMALESILRRRFTHQSDVWSYGVTV
YQNLSENNQCAATEPKKKELCAGSPKTNRHCTENDLERFRNRKSHEAKFISELNTETEDSEIDTQYLRNMFR

ASAADKTKNIEHAEQTTSHLLHNYKRDISCSLTSTLKEIKETYLQNEQHLFSRYARSQKQPLNKAVLYNGQSTV
NKEDDPNEDWCAVCQNGGDLLCCEKCPKVFHLTCHVPTLLSFPSGEWICTFCRDLSKPEVEYDCDNLQ
PAKTNDTMTKVPFLQGFGYPIPSGSYYAPYGMPYTSMPMMNLGYYGQYPAPLYLSHTLGAASPFMRPAVPR
HLSCHIPSLMTFPRGEWICTFCRDLSKPEVEYDCDNPNQSSEKRKLEGYMGLAPIDRRKCERLLLYLYCHE

VLRINNFEPNCLRRRPSDENNITPSEVTQWTNHRVMEWLRSVDLAEYAPNLRGSGVHGGLMVLEPRFNV
SNLEQEEERAAEQSPCPSTGTLFHFALPKEGDVIQPLTSVTPSLIGQLKMGPRRHSTPIVDGSCPDRTIVTSDL
NQNILEIKGNYIDSEAIKGEVLKVGNRSCESIRLQSESIFCMVPRDLLKSHSELNIEWKQEVLSTLIGKVLVLQI
MGFIHRDIKPDNILIDRDGHIKLTDFGLCTGFRWTHDSKYYQSGDHSRQDSMDFSNEWGDPANCRCG
AGPGDPEQCEAEATRGCGNGTSSRDDRKEVGEASSAMGDGPSKMATIASSEFLPGGMEDVGKHDDDD
PHRCISSCQSYSTFSSENFSVSDGEEGNTSDHSNSPDEMANRLEDQLTEKLEEMLSQTPEIPIEISTQSDGLS

IRFMQFRAKDCYSLALSKLEKKEREKGSNLAFMFRLPFAAGRVFSISMLDTLLYQSFVKDYMISITRLLLGLDT

LEDRKCKPNGECSLGILKVFSNERYRILPDQGTNFQPGNSSNIKKGNKRLLSASMSNPRPQNESQEDHT
GPPKYPCANDTSICLQPDKLCNGRRDCPDGSDEGDLCDECSLNNGGCSHQCSVVPGRRIVCFCPPG

LTCSLLDSGGVRNEVLQLLKPMCEVKTDFCQTLLPYLIHDILLHDSNESWRNLLSMYIQRFFTTCCRYSSSS

NVEKLSVSVSLWDDVLTIGDEIDVWCSNVISQLNEGISNLNSSQKMEAFLKEFQSEVINKGQKLEQLSSKV
AGGTVLSIQDEMENVFVWEHLQAYESQSKGAWLGMTFNPKGGTLVWHDDTAMNYSNWGQHDTGPS

NSNLAIENQHILGDTKCRPKQFENMDIWDLYEKDIEKDVLETLVPWEDSVLSYRCSDFSSSNAGEDLVVSP

SISNLGIASKGLVIEGRGRVLLTYTNGSVCISADGKRTSYTTIIHFTCSKGTLNSSPRFIDNRECVVTFLWETEA

SQYTFCGLDSLQVLKMAGNSFKDNTLGDSFMKLTQLLTLDVSNCKLIQVSLSTFSSLGKLQNLNISHNKLLT
ITWEKDNKQLLSSVHITIAPYGYMKIHRLKPSDAGTYTCTAGPVREHFVIKLIGGNNKLIINQPAGIREEEGTR

ESVSSTAGVEAAPKAEKKESHKPGKAAGLNPDSDSSATCSADEMDEQDAVDKNRLLSPRPSLLSAATDARL

EKLDTWPIGKARTATLGIDLNAKELKSPRLQ

LAQKLGLGILSNAFRLSPTPAKTLLDNYEVSGGTVKELVDALRLMDYTEAIEIIQKVLSIPLNQSSNEDNVLKAP

YSPNGLETCESCPLGTYQPVFGSKNCIACPENMSTVKRGAVDISACGVPCPAGEFSRSGLMPCYPCPRDY
RSPASTRAYLQHLKVIDNQKELSRLSRELES

FQPLQRVQPQLRRFFHQEYLGGREPSDHNVRVATHSLITGLQEYIRESFASVRVQDGVDITRTNLEFLQEHI

SCLLEHVGRMDKIHGLPPRLGAPSRHPELSSALPTLTPAPWSRWRSIRLFISSPVRDMQGERDVLLRAVLPA

ITSGYMTDGGLGLYARRLNRLPDGMAAVRETLQRNTSLGLGDADSWDDSSSVSSGISDTIDNLSTDDINT

PFTVIRRKHHCRSCGKIFCSRCSSHSAPLPRYGQMKPVRVCTHCYMFHVTPFYSDKAGI

IPLVLINGAEGIGTGWSSRIPNFDIREVVNNIRLLMDGEEPLPMLPSYKNFKGTIDELGPNQYLINGEVSILDS
SSVDSDDSIEQEIRKFLAEKAKESANSMEIRRDNLTADSLRVTKPRANKGKTKHQLIKNETNILSGQSEKTKKV

RRAWDYPLPSCIAECGGRFKGESSGRILSPGYPFPYDNNLRCMWMIEVDPGNIVSLQFLAFDTEASHDIL

PKNMQADLIIKPSMAIEFITHVGANIPAFARSGVQMISKIYHESGIEARVGLRAGQLKLSIPAPKTPTKLFSFSNT
SQSPPVSEGSVDDILRLGALCLSDKAGADSGSPQLSPDSALVGRASTGGAEPPAAEESEDPRSLLDEGAALI

SGRGAASPKEFYARNSRWTEGLISASKAVGWGATVMVDAADLVVQGKGKFEELMVCSHEIAASTAQLVAAS

VRCMLDRDEDMLITGGRHSFLGRYKVGFMKNGKINSLEVSYYSNGGNSEDLSVAVMDRALFHLDNSY

SQLWIIFDQNKRQMGSGDAYPHQYSLKLEKGDYTIRLQIRHEQNSELDRIKDLPFVVSHRLSNTLSLDIYE
ARELIEQIGRPLELDTDGIWCVLPNSFPENFVIKSTNAKKPKVTISYPGAMLNILVKEGFTNDQYQELVDPAS

FLPKGWEVRHAPNGRPFFIDHNTKTTTWEDPRLKIPAHLRKKTSLDPVDLGPLPPGWEERTHTDGRIFYIN

SPSSPVPSVTQEPTYLQSCSPTHSSIMGQQQHQLPKVQRNESPTNQPLSLPELHEDSNHNLAPIPVMIKRE

IMPSGRSRLLEDFRNNRFPNLQLRDLIGHIVEFSQDQHGSRFIQQKLERATPAERQMVFAEILQAAYQLMT

EVRYPRFLSTEAISIMRRLLRRNPERRLGAGEKDAEDVKKHHFFRVIDWNALLAKKVKPPFVPTIRGREDVS

LTPLGRILAKLPIEPRLGKMMIMGCIFYVGDAVCTISAATCFPEPFISEGKRLGYVHRNFAGNRFSDHVALL

PPLVIDGAEFWFVPDKNLSYQEAAVYCANNDSDLASVDSYTKLRALLTKIEKLSGAEQKWWLKYMDRGTS

LSLKGETVNDCHAEIISRRGFVRFLYSELMKYNPTDPSSAEQSIFESAGENRLKIKDDVTFHLYVSTAPCGD
RWLEHMAKGKTSEALSRLISLQATEATLVNLGPGNLLLSEDQVDVELVQRGDIVKVVPGGKFPVDGCVIEG

TFWLNFLKSLVPVEEPIAFGGRLKNPLHVVLVATHADIVNLPRPVGGEFGYDKDMSLLKEIRNRFGNDLQI

CVDGDNWYRCECAPGFAGPDCRININECQSSPCAFGATCVDEINGYRCICPPGRSGPRCQEVTGRPCI

DE

QIHFHQNTSQQLLVSPSTRLEVVLELTNRGEDAYHTAVHLPQLPGLSFRKASVLESSVQARVSCNGLETPDT

GISVHKKAAPQNKTATLETLYQSPLNSGTYGDTNNGEELKTQQLLMAEIMVSDEDDCAVLECNVNAQCVPL

DGICLFKGSASAQQGLQSCKFQNLAGSSVASVTFRILPLELGLHTINFTLLTEQNSEVLVKTLRVVPEGIKREL

SPQRSGPVVKSHSITNIDNGALKIFDIVNDNGSQESSSVVKSASAGPDGKNIVRSKSATLLYDQSLQVFPGS
SPLKSETHPDNRAIFYEETRYSKKKITDFLSALEGFKVMLETIAIMEEIAADFKSKILKQLVTLKSKNPEGCFPDL

SALGTAGMADQGPTVKPNSAGVVVASKASTEEPRMRRLKQTSESSGKDKVKLSKLKPAPPPPLSSASIGKPG

ECLQAGYRHVPLQSLTGEILAHAFLFVHVAITNRRGGGKPHKRGLSVRKGKKSREYASMRTLWIKTIDEVF

VMLLPPFLSDQAQHVSVHGTFHLFCQATGPSDAHFVWKKNGQKMEMCVSEQSHALADGRVHILSWVK

ESLHSSIENCKENEKEEVLEHTDFEEETKINPYLKNSLIQRNF
SGHYHCRARNQISYGDSSPIRLTVYFSSMTIAKNSVLGVGVVVAFITLLLALGFTLKRWRKRRLPELVVAPLG

VELEVYKISPKFSFLVTSGPFVYTAIAVINVLMNSSLLRGQAPVILSLQPSFTMPDPR
ISQYGPQQAGPSMSPHPSPGGQMHPGIGSFQQSNSSGAYGPQMSQYGPQGNYSRPSTYSGVPNTSYS

SSHFHILWNTSTTVEWGTVFYCVESKVLQTRFVDSERCLPPWNLTVPSYKVKGLKPYMQFDFTVTPYTYWG

IHRKAYNSDTSLENVDAKIAKLMGEGFSFEEVKRALEIAQNNVDVARSILREFACFPPPVSPRLNL
NLKSPQTPSQLAGMLAGPTAAAAAASIKSPPVLGSAAASPVHLKSPSLPAPSPGWTSSPKPPLQSPGIPPNHKA
TEEIYFVMKSCWALDSRKRPSFSQLRSLLARQLADAEGAVYQNMESNISTYPSNSKTRPPVSRKNERLPSTD
SDASSFVKRENLSRAHTLLRLKKAYTSLSNDSLNSGSTSDGYCTKEHMKPCTRASFLDCKEELQRYQKRPS
ANSGRLSSTSEASSLSSKQDFTKTARTSMPDTKNSQMASLLEKSVVFFSEENMDLTGSCVVNVPDTVLTERK

NRLPGSSLLLEPSSLTASVKEVPLFRLRHFPCGNISYGYQQQGLPLEASAPPCASNYEDSVHRNKSHVARQG

ASAPQATAAAGGGEGPSQLEDEALGPPRLDSETESDHDDPFLSIVSPEIQLPLQPGKRRTQSLSALPKDRDS

IQRCLAHSLVGTPNYIAPEVLLRKGYTQLCDWWSVGVILFEMLVGQPPFLAPTPTETQLKVINWENTLHIPLQ
SGWQGQTCEIDINECVKSPCRNGATCQNTDGSYHCNCKAGYSGHNCDTDIDDCKPNPCHNGGSCS
PQTPLLPSVKVQSQPQPPLQPPAHPSVQQLQQQQRPIHMQPMQFPPHIPQPQPPPQQHPPPQPQPAK
IARLEPLDGSRDETMQKTPVIEVFQRALAATRSWFREVLKQHLLQGHSDSIVTFTFPDELTVWSQFVRISFPD
HDDREGEGGEKWVNRGRGRGSFPRGRGRFMFRKSSTSPKWAHDKFSGEEGEIEDDESGTENREEKD
IGPDLEKGKPSFKNEHSYYLEVPVEHLEKEKPAEESSTIFTEKIRSDFKDQSLEVSVGNDEENESVVVSAFSEV

QYRFSIAAFTKYDLVDPSVSFTAFSAAGAGADPSPVTVPVVAGIVAGFLLTLGVIFGWVYWKQVRSKRTKTSN
GSLSMELGMTSADSLQKDDASSLPHCTDSLAETDDSLLFEPGAGAKGEGEKGRRRSSPGRSRLKQGRSS
CQNPSASESSLSCSTTTKITGRSTNKREPTDPALSGTNGTNCISRHSCSQSKQSSKLADNATSVRKDLKLEG

DMTAGKTKMPKKGRGGLEKSDLDLSPTVLPLEKDKGLRLSAPSSSTVKHSTSSISSMLAQADKLPMTDKRVA
NKKGSTSKKKIINLKKQQVESSDDAEKSSLEKEETNHSSEDKKIRKGTAVNEKKNRDLKERTSKGKHDGSSD

VWELMTFGAKPYDGIPAREIPDLLEKGERLPQPPICTIDVYMIMVKCWMIDSECRPKFRELVTEFSRMARD
RHSKRLSFTLHPSPVKESVAESTASETLRKSGASCVEYRDNNRDLKNEPTYVKKTVGLDEVCEREEPKTSDSA

VSEPHDKSVNTTSGSYRSHVSQRDTKSLSVSVPNAKSCSGHTTKKNSLYDKVTIATCFNNSCNVHQTSSKLS
HSKKGKTAQGLSPVDQRKCERLLLYLYCHELSIEFQEPVPASIPNYYKIIKKPMDLSTVKKKLQKKHSQHYQT
PPQFHSSSHMKMSTTAKHKAKHGVHLQAPVGMSLGDMQSSLAPPKVGGTSLSSGRLHKRKHKHKHKHKD
EMSLAFQDPVSPTVPEYYKIIKKPMDLSTIKKRLQVNHPLYMKPEDFVADFRLIFQNCAEFNEPDSEVADAGN

ETMAQLLNIPPNKTLLRRHLATHFNLLVGQDAQHQKREAMESPDYVLLTATAKVKPKKLAFSNFGSLRKKKQ
LTAEGSIGTNNVTVESAMASADVLEECEKGDSGAASEADGKLCLRMSLATPVSEESEGSLPFSLEKPAASERK
DQNFTGLIAGVVSTSVLFLLLLGGFLWMKRKKQIKDLGSDLVRYDGRVHTPHLDRLVSARSVSPTTEMVSSD
GDRLKPLERRAARQHQRCLAHSLVGTPNYIAPEVLLRTGYTQLCDWWSVGVILFEMLVGQPPFLAQTPLETQ
EDDEELSIFSSLLASKLHLSSPEGPVGSVLALRDRVGTDSRALSPTWEPVRWDPMRRHSTRLCPGGVLDGS
SDKECAVRRVKTQMSLGKLYADDHSCENQAQFGESDCDSSEGECSDATVRTNNPCSSATW

TPGSGFLCSMKITEDDLWIRTYARLYQKLCSSTGDIPIGIYRTESQKLTTSESREMASQSQISISVEEWEDTKE

QTKEVNGDVVKYSARKRPNLGKGKENREVTFLTKSEAGSGATTQEFPTEKRKRPTLNFTSWKTSKASGCRRT
FYLNPDNKTCEIVDYCTKHLKCSQVCEQHKHNVKCSCYEGWKINNDGESCTSIDPFEAFIIFSIRHEIRKID

RSTTPANSNSEPETFVSGGLDKVSRRTMLDVVDYLRSQKRSLSGTIFDNSFWLDLNYLEVAMVAQSCAAHF

SDLKELTNSQESTADLQFLEADLRQKLEHAKEMSETAKETLEDFSAQKMQLLKFIDHMTDWLTKVEESLLNC
MLSQNSCYWIQSSNGVWHLGSCSNVTMGVICKIPRVEESSFSRAAVPDNTTAIAVVVLSALALCALIALAVYLY

PDTNYSTSDSYISPTFIGDERENEDKHFDRETIFDGMINSNSDEPKILEETDKESLPQPSTRSMPFLSSRNTE

ACPVTTTKDETQSCSVRDPNSGFLFNMQPLASEKGYITSGIGKTFMLNICGPAPDCGGGGSDGKPAGGC

DHSAYKPLQALTVLDFSSNQLTVLTEKALESLPSSLVHLDLSYNLLDCSCNHLSFLKWAKEHGELLRSTELM
KSSLNEALHTQEKHQNGILFNRSKAEKRGHLADPSSQYDDIVSRLLKQRGWPGENLESWEAQESTERNA

SSPQKPLDLKQLKQRAAAIPPIISEGFPDGTLQSGSMKPQVPHHALALYQQQITKAHESARDDVAQNKQP

PSSPSAFATKEETGELYNVKLQDNESMCDSGVETSFRRLSFTYSESLNSKSSLTLSKMALGYGQEGSVQGK

YQPDPGKSYCLSCPFYGTTTIIGARSITDCSSFGSTFTAAEESVMVPVSPENISKQYKVSSQVFHECFLEPC

FSRIATHILRCADPTFGCRLLELCNQGLFECLALNLYCLRGERGALTAVINDRIRRLSADVNPSLVSWLTAM

LRARAAPHRLALQEIDLRWGITEQEARQERQLELCLSEVARSQLFVGILGERYGYVPREYSLPDAPQYEWV

SSSISSYANTPASSRKNVDAQTDAEKHSQIERNSLWSSDEVKKSDGGSDSGIKMESGSKWRRNPSDVSD

TTIEISELPIRTWTQTYKEQVLEPMLNGTEKTPPLITDYREYHTDTTVKFIVKMSEQKLAEAEAAGLHKVFKLQT
SQQTGELKSSRRTVGESAIMHDGGKSASYAENMYLHTTLEFKTEQGTVWSKGLAAGALSVKGNPLSKKSTI

RVWDGPPENEMLLKEISGSLIPESIHSTLNVVTIQFDTDFYISKSGFAIQFSSSVATACRDPGVPMNGTRNG

LYLVSPTKTEVIPPLIENRESWTSCNPFFVGLNYCTKVAYSNASYTDAAPYYPLTGESRIEIEIQPTGSVHEYSA
PSSSISLGVEFTDTGTLYAVTPLPWCPHVDSLQPLPAAGLDALAPCEECGSQGENWVCLACYRVLCGRYV

KVKADKDSANLARLRQASRGVNQATAGVVASTKSGKSQIEEKDSMDFSSMTLTQIKRQEMDSQVRVLELE

IIPNVKGVGTVCKTNLSSNTAFRGFGGPQGMMVAECWMSDVILTCGLPAEQVRKLNLYNEGDLTHFNQK

SHSLALLGKKKSNSLTLPPKHSHPFFVTSLPDDKIPKGAGPGCYLVGALTLSKTELGKKAGQSAAKRQGKFKK
LTYVTRSENSIFFEVDGPYLAMILPASKEEGKKLKKRYAVFNEDGSLAELKGFEVKRRGELQLVKIFQSSVFEA

IHNTKKTQWEDPRLQNVAITGPAVPYSRDYKRKYEFFRKKLKKQSDIPNRFEMKLHRTTILEDSYRRIIAVKR

PEELDQLYLDDALNAVSHCRTSCMTNSYNAIILTMWSWFLPMPVTCGMAVKTYLTLSCS

DVFGNYVIQKFFEFGSPEQKLALATRIRGHVLPLALQMYGCRVIQKALESISPDQQVINEMVKELDGHVLK

FDDEFTSETPILTPPREPRILSEEEQETFRDFDYIADWC

SVFQAWDDARMGGEDAEIRFCEHKRLNMATLRMTWEAKVQLKDILINSGFPEECLMTQIFNNTGPDNNL

HRSLLQLFPRFHDRFWRDCWYISSKGWYRDYPINCNLKLPFICEKSNASLLEKHEPEYEPPKGGCPKEWI

GALFDKSCSDQASVAGESQHQPLFENPKQGKLRTKVENGEGTIPVESSDIVPTWDGIQHGERLRTMSCSD
QSMVDESLITGEAMPVAKKPGSTVIAGSINQNGSLLISATHVGADTTLSQIVKLVEEAQTSKAPIQQFADKLSGY

LDKLFVLDGGASGSKDMKLLRNHLQELRSQIISICPPMTPLCEKMISTLPSWRKINGPNQLMSLQQFVYDVG

NNGRVMSDGAKWDDDCNTCQCWNGKVACSKVWCGPRPCLIHAKGHECPAGHTCVPIKDDHCFTHF

NSRNLSCNVSHPIYWGNSRVLIQLLFDILTNSSWGDYLELDVMASSDNDMNGTLVDNQDSYRIPVLYPIN

EDGAICQCLKGFTREGKSCYDVDECARNMDRCNQHLSDCINTEGGYVCKCLEGYRGDGLHCYDIDE[illegible]

[illegible]AGFSLDPQGVYGIIRRRQEFRHRTPLDLVPKTKIDRTVSIKGQFLGEVISAVLKREGVQFLTNLPKGSAEAEL

SSSSDLVSCTKTLFKFDSNHNPEGANVVRGPVPSGAQMFSVPQYNIQYSSSTTAKDTLWLPKQNAQIEQGS
[illegible]STELKRWDTAFDHNEARKTGVITPKAGFDPDYDKALKDIIDIEENLREYLDKQRKRLGFKAMLYWGAGKN[illegible]
KSSHGPSQEAADSGSGAKSKQLTQVADTISGDGAKPNQLGEGVKKPGLPSVPKPQSSTKLLLTTAAPAPSSSS

[illegible]NALQPIRDATDLRENMQNAVVSFKELCGLSPVANLMQCILAVSTRLVGPDSTPLVLLNLNDQYPTMEIQGT

[illegible]AVAENSEYCCSVLSKAGNKTSKVLITVEGKDSTGQAEWVKEYDSWKSAVSEHHAMMQNWKTTWVSFLT[illegible]

[illegible]RQRRSFFPRRREPLKTQRPTSSVGCLNGVSEDAVNYATLQFPPSHLQGPTAPASVPGNSRQRVKPGPADE[illegible]

VELEVYKISPKFSFLVTSGPFVYTAIAVINVLMNSSLLRGQAPVILSLQPSFTMPDPR
[illegible]SQYGPQQAGPSMSPHPSPGGQMHPGIGSFQQSNSSGAYGPQMSQYGPQGNYSRPSTYSGVPNTSYS[illegible]

SSHFHILWNTSTTVEWGTVFYCVESKVLQTRFVDSERCLPPWNLTVPSYKVKGLKPYMQFDFTVTPYTYWG[illegible]

[illegible]HRKAYNSDTSLENVDAKIAKLMGEGFSFEEVKRALEIAQNNVDVARSILREFACFPPPVSPRLNL
[illegible]LKSPQTPSQLAGMLAGPTAAAAAASIKSPPVLGSAAASPVHLKSPSLPAPSPGWTSSPKPPLQSPGIPPNHKA
[illegible]EEIYFVMKSCWALDSRKRPSFSQLRSLLARQLADAEGAVYQNMESNISTYPSNSKTRPPVSRKNERLPSTD

SDASSFVKRENLSRAHTLLRLKKAYTSLSNDSLNSGSTSDGYCTKEHMKPCTRASFLDCKEELQRYQKRPSI
ANSGRLSSTSEASSLSSKQDFTKTARTSMPDTKNSQMASLLEKSVVFFSEENMDLTGSCVVNVPDTVLTERK

NRLPGSSLLLEPSSLTASVKEVPLFRLRHFPCGNISYGYQQQGLPLEASAPPCASNYEDSVHRNKSHVARQG

ASAPQATAAAGGGEGPSQLEDEALGPPRLDSETESDHDDPFLSIVSPEIQLPLQPGKRRTQSLSALPKDRDS

IQRCLAHSLVGTPNYIAPEVLLRKGYTQLCDWWSVGVILFEMLVGQPPFLAPTPTETQLKVINWENTLHIPLQ
SGWQGQTCEIDINECVKSPCRNGATCQNTDGSYHCNCKAGYSGHNCDTDIDDCKPNPCHNGGSCS
QPQTPLLPSVKVQSQPQPPLQPPAHPSVQQLQQQQQRPIHMQPMQFPPHIPQPQPPPQQHPPPQPQPAK
IARLEPLDGSRDETMQKTPVIEVFQRALAATRSWFREVLKQHLLQGHSDSIVTFTFPDELTVWSQFVRISFPD
HDDREGEGGEKWVNRGRGRGSFPRGRGRFMFRKSSTSPKWAHDKFSGEEGEIEDDESGTENREEKDN
IGPDLEKGKPSFKNEHSYYLEVPVEHLEKEKPAEESSTIFTEKIRSDFKDQSLEVSVGNDEENESVVVSAFSEV

QYRFSIAAFTKYDLVDPSVSFTAFSAAGAGADPSPVTVPVVAGIVAGFLLTLGVIFGWVYWKQVRSKRTKTSNL
GSLSMELGMTSADSLQKDDASSLPHCTDSLAETDDSLLFEPGAGAKGEGEKGRRRSSPGRSRLKQGRSSS
CQNPSASESSLSCSTTTKITGRSTNKREPTDPALSGTNGTNCISRHSCSQSKQSSKLADNATSVRKDLKLEG

VDMTAGKTKMPKKGRGGLEKSDLDLSPTVLPLEKDKGLRLSAPSSSTVKHSTSSISSMLAQADKLPMTDKRVA
QNKKGSTSKKKIINLKKQQVESSDDAEKSSLEKEETNHSSEDKKIRKGTAVNEKKNRDLKERTSKGKHDGSSD

VWELMTFGAKPYDGIPAREIPDLLEKGERLPQPPICTIDVYMIMVKCWMIDSECRPKFRELVTEFSRMARDF
RHSKRLSFTLHPSPVKESVAESTASETLRKSGASCVEYRDNNRDLKNEPTYVKKTVGLDEVCEREEPKTSDSA

VSEPHDKSVNTTSGSYRSHVSQRDTKSLSVSVPNAKSCSGHTTKKNSLYDKVTIATCFNNSCNVHQTSSKLS
HSKKGKTAQGLSPVDQRKCERLLLYLYCHELSIEFQEPVPASIPNYYKIIKKPMDLSTVKKKLQKKHSQHYQT
PPQFHSSSHMKMSTTAKHKAKHGVHLQAPVGMSLGDMQSSLAPPKVGGTSLSSGRLHKRKHKHKHKHKD
EMSLAFQDPVSPTVPEYYKIIKKPMDLSTIKKRLQVNHPLYMKPEDFVADFRLIFQNCAEFNEPDSEVADAGN

ETMAQLLNIPPNKTLLRRHLATHFNLLVGQDAQHQKREAMESPDYVLLTATAKVKPKKLAFSNFGSLRKKKQ
LTAEGSIGTNNVTVESAMASADVLEECEKGDSGAASEADGKLCLRMSLATPVSEESEGSLPFSLEKPAASERK
DQNFTGLIAGVVSTSVLFLLLLGGFLWMKRKKQIKDLGSDLVRYDGRVHTPHLDRLVSARSVSPTTEMVSSD
GDRLKPLERRAARQHQRCLAHSLVGTPNYIAPEVLLRTGYTQLCDWWSVGVILFEMLVGQPPFLAQTPLETQ
EDDEELSIFSSLLASKLHLSSPEGPVGSVLALRDRVGTDSRALSPTWEPVRWDPMRRHSTRLCPGGVLDGS
SDKECAVRRVKTQMSLGKLYADDHSCENQAQFGESDCDSSEGECSDATVRTNNPCSSATW

TPGSGFLCSMKITEDDLWIRTYARLYQKLCSSTGDIPIGIYRTESQKLTTSESREMASQSQISISVEEWEDTKE

TKEVNGDVVKYSARKRPNLGKGKENREVTFLTKSEAGSGATTQEFPTEKRKRPTLNFTSWKTSKASGCRRT
YLNPDNKTCEIVDYCTKHLKCSQVCEQHKHNVKCSCYEGWKINNDGESCTSIDPFEAFIIFSIRHEIRKID

RSTTPANSNSEPETFVSGGLDKVSRRTMLDVVDYLRSQKRSLSGTIFDNSFWLDLNYLEVAMVAQSCAAH

SDLKELTNSQESTADLQFLEADLRQKLEHAKEMSETAKETLEDFSAQKMQLLKFIDHMTDWLTKVEESLLNC
LSQNSCYWIQSSNGVWHLGSCSNVTMGVICKIPRVEESSFSRAAVPDNTTAIAVVVLSALALCALIALAVYL

PDTNYSTSDSYISPTFIGDERENEDKHFDRETIFDGMINSNSDEPKILEETDKESLPQPSTRSMPFLSSRNT

ACPVTTTKDETQSCSVRDPNSGFLFNMQPLASEKGYITSGIGKTFMLNICGPAPDCGGGGSDGKPAGGC

DHSAYKPLQALTVLDFSSNQLTVLTEKALESLPSSLVHLDLSYNLLDCSCNHLSFLKWAKEHGELLRSTELM
KSSLNEALHTQEKHQNGILFNRSKAEKRGHLADPSSQYDDIVSRLLKQRGWPGENLESWEAQESTERNA

SSPQKPLDLKQLKQRAAAIPPIISEGFPDGTLQSGSMKPQVPHHALALYQQQITKAHESARDDVAQNKQP

PSSPSAFATKEETGELYNVKLQDNESMCDSGVETSFRRLSFTYSESLNSKSSLTLSKMALGYGQEGSVQGK

YQPDPGKSYCLSCPFYGTTTIIGARSITDCSSFGSTFTAAEESVMVPVSPENISKQYKVSSQVFHECFLEPC

FSRIATHILRCADPTFGCRLLELCNQGLFECLALNLYCLRGERGALTAVINDRIRRLSADVNPSLVSWLTAM

LRARAAPHRLALQEIDLRWGITEQEARQERQLELCLSEVARSQLFVGILGERYGYVPREYSLPDAPQYEWV

SSSISSYANTPASSRKNVDAQTDAEKHSQIERNSLWSSDEVKKSDGGSDSGIKMESGSKWRRNPSDVSD

TTIEISELPIRTWTQTYKEQVLEPMLNGTEKTPPLITDYREYHTDTTVKFIVKMSEQKLAEAEAAGLHKVFKLQT
SQQTGELKSSRRTVGESAIMHDGGKSASYAENMYLHTTLEFKTEQGTVWSKGLAAGALSVKGNPLSKKSTI

RVWDGPPENEMLLKEISGSLIPESIHSTLNVVTIQFDTDFYISKSGFAIQFSSSVATACRDPGVPMNGTRNG

LYLVSPTKTEVIPPLIENRESWTSCNPFFVGLNYCTKVAYSNASYTDAAPYYPLTGESRIEIEIQPTGSVHEYSA
LPSSSISLGVEFTDTGTLYAVTPLPWCPHVDSLQPLPAAGLDALAPCEECGSQGENWVCLACYRVLCGRYV

SKVKADKDSANLARLRQASRGVNQATAGVVASTKSGKSQIEEKDSMDFSSMTLTQIKRQEMDSQVRVLELE

IIPNVKGVGTVCKTNLSSNTAFRGFGGPQGMMVAECWMSDVILTCGLPAEQVRKLNLYNEGDLTHFNQK

SHSLALLGKKKSNSLTLPPKHSHPFFVTSLPDDKIPKGAGPGCYLVGALTLSKTELGKKAGQSAAKRQGKFKK
SLTYVTRSENSIFFEVDGPYLAMILPASKEEGKKLKKRYAVFNEDGSLAELKGFEVKRRGELQLVKIFQSSVFEA

IHNTKKTQWEDPRLQNVAITGPAVPYSRDYKRKYEFFRKKLKKQSDIPNRFEMKLHRTTILEDSYRRIIAVKRA

EPEELDQLYLDDALNAVSHCRTSCMTNSYNAIILTMWSWFLPMPVTCGMAVKTYLTLSCS

TDVFGNYVIQKFFEFGSPEQKLALATRIRGHVLPLALQMYGCRVIQKALESISPDQQVINEMVKELDGHVLKC

NFDDEFTSETPILTPPREPRILSEEEQETFRDFDYIADWC

SVFQAWDDARMGGEDAEIRFCEHKRLNMATLRMTWEAKVQLKDILINSGFPEECLMTQIFNNTGPDNNL

HRSLLQLFPRFHDRFWRDCWYISSKGWYRDYPINCNLKLPFICEKSNASLLEKHEPEYEPPKGGCPKEWI

GALFDKSCSDQASVAGESQHQPLFENPKQGKLRTKVENGEGTIPVESSDIVPTWDGIQHGERLRTMSCSD
QSMVDESLITGEAMPVAKKPGSTVIAGSINQNGSLLISATHVGADTTLSQIVKLVEEAQTSKAPIQQFADKLSGY

LDKLFVLDGGASGSKDMKLLRNHLQELRSQIISICPPMTPLCEKMISTLPSWRKINGPNQLMSLQQFVYDVC

NNGRVMSDGAKWDDDCNTCQCWNGKVACSKVWCGPRPCLIHAKGHECPAGHTCVPIKDDHCFTHP

NSRNLSCNVSHPIYWGNSRVLIQLLFDILTNSSWGDYLELDVMASSDNDMNGTLVDNQDSYRIPVLYPIN

EDGAICQCLKGFTREGKSCYDVDECARNMDRCNQHLSDCINTEGGYVCKCLEGYRGDGLHCYDIDEC

HAGFSLDPQGVYGIIRRRQEFRHRTPLDLVPKTKIDRTVSIKGQFLGEVISAVLKREGVQFLTNLPKGSAEAEL

SSSSDLVSCTKTLFKFDSNHNPEGANVVRGPVPSGAQMFSVPQYNIQYSSSTTAKDTLWLPKQNAQIEQGS
LSTELKRWDTAFDHNEARKTGVITPKAGFDPDYDKALKDIIDIEENLREYLDKQRKRLGFKAMLYWGAGKNP

KSSHGPSQEAADSGSGAKSKQLTQVADTISGDGAKPNQLGEGVKKPGLPSVPKPQSSTKLLLTTAAPAPSSS

NALQPIRDATDLRENMQNAVVSFKELCGLSPVANLMQCILAVSTRLVGPDSTPLVLLNLNDQYPTMEIQGT

AVAENSEYCCSVLSKAGNKTSKVLITVEGKDSTGQAEWVKEYDSWKSAVSEHHAMMQNWKTTWVSFLTC

RQRRSFFPRRREPLKTQRPTSSVGCLNGVSEDAVNYATLQFPPSHLQGPTAPASVPGNSRQRVKPGPADE

PGPGMGINANSQIHGQGPSQPCGTMPLGRMPSAGMQSRPFPGNMGSMTPNSPGMSLQGGPGMG

EGPTTSISLRSPEEVPSAPANPRIYVSHN

SLTMSSPAMLGNVESGGPPPSTVSQSAPVTLPGNLPSSSPYTMPPEPTLSQNPLSIMMSRMSKFAMPSSTPL
ASQDEDSQIKK
LDLKKILIHRPERMEKPGVKVKESGEPDKLEPRDTHD
DTLFPKKKDLTSCPTIIAADSLQKQGCDQCPLKPLSLSN

P

SSEKDGRSPNKREKDHIRRPMNAFMIFSKRHRALVHQRH

IKLSPEASDLITKLCCAAEERLGRSGADDIKAHPFFNSIDFSTDIRRQTAPY
DSIDAFFCTCLAGFRGPKCEEDINECASNPCKNGANCTDCVNSYTCTCPSGFSGIHCENNTPDCTESS
PQQVIQHHPSPRHHKPDPYSTGHLREAPSPLMMHSPQM
ERFTEEWKKTLLSDMEGRIKQEQPVRQIAAYCRHHDKFET
NLQSAAD
LFPSCDVIVTAEETETVVQAA

LSQEMTTYSLRNVHRPIPIQSFKQYYEAKTTNANHGFFQDFEELKEV
SFPGRRRPRGGTQGGRGRGRARLKSTTSSIETLALADIDSSPSKEDDEEEDDTMQNTVVLFS
LSKLLNQVSSNIAEKASLPCRDMQGLVTSLVYNEIEETARKESL

SLLKKAKAQLYKIEKSKSLKQ
TEKSSLEKEESCHSSDGKKRAEAKEKKNRNLKKRKVQND

PQRFVVIQNDDMMGPASPIDSTFYRALLEEEDMQDLVDAEEYLVPHQ
KGISGHRPGTHFSRDTEYGHISDDGKTISQVPDQVQLFSAPRAATAKI

SQFEKNKAYNDLNLKTNFAMKEGGIH
PDDFVADVRLIFKNCERFNEMMKVVQVYAETQEINLKADSEVAQAGKAVA
DERILGAHEDLSGLFASKSTGFSSHAIS
MKLEAYFEELLKTLYPEREFPIQPYCRSERENPEFSDD

DDVEEYVCPMELGQASGSGSKKGFKAGLDIQVYDDDDLDRLEQME
NGSSAVAGAVASSQKTLSVFSRVCEVRREDEARGQD
DSVDYRSTFLEDQLPNLSQNGSCRP
QMKVINWQTALHIPPQAKLTPEASDLIIKLCRGPEDRLGKNGTDEIKAHPFFKAIDFS
SHAMDAGVQLSHKRRNDSMAARRSKRLRSQ

EQINNRNNHRNSTSSDQSD

MDTSFSKIEGKSAVSKQHQLLLDEPR
LHKRDYSLLVPGLRNTIALDFHFNQSLLYWTDVVEDKIYRGKLSETGGVSAIEVVVQHGLATPEGLTVDWIA

FTALLYTEIYADKINLDKQQERSAFKASRN

SCNQDPKSLGKVKETQKELQLQQSSIDSTQ
YKRRQSSERGAFESARYSRTTSNPSESAEKNILVSDMEMNEQQD

EMWNVSLNNVSELKAENSEITALPNTCTVSLN

EVEGLKSVRIVGLDKTLTLSSEGFLTLTYRGNFEVKSGTSD

VCNSPEHLKNANMLNFDL
SSEEEQSQEYSLPFTMVTEQKRLDDIIRNLSQQPEELKDVYSKQFVVQLAD

QQQQQTEQEQQPSSSPRGKSRSPAVGDKEEKPVPFPSLAEGPKLSGE

M

HNNGTCKQLGSGYICMCPPGYTGLKCEKEIDECKSTPCLNNGVCKDG

MGLRLQVVLEHMPVTEEQVLPYVRRVGEAPQPAPDEPMEVQAA

KSYPRGRSITELEVMQFLSSRRNSGSAAQAFFYLREPDFLG

ESDKSTSGRKNTVISQTGSWRRGMSAQVGITTPRTKPSTTS

NLTCNSMVLFDSVGFLKKYETPLDILR
EPKQKSLSSTHSKGDGCKLQNYFNAKSNSKR

DGREPGDTVIFQCDPGYEIQGDERITCIQVENRYFWQPSPPVCI

NANYELQREENGLVD
NQHMLAHGSISGHPLVLSYADLSVWCYTCQSYVHHPTLLPAKTLAHRMKFGEDP

NQLQKERQKLGELRKKHYKLAGVAEGWDEDARSQ

LEEFTLRRCWEECLVNSEYYHRRKIIDEFNRQNRWKKRGI

DVVTVHYHLIPSPTKTKNGSKEKEREPE
FLKGTTLEEVYASVAKVADYWLDVLYSKAASMP

ADFLKARLWIEFDGEKGLDYGGVAREWFFLLSKEMF

VKDQNGNHVVQKCIECVQPQSLQFIIDAFKGQVFVLSTHPYGCRVIQRILEHC

DVVISLLAFGVYPNVCYHKEKRKILTTEGRNALI

LFENKCFLKIEHRFLTFKEANEVCETFGGTLPSISSQAEQDFLISLLPGLEENIWIGLKFFFSIRENKWV

KILRWNVLGLQGALLSHFMQPVYLSSVTLGYLYSQGHL
YFVPFIVAVSVVTLFAWIVIGF

QDQLNPLASEDDLRHIAQQLHSIGEINIM

PCSGVGVCWPSNQQPVKTKCNSDSYYQDNCANITFTFNKELMVSGLTTEHICSDLRNLNILKNVSVEYSIY

IIITKGADGSTQYISFSSPNPEIKSVQHVYQVENLLPGSFQQPEV

CKLGTHTCGENTVCTNTEGNYTCTCINGLSGTGDNCSESAMPSTTSSGDNV

MSIAPVFYV

SLPPQRLSRSDSTESSNYVKHSASMN
RYQMEIPENAISHNLPEEYELKSTRKGYKRYWTKEIEKML
SAPSAPSGDQPSSTAFIPLISTRVSLRKTRQPPERI

VPEVLKKIVTAYDMMIQTIKTLIETANSVYEKIVQCQKA

CMLSGIILL

TVIYSVVKKPHLPPKGEAKGDYENVGAQLEEDEE

GPGPGMGINANSQIHGQGPSQPCGTMPLGRMPSAGMQSRPFPGNMGSMTPNSPGMSLQGGPGMGP

EGPTTSISLRSPEEVPSAPANPRIYVSHN

SLTMSSPAMLGNVESGGPPPSTVSQSAPVTLPGNLPSSSPYTMPPEPTLSQNPLSIMMSRMSKFAMPSSTPL
ASQDEDSQIKK

RLDLKKILIHRPERMEKPGVKVKESGEPDKLEPRDTHD
DTLFPKKKDLTSCPTIIAADSLQKQGCDQCPLKPLSLSN

GP

SSEKDGRSPNKREKDHIRRPMNAFMIFSKRHRALVHQRH

QIKLSPEASDLITKLCCAAEERLGRSGADDIKAHPFFNSIDFSTDIRRQTAPY
SDSIDAFFCTCLAGFRGPKCEEDINECASNPCKNGANCTDCVNSYTCTCPSGFSGIHCENNTPDCTESS
PQQVIQHHPSPRHHKPDPYSTGHLREAPSPLMMHSPQM
ERFTEEWKKTLLSDMEGRIKQEQPVRQIAAYCRHHDKFET
NLQSAAD
TLFPSCDVIVTAEETETVVQAA

LSQEMTTYSLRNVHRPIPIQSFKQYYEAKTTNANHGFFQDFEELKEV
SFPGRRRPRGGTQGGRGRGRARLKSTTSSIETLALADIDSSPSKEDDEEEDDTMQNTVVLFS
LSKLLNQVSSNIAEKASLPCRDMQGLVTSLVYNEIEETARKESL

ASLLKKAKAQLYKIEKSKSLKQ
TEKSSLEKEESCHSSDGKKRAEAKEKKNRNLKKRKVQND

PQRFVVIQNDDMMGPASPIDSTFYRALLEEEDMQDLVDAEEYLVPHQ
AKGISGHRPGTHFSRDTEYGHISDDGKTISQVPDQVQLFSAPRAATAKI

SQFEKNKAYNDLNLKTNFAMKEGGIH
PDDFVADVRLIFKNCERFNEMMKVVQVYAETQEINLKADSEVAQAGKAVA
ERILGAHEDLSGLFASKSTGFSSHAIS
MKLEAYFEELLKTLYPEREFPIQPYCRSERENPEFSDD

DDVEEYVCPMELGQASGSGSKKGFKAGLDIQVYDDDDLDRLEQME
NGSSAVAGAVASSQKTLSVFSRVCEVRREDEARGQD
SVDYRSTFLEDQLPNLSQNGSCRP
QMKVINWQTALHIPPQAKLTPEASDLIIKLCRGPEDRLGKNGTDEIKAHPFFKAIDFS
SHAMDAGVQLSHKRRNDSMAARRSKRLRSQ

EQINNRNNHRNSTSSDQSD

MDTSFSKIEGKSAVSKQHQLLLDEPR
LHKRDYSLLVPGLRNTIALDFHFNQSLLYWTDVVEDKIYRGKLSETGGVSAIEVVVQHGLATPEGLTVDWIA[illegible]

FTALLYTEIYADKINLDKQQERSAFKASRN

SCNQDPKSLGKVKETQKELQLQQSSIDSTQ
YKRRQSSERGAFESARYSRTTSNPSESAEKNILVSDMEMNEQQD

EMWNVSLNNVSELKAENSEITALPNTCTVSLN

EVEGLKSVRIVGLDKTLTLSSEGFLTLTYRGNFEVKSGTSD

VCNSPEHLKNANMLNFDL
SSEEEQSQEYSLPFTMVTEQKRLDDIIRNLSQQPEELKDVYSKQFVVQLAD

QQQQTEQEQQPSSSPRGKSRSPAVGDKEEKPVPFPSLAEGPKLSGE

M

NNGTCKQLGSGYICMCPPGYTGLKCEKEIDECKSTPCLNNGVCKDG

GLRLQVVLEHMPVTEEQVLPYVRRVGEAPQPAPDEPMEVQAA

SYPRGRSITELEVMQFLSSRRNSGSAAQAFFYLREPDFLG

SDKSTSGRKNTVISQTGSWRRGMSAQVGITTPRTKPSTTS

LTCNSMVLFDSVGFLKKYETPLDILR
PKQKSLSSTHSKGDGCKLQNYFNAKSNSKR

DGREPGDTVIFQCDPGYEIQGDERITCIQVENRYFWQPSPPVCI

NANYELQREENGLVD
NQHMLAHGSISGHPLVLSYADLSVWCYTCQSYVHHPTLLPAKTLAHRMKFGEDP

NQLQKERQKLGELRKKHYKLAGVAEGWDEDARSQ

LEEFTLRRCWEECLVNSEYYHRRKIIDEFNRQNRWKKRGI

DVVTVHYHLIPSPTKTKNGSKEKEREPE
FLKGTTLEEVYASVAKVADYWLDVLYSKAASMP

ADFLKARLWIEFDGEKGLDYGGVAREWFFLLSKEMF

VKDQNGNHVVQKCIECVQPQSLQFIIDAFKGQVFVLSTHPYGCRVIQRILEHC

DVVISLLAFGVYPNVCYHKEKRKILTTEGRNALI

LFENKCFLKIEHRFLTFKEANEVCETFGGTLPSISSQAEQDFLISLLPGLEENIWIGLKFFFSIRENKWV

KILRWNVLGLQGALLSHFMQPVYLSSVTLGYLYSQGHL
YFVPFIVAVSVVTLFAWIVIGF

QDQLNPLASEDDLRHIAQQLHSIGEINIM

PCSGVGVCWPSNQQPVKTKCNSDSYYQDNCANITFTFNKELMVSGLTTEHICSDLRNLNILKNVSVEYSIY

IIITKGADGSTQYISFSSPNPEIKSVQHVYQVENLLPGSFQQPEV

CKLGTHTCGENTVCTNTEGNYTCTCINGLSGTGDNCSESAMPSTTSSGDNV

MSIAPVFYV

SLPPQRLSRSDSTESSNYVKHSASMN
RYQMEIPENAISHNLPEEYELKSTRKGYKRYWTKEIEKML

SAPSAPSGDQPSSTAFIPLISTRVSLRKTRQPPERI

VPEVLKKIVTAYDMMIQTIKTLIETANSVYEKIVQCQKA

CMLSGIILL

TVIYSVVKKPHLPPKGEAKGDYENVGAQLEEDEE

PPMPTVNRKAQEAAAAVMQAAANSAQSRPPYIRSPAY

LY

CFNGGTCVDGINTFTCVCPPGFTGSYCEHDINE

GNIYWIDS

IICEPSHLAHNEIHVAISAEDIGKDENPIK

Y

ᑊPMPTVNRKAQEAAAAVMQAAANSAQSRPPYIRSPAY

.Y

CFNGGTCVDGINTFTCVCPPGFTGSYCEHDINE

GNIYWIDS

IICEPSHLAHNEIHVAISAEDIGKDENPIK

**Supplementary Table S5**

| | |
|---|---|
| **A** | Top genes based on models and LongevityMAP and GenAge |
| **B** | Top genes and human variants |
| **C** | Human average VESPA score (only those with equal or great |
| **D** | Human average Tranception final score |
| **E** | Human average ESM1v ensemble score |
| **F** | Human average Progen2 final score |
| **G** | Human average RITA final score |

**A** Top genes based on models and LongevityMAP and GenAge

| PA1, Trancepti | LongevityMAF | GenAge | lAge (affect sen) |
|---|---|---|---|
| IFNAR1 (5) | | | |
| ERCC1 (4) | yes/non-sig | yes | nhibits unclear |
| DHX58 (4) | | | |
| TNIP3 (4) | | | |
| EGF (4) | yes/sig | yes | |
| PLCL2 (4) | | | |
| APOB (4) | yes/sig | | |
| FOXO3 (3) | yes/sig | yes | nhibits unclear |
| TP53BP1 (3) | | yes | nduces unclear |
| CIC (3) | yes/non-sig | | |
| elomerase/sh | yes/sig | | |
| MALSU1 (3) | | | |
| TRIM25 (3) | yes/sig | | |
| CDH11 (3) | | | |
| MAP3KB (3) | | | |
| ZPLD1 (3) | | | |
| PPP1R26 (3) | yes/sig | | |
| DHX9 (3) | | | Inhibits replicative |
| IL1RL2 (3) | | | |
| BAG6 (3) | | | |
| TSFM (3) | yes/sig | | |
| CD302 (3) | | | |
| POT1 (2) | yes/sig | | Inhibits replicative |
| BCL9 (2) | | | |
| AFF3 (2) | | | |
| SETBP1 (2) | | | |
| CFP (2) | | | |
| TRAT1 (2) | | | |
| CGN21 (2) | | | |
| HDAC6 (2) | | | |
| CFH (2) | | | |
| DAPKI (2) | | | |
| JAG1 (2) | | | |
| ITGAL (2) | | | |

IL18RAP (2)
SEMA4D (2)
ZBTB20 (2) yes/sig
POU2F2 (2)
CASC5 (2)
LPXN (2)
IFNGR2 (2)

### B Top genes and human variants

| PA1, Tranceptı | **nathan Varia** | **an variant/un** | **Variant ID** | **an variant/un** | **ain/affected** | **Gene Set** |
|---|---|---|---|---|---|---|
| IFNAR1 (5) | T68E | (**no variant found**) | | T71 | ronectin type | immune |
| ERCC1 (4) | L69S | **L69P** | s1319067405 | | --- | **DNA repair** |
| DHX58 (4) | T395S | (**no variant found**) | | | Helicase C-termi | immune |
| TNIP3 (4) | G114T | | | | disordered | immune |
| EGF (4) | R1082S | **R1082K,T** | s776085111 | | disordered | immune |
| PLCL2 (4) | S86G | (**no variant found**) | | | disordered | immune |
| APOB (4) | 2012Q; A428€ | (**no variant found**) | A4286 (**no variant f** | | **---** | **age-related** |
| FOXO3 (3) | A98G | **A98T,V** | 15798;rs752987992 | | disordered | **age-related** |
| TP53BP1 (3) | M721V | **M716V,T** | 7425;rs1173587915 | | --- | **DNA repair** |
| CIC (3) | 1258A; L2144 | **S1258G** | s77401421( | **L2144V** | --- | **cancer** |
| elomerase/sl | S429P | **S424P** | s763749808 | | --- | **age-related** |
| MALSU1 (3) | V28S | **V28I,L** | s754299384 | | --- | s-selected in |
| TRIM25 (3) | K332T | **K332R** | s767391521 | | --- | **age-related** |
| CDH11 (3) | M382V | **M382I** | s1227122026 | | Cadherin doma | **cancer** |
| MAP3KB (3) | I745L | (**no variant found**) | | | disordered | **cancer** |
| ZPLD1 (3) | T38I | **T50M,R** | s551852767 | | --- | **age-related** |
| PPP1R26 (3) | **D503G** | **D503G,E** | 1401;rs1049541401 | | disordered | **age-related** |
| DHX9 (3) | T108A | (**no variant found**) | | | disordered | immune |
| IL1RL2 (3) | G127S | **G123S,D** | 79884;rs124 | **G123S,D** | -like C2-type | immune |
| BAG6 (3) | V1055L | **V1067** | | | ER localization do | **age-related** |
| TSFM (3) | L143A | **L143I** | s770881526 | | --- | **age-related** |
| CD302 (3) | **M110V** | **M110V** | s1352505150 | | C-type lectin | immune |
| POT1 (2) | K440E | (**no variant found**) | | | --- | **age-related** |
| BCL9 (2) | M1074V | **M1065I,LR** | 32529;rs1591976450 | | --- | **cancer** |
| AFF3 (2) | K978R | K985 | s131630763 | K967Q | **---** | **cancer** |
| SETBP1 (2) | L556I | **L556P,V** | 28541; rs773986475 | | --- | **cancer** |
| CFP (2) | K44E | (**no variant found**) | | | TSP type -10 | immune |
| TRAT1 (2) | K92D | (**no variant found**) | | | --- | immune |
| CGN21 (2) | Q27D; C234D | Q27; **C234S,Y** | s1271106835 | | --- | **age-related** |
| HDAC6 (2) | N60R | | rs782651797; | | --- | **age-related** |
| CFH (2) | ,S720D,S722F | **o variants found** | | | Sushi 12 domai | immune |
| H728S, S730K,T732E,H735D,V745T | | | | | | |
| DAPKI (2) | **P1355S** | **P1355S** | s123407757 | **e impact (NC** | Death domain | immune |
| JAG1 (2) | G1100T | **G1100D** | 1543;rs1354062495 | | --- | immune |
| ITGAL (2) | M1125L | **M1125I** | s764924318 | | --- | immune |
| IL18RAP (2) | Q66R,G117E, | **5T,Q66K,G11** | 7995;COSM | **55T high impa** | -like C2-type | immune |

| | | | | | |
|---|---|---|---|---|---|
| SEMA4D (2) | G602H | **G602S** | s907595463 | Ig-like C2-type | immune |
| ZBTB20 (2) | N437D | ' (**no variant found**) | | disordered | **age-related** |
| POU2F2 (2) | G528A | **G528C; G528[illegible]98305; rs202057376** | | --- | immune |
| CASC5 (2) | I415D,N2321 | **Y**;N2321 (**no** [illegible] | 0896; COSM1372674 | interaction with BUB1 | **cancer** |
| LPXN (2) | P383T | **P385A** | s131640337378 (no varian | zinc-binding | immune |
| IFNGR2 (2) | S246G,L314Q,D343G | | | --- | immune |

**C** Human average VESPA score (only those with equal or great

erage VESPA1 down to 0.6

ERCC5
CNOT3
RAD50
APEX2
MID2
CCL5
BCL11B
KLRF1
PLEKHA1
MADCAM1
NAB2
CD22
APOA5
PPM1D
BAG6
XRCC3
LPXN
ST6GAL1
CYC1
VAV3
TRIM8
KMT2D
GZMB
DAXX
CD207
ZPLD1
IFNGR2
SYT13
NCOR2
TINAGL1
BRCA1
CASC5
CD302
SH2D4A
LEP
CPA2
HSF5

| |
|---|
| TP53BP1 |
| HAVCR2 |
| ERBB2IP |
| HLA-DRB1 |
| APOB |
| ATF6B |
| GASL |
| HOXD12 |
| CD200 |
| IFNAR1 |
| LAMP1 |
| TSFM |

**D** Human average Tranception final score

| ranception fin | ranception final score (negative) |
|---|---|
| RB1 | yes |
| BCL9 | |
| CIC | yes |
| SETD2 | |
| CDH11 | yes |
| RALGDS | |
| TP53BP1 | yes |
| MAP3K13 | yes |
| AFF3 | yes |
| KCNT2 | yes |
| FOXO3 | |
| POT1 | |
| IFNGR2 | |
| PRUNE2 | |
| CGNL1 | yes |
| RAPGEF3 | yes |
| IFNAR1 | yes |
| BAG6 | |
| ZPLD1 | |
| PPP1R26 | |
| ERCC1 | yes |
| APOB | yes |
| HDAC6 | yes |
| CFH | |
| CD302 | yes |
| DHX9 | yes |
| IL1RL2 | yes |
| ATF6 | yes |
| DAPK1 | yes |
| NFKBIZ | yes |
| TRAT1 | yes |

| | |
|---|---|
| DHX58 | yes |
| JAG1 | yes |
| TNIP3 | yes |
| ITGAL | yes |
| IL18RAP | yes |
| EGF | yes |
| CYP27B1 | |
| PLCL2 | yes |
| SEMA4D | yes |

### E Human average ESM1v ensemble score

| e ESM1v ense | ESM1v score | ge ESM1v ensemble (negative) |
|---|---|---|
| ARID1B | 1.035 | yes (-0.76) |
| ROS1 | 1.51 | yes (-0.1) |
| CBLB | 2.104 | yes |
| BCL9 | | |
| CIC | 2.104 | yes (-2.84) |
| THRAP3 | 1.034 | yes |
| BRCA1 | 1.355 | yes |
| TRIM24 | 1.19 | yes |
| CDH11 | 2.86 | yes (-2.68) |
| TP53BP1 | | yes |
| LATS1 | | yes |
| MAP3K13 | 1.42 | yes(-1.10) |
| AFF3 | 1.001 | yes |
| FOXO3 | | |
| POT1 | | |
| ACD | | |
| LMX1B | | yes |
| MRC2 | | yes |
| MALSU1 | | yes |
| CGNL1 | 1.216 | yes (-1.46) |
| TRIM25 | | |
| F7 | | yes |
| TSFM | 1.36 | yes (-0.88) |
| NFKB1 | | yes |
| ZBTB20 | | yes |
| RAPGEF3 | | yes |
| IFNAR1 | 0.844 | yes (-3.93) |
| BAG6 | | yes |
| TOP2A | | yes |
| ERCC1 | | yes |
| APOB | | yes |
| HDAC6 | | yes |
| NEDD4 | 1.33 | yes (-1.32) |
| CFH | 0.74 | yes(-2.51) |

| | | |
|---|---|---|
| DHX33 | | |
| PUM2 | | yes |
| CD302 | 1.309 | yes (-0.96) |
| DHX9 | | yes |
| IL1RL2 | | yes |
| ADAR | | yes |
| DAPK1 | | yes |
| DHX58 | 1.124 | yes |
| MAP4K2 | 1.1 | yes (-1.17) |
| JAG1 | | yes |
| TNIP3 | | yes |
| ITGAL | | yes |
| POU2F2 | | yes |
| IL18RAP | 1.4 | yes (-0.98) |
| EGF | | yes (-1.51) |
| PLCL2 | | yes |
| SPPL2A | | yes |
| PRKCG | 2.87 | yes (-2.85) |
| SEMA4D | 1.64 | yes (-2.46) |

### F Human average Progen2 final score

| Progen2 final: | Progen2 final score (negative) |
|---|---|
| KIAA1549 | |
| CASC5 | yes |
| CIC | yes |
| NSD1 | |
| TET1 | |
| SETBP1 | yes |
| CDH11 | yes |
| MAP3K13 | yes |
| SSPN | yes |
| ACD | |
| MALSU1 | yes |
| TRIM25 | yes |
| ZBTB20 | yes |
| IFNAR1 | yes |
| ZPLD1 | |
| PPP1R26 | |
| ERCC1 | yes |
| DHX33 | |
| DHX9 | |
| CD38 | |
| DHX58 | yes |
| TNIP3 | yes |
| TMEM173 | |
| LPXN | yes |

| | |
|---|---|
| THEMIS | yes |
| CFP | |
| POU2F2 | yes |
| EGF | yes |
| PLCL2 | |

**G** Human average RITA final score

| ge RITA final s | erage RITA score (negative) |
|---|---|
| SETBP1 | yes |
| FOXO3 | |
| ACD | yes |
| MALSU1 | yes |
| TRIM25 | |
| TSFM | |
| IFNAR1 | yes |
| ERCC1 | yes |
| APOB | yes |
| IL1RL2 | yes |
| APOBEC2 | yes |
| TRAT1 | yes |
| DHX58 | yes |
| TNIP3 | |
| THEMIS | |
| CFP | yes |
| EGF | yes |
| PLCL2 | yes |

er than 0.6)

LG

er than 0.6)

**Supplementary Table S6**

**A** Tortoise-specific gene expansion (from Quesada et al. Nat Ec

**B** Jonathan and Tank gene expansion

### A Tortoise-specific gene expansion (from Quesada et al. Nat Ec

| Tortoise-spec | Gene | Expansions | | | | |
|---|---|---|---|---|---|---|
| **Family** | **Human** | C. abingdonii | G. agassizii | P. Sinensis | Tank | Jonathan |
| OTX2 | 1 | 2 | 2 | 1 | 2 | 2 |
| ATP6V1A | 1 | 2 | 2 | 1 | 2 | 2 |
| LAMTOR4 | 1 | 2 | 2 | 1 | 2 | 2 |
| STXBP1 \ STX | 2 | 4 | 6 | 2 | 6 | 6 |
| TPT1 | 1 | 2 | 1 | 1 | 1 | 1 |
| VCP | 1 | 2 | 2 | 1 | 2 | 2 |
| RPL11 | 1 | 2 | 2 | 1 | 1 | 1 |
| EEF2 | 1 | 2 | 2 | 1 | 2 | 2 |
| GJD2 | 1 | 2 | 2 | 1 | 2 | 2 |
| RPS25 | 1 | 2 | 1 | 1 | 2 | 2 |
| POLR2L | 1 | 2 | 2 | 1 | 2 | 2 |
| EEF1A1 \ EEF | 2 | 6 | 5 | 3 | 5 | 5 |

### B Jonathan and Tank gene expansion

| Family | Human | Tank | Jon | List |
|---|---|---|---|---|
| USP12 | 1 | 2 | 2 | Cancer |
| CHIA | 1 | 4 | 3 | Immunity |
| GZMB | 1 | 4 | 4 | Immunity |
| GZMH | 1 | 4 | 4 | Immunity |
| CTSG | 1 | 9 | 8 | Immunity |
| **PRF1** | 1 | 11 | 10 | Immunity |
| **TLR2** | 1 | 4 | 4 | Immunity |
| **TLR5** | 1 | 2 | 2 | Immunity |
| **TLR8** | 1 | 2 | 2 | Immunity |
| **TLR13 - mou** | 0 | 3 | 3 | Immunity |
| NEIL1 | 1 | 2 | 2 | DNA repair |
| MYCN | 1 | 2 | 2 | Cancer |
| P2RY8 | 1 | 2 | 2 | Cancer |
| NF2 | 1 | 2 | 2 | Cancer |
| PML | 1 | 2 | 2 | Cancer |
| SMAD4 | 1 | 2 | 2 | Development |
| EEF1A1 /EEF | 1 | 2 | 2 | Proteostasis |
| U2AF1 | 1 | 2 | 2 | Cancer |
| RPS25 | 1 | 2 | 2 | Ribsomal |
| STXBP1 | 1 | 2 | 2 | Synaptic vesicle |
| STXBP2 | 1 | 4 | 4 | Synaptic vesicle |

From Quesada et al. Nat Ecol Evol, 2019 (

:ol Evol, 2019)

:ol Evol, 2019)

or from GeneCards

**Supplementary Table S7**

**A** List genes affected by hypermethylation DMRs in Jonathan
**B** DAVID gene name of  list of affected genes from tab "A"
**C** List of genes whose promoter or coding region contains the h
**D** DAVID gene names of affected genes from tab "C"
**E** PRC2 targets identified in "A" tab
**F** GO analysis of genes in tab "A"

**A** List genes affected by hypermethylation DMRs in Jonathan

CFAP54
HIBCH
LRRC28
MED21
ZCCHC7
JAZF1
F9
PINX1
CEP44
ST6GALNAC1
SNRNP70
PRSS16
SPATA16
PLXNC1
TJP1
WRN
CAPRIN2
ZNF410
LRCH3
SEMA3G
PCDHGC5
SYN2
FOXK2
POLE3
CTNNBL1
HDAC7
SKIL
SLC16A13
SNX2
CYP26C1
ALDH8A1
HEATR5B
PCSK6
MAP3K5
PTPRM

FGFRL1
PDE4B
IGFBP2
LUZP1
STARD10
MUM1
ARIH1
CHRND
PDE9A
NPHP1
IPO11
VAV3
CLASP2
FOXC2
GRIA3
INPP4A
ABCA8
COL4A1
MBNL1
UBE2I
FGFR1OP
SERPINC1
INTS4
IPO5
LPPR1
PPP3R1
TBC1D22A
SH3RF3
MTM1
OSTN
COL18A1
THOC3
VIT
TNIK
MSLN
ATP6V1H
SH3GLB2
FLT3
IMPDH1
TLX1
EXD3
MED24
PROM2
ITPR2
IDO1

C12orf4
RHOT2
DCBLD2
DAZL
SCN5A
GABPA
TSPAN18
OSBPL6
RBM6
KIAA1549
JPH3
BCL6
MRPL48
MED13
SNX18
BRIP1
SLC15A4
PCCB
ST8SIA1
WDR17
GK5
FTH1
HEATR9
ZFAND3
NDE1
SRF
TRIM47
ZCCHC2
LDLRAD3
TBX5
INTS12
TMEM150C
ALKBH8
SLC9A8
ARAP2
SYNJ1
TCERG1
RPF2
FGD4
ZNF740
ETFA
TBL1XR1
GYG1
KIF6
BANK1

PTPN4
CHMP2B
AMER1
TMEM59L
DRD3
DAW1
KMO
PAPOLA
ARHGEF17
PDK1
SLC25A27
ZNF516
TCP11
GIMAP8
MTX2
SOX17
CAPN5
BACH2
SEMA3B
NPDC1
ZNF462
WIPF3
OPTC
FAM129B
EAPP
ZNF618
TDRD5
PKD2L2
CAV1
RNF125
PSIP1
ST5
CHD8
GSPT1
DSCAML1
CDK19
DDX59
GPATCH1
CSE1L
CTSB
CRISP2
LCK
KAT6B
ATG7
SRGAP3

TAMM41
CTNNAL1
CYB5R3
PPP2R2C
ENOX2
UTP18
DYNC2H1
RCOR1
MIOX
EXT1
SPATA6L
ACBD5
HCN4
CLIP4
HHATL
MED9
ZNHIT6
FRMD4A
CASR
C17orf53
PLB1
SLC35F1
ISG20L2
AKAP2
OR5D18
ADSS
PIGQ
GCAT
KIAA1377
CCDC60
GPR61
TRIO
AOC2
RSU1
COPB2
SNTB1
DDAH1
PHF10
ZNF646
FADS2
SERPINB6
SUPT6H
AGRN
ECHDC1
PLA2G4E

VAX2
FAM19A4
VCAN
TM9SF2
USP20
SPIDR
L2HGDH
RAB3GAP1
ERCC6L2
PLEKHA1
RALGDS
LUC7L3
ORC3
KCMF1
SSH2
HSPBAP1
CD81
ADIPOR1
NDUFAF2
KATNAL2
LOC400499
GUK1
ITGA8
NMT1
NARS2
ZNF704
ZC3H14
PDSS1
CADM1
NCOA3
ING4
RASAL1
INTS2
TTC6
MCF2
TIMMDC1
WDR37
DARS
SPECC1L
SORL1
STAG1
CPNE7
ADK
TENM3
VIPR1

UBE2H
GRAMD1C
WDR59
GAB2
RDH10
TMEM9B
GBE1
TG
SPOCD1
DCTN1
TOX2
COMT
MTCL1
CARM1
SLC9A4
CASKIN1
TBATA
PHF20
NEURL1
PCSK7
OLR1
ARHGEF2
THSD7A
GNG2
SQRDL
MPZ
TGM1
NCKAP1L
TBC1D5
AKT1
KIAA0226
TRIM29
SRFBP1
OCSTAMP
RANBP3
NCOA7
RASGEF1B
RANBP10
MVD
GMDS
NUDT9
MATK
CHPT1
WDR48
CACNG4

LCOR
CDC14A
STAU2
DCTN6
ANG
WDR4
PDLIM4
TIMM50
LHX8
HYDIN
C17orf104
ENDOD1
TBC1D20
ERP27
TNXB
DIAPH1
ZNF512B
ORC5
LCP1
SPON1
ADRB3
ADAMTS18
TRDMT1
DCST1
PM20D2
PPP6R3
ASCC1
GEMIN2
GPCPD1
BTNL2
RPS11
STIM2
IGSF9B
PLCL2
RCBTB1
PPA2
ZAK
TRIQK
STAT6
PTPRU
BSG
PLCD3
PPP1R9B
TSPAN9
KCNT2

PRKDC
SEZ6
TACR1
PKP2
KAL1
NDUFV2
COL6A1
LIMCH1
KIF3C
TNNT2
AFF2
USP46
RP2
CLTA
CUBN
ST3GAL3
UBTF
CHTF18
FST
TMEM63B
SPON2
DRC1
SEMA6B
ROS1
MEX3D
THBS3
B3GALTL
SAMD15
SNAP91
ZDHHC21
FAF2
CRMP1
APLF
RAD50
IRS2
IRAK2
KMT2C
SNRPA1
TAF5L
TACC3
SUCLG2
SHC2
WSCD2
DAAM1
KIAA0895

ARHGAP19
GRWD1
MBNL2
CNTN2
DNAH9
ADAMTS8
KCNH1
DCAKD
NPY2R
COL5A1
SPOCK3
KCNU1
PPP2R3A
MRPS36
COMMD1
MYBPC2
C20orf27
SNAPC5
PITRM1
RNF126
NUDT3
DCTN2
ARHGAP21
FAM171B
ARL2BP
RAB37
KLHL36
P3H3
ABCG1
HR
ABCC1
CLSPN
MBD5
ZNF438
PRKCQ
EHMT1
TPM3
VPS53
RGS9
ENTPD8
ITIH2
CHCHD3
ALPK3
TRIM8
ZNF407

KAT2B
FAM179A
PFKL
PTDSS2
TNFRSF21
WDR43
NBEAL1
SYT14
DCST2
BCAS3
VRK2
SLC35F6
MUTYH
TRAPPC11
SPATA7
TNS4
POR
ADAMTS9
PCGF5
PDLIM7
RASA3
CHD9
AIRE
CRNKL1
TRIM9
SIDT2
TNRC6B
SLC12A4
UCKL1
SEMA5A
ACVR2B
SLC30A9
PDE3A
DGAT2
CA4
AARS
CSNK1D
TMEM44
TMED8
RANBP1
FLII
SLIT3
SND1
ZNRF2
BAG3

OXR1
TP53BP1
FBXO31
C16orf96
PAX8
BRD8
COL4A6
TMCC3
MBD3
VTI1A
CENPC
EPSTI1
MED13L
TSTD1
TRIM44
NDUFAF6
NPSR1
GBA
IGFN1
NDUFA2
CC2D1A
TLCD2
ZFAT
WNK1
MAP4K4
LPP
VWA3A
CERS1
SLC25A11
ENO4
DLG5
KDSR
BRWD1
CASP3
CFAP44
MXD4
PRKD1
RBFOX2
ENSA
KIAA0430
PLEKHA7
AAMDC
JAG2
TSPAN10
FOXO1

FCRL3
ZNF316
TRIM65
SMCHD1
STRN
IQCB1
HARS
SYNE2
SEC31A
PGAP2
SDC3
DUSP8
GPRC6A
HEPACAM
OSTF1
MCMBP
MROH1
FAF1
SUFU
THG1L
GRM4
LIG3
ABI1
PPP1R16B
TBP
BAZ2B
CUEDC2
SMARCA2
CSNK2B
GLG1
DEGS2
RP1L1
LRRC6
EIF2AK4
IARS
ANKRD28
SRRM4
IGSF21
FKBP11
LZTS3
TM9SF4
CA10
GABBR2
U2AF2
CNTFR

MORC3
GCC2
TLK1
ERBB2IP
ISL2
ARL3
DGKQ
EXOC3
CABIN1
PRDX3
BNIP3
NRDE2
RAB39A
GNAI2
PLEKHG2
STK17B
EPHA5
EXOC1
PJA2
NCKAP5
HCN2
CDA
HNRNPM
CBL
CPM
IQGAP3
CACNG1
ADD2
HS2ST1
PUS7L
ANKRD11
NAALAD2
B4GALNT2
KLHDC4
VGLL3
LRWD1
MGRN1
GIT1
KANSL1
UQCRC1
PFKP
CRYM
IRF4
ATP11C
KIAA2012

ITGA11
SRSF4
BRE
TRIM66
DYM
ARMC8
NDRG4
STON2
WDR33
NAALADL2
PIGN
DPYS
CYB5D1
MICALCL
EPN2
TM7SF3
C9orf41
CABLES1
TUBGCP2
LOC100996634
MYO10
SLC39A8
ADIPOR2
RIMBP2
HDAC5
FARS2
PLEKHO2
ALG6
FAM129A
VSTM2B
KCNQ4
ADGRA3
TBC1D2
GCNT1
SORCS3
KATNB1
TGFBRAP1
CAPS2
CARS
PKDCC
PAPSS1
PLD4
AFG3L2
PRDX6
GAN

MYZAP
PPP5C
SETD4
AKAP13
HTT
KIF2C
BBS7
FXR2
TESK2
AGAP2
HERC2
PIAS2
SESN2
UBXN1
FAM69B
LATS2
SHB
GPT2
CBLB
PTPRZ1
COMMD10
TIAL1
CYP27A1
PARL
MEX3C
PTAR1
GRIK1
WNT11
TRPM5
PSD3
GNAT2
PDK3
UHRF2
LEFTY2
RELB
TNIP3
ALPK2
HIBADH
CENPT
COL16A1
THOC2
FBXO18
DAPK1
SLC17A7
MED17

RSF1
WAC
SCO2
USP2
TMC3
INF2
STAB2
IK
PRPSAP2
OR6P1
USP33
ROBO4
MAP3K10
MICU1
PPP2R2D
LHPP
PPARGC1A
FAT1
RAB38
SNCAIP
EFCAB6
F7
TRIP11
ADGRB2
TRPM4
MTA3
DLGAP4
MAP2K1
DUOXA1
SNX29
ATP1A3
CLPTM1L
ASPSCR1
PDZD11
SELP
LCP2
MITF
PTPN1
PLEKHH1
EXPH5
IER3
GDAP2
LRP8
CPS1
ANXA2

GDI2
ARMC9
CRNN
RABEP1
RB1CC1
COPA
PDS5B
CLEC18B
STYXL1
SERPINA10
TIE1
RNASEH2B
TIGD1
MDFIC
FNIP1
CPNE4
TRAPPC10
THSD7B
SOCS7
RASL10B
MIPOL1
APBB1IP
PPP2R2B
SNX11
ZNF436
PLD1
ZFYVE21
MYH11
NPC1
AMBRA1
PPP1R13B
SLC25A25
FOCAD
RET
CARS2
KCNK17
KIF13A
TMEM67
FA2H
MYO6
NAT14
UBE2F
GUCY2D
WNT5A
PIWIL4

RBBP6
TRAM1
ARHGAP33
KLHL12
MAST2
SLC22A18
IQSEC3
DNAH1
MXRA7
CA2
C8orf48
PADI3
SLC6A11
F11R
RPS8
EAF1
DNM1L
IGF2R
PPP2R5C
STK36
POLR3F
TSPAN31
STAB1
CCDC178
CACYBP
C10orf2
SLAIN2
TMEM79
KTN1
STX12
GFOD2
PDZK1IP1
EIF4A3
KIN
PRKCB
LYRM4
TRPM6
FBXO42
NUS1
APLP2
CACNA1I
TXNDC11
MBOAT2
CDH12
DDX39A

ZBTB20
ASB13
PRRC2C
RHOJ
ITGAV
ATE1
NNMT
NLK
11-Mar
PSMA1
EHBP1L1
FAM53C
PDZD2
C8orf88
PPFIBP1
PLS3
SKOR2
KIF5B
ACSS2
MBOAT7
CDH20
TOX
RIMS3
DDX24
EML5
TMEFF1
ESRRB
SH3TC1
SDE2
ZCCHC14
SLC9A5
MAB21L2
SUSD6
TMTC4
SLC22A5
AGPAT6
ABHD5
FGD3
PHACTR3
SLC5A9
HYOU1
ZNF22
SRL
GINS2
SIM2

DUS2
CD300LF
CRYBG3
PIK3R3
FAM13B
NEDD4
TSNAXIP1
EIF3A
DENND5B
MOGAT2
UBASH3B
ALCAM
ARHGAP17
NOXA1
SSU72
IDE
CDH6
STAT1
FASLG
KIAA1524
CSNK1G1
TP63
WDR26
SEC23B
EPB41L2
C1orf168
CLSTN1
RHOT1
POF1B
PCDH11X
MUC3A
PSMF1
FAM76B
TTC23
ARHGEF10L
TMEM63C
C11orf97
GNB1
MOB3C
MCM8
RGS3
COG6
DDX26B
PACS1
TTC21A

IFNGR2
CEP164
IKBKB
RAB15
MYO15A
PPIP5K1
RPAP1
C11orf16
SLC16A7
TOE1
CALB2
GABBR1
RPN2
TCF21
SORCS2
SLC25A1
RARS
DBNL
RAPGEF4
PTPRE
NFAM1
ATP2C2
CRYGS
NOL4L
UBE2V2
MTMR4
RHPN1
ACTR2
ABLIM2
FAM83C
CPA2
SLC15A1
GPNMB
MXD1
FZD8
LOC102725191
ATP5SL
MCHR2
GART
TNS2
AFF1
LAMA4
ALPK1
GSTO1
CNTLN

CDC123
PLCB4
INADL
CDC73
CNOT7
FAT3
PECAM1
GMNC
TBX15
NME8
EFCAB3
OXSR1
FAM92B
PALM
FNBP4
CNTNAP1
DCUN1D4
CRADD
PAPSS2
FAM183A
ATP2C1
BMPER
OBFC1
PPP1R37
NSMCE2
KIFAP3
DIAPH2
EPHA7
MROH7
SERAC1
PUM1
PTCH1
SNX6
GPATCH2
CCM2L
AAK1
MLKL
DEPDC1B
CPEB2
ARHGAP12
TET3
SPPL2B
PALLD
CDCA4
EXOSC10

GYS2
VWA7
CLASRP
DPF3
RPL38
CERS3
PSMD1
NT5C3B
MAP7
PDE3B
ATP6V1B1
GAREML
CD274
NSUN3
SLK
ABCC3
ASL
BTBD19
PSMD12
LIMK2
RAB8A
AGK
FZD2
KCTD19
KANSL1L
EIF3H
DMD
KIAA1522
GNPTAB
HSD11B2
RCAN2
SLC22A23
ADGRA1
CLGN
RND2
ASTN1
RBPJL
GAS2L2
DNMT3A
FBXW8
NEB
POPDC2
SUPT3H
RNF145
DOCK7

EEF2KMT
TSPEAR
SGOL1
SLC16A10
TNPO3
DTNBP1
VAV2
PTPN13
TRA2A
RBM19
TMEM38B
PHF21B
TECTA
BLVRB
DAPK2
OXCT1
EEPD1
USF1
PLEKHA8
CCDC150
SRXN1
DNAJC9
IRX6
SNAP47
CASK
SLC30A6
EEA1
CTSD
LIFR
CTH
MTA1
PGM2L1
MFSD7
ZNF70
RPGRIP1L
GRM5
USP10
MAP4
REM1
SMARCC1
ADGRG2
USP25
TPPP
OTOG
PPIL1

KIAA0232
TBCK
SMURF2
UBE2D1
SAG
RALGAPA1
NIPBL
CCDC103
CD300LB
ADGRE1
ENTPD6
TNFRSF11A
ZFYVE1
GALNT1
GABRA5
SAP130
MAP6
SYVN1
CNOT2
MARK4
BMPR2
SKA3
ARHGAP30
LOC100130705
LRP12
SEZ6L
STT3B
SLC9A3R2
RNF121
COL27A1
GFPT2
CD80
GALNS
SNX25
TMEM86A
NPAS3
OTUD5
ST18
BCAR1
OCA2
POLR1A
CMTM7
AGAP1
RELN
CASQ1

HYKK
DYNC1H1
EGF
CCDC38
LPPR5
SMARCA1
PCNT
KNDC1
ECE1
NDC80
UNC45B
CEP112
CCDC73
NT5DC3
BCAT1
TTC39A
SLC48A1
PIEZO1
ARPC1B
SNTB2
FEZ1
ERICH1
MCU
TES
MRPL27
RPF1
RAB20
PLEKHM2
FBXW10
DENND1A
TRPV3
PRPF4B
SLIT2
CYBA
MTMR1
CCDC169
MAST4
ACER3
OLFML2B
SPSB3
KLHL4
TYMS
PALD1
TOP1
MMACHC

CCDC93
USP34
6-Mar
CUL5
RFTN1
DUX4
COLEC12
ZNFX1
PRKCZ
PGM5
DPYSL5
UVRAG
GIPC3
NSD1
GRK6
LRRC20
ETV6
ZNF451
NPNT
CRHR1
SKAP1
SLFN14
TBC1D16
NUDT8
COPE
MAN1A1
NOTCH1
MON2
ACSF2
USP43
MED30
GPC4
SERPING1
ST8SIA5
SCUBE3
SF3B3
FRMD3
ANKRD16
STARD8
OCRL
PHF3
DNMT3B
AFAP1
TTLL1
POLR1E

DAP
RRN3
TSPAN15
FBXW11
SYPL1
GRHPR
FRAS1
VGLL4
MICU2
ATP6V0A4
ECI2
APBB2
RYK
TBC1D30
TSC22D2
RAB11FIP5
INTS3
MEGF9
C14orf80
NHLRC3
B2M
KCNH2
RASAL2
STXBP4
ATP6V1G1
ADAM32
IKZF1
C14orf2
ARID3A
SLC38A9
FBXL6
NOVA1
APBB3
C19orf54
STX8
ADAMTSL3
TRHDE
FAM135B
MFSD8
DHX37
ARNTL
TDRD9
SLC4A4
LRIG1
RPGR

CHURC1
OTOF
FAM155B
ACSS1
CLDN16
PHKB
SCARA5
YTHDF2
XPNPEP1
DHRSX
KIAA1598
BRINP2
HFE2
PLCD1
ASB7
NHSL2
LAMA3
MYH10
MID2
CRB2
KCNS2
MYO16
FAM205A
HDAC4
AK8
DNER
SPAG9
AIG1
WDR25
SLC26A7
SCAI
TRMT11
IMPG1
SPTLC1
BBS9
ITM2C
INTS6
FMO5
HHAT
ALS2CL
COG2
ZDHHC14
LRP5
MAML3
UXS1

SHBG
DROSHA
CDK5RAP2
LAPTM5
MPP2
LMNA
NAA50
PLD3
MMP28
VMP1
PI4KA
PPP3CA
TRPM7
UBQLN4
LRP2
GPHN
RHBDF2
FREM2
TMEM240
FOXO3
DHRS7C
HNRNPH1
LRRC69
LRRC4
TM6SF1
UAP1
XAF1
ALOX12B
PDXDC1
NLRP12
P3H1
LMNTD1
SFXN5
FRMPD1
RFC1
OPHN1
LRFN2
RALGAPB
CDYL
SEC23A
TUBGCP3
OVCH1
BRD1
MRPL21
MURC

TNPO1
SIX2
CHD6
CYB5A
SLC25A17
MBTPS1
AIFM2
KCP
LLGL1
SNX30
ATRX
NFATC3
CRHR2
ARHGEF4
LOC200726
PNPLA6
C16orf62
YES1
TEKT3
RRP12
STK33
ZNF641
ALDH9A1
CFAP99
CHRNA1
VWA1
MAP4K3
FAM126A
ARCN1
PBX3
C2orf40
RABAC1
BIN1
TBC1D1
WARS2
POLR1B
TMEM41B
GALNT2
TTC1
TRMT61A
POU2F3
LPAR1
RMDN2
NECAB2
LONRF2

C2CD3
INPP5E
GPN1
IFT122
PREX2
PARP4
PLA2G12A
CDKL3
CNTN6
UBA3
ADAMTS14
PYCARD
SIDT1
ALDH4A1
NTAN1
RAF1
CORO1C
WWOX
KIAA0586
DHRS7B
ARHGAP44
WNT3
HTR2A
PDE7B
RNF4
ANKRD31
ARAP3
ATP10B
KLRF1
C14orf37
PASK
SLC1A6
FBXL17
SLC38A3
BDH1
RNF165
FAM98B
IQCE
VSIG1
PTPN3
AMOTL1
PLD6
DHRS3
EPC1
MMD2

TOP1MT
U2SURP
METTL24
OASL
MCM5
ANKRD17
TSG101
OSBPL1A
TLE1
ACAP2
YWHAZ
IFI44L
ACSF3
PPIL4
UHMK1
ADAMTSL1
GRHL1
SLC2A2
STPG2
TLL2
RAB5A
UGT8
MAP2K6
E2F6
MRPL15
CCNI
RP1
FSIP1
NCBP1
RIF1
ADH4
TEX264
GRM1
WDR49
CCDC34
IRX2
CNDP1
CMSS1
DYTN
FABP3
RPS6KA2
FAM160B1
ERI1
CEP192
ZFAND6

KIAA0408
FRMD1
SCARF2
PIK3CB
MRPS35
TEX9
RIC3
LRRC7
RBKS
CENPO
ITFG1
ATP1A2
PDLIM1
NAV2
EXT2
COL9A2
ATP2B2
CAPN7
TAF1A
FAM166A
AP1M1
FAM175A
GALT
KDM6A
GRIP2
VPS11
GRB2
TMEM47
SLC2A6
UBE2G2
FOXP4
ANK2
PDE6B
AVL9
POU6F2
NDUFS4
TAB1
MMP24
NEDD4L
TRPM2
CEP170B
PRUNE2
DHX35
STXBP5
GAD2

FAM20C
TALDO1
MICU3
ATF6
PYROXD2
ARHGAP15
RBM45
FAH
MASP2
SAMD11
SYPL2
CRBN
RALGPS2
GRIA4
IL1RL1
LRRC16A
PAXBP1
NR2C1
AKAP11
ITGB2
ZC3H6
GATAD2B
TRPA1
NAGK
MAP3K8
DKK3
ATG4C
MIB1
RYR1
C3
SLC39A6
NARFL
NDRG1
CAPNS1
EDEM2
CNNM1
INPP5A
NUP133
SPTLC2
LSS
TM2D3
RAB10
CCDC15
PCGF6
LURAP1L

FKBP10
RNLS
C2CD5
PAX2
HS3ST3B1
POC1B
IPO13
PRKRIR
B3GNTL1
KCNAB2
ABHD6
KIAA0368
RUNDC3B
RGL3
MAML2
TOPAZ1
KIF9
VWA3B
MIA3
C9orf91
USP42
FMNL2
SLC9C1
CYP39A1
MKX
FNDC3B
IGFBPL1
ISPD
MATN4
ZNRF1
NCAM1
PDK2
KRT222
PTPRO
CAST
DDX17
KPNA5
DYNC1LI1
FAM65A
ADGRG1
IL17RA
ITSN1
HOOK3
PPP6R2
SAP30BP

KIAA1217
TRIM37
TBC1D31
IGHMBP2
USH1C
LPCAT2
HMBOX1
EPHA8
PDPR
NRBP1
FAM178A
COL11A1
INPPL1
INSC
SFMBT1
FHIT
SPSB4
CFAP53
PDE1C
MTMR8
GLTSCR1
SH2D4B
SNAPC2
ARHGEF12
MKKS
NPTX1
SCRT2
ISCA2
CAMKMT
PRKD2
MYO5B
SHCBP1
TCTE3
CRIM1
RNF212B
UFSP2
PAX1
HSF1
PIGU
CXCL11
ENOX1
TRAK2
FAM161B
DOT1L
CXorf36

MYBPC1
USP22
REC114
C20orf196
PDE2A
ZDHHC17
CLU
NDUFA6
ATG4B
ENPP6
ARHGEF38
ELP2
TMEM71
VPS13B
AMFR
HK1
CYP4V2
ABTB2
AZIN2
ABCB11
NHSL1
FAT4
LRRK1
ERO1LB
SAMHD1
WDFY3
VAT1
TRIM28
SLC25A32
ADAP1
ALK
USP5
RXRA
SIRPA
DMPK
LGMN
TRMT44
ANKS6
FTO
FAM172A
ZSWIM8
LOH12CR1
KCNH4
CASP1
SPO11

STAT2
ASCC3
MAX
MAPK11
CLCN3
RADIL
LETM1
HS3ST2
ERG
DCAF13
NUDT14
CHL1
SLC35G1
HAT1
HIP1
NAV3
ALKBH3
NFE2L2
GLRX3
CDC16
ELP4
CUL1
FRYL
PRSS55
SPEN
SPHK2
SRPX2
TIPIN
RNF167
ZC3H7B
PDCL2
TTLL10
UTP6
TARSL2
FNDC7
RRAS2
PPP2R4
NADK2
MEST
FGFR2
SP5
OXNAD1
PDE4DIP
COL12A1
PQLC2

DOCK3
FCHO2
ACO1
DGKD
NDOR1
EIF5B
HIC1
USP9X
REV1
GGT7
ACTR5
STK26
MED25
ADCY2
CD226
ARHGEF9
KCNC3
EIF2B1
SLC8A2
FBXL18
THOC7
ADCY1
ACADM
TEAD1
CMTM6
MYLK2
KIAA1429
PARD6G
ALX4
ALG1
ANKRD33B
CBFA2T3
RAP1GAP2
NR2C2
MOCOS
PXDN
IGDCC3
WFDC8
DERA
JAKMIP1
CLN5
BICD2
SNTA1
IGFBP5
PCOLCE2

EDN1
ZBBX
TMEM184B
MAPKAP1
TRAF1
STX18
COL4A2
MTMR2
COL24A1
SPAG17
FOPNL
RIMS2
HK2
TCEA3
FTCD
AGO2
SLC4A11
NCAPD3
RYR3
MAP4K2
FRMD4B
XYLT2
GGCT
IPO7
LOC101928991
GFI1B
RAPGEF6
ASB18
MSANTD1
MED23
FAM63B
TOPORS
NCAPD2
LRP6
WDR63
DOC2B
VMAC
TMEM245
METRN
ABCA2
PPM1H
MRPL38
UBE2L3
C11orf30
DNAJB12

SYT13
SPTBN5
CIZ1
LSP1
STIL
RHOBTB1
XPO4
ABLIM1
GPC1
DUSP22
SERGEF
GKN1
C11orf80
TRMT1
SIGLEC15
ZFPM1
ANO6
RTTN
CACNG8
HELQ
GRIP1
IDO2
POLR2B
HECTD2
KIAA1211L
DNAJC13
GFAP
ESRP1
PLEKHM1
LOXHD1
CAMTA1
MYCBP2
TMEM8A
C1orf53
ZFYVE28
ADRBK2
GORASP1
SLC24A3
LYPD1
BANP
CC2D2A
FAM64A
LRRC75B
EPM2A
MYRF

MDN1
DNAH2
ANTXR2
NUP210L
JPH1
C6orf229
CROT
AGPAT2
LRRC49
TFDP2
DBX1
TAF1B
PIK3CG
MEDAG
KDM1A
KHDRBS3
CCDC25
TIMM17A
CCDC47
PLLP
ESAM
FAM53B
NUBPL
NCAN
GPT
DTD1
ST6GAL1
MMP16
PTBP3
UMODL1
AP2B1
SLC25A15
GYS1
DSCAM
ARFGAP3
TRAPPC12
EXOC5
FDFT1
CDH23
SBNO1
TRIM54
NTRK3
DYSF
TIAM2
TCF3

PCSK5
PDGFD
SLC37A2
TDRD1
CCDC12
POLE4
TTC12
NSF
AVEN
ADGB
YTHDF3
NOXRED1
GPR137C
REV3L
CATSPERG
CCDC152
MAPK10
SNPH
DOCK4
DRC7
FAM120A
HSD17B12
RCN1
SRBD1
TSKU
CECR5
DHX9
RAD51B
NCR3LG1
MTMR3
WDR7
CROCC
GABRG1
ADAM19
ADAMTS15
EIF2S2
SLC6A6
ENOPH1
XRCC4
TMEM170A
PKP3
CCSER1
C20orf194
LVRN
TET1

MCM3AP
ASAP1
GRIN1
HRAS
WFS1
CACNB2
PDE6D
TCF12
HERPUD2
HPSE2
ULK2
CCR10
RBL2
HSF2BP
CDH1
NWD2
SORCS1
SPATC1
IQGAP1
BCL7A
SEC16B
TRPS1
SPHK1
CDKL1
OSBPL10
GPBP1
SAPCD2
SCIN
ERBB4
SUCO
LRCH1
SLC30A3
MGMT
CAPN2
CDH2
CDS1
SLCO1C1
PRLHR
UGCG
FBLN2
MRPS28
DENND4C
LRP3
NETO2
SMOX

FZR1
2-Mar
SOGA1
CAPZB
EGR3
MAPK8IP3
ESD
L3MBTL1
CAP2
CD38
POLQ
CHRDL1
REEP1
DNAJB5
UACA
UBP1
SH3GLB1
PRRC2B
VWF
SERINC2
SLC31A2
RALBP1
ARHGEF5
CACNA1A
AR
FBXO33
SLC24A2
LIMS2
SUN1
RELL1
PIK3AP1
MAMDC2
SDCBP2
STK35
TRRAP
HMCN2
TOP3B
DMAP1
SCUBE2
HAGH
DPY19L4
BRCA2
FBXO10
THAP9
NUCB2

SLC39A10
C21orf2
ABHD8
SPOCK2
LARS2
GSE1
IDUA
HIVEP3
UHRF1BP1
KIAA0355
PLEKHG1
FAM208B
FJX1
VPS13A
SYNPO2
EIF3I
OPLAH
HELZ
KIF26B
CHEK2
MAP3K13
RAB11FIP4
IDH3A
UNC93A
CTDP1
ZNF236
QRSL1
NUP160
YPEL4
GRIK4
ZCCHC6
GABRP
KCNH6
CFAP46
AMPD2
GALNT7
VPS8
GATA2
TDRD12
SNAPC3
DRD2
INTS9
SLC12A3
PRDM5
WNT5B

RABL3
NLRC5
KLHL5
PRPF40B
VPS41
B4GALNT3
ARMC4
CD93
RFXANK
SHC1
STAC2
KCNT1
WSCD1
ARHGEF7
MLLT4
RPP30
NATD1
HGFAC
TNFRSF8
ZMYND10
RSPO3
ISY1
COPG2
EFHC2
UQCC1
LDB3
PHYHD1
ANKLE1
FCHSD2
GNAI3
DNAJC28
FOXP1
SEC61A1
SSB
YME1L1
CHUK
ATG16L1
PRDM15
CCDC82
NCOR2
PEAR1
CYP2R1
RCSD1
ARHGAP42
STOML2

CDH13
SYNRG
CUL9
IGDCC4
PHRF1
MKRN2
SDK2
JADE1
GABRR2
ADAM10
AK7
PTPRT
PPAP2A
ZNF511
PARD3B
ADAT1
G6PC3
RFX1
BRINP1
GRIN3A
SLAIN1
NAT1
NUF2
ASTN2
LMF2
MEP1B
ATRN
BOC
BTF3
PRKAG2
CARKD
WDR19
ARMC2
BEND5
HDHD2
OFCC1
EHMT2
PLCB1
DENND5A
NIPSNAP3A
SIM1
SMYD1
RAB11FIP3
BRD4
CAPN9

PRKX
GUCA1A
SCHIP1
NLE1
IFT52
EPB41
EMX2
PSAP
ETV5
C5orf22
ITPRIPL2
PLA2G12B
SLC6A14
HPRT1
ASB11
PPM1F
EML4
RBFOX3
DPY19L3
SOAT1
ARID1B
PLCL1
NFIX
LOXL4
CASC4
HDDC3
FRY
CHAF1B
CHRNA3
SLC44A5
BHMG1
AKNA
RAPGEF2
FBXL20
IL1RAP
MAPKBP1
PHF2
SLC39A9
MANSC4
PDGFA
TRPC4AP
SUSD1
DDX46
ROCK2
ADAM11

SVEP1
TTYH3
VSTM2L
DNAAF1
CACNA2D2
CPNE2
NOC2L
ZBTB7C
MAB21L1
WDR62
ESR2
RPGRIP1
DIS3L2
RNF157
ANKRD10
UBTD1
CNTNAP5
IDH1
BIRC6
NOS1AP
EHD2
KIFC3
NOLC1
OBSCN
C9orf85
ADAMTS7
ECI1
CLSTN2
RUFY2
PVRL1
B4GALNT4
DHX30
DSG2
TRIM33
BARX1
TGFBR2
SECISBP2
CEP97
TDRD3
DTWD1
TRIM24
NDRG3
RNGTT
PPP1R9A
EIF2B3

DGKB
ERCC5
NOBOX
KCNK10
SRP72
PLK4
NUP85
ARNT2
EPHB2
CHRNG
COL6A2
CAPN3
ZBTB44
PGAP1
PILRA
ABCA3
AQR
FBXL3
DDX4
CD101
ENPP3
MCF2L
NUP188
ZNF827
ISL1
MPP5
RANBP9
KPNA4
HMHA1
TBX22
AMPH
ACE
WDR88
MISP
AFAP1L1
CTNND2
GSR
SBF2
TMOD1
PHLDB2
EME1
ATP11A
IL18RAP
CFAP47
FMR1

ATAD3A
PSME4
FIGLA
PRMT8
DPY19L1
NAP1L4
EIF2AK3
SLC13A3
SLC25A26
DOCK11
TMEM62
SCAF11
SLC39A11
CLUAP1
SGK494
FAM227A
TRPC6
C2orf72
MAP2K4
THEMIS
PCYT1A
UBE2E3
CACNA1G
CEP295
RTFDC1
FADS1
TMEM220
TBXAS1
ATP6AP1L
TNR
MKLN1
SMO
DMXL1
KLHL41
ADAM23
KLHL18
PHKA1
ABCD3
VPS4B
IGF2BP1
DNAI1
UBE2D3
C16orf70
GRIK5
SMAD2

FLNC
FAM117B
TNFRSF14
PTK2
AP2A2
TNRC6A
COL17A1
HELLS
FSCN3
TRAPPC9
UBE4A
STIM1
CARF
POT1
MYO3B
ANTXR1
VEPH1
CCDC171
LHX5
ZDHHC11B
RBM46
AHCYL2
DBR1
MSL2
EZH1
RWDD1
WNK2
PLBD1
OGFOD1
SAMD3
MTF1
CFDP1
CFAP43
ADGRL4
SCYL3
APP
GATA4
GSTCD
CTSL
THOC5
CPNE1
GTF2F2
SNX31
STXBP1
GNPAT

TBX1
INPP5F
MSTO1
ENAH
CLRN3
UBAC1
FAM49B
NFIB
RFX3
ADGRA2
SLCO4A1
RABGAP1
TXNRD1
FGD5
PIGK
ZBTB22
FAM189A1
CDC25B
RBM33
DEPDC1
DCDC2B
ATP8B1
LMAN1
ACADSB
NUP93
EIF4E3
KCTD12
ENOSF1
CHM
CENPF
KLF13
ERI3
GTF2A1
MOV10
PMPCA
RPS6KA5
IRX4
POU2F1
CDH22
TP53BP2
TAF4B
MPP7
FAM151B
DCAF17
BTBD11

NF2
SLC6A2
ANKRD6
PRMT7
TMEM128
TMEM132E
SLC19A3
RAB1A
MCTP2
STXBP6
DFNB31
IKZF2
IKBKAP
ACAP3
RFX2
NKX2-6
CBWD1
CDC23
DHRS7
ADD3
TNN
TECPR2
EDAR
CACUL1
TMEM68
FGGY
ATP11B
LHX9
QSER1
SH3YL1
LMBR1
C16orf86
PTGR1
SPDL1
ARAP1
GJA4
PCNXL2
NKX2-8
SOS2
RNF166
PHLPP1
ZDHHC9
MTMR9
CEP89
FIS1

PARP8
CDKN2AIP
EMILIN1
MYT1L
TRIM62
ADAMTSL4
TMEM255A
ME3
RNF217
POLR2E
RRM2B
TBC1D22B
STOX2
ADCY9
PPM1L
ARL6IP1
BLK
KLHDC10
MINK1
MUSK
9-Sep
CPPED1
GANC
TRIM36
ZNF654
DPM2
UBE3B
DDX10
MGAT4A
FAM193B
SNX3
CPXM2
NINJ2
SGSM1
PLOD1
DTX3L
MAP2K3
UBFD1
FBXW9
HMP19
CPNE8
CACNA1C
UBR4
SLC4A5
GALNT6

UBE2E1
EXOC4
PXDC1
GDPD5
MTSS1L
ASXL3
KDM6B
ATAT1
PAPD7
AKAP6
ATF2
SETBP1
JUP
LRBA
CNGA4
TMPRSS6
EVA1A
NUMA1
WDR27
SLC38A10
KIZ
OSBPL5
TFEC
ERCC6
HERC1
SLC35A5
SON
ANK1
GBF1
PWP2
CCDC87
HIPK2
EPS15
CMIP
PTPRN
CCDC50
ADAT2
SHQ1
MYBL2
L3MBTL3
SEC63
ANKEF1
KLHL14
ABCC8
MAP7D3

TRPC5
CC2D1B
CATSPER4
METTL15
RNF122
KDM3B
EHBP1
CABP2
ZNRD1
CACNA1B
GRAP
SMOC2
PIK3C2A
NFASC
HLTF
MASTL
GFRA2
SIK3
STK24
NBAS
WDR18
MUC16
MET
ABCB10
RPRD1B
HCK
ICE2
CXADR
BRMS1L
FAM102A
PLEKHN1
RASAL3
RPRD2
LRRC2
PTGFR
PIGO
CORO2A
STK4
RNF115
SDK1
CNTN1
PLK1
CRLF1
CCDC88C
POLR2A

DDX49
ANKS1A
RCL1
GXYLT2
REXO1
DNAJC2
ATP8A1
FXYD2
CEBPZ
CAMK4
RSPRY1
DCAF7
TPTE2
DPH6
THADA
RHOG
KIF4A
PIEZO2
ADAM12
ADCY10
SH2D3C
FHAD1
RGS19
KAT6A
EYA2
PREP
NBN
SPECC1
FANCC
GAL3ST1
TMEM39B
VAC14
SETDB2
ING1
KDM4A
PSMB9
ATAD1
FAM45A
BCL10
PKP4
ABCC9
RSRC1
TOP2A
MRPS18C
ENTHD1

RRNAD1
HAGHL
ARHGEF28
QDPR
UBR5
CDC27
CTAGE5
CIDEC
NEK8
UNC80
SMC4
PMEPA1
RASL12
MPPED1
FAM184A
C18orf54
DCX
PAPD5
HADHA
KIF13B
CHD3
CDC40
AGBL1
METTL4
XXYLT1
IMMP2L
CEP76
CHFR
FPGT-TNNI3K
MFSD12
VWCE
LRCH2
HNRNPR
HSPA4L
GALNT12
COL6A6
POLA1
TGDS
GMPR2
CELSR3
OSBPL9
NMNAT3
LAPTM4B
KHDRBS1
EOGT

VANGL2
MGAT5B
CNDP2
NDUFS7
SETD1B

### B DAVID gene name of list of affected genes from tab "A"

| From | To | Species | David Gene Name |
|---|---|---|---|
| ATF2 | 1386 | Homo sapiens | activating transcription factor 2(ATF2) |
| ITSN1 | 6453 | Homo sapiens | intersectin 1(ITSN1) |
| TESK2 | 10420 | Homo sapiens | testis associated actin remodelling kinase 2(TESK2) |
| F11R | 50848 | Homo sapiens | F11 receptor(F11R) |
| NOC2L | 26155 | Homo sapiens | NOC2 like nucleolar associated transcriptional repressor(NOC2L) |
| TSKU | 25987 | Homo sapiens | tsukushi, small leucine rich proteoglycan(TSKU) |
| ALKBH8 | 91801 | Homo sapiens | alkB homolog 8, tRNA methyltransferase(ALKBH8) |
| ALKBH3 | 221120 | Homo sapiens | alkB homolog 3, alpha-ketoglutarate dependent dioxygenase(A |
| CAPNS1 | 826 | Homo sapiens | calpain small subunit 1(CAPNS1) |
| LURAP1L | 286343 | Homo sapiens | leucine rich adaptor protein 1 like(LURAP1L) |
| DPYSL5 | 56896 | Homo sapiens | dihydropyrimidinase like 5(DPYSL5) |
| GPBP1 | 65056 | Homo sapiens | GC-rich promoter binding protein 1(GPBP1) |
| EPSTI1 | 94240 | Homo sapiens | epithelial stromal interaction 1(EPSTI1) |
| AKT1 | 207 | Homo sapiens | AKT serine/threonine kinase 1(AKT1) |
| FNBP4 | 23360 | Homo sapiens | formin binding protein 4(FNBP4) |
| SAP30BP | 29115 | Homo sapiens | SAP30 binding protein(SAP30BP) |
| CXADR | 1525 | Homo sapiens | CXADR Ig-like cell adhesion molecule(CXADR) |
| TSTD1 | 100131187 | Homo sapiens | thiosulfate sulfurtransferase like domain containing 1(TSTD1) |
| SECISBP2 | 79048 | Homo sapiens | SECIS binding protein 2(SECISBP2) |
| CACNA2D2 | 9254 | Homo sapiens | calcium voltage-gated channel auxiliary subunit alpha2delta 2(C |
| PHKA1 | 5255 | Homo sapiens | phosphorylase kinase regulatory subunit alpha 1(PHKA1) |
| ISL1 | 3670 | Homo sapiens | ISL LIM homeobox 1(ISL1) |
| LARS2 | 23395 | Homo sapiens | leucyl-tRNA synthetase 2, mitochondrial(LARS2) |
| ISL2 | 64843 | Homo sapiens | ISL LIM homeobox 2(ISL2) |
| SCO2 | 9997 | Homo sapiens | synthesis of cytochrome C oxidase 2(SCO2) |
| OPTC | 26254 | Homo sapiens | opticin(OPTC) |
| FEZ1 | 9638 | Homo sapiens | fasciculation and elongation protein zeta 1(FEZ1) |
| RABL3 | 285282 | Homo sapiens | RAB, member of RAS oncogene family like 3(RABL3) |
| PLBD1 | 79887 | Homo sapiens | phospholipase B domain containing 1(PLBD1) |
| NOBOX | 135935 | Homo sapiens | NOBOX oogenesis homeobox(NOBOX) |
| ATF6 | 22926 | Homo sapiens | activating transcription factor 6(ATF6) |
| SKAP1 | 8631 | Homo sapiens | src kinase associated phosphoprotein 1(SKAP1) |
| EXD3 | 54932 | Homo sapiens | exonuclease 3'-5' domain containing 3(EXD3) |
| CDCA4 | 55038 | Homo sapiens | cell division cycle associated 4(CDCA4) |
| LYPD1 | 116372 | Homo sapiens | LY6/PLAUR domain containing 1(LYPD1) |
| PDGFA | 5154 | Homo sapiens | platelet derived growth factor subunit A(PDGFA) |
| ZNF22 | 7570 | Homo sapiens | zinc finger protein 22(ZNF22) |
| GATA4 | 2626 | Homo sapiens | GATA binding protein 4(GATA4) |

| | | | |
|---|---|---|---|
| BAZ2B | 29994 | Homo sapiens | bromodomain adjacent to zinc finger domain 2B(BAZ2B) |
| GATA2 | 2624 | Homo sapiens | GATA binding protein 2(GATA2) |
| ADAMTS15 | 170689 | Homo sapiens | ADAM metallopeptidase with thrombospondin type 1 motif 15(A |
| RHOT2 | 89941 | Homo sapiens | ras homolog family member T2(RHOT2) |
| ADAMTS14 | 140766 | Homo sapiens | ADAM metallopeptidase with thrombospondin type 1 motif 14(A |
| PRPSAP2 | 5636 | Homo sapiens | phosphoribosyl pyrophosphate synthetase associated protein 2( |
| RHOT1 | 55288 | Homo sapiens | ras homolog family member T1(RHOT1) |
| RDH10 | 157506 | Homo sapiens | retinol dehydrogenase 10(RDH10) |
| EPB41L2 | 2037 | Homo sapiens | erythrocyte membrane protein band 4.1 like 2(EPB41L2) |
| ADAMTS18 | 170692 | Homo sapiens | ADAM metallopeptidase with thrombospondin type 1 motif 18(A |
| PDGFD | 80310 | Homo sapiens | platelet derived growth factor D(PDGFD) |
| RNF217 | 154214 | Homo sapiens | ring finger protein 217(RNF217) |
| ATG7 | 10533 | Homo sapiens | autophagy related 7(ATG7) |
| JAG2 | 3714 | Homo sapiens | jagged canonical Notch ligand 2(JAG2) |
| TOE1 | 114034 | Homo sapiens | target of EGR1, exonuclease(TOE1) |
| BTBD19 | 149478 | Homo sapiens | BTB domain containing 19(BTBD19) |
| JUP | 3728 | Homo sapiens | junction plakoglobin(JUP) |
| PBX3 | 5090 | Homo sapiens | PBX homeobox 3(PBX3) |
| EIF2AK3 | 9451 | Homo sapiens | eukaryotic translation initiation factor 2 alpha kinase 3(EIF2AK3 |
| EIF2AK4 | 440275 | Homo sapiens | eukaryotic translation initiation factor 2 alpha kinase 4(EIF2AK4 |
| MOV10 | 4343 | Homo sapiens | Mov10 RNA helicase(MOV10) |
| PPFIBP1 | 8496 | Homo sapiens | PPFIA binding protein 1(PPFIBP1) |
| VANGL2 | 57216 | Homo sapiens | VANGL planar cell polarity protein 2(VANGL2) |
| FXYD2 | 486 | Homo sapiens | FXYD domain containing ion transport regulator 2(FXYD2) |
| AGO2 | 27161 | Homo sapiens | argonaute RISC catalytic component 2(AGO2) |
| NMT1 | 4836 | Homo sapiens | N-myristoyltransferase 1(NMT1) |
| FAT1 | 2195 | Homo sapiens | FAT atypical cadherin 1(FAT1) |
| HYOU1 | 10525 | Homo sapiens | hypoxia up-regulated 1(HYOU1) |
| STRN | 6801 | Homo sapiens | striatin(STRN) |
| FAT3 | 120114 | Homo sapiens | FAT atypical cadherin 3(FAT3) |
| FAT4 | 79633 | Homo sapiens | FAT atypical cadherin 4(FAT4) |
| SFXN5 | 94097 | Homo sapiens | sideroflexin 5(SFXN5) |
| LGMN | 5641 | Homo sapiens | legumain(LGMN) |
| KCNC3 | 3748 | Homo sapiens | potassium voltage-gated channel subfamily C member 3(KCNC |
| DPYS | 1807 | Homo sapiens | dihydropyrimidinase(DPYS) |
| PRKAG2 | 51422 | Homo sapiens | protein kinase AMP-activated non-catalytic subunit gamma 2(PI |
| SNX11 | 29916 | Homo sapiens | sorting nexin 11(SNX11) |
| SHB | 6461 | Homo sapiens | SH2 domain containing adaptor protein B(SHB) |
| PEAR1 | 375033 | Homo sapiens | platelet endothelial aggregation receptor 1(PEAR1) |
| OSBPL10 | 114884 | Homo sapiens | oxysterol binding protein like 10(OSBPL10) |
| MED17 | 9440 | Homo sapiens | mediator complex subunit 17(MED17) |
| CHAF1B | 8208 | Homo sapiens | chromatin assembly factor 1 subunit B(CHAF1B) |
| NHSL2 | 340527 | Homo sapiens | NHS like 2(NHSL2) |
| NHSL1 | 57224 | Homo sapiens | NHS like 1(NHSL1) |
| MED13 | 9969 | Homo sapiens | mediator complex subunit 13(MED13) |

| | | | |
|---|---|---|---|
| MMP24 | 10893 | Homo sapiens | matrix metallopeptidase 24(MMP24) |
| MMP28 | 79148 | Homo sapiens | matrix metallopeptidase 28(MMP28) |
| OLR1 | 4973 | Homo sapiens | oxidized low density lipoprotein receptor 1(OLR1) |
| ENOPH1 | 58478 | Homo sapiens | enolase-phosphatase 1(ENOPH1) |
| DLGAP4 | 22839 | Homo sapiens | DLG associated protein 4(DLGAP4) |
| MYBPC1 | 4604 | Homo sapiens | myosin binding protein C1(MYBPC1) |
| MYBPC2 | 4606 | Homo sapiens | myosin binding protein C2(MYBPC2) |
| GSTO1 | 9446 | Homo sapiens | glutathione S-transferase omega 1(GSTO1) |
| SPHK2 | 56848 | Homo sapiens | sphingosine kinase 2(SPHK2) |
| METRN | 79006 | Homo sapiens | meteorin, glial cell differentiation regulator(METRN) |
| SPHK1 | 8877 | Homo sapiens | sphingosine kinase 1(SPHK1) |
| MAMDC2 | 256691 | Homo sapiens | MAM domain containing 2(MAMDC2) |
| SLC39A11 | 201266 | Homo sapiens | solute carrier family 39 member 11(SLC39A11) |
| SLC39A10 | 57181 | Homo sapiens | solute carrier family 39 member 10(SLC39A10) |
| DUSP8 | 1850 | Homo sapiens | dual specificity phosphatase 8(DUSP8) |
| PGM2L1 | 283209 | Homo sapiens | phosphoglucomutase 2 like 1(PGM2L1) |
| MED23 | 9439 | Homo sapiens | mediator complex subunit 23(MED23) |
| TMEFF1 | 8577 | Homo sapiens | transmembrane protein with EGF like and two follistatin like do[illegible] |
| MED25 | 81857 | Homo sapiens | mediator complex subunit 25(MED25) |
| MED24 | 9862 | Homo sapiens | mediator complex subunit 24(MED24) |
| MMP16 | 4325 | Homo sapiens | matrix metallopeptidase 16(MMP16) |
| MED21 | 9412 | Homo sapiens | mediator complex subunit 21(MED21) |
| PXDN | 7837 | Homo sapiens | peroxidasin(PXDN) |
| POT1 | 25913 | Homo sapiens | protection of telomeres 1(POT1) |
| RAPGEF2 | 9693 | Homo sapiens | Rap guanine nucleotide exchange factor 2(RAPGEF2) |
| L2HGDH | 79944 | Homo sapiens | L-2-hydroxyglutarate dehydrogenase(L2HGDH) |
| CACUL1 | 143384 | Homo sapiens | CDK2 associated cullin domain 1(CACUL1) |
| RAPGEF6 | 51735 | Homo sapiens | Rap guanine nucleotide exchange factor 6(RAPGEF6) |
| RAPGEF4 | 11069 | Homo sapiens | Rap guanine nucleotide exchange factor 4(RAPGEF4) |
| CRB2 | 286204 | Homo sapiens | crumbs cell polarity complex component 2(CRB2) |
| SYNPO2 | 171024 | Homo sapiens | synaptopodin 2(SYNPO2) |
| CCM2L | 140706 | Homo sapiens | CCM2 like scaffold protein(CCM2L) |
| SNX30 | 401548 | Homo sapiens | sorting nexin family member 30(SNX30) |
| ABCB11 | 8647 | Homo sapiens | ATP binding cassette subfamily B member 11(ABCB11) |
| SNX31 | 169166 | Homo sapiens | sorting nexin 31(SNX31) |
| FAM117B | 150864 | Homo sapiens | family with sequence similarity 117 member B(FAM117B) |
| CENPC | 1060 | Homo sapiens | centromere protein C(CENPC) |
| SNX29 | 92017 | Homo sapiens | sorting nexin 29(SNX29) |
| WRN | 7486 | Homo sapiens | WRN RecQ like helicase(WRN) |
| EXOSC10 | 5394 | Homo sapiens | exosome component 10(EXOSC10) |
| SNX25 | 83891 | Homo sapiens | sorting nexin 25(SNX25) |
| ADAM23 | 8745 | Homo sapiens | ADAM metallopeptidase domain 23(ADAM23) |
| ZSWIM8 | 23053 | Homo sapiens | zinc finger SWIM-type containing 8(ZSWIM8) |
| FPGT-TNNI3K | 100526835 | Homo sapiens | FPGT-TNNI3K readthrough(FPGT-TNNI3K) |
| SAMD3 | 154075 | Homo sapiens | sterile alpha motif domain containing 3(SAMD3) |

| | | | |
|---|---|---|---|
| CENPT | 80152 | Homo sapiens | centromere protein T(CENPT) |
| ATAD1 | 84896 | Homo sapiens | ATPase family AAA domain containing 1(ATAD1) |
| MBD3 | 53615 | Homo sapiens | methyl-CpG binding domain protein 3(MBD3) |
| ZNF462 | 58499 | Homo sapiens | zinc finger protein 462(ZNF462) |
| RIC3 | 79608 | Homo sapiens | RIC3 acetylcholine receptor chaperone(RIC3) |
| MBD5 | 55777 | Homo sapiens | methyl-CpG binding domain protein 5(MBD5) |
| CRBN | 51185 | Homo sapiens | cereblon(CRBN) |
| TRAPPC10 | 7109 | Homo sapiens | trafficking protein particle complex subunit 10(TRAPPC10) |
| KIAA2012 | 100652824 | Homo sapiens | KIAA2012(KIAA2012) |
| ADAM32 | 203102 | Homo sapiens | ADAM metallopeptidase domain 32(ADAM32) |
| DCBLD2 | 131566 | Homo sapiens | discoidin, CUB and LCCL domain containing 2(DCBLD2) |
| ARCN1 | 372 | Homo sapiens | archain 1(ARCN1) |
| CENPF | 1063 | Homo sapiens | centromere protein F(CENPF) |
| SNX18 | 112574 | Homo sapiens | sorting nexin 18(SNX18) |
| ZNF70 | 7621 | Homo sapiens | zinc finger protein 70(ZNF70) |
| ABCB10 | 23456 | Homo sapiens | ATP binding cassette subfamily B member 10(ABCB10) |
| NCAPD2 | 9918 | Homo sapiens | non-SMC condensin I complex subunit D2(NCAPD2) |
| CENPO | 79172 | Homo sapiens | centromere protein O(CENPO) |
| NCAPD3 | 23310 | Homo sapiens | non-SMC condensin II complex subunit D3(NCAPD3) |
| ADGRL4 | 64123 | Homo sapiens | adhesion G protein-coupled receptor L4(ADGRL4) |
| AGRN | 375790 | Homo sapiens | agrin(AGRN) |
| YPEL4 | 219539 | Homo sapiens | yippee like 4(YPEL4) |
| FRYL | 285527 | Homo sapiens | FRY like transcription coactivator(FRYL) |
| CRLF1 | 9244 | Homo sapiens | cytokine receptor like factor 1(CRLF1) |
| LIMS2 | 55679 | Homo sapiens | LIM zinc finger domain containing 2(LIMS2) |
| ZNF451 | 26036 | Homo sapiens | zinc finger protein 451(ZNF451) |
| OLFML2B | 25903 | Homo sapiens | olfactomedin like 2B(OLFML2B) |
| RIF1 | 55183 | Homo sapiens | replication timing regulatory factor 1(RIF1) |
| PLOD1 | 5351 | Homo sapiens | procollagen-lysine,2-oxoglutarate 5-dioxygenase 1(PLOD1) |
| IL1RAP | 3556 | Homo sapiens | interleukin 1 receptor accessory protein(IL1RAP) |
| FRY | 10129 | Homo sapiens | FRY microtubule binding protein(FRY) |
| FADS2 | 9415 | Homo sapiens | fatty acid desaturase 2(FADS2) |
| LAPTM4B | 55353 | Homo sapiens | lysosomal protein transmembrane 4 beta(LAPTM4B) |
| PTAR1 | 375743 | Homo sapiens | protein prenyltransferase alpha subunit repeat containing 1(PTA |
| HHAT | 55733 | Homo sapiens | hedgehog acyltransferase(HHAT) |
| SLK | 9748 | Homo sapiens | STE20 like kinase(SLK) |
| TNFRSF8 | 943 | Homo sapiens | TNF receptor superfamily member 8(TNFRSF8) |
| MAP3K8 | 1326 | Homo sapiens | mitogen-activated protein kinase kinase kinase 8(MAP3K8) |
| NUDT14 | 256281 | Homo sapiens | nudix hydrolase 14(NUDT14) |
| FADS1 | 3992 | Homo sapiens | fatty acid desaturase 1(FADS1) |
| KCNH1 | 3756 | Homo sapiens | potassium voltage-gated channel subfamily H member 1(KCNH |
| MAP3K5 | 4217 | Homo sapiens | mitogen-activated protein kinase kinase kinase 5(MAP3K5) |
| KCNH2 | 3757 | Homo sapiens | potassium voltage-gated channel subfamily H member 2(KCNH |
| CDKL3 | 51265 | Homo sapiens | cyclin dependent kinase like 3(CDKL3) |
| KCNH4 | 23415 | Homo sapiens | potassium voltage-gated channel subfamily H member 4(KCNH |

| | | | |
|---|---|---|---|
| DGAT2 | 84649 | Homo sapiens | diacylglycerol O-acyltransferase 2(DGAT2) |
| FBXW8 | 26259 | Homo sapiens | F-box and WD repeat domain containing 8(FBXW8) |
| KCNH6 | 81033 | Homo sapiens | potassium voltage-gated channel subfamily H member 6(KCNH |
| FST | 10468 | Homo sapiens | follistatin(FST) |
| STX8 | 9482 | Homo sapiens | syntaxin 8(STX8) |
| FBXW9 | 84261 | Homo sapiens | F-box and WD repeat domain containing 9(FBXW9) |
| ATP11C | 286410 | Homo sapiens | ATPase phospholipid transporting 11C(ATP11C) |
| ATP11B | 23200 | Homo sapiens | ATPase phospholipid transporting 11B (putative)(ATP11B) |
| ATP11A | 23250 | Homo sapiens | ATPase phospholipid transporting 11A(ATP11A) |
| FBXO10 | 26267 | Homo sapiens | F-box protein 10(FBXO10) |
| SDCBP2 | 27111 | Homo sapiens | syndecan binding protein 2(SDCBP2) |
| PIAS2 | 9063 | Homo sapiens | protein inhibitor of activated STAT 2(PIAS2) |
| CDC25B | 994 | Homo sapiens | cell division cycle 25B(CDC25B) |
| VMAC | 400673 | Homo sapiens | vimentin type intermediate filament associated coiled-coil prote |
| SMO | 6608 | Homo sapiens | smoothened, frizzled class receptor(SMO) |
| ZNF438 | 220929 | Homo sapiens | zinc finger protein 438(ZNF438) |
| PKP2 | 5318 | Homo sapiens | plakophilin 2(PKP2) |
| SRSF4 | 6429 | Homo sapiens | serine and arginine rich splicing factor 4(SRSF4) |
| GMNC | 647309 | Homo sapiens | geminin coiled-coil domain containing(GMNC) |
| ZNF436 | 80818 | Homo sapiens | zinc finger protein 436(ZNF436) |
| PKP4 | 8502 | Homo sapiens | plakophilin 4(PKP4) |
| PKP3 | 11187 | Homo sapiens | plakophilin 3(PKP3) |
| CDKL1 | 8814 | Homo sapiens | cyclin dependent kinase like 1(CDKL1) |
| BLK | 640 | Homo sapiens | BLK proto-oncogene, Src family tyrosine kinase(BLK) |
| FTO | 79068 | Homo sapiens | FTO alpha-ketoglutarate dependent dioxygenase(FTO) |
| COL17A1 | 1308 | Homo sapiens | collagen type XVII alpha 1 chain(COL17A1) |
| CD274 | 29126 | Homo sapiens | CD274 molecule(CD274) |
| KHDRBS1 | 10657 | Homo sapiens | KH RNA binding domain containing, signal transduction associat |
| SHC2 | 25759 | Homo sapiens | SHC adaptor protein 2(SHC2) |
| KHDRBS3 | 10656 | Homo sapiens | KH RNA binding domain containing, signal transduction associat |
| SHC1 | 6464 | Homo sapiens | SHC adaptor protein 1(SHC1) |
| KMT2C | 58508 | Homo sapiens | lysine methyltransferase 2C(KMT2C) |
| UAP1 | 6675 | Homo sapiens | UDP-N-acetylglucosamine pyrophosphorylase 1(UAP1) |
| SPATA7 | 55812 | Homo sapiens | spermatogenesis associated 7(SPATA7) |
| NPAS3 | 64067 | Homo sapiens | neuronal PAS domain protein 3(NPAS3) |
| FBXO42 | 54455 | Homo sapiens | F-box protein 42(FBXO42) |
| CORO1C | 23603 | Homo sapiens | coronin 1C(CORO1C) |
| IGSF21 | 84966 | Homo sapiens | immunoglobin superfamily member 21(IGSF21) |
| LOXHD1 | 125336 | Homo sapiens | lipoxygenase homology PLAT domains 1(LOXHD1) |
| NDOR1 | 27158 | Homo sapiens | NADPH dependent diflavin oxidoreductase 1(NDOR1) |
| MED30 | 90390 | Homo sapiens | mediator complex subunit 30(MED30) |
| SMYD1 | 150572 | Homo sapiens | SET and MYND domain containing 1(SMYD1) |
| TNFRSF14 | 8764 | Homo sapiens | TNF receptor superfamily member 14(TNFRSF14) |
| KDSR | 2531 | Homo sapiens | 3-ketodihydrosphingosine reductase(KDSR) |
| TSNAXIP1 | 55815 | Homo sapiens | translin associated factor X interacting protein 1(TSNAXIP1) |

| | | | |
|---|---|---|---|
| GRIA3 | 2892 | Homo sapiens | glutamate ionotropic receptor AMPA type subunit 3(GRIA3) |
| GRIA4 | 2893 | Homo sapiens | glutamate ionotropic receptor AMPA type subunit 4(GRIA4) |
| NEK8 | 284086 | Homo sapiens | NIMA related kinase 8(NEK8) |
| HHATL | 57467 | Homo sapiens | hedgehog acyltransferase like(HHATL) |
| RASL10B | 91608 | Homo sapiens | RAS like family 10 member B(RASL10B) |
| RFXANK | 8625 | Homo sapiens | regulatory factor X associated ankyrin containing protein(RFXAN |
| CORO2A | 7464 | Homo sapiens | coronin 2A(CORO2A) |
| FBXO33 | 254170 | Homo sapiens | F-box protein 33(FBXO33) |
| NDC80 | 10403 | Homo sapiens | NDC80 kinetochore complex component(NDC80) |
| UCKL1 | 54963 | Homo sapiens | uridine-cytidine kinase 1 like 1(UCKL1) |
| FBXO31 | 79791 | Homo sapiens | F-box protein 31(FBXO31) |
| LRCH2 | 57631 | Homo sapiens | leucine rich repeats and calponin homology domain containing 2 |
| G6PC3 | 92579 | Homo sapiens | glucose-6-phosphatase catalytic subunit 3(G6PC3) |
| SON | 6651 | Homo sapiens | SON DNA and RNA binding protein(SON) |
| LRCH1 | 23143 | Homo sapiens | leucine rich repeats and calponin homology domain containing 1 |
| DMXL1 | 1657 | Homo sapiens | Dmx like 1(DMXL1) |
| NOXA1 | 10811 | Homo sapiens | NADPH oxidase activator 1(NOXA1) |
| TBXAS1 | 6916 | Homo sapiens | thromboxane A synthase 1(TBXAS1) |
| SP5 | 389058 | Homo sapiens | Sp5 transcription factor(SP5) |
| ACO1 | 48 | Homo sapiens | aconitase 1(ACO1) |
| CLPTM1L | 81037 | Homo sapiens | CLPTM1 like(CLPTM1L) |
| OSBPL1A | 114876 | Homo sapiens | oxysterol binding protein like 1A(OSBPL1A) |
| SHBG | 6462 | Homo sapiens | sex hormone binding globulin(SHBG) |
| LRCH3 | 84859 | Homo sapiens | leucine rich repeats and calponin homology domain containing 3 |
| ZNF410 | 57862 | Homo sapiens | zinc finger protein 410(ZNF410) |
| ISY1 | 57461 | Homo sapiens | ISY1 splicing factor homolog(ISY1) |
| PIWIL4 | 143689 | Homo sapiens | piwi like RNA-mediated gene silencing 4(PIWIL4) |
| NUMA1 | 4926 | Homo sapiens | nuclear mitotic apparatus protein 1(NUMA1) |
| MTCL1 | 23255 | Homo sapiens | microtubule crosslinking factor 1(MTCL1) |
| ITGB2 | 3689 | Homo sapiens | integrin subunit beta 2(ITGB2) |
| DYSF | 8291 | Homo sapiens | dysferlin(DYSF) |
| ECE1 | 1889 | Homo sapiens | endothelin converting enzyme 1(ECE1) |
| ETFA | 2108 | Homo sapiens | electron transfer flavoprotein subunit alpha(ETFA) |
| HTR2A | 3356 | Homo sapiens | 5-hydroxytryptamine receptor 2A(HTR2A) |
| MSLN | 10232 | Homo sapiens | mesothelin(MSLN) |
| TSPAN10 | 83882 | Homo sapiens | tetraspanin 10(TSPAN10) |
| CDC73 | 79577 | Homo sapiens | cell division cycle 73(CDC73) |
| BICD2 | 23299 | Homo sapiens | BICD cargo adaptor 2(BICD2) |
| UNC80 | 285175 | Homo sapiens | unc-80 homolog, NALCN channel complex subunit(UNC80) |
| KIAA1549 | 57670 | Homo sapiens | KIAA1549(KIAA1549) |
| PPP6R2 | 9701 | Homo sapiens | protein phosphatase 6 regulatory subunit 2(PPP6R2) |
| ZNF407 | 55628 | Homo sapiens | zinc finger protein 407(ZNF407) |
| BOC | 91653 | Homo sapiens | BOC cell adhesion associated, oncogene regulated(BOC) |
| PPP6R3 | 55291 | Homo sapiens | protein phosphatase 6 regulatory subunit 3(PPP6R3) |
| PRLHR | 2834 | Homo sapiens | prolactin releasing hormone receptor(PRLHR) |

| | | | |
|---|---|---|---|
| PGM5 | 5239 | Homo sapiens | phosphoglucomutase 5(PGM5) |
| ITGAV | 3685 | Homo sapiens | integrin subunit alpha V(ITGAV) |
| SPDL1 | 54908 | Homo sapiens | spindle apparatus coiled-coil protein 1(SPDL1) |
| TRIM65 | 201292 | Homo sapiens | tripartite motif containing 65(TRIM65) |
| TRIM66 | 9866 | Homo sapiens | tripartite motif containing 66(TRIM66) |
| CERS3 | 204219 | Homo sapiens | ceramide synthase 3(CERS3) |
| BCAS3 | 54828 | Homo sapiens | BCAS3 microtubule associated cell migration factor(BCAS3) |
| UNC45B | 146862 | Homo sapiens | unc-45 myosin chaperone B(UNC45B) |
| EIF5B | 9669 | Homo sapiens | eukaryotic translation initiation factor 5B(EIF5B) |
| ACTR2 | 10097 | Homo sapiens | actin related protein 2(ACTR2) |
| TRIM62 | 55223 | Homo sapiens | tripartite motif containing 62(TRIM62) |
| TMEM255A | 55026 | Homo sapiens | transmembrane protein 255A(TMEM255A) |
| CEP295 | 85459 | Homo sapiens | centrosomal protein 295(CEP295) |
| ACTR5 | 79913 | Homo sapiens | actin related protein 5(ACTR5) |
| COMMD10 | 51397 | Homo sapiens | COMM domain containing 10(COMMD10) |
| ATRX | 546 | Homo sapiens | ATRX chromatin remodeler(ATRX) |
| VPS13A | 23230 | Homo sapiens | vacuolar protein sorting 13 homolog A(VPS13A) |
| MSL2 | 55167 | Homo sapiens | MSL complex subunit 2(MSL2) |
| VPS13B | 157680 | Homo sapiens | vacuolar protein sorting 13 homolog B(VPS13B) |
| SEZ6L | 23544 | Homo sapiens | seizure related 6 homolog like(SEZ6L) |
| ARFGAP3 | 26286 | Homo sapiens | ADP ribosylation factor GTPase activating protein 3(ARFGAP3) |
| ATRN | 8455 | Homo sapiens | attractin(ATRN) |
| RRM2B | 50484 | Homo sapiens | ribonucleotide reductase regulatory TP53 inducible subunit M2B( |
| MYO3B | 140469 | Homo sapiens | myosin IIIB(MYO3B) |
| SPATC1 | 375686 | Homo sapiens | spermatogenesis and centriole associated 1(SPATC1) |
| PCCB | 5096 | Homo sapiens | propionyl-CoA carboxylase subunit beta(PCCB) |
| ITGA8 | 8516 | Homo sapiens | integrin subunit alpha 8(ITGA8) |
| LCOR | 84458 | Homo sapiens | ligand dependent nuclear receptor corepressor(LCOR) |
| LCP2 | 3937 | Homo sapiens | lymphocyte cytosolic protein 2(LCP2) |
| LCP1 | 3936 | Homo sapiens | lymphocyte cytosolic protein 1(LCP1) |
| TOP1 | 7150 | Homo sapiens | DNA topoisomerase I(TOP1) |
| BCAT1 | 586 | Homo sapiens | branched chain amino acid transaminase 1(BCAT1) |
| TRIM54 | 57159 | Homo sapiens | tripartite motif containing 54(TRIM54) |
| CERS1 | 10715 | Homo sapiens | ceramide synthase 1(CERS1) |
| TNFRSF21 | 27242 | Homo sapiens | TNF receptor superfamily member 21(TNFRSF21) |
| SNAP47 | 116841 | Homo sapiens | synaptosome associated protein 47(SNAP47) |
| UHRF2 | 115426 | Homo sapiens | ubiquitin like with PHD and ring finger domains 2(UHRF2) |
| ROCK2 | 9475 | Homo sapiens | Rho associated coiled-coil containing protein kinase 2(ROCK2) |
| MGRN1 | 23295 | Homo sapiens | mahogunin ring finger 1(MGRN1) |
| SRF | 6722 | Homo sapiens | serum response factor(SRF) |
| SRBD1 | 55133 | Homo sapiens | S1 RNA binding domain 1(SRBD1) |
| ATP10B | 23120 | Homo sapiens | ATPase phospholipid transporting 10B (putative)(ATP10B) |
| SRL | 6345 | Homo sapiens | sarcalumenin(SRL) |
| EHBP1 | 23301 | Homo sapiens | EH domain binding protein 1(EHBP1) |
| TSPAN31 | 6302 | Homo sapiens | tetraspanin 31(TSPAN31) |

| | | | |
|---|---|---|---|
| ZNRF1 | 84937 | Homo sapiens | zinc and ring finger 1(ZNRF1) |
| ZNRF2 | 223082 | Homo sapiens | zinc and ring finger 2(ZNRF2) |
| U2AF2 | 11338 | Homo sapiens | U2 small nuclear RNA auxiliary factor 2(U2AF2) |
| IGF2BP1 | 10642 | Homo sapiens | insulin like growth factor 2 mRNA binding protein 1(IGF2BP1) |
| TRIM47 | 91107 | Homo sapiens | tripartite motif containing 47(TRIM47) |
| E2F6 | 1876 | Homo sapiens | E2F transcription factor 6(E2F6) |
| UBQLN4 | 56893 | Homo sapiens | ubiquilin 4(UBQLN4) |
| TRIM44 | 54765 | Homo sapiens | tripartite motif containing 44(TRIM44) |
| TBX1 | 6899 | Homo sapiens | T-box transcription factor 1(TBX1) |
| CIZ1 | 25792 | Homo sapiens | CDKN1A interacting zinc finger protein 1(CIZ1) |
| SSB | 6741 | Homo sapiens | small RNA binding exonuclease protection factor La(SSB) |
| MYO10 | 4651 | Homo sapiens | myosin X(MYO10) |
| NTRK3 | 4916 | Homo sapiens | neurotrophic receptor tyrosine kinase 3(NTRK3) |
| CABLES1 | 91768 | Homo sapiens | Cdk5 and Abl enzyme substrate 1(CABLES1) |
| SYT14 | 255928 | Homo sapiens | synaptotagmin 14(SYT14) |
| SYT13 | 57586 | Homo sapiens | synaptotagmin 13(SYT13) |
| KIAA0232 | 9778 | Homo sapiens | KIAA0232(KIAA0232) |
| TM6SF1 | 53346 | Homo sapiens | transmembrane 6 superfamily member 1(TM6SF1) |
| TBX5 | 6910 | Homo sapiens | T-box transcription factor 5(TBX5) |
| SORL1 | 6653 | Homo sapiens | sortilin related receptor 1(SORL1) |
| MYO16 | 23026 | Homo sapiens | myosin XVI(MYO16) |
| TTC39A | 22996 | Homo sapiens | tetratricopeptide repeat domain 39A(TTC39A) |
| DYTN | 391475 | Homo sapiens | dystrotelin(DYTN) |
| TSPAN15 | 23555 | Homo sapiens | tetraspanin 15(TSPAN15) |
| MCM3AP | 8888 | Homo sapiens | minichromosome maintenance complex component 3 associate |
| NAGK | 55577 | Homo sapiens | N-acetylglucosamine kinase(NAGK) |
| TSPAN18 | 90139 | Homo sapiens | tetraspanin 18(TSPAN18) |
| EIF3I | 8668 | Homo sapiens | eukaryotic translation initiation factor 3 subunit I(EIF3I) |
| APLF | 200558 | Homo sapiens | aprataxin and PNKP like factor(APLF) |
| TRIM36 | 55521 | Homo sapiens | tripartite motif containing 36(TRIM36) |
| EIF3H | 8667 | Homo sapiens | eukaryotic translation initiation factor 3 subunit H(EIF3H) |
| KCP | 375616 | Homo sapiens | kielin cysteine rich BMP regulator(KCP) |
| TRIM37 | 4591 | Homo sapiens | tripartite motif containing 37(TRIM37) |
| PIK3AP1 | 118788 | Homo sapiens | phosphoinositide-3-kinase adaptor protein 1(PIK3AP1) |
| TBATA | 219793 | Homo sapiens | thymus, brain and testes associated(TBATA) |
| BCAR1 | 9564 | Homo sapiens | BCAR1 scaffold protein, Cas family member(BCAR1) |
| TRIM33 | 51592 | Homo sapiens | tripartite motif containing 33(TRIM33) |
| EIF3A | 8661 | Homo sapiens | eukaryotic translation initiation factor 3 subunit A(EIF3A) |
| CPNE8 | 144402 | Homo sapiens | copine 8(CPNE8) |
| THSD7B | 80731 | Homo sapiens | thrombospondin type 1 domain containing 7B(THSD7B) |
| CPNE4 | 131034 | Homo sapiens | copine 4(CPNE4) |
| ECI1 | 1632 | Homo sapiens | enoyl-CoA delta isomerase 1(ECI1) |
| CPNE7 | 27132 | Homo sapiens | copine 7(CPNE7) |
| ECI2 | 10455 | Homo sapiens | enoyl-CoA delta isomerase 2(ECI2) |
| LDLRAD3 | 143458 | Homo sapiens | low density lipoprotein receptor class A domain containing 3(LDI |

| | | | |
|---|---|---|---|
| IKZF1 | 10320 | Homo sapiens | IKAROS family zinc finger 1(IKZF1) |
| IKZF2 | 22807 | Homo sapiens | IKAROS family zinc finger 2(IKZF2) |
| RCBTB1 | 55213 | Homo sapiens | RCC1 and BTB domain containing protein 1(RCBTB1) |
| NUDT3 | 11165 | Homo sapiens | nudix hydrolase 3(NUDT3) |
| HYKK | 123688 | Homo sapiens | hydroxylysine kinase(HYKK) |
| CDC23 | 8697 | Homo sapiens | cell division cycle 23(CDC23) |
| HERC2 | 8924 | Homo sapiens | HECT and RLD domain containing E3 ubiquitin protein ligase 2(H |
| ADGRA1 | 84435 | Homo sapiens | adhesion G protein-coupled receptor A1(ADGRA1) |
| TRIM28 | 10155 | Homo sapiens | tripartite motif containing 28(TRIM28) |
| OPHN1 | 4983 | Homo sapiens | oligophrenin 1(OPHN1) |
| HERC1 | 8925 | Homo sapiens | HECT and RLD domain containing E3 ubiquitin protein ligase fan |
| ADGRA2 | 25960 | Homo sapiens | adhesion G protein-coupled receptor A2(ADGRA2) |
| TRIM29 | 23650 | Homo sapiens | tripartite motif containing 29(TRIM29) |
| TOPAZ1 | 375337 | Homo sapiens | testis and ovary specific TOPAZ 1(TOPAZ1) |
| BSG | 682 | Homo sapiens | basigin (Ok blood group)(BSG) |
| CDC27 | 996 | Homo sapiens | cell division cycle 27(CDC27) |
| TRIM24 | 8805 | Homo sapiens | tripartite motif containing 24(TRIM24) |
| PIEZO2 | 63895 | Homo sapiens | piezo type mechanosensitive ion channel component 2(PIEZO2) |
| CPNE1 | 8904 | Homo sapiens | copine 1(CPNE1) |
| PIEZO1 | 9780 | Homo sapiens | piezo type mechanosensitive ion channel component 1 (Er bloo |
| EPC1 | 80314 | Homo sapiens | enhancer of polycomb homolog 1(EPC1) |
| THSD7A | 221981 | Homo sapiens | thrombospondin type 1 domain containing 7A(THSD7A) |
| CPNE2 | 221184 | Homo sapiens | copine 2(CPNE2) |
| CCDC93 | 54520 | Homo sapiens | coiled-coil domain containing 93(CCDC93) |
| TBC1D22A | 25771 | Homo sapiens | TBC1 domain family member 22A(TBC1D22A) |
| TP63 | 8626 | Homo sapiens | tumor protein p63(TP63) |
| TBC1D22B | 55633 | Homo sapiens | TBC1 domain family member 22B(TBC1D22B) |
| ATP6V1G1 | 9550 | Homo sapiens | ATPase H+ transporting V1 subunit G1(ATP6V1G1) |
| SMARCC1 | 6599 | Homo sapiens | SWI/SNF related, matrix associated, actin dependent regulator |
| KCTD19 | 146212 | Homo sapiens | potassium channel tetramerization domain containing 19(KCTD |
| ALG6 | 29929 | Homo sapiens | ALG6 alpha-1,3-glucosyltransferase(ALG6) |
| KCNK10 | 54207 | Homo sapiens | potassium two pore domain channel subfamily K member 10(K |
| LIMCH1 | 22998 | Homo sapiens | LIM and calponin homology domains 1(LIMCH1) |
| TALDO1 | 6888 | Homo sapiens | transaldolase 1(TALDO1) |
| ADGRA3 | 166647 | Homo sapiens | adhesion G protein-coupled receptor A3(ADGRA3) |
| CSNK1D | 1453 | Homo sapiens | casein kinase 1 delta(CSNK1D) |
| UNC93A | 54346 | Homo sapiens | unc-93 homolog A(UNC93A) |
| ALG1 | 56052 | Homo sapiens | ALG1 chitobiosyldiphosphodolichol beta-mannosyltransferase(AL |
| KCNK17 | 89822 | Homo sapiens | potassium two pore domain channel subfamily K member 17(K |
| MFSD12 | 126321 | Homo sapiens | major facilitator superfamily domain containing 12(MFSD12) |
| TMEM170A | 124491 | Homo sapiens | transmembrane protein 170A(TMEM170A) |
| CDC40 | 51362 | Homo sapiens | cell division cycle 40(CDC40) |
| CYP39A1 | 51302 | Homo sapiens | cytochrome P450 family 39 subfamily A member 1(CYP39A1) |
| CFDP1 | 10428 | Homo sapiens | craniofacial development protein 1(CFDP1) |
| PPA2 | 27068 | Homo sapiens | inorganic pyrophosphatase 2(PPA2) |

| | | | |
|---|---|---|---|
| ADGRB2 | 576 | Homo sapiens | adhesion G protein-coupled receptor B2(ADGRB2) |
| SOAT1 | 6646 | Homo sapiens | sterol O-acyltransferase 1(SOAT1) |
| NINJ2 | 4815 | Homo sapiens | ninjurin 2(NINJ2) |
| CSNK2B | 1460 | Homo sapiens | casein kinase 2 beta(CSNK2B) |
| KCNQ4 | 9132 | Homo sapiens | potassium voltage-gated channel subfamily Q member 4(KCNQ |
| FAM76B | 143684 | Homo sapiens | family with sequence similarity 76 member B(FAM76B) |
| CD226 | 10666 | Homo sapiens | CD226 molecule(CD226) |
| CCDC87 | 55231 | Homo sapiens | coiled-coil domain containing 87(CCDC87) |
| KCTD12 | 115207 | Homo sapiens | potassium channel tetramerization domain containing 12(KCTD1 |
| CCDC82 | 79780 | Homo sapiens | coiled-coil domain containing 82(CCDC82) |
| TMEM63B | 55362 | Homo sapiens | transmembrane protein 63B(TMEM63B) |
| EPB41 | 2035 | Homo sapiens | erythrocyte membrane protein band 4.1(EPB41) |
| TMEM63C | 57156 | Homo sapiens | transmembrane protein 63C(TMEM63C) |
| DHX9 | 1660 | Homo sapiens | DExH-box helicase 9(DHX9) |
| SDC3 | 9672 | Homo sapiens | syndecan 3(SDC3) |
| MYCBP2 | 23077 | Homo sapiens | MYC binding protein 2(MYCBP2) |
| LPAR1 | 1902 | Homo sapiens | lysophosphatidic acid receptor 1(LPAR1) |
| CDKN2AIP | 55602 | Homo sapiens | CDKN2A interacting protein(CDKN2AIP) |
| GSPT1 | 2935 | Homo sapiens | G1 to S phase transition 1(GSPT1) |
| BRIP1 | 83990 | Homo sapiens | BRCA1 interacting helicase 1(BRIP1) |
| HLTF | 6596 | Homo sapiens | helicase like transcription factor(HLTF) |
| CEP192 | 55125 | Homo sapiens | centrosomal protein 192(CEP192) |
| B4GALNT3 | 283358 | Homo sapiens | beta-1,4-N-acetyl-galactosaminyltransferase 3(B4GALNT3) |
| B4GALNT2 | 124872 | Homo sapiens | beta-1,4-N-acetyl-galactosaminyltransferase 2 (SID blood grou |
| B4GALNT4 | 338707 | Homo sapiens | beta-1,4-N-acetyl-galactosaminyltransferase 4(B4GALNT4) |
| YES1 | 7525 | Homo sapiens | YES proto-oncogene 1, Src family tyrosine kinase(YES1) |
| EGR3 | 1960 | Homo sapiens | early growth response 3(EGR3) |
| TGDS | 23483 | Homo sapiens | TDP-glucose 4,6-dehydratase(TGDS) |
| NBEAL1 | 65065 | Homo sapiens | neurobeachin like 1(NBEAL1) |
| SUCO | 51430 | Homo sapiens | SUN domain containing ossification factor(SUCO) |
| SMARCA1 | 6594 | Homo sapiens | SWI/SNF related, matrix associated, actin dependent regulator |
| NTAN1 | 123803 | Homo sapiens | N-terminal asparagine amidase(NTAN1) |
| SMARCA2 | 6595 | Homo sapiens | SWI/SNF related, matrix associated, actin dependent regulator |
| PTK2 | 5747 | Homo sapiens | protein tyrosine kinase 2(PTK2) |
| MED13L | 23389 | Homo sapiens | mediator complex subunit 13L(MED13L) |
| EXT1 | 2131 | Homo sapiens | exostosin glycosyltransferase 1(EXT1) |
| CRNN | 49860 | Homo sapiens | cornulin(CRNN) |
| EXT2 | 2132 | Homo sapiens | exostosin glycosyltransferase 2(EXT2) |
| POLA1 | 5422 | Homo sapiens | DNA polymerase alpha 1, catalytic subunit(POLA1) |
| NDUFAF6 | 137682 | Homo sapiens | NADH:ubiquinone oxidoreductase complex assembly factor 6(N |
| MYO5B | 4645 | Homo sapiens | myosin VB(MYO5B) |
| CDC16 | 8881 | Homo sapiens | cell division cycle 16(CDC16) |
| ECHDC1 | 55862 | Homo sapiens | ethylmalonyl-CoA decarboxylase 1(ECHDC1) |
| NDUFAF2 | 91942 | Homo sapiens | NADH:ubiquinone oxidoreductase complex assembly factor 2(N |
| RANBP10 | 57610 | Homo sapiens | RAN binding protein 10(RANBP10) |

| | | | |
|---|---|---|---|
| FBXL3 | 26224 | Homo sapiens | F-box and leucine rich repeat protein 3(FBXL3) |
| GRB2 | 2885 | Homo sapiens | growth factor receptor bound protein 2(GRB2) |
| GRAP | 10750 | Homo sapiens | GRB2 related adaptor protein(GRAP) |
| IGSF9B | 22997 | Homo sapiens | immunoglobulin superfamily member 9B(IGSF9B) |
| FBXL6 | 26233 | Homo sapiens | F-box and leucine rich repeat protein 6(FBXL6) |
| WFDC8 | 90199 | Homo sapiens | WAP four-disulfide core domain 8(WFDC8) |
| HERPUD2 | 64224 | Homo sapiens | HERPUD family member 2(HERPUD2) |
| SPAG17 | 200162 | Homo sapiens | sperm associated antigen 17(SPAG17) |
| HIBADH | 11112 | Homo sapiens | 3-hydroxyisobutyrate dehydrogenase(HIBADH) |
| GRIK5 | 2901 | Homo sapiens | glutamate ionotropic receptor kainate type subunit 5(GRIK5) |
| GRIK4 | 2900 | Homo sapiens | glutamate ionotropic receptor kainate type subunit 4(GRIK4) |
| GRIK1 | 2897 | Homo sapiens | glutamate ionotropic receptor kainate type subunit 1(GRIK1) |
| CROT | 54677 | Homo sapiens | carnitine O-octanoyltransferase(CROT) |
| NAT14 | 57106 | Homo sapiens | N-acetyltransferase 14 (putative)(NAT14) |
| B3GNTL1 | 146712 | Homo sapiens | UDP-GlcNAc:betaGal beta-1,3-N-acetylglucosaminyltransferase |
| MEDAG | 84935 | Homo sapiens | mesenteric estrogen dependent adipogenesis(MEDAG) |
| AKAP13 | 11214 | Homo sapiens | A-kinase anchoring protein 13(AKAP13) |
| PPP3CA | 5530 | Homo sapiens | protein phosphatase 3 catalytic subunit alpha(PPP3CA) |
| KCNT1 | 57582 | Homo sapiens | potassium sodium-activated channel subfamily T member 1(KC |
| AKAP11 | 11215 | Homo sapiens | A-kinase anchoring protein 11(AKAP11) |
| KCNT2 | 343450 | Homo sapiens | potassium sodium-activated channel subfamily T member 2(KC |
| SUFU | 51684 | Homo sapiens | SUFU negative regulator of hedgehog signaling(SUFU) |
| BBS9 | 27241 | Homo sapiens | Bardet-Biedl syndrome 9(BBS9) |
| BBS7 | 55212 | Homo sapiens | Bardet-Biedl syndrome 7(BBS7) |
| GIT1 | 28964 | Homo sapiens | GIT ArfGAP 1(GIT1) |
| MRPS28 | 28957 | Homo sapiens | mitochondrial ribosomal protein S28(MRPS28) |
| PCYT1A | 5130 | Homo sapiens | phosphate cytidylyltransferase 1A, choline(PCYT1A) |
| PRPF40B | 25766 | Homo sapiens | pre-mRNA processing factor 40 homolog B(PRPF40B) |
| CACYBP | 27101 | Homo sapiens | calcyclin binding protein(CACYBP) |
| GAB2 | 9846 | Homo sapiens | GRB2 associated binding protein 2(GAB2) |
| SYN2 | 6854 | Homo sapiens | synapsin II(SYN2) |
| KIF6 | 221458 | Homo sapiens | kinesin family member 6(KIF6) |
| DUS2 | 54920 | Homo sapiens | dihydrouridine synthase 2(DUS2) |
| KIN | 22944 | Homo sapiens | Kin17 DNA and RNA binding protein(KIN) |
| KCNU1 | 157855 | Homo sapiens | potassium calcium-activated channel subfamily U member 1(KC |
| KIF9 | 64147 | Homo sapiens | kinesin family member 9(KIF9) |
| ARHGEF9 | 23229 | Homo sapiens | Cdc42 guanine nucleotide exchange factor 9(ARHGEF9) |
| TOX2 | 84969 | Homo sapiens | TOX high mobility group box family member 2(TOX2) |
| GAN | 8139 | Homo sapiens | gigaxonin(GAN) |
| DEPDC1 | 55635 | Homo sapiens | DEP domain containing 1(DEPDC1) |
| SDE2 | 163859 | Homo sapiens | SDE2 telomere maintenance homolog(SDE2) |
| ARHGEF4 | 50649 | Homo sapiens | Rho guanine nucleotide exchange factor 4(ARHGEF4) |
| MICU3 | 286097 | Homo sapiens | mitochondrial calcium uptake family member 3(MICU3) |
| TLCD2 | 727910 | Homo sapiens | TLC domain containing 2(TLCD2) |
| FAM193B | 54540 | Homo sapiens | family with sequence similarity 193 member B(FAM193B) |

| | | | |
|---|---|---|---|
| MICU2 | 221154 | Homo sapiens | mitochondrial calcium uptake 2(MICU2) |
| ARHGEF2 | 9181 | Homo sapiens | Rho/Rac guanine nucleotide exchange factor 2(ARHGEF2) |
| MICU1 | 10367 | Homo sapiens | mitochondrial calcium uptake 1(MICU1) |
| ARHGEF7 | 8874 | Homo sapiens | Rho guanine nucleotide exchange factor 7(ARHGEF7) |
| FJX1 | 24147 | Homo sapiens | four-jointed box kinase 1(FJX1) |
| KIZ | 55857 | Homo sapiens | kizuna centrosomal protein(KIZ) |
| ARHGEF5 | 7984 | Homo sapiens | Rho guanine nucleotide exchange factor 5(ARHGEF5) |
| MRPS35 | 60488 | Homo sapiens | mitochondrial ribosomal protein S35(MRPS35) |
| GNAI3 | 2773 | Homo sapiens | G protein subunit alpha i3(GNAI3) |
| RELB | 5971 | Homo sapiens | RELB proto-oncogene, NF-kB subunit(RELB) |
| GNAI2 | 2771 | Homo sapiens | G protein subunit alpha i2(GNAI2) |
| REM1 | 28954 | Homo sapiens | RRAD and GEM like GTPase 1(REM1) |
| RELN | 5649 | Homo sapiens | reelin(RELN) |
| PDZD11 | 51248 | Homo sapiens | PDZ domain containing 11(PDZD11) |
| PCNT | 5116 | Homo sapiens | pericentrin(PCNT) |
| EPS15 | 2060 | Homo sapiens | epidermal growth factor receptor pathway substrate 15(EPS15) |
| CAP2 | 10486 | Homo sapiens | cyclase associated actin cytoskeleton regulatory protein 2(CAP2 |
| OFCC1 | 266553 | Homo sapiens | orofacial cleft 1 candidate 1(OFCC1) |
| GABBR2 | 9568 | Homo sapiens | gamma-aminobutyric acid type B receptor subunit 2(GABBR2) |
| DYNC1H1 | 1778 | Homo sapiens | dynein cytoplasmic 1 heavy chain 1(DYNC1H1) |
| GABBR1 | 2550 | Homo sapiens | gamma-aminobutyric acid type B receptor subunit 1(GABBR1) |
| MON2 | 23041 | Homo sapiens | MON2 homolog, regulator of endosome-to-Golgi trafficking(MOI |
| TTC1 | 7265 | Homo sapiens | tetratricopeptide repeat domain 1(TTC1) |
| GAD2 | 2572 | Homo sapiens | glutamate decarboxylase 2(GAD2) |
| KLHL4 | 56062 | Homo sapiens | kelch like family member 4(KLHL4) |
| GRIN1 | 2902 | Homo sapiens | glutamate ionotropic receptor NMDA type subunit 1(GRIN1) |
| SGSM1 | 129049 | Homo sapiens | small G protein signaling modulator 1(SGSM1) |
| RP1L1 | 94137 | Homo sapiens | RP1 like 1(RP1L1) |
| NUDT9 | 53343 | Homo sapiens | nudix hydrolase 9(NUDT9) |
| VMP1 | 81671 | Homo sapiens | vacuole membrane protein 1(VMP1) |
| SH3RF3 | 344558 | Homo sapiens | SH3 domain containing ring finger 3(SH3RF3) |
| PDCL2 | 132954 | Homo sapiens | phosducin like 2(PDCL2) |
| NUDT8 | 254552 | Homo sapiens | nudix hydrolase 8(NUDT8) |
| NOL4L | 140688 | Homo sapiens | nucleolar protein 4 like(NOL4L) |
| TTC6 | 319089 | Homo sapiens | tetratricopeptide repeat domain 6(TTC6) |
| KLHL5 | 51088 | Homo sapiens | kelch like family member 5(KLHL5) |
| KCNS2 | 3788 | Homo sapiens | potassium voltage-gated channel modifier subfamily S member |
| TCEA3 | 6920 | Homo sapiens | transcription elongation factor A3(TCEA3) |
| TACC3 | 10460 | Homo sapiens | transforming acidic coiled-coil containing protein 3(TACC3) |
| PNPLA6 | 10908 | Homo sapiens | patatin like phospholipase domain containing 6(PNPLA6) |
| OTUD5 | 55593 | Homo sapiens | OTU deubiquitinase 5(OTUD5) |
| NUP188 | 23511 | Homo sapiens | nucleoporin 188(NUP188) |
| ICE2 | 79664 | Homo sapiens | interactor of little elongation complex ELL subunit 2(ICE2) |
| DENND5B | 160518 | Homo sapiens | DENN domain containing 5B(DENND5B) |
| ZFAND3 | 60685 | Homo sapiens | zinc finger AN1-type containing 3(ZFAND3) |

| | | | |
|---|---|---|---|
| ZFAND6 | 54469 | Homo sapiens | zinc finger AN1-type containing 6(ZFAND6) |
| RPP30 | 10556 | Homo sapiens | ribonuclease P/MRP subunit p30(RPP30) |
| PRDM5 | 11107 | Homo sapiens | PR/SET domain 5(PRDM5) |
| PCDH11X | 27328 | Homo sapiens | protocadherin 11 X-linked(PCDH11X) |
| INPPL1 | 3636 | Homo sapiens | inositol polyphosphate phosphatase like 1(INPPL1) |
| LOXL4 | 84171 | Homo sapiens | lysyl oxidase like 4(LOXL4) |
| UBE3B | 89910 | Homo sapiens | ubiquitin protein ligase E3B(UBE3B) |
| DUX4 | 100288687 | Homo sapiens | double homeobox 4(DUX4) |
| CARF | 79800 | Homo sapiens | calcium responsive transcription factor(CARF) |
| CELSR3 | 1951 | Homo sapiens | cadherin EGF LAG seven-pass G-type receptor 3(CELSR3) |
| PLB1 | 151056 | Homo sapiens | phospholipase B1(PLB1) |
| TGM1 | 7051 | Homo sapiens | transglutaminase 1(TGM1) |
| GRIP1 | 23426 | Homo sapiens | glutamate receptor interacting protein 1(GRIP1) |
| GNPTAB | 79158 | Homo sapiens | N-acetylglucosamine-1-phosphate transferase subunits alpha ar |
| TBC1D31 | 93594 | Homo sapiens | TBC1 domain family member 31(TBC1D31) |
| PPIP5K1 | 9677 | Homo sapiens | diphosphoinositol pentakisphosphate kinase 1(PPIP5K1) |
| TBC1D30 | 23329 | Homo sapiens | TBC1 domain family member 30(TBC1D30) |
| TEKT3 | 64518 | Homo sapiens | tektin 3(TEKT3) |
| DENND5A | 23258 | Homo sapiens | DENN domain containing 5A(DENND5A) |
| ENPP3 | 5169 | Homo sapiens | ectonucleotide pyrophosphatase/phosphodiesterase 3(ENPP3) |
| ENPP6 | 133121 | Homo sapiens | ectonucleotide pyrophosphatase/phosphodiesterase 6(ENPP6) |
| TPTE2 | 93492 | Homo sapiens | transmembrane phosphoinositide 3-phosphatase and tensin hor |
| CAST | 831 | Homo sapiens | calpastatin(CAST) |
| TMOD1 | 7111 | Homo sapiens | tropomodulin 1(TMOD1) |
| CASR | 846 | Homo sapiens | calcium sensing receptor(CASR) |
| MORC3 | 23515 | Homo sapiens | MORC family CW-type zinc finger 3(MORC3) |
| FBXW10 | 10517 | Homo sapiens | F-box and WD repeat domain containing 10(FBXW10) |
| FBXW11 | 23291 | Homo sapiens | F-box and WD repeat domain containing 11(FBXW11) |
| ADAM11 | 4185 | Homo sapiens | ADAM metallopeptidase domain 11(ADAM11) |
| DNAJB12 | 54788 | Homo sapiens | DnaJ heat shock protein family (Hsp40) member B12(DNAJB12 |
| UBE4A | 9354 | Homo sapiens | ubiquitination factor E4A(UBE4A) |
| ADAM10 | 102 | Homo sapiens | ADAM metallopeptidase domain 10(ADAM10) |
| CASK | 8573 | Homo sapiens | calcium/calmodulin dependent serine protein kinase(CASK) |
| KMO | 8564 | Homo sapiens | kynurenine 3-monooxygenase(KMO) |
| MUC3A | 4584 | Homo sapiens | mucin 3A, cell surface associated(MUC3A) |
| GRIP2 | 80852 | Homo sapiens | glutamate receptor interacting protein 2(GRIP2) |
| ADAM19 | 8728 | Homo sapiens | ADAM metallopeptidase domain 19(ADAM19) |
| SPOCD1 | 90853 | Homo sapiens | SPOC domain containing 1(SPOCD1) |
| SHQ1 | 55164 | Homo sapiens | SHQ1, H/ACA ribonucleoprotein assembly factor(SHQ1) |
| TBC1D20 | 128637 | Homo sapiens | TBC1 domain family member 20(TBC1D20) |
| MAP7D3 | 79649 | Homo sapiens | MAP7 domain containing 3(MAP7D3) |
| SLFN14 | 342618 | Homo sapiens | schlafen family member 14(SLFN14) |
| EHBP1L1 | 254102 | Homo sapiens | EH domain binding protein 1 like 1(EHBP1L1) |
| ADAM12 | 8038 | Homo sapiens | ADAM metallopeptidase domain 12(ADAM12) |
| SIDT1 | 54847 | Homo sapiens | SID1 transmembrane family member 1(SIDT1) |

| | | | |
|---|---|---|---|
| ANKS6 | 203286 | Homo sapiens | ankyrin repeat and sterile alpha motif domain containing 6(ANK |
| WDFY3 | 23001 | Homo sapiens | WD repeat and FYVE domain containing 3(WDFY3) |
| RAF1 | 5894 | Homo sapiens | Raf-1 proto-oncogene, serine/threonine kinase(RAF1) |
| CARS2 | 79587 | Homo sapiens | cysteinyl-tRNA synthetase 2, mitochondrial(CARS2) |
| SIDT2 | 51092 | Homo sapiens | SID1 transmembrane family member 2(SIDT2) |
| TSG101 | 7251 | Homo sapiens | tumor susceptibility 101(TSG101) |
| NNMT | 4837 | Homo sapiens | nicotinamide N-methyltransferase(NNMT) |
| DDX4 | 54514 | Homo sapiens | DEAD-box helicase 4(DDX4) |
| PKDCC | 91461 | Homo sapiens | protein kinase domain containing, cytoplasmic(PKDCC) |
| CACNA1B | 774 | Homo sapiens | calcium voltage-gated channel subunit alpha1 B(CACNA1B) |
| PLD4 | 122618 | Homo sapiens | phospholipase D family member 4(PLD4) |
| CACNA1A | 773 | Homo sapiens | calcium voltage-gated channel subunit alpha1 A(CACNA1A) |
| PLD6 | 201164 | Homo sapiens | phospholipase D family member 6(PLD6) |
| CACNA1C | 775 | Homo sapiens | calcium voltage-gated channel subunit alpha1 C(CACNA1C) |
| PLD1 | 5337 | Homo sapiens | phospholipase D1(PLD1) |
| TMEM240 | 339453 | Homo sapiens | transmembrane protein 240(TMEM240) |
| NUP160 | 23279 | Homo sapiens | nucleoporin 160(NUP160) |
| PLD3 | 23646 | Homo sapiens | phospholipase D family member 3(PLD3) |
| CACNA1G | 8913 | Homo sapiens | calcium voltage-gated channel subunit alpha1 G(CACNA1G) |
| TIMM50 | 92609 | Homo sapiens | translocase of inner mitochondrial membrane 50(TIMM50) |
| CACNA1I | 8911 | Homo sapiens | calcium voltage-gated channel subunit alpha1 I(CACNA1I) |
| ADGRG1 | 9289 | Homo sapiens | adhesion G protein-coupled receptor G1(ADGRG1) |
| TMEM245 | 23731 | Homo sapiens | transmembrane protein 245(TMEM245) |
| ADGRG2 | 10149 | Homo sapiens | adhesion G protein-coupled receptor G2(ADGRG2) |
| IRAK2 | 3656 | Homo sapiens | interleukin 1 receptor associated kinase 2(IRAK2) |
| DHRS7B | 25979 | Homo sapiens | dehydrogenase/reductase 7B(DHRS7B) |
| DHRS7C | 201140 | Homo sapiens | dehydrogenase/reductase 7C(DHRS7C) |
| GRK6 | 2870 | Homo sapiens | G protein-coupled receptor kinase 6(GRK6) |
| RHPN1 | 114822 | Homo sapiens | rhophilin Rho GTPase binding protein 1(RHPN1) |
| QSER1 | 79832 | Homo sapiens | glutamine and serine rich 1(QSER1) |
| TBC1D16 | 125058 | Homo sapiens | TBC1 domain family member 16(TBC1D16) |
| CNTLN | 54875 | Homo sapiens | centlein(CNTLN) |
| UBE2F | 140739 | Homo sapiens | ubiquitin conjugating enzyme E2 F (putative)(UBE2F) |
| UBE2H | 7328 | Homo sapiens | ubiquitin conjugating enzyme E2 H(UBE2H) |
| UBE2I | 7329 | Homo sapiens | ubiquitin conjugating enzyme E2 I(UBE2I) |
| MUC16 | 94025 | Homo sapiens | mucin 16, cell surface associated(MUC16) |
| CNTN6 | 27255 | Homo sapiens | contactin 6(CNTN6) |
| CAV1 | 857 | Homo sapiens | caveolin 1(CAV1) |
| DENND4C | 55667 | Homo sapiens | DENN domain containing 4C(DENND4C) |
| HOOK3 | 84376 | Homo sapiens | hook microtubule tethering protein 3(HOOK3) |
| IGF2R | 3482 | Homo sapiens | insulin like growth factor 2 receptor(IGF2R) |
| SDK1 | 221935 | Homo sapiens | sidekick cell adhesion molecule 1(SDK1) |
| NAT1 | 9 | Homo sapiens | N-acetyltransferase 1(NAT1) |
| FAM151B | 167555 | Homo sapiens | family with sequence similarity 151 member B(FAM151B) |
| GNB1 | 2782 | Homo sapiens | G protein subunit beta 1(GNB1) |

| | | | |
|---|---|---|---|
| MANSC4 | 100287284 | Homo sapiens | MANSC domain containing 4(MANSC4) |
| MKRN2 | 23609 | Homo sapiens | makorin ring finger protein 2(MKRN2) |
| UBA3 | 9039 | Homo sapiens | ubiquitin like modifier activating enzyme 3(UBA3) |
| CNTN1 | 1272 | Homo sapiens | contactin 1(CNTN1) |
| CNTN2 | 6900 | Homo sapiens | contactin 2(CNTN2) |
| PYROXD2 | 84795 | Homo sapiens | pyridine nucleotide-disulphide oxidoreductase domain 2(PYROXI |
| AMOTL1 | 154810 | Homo sapiens | angiomotin like 1(AMOTL1) |
| TCF3 | 6929 | Homo sapiens | transcription factor 3(TCF3) |
| DEGS2 | 123099 | Homo sapiens | delta 4-desaturase, sphingolipid 2(DEGS2) |
| MXD1 | 4084 | Homo sapiens | MAX dimerization protein 1(MXD1) |
| SDK2 | 54549 | Homo sapiens | sidekick cell adhesion molecule 2(SDK2) |
| MXD4 | 10608 | Homo sapiens | MAX dimerization protein 4(MXD4) |
| CUEDC2 | 79004 | Homo sapiens | CUE domain containing 2(CUEDC2) |
| TTC23 | 64927 | Homo sapiens | tetratricopeptide repeat domain 23(TTC23) |
| PINX1 | 54984 | Homo sapiens | PIN2 (TERF1) interacting telomerase inhibitor 1(PINX1) |
| LDB3 | 11155 | Homo sapiens | LIM domain binding 3(LDB3) |
| RND2 | 8153 | Homo sapiens | Rho family GTPase 2(RND2) |
| GRM1 | 2911 | Homo sapiens | glutamate metabotropic receptor 1(GRM1) |
| CABIN1 | 23523 | Homo sapiens | calcineurin binding protein 1(CABIN1) |
| TMEM220 | 388335 | Homo sapiens | transmembrane protein 220(TMEM220) |
| ADGRE1 | 2015 | Homo sapiens | adhesion G protein-coupled receptor E1(ADGRE1) |
| IFT52 | 51098 | Homo sapiens | intraflagellar transport 52(IFT52) |
| GRM5 | 2915 | Homo sapiens | glutamate metabotropic receptor 5(GRM5) |
| GRM4 | 2914 | Homo sapiens | glutamate metabotropic receptor 4(GRM4) |
| CA2 | 760 | Homo sapiens | carbonic anhydrase 2(CA2) |
| RASGEF1B | 153020 | Homo sapiens | RasGEF domain family member 1B(RASGEF1B) |
| RB1CC1 | 9821 | Homo sapiens | RB1 inducible coiled-coil 1(RB1CC1) |
| CA4 | 762 | Homo sapiens | carbonic anhydrase 4(CA4) |
| MAN1A1 | 4121 | Homo sapiens | mannosidase alpha class 1A member 1(MAN1A1) |
| MYBL2 | 4605 | Homo sapiens | MYB proto-oncogene like 2(MYBL2) |
| SNTA1 | 6640 | Homo sapiens | syntrophin alpha 1(SNTA1) |
| MKKS | 8195 | Homo sapiens | MKKS centrosomal shuttling protein(MKKS) |
| SUN1 | 23353 | Homo sapiens | Sad1 and UNC84 domain containing 1(SUN1) |
| TBP | 6908 | Homo sapiens | TATA-box binding protein(TBP) |
| NUP133 | 55746 | Homo sapiens | nucleoporin 133(NUP133) |
| FIGLA | 344018 | Homo sapiens | folliculogenesis specific bHLH transcription factor(FIGLA) |
| DCAKD | 79877 | Homo sapiens | dephospho-CoA kinase domain containing(DCAKD) |
| NAV2 | 89797 | Homo sapiens | neuron navigator 2(NAV2) |
| QRSL1 | 55278 | Homo sapiens | glutaminyl-tRNA amidotransferase subunit QRSL1(QRSL1) |
| SPIDR | 23514 | Homo sapiens | scaffold protein involved in DNA repair(SPIDR) |
| ETV5 | 2119 | Homo sapiens | ETS variant transcription factor 5(ETV5) |
| EPN2 | 22905 | Homo sapiens | epsin 2(EPN2) |
| ETV6 | 2120 | Homo sapiens | ETS variant transcription factor 6(ETV6) |
| CYP26C1 | 340665 | Homo sapiens | cytochrome P450 family 26 subfamily C member 1(CYP26C1) |
| NAV3 | 89795 | Homo sapiens | neuron navigator 3(NAV3) |

| | | | |
|---|---|---|---|
| SETBP1 | 26040 | Homo sapiens | SET binding protein 1(SETBP1) |
| GFOD2 | 81577 | Homo sapiens | Gfo/Idh/MocA-like oxidoreductase domain containing 2(GFOD2) |
| ZMYND10 | 51364 | Homo sapiens | zinc finger MYND-type containing 10(ZMYND10) |
| CLRN3 | 119467 | Homo sapiens | clarin 3(CLRN3) |
| ATP6V1B1 | 525 | Homo sapiens | ATPase H+ transporting V1 subunit B1(ATP6V1B1) |
| DNAH1 | 25981 | Homo sapiens | dynein axonemal heavy chain 1(DNAH1) |
| RRN3 | 54700 | Homo sapiens | RRN3 homolog, RNA polymerase I transcription factor(RRN3) |
| GALT | 2592 | Homo sapiens | galactose-1-phosphate uridylyltransferase(GALT) |
| DNAH2 | 146754 | Homo sapiens | dynein axonemal heavy chain 2(DNAH2) |
| ASB13 | 79754 | Homo sapiens | ankyrin repeat and SOCS box containing 13(ASB13) |
| DNAH9 | 1770 | Homo sapiens | dynein axonemal heavy chain 9(DNAH9) |
| ASB11 | 140456 | Homo sapiens | ankyrin repeat and SOCS box containing 11(ASB11) |
| ASAP1 | 50807 | Homo sapiens | ArfGAP with SH3 domain, ankyrin repeat and PH domain 1(ASAI |
| CBL | 867 | Homo sapiens | Cbl proto-oncogene(CBL) |
| RNF4 | 6047 | Homo sapiens | ring finger protein 4(RNF4) |
| ADD3 | 120 | Homo sapiens | adducin 3(ADD3) |
| AGPAT2 | 10555 | Homo sapiens | 1-acylglycerol-3-phosphate O-acyltransferase 2(AGPAT2) |
| ASB18 | 401036 | Homo sapiens | ankyrin repeat and SOCS box containing 18(ASB18) |
| ADD2 | 119 | Homo sapiens | adducin 2(ADD2) |
| GK5 | 256356 | Homo sapiens | glycerol kinase 5(GK5) |
| IL1RL1 | 9173 | Homo sapiens | interleukin 1 receptor like 1(IL1RL1) |
| MTA1 | 9112 | Homo sapiens | metastasis associated 1(MTA1) |
| GNG2 | 54331 | Homo sapiens | G protein subunit gamma 2(GNG2) |
| NWD2 | 57495 | Homo sapiens | NACHT and WD repeat domain containing 2(NWD2) |
| TBX22 | 50945 | Homo sapiens | T-box transcription factor 22(TBX22) |
| OXCT1 | 5019 | Homo sapiens | 3-oxoacid CoA-transferase 1(OXCT1) |
| SPOCK2 | 9806 | Homo sapiens | SPARC (osteonectin), cwcv and kazal like domains proteoglycar |
| TRA2A | 29896 | Homo sapiens | transformer 2 alpha homolog(TRA2A) |
| SPOCK3 | 50859 | Homo sapiens | SPARC (osteonectin), cwcv and kazal like domains proteoglycar |
| DROSHA | 29102 | Homo sapiens | drosha ribonuclease III(DROSHA) |
| SLIT3 | 6586 | Homo sapiens | slit guidance ligand 3(SLIT3) |
| SLIT2 | 9353 | Homo sapiens | slit guidance ligand 2(SLIT2) |
| CRYM | 1428 | Homo sapiens | crystallin mu(CRYM) |
| MTA3 | 57504 | Homo sapiens | metastasis associated 1 family member 3(MTA3) |
| DNAI1 | 27019 | Homo sapiens | dynein axonemal intermediate chain 1(DNAI1) |
| TRDMT1 | 1787 | Homo sapiens | tRNA aspartic acid methyltransferase 1(TRDMT1) |
| TTC12 | 54970 | Homo sapiens | tetratricopeptide repeat domain 12(TTC12) |
| NIPSNAP3A | 25934 | Homo sapiens | nipsnap homolog 3A(NIPSNAP3A) |
| EYA2 | 2139 | Homo sapiens | EYA transcriptional coactivator and phosphatase 2(EYA2) |
| BNIP3 | 664 | Homo sapiens | BCL2 interacting protein 3(BNIP3) |
| USF1 | 7391 | Homo sapiens | upstream transcription factor 1(USF1) |
| WARS2 | 10352 | Homo sapiens | tryptophanyl tRNA synthetase 2, mitochondrial(WARS2) |
| HS3ST2 | 9956 | Homo sapiens | heparan sulfate-glucosamine 3-sulfotransferase 2(HS3ST2) |
| GALNS | 2588 | Homo sapiens | galactosamine (N-acetyl)-6-sulfatase(GALNS) |
| POLE4 | 56655 | Homo sapiens | DNA polymerase epsilon 4, accessory subunit(POLE4) |

| | | | |
|---|---|---|---|
| TBX15 | 6913 | Homo sapiens | T-box transcription factor 15(TBX15) |
| MYO15A | 51168 | Homo sapiens | myosin XVA(MYO15A) |
| TNIP3 | 79931 | Homo sapiens | TNFAIP3 interacting protein 3(TNIP3) |
| CARM1 | 10498 | Homo sapiens | coactivator associated arginine methyltransferase 1(CARM1) |
| POLE3 | 54107 | Homo sapiens | DNA polymerase epsilon 3, accessory subunit(POLE3) |
| ERCC5 | 2073 | Homo sapiens | ERCC excision repair 5, endonuclease(ERCC5) |
| ERCC6 | 2074 | Homo sapiens | ERCC excision repair 6, chromatin remodeling factor(ERCC6) |
| NRDE2 | 55051 | Homo sapiens | NRDE-2, necessary for RNA interference, domain containing(NR |
| SNTB1 | 6641 | Homo sapiens | syntrophin beta 1(SNTB1) |
| SNTB2 | 6645 | Homo sapiens | syntrophin beta 2(SNTB2) |
| IPO13 | 9670 | Homo sapiens | importin 13(IPO13) |
| CDA | 978 | Homo sapiens | cytidine deaminase(CDA) |
| IPO11 | 51194 | Homo sapiens | importin 11(IPO11) |
| ANKLE1 | 126549 | Homo sapiens | ankyrin repeat and LEM domain containing 1(ANKLE1) |
| TES | 26136 | Homo sapiens | testin LIM domain protein(TES) |
| FRMPD1 | 22844 | Homo sapiens | FERM and PDZ domain containing 1(FRMPD1) |
| WIPF3 | 644150 | Homo sapiens | WAS/WASL interacting protein family member 3(WIPF3) |
| XYLT2 | 64132 | Homo sapiens | xylosyltransferase 2(XYLT2) |
| GPATCH2 | 55105 | Homo sapiens | G-patch domain containing 2(GPATCH2) |
| GPATCH1 | 55094 | Homo sapiens | G-patch domain containing 1(GPATCH1) |
| HNRNPR | 10236 | Homo sapiens | heterogeneous nuclear ribonucleoprotein R(HNRNPR) |
| SMC4 | 10051 | Homo sapiens | structural maintenance of chromosomes 4(SMC4) |
| SLC4A4 | 8671 | Homo sapiens | solute carrier family 4 member 4(SLC4A4) |
| SLC4A5 | 57835 | Homo sapiens | solute carrier family 4 member 5(SLC4A5) |
| FGFRL1 | 53834 | Homo sapiens | fibroblast growth factor receptor like 1(FGFRL1) |
| WDR88 | 126248 | Homo sapiens | WD repeat domain 88(WDR88) |
| ALCAM | 214 | Homo sapiens | activated leukocyte cell adhesion molecule(ALCAM) |
| EVA1A | 84141 | Homo sapiens | eva-1 homolog A, regulator of programmed cell death(EVA1A) |
| MSANTD1 | 345222 | Homo sapiens | Myb/SANT DNA binding domain containing 1(MSANTD1) |
| GJA4 | 2701 | Homo sapiens | gap junction protein alpha 4(GJA4) |
| PXDC1 | 221749 | Homo sapiens | PX domain containing 1(PXDC1) |
| CEP97 | 79598 | Homo sapiens | centrosomal protein 97(CEP97) |
| NCKAP5 | 344148 | Homo sapiens | NCK associated protein 5(NCKAP5) |
| CDK5RAP2 | 55755 | Homo sapiens | CDK5 regulatory subunit associated protein 2(CDK5RAP2) |
| DDX17 | 10521 | Homo sapiens | DEAD-box helicase 17(DDX17) |
| ADGB | 79747 | Homo sapiens | androglobin(ADGB) |
| MBNL1 | 4154 | Homo sapiens | muscleblind like splicing regulator 1(MBNL1) |
| MBNL2 | 10150 | Homo sapiens | muscleblind like splicing regulator 2(MBNL2) |
| PRKCB | 5579 | Homo sapiens | protein kinase C beta(PRKCB) |
| DNAAF1 | 123872 | Homo sapiens | dynein axonemal assembly factor 1(DNAAF1) |
| ARMC9 | 80210 | Homo sapiens | armadillo repeat containing 9(ARMC9) |
| REV1 | 51455 | Homo sapiens | REV1 DNA directed polymerase(REV1) |
| FNDC3B | 64778 | Homo sapiens | fibronectin type III domain containing 3B(FNDC3B) |
| AFAP1 | 60312 | Homo sapiens | actin filament associated protein 1(AFAP1) |
| DDX10 | 1662 | Homo sapiens | DEAD-box helicase 10(DDX10) |

| | | | |
|---|---|---|---|
| ARMC8 | 25852 | Homo sapiens | armadillo repeat containing 8(ARMC8) |
| GANC | 2595 | Homo sapiens | glucosidase alpha, neutral C(GANC) |
| GABRG1 | 2565 | Homo sapiens | gamma-aminobutyric acid type A receptor subunit gamma1(GA |
| NUP93 | 9688 | Homo sapiens | nucleoporin 93(NUP93) |
| ARMC2 | 84071 | Homo sapiens | armadillo repeat containing 2(ARMC2) |
| DEPDC1B | 55789 | Homo sapiens | DEP domain containing 1B(DEPDC1B) |
| ADCY9 | 115 | Homo sapiens | adenylate cyclase 9(ADCY9) |
| DAAM1 | 23002 | Homo sapiens | dishevelled associated activator of morphogenesis 1(DAAM1) |
| PRKD2 | 25865 | Homo sapiens | protein kinase D2(PRKD2) |
| PRKCQ | 5588 | Homo sapiens | protein kinase C theta(PRKCQ) |
| SRFBP1 | 153443 | Homo sapiens | serum response factor binding protein 1(SRFBP1) |
| PRKD1 | 5587 | Homo sapiens | protein kinase D1(PRKD1) |
| ASTN2 | 23245 | Homo sapiens | astrotactin 2(ASTN2) |
| FREM2 | 341640 | Homo sapiens | FRAS1 related extracellular matrix 2(FREM2) |
| ASTN1 | 460 | Homo sapiens | astrotactin 1(ASTN1) |
| PDXDC1 | 23042 | Homo sapiens | pyridoxal dependent decarboxylase domain containing 1(PDXD |
| PRKDC | 5591 | Homo sapiens | protein kinase, DNA-activated, catalytic subunit(PRKDC) |
| PRDM15 | 63977 | Homo sapiens | PR/SET domain 15(PRDM15) |
| BCL10 | 8915 | Homo sapiens | BCL10 immune signaling adaptor(BCL10) |
| WDR62 | 284403 | Homo sapiens | WD repeat domain 62(WDR62) |
| PRKCZ | 5590 | Homo sapiens | protein kinase C zeta(PRKCZ) |
| ADH4 | 127 | Homo sapiens | alcohol dehydrogenase 4 (class II), pi polypeptide(ADH4) |
| NUP85 | 79902 | Homo sapiens | nucleoporin 85(NUP85) |
| VPS11 | 55823 | Homo sapiens | VPS11 core subunit of CORVET and HOPS complexes(VPS11) |
| ALX4 | 60529 | Homo sapiens | ALX homeobox 4(ALX4) |
| MCMBP | 79892 | Homo sapiens | minichromosome maintenance complex binding protein(MCMBP |
| NCKAP1L | 3071 | Homo sapiens | NCK associated protein 1 like(NCKAP1L) |
| CEP76 | 79959 | Homo sapiens | centrosomal protein 76(CEP76) |
| FDFT1 | 2222 | Homo sapiens | farnesyl-diphosphate farnesyltransferase 1(FDFT1) |
| ST3GAL3 | 6487 | Homo sapiens | ST3 beta-galactoside alpha-2,3-sialyltransferase 3(ST3GAL3) |
| PROM2 | 150696 | Homo sapiens | prominin 2(PROM2) |
| LHPP | 64077 | Homo sapiens | phospholysine phosphohistidine inorganic pyrophosphate phosph |
| NAA50 | 80218 | Homo sapiens | N-alpha-acetyltransferase 50, NatE catalytic subunit(NAA50) |
| BTF3 | 689 | Homo sapiens | basic transcription factor 3(BTF3) |
| PLK4 | 10733 | Homo sapiens | polo like kinase 4(PLK4) |
| WDR59 | 79726 | Homo sapiens | WD repeat domain 59(WDR59) |
| AFAP1L1 | 134265 | Homo sapiens | actin filament associated protein 1 like 1(AFAP1L1) |
| DMPK | 1760 | Homo sapiens | DM1 protein kinase(DMPK) |
| SETD1B | 23067 | Homo sapiens | SET domain containing 1B, histone lysine methyltransferase(SE |
| STARD10 | 10809 | Homo sapiens | StAR related lipid transfer domain containing 10(STARD10) |
| PLK1 | 5347 | Homo sapiens | polo like kinase 1(PLK1) |
| POU6F2 | 11281 | Homo sapiens | POU class 6 homeobox 2(POU6F2) |
| NFATC3 | 4775 | Homo sapiens | nuclear factor of activated T cells 3(NFATC3) |
| RMDN2 | 151393 | Homo sapiens | regulator of microtubule dynamics 2(RMDN2) |
| MEX3D | 399664 | Homo sapiens | mex-3 RNA binding family member D(MEX3D) |

| | | | |
|---|---|---|---|
| MEX3C | 51320 | Homo sapiens | mex-3 RNA binding family member C(MEX3C) |
| EHD2 | 30846 | Homo sapiens | EH domain containing 2(EHD2) |
| HNRNPM | 4670 | Homo sapiens | heterogeneous nuclear ribonucleoprotein M(HNRNPM) |
| ARL2BP | 23568 | Homo sapiens | ADP ribosylation factor like GTPase 2 binding protein(ARL2BP) |
| REC114 | 283677 | Homo sapiens | REC114 meiotic recombination protein(REC114) |
| CEP89 | 84902 | Homo sapiens | centrosomal protein 89(CEP89) |
| PHF3 | 23469 | Homo sapiens | PHD finger protein 3(PHF3) |
| WDR49 | 151790 | Homo sapiens | WD repeat domain 49(WDR49) |
| SETD4 | 54093 | Homo sapiens | SET domain containing 4(SETD4) |
| PHF2 | 5253 | Homo sapiens | PHD finger protein 2(PHF2) |
| WDR48 | 57599 | Homo sapiens | WD repeat domain 48(WDR48) |
| PHF20 | 51230 | Homo sapiens | PHD finger protein 20(PHF20) |
| DENND1A | 57706 | Homo sapiens | DENN domain containing 1A(DENND1A) |
| ARPC1B | 10095 | Homo sapiens | actin related protein 2/3 complex subunit 1B(ARPC1B) |
| SERPINA10 | 51156 | Homo sapiens | serpin family A member 10(SERPINA10) |
| CHM | 1121 | Homo sapiens | CHM Rab escort protein(CHM) |
| GPT | 2875 | Homo sapiens | glutamic--pyruvic transaminase(GPT) |
| STON2 | 85439 | Homo sapiens | stonin 2(STON2) |
| WDR43 | 23160 | Homo sapiens | WD repeat domain 43(WDR43) |
| FARS2 | 10667 | Homo sapiens | phenylalanyl-tRNA synthetase 2, mitochondrial(FARS2) |
| ADAMTSL1 | 92949 | Homo sapiens | ADAMTS like 1(ADAMTSL1) |
| ADAMTSL4 | 54507 | Homo sapiens | ADAMTS like 4(ADAMTSL4) |
| AIFM2 | 84883 | Homo sapiens | apoptosis inducing factor mitochondria associated 2(AIFM2) |
| ADAMTSL3 | 57188 | Homo sapiens | ADAMTS like 3(ADAMTSL3) |
| PLEKHN1 | 84069 | Homo sapiens | pleckstrin homology domain containing N1(PLEKHN1) |
| RRP12 | 23223 | Homo sapiens | ribosomal RNA processing 12 homolog(RRP12) |
| TEAD1 | 7003 | Homo sapiens | TEA domain transcription factor 1(TEAD1) |
| HELLS | 3070 | Homo sapiens | helicase, lymphoid specific(HELLS) |
| WDR37 | 22884 | Homo sapiens | WD repeat domain 37(WDR37) |
| PLLP | 51090 | Homo sapiens | plasmolipin(PLLP) |
| WDR33 | 55339 | Homo sapiens | WD repeat domain 33(WDR33) |
| NCOR2 | 9612 | Homo sapiens | nuclear receptor corepressor 2(NCOR2) |
| APBB1IP | 54518 | Homo sapiens | amyloid beta precursor protein binding family B member 1 inter |
| KAT2B | 8850 | Homo sapiens | lysine acetyltransferase 2B(KAT2B) |
| HADHA | 3030 | Homo sapiens | hydroxyacyl-CoA dehydrogenase trifunctional multienzyme com |
| HEATR5B | 54497 | Homo sapiens | HEAT repeat containing 5B(HEATR5B) |
| MTF1 | 4520 | Homo sapiens | metal regulatory transcription factor 1(MTF1) |
| EXOC4 | 60412 | Homo sapiens | exocyst complex component 4(EXOC4) |
| EXOC3 | 11336 | Homo sapiens | exocyst complex component 3(EXOC3) |
| MYZAP | 100820829 | Homo sapiens | myocardial zonula adherens protein(MYZAP) |
| PLEKHO2 | 80301 | Homo sapiens | pleckstrin homology domain containing O2(PLEKHO2) |
| EXOC5 | 10640 | Homo sapiens | exocyst complex component 5(EXOC5) |
| KIFC3 | 3801 | Homo sapiens | kinesin family member C3(KIFC3) |
| GART | 2618 | Homo sapiens | phosphoribosylglycinamide formyltransferase, phosphoribosylgly |
| EXOC1 | 55763 | Homo sapiens | exocyst complex component 1(EXOC1) |

| | | | |
|---|---|---|---|
| WDR27 | 253769 | Homo sapiens | WD repeat domain 27(WDR27) |
| GRAMD1C | 54762 | Homo sapiens | GRAM domain containing 1C(GRAMD1C) |
| CDC123 | 8872 | Homo sapiens | cell division cycle 123(CDC123) |
| NBAS | 51594 | Homo sapiens | NBAS subunit of NRZ tethering complex(NBAS) |
| WDR25 | 79446 | Homo sapiens | WD repeat domain 25(WDR25) |
| NFIX | 4784 | Homo sapiens | nuclear factor I X(NFIX) |
| WDR26 | 80232 | Homo sapiens | WD repeat domain 26(WDR26) |
| PRUNE2 | 158471 | Homo sapiens | prune homolog 2 with BCH domain(PRUNE2) |
| ADCY2 | 108 | Homo sapiens | adenylate cyclase 2(ADCY2) |
| ADCY1 | 107 | Homo sapiens | adenylate cyclase 1(ADCY1) |
| PAPSS2 | 9060 | Homo sapiens | 3'-phosphoadenosine 5'-phosphosulfate synthase 2(PAPSS2) |
| PAPSS1 | 9061 | Homo sapiens | 3'-phosphoadenosine 5'-phosphosulfate synthase 1(PAPSS1) |
| FGD3 | 89846 | Homo sapiens | FYVE, RhoGEF and PH domain containing 3(FGD3) |
| FGD4 | 121512 | Homo sapiens | FYVE, RhoGEF and PH domain containing 4(FGD4) |
| FGD5 | 152273 | Homo sapiens | FYVE, RhoGEF and PH domain containing 5(FGD5) |
| DHRSX | 207063 | Homo sapiens | dehydrogenase/reductase X-linked(DHRSX) |
| GPNMB | 10457 | Homo sapiens | glycoprotein nmb(GPNMB) |
| NSD1 | 64324 | Homo sapiens | nuclear receptor binding SET domain protein 1(NSD1) |
| SLC17A7 | 57030 | Homo sapiens | solute carrier family 17 member 7(SLC17A7) |
| WDR17 | 116966 | Homo sapiens | WD repeat domain 17(WDR17) |
| PCGF6 | 84108 | Homo sapiens | polycomb group ring finger 6(PCGF6) |
| YTHDF2 | 51441 | Homo sapiens | YTH N6-methyladenosine RNA binding protein F2(YTHDF2) |
| YTHDF3 | 253943 | Homo sapiens | YTH N6-methyladenosine RNA binding protein F3(YTHDF3) |
| NOXRED1 | 122945 | Homo sapiens | NADP dependent oxidoreductase domain containing 1(NOXRED1 |
| MGMT | 4255 | Homo sapiens | O-6-methylguanine-DNA methyltransferase(MGMT) |
| PCGF5 | 84333 | Homo sapiens | polycomb group ring finger 5(PCGF5) |
| RNLS | 55328 | Homo sapiens | renalase, FAD dependent amine oxidase(RNLS) |
| GABRA5 | 2558 | Homo sapiens | gamma-aminobutyric acid type A receptor subunit alpha5(GABR |
| MCF2 | 4168 | Homo sapiens | MCF.2 cell line derived transforming sequence(MCF2) |
| WDR18 | 57418 | Homo sapiens | WD repeat domain 18(WDR18) |
| MOCOS | 55034 | Homo sapiens | molybdenum cofactor sulfurase(MOCOS) |
| GSR | 2936 | Homo sapiens | glutathione-disulfide reductase(GSR) |
| WDR19 | 57728 | Homo sapiens | WD repeat domain 19(WDR19) |
| SYVN1 | 84447 | Homo sapiens | synoviolin 1(SYVN1) |
| ELP2 | 55250 | Homo sapiens | elongator acetyltransferase complex subunit 2(ELP2) |
| ELP4 | 26610 | Homo sapiens | elongator acetyltransferase complex subunit 4(ELP4) |
| SNCAIP | 9627 | Homo sapiens | synuclein alpha interacting protein(SNCAIP) |
| DHRS3 | 9249 | Homo sapiens | dehydrogenase/reductase 3(DHRS3) |
| HIPK2 | 28996 | Homo sapiens | homeodomain interacting protein kinase 2(HIPK2) |
| MKLN1 | 4289 | Homo sapiens | muskelin 1(MKLN1) |
| SMOC2 | 64094 | Homo sapiens | SPARC related modular calcium binding 2(SMOC2) |
| DHRS7 | 51635 | Homo sapiens | dehydrogenase/reductase 7(DHRS7) |
| NFIB | 4781 | Homo sapiens | nuclear factor I B(NFIB) |
| TMEM39B | 55116 | Homo sapiens | transmembrane protein 39B(TMEM39B) |
| CEP44 | 80817 | Homo sapiens | centrosomal protein 44(CEP44) |

| | | | |
|---|---|---|---|
| NUCB2 | 4925 | Homo sapiens | nucleobindin 2(NUCB2) |
| OXNAD1 | 92106 | Homo sapiens | oxidoreductase NAD binding domain containing 1(OXNAD1) |
| C8ORF48 | 157773 | Homo sapiens | chromosome 8 open reading frame 48(C8orf48) |
| UBP1 | 7342 | Homo sapiens | upstream binding protein 1(UBP1) |
| TM7SF3 | 51768 | Homo sapiens | transmembrane 7 superfamily member 3(TM7SF3) |
| IPO7 | 10527 | Homo sapiens | importin 7(IPO7) |
| CLU | 1191 | Homo sapiens | clusterin(CLU) |
| CDC14A | 8556 | Homo sapiens | cell division cycle 14A(CDC14A) |
| IPO5 | 3843 | Homo sapiens | importin 5(IPO5) |
| THG1L | 54974 | Homo sapiens | tRNA-histidine guanylyltransferase 1 like(THG1L) |
| SLC8A2 | 6543 | Homo sapiens | solute carrier family 8 member A2(SLC8A2) |
| RIMS2 | 9699 | Homo sapiens | regulating synaptic membrane exocytosis 2(RIMS2) |
| RIMS3 | 9783 | Homo sapiens | regulating synaptic membrane exocytosis 3(RIMS3) |
| SLC39A9 | 55334 | Homo sapiens | solute carrier family 39 member 9(SLC39A9) |
| TNN | 63923 | Homo sapiens | tenascin N(TNN) |
| PHYHD1 | 254295 | Homo sapiens | phytanoyl-CoA dioxygenase domain containing 1(PHYHD1) |
| SLC39A6 | 25800 | Homo sapiens | solute carrier family 39 member 6(SLC39A6) |
| PAPOLA | 10914 | Homo sapiens | poly(A) polymerase alpha(PAPOLA) |
| TNR | 7143 | Homo sapiens | tenascin R(TNR) |
| PLS3 | 5358 | Homo sapiens | plastin 3(PLS3) |
| SLC39A8 | 64116 | Homo sapiens | solute carrier family 39 member 8(SLC39A8) |
| PLXNC1 | 10154 | Homo sapiens | plexin C1(PLXNC1) |
| KCMF1 | 56888 | Homo sapiens | potassium channel modulatory factor 1(KCMF1) |
| MAP2K3 | 5606 | Homo sapiens | mitogen-activated protein kinase kinase 3(MAP2K3) |
| MAP2K4 | 6416 | Homo sapiens | mitogen-activated protein kinase kinase 4(MAP2K4) |
| MAP2K1 | 5604 | Homo sapiens | mitogen-activated protein kinase kinase 1(MAP2K1) |
| TLE1 | 7088 | Homo sapiens | TLE family member 1, transcriptional corepressor(TLE1) |
| COL27A1 | 85301 | Homo sapiens | collagen type XXVII alpha 1 chain(COL27A1) |
| USP5 | 8078 | Homo sapiens | ubiquitin specific peptidase 5(USP5) |
| RALGAPA1 | 253959 | Homo sapiens | Ral GTPase activating protein catalytic subunit alpha 1(RALGAPA |
| TPM3 | 7170 | Homo sapiens | tropomyosin 3(TPM3) |
| USP2 | 9099 | Homo sapiens | ubiquitin specific peptidase 2(USP2) |
| TTC21A | 199223 | Homo sapiens | tetratricopeptide repeat domain 21A(TTC21A) |
| FRMD4A | 55691 | Homo sapiens | FERM domain containing 4A(FRMD4A) |
| THOC3 | 84321 | Homo sapiens | THO complex subunit 3(THOC3) |
| FRMD4B | 23150 | Homo sapiens | FERM domain containing 4B(FRMD4B) |
| VRK2 | 7444 | Homo sapiens | VRK serine/threonine kinase 2(VRK2) |
| THOC2 | 57187 | Homo sapiens | THO complex subunit 2(THOC2) |
| THOC5 | 8563 | Homo sapiens | THO complex subunit 5(THOC5) |
| HIC1 | 3090 | Homo sapiens | HIC ZBTB transcriptional repressor 1(HIC1) |
| NRBP1 | 29959 | Homo sapiens | nuclear receptor binding protein 1(NRBP1) |
| THOC7 | 80145 | Homo sapiens | THO complex subunit 7(THOC7) |
| EDAR | 10913 | Homo sapiens | ectodysplasin A receptor(EDAR) |
| OBSCN | 84033 | Homo sapiens | obscurin, cytoskeletal calmodulin and titin-interacting RhoGEF(C |
| KAT6B | 23522 | Homo sapiens | lysine acetyltransferase 6B(KAT6B) |

| | | | |
|---|---|---|---|
| RAB37 | 326624 | Homo sapiens | RAB37, member RAS oncogene family(RAB37) |
| KAT6A | 7994 | Homo sapiens | lysine acetyltransferase 6A(KAT6A) |
| ZNF316 | 100131017 | Homo sapiens | zinc finger protein 316(ZNF316) |
| RAB38 | 23682 | Homo sapiens | RAB38, member RAS oncogene family(RAB38) |
| TOX | 9760 | Homo sapiens | thymocyte selection associated high mobility group box(TOX) |
| CSNK1G1 | 53944 | Homo sapiens | casein kinase 1 gamma 1(CSNK1G1) |
| DRC7 | 84229 | Homo sapiens | dynein regulatory complex subunit 7(DRC7) |
| LOC400499 | 400499 | Homo sapiens | putative uncharacterized protein LOC400499(LOC400499) |
| CCDC25 | 55246 | Homo sapiens | coiled-coil domain containing 25(CCDC25) |
| UBR4 | 23352 | Homo sapiens | ubiquitin protein ligase E3 component n-recognin 4(UBR4) |
| SLC38A10 | 124565 | Homo sapiens | solute carrier family 38 member 10(SLC38A10) |
| MTM1 | 4534 | Homo sapiens | myotubularin 1(MTM1) |
| ORC5 | 5001 | Homo sapiens | origin recognition complex subunit 5(ORC5) |
| SCRT2 | 85508 | Homo sapiens | scratch family transcriptional repressor 2(SCRT2) |
| FGGY | 55277 | Homo sapiens | FGGY carbohydrate kinase domain containing(FGGY) |
| ORC3 | 23595 | Homo sapiens | origin recognition complex subunit 3(ORC3) |
| UBR5 | 51366 | Homo sapiens | ubiquitin protein ligase E3 component n-recognin 5(UBR5) |
| CD300LB | 124599 | Homo sapiens | CD300 molecule like family member b(CD300LB) |
| SRGAP3 | 9901 | Homo sapiens | SLIT-ROBO Rho GTPase activating protein 3(SRGAP3) |
| TPPP | 11076 | Homo sapiens | tubulin polymerization promoting protein(TPPP) |
| CD300LF | 146722 | Homo sapiens | CD300 molecule like family member f(CD300LF) |
| DRD2 | 1813 | Homo sapiens | dopamine receptor D2(DRD2) |
| DRD3 | 1814 | Homo sapiens | dopamine receptor D3(DRD3) |
| AP1M1 | 8907 | Homo sapiens | adaptor related protein complex 1 subunit mu 1(AP1M1) |
| MAP2K6 | 5608 | Homo sapiens | mitogen-activated protein kinase kinase 6(MAP2K6) |
| OSBPL9 | 114883 | Homo sapiens | oxysterol binding protein like 9(OSBPL9) |
| UVRAG | 7405 | Homo sapiens | UV radiation resistance associated(UVRAG) |
| SEC16B | 89866 | Homo sapiens | SEC16 homolog B, endoplasmic reticulum export factor(SEC16B |
| CEP112 | 201134 | Homo sapiens | centrosomal protein 112(CEP112) |
| EIF2B3 | 8891 | Homo sapiens | eukaryotic translation initiation factor 2B subunit gamma(EIF2B3 |
| TMEM132E | 124842 | Homo sapiens | transmembrane protein 132E(TMEM132E) |
| OSBPL6 | 114880 | Homo sapiens | oxysterol binding protein like 6(OSBPL6) |
| OSBPL5 | 114879 | Homo sapiens | oxysterol binding protein like 5(OSBPL5) |
| CCDC12 | 151903 | Homo sapiens | coiled-coil domain containing 12(CCDC12) |
| CCDC15 | 80071 | Homo sapiens | coiled-coil domain containing 15(CCDC15) |
| SLC4A11 | 83959 | Homo sapiens | solute carrier family 4 member 11(SLC4A11) |
| PHKB | 5257 | Homo sapiens | phosphorylase kinase regulatory subunit beta(PHKB) |
| FANCC | 2176 | Homo sapiens | FA complementation group C(FANCC) |
| TRAPPC9 | 83696 | Homo sapiens | trafficking protein particle complex subunit 9(TRAPPC9) |
| GFAP | 2670 | Homo sapiens | glial fibrillary acidic protein(GFAP) |
| REEP1 | 65055 | Homo sapiens | receptor accessory protein 1(REEP1) |
| LRFN2 | 57497 | Homo sapiens | leucine rich repeat and fibronectin type III domain containing 2( |
| ADCY10 | 55811 | Homo sapiens | adenylate cyclase 10(ADCY10) |
| BCL6 | 604 | Homo sapiens | BCL6 transcription repressor(BCL6) |
| SYNJ1 | 8867 | Homo sapiens | synaptojanin 1(SYNJ1) |

| | | | |
|---|---|---|---|
| PLEKHM1 | 9842 | Homo sapiens | pleckstrin homology and RUN domain containing M1(PLEKHM1) |
| MAP3K10 | 4294 | Homo sapiens | mitogen-activated protein kinase kinase kinase 10(MAP3K10) |
| PLEKHM2 | 23207 | Homo sapiens | pleckstrin homology and RUN domain containing M2(PLEKHM2) |
| SLC26A7 | 115111 | Homo sapiens | solute carrier family 26 member 7(SLC26A7) |
| LHX5 | 64211 | Homo sapiens | LIM homeobox 5(LHX5) |
| DRC1 | 92749 | Homo sapiens | dynein regulatory complex subunit 1(DRC1) |
| MAP3K13 | 9175 | Homo sapiens | mitogen-activated protein kinase kinase kinase 13(MAP3K13) |
| RAB5A | 5868 | Homo sapiens | RAB5A, member RAS oncogene family(RAB5A) |
| LHX9 | 56956 | Homo sapiens | LIM homeobox 9(LHX9) |
| LHX8 | 431707 | Homo sapiens | LIM homeobox 8(LHX8) |
| SLC48A1 | 55652 | Homo sapiens | solute carrier family 48 member 1(SLC48A1) |
| CPM | 1368 | Homo sapiens | carboxypeptidase M(CPM) |
| BMPR2 | 659 | Homo sapiens | bone morphogenetic protein receptor type 2(BMPR2) |
| GAL3ST1 | 9514 | Homo sapiens | galactose-3-O-sulfotransferase 1(GAL3ST1) |
| MCM8 | 84515 | Homo sapiens | minichromosome maintenance 8 homologous recombination rep |
| CCDC47 | 57003 | Homo sapiens | coiled-coil domain containing 47(CCDC47) |
| WDR4 | 10785 | Homo sapiens | WD repeat domain 4(WDR4) |
| CEP164 | 22897 | Homo sapiens | centrosomal protein 164(CEP164) |
| UFSP2 | 55325 | Homo sapiens | UFM1 specific peptidase 2(UFSP2) |
| ZC3H6 | 376940 | Homo sapiens | zinc finger CCCH-type containing 6(ZC3H6) |
| PIK3C2A | 5286 | Homo sapiens | phosphatidylinositol-4-phosphate 3-kinase catalytic subunit type |
| FTCD | 10841 | Homo sapiens | formimidoyltransferase cyclodeaminase(FTCD) |
| SYNE2 | 23224 | Homo sapiens | spectrin repeat containing nuclear envelope protein 2(SYNE2) |
| SNRNP70 | 6625 | Homo sapiens | small nuclear ribonucleoprotein U1 subunit 70(SNRNP70) |
| CEP170B | 283638 | Homo sapiens | centrosomal protein 170B(CEP170B) |
| UBTF | 7343 | Homo sapiens | upstream binding transcription factor(UBTF) |
| PHACTR3 | 116154 | Homo sapiens | phosphatase and actin regulator 3(PHACTR3) |
| TLK1 | 9874 | Homo sapiens | tousled like kinase 1(TLK1) |
| WDR7 | 23335 | Homo sapiens | WD repeat domain 7(WDR7) |
| EMILIN1 | 11117 | Homo sapiens | elastin microfibril interfacer 1(EMILIN1) |
| SERAC1 | 84947 | Homo sapiens | serine active site containing 1(SERAC1) |
| TNPO1 | 3842 | Homo sapiens | transportin 1(TNPO1) |
| EIF2B1 | 1967 | Homo sapiens | eukaryotic translation initiation factor 2B subunit alpha(EIF2B1) |
| ADAMTS8 | 11095 | Homo sapiens | ADAM metallopeptidase with thrombospondin type 1 motif 8(AD |
| ADAMTS9 | 56999 | Homo sapiens | ADAM metallopeptidase with thrombospondin type 1 motif 9(AD |
| HGFAC | 3083 | Homo sapiens | HGF activator(HGFAC) |
| TNPO3 | 23534 | Homo sapiens | transportin 3(TNPO3) |
| CCDC50 | 152137 | Homo sapiens | coiled-coil domain containing 50(CCDC50) |
| C8ORF88 | 100127983 | Homo sapiens | chromosome 8 open reading frame 88(C8orf88) |
| IDH3A | 3419 | Homo sapiens | isocitrate dehydrogenase (NAD(+)) 3 catalytic subunit alpha(ID |
| ADAMTS7 | 11173 | Homo sapiens | ADAM metallopeptidase with thrombospondin type 1 motif 7(AD |
| CCDC38 | 120935 | Homo sapiens | coiled-coil domain containing 38(CCDC38) |
| TRPC5 | 7224 | Homo sapiens | transient receptor potential cation channel subfamily C member |
| PLA2G12A | 81579 | Homo sapiens | phospholipase A2 group XIIA(PLA2G12A) |
| PLEKHG1 | 57480 | Homo sapiens | pleckstrin homology and RhoGEF domain containing G1(PLEKHG |

| | | | |
|---|---|---|---|
| SLC13A3 | 64849 | Homo sapiens | solute carrier family 13 member 3(SLC13A3) |
| TRPC6 | 7225 | Homo sapiens | transient receptor potential cation channel subfamily C member |
| PLA2G12B | 84647 | Homo sapiens | phospholipase A2 group XIIB(PLA2G12B) |
| PLEKHG2 | 64857 | Homo sapiens | pleckstrin homology and RhoGEF domain containing G2(PLEKHG |
| RFC1 | 5981 | Homo sapiens | replication factor C subunit 1(RFC1) |
| CCDC34 | 91057 | Homo sapiens | coiled-coil domain containing 34(CCDC34) |
| DNAJC2 | 27000 | Homo sapiens | DnaJ heat shock protein family (Hsp40) member C2(DNAJC2) |
| SAPCD2 | 89958 | Homo sapiens | suppressor APC domain containing 2(SAPCD2) |
| TFDP2 | 7029 | Homo sapiens | transcription factor Dp-2(TFDP2) |
| MEP1B | 4225 | Homo sapiens | meprin A subunit beta(MEP1B) |
| LCK | 3932 | Homo sapiens | LCK proto-oncogene, Src family tyrosine kinase(LCK) |
| POLR1A | 25885 | Homo sapiens | RNA polymerase I subunit A(POLR1A) |
| CATSPERG | 57828 | Homo sapiens | cation channel sperm associated auxiliary subunit gamma(CATS |
| POLR1B | 84172 | Homo sapiens | RNA polymerase I subunit B(POLR1B) |
| NOS1AP | 9722 | Homo sapiens | nitric oxide synthase 1 adaptor protein(NOS1AP) |
| PECAM1 | 5175 | Homo sapiens | platelet and endothelial cell adhesion molecule 1(PECAM1) |
| POLR1E | 64425 | Homo sapiens | RNA polymerase I subunit E(POLR1E) |
| MCM5 | 4174 | Homo sapiens | minichromosome maintenance complex component 5(MCM5) |
| DNAJC9 | 23234 | Homo sapiens | DnaJ heat shock protein family (Hsp40) member C9(DNAJC9) |
| ALDH9A1 | 223 | Homo sapiens | aldehyde dehydrogenase 9 family member A1(ALDH9A1) |
| HDAC4 | 9759 | Homo sapiens | histone deacetylase 4(HDAC4) |
| HDAC5 | 10014 | Homo sapiens | histone deacetylase 5(HDAC5) |
| RAB1A | 5861 | Homo sapiens | RAB1A, member RAS oncogene family(RAB1A) |
| CATSPER4 | 378807 | Homo sapiens | cation channel sperm associated 4(CATSPER4) |
| VPS4B | 9525 | Homo sapiens | vacuolar protein sorting 4 homolog B(VPS4B) |
| NLRC5 | 84166 | Homo sapiens | NLR family CARD domain containing 5(NLRC5) |
| LMF2 | 91289 | Homo sapiens | lipase maturation factor 2(LMF2) |
| HDAC7 | 51564 | Homo sapiens | histone deacetylase 7(HDAC7) |
| FBXL20 | 84961 | Homo sapiens | F-box and leucine rich repeat protein 20(FBXL20) |
| STOX2 | 56977 | Homo sapiens | storkhead box 2(STOX2) |
| ABLIM1 | 3983 | Homo sapiens | actin binding LIM protein 1(ABLIM1) |
| ABLIM2 | 84448 | Homo sapiens | actin binding LIM protein family member 2(ABLIM2) |
| RAB20 | 55647 | Homo sapiens | RAB20, member RAS oncogene family(RAB20) |
| HECTD2 | 143279 | Homo sapiens | HECT domain E3 ubiquitin protein ligase 2(HECTD2) |
| POLR2A | 5430 | Homo sapiens | RNA polymerase II subunit A(POLR2A) |
| MSTO1 | 55154 | Homo sapiens | misato mitochondrial distribution and morphology regulator 1(M |
| POLR2B | 5431 | Homo sapiens | RNA polymerase II subunit B(POLR2B) |
| DNAJB5 | 25822 | Homo sapiens | DnaJ heat shock protein family (Hsp40) member B5(DNAJB5) |
| VPS53 | 55275 | Homo sapiens | VPS53 subunit of GARP complex(VPS53) |
| POLR2E | 5434 | Homo sapiens | RNA polymerase II, I and III subunit E(POLR2E) |
| ASCC3 | 10973 | Homo sapiens | activating signal cointegrator 1 complex subunit 3(ASCC3) |
| PLEKHH1 | 57475 | Homo sapiens | pleckstrin homology, MyTH4 and FERM domain containing H1(P |
| ZNF740 | 283337 | Homo sapiens | zinc finger protein 740(ZNF740) |
| JAZF1 | 221895 | Homo sapiens | JAZF zinc finger 1(JAZF1) |
| CCDC73 | 493860 | Homo sapiens | coiled-coil domain containing 73(CCDC73) |

| | | | |
|---|---|---|---|
| PPIL1 | 51645 | Homo sapiens | peptidylprolyl isomerase like 1(PPIL1) |
| RNGTT | 8732 | Homo sapiens | RNA guanylyltransferase and 5'-phosphatase(RNGTT) |
| TRPA1 | 8989 | Homo sapiens | transient receptor potential cation channel subfamily A member |
| SMOX | 54498 | Homo sapiens | spermine oxidase(SMOX) |
| ASCC1 | 51008 | Homo sapiens | activating signal cointegrator 1 complex subunit 1(ASCC1) |
| AMFR | 267 | Homo sapiens | autocrine motility factor receptor(AMFR) |
| FBXL17 | 64839 | Homo sapiens | F-box and leucine rich repeat protein 17(FBXL17) |
| FBXL18 | 80028 | Homo sapiens | F-box and leucine rich repeat protein 18(FBXL18) |
| RPRD2 | 23248 | Homo sapiens | regulation of nuclear pre-mRNA domain containing 2(RPRD2) |
| CPPED1 | 55313 | Homo sapiens | calcineurin like phosphoesterase domain containing 1(CPPED1) |
| RAB10 | 10890 | Homo sapiens | RAB10, member RAS oncogene family(RAB10) |
| FRAS1 | 80144 | Homo sapiens | Fraser extracellular matrix complex subunit 1(FRAS1) |
| KIAA0586 | 9786 | Homo sapiens | KIAA0586(KIAA0586) |
| RAB15 | 376267 | Homo sapiens | RAB15, member RAS oncogene family(RAB15) |
| CTH | 1491 | Homo sapiens | cystathionine gamma-lyase(CTH) |
| ASXL3 | 80816 | Homo sapiens | ASXL transcriptional regulator 3(ASXL3) |
| MAB21L1 | 4081 | Homo sapiens | mab-21 like 1(MAB21L1) |
| MAB21L2 | 10586 | Homo sapiens | mab-21 like 2(MAB21L2) |
| VPS41 | 27072 | Homo sapiens | VPS41 subunit of HOPS complex(VPS41) |
| POLR3F | 10621 | Homo sapiens | RNA polymerase III subunit F(POLR3F) |
| TLL2 | 7093 | Homo sapiens | tolloid like 2(TLL2) |
| CEBPZ | 10153 | Homo sapiens | CCAAT enhancer binding protein zeta(CEBPZ) |
| PPIL4 | 85313 | Homo sapiens | peptidylprolyl isomerase like 4(PPIL4) |
| CCDC60 | 160777 | Homo sapiens | coiled-coil domain containing 60(CCDC60) |
| TNRC6A | 27327 | Homo sapiens | trinucleotide repeat containing adaptor 6A(TNRC6A) |
| TNRC6B | 23112 | Homo sapiens | trinucleotide repeat containing adaptor 6B(TNRC6B) |
| UMODL1 | 89766 | Homo sapiens | uromodulin like 1(UMODL1) |
| ISCA2 | 122961 | Homo sapiens | iron-sulfur cluster assembly 2(ISCA2) |
| TCERG1 | 10915 | Homo sapiens | transcription elongation regulator 1(TCERG1) |
| CBLB | 868 | Homo sapiens | Cbl proto-oncogene B(CBLB) |
| TIAL1 | 7073 | Homo sapiens | TIA1 cytotoxic granule associated RNA binding protein like 1(TI/ |
| OR5D18 | 219438 | Homo sapiens | olfactory receptor family 5 subfamily D member 18(OR5D18) |
| RPS6KA5 | 9252 | Homo sapiens | ribosomal protein S6 kinase A5(RPS6KA5) |
| FTH1 | 2495 | Homo sapiens | ferritin heavy chain 1(FTH1) |
| RPS6KA2 | 6196 | Homo sapiens | ribosomal protein S6 kinase A2(RPS6KA2) |
| HMCN2 | 256158 | Homo sapiens | hemicentin 2(HMCN2) |
| PDK3 | 5165 | Homo sapiens | pyruvate dehydrogenase kinase 3(PDK3) |
| RPS11 | 6205 | Homo sapiens | ribosomal protein S11(RPS11) |
| SLC35A5 | 55032 | Homo sapiens | solute carrier family 35 member A5(SLC35A5) |
| PDK2 | 5164 | Homo sapiens | pyruvate dehydrogenase kinase 2(PDK2) |
| PDK1 | 5163 | Homo sapiens | pyruvate dehydrogenase kinase 1(PDK1) |
| DYNC2H1 | 79659 | Homo sapiens | dynein cytoplasmic 2 heavy chain 1(DYNC2H1) |
| SRRM4 | 84530 | Homo sapiens | serine/arginine repetitive matrix 4(SRRM4) |
| COG6 | 57511 | Homo sapiens | component of oligomeric golgi complex 6(COG6) |
| COG2 | 22796 | Homo sapiens | component of oligomeric golgi complex 2(COG2) |

| | | | |
|---|---|---|---|
| P3H1 | 64175 | Homo sapiens | prolyl 3-hydroxylase 1(P3H1) |
| P3H3 | 10536 | Homo sapiens | prolyl 3-hydroxylase 3(P3H3) |
| MAPK8IP3 | 23162 | Homo sapiens | mitogen-activated protein kinase 8 interacting protein 3(MAPK8 |
| CRNKL1 | 51340 | Homo sapiens | crooked neck pre-mRNA splicing factor 1(CRNKL1) |
| UHMK1 | 127933 | Homo sapiens | U2AF homology motif kinase 1(UHMK1) |
| TIAM2 | 26230 | Homo sapiens | TIAM Rac1 associated GEF 2(TIAM2) |
| POR | 5447 | Homo sapiens | cytochrome p450 oxidoreductase(POR) |
| KATNAL2 | 83473 | Homo sapiens | katanin catalytic subunit A1 like 2(KATNAL2) |
| SIK3 | 23387 | Homo sapiens | SIK family kinase 3(SIK3) |
| SUCLG2 | 8801 | Homo sapiens | succinate-CoA ligase GDP-forming subunit beta(SUCLG2) |
| SPTBN5 | 51332 | Homo sapiens | spectrin beta, non-erythrocytic 5(SPTBN5) |
| SLC22A5 | 6584 | Homo sapiens | solute carrier family 22 member 5(SLC22A5) |
| STXBP4 | 252983 | Homo sapiens | syntaxin binding protein 4(STXBP4) |
| STXBP1 | 6812 | Homo sapiens | syntaxin binding protein 1(STXBP1) |
| DCUN1D4 | 23142 | Homo sapiens | defective in cullin neddylation 1 domain containing 4(DCUN1D4 |
| RPAP1 | 26015 | Homo sapiens | RNA polymerase II associated protein 1(RPAP1) |
| CRHR1 | 1394 | Homo sapiens | corticotropin releasing hormone receptor 1(CRHR1) |
| CRHR2 | 1395 | Homo sapiens | corticotropin releasing hormone receptor 2(CRHR2) |
| TRPM2 | 7226 | Homo sapiens | transient receptor potential cation channel subfamily M member |
| UGCG | 7357 | Homo sapiens | UDP-glucose ceramide glucosyltransferase(UGCG) |
| FXR2 | 9513 | Homo sapiens | FMR1 autosomal homolog 2(FXR2) |
| ZNF827 | 152485 | Homo sapiens | zinc finger protein 827(ZNF827) |
| ZNHIT6 | 54680 | Homo sapiens | zinc finger HIT-type containing 6(ZNHIT6) |
| TP53BP2 | 7159 | Homo sapiens | tumor protein p53 binding protein 2(TP53BP2) |
| GSE1 | 23199 | Homo sapiens | Gse1 coiled-coil protein(GSE1) |
| TRPM7 | 54822 | Homo sapiens | transient receptor potential cation channel subfamily M member |
| ATP6V0A4 | 50617 | Homo sapiens | ATPase H+ transporting V0 subunit a4(ATP6V0A4) |
| STXBP5 | 134957 | Homo sapiens | syntaxin binding protein 5(STXBP5) |
| TP53BP1 | 7158 | Homo sapiens | tumor protein p53 binding protein 1(TP53BP1) |
| STXBP6 | 29091 | Homo sapiens | syntaxin binding protein 6(STXBP6) |
| TRPM5 | 29850 | Homo sapiens | transient receptor potential cation channel subfamily M member |
| TRPM6 | 140803 | Homo sapiens | transient receptor potential cation channel subfamily M member |
| RALGDS | 5900 | Homo sapiens | ral guanine nucleotide dissociation stimulator(RALGDS) |
| CCR10 | 2826 | Homo sapiens | C-C motif chemokine receptor 10(CCR10) |
| TRPM4 | 54795 | Homo sapiens | transient receptor potential cation channel subfamily M member |
| VAT1 | 10493 | Homo sapiens | vesicle amine transport 1(VAT1) |
| TMEM184B | 25829 | Homo sapiens | transmembrane protein 184B(TMEM184B) |
| SYNRG | 11276 | Homo sapiens | synergin gamma(SYNRG) |
| SETDB2 | 83852 | Homo sapiens | SET domain bifurcated histone lysine methyltransferase 2(SETD |
| PUM1 | 9698 | Homo sapiens | pumilio RNA binding family member 1(PUM1) |
| SNAP91 | 9892 | Homo sapiens | synaptosome associated protein 91(SNAP91) |
| DERA | 51071 | Homo sapiens | deoxyribose-phosphate aldolase(DERA) |
| PM20D2 | 135293 | Homo sapiens | peptidase M20 domain containing 2(PM20D2) |
| CCSER1 | 401145 | Homo sapiens | coiled-coil serine rich protein 1(CCSER1) |
| SIM1 | 6492 | Homo sapiens | SIM bHLH transcription factor 1(SIM1) |

| | | | |
|---|---|---|---|
| TECPR2 | 9895 | Homo sapiens | tectonin beta-propeller repeat containing 2(TECPR2) |
| SIM2 | 6493 | Homo sapiens | SIM bHLH transcription factor 2(SIM2) |
| TOP2A | 7153 | Homo sapiens | DNA topoisomerase II alpha(TOP2A) |
| SLC44A5 | 204962 | Homo sapiens | solute carrier family 44 member 5(SLC44A5) |
| COL18A1 | 80781 | Homo sapiens | collagen type XVIII alpha 1 chain(COL18A1) |
| SLC35F6 | 54978 | Homo sapiens | solute carrier family 35 member F6(SLC35F6) |
| C2CD5 | 9847 | Homo sapiens | C2 calcium dependent domain containing 5(C2CD5) |
| ROBO4 | 54538 | Homo sapiens | roundabout guidance receptor 4(ROBO4) |
| UBXN1 | 51035 | Homo sapiens | UBX domain protein 1(UBXN1) |
| TENM3 | 55714 | Homo sapiens | teneurin transmembrane protein 3(TENM3) |
| HIP1 | 3092 | Homo sapiens | huntingtin interacting protein 1(HIP1) |
| C2CD3 | 26005 | Homo sapiens | C2 domain containing 3 centriole elongation regulator(C2CD3) |
| GBE1 | 2632 | Homo sapiens | 1,4-alpha-glucan branching enzyme 1(GBE1) |
| SLC35F1 | 222553 | Homo sapiens | solute carrier family 35 member F1(SLC35F1) |
| RSF1 | 51773 | Homo sapiens | remodeling and spacing factor 1(RSF1) |
| ZCCHC14 | 23174 | Homo sapiens | zinc finger CCHC-type containing 14(ZCCHC14) |
| GABRR2 | 2570 | Homo sapiens | gamma-aminobutyric acid type A receptor subunit rho2(GABRR2 |
| XPO4 | 64328 | Homo sapiens | exportin 4(XPO4) |
| SLC22A18 | 5002 | Homo sapiens | solute carrier family 22 member 18(SLC22A18) |
| DNER | 92737 | Homo sapiens | delta/notch like EGF repeat containing(DNER) |
| HRAS | 3265 | Homo sapiens | HRas proto-oncogene, GTPase(HRAS) |
| GABPA | 2551 | Homo sapiens | GA binding protein transcription factor subunit alpha(GABPA) |
| RAB8A | 4218 | Homo sapiens | RAB8A, member RAS oncogene family(RAB8A) |
| RALGPS2 | 55103 | Homo sapiens | Ral GEF with PH domain and SH3 binding motif 2(RALGPS2) |
| VAV3 | 10451 | Homo sapiens | vav guanine nucleotide exchange factor 3(VAV3) |
| TOP3B | 8940 | Homo sapiens | DNA topoisomerase III beta(TOP3B) |
| EDN1 | 1906 | Homo sapiens | endothelin 1(EDN1) |
| TMED8 | 283578 | Homo sapiens | transmembrane p24 trafficking protein family member 8(TMED8 |
| PARP4 | 143 | Homo sapiens | poly(ADP-ribose) polymerase family member 4(PARP4) |
| SLC35G1 | 159371 | Homo sapiens | solute carrier family 35 member G1(SLC35G1) |
| ZDHHC17 | 23390 | Homo sapiens | zinc finger DHHC-type palmitoyltransferase 17(ZDHHC17) |
| GFRA2 | 2675 | Homo sapiens | GDNF family receptor alpha 2(GFRA2) |
| CBFA2T3 | 863 | Homo sapiens | CBFA2/RUNX1 partner transcriptional co-repressor 3(CBFA2T3) |
| ZDHHC14 | 79683 | Homo sapiens | zinc finger DHHC-type palmitoyltransferase 14(ZDHHC14) |
| ITFG1 | 81533 | Homo sapiens | integrin alpha FG-GAP repeat containing 1(ITFG1) |
| TOPORS | 10210 | Homo sapiens | TOP1 binding arginine/serine rich protein, E3 ubiquitin ligase(TO |
| VAV2 | 7410 | Homo sapiens | vav guanine nucleotide exchange factor 2(VAV2) |
| PARP8 | 79668 | Homo sapiens | poly(ADP-ribose) polymerase family member 8(PARP8) |
| SLC22A23 | 63027 | Homo sapiens | solute carrier family 22 member 23(SLC22A23) |
| HCK | 3055 | Homo sapiens | HCK proto-oncogene, Src family tyrosine kinase(HCK) |
| HNRNPH1 | 3187 | Homo sapiens | heterogeneous nuclear ribonucleoprotein H1(HNRNPH1) |
| VWA1 | 64856 | Homo sapiens | von Willebrand factor A domain containing 1(VWA1) |
| CLDN16 | 10686 | Homo sapiens | claudin 16(CLDN16) |
| MATN4 | 8785 | Homo sapiens | matrilin 4(MATN4) |
| VGLL4 | 9686 | Homo sapiens | vestigial like family member 4(VGLL4) |

| | | | |
|---|---|---|---|
| VGLL3 | 389136 | Homo sapiens | vestigial like family member 3(VGLL3) |
| VWA7 | 80737 | Homo sapiens | von Willebrand factor A domain containing 7(VWA7) |
| AMER1 | 139285 | Homo sapiens | APC membrane recruitment protein 1(AMER1) |
| GUCY2D | 3000 | Homo sapiens | guanylate cyclase 2D, retinal(GUCY2D) |
| ZDHHC21 | 340481 | Homo sapiens | zinc finger DHHC-type palmitoyltransferase 21(ZDHHC21) |
| VWCE | 220001 | Homo sapiens | von Willebrand factor C and EGF domains(VWCE) |
| TYMS | 7298 | Homo sapiens | thymidylate synthetase(TYMS) |
| GDPD5 | 81544 | Homo sapiens | glycerophosphodiester phosphodiesterase domain containing 5( |
| DCST1 | 149095 | Homo sapiens | DC-STAMP domain containing 1(DCST1) |
| DCST2 | 127579 | Homo sapiens | DC-STAMP domain containing 2(DCST2) |
| PDPR | 55066 | Homo sapiens | pyruvate dehydrogenase phosphatase regulatory subunit(PDPR) |
| LMNA | 4000 | Homo sapiens | lamin A/C(LMNA) |
| METTL4 | 64863 | Homo sapiens | methyltransferase 4, N6-adenosine(METTL4) |
| UBTD1 | 80019 | Homo sapiens | ubiquitin domain containing 1(UBTD1) |
| APBB2 | 323 | Homo sapiens | amyloid beta precursor protein binding family B member 2(APBI |
| SUSD6 | 9766 | Homo sapiens | sushi domain containing 6(SUSD6) |
| KANSL1L | 151050 | Homo sapiens | KAT8 regulatory NSL complex subunit 1 like(KANSL1L) |
| APBB3 | 10307 | Homo sapiens | amyloid beta precursor protein binding family B member 3(APBI |
| GABRP | 2568 | Homo sapiens | gamma-aminobutyric acid type A receptor subunit pi(GABRP) |
| GINS2 | 51659 | Homo sapiens | GINS complex subunit 2(GINS2) |
| MPP2 | 4355 | Homo sapiens | MAGUK p55 scaffold protein 2(MPP2) |
| SLC31A2 | 1318 | Homo sapiens | solute carrier family 31 member 2(SLC31A2) |
| MTX2 | 10651 | Homo sapiens | metaxin 2(MTX2) |
| SUSD1 | 64420 | Homo sapiens | sushi domain containing 1(SUSD1) |
| GBF1 | 8729 | Homo sapiens | golgi brefeldin A resistant guanine nucleotide exchange factor 1( |
| GRHL1 | 29841 | Homo sapiens | grainyhead like transcription factor 1(GRHL1) |
| AIG1 | 51390 | Homo sapiens | androgen induced 1(AIG1) |
| VAX2 | 25806 | Homo sapiens | ventral anterior homeobox 2(VAX2) |
| MPP7 | 143098 | Homo sapiens | MAGUK p55 scaffold protein 7(MPP7) |
| ATAT1 | 79969 | Homo sapiens | alpha tubulin acetyltransferase 1(ATAT1) |
| COPG2 | 26958 | Homo sapiens | COPI coat complex subunit gamma 2(COPG2) |
| MEGF9 | 1955 | Homo sapiens | multiple EGF like domains 9(MEGF9) |
| KDM1A | 23028 | Homo sapiens | lysine demethylase 1A(KDM1A) |
| TAMM41 | 132001 | Homo sapiens | TAM41 mitochondrial translocator assembly and maintenance ho |
| SPPL2B | 56928 | Homo sapiens | signal peptide peptidase like 2B(SPPL2B) |
| CLEC18B | 497190 | Homo sapiens | C-type lectin domain family 18 member B(CLEC18B) |
| COMT | 1312 | Homo sapiens | catechol-O-methyltransferase(COMT) |
| CNDP1 | 84735 | Homo sapiens | carnosine dipeptidase 1(CNDP1) |
| CNDP2 | 55748 | Homo sapiens | carnosine dipeptidase 2(CNDP2) |
| DOCK11 | 139818 | Homo sapiens | dedicator of cytokinesis 11(DOCK11) |
| BMPER | 168667 | Homo sapiens | BMP binding endothelial regulator(BMPER) |
| EME1 | 146956 | Homo sapiens | essential meiotic structure-specific endonuclease 1(EME1) |
| NARS2 | 79731 | Homo sapiens | asparaginyl-tRNA synthetase 2, mitochondrial(NARS2) |
| TRAPPC12 | 51112 | Homo sapiens | trafficking protein particle complex subunit 12(TRAPPC12) |
| TRAPPC11 | 60684 | Homo sapiens | trafficking protein particle complex subunit 11(TRAPPC11) |

| | | | |
|---|---|---|---|
| SVEP1 | 79987 | Homo sapiens | sushi, von Willebrand factor type A, EGF and pentraxin domain |
| TMEM38B | 55151 | Homo sapiens | transmembrane protein 38B(TMEM38B) |
| RALBP1 | 10928 | Homo sapiens | ralA binding protein 1(RALBP1) |
| WNT5B | 81029 | Homo sapiens | Wnt family member 5B(WNT5B) |
| WNT5A | 7474 | Homo sapiens | Wnt family member 5A(WNT5A) |
| LIFR | 3977 | Homo sapiens | LIF receptor subunit alpha(LIFR) |
| LIG3 | 3980 | Homo sapiens | DNA ligase 3(LIG3) |
| TRPC4AP | 26133 | Homo sapiens | transient receptor potential cation channel subfamily C member |
| SPECC1L | 23384 | Homo sapiens | sperm antigen with calponin homology and coiled-coil domains |
| AMPH | 273 | Homo sapiens | amphiphysin(AMPH) |
| FAM161B | 145483 | Homo sapiens | FAM161 centrosomal protein B(FAM161B) |
| CHRNA1 | 1134 | Homo sapiens | cholinergic receptor nicotinic alpha 1 subunit(CHRNA1) |
| COPB2 | 9276 | Homo sapiens | COPI coat complex subunit beta 2(COPB2) |
| ZCCHC7 | 84186 | Homo sapiens | zinc finger CCHC-type containing 7(ZCCHC7) |
| RASL12 | 51285 | Homo sapiens | RAS like family 12(RASL12) |
| CHRNA3 | 1136 | Homo sapiens | cholinergic receptor nicotinic alpha 3 subunit(CHRNA3) |
| COPA | 1314 | Homo sapiens | COPI coat complex subunit alpha(COPA) |
| GSTCD | 79807 | Homo sapiens | glutathione S-transferase C-terminal domain containing(GSTCD |
| CABP2 | 51475 | Homo sapiens | calcium binding protein 2(CABP2) |
| FBLN2 | 2199 | Homo sapiens | fibulin 2(FBLN2) |
| TRHDE | 29953 | Homo sapiens | thyrotropin releasing hormone degrading enzyme(TRHDE) |
| LPP | 4026 | Homo sapiens | LIM domain containing preferred translocation partner in lipoma |
| SCUBE2 | 57758 | Homo sapiens | signal peptide, CUB domain and EGF like domain containing 2(S |
| SCUBE3 | 222663 | Homo sapiens | signal peptide, CUB domain and EGF like domain containing 3(S |
| CDH20 | 28316 | Homo sapiens | cadherin 20(CDH20) |
| CDH22 | 64405 | Homo sapiens | cadherin 22(CDH22) |
| CDH23 | 64072 | Homo sapiens | cadherin related 23(CDH23) |
| RPRD1B | 58490 | Homo sapiens | regulation of nuclear pre-mRNA domain containing 1B(RPRD1B) |
| CLSPN | 63967 | Homo sapiens | claspin(CLSPN) |
| FAM83C | 128876 | Homo sapiens | family with sequence similarity 83 member C(FAM83C) |
| GAS2L2 | 246176 | Homo sapiens | growth arrest specific 2 like 2(GAS2L2) |
| LONRF2 | 164832 | Homo sapiens | LON peptidase N-terminal domain and ring finger 2(LONRF2) |
| KLHDC10 | 23008 | Homo sapiens | kelch domain containing 10(KLHDC10) |
| SPECC1 | 92521 | Homo sapiens | sperm antigen with calponin homology and coiled-coil domains |
| KLHDC4 | 54758 | Homo sapiens | kelch domain containing 4(KLHDC4) |
| POU2F1 | 5451 | Homo sapiens | POU class 2 homeobox 1(POU2F1) |
| HELQ | 113510 | Homo sapiens | helicase, POLQ like(HELQ) |
| MLKL | 197259 | Homo sapiens | mixed lineage kinase domain like pseudokinase(MLKL) |
| UQCC1 | 55245 | Homo sapiens | ubiquinol-cytochrome c reductase complex assembly factor 1(U |
| IDH1 | 3417 | Homo sapiens | isocitrate dehydrogenase (NADP(+)) 1(IDH1) |
| TRPV3 | 162514 | Homo sapiens | transient receptor potential cation channel subfamily V member |
| HELZ | 9931 | Homo sapiens | helicase with zinc finger(HELZ) |
| EIF2S2 | 8894 | Homo sapiens | eukaryotic translation initiation factor 2 subunit beta(EIF2S2) |
| POU2F3 | 25833 | Homo sapiens | POU class 2 homeobox 3(POU2F3) |
| SELP | 6403 | Homo sapiens | selectin P(SELP) |

| | | | |
|---|---|---|---|
| DIAPH1 | 1729 | Homo sapiens | diaphanous related formin 1(DIAPH1) |
| GRHPR | 9380 | Homo sapiens | glyoxylate and hydroxypyruvate reductase(GRHPR) |
| DAP | 1611 | Homo sapiens | death associated protein(DAP) |
| DIAPH2 | 1730 | Homo sapiens | diaphanous related formin 2(DIAPH2) |
| TLX1 | 3195 | Homo sapiens | T cell leukemia homeobox 1(TLX1) |
| CHMP2B | 25978 | Homo sapiens | charged multivesicular body protein 2B(CHMP2B) |
| CAPRIN2 | 65981 | Homo sapiens | caprin family member 2(CAPRIN2) |
| PI4KA | 5297 | Homo sapiens | phosphatidylinositol 4-kinase alpha(PI4KA) |
| BRWD1 | 54014 | Homo sapiens | bromodomain and WD repeat domain containing 1(BRWD1) |
| COPE | 11316 | Homo sapiens | COPI coat complex subunit epsilon(COPE) |
| ZCCHC2 | 54877 | Homo sapiens | zinc finger CCHC-type containing 2(ZCCHC2) |
| FGFR2 | 2263 | Homo sapiens | fibroblast growth factor receptor 2(FGFR2) |
| LRP12 | 29967 | Homo sapiens | LDL receptor related protein 12(LRP12) |
| AVEN | 57099 | Homo sapiens | apoptosis and caspase activation inhibitor(AVEN) |
| OPLAH | 26873 | Homo sapiens | 5-oxoprolinase, ATP-hydrolysing(OPLAH) |
| HK2 | 3099 | Homo sapiens | hexokinase 2(HK2) |
| OASL | 8638 | Homo sapiens | 2'-5'-oligoadenylate synthetase like(OASL) |
| HK1 | 3098 | Homo sapiens | hexokinase 1(HK1) |
| HS2ST1 | 9653 | Homo sapiens | heparan sulfate 2-O-sulfotransferase 1(HS2ST1) |
| MFSD8 | 256471 | Homo sapiens | major facilitator superfamily domain containing 8(MFSD8) |
| SNAPC5 | 10302 | Homo sapiens | small nuclear RNA activating complex polypeptide 5(SNAPC5) |
| FHAD1 | 114827 | Homo sapiens | forkhead associated phosphopeptide binding domain 1(FHAD1) |
| SNAPC2 | 6618 | Homo sapiens | small nuclear RNA activating complex polypeptide 2(SNAPC2) |
| TRPS1 | 7227 | Homo sapiens | transcriptional repressor GATA binding 1(TRPS1) |
| SNAPC3 | 6619 | Homo sapiens | small nuclear RNA activating complex polypeptide 3(SNAPC3) |
| SIX2 | 10736 | Homo sapiens | SIX homeobox 2(SIX2) |
| TOP1MT | 116447 | Homo sapiens | DNA topoisomerase I mitochondrial(TOP1MT) |
| CASP3 | 836 | Homo sapiens | caspase 3(CASP3) |
| MCF2L | 23263 | Homo sapiens | MCF.2 cell line derived transforming sequence like(MCF2L) |
| CYP2R1 | 120227 | Homo sapiens | cytochrome P450 family 2 subfamily R member 1(CYP2R1) |
| CASP1 | 834 | Homo sapiens | caspase 1(CASP1) |
| RBM6 | 10180 | Homo sapiens | RNA binding motif protein 6(RBM6) |
| KDM6A | 7403 | Homo sapiens | lysine demethylase 6A(KDM6A) |
| KDM6B | 23135 | Homo sapiens | lysine demethylase 6B(KDM6B) |
| STPG2 | 285555 | Homo sapiens | sperm tail PG-rich repeat containing 2(STPG2) |
| RHBDF2 | 79651 | Homo sapiens | rhomboid 5 homolog 2(RHBDF2) |
| IQCE | 23288 | Homo sapiens | IQ motif containing E(IQCE) |
| OXSR1 | 9943 | Homo sapiens | oxidative stress responsive kinase 1(OXSR1) |
| PPP2R5C | 5527 | Homo sapiens | protein phosphatase 2 regulatory subunit B'gamma(PPP2R5C) |
| LSS | 4047 | Homo sapiens | lanosterol synthase(LSS) |
| RBL2 | 5934 | Homo sapiens | RB transcriptional corepressor like 2(RBL2) |
| RABEP1 | 9135 | Homo sapiens | rabaptin, RAB GTPase binding effector protein 1(RABEP1) |
| FCHSD2 | 9873 | Homo sapiens | FCH and double SH3 domains 2(FCHSD2) |
| DDAH1 | 23576 | Homo sapiens | dimethylarginine dimethylaminohydrolase 1(DDAH1) |
| IGDCC4 | 57722 | Homo sapiens | immunoglobulin superfamily DCC subclass member 4(IGDCC4) |

| | | | |
|---|---|---|---|
| RASA3 | 22821 | Homo sapiens | RAS p21 protein activator 3(RASA3) |
| COL6A2 | 1292 | Homo sapiens | collagen type VI alpha 2 chain(COL6A2) |
| IGDCC3 | 9543 | Homo sapiens | immunoglobulin superfamily DCC subclass member 3(IGDCC3) |
| COL6A1 | 1291 | Homo sapiens | collagen type VI alpha 1 chain(COL6A1) |
| DCX | 1641 | Homo sapiens | doublecortin(DCX) |
| CDH12 | 1010 | Homo sapiens | cadherin 12(CDH12) |
| CDH13 | 1012 | Homo sapiens | cadherin 13(CDH13) |
| COL6A6 | 131873 | Homo sapiens | collagen type VI alpha 6 chain(COL6A6) |
| HAGH | 3029 | Homo sapiens | hydroxyacylglutathione hydrolase(HAGH) |
| CASQ1 | 844 | Homo sapiens | calsequestrin 1(CASQ1) |
| RBKS | 64080 | Homo sapiens | ribokinase(RBKS) |
| COLEC12 | 81035 | Homo sapiens | collectin subfamily member 12(COLEC12) |
| VAC14 | 55697 | Homo sapiens | VAC14 component of PIKFYVE complex(VAC14) |
| STOML2 | 30968 | Homo sapiens | stomatin like 2(STOML2) |
| KDM3B | 51780 | Homo sapiens | lysine demethylase 3B(KDM3B) |
| ATP1A3 | 478 | Homo sapiens | ATPase Na+/K+ transporting subunit alpha 3(ATP1A3) |
| ATP1A2 | 477 | Homo sapiens | ATPase Na+/K+ transporting subunit alpha 2(ATP1A2) |
| NR2C2 | 7182 | Homo sapiens | nuclear receptor subfamily 2 group C member 2(NR2C2) |
| NR2C1 | 7181 | Homo sapiens | nuclear receptor subfamily 2 group C member 1(NR2C1) |
| THBS3 | 7059 | Homo sapiens | thrombospondin 3(THBS3) |
| WNT11 | 7481 | Homo sapiens | Wnt family member 11(WNT11) |
| ERICH1 | 157697 | Homo sapiens | glutamate rich 1(ERICH1) |
| FAM120A | 23196 | Homo sapiens | family with sequence similarity 120 member A(FAM120A) |
| POC1B | 282809 | Homo sapiens | POC1 centriolar protein B(POC1B) |
| PDSS1 | 23590 | Homo sapiens | decaprenyl diphosphate synthase subunit 1(PDSS1) |
| CRYGS | 1427 | Homo sapiens | crystallin gamma S(CRYGS) |
| KDM4A | 9682 | Homo sapiens | lysine demethylase 4A(KDM4A) |
| LIMK2 | 3985 | Homo sapiens | LIM domain kinase 2(LIMK2) |
| DAW1 | 164781 | Homo sapiens | dynein assembly factor with WD repeats 1(DAW1) |
| PPP2R3A | 5523 | Homo sapiens | protein phosphatase 2 regulatory subunit B''alpha(PPP2R3A) |
| PPP2R2C | 5522 | Homo sapiens | protein phosphatase 2 regulatory subunit Bgamma(PPP2R2C) |
| KANSL1 | 284058 | Homo sapiens | KAT8 regulatory NSL complex subunit 1(KANSL1) |
| PPP2R2B | 5521 | Homo sapiens | protein phosphatase 2 regulatory subunit Bbeta(PPP2R2B) |
| PPP2R2D | 55844 | Homo sapiens | protein phosphatase 2 regulatory subunit Bdelta(PPP2R2D) |
| FSIP1 | 161835 | Homo sapiens | fibrous sheath interacting protein 1(FSIP1) |
| C2ORF72 | 257407 | Homo sapiens | chromosome 2 open reading frame 72(C2orf72) |
| PAXBP1 | 94104 | Homo sapiens | PAX3 and PAX7 binding protein 1(PAXBP1) |
| C5ORF22 | 55322 | Homo sapiens | chromosome 5 open reading frame 22(C5orf22) |
| MAPKBP1 | 23005 | Homo sapiens | mitogen-activated protein kinase binding protein 1(MAPKBP1) |
| PGAP1 | 80055 | Homo sapiens | post-GPI attachment to proteins inositol deacylase 1(PGAP1) |
| PGAP2 | 27315 | Homo sapiens | post-GPI attachment to proteins 2(PGAP2) |
| EHMT2 | 10919 | Homo sapiens | euchromatic histone lysine methyltransferase 2(EHMT2) |
| UXS1 | 80146 | Homo sapiens | UDP-glucuronate decarboxylase 1(UXS1) |
| EHMT1 | 79813 | Homo sapiens | euchromatic histone lysine methyltransferase 1(EHMT1) |
| ZFYVE1 | 53349 | Homo sapiens | zinc finger FYVE-type containing 1(ZFYVE1) |

| | | | |
|---|---|---|---|
| PREX2 | 80243 | Homo sapiens | phosphatidylinositol-3,4,5-trisphosphate dependent Rac exchan |
| ZFYVE28 | 57732 | Homo sapiens | zinc finger FYVE-type containing 28(ZFYVE28) |
| SAMD15 | 161394 | Homo sapiens | sterile alpha motif domain containing 15(SAMD15) |
| UBASH3B | 84959 | Homo sapiens | ubiquitin associated and SH3 domain containing B(UBASH3B) |
| PSMD1 | 5707 | Homo sapiens | proteasome 26S subunit, non-ATPase 1(PSMD1) |
| SAMD11 | 148398 | Homo sapiens | sterile alpha motif domain containing 11(SAMD11) |
| RBFOX2 | 23543 | Homo sapiens | RNA binding fox-1 homolog 2(RBFOX2) |
| RBFOX3 | 146713 | Homo sapiens | RNA binding fox-1 homolog 3(RBFOX3) |
| STARD8 | 9754 | Homo sapiens | StAR related lipid transfer domain containing 8(STARD8) |
| RFX2 | 5990 | Homo sapiens | regulatory factor X2(RFX2) |
| ZFYVE21 | 79038 | Homo sapiens | zinc finger FYVE-type containing 21(ZFYVE21) |
| RFX3 | 5991 | Homo sapiens | regulatory factor X3(RFX3) |
| RFX1 | 5989 | Homo sapiens | regulatory factor X1(RFX1) |
| HDHD2 | 84064 | Homo sapiens | haloacid dehalogenase like hydrolase domain containing 2(HDH |
| SND1 | 27044 | Homo sapiens | staphylococcal nuclease and tudor domain containing 1(SND1) |
| EML4 | 27436 | Homo sapiens | EMAP like 4(EML4) |
| AR | 367 | Homo sapiens | androgen receptor(AR) |
| EML5 | 161436 | Homo sapiens | EMAP like 5(EML5) |
| DDX39A | 10212 | Homo sapiens | DExD-box helicase 39A(DDX39A) |
| SLC5A9 | 200010 | Homo sapiens | solute carrier family 5 member 9(SLC5A9) |
| NUP210L | 91181 | Homo sapiens | nucleoporin 210 like(NUP210L) |
| PSME4 | 23198 | Homo sapiens | proteasome activator subunit 4(PSME4) |
| TMEM79 | 84283 | Homo sapiens | transmembrane protein 79(TMEM79) |
| RUFY2 | 55680 | Homo sapiens | RUN and FYVE domain containing 2(RUFY2) |
| UQCRC1 | 7384 | Homo sapiens | ubiquinol-cytochrome c reductase core protein 1(UQCRC1) |
| PADI3 | 51702 | Homo sapiens | peptidyl arginine deiminase 3(PADI3) |
| ENTHD1 | 150350 | Homo sapiens | ENTH domain containing 1(ENTHD1) |
| IDO2 | 169355 | Homo sapiens | indoleamine 2,3-dioxygenase 2(IDO2) |
| CFAP99 | 402160 | Homo sapiens | cilia and flagella associated protein 99(CFAP99) |
| IDO1 | 3620 | Homo sapiens | indoleamine 2,3-dioxygenase 1(IDO1) |
| MTMR1 | 8776 | Homo sapiens | myotubularin related protein 1(MTMR1) |
| MTMR2 | 8898 | Homo sapiens | myotubularin related protein 2(MTMR2) |
| TTYH3 | 80727 | Homo sapiens | tweety family member 3(TTYH3) |
| MTMR3 | 8897 | Homo sapiens | myotubularin related protein 3(MTMR3) |
| AIRE | 326 | Homo sapiens | autoimmune regulator(AIRE) |
| FLII | 2314 | Homo sapiens | FLII actin remodeling protein(FLII) |
| AGAP2 | 116986 | Homo sapiens | ArfGAP with GTPase domain, ankyrin repeat and PH domain 2(A |
| MTMR8 | 55613 | Homo sapiens | myotubularin related protein 8(MTMR8) |
| AGAP1 | 116987 | Homo sapiens | ArfGAP with GTPase domain, ankyrin repeat and PH domain 1(A |
| MTMR9 | 66036 | Homo sapiens | myotubularin related protein 9(MTMR9) |
| IQGAP1 | 8826 | Homo sapiens | IQ motif containing GTPase activating protein 1(IQGAP1) |
| MRPL15 | 29088 | Homo sapiens | mitochondrial ribosomal protein L15(MRPL15) |
| MTMR4 | 9110 | Homo sapiens | myotubularin related protein 4(MTMR4) |
| IQGAP3 | 128239 | Homo sapiens | IQ motif containing GTPase activating protein 3(IQGAP3) |
| C3 | 718 | Homo sapiens | complement C3(C3) |

| | | | |
|---|---|---|---|
| HSD11B2 | 3291 | Homo sapiens | hydroxysteroid 11-beta dehydrogenase 2(HSD11B2) |
| UGT8 | 7368 | Homo sapiens | UDP glycosyltransferase 8(UGT8) |
| CYB5R3 | 1727 | Homo sapiens | cytochrome b5 reductase 3(CYB5R3) |
| MYO6 | 4646 | Homo sapiens | myosin VI(MYO6) |
| VTI1A | 143187 | Homo sapiens | vesicle transport through interaction with t-SNAREs 1A(VTI1A) |
| PSMF1 | 9491 | Homo sapiens | proteasome inhibitor subunit 1(PSMF1) |
| RSU1 | 6251 | Homo sapiens | Ras suppressor protein 1(RSU1) |
| PPARGC1A | 10891 | Homo sapiens | PPARG coactivator 1 alpha(PPARGC1A) |
| EXPH5 | 23086 | Homo sapiens | exophilin 5(EXPH5) |
| ZBTB7C | 201501 | Homo sapiens | zinc finger and BTB domain containing 7C(ZBTB7C) |
| ABCA2 | 20 | Homo sapiens | ATP binding cassette subfamily A member 2(ABCA2) |
| CADM1 | 23705 | Homo sapiens | cell adhesion molecule 1(CADM1) |
| COL24A1 | 255631 | Homo sapiens | collagen type XXIV alpha 1 chain(COL24A1) |
| ABCA3 | 21 | Homo sapiens | ATP binding cassette subfamily A member 3(ABCA3) |
| LRBA | 987 | Homo sapiens | LPS responsive beige-like anchor protein(LRBA) |
| EDEM2 | 55741 | Homo sapiens | ER degradation enhancing alpha-mannosidase like protein 2(ED |
| MRPL27 | 51264 | Homo sapiens | mitochondrial ribosomal protein L27(MRPL27) |
| ABCA8 | 10351 | Homo sapiens | ATP binding cassette subfamily A member 8(ABCA8) |
| NETO2 | 81831 | Homo sapiens | neuropilin and tolloid like 2(NETO2) |
| MRPL21 | 219927 | Homo sapiens | mitochondrial ribosomal protein L21(MRPL21) |
| ABTB2 | 25841 | Homo sapiens | ankyrin repeat and BTB domain containing 2(ABTB2) |
| DAZL | 1618 | Homo sapiens | deleted in azoospermia like(DAZL) |
| FAM135B | 51059 | Homo sapiens | family with sequence similarity 135 member B(FAM135B) |
| GRIN3A | 116443 | Homo sapiens | glutamate ionotropic receptor NMDA type subunit 3A(GRIN3A) |
| ZFAT | 57623 | Homo sapiens | zinc finger and AT-hook domain containing(ZFAT) |
| C11ORF16 | 56673 | Homo sapiens | chromosome 11 open reading frame 16(C11orf16) |
| AVL9 | 23080 | Homo sapiens | AVL9 cell migration associated(AVL9) |
| TMEM9B | 56674 | Homo sapiens | TMEM9 domain family member B(TMEM9B) |
| EZH1 | 2145 | Homo sapiens | enhancer of zeste 1 polycomb repressive complex 2 subunit(EZI |
| PIGU | 128869 | Homo sapiens | phosphatidylinositol glycan anchor biosynthesis class U(PIGU) |
| PIGO | 84720 | Homo sapiens | phosphatidylinositol glycan anchor biosynthesis class O(PIGO) |
| PIGN | 23556 | Homo sapiens | phosphatidylinositol glycan anchor biosynthesis class N(PIGN) |
| PIGQ | 9091 | Homo sapiens | phosphatidylinositol glycan anchor biosynthesis class Q(PIGQ) |
| NCAN | 1463 | Homo sapiens | neurocan(NCAN) |
| LRRC4 | 64101 | Homo sapiens | leucine rich repeat containing 4(LRRC4) |
| MRPL38 | 64978 | Homo sapiens | mitochondrial ribosomal protein L38(MRPL38) |
| EFCAB3 | 146779 | Homo sapiens | EF-hand calcium binding domain 3(EFCAB3) |
| EFCAB6 | 64800 | Homo sapiens | EF-hand calcium binding domain 6(EFCAB6) |
| RCSD1 | 92241 | Homo sapiens | RCSD domain containing 1(RCSD1) |
| JPH1 | 56704 | Homo sapiens | junctophilin 1(JPH1) |
| LRRC2 | 79442 | Homo sapiens | leucine rich repeat containing 2(LRRC2) |
| EEA1 | 8411 | Homo sapiens | early endosome antigen 1(EEA1) |
| TMEM47 | 83604 | Homo sapiens | transmembrane protein 47(TMEM47) |
| GPRC6A | 222545 | Homo sapiens | G protein-coupled receptor class C group 6 member A(GPRC6A) |
| IKBKB | 3551 | Homo sapiens | inhibitor of nuclear factor kappa B kinase subunit beta(IKBKB) |

| | | | |
|---|---|---|---|
| LMBR1 | 64327 | Homo sapiens | limb development membrane protein 1(LMBR1) |
| NIPBL | 25836 | Homo sapiens | NIPBL cohesin loading factor(NIPBL) |
| JPH3 | 57338 | Homo sapiens | junctophilin 3(JPH3) |
| TMEM44 | 93109 | Homo sapiens | transmembrane protein 44(TMEM44) |
| LRRC49 | 54839 | Homo sapiens | leucine rich repeat containing 49(LRRC49) |
| NSUN3 | 63899 | Homo sapiens | NOP2/Sun RNA methyltransferase 3(NSUN3) |
| RRAS2 | 22800 | Homo sapiens | RAS related 2(RRAS2) |
| MRPL48 | 51642 | Homo sapiens | mitochondrial ribosomal protein L48(MRPL48) |
| PTGR1 | 22949 | Homo sapiens | prostaglandin reductase 1(PTGR1) |
| TMC3 | 342125 | Homo sapiens | transmembrane channel like 3(TMC3) |
| F7 | 2155 | Homo sapiens | coagulation factor VII(F7) |
| F9 | 2158 | Homo sapiens | coagulation factor IX(F9) |
| PSMA1 | 5682 | Homo sapiens | proteasome 20S subunit alpha 1(PSMA1) |
| BDH1 | 622 | Homo sapiens | 3-hydroxybutyrate dehydrogenase 1(BDH1) |
| LRRC7 | 57554 | Homo sapiens | leucine rich repeat containing 7(LRRC7) |
| PIGK | 10026 | Homo sapiens | phosphatidylinositol glycan anchor biosynthesis class K(PIGK) |
| ZNF236 | 7776 | Homo sapiens | zinc finger protein 236(ZNF236) |
| DSCAML1 | 57453 | Homo sapiens | DS cell adhesion molecule like 1(DSCAML1) |
| MYRF | 745 | Homo sapiens | myelin regulatory factor(MYRF) |
| STAU2 | 27067 | Homo sapiens | staufen double-stranded RNA binding protein 2(STAU2) |
| IDUA | 3425 | Homo sapiens | alpha-L-iduronidase(IDUA) |
| COL11A1 | 1301 | Homo sapiens | collagen type XI alpha 1 chain(COL11A1) |
| SEZ6 | 124925 | Homo sapiens | seizure related 6 homolog(SEZ6) |
| HTT | 3064 | Homo sapiens | huntingtin(HTT) |
| STYXL1 | 51657 | Homo sapiens | serine/threonine/tyrosine interacting like 1(STYXL1) |
| TMEM71 | 137835 | Homo sapiens | transmembrane protein 71(TMEM71) |
| SRPX2 | 27286 | Homo sapiens | sushi repeat containing protein X-linked 2(SRPX2) |
| TMEM67 | 91147 | Homo sapiens | transmembrane protein 67(TMEM67) |
| TMEM68 | 137695 | Homo sapiens | transmembrane protein 68(TMEM68) |
| STAT6 | 6778 | Homo sapiens | signal transducer and activator of transcription 6(STAT6) |
| DMD | 1756 | Homo sapiens | dystrophin(DMD) |
| FLNC | 2318 | Homo sapiens | filamin C(FLNC) |
| RABAC1 | 10567 | Homo sapiens | Rab acceptor 1(RABAC1) |
| LEFTY2 | 7044 | Homo sapiens | left-right determination factor 2(LEFTY2) |
| UTP6 | 55813 | Homo sapiens | UTP6 small subunit processome component(UTP6) |
| USP9X | 8239 | Homo sapiens | ubiquitin specific peptidase 9 X-linked(USP9X) |
| STAT1 | 6772 | Homo sapiens | signal transducer and activator of transcription 1(STAT1) |
| STAT2 | 6773 | Homo sapiens | signal transducer and activator of transcription 2(STAT2) |
| LRRC28 | 123355 | Homo sapiens | leucine rich repeat containing 28(LRRC28) |
| TMEM62 | 80021 | Homo sapiens | transmembrane protein 62(TMEM62) |
| VWA3A | 146177 | Homo sapiens | von Willebrand factor A domain containing 3A(VWA3A) |
| VWA3B | 200403 | Homo sapiens | von Willebrand factor A domain containing 3B(VWA3B) |
| GATAD2B | 57459 | Homo sapiens | GATA zinc finger domain containing 2B(GATAD2B) |
| PSMB9 | 5698 | Homo sapiens | proteasome 20S subunit beta 9(PSMB9) |
| LRRC20 | 55222 | Homo sapiens | leucine rich repeat containing 20(LRRC20) |

| | | | |
|---|---|---|---|
| LETM1 | 3954 | Homo sapiens | leucine zipper and EF-hand containing transmembrane protein 1 |
| SPSB3 | 90864 | Homo sapiens | splA/ryanodine receptor domain and SOCS box containing 3(SPS |
| FABP3 | 2170 | Homo sapiens | fatty acid binding protein 3(FABP3) |
| LYRM4 | 57128 | Homo sapiens | LYR motif containing 4(LYRM4) |
| TTLL1 | 25809 | Homo sapiens | TTL family tubulin polyglutamylase complex subunit L1(TTLL1) |
| RIMBP2 | 23504 | Homo sapiens | RIMS binding protein 2(RIMBP2) |
| SPSB4 | 92369 | Homo sapiens | splA/ryanodine receptor domain and SOCS box containing 4(SPS |
| MYLK2 | 85366 | Homo sapiens | myosin light chain kinase 2(MYLK2) |
| FAM13B | 51306 | Homo sapiens | family with sequence similarity 13 member B(FAM13B) |
| CSE1L | 1434 | Homo sapiens | chromosome segregation 1 like(CSE1L) |
| UBE2D3 | 7323 | Homo sapiens | ubiquitin conjugating enzyme E2 D3(UBE2D3) |
| ZBTB20 | 26137 | Homo sapiens | zinc finger and BTB domain containing 20(ZBTB20) |
| UBE2D1 | 7321 | Homo sapiens | ubiquitin conjugating enzyme E2 D1(UBE2D1) |
| HR | 55806 | Homo sapiens | HR lysine demethylase and nuclear receptor corepressor(HR) |
| GPR61 | 83873 | Homo sapiens | G protein-coupled receptor 61(GPR61) |
| ZBTB22 | 9278 | Homo sapiens | zinc finger and BTB domain containing 22(ZBTB22) |
| SLC9C1 | 285335 | Homo sapiens | solute carrier family 9 member C1(SLC9C1) |
| HDDC3 | 374659 | Homo sapiens | HD domain containing 3(HDDC3) |
| SMCHD1 | 23347 | Homo sapiens | structural maintenance of chromosomes flexible hinge domain c |
| SESN2 | 83667 | Homo sapiens | sestrin 2(SESN2) |
| SLC16A7 | 9194 | Homo sapiens | solute carrier family 16 member 7(SLC16A7) |
| KPNA4 | 3840 | Homo sapiens | karyopherin subunit alpha 4(KPNA4) |
| PMPCA | 23203 | Homo sapiens | peptidase, mitochondrial processing subunit alpha(PMPCA) |
| KPNA5 | 3841 | Homo sapiens | karyopherin subunit alpha 5(KPNA5) |
| ROS1 | 6098 | Homo sapiens | ROS proto-oncogene 1, receptor tyrosine kinase(ROS1) |
| TCP11 | 6954 | Homo sapiens | t-complex 11(TCP11) |
| PPP1R16B | 26051 | Homo sapiens | protein phosphatase 1 regulatory subunit 16B(PPP1R16B) |
| SH3GLB1 | 51100 | Homo sapiens | SH3 domain containing GRB2 like, endophilin B1(SH3GLB1) |
| EPM2A | 7957 | Homo sapiens | EPM2A glucan phosphatase, laforin(EPM2A) |
| CPA2 | 1358 | Homo sapiens | carboxypeptidase A2(CPA2) |
| SH3GLB2 | 56904 | Homo sapiens | SH3 domain containing GRB2 like, endophilin B2(SH3GLB2) |
| IK | 3550 | Homo sapiens | IK cytokine(IK) |
| UBE2E3 | 10477 | Homo sapiens | ubiquitin conjugating enzyme E2 E3(UBE2E3) |
| UBE2E1 | 7324 | Homo sapiens | ubiquitin conjugating enzyme E2 E1(UBE2E1) |
| FOXP4 | 116113 | Homo sapiens | forkhead box P4(FOXP4) |
| FRMD1 | 79981 | Homo sapiens | FERM domain containing 1(FRMD1) |
| FOXP1 | 27086 | Homo sapiens | forkhead box P1(FOXP1) |
| FAM227A | 646851 | Homo sapiens | family with sequence similarity 227 member A(FAM227A) |
| FRMD3 | 257019 | Homo sapiens | FERM domain containing 3(FRMD3) |
| ACAP3 | 116983 | Homo sapiens | ArfGAP with coiled-coil, ankyrin repeat and PH domains 3(ACAP |
| ACAP2 | 23527 | Homo sapiens | ArfGAP with coiled-coil, ankyrin repeat and PH domains 2(ACAP |
| SLC9A4 | 389015 | Homo sapiens | solute carrier family 9 member A4(SLC9A4) |
| SLC9A5 | 6553 | Homo sapiens | solute carrier family 9 member A5(SLC9A5) |
| SLC9A8 | 23315 | Homo sapiens | solute carrier family 9 member A8(SLC9A8) |
| HAT1 | 8520 | Homo sapiens | histone acetyltransferase 1(HAT1) |

| | | | |
|---|---|---|---|
| C18ORF54 | 162681 | Homo sapiens | chromosome 18 open reading frame 54(C18orf54) |
| FSCN3 | 29999 | Homo sapiens | fascin actin-bundling protein 3(FSCN3) |
| KIAA0408 | 9729 | Homo sapiens | KIAA0408(KIAA0408) |
| RNASEH2B | 79621 | Homo sapiens | ribonuclease H2 subunit B(RNASEH2B) |
| C11ORF97 | 643037 | Homo sapiens | chromosome 11 open reading frame 97(C11orf97) |
| MAX | 4149 | Homo sapiens | MYC associated factor X(MAX) |
| TUBGCP2 | 10844 | Homo sapiens | tubulin gamma complex component 2(TUBGCP2) |
| ARL3 | 403 | Homo sapiens | ADP ribosylation factor like GTPase 3(ARL3) |
| ISG20L2 | 81875 | Homo sapiens | interferon stimulated exonuclease gene 20 like 2(ISG20L2) |
| UBFD1 | 56061 | Homo sapiens | ubiquitin family domain containing 1(UBFD1) |
| TNFRSF11A | 8792 | Homo sapiens | TNF receptor superfamily member 11a(TNFRSF11A) |
| THADA | 63892 | Homo sapiens | THADA armadillo repeat containing(THADA) |
| FOXO3 | 2309 | Homo sapiens | forkhead box O3(FOXO3) |
| FHIT | 2272 | Homo sapiens | fragile histidine triad diadenosine triphosphatase(FHIT) |
| FOXO1 | 2308 | Homo sapiens | forkhead box O1(FOXO1) |
| SBNO1 | 55206 | Homo sapiens | strawberry notch homolog 1(SBNO1) |
| STK33 | 65975 | Homo sapiens | serine/threonine kinase 33(STK33) |
| STK35 | 140901 | Homo sapiens | serine/threonine kinase 35(STK35) |
| STK36 | 27148 | Homo sapiens | serine/threonine kinase 36(STK36) |
| ZBBX | 79740 | Homo sapiens | zinc finger B-box domain containing(ZBBX) |
| SLC38A9 | 153129 | Homo sapiens | solute carrier family 38 member 9(SLC38A9) |
| NPTX1 | 4884 | Homo sapiens | neuronal pentraxin 1(NPTX1) |
| SLC38A3 | 10991 | Homo sapiens | solute carrier family 38 member 3(SLC38A3) |
| SOCS7 | 30837 | Homo sapiens | suppressor of cytokine signaling 7(SOCS7) |
| MDN1 | 23195 | Homo sapiens | midasin AAA ATPase 1(MDN1) |
| LRRC69 | 100130742 | Homo sapiens | leucine rich repeat containing 69(LRRC69) |
| LVRN | 206338 | Homo sapiens | laeverin(LVRN) |
| GFPT2 | 9945 | Homo sapiens | glutamine-fructose-6-phosphate transaminase 2(GFPT2) |
| GGCT | 79017 | Homo sapiens | gamma-glutamylcyclotransferase(GGCT) |
| PTPRE | 5791 | Homo sapiens | protein tyrosine phosphatase receptor type E(PTPRE) |
| CXCL11 | 6373 | Homo sapiens | C-X-C motif chemokine ligand 11(CXCL11) |
| STK24 | 8428 | Homo sapiens | serine/threonine kinase 24(STK24) |
| STK26 | 51765 | Homo sapiens | serine/threonine kinase 26(STK26) |
| AAK1 | 22848 | Homo sapiens | AP2 associated kinase 1(AAK1) |
| TUBGCP3 | 10426 | Homo sapiens | tubulin gamma complex component 3(TUBGCP3) |
| ZNF654 | 55279 | Homo sapiens | zinc finger protein 654(ZNF654) |
| MCU | 90550 | Homo sapiens | mitochondrial calcium uniporter(MCU) |
| NFE2L2 | 4780 | Homo sapiens | NFE2 like bZIP transcription factor 2(NFE2L2) |
| SLC25A1 | 6576 | Homo sapiens | solute carrier family 25 member 1(SLC25A1) |
| ZNF512B | 57473 | Homo sapiens | zinc finger protein 512B(ZNF512B) |
| CNTNAP1 | 8506 | Homo sapiens | contactin associated protein 1(CNTNAP1) |
| ST6GALNAC1 | 55808 | Homo sapiens | ST6 N-acetylgalactosaminide alpha-2,6-sialyltransferase 1(ST6( |
| PPP1R13B | 23368 | Homo sapiens | protein phosphatase 1 regulatory subunit 13B(PPP1R13B) |
| SH2D4B | 387694 | Homo sapiens | SH2 domain containing 4B(SH2D4B) |
| FLT3 | 2322 | Homo sapiens | fms related receptor tyrosine kinase 3(FLT3) |

| | | | |
|---|---|---|---|
| CRYBG3 | 131544 | Homo sapiens | crystallin beta-gamma domain containing 3(CRYBG3) |
| SCHIP1 | 29970 | Homo sapiens | schwannomin interacting protein 1(SCHIP1) |
| LRRK1 | 79705 | Homo sapiens | leucine rich repeat kinase 1(LRRK1) |
| FASLG | 356 | Homo sapiens | Fas ligand(FASLG) |
| PIK3CB | 5291 | Homo sapiens | phosphatidylinositol-4,5-bisphosphate 3-kinase catalytic subunit |
| PIK3CG | 5294 | Homo sapiens | phosphatidylinositol-4,5-bisphosphate 3-kinase catalytic subunit |
| NADK2 | 133686 | Homo sapiens | NAD kinase 2, mitochondrial(NADK2) |
| CAPZB | 832 | Homo sapiens | capping actin protein of muscle Z-line subunit beta(CAPZB) |
| ZNF646 | 9726 | Homo sapiens | zinc finger protein 646(ZNF646) |
| PMEPA1 | 56937 | Homo sapiens | prostate transmembrane protein, androgen induced 1(PMEPA1) |
| ME3 | 10873 | Homo sapiens | malic enzyme 3(ME3) |
| DNM1L | 10059 | Homo sapiens | dynamin 1 like(DNM1L) |
| ZNF641 | 121274 | Homo sapiens | zinc finger protein 641(ZNF641) |
| CTNNBL1 | 56259 | Homo sapiens | catenin beta like 1(CTNNBL1) |
| DIS3L2 | 129563 | Homo sapiens | DIS3 like 3'-5' exoribonuclease 2(DIS3L2) |
| EMX2 | 2018 | Homo sapiens | empty spiracles homeobox 2(EMX2) |
| OVCH1 | 341350 | Homo sapiens | ovochymase 1(OVCH1) |
| GKN1 | 56287 | Homo sapiens | gastrokine 1(GKN1) |
| SH2D3C | 10044 | Homo sapiens | SH2 domain containing 3C(SH2D3C) |
| PARD3B | 117583 | Homo sapiens | par-3 family cell polarity regulator beta(PARD3B) |
| SLCO1C1 | 53919 | Homo sapiens | solute carrier organic anion transporter family member 1C1(SLC |
| SLC25A15 | 10166 | Homo sapiens | solute carrier family 25 member 15(SLC25A15) |
| NT5C3B | 115024 | Homo sapiens | 5'-nucleotidase, cytosolic IIIB(NT5C3B) |
| SLC25A17 | 10478 | Homo sapiens | solute carrier family 25 member 17(SLC25A17) |
| NPC1 | 4864 | Homo sapiens | NPC intracellular cholesterol transporter 1(NPC1) |
| SNPH | 9751 | Homo sapiens | syntaphilin(SNPH) |
| BIRC6 | 57448 | Homo sapiens | baculoviral IAP repeat containing 6(BIRC6) |
| DSG2 | 1829 | Homo sapiens | desmoglein 2(DSG2) |
| SLC25A11 | 8402 | Homo sapiens | solute carrier family 25 member 11(SLC25A11) |
| MET | 4233 | Homo sapiens | MET proto-oncogene, receptor tyrosine kinase(MET) |
| CNTNAP5 | 129684 | Homo sapiens | contactin associated protein family member 5(CNTNAP5) |
| RBPJL | 11317 | Homo sapiens | recombination signal binding protein for immunoglobulin kappa |
| FOCAD | 54914 | Homo sapiens | focadhesin(FOCAD) |
| SEC23A | 10484 | Homo sapiens | SEC23 homolog A, COPII coat complex component(SEC23A) |
| ITIH2 | 3698 | Homo sapiens | inter-alpha-trypsin inhibitor heavy chain 2(ITIH2) |
| PCOLCE2 | 26577 | Homo sapiens | procollagen C-endopeptidase enhancer 2(PCOLCE2) |
| PPM1L | 151742 | Homo sapiens | protein phosphatase, Mg2+/Mn2+ dependent 1L(PPM1L) |
| LRP5 | 4041 | Homo sapiens | LDL receptor related protein 5(LRP5) |
| USH1C | 10083 | Homo sapiens | USH1 protein network component harmonin(USH1C) |
| PPM1H | 57460 | Homo sapiens | protein phosphatase, Mg2+/Mn2+ dependent 1H(PPM1H) |
| FOXK2 | 3607 | Homo sapiens | forkhead box K2(FOXK2) |
| KATNB1 | 10300 | Homo sapiens | katanin regulatory subunit B1(KATNB1) |
| PARD6G | 84552 | Homo sapiens | par-6 family cell polarity regulator gamma(PARD6G) |
| LRP3 | 4037 | Homo sapiens | LDL receptor related protein 3(LRP3) |
| HSD17B12 | 51144 | Homo sapiens | hydroxysteroid 17-beta dehydrogenase 12(HSD17B12) |

| | | | |
|---|---|---|---|
| LRP2 | 4036 | Homo sapiens | LDL receptor related protein 2(LRP2) |
| ZBTB44 | 29068 | Homo sapiens | zinc finger and BTB domain containing 44(ZBTB44) |
| BARX1 | 56033 | Homo sapiens | BARX homeobox 1(BARX1) |
| LRP8 | 7804 | Homo sapiens | LDL receptor related protein 8(LRP8) |
| PPM1F | 9647 | Homo sapiens | protein phosphatase, Mg2+/Mn2+ dependent 1F(PPM1F) |
| LRP6 | 4040 | Homo sapiens | LDL receptor related protein 6(LRP6) |
| SLC25A27 | 9481 | Homo sapiens | solute carrier family 25 member 27(SLC25A27) |
| SLC25A26 | 115286 | Homo sapiens | solute carrier family 25 member 26(SLC25A26) |
| CNNM1 | 26507 | Homo sapiens | cyclin and CBS domain divalent metal cation transport mediator |
| PTPRZ1 | 5803 | Homo sapiens | protein tyrosine phosphatase receptor type Z1(PTPRZ1) |
| FCHO2 | 115548 | Homo sapiens | FCH and mu domain containing endocytic adaptor 2(FCHO2) |
| SEC23B | 10483 | Homo sapiens | SEC23 homolog B, COPII coat complex component(SEC23B) |
| SLC25A25 | 114789 | Homo sapiens | solute carrier family 25 member 25(SLC25A25) |
| BEND5 | 79656 | Homo sapiens | BEN domain containing 5(BEND5) |
| CYB5A | 1528 | Homo sapiens | cytochrome b5 type A(CYB5A) |
| PTPN1 | 5770 | Homo sapiens | protein tyrosine phosphatase non-receptor type 1(PTPN1) |
| SLC12A3 | 6559 | Homo sapiens | solute carrier family 12 member 3(SLC12A3) |
| SLC12A4 | 6560 | Homo sapiens | solute carrier family 12 member 4(SLC12A4) |
| NDUFA6 | 4700 | Homo sapiens | NADH:ubiquinone oxidoreductase subunit A6(NDUFA6) |
| MBOAT7 | 79143 | Homo sapiens | membrane bound O-acyltransferase domain containing 7(MBOA |
| TIE1 | 7075 | Homo sapiens | tyrosine kinase with immunoglobulin like and EGF like domains |
| NDUFA2 | 4695 | Homo sapiens | NADH:ubiquinone oxidoreductase subunit A2(NDUFA2) |
| MBOAT2 | 129642 | Homo sapiens | membrane bound O-acyltransferase domain containing 2(MBOA |
| ZNF618 | 114991 | Homo sapiens | zinc finger protein 618(ZNF618) |
| NOVA1 | 4857 | Homo sapiens | NOVA alternative splicing regulator 1(NOVA1) |
| SLC25A32 | 81034 | Homo sapiens | solute carrier family 25 member 32(SLC25A32) |
| PTPN4 | 5775 | Homo sapiens | protein tyrosine phosphatase non-receptor type 4(PTPN4) |
| ALDH8A1 | 64577 | Homo sapiens | aldehyde dehydrogenase 8 family member A1(ALDH8A1) |
| PTPN3 | 5774 | Homo sapiens | protein tyrosine phosphatase non-receptor type 3(PTPN3) |
| SPON2 | 10417 | Homo sapiens | spondin 2(SPON2) |
| AHCYL2 | 23382 | Homo sapiens | adenosylhomocysteinase like 2(AHCYL2) |
| SPON1 | 10418 | Homo sapiens | spondin 1(SPON1) |
| CCNI | 10983 | Homo sapiens | cyclin I(CCNI) |
| TMEM41B | 440026 | Homo sapiens | transmembrane protein 41B(TMEM41B) |
| EIF4A3 | 9775 | Homo sapiens | eukaryotic translation initiation factor 4A3(EIF4A3) |
| PREP | 5550 | Homo sapiens | prolyl endopeptidase(PREP) |
| TXNDC11 | 51061 | Homo sapiens | thioredoxin domain containing 11(TXNDC11) |
| GCC2 | 9648 | Homo sapiens | GRIP and coiled-coil domain containing 2(GCC2) |
| CYB5D1 | 124637 | Homo sapiens | cytochrome b5 domain containing 1(CYB5D1) |
| UBE2L3 | 7332 | Homo sapiens | ubiquitin conjugating enzyme E2 L3(UBE2L3) |
| POF1B | 79983 | Homo sapiens | POF1B actin binding protein(POF1B) |
| OGFOD1 | 55239 | Homo sapiens | 2-oxoglutarate and iron dependent oxygenase domain containi |
| FAM53C | 51307 | Homo sapiens | family with sequence similarity 53 member C(FAM53C) |
| FAM53B | 9679 | Homo sapiens | family with sequence similarity 53 member B(FAM53B) |
| EPHB2 | 2048 | Homo sapiens | EPH receptor B2(EPHB2) |

| | | | |
|---|---|---|---|
| SEC63 | 11231 | Homo sapiens | SEC63 homolog, protein translocation regulator(SEC63) |
| EPHA5 | 2044 | Homo sapiens | EPH receptor A5(EPHA5) |
| EPHA7 | 2045 | Homo sapiens | EPH receptor A7(EPHA7) |
| ST6GAL1 | 6480 | Homo sapiens | ST6 beta-galactoside alpha-2,6-sialyltransferase 1(ST6GAL1) |
| MUSK | 4593 | Homo sapiens | muscle associated receptor tyrosine kinase(MUSK) |
| SFMBT1 | 51460 | Homo sapiens | Scm like with four mbt domains 1(SFMBT1) |
| EPHA8 | 2046 | Homo sapiens | EPH receptor A8(EPHA8) |
| SCAF11 | 9169 | Homo sapiens | SR-related CTD associated factor 11(SCAF11) |
| ENTPD6 | 955 | Homo sapiens | ectonucleoside triphosphate diphosphohydrolase 6(ENTPD6) |
| NSMCE2 | 286053 | Homo sapiens | NSE2 (MMS21) homolog, SMC5-SMC6 complex SUMO ligase(NS |
| ENTPD8 | 377841 | Homo sapiens | ectonucleoside triphosphate diphosphohydrolase 8(ENTPD8) |
| MMD2 | 221938 | Homo sapiens | monocyte to macrophage differentiation associated 2(MMD2) |
| MMD2 | 100505381 | Homo sapiens | Miyoshi muscular dystrophy 2(MMD2) |
| ANK2 | 287 | Homo sapiens | ankyrin 2(ANK2) |
| ANK1 | 286 | Homo sapiens | ankyrin 1(ANK1) |
| FNDC7 | 163479 | Homo sapiens | fibronectin type III domain containing 7(FNDC7) |
| GCAT | 23464 | Homo sapiens | glycine C-acetyltransferase(GCAT) |
| GNPAT | 8443 | Homo sapiens | glyceronephosphate O-acyltransferase(GNPAT) |
| SPATA16 | 83893 | Homo sapiens | spermatogenesis associated 16(SPATA16) |
| THAP9 | 79725 | Homo sapiens | THAP domain containing 9(THAP9) |
| ULK2 | 9706 | Homo sapiens | unc-51 like autophagy activating kinase 2(ULK2) |
| ALPK3 | 57538 | Homo sapiens | alpha kinase 3(ALPK3) |
| MXRA7 | 439921 | Homo sapiens | matrix remodeling associated 7(MXRA7) |
| ALPK2 | 115701 | Homo sapiens | alpha kinase 2(ALPK2) |
| ALPK1 | 80216 | Homo sapiens | alpha kinase 1(ALPK1) |
| L3MBTL1 | 26013 | Homo sapiens | L3MBTL histone methyl-lysine binding protein 1(L3MBTL1) |
| PPP1R37 | 284352 | Homo sapiens | protein phosphatase 1 regulatory subunit 37(PPP1R37) |
| L3MBTL3 | 84456 | Homo sapiens | L3MBTL histone methyl-lysine binding protein 3(L3MBTL3) |
| PIK3R3 | 8503 | Homo sapiens | phosphoinositide-3-kinase regulatory subunit 3(PIK3R3) |
| SNX3 | 8724 | Homo sapiens | sorting nexin 3(SNX3) |
| DHX30 | 22907 | Homo sapiens | DExH-box helicase 30(DHX30) |
| SNX2 | 6643 | Homo sapiens | sorting nexin 2(SNX2) |
| ERI1 | 90459 | Homo sapiens | exoribonuclease 1(ERI1) |
| TSPAN9 | 10867 | Homo sapiens | tetraspanin 9(TSPAN9) |
| ZNF704 | 619279 | Homo sapiens | zinc finger protein 704(ZNF704) |
| DHX35 | 60625 | Homo sapiens | DEAH-box helicase 35(DHX35) |
| ERI3 | 79033 | Homo sapiens | ERI1 exoribonuclease family member 3(ERI3) |
| DHX37 | 57647 | Homo sapiens | DEAH-box helicase 37(DHX37) |
| ST8SIA5 | 29906 | Homo sapiens | ST8 alpha-N-acetyl-neuraminide alpha-2,8-sialyltransferase 5(S |
| DYM | 54808 | Homo sapiens | dymeclin(DYM) |
| HIVEP3 | 59269 | Homo sapiens | HIVEP zinc finger 3(HIVEP3) |
| RAB11FIP3 | 9727 | Homo sapiens | RAB11 family interacting protein 3(RAB11FIP3) |
| RAB11FIP4 | 84440 | Homo sapiens | RAB11 family interacting protein 4(RAB11FIP4) |
| SEC31A | 22872 | Homo sapiens | SEC31 homolog A, COPII coat complex component(SEC31A) |
| RAB11FIP5 | 26056 | Homo sapiens | RAB11 family interacting protein 5(RAB11FIP5) |

| | | | |
|---|---|---|---|
| SNX6 | 58533 | Homo sapiens | sorting nexin 6(SNX6) |
| TMEM86A | 144110 | Homo sapiens | transmembrane protein 86A(TMEM86A) |
| CRADD | 8738 | Homo sapiens | CASP2 and RIPK1 domain containing adaptor with death domai |
| XRCC4 | 7518 | Homo sapiens | X-ray repair cross complementing 4(XRCC4) |
| ST8SIA1 | 6489 | Homo sapiens | ST8 alpha-N-acetyl-neuraminide alpha-2,8-sialyltransferase 1(S |
| INTS3 | 65123 | Homo sapiens | integrator complex subunit 3(INTS3) |
| IQSEC3 | 440073 | Homo sapiens | IQ motif and Sec7 domain ArfGEF 3(IQSEC3) |
| INTS2 | 57508 | Homo sapiens | integrator complex subunit 2(INTS2) |
| EAF1 | 85403 | Homo sapiens | ELL associated factor 1(EAF1) |
| PILRA | 29992 | Homo sapiens | paired immunoglobin like type 2 receptor alpha(PILRA) |
| TG | 7038 | Homo sapiens | thyroglobulin(TG) |
| WNK1 | 65125 | Homo sapiens | WNK lysine deficient protein kinase 1(WNK1) |
| WNK2 | 65268 | Homo sapiens | WNK lysine deficient protein kinase 2(WNK2) |
| ACER3 | 55331 | Homo sapiens | alkaline ceramidase 3(ACER3) |
| INTS4 | 92105 | Homo sapiens | integrator complex subunit 4(INTS4) |
| RGL3 | 57139 | Homo sapiens | ral guanine nucleotide dissociation stimulator like 3(RGL3) |
| COL9A2 | 1298 | Homo sapiens | collagen type IX alpha 2 chain(COL9A2) |
| INTS6 | 26512 | Homo sapiens | integrator complex subunit 6(INTS6) |
| MKX | 283078 | Homo sapiens | mohawk homeobox(MKX) |
| INTS9 | 55756 | Homo sapiens | integrator complex subunit 9(INTS9) |
| RCOR1 | 23186 | Homo sapiens | REST corepressor 1(RCOR1) |
| FKBP11 | 51303 | Homo sapiens | FKBP prolyl isomerase 11(FKBP11) |
| PTPRT | 11122 | Homo sapiens | protein tyrosine phosphatase receptor type T(PTPRT) |
| FKBP10 | 60681 | Homo sapiens | FKBP prolyl isomerase 10(FKBP10) |
| INTS12 | 57117 | Homo sapiens | integrator complex subunit 12(INTS12) |
| PTPRU | 10076 | Homo sapiens | protein tyrosine phosphatase receptor type U(PTPRU) |
| MOGAT2 | 80168 | Homo sapiens | monoacylglycerol O-acyltransferase 2(MOGAT2) |
| PTPRN | 5798 | Homo sapiens | protein tyrosine phosphatase receptor type N(PTPRN) |
| PTPRO | 5800 | Homo sapiens | protein tyrosine phosphatase receptor type O(PTPRO) |
| PTPRM | 5797 | Homo sapiens | protein tyrosine phosphatase receptor type M(PTPRM) |
| MIA3 | 375056 | Homo sapiens | MIA SH3 domain ER export factor 3(MIA3) |
| DOC2B | 8447 | Homo sapiens | double C2 domain beta(DOC2B) |
| IDE | 3416 | Homo sapiens | insulin degrading enzyme(IDE) |
| PCSK7 | 9159 | Homo sapiens | proprotein convertase subtilisin/kexin type 7(PCSK7) |
| BRCA2 | 675 | Homo sapiens | BRCA2 DNA repair associated(BRCA2) |
| PCSK6 | 5046 | Homo sapiens | proprotein convertase subtilisin/kexin type 6(PCSK6) |
| PCSK5 | 5125 | Homo sapiens | proprotein convertase subtilisin/kexin type 5(PCSK5) |
| CHRDL1 | 91851 | Homo sapiens | chordin like 1(CHRDL1) |
| AP2A2 | 161 | Homo sapiens | adaptor related protein complex 2 subunit alpha 2(AP2A2) |
| IMMP2L | 83943 | Homo sapiens | inner mitochondrial membrane peptidase subunit 2(IMMP2L) |
| SPTLC1 | 10558 | Homo sapiens | serine palmitoyltransferase long chain base subunit 1(SPTLC1) |
| SPTLC2 | 9517 | Homo sapiens | serine palmitoyltransferase long chain base subunit 2(SPTLC2) |
| CTSL | 1514 | Homo sapiens | cathepsin L(CTSL) |
| RSRC1 | 51319 | Homo sapiens | arginine and serine rich coiled-coil 1(RSRC1) |
| REV3L | 5980 | Homo sapiens | REV3 like, DNA directed polymerase zeta catalytic subunit(REV3 |

| | | | |
|---|---|---|---|
| CD38 | 952 | Homo sapiens | CD38 molecule(CD38) |
| ARIH1 | 25820 | Homo sapiens | ariadne RBR E3 ubiquitin protein ligase 1(ARIH1) |
| ACADM | 34 | Homo sapiens | acyl-CoA dehydrogenase medium chain(ACADM) |
| CTSD | 1509 | Homo sapiens | cathepsin D(CTSD) |
| CTSB | 1508 | Homo sapiens | cathepsin B(CTSB) |
| DBNL | 28988 | Homo sapiens | drebrin like(DBNL) |
| NCOA3 | 8202 | Homo sapiens | nuclear receptor coactivator 3(NCOA3) |
| MINK1 | 50488 | Homo sapiens | misshapen like kinase 1(MINK1) |
| ANO6 | 196527 | Homo sapiens | anoctamin 6(ANO6) |
| SORCS1 | 114815 | Homo sapiens | sortilin related VPS10 domain containing receptor 1(SORCS1) |
| COMMD1 | 150684 | Homo sapiens | copper metabolism domain containing 1(COMMD1) |
| SORCS3 | 22986 | Homo sapiens | sortilin related VPS10 domain containing receptor 3(SORCS3) |
| TMEM150C | 441027 | Homo sapiens | transmembrane protein 150C(TMEM150C) |
| SORCS2 | 57537 | Homo sapiens | sortilin related VPS10 domain containing receptor 2(SORCS2) |
| SUPT3H | 8464 | Homo sapiens | SPT3 homolog, SAGA and STAGA complex component(SUPT3H) |
| ENAH | 55740 | Homo sapiens | ENAH actin regulator(ENAH) |
| BANK1 | 55024 | Homo sapiens | B cell scaffold protein with ankyrin repeats 1(BANK1) |
| NCOA7 | 135112 | Homo sapiens | nuclear receptor coactivator 7(NCOA7) |
| KIF2C | 11004 | Homo sapiens | kinesin family member 2C(KIF2C) |
| CAPS2 | 84698 | Homo sapiens | calcyphosine 2(CAPS2) |
| POPDC2 | 64091 | Homo sapiens | popeye domain containing 2(POPDC2) |
| FOXC2 | 2303 | Homo sapiens | forkhead box C2(FOXC2) |
| STX12 | 23673 | Homo sapiens | syntaxin 12(STX12) |
| RABGAP1 | 23637 | Homo sapiens | RAB GTPase activating protein 1(RABGAP1) |
| TMPRSS6 | 164656 | Homo sapiens | transmembrane serine protease 6(TMPRSS6) |
| CCDC103 | 388389 | Homo sapiens | coiled-coil domain containing 103(CCDC103) |
| DOT1L | 84444 | Homo sapiens | DOT1 like histone lysine methyltransferase(DOT1L) |
| STX18 | 53407 | Homo sapiens | syntaxin 18(STX18) |
| IGHMBP2 | 3508 | Homo sapiens | immunoglobulin mu DNA binding protein 2(IGHMBP2) |
| PRKX | 5613 | Homo sapiens | protein kinase cAMP-dependent X-linked catalytic subunit(PRKX |
| RXRA | 6256 | Homo sapiens | retinoid X receptor alpha(RXRA) |
| ALS2CL | 259173 | Homo sapiens | ALS2 C-terminal like(ALS2CL) |
| GPC1 | 2817 | Homo sapiens | glypican 1(GPC1) |
| HSF1 | 3297 | Homo sapiens | heat shock transcription factor 1(HSF1) |
| GPC4 | 2239 | Homo sapiens | glypican 4(GPC4) |
| HEATR9 | 256957 | Homo sapiens | HEAT repeat containing 9(HEATR9) |
| NPDC1 | 56654 | Homo sapiens | neural proliferation, differentiation and control 1(NPDC1) |
| PLEKHA1 | 59338 | Homo sapiens | pleckstrin homology domain containing A1(PLEKHA1) |
| ERCC6L2 | 375748 | Homo sapiens | ERCC excision repair 6 like 2(ERCC6L2) |
| NUBPL | 80224 | Homo sapiens | NUBP iron-sulfur cluster assembly factor, mitochondrial(NUBPL) |
| RFTN1 | 23180 | Homo sapiens | raftlin, lipid raft linker 1(RFTN1) |
| UBE2G2 | 7327 | Homo sapiens | ubiquitin conjugating enzyme E2 G2(UBE2G2) |
| AP2B1 | 163 | Homo sapiens | adaptor related protein complex 2 subunit beta 1(AP2B1) |
| PLEKHA7 | 144100 | Homo sapiens | pleckstrin homology domain containing A7(PLEKHA7) |
| NLE1 | 54475 | Homo sapiens | notchless homolog 1(NLE1) |

| | | | |
|---|---|---|---|
| PLEKHA8 | 84725 | Homo sapiens | pleckstrin homology domain containing A8(PLEKHA8) |
| MIB1 | 57534 | Homo sapiens | MIB E3 ubiquitin protein ligase 1(MIB1) |
| ALDH4A1 | 8659 | Homo sapiens | aldehyde dehydrogenase 4 family member A1(ALDH4A1) |
| DPY19L1 | 23333 | Homo sapiens | dpy-19 like C-mannosyltransferase 1(DPY19L1) |
| GFI1B | 8328 | Homo sapiens | growth factor independent 1B transcriptional repressor(GFI1B) |
| NLRP12 | 91662 | Homo sapiens | NLR family pyrin domain containing 12(NLRP12) |
| CMSS1 | 84319 | Homo sapiens | cms1 ribosomal small subunit homolog(CMSS1) |
| DPY19L3 | 147991 | Homo sapiens | dpy-19 like C-mannosyltransferase 3(DPY19L3) |
| ZDHHC9 | 51114 | Homo sapiens | zinc finger DHHC-type palmitoyltransferase 9(ZDHHC9) |
| DPY19L4 | 286148 | Homo sapiens | dpy-19 like 4(DPY19L4) |
| NEDD4 | 4734 | Homo sapiens | NEDD4 E3 ubiquitin protein ligase(NEDD4) |
| KIF26B | 55083 | Homo sapiens | kinesin family member 26B(KIF26B) |
| STK17B | 9262 | Homo sapiens | serine/threonine kinase 17b(STK17B) |
| FAM171B | 165215 | Homo sapiens | family with sequence similarity 171 member B(FAM171B) |
| DBR1 | 51163 | Homo sapiens | debranching RNA lariats 1(DBR1) |
| KLHL36 | 79786 | Homo sapiens | kelch like family member 36(KLHL36) |
| CD81 | 975 | Homo sapiens | CD81 molecule(CD81) |
| CD80 | 941 | Homo sapiens | CD80 molecule(CD80) |
| GEMIN2 | 8487 | Homo sapiens | gem nuclear organelle associated protein 2(GEMIN2) |
| CCDC169 | 728591 | Homo sapiens | coiled-coil domain containing 169(CCDC169) |
| JADE1 | 79960 | Homo sapiens | jade family PHD finger 1(JADE1) |
| KNDC1 | 85442 | Homo sapiens | kinase non-catalytic C-lobe domain containing 1(KNDC1) |
| GXYLT2 | 727936 | Homo sapiens | glucoside xylosyltransferase 2(GXYLT2) |
| PHRF1 | 57661 | Homo sapiens | PHD and ring finger domains 1(PHRF1) |
| IFI44L | 10964 | Homo sapiens | interferon induced protein 44 like(IFI44L) |
| GIMAP8 | 155038 | Homo sapiens | GTPase, IMAP family member 8(GIMAP8) |
| CLGN | 1047 | Homo sapiens | calmegin(CLGN) |
| CAPN9 | 10753 | Homo sapiens | calpain 9(CAPN9) |
| RGS3 | 5998 | Homo sapiens | regulator of G protein signaling 3(RGS3) |
| CAPN7 | 23473 | Homo sapiens | calpain 7(CAPN7) |
| DUOXA1 | 90527 | Homo sapiens | dual oxidase maturation factor 1(DUOXA1) |
| KIF5B | 3799 | Homo sapiens | kinesin family member 5B(KIF5B) |
| MPZ | 4359 | Homo sapiens | myelin protein zero(MPZ) |
| CAPN5 | 726 | Homo sapiens | calpain 5(CAPN5) |
| PALM | 5064 | Homo sapiens | paralemmin(PALM) |
| CAPN2 | 824 | Homo sapiens | calpain 2(CAPN2) |
| CAPN3 | 825 | Homo sapiens | calpain 3(CAPN3) |
| OSTF1 | 26578 | Homo sapiens | osteoclast stimulating factor 1(OSTF1) |
| SBF2 | 81846 | Homo sapiens | SET binding factor 2(SBF2) |
| RGS9 | 8787 | Homo sapiens | regulator of G protein signaling 9(RGS9) |
| SLC30A6 | 55676 | Homo sapiens | solute carrier family 30 member 6(SLC30A6) |
| SLC30A9 | 10463 | Homo sapiens | solute carrier family 30 member 9(SLC30A9) |
| CD93 | 22918 | Homo sapiens | CD93 molecule(CD93) |
| SLC30A3 | 7781 | Homo sapiens | solute carrier family 30 member 3(SLC30A3) |
| SCARA5 | 286133 | Homo sapiens | scavenger receptor class A member 5(SCARA5) |

| | | | |
|---|---|---|---|
| THEMIS | 387357 | Homo sapiens | thymocyte selection associated(THEMIS) |
| SSU72 | 29101 | Homo sapiens | SSU72 homolog, RNA polymerase II CTD phosphatase(SSU72) |
| CYBA | 1535 | Homo sapiens | cytochrome b-245 alpha chain(CYBA) |
| RPGRIP1L | 23322 | Homo sapiens | RPGRIP1 like(RPGRIP1L) |
| RUNDC3B | 154661 | Homo sapiens | RUN domain containing 3B(RUNDC3B) |
| NPNT | 255743 | Homo sapiens | nephronectin(NPNT) |
| ATAD3A | 55210 | Homo sapiens | ATPase family AAA domain containing 3A(ATAD3A) |
| KLHL41 | 10324 | Homo sapiens | kelch like family member 41(KLHL41) |
| MID2 | 11043 | Homo sapiens | midline 2(MID2) |
| CAMKMT | 79823 | Homo sapiens | calmodulin-lysine N-methyltransferase(CAMKMT) |
| UBE2V2 | 7336 | Homo sapiens | ubiquitin conjugating enzyme E2 V2(UBE2V2) |
| MASP2 | 10747 | Homo sapiens | MBL associated serine protease 2(MASP2) |
| CCDC152 | 100129792 | Homo sapiens | coiled-coil domain containing 152(CCDC152) |
| CCDC150 | 284992 | Homo sapiens | coiled-coil domain containing 150(CCDC150) |
| KLHL18 | 23276 | Homo sapiens | kelch like family member 18(KLHL18) |
| NDRG4 | 65009 | Homo sapiens | NDRG family member 4(NDRG4) |
| PSMD12 | 5718 | Homo sapiens | proteasome 26S subunit, non-ATPase 12(PSMD12) |
| NDRG3 | 57446 | Homo sapiens | NDRG family member 3(NDRG3) |
| KLHL14 | 57565 | Homo sapiens | kelch like family member 14(KLHL14) |
| ARHGEF28 | 64283 | Homo sapiens | Rho guanine nucleotide exchange factor 28(ARHGEF28) |
| KLHL12 | 59349 | Homo sapiens | kelch like family member 12(KLHL12) |
| PDS5B | 23047 | Homo sapiens | PDS5 cohesin associated factor B(PDS5B) |
| SLC1A6 | 6511 | Homo sapiens | solute carrier family 1 member 6(SLC1A6) |
| RPF2 | 84154 | Homo sapiens | ribosome production factor 2 homolog(RPF2) |
| NDRG1 | 10397 | Homo sapiens | N-myc downstream regulated 1(NDRG1) |
| SUPT6H | 6830 | Homo sapiens | SPT6 homolog, histone chaperone and transcription elongation f |
| TAF5L | 27097 | Homo sapiens | TATA-box binding protein associated factor 5 like(TAF5L) |
| MCHR2 | 84539 | Homo sapiens | melanin concentrating hormone receptor 2(MCHR2) |
| RPF1 | 80135 | Homo sapiens | ribosome production factor 1 homolog(RPF1) |
| BRINP2 | 57795 | Homo sapiens | BMP/retinoic acid inducible neural specific 2(BRINP2) |
| BRINP1 | 1620 | Homo sapiens | BMP/retinoic acid inducible neural specific 1(BRINP1) |
| NPHP1 | 4867 | Homo sapiens | nephrocystin 1(NPHP1) |
| CAMTA1 | 23261 | Homo sapiens | calmodulin binding transcription activator 1(CAMTA1) |
| KIF3C | 3797 | Homo sapiens | kinesin family member 3C(KIF3C) |
| NDUFV2 | 4729 | Homo sapiens | NADH:ubiquinone oxidoreductase core subunit V2(NDUFV2) |
| WNT3 | 7473 | Homo sapiens | Wnt family member 3(WNT3) |
| GGT7 | 2686 | Homo sapiens | gamma-glutamyltransferase 7(GGT7) |
| SPATA6L | 55064 | Homo sapiens | spermatogenesis associated 6 like(SPATA6L) |
| PHF10 | 55274 | Homo sapiens | PHD finger protein 10(PHF10) |
| ARHGEF38 | 54848 | Homo sapiens | Rho guanine nucleotide exchange factor 38(ARHGEF38) |
| CRIM1 | 51232 | Homo sapiens | cysteine rich transmembrane BMP regulator 1(CRIM1) |
| SSH2 | 85464 | Homo sapiens | slingshot protein phosphatase 2(SSH2) |
| QDPR | 5860 | Homo sapiens | quinoid dihydropteridine reductase(QDPR) |
| RCL1 | 10171 | Homo sapiens | RNA terminal phosphate cyclase like 1(RCL1) |
| CPS1 | 1373 | Homo sapiens | carbamoyl-phosphate synthase 1(CPS1) |

| | | | |
|---|---|---|---|
| COL5A1 | 1289 | Homo sapiens | collagen type V alpha 1 chain(COL5A1) |
| KIF4A | 24137 | Homo sapiens | kinesin family member 4A(KIF4A) |
| TAF4B | 6875 | Homo sapiens | TATA-box binding protein associated factor 4b(TAF4B) |
| TAB1 | 10454 | Homo sapiens | TGF-beta activated kinase 1 (MAP3K7) binding protein 1(TAB1) |
| STT3B | 201595 | Homo sapiens | STT3 oligosaccharyltransferase complex catalytic subunit B(STT3 |
| OSTN | 344901 | Homo sapiens | osteocrin(OSTN) |
| SERGEF | 26297 | Homo sapiens | secretion regulating guanine nucleotide exchange factor(SERGEI |
| DDX49 | 54555 | Homo sapiens | DEAD-box helicase 49(DDX49) |
| DDX46 | 9879 | Homo sapiens | DEAD-box helicase 46(DDX46) |
| MAST4 | 375449 | Homo sapiens | microtubule associated serine/threonine kinase family member |
| CPXM2 | 119587 | Homo sapiens | carboxypeptidase X, M14 family member 2(CPXM2) |
| CHD9 | 80205 | Homo sapiens | chromodomain helicase DNA binding protein 9(CHD9) |
| MAST2 | 23139 | Homo sapiens | microtubule associated serine/threonine kinase 2(MAST2) |
| CHD8 | 57680 | Homo sapiens | chromodomain helicase DNA binding protein 8(CHD8) |
| CHD6 | 84181 | Homo sapiens | chromodomain helicase DNA binding protein 6(CHD6) |
| PSIP1 | 11168 | Homo sapiens | PC4 and SRSF1 interacting protein 1(PSIP1) |
| ARHGEF10L | 55160 | Homo sapiens | Rho guanine nucleotide exchange factor 10 like(ARHGEF10L) |
| CHD3 | 1107 | Homo sapiens | chromodomain helicase DNA binding protein 3(CHD3) |
| PPP1R9A | 55607 | Homo sapiens | protein phosphatase 1 regulatory subunit 9A(PPP1R9A) |
| AFF1 | 4299 | Homo sapiens | ALF transcription elongation factor 1(AFF1) |
| GPHN | 10243 | Homo sapiens | gephyrin(GPHN) |
| PPP1R9B | 84687 | Homo sapiens | protein phosphatase 1 regulatory subunit 9B(PPP1R9B) |
| RPGR | 6103 | Homo sapiens | retinitis pigmentosa GTPase regulator(RPGR) |
| AFF2 | 2334 | Homo sapiens | ALF transcription elongation factor 2(AFF2) |
| CALB2 | 794 | Homo sapiens | calbindin 2(CALB2) |
| SEC61A1 | 29927 | Homo sapiens | SEC61 translocon subunit alpha 1(SEC61A1) |
| RNF212B | 100507650 | Homo sapiens | ring finger protein 212B(RNF212B) |
| BCL7A | 605 | Homo sapiens | BAF chromatin remodeling complex subunit BCL7A(BCL7A) |
| CFAP47 | 286464 | Homo sapiens | cilia and flagella associated protein 47(CFAP47) |
| TIMM17A | 10440 | Homo sapiens | translocase of inner mitochondrial membrane 17A(TIMM17A) |
| CFAP46 | 54777 | Homo sapiens | cilia and flagella associated protein 46(CFAP46) |
| SCN5A | 6331 | Homo sapiens | sodium voltage-gated channel alpha subunit 5(SCN5A) |
| CFAP43 | 80217 | Homo sapiens | cilia and flagella associated protein 43(CFAP43) |
| SCARF2 | 91179 | Homo sapiens | scavenger receptor class F member 2(SCARF2) |
| CFAP44 | 55779 | Homo sapiens | cilia and flagella associated protein 44(CFAP44) |
| RBM33 | 155435 | Homo sapiens | RNA binding motif protein 33(RBM33) |
| KRT222 | 125113 | Homo sapiens | keratin 222(KRT222) |
| HS3ST3B1 | 9953 | Homo sapiens | heparan sulfate-glucosamine 3-sulfotransferase 3B1(HS3ST3B1) |
| DDX59 | 83479 | Homo sapiens | DEAD-box helicase 59(DDX59) |
| ARHGEF12 | 23365 | Homo sapiens | Rho guanine nucleotide exchange factor 12(ARHGEF12) |
| NCBP1 | 4686 | Homo sapiens | nuclear cap binding protein subunit 1(NCBP1) |
| GLRX3 | 10539 | Homo sapiens | glutaredoxin 3(GLRX3) |
| PLA2G4E | 123745 | Homo sapiens | phospholipase A2 group IVE(PLA2G4E) |
| ARHGEF17 | 9828 | Homo sapiens | Rho guanine nucleotide exchange factor 17(ARHGEF17) |
| RBM19 | 9904 | Homo sapiens | RNA binding motif protein 19(RBM19) |

| | | | |
|---|---|---|---|
| GPCPD1 | 56261 | Homo sapiens | glycerophosphocholine phosphodiesterase 1(GPCPD1) |
| MASTL | 84930 | Homo sapiens | microtubule associated serine/threonine kinase like(MASTL) |
| UTP18 | 51096 | Homo sapiens | UTP18 small subunit processome component(UTP18) |
| YWHAZ | 7534 | Homo sapiens | tyrosine 3-monooxygenase/tryptophan 5-monooxygenase activ |
| PJA2 | 9867 | Homo sapiens | praja ring finger ubiquitin ligase 2(PJA2) |
| TBC1D1 | 23216 | Homo sapiens | TBC1 domain family member 1(TBC1D1) |
| VCAN | 1462 | Homo sapiens | versican(VCAN) |
| TBC1D2 | 55357 | Homo sapiens | TBC1 domain family member 2(TBC1D2) |
| RCN1 | 5954 | Homo sapiens | reticulocalbin 1(RCN1) |
| NDUFS7 | 374291 | Homo sapiens | NADH:ubiquinone oxidoreductase core subunit S7(NDUFS7) |
| TBC1D5 | 9779 | Homo sapiens | TBC1 domain family member 5(TBC1D5) |
| IRF4 | 3662 | Homo sapiens | interferon regulatory factor 4(IRF4) |
| GORASP1 | 64689 | Homo sapiens | golgi reassembly stacking protein 1(GORASP1) |
| NDUFS4 | 4724 | Homo sapiens | NADH:ubiquinone oxidoreductase subunit S4(NDUFS4) |
| ATG4C | 84938 | Homo sapiens | autophagy related 4C cysteine peptidase(ATG4C) |
| BLVRB | 645 | Homo sapiens | biliverdin reductase B(BLVRB) |
| ATG4B | 23192 | Homo sapiens | autophagy related 4B cysteine peptidase(ATG4B) |
| MVD | 4597 | Homo sapiens | mevalonate diphosphate decarboxylase(MVD) |
| CFAP54 | 144535 | Homo sapiens | cilia and flagella associated protein 54(CFAP54) |
| EAPP | 55837 | Homo sapiens | E2F associated phosphoprotein(EAPP) |
| CFAP53 | 220136 | Homo sapiens | cilia and flagella associated protein 53(CFAP53) |
| DCTN6 | 10671 | Homo sapiens | dynactin subunit 6(DCTN6) |
| DDX24 | 57062 | Homo sapiens | DEAD-box helicase 24(DDX24) |
| DCTN2 | 10540 | Homo sapiens | dynactin subunit 2(DCTN2) |
| DCTN1 | 1639 | Homo sapiens | dynactin subunit 1(DCTN1) |
| LAMA4 | 3910 | Homo sapiens | laminin subunit alpha 4(LAMA4) |
| LAMA3 | 3909 | Homo sapiens | laminin subunit alpha 3(LAMA3) |
| TGFBRAP1 | 9392 | Homo sapiens | transforming growth factor beta receptor associated protein 1(T |
| DBX1 | 120237 | Homo sapiens | developing brain homeobox 1(DBX1) |
| CRMP1 | 1400 | Homo sapiens | collapsin response mediator protein 1(CRMP1) |
| ATP2C2 | 9914 | Homo sapiens | ATPase secretory pathway Ca2+ transporting 2(ATP2C2) |
| ATP2C1 | 27032 | Homo sapiens | ATPase secretory pathway Ca2+ transporting 1(ATP2C1) |
| SAMHD1 | 25939 | Homo sapiens | SAM and HD domain containing deoxynucleoside triphosphate tr |
| CLCN3 | 1182 | Homo sapiens | chloride voltage-gated channel 3(CLCN3) |
| PARL | 55486 | Homo sapiens | presenilin associated rhomboid like(PARL) |
| CLN5 | 1203 | Homo sapiens | CLN5 intracellular trafficking protein(CLN5) |
| PDLIM1 | 9124 | Homo sapiens | PDZ and LIM domain 1(PDLIM1) |
| TAF1B | 9014 | Homo sapiens | TATA-box binding protein associated factor, RNA polymerase I s |
| SH3TC1 | 54436 | Homo sapiens | SH3 domain and tetratricopeptide repeats 1(SH3TC1) |
| DCDC2B | 149069 | Homo sapiens | doublecortin domain containing 2B(DCDC2B) |
| TAF1A | 9015 | Homo sapiens | TATA-box binding protein associated factor, RNA polymerase I s |
| PDLIM4 | 8572 | Homo sapiens | PDZ and LIM domain 4(PDLIM4) |
| PDLIM7 | 9260 | Homo sapiens | PDZ and LIM domain 7(PDLIM7) |
| CCDC178 | 374864 | Homo sapiens | coiled-coil domain containing 178(CCDC178) |
| GALNT7 | 51809 | Homo sapiens | polypeptide N-acetylgalactosaminyltransferase 7(GALNT7) |

| | | | |
|---|---|---|---|
| GALNT6 | 11226 | Homo sapiens | polypeptide N-acetylgalactosaminyltransferase 6(GALNT6) |
| MBTPS1 | 8720 | Homo sapiens | membrane bound transcription factor peptidase, site 1(MBTPS1) |
| CHURC1 | 91612 | Homo sapiens | churchill domain containing 1(CHURC1) |
| EGF | 1950 | Homo sapiens | epidermal growth factor(EGF) |
| GALNT2 | 2590 | Homo sapiens | polypeptide N-acetylgalactosaminyltransferase 2(GALNT2) |
| GALNT1 | 2589 | Homo sapiens | polypeptide N-acetylgalactosaminyltransferase 1(GALNT1) |
| ATP2B2 | 491 | Homo sapiens | ATPase plasma membrane Ca2+ transporting 2(ATP2B2) |
| LRWD1 | 222229 | Homo sapiens | leucine rich repeats and WD repeat domain containing 1(LRWD1 |
| ACVR2B | 93 | Homo sapiens | activin A receptor type 2B(ACVR2B) |
| CHFR | 55743 | Homo sapiens | checkpoint with forkhead and ring finger domains(CHFR) |
| U2SURP | 23350 | Homo sapiens | U2 snRNP associated SURP domain containing(U2SURP) |
| MAPK10 | 5602 | Homo sapiens | mitogen-activated protein kinase 10(MAPK10) |
| EEPD1 | 80820 | Homo sapiens | endonuclease/exonuclease/phosphatase family domain containi |
| MAPK11 | 5600 | Homo sapiens | mitogen-activated protein kinase 11(MAPK11) |
| BRMS1L | 84312 | Homo sapiens | BRMS1 like transcriptional repressor(BRMS1L) |
| ASB7 | 140460 | Homo sapiens | ankyrin repeat and SOCS box containing 7(ASB7) |
| CHPT1 | 56994 | Homo sapiens | choline phosphotransferase 1(CHPT1) |
| FAM98B | 283742 | Homo sapiens | family with sequence similarity 98 member B(FAM98B) |
| RBM45 | 129831 | Homo sapiens | RNA binding motif protein 45(RBM45) |
| CLUAP1 | 23059 | Homo sapiens | clusterin associated protein 1(CLUAP1) |
| RBM46 | 166863 | Homo sapiens | RNA binding motif protein 46(RBM46) |
| CCDC171 | 203238 | Homo sapiens | coiled-coil domain containing 171(CCDC171) |
| VIT | 5212 | Homo sapiens | vitrin(VIT) |
| CNTFR | 1271 | Homo sapiens | ciliary neurotrophic factor receptor(CNTFR) |
| TRIO | 7204 | Homo sapiens | trio Rho guanine nucleotide exchange factor(TRIO) |
| MAML2 | 84441 | Homo sapiens | mastermind like transcriptional coactivator 2(MAML2) |
| MYT1L | 23040 | Homo sapiens | myelin transcription factor 1 like(MYT1L) |
| TRRAP | 8295 | Homo sapiens | transformation/transcription domain associated protein(TRRAP) |
| COL12A1 | 1303 | Homo sapiens | collagen type XII alpha 1 chain(COL12A1) |
| DCAF7 | 10238 | Homo sapiens | DDB1 and CUL4 associated factor 7(DCAF7) |
| BACH2 | 60468 | Homo sapiens | BTB domain and CNC homolog 2(BACH2) |
| ENO4 | 387712 | Homo sapiens | enolase 4(ENO4) |
| RNF115 | 27246 | Homo sapiens | ring finger protein 115(RNF115) |
| GNAT2 | 2780 | Homo sapiens | G protein subunit alpha transducin 2(GNAT2) |
| SOX17 | 64321 | Homo sapiens | SRY-box transcription factor 17(SOX17) |
| RPL38 | 6169 | Homo sapiens | ribosomal protein L38(RPL38) |
| RTTN | 25914 | Homo sapiens | rotatin(RTTN) |
| ANKS1A | 23294 | Homo sapiens | ankyrin repeat and sterile alpha motif domain containing 1A(AN |
| IER3 | 8870 | Homo sapiens | immediate early response 3(IER3) |
| CUBN | 8029 | Homo sapiens | cubilin(CUBN) |
| NKX2-8 | 26257 | Homo sapiens | NK2 homeobox 8(NKX2-8) |
| TRIQK | 286144 | Homo sapiens | triple QxxK/R motif containing(TRIQK) |
| NKX2-6 | 137814 | Homo sapiens | NK2 homeobox 6(NKX2-6) |
| NHLRC3 | 387921 | Homo sapiens | NHL repeat containing 3(NHLRC3) |
| DAPK1 | 1612 | Homo sapiens | death associated protein kinase 1(DAPK1) |

| | | | |
|---|---|---|---|
| DAPK2 | 23604 | Homo sapiens | death associated protein kinase 2(DAPK2) |
| MED9 | 55090 | Homo sapiens | mediator complex subunit 9(MED9) |
| PASK | 23178 | Homo sapiens | PAS domain containing serine/threonine kinase(PASK) |
| DKK3 | 27122 | Homo sapiens | dickkopf WNT signaling pathway inhibitor 3(DKK3) |
| KIAA1217 | 56243 | Homo sapiens | KIAA1217(KIAA1217) |
| DYNC1LI1 | 51143 | Homo sapiens | dynein cytoplasmic 1 light intermediate chain 1(DYNC1LI1) |
| RNF126 | 55658 | Homo sapiens | ring finger protein 126(RNF126) |
| RNF125 | 54941 | Homo sapiens | ring finger protein 125(RNF125) |
| BIN1 | 274 | Homo sapiens | bridging integrator 1(BIN1) |
| YME1L1 | 10730 | Homo sapiens | YME1 like 1 ATPase(YME1L1) |
| SERPING1 | 710 | Homo sapiens | serpin family G member 1(SERPING1) |
| MAML3 | 55534 | Homo sapiens | mastermind like transcriptional coactivator 3(MAML3) |
| HPRT1 | 3251 | Homo sapiens | hypoxanthine phosphoribosyltransferase 1(HPRT1) |
| RNF122 | 79845 | Homo sapiens | ring finger protein 122(RNF122) |
| RNF121 | 55298 | Homo sapiens | ring finger protein 121(RNF121) |
| RPN2 | 6185 | Homo sapiens | ribophorin II(RPN2) |
| MAPKAP1 | 79109 | Homo sapiens | MAPK associated protein 1(MAPKAP1) |
| HSPA4L | 22824 | Homo sapiens | heat shock protein family A (Hsp70) member 4 like(HSPA4L) |
| ENDOD1 | 23052 | Homo sapiens | endonuclease domain containing 1(ENDOD1) |
| NEDD4L | 23327 | Homo sapiens | NEDD4 like E3 ubiquitin protein ligase(NEDD4L) |
| NOLC1 | 9221 | Homo sapiens | nucleolar and coiled-body phosphoprotein 1(NOLC1) |
| SLAIN2 | 57606 | Homo sapiens | SLAIN motif family member 2(SLAIN2) |
| SLAIN1 | 122060 | Homo sapiens | SLAIN motif family member 1(SLAIN1) |
| SKA3 | 221150 | Homo sapiens | spindle and kinetochore associated complex subunit 3(SKA3) |
| GLG1 | 2734 | Homo sapiens | golgi glycoprotein 1(GLG1) |
| PRDX3 | 10935 | Homo sapiens | peroxiredoxin 3(PRDX3) |
| PPP3R1 | 5534 | Homo sapiens | protein phosphatase 3 regulatory subunit B, alpha(PPP3R1) |
| OCSTAMP | 128506 | Homo sapiens | osteoclast stimulatory transmembrane protein(OCSTAMP) |
| SLC19A3 | 80704 | Homo sapiens | solute carrier family 19 member 3(SLC19A3) |
| CDK19 | 23097 | Homo sapiens | cyclin dependent kinase 19(CDK19) |
| FZD2 | 2535 | Homo sapiens | frizzled class receptor 2(FZD2) |
| ACE | 1636 | Homo sapiens | angiotensin I converting enzyme(ACE) |
| PRRC2C | 23215 | Homo sapiens | proline rich coiled-coil 2C(PRRC2C) |
| FZD8 | 8325 | Homo sapiens | frizzled class receptor 8(FZD8) |
| SLC16A10 | 117247 | Homo sapiens | solute carrier family 16 member 10(SLC16A10) |
| PRDX6 | 9588 | Homo sapiens | peroxiredoxin 6(PRDX6) |
| GPN1 | 11321 | Homo sapiens | GPN-loop GTPase 1(GPN1) |
| ST18 | 9705 | Homo sapiens | ST18 C2H2C-type zinc finger transcription factor(ST18) |
| PRRC2B | 84726 | Homo sapiens | proline rich coiled-coil 2B(PRRC2B) |
| C9ORF85 | 138241 | Homo sapiens | chromosome 9 open reading frame 85(C9orf85) |
| GPR137C | 283554 | Homo sapiens | G protein-coupled receptor 137C(GPR137C) |
| SLC16A13 | 201232 | Homo sapiens | solute carrier family 16 member 13(SLC16A13) |
| FA2H | 79152 | Homo sapiens | fatty acid 2-hydroxylase(FA2H) |
| SLC6A6 | 6533 | Homo sapiens | solute carrier family 6 member 6(SLC6A6) |
| PPP5C | 5536 | Homo sapiens | protein phosphatase 5 catalytic subunit(PPP5C) |

| | | | |
|---|---|---|---|
| GIPC3 | 126326 | Homo sapiens | GIPC PDZ domain containing family member 3(GIPC3) |
| CPEB2 | 132864 | Homo sapiens | cytoplasmic polyadenylation element binding protein 2(CPEB2) |
| RET | 5979 | Homo sapiens | ret proto-oncogene(RET) |
| ADK | 132 | Homo sapiens | adenosine kinase(ADK) |
| TCF21 | 6943 | Homo sapiens | transcription factor 21(TCF21) |
| CLTA | 1211 | Homo sapiens | clathrin light chain A(CLTA) |
| ING1 | 3621 | Homo sapiens | inhibitor of growth family member 1(ING1) |
| SLC6A2 | 6530 | Homo sapiens | solute carrier family 6 member 2(SLC6A2) |
| TRIM8 | 81603 | Homo sapiens | tripartite motif containing 8(TRIM8) |
| ING4 | 51147 | Homo sapiens | inhibitor of growth family member 4(ING4) |
| RNF157 | 114804 | Homo sapiens | ring finger protein 157(RNF157) |
| TRIM9 | 114088 | Homo sapiens | tripartite motif containing 9(TRIM9) |
| CHL1 | 10752 | Homo sapiens | cell adhesion molecule L1 like(CHL1) |
| NUF2 | 83540 | Homo sapiens | NUF2 component of NDC80 kinetochore complex(NUF2) |
| ENSA | 2029 | Homo sapiens | endosulfine alpha(ENSA) |
| RSPO3 | 84870 | Homo sapiens | R-spondin 3(RSPO3) |
| XXYLT1 | 152002 | Homo sapiens | xyloside xylosyltransferase 1(XXYLT1) |
| TIGD1 | 200765 | Homo sapiens | tigger transposable element derived 1(TIGD1) |
| MIOX | 55586 | Homo sapiens | myo-inositol oxygenase(MIOX) |
| BRD1 | 23774 | Homo sapiens | bromodomain containing 1(BRD1) |
| ANXA2 | 302 | Homo sapiens | annexin A2(ANXA2) |
| FCRL3 | 115352 | Homo sapiens | Fc receptor like 3(FCRL3) |
| NFAM1 | 150372 | Homo sapiens | NFAT activating protein with ITAM motif 1(NFAM1) |
| IFNGR2 | 3460 | Homo sapiens | interferon gamma receptor 2(IFNGR2) |
| TCF12 | 6938 | Homo sapiens | transcription factor 12(TCF12) |
| TET1 | 80312 | Homo sapiens | tet methylcytosine dioxygenase 1(TET1) |
| PRSS16 | 10279 | Homo sapiens | serine protease 16(PRSS16) |
| TACR1 | 6869 | Homo sapiens | tachykinin receptor 1(TACR1) |
| PAX2 | 5076 | Homo sapiens | paired box 2(PAX2) |
| PAX1 | 5075 | Homo sapiens | paired box 1(PAX1) |
| SERPINB6 | 5269 | Homo sapiens | serpin family B member 6(SERPINB6) |
| RNF167 | 26001 | Homo sapiens | ring finger protein 167(RNF167) |
| INF2 | 64423 | Homo sapiens | inverted formin 2(INF2) |
| PLCB4 | 5332 | Homo sapiens | phospholipase C beta 4(PLCB4) |
| PAX8 | 7849 | Homo sapiens | paired box 8(PAX8) |
| SLCO4A1 | 28231 | Homo sapiens | solute carrier organic anion transporter family member 4A1(SLC |
| TET3 | 200424 | Homo sapiens | tet methylcytosine dioxygenase 3(TET3) |
| PLCB1 | 23236 | Homo sapiens | phospholipase C beta 1(PLCB1) |
| RNF166 | 115992 | Homo sapiens | ring finger protein 166(RNF166) |
| NPSR1 | 387129 | Homo sapiens | neuropeptide S receptor 1(NPSR1) |
| OCA2 | 4948 | Homo sapiens | OCA2 melanosomal transmembrane protein(OCA2) |
| SF3B3 | 23450 | Homo sapiens | splicing factor 3b subunit 3(SF3B3) |
| PHLPP1 | 23239 | Homo sapiens | PH domain and leucine rich repeat protein phosphatase 1(PHLPI |
| CTDP1 | 9150 | Homo sapiens | CTD phosphatase subunit 1(CTDP1) |
| SERPINC1 | 462 | Homo sapiens | serpin family C member 1(SERPINC1) |

| | | | |
|---|---|---|---|
| GCNT1 | 2650 | Homo sapiens | glucosaminyl (N-acetyl) transferase 1(GCNT1) |
| TM9SF4 | 9777 | Homo sapiens | transmembrane 9 superfamily member 4(TM9SF4) |
| MIPOL1 | 145282 | Homo sapiens | mirror-image polydactyly 1(MIPOL1) |
| ATP6AP1L | 92270 | Homo sapiens | ATPase H+ transporting accessory protein 1 like (pseudogene)( |
| CHRND | 1144 | Homo sapiens | cholinergic receptor nicotinic delta subunit(CHRND) |
| LMAN1 | 3998 | Homo sapiens | lectin, mannose binding 1(LMAN1) |
| CHRNG | 1146 | Homo sapiens | cholinergic receptor nicotinic gamma subunit(CHRNG) |
| SRP72 | 6731 | Homo sapiens | signal recognition particle 72(SRP72) |
| BRD8 | 10902 | Homo sapiens | bromodomain containing 8(BRD8) |
| C16ORF96 | 342346 | Homo sapiens | chromosome 16 open reading frame 96(C16orf96) |
| MISP | 126353 | Homo sapiens | mitotic spindle positioning(MISP) |
| DTD1 | 92675 | Homo sapiens | D-aminoacyl-tRNA deacylase 1(DTD1) |
| BRD4 | 23476 | Homo sapiens | bromodomain containing 4(BRD4) |
| WWOX | 51741 | Homo sapiens | WW domain containing oxidoreductase(WWOX) |
| AOC2 | 314 | Homo sapiens | amine oxidase copper containing 2(AOC2) |
| TIMMDC1 | 51300 | Homo sapiens | translocase of inner mitochondrial membrane domain containing |
| AGK | 55750 | Homo sapiens | acylglycerol kinase(AGK) |
| NDE1 | 54820 | Homo sapiens | nudE neurodevelopment protein 1(NDE1) |
| LAPTM5 | 7805 | Homo sapiens | lysosomal protein transmembrane 5(LAPTM5) |
| ADAT1 | 23536 | Homo sapiens | adenosine deaminase tRNA specific 1(ADAT1) |
| NMNAT3 | 349565 | Homo sapiens | nicotinamide nucleotide adenylyltransferase 3(NMNAT3) |
| TRMT1 | 55621 | Homo sapiens | tRNA methyltransferase 1(TRMT1) |
| FMO5 | 2330 | Homo sapiens | flavin containing dimethylaniline monoxygenase 5(FMO5) |
| ADAT2 | 134637 | Homo sapiens | adenosine deaminase tRNA specific 2(ADAT2) |
| KTN1 | 3895 | Homo sapiens | kinectin 1(KTN1) |
| TM9SF2 | 9375 | Homo sapiens | transmembrane 9 superfamily member 2(TM9SF2) |
| RNF145 | 153830 | Homo sapiens | ring finger protein 145(RNF145) |
| ZNFX1 | 57169 | Homo sapiens | zinc finger NFX1-type containing 1(ZNFX1) |
| AGBL1 | 123624 | Homo sapiens | AGBL carboxypeptidase 1(AGBL1) |
| VEPH1 | 79674 | Homo sapiens | ventricular zone expressed PH domain containing 1(VEPH1) |
| ABI1 | 10006 | Homo sapiens | abl interactor 1(ABI1) |
| LPCAT2 | 54947 | Homo sapiens | lysophosphatidylcholine acyltransferase 2(LPCAT2) |
| C16ORF86 | 388284 | Homo sapiens | chromosome 16 open reading frame 86(C16orf86) |
| PLCD3 | 113026 | Homo sapiens | phospholipase C delta 3(PLCD3) |
| PLCD1 | 5333 | Homo sapiens | phospholipase C delta 1(PLCD1) |
| SEMA5A | 9037 | Homo sapiens | semaphorin 5A(SEMA5A) |
| DGKD | 8527 | Homo sapiens | diacylglycerol kinase delta(DGKD) |
| COL16A1 | 1307 | Homo sapiens | collagen type XVI alpha 1 chain(COL16A1) |
| CLSTN2 | 64084 | Homo sapiens | calsyntenin 2(CLSTN2) |
| ATP8A1 | 10396 | Homo sapiens | ATPase phospholipid transporting 8A1(ATP8A1) |
| DGKB | 1607 | Homo sapiens | diacylglycerol kinase beta(DGKB) |
| FMR1 | 2332 | Homo sapiens | fragile X messenger ribonucleoprotein 1(FMR1) |
| CTNND2 | 1501 | Homo sapiens | catenin delta 2(CTNND2) |
| CLSTN1 | 22883 | Homo sapiens | calsyntenin 1(CLSTN1) |
| GUCA1A | 2978 | Homo sapiens | guanylate cyclase activator 1A(GUCA1A) |

| | | | |
|---|---|---|---|
| IRS2 | 8660 | Homo sapiens | insulin receptor substrate 2(IRS2) |
| CDH6 | 1004 | Homo sapiens | cadherin 6(CDH6) |
| PYCARD | 29108 | Homo sapiens | PYD and CARD domain containing(PYCARD) |
| CDH2 | 1000 | Homo sapiens | cadherin 2(CDH2) |
| CDH1 | 999 | Homo sapiens | cadherin 1(CDH1) |
| ANKRD31 | 256006 | Homo sapiens | ankyrin repeat domain 31(ANKRD31) |
| NBN | 4683 | Homo sapiens | nibrin(NBN) |
| GYG1 | 2992 | Homo sapiens | glycogenin 1(GYG1) |
| TECTA | 7007 | Homo sapiens | tectorin alpha(TECTA) |
| LZTS3 | 9762 | Homo sapiens | leucine zipper tumor suppressor family member 3(LZTS3) |
| SEMA6B | 10501 | Homo sapiens | semaphorin 6B(SEMA6B) |
| SLC15A1 | 6564 | Homo sapiens | solute carrier family 15 member 1(SLC15A1) |
| IGFBP5 | 3488 | Homo sapiens | insulin like growth factor binding protein 5(IGFBP5) |
| RPS8 | 6202 | Homo sapiens | ribosomal protein S8(RPS8) |
| GPT2 | 84706 | Homo sapiens | glutamic--pyruvic transaminase 2(GPT2) |
| IGFBP2 | 3485 | Homo sapiens | insulin like growth factor binding protein 2(IGFBP2) |
| MITF | 4286 | Homo sapiens | melanocyte inducing transcription factor(MITF) |
| SYPL2 | 284612 | Homo sapiens | synaptophysin like 2(SYPL2) |
| GTF2F2 | 2963 | Homo sapiens | general transcription factor IIF subunit 2(GTF2F2) |
| MOB3C | 148932 | Homo sapiens | MOB kinase activator 3C(MOB3C) |
| CACNB2 | 783 | Homo sapiens | calcium voltage-gated channel auxiliary subunit beta 2(CACNB2 |
| RALGAPB | 57148 | Homo sapiens | Ral GTPase activating protein non-catalytic subunit beta(RALGAI |
| SYPL1 | 6856 | Homo sapiens | synaptophysin like 1(SYPL1) |
| STIM1 | 6786 | Homo sapiens | stromal interaction molecule 1(STIM1) |
| TEX9 | 374618 | Homo sapiens | testis expressed 9(TEX9) |
| STIM2 | 57620 | Homo sapiens | stromal interaction molecule 2(STIM2) |
| DGKQ | 1609 | Homo sapiens | diacylglycerol kinase theta(DGKQ) |
| CLIP4 | 79745 | Homo sapiens | CAP-Gly domain containing linker protein family member 4(CLII |
| ERP27 | 121506 | Homo sapiens | endoplasmic reticulum protein 27(ERP27) |
| TNIK | 23043 | Homo sapiens | TRAF2 and NCK interacting kinase(TNIK) |
| ERG | 2078 | Homo sapiens | ETS transcription factor ERG(ERG) |
| PHLDB2 | 90102 | Homo sapiens | pleckstrin homology like domain family B member 2(PHLDB2) |
| ANKRD17 | 26057 | Homo sapiens | ankyrin repeat domain 17(ANKRD17) |
| CUL9 | 23113 | Homo sapiens | cullin 9(CUL9) |
| NOTCH1 | 4851 | Homo sapiens | notch receptor 1(NOTCH1) |
| ANKRD16 | 54522 | Homo sapiens | ankyrin repeat domain 16(ANKRD16) |
| CUL5 | 8065 | Homo sapiens | cullin 5(CUL5) |
| ANKRD11 | 29123 | Homo sapiens | ankyrin repeat domain containing 11(ANKRD11) |
| NATD1 | 256302 | Homo sapiens | N-acetyltransferase domain containing 1(NATD1) |
| CUL1 | 8454 | Homo sapiens | cullin 1(CUL1) |
| ABHD8 | 79575 | Homo sapiens | abhydrolase domain containing 8(ABHD8) |
| ITPR2 | 3709 | Homo sapiens | inositol 1,4,5-trisphosphate receptor type 2(ITPR2) |
| RASAL1 | 8437 | Homo sapiens | RAS protein activator like 1(RASAL1) |
| ABHD6 | 57406 | Homo sapiens | abhydrolase domain containing 6, acylglycerol lipase(ABHD6) |
| RASAL2 | 9462 | Homo sapiens | RAS protein activator like 2(RASAL2) |

| | | | |
|---|---|---|---|
| ABHD5 | 51099 | Homo sapiens | abhydrolase domain containing 5, lysophosphatidic acid acyltrar |
| RASAL3 | 64926 | Homo sapiens | RAS protein activator like 3(RASAL3) |
| AK7 | 122481 | Homo sapiens | adenylate kinase 7(AK7) |
| C1ORF53 | 388722 | Homo sapiens | chromosome 1 open reading frame 53(C1orf53) |
| AK8 | 158067 | Homo sapiens | adenylate kinase 8(AK8) |
| RHOBTB1 | 9886 | Homo sapiens | Rho related BTB domain containing 1(RHOBTB1) |
| PHF21B | 112885 | Homo sapiens | PHD finger protein 21B(PHF21B) |
| ITPRIPL2 | 162073 | Homo sapiens | ITPRIP like 2(ITPRIPL2) |
| HMBOX1 | 79618 | Homo sapiens | homeobox containing 1(HMBOX1) |
| PITRM1 | 10531 | Homo sapiens | pitrilysin metallopeptidase 1(PITRM1) |
| BAG3 | 9531 | Homo sapiens | BAG cochaperone 3(BAG3) |
| PSAP | 5660 | Homo sapiens | prosaposin(PSAP) |
| ESD | 2098 | Homo sapiens | esterase D(ESD) |
| ANKRD10 | 55608 | Homo sapiens | ankyrin repeat domain 10(ANKRD10) |
| SLC15A4 | 121260 | Homo sapiens | solute carrier family 15 member 4(SLC15A4) |
| SKIL | 6498 | Homo sapiens | SKI like proto-oncogene(SKIL) |
| IMPG1 | 3617 | Homo sapiens | interphotoreceptor matrix proteoglycan 1(IMPG1) |
| ANKRD28 | 23243 | Homo sapiens | ankyrin repeat domain 28(ANKRD28) |
| SLC37A2 | 219855 | Homo sapiens | solute carrier family 37 member 2(SLC37A2) |
| ATP8B1 | 5205 | Homo sapiens | ATPase phospholipid transporting 8B1(ATP8B1) |
| WFS1 | 7466 | Homo sapiens | wolframin ER transmembrane glycoprotein(WFS1) |
| ESRRB | 2103 | Homo sapiens | estrogen related receptor beta(ESRRB) |
| NEB | 4703 | Homo sapiens | nebulin(NEB) |
| ADAP1 | 11033 | Homo sapiens | ArfGAP with dual PH domains 1(ADAP1) |
| CNOT7 | 29883 | Homo sapiens | CCR4-NOT transcription complex subunit 7(CNOT7) |
| CNOT2 | 4848 | Homo sapiens | CCR4-NOT transcription complex subunit 2(CNOT2) |
| RCAN2 | 10231 | Homo sapiens | regulator of calcineurin 2(RCAN2) |
| ITGA11 | 22801 | Homo sapiens | integrin subunit alpha 11(ITGA11) |
| TDRD12 | 91646 | Homo sapiens | tudor domain containing 12(TDRD12) |
| ESAM | 90952 | Homo sapiens | endothelial cell adhesion molecule(ESAM) |
| NF2 | 4771 | Homo sapiens | NF2, moesin-ezrin-radixin like (MERLIN) tumor suppressor(NF2 |
| XAF1 | 54739 | Homo sapiens | XIAP associated factor 1(XAF1) |
| SERINC2 | 347735 | Homo sapiens | serine incorporator 2(SERINC2) |
| ALK | 238 | Homo sapiens | ALK receptor tyrosine kinase(ALK) |
| EIF4E3 | 317649 | Homo sapiens | eukaryotic translation initiation factor 4E family member 3(EIF4 |
| ANKRD33B | 651746 | Homo sapiens | ankyrin repeat domain 33B(ANKRD33B) |
| SRXN1 | 140809 | Homo sapiens | sulfiredoxin 1(SRXN1) |
| LSP1 | 4046 | Homo sapiens | lymphocyte specific protein 1(LSP1) |
| PSD3 | 23362 | Homo sapiens | pleckstrin and Sec7 domain containing 3(PSD3) |
| HYDIN | 54768 | Homo sapiens | HYDIN axonemal central pair apparatus protein(HYDIN) |
| DNMT3B | 1789 | Homo sapiens | DNA methyltransferase 3 beta(DNMT3B) |
| CNGA4 | 1262 | Homo sapiens | cyclic nucleotide gated channel subunit alpha 4(CNGA4) |
| METTL15 | 196074 | Homo sapiens | methyltransferase 15, mitochondrial 12S rRNA N4-cytidine(MET |
| IQCB1 | 9657 | Homo sapiens | IQ motif containing B1(IQCB1) |
| REXO1 | 57455 | Homo sapiens | RNA exonuclease 1 homolog(REXO1) |

| | | | |
|---|---|---|---|
| VWF | 7450 | Homo sapiens | von Willebrand factor(VWF) |
| CHUK | 1147 | Homo sapiens | component of inhibitor of nuclear factor kappa B kinase comple: |
| DTX3L | 151636 | Homo sapiens | deltex E3 ubiquitin ligase 3L(DTX3L) |
| DNMT3A | 1788 | Homo sapiens | DNA methyltransferase 3 alpha(DNMT3A) |
| VSTM2L | 128434 | Homo sapiens | V-set and transmembrane domain containing 2 like(VSTM2L) |
| KCNAB2 | 8514 | Homo sapiens | potassium voltage-gated channel subfamily A regulatory beta su |
| TRAF1 | 7185 | Homo sapiens | TNF receptor associated factor 1(TRAF1) |
| PTBP3 | 9991 | Homo sapiens | polypyrimidine tract binding protein 3(PTBP3) |
| DUSP22 | 56940 | Homo sapiens | dual specificity phosphatase 22(DUSP22) |
| RP1 | 6101 | Homo sapiens | RP1 axonemal microtubule associated(RP1) |
| METTL24 | 728464 | Homo sapiens | methyltransferase like 24(METTL24) |
| RP2 | 6102 | Homo sapiens | RP2 activator of ARL3 GTPase(RP2) |
| ESRP1 | 54845 | Homo sapiens | epithelial splicing regulatory protein 1(ESRP1) |
| ZNF516 | 9658 | Homo sapiens | zinc finger protein 516(ZNF516) |
| ANG | 283 | Homo sapiens | angiogenin(ANG) |
| SNRPA1 | 6627 | Homo sapiens | small nuclear ribonucleoprotein polypeptide A'(SNRPA1) |
| EFHC2 | 80258 | Homo sapiens | EF-hand domain containing 2(EFHC2) |
| ZNF511 | 118472 | Homo sapiens | zinc finger protein 511(ZNF511) |
| SOS2 | 6655 | Homo sapiens | SOS Ras/Rho guanine nucleotide exchange factor 2(SOS2) |
| VSTM2B | 342865 | Homo sapiens | V-set and transmembrane domain containing 2B(VSTM2B) |
| SLC24A2 | 25769 | Homo sapiens | solute carrier family 24 member 2(SLC24A2) |
| GTF2A1 | 2957 | Homo sapiens | general transcription factor IIA subunit 1(GTF2A1) |
| SLC24A3 | 57419 | Homo sapiens | solute carrier family 24 member 3(SLC24A3) |
| ACSS2 | 55902 | Homo sapiens | acyl-CoA synthetase short chain family member 2(ACSS2) |
| LUZP1 | 7798 | Homo sapiens | leucine zipper protein 1(LUZP1) |
| SEMA3B | 7869 | Homo sapiens | semaphorin 3B(SEMA3B) |
| SEMA3G | 56920 | Homo sapiens | semaphorin 3G(SEMA3G) |
| AMBRA1 | 55626 | Homo sapiens | autophagy and beclin 1 regulator 1(AMBRA1) |
| JAKMIP1 | 152789 | Homo sapiens | janus kinase and microtubule interacting protein 1(JAKMIP1) |
| MROH1 | 727957 | Homo sapiens | maestro heat like repeat family member 1(MROH1) |
| ENOX2 | 10495 | Homo sapiens | ecto-NOX disulfide-thiol exchanger 2(ENOX2) |
| FZR1 | 51343 | Homo sapiens | fizzy and cell division cycle 20 related 1(FZR1) |
| DCAF17 | 80067 | Homo sapiens | DDB1 and CUL4 associated factor 17(DCAF17) |
| CACNG8 | 59283 | Homo sapiens | calcium voltage-gated channel auxiliary subunit gamma 8(CACI |
| MGAT5B | 146664 | Homo sapiens | alpha-1,6-mannosylglycoprotein 6-beta-N-acetylglucosaminyltra |
| EEF2KMT | 196483 | Homo sapiens | eukaryotic elongation factor 2 lysine methyltransferase(EEF2KM |
| RBBP6 | 5930 | Homo sapiens | RB binding protein 6, ubiquitin ligase(RBBP6) |
| MROH7 | 374977 | Homo sapiens | maestro heat like repeat family member 7(MROH7) |
| CACNG1 | 786 | Homo sapiens | calcium voltage-gated channel auxiliary subunit gamma 1(CACI |
| FNIP1 | 96459 | Homo sapiens | folliculin interacting protein 1(FNIP1) |
| ACSS1 | 84532 | Homo sapiens | acyl-CoA synthetase short chain family member 1(ACSS1) |
| MEST | 4232 | Homo sapiens | mesoderm specific transcript(MEST) |
| CTNNAL1 | 8727 | Homo sapiens | catenin alpha like 1(CTNNAL1) |
| DCAF13 | 25879 | Homo sapiens | DDB1 and CUL4 associated factor 13(DCAF13) |
| CACNG4 | 27092 | Homo sapiens | calcium voltage-gated channel auxiliary subunit gamma 4(CACI |

| | | | |
|---|---|---|---|
| SPAG9 | 9043 | Homo sapiens | sperm associated antigen 9(SPAG9) |
| IRX2 | 153572 | Homo sapiens | iroquois homeobox 2(IRX2) |
| SKOR2 | 652991 | Homo sapiens | SKI family transcriptional corepressor 2(SKOR2) |
| IRX6 | 79190 | Homo sapiens | iroquois homeobox 6(IRX6) |
| TXNRD1 | 7296 | Homo sapiens | thioredoxin reductase 1(TXNRD1) |
| PLCL1 | 5334 | Homo sapiens | phospholipase C like 1 (inactive)(PLCL1) |
| PLCL2 | 23228 | Homo sapiens | phospholipase C like 2(PLCL2) |
| IRX4 | 50805 | Homo sapiens | iroquois homeobox 4(IRX4) |
| PTPN13 | 5783 | Homo sapiens | protein tyrosine phosphatase non-receptor type 13(PTPN13) |
| OXR1 | 55074 | Homo sapiens | oxidation resistance 1(OXR1) |
| ENOX1 | 55068 | Homo sapiens | ecto-NOX disulfide-thiol exchanger 1(ENOX1) |
| RPGRIP1 | 57096 | Homo sapiens | RPGR interacting protein 1(RPGRIP1) |
| RAD50 | 10111 | Homo sapiens | RAD50 double strand break repair protein(RAD50) |
| NFASC | 23114 | Homo sapiens | neurofascin(NFASC) |
| PFKL | 5211 | Homo sapiens | phosphofructokinase, liver type(PFKL) |
| IMPDH1 | 3614 | Homo sapiens | inosine monophosphate dehydrogenase 1(IMPDH1) |
| CROCC | 9696 | Homo sapiens | ciliary rootlet coiled-coil, rootletin(CROCC) |
| MGAT4A | 11320 | Homo sapiens | alpha-1,3-mannosyl-glycoprotein 4-beta-N-acetylglucosaminylt |
| MMACHC | 25974 | Homo sapiens | metabolism of cobalamin associated C(MMACHC) |
| TRMT61A | 115708 | Homo sapiens | tRNA methyltransferase 61A(TRMT61A) |
| PFKP | 5214 | Homo sapiens | phosphofructokinase, platelet(PFKP) |
| APP | 351 | Homo sapiens | amyloid beta precursor protein(APP) |
| VIPR1 | 7433 | Homo sapiens | vasoactive intestinal peptide receptor 1(VIPR1) |
| HPSE2 | 60495 | Homo sapiens | heparanase 2 (inactive)(HPSE2) |
| ARL6IP1 | 23204 | Homo sapiens | ADP ribosylation factor like GTPase 6 interacting protein 1(ARL6 |
| TTLL10 | 254173 | Homo sapiens | tubulin tyrosine ligase like 10(TTLL10) |
| HSF2BP | 11077 | Homo sapiens | heat shock transcription factor 2 binding protein(HSF2BP) |
| PKD2L2 | 27039 | Homo sapiens | polycystin 2 like 2, transient receptor potential cation channel(P |
| PWP2 | 5822 | Homo sapiens | PWP2 small subunit processome component(PWP2) |
| ANTXR2 | 118429 | Homo sapiens | ANTXR cell adhesion molecule 2(ANTXR2) |
| ANTXR1 | 84168 | Homo sapiens | ANTXR cell adhesion molecule 1(ANTXR1) |
| VSIG1 | 340547 | Homo sapiens | V-set and immunoglobulin domain containing 1(VSIG1) |
| CHEK2 | 11200 | Homo sapiens | checkpoint kinase 2(CHEK2) |
| INSC | 387755 | Homo sapiens | INSC spindle orientation adaptor protein(INSC) |
| SAP130 | 79595 | Homo sapiens | Sin3A associated protein 130(SAP130) |
| DPF3 | 8110 | Homo sapiens | double PHD fingers 3(DPF3) |
| TRMT44 | 152992 | Homo sapiens | tRNA methyltransferase 44 homolog(TRMT44) |
| TNS4 | 84951 | Homo sapiens | tensin 4(TNS4) |
| KIFAP3 | 22920 | Homo sapiens | kinesin associated protein 3(KIFAP3) |
| B2M | 567 | Homo sapiens | beta-2-microglobulin(B2M) |
| TNS2 | 23371 | Homo sapiens | tensin 2(TNS2) |
| AQR | 9716 | Homo sapiens | aquarius intron-binding spliceosomal factor(AQR) |
| TIPIN | 54962 | Homo sapiens | TIMELESS interacting protein(TIPIN) |
| SLC6A14 | 11254 | Homo sapiens | solute carrier family 6 member 14(SLC6A14) |
| SLC6A11 | 6538 | Homo sapiens | solute carrier family 6 member 11(SLC6A11) |

| | | | |
|---|---|---|---|
| MRPS18C | 51023 | Homo sapiens | mitochondrial ribosomal protein S18C(MRPS18C) |
| CYP27A1 | 1593 | Homo sapiens | cytochrome P450 family 27 subfamily A member 1(CYP27A1) |
| INPP4A | 3631 | Homo sapiens | inositol polyphosphate-4-phosphatase type I A(INPP4A) |
| AFG3L2 | 10939 | Homo sapiens | AFG3 like matrix AAA peptidase subunit 2(AFG3L2) |
| GRWD1 | 83743 | Homo sapiens | glutamate rich WD repeat containing 1(GRWD1) |
| ZFPM1 | 161882 | Homo sapiens | zinc finger protein, FOG family member 1(ZFPM1) |
| LLGL1 | 3996 | Homo sapiens | LLGL scribble cell polarity complex component 1(LLGL1) |
| CD101 | 9398 | Homo sapiens | CD101 molecule(CD101) |
| CDS1 | 1040 | Homo sapiens | CDP-diacylglycerol synthase 1(CDS1) |
| NECAB2 | 54550 | Homo sapiens | N-terminal EF-hand calcium binding protein 2(NECAB2) |
| PTGFR | 5737 | Homo sapiens | prostaglandin F receptor(PTGFR) |
| PDE1C | 5137 | Homo sapiens | phosphodiesterase 1C(PDE1C) |
| TEX264 | 51368 | Homo sapiens | testis expressed 264, ER-phagy receptor(TEX264) |
| HEPACAM | 220296 | Homo sapiens | hepatic and glial cell adhesion molecule(HEPACAM) |
| NLK | 51701 | Homo sapiens | nemo like kinase(NLK) |
| TMTC4 | 84899 | Homo sapiens | transmembrane O-mannosyltransferase targeting cadherins 4(T |
| PTDSS2 | 81490 | Homo sapiens | phosphatidylserine synthase 2(PTDSS2) |
| INPP5A | 3632 | Homo sapiens | inositol polyphosphate-5-phosphatase A(INPP5A) |
| INPP5F | 22876 | Homo sapiens | inositol polyphosphate-5-phosphatase F(INPP5F) |
| INPP5E | 56623 | Homo sapiens | inositol polyphosphate-5-phosphatase E(INPP5E) |
| VPS8 | 23355 | Homo sapiens | VPS8 subunit of CORVET complex(VPS8) |
| MAP7 | 9053 | Homo sapiens | microtubule associated protein 7(MAP7) |
| OTOG | 340990 | Homo sapiens | otogelin(OTOG) |
| MAP6 | 4135 | Homo sapiens | microtubule associated protein 6(MAP6) |
| OTOF | 9381 | Homo sapiens | otoferlin(OTOF) |
| NAALAD2 | 10003 | Homo sapiens | N-acetylated alpha-linked acidic dipeptidase 2(NAALAD2) |
| DPH6 | 89978 | Homo sapiens | diphthamine biosynthesis 6(DPH6) |
| ASL | 435 | Homo sapiens | argininosuccinate lyase(ASL) |
| MAP4 | 4134 | Homo sapiens | microtubule associated protein 4(MAP4) |
| SCAI | 286205 | Homo sapiens | suppressor of cancer cell invasion(SCAI) |
| BANP | 54971 | Homo sapiens | BTG3 associated nuclear protein(BANP) |
| NUS1 | 116150 | Homo sapiens | NUS1 dehydrodolichyl diphosphate synthase subunit(NUS1) |
| FIS1 | 51024 | Homo sapiens | fission, mitochondrial 1(FIS1) |
| SIGLEC15 | 284266 | Homo sapiens | sialic acid binding Ig like lectin 15(SIGLEC15) |
| PDE2A | 5138 | Homo sapiens | phosphodiesterase 2A(PDE2A) |
| RAB39A | 54734 | Homo sapiens | RAB39A, member RAS oncogene family(RAB39A) |
| ANKEF1 | 63926 | Homo sapiens | ankyrin repeat and EF-hand domain containing 1(ANKEF1) |
| HSPBAP1 | 79663 | Homo sapiens | HSPB1 associated protein 1(HSPBAP1) |
| ACADSB | 36 | Homo sapiens | acyl-CoA dehydrogenase short/branched chain(ACADSB) |
| CIDEC | 63924 | Homo sapiens | cell death inducing DFFA like effector c(CIDEC) |
| UBAC1 | 10422 | Homo sapiens | UBA domain containing 1(UBAC1) |
| STAG1 | 10274 | Homo sapiens | STAG1 cohesin complex component(STAG1) |
| CCDC88C | 440193 | Homo sapiens | coiled-coil domain containing 88C(CCDC88C) |
| PDE3A | 5139 | Homo sapiens | phosphodiesterase 3A(PDE3A) |
| TMCC3 | 57458 | Homo sapiens | transmembrane and coiled-coil domain family 3(TMCC3) |

| | | | |
|---|---|---|---|
| TRMT11 | 60487 | Homo sapiens | tRNA methyltransferase 11 homolog(TRMT11) |
| CC2D1A | 54862 | Homo sapiens | coiled-coil and C2 domain containing 1A(CC2D1A) |
| ITM2C | 81618 | Homo sapiens | integral membrane protein 2C(ITM2C) |
| CC2D1B | 200014 | Homo sapiens | coiled-coil and C2 domain containing 1B(CC2D1B) |
| RYR1 | 6261 | Homo sapiens | ryanodine receptor 1(RYR1) |
| TRAM1 | 23471 | Homo sapiens | translocation associated membrane protein 1(TRAM1) |
| FAF1 | 11124 | Homo sapiens | Fas associated factor 1(FAF1) |
| GDI2 | 2665 | Homo sapiens | GDP dissociation inhibitor 2(GDI2) |
| FAF2 | 23197 | Homo sapiens | Fas associated factor family member 2(FAF2) |
| STAB2 | 55576 | Homo sapiens | stabilin 2(STAB2) |
| RYR3 | 6263 | Homo sapiens | ryanodine receptor 3(RYR3) |
| GYS2 | 2998 | Homo sapiens | glycogen synthase 2(GYS2) |
| GYS1 | 2997 | Homo sapiens | glycogen synthase 1(GYS1) |
| GUK1 | 2987 | Homo sapiens | guanylate kinase 1(GUK1) |
| STAB1 | 23166 | Homo sapiens | stabilin 1(STAB1) |
| KIF13A | 63971 | Homo sapiens | kinesin family member 13A(KIF13A) |
| KIF13B | 23303 | Homo sapiens | kinesin family member 13B(KIF13B) |
| CC2D2A | 57545 | Homo sapiens | coiled-coil and C2 domain containing 2A(CC2D2A) |
| RSPRY1 | 89970 | Homo sapiens | ring finger and SPRY domain containing 1(RSPRY1) |
| ANKRD6 | 22881 | Homo sapiens | ankyrin repeat domain 6(ANKRD6) |
| MAP4K2 | 5871 | Homo sapiens | mitogen-activated protein kinase kinase kinase kinase 2(MAP4K |
| PRMT8 | 56341 | Homo sapiens | protein arginine methyltransferase 8(PRMT8) |
| PRMT7 | 54496 | Homo sapiens | protein arginine methyltransferase 7(PRMT7) |
| TSC22D2 | 9819 | Homo sapiens | TSC22 domain family member 2(TSC22D2) |
| RHOG | 391 | Homo sapiens | ras homolog family member G(RHOG) |
| AMPD2 | 271 | Homo sapiens | adenosine monophosphate deaminase 2(AMPD2) |
| IGFN1 | 91156 | Homo sapiens | immunoglobulin like and fibronectin type III domain containing |
| NAALADL2 | 254827 | Homo sapiens | N-acetylated alpha-linked acidic dipeptidase like 2(NAALADL2) |
| PALD1 | 27143 | Homo sapiens | phosphatase domain containing paladin 1(PALD1) |
| IL17RA | 23765 | Homo sapiens | interleukin 17 receptor A(IL17RA) |
| RAD51B | 5890 | Homo sapiens | RAD51 paralog B(RAD51B) |
| ADRB3 | 155 | Homo sapiens | adrenoceptor beta 3(ADRB3) |
| MPPED1 | 758 | Homo sapiens | metallophosphoesterase domain containing 1(MPPED1) |
| MDFIC | 29969 | Homo sapiens | MyoD family inhibitor domain containing(MDFIC) |
| PACS1 | 55690 | Homo sapiens | phosphofurin acidic cluster sorting protein 1(PACS1) |
| NME8 | 51314 | Homo sapiens | NME/NM23 family member 8(NME8) |
| TNNT2 | 7139 | Homo sapiens | troponin T2, cardiac type(TNNT2) |
| TFEC | 22797 | Homo sapiens | transcription factor EC(TFEC) |
| RHOJ | 57381 | Homo sapiens | ras homolog family member J(RHOJ) |
| WAC | 51322 | Homo sapiens | WW domain containing adaptor with coiled-coil(WAC) |
| FAM20C | 56975 | Homo sapiens | FAM20C golgi associated secretory pathway kinase(FAM20C) |
| TNXB | 7148 | Homo sapiens | tenascin XB(TNXB) |
| ALOX12B | 242 | Homo sapiens | arachidonate 12-lipoxygenase, 12R type(ALOX12B) |
| AKAP6 | 9472 | Homo sapiens | A-kinase anchoring protein 6(AKAP6) |
| TRAK2 | 66008 | Homo sapiens | trafficking kinesin protein 2(TRAK2) |

| | | | |
|---|---|---|---|
| FAM184A | 79632 | Homo sapiens | family with sequence similarity 184 member A(FAM184A) |
| NCR3LG1 | 374383 | Homo sapiens | natural killer cell cytotoxicity receptor 3 ligand 1(NCR3LG1) |
| MAP4K3 | 8491 | Homo sapiens | mitogen-activated protein kinase kinase kinase kinase 3(MAP4K |
| AAMDC | 28971 | Homo sapiens | adipogenesis associated Mth938 domain containing(AAMDC) |
| MAP4K4 | 9448 | Homo sapiens | mitogen-activated protein kinase kinase kinase kinase 4(MAP4K |
| RANBP1 | 5902 | Homo sapiens | RAN binding protein 1(RANBP1) |
| ARNT2 | 9915 | Homo sapiens | aryl hydrocarbon receptor nuclear translocator 2(ARNT2) |
| CA10 | 56934 | Homo sapiens | carbonic anhydrase 10(CA10) |
| RANBP3 | 8498 | Homo sapiens | RAN binding protein 3(RANBP3) |
| RYK | 6259 | Homo sapiens | receptor like tyrosine kinase(RYK) |
| RANBP9 | 10048 | Homo sapiens | RAN binding protein 9(RANBP9) |
| PDZK1IP1 | 10158 | Homo sapiens | PDZK1 interacting protein 1(PDZK1IP1) |
| PUS7L | 83448 | Homo sapiens | pseudouridine synthase 7 like(PUS7L) |
| FAH | 2184 | Homo sapiens | fumarylacetoacetate hydrolase(FAH) |
| ZDHHC11B | 653082 | Homo sapiens | zinc finger DHHC-type containing 11B(ZDHHC11B) |
| TJP1 | 7082 | Homo sapiens | tight junction protein 1(TJP1) |
| CAMK4 | 814 | Homo sapiens | calcium/calmodulin dependent protein kinase IV(CAMK4) |
| STAC2 | 342667 | Homo sapiens | SH3 and cysteine rich domain 2(STAC2) |
| TRIP11 | 9321 | Homo sapiens | thyroid hormone receptor interactor 11(TRIP11) |
| RAB3GAP1 | 22930 | Homo sapiens | RAB3 GTPase activating protein catalytic subunit 1(RAB3GAP1) |
| GALNT12 | 79695 | Homo sapiens | polypeptide N-acetylgalactosaminyltransferase 12(GALNT12) |
| NPY2R | 4887 | Homo sapiens | neuropeptide Y receptor Y2(NPY2R) |
| CMIP | 80790 | Homo sapiens | c-Maf inducing protein(CMIP) |
| ARHGAP44 | 9912 | Homo sapiens | Rho GTPase activating protein 44(ARHGAP44) |
| ARHGAP42 | 143872 | Homo sapiens | Rho GTPase activating protein 42(ARHGAP42) |
| IL18RAP | 8807 | Homo sapiens | interleukin 18 receptor accessory protein(IL18RAP) |
| CHCHD3 | 54927 | Homo sapiens | coiled-coil-helix-coiled-coil-helix domain containing 3(CHCHD3) |
| SIRPA | 140885 | Homo sapiens | signal regulatory protein alpha(SIRPA) |
| NSF | 4905 | Homo sapiens | N-ethylmaleimide sensitive factor, vesicle fusing ATPase(NSF) |
| DSCAM | 1826 | Homo sapiens | DS cell adhesion molecule(DSCAM) |
| APLP2 | 334 | Homo sapiens | amyloid beta precursor like protein 2(APLP2) |
| MATK | 4145 | Homo sapiens | megakaryocyte-associated tyrosine kinase(MATK) |
| OCRL | 4952 | Homo sapiens | OCRL inositol polyphosphate-5-phosphatase(OCRL) |
| CDYL | 9425 | Homo sapiens | chromodomain Y like(CDYL) |
| PRSS55 | 203074 | Homo sapiens | serine protease 55(PRSS55) |
| DPM2 | 8818 | Homo sapiens | dolichyl-phosphate mannosyltransferase subunit 2, regulatory(D |
| COL4A2 | 1284 | Homo sapiens | collagen type IV alpha 2 chain(COL4A2) |
| COL4A1 | 1282 | Homo sapiens | collagen type IV alpha 1 chain(COL4A1) |
| TBL1XR1 | 79718 | Homo sapiens | TBL1X/Y related 1(TBL1XR1) |
| EOGT | 285203 | Homo sapiens | EGF domain specific O-linked N-acetylglucosamine transferase( |
| COL4A6 | 1288 | Homo sapiens | collagen type IV alpha 6 chain(COL4A6) |
| KLRF1 | 51348 | Homo sapiens | killer cell lectin like receptor F1(KLRF1) |
| RELL1 | 768211 | Homo sapiens | RELT like 1(RELL1) |
| PDE9A | 5152 | Homo sapiens | phosphodiesterase 9A(PDE9A) |
| ABCG1 | 9619 | Homo sapiens | ATP binding cassette subfamily G member 1(ABCG1) |

| | | | |
|---|---|---|---|
| AKNA | 80709 | Homo sapiens | AT-hook transcription factor(AKNA) |
| TMEM59L | 25789 | Homo sapiens | transmembrane protein 59 like(TMEM59L) |
| ARHGAP19 | 84986 | Homo sapiens | Rho GTPase activating protein 19(ARHGAP19) |
| ARHGAP17 | 55114 | Homo sapiens | Rho GTPase activating protein 17(ARHGAP17) |
| ADIPOR1 | 51094 | Homo sapiens | adiponectin receptor 1(ADIPOR1) |
| ARHGAP15 | 55843 | Homo sapiens | Rho GTPase activating protein 15(ARHGAP15) |
| STK4 | 6789 | Homo sapiens | serine/threonine kinase 4(STK4) |
| ADIPOR2 | 79602 | Homo sapiens | adiponectin receptor 2(ADIPOR2) |
| DNAJC28 | 54943 | Homo sapiens | DnaJ heat shock protein family (Hsp40) member C28(DNAJC28 |
| ARHGAP12 | 94134 | Homo sapiens | Rho GTPase activating protein 12(ARHGAP12) |
| ARHGAP21 | 57584 | Homo sapiens | Rho GTPase activating protein 21(ARHGAP21) |
| SCIN | 85477 | Homo sapiens | scinderin(SCIN) |
| PDZD2 | 23037 | Homo sapiens | PDZ domain containing 2(PDZD2) |
| TMEM128 | 85013 | Homo sapiens | transmembrane protein 128(TMEM128) |
| CASKIN1 | 57524 | Homo sapiens | CASK interacting protein 1(CASKIN1) |
| ATP6V1H | 51606 | Homo sapiens | ATPase H+ transporting V1 subunit H(ATP6V1H) |
| MYH11 | 4629 | Homo sapiens | myosin heavy chain 11(MYH11) |
| DMAP1 | 55929 | Homo sapiens | DNA methyltransferase 1 associated protein 1(DMAP1) |
| MARK4 | 57787 | Homo sapiens | microtubule affinity regulating kinase 4(MARK4) |
| ZC3H14 | 79882 | Homo sapiens | zinc finger CCCH-type containing 14(ZC3H14) |
| MYH10 | 4628 | Homo sapiens | myosin heavy chain 10(MYH10) |
| SMAD2 | 4087 | Homo sapiens | SMAD family member 2(SMAD2) |
| RAP1GAP2 | 23108 | Homo sapiens | RAP1 GTPase activating protein 2(RAP1GAP2) |
| ACBD5 | 91452 | Homo sapiens | acyl-CoA binding domain containing 5(ACBD5) |
| STIL | 6491 | Homo sapiens | STIL centriolar assembly protein(STIL) |
| LRRC75B | 388886 | Homo sapiens | leucine rich repeat containing 75B(LRRC75B) |
| WSCD1 | 23302 | Homo sapiens | WSC domain containing 1(WSCD1) |
| SMURF2 | 64750 | Homo sapiens | SMAD specific E3 ubiquitin protein ligase 2(SMURF2) |
| WSCD2 | 9671 | Homo sapiens | WSC domain containing 2(WSCD2) |
| PTCH1 | 5727 | Homo sapiens | patched 1(PTCH1) |
| ASPSCR1 | 79058 | Homo sapiens | ASPSCR1 tether for SLC2A4, UBX domain containing(ASPSCR1) |
| TDRD9 | 122402 | Homo sapiens | tudor domain containing 9(TDRD9) |
| IFT122 | 55764 | Homo sapiens | intraflagellar transport 122(IFT122) |
| AZIN2 | 113451 | Homo sapiens | antizyme inhibitor 2(AZIN2) |
| DNAJC13 | 23317 | Homo sapiens | DnaJ heat shock protein family (Hsp40) member C13(DNAJC13 |
| BTNL2 | 56244 | Homo sapiens | butyrophilin like 2(BTNL2) |
| TDRD5 | 163589 | Homo sapiens | tudor domain containing 5(TDRD5) |
| ACSF3 | 197322 | Homo sapiens | acyl-CoA synthetase family member 3(ACSF3) |
| TDRD3 | 81550 | Homo sapiens | tudor domain containing 3(TDRD3) |
| ACSF2 | 80221 | Homo sapiens | acyl-CoA synthetase family member 2(ACSF2) |
| ESR2 | 2100 | Homo sapiens | estrogen receptor 2(ESR2) |
| TDRD1 | 56165 | Homo sapiens | tudor domain containing 1(TDRD1) |
| ARHGAP33 | 115703 | Homo sapiens | Rho GTPase activating protein 33(ARHGAP33) |
| TBCK | 93627 | Homo sapiens | TBC1 domain containing kinase(TBCK) |
| ARHGAP30 | 257106 | Homo sapiens | Rho GTPase activating protein 30(ARHGAP30) |

| | | | |
|---|---|---|---|
| ATG16L1 | 55054 | Homo sapiens | autophagy related 16 like 1(ATG16L1) |
| DLG5 | 9231 | Homo sapiens | discs large MAGUK scaffold protein 5(DLG5) |
| RADIL | 55698 | Homo sapiens | Rap associating with DIL domain(RADIL) |
| CLASRP | 11129 | Homo sapiens | CLK4 associating serine/arginine rich protein(CLASRP) |
| CRISP2 | 7180 | Homo sapiens | cysteine rich secretory protein 2(CRISP2) |
| SCYL3 | 57147 | Homo sapiens | SCY1 like pseudokinase 3(SCYL3) |
| DOCK4 | 9732 | Homo sapiens | dedicator of cytokinesis 4(DOCK4) |
| DOCK3 | 1795 | Homo sapiens | dedicator of cytokinesis 3(DOCK3) |
| ABCD3 | 5825 | Homo sapiens | ATP binding cassette subfamily D member 3(ABCD3) |
| SPO11 | 23626 | Homo sapiens | SPO11 initiator of meiotic double strand breaks(SPO11) |
| USP33 | 23032 | Homo sapiens | ubiquitin specific peptidase 33(USP33) |
| USP34 | 9736 | Homo sapiens | ubiquitin specific peptidase 34(USP34) |
| DOCK7 | 85440 | Homo sapiens | dedicator of cytokinesis 7(DOCK7) |
| PDE3B | 5140 | Homo sapiens | phosphodiesterase 3B(PDE3B) |
| SLC2A2 | 6514 | Homo sapiens | solute carrier family 2 member 2(SLC2A2) |
| CHTF18 | 63922 | Homo sapiens | chromosome transmission fidelity factor 18(CHTF18) |
| SLC2A6 | 11182 | Homo sapiens | solute carrier family 2 member 6(SLC2A6) |
| DTWD1 | 56986 | Homo sapiens | DTW domain containing 1(DTWD1) |
| ZC3H7B | 23264 | Homo sapiens | zinc finger CCCH-type containing 7B(ZC3H7B) |
| ENOSF1 | 55556 | Homo sapiens | enolase superfamily member 1(ENOSF1) |
| CYP4V2 | 285440 | Homo sapiens | cytochrome P450 family 4 subfamily V member 2(CYP4V2) |
| PDE4B | 5142 | Homo sapiens | phosphodiesterase 4B(PDE4B) |
| HIBCH | 26275 | Homo sapiens | 3-hydroxyisobutyryl-CoA hydrolase(HIBCH) |
| USP46 | 64854 | Homo sapiens | ubiquitin specific peptidase 46(USP46) |
| ABCC3 | 8714 | Homo sapiens | ATP binding cassette subfamily C member 3(ABCC3) |
| ABCC1 | 4363 | Homo sapiens | ATP binding cassette subfamily C member 1 (ABCC1 blood grou |
| KLF13 | 51621 | Homo sapiens | KLF transcription factor 13(KLF13) |
| USP42 | 84132 | Homo sapiens | ubiquitin specific peptidase 42(USP42) |
| USP43 | 124739 | Homo sapiens | ubiquitin specific peptidase 43(USP43) |
| ABCC8 | 6833 | Homo sapiens | ATP binding cassette subfamily C member 8(ABCC8) |
| LMNTD1 | 160492 | Homo sapiens | lamin tail domain containing 1(LMNTD1) |
| ARAP2 | 116984 | Homo sapiens | ArfGAP with RhoGAP domain, ankyrin repeat and PH domain 2( |
| ARAP3 | 64411 | Homo sapiens | ArfGAP with RhoGAP domain, ankyrin repeat and PH domain 3( |
| ABCC9 | 10060 | Homo sapiens | ATP binding cassette subfamily C member 9(ABCC9) |
| ARAP1 | 116985 | Homo sapiens | ArfGAP with RhoGAP domain, ankyrin repeat and PH domain 1( |
| OR6P1 | 128366 | Homo sapiens | olfactory receptor family 6 subfamily P member 1(OR6P1) |
| ARID1B | 57492 | Homo sapiens | AT-rich interaction domain 1B(ARID1B) |
| TGFBR2 | 7048 | Homo sapiens | transforming growth factor beta receptor 2(TGFBR2) |
| LATS2 | 26524 | Homo sapiens | large tumor suppressor kinase 2(LATS2) |
| PALLD | 23022 | Homo sapiens | palladin, cytoskeletal associated protein(PALLD) |
| SAG | 6295 | Homo sapiens | S-antigen visual arrestin(SAG) |
| LUC7L3 | 51747 | Homo sapiens | LUC7 like 3 pre-mRNA splicing factor(LUC7L3) |
| MCTP2 | 55784 | Homo sapiens | multiple C2 and transmembrane domain containing 2(MCTP2) |
| HCN4 | 10021 | Homo sapiens | hyperpolarization activated cyclic nucleotide gated potassium ch |
| RGS19 | 10287 | Homo sapiens | regulator of G protein signaling 19(RGS19) |

| | | | |
|---|---|---|---|
| USP10 | 9100 | Homo sapiens | ubiquitin specific peptidase 10(USP10) |
| GMPR2 | 51292 | Homo sapiens | guanosine monophosphate reductase 2(GMPR2) |
| TM2D3 | 80213 | Homo sapiens | TM2 domain containing 3(TM2D3) |
| UACA | 55075 | Homo sapiens | uveal autoantigen with coiled-coil domains and ankyrin repeats |
| IGFBPL1 | 347252 | Homo sapiens | insulin like growth factor binding protein like 1(IGFBPL1) |
| GMDS | 2762 | Homo sapiens | GDP-mannose 4,6-dehydratase(GMDS) |
| ERBB4 | 2066 | Homo sapiens | erb-b2 receptor tyrosine kinase 4(ERBB4) |
| NT5DC3 | 51559 | Homo sapiens | 5'-nucleotidase domain containing 3(NT5DC3) |
| PDE6D | 5147 | Homo sapiens | phosphodiesterase 6D(PDE6D) |
| LRIG1 | 26018 | Homo sapiens | leucine rich repeats and immunoglobulin like domains 1(LRIG1) |
| PDE6B | 5158 | Homo sapiens | phosphodiesterase 6B(PDE6B) |
| NCAM1 | 4684 | Homo sapiens | neural cell adhesion molecule 1(NCAM1) |
| MUTYH | 4595 | Homo sapiens | mutY DNA glycosylase(MUTYH) |
| CLASP2 | 23122 | Homo sapiens | cytoplasmic linker associated protein 2(CLASP2) |
| RWDD1 | 51389 | Homo sapiens | RWD domain containing 1(RWDD1) |
| CMTM7 | 112616 | Homo sapiens | CKLF like MARVEL transmembrane domain containing 7(CMTM7 |
| SPEN | 23013 | Homo sapiens | spen family transcriptional repressor(SPEN) |
| USP25 | 29761 | Homo sapiens | ubiquitin specific peptidase 25(USP25) |
| POLQ | 10721 | Homo sapiens | DNA polymerase theta(POLQ) |
| ATE1 | 11101 | Homo sapiens | arginyltransferase 1(ATE1) |
| USP20 | 10868 | Homo sapiens | ubiquitin specific peptidase 20(USP20) |
| PCDHGC5 | 56097 | Homo sapiens | protocadherin gamma subfamily C, 5(PCDHGC5) |
| USP22 | 23326 | Homo sapiens | ubiquitin specific peptidase 22(USP22) |
| HAGHL | 84264 | Homo sapiens | hydroxyacylglutathione hydrolase like(HAGHL) |
| CMTM6 | 54918 | Homo sapiens | CKLF like MARVEL transmembrane domain containing 6(CMTM6 |
| DTNBP1 | 84062 | Homo sapiens | dystrobrevin binding protein 1(DTNBP1) |
| PDE4DIP | 9659 | Homo sapiens | phosphodiesterase 4D interacting protein(PDE4DIP) |
| ARID3A | 1820 | Homo sapiens | AT-rich interaction domain 3A(ARID3A) |
| TSPEAR | 54084 | Homo sapiens | thrombospondin type laminin G domain and EAR repeats(TSPEA |
| SHCBP1 | 79801 | Homo sapiens | SHC binding and spindle associated 1(SHCBP1) |
| GDAP2 | 54834 | Homo sapiens | ganglioside induced differentiation associated protein 2(GDAP2) |
| NAP1L4 | 4676 | Homo sapiens | nucleosome assembly protein 1 like 4(NAP1L4) |
| SH3YL1 | 26751 | Homo sapiens | SH3 and SYLF domain containing 1(SH3YL1) |
| FMNL2 | 114793 | Homo sapiens | formin like 2(FMNL2) |
| XPNPEP1 | 7511 | Homo sapiens | X-prolyl aminopeptidase 1(XPNPEP1) |
| NEURL1 | 9148 | Homo sapiens | neuralized E3 ubiquitin protein ligase 1(NEURL1) |
| PDE7B | 27115 | Homo sapiens | phosphodiesterase 7B(PDE7B) |
| HCN2 | 610 | Homo sapiens | hyperpolarization activated cyclic nucleotide gated potassium ar |

### **C** List of genes whose promoter or coding region contains the h

SEZ6
LRFN1
INTS5
F9
C3

SHCBP1L
RNASEL
SDC3
MED9
DUSP12
IRX6
PDYN
FOSL2
SPO11
TBRG1
TTC23
TMEM63C
OLIG3
SHQ1
LAMB4
GPR61
NUDT14
MTHFS
NUP210L
DDX46
TMEM17
HIP1
PKD2
ZNF524
NKX1-2
FNDC4
BAX
ARID3A
GABBR2
MAB21L1
DOHH
HECW2
CCDC102A
STBD1
PRSS55
PTPN2
TMEM230
TCF21
ASPH
ADIPOR1
PLEKHG2
ALOX5
GDF3
WDR36
IL7

MKX
GIMAP7
PLEKHN1
SNAPC5
ISPD
CLEC2D
NSUN2
DSG2
CNKSR3
CA2
CHRNA5
TOP1MT
GATA2
TDRD12
GZMH
SYNGR1
NHSL2
VSIG4
TSSK3
WNT5B
PNCK
C20orf96
MTHFSD
BCAN
RGCC
HNF1B
CHCHD3
NAA38
NCS1
FAM229A
GDNF
NAT8L
VSTM2B
TMEM184B
ZBTB20
CNTD1
PILRA
TAF9
BSCL2
ASCL3
TCTA
CDCA7L
RNF186
ATP5A1
KIAA0907

PPP1R3D
TBX22
SPINK6
ABCA5
WDR88
HLA-DPA1
TRAV20
G2E3
RNFT2
CWF19L1
OTUD7B
VEGFB
ZBED2
RIMS4
CYBB
MAB21L2
SND1
PIWIL1
SLC11A2
MTMR7
USP36
WDR4
AVL9
SH3BP5
COA3
VMAC
HOXA7
SRD5A1
GIPC3
TRIM36
METRN
C20orf196
INPP5D
CYP26A1
COX4I2
MRPL39
PPFIA3
EIF3H
TBC1D16
CACFD1
KLHL11
LOC100996758
EAPP
SFMBT2
CD74

CLEC11A
FKBP1B
COX16
KCNK13
C1QA
GOT1
TEKT3
TIMM44
MRPS30
GNE
GPR21
RPL18
RHBDD2
AP2A2
RUNX1
RPL3L
PTGES
THUMPD3
ADAM20

### D DAVID gene names of affected genes from tab "C"

| From | To | Species | David Gene Name |
|---|---|---|---|
| THUMPD3 | 25917 | Homo sapiens | THUMP domain containing 3(THUMPD3) |
| ZBTB20 | 26137 | Homo sapiens | zinc finger and BTB domain containing 20(ZBTB20) |
| GPR61 | 83873 | Homo sapiens | G protein-coupled receptor 61(GPR61) |
| CCDC102A | 92922 | Homo sapiens | coiled-coil domain containing 102A(CCDC102A) |
| GIMAP7 | 168537 | Homo sapiens | GTPase, IMAP family member 7(GIMAP7) |
| WDR88 | 126248 | Homo sapiens | WD repeat domain 88(WDR88) |
| RIMS4 | 140730 | Homo sapiens | regulating synaptic membrane exocytosis 4(RIMS4) |
| CHCHD3 | 54927 | Homo sapiens | coiled-coil-helix-coiled-coil-helix domain containing 3(CHCHD3) |
| NKX1-2 | 390010 | Homo sapiens | NK1 homeobox 2(NKX1-2) |
| TEKT3 | 64518 | Homo sapiens | tektin 3(TEKT3) |
| VSIG4 | 11326 | Homo sapiens | V-set and immunoglobulin domain containing 4(VSIG4) |
| HOXA7 | 3204 | Homo sapiens | homeobox A7(HOXA7) |
| NUDT14 | 256281 | Homo sapiens | nudix hydrolase 14(NUDT14) |
| PPFIA3 | 8541 | Homo sapiens | PTPRF interacting protein alpha 3(PPFIA3) |
| WNT5B | 81029 | Homo sapiens | Wnt family member 5B(WNT5B) |
| SPINK6 | 404203 | Homo sapiens | serine peptidase inhibitor Kazal type 6(SPINK6) |
| SFMBT2 | 57713 | Homo sapiens | Scm like with four mbt domains 2(SFMBT2) |
| KCNK13 | 56659 | Homo sapiens | potassium two pore domain channel subfamily K member 13(K |
| CLEC11A | 6320 | Homo sapiens | C-type lectin domain containing 11A(CLEC11A) |
| SLC11A2 | 4891 | Homo sapiens | solute carrier family 11 member 2(SLC11A2) |
| CYBB | 1536 | Homo sapiens | cytochrome b-245 beta chain(CYBB) |
| MED9 | 55090 | Homo sapiens | mediator complex subunit 9(MED9) |

| | | | |
|---|---|---|---|
| HNF1B | 6928 | Homo sapiens | HNF1 homeobox B(HNF1B) |
| PDYN | 5173 | Homo sapiens | prodynorphin(PDYN) |
| PRSS55 | 203074 | Homo sapiens | serine protease 55(PRSS55) |
| SND1 | 27044 | Homo sapiens | staphylococcal nuclease and tudor domain containing 1(SND1) |
| RUNX1 | 861 | Homo sapiens | RUNX family transcription factor 1(RUNX1) |
| BSCL2 | 26580 | Homo sapiens | BSCL2 lipid droplet biogenesis associated, seipin(BSCL2) |
| BCAN | 63827 | Homo sapiens | brevican(BCAN) |
| VMAC | 400673 | Homo sapiens | vimentin type intermediate filament associated coiled-coil prote |
| SHQ1 | 55164 | Homo sapiens | SHQ1, H/ACA ribonucleoprotein assembly factor(SHQ1) |
| FNDC4 | 64838 | Homo sapiens | fibronectin type III domain containing 4(FNDC4) |
| NUP210L | 91181 | Homo sapiens | nucleoporin 210 like(NUP210L) |
| HECW2 | 57520 | Homo sapiens | HECT, C2 and WW domain containing E3 ubiquitin protein ligase |
| PPP1R3D | 5509 | Homo sapiens | protein phosphatase 1 regulatory subunit 3D(PPP1R3D) |
| TSSK3 | 81629 | Homo sapiens | testis specific serine kinase 3(TSSK3) |
| TCTA | 6988 | Homo sapiens | T cell leukemia translocation altered(TCTA) |
| GNE | 10020 | Homo sapiens | glucosamine (UDP-N-acetyl)-2-epimerase/N-acetylmannosamir |
| PTGES | 9536 | Homo sapiens | prostaglandin E synthase(PTGES) |
| COX16 | 51241 | Homo sapiens | cytochrome c oxidase assembly factor COX16(COX16) |
| CHRNA5 | 1138 | Homo sapiens | cholinergic receptor nicotinic alpha 5 subunit(CHRNA5) |
| TMEM63C | 57156 | Homo sapiens | transmembrane protein 63C(TMEM63C) |
| SDC3 | 9672 | Homo sapiens | syndecan 3(SDC3) |
| TRAV20 | 28663 | Homo sapiens | T cell receptor alpha variable 20(TRAV20) |
| KLHL11 | 55175 | Homo sapiens | kelch like family member 11(KLHL11) |
| ADIPOR1 | 51094 | Homo sapiens | adiponectin receptor 1(ADIPOR1) |
| GATA2 | 2624 | Homo sapiens | GATA binding protein 2(GATA2) |
| MTMR7 | 9108 | Homo sapiens | myotubularin related protein 7(MTMR7) |
| C3 | 718 | Homo sapiens | complement C3(C3) |
| SYNGR1 | 9145 | Homo sapiens | synaptogyrin 1(SYNGR1) |
| INPP5D | 3635 | Homo sapiens | inositol polyphosphate-5-phosphatase D(INPP5D) |
| ZBED2 | 79413 | Homo sapiens | zinc finger BED-type containing 2(ZBED2) |
| TBC1D16 | 125058 | Homo sapiens | TBC1 domain family member 16(TBC1D16) |
| COA3 | 28958 | Homo sapiens | cytochrome c oxidase assembly factor 3(COA3) |
| TMEM184B | 25829 | Homo sapiens | transmembrane protein 184B(TMEM184B) |
| GOT1 | 2805 | Homo sapiens | glutamic-oxaloacetic transaminase 1(GOT1) |
| ABCA5 | 23461 | Homo sapiens | ATP binding cassette subfamily A member 5(ABCA5) |
| SRD5A1 | 6715 | Homo sapiens | steroid 5 alpha-reductase 1(SRD5A1) |
| RHBDD2 | 57414 | Homo sapiens | rhomboid domain containing 2(RHBDD2) |
| CACFD1 | 11094 | Homo sapiens | calcium channel flower domain containing 1(CACFD1) |
| FAM229A | 100128071 | Homo sapiens | family with sequence similarity 229 member A(FAM229A) |
| TIMM44 | 10469 | Homo sapiens | translocase of inner mitochondrial membrane 44(TIMM44) |
| MTHFS | 10588 | Homo sapiens | methenyltetrahydrofolate synthetase(MTHFS) |
| PILRA | 29992 | Homo sapiens | paired immunoglobin like type 2 receptor alpha(PILRA) |
| GDF3 | 9573 | Homo sapiens | growth differentiation factor 3(GDF3) |
| RPL3L | 6123 | Homo sapiens | ribosomal protein L3 like(RPL3L) |
| TMEM230 | 29058 | Homo sapiens | transmembrane protein 230(TMEM230) |

| | | | |
|---|---|---|---|
| FOSL2 | 2355 | Homo sapiens | FOS like 2, AP-1 transcription factor subunit(FOSL2) |
| CYP26A1 | 1592 | Homo sapiens | cytochrome P450 family 26 subfamily A member 1(CYP26A1) |
| RGCC | 28984 | Homo sapiens | regulator of cell cycle(RGCC) |
| TMEM17 | 200728 | Homo sapiens | transmembrane protein 17(TMEM17) |
| GDNF | 2668 | Homo sapiens | glial cell derived neurotrophic factor(GDNF) |
| GIPC3 | 126326 | Homo sapiens | GIPC PDZ domain containing family member 3(GIPC3) |
| IL7 | 3574 | Homo sapiens | interleukin 7(IL7) |
| LRFN1 | 57622 | Homo sapiens | leucine rich repeat and fibronectin type III domain containing 1( |
| CWF19L1 | 55280 | Homo sapiens | CWF19 like cell cycle control factor 1(CWF19L1) |
| TDRD12 | 91646 | Homo sapiens | tudor domain containing 12(TDRD12) |
| INTS5 | 80789 | Homo sapiens | integrator complex subunit 5(INTS5) |
| MKX | 283078 | Homo sapiens | mohawk homeobox(MKX) |
| RNF186 | 54546 | Homo sapiens | ring finger protein 186(RNF186) |
| AVL9 | 23080 | Homo sapiens | AVL9 cell migration associated(AVL9) |
| USP36 | 57602 | Homo sapiens | ubiquitin specific peptidase 36(USP36) |
| PIWIL1 | 9271 | Homo sapiens | piwi like RNA-mediated gene silencing 1(PIWIL1) |
| TTC23 | 64927 | Homo sapiens | tetratricopeptide repeat domain 23(TTC23) |
| HIP1 | 3092 | Homo sapiens | huntingtin interacting protein 1(HIP1) |
| DDX46 | 9879 | Homo sapiens | DEAD-box helicase 46(DDX46) |
| SPO11 | 23626 | Homo sapiens | SPO11 initiator of meiotic double strand breaks(SPO11) |
| GPR21 | 2844 | Homo sapiens | G protein-coupled receptor 21(GPR21) |
| CDCA7L | 55536 | Homo sapiens | cell division cycle associated 7 like(CDCA7L) |
| WDR4 | 10785 | Homo sapiens | WD repeat domain 4(WDR4) |
| COX4I2 | 84701 | Homo sapiens | cytochrome c oxidase subunit 4I2(COX4I2) |
| TCF21 | 6943 | Homo sapiens | transcription factor 21(TCF21) |
| MRPL39 | 54148 | Homo sapiens | mitochondrial ribosomal protein L39(MRPL39) |
| OTUD7B | 56957 | Homo sapiens | OTU deubiquitinase 7B(OTUD7B) |
| PKD2 | 5311 | Homo sapiens | polycystin 2, transient receptor potential cation channel(PKD2) |
| AP2A2 | 161 | Homo sapiens | adaptor related protein complex 2 subunit alpha 2(AP2A2) |
| SNAPC5 | 10302 | Homo sapiens | small nuclear RNA activating complex polypeptide 5(SNAPC5) |
| DUSP12 | 11266 | Homo sapiens | dual specificity phosphatase 12(DUSP12) |
| NHSL2 | 340527 | Homo sapiens | NHS like 2(NHSL2) |
| CA2 | 760 | Homo sapiens | carbonic anhydrase 2(CA2) |
| TOP1MT | 116447 | Homo sapiens | DNA topoisomerase I mitochondrial(TOP1MT) |
| STBD1 | 8987 | Homo sapiens | starch binding domain 1(STBD1) |
| NCS1 | 23413 | Homo sapiens | neuronal calcium sensor 1(NCS1) |
| ZNF524 | 147807 | Homo sapiens | zinc finger protein 524(ZNF524) |
| PLEKHN1 | 84069 | Homo sapiens | pleckstrin homology domain containing N1(PLEKHN1) |
| HLA-DPA1 | 3113 | Homo sapiens | major histocompatibility complex, class II, DP alpha 1(HLA-DPA |
| WDR36 | 134430 | Homo sapiens | WD repeat domain 36(WDR36) |
| CNTD1 | 124817 | Homo sapiens | cyclin N-terminal domain containing 1(CNTD1) |
| PLEKHG2 | 64857 | Homo sapiens | pleckstrin homology and RhoGEF domain containing G2(PLEKHG |
| OLIG3 | 167826 | Homo sapiens | oligodendrocyte transcription factor 3(OLIG3) |
| METRN | 79006 | Homo sapiens | meteorin, glial cell differentiation regulator(METRN) |
| NSUN2 | 54888 | Homo sapiens | NOP2/Sun RNA methyltransferase 2(NSUN2) |

| | | | |
|---|---|---|---|
| C20ORF96 | 140680 | Homo sapiens | chromosome 20 open reading frame 96(C20orf96) |
| F9 | 2158 | Homo sapiens | coagulation factor IX(F9) |
| ASPH | 444 | Homo sapiens | aspartate beta-hydroxylase(ASPH) |
| CNKSR3 | 154043 | Homo sapiens | CNKSR family member 3(CNKSR3) |
| G2E3 | 55632 | Homo sapiens | G2/M-phase specific E3 ubiquitin protein ligase(G2E3) |
| DSG2 | 1829 | Homo sapiens | desmoglein 2(DSG2) |
| NAA38 | 84316 | Homo sapiens | N-alpha-acetyltransferase 38, NatC auxiliary subunit(NAA38) |
| EAPP | 55837 | Homo sapiens | E2F associated phosphoprotein(EAPP) |
| NAT8L | 339983 | Homo sapiens | N-acetyltransferase 8 like(NAT8L) |
| VSTM2B | 342865 | Homo sapiens | V-set and transmembrane domain containing 2B(VSTM2B) |
| RNFT2 | 84900 | Homo sapiens | ring finger protein, transmembrane 2(RNFT2) |
| C1QA | 712 | Homo sapiens | complement C1q A chain(C1QA) |
| RNASEL | 6041 | Homo sapiens | ribonuclease L(RNASEL) |
| ASCL3 | 56676 | Homo sapiens | achaete-scute family bHLH transcription factor 3(ASCL3) |
| PNCK | 139728 | Homo sapiens | pregnancy up-regulated nonubiquitous CaM kinase(PNCK) |
| SHCBP1L | 81626 | Homo sapiens | SHC binding and spindle associated 1 like(SHCBP1L) |
| TAF9 | 6880 | Homo sapiens | TATA-box binding protein associated factor 9(TAF9) |
| SEZ6 | 124925 | Homo sapiens | seizure related 6 homolog(SEZ6) |
| ADAM20 | 8748 | Homo sapiens | ADAM metallopeptidase domain 20(ADAM20) |
| MRPS30 | 10884 | Homo sapiens | mitochondrial ribosomal protein S30(MRPS30) |
| MTHFSD | 64779 | Homo sapiens | methenyltetrahydrofolate synthetase domain containing(MTHFS |
| TBRG1 | 84897 | Homo sapiens | transforming growth factor beta regulator 1(TBRG1) |
| TBX22 | 50945 | Homo sapiens | T-box transcription factor 22(TBX22) |
| ALOX5 | 240 | Homo sapiens | arachidonate 5-lipoxygenase(ALOX5) |
| SH3BP5 | 9467 | Homo sapiens | SH3 domain binding protein 5(SH3BP5) |
| RPL18 | 6141 | Homo sapiens | ribosomal protein L18(RPL18) |
| CD74 | 972 | Homo sapiens | CD74 molecule(CD74) |
| GABBR2 | 9568 | Homo sapiens | gamma-aminobutyric acid type B receptor subunit 2(GABBR2) |
| IRX6 | 79190 | Homo sapiens | iroquois homeobox 6(IRX6) |
| LAMB4 | 22798 | Homo sapiens | laminin subunit beta 4(LAMB4) |
| VEGFB | 7423 | Homo sapiens | vascular endothelial growth factor B(VEGFB) |
| ARID3A | 1820 | Homo sapiens | AT-rich interaction domain 3A(ARID3A) |
| DOHH | 83475 | Homo sapiens | deoxyhypusine hydroxylase(DOHH) |
| GZMH | 2999 | Homo sapiens | granzyme H(GZMH) |
| MAB21L1 | 4081 | Homo sapiens | mab-21 like 1(MAB21L1) |
| MAB21L2 | 10586 | Homo sapiens | mab-21 like 2(MAB21L2) |
| FKBP1B | 2281 | Homo sapiens | FKBP prolyl isomerase 1B(FKBP1B) |
| EIF3H | 8667 | Homo sapiens | eukaryotic translation initiation factor 3 subunit H(EIF3H) |
| TRIM36 | 55521 | Homo sapiens | tripartite motif containing 36(TRIM36) |
| BAX | 581 | Homo sapiens | BCL2 associated X, apoptosis regulator(BAX) |
| CLEC2D | 29121 | Homo sapiens | C-type lectin domain family 2 member D(CLEC2D) |
| PTPN2 | 5771 | Homo sapiens | protein tyrosine phosphatase non-receptor type 2(PTPN2) |
| ISPD/CRPPA | | 2-C-methyl-D-erythritol 4-phosphate cytidylyltransferase-like protein | |
| ATP5A1/ATP5F1A | | ATP Synthase F1 Subunit Alpha | |
| LOC100996758 | | | |

| | |
|---|---|
| C20ORF196/SHLD1 | Component of the shieldin complex, which plays an important role in repair of D |
| KIAA0907/KHDC4 | KH Domain Containing 4, Pre-MRNA Splicing Factor |

### E PRC2 targets identified in "A" tab

IGF2R
SOX17
WNT11
WNT5A
WNT3
WNT5b
CDKN2AIP
LHX5
LHX9
LHX8
ISL1
ISL2
STAT1
STAT2
STAT6
SKIL
SETD4
SETBP1
SETDB2
SETD1B
BARX1
FBXO42
FBXO31
FBXO18
FBXO33
FBXO10

### F GO analysis of genes in tab "A"

| | |
|---|---|
| Analysis Type | PANTHER Overrepresentation Test (Released 20240807) |
| Annotation V | GO Ontology database DOI: 10.5281/zenodo.12173881 Released 2024-06-17 |
| Analyzed List | upload_1 (Homo sapiens) |
| Reference Lis | Homo sapiens (all genes in database) |
| Test Type: | FISHER |
| Correction: | FDR |

| GO biologica | Homo sapien | upload_1 (28 | upload_1 (ex | upload_1 (ov | upload_1 (fol | upload_1 (rav |
|---|---|---|---|---|---|---|
| membrane de | 5 | 5 | 0.7 | + | 7.15 | 5.32E-05 |
| positive regul | 4 | 4 | 0.56 | + | 7.15 | 3.81E-04 |
| insulin recep | 4 | 4 | 0.56 | + | 7.15 | 3.81E-04 |
| positive regul | 4 | 4 | 0.56 | + | 7.15 | 3.81E-04 |
| positive regul | 4 | 4 | 0.56 | + | 7.15 | 3.81E-04 |
| cellular respo | 7 | 6 | 0.98 | + | 6.13 | 4.58E-05 |

| | | | | | | |
|---|---|---|---|---|---|---|
| regulation of | 6 | 5 | 0.84 | + | 5.96 | 2.82E-04 |
| negative regu | 6 | 5 | 0.84 | + | 5.96 | 2.82E-04 |
| regulation of | 9 | 7 | 1.26 | + | 5.56 | 2.88E-05 |
| regulation of | 9 | 7 | 1.26 | + | 5.56 | 2.88E-05 |
| regulation of | 8 | 6 | 1.12 | + | 5.36 | 1.61E-04 |
| mesenchyma | 7 | 5 | 0.98 | + | 5.11 | 8.73E-04 |
| mesenchyma | 7 | 5 | 0.98 | + | 5.11 | 8.73E-04 |
| AV node cell t | 7 | 5 | 0.98 | + | 5.11 | 8.73E-04 |
| AV node cell a | 7 | 5 | 0.98 | + | 5.11 | 8.73E-04 |
| response to fl | 7 | 5 | 0.98 | + | 5.11 | 8.73E-04 |
| mesenchyma | 7 | 5 | 0.98 | + | 5.11 | 8.73E-04 |
| response to c | 9 | 6 | 1.26 | + | 4.77 | 4.26E-04 |
| regulatory nc | 18 | 12 | 2.52 | + | 4.77 | 4.45E-07 |
| metanephric | 10 | 6 | 1.4 | + | 4.29 | 9.40E-04 |
| siRNA-media | 10 | 6 | 1.4 | + | 4.29 | 9.40E-04 |
| siRNA-media | 10 | 6 | 1.4 | + | 4.29 | 9.40E-04 |
| mitochondria | 10 | 6 | 1.4 | + | 4.29 | 9.40E-04 |
| AV node cell t | 10 | 6 | 1.4 | + | 4.29 | 9.40E-04 |
| desmosome ( | 10 | 6 | 1.4 | + | 4.29 | 9.40E-04 |
| regulation of | 12 | 7 | 1.68 | + | 4.17 | 4.30E-04 |
| dendrite self- | 14 | 8 | 1.96 | + | 4.09 | 1.97E-04 |
| commissural | 13 | 7 | 1.82 | + | 3.85 | 8.19E-04 |
| membrane de | 17 | 9 | 2.38 | + | 3.79 | 1.69E-04 |
| phosphatidyl | 29 | 15 | 4.05 | + | 3.7 | 1.63E-06 |
| retrotranspo | 16 | 8 | 2.24 | + | 3.58 | 6.50E-04 |
| regulation of | 19 | 9 | 2.66 | + | 3.39 | 4.92E-04 |
| membrane de | 22 | 10 | 3.08 | + | 3.25 | 3.62E-04 |
| centrosome ( | 36 | 16 | 5.03 | + | 3.18 | 9.20E-06 |
| regulation of | 25 | 11 | 3.49 | + | 3.15 | 2.62E-04 |
| regulation of | 34 | 14 | 4.75 | + | 2.95 | 9.30E-05 |
| adipose tissu | 44 | 18 | 6.15 | + | 2.93 | 1.06E-05 |
| insulin-like gr | 32 | 13 | 4.47 | + | 2.91 | 1.94E-04 |
| regulation of | 40 | 16 | 5.59 | + | 2.86 | 4.55E-05 |
| regulation of | 33 | 13 | 4.61 | + | 2.82 | 2.80E-04 |
| regulation of | 28 | 11 | 3.91 | + | 2.81 | 8.43E-04 |
| calcium ion i | 28 | 11 | 3.91 | + | 2.81 | 8.43E-04 |
| negative regu | 31 | 12 | 4.33 | + | 2.77 | 5.77E-04 |
| phospholipid | 42 | 16 | 5.87 | + | 2.73 | 9.13E-05 |
| calcium ion i | 29 | 11 | 4.05 | + | 2.71 | 1.19E-03 |
| regulation of | 40 | 15 | 5.59 | + | 2.68 | 1.85E-04 |
| response to a | 38 | 14 | 5.31 | + | 2.64 | 3.74E-04 |
| calcium ion i | 41 | 15 | 5.73 | + | 2.62 | 2.55E-04 |
| membrane de | 55 | 20 | 7.69 | + | 2.6 | 2.78E-05 |
| peptidyl-tyro | 33 | 12 | 4.61 | + | 2.6 | 1.11E-03 |
| regulation of | 33 | 12 | 4.61 | + | 2.6 | 1.11E-03 |

| | | | | | | |
|---|---|---|---|---|---|---|
| animal organ | 39 | 14 | 5.45 | + | 2.57 | 5.09E-04 |
| insulin recep | 68 | 24 | 9.51 | + | 2.52 | 8.24E-06 |
| regulation of | 40 | 14 | 5.59 | + | 2.5 | 6.83E-04 |
| cAMP-mediat | 40 | 14 | 5.59 | + | 2.5 | 6.83E-04 |
| negative regu | 41 | 14 | 5.73 | + | 2.44 | 9.05E-04 |
| regulation of | 50 | 17 | 6.99 | + | 2.43 | 2.81E-04 |
| establishmer | 42 | 14 | 5.87 | + | 2.38 | 1.18E-03 |
| negative regu | 42 | 14 | 5.87 | + | 2.38 | 1.18E-03 |
| cell commun | 42 | 14 | 5.87 | + | 2.38 | 1.18E-03 |
| receptor loca | 45 | 15 | 6.29 | + | 2.38 | 8.00E-04 |
| extracellular | 46 | 15 | 6.43 | + | 2.33 | 1.04E-03 |
| heart looping | 65 | 21 | 9.09 | + | 2.31 | 1.85E-04 |
| cardiac cond | 67 | 21 | 9.37 | + | 2.24 | 2.51E-04 |
| regulation of | 64 | 20 | 8.95 | + | 2.24 | 3.72E-04 |
| negative regu | 61 | 19 | 8.53 | + | 2.23 | 5.53E-04 |
| negative regu | 58 | 18 | 8.11 | + | 2.22 | 8.23E-04 |
| embryonic he | 85 | 26 | 11.88 | + | 2.19 | 1.02E-04 |
| calcium ion t | 95 | 29 | 13.28 | + | 2.18 | 3.49E-05 |
| phosphatidyl | 69 | 21 | 9.65 | + | 2.18 | 3.61E-04 |
| regulation of | 66 | 20 | 9.23 | + | 2.17 | 5.31E-04 |
| calcium ion t | 212 | 64 | 29.64 | + | 2.16 | 8.64E-10 |
| determinatio | 70 | 21 | 9.79 | + | 2.15 | 4.37E-04 |
| embryonic he | 71 | 21 | 9.93 | + | 2.12 | 7.88E-04 |
| regulatory nc | 105 | 31 | 14.68 | + | 2.11 | 3.03E-05 |
| regulation of | 61 | 18 | 8.53 | + | 2.11 | 1.38E-03 |
| calcium ion t | 262 | 76 | 36.63 | + | 2.07 | 2.24E-10 |
| DNA conform | 89 | 25 | 12.44 | + | 2.01 | 5.52E-04 |
| regulation of | 82 | 23 | 11.46 | + | 2.01 | 1.08E-03 |
| DNA duplex u | 75 | 21 | 10.48 | + | 2 | 1.28E-03 |
| inorganic ion | 104 | 29 | 14.54 | + | 1.99 | 1.78E-04 |
| inorganic cat | 104 | 29 | 14.54 | + | 1.99 | 1.78E-04 |
| phosphatidyl | 126 | 35 | 17.61 | + | 1.99 | 4.59E-05 |
| axoneme ass | 101 | 28 | 14.12 | + | 1.98 | 2.57E-04 |
| microtubule | 130 | 36 | 18.17 | + | 1.98 | 3.63E-05 |
| axon guidanc | 224 | 62 | 31.31 | + | 1.98 | 9.17E-08 |
| protein K48-l | 94 | 26 | 13.14 | + | 1.98 | 4.69E-04 |
| neuron proje | 225 | 62 | 31.45 | + | 1.97 | 9.80E-08 |
| regulation of | 102 | 28 | 14.26 | + | 1.96 | 4.49E-04 |
| determinatio | 135 | 37 | 18.87 | + | 1.96 | 4.69E-05 |
| vesicle-medi | 136 | 37 | 19.01 | + | 1.95 | 5.09E-05 |
| positive regul | 92 | 25 | 12.86 | + | 1.94 | 7.66E-04 |
| left/right patt | 141 | 38 | 19.71 | + | 1.93 | 4.42E-05 |
| regulation of | 167 | 45 | 23.35 | + | 1.93 | 9.00E-06 |
| negative regu | 104 | 28 | 14.54 | + | 1.93 | 5.18E-04 |
| regulation of | 131 | 35 | 18.31 | + | 1.91 | 1.14E-04 |

| | | | | | | |
|---|---|---|---|---|---|---|
| phosphatidyl | 158 | 42 | 22.09 | + | 1.9 | 2.57E-05 |
| protein locali | 196 | 52 | 27.4 | + | 1.9 | 3.55E-06 |
| protein depho | 137 | 36 | 19.15 | + | 1.88 | 1.59E-04 |
| determinatio | 141 | 37 | 19.71 | + | 1.88 | 1.21E-04 |
| specification | 141 | 37 | 19.71 | + | 1.88 | 1.21E-04 |
| protein locali | 237 | 62 | 33.13 | + | 1.87 | 6.43E-07 |
| regulation of | 157 | 41 | 21.95 | + | 1.87 | 6.41E-05 |
| establishmer | 203 | 53 | 28.38 | + | 1.87 | 5.31E-06 |
| heterochrom | 112 | 29 | 15.66 | + | 1.85 | 8.51E-04 |
| cellular respo | 148 | 38 | 20.69 | + | 1.84 | 1.78E-04 |
| regulation of | 226 | 58 | 31.59 | + | 1.84 | 2.72E-06 |
| formation of p | 121 | 31 | 16.92 | + | 1.83 | 8.36E-04 |
| mesenchyma | 172 | 44 | 24.04 | + | 1.83 | 5.41E-05 |
| peptidyl-seri | 157 | 40 | 21.95 | + | 1.82 | 1.14E-04 |
| peptidyl-seri | 169 | 43 | 23.63 | + | 1.82 | 7.56E-05 |
| cell-cell junc | 126 | 32 | 17.61 | + | 1.82 | 6.67E-04 |
| phosphorylat | 548 | 137 | 76.61 | + | 1.79 | 3.53E-12 |
| establishmer | 132 | 33 | 18.45 | + | 1.79 | 8.93E-04 |
| dephosphory | 193 | 48 | 26.98 | + | 1.78 | 5.85E-05 |
| phospholipid | 242 | 60 | 33.83 | + | 1.77 | 6.09E-06 |
| glycerophosp | 202 | 50 | 28.24 | + | 1.77 | 3.79E-05 |
| positive regu | 150 | 37 | 20.97 | + | 1.76 | 5.11E-04 |
| intracellular | 199 | 49 | 27.82 | + | 1.76 | 5.23E-05 |
| lipid modifica | 167 | 41 | 23.35 | + | 1.76 | 2.79E-04 |
| positive regu | 139 | 34 | 19.43 | + | 1.75 | 8.33E-04 |
| calcium-mec | 160 | 39 | 22.37 | + | 1.74 | 5.04E-04 |
| protein phosp | 489 | 119 | 68.36 | + | 1.74 | 5.65E-10 |
| glycerolipid b | 224 | 54 | 31.31 | + | 1.72 | 4.10E-05 |
| regulation of | 303 | 73 | 42.36 | + | 1.72 | 2.27E-06 |
| plasma meml | 437 | 105 | 61.09 | + | 1.72 | 1.68E-08 |
| cell-cell junc | 179 | 43 | 25.02 | + | 1.72 | 3.03E-04 |
| cell projectic | 450 | 108 | 62.91 | + | 1.72 | 8.40E-09 |
| negative regu | 151 | 36 | 21.11 | + | 1.71 | 1.29E-03 |
| cilium organi | 374 | 89 | 52.28 | + | 1.7 | 3.43E-07 |
| axonogenesis | 366 | 87 | 51.17 | + | 1.7 | 4.00E-07 |
| cilium assem | 342 | 81 | 47.81 | + | 1.69 | 1.26E-06 |
| glycerophosp | 293 | 69 | 40.96 | + | 1.68 | 8.71E-06 |
| cell morphog | 443 | 104 | 61.93 | + | 1.68 | 7.11E-08 |
| negative regu | 179 | 42 | 25.02 | + | 1.68 | 6.91E-04 |
| regulation of | 243 | 57 | 33.97 | + | 1.68 | 7.81E-05 |
| kidney develo | 304 | 71 | 42.5 | + | 1.67 | 8.69E-06 |
| neuron proje | 485 | 113 | 67.8 | + | 1.67 | 2.78E-08 |
| plasma meml | 490 | 114 | 68.5 | + | 1.66 | 2.27E-08 |
| renal system | 314 | 73 | 43.9 | + | 1.66 | 8.05E-06 |
| import across | 164 | 38 | 22.93 | + | 1.66 | 1.41E-03 |

| | | | | | | |
|---|---|---|---|---|---|---|
| cell projectic | 495 | 114 | 69.2 | + | 1.65 | 3.99E-08 |
| cellular respo | 244 | 56 | 34.11 | + | 1.64 | 1.77E-04 |
| calcium ion h | 227 | 52 | 31.73 | + | 1.64 | 2.36E-04 |
| phospholipid | 364 | 83 | 50.89 | + | 1.63 | 5.62E-06 |
| regulation of | 229 | 52 | 32.01 | + | 1.62 | 3.48E-04 |
| microtubule- | 209 | 47 | 29.22 | + | 1.61 | 8.49E-04 |
| external enca | 277 | 62 | 38.72 | + | 1.6 | 1.55E-04 |
| modulation o | 479 | 107 | 66.96 | + | 1.6 | 6.58E-07 |
| positive regul | 215 | 48 | 30.06 | + | 1.6 | 9.99E-04 |
| cell morphog | 686 | 153 | 95.9 | + | 1.6 | 2.19E-09 |
| regulation of | 480 | 107 | 67.1 | + | 1.59 | 6.78E-07 |
| regulation of | 378 | 84 | 52.84 | + | 1.59 | 1.22E-05 |
| extracellular | 275 | 61 | 38.44 | + | 1.59 | 2.12E-04 |
| extracellular | 276 | 61 | 38.58 | + | 1.58 | 2.21E-04 |
| glycerolipid r | 359 | 79 | 50.19 | + | 1.57 | 3.03E-05 |
| cell division ( | 520 | 114 | 72.69 | + | 1.57 | 6.80E-07 |
| intracellular | 826 | 181 | 115.47 | + | 1.57 | 3.10E-10 |
| localization v | 548 | 120 | 76.61 | + | 1.57 | 3.54E-07 |
| microtubule ( | 557 | 121 | 77.87 | + | 1.55 | 4.53E-07 |
| cell surface r | 420 | 91 | 58.71 | + | 1.55 | 1.74E-05 |
| axon develop | 420 | 91 | 58.71 | + | 1.55 | 1.74E-05 |
| regulation of | 453 | 98 | 63.33 | + | 1.55 | 7.45E-06 |
| heart develop | 564 | 122 | 78.84 | + | 1.55 | 5.28E-07 |
| cellular respo | 278 | 60 | 38.86 | + | 1.54 | 4.70E-04 |
| regulation of | 297 | 64 | 41.52 | + | 1.54 | 3.65E-04 |
| Golgi vesicle | 288 | 62 | 40.26 | + | 1.54 | 4.21E-04 |
| metal ion tra | 646 | 139 | 90.31 | + | 1.54 | 1.29E-07 |
| plasma meml | 1148 | 247 | 160.49 | + | 1.54 | 9.19E-13 |
| regulation of | 265 | 57 | 37.05 | + | 1.54 | 8.93E-04 |
| import into c | 690 | 148 | 96.46 | + | 1.53 | 5.20E-08 |
| response to p | 374 | 80 | 52.28 | + | 1.53 | 8.26E-05 |
| cell projectic | 1196 | 255 | 167.2 | + | 1.53 | 9.93E-13 |
| intracellular | 1694 | 361 | 236.81 | + | 1.52 | 9.75E-18 |
| intracellular | 437 | 93 | 61.09 | + | 1.52 | 2.59E-05 |
| endocytosis ( | 503 | 107 | 70.32 | + | 1.52 | 6.29E-06 |
| intracellular | 428 | 91 | 59.83 | + | 1.52 | 3.94E-05 |
| regulation of | 302 | 64 | 42.22 | + | 1.52 | 5.68E-04 |
| cellular respo | 497 | 105 | 69.48 | + | 1.51 | 1.04E-05 |
| sensory syste | 393 | 83 | 54.94 | + | 1.51 | 9.19E-05 |
| actin cytoske | 550 | 116 | 76.89 | + | 1.51 | 4.72E-06 |
| cellular home | 679 | 143 | 94.92 | + | 1.51 | 3.49E-07 |
| telencephalo | 276 | 58 | 38.58 | + | 1.5 | 1.18E-03 |
| monoatomic | 514 | 108 | 71.86 | + | 1.5 | 1.06E-05 |
| establishmer | 343 | 72 | 47.95 | + | 1.5 | 3.79E-04 |
| phosphate-c | 1579 | 331 | 220.74 | + | 1.5 | 2.81E-15 |

| | | | | | | |
|---|---|---|---|---|---|---|
| neuron devel | 859 | 180 | 120.08 | + | 1.5 | 1.14E-08 |
| cellular respo | 573 | 120 | 80.1 | + | 1.5 | 4.07E-06 |
| phosphorus r | 1610 | 337 | 225.07 | + | 1.5 | 1.97E-15 |
| visual system | 387 | 81 | 54.1 | + | 1.5 | 1.51E-04 |
| Wnt signaling | 282 | 59 | 39.42 | + | 1.5 | 1.30E-03 |
| microtubule- | 398 | 83 | 55.64 | + | 1.49 | 1.40E-04 |
| actin filamen | 619 | 129 | 86.53 | + | 1.49 | 2.22E-06 |
| monoatomic | 504 | 105 | 70.46 | + | 1.49 | 2.14E-05 |
| regulation of | 456 | 95 | 63.75 | + | 1.49 | 5.15E-05 |
| monoatomic | 653 | 136 | 91.29 | + | 1.49 | 1.27E-06 |
| proteasome- | 303 | 63 | 42.36 | + | 1.49 | 1.07E-03 |
| intracellular | 569 | 118 | 79.54 | + | 1.48 | 9.08E-06 |
| circulatory sy | 926 | 192 | 129.45 | + | 1.48 | 8.88E-09 |
| protein locali | 459 | 95 | 64.17 | + | 1.48 | 7.21E-05 |
| regulation of | 363 | 75 | 50.75 | + | 1.48 | 4.20E-04 |
| eye developn | 383 | 79 | 53.54 | + | 1.48 | 3.33E-04 |
| cytoskeleton | 1256 | 259 | 175.58 | + | 1.48 | 2.67E-11 |
| proteasomal | 390 | 80 | 54.52 | + | 1.47 | 3.76E-04 |
| monoatomic | 815 | 167 | 113.93 | + | 1.47 | 2.22E-07 |
| peptidyl-ami | 523 | 107 | 73.11 | + | 1.46 | 4.00E-05 |
| microtubule- | 824 | 168 | 115.19 | + | 1.46 | 2.61E-07 |
| regulation of | 633 | 129 | 88.49 | + | 1.46 | 6.84E-06 |
| inorganic cat | 638 | 130 | 89.19 | + | 1.46 | 7.22E-06 |
| neuron proje | 692 | 141 | 96.74 | + | 1.46 | 2.57E-06 |
| regulation of | 437 | 89 | 61.09 | + | 1.46 | 2.07E-04 |
| regulation of | 649 | 132 | 90.73 | + | 1.45 | 6.57E-06 |
| membrane or | 744 | 151 | 104.01 | + | 1.45 | 1.48E-06 |
| regionalizatio | 409 | 83 | 57.18 | + | 1.45 | 3.94E-04 |
| mitotic cell c | 513 | 104 | 71.72 | + | 1.45 | 7.78E-05 |
| cell junction | 533 | 108 | 74.51 | + | 1.45 | 6.17E-05 |
| regulation of | 1017 | 206 | 142.17 | + | 1.45 | 1.74E-08 |
| anatomical s | 969 | 196 | 135.46 | + | 1.45 | 4.29E-08 |
| ubiquitin-dep | 490 | 99 | 68.5 | + | 1.45 | 1.24E-04 |
| monoatomic | 787 | 159 | 110.02 | + | 1.45 | 9.90E-07 |
| modification- | 505 | 102 | 70.6 | + | 1.44 | 1.18E-04 |
| regulation of | 828 | 167 | 115.75 | + | 1.44 | 6.24E-07 |
| embryonic or | 452 | 91 | 63.19 | + | 1.44 | 2.65E-04 |
| regulation of | 449 | 90 | 62.77 | + | 1.43 | 3.32E-04 |
| monoatomic | 980 | 196 | 137 | + | 1.43 | 1.11E-07 |
| organelle ass | 865 | 173 | 120.92 | + | 1.43 | 6.42E-07 |
| protein modi | 2055 | 411 | 287.28 | + | 1.43 | 2.48E-15 |
| neurogenesis | 1336 | 267 | 186.77 | + | 1.43 | 4.72E-10 |
| neuron differ | 1082 | 216 | 151.26 | + | 1.43 | 2.67E-08 |
| establishmer | 1298 | 259 | 181.46 | + | 1.43 | 1.03E-09 |
| protein ubiqu | 627 | 125 | 87.65 | + | 1.43 | 3.07E-05 |

| | | | | | | |
|---|---|---|---|---|---|---|
| proteolysis ir | 607 | 121 | 84.86 | + | 1.43 | 3.93E-05 |
| positive regul | 522 | 104 | 72.97 | + | 1.43 | 1.53E-04 |
| establishmer | 1628 | 324 | 227.59 | + | 1.42 | 7.68E-12 |
| intracellular | 558 | 111 | 78.01 | + | 1.42 | 9.20E-05 |
| cell-cell adhe | 549 | 109 | 76.75 | + | 1.42 | 1.33E-04 |
| protein locali | 1950 | 387 | 272.6 | + | 1.42 | 5.82E-14 |
| response to c | 902 | 179 | 126.1 | + | 1.42 | 6.56E-07 |
| cellular macr | 1958 | 388 | 273.72 | + | 1.42 | 6.54E-14 |
| forebrain dev | 409 | 81 | 57.18 | + | 1.42 | 1.13E-03 |
| regulation of | 510 | 101 | 71.3 | + | 1.42 | 2.77E-04 |
| modification- | 500 | 99 | 69.9 | + | 1.42 | 3.11E-04 |
| intracellular | 1193 | 236 | 166.78 | + | 1.42 | 1.22E-08 |
| protein modi | 704 | 139 | 98.42 | + | 1.41 | 1.91E-05 |
| vesicle-medi | 1298 | 256 | 181.46 | + | 1.41 | 3.79E-09 |
| pattern speci | 457 | 90 | 63.89 | + | 1.41 | 6.26E-04 |
| cellular resp | 508 | 100 | 71.02 | + | 1.41 | 3.48E-04 |
| inorganic ion | 432 | 85 | 60.39 | + | 1.41 | 9.46E-04 |
| organophosp | 549 | 108 | 76.75 | + | 1.41 | 2.20E-04 |
| generation of | 1160 | 228 | 162.16 | + | 1.41 | 4.52E-08 |
| protein trans | 1094 | 215 | 152.94 | + | 1.41 | 1.12E-07 |
| positive regul | 835 | 164 | 116.73 | + | 1.4 | 4.19E-06 |
| cellular local | 2659 | 522 | 371.72 | + | 1.4 | 5.82E-18 |
| protein catab | 724 | 142 | 101.21 | + | 1.4 | 1.97E-05 |
| regulation of | 459 | 90 | 64.17 | + | 1.4 | 8.13E-04 |
| brain develop | 730 | 143 | 102.05 | + | 1.4 | 2.10E-05 |
| organelle loc | 537 | 105 | 75.07 | + | 1.4 | 3.10E-04 |
| cell cycle pro | 881 | 172 | 123.16 | + | 1.4 | 3.58E-06 |
| macromolecu | 2239 | 437 | 313 | + | 1.4 | 1.60E-14 |
| negative regu | 683 | 133 | 95.48 | + | 1.39 | 5.14E-05 |
| central nervo | 1000 | 194 | 139.8 | + | 1.39 | 1.35E-06 |
| regulation of | 978 | 189 | 136.72 | + | 1.38 | 2.15E-06 |
| regulation of | 1186 | 229 | 165.8 | + | 1.38 | 1.66E-07 |
| cellular resp | 1010 | 195 | 141.19 | + | 1.38 | 1.54E-06 |
| protein modi | 808 | 156 | 112.96 | + | 1.38 | 2.06E-05 |
| chemical hor | 887 | 171 | 124 | + | 1.38 | 7.93E-06 |
| positive regul | 509 | 98 | 71.16 | + | 1.38 | 9.25E-04 |
| inorganic ion | 733 | 141 | 102.47 | + | 1.38 | 5.76E-05 |
| head develop | 780 | 150 | 109.04 | + | 1.38 | 3.80E-05 |
| lipid biosynth | 598 | 115 | 83.6 | + | 1.38 | 3.16E-04 |
| response to r | 1009 | 194 | 141.05 | + | 1.38 | 2.39E-06 |
| nervous syste | 2193 | 421 | 306.57 | + | 1.37 | 8.15E-13 |
| post-translat | 865 | 166 | 120.92 | + | 1.37 | 1.55E-05 |
| cell adhesior | 959 | 184 | 134.06 | + | 1.37 | 5.48E-06 |
| sensory orgar | 584 | 112 | 81.64 | + | 1.37 | 4.28E-04 |
| blood vessel | 527 | 101 | 73.67 | + | 1.37 | 9.00E-04 |

| | | | | | | |
|---|---|---|---|---|---|---|
| mitotic cell c | 595 | 114 | 83.18 | + | 1.37 | 3.86E-04 |
| embryonic m | 588 | 112 | 82.2 | + | 1.36 | 5.62E-04 |
| organelle org | 3099 | 590 | 433.23 | + | 1.36 | 2.09E-17 |
| anatomical s | 2220 | 422 | 310.35 | + | 1.36 | 3.50E-12 |
| regulation of | 979 | 186 | 136.86 | + | 1.36 | 8.51E-06 |
| enzyme-linke | 611 | 116 | 85.42 | + | 1.36 | 4.65E-04 |
| macromolecu | 2391 | 453 | 334.25 | + | 1.36 | 7.30E-13 |
| positive regul | 1115 | 211 | 155.87 | + | 1.35 | 2.39E-06 |
| protein-DNA | 878 | 166 | 122.74 | + | 1.35 | 3.49E-05 |
| homeostatic | 1429 | 270 | 199.77 | + | 1.35 | 8.74E-08 |
| embryo devel | 1043 | 197 | 145.81 | + | 1.35 | 6.68E-06 |
| tissue morph | 567 | 107 | 79.26 | + | 1.35 | 1.10E-03 |
| regulation of | 2413 | 455 | 337.33 | + | 1.35 | 1.42E-12 |
| regulation of | 1576 | 297 | 220.32 | + | 1.35 | 2.52E-08 |
| chromatin re | 637 | 120 | 89.05 | + | 1.35 | 5.94E-04 |
| chromatin or | 786 | 148 | 109.88 | + | 1.35 | 1.24E-04 |
| protein locali | 739 | 139 | 103.31 | + | 1.35 | 2.29E-04 |
| cell cycle (GO | 973 | 183 | 136.02 | + | 1.35 | 1.92E-05 |
| behavior (GO | 618 | 116 | 86.39 | + | 1.34 | 7.67E-04 |
| tube morphog | 682 | 128 | 95.34 | + | 1.34 | 3.94E-04 |
| multicellular | 3945 | 740 | 551.49 | + | 1.34 | 1.11E-20 |
| regulation of | 1984 | 372 | 277.36 | + | 1.34 | 5.15E-10 |
| cell developr | 2226 | 417 | 311.19 | + | 1.34 | 4.44E-11 |
| cellular comp | 5623 | 1052 | 786.07 | + | 1.34 | 1.20E-31 |
| regulation of | 868 | 162 | 121.34 | + | 1.34 | 9.20E-05 |
| system devel | 3527 | 657 | 493.06 | + | 1.33 | 2.81E-17 |
| establishmer | 3935 | 733 | 550.1 | + | 1.33 | 1.42E-19 |
| organophosp | 940 | 175 | 131.41 | + | 1.33 | 5.07E-05 |
| animal organ | 2843 | 529 | 397.44 | + | 1.33 | 1.19E-13 |
| localization ( | 4498 | 835 | 628.8 | + | 1.33 | 2.12E-22 |
| regulation of | 3423 | 633 | 478.52 | + | 1.32 | 7.22E-16 |
| transport (GO | 3665 | 677 | 512.35 | + | 1.32 | 5.36E-17 |
| cellular comp | 5838 | 1078 | 816.13 | + | 1.32 | 3.64E-30 |
| regulation of | 3428 | 631 | 479.22 | + | 1.32 | 2.26E-15 |
| DNA metabol | 735 | 135 | 102.75 | + | 1.31 | 7.63E-04 |
| response to c | 1507 | 276 | 210.67 | + | 1.31 | 1.11E-06 |
| cellular lipid | 919 | 168 | 128.47 | + | 1.31 | 2.10E-04 |
| cellular comp | 2459 | 448 | 343.76 | + | 1.3 | 4.28E-10 |
| nitrogen com | 1526 | 278 | 213.33 | + | 1.3 | 1.56E-06 |
| proteolysis (G | 1208 | 219 | 168.87 | + | 1.3 | 3.26E-05 |
| DNA damage | 780 | 141 | 109.04 | + | 1.29 | 1.08E-03 |
| regulation of | 2822 | 510 | 394.5 | + | 1.29 | 6.54E-11 |
| negative regu | 1400 | 253 | 195.71 | + | 1.29 | 9.10E-06 |
| negative regu | 1401 | 253 | 195.85 | + | 1.29 | 1.09E-05 |
| response to h | 787 | 142 | 110.02 | + | 1.29 | 1.14E-03 |

| | | | | | | |
|---|---|---|---|---|---|---|
| negative regu | 1309 | 236 | 182.99 | + | 1.29 | 2.56E-05 |
| transmembra | 1283 | 231 | 179.36 | + | 1.29 | 3.13E-05 |
| regulation of | 2409 | 432 | 336.77 | + | 1.28 | 8.19E-09 |
| regulation of | 3017 | 541 | 421.76 | + | 1.28 | 4.77E-11 |
| organonitrog | 1116 | 200 | 156.01 | + | 1.28 | 1.57E-04 |
| regulation of | 1122 | 201 | 156.85 | + | 1.28 | 1.63E-04 |
| positive regu | 1140 | 204 | 159.37 | + | 1.28 | 1.53E-04 |
| regulation of | 1121 | 200 | 156.71 | + | 1.28 | 1.95E-04 |
| regulation of | 1940 | 345 | 271.2 | + | 1.27 | 8.34E-07 |
| anatomical s | 5208 | 926 | 728.06 | + | 1.27 | 3.71E-19 |
| tube develop | 900 | 160 | 125.82 | + | 1.27 | 1.15E-03 |
| positive regu | 1743 | 309 | 243.66 | + | 1.27 | 4.42E-06 |
| nucleic acid | 2140 | 379 | 299.16 | + | 1.27 | 3.19E-07 |
| RNA metabol | 1519 | 268 | 212.35 | + | 1.26 | 3.21E-05 |
| regulation of | 941 | 166 | 131.55 | + | 1.26 | 1.25E-03 |
| positive regu | 1678 | 296 | 234.58 | + | 1.26 | 1.20E-05 |
| cellular comp | 2705 | 477 | 378.15 | + | 1.26 | 1.06E-08 |
| positive regu | 1681 | 296 | 235 | + | 1.26 | 1.44E-05 |
| nucleic acid | 1455 | 256 | 203.4 | + | 1.26 | 6.18E-05 |
| development | 5711 | 1003 | 798.37 | + | 1.26 | 2.27E-19 |
| regulation of | 1042 | 183 | 145.67 | + | 1.26 | 9.45E-04 |
| cellular resp | 1589 | 279 | 222.14 | + | 1.26 | 3.34E-05 |
| cellular resp | 1112 | 195 | 155.45 | + | 1.25 | 6.10E-04 |
| nucleobase-c | 2697 | 472 | 377.03 | + | 1.25 | 3.44E-08 |
| positive regu | 1741 | 304 | 243.38 | + | 1.25 | 1.98E-05 |
| positive regu | 1323 | 231 | 184.95 | + | 1.25 | 2.59E-04 |
| protein meta | 3530 | 616 | 493.48 | + | 1.25 | 1.75E-10 |
| positive regu | 1806 | 315 | 252.47 | + | 1.25 | 1.67E-05 |
| positive regu | 2009 | 350 | 280.85 | + | 1.25 | 5.50E-06 |
| protein-cont | 2044 | 356 | 285.74 | + | 1.25 | 4.71E-06 |
| cellular deve | 3641 | 634 | 509 | + | 1.25 | 1.23E-10 |
| RNA biosynth | 1367 | 238 | 191.1 | + | 1.25 | 2.35E-04 |
| regulation of | 1082 | 188 | 151.26 | + | 1.24 | 1.36E-03 |
| cell differenti | 3638 | 632 | 508.58 | + | 1.24 | 2.04E-10 |
| cellular resp | 1908 | 331 | 266.73 | + | 1.24 | 1.44E-05 |
| primary meta | 6886 | 1192 | 962.63 | + | 1.24 | 6.06E-22 |
| positive regu | 5601 | 968 | 783 | + | 1.24 | 2.60E-16 |
| cellular meta | 5659 | 978 | 791.11 | + | 1.24 | 1.52E-16 |
| regulation of | 1384 | 239 | 193.48 | + | 1.24 | 3.52E-04 |
| nucleobase-c | 1744 | 301 | 243.8 | + | 1.23 | 6.03E-05 |
| positive regu | 6177 | 1065 | 863.52 | + | 1.23 | 3.97E-18 |
| tissue develo | 1748 | 301 | 244.36 | + | 1.23 | 7.15E-05 |
| regulation of | 1534 | 264 | 214.45 | + | 1.23 | 2.34E-04 |
| lipid metabol | 1227 | 211 | 171.53 | + | 1.23 | 1.07E-03 |
| macromolecu | 5602 | 958 | 783.14 | + | 1.22 | 8.62E-15 |

| | | | | | | |
|---|---|---|---|---|---|---|
| response to e | 1375 | 235 | 192.22 | + | 1.22 | 8.22E-04 |
| negative regu | 1656 | 283 | 231.5 | + | 1.22 | 2.16E-04 |
| positive regul | 1558 | 265 | 217.8 | + | 1.22 | 4.66E-04 |
| regulation of | 2110 | 357 | 294.97 | + | 1.21 | 5.98E-05 |
| regulation of | 4010 | 677 | 560.58 | + | 1.21 | 6.94E-09 |
| regulation of | 1665 | 281 | 232.76 | + | 1.21 | 5.24E-04 |
| catabolic pro | 1944 | 328 | 271.76 | + | 1.21 | 1.55E-04 |
| metabolic pro | 7829 | 1319 | 1094.46 | + | 1.21 | 3.29E-20 |
| organonitrog | 4641 | 781 | 648.79 | + | 1.2 | 4.47E-10 |
| cell surface r | 2057 | 345 | 287.56 | + | 1.2 | 1.72E-04 |
| negative regu | 4749 | 795 | 663.89 | + | 1.2 | 8.34E-10 |
| positive regul | 2241 | 375 | 313.28 | + | 1.2 | 9.36E-05 |
| positive regul | 3677 | 615 | 514.03 | + | 1.2 | 2.00E-07 |
| positive regul | 3124 | 522 | 436.72 | + | 1.2 | 2.84E-06 |
| positive regul | 3364 | 561 | 470.27 | + | 1.19 | 1.29E-06 |
| negative regu | 5171 | 859 | 722.88 | + | 1.19 | 5.85E-10 |
| cellular biosy | 3530 | 584 | 493.48 | + | 1.18 | 2.31E-06 |
| negative regu | 2815 | 464 | 393.53 | + | 1.18 | 5.35E-05 |
| negative regu | 2602 | 425 | 363.75 | + | 1.17 | 2.81E-04 |
| multicellular | 6239 | 1015 | 872.19 | + | 1.16 | 6.74E-10 |
| positive regul | 2668 | 434 | 372.98 | + | 1.16 | 3.28E-04 |
| macromolecu | 2948 | 479 | 412.12 | + | 1.16 | 1.69E-04 |
| negative regu | 2425 | 394 | 339.01 | + | 1.16 | 7.55E-04 |
| biosynthetic | 4002 | 649 | 559.46 | + | 1.16 | 7.73E-06 |
| positive regul | 2568 | 416 | 359 | + | 1.16 | 6.54E-04 |
| positive regul | 2625 | 425 | 366.96 | + | 1.16 | 5.89E-04 |
| regulation of | 2931 | 473 | 409.74 | + | 1.15 | 3.59E-04 |
| cellular proc | 14769 | 2375 | 2064.65 | + | 1.15 | 3.97E-47 |
| cellular respo | 6306 | 996 | 881.55 | + | 1.13 | 8.21E-07 |
| cell commun | 5158 | 814 | 721.07 | + | 1.13 | 1.97E-05 |
| signaling (GC | 5116 | 806 | 715.2 | + | 1.13 | 3.12E-05 |
| regulation of | 11711 | 1838 | 1637.15 | + | 1.12 | 2.59E-16 |
| biological reg | 12149 | 1903 | 1698.38 | + | 1.12 | 3.38E-17 |
| regulation of | 11103 | 1737 | 1552.15 | + | 1.12 | 8.04E-14 |
| signal transdu | 4794 | 741 | 670.18 | + | 1.11 | 8.69E-04 |
| regulation of | 6519 | 1003 | 911.33 | + | 1.1 | 8.39E-05 |
| regulation of | 5837 | 896 | 815.99 | + | 1.1 | 3.92E-04 |
| regulation of | 5544 | 851 | 775.03 | + | 1.1 | 6.75E-04 |
| regulation of | 5998 | 918 | 838.5 | + | 1.09 | 4.74E-04 |
| response to s | 8080 | 1230 | 1129.55 | + | 1.09 | 3.83E-05 |
| biological_pr | 17791 | 2697 | 2487.11 | + | 1.08 | 9.03E-41 |
| immune syste | 2301 | 265 | 321.67 | - | 0.82 | 2.42E-04 |
| immune resp | 1662 | 157 | 232.34 | - | 0.68 | 7.83E-09 |
| sensory perce | 976 | 89 | 136.44 | - | 0.65 | 2.69E-06 |
| G protein-co | 1255 | 104 | 175.44 | - | 0.59 | 2.44E-10 |

| | | | | | | |
|---|---|---|---|---|---|---|
| adaptive imm | 677 | 49 | 94.64 | - | 0.52 | 3.76E-08 |
| Unclassified ( | 2789 | 180 | 389.89 | - | 0.46 | 9.03E-41 |
| defense resp | 302 | 19 | 42.22 | - | 0.45 | 3.58E-05 |
| cell killing (G | 138 | 7 | 19.29 | - | 0.36 | 1.21E-03 |
| detection of s | 675 | 28 | 94.36 | - | 0.3 | 2.04E-17 |
| antimicrobia | 144 | 4 | 20.13 | - | 0.2 | 9.48E-06 |
| disruption of | 84 | 2 | 11.74 | - | 0.17 | 7.07E-04 |
| antimicrobia | 97 | 2 | 13.56 | - | 0.15 | 1.00E-04 |
| sensory perce | 542 | 11 | 75.77 | - | 0.15 | 1.86E-22 |
| detection of s | 558 | 11 | 78.01 | - | 0.14 | 1.73E-23 |
| sensory perce | 466 | 8 | 65.14 | - | 0.12 | 1.00E-20 |
| disruption of | 81 | 1 | 11.32 | - | 0.09 | 1.42E-04 |
| killing of cells | 81 | 1 | 11.32 | - | 0.09 | 1.42E-04 |
| detection of ( | 520 | 6 | 72.69 | - | 0.08 | 4.63E-26 |
| detection of ( | 488 | 4 | 68.22 | - | 0.06 | 1.71E-26 |
| detection of ( | 440 | 2 | 61.51 | - | 0.03 | 3.25E-26 |

ypermethylated DMRs in Jonathan

LKBH3)

CACNA2D2)

DAMTS15)

DAMTS14)
PRPSAP2)

DAMTS18)

)
)

:3)

RKAG2)

nains 1(TMEFF1)

)

)

R1)

1)

2)

4)

l6)

in(VMAC)

ed 1(KHDRBS1)

ed 3(KHDRBS3)
