## Supplemental document for "Epigenomic insights into extreme longevity in the world’s oldest terrestrial animal, Jonathan"

**Jonathan’s estimated age**

Although Jonathan's exact age is unknown, we estimate his age to be at least 192 based on the following work.

Information about the existence of giant tortoises on St. Helena comes from the work of Dr. Authur Loveridge. Dr. Loveridge was a Professor of Biology at Harvard University and joined its Museum of Comparative Zoology in 1924 as Curator of Herpetology until 1957^1^. Upon his retirement, he moved to St. Helena and contributed several articles on the wildlife on the island. He cites two bodies of work, in his published articles in the island’s *Wirebird* magazine: Work by Dr. Albert Gunther, who states the Island had a persistent occupation of giant tortoises preceding Napoleon's exile (1815-1821), with the first arriving via the Mauritius island in 1776^2,3^. He cites “Plantation Notes” from St. Helena, where a letter was found from St. Helena’s governor Grey Wilson (1890-1897) addressed to Captain P. Oliver and dated 10/28/1894, stating that a fully grown male giant tortoise was imported to the island in 1882^4^. After this male arrived, only it and a female existed on the island until the female died in 1918. Her body was examined and was found to contain eggs. The lone existing male, in 1918 would later be named Jonathan^2^.

An old photograph of Jonathan and another tortoise has been circulating in the news for at least ten years, with a publication in 2014 by the BBC and again in 2016 by the Daily Mail (**Extended Data Fig. 1A**)^5,6^. The BBC article claimed the photograph was from an 1882 celebration of Jonathan’s arrival. Jonathan can be identified in this picture because he has a unique marking on one of his scutes, which is apparent in recent photographs of Jonathan and his veterinarian Joe Hollins (**Extended Data Fig. 1B**). This photograph was examined by Geoffrey Batchen, professor of the History of Art at Oxford University, with a special expertise on the history of photography. Dr. Batchen believes this photograph is an albumen print from a glass collodion negative and was likely taken between 1850 and 1900. The earliest record of this photograph identified was a postcard dated December 1907 (**Extended Data Fig. 1C**)^7^. If we assume this photograph was taken in 1907, then Jonathan is at least 167 years old, however, we believe, as detailed below, he is much older.

After the Daily Mail circulation of this photograph in 2016, an English lady, named Rosemary Rees, contacted the island to inform them that her paternal grandfather had the same photograph in his possession at the time of his death in 1886 (**Extended Data Fig. 2A**). According to Mrs. Rees, her grandfather lived at Bedford in the Cape colony of South Africa, where he was a medical officer aboard the Walmer and Dublin Castle ship, which frequently called upon St. Helena. On the back of her photograph, there is a stamp that reads: “A.L. Innes, Jamestown, St. Helena Studio” (**Extended Data Fig. 2B**). In addition, there was a description of the dimensions of Jonathan that match his current dimensions and thus indicate that at the time of the photograph, he was fully grown. In addition, we found a collection of photographs at Christie's auction house, auctioned on April 8, 2004 (**Extended Data Fig. 2C**)^8^. The collection is labeled “St. Helena, 1889-1890 – An album of 76 photographs of St. Helena by B. Wood, B. Grant, and others, printed by A.L. Innes, St. Helena Studio, Jamestown…”. We have not determined if the photograph of Jonathan is part of this collection, as Christie’s reached out to the buyer in 2024 without reply, but this establishes that “A.L. Innes” was printing photographs at least as early as 1889. Regardless, we believe the above evidence is sufficient to estimate the photograph was taken, likely in celebration of Jonathan’s arrival on the island, between 1882 and 1886.

A paucity of data limits our understanding of Aldabra shell growth rates. Dr. P. Grubbs, of the Department of Zoology, University of Ghana mostly completed data relating carapace length to age. Dr. Grubb took measurements from the Aldabra atoll at three locations: Cinq and Takamaka locations on the Grande-Terre island, and one location on Malabar island^9^. The range of ages in this cohort was from one to 31 years. In this study, a carapace length (**Extended Data Fig. 3A**) was measured on each tortoise and a presumed age was determined by counting the number of annuli present in one scute (**Extended Data Fig. 3B**). The scutes are bony units that cover and comprise the tortoise shell, and the annuli are growth rings within the scutes^10^. Data from Dr. Gubb's paper used the annuli to determine the age of the tortoises. However, he noted that beyond 25 annuli, it becomes difficult to count as the annuli get compressed together. Thus, for this reason, the ages of the Aldabras were all less than 32. A total of 368 Aldabras were measured from Grand Terre with ages ranging from one to 24, and 58 Aldabras were measured from Malabar with ages 12 to 31 (**Extended Data Fig.4**)^9^. The longest carapace length found was 95 cm from a 30-year-old Malabar Aldabra and the largest standard deviation in carapace length was 10 cm, found within a 12-year-old group from Grande-Terre.

Next, from the photograph containing Jonathan (**Extended Data Fig. 5)**, we estimated his carapace length. Based on historical data we assumed the gentleman standing closest to Jonathan was 66.8 inches or approximately 170.18 cm^11^. From the photograph of Jonathan and the use of a photo measuring technique, we determined Jonathan's carapace length is 0.65 that of the gentleman's height (**Extended Data Fig. 5**)^12^. Thus, we calculate Joanthan’s carapace length to be approximately 110 cm. Jonathan’s current carapace length is also 110 cm, implying he was fully grown in the photograph. Based on our work regarding Jonathan’s biogeography (**Fig. 1B-D**), we believe Jonathan likely arose from the Grand Terre island of the Aldabra atoll as he is more proximal to this island than Malabar. Using linear regression of Dr. Grubb’s data we produced three plots (size vs age) for all sites of measurements (**Extended Data Fig. 4**). We extrapolated from these plots and based on the size of 110 cm, Jonathan is predicted to be 45 with the Takamaka data, 62.2 from the Cinq data, and 46 from the Malabar data. If we average the two Grand Terre sites (Takamaka and Cinq) we get an average age of 53.6 and if we average all three sites (Takamaka, Cinq, and Malabar) we get an average age of 51.1 years. From the residual plots we can tell that growth is not continuous and thus, a linear regression plot does not faithfully represent the data and likely underestimates his age when he is 110 cm, as growth rates are declining in the longer animals. We think the differences in the three sites are likely due to population density and access to food. This data is limited in three ways: first, annuli were used to estimate age, even though Dr. Grubb states that beyond 25 annuli it becomes difficult to count the annuli (four animals from Malabar are estimated to be older than 25), secondly, several of the “longer” specimens only had a measurement of one individual and finally, these measurements were made in 1967 and 1968. Since Jonathan was likely present on one of these islands 136 years earlier, the population density and food access at that time were likely different. It is also unlikely Jonathan was captured in his 50th year, and he likely remained on the island several years after his 50th birthday. Thus, we conservatively estimate it took him 50 years to reach 110 cm and we conservatively estimate he was 50 years old in the Jonathan photograph. If he is 50 in the Jonathan photograph and it was taken in 1882, Jonathan’s birthdate would be approximately 1832, and his current age would be 192. However, Jonathan could be older than 50 in the photograph and his current age could be older than 192.

In conclusion, Jonathan’s exact age cannot be determined with absolute certainty using these data. Recent work in epigenetics, especially DNA methylation, has produced epigenetic clocks that closely estimate chronological age. For instance, DNA methylation data in bats has been used to construct an epigenetic clock that can accurately predict their chronological age^13^. Hopefully, an epigenetic clock can be generated for Aldabra, and from such a clock we could obtain a more reliable estimate of Jonathan’s age.

**Insertions and deletions**

Lastly, we examined our gene set for insertion and deletions. Ten genes had one amino-acid change (*ATRX*, *FOSL2*, *FUS*, *HLA-DRB1*, *HNRNPD*, *IFNL3*, *IGF1R*, *LY9*, *PPP1R26*, and *TDG*), seven with insertions, and three with deletions (*HNRNPD*, *LY9*, and *TDG*)(**Supplementary Table 9**). Interestingly, previous work on Lonesome George found that in both him and the Aldabra, a single amino acid deletion was identified in IGF2R and a variant was found in IGF1R, both of which were verified^14^. Thus, Jonathan has alterations in both *IGF1R* and *IGF2R*.

**Extended Data Figures/Table Legends**

**Extended Data Figure 1.** **Jonathan photograph.**  **A.** The photograph shows two giant tortoises, with Jonathan being on the left.   The yellow arrow points to a unique scute on Jonathan’s shell.   The gentleman behind Jonathan is thought to be Helenians based on the writing on the back of Rosemary Rees’s copy.  The location of the photograph is at the corner of Plantation paddock, near the public gate to the tortoise corridor on the island of St. Helena.  We believe this photograph was taken between 1882 and 1886 and likely in 1882.  **B.** A more recent photograph of Jonathan and his vet, Joe Hollins.  The yellow arrow points to a unique scute on Jonathan’s shell as in 1A.  The location of the photograph is the grounds of the Plantation House, the official residence of the governor of St. Helena.  **C.**  A postcard reproducing the Jonathan photograph.  The date on the postcard is “Xmas 1907”, which would be sometime in December of 1907^7^.

**Note:** **Extended Data Figure 1A: The source of the “Jonathan photograph” is from the personal collection of Joe Hollins and Rosemary Rees and not from a database. Extended Data Figure 1B includes a photograph of author Joe Hollins with Jonathan.**

**Extended Data Figure 2. Determining the age of Jonathan's photograph.** **A.** The copy of the Jonathan photograph, presented by Rosemary Rees.  The back of the photograph is shown as well with the printer’s stamp and the writing on her copy.  **B.**  A close-up of the printer’s stamp is shown as in “A”.  The top reads “A.L. INNES”, the bottom reads, “JAMESTOWN”, and in the center of the stamp reads “ST. HELENA STUDIOS”.  **c.**  The Christie’s auction web page depicts “THE QUENTIN KEYNES COLLECTION, Lot 93”.  The red underlined portion of the collection description highlights the date “1889-1890” of the collection and that it was printed by “A.L. Innes, St. Helena Studio, Jamestown”^8^.

**Extended Data Figure 3.   Carapace length and sute size with annulus. A.**  A reproduction of “Figure 1” in Dr. Grubbs's paper^9^.  The left figure depicts a ventral view of the Aldabra shell, with “A” being the carapace length.  **B.**  A dorsal view of a scute, with “D” being the fourth annulus (4th year) of the scute and “E” being the width of the scute. **C.** Photograph of 3rd dorsal scute of an Aldarba.

**Extended Data Figure 4.**  **Replotting of Dr. Grubbs growth data for Aldabras on the Aldabra atoll.**   A replot of Dr. Grubbs's data, with the x-axis representing the carapace length in millimeters (mm) and the y-axis representing the “presumed age” in years of the tortoises^9^.  The age was calculated by counting the number of annuli of a scute.  Three separate populations are plotted, one population from Malabar and two populations from Grande Terre (Takamaka and Cinq).  Based on linear regression, a 110 cm carapace length tortoise would have an age of 45 years old with the Takamaka data, 62.2 years old with the Cinq data, and 46 years old with the data from Malabar.   The average of these populations gives a predicted age of 51.3 years.  Open red dots = residuals.  A residual in linear regression is the difference in the actual value and the value predicted by the regression of the line to that point and equals observed value minus (-) predicted value.

**Extended Data Figure 5.** **Carapace length of Jonathan in Jonathan's photograph.** The Jonathan photograph is shown with markings used to measure his carapace length.  The closest person to Jonathan is set as 1.0 (red markings) and then Jonathan’s carapace measurement is determined by a photo measuring technique to be 0.65 in relation to the human measurement.   Based on historical data we assumed the gentleman standing closest to Jonathan was 66.8 inches or approximately 170.18 cm^11^ and based on this we calculated Joanthan’s carapace length to be 110 cm.

**Extended Data Figure 6. Geographic location of St. Helena and Aldabra Atoll.**

**A.** Location of St. Helena in the South Atlantic Ocean.  **B.** Location of the Aldabra Atoll in the Indian Ocean. Maps generated from Google Earth.  Bar = 1000 km, White Bars = 3.2 km.

**Extended Data Figure 7**. **The *de novo* sequencing and assembly of the reference genome of Tank and the consensus genome of Jonathan**. **A**. Assembly statistics of the reference genome of Jonathan and Tank. **B.** Assembly metrics, sequence composition, and BUSCO completeness scores for the Sauropsida dataset and Tank. The number of annotated coding and non-coding genes in Tank and Jonathan’s genome.

**Extended Data Figure 8**. **Lonesome George’s location in principal component plot.** Principal component analysis plot of 32 *Aldabrachelys gigantea* previously plotted from the Islands of Malabar and Grande Terre of the Aldabra atoll.  Included are the two Zurich Zoo Aldabras, Tank, and Jonathan, and as a negative control, Lonesome George (LG).

**Extended Data Figure 9.**  **GO analysis of Jonathan’s positively selected genes.** Categories include aging, development, immunity, and signaling.

**Extended Data Figure 10.   STRING analysis of Jonathan’s positively selected genes.**  Using the STRING algorithm protein-protein connectivity networks were identified.  Listed are networks of interest.

**Extended Data Figure 11. Variants found in Lonesome George, were also found in Jonathan.** **A.** nsNSV in the gene *XRCC6* (K556R). **B.** nsNSV in *DCLRE1B* (R498C) **C.** nsNSV in *IGF1R* (N724D).  **D.** nsNSV in *ALDH2* (M487T).  **E.** nsNSV in *IGF2R* (Y1758del).  **F.** nsNSV in *GSK3A* (R272Q).  **G.** nsNSV in *MIF* (N111C).   TORT, Jonathan; CABP, Lonesome George.

**Extended Data Figure 12. Expansion of *LAMTOR4* gene in giant tortoises. A.** Comparison of both copies of LAMTOR4 in Jonathan to a single copy in humans.  LAMTOR4JON1, Jonathan LAMTOR4 copy 1; LAMTOR4JON2, Jonathan LAMTOR4 copy 2.  nsNSVs in copies were evaluated by protein function algorithms, Provean, Polyphen2, and SIFT.  **B.**   Comparison of normal copy of LAMTOR4 between human (H), Lonesome George (LG), Hermania (Herm), Tank, and Jon (Jonathan).   Comparison of copy #2 in the above tortoises to humans.  **C.**  Phylogenetic tree relationship between the LAMTOR4 copies (millions of years).

**Extended Data Figure 13. GO analysis of Jonathan’s LERs (169 genes) using STRING database.**

**Supplementary Table 1**. **Jonathan’s and Tank’s genome assembly metrics.**

**Supplementary Table 2.** **Jonathan’s positively selected genes (PSGs).**  Provean and SIFT analysis of each variant. Only genes with omega values greater than one are shown.  Jonathan’s PSGs shared with other long-lived species and with LongevityMap and GeneAge.

**Supplementary Table 3.**  **Unique nsNSVs found only in Jonathan.**

**Supplementary Table 4.**  **Evaluation of Jonathan’s unique nsNSVs by protein language models.** Protein Language models predictions (Tranception L no retrieval, ESM1v ensemble, Progen2 base, RITA XL, and VESPA1). Ratio A and Ratio B.

**Supplementary Table 5. Genes that are found in more than one protein language model.**

**Supplementary Table 6. Gene expansion in Jonathan.** Tortoise-specific gene expansions.  Comparing Jonathan, Lonesome George, and other tortoises.

**Supplementary Table 7. Genes within Jonathan’s hypermethylated DMRs and GO analysis**

**Supplementary Table 8. Genes within Jonathan’s low-entropy regions (LERs).**

**Supplementary Table 9.  Insertions and deletions in Jonathan.**

1.	History | Museum of Comparative Zoology. https://www.mcz.harvard.edu/herpetology-history.

2.	Loveridge, A. Our tortoises. Part 1. The truth about Jonathan. *Wirebird* vol. 3 (1962).

3.	Gunther, A. One hundred and tenth session, 1897-98,. **110**, 14–29 (1898).

4.	Galway, S. H. Plantation notes, St. Helena (1902-1910). (1910).

5.	Kettle, S. Meet Jonathan, St Helena’s 182-year-old giant tortoise. *BBC* (2014).

6.	Greenhill, S. Still sprightly at 183, the oldest creature on Earth. *Daily Mail* (2016).

7.	Richardson, R. Early Postcards of St. Helena, 1900-1910. http://www.tpa-project.info/TPA20_3_St_Helena.pdf.

8.	Christie’s. *The Quentin Keynes Collection-Live Auction 6890*. (1889).

9.	Grubb, P. The growth, ecology, and population of structure of giant tortoises on aldabra. *Phil Trans R Soc Lond B* **260**, 327–392 (1971).

10.	Moustakas-Verho, J. E., Cebra-Thomas, J. & Gilbert, S. F. Patterning of the turtle shell. *Curr Opin Genet Dev* **45**, 124–131 (2017).

11.	Floud, R. & Harris, B. Health, Height, and Welfare: Britain, 1700-1980. in *Health and Welfare during Industrialization* 91–126 (University of Chicago Press, 1997).

12.	Photomeasure. Eleif.

13.	Wilkinson, G. S. *et al.* DNA methylation predicts age and provides insight into exceptional longevity of bats. *Nat Commun* **12**, 1615 (2021).

14.	Quesada, V. *et al.* Giant tortoise genomes provide insights into longevity and age-related disease. *Nat Ecol Evol* **3**, 87–95 (2019).
